# Protein Structure Description with *ρ, θ* and *ϕ*: A Case Study with Caenopore-5

**DOI:** 10.1101/2025.01.08.631847

**Authors:** Wei Li

## Abstract

Since its establishment in 1971, the Protein Data Bank (PDB) has been using Cartesian coordinate system (CCS) as the standard framework for protein structure description with *x, y, z*. Despite the interconvertibility of CCS and spherical coordinate systems (SCS, *ρ, θ* and *ϕ*), CCS remains to date the default and the only framework for protein structure description in PDB. Recent advances in protein structure prediction (e.g., AlphaFold) revolutionized the field by integrating deep learning algorithms with experimental structural data, achieving unprecedented accuracy of protein structure prediction and relying on Cartesian representation of protein structures to extract geometric features. To this end, questions remain about what drives the next stage of continued performance improvement of protein structure prediction. Therefore, this article introduces an alternative coordinate system for protein structure description and feature extraction. Using Caenopore-5 as an example, this article redefines protein backbone structures using atomic bonding networks (ABN) within the SCS framework (ABN-SCS), leading to the extraction of a set of spherical parameters (*ρ, θ* and *ϕ*) from the NMR ensemble of Caenopore-5, encompassing 477 covalent bonds and 80 peptide bonds within its backbone for each structural model in its NMR ensemble. Finally, this work demonstrates that ABN-SCS enables characterization of spherical bond-level geometries, expanding the feature space available for computational pipelines such as AlphaFold2, and argues that integrating ABN-SCS features into protein structure prediction pipelines can enhance geometric fidelity, and that the time is now ripe for the trapped spherical features [*ρ, θ, ϕ*] to be integrated into algorithms such as AF2 towards protein structure prediction with improved performance.

## Introduction

### [*x, y, z*] and [*ρ, θ, ϕ*] for protein structure description

In 1971, Protein Data Bank (PDB) was established as the global repository for experimentally determined biomolecular 3D structures [1–4]. Since then, it has been acting as a catalyst [5] for structural biology, i.e., understanding [6–8], determination [9–12], description [13–18] and prediction [19–24] of the structure of protein, the essential building block of life. Since 1971 [1, 2], Cartesian coordinate system (CCS) has served as the default geometric framework to specify atomic positions as [*x, y, z*] (Figure 1), primarily because it directly corresponds to experimental data from techniques such as X-ray crystallography and NMR spectroscopy [25, 26], which inherently produce atomic positions as [*x, y, z*]. By definition, CCS treats each atom as an independent point in space and conceptualizes protein structure as a set of discrete points, without explicitly encoding their underlying chemical connectivity or bonding network in the PDB-format text. This conceptual aspect suggests that incorporating alternative coordinate systems or representations that explicitly capture atomic bonding network and chemical connectivity—such as graph-based, network-based, or spherical coordinate systems tied to atomic bonding patterns—could complement or enhance protein structure description, feature extraction and prediction beyond what is currently achievable with the traditional CCS approach.

**Fig 1.**
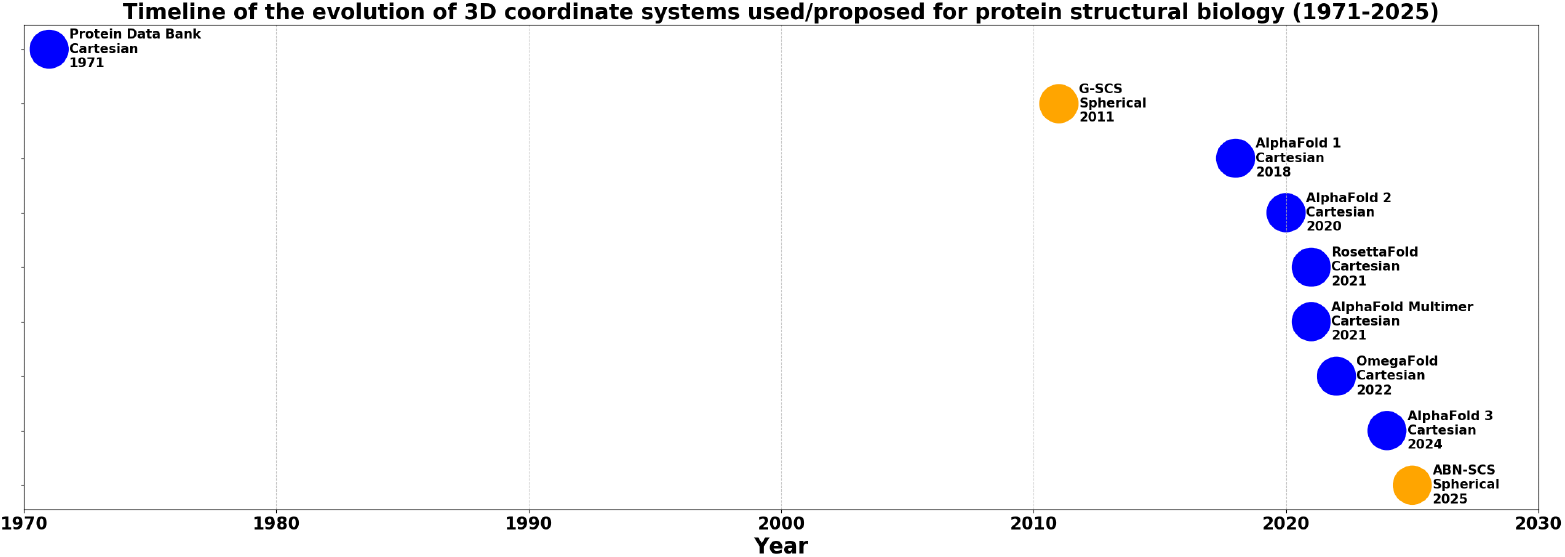
Over half a century burial of *ρ, θ, ϕ* in PDB [3, 5]. In this figure, blue and orange solid circles represent Cartesian and spherical approaches for protein structure description, respectively, with **Local SCS** (the orange solid circle at the bottom) representing the ABN-SCS framework described in this manuscript, and **Global SCS** (the orange solid circle at the up side) representing a global spherical coordinate system, which anchors atomic coordinates to an imaginary geometric centroid [14, 15].

For example, Cartesian coordinate system and spherical coordinate system (SCS)—[*ρ, θ, ϕ*]—are mathematically interconvertible [18], underscoring the technical and geometric feasibility of specifying atomic positions using SCS—[*ρ, θ, ϕ*], in addition to the default [*x, y, z*] approach since 1971 [1]. In 2011, for instance, it was proposed for the first time that protein structure be represented in spherical coordinates [14, 15], where a global spherical coordinate system (G-SCS, Figure 1) was proposed and used for the separation of the protein outer layer from its inner core, and for the identification of protrusions and invaginations on the protein surface [14, 15]. By definition, G-SCS (Figure 1) requires three parameters (*R, θ* and *ϕ*), and takes the geometric centroid (an imaginary point) of the protein structure as origin ([0, 0, 0]), where *R* is used to signify the distance between the imaginary centroid and any atom of the protein [14, 15].

### A Cartesian PDB for protein structure prediction

As of August 29, 2025, ~ 241,055 biomolecular structures have been deposited in PDB [27], the majority of them resolved by experimental techniques such as X-ray crystallography [9, 12], NMR spectroscopy [10, 12], cryo-electron microscopy [11, 12, 28], etc [12, 29]. This number, however, represents only a minute fraction of the billions of known protein sequences, highlighting a fundamental gap between sequence discovery and structural characterization [30]. In recent years, advances in computational methodologies—particularly deep learning—have addressed this disparity, enabling accurate large-scale protein structure prediction [31, 32]. Notably, AlphaFold [33] and RoseTTAFold [31] (Figure 1) have leveraged neural networks to replace traditional energy models and sampling procedures, yielding substantial gains in predictive accuracy [23, 24, 34–37]. Of further interest, geometric feature extraction is one common cornerstone for both experimental structure measurement and protein structure prediction, where experimentally measured raw data or computationally extracted structural features are transformed into geometric descriptors that underpin the construction of the final structural model or NMR ensemble of the molecule of interest. In other words, to neuralized protein structure prediction (e.g., by AF2 [33]), protein structural feature extraction is similar to what experimental measurements of chemical shifts or X-ray diffraction patterns are to protein structure determination using NMR spectroscopy or X-ray crystallography, respectively.

As the primary source of training and validation data, PDB underpins the performance of most protein structure prediction tools [23, 31–33, 38]. For example, AlphaFold2 (AF2) both explicitly and implicitly extracts diverse geometric features (see supplementary **S2.pdf**) from Cartesian-format PDB texts, including atomic coordinates, masks for valid atoms, amino acid encodings, etc. Subsequently, AF2 computes backbone atomic positions (N, C_*α*_, C, O), torsion angles (*ϕ, ψ, ω*), masks for valid residue pairs, etc. These structural template-derived geometric features are supplemented by implicit geometric data generated by the **Evoformer** module during inference, such as inter-residue Euclidean distance maps from predicted C_*β*_ coordinates, backbone torsions angles (*ϕ, ψ, ω*), side-chain torsion *χ* angles and pairwise orientation vectors. Afterwards, AF2 constructs residue-specific rigid-body frames and frame-to-frame transformations to represent local coordinate systems, while geometric validity masks exclude physically implausible conformations, and attention biases based on distance and orientation restraints guide generations of biologically plausible 3D coordinates in the format of [*x, y, z*] [39, 40]. Of note here, by definition, AF2’s backbone torsions (*ϕ, ψ*) differ from the spherical coordinates [*ρ, θ, ϕ*] as described in the Graphical Abstract (**grafic.pdf**) and in [41].

Collectively, these diverse geometric descriptors are critical ingredients for AF2’s accurate structural predictions. While this Cartesian-based prediction pipeline has proven transformative in its performance [33], geometrically, it encodes one viewpoint of protein 3D structure, representing one side of the coin (Ref: **Graphical abstract**). As mentioned above, Cartesian coordinate system and spherical coordinate system (SCS) are mathematically interconvertible [18]. Hence, this article proposes an atomic bonding network–based spherical coordinate system (ABN-SCS) for protein structure representation and feature extraction.

## Motivation

Protein structural modeling has evolved from template-based methods [42–44] to molecular dynamics and energy minimization techniques [45]. The recent emergence of AlphaFold has revolutionized the field, employing deep learning to achieve remarkable accuracy in structure prediction [20–22, 33, 46, 47]. Despite these advances, further improvements in accuracy and efficiency remain an open challenge, raising questions about the next phase of development in this field. Therefore, this article introduces an atomic bonding network-based spherical coordinate system (ABN-SCS) [16–18], which describes protein structures using [*ρ, θ, ϕ*] instead of Cartesian coordinates [*x, y, z*]. Rooted in the intrinsic geometry of atomic bonding networks, ABN-SCS provides an alternative framework for structural representation. Using Caenopore-5 as a case study [48–50], this work demonstrates the application of ABN-SCS in redefining protein structures and extracting spherical structural features. Moreover, with these spherical features, this article puts forward an ABN-SCS framework to complement existing Cartesian-based descriptors employed in current protein structure prediction tools such as AlphaFold (Figure 1) [33] and RoseTTAFold (Figure 1) [31], for the continued improvement of their performances.

## Materials and Methods

### Two geometrically inter-convertabile coordinate systems

As mentioned above, CCS and spherical coordinate system (SCS, [*ρ, θ, ϕ*]) are interconvertible, both applicable in the specification of atomic positions [17, 18, 51].

By transforming the Cartesian coordinates into spherical coordinates (Equations 1, 2 and 3), this ABN-SCS approach decouples the spatial information of atoms into radial distance (*ρ*), polar angle (*θ*), and azimuthal angle (*ϕ*), where *ρ* is defined as the equilibrium inter-atomic bond length (a physical constant), i.e., the inter-nuclear distance at which the system energy minimum occurs [52, 53],

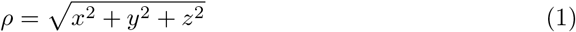

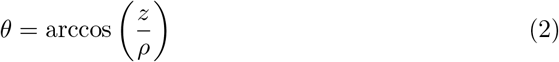

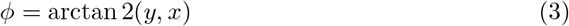

where:

- *ρ* is the radial distance from the origin to the point, Figure 2.
- *θ* is the polar angle (angle from the positive *z*-axis, Figure 2).
- *ϕ* is the azimuthal angle (angle from the positive *x*-axis in the *xy*-plane, Figure 2).

Here in this manuscript, the ABN-SCS framework is defined as a chemically grounded geometric framework for the representation of protein structures by integrating covalent atomic bonding information with spherical coordinates [*ρ, θ, ϕ*]. Its core principles and methodology are outlined as follows:

**Fig 2.**
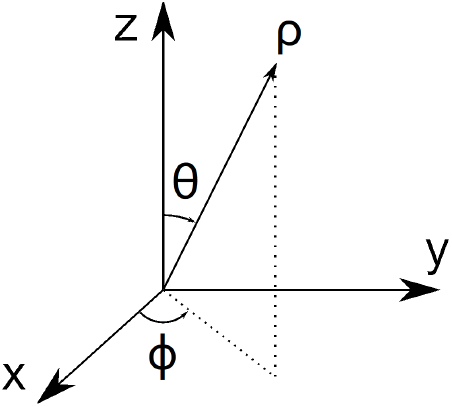
Geometrically, Cartesian and spherical coordinate systems are like two sides of one coin (Ref: **Graphical abstract, grafic.pdf**), and are both applicable in the specifications of atomic positions for protein structure description.

1. **foundation: inter-atomic bonding network (ABN)** ABN-SCS anchors the coordinate system to the covalent bonding network of the protein rather than an arbitrary geometric centroid [14, 15]. Each atom’s position is defined relative to its bonded neighbor, ensuring that chemical connectivity is inherently encoded in the ABN-SCS framework.
2. **spherical coordinate transformation** In ABN-SCS, atomic positions are expressed in spherical coordinates [*ρ, θ, ϕ*], which can be reversibly converted back to [*x, y, z*] (Figure 2).
3. **equilibrium bond lengths as radial distances** In ABN-SCS, the radial coordinate *ρ* (Figure 2) is defined as the equilibrium inter-atomic bond length, i.e., the inter-nuclear distance at which the system energy minimum occurs [52, 53]. This ensures that the ABN-SCS framework captures chemically relevant spacing between atoms, and that the value of *ρ* is physically and structurally restrained [41], making it a more physically grounded paradigm compared to the default Cartesian approach.
4. **chemical significance of spherical coordinates** [*θ, ϕ*] As two spherical angles, *θ* and *ϕ* capture local spherical bond-level geometric features, enabling an ABN-SCS perspective for the extraction of spherical protein structure features, in addition to Cartesian features (see supplementary **supps.pdf**) as used by computational pipelines such as AF2 [33].
5. **local SCS frame definition for protein structure feature extraction** For each covalently bonded atom pair, a local spherical coordinate frame is able to be established based on its covalently bonded neighbors, providing a consistent reference for measuring angles and distances across the protein, where global rotations and translations does not affect protein structure feature extraction.

### An ABN-SCS description of a model pentapeptide’s structure

As proteins are essential building blocks of life, amino acids are building blocks of protein. Thus, for an ABN-SCS description of protein structure, it is necessary to first use the ABN-SCS framework in the descriptions of the chemical connectivity (atomic bonding network, ABN) of twenty stand-alone natural amino acids, for which all details are included in supplementary file **S1.pdf**. With supplementary file **S1.pdf** in place, next step is to use the ABN-SCS framework to define the formation of a peptide bond, which is described in section 1.4 of supplementary file **S2.pdf**. Afterwards, the entire inter-atomic covalent bonding network of the backbone of the short peptide is also defined sequentially as in section 1.4 of supplementary file **S2.pdf**.

With this generalizable ABN-SCS framework, the complete covalent chemical connectivity of any protein structure—including both backbone and side-chain atoms—can be fully and reversibly mapped into spherical coordinates (*ρ, θ, ϕ*), thereby embedding intrinsic chemical topology directly into protein structure description with the ABN-SCS framework described in this manuscript.

### An ABN-SCS description of protein backbone and peptide bond structures: Caenopore-5 as an example

To illustrate the redefinition of protein backbone structure with *ρ, θ* and *ϕ*, this article takes Caenopore-5 (Figure 3) as an example, which adopts a helical conformation, forming oligomeric pores upon interaction with lipid bilayers, with five helices stabilized by three disulfide bridges (Figure 4).

**Fig 3.**
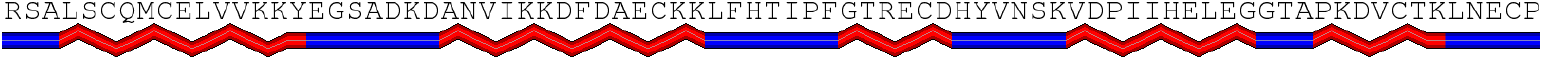
Amino acid sequence and secondary structure of Caenopore-5 (81 amino acids) [48–50]. This figure was prepared by the PSIPRED server [54].

**Fig 4.**
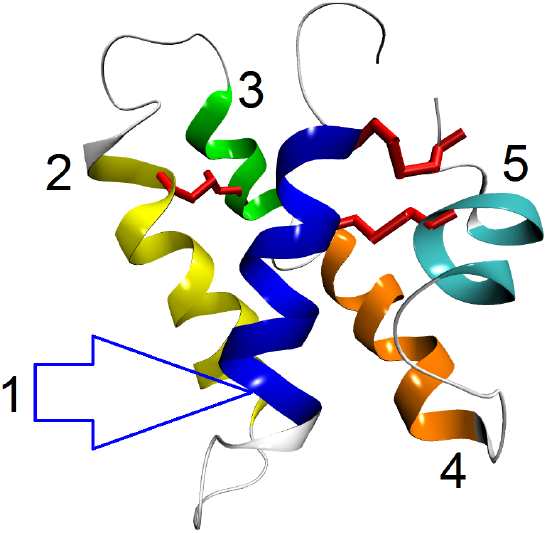
Three-dimensional structure of Caenopore-5 (PDB ID: 2JSA) stabilized by three disulfide bonds (red sticks) [48–50]. This figure is prepared by VMD [55].

As the structure of Caenopore-5 was determined by liquid-state NMR spectroscopy [48, 49], the structural data deposited in PDB [2] is an NMR ensemble of 15, instead of 1, structural models calculated with restraints from experimental NMR data. As a result, the redefinition of the backbone structure of Caenopore-5 with [*ρ, θ, ϕ*] is for all 15 NMR structural models [48–50]. Specifically, the peptide bond and the backbone structures of Caenopore-5 are described with [*ρ, θ, ϕ*] according to section 1.4 of supplementary file **S2.pdf**, where an in-house python script (supplementary file **S3.pdf**) was used to extract covalent bond-level spherical features from the NMR ensemble of Caenopore-5 [48, 49].

## Results

This article used Caenopore-5 [48–50] as an example to define an alternative framework, i.e., ABN-SCS, for protein structure description with [*ρ, θ, ϕ*] [56] and the extraction of spherical structural features in addition to those Cartesian structural features (sections 3.2 and 3.3 of supplementary file **S2.pdf**) used by current protein structure prediction tools [57–63]. In other words, this ABN-SCS framework offers an alternative approach for extracting essential structural features from protein backbones by transforming Cartesian coordinates (*x, y*, and *z*) into spherical coordinates *ρ, θ* and *ϕ*. Using Caenopore-5 [48–50] as an example here, two sets of spherical parameters were extracted from its NMR ensemble (PDB ID: 2JSA) and summarized as two long tables (sections 2.5 and 2.6 of supplementary file **S2.pdf**). To sum up, for its backbone structure [48–50], a 3 *×* 477 matrix of spherical parameters were extracted and visualized in Figures 4, 5 and 6 in supplementary file **S2.pdf** for *ρ, θ* and *ϕ*, respectively; For its peptide bond structure [48–50], a 3*×* 80 matrix of spherical parameters were extracted and visualized in Figures 7, 8 and 9 in supplementary file **S2.pdf** for *ρ, θ* and *ϕ*, respectively.

**Fig 5.**
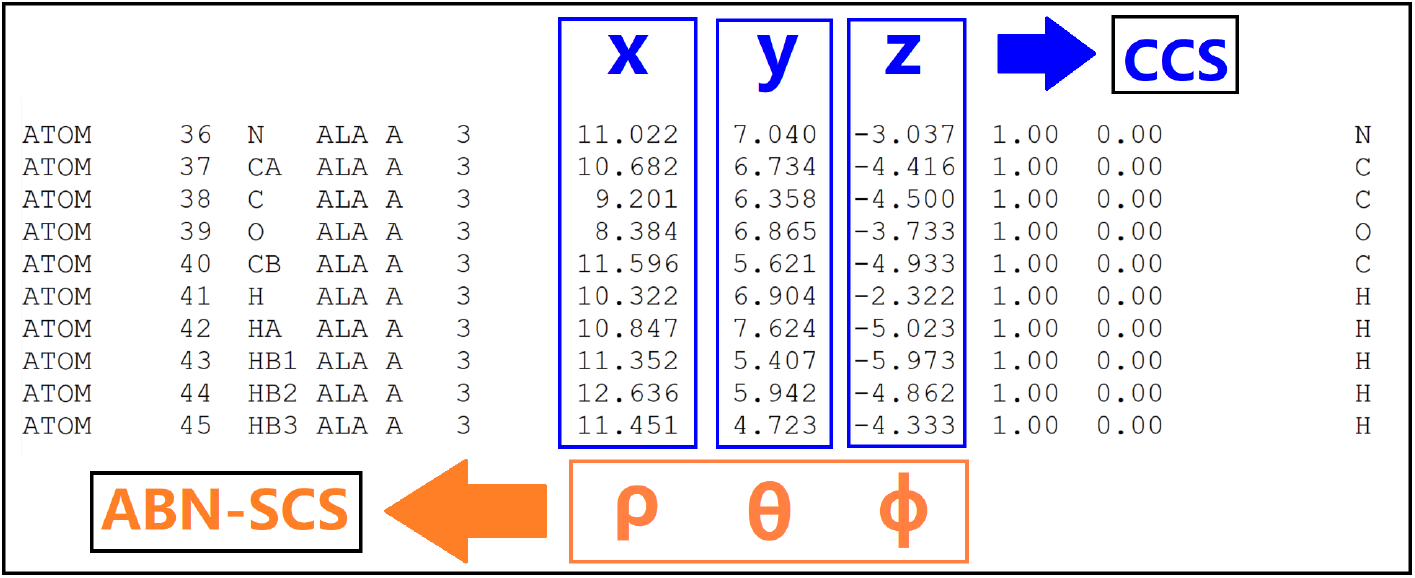
Towards a spherical Protein Data Bank in addition to its Cartesian counterpart [1, 2]. This figure represents a simplified example of the PDB-format ATOM record with Cartesian coordinates ([*x, y, z*]) replaceable by ABN-SCS coordinates [*ρ, θ, ϕ*].

Specifically, an in-house python script (**S3.pdf**) was used to extract spherical features from the 15 NMR structural models of Caenopore-5. While these spherical features (*ρ, θ* and *ϕ*) are tabulated in two long tables in sections 2.5 and 2.6 of supplementary file **S2.pdf**, six figures were also prepared to visualize the distributions of these spherical features. For instance, it can be seen from Figure 4 in **S2.pdf** that the distribution pattern of the lengths (*ρ*) of 477 covalent bonds appears rather stable for the NMR ensemble [48–50], oscillating approximately between 1.0-1.5 Å due to the existence of various types of covalent bonds in Caenopore-5’s backbone. Thus, another figure was prepared to visualize the distribution of the lengths of only one type of covalent bond, i.e., the peptide bond of Caenopore-5, where Figure 13 in **S2.pdf** illustrates clearly the distribution patterns of the lengths (*ρ*) of 80 peptide bonds of Caenopore-5 [48–50], and that the lengths (*ρ*) of the 80 peptide bonds are almost in the shape of a straight line (Figures 7 and 13 in **S2.pdf**). Quantitatively, nonetheless, there is still slight variations of the values of *ρ* to a certain degree across its 81 amino acid residues, as shown in Figures 7 and 13 in **S2.pdf**, where the lengths of the 80 peptide bonds possess an average at 1.326 Å and a standard deviation at 0.009 Å (Table 5 in supplementary file **S2.pdf**).

Apart from Figures 4 and 7 in **S2.pdf**, the spherical structural features of Caenopore-5 extracted by the python script (**S3.pdf**) were also visualized in another four figures, namely, Figures 5, 6, 8 and 9 in **S2.pdf**. For the four supplementary figures, the underlying raw data are included in two long tables in sections 2.5 and 2.6 of supplementary file **S2.pdf**. As a matter of fact, it is the raw data of these four figures that are able to act as two sets of ABN-SCS-based spherical fingerprints or descriptors (two sets of angles of *θ* and *ϕ*) as a function of the sequence of Caenopore-5, because with the ABN-SCS framework here, the 3D backbone structure Caenopore-5 is reversibly converted from [*x, y, z*] to [*ρ, θ, ϕ*] with the in-house python script (**S3.pdf**). Also, it is the raw data of these four figures that are different from the current Cartesian structural features (sections 3.2 and 3.3 of supplementary file **S2.pdf**) used by current protein structure prediction tools [57–60], offering addtional alternative protein-specific structural features to be plugged into deep learning algorithms for protein structure prediction with improved performance in future.

Moreover, for Figures 5, 6, 8 and 9 in **S2.pdf**, a straightforward naked-eye inspection reveals that the value of *θ* oscillates approximately between 0° and 180°, while the value of *ϕ* oscillates approximately between −180° and 180°. Geometrically, a *θ* ranging from 0° to 180° and a *ϕ* ranging from −180° to 180° are both necessary and sufficient to cover the entire 3D space as defined by Cartesian coordinates [*x, y, z*], while a *θ* ranging from −180° to 180° and a *ϕ* ranging from −180° to 180° are sufficient yet unnecessary to cover the entire 3D space as defined by [*x, y, z*]. Thus, the fact that the value of *θ* oscillates approximately between 0° and 180°, while the value of *ϕ* oscillates approximately between −180° and 180° supports the robustness of the in-house python script (**S3.pdf**) for the conversion from [*x, y, z*] to [*ρ, θ, ϕ*], including in particular the two python functions (**conv_1** and **conv_2** in **S3.pdf**) for the reversible conversion between Cartesian and spherical coordinate systems.

Further details of the results and analysis are included in supplementary result section in **S2.pdf**.

## Conclusion

To sum up, this article used Caenopore-5 [48–50] as an example for protein 3D structure description with a spherical coordinate system based on the intrinsic inter-atomic bonding (ABN-SCS) network of protein. In addition to Caenopore-5, this ABN-SCS-based description is also applicable for other proteins in PDB [2], regardless of whether X-ray diffraction, NMR spectroscopy or Cryo-EM is the tool for experimental structure determination, since for all experimental biomolecular structures, the Protein Data Bank (PDB) requires a standardized data format before deposition, and that the two coordinate systems are geometrically interconvertible. Moreover, this manuscript provides a supplementary file **S3**, a python script for ABN-SCS-based extraction of spherical geometric features, which is in fact an implementation of Equations 1, 2 and 3 (Figure 2).

Geometrically, this spherical ABN-SCS framework is an uncharted territory, different from the default Cartesian approach since 1971 [1–4], in the sense that ABN-SCS (*ρ, θ* and *ϕ*) not only is a new language for protein structure description [13–18] and understanding [6–8], but also paves a new way for the extraction of protein structure features beyond conventional Cartesian feature extraction (e.g., inter-atomic and inter-residue distances) by computational tools such as AlphaFold [33]. Essentially, this spherical ABN-SCS approach is more of a geometric algorithm, through which spherical structural features (*ρ, θ* and *ϕ*) of protein can be extracted from the entire collection of experimentally determined protein structures inside PDB [1–4]. Therefore, this article here proposes a spherical Protein Data Bank (Figure 5) on top of its 54-year-old Cartesian counterpart [1–4], such that additional spherical structural features are extracted and can be incorporated into deep learning algorithms for continued improvement of the performance of protein structure prediction tools [19–22].

### Discussion: towards a spherical PDB for protein structure feature extraction

In recent years, the application of artificial intelligence (AI) has expanded rapidly across many scientific disciplines [64], including neural network–based protein structure prediction by tools such as AF2 [33]. Nevertheless, the adoption of AI in science is not without limitations; epistemological concerns have been raised regarding its widespread use [64]. Take astronomy as an analogy [64]. The model of the Universe with Earth at its centre was extremely accurate at predicting planetary motions, because of tricks such as ‘epicycles’ — the assumption that planets move in circles whose centres revolve around Earth along a larger circular path. This analogy suggests that extremely accurate predictions do not necessarily equate to genuine scientific understanding in an unbiased manner, which is critical for continued improvement of the performance of technologies and computational tools (e.g., AF2) built upon the way protein structure is conceptualized and described in the central global repository [1]. Today, AI excels at producing the equivalent of ‘epicycles’ [64]. All else being equal, being able to squeeze more predictive juice [65] out of flawed theories or inadequate paradigms will help them to stick around for longer, impeding true scientific progress [64] and continued improvement of the performance of protein structure prediction pipelines such as AF2 [33]. To this end, for protein structure description with [*x, y, z*], CCS is an inadequate paradigm, in the sense that CCS treats each atom as an independent point in three-dimensional space, representing a protein as a collection of discrete (*x, y, z*) points without explicitly encoding its underlying covalent bonding network or chemical connectivity in the PDB text format. In other words, Cartesian coordinates have been the default “language” of protein structure description since 1971 (Figure 1) [1, 2], reflecting a geometric rather than a chemically and physically grounded perspective of how protein structure is conceptualized and described in the central global repository [2].

Yet, this by no means is to suggest that CCS is wrong or flawed. As one of the foundational paradigms of structural biology, CCS has been not only geometrically correct, but also proven indispensable and immensely useful in its real world applications. Nevertheless, it is conceptually inadequate, as it omits explicit representation of chemical connectivity within protein structure. From a mathematical standpoint, CCS and spherical coordinate systems (SCS; *ρ, θ, ϕ*) are fully interconvertible [18]. Thus, specifying atomic positions using spherical coordinates alongside the conventional (*x, y, z*) representation has been both technically and geometrically feasible and enforceable. Building on this geometric equivalence (Ref: **Graphical Abstract**), this manuscript puts forward an ABN-SCS framework for protein structure representation and feature extraction. Unlike earlier global SCS approaches (e.g., the first orange circle at the top of Figure 1) [14, 15], which anchor coordinates to an imaginary geometric centroid, ABN-SCS is intrinsically rooted in the covalent bonding network of the target protein. This allows the derivation of chemically meaningful geometric descriptors that can augment conventional Cartesian-based features used by computational pipelines such as AF2 [33].

Taken together, this manuscript advocates an alternative structural biology perspective (Ref **Graphical Abstract**), and with this manuscript, I argue that the time is now ripe for the computational structural biology and bioinformatics community to take a look at the other side of the coin (Ref: **Graphical abstract**), and also for the trapped spherical features [*ρ, θ*, and *ϕ*] to be released from PDB and integrated into algorithms such as AF2 towards protein structure prediction with improved performance, as this dual-coordinate system strategy is not just geometrically and technically feasible and enforceable, but also paves a way to further narrow the gap between structure predictions and the physical–chemical reality not just of proteins, but also of biomolecular architectures in general.

## Supporting information

Graphical Abstract

## Declaration of generative AI and AI-assisted technologies in the writing process

During the preparation of this work, the author used OpenAI’s ChatGPT in order to improve the readability of the manuscript. After using this tool, the author reviewed and edited the content as needed and takes full responsibility for the content of the publication.

## Supporting information

**S1 File. S1.pdf** ABN-SCS-based descriptions of the chemical connectivities of twenty stand-alone natural amino acids.

**S2 File. S2.pdf** Supplementary Materials and Methods, Results and Discussions.

**S3 File. S3.pdf** A python script for ABN-SCS-based extraction of spherical geometric features, i.e., [*ρ, θ, ϕ*].

