## Supplementary material for "Protein Structure Description with *ρ, θ* and *ϕ*: A Case Study with Caenopore-5": Graphical Abstract

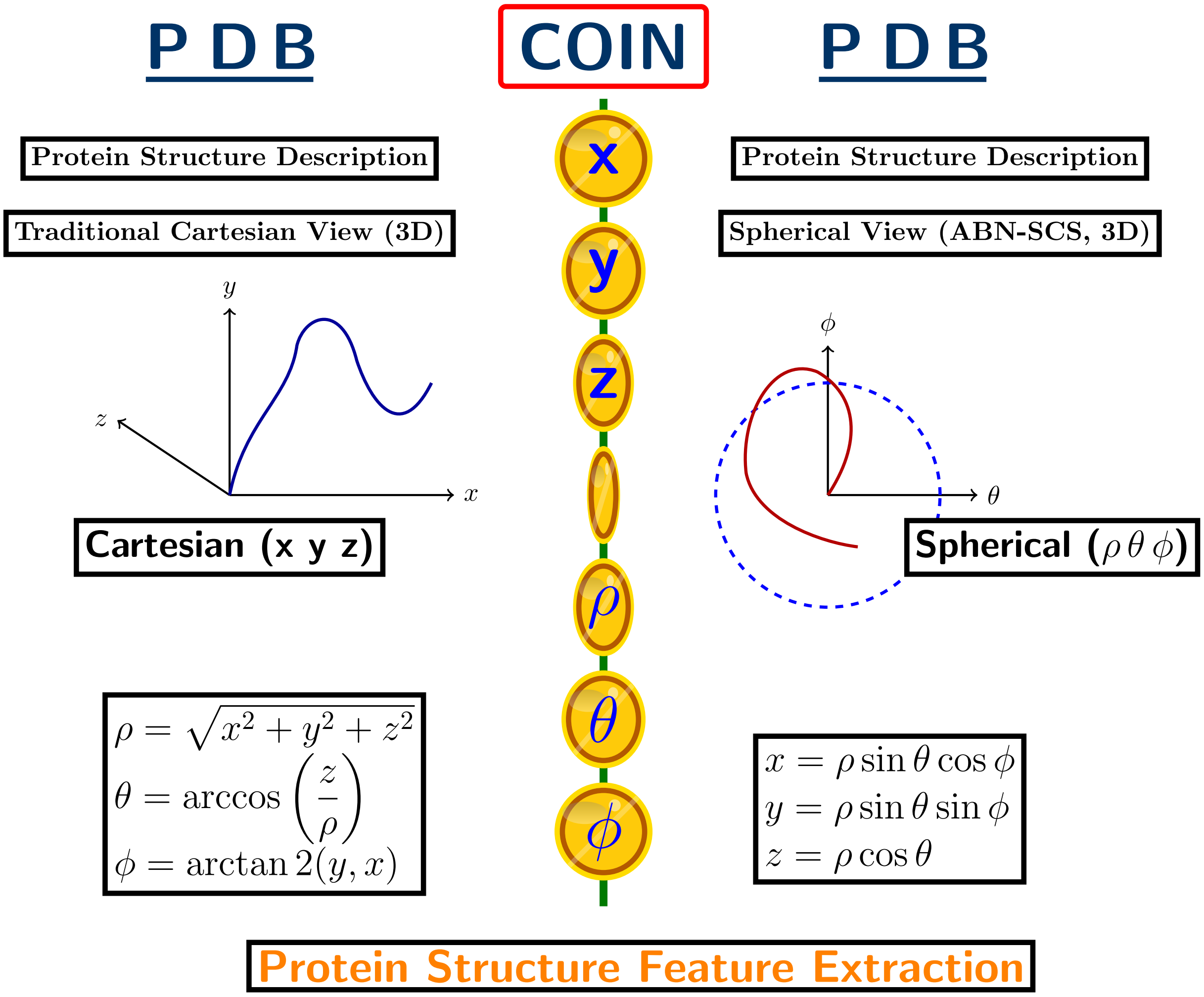



**Graphical Abstract.** Since 1971, experimentally measured protein structures have been deposited in the Protein Data Bank (PDB) with atomic positions expressed in Cartesian coordinates $[x, y, z]$. In light of the geometric inter-convertibility of Cartesian and spherical (SCS, $[\rho,\ \theta,\ \phi]$) coordinate systems, this figure presents a conceptual shift towards a spherical view of protein structure. As an alternative to the Cartesian default since 1971, the ABN-SCS framework described in this manuscript offers a radial, angular description of protein structure with $[\rho,\ \theta,\ \phi]$, where $\rho$ is defined as the equilibrium inter-atomic bond length (a physical constant), i.e., the inter-nuclear distance at which the system energy minimum occurs. In the center of this figure are seven flipping golden coins, which constitutes a metaphor for geometric inter-convertibility of Cartesian and spherical coordinate systems, advocating for the extraction of $[\rho,\ \theta,\ \phi]$ rooted in the atomic bonding network (ABN) within protein structure, and also for the integration of spherical structural features $[\rho,\ \theta,\ \phi]$ into protein structure prediction algorithms like AlphaFold.
