## Supplementary file 1 for "Protein Structure Description with *ρ, θ* and *ϕ*: A Case Study with Caenopore-5"

### Protein Structure Description with ρ, θ and ϕ: Supplementary Information

Wei Li *
Independent Researcher
Room 902, Building No. 41, Fourth Community, Yuhongjiayuan Village
No. 1168, Antong Road, Chenjia Town, Chongming Island
Shanghai, People's Republic of China

August 29, 2025

*

### Contents


1 An ABN-SCS Description with ρ, θ and ϕ for alanine 3

2 An ABN-SCS Description with ρ, θ and ϕ for arginine 5

3 An ABN-SCS Description with ρ, θ and ϕ for asparagine 9

4 An ABN-SCS Description with ρ, θ and ϕ for aspartate 12

5 An ABN-SCS Description with ρ, θ and ϕ for cysteine 15

6 An ABN-SCS Description with ρ, θ and ϕ for glutamate 17

7 An ABN-SCS Description with ρ, θ and ϕ for glutamine 20

8 An ABN-SCS Description with ρ, θ and ϕ for glycine 23

9 An ABN-SCS Description with ρ, θ and ϕ for histidine 25

10 An ABN-SCS Description with ρ, θ and ϕ for isoleucine 29

11 An ABN-SCS Description with ρ, θ and ϕ for leucine 33

12 An ABN-SCS Description with ρ, θ and ϕ for lysine 37

13 An ABN-SCS Description with ρ, θ and ϕ for methionine 41

14 An ABN-SCS Description with ρ, θ and ϕ for phenylalanine 44

15 An ABN-SCS Description with ρ, θ and ϕ for proline 48

16 An ABN-SCS Description with ρ, θ and ϕ for serine 51

17 An ABN-SCS Description with ρ, θ and ϕ for threonine 53

18 An ABN-SCS Description with ρ, θ and ϕ for tryptophan 56

19 An ABN-SCS Description with ρ, θ and ϕ for tyrosine 61

20 An ABN-SCS Description with ρ, θ and ϕ for valine 65

### 1 An ABN-SCS Description with ρ, θ and ϕ for alanine

#### Atoms in Alanine (Ala, A)

1. **Backbone Atoms:**
   - N (Amine nitrogen)
   - $H_1$ (First hydrogen attached to N)
   - $H_2$ (Second hydrogen attached to N)
   - $C\alpha$ (Alpha carbon)
   - $H\alpha$ (Hydrogen attached to $C\alpha$)
   - C (Carbonyl carbon)
   - O (Carbonyl oxygen)
   - $O^-$ (Carbonyl oxygen in the ionized state)
2. **Side Chain Atoms:**
   - $C\beta$ (Beta carbon)
   - $H\beta_1$ (First hydrogen attached to $C\beta$)
   - $H\beta_2$ (Second hydrogen attached to $C\beta$)
   - $H\beta_3$ (Third hydrogen attached to $C\beta$)

#### An ABN-SCS definition of the atomic bonding network of Ala

1. **N to C**$\alpha$:

$$\begin{aligned} \rho_{\mathrm{N-C}\alpha} &: \text{Distance} \\ \theta_{\mathrm{N-C}\alpha} &: \text{Polar angle} \\ \phi_{\mathrm{N-C}\alpha} &: \text{Azimuthal angle} \end{aligned}$$

2. **C**$\alpha$ **to C**$\beta$:

$$\begin{aligned} \rho_{\mathrm{C}\alpha-\mathrm{C}\beta} &: \text{Distance} \\ \theta_{\mathrm{C}\alpha-\mathrm{C}\beta} &: \text{Polar angle} \\ \phi_{\mathrm{C}\alpha-\mathrm{C}\beta} &: \text{Azimuthal angle} \end{aligned}$$

3. **C**$\alpha$ **to C**:

$$\begin{aligned} \rho_{\mathrm{C}\alpha-\mathrm{C}} &: \text{Distance} \\ \theta_{\mathrm{C}\alpha-\mathrm{C}} &: \text{Polar angle} \\ \phi_{\mathrm{C}\alpha-\mathrm{C}} &: \text{Azimuthal angle} \end{aligned}$$

4. **C to O (Carbonyl Oxygen)**:

$$\begin{aligned} \rho_{\mathrm{C-O}} &: \text{Distance} \\ \theta_{\mathrm{C-O}} &: \text{Polar angle} \\ \phi_{\mathrm{C-O}} &: \text{Azimuthal angle} \end{aligned}$$

5. **C to $\mathbf{O_H}$ (Hydroxyl Oxygen)**:

$$\begin{aligned} \rho_{C-OH} &: \text{Distance} \\ \theta_{C-OH} &: \text{Polar angle} \\ \phi_{C-OH} &: \text{Azimuthal angle} \end{aligned}$$

6. **O to $\mathbf{H_{OH}}$ (Hydroxyl Hydrogen)**:

$$\begin{aligned} \rho_{O-HOH} &: \text{Distance} \\ \theta_{O-HOH} &: \text{Polar angle} \\ \phi_{O-HOH} &: \text{Azimuthal angle} \end{aligned}$$

7. **N to $\mathbf{H}_1$**:

$$\begin{aligned} \rho_{N-H_1} &: \text{Distance} \\ \theta_{N-H_1} &: \text{Polar angle} \\ \phi_{N-H_1} &: \text{Azimuthal angle} \end{aligned}$$

8. **N to $\mathbf{H}_2$**:

$$\begin{aligned} \rho_{N-H_2} &: \text{Distance} \\ \theta_{N-H_2} &: \text{Polar angle} \\ \phi_{N-H_2} &: \text{Azimuthal angle} \end{aligned}$$

9. **C$\alpha$ to H$\alpha$**:

$$\begin{aligned} \rho_{C\alpha-H\alpha} &: \text{Distance} \\ \theta_{C\alpha-H\alpha} &: \text{Polar angle} \\ \phi_{C\alpha-H\alpha} &: \text{Azimuthal angle} \end{aligned}$$

10. **C$\beta$ to H$\beta_1$**:

$$\begin{aligned} \rho_{C\beta-H\beta_1} &: \text{Distance} \\ \theta_{C\beta-H\beta_1} &: \text{Polar angle} \\ \phi_{C\beta-H\beta_1} &: \text{Azimuthal angle} \end{aligned}$$

11. **C$\beta$ to H$\beta_2$**:

$$\begin{aligned} \rho_{C\beta-H\beta_2} &: \text{Distance} \\ \theta_{C\beta-H\beta_2} &: \text{Polar angle} \\ \phi_{C\beta-H\beta_2} &: \text{Azimuthal angle} \end{aligned}$$

12. **C$\beta$ to H$\beta_3$**:

$$\begin{aligned} \rho_{C\beta-H\beta_3} &: \text{Distance} \\ \theta_{C\beta-H\beta_3} &: \text{Polar angle} \\ \phi_{C\beta-H\beta_3} &: \text{Azimuthal angle} \end{aligned}$$

### 2 An ABN-SCS Description with ρ, θ and ϕ for arginine

#### Atoms in Arginine (Arg, R)

1. **Backbone Atoms:**
   - N (Amine nitrogen)
   - H (Hydrogen attached to N)
   - C$\alpha$ (Alpha carbon)
   - H$\alpha$ (Hydrogen attached to C$\alpha$)
   - C (Carbonyl carbon)
   - O (Carbonyl oxygen)
   - $O^-$ (Carbonyl oxygen in the ionized state)
2. **Side Chain Atoms:**
   - C$\beta$ (Beta carbon)
   - H$\beta_1$ (First hydrogen attached to C$\beta$)
   - H$\beta_2$ (Second hydrogen attached to C$\beta$)
   - C$\gamma$ (Gamma carbon)
   - H$\gamma_1$ (First hydrogen attached to C$\gamma$)
   - H$\gamma_2$ (Second hydrogen attached to C$\gamma$)
   - C$\delta$ (Delta carbon)
   - H$\delta_1$ (First hydrogen attached to C$\delta$)
   - H$\delta_2$ (Second hydrogen attached to C$\delta$)
   - N$\epsilon$ (Epsilon nitrogen)
   - H$\epsilon$ (Hydrogen attached to N$\epsilon$)
   - C$\zeta$ (Zeta carbon)
   - N$\eta_1$ (First eta nitrogen)
   - H$\eta_1$ (First hydrogen attached to N$\eta_1$)
   - H$\eta_1$ (Second hydrogen attached to N$\eta_1$)
   - N$\eta_2$ (Second eta nitrogen)
   - H$\eta_2$ (First hydrogen attached to N$\eta_2$)
   - H$\eta_2$ (Second hydrogen attached to N$\eta_2$)

#### An ABN-SCS definition of the atomic bonding network of Arg

1. **N to C$\alpha$**:

$$\rho_{\mathrm{N-C}\alpha} : \text{Distance}$$
$$\theta_{\mathrm{N-C}\alpha} : \text{Polar angle}$$
$$\phi_{\mathrm{N-C}\alpha} : \text{Azimuthal angle}$$

2. **C$\alpha$ to C$\beta$**:

$$\rho_{C\alpha-C\beta} : \text{Distance}$$
$$\theta_{C\alpha-C\beta} : \text{Polar angle}$$
$$\phi_{C\alpha-C\beta} : \text{Azimuthal angle}$$

3. **C$\alpha$ to C**:

$$\rho_{C\alpha-C} : \text{Distance}$$
$$\theta_{C\alpha-C} : \text{Polar angle}$$
$$\phi_{C\alpha-C} : \text{Azimuthal angle}$$

4. **C to O (Carbonyl Oxygen)**:

$$\rho_{C-O} : \text{Distance}$$
$$\theta_{C-O} : \text{Polar angle}$$
$$\phi_{C-O} : \text{Azimuthal angle}$$

5. **C to $O_H$ (Hydroxyl Oxygen)**:

$$\rho_{C-OH} : \text{Distance}$$
$$\theta_{C-OH} : \text{Polar angle}$$
$$\phi_{C-OH} : \text{Azimuthal angle}$$

6. **O to $H_{OH}$ (Hydroxyl Hydrogen)**:

$$\rho_{O-HOH} : \text{Distance}$$
$$\theta_{O-HOH} : \text{Polar angle}$$
$$\phi_{O-HOH} : \text{Azimuthal angle}$$

7. **C$\beta$ to C$\gamma$**:

$$\rho_{C\beta-C\gamma} : \text{Distance}$$
$$\theta_{C\beta-C\gamma} : \text{Polar angle}$$
$$\phi_{C\beta-C\gamma} : \text{Azimuthal angle}$$

8. **C$\gamma$ to C$\delta$**:

$$\rho_{C\gamma-C\delta} : \text{Distance}$$
$$\theta_{C\gamma-C\delta} : \text{Polar angle}$$
$$\phi_{C\gamma-C\delta} : \text{Azimuthal angle}$$

9. **C$\delta$ to N$\epsilon$**:

$$\rho_{C\delta-N\epsilon} : \text{Distance}$$
$$\theta_{C\delta-N\epsilon} : \text{Polar angle}$$
$$\phi_{C\delta-N\epsilon} : \text{Azimuthal angle}$$

10. $\mathbf{N}\epsilon$ **to** $\mathbf{C}\zeta$:

$$\rho_{N\epsilon-C\zeta} : \text{Distance}$$
$$\theta_{N\epsilon-C\zeta} : \text{Polar angle}$$
$$\phi_{N\epsilon-C\zeta} : \text{Azimuthal angle}$$

11. $\mathbf{C}\zeta$ **to** $\mathbf{N}\eta_1$:

$$\rho_{C\zeta-N\eta_1} : \text{Distance}$$
$$\theta_{C\zeta-N\eta_1} : \text{Polar angle}$$
$$\phi_{C\zeta-N\eta_1} : \text{Azimuthal angle}$$

12. $\mathbf{C}\zeta$ **to** $\mathbf{N}\eta_2$:

$$\rho_{C\zeta-N\eta_2} : \text{Distance}$$
$$\theta_{C\zeta-N\eta_2} : \text{Polar angle}$$
$$\phi_{C\zeta-N\eta_2} : \text{Azimuthal angle}$$

13. **N to** $\mathbf{H_{N1}}$:

$$\rho_{N-H_{N1}} : \text{Distance}$$
$$\theta_{N-H_{N1}} : \text{Polar angle}$$
$$\phi_{N-H_{N1}} : \text{Azimuthal angle}$$

14. **N to** $\mathbf{H_{N2}}$:

$$\rho_{N-H_{N2}} : \text{Distance}$$
$$\theta_{N-H_{N2}} : \text{Polar angle}$$
$$\phi_{N-H_{N2}} : \text{Azimuthal angle}$$

15. $\mathbf{C}\alpha$ **to** $\mathbf{H}\alpha$:

$$\rho_{C\alpha-H\alpha} : \text{Distance}$$
$$\theta_{C\alpha-H\alpha} : \text{Polar angle}$$
$$\phi_{C\alpha-H\alpha} : \text{Azimuthal angle}$$

16. $\mathbf{C}\beta$ **to** $\mathbf{H}\beta_1$:

$$\rho_{C\beta-H\beta_1} : \text{Distance}$$
$$\theta_{C\beta-H\beta_1} : \text{Polar angle}$$
$$\phi_{C\beta-H\beta_1} : \text{Azimuthal angle}$$

17. $\mathbf{C}\beta$ **to** $\mathbf{H}\beta_2$:

$$\rho_{C\beta-H\beta_2} : \text{Distance}$$
$$\theta_{C\beta-H\beta_2} : \text{Polar angle}$$
$$\phi_{C\beta-H\beta_2} : \text{Azimuthal angle}$$

18. **C$\gamma$ to H$\gamma_1$**:

$$\rho_{\mathrm{C}\gamma-\mathrm{H}\gamma_1} : \text{Distance}$$
$$\theta_{\mathrm{C}\gamma-\mathrm{H}\gamma_1} : \text{Polar angle}$$
$$\phi_{\mathrm{C}\gamma-\mathrm{H}\gamma_1} : \text{Azimuthal angle}$$

19. **C$\gamma$ to H$\gamma_2$**:

$$\rho_{\mathrm{C}\gamma-\mathrm{H}\gamma_2} : \text{Distance}$$
$$\theta_{\mathrm{C}\gamma-\mathrm{H}\gamma_2} : \text{Polar angle}$$
$$\phi_{\mathrm{C}\gamma-\mathrm{H}\gamma_2} : \text{Azimuthal angle}$$

20. **C$\delta$ to H$\delta_1$**:

$$\rho_{\mathrm{C}\delta-\mathrm{H}\delta_1} : \text{Distance}$$
$$\theta_{\mathrm{C}\delta-\mathrm{H}\delta_1} : \text{Polar angle}$$
$$\phi_{\mathrm{C}\delta-\mathrm{H}\delta_1} : \text{Azimuthal angle}$$

21. **C$\delta$ to H$\delta_2$**:

$$\rho_{\mathrm{C}\delta-\mathrm{H}\delta_2} : \text{Distance}$$
$$\theta_{\mathrm{C}\delta-\mathrm{H}\delta_2} : \text{Polar angle}$$
$$\phi_{\mathrm{C}\delta-\mathrm{H}\delta_2} : \text{Azimuthal angle}$$

$$\rho_{\mathrm{N}\epsilon-\mathrm{H}\epsilon} : \text{Distance}$$
$$\theta_{\mathrm{N}\epsilon-\mathrm{H}\epsilon} : \text{Polar angle}$$
$$\phi_{\mathrm{N}\epsilon-\mathrm{H}\epsilon} : \text{Azimuthal angle}$$

22. **N$\eta_1$ to H$\eta_{1,1}$**:

$$\rho_{\mathrm{N}\eta_1-\mathrm{H}\eta_{1,1}} : \text{Distance}$$
$$\theta_{\mathrm{N}\eta_1-\mathrm{H}\eta_{1,1}} : \text{Polar angle}$$
$$\phi_{\mathrm{N}\eta_1-\mathrm{H}\eta_{1,1}} : \text{Azimuthal angle}$$

23. **N$\eta_1$ to H$\eta_{1,2}$**:

$$\rho_{\mathrm{N}\eta_1-\mathrm{H}\eta_{1,2}} : \text{Distance}$$
$$\theta_{\mathrm{N}\eta_1-\mathrm{H}\eta_{1,2}} : \text{Polar angle}$$
$$\phi_{\mathrm{N}\eta_1-\mathrm{H}\eta_{1,2}} : \text{Azimuthal angle}$$

24. **N$\eta_2$ to H$\eta_{2,1}$**:

$$\rho_{\mathrm{N}\eta_2-\mathrm{H}\eta_{2,1}} : \text{Distance}$$
$$\theta_{\mathrm{N}\eta_2-\mathrm{H}\eta_{2,1}} : \text{Polar angle}$$
$$\phi_{\mathrm{N}\eta_2-\mathrm{H}\eta_{2,1}} : \text{Azimuthal angle}$$

25. **N$\eta_2$ to H$\eta_{2,2}$**:

$$\rho_{\mathrm{N}\eta_2-\mathrm{H}\eta_{2,2}} : \text{Distance}$$
$$\theta_{\mathrm{N}\eta_2-\mathrm{H}\eta_{2,2}} : \text{Polar angle}$$
$$\phi_{\mathrm{N}\eta_2-\mathrm{H}\eta_{2,2}} : \text{Azimuthal angle}$$

### 3 An ABN-SCS Description with $\rho$, $\theta$ and $\phi$ for asparagine

#### Atoms in Asparagine (Asn, N)

1. **Backbone Atoms:**
   - N (Amine nitrogen)
   - H (Hydrogen attached to N)
   - $C\alpha$ (Alpha carbon)
   - $H\alpha$ (Hydrogen attached to $C\alpha$)
   - C (Carbonyl carbon)
   - O (Carbonyl oxygen)
   - $O^-$ (Carbonyl oxygen in the ionized state)
2. **Side Chain Atoms:**
   - $C\beta$ (Beta carbon)
   - $H\beta_1$ (First hydrogen attached to $C\beta$)
   - $H\beta_2$ (Second hydrogen attached to $C\beta$)
   - $C\gamma$ (Gamma carbon)
   - $O\delta$ (Oxygen attached to $C\gamma$)
   - $N\delta$ (Amide nitrogen)
   - $H\delta_1$ (First hydrogen attached to $N\delta$)
   - $H\delta_2$ (Second hydrogen attached to $N\delta$)

#### An ABN-SCS definition of the atomic bonding network of Asn

1. **N to C$\alpha$**:

$$\begin{aligned}\rho_{N-C\alpha} &: \text{Distance}\\ \theta_{N-C\alpha} &: \text{Polar angle}\\ \phi_{N-C\alpha} &: \text{Azimuthal angle}\end{aligned}$$

2. **C$\alpha$ to C**:

$$\begin{aligned}\rho_{C\alpha-C} &: \text{Distance}\\ \theta_{C\alpha-C} &: \text{Polar angle}\\ \phi_{C\alpha-C} &: \text{Azimuthal angle}\end{aligned}$$

3. **C to O (Carbonyl Oxygen)**:

$$\begin{aligned}\rho_{C-O} &: \text{Distance}\\ \theta_{C-O} &: \text{Polar angle}\\ \phi_{C-O} &: \text{Azimuthal angle}\end{aligned}$$

4. **C to $O_H$ (Hydroxyl Oxygen)**:

$$\rho_{\mathrm{C-OH}} : \text{Distance}$$
$$\theta_{\mathrm{C-OH}} : \text{Polar angle}$$
$$\phi_{\mathrm{C-OH}} : \text{Azimuthal angle}$$

5. **O to $H_{OH}$ (Hydroxyl Hydrogen)**:

$$\rho_{\mathrm{O-HOH}} : \text{Distance}$$
$$\theta_{\mathrm{O-HOH}} : \text{Polar angle}$$
$$\phi_{\mathrm{O-HOH}} : \text{Azimuthal angle}$$

6. **C$\alpha$ to C$\beta$**:

$$\rho_{\mathrm{C}\alpha-\mathrm{C}\beta} : \text{Distance}$$
$$\theta_{\mathrm{C}\alpha-\mathrm{C}\beta} : \text{Polar angle}$$
$$\phi_{\mathrm{C}\alpha-\mathrm{C}\beta} : \text{Azimuthal angle}$$

7. **C$\beta$ to C$\gamma$**:

$$\rho_{\mathrm{C}\beta-\mathrm{C}\gamma} : \text{Distance}$$
$$\theta_{\mathrm{C}\beta-\mathrm{C}\gamma} : \text{Polar angle}$$
$$\phi_{\mathrm{C}\beta-\mathrm{C}\gamma} : \text{Azimuthal angle}$$

8. **C$\gamma$ to O$\delta$**:

$$\rho_{\mathrm{C}\gamma-\mathrm{O}\delta} : \text{Distance}$$
$$\theta_{\mathrm{C}\gamma-\mathrm{O}\delta} : \text{Polar angle}$$
$$\phi_{\mathrm{C}\gamma-\mathrm{O}\delta} : \text{Azimuthal angle}$$

9. **C$\gamma$ to N$\delta$**:

$$\rho_{\mathrm{C}\gamma-\mathrm{N}\delta} : \text{Distance}$$
$$\theta_{\mathrm{C}\gamma-\mathrm{N}\delta} : \text{Polar angle}$$
$$\phi_{\mathrm{C}\gamma-\mathrm{N}\delta} : \text{Azimuthal angle}$$

10. **N$\delta$ to H$\delta_1$**:

$$\rho_{\mathrm{N}\delta-\mathrm{H}\delta_1} : \text{Distance}$$
$$\theta_{\mathrm{N}\delta-\mathrm{H}\delta_1} : \text{Polar angle}$$
$$\phi_{\mathrm{N}\delta-\mathrm{H}\delta_1} : \text{Azimuthal angle}$$

11. **N$\delta$ to H$\delta_2$**:

$$\rho_{\mathrm{N}\delta-\mathrm{H}\delta_2} : \text{Distance}$$
$$\theta_{\mathrm{N}\delta-\mathrm{H}\delta_2} : \text{Polar angle}$$
$$\phi_{\mathrm{N}\delta-\mathrm{H}\delta_2} : \text{Azimuthal angle}$$

12. **N to $\mathbf{H_{N1}}$**:

$$\rho_{N-H_{N1}} : \text{Distance}$$
$$\theta_{N-H_{N1}} : \text{Polar angle}$$
$$\phi_{N-H_{N1}} : \text{Azimuthal angle}$$

13. **N to $\mathbf{H_{N2}}$**:

$$\rho_{N-H_{N2}} : \text{Distance}$$
$$\theta_{N-H_{N2}} : \text{Polar angle}$$
$$\phi_{N-H_{N2}} : \text{Azimuthal angle}$$

14. **C$\alpha$ to H$\alpha$**:

$$\rho_{C\alpha-H\alpha} : \text{Distance}$$
$$\theta_{C\alpha-H\alpha} : \text{Polar angle}$$
$$\phi_{C\alpha-H\alpha} : \text{Azimuthal angle}$$

15. **C$\beta$ to H$\beta_1$**:

$$\rho_{C\beta-H\beta_1} : \text{Distance}$$
$$\theta_{C\beta-H\beta_1} : \text{Polar angle}$$
$$\phi_{C\beta-H\beta_1} : \text{Azimuthal angle}$$

16. **C$\beta$ to H$\beta_2$**:

$$\rho_{C\beta-H\beta_2} : \text{Distance}$$
$$\theta_{C\beta-H\beta_2} : \text{Polar angle}$$
$$\phi_{C\beta-H\beta_2} : \text{Azimuthal angle}$$

### 4 An ABN-SCS Description with ρ, θ and ϕ for aspartate

#### Atoms in Aspartic Acid (Asp, D)

1. **Backbone Atoms:**
   - N (Amine nitrogen)
   - H (Hydrogen attached to N)
   - C$\alpha$ (Alpha carbon)
   - H$\alpha$ (Hydrogen attached to C$\alpha$)
   - C (Carbonyl carbon)
   - O (Carbonyl oxygen)
   - $O^-$ (Carbonyl oxygen in the ionized state)
2. **Side Chain Atoms:**
   - C$\beta$ (Beta carbon)
   - H$\beta_1$ (First hydrogen attached to C$\beta$)
   - H$\beta_2$ (Second hydrogen attached to C$\beta$)
   - C$\gamma$ (Gamma carbon)
   - O$\delta_1$ (First delta oxygen, part of the carboxyl group)
   - O$\delta_2^-$ (Second delta oxygen, ionized form in the carboxyl group)
   - H$\delta_2$ (Hydroxyl hydrogen attached to O$\delta_2^-$)

#### An ABN-SCS definition of the atomic bonding network of Asp

1. **N to C$\alpha$**:

$$\rho_{\mathrm{N-C}\alpha} : \text{Distance}$$
$$\theta_{\mathrm{N-C}\alpha} : \text{Polar angle}$$
$$\phi_{\mathrm{N-C}\alpha} : \text{Azimuthal angle}$$

2. **C$\alpha$ to C**:

$$\rho_{\mathrm{C}\alpha-\mathrm{C}} : \text{Distance}$$
$$\theta_{\mathrm{C}\alpha-\mathrm{C}} : \text{Polar angle}$$
$$\phi_{\mathrm{C}\alpha-\mathrm{C}} : \text{Azimuthal angle}$$

3. **C to O (Carbonyl Oxygen)**:

$$\rho_{\mathrm{C-O}} : \text{Distance}$$
$$\theta_{\mathrm{C-O}} : \text{Polar angle}$$
$$\phi_{\mathrm{C-O}} : \text{Azimuthal angle}$$

4. **C to $\mathbf{O_H}$ (Hydroxyl Oxygen)**:

$$\rho_{C-OH} : \text{Distance}$$
$$\theta_{C-OH} : \text{Polar angle}$$
$$\phi_{C-OH} : \text{Azimuthal angle}$$

5. **O to $\mathbf{H_{OH}}$ (Hydroxyl Hydrogen)**:

$$\rho_{O-HOH} : \text{Distance}$$
$$\theta_{O-HOH} : \text{Polar angle}$$
$$\phi_{O-HOH} : \text{Azimuthal angle}$$

6. **C$\alpha$ to C$\beta$**:

$$\rho_{C\alpha-C\beta} : \text{Distance}$$
$$\theta_{C\alpha-C\beta} : \text{Polar angle}$$
$$\phi_{C\alpha-C\beta} : \text{Azimuthal angle}$$

7. **C$\beta$ to H$\beta_1$**:

$$\rho_{C\beta-H\beta_1} : \text{Distance}$$
$$\theta_{C\beta-H\beta_1} : \text{Polar angle}$$
$$\phi_{C\beta-H\beta_1} : \text{Azimuthal angle}$$

8. **C$\beta$ to H$\beta_2$**:

$$\rho_{C\beta-H\beta_2} : \text{Distance}$$
$$\theta_{C\beta-H\beta_2} : \text{Polar angle}$$
$$\phi_{C\beta-H\beta_2} : \text{Azimuthal angle}$$

9. **C$\beta$ to C$\gamma$**:

$$\rho_{C\beta-C\gamma} : \text{Distance}$$
$$\theta_{C\beta-C\gamma} : \text{Polar angle}$$
$$\phi_{C\beta-C\gamma} : \text{Azimuthal angle}$$

10. **C$\gamma$ to O$\delta_1$**:

$$\rho_{C\gamma-O\delta_1} : \text{Distance}$$
$$\theta_{C\gamma-O\delta_1} : \text{Polar angle}$$
$$\phi_{C\gamma-O\delta_1} : \text{Azimuthal angle}$$

11. **C$\gamma$ to O$\delta_2^-$**:

$$\rho_{C\gamma-O\delta_2^-} : \text{Distance}$$
$$\theta_{C\gamma-O\delta_2^-} : \text{Polar angle}$$
$$\phi_{C\gamma-O\delta_2^-} : \text{Azimuthal angle}$$

12. **N to $\mathbf{H_{N1}}$**:

$$\rho_{N-H_{N1}} : \text{Distance}$$
$$\theta_{N-H_{N1}} : \text{Polar angle}$$
$$\phi_{N-H_{N1}} : \text{Azimuthal angle}$$

13. **N to $\mathbf{H_{N2}}$**:

$$\rho_{N-H_{N2}} : \text{Distance}$$
$$\theta_{N-H_{N2}} : \text{Polar angle}$$
$$\phi_{N-H_{N2}} : \text{Azimuthal angle}$$

14. **C$\alpha$ to H$\alpha$**:

$$\rho_{C\alpha-H\alpha} : \text{Distance}$$
$$\theta_{C\alpha-H\alpha} : \text{Polar angle}$$
$$\phi_{C\alpha-H\alpha} : \text{Azimuthal angle}$$

### 5 An ABN-SCS Description with $\rho$, $\theta$ and $\phi$ for cysteine

#### Atoms in Cysteine (Cys, C)

1. **Backbone Atoms:**
    - N (Amine nitrogen)
    - H (Hydrogen attached to N)
    - C$\alpha$ (Alpha carbon)
    - H$\alpha$ (Hydrogen attached to C$\alpha$)
    - C (Carbonyl carbon)
    - O (Carbonyl oxygen)
    - $O^-$ (Carbonyl oxygen in the ionized state)
2. **Side Chain Atoms:**
    - C$\beta$ (Beta carbon)
    - H$\beta_1$ (First hydrogen attached to C$\beta$)
    - H$\beta_2$ (Second hydrogen attached to C$\beta$)
    - S$\gamma$ (Gamma sulfur)
    - H$\gamma$ (Hydrogen attached to S$\gamma$)

#### An ABN-SCS definition of the atomic bonding network of Cys

1. **N to C$\alpha$**:

$$\begin{aligned} \rho_{\mathrm{N-C\alpha}} &: \text{Distance} \\ \theta_{\mathrm{N-C\alpha}} &: \text{Polar angle} \\ \phi_{\mathrm{N-C\alpha}} &: \text{Azimuthal angle} \end{aligned}$$

2. **C$\alpha$ to C$\beta$**:

$$\begin{aligned} \rho_{\mathrm{C\alpha-C\beta}} &: \text{Distance} \\ \theta_{\mathrm{C\alpha-C\beta}} &: \text{Polar angle} \\ \phi_{\mathrm{C\alpha-C\beta}} &: \text{Azimuthal angle} \end{aligned}$$

3. **C$\alpha$ to C**:

$$\begin{aligned} \rho_{\mathrm{C\alpha-C}} &: \text{Distance} \\ \theta_{\mathrm{C\alpha-C}} &: \text{Polar angle} \\ \phi_{\mathrm{C\alpha-C}} &: \text{Azimuthal angle} \end{aligned}$$

4. **C to O (Carbonyl Oxygen)**:

$$\begin{aligned} \rho_{\mathrm{C-O}} &: \text{Distance} \\ \theta_{\mathrm{C-O}} &: \text{Polar angle} \\ \phi_{\mathrm{C-O}} &: \text{Azimuthal angle} \end{aligned}$$

5. **C to $\mathbf{O_H}$ (Hydroxyl Oxygen)**:

$$\rho_{C-OH} : \text{Distance}$$
$$\theta_{C-OH} : \text{Polar angle}$$
$$\phi_{C-OH} : \text{Azimuthal angle}$$

6. **O to $\mathbf{H_{OH}}$ (Hydroxyl Hydrogen)**:

$$\rho_{O-HOH} : \text{Distance}$$
$$\theta_{O-HOH} : \text{Polar angle}$$
$$\phi_{O-HOH} : \text{Azimuthal angle}$$

7. **N to $\mathbf{H_{N1}}$**:

$$\rho_{N-H_{N1}} : \text{Distance}$$
$$\theta_{N-H_{N1}} : \text{Polar angle}$$
$$\phi_{N-H_{N1}} : \text{Azimuthal angle}$$

8. **N to $\mathbf{H_{N2}}$**:

$$\rho_{N-H_{N2}} : \text{Distance}$$
$$\theta_{N-H_{N2}} : \text{Polar angle}$$
$$\phi_{N-H_{N2}} : \text{Azimuthal angle}$$

9. **C$\alpha$ to H$\alpha$**:

$$\rho_{C\alpha-H\alpha} : \text{Distance}$$
$$\theta_{C\alpha-H\alpha} : \text{Polar angle}$$
$$\phi_{C\alpha-H\alpha} : \text{Azimuthal angle}$$

10. **C$\beta$ to H$\beta_1$**:

$$\rho_{C\beta-H\beta_1} : \text{Distance}$$
$$\theta_{C\beta-H\beta_1} : \text{Polar angle}$$
$$\phi_{C\beta-H\beta_1} : \text{Azimuthal angle}$$

11. **C$\beta$ to H$\beta_2$**:

$$\rho_{C\beta-H\beta_2} : \text{Distance}$$
$$\theta_{C\beta-H\beta_2} : \text{Polar angle}$$
$$\phi_{C\beta-H\beta_2} : \text{Azimuthal angle}$$

12. **C$\beta$ to S$\gamma$**:

$$\rho_{C\beta-S\gamma} : \text{Distance}$$
$$\theta_{C\beta-S\gamma} : \text{Polar angle}$$
$$\phi_{C\beta-S\gamma} : \text{Azimuthal angle}$$

13. **S$\gamma$ to H$\gamma$**:

$$\rho_{S\gamma-H\gamma} : \text{Distance}$$
$$\theta_{S\gamma-H\gamma} : \text{Polar angle}$$
$$\phi_{S\gamma-H\gamma} : \text{Azimuthal angle}$$

### 6 An ABN-SCS Description with ρ, θ and φ for glutamate

#### Atoms in Glutamic Acid (Glu, E)

1. **Backbone Atoms:**
   - N (Amine nitrogen)
   - H (Hydrogen attached to N)
   - C$\alpha$ (Alpha carbon)
   - H$\alpha$ (Hydrogen attached to C$\alpha$)
   - C (Carbonyl carbon)
   - O (Carbonyl oxygen)
   - $O^-$ (Carbonyl oxygen in the ionized state)
2. **Side Chain Atoms:**
   - C$\beta$ (Beta carbon)
   - H$\beta_1$ (First hydrogen attached to C$\beta$)
   - H$\beta_2$ (Second hydrogen attached to C$\beta$)
   - C$\gamma$ (Gamma carbon)
   - H$\gamma_1$ (First hydrogen attached to C$\gamma$)
   - H$\gamma_2$ (Second hydrogen attached to C$\gamma$)
   - C$\delta$ (Delta carbon)
   - O$\delta_1^-$ (Ionized first oxygen attached to C$\delta$)
   - O$\delta_2^-$ (Ionized second oxygen attached to C$\delta$)
   - H$\delta_1$ (Hydroxyl hydrogen attached to O$\delta_1$)
   - H$\delta_2$ (Hydroxyl hydrogen attached to O$\delta_2$)

#### An ABN-SCS definition of the atomic bonding network of Glu

1. **N to C$\alpha$**:

$$\rho_{\mathrm{N-C\alpha}} : \text{Distance}$$
$$\theta_{\mathrm{N-C\alpha}} : \text{Polar angle}$$
$$\phi_{\mathrm{N-C\alpha}} : \text{Azimuthal angle}$$

2. **C$\alpha$ to C**:

$$\rho_{\mathrm{C\alpha-C}} : \text{Distance}$$
$$\theta_{\mathrm{C\alpha-C}} : \text{Polar angle}$$
$$\phi_{\mathrm{C\alpha-C}} : \text{Azimuthal angle}$$

3. **C to O (Carbonyl Oxygen)**:

$$\rho_{\mathrm{C-O}} : \text{Distance}$$
$$\theta_{\mathrm{C-O}} : \text{Polar angle}$$
$$\phi_{\mathrm{C-O}} : \text{Azimuthal angle}$$

4. **C to $\mathbf{O_H}$ (Hydroxyl Oxygen)**:

$$\rho_{\mathrm{C-OH}} : \text{Distance}$$
$$\theta_{\mathrm{C-OH}} : \text{Polar angle}$$
$$\phi_{\mathrm{C-OH}} : \text{Azimuthal angle}$$

5. **O to $\mathbf{H_{OH}}$ (Hydroxyl Hydrogen)**:

$$\rho_{\mathrm{O-HOH}} : \text{Distance}$$
$$\theta_{\mathrm{O-HOH}} : \text{Polar angle}$$
$$\phi_{\mathrm{O-HOH}} : \text{Azimuthal angle}$$

6. **C$\alpha$ to C$\beta$**:

$$\rho_{\mathrm{C}\alpha-\mathrm{C}\beta} : \text{Distance}$$
$$\theta_{\mathrm{C}\alpha-\mathrm{C}\beta} : \text{Polar angle}$$
$$\phi_{\mathrm{C}\alpha-\mathrm{C}\beta} : \text{Azimuthal angle}$$

7. **C$\beta$ to C$\gamma$**:

$$\rho_{\mathrm{C}\beta-\mathrm{C}\gamma} : \text{Distance}$$
$$\theta_{\mathrm{C}\beta-\mathrm{C}\gamma} : \text{Polar angle}$$
$$\phi_{\mathrm{C}\beta-\mathrm{C}\gamma} : \text{Azimuthal angle}$$

8. **C$\gamma$ to C$\delta$**:

$$\rho_{\mathrm{C}\gamma-\mathrm{C}\delta} : \text{Distance}$$
$$\theta_{\mathrm{C}\gamma-\mathrm{C}\delta} : \text{Polar angle}$$
$$\phi_{\mathrm{C}\gamma-\mathrm{C}\delta} : \text{Azimuthal angle}$$

9. **C$\delta$ to O$\delta_1^-$**:

$$\rho_{\mathrm{C}\delta-\mathrm{O}\delta_1^-} : \text{Distance}$$
$$\theta_{\mathrm{C}\delta-\mathrm{O}\delta_1^-} : \text{Polar angle}$$
$$\phi_{\mathrm{C}\delta-\mathrm{O}\delta_1^-} : \text{Azimuthal angle}$$

10. **C$\delta$ to O$\delta_2^-$**:

$$\rho_{\mathrm{C}\delta-\mathrm{O}\delta_2^-} : \text{Distance}$$
$$\theta_{\mathrm{C}\delta-\mathrm{O}\delta_2^-} : \text{Polar angle}$$
$$\phi_{\mathrm{C}\delta-\mathrm{O}\delta_2^-} : \text{Azimuthal angle}$$

11. **O$\delta_1$ to H$\delta_1$**:

$$\rho_{\mathrm{O}\delta_1-\mathrm{H}\delta_1} : \text{Distance}$$
$$\theta_{\mathrm{O}\delta_1-\mathrm{H}\delta_1} : \text{Polar angle}$$
$$\phi_{\mathrm{O}\delta_1-\mathrm{H}\delta_1} : \text{Azimuthal angle}$$

12. **N to $\mathbf{H_{N1}}$**:

$$\rho_{N-H_{N1}} : \text{Distance}$$
$$\theta_{N-H_{N1}} : \text{Polar angle}$$
$$\phi_{N-H_{N1}} : \text{Azimuthal angle}$$

13. **N to $\mathbf{H_{N2}}$**:

$$\rho_{N-H_{N2}} : \text{Distance}$$
$$\theta_{N-H_{N2}} : \text{Polar angle}$$
$$\phi_{N-H_{N2}} : \text{Azimuthal angle}$$

14. **C$\alpha$ to H$\alpha$**:

$$\rho_{C\alpha-H\alpha} : \text{Distance}$$
$$\theta_{C\alpha-H\alpha} : \text{Polar angle}$$
$$\phi_{C\alpha-H\alpha} : \text{Azimuthal angle}$$

15. **C$\beta$ to H$\beta_1$**:

$$\rho_{C\beta-H\beta_1} : \text{Distance}$$
$$\theta_{C\beta-H\beta_1} : \text{Polar angle}$$
$$\phi_{C\beta-H\beta_1} : \text{Azimuthal angle}$$

16. **C$\beta$ to H$\beta_2$**:

$$\rho_{C\beta-H\beta_2} : \text{Distance}$$
$$\theta_{C\beta-H\beta_2} : \text{Polar angle}$$
$$\phi_{C\beta-H\beta_2} : \text{Azimuthal angle}$$

17. **C$\gamma$ to H$\gamma_1$**:

$$\rho_{C\gamma-H\gamma_1} : \text{Distance}$$
$$\theta_{C\gamma-H\gamma_1} : \text{Polar angle}$$
$$\phi_{C\gamma-H\gamma_1} : \text{Azimuthal angle}$$

18. **C$\gamma$ to H$\gamma_2$**:

$$\rho_{C\gamma-H\gamma_2} : \text{Distance}$$
$$\theta_{C\gamma-H\gamma_2} : \text{Polar angle}$$
$$\phi_{C\gamma-H\gamma_2} : \text{Azimuthal angle}$$

### 7 An ABN-SCS Description with ρ, θ and ϕ for glutamine

#### Atoms in Glutamine (Gln, Q)

1. **Backbone Atoms:**
   - N (Amine nitrogen)
   - H (Hydrogen attached to N)
   - C$\alpha$ (Alpha carbon)
   - H$\alpha$ (Hydrogen attached to C$\alpha$)
   - C (Carbonyl carbon)
   - O (Carbonyl oxygen)
   - $O^-$ (Carbonyl oxygen in the ionized state)
2. **Side Chain Atoms:**
   - C$\beta$ (Beta carbon)
   - H$\beta_1$ (First hydrogen attached to C$\beta$)
   - H$\beta_2$ (Second hydrogen attached to C$\beta$)
   - C$\gamma$ (Gamma carbon)
   - H$\gamma_1$ (First hydrogen attached to C$\gamma$)
   - H$\gamma_2$ (Second hydrogen attached to C$\gamma$)
   - C$\delta_1$ (Delta carbon 1)
   - O$\delta_1$ (Oxygen attached to C$\delta_1$)
   - N$\delta$ (Amide nitrogen)
   - H$\delta_1$ (First hydrogen attached to N$\delta$)
   - H$\delta_2$ (Second hydrogen attached to N$\delta$)

#### An ABN-SCS definition of the atomic bonding network of Gln

1. **N to C$\alpha$**:

$$\rho_{\mathrm{N-C}\alpha} : \text{Distance}$$
$$\theta_{\mathrm{N-C}\alpha} : \text{Polar angle}$$
$$\phi_{\mathrm{N-C}\alpha} : \text{Azimuthal angle}$$

2. **C$\alpha$ to C**:

$$\rho_{\mathrm{C}\alpha-\mathrm{C}} : \text{Distance}$$
$$\theta_{\mathrm{C}\alpha-\mathrm{C}} : \text{Polar angle}$$
$$\phi_{\mathrm{C}\alpha-\mathrm{C}} : \text{Azimuthal angle}$$

3. **C to O (Carbonyl Oxygen)**:

$$\rho_{\mathrm{C-O}} : \text{Distance}$$
$$\theta_{\mathrm{C-O}} : \text{Polar angle}$$
$$\phi_{\mathrm{C-O}} : \text{Azimuthal angle}$$

4. **C to $\mathbf{O_H}$ (Hydroxyl Oxygen)**:

$$\rho_{\text{C}-\text{OH}} : \text{Distance}$$
$$\theta_{\text{C}-\text{OH}} : \text{Polar angle}$$
$$\phi_{\text{C}-\text{OH}} : \text{Azimuthal angle}$$

5. **O to $\mathbf{H_{OH}}$ (Hydroxyl Hydrogen)**:

$$\rho_{\text{O}-\text{HOH}} : \text{Distance}$$
$$\theta_{\text{O}-\text{HOH}} : \text{Polar angle}$$
$$\phi_{\text{O}-\text{HOH}} : \text{Azimuthal angle}$$

6. **C$\alpha$ to C$\beta$**:

$$\rho_{\text{C}\alpha-\text{C}\beta} : \text{Distance}$$
$$\theta_{\text{C}\alpha-\text{C}\beta} : \text{Polar angle}$$
$$\phi_{\text{C}\alpha-\text{C}\beta} : \text{Azimuthal angle}$$

7. **C$\beta$ to C$\gamma$**:

$$\rho_{\text{C}\beta-\text{C}\gamma} : \text{Distance}$$
$$\theta_{\text{C}\beta-\text{C}\gamma} : \text{Polar angle}$$
$$\phi_{\text{C}\beta-\text{C}\gamma} : \text{Azimuthal angle}$$

8. **C$\gamma$ to H$\gamma_1$**:

$$\rho_{\text{C}\gamma-\text{H}\gamma_1} : \text{Distance}$$
$$\theta_{\text{C}\gamma-\text{H}\gamma_1} : \text{Polar angle}$$
$$\phi_{\text{C}\gamma-\text{H}\gamma_1} : \text{Azimuthal angle}$$

9. **C$\gamma$ to H$\gamma_2$**:

$$\rho_{\text{C}\gamma-\text{H}\gamma_2} : \text{Distance}$$
$$\theta_{\text{C}\gamma-\text{H}\gamma_2} : \text{Polar angle}$$
$$\phi_{\text{C}\gamma-\text{H}\gamma_2} : \text{Azimuthal angle}$$

10. **C$\gamma$ to C$\delta_1$**:

$$\rho_{\text{C}\gamma-\text{C}\delta_1} : \text{Distance}$$
$$\theta_{\text{C}\gamma-\text{C}\delta_1} : \text{Polar angle}$$
$$\phi_{\text{C}\gamma-\text{C}\delta_1} : \text{Azimuthal angle}$$

11. **C$\delta_1$ to O$\delta_1$**:

$$\rho_{\text{C}\delta_1-\text{O}\delta_1} : \text{Distance}$$
$$\theta_{\text{C}\delta_1-\text{O}\delta_1} : \text{Polar angle}$$
$$\phi_{\text{C}\delta_1-\text{O}\delta_1} : \text{Azimuthal angle}$$

12. **C**$\delta_1$ **to N**$\delta$:

$\rho_{\mathrm{C}\delta_1-\mathrm{N}\delta}$ : Distance
$\theta_{\mathrm{C}\delta_1-\mathrm{N}\delta}$ : Polar angle
$\phi_{\mathrm{C}\delta_1-\mathrm{N}\delta}$ : Azimuthal angle

13. **N**$\delta$ **to H**$\delta_1$:

$\rho_{\mathrm{N}\delta-\mathrm{H}\delta_1}$ : Distance
$\theta_{\mathrm{N}\delta-\mathrm{H}\delta_1}$ : Polar angle
$\phi_{\mathrm{N}\delta-\mathrm{H}\delta_1}$ : Azimuthal angle

14. **N**$\delta$ **to H**$\delta_2$:

$\rho_{\mathrm{N}\delta-\mathrm{H}\delta_2}$ : Distance
$\theta_{\mathrm{N}\delta-\mathrm{H}\delta_2}$ : Polar angle
$\phi_{\mathrm{N}\delta-\mathrm{H}\delta_2}$ : Azimuthal angle

15. **N to H**$_{\mathbf{N1}}$:

$\rho_{\mathrm{N}-\mathrm{H}_{\mathrm{N1}}}$ : Distance
$\theta_{\mathrm{N}-\mathrm{H}_{\mathrm{N1}}}$ : Polar angle
$\phi_{\mathrm{N}-\mathrm{H}_{\mathrm{N1}}}$ : Azimuthal angle

16. **N to H**$_{\mathbf{N2}}$:

$\rho_{\mathrm{N}-\mathrm{H}_{\mathrm{N2}}}$ : Distance
$\theta_{\mathrm{N}-\mathrm{H}_{\mathrm{N2}}}$ : Polar angle
$\phi_{\mathrm{N}-\mathrm{H}_{\mathrm{N2}}}$ : Azimuthal angle

17. **C**$\alpha$ **to H**$\alpha$:

$\rho_{\mathrm{C}\alpha-\mathrm{H}\alpha}$ : Distance
$\theta_{\mathrm{C}\alpha-\mathrm{H}\alpha}$ : Polar angle
$\phi_{\mathrm{C}\alpha-\mathrm{H}\alpha}$ : Azimuthal angle

18. **C**$\beta$ **to H**$\beta_1$:

$\rho_{\mathrm{C}\beta-\mathrm{H}\beta_1}$ : Distance
$\theta_{\mathrm{C}\beta-\mathrm{H}\beta_1}$ : Polar angle
$\phi_{\mathrm{C}\beta-\mathrm{H}\beta_1}$ : Azimuthal angle

19. **C**$\beta$ **to H**$\beta_2$:

$\rho_{\mathrm{C}\beta-\mathrm{H}\beta_2}$ : Distance
$\theta_{\mathrm{C}\beta-\mathrm{H}\beta_2}$ : Polar angle
$\phi_{\mathrm{C}\beta-\mathrm{H}\beta_2}$ : Azimuthal angle

### 8 An ABN-SCS Description with $\rho$, $\theta$ and $\phi$ for glycine

#### Atoms in Glycine (Gly, G)

1. **Backbone Atoms:**
   - N (Amine nitrogen)
   - H (Hydrogen attached to N)
   - $C\alpha$ (Alpha carbon)
   - $H\alpha_1$ (First hydrogen attached to $C\alpha$)
   - $H\alpha_2$ (Second hydrogen attached to $C\alpha$)
   - C (Carbonyl carbon)
   - O (Carbonyl oxygen)
   - $O^-$ (Carbonyl oxygen in the ionized state)
2. **Side Chain Atoms:**
   - Glycine does not have a side chain. The alpha carbon ($C\alpha$) is directly bonded to two hydrogens ($H\alpha_1$ and $H\alpha_2$).

#### An ABN-SCS definition of the atomic bonding network of Gly

1. **N to $C\alpha$**:

$$\rho_{N-C\alpha} : \text{Distance}$$
$$\theta_{N-C\alpha} : \text{Polar angle}$$
$$\phi_{N-C\alpha} : \text{Azimuthal angle}$$

2. **$C\alpha$ to C**:

$$\rho_{C\alpha-C} : \text{Distance}$$
$$\theta_{C\alpha-C} : \text{Polar angle}$$
$$\phi_{C\alpha-C} : \text{Azimuthal angle}$$

3. **C to O (Carbonyl Oxygen)**:

$$\rho_{C-O} : \text{Distance}$$
$$\theta_{C-O} : \text{Polar angle}$$
$$\phi_{C-O} : \text{Azimuthal angle}$$

4. **C to $O_H$ (Hydroxyl Oxygen)**:

$$\rho_{C-OH} : \text{Distance}$$
$$\theta_{C-OH} : \text{Polar angle}$$
$$\phi_{C-OH} : \text{Azimuthal angle}$$

5. **O to $H_{OH}$ (Hydroxyl Hydrogen)**:

$$\rho_{O-HOH} : \text{Distance}$$
$$\theta_{O-HOH} : \text{Polar angle}$$
$$\phi_{O-HOH} : \text{Azimuthal angle}$$

6. **N to $\mathbf{H_{N1}}$**:

$$\rho_{\mathrm{N-H_{N1}}} : \text{Distance}$$
$$\theta_{\mathrm{N-H_{N1}}} : \text{Polar angle}$$
$$\phi_{\mathrm{N-H_{N1}}} : \text{Azimuthal angle}$$

7. **N to $\mathbf{H_{N2}}$**:

$$\rho_{\mathrm{N-H_{N2}}} : \text{Distance}$$
$$\theta_{\mathrm{N-H_{N2}}} : \text{Polar angle}$$
$$\phi_{\mathrm{N-H_{N2}}} : \text{Azimuthal angle}$$

8. **C$\alpha$ to H$\alpha_{\mathbf{1}}$**:

$$\rho_{\mathrm{C\alpha-H\alpha_1}} : \text{Distance}$$
$$\theta_{\mathrm{C\alpha-H\alpha_1}} : \text{Polar angle}$$
$$\phi_{\mathrm{C\alpha-H\alpha_1}} : \text{Azimuthal angle}$$

9. **C$\alpha$ to H$\alpha_{\mathbf{2}}$**:

$$\rho_{\mathrm{C\alpha-H\alpha_2}} : \text{Distance}$$
$$\theta_{\mathrm{C\alpha-H\alpha_2}} : \text{Polar angle}$$
$$\phi_{\mathrm{C\alpha-H\alpha_2}} : \text{Azimuthal angle}$$

### 9 An ABN-SCS Description with ρ, θ and ϕ for histidine

#### Atoms in Histidine (His, H)

1. **Backbone Atoms:**
   - N (Amine nitrogen)
   - H (Hydrogen attached to N)
   - C$\alpha$ (Alpha carbon)
   - H$\alpha$ (Hydrogen attached to C$\alpha$)
   - C (Carbonyl carbon)
   - O (Carbonyl oxygen)
   - $O^-$ (Carbonyl oxygen in the ionized state)
2. **Side Chain Atoms:**
   - C$\beta$ (Beta carbon)
   - H$\beta_1$ (First hydrogen attached to C$\beta$)
   - H$\beta_2$ (Second hydrogen attached to C$\beta$)
   - C$\gamma$ (Gamma carbon)
   - N$\delta_1$ (Delta nitrogen)
   - H$\delta_1$ (Hydrogen attached to N$\delta_1$)
   - C$\epsilon_1$ (Epsilon carbon)
   - H$\epsilon_1$ (Hydrogen attached to C$\epsilon_1$)
   - N$\epsilon_2$ (Epsilon nitrogen)
   - H$\epsilon_2$ (Hydrogen attached to N$\epsilon_2$)
   - C$\delta_2$ (Delta carbon)
   - H$\delta_2$ (Hydrogen attached to C$\delta_2$)

#### An ABN-SCS definition of the atomic bonding network of His

1. **N to C$\alpha$**:

$$\rho_{\mathrm{N-C}\alpha} : \text{Distance}$$
$$\theta_{\mathrm{N-C}\alpha} : \text{Polar angle}$$
$$\phi_{\mathrm{N-C}\alpha} : \text{Azimuthal angle}$$

2. **C$\alpha$ to C**:

$$\rho_{\mathrm{C}\alpha\mathrm{-C}} : \text{Distance}$$
$$\theta_{\mathrm{C}\alpha\mathrm{-C}} : \text{Polar angle}$$
$$\phi_{\mathrm{C}\alpha\mathrm{-C}} : \text{Azimuthal angle}$$

3. **C to O (Carbonyl Oxygen)**:

$$\rho_{\mathrm{C-O}} : \text{Distance}$$
$$\theta_{\mathrm{C-O}} : \text{Polar angle}$$
$$\phi_{\mathrm{C-O}} : \text{Azimuthal angle}$$

4. **C to $\mathbf{O_H}$ (Hydroxyl Oxygen)**:

$$\rho_{C-OH} : \text{Distance}$$
$$\theta_{C-OH} : \text{Polar angle}$$
$$\phi_{C-OH} : \text{Azimuthal angle}$$

5. **O to $\mathbf{H_{OH}}$ (Hydroxyl Hydrogen)**:

$$\rho_{O-HOH} : \text{Distance}$$
$$\theta_{O-HOH} : \text{Polar angle}$$
$$\phi_{O-HOH} : \text{Azimuthal angle}$$

6. **C$\alpha$ to C$\beta$**:

$$\rho_{C\alpha-C\beta} : \text{Distance}$$
$$\theta_{C\alpha-C\beta} : \text{Polar angle}$$
$$\phi_{C\alpha-C\beta} : \text{Azimuthal angle}$$

7. **C$\beta$ to C$\gamma$**:

$$\rho_{C\beta-C\gamma} : \text{Distance}$$
$$\theta_{C\beta-C\gamma} : \text{Polar angle}$$
$$\phi_{C\beta-C\gamma} : \text{Azimuthal angle}$$

8. **C$\gamma$ to N$\delta_1$**:

$$\rho_{C\gamma-N\delta_1} : \text{Distance}$$
$$\theta_{C\gamma-N\delta_1} : \text{Polar angle}$$
$$\phi_{C\gamma-N\delta_1} : \text{Azimuthal angle}$$

9. **N$\delta_1$ to H$\delta_1$**:

$$\rho_{N\delta_1-H\delta_1} : \text{Distance}$$
$$\theta_{N\delta_1-H\delta_1} : \text{Polar angle}$$
$$\phi_{N\delta_1-H\delta_1} : \text{Azimuthal angle}$$

10. **N$\delta_1$ to C$\epsilon_1$**:

$$\rho_{N\delta_1-C\epsilon_1} : \text{Distance}$$
$$\theta_{N\delta_1-C\epsilon_1} : \text{Polar angle}$$
$$\phi_{N\delta_1-C\epsilon_1} : \text{Azimuthal angle}$$

11. **C$\epsilon_1$ to H$\epsilon_1$**:

$$\rho_{C\epsilon_1-H\epsilon_1} : \text{Distance}$$
$$\theta_{C\epsilon_1-H\epsilon_1} : \text{Polar angle}$$
$$\phi_{C\epsilon_1-H\epsilon_1} : \text{Azimuthal angle}$$

12. $\mathbf{C\epsilon_1}$ **to** $\mathbf{N\epsilon_2}$:

$$\rho_{\mathrm{C}\epsilon_1-\mathrm{N}\epsilon_2} : \text{Distance}$$
$$\theta_{\mathrm{C}\epsilon_1-\mathrm{N}\epsilon_2} : \text{Polar angle}$$
$$\phi_{\mathrm{C}\epsilon_1-\mathrm{N}\epsilon_2} : \text{Azimuthal angle}$$

13. $\mathbf{N\epsilon_2}$ **to** $\mathbf{H\epsilon_2}$:

$$\rho_{\mathrm{N}\epsilon_2-\mathrm{H}\epsilon_2} : \text{Distance}$$
$$\theta_{\mathrm{N}\epsilon_2-\mathrm{H}\epsilon_2} : \text{Polar angle}$$
$$\phi_{\mathrm{N}\epsilon_2-\mathrm{H}\epsilon_2} : \text{Azimuthal angle}$$

14. $\mathbf{N\epsilon_2}$ **to** $\mathbf{C\delta_2}$:

$$\rho_{\mathrm{N}\epsilon_2-\mathrm{C}\delta_2} : \text{Distance}$$
$$\theta_{\mathrm{N}\epsilon_2-\mathrm{C}\delta_2} : \text{Polar angle}$$
$$\phi_{\mathrm{N}\epsilon_2-\mathrm{C}\delta_2} : \text{Azimuthal angle}$$

15. $\mathbf{C\delta_2}$ **to** $\mathbf{H\delta_2}$:

$$\rho_{\mathrm{C}\delta_2-\mathrm{H}\delta_2} : \text{Distance}$$
$$\theta_{\mathrm{C}\delta_2-\mathrm{H}\delta_2} : \text{Polar angle}$$
$$\phi_{\mathrm{C}\delta_2-\mathrm{H}\delta_2} : \text{Azimuthal angle}$$

16. $\mathbf{C\delta_2}$ **to** $\mathbf{C\gamma}$:

$$\rho_{\mathrm{C}\delta_2-\mathrm{C}\gamma} : \text{Distance}$$
$$\theta_{\mathrm{C}\delta_2-\mathrm{C}\gamma} : \text{Polar angle}$$
$$\phi_{\mathrm{C}\delta_2-\mathrm{C}\gamma} : \text{Azimuthal angle}$$

17. **N to** $\mathbf{H_{N1}}$:

$$\rho_{\mathrm{N}-\mathrm{H}_{\mathrm{N1}}} : \text{Distance}$$
$$\theta_{\mathrm{N}-\mathrm{H}_{\mathrm{N1}}} : \text{Polar angle}$$
$$\phi_{\mathrm{N}-\mathrm{H}_{\mathrm{N1}}} : \text{Azimuthal angle}$$

18. **N to** $\mathbf{H_{N2}}$:

$$\rho_{\mathrm{N}-\mathrm{H}_{\mathrm{N2}}} : \text{Distance}$$
$$\theta_{\mathrm{N}-\mathrm{H}_{\mathrm{N2}}} : \text{Polar angle}$$
$$\phi_{\mathrm{N}-\mathrm{H}_{\mathrm{N2}}} : \text{Azimuthal angle}$$

19. $\mathbf{C\alpha}$ **to** $\mathbf{H\alpha}$:

$$\rho_{\mathrm{C}\alpha-\mathrm{H}\alpha} : \text{Distance}$$
$$\theta_{\mathrm{C}\alpha-\mathrm{H}\alpha} : \text{Polar angle}$$
$$\phi_{\mathrm{C}\alpha-\mathrm{H}\alpha} : \text{Azimuthal angle}$$

20. **N to $\mathbf{H_{N1}}$**:

$$\rho_{\mathrm{N-H_{N1}}} : \text{Distance}$$
$$\theta_{\mathrm{N-H_{N1}}} : \text{Polar angle}$$
$$\phi_{\mathrm{N-H_{N1}}} : \text{Azimuthal angle}$$

21. **N to $\mathbf{H_{N2}}$**:

$$\rho_{\mathrm{N-H_{N2}}} : \text{Distance}$$
$$\theta_{\mathrm{N-H_{N2}}} : \text{Polar angle}$$
$$\phi_{\mathrm{N-H_{N2}}} : \text{Azimuthal angle}$$

22. **C$\alpha$ to H$\alpha$**:

$$\rho_{\mathrm{C\alpha-H\alpha}} : \text{Distance}$$
$$\theta_{\mathrm{C\alpha-H\alpha}} : \text{Polar angle}$$
$$\phi_{\mathrm{C\alpha-H\alpha}} : \text{Azimuthal angle}$$

### 10 An ABN-SCS Description with ρ, θ and ϕ for isoleucine

#### Atoms in Isoleucine (Ile, I)

1. **Backbone Atoms:**
   - N (Amine nitrogen)
   - H (Hydrogen attached to N)
   - C$\alpha$ (Alpha carbon)
   - H$\alpha$ (Hydrogen attached to C$\alpha$)
   - C (Carbonyl carbon)
   - O (Carbonyl oxygen)
   - $O^-$ (Carbonyl oxygen in the ionized state)
2. **Side Chain Atoms:**
   - C$\beta$ (Beta carbon)
   - H$\beta_1$ (First hydrogen attached to C$\beta$)
   - C$\gamma_1$ (Gamma carbon)
   - H$\gamma_1 a$ (Hydrogen attached to C$\gamma_1$)
   - H$\gamma_1 b$ (Hydrogen attached to C$\gamma_1$)
   - C$\gamma_2$ (Gamma carbon)
   - H$\gamma_1 a$ (Hydrogen attached to C$\gamma_2$)
   - H$\gamma_1 b$ (Hydrogen attached to C$\gamma_2$)
   - H$\gamma_1 c$ (Hydrogen attached to C$\gamma_2$)
   - C$\delta$ (Delta carbon)
   - H$\delta_1$ (First hydrogen attached to C$\delta$)
   - H$\delta_2$ (Second hydrogen attached to C$\delta$)
   - H$\delta_3$ (Third hydrogen attached to C$\delta$)

#### An ABN-SCS definition of the atomic bonding network of Ile

1. **N to C$\alpha$**:

$$\begin{aligned} \rho_{\mathrm{N-C}\alpha} &: \text{Distance} \\ \theta_{\mathrm{N-C}\alpha} &: \text{Polar angle} \\ \phi_{\mathrm{N-C}\alpha} &: \text{Azimuthal angle} \end{aligned}$$

2. **C$\alpha$ to C**:

$$\begin{aligned} \rho_{\mathrm{C}\alpha-\mathrm{C}} &: \text{Distance} \\ \theta_{\mathrm{C}\alpha-\mathrm{C}} &: \text{Polar angle} \\ \phi_{\mathrm{C}\alpha-\mathrm{C}} &: \text{Azimuthal angle} \end{aligned}$$

3. **C to O (Carbonyl Oxygen)**:

$$\rho_{C-O} : \text{Distance}$$
$$\theta_{C-O} : \text{Polar angle}$$
$$\phi_{C-O} : \text{Azimuthal angle}$$

4. **C to $O_H$ (Hydroxyl Oxygen)**:

$$\rho_{C-OH} : \text{Distance}$$
$$\theta_{C-OH} : \text{Polar angle}$$
$$\phi_{C-OH} : \text{Azimuthal angle}$$

5. **O to $H_{OH}$ (Hydroxyl Hydrogen)**:

$$\rho_{O-HOH} : \text{Distance}$$
$$\theta_{O-HOH} : \text{Polar angle}$$
$$\phi_{O-HOH} : \text{Azimuthal angle}$$

6. **C$\alpha$ to C$\beta$**:

$$\rho_{C\alpha-C\beta} : \text{Distance}$$
$$\theta_{C\alpha-C\beta} : \text{Polar angle}$$
$$\phi_{C\alpha-C\beta} : \text{Azimuthal angle}$$

7. **C$\beta$ to C$\gamma_1$**:

$$\rho_{C\beta-C\gamma_1} : \text{Distance}$$
$$\theta_{C\beta-C\gamma_1} : \text{Polar angle}$$
$$\phi_{C\beta-C\gamma_1} : \text{Azimuthal angle}$$

8. **C$\gamma_1$ to H$\gamma_1 a$**:

$$\rho_{C\gamma_1-H\gamma_1 a} : \text{Distance}$$
$$\theta_{C\gamma_1-H\gamma_1 a} : \text{Polar angle}$$
$$\phi_{C\gamma_1-H\gamma_1 a} : \text{Azimuthal angle}$$

9. **C$\gamma_1$ to H$\gamma_1 b$**:

$$\rho_{C\gamma_1-H\gamma_1 b} : \text{Distance}$$
$$\theta_{C\gamma_1-H\gamma_1 b} : \text{Polar angle}$$
$$\phi_{C\gamma_1-H\gamma_1 b} : \text{Azimuthal angle}$$

10. **C$\beta$ to C$\gamma_2$**:

$$\rho_{C\beta-C\gamma_2} : \text{Distance}$$
$$\theta_{C\beta-C\gamma_2} : \text{Polar angle}$$
$$\phi_{C\beta-C\gamma_2} : \text{Azimuthal angle}$$

11. **C**$\gamma_2$ **to H**$\gamma_2 a$:

$$\rho_{C\gamma_2-H\gamma_2 a} : \text{Distance}$$
$$\theta_{C\gamma_2-H\gamma_2 a} : \text{Polar angle}$$
$$\phi_{C\gamma_2-H\gamma_2 a} : \text{Azimuthal angle}$$

12. **C**$\gamma_2$ **to H**$\gamma_2 b$:

$$\rho_{C\gamma_2-H\gamma_2 b} : \text{Distance}$$
$$\theta_{C\gamma_2-H\gamma_2 b} : \text{Polar angle}$$
$$\phi_{C\gamma_2-H\gamma_2 b} : \text{Azimuthal angle}$$

13. **C**$\gamma_2$ **to H**$\gamma_2 c$:

$$\rho_{C\gamma_2-H\gamma_2 c} : \text{Distance}$$
$$\theta_{C\gamma_2-H\gamma_2 c} : \text{Polar angle}$$
$$\phi_{C\gamma_2-H\gamma_2 c} : \text{Azimuthal angle}$$

14. **C**$\gamma_1$ **to C**$\delta$:

$$\rho_{C\gamma_1-C\delta} : \text{Distance}$$
$$\theta_{C\gamma_1-C\delta} : \text{Polar angle}$$
$$\phi_{C\gamma_1-C\delta} : \text{Azimuthal angle}$$

15. **C**$\delta$ **to H**$\delta_1$:

$$\rho_{C\delta-H\delta_1} : \text{Distance}$$
$$\theta_{C\delta-H\delta_1} : \text{Polar angle}$$
$$\phi_{C\delta-H\delta_1} : \text{Azimuthal angle}$$

16. **C**$\delta$ **to H**$\delta_2$:

$$\rho_{C\delta-H\delta_2} : \text{Distance}$$
$$\theta_{C\delta-H\delta_2} : \text{Polar angle}$$
$$\phi_{C\delta-H\delta_2} : \text{Azimuthal angle}$$

17. **C**$\delta$ **to H**$\delta_3$:

$$\rho_{C\delta-H\delta_3} : \text{Distance}$$
$$\theta_{C\delta-H\delta_3} : \text{Polar angle}$$
$$\phi_{C\delta-H\delta_3} : \text{Azimuthal angle}$$

18. **N to H$_{\mathbf{N1}}$**:

$$\rho_{N-H_{N1}} : \text{Distance}$$
$$\theta_{N-H_{N1}} : \text{Polar angle}$$
$$\phi_{N-H_{N1}} : \text{Azimuthal angle}$$

19. **N to $\mathbf{H_{N2}}$**:

$$\rho_{\mathrm{N-H_{N2}}} : \text{Distance}$$
$$\theta_{\mathrm{N-H_{N2}}} : \text{Polar angle}$$
$$\phi_{\mathrm{N-H_{N2}}} : \text{Azimuthal angle}$$

20. **C$\alpha$ to H$\alpha$**:

$$\rho_{\mathrm{C}\alpha-\mathrm{H}\alpha} : \text{Distance}$$
$$\theta_{\mathrm{C}\alpha-\mathrm{H}\alpha} : \text{Polar angle}$$
$$\phi_{\mathrm{C}\alpha-\mathrm{H}\alpha} : \text{Azimuthal angle}$$

21. **C$\beta$ to H$\alpha$**:

$$\rho_{\mathrm{C}\beta-\mathrm{H}\alpha} : \text{Distance}$$
$$\theta_{\mathrm{C}\beta-\mathrm{H}\alpha} : \text{Polar angle}$$
$$\phi_{\mathrm{C}\beta-\mathrm{H}\alpha} : \text{Azimuthal angle}$$

### 11 An ABN-SCS Description with ρ, θ and ϕ for leucine

#### Atoms in Leucine (Leu, L)

1. **Backbone Atoms:**
   - N (Amine nitrogen)
   - H (Hydrogen attached to N)
   - C$\alpha$ (Alpha carbon)
   - H$\alpha$ (Hydrogen attached to C$\alpha$)
   - C (Carbonyl carbon)
   - O (Carbonyl oxygen)
   - $O^-$ (Carbonyl oxygen in the ionized state)
2. **Side Chain Atoms:**
   - C$\beta$ (Beta carbon)
   - H$\beta_1$ (First hydrogen attached to C$\beta$)
   - H$\beta_2$ (Second hydrogen attached to C$\beta$)
   - C$\gamma$ (Gamma carbon)
   - H$\gamma$ (Hydrogen attached to C$\gamma$)
   - C$\delta_1$ (Delta carbon 1)
   - H$\delta_1 a$ (First hydrogen attached to C$\delta_1$)
   - H$\delta_1 b$ (Second hydrogen attached to C$\delta_1$)
   - H$\delta_1 c$ (Third hydrogen attached to C$\delta_1$)
   - C$\delta_2$ (Delta carbon 2)
   - H$\delta_2 a$ (First hydrogen attached to C$\delta_2$)
   - H$\delta_2 b$ (Second hydrogen attached to C$\delta_2$)
   - H$\delta_2 c$ (Third hydrogen attached to C$\delta_2$)

#### An ABN-SCS definition of the atomic bonding network of Leu

1. **N to C$\alpha$**:

$$\rho_{\mathrm{N-C\alpha}} : \text{Distance}$$
$$\theta_{\mathrm{N-C\alpha}} : \text{Polar angle}$$
$$\phi_{\mathrm{N-C\alpha}} : \text{Azimuthal angle}$$

2. **C$\alpha$ to C**:

$$\rho_{\mathrm{C\alpha-C}} : \text{Distance}$$
$$\theta_{\mathrm{C\alpha-C}} : \text{Polar angle}$$
$$\phi_{\mathrm{C\alpha-C}} : \text{Azimuthal angle}$$

3. **C to O (Carbonyl Oxygen)**:

$$\rho_{C-O} : \text{Distance}$$
$$\theta_{C-O} : \text{Polar angle}$$
$$\phi_{C-O} : \text{Azimuthal angle}$$

4. **C to $O_H$ (Hydroxyl Oxygen)**:

$$\rho_{C-OH} : \text{Distance}$$
$$\theta_{C-OH} : \text{Polar angle}$$
$$\phi_{C-OH} : \text{Azimuthal angle}$$

5. **O to $H_{OH}$ (Hydroxyl Hydrogen)**:

$$\rho_{O-HOH} : \text{Distance}$$
$$\theta_{O-HOH} : \text{Polar angle}$$
$$\phi_{O-HOH} : \text{Azimuthal angle}$$

6. **C$\alpha$ to C$\beta$**:

$$\rho_{C\alpha-C\beta} : \text{Distance}$$
$$\theta_{C\alpha-C\beta} : \text{Polar angle}$$
$$\phi_{C\alpha-C\beta} : \text{Azimuthal angle}$$

7. **C$\beta$ to C$\gamma$**:

$$\rho_{C\beta-C\gamma} : \text{Distance}$$
$$\theta_{C\beta-C\gamma} : \text{Polar angle}$$
$$\phi_{C\beta-C\gamma} : \text{Azimuthal angle}$$

8. **C$\beta$ to H$\beta_1$**:

$$\rho_{C\beta-H\beta_1} : \text{Distance}$$
$$\theta_{C\beta-H\beta_1} : \text{Polar angle}$$
$$\phi_{C\beta-H\beta_1} : \text{Azimuthal angle}$$

9. **C$\beta$ to H$\beta_2$**:

$$\rho_{C\beta-H\beta_2} : \text{Distance}$$
$$\theta_{C\beta-H\beta_2} : \text{Polar angle}$$
$$\phi_{C\beta-H\beta_2} : \text{Azimuthal angle}$$

10. **C$\gamma$ to H$\gamma$**:

$$\rho_{C\gamma-H\gamma} : \text{Distance}$$
$$\theta_{C\gamma-H\gamma} : \text{Polar angle}$$
$$\phi_{C\gamma-H\gamma} : \text{Azimuthal angle}$$

11. $\mathbf{C\gamma}$ **to** $\mathbf{C\delta_1}$:

$$\rho_{\mathrm{C}\gamma-\mathrm{C}\delta_1} : \text{Distance}$$
$$\theta_{\mathrm{C}\gamma-\mathrm{C}\delta_1} : \text{Polar angle}$$
$$\phi_{\mathrm{C}\gamma-\mathrm{C}\delta_1} : \text{Azimuthal angle}$$

12. $\mathbf{C\gamma}$ **to** $\mathbf{C\delta_2}$:

$$\rho_{\mathrm{C}\gamma-\mathrm{C}\delta_2} : \text{Distance}$$
$$\theta_{\mathrm{C}\gamma-\mathrm{C}\delta_2} : \text{Polar angle}$$
$$\phi_{\mathrm{C}\gamma-\mathrm{C}\delta_2} : \text{Azimuthal angle}$$

13. $\mathbf{C\delta_1}$ **to** $\mathbf{H\delta_1}a$:

$$\rho_{\mathrm{C}\delta_1-\mathrm{H}\delta_1 a} : \text{Distance}$$
$$\theta_{\mathrm{C}\delta_1-\mathrm{H}\delta_1 a} : \text{Polar angle}$$
$$\phi_{\mathrm{C}\delta_1-\mathrm{H}\delta_1 a} : \text{Azimuthal angle}$$

14. $\mathbf{C\delta_1}$ **to** $\mathbf{H\delta_1}b$:

$$\rho_{\mathrm{C}\delta_1-\mathrm{H}\delta_1 b} : \text{Distance}$$
$$\theta_{\mathrm{C}\delta_1-\mathrm{H}\delta_1 b} : \text{Polar angle}$$
$$\phi_{\mathrm{C}\delta_1-\mathrm{H}\delta_1 b} : \text{Azimuthal angle}$$

15. $\mathbf{C\delta_1}$ **to** $\mathbf{H\delta_1}c$:

$$\rho_{\mathrm{C}\delta_1-\mathrm{H}\delta_1 c} : \text{Distance}$$
$$\theta_{\mathrm{C}\delta_1-\mathrm{H}\delta_1 c} : \text{Polar angle}$$
$$\phi_{\mathrm{C}\delta_1-\mathrm{H}\delta_1 c} : \text{Azimuthal angle}$$

16. $\mathbf{C\delta_2}$ **to** $\mathbf{H\delta_2}a$:

$$\rho_{\mathrm{C}\delta_2-\mathrm{H}\delta_2 a} : \text{Distance}$$
$$\theta_{\mathrm{C}\delta_2-\mathrm{H}\delta_2 a} : \text{Polar angle}$$
$$\phi_{\mathrm{C}\delta_2-\mathrm{H}\delta_2 a} : \text{Azimuthal angle}$$

17. $\mathbf{C\delta_2}$ **to** $\mathbf{H\delta_2}b$:

$$\rho_{\mathrm{C}\delta_2-\mathrm{H}\delta_2 b} : \text{Distance}$$
$$\theta_{\mathrm{C}\delta_2-\mathrm{H}\delta_2 b} : \text{Polar angle}$$
$$\phi_{\mathrm{C}\delta_2-\mathrm{H}\delta_2 b} : \text{Azimuthal angle}$$

18. $\mathbf{C\delta_2}$ **to** $\mathbf{H\delta_2}c$:

$$\rho_{\mathrm{C}\delta_2-\mathrm{H}\delta_2 c} : \text{Distance}$$
$$\theta_{\mathrm{C}\delta_2-\mathrm{H}\delta_2 c} : \text{Polar angle}$$
$$\phi_{\mathrm{C}\delta_2-\mathrm{H}\delta_2 c} : \text{Azimuthal angle}$$

19. **N to $\mathbf{H_{N1}}$**:

$$\rho_{\mathrm{N-H_{N1}}} : \text{Distance}$$
$$\theta_{\mathrm{N-H_{N1}}} : \text{Polar angle}$$
$$\phi_{\mathrm{N-H_{N1}}} : \text{Azimuthal angle}$$

20. **N to $\mathbf{H_{N2}}$**:

$$\rho_{\mathrm{N-H_{N2}}} : \text{Distance}$$
$$\theta_{\mathrm{N-H_{N2}}} : \text{Polar angle}$$
$$\phi_{\mathrm{N-H_{N2}}} : \text{Azimuthal angle}$$

21. **C$\alpha$ to H$\alpha$**:

$$\rho_{\mathrm{C\alpha-H\alpha}} : \text{Distance}$$
$$\theta_{\mathrm{C\alpha-H\alpha}} : \text{Polar angle}$$
$$\phi_{\mathrm{C\alpha-H\alpha}} : \text{Azimuthal angle}$$

### 12 An ABN-SCS Description with ρ, θ and ϕ for lysine

#### Atoms in Lysine (Lys, K)

1. **Backbone Atoms:**
    - N (Amine nitrogen)
    - H (Hydrogen attached to N)
    - C$\alpha$ (Alpha carbon)
    - H$\alpha$ (Hydrogen attached to C$\alpha$)
    - C (Carbonyl carbon)
    - O (Carbonyl oxygen)
    - $O^-$ (Carbonyl oxygen in the ionized state)
2. **Side Chain Atoms:**
    - C$\beta$ (Beta carbon)
    - H$\beta_1$ (First hydrogen attached to C$\beta$)
    - H$\beta_2$ (Second hydrogen attached to C$\beta$)
    - C$\gamma$ (Gamma carbon)
    - H$\gamma_1$ (First hydrogen attached to C$\gamma$)
    - H$\gamma_2$ (Second hydrogen attached to C$\gamma$)
    - C$\delta$ (Delta carbon)
    - H$\delta_1$ (First hydrogen attached to C$\delta$)
    - H$\delta_2$ (Second hydrogen attached to C$\delta$)
    - C$\epsilon$ (Epsilon carbon)
    - H$\epsilon_1$ (First hydrogen attached to C$\epsilon$)
    - H$\epsilon_2$ (Second hydrogen attached to C$\epsilon$)
    - N$\zeta$ (Zeta nitrogen)
    - H$\zeta_1$ (First hydrogen attached to N$\zeta$) (if protonated)
    - H$\zeta_2$ (Second hydrogen attached to N$\zeta$) (if protonated)
    - H$\zeta_3$ (Third hydrogen attached to N$\zeta$) (if protonated)

#### An ABN-SCS definition of the atomic bonding network of Lys

1. **N to C$\alpha$**:

$$\rho_{N-C\alpha} : \text{Distance}$$
$$\theta_{N-C\alpha} : \text{Polar angle}$$
$$\phi_{N-C\alpha} : \text{Azimuthal angle}$$

2. **C$\alpha$ to C**:

$$\rho_{C\alpha-C} : \text{Distance}$$
$$\theta_{C\alpha-C} : \text{Polar angle}$$
$$\phi_{C\alpha-C} : \text{Azimuthal angle}$$

3. **C to O (Carbonyl Oxygen)**:

$$\rho_{C-O} : \text{Distance}$$
$$\theta_{C-O} : \text{Polar angle}$$
$$\phi_{C-O} : \text{Azimuthal angle}$$

4. **C to $\mathbf{O_H}$ (Hydroxyl Oxygen)**:

$$\rho_{C-OH} : \text{Distance}$$
$$\theta_{C-OH} : \text{Polar angle}$$
$$\phi_{C-OH} : \text{Azimuthal angle}$$

5. **O to $\mathbf{H_{OH}}$ (Hydroxyl Hydrogen)**:

$$\rho_{O-HOH} : \text{Distance}$$
$$\theta_{O-HOH} : \text{Polar angle}$$
$$\phi_{O-HOH} : \text{Azimuthal angle}$$

6. **C$\alpha$ to C$\beta$**:

$$\rho_{C\alpha-C\beta} : \text{Distance}$$
$$\theta_{C\alpha-C\beta} : \text{Polar angle}$$
$$\phi_{C\alpha-C\beta} : \text{Azimuthal angle}$$

7. **C$\beta$ to C$\gamma$**:

$$\rho_{C\beta-C\gamma} : \text{Distance}$$
$$\theta_{C\beta-C\gamma} : \text{Polar angle}$$
$$\phi_{C\beta-C\gamma} : \text{Azimuthal angle}$$

8. **C$\gamma$ to C$\delta$**:

$$\rho_{C\gamma-C\delta} : \text{Distance}$$
$$\theta_{C\gamma-C\delta} : \text{Polar angle}$$
$$\phi_{C\gamma-C\delta} : \text{Azimuthal angle}$$

9. **C$\delta$ to C$\epsilon$**:

$$\rho_{C\delta-C\epsilon} : \text{Distance}$$
$$\theta_{C\delta-C\epsilon} : \text{Polar angle}$$
$$\phi_{C\delta-C\epsilon} : \text{Azimuthal angle}$$

10. **C$\epsilon$ to N$\zeta$**:

$$\rho_{C\epsilon-N\zeta} : \text{Distance}$$
$$\theta_{C\epsilon-N\zeta} : \text{Polar angle}$$
$$\phi_{C\epsilon-N\zeta} : \text{Azimuthal angle}$$

11. **N to $\mathbf{H_{N1}}$**:

$$\rho_{N-H_{N1}} : \text{Distance}$$
$$\theta_{N-H_{N1}} : \text{Polar angle}$$
$$\phi_{N-H_{N1}} : \text{Azimuthal angle}$$

12. **N to $\mathbf{H_{N2}}$**:

$$\rho_{N-H_{N2}} : \text{Distance}$$
$$\theta_{N-H_{N2}} : \text{Polar angle}$$
$$\phi_{N-H_{N2}} : \text{Azimuthal angle}$$

13. **C$\alpha$ to H$\alpha$**:

$$\rho_{C\alpha-H\alpha} : \text{Distance}$$
$$\theta_{C\alpha-H\alpha} : \text{Polar angle}$$
$$\phi_{C\alpha-H\alpha} : \text{Azimuthal angle}$$

14. **C$\beta$ to H$\beta_1$**:

$$\rho_{C\beta-H\beta_1} : \text{Distance}$$
$$\theta_{C\beta-H\beta_1} : \text{Polar angle}$$
$$\phi_{C\beta-H\beta_1} : \text{Azimuthal angle}$$

15. **C$\beta$ to H$\beta_2$**:

$$\rho_{C\beta-H\beta_2} : \text{Distance}$$
$$\theta_{C\beta-H\beta_2} : \text{Polar angle}$$
$$\phi_{C\beta-H\beta_2} : \text{Azimuthal angle}$$

16. **C$\gamma$ to H$\gamma_1$**:

$$\rho_{C\gamma-H\gamma_1} : \text{Distance}$$
$$\theta_{C\gamma-H\gamma_1} : \text{Polar angle}$$
$$\phi_{C\gamma-H\gamma_1} : \text{Azimuthal angle}$$

17. **C$\gamma$ to H$\gamma_2$**:

$$\rho_{C\gamma-H\gamma_2} : \text{Distance}$$
$$\theta_{C\gamma-H\gamma_2} : \text{Polar angle}$$
$$\phi_{C\gamma-H\gamma_2} : \text{Azimuthal angle}$$

18. **C$\delta$ to H$\delta_1$**:

$$\rho_{C\delta-H\delta_1} : \text{Distance}$$
$$\theta_{C\delta-H\delta_1} : \text{Polar angle}$$
$$\phi_{C\delta-H\delta_1} : \text{Azimuthal angle}$$

19. **C$\delta$ to H$\delta_2$**:

$$\rho_{C\delta-H\delta_2} : \text{Distance}$$
$$\theta_{C\delta-H\delta_2} : \text{Polar angle}$$
$$\phi_{C\delta-H\delta_2} : \text{Azimuthal angle}$$

20. **C$\epsilon$ to H$\epsilon_1$**:

$$\rho_{C\epsilon-H\epsilon_1} : \text{Distance}$$
$$\theta_{C\epsilon-H\epsilon_1} : \text{Polar angle}$$
$$\phi_{C\epsilon-H\epsilon_1} : \text{Azimuthal angle}$$

21. **C$\epsilon$ to H$\epsilon_2$**:

$$\rho_{C\epsilon-H\epsilon_2} : \text{Distance}$$
$$\theta_{C\epsilon-H\epsilon_2} : \text{Polar angle}$$
$$\phi_{C\epsilon-H\epsilon_2} : \text{Azimuthal angle}$$

22. **N$\zeta$ to H$\zeta_1$**:

$$\rho_{N\zeta-H\zeta_1} : \text{Distance}$$
$$\theta_{N\zeta-H\zeta_1} : \text{Polar angle}$$
$$\phi_{N\zeta-H\zeta_1} : \text{Azimuthal angle}$$

23. **N$\zeta$ to H$\zeta_2$**:

$$\rho_{N\zeta-H\zeta_2} : \text{Distance}$$
$$\theta_{N\zeta-H\zeta_2} : \text{Polar angle}$$
$$\phi_{N\zeta-H\zeta_2} : \text{Azimuthal angle}$$

24. **N$\zeta$ to H$\zeta_3$**:

$$\rho_{N\zeta-H\zeta_3} : \text{Distance}$$
$$\theta_{N\zeta-H\zeta_3} : \text{Polar angle}$$
$$\phi_{N\zeta-H\zeta_3} : \text{Azimuthal angle}$$

### 13 An ABN-SCS Description with $\rho$, $\theta$ and $\phi$ for methionine

#### Atoms in Methionine (Met, M)

1. **Backbone Atoms:**
   - N (Amine nitrogen)
   - H (Hydrogen attached to N)
   - $C\alpha$ (Alpha carbon)
   - $H\alpha$ (Hydrogen attached to $C\alpha$)
   - C (Carbonyl carbon)
   - O (Carbonyl oxygen)
2. **Side Chain Atoms:**
   - $C\beta$ (Beta carbon)
   - $H\beta_1$ (First hydrogen attached to $C\beta$)
   - $H\beta_2$ (Second hydrogen attached to $C\beta$)
   - $C\gamma$ (Gamma carbon)
   - $H\gamma_1$ (First hydrogen attached to $C\gamma$)
   - $H\gamma_2$ (Second hydrogen attached to $C\gamma$)
   - $S\delta$ (Sulfur atom)
   - $C\epsilon$ (Epsilon carbon)
   - $H\epsilon_1$ (First hydrogen attached to $C\epsilon$)
   - $H\epsilon_2$ (Second hydrogen attached to $C\epsilon$)
   - $H\epsilon_3$ (Third hydrogen attached to $C\epsilon$)

#### An ABN-SCS definition of the atomic bonding network of Met

1. **N to C$\alpha$**:

$$\rho_{\mathrm{N-C}\alpha} : \text{Distance}$$
$$\theta_{\mathrm{N-C}\alpha} : \text{Polar angle}$$
$$\phi_{\mathrm{N-C}\alpha} : \text{Azimuthal angle}$$

2. **C$\alpha$ to C**:

$$\rho_{\mathrm{C}\alpha-\mathrm{C}} : \text{Distance}$$
$$\theta_{\mathrm{C}\alpha-\mathrm{C}} : \text{Polar angle}$$
$$\phi_{\mathrm{C}\alpha-\mathrm{C}} : \text{Azimuthal angle}$$

3. **C to O (Carbonyl Oxygen)**:

$$\rho_{\mathrm{C-O}} : \text{Distance}$$
$$\theta_{\mathrm{C-O}} : \text{Polar angle}$$
$$\phi_{\mathrm{C-O}} : \text{Azimuthal angle}$$

4. **C to $\mathbf{O_H}$ (Hydroxyl Oxygen)**:

$$\rho_{\text{C}-\text{OH}} : \text{Distance}$$
$$\theta_{\text{C}-\text{OH}} : \text{Polar angle}$$
$$\phi_{\text{C}-\text{OH}} : \text{Azimuthal angle}$$

5. **O to $\mathbf{H_{OH}}$ (Hydroxyl Hydrogen)**:

$$\rho_{\text{O}-\text{HOH}} : \text{Distance}$$
$$\theta_{\text{O}-\text{HOH}} : \text{Polar angle}$$
$$\phi_{\text{O}-\text{HOH}} : \text{Azimuthal angle}$$

6. **C$\alpha$ to C$\beta$**:

$$\rho_{\text{C}\alpha-\text{C}\beta} : \text{Distance}$$
$$\theta_{\text{C}\alpha-\text{C}\beta} : \text{Polar angle}$$
$$\phi_{\text{C}\alpha-\text{C}\beta} : \text{Azimuthal angle}$$

7. **C$\beta$ to C$\gamma$**:

$$\rho_{\text{C}\beta-\text{C}\gamma} : \text{Distance}$$
$$\theta_{\text{C}\beta-\text{C}\gamma} : \text{Polar angle}$$
$$\phi_{\text{C}\beta-\text{C}\gamma} : \text{Azimuthal angle}$$

8. **C$\beta$ to H$\beta_1$**:

$$\rho_{\text{C}\beta-\text{H}\beta_1} : \text{Distance}$$
$$\theta_{\text{C}\beta-\text{H}\beta_1} : \text{Polar angle}$$
$$\phi_{\text{C}\beta-\text{H}\beta_1} : \text{Azimuthal angle}$$

9. **C$\beta$ to H$\beta_2$**:

$$\rho_{\text{C}\beta-\text{H}\beta_2} : \text{Distance}$$
$$\theta_{\text{C}\beta-\text{H}\beta_2} : \text{Polar angle}$$
$$\phi_{\text{C}\beta-\text{H}\beta_2} : \text{Azimuthal angle}$$

10. **C$\gamma$ to H$\gamma_1$**:

$$\rho_{\text{C}\gamma-\text{H}\gamma_1} : \text{Distance}$$
$$\theta_{\text{C}\gamma-\text{H}\gamma_1} : \text{Polar angle}$$
$$\phi_{\text{C}\gamma-\text{H}\gamma_1} : \text{Azimuthal angle}$$

11. **C$\gamma$ to H$\gamma_2$**:

$$\rho_{\text{C}\gamma-\text{H}\gamma_2} : \text{Distance}$$
$$\theta_{\text{C}\gamma-\text{H}\gamma_2} : \text{Polar angle}$$
$$\phi_{\text{C}\gamma-\text{H}\gamma_2} : \text{Azimuthal angle}$$

12. **C$\gamma$ to S$\delta$**:

$$\rho_{\mathrm{C}\gamma-\mathrm{S}\delta} : \text{Distance}$$
$$\theta_{\mathrm{C}\gamma-\mathrm{S}\delta} : \text{Polar angle}$$
$$\phi_{\mathrm{C}\gamma-\mathrm{S}\delta} : \text{Azimuthal angle}$$

13. **S$\delta$ to C$\epsilon$**:

$$\rho_{\mathrm{S}\delta-\mathrm{C}\epsilon} : \text{Distance}$$
$$\theta_{\mathrm{S}\delta-\mathrm{C}\epsilon} : \text{Polar angle}$$
$$\phi_{\mathrm{S}\delta-\mathrm{C}\epsilon} : \text{Azimuthal angle}$$

14. **C$\epsilon$ to H$\epsilon_1$**:

$$\rho_{\mathrm{C}\epsilon-\mathrm{H}\epsilon_1} : \text{Distance}$$
$$\theta_{\mathrm{C}\epsilon-\mathrm{H}\epsilon_1} : \text{Polar angle}$$
$$\phi_{\mathrm{C}\epsilon-\mathrm{H}\epsilon_1} : \text{Azimuthal angle}$$

15. **C$\epsilon$ to H$\epsilon_2$**:

$$\rho_{\mathrm{C}\epsilon-\mathrm{H}\epsilon_2} : \text{Distance}$$
$$\theta_{\mathrm{C}\epsilon-\mathrm{H}\epsilon_2} : \text{Polar angle}$$
$$\phi_{\mathrm{C}\epsilon-\mathrm{H}\epsilon_2} : \text{Azimuthal angle}$$

16. **C$\epsilon$ to H$\epsilon_3$**:

$$\rho_{\mathrm{C}\epsilon-\mathrm{H}\epsilon_3} : \text{Distance}$$
$$\theta_{\mathrm{C}\epsilon-\mathrm{H}\epsilon_3} : \text{Polar angle}$$
$$\phi_{\mathrm{C}\epsilon-\mathrm{H}\epsilon_3} : \text{Azimuthal angle}$$

17. **N to $\mathbf{H_{N1}}$**:

$$\rho_{\mathrm{N}-\mathrm{H_{N1}}} : \text{Distance}$$
$$\theta_{\mathrm{N}-\mathrm{H_{N1}}} : \text{Polar angle}$$
$$\phi_{\mathrm{N}-\mathrm{H_{N1}}} : \text{Azimuthal angle}$$

18. **N to $\mathbf{H_{N2}}$**:

$$\rho_{\mathrm{N}-\mathrm{H_{N2}}} : \text{Distance}$$
$$\theta_{\mathrm{N}-\mathrm{H_{N2}}} : \text{Polar angle}$$
$$\phi_{\mathrm{N}-\mathrm{H_{N2}}} : \text{Azimuthal angle}$$

19. **C$\alpha$ to H$\alpha$**:

$$\rho_{\mathrm{C}\alpha-\mathrm{H}\alpha} : \text{Distance}$$
$$\theta_{\mathrm{C}\alpha-\mathrm{H}\alpha} : \text{Polar angle}$$
$$\phi_{\mathrm{C}\alpha-\mathrm{H}\alpha} : \text{Azimuthal angle}$$

### 14 An ABN-SCS Description with ρ, θ and ϕ for phenylalanine

#### Atoms in Phenylalanine (Phe, F)

1. **Backbone Atoms:**
   - N (Amine nitrogen)
   - H (Hydrogen attached to N)
   - C$\alpha$ (Alpha carbon)
   - H$\alpha$ (Hydrogen attached to C$\alpha$)
   - C (Carbonyl carbon)
   - O (Carbonyl oxygen)
   - $O^-$ (Carbonyl oxygen in the ionized state)
2. **Side Chain Atoms:**
   - C$\beta$ (Beta carbon)
   - H$\beta_1$ (First hydrogen attached to C$\beta$)
   - H$\beta_2$ (Second hydrogen attached to C$\beta$)
   - C$\gamma$ (Gamma carbon)
   - C$\delta_1$ (Delta carbon 1)
   - H$\delta_1$ (Hydrogen attached to C$\delta_1$)
   - C$\delta_2$ (Delta carbon 2)
   - H$\delta_2$ (Hydrogen attached to C$\delta_2$)
   - C$\epsilon_1$ (Epsilon carbon 1)
   - H$\epsilon_1$ (Hydrogen attached to C$\epsilon_1$)
   - C$\epsilon_2$ (Epsilon carbon 2)
   - H$\epsilon_2$ (Hydrogen attached to C$\epsilon_2$)
   - C$\zeta$ (Zeta carbon)
   - H$\zeta$ (Hydrogen attached to C$\zeta$)

#### An ABN-SCS definition of the atomic bonding network of Phe

1. **N to C$\alpha$**:

$$\rho_{\mathrm{N-C}\alpha} : \text{Distance}$$
$$\theta_{\mathrm{N-C}\alpha} : \text{Polar angle}$$
$$\phi_{\mathrm{N-C}\alpha} : \text{Azimuthal angle}$$

2. **C$\alpha$ to C**:

$$\rho_{\mathrm{C}\alpha-\mathrm{C}} : \text{Distance}$$
$$\theta_{\mathrm{C}\alpha-\mathrm{C}} : \text{Polar angle}$$
$$\phi_{\mathrm{C}\alpha-\mathrm{C}} : \text{Azimuthal angle}$$

3. **C to O (Carbonyl Oxygen)**:

$$\rho_{C-O} : \text{Distance}$$
$$\theta_{C-O} : \text{Polar angle}$$
$$\phi_{C-O} : \text{Azimuthal angle}$$

4. **C to $O_H$ (Hydroxyl Oxygen)**:

$$\rho_{C-OH} : \text{Distance}$$
$$\theta_{C-OH} : \text{Polar angle}$$
$$\phi_{C-OH} : \text{Azimuthal angle}$$

5. **O to $H_{OH}$ (Hydroxyl Hydrogen)**:

$$\rho_{O-HOH} : \text{Distance}$$
$$\theta_{O-HOH} : \text{Polar angle}$$
$$\phi_{O-HOH} : \text{Azimuthal angle}$$

6. **C$\alpha$ to C$\beta$**:

$$\rho_{C\alpha-C\beta} : \text{Distance}$$
$$\theta_{C\alpha-C\beta} : \text{Polar angle}$$
$$\phi_{C\alpha-C\beta} : \text{Azimuthal angle}$$

7. **C$\beta$ to C$\gamma$**:

$$\rho_{C\beta-C\gamma} : \text{Distance}$$
$$\theta_{C\beta-C\gamma} : \text{Polar angle}$$
$$\phi_{C\beta-C\gamma} : \text{Azimuthal angle}$$

8. **C$\gamma$ to C$\delta_1$**:

$$\rho_{C\gamma-C\delta_1} : \text{Distance}$$
$$\theta_{C\gamma-C\delta_1} : \text{Polar angle}$$
$$\phi_{C\gamma-C\delta_1} : \text{Azimuthal angle}$$

9. **C$\gamma$ to C$\delta_2$**:

$$\rho_{C\gamma-C\delta_2} : \text{Distance}$$
$$\theta_{C\gamma-C\delta_2} : \text{Polar angle}$$
$$\phi_{C\gamma-C\delta_2} : \text{Azimuthal angle}$$

10. **C$\delta_1$ to H$\delta_1$**:

$$\rho_{C\delta_1-H\delta_1} : \text{Distance}$$
$$\theta_{C\delta_1-H\delta_1} : \text{Polar angle}$$
$$\phi_{C\delta_1-H\delta_1} : \text{Azimuthal angle}$$

11. **C$\delta_2$ to H$\delta_2$**:

$$\rho_{\mathrm{C}\delta_2-\mathrm{H}\delta_2} : \text{Distance}$$
$$\theta_{\mathrm{C}\delta_2-\mathrm{H}\delta_2} : \text{Polar angle}$$
$$\phi_{\mathrm{C}\delta_2-\mathrm{H}\delta_2} : \text{Azimuthal angle}$$

12. **C$\delta_1$ to C$\epsilon_1$**:

$$\rho_{\mathrm{C}\delta_1-\mathrm{C}\epsilon_1} : \text{Distance}$$
$$\theta_{\mathrm{C}\delta_1-\mathrm{C}\epsilon_1} : \text{Polar angle}$$
$$\phi_{\mathrm{C}\delta_1-\mathrm{C}\epsilon_1} : \text{Azimuthal angle}$$

13. **C$\delta_2$ to C$\epsilon_2$**:

$$\rho_{\mathrm{C}\delta_2-\mathrm{C}\epsilon_2} : \text{Distance}$$
$$\theta_{\mathrm{C}\delta_2-\mathrm{C}\epsilon_2} : \text{Polar angle}$$
$$\phi_{\mathrm{C}\delta_2-\mathrm{C}\epsilon_2} : \text{Azimuthal angle}$$

14. **C$\epsilon_1$ to H$\epsilon_1$**:

$$\rho_{\mathrm{C}\epsilon_1-\mathrm{H}\epsilon_1} : \text{Distance}$$
$$\theta_{\mathrm{C}\epsilon_1-\mathrm{H}\epsilon_1} : \text{Polar angle}$$
$$\phi_{\mathrm{C}\epsilon_1-\mathrm{H}\epsilon_1} : \text{Azimuthal angle}$$

15. **C$\epsilon_2$ to H$\epsilon_2$**:

$$\rho_{\mathrm{C}\epsilon_2-\mathrm{H}\epsilon_2} : \text{Distance}$$
$$\theta_{\mathrm{C}\epsilon_2-\mathrm{H}\epsilon_2} : \text{Polar angle}$$
$$\phi_{\mathrm{C}\epsilon_2-\mathrm{H}\epsilon_2} : \text{Azimuthal angle}$$

16. **C$\epsilon_1$ to C$\zeta$**:

$$\rho_{\mathrm{C}\epsilon_1-\mathrm{C}\zeta} : \text{Distance}$$
$$\theta_{\mathrm{C}\epsilon_1-\mathrm{C}\zeta} : \text{Polar angle}$$
$$\phi_{\mathrm{C}\epsilon_1-\mathrm{C}\zeta} : \text{Azimuthal angle}$$

17. **C$\epsilon_2$ to C$\zeta$**:

$$\rho_{\mathrm{C}\epsilon_2-\mathrm{C}\zeta} : \text{Distance}$$
$$\theta_{\mathrm{C}\epsilon_2-\mathrm{C}\zeta} : \text{Polar angle}$$
$$\phi_{\mathrm{C}\epsilon_2-\mathrm{C}\zeta} : \text{Azimuthal angle}$$

18. **C$\zeta$ to H$\zeta$**:

$$\rho_{\mathrm{C}\zeta-\mathrm{H}\zeta} : \text{Distance}$$
$$\theta_{\mathrm{C}\zeta-\mathrm{H}\zeta} : \text{Polar angle}$$
$$\phi_{\mathrm{C}\zeta-\mathrm{H}\zeta} : \text{Azimuthal angle}$$

19. **N to $\mathbf{H_{N1}}$**:

$$\rho_{N-H_{N1}} : \text{Distance}$$
$$\theta_{N-H_{N1}} : \text{Polar angle}$$
$$\phi_{N-H_{N1}} : \text{Azimuthal angle}$$

20. **N to $\mathbf{H_{N2}}$**:

$$\rho_{N-H_{N2}} : \text{Distance}$$
$$\theta_{N-H_{N2}} : \text{Polar angle}$$
$$\phi_{N-H_{N2}} : \text{Azimuthal angle}$$

21. **C$\alpha$ to H$\alpha$**:

$$\rho_{C\alpha-H\alpha} : \text{Distance}$$
$$\theta_{C\alpha-H\alpha} : \text{Polar angle}$$
$$\phi_{C\alpha-H\alpha} : \text{Azimuthal angle}$$

### 15 An ABN-SCS Description with $\rho$, $\theta$ and $\phi$ for proline

#### Atoms in Proline (Pro, P)

1. **Backbone Atoms:**
   - N (Amine nitrogen)
   - H (Hydrogen attached to N)
   - C$\alpha$ (Alpha carbon)
   - H$\alpha$ (Hydrogen attached to C$\alpha$)
   - C (Carbonyl carbon)
   - O (Carbonyl oxygen)
   - $O^-$ (Carbonyl oxygen in the ionized state)
2. **Side Chain Atoms:**
   - C$\beta$ (Beta carbon)
   - H$\beta_1$ (First hydrogen attached to C$\beta$)
   - H$\beta_2$ (Second hydrogen attached to C$\beta$)
   - C$\gamma$ (Gamma carbon)
   - H$\gamma_1$ (First hydrogen attached to C$\gamma$)
   - H$\gamma_2$ (Second hydrogen attached to C$\gamma$)
   - C$\delta$ (Delta carbon)
   - H$\delta_1$ (First hydrogen attached to C$\delta$)
   - H$\delta_2$ (Second hydrogen attached to C$\delta$)

#### An ABN-SCS definition of the atomic bonding network of Pro

1. **N to C$\alpha$**:

$$\begin{aligned} \rho_{\mathrm{N-C}\alpha} &: \text{Distance} \\ \theta_{\mathrm{N-C}\alpha} &: \text{Polar angle} \\ \phi_{\mathrm{N-C}\alpha} &: \text{Azimuthal angle} \end{aligned}$$

2. **C$\alpha$ to C**:

$$\begin{aligned} \rho_{\mathrm{C}\alpha-\mathrm{C}} &: \text{Distance} \\ \theta_{\mathrm{C}\alpha-\mathrm{C}} &: \text{Polar angle} \\ \phi_{\mathrm{C}\alpha-\mathrm{C}} &: \text{Azimuthal angle} \end{aligned}$$

3. **C to O (Carbonyl Oxygen)**:

$$\begin{aligned} \rho_{\mathrm{C-O}} &: \text{Distance} \\ \theta_{\mathrm{C-O}} &: \text{Polar angle} \\ \phi_{\mathrm{C-O}} &: \text{Azimuthal angle} \end{aligned}$$

4. **C to $\mathbf{O_H}$ (Hydroxyl Oxygen)**:

$$\rho_{C-OH} : \text{Distance}$$
$$\theta_{C-OH} : \text{Polar angle}$$
$$\phi_{C-OH} : \text{Azimuthal angle}$$

5. **O to $\mathbf{H_{OH}}$ (Hydroxyl Hydrogen)**:

$$\rho_{O-HOH} : \text{Distance}$$
$$\theta_{O-HOH} : \text{Polar angle}$$
$$\phi_{O-HOH} : \text{Azimuthal angle}$$

6. **C$\alpha$ to C$\beta$**:

$$\rho_{C\alpha-C\beta} : \text{Distance}$$
$$\theta_{C\alpha-C\beta} : \text{Polar angle}$$
$$\phi_{C\alpha-C\beta} : \text{Azimuthal angle}$$

7. **C$\beta$ to C$\gamma$**:

$$\rho_{C\beta-C\gamma} : \text{Distance}$$
$$\theta_{C\beta-C\gamma} : \text{Polar angle}$$
$$\phi_{C\beta-C\gamma} : \text{Azimuthal angle}$$

8. **C$\gamma$ to C$\delta$**:

$$\rho_{C\gamma-C\delta} : \text{Distance}$$
$$\theta_{C\gamma-C\delta} : \text{Polar angle}$$
$$\phi_{C\gamma-C\delta} : \text{Azimuthal angle}$$

9. **C$\beta$ to H$\beta_1$**:

$$\rho_{C\beta-H\beta_1} : \text{Distance}$$
$$\theta_{C\beta-H\beta_1} : \text{Polar angle}$$
$$\phi_{C\beta-H\beta_1} : \text{Azimuthal angle}$$

10. **C$\beta$ to H$\beta_2$**:

$$\rho_{C\beta-H\beta_2} : \text{Distance}$$
$$\theta_{C\beta-H\beta_2} : \text{Polar angle}$$
$$\phi_{C\beta-H\beta_2} : \text{Azimuthal angle}$$

11. **C$\gamma$ to H$\gamma_1$**:

$$\rho_{C\gamma-H\gamma_1} : \text{Distance}$$
$$\theta_{C\gamma-H\gamma_1} : \text{Polar angle}$$
$$\phi_{C\gamma-H\gamma_1} : \text{Azimuthal angle}$$

12. **C$\gamma$ to H$\gamma_2$**:

$$\rho_{C\gamma-H\gamma_2} : \text{Distance}$$
$$\theta_{C\gamma-H\gamma_2} : \text{Polar angle}$$
$$\phi_{C\gamma-H\gamma_2} : \text{Azimuthal angle}$$

13. **C$\delta$ to H$\delta_1$**:

$$\rho_{C\delta-H\delta_1} : \text{Distance}$$
$$\theta_{C\delta-H\delta_1} : \text{Polar angle}$$
$$\phi_{C\delta-H\delta_1} : \text{Azimuthal angle}$$

14. **C$\delta$ to H$\delta_2$**:

$$\rho_{C\delta-H\delta_2} : \text{Distance}$$
$$\theta_{C\delta-H\delta_2} : \text{Polar angle}$$
$$\phi_{C\delta-H\delta_2} : \text{Azimuthal angle}$$

15. **N to $\mathbf{H_{N1}}$**:

$$\rho_{N-H_{N1}} : \text{Distance}$$
$$\theta_{N-H_{N1}} : \text{Polar angle}$$
$$\phi_{N-H_{N1}} : \text{Azimuthal angle}$$

16. **N to $\mathbf{H_{N2}}$**:

$$\rho_{N-H_{N2}} : \text{Distance}$$
$$\theta_{N-H_{N2}} : \text{Polar angle}$$
$$\phi_{N-H_{N2}} : \text{Azimuthal angle}$$

17. **C$\alpha$ to H$\alpha$**:

$$\rho_{C\alpha-H\alpha} : \text{Distance}$$
$$\theta_{C\alpha-H\alpha} : \text{Polar angle}$$
$$\phi_{C\alpha-H\alpha} : \text{Azimuthal angle}$$

18. **C$\delta$ to N**:

$$\rho_{C\delta-N} : \text{Distance}$$
$$\theta_{C\delta-N} : \text{Polar angle}$$
$$\phi_{C\delta-N} : \text{Azimuthal angle}$$

### 16 An ABN-SCS Description with ρ, θ and ϕ for serine

#### Atoms in Serine (Ser, S)

1. **Backbone Atoms:**
   - N (Amine nitrogen)
   - H (Hydrogen attached to N)
   - C$\alpha$ (Alpha carbon)
   - H$\alpha$ (Hydrogen attached to C$\alpha$)
   - C (Carbonyl carbon)
   - O (Carbonyl oxygen)
   - $O^-$ (Carbonyl oxygen in the ionized state)
2. **Side Chain Atoms:**
   - C$\beta$ (Beta carbon)
   - H$\beta_1$ (First hydrogen attached to C$\beta$)
   - H$\beta_2$ (Second hydrogen attached to C$\beta$)
   - O$\gamma$ (Gamma oxygen, hydroxyl group)
   - H$\gamma$ (Hydrogen attached to O$\gamma$)

#### An ABN-SCS definition of the atomic bonding network of Ser

1. **N to C$\alpha$**:

$$\rho_{\mathrm{N-C\alpha}} : \text{Distance}$$
$$\theta_{\mathrm{N-C\alpha}} : \text{Polar angle}$$
$$\phi_{\mathrm{N-C\alpha}} : \text{Azimuthal angle}$$

2. **C$\alpha$ to C**:

$$\rho_{\mathrm{C\alpha-C}} : \text{Distance}$$
$$\theta_{\mathrm{C\alpha-C}} : \text{Polar angle}$$
$$\phi_{\mathrm{C\alpha-C}} : \text{Azimuthal angle}$$

3. **C to O (Carbonyl Oxygen)**:

$$\rho_{\mathrm{C-O}} : \text{Distance}$$
$$\theta_{\mathrm{C-O}} : \text{Polar angle}$$
$$\phi_{\mathrm{C-O}} : \text{Azimuthal angle}$$

4. **C to $O_H$ (Hydroxyl Oxygen)**:

$$\rho_{\mathrm{C-OH}} : \text{Distance}$$
$$\theta_{\mathrm{C-OH}} : \text{Polar angle}$$
$$\phi_{\mathrm{C-OH}} : \text{Azimuthal angle}$$

5. **O to $\mathbf{H_{OH}}$ (Hydroxyl Hydrogen)**:

$$\rho_{\mathrm{O-HOH}} : \text{Distance}$$
$$\theta_{\mathrm{O-HOH}} : \text{Polar angle}$$
$$\phi_{\mathrm{O-HOH}} : \text{Azimuthal angle}$$

6. **C$\alpha$ to C$\beta$**:

$$\rho_{\mathrm{C}\alpha-\mathrm{C}\beta} : \text{Distance}$$
$$\theta_{\mathrm{C}\alpha-\mathrm{C}\beta} : \text{Polar angle}$$
$$\phi_{\mathrm{C}\alpha-\mathrm{C}\beta} : \text{Azimuthal angle}$$

7. **C$\beta$ to H$\beta_1$**:

$$\rho_{\mathrm{C}\beta-\mathrm{H}\beta_1} : \text{Distance}$$
$$\theta_{\mathrm{C}\beta-\mathrm{H}\beta_1} : \text{Polar angle}$$
$$\phi_{\mathrm{C}\beta-\mathrm{H}\beta_1} : \text{Azimuthal angle}$$

8. **C$\beta$ to H$\beta_2$**:

$$\rho_{\mathrm{C}\beta-\mathrm{H}\beta_2} : \text{Distance}$$
$$\theta_{\mathrm{C}\beta-\mathrm{H}\beta_2} : \text{Polar angle}$$
$$\phi_{\mathrm{C}\beta-\mathrm{H}\beta_2} : \text{Azimuthal angle}$$

9. **C$\beta$ to O$\gamma$**:

$$\rho_{\mathrm{C}\beta-\mathrm{O}\gamma} : \text{Distance}$$
$$\theta_{\mathrm{C}\beta-\mathrm{O}\gamma} : \text{Polar angle}$$
$$\phi_{\mathrm{C}\beta-\mathrm{O}\gamma} : \text{Azimuthal angle}$$

10. **O$\gamma$ to H$\gamma$**:

$$\rho_{\mathrm{O}\gamma-\mathrm{H}\gamma} : \text{Distance}$$
$$\theta_{\mathrm{O}\gamma-\mathrm{H}\gamma} : \text{Polar angle}$$
$$\phi_{\mathrm{O}\gamma-\mathrm{H}\gamma} : \text{Azimuthal angle}$$

11. **N to $\mathbf{H_{N1}}$**:

$$\rho_{\mathrm{N-H_{N1}}} : \text{Distance}$$
$$\theta_{\mathrm{N-H_{N1}}} : \text{Polar angle}$$
$$\phi_{\mathrm{N-H_{N1}}} : \text{Azimuthal angle}$$

12. **N to $\mathbf{H_{N2}}$**:

$$\rho_{\mathrm{N-H_{N2}}} : \text{Distance}$$
$$\theta_{\mathrm{N-H_{N2}}} : \text{Polar angle}$$
$$\phi_{\mathrm{N-H_{N2}}} : \text{Azimuthal angle}$$

13. **C$\alpha$ to H$\alpha$**:

$$\rho_{\mathrm{C}\alpha-\mathrm{H}\alpha} : \text{Distance}$$
$$\theta_{\mathrm{C}\alpha-\mathrm{H}\alpha} : \text{Polar angle}$$
$$\phi_{\mathrm{C}\alpha-\mathrm{H}\alpha} : \text{Azimuthal angle}$$

### 17 An ABN-SCS Description with ρ, θ and ϕ for threonine

#### Atoms in Threonine (Thr, T)

1. **Backbone Atoms:**
   - N (Amine nitrogen)
   - H (Hydrogen attached to N)
   - C$\alpha$ (Alpha carbon)
   - H$\alpha$ (Hydrogen attached to C$\alpha$)
   - C (Carbonyl carbon)
   - O (Carbonyl oxygen)
   - $O^-$ (Carbonyl oxygen in the ionized state)
2. **Side Chain Atoms:**
   - C$\beta$ (Beta carbon)
   - H$\beta$ (Hydrogen attached to C$\beta$)
   - O$\gamma$ (Gamma oxygen, hydroxyl group)
   - H$\gamma$ (Hydrogen attached to O$\gamma$)
   - C$\gamma$ (Gamma carbon, methyl group)
   - H$\gamma_1$ (First hydrogen attached to C$\gamma$)
   - H$\gamma_2$ (Second hydrogen attached to C$\gamma$)
   - H$\gamma_3$ (Third hydrogen attached to C$\gamma$)

#### An ABN-SCS definition of the atomic bonding network of Thr

1. **N to C$\alpha$**:

$$\rho_{\mathrm{N-C}\alpha} : \text{Distance}$$
$$\theta_{\mathrm{N-C}\alpha} : \text{Polar angle}$$
$$\phi_{\mathrm{N-C}\alpha} : \text{Azimuthal angle}$$

2. **C$\alpha$ to C**:

$$\rho_{\mathrm{C}\alpha-\mathrm{C}} : \text{Distance}$$
$$\theta_{\mathrm{C}\alpha-\mathrm{C}} : \text{Polar angle}$$
$$\phi_{\mathrm{C}\alpha-\mathrm{C}} : \text{Azimuthal angle}$$

3. **C to O (Carbonyl Oxygen)**:

$$\rho_{\mathrm{C-O}} : \text{Distance}$$
$$\theta_{\mathrm{C-O}} : \text{Polar angle}$$
$$\phi_{\mathrm{C-O}} : \text{Azimuthal angle}$$

4. **C to $\mathbf{O_H}$ (Hydroxyl Oxygen)**:

$$\rho_{C-OH} : \text{Distance}$$
$$\theta_{C-OH} : \text{Polar angle}$$
$$\phi_{C-OH} : \text{Azimuthal angle}$$

5. **O to $\mathbf{H_{OH}}$ (Hydroxyl Hydrogen)**:

$$\rho_{O-HOH} : \text{Distance}$$
$$\theta_{O-HOH} : \text{Polar angle}$$
$$\phi_{O-HOH} : \text{Azimuthal angle}$$

6. **C$\alpha$ to C$\beta$**:

$$\rho_{C\alpha-C\beta} : \text{Distance}$$
$$\theta_{C\alpha-C\beta} : \text{Polar angle}$$
$$\phi_{C\alpha-C\beta} : \text{Azimuthal angle}$$

7. **C$\beta$ to O$\gamma$**:

$$\rho_{C\beta-O\gamma} : \text{Distance}$$
$$\theta_{C\beta-O\gamma} : \text{Polar angle}$$
$$\phi_{C\beta-O\gamma} : \text{Azimuthal angle}$$

8. **O$\gamma$ to H$\gamma$**:

$$\rho_{O\gamma-H\gamma} : \text{Distance}$$
$$\theta_{O\gamma-H\gamma} : \text{Polar angle}$$
$$\phi_{O\gamma-H\gamma} : \text{Azimuthal angle}$$

9. **C$\beta$ to C$\gamma$**:

$$\rho_{C\beta-C\gamma} : \text{Distance}$$
$$\theta_{C\beta-C\gamma} : \text{Polar angle}$$
$$\phi_{C\beta-C\gamma} : \text{Azimuthal angle}$$

10. **C$\gamma$ to H$\gamma_1$**:

$$\rho_{C\gamma-H\gamma_1} : \text{Distance}$$
$$\theta_{C\gamma-H\gamma_1} : \text{Polar angle}$$
$$\phi_{C\gamma-H\gamma_1} : \text{Azimuthal angle}$$

11. **C$\gamma$ to H$\gamma_2$**:

$$\rho_{C\gamma-H\gamma_2} : \text{Distance}$$
$$\theta_{C\gamma-H\gamma_2} : \text{Polar angle}$$
$$\phi_{C\gamma-H\gamma_2} : \text{Azimuthal angle}$$

12. $\mathbf{C\gamma}$ **to** $\mathbf{H\gamma_3}$:

$$\rho_{\mathrm{C}\gamma-\mathrm{H}\gamma_3} : \text{Distance}$$
$$\theta_{\mathrm{C}\gamma-\mathrm{H}\gamma_3} : \text{Polar angle}$$
$$\phi_{\mathrm{C}\gamma-\mathrm{H}\gamma_3} : \text{Azimuthal angle}$$

13. **N to** $\mathbf{H_{N1}}$:

$$\rho_{\mathrm{N}-\mathrm{H}_{\mathrm{N1}}} : \text{Distance}$$
$$\theta_{\mathrm{N}-\mathrm{H}_{\mathrm{N1}}} : \text{Polar angle}$$
$$\phi_{\mathrm{N}-\mathrm{H}_{\mathrm{N1}}} : \text{Azimuthal angle}$$

14. **N to** $\mathbf{H_{N2}}$:

$$\rho_{\mathrm{N}-\mathrm{H}_{\mathrm{N2}}} : \text{Distance}$$
$$\theta_{\mathrm{N}-\mathrm{H}_{\mathrm{N2}}} : \text{Polar angle}$$
$$\phi_{\mathrm{N}-\mathrm{H}_{\mathrm{N2}}} : \text{Azimuthal angle}$$

15. $\mathbf{C\alpha}$ **to** $\mathbf{H\alpha}$:

$$\rho_{\mathrm{C}\alpha-\mathrm{H}\alpha} : \text{Distance}$$
$$\theta_{\mathrm{C}\alpha-\mathrm{H}\alpha} : \text{Polar angle}$$
$$\phi_{\mathrm{C}\alpha-\mathrm{H}\alpha} : \text{Azimuthal angle}$$

### 18 An ABN-SCS Description with $\rho$, $\theta$ and $\phi$ for tryptophan

#### Atoms in Tryptophan (Trp, W)

1. **Backbone Atoms:**
   - N (Amine nitrogen)
   - H (Hydrogen attached to N)
   - $C\alpha$ (Alpha carbon)
   - $H\alpha$ (Hydrogen attached to $C\alpha$)
   - C (Carbonyl carbon)
   - O (Carbonyl oxygen)
   - $O^-$ (Carbonyl oxygen in the ionized state)

2. **Side Chain Atoms:**
   - $C\beta$ (Beta carbon)
   - $H\beta$ (Hydrogen attached to $C\beta$)
   - $C\gamma$ (Gamma carbon)
   - $C\delta_1$ (Delta-1 carbon)
   - $C\delta_2$ (Delta-2 carbon)
   - $H\delta_1$ (Hydrogen attached to $C\delta_1$)
   - $N\epsilon$ (Epsilon nitrogen)
   - $H\epsilon$ (Hydrogen attached to $N\epsilon$)
   - $C\epsilon_2$ (Epsilon-2 carbon)
   - $C\epsilon_3$ (Epsilon-3 carbon)
   - $H\epsilon_3$ (Hydrogen attached to $C\epsilon_3$)
   - $C\zeta_2$ (Zeta-2 carbon)
   - $C\zeta_3$ (Zeta-3 carbon)
   - $H\zeta_2$ (Hydrogen attached to $C\zeta_2$)
   - $H\zeta_3$ (Hydrogen attached to $C\zeta_3$)
   - $C\eta_2$ (Eta-2 carbon)
   - $H\eta_2$ (Hydrogen attached to $C\eta_2$)

#### An ABN-SCS definition of the atomic bonding network of Trp

1. **N to $C\alpha$**:

$$\rho_{N-C\alpha} : \text{Distance}$$
$$\theta_{N-C\alpha} : \text{Polar angle}$$
$$\phi_{N-C\alpha} : \text{Azimuthal angle}$$

2. **C$\alpha$ to C**:

$$\rho_{C\alpha-C} : \text{Distance}$$
$$\theta_{C\alpha-C} : \text{Polar angle}$$
$$\phi_{C\alpha-C} : \text{Azimuthal angle}$$

3. **C to O (Carbonyl Oxygen)**:

$$\rho_{C-O} : \text{Distance}$$
$$\theta_{C-O} : \text{Polar angle}$$
$$\phi_{C-O} : \text{Azimuthal angle}$$

4. **C to $O_H$ (Hydroxyl Oxygen)**:

$$\rho_{C-OH} : \text{Distance}$$
$$\theta_{C-OH} : \text{Polar angle}$$
$$\phi_{C-OH} : \text{Azimuthal angle}$$

5. **O to $H_{OH}$ (Hydroxyl Hydrogen)**:

$$\rho_{O-HOH} : \text{Distance}$$
$$\theta_{O-HOH} : \text{Polar angle}$$
$$\phi_{O-HOH} : \text{Azimuthal angle}$$

6. **C$\alpha$ to C$\beta$**:

$$\rho_{C\alpha-C\beta} : \text{Distance}$$
$$\theta_{C\alpha-C\beta} : \text{Polar angle}$$
$$\phi_{C\alpha-C\beta} : \text{Azimuthal angle}$$

7. **C$\beta$ to H$\beta_1$**:

$$\rho_{C\beta-H\beta_1} : \text{Distance}$$
$$\theta_{C\beta-H\beta_1} : \text{Polar angle}$$
$$\phi_{C\beta-H\beta_1} : \text{Azimuthal angle}$$

8. **C$\beta$ to H$\beta_2$**:

$$\rho_{C\beta-H\beta_2} : \text{Distance}$$
$$\theta_{C\beta-H\beta_2} : \text{Polar angle}$$
$$\phi_{C\beta-H\beta_2} : \text{Azimuthal angle}$$

9. **C$\beta$ to C$\gamma$**:

$$\rho_{C\beta-C\gamma} : \text{Distance}$$
$$\theta_{C\beta-C\gamma} : \text{Polar angle}$$
$$\phi_{C\beta-C\gamma} : \text{Azimuthal angle}$$

10. **C$\gamma$ to C$\delta_1$**:

$\rho_{C\gamma-C\delta_1}$ : Distance
$\theta_{C\gamma-C\delta_1}$ : Polar angle
$\phi_{C\gamma-C\delta_1}$ : Azimuthal angle

11. **C$\gamma$ to C$\delta_2$**:

$\rho_{C\gamma-C\delta_2}$ : Distance
$\theta_{C\gamma-C\delta_2}$ : Polar angle
$\phi_{C\gamma-C\delta_2}$ : Azimuthal angle

12. **C$\delta_1$ to H$\delta_1$**:

$\rho_{C\delta_1-H\delta_1}$ : Distance
$\theta_{C\delta_1-H\delta_1}$ : Polar angle
$\phi_{C\delta_1-H\delta_1}$ : Azimuthal angle

13. **C$\delta_1$ to N$\epsilon$**:

$\rho_{C\delta_1-N\epsilon}$ : Distance
$\theta_{C\delta_1-N\epsilon}$ : Polar angle
$\phi_{C\delta_1-N\epsilon}$ : Azimuthal angle

14. **N$\epsilon$ to H$\epsilon$**:

$\rho_{N\epsilon-H\epsilon}$ : Distance
$\theta_{N\epsilon-H\epsilon}$ : Polar angle
$\phi_{N\epsilon-H\epsilon}$ : Azimuthal angle

15. **N$\epsilon$ to C$\epsilon_2$**:

$\rho_{N\epsilon-C\epsilon_2}$ : Distance
$\theta_{N\epsilon-C\epsilon_2}$ : Polar angle
$\phi_{N\epsilon-C\epsilon_2}$ : Azimuthal angle

16. **C$\delta_2$ to C$\epsilon_3$**:

$\rho_{C\delta_2-C\epsilon_3}$ : Distance
$\theta_{C\delta_2-C\epsilon_3}$ : Polar angle
$\phi_{C\delta_2-C\epsilon_3}$ : Azimuthal angle

17. **C$\epsilon_3$ to H$\epsilon_3$**:

$\rho_{C\epsilon_3-H\epsilon_3}$ : Distance
$\theta_{C\epsilon_3-H\epsilon_3}$ : Polar angle
$\phi_{C\epsilon_3-H\epsilon_3}$ : Azimuthal angle

18. **C$\epsilon_3$ to C$\zeta_3$**:

$$\rho_{C\epsilon_3-C\zeta_3} : \text{Distance}$$
$$\theta_{C\epsilon_3-C\zeta_3} : \text{Polar angle}$$
$$\phi_{C\epsilon_3-Z\zeta_3} : \text{Azimuthal angle}$$

19. **C$\zeta_3$ to H$\zeta_3$**:

$$\rho_{C\zeta_3-H\zeta_3} : \text{Distance}$$
$$\theta_{C\zeta_3-H\zeta_3} : \text{Polar angle}$$
$$\phi_{C\zeta_3-H\zeta_3} : \text{Azimuthal angle}$$

20. **C$\zeta_3$ to C$\eta_2$**:

$$\rho_{C\zeta_3-C\eta_2} : \text{Distance}$$
$$\theta_{C\zeta_3-C\eta_2} : \text{Polar angle}$$
$$\phi_{C\zeta_3-C\eta_2} : \text{Azimuthal angle}$$

21. **C$\eta_2$ to H$\eta_2$**:

$$\rho_{C\eta_2-H\eta_2} : \text{Distance}$$
$$\theta_{C\eta_2-H\eta_2} : \text{Polar angle}$$
$$\phi_{C\eta_2-H\eta_2} : \text{Azimuthal angle}$$

22. **C$\epsilon_2$ to C$\zeta_2$**:

$$\rho_{C\epsilon_2-C\zeta_2} : \text{Distance}$$
$$\theta_{C\epsilon_2-C\zeta_2} : \text{Polar angle}$$
$$\phi_{C\epsilon_2-Z\zeta_2} : \text{Azimuthal angle}$$

23. **C$\zeta_2$ to H$\zeta_2$**:

$$\rho_{C\zeta_2-H\zeta_2} : \text{Distance}$$
$$\theta_{C\zeta_2-H\zeta_2} : \text{Polar angle}$$
$$\phi_{C\zeta_2-H\zeta_2} : \text{Azimuthal angle}$$

24. **C$\zeta_2$ to C$\eta_2$**:

$$\rho_{C\zeta_2-C\eta_2} : \text{Distance}$$
$$\theta_{C\zeta_2-C\eta_2} : \text{Polar angle}$$
$$\phi_{C\zeta_2-C\eta_2} : \text{Azimuthal angle}$$

25. **N to $\mathbf{H_{N1}}$**:

$$\rho_{N-H_{N1}} : \text{Distance}$$
$$\theta_{N-H_{N1}} : \text{Polar angle}$$
$$\phi_{N-H_{N1}} : \text{Azimuthal angle}$$

26. **N to $\mathbf{H_{N2}}$**:

$$\rho_{\mathrm{N-H_{N2}}} : \text{Distance}$$
$$\theta_{\mathrm{N-H_{N2}}} : \text{Polar angle}$$
$$\phi_{\mathrm{N-H_{N2}}} : \text{Azimuthal angle}$$

27. **C$\alpha$ to H$\alpha$**:

$$\rho_{\mathrm{C\alpha-H\alpha}} : \text{Distance}$$
$$\theta_{\mathrm{C\alpha-H\alpha}} : \text{Polar angle}$$
$$\phi_{\mathrm{C\alpha-H\alpha}} : \text{Azimuthal angle}$$

### 19 An ABN-SCS Description with $\rho$, $\theta$ and $\phi$ for tyrosine

#### Atoms in Tyrosine (Tyr, Y)

1. **Backbone Atoms:**
   - N (Amine nitrogen)
   - H (Hydrogen attached to N)
   - $C\alpha$ (Alpha carbon)
   - $H\alpha$ (Hydrogen attached to $C\alpha$)
   - C (Carbonyl carbon)
   - O (Carbonyl oxygen)
   - $O^-$ (Carbonyl oxygen in the ionized state)
2. **Side Chain Atoms:**
   - $C\beta$ (Beta carbon)
   - $H\beta_1$ (First hydrogen attached to $C\beta$)
   - $H\beta_2$ (Second hydrogen attached to $C\beta$)
   - $C\gamma$ (Gamma carbon)
   - $C\delta_1$ (First delta carbon in the benzene ring)
   - $H\delta_1$ (Hydrogen attached to $C\delta_1$)
   - $C\delta_2$ (Second delta carbon in the benzene ring)
   - $H\delta_2$ (Hydrogen attached to $C\delta_2$)
   - $C\epsilon_1$ (First epsilon carbon in the benzene ring)
   - $H\epsilon_1$ (Hydrogen attached to $C\epsilon_1$)
   - $C\epsilon_2$ (Second epsilon carbon in the benzene ring)
   - $H\epsilon_2$ (Hydrogen attached to $C\epsilon_2$)
   - $C\zeta$ (Zeta carbon)
   - $O\zeta$ (Hydroxyl oxygen attached to $C\zeta$)
   - $H\zeta$ (Hydroxyl hydrogen attached to $O\zeta$)

#### An ABN-SCS definition of the atomic bonding network of Tyr

1. **N to $C\alpha$**:

$$\rho_{N-C\alpha} : \text{Distance}$$
$$\theta_{N-C\alpha} : \text{Polar angle}$$
$$\phi_{N-C\alpha} : \text{Azimuthal angle}$$

2. **$C\alpha$ to C**:

$$\rho_{C\alpha-C} : \text{Distance}$$
$$\theta_{C\alpha-C} : \text{Polar angle}$$
$$\phi_{C\alpha-C} : \text{Azimuthal angle}$$

3. **C to O (Carbonyl Oxygen)**:

$$\rho_{C-O}: \text{Distance}$$
$$\theta_{C-O}: \text{Polar angle}$$
$$\phi_{C-O}: \text{Azimuthal angle}$$

4. **C to $\mathbf{O_H}$ (Hydroxyl Oxygen)**:

$$\rho_{C-OH}: \text{Distance}$$
$$\theta_{C-OH}: \text{Polar angle}$$
$$\phi_{C-OH}: \text{Azimuthal angle}$$

5. **O to $\mathbf{H_{OH}}$ (Hydroxyl Hydrogen)**:

$$\rho_{O-HOH}: \text{Distance}$$
$$\theta_{O-HOH}: \text{Polar angle}$$
$$\phi_{O-HOH}: \text{Azimuthal angle}$$

6. **C$\alpha$ to C$\beta$**:

$$\rho_{C\alpha-C\beta}: \text{Distance}$$
$$\theta_{C\alpha-C\beta}: \text{Polar angle}$$
$$\phi_{C\alpha-C\beta}: \text{Azimuthal angle}$$

7. **C$\beta$ to C$\gamma$**:

$$\rho_{C\beta-C\gamma}: \text{Distance}$$
$$\theta_{C\beta-C\gamma}: \text{Polar angle}$$
$$\phi_{C\beta-C\gamma}: \text{Azimuthal angle}$$

8. **C$\gamma$ to C$\delta_1$**:

$$\rho_{C\gamma-C\delta_1}: \text{Distance}$$
$$\theta_{C\gamma-C\delta_1}: \text{Polar angle}$$
$$\phi_{C\gamma-C\delta_1}: \text{Azimuthal angle}$$

9. **C$\gamma$ to C$\delta_2$**:

$$\rho_{C\gamma-C\delta_2}: \text{Distance}$$
$$\theta_{C\gamma-C\delta_2}: \text{Polar angle}$$
$$\phi_{C\gamma-C\delta_2}: \text{Azimuthal angle}$$

10. **C$\delta_1$ to H$\delta_1$**:

$$\rho_{C\delta_1-H\delta_1}: \text{Distance}$$
$$\theta_{C\delta_1-H\delta_1}: \text{Polar angle}$$
$$\phi_{C\delta_1-H\delta_1}: \text{Azimuthal angle}$$

11. **C$\delta_2$ to H$\delta_2$:**

$\rho_{C\delta_2 - H\delta_2}$ : Distance
$\theta_{C\delta_2 - H\delta_2}$ : Polar angle
$\phi_{C\delta_2 - H\delta_2}$ : Azimuthal angle

12. **C$\delta_1$ to C$\epsilon_1$:**

$\rho_{C\delta_1 - C\epsilon_1}$ : Distance
$\theta_{C\delta_1 - C\epsilon_1}$ : Polar angle
$\phi_{C\delta_1 - C\epsilon_1}$ : Azimuthal angle

13. **C$\delta_2$ to C$\epsilon_2$:**

$\rho_{C\delta_2 - C\epsilon_2}$ : Distance
$\theta_{C\delta_2 - C\epsilon_2}$ : Polar angle
$\phi_{C\delta_2 - C\epsilon_2}$ : Azimuthal angle

14. **C$\epsilon_1$ to H$\epsilon_1$:**

$\rho_{C\epsilon_1 - H\epsilon_1}$ : Distance
$\theta_{C\epsilon_1 - H\epsilon_1}$ : Polar angle
$\phi_{C\epsilon_1 - H\epsilon_1}$ : Azimuthal angle

15. **C$\epsilon_2$ to H$\epsilon_2$:**

$\rho_{C\epsilon_2 - H\epsilon_2}$ : Distance
$\theta_{C\epsilon_2 - H\epsilon_2}$ : Polar angle
$\phi_{C\epsilon_2 - H\epsilon_2}$ : Azimuthal angle

16. **C$\epsilon_1$ to C$\zeta$:**

$\rho_{C\epsilon_1 - C\zeta}$ : Distance
$\theta_{C\epsilon_1 - C\zeta}$ : Polar angle
$\phi_{C\epsilon_1 - C\zeta}$ : Azimuthal angle

17. **C$\epsilon_2$ to C$\zeta$:**

$\rho_{C\epsilon_2 - C\zeta}$ : Distance
$\theta_{C\epsilon_2 - C\zeta}$ : Polar angle
$\phi_{C\epsilon_2 - C\zeta}$ : Azimuthal angle

18. **C$\zeta$ to O$\zeta$:**

$\rho_{C\zeta - O\zeta}$ : Distance
$\theta_{C\zeta - O\zeta}$ : Polar angle
$\phi_{C\zeta - O\zeta}$ : Azimuthal angle

19. $\mathbf{O\zeta}$ **to** $\mathbf{H\zeta}$:

$$\rho_{O\zeta-H\zeta} : \text{Distance}$$
$$\theta_{O\zeta-H\zeta} : \text{Polar angle}$$
$$\phi_{O\zeta-H\zeta} : \text{Azimuthal angle}$$

20. **N to** $\mathbf{H_{N1}}$:

$$\rho_{N-H_{N1}} : \text{Distance}$$
$$\theta_{N-H_{N1}} : \text{Polar angle}$$
$$\phi_{N-H_{N1}} : \text{Azimuthal angle}$$

21. **N to** $\mathbf{H_{N2}}$:

$$\rho_{N-H_{N2}} : \text{Distance}$$
$$\theta_{N-H_{N2}} : \text{Polar angle}$$
$$\phi_{N-H_{N2}} : \text{Azimuthal angle}$$

22. $\mathbf{C\alpha}$ **to** $\mathbf{H\alpha}$:

$$\rho_{C\alpha-H\alpha} : \text{Distance}$$
$$\theta_{C\alpha-H\alpha} : \text{Polar angle}$$
$$\phi_{C\alpha-H\alpha} : \text{Azimuthal angle}$$

### 20 An ABN-SCS Description with ρ, θ and ϕ for valine

#### Atoms in Valine (Val, V)

1. **Backbone Atoms:**
   - N (Amine nitrogen)
   - H (Hydrogen attached to N)
   - C$\alpha$ (Alpha carbon)
   - H$\alpha$ (Hydrogen attached to C$\alpha$)
   - C (Carbonyl carbon)
   - O (Carbonyl oxygen)
   - $O^-$ (Carbonyl oxygen in the ionized state)
2. **Side Chain Atoms:**
   - C$\beta$ (Beta carbon)
   - H$\beta$ (Hydrogen attached to C$\beta$)
   - C$\gamma_1$ (First gamma carbon)
   - H$\gamma_{1,1}$ (First hydrogen attached to C$\gamma_1$)
   - H$\gamma_{1,2}$ (Second hydrogen attached to C$\gamma_1$)
   - H$\gamma_{1,3}$ (Third hydrogen attached to C$\gamma_1$)
   - C$\gamma_2$ (Second gamma carbon)
   - H$\gamma_{2,1}$ (First hydrogen attached to C$\gamma_2$)
   - H$\gamma_{2,2}$ (Second hydrogen attached to C$\gamma_2$)
   - H$\gamma_{2,3}$ (Third hydrogen attached to C$\gamma_2$)

#### An ABN-SCS definition of the atomic bonding network of Val

1. **N to C$\alpha$**:

$$\rho_{\mathrm{N-C\alpha}} : \text{Distance}$$
$$\theta_{\mathrm{N-C\alpha}} : \text{Polar angle}$$
$$\phi_{\mathrm{N-C\alpha}} : \text{Azimuthal angle}$$

2. **C$\alpha$ to C**:

$$\rho_{\mathrm{C\alpha-C}} : \text{Distance}$$
$$\theta_{\mathrm{C\alpha-C}} : \text{Polar angle}$$
$$\phi_{\mathrm{C\alpha-C}} : \text{Azimuthal angle}$$

3. **C to O (Carbonyl Oxygen)**:

$$\rho_{\mathrm{C-O}} : \text{Distance}$$
$$\theta_{\mathrm{C-O}} : \text{Polar angle}$$
$$\phi_{\mathrm{C-O}} : \text{Azimuthal angle}$$

4. **C to $O_H$ (Hydroxyl Oxygen)**:

$\rho_{C-OH}$ : Distance
$\theta_{C-OH}$ : Polar angle
$\phi_{C-OH}$ : Azimuthal angle

5. **O to $H_{OH}$ (Hydroxyl Hydrogen)**:

$\rho_{O-HOH}$ : Distance
$\theta_{O-HOH}$ : Polar angle
$\phi_{O-HOH}$ : Azimuthal angle

6. **C$\alpha$ to C$\beta$**:

$\rho_{C\alpha-C\beta}$ : Distance
$\theta_{C\alpha-C\beta}$ : Polar angle
$\phi_{C\alpha-C\beta}$ : Azimuthal angle

7. **C$\beta$ to C$\gamma_1$**:

$\rho_{C\beta-C\gamma_1}$ : Distance
$\theta_{C\beta-C\gamma_1}$ : Polar angle
$\phi_{C\beta-C\gamma_1}$ : Azimuthal angle

8. **C$\beta$ to C$\gamma_2$**:

$\rho_{C\beta-C\gamma_2}$ : Distance
$\theta_{C\beta-C\gamma_2}$ : Polar angle
$\phi_{C\beta-C\gamma_2}$ : Azimuthal angle

9. **N to $H_{N1}$**:

$\rho_{N-H_{N1}}$ : Distance
$\theta_{N-H_{N1}}$ : Polar angle
$\phi_{N-H_{N1}}$ : Azimuthal angle

10. **N to $H_{N2}$**:

$\rho_{N-H_{N2}}$ : Distance
$\theta_{N-H_{N2}}$ : Polar angle
$\phi_{N-H_{N2}}$ : Azimuthal angle

11. **C$\alpha$ to H$\alpha$**:

$\rho_{C\alpha-H\alpha}$ : Distance
$\theta_{C\alpha-H\alpha}$ : Polar angle
$\phi_{C\alpha-H\alpha}$ : Azimuthal angle

12. **C$\beta$ to H$\beta$**:

$$\rho_{C\beta-H\beta} : \text{Distance}$$
$$\theta_{C\beta-H\beta} : \text{Polar angle}$$
$$\phi_{C\beta-H\beta} : \text{Azimuthal angle}$$

13. **C$\gamma_1$ to H$\gamma_{1,1}$**:

$$\rho_{C\gamma_1-H\gamma_{1,1}} : \text{Distance}$$
$$\theta_{C\gamma_1-H\gamma_{1,1}} : \text{Polar angle}$$
$$\phi_{C\gamma_1-H\gamma_{1,1}} : \text{Azimuthal angle}$$

14. **C$\gamma_1$ to H$\gamma_{1,2}$**:

$$\rho_{C\gamma_1-H\gamma_{1,2}} : \text{Distance}$$
$$\theta_{C\gamma_1-H\gamma_{1,2}} : \text{Polar angle}$$
$$\phi_{C\gamma_1-H\gamma_{1,2}} : \text{Azimuthal angle}$$

15. **C$\gamma_1$ to H$\gamma_{1,3}$**:

$$\rho_{C\gamma_1-H\gamma_{1,3}} : \text{Distance}$$
$$\theta_{C\gamma_1-H\gamma_{1,3}} : \text{Polar angle}$$
$$\phi_{C\gamma_1-H\gamma_{1,3}} : \text{Azimuthal angle}$$

16. **C$\gamma_2$ to H$\gamma_{2,1}$**:

$$\rho_{C\gamma_2-H\gamma_{2,1}} : \text{Distance}$$
$$\theta_{C\gamma_2-H\gamma_{2,1}} : \text{Polar angle}$$
$$\phi_{C\gamma_2-H\gamma_{2,1}} : \text{Azimuthal angle}$$

17. **C$\gamma_2$ to H$\gamma_{2,2}$**:

$$\rho_{C\gamma_2-H\gamma_{2,2}} : \text{Distance}$$
$$\theta_{C\gamma_2-H\gamma_{2,2}} : \text{Polar angle}$$
$$\phi_{C\gamma_2-H\gamma_{2,2}} : \text{Azimuthal angle}$$

18. **C$\gamma_2$ to H$\gamma_{2,3}$**:

$$\rho_{C\gamma_2-H\gamma_{2,3}} : \text{Distance}$$
$$\theta_{C\gamma_2-H\gamma_{2,3}} : \text{Polar angle}$$
$$\phi_{C\gamma_2-H\gamma_{2,3}} : \text{Azimuthal angle}$$
