## Supplementary file 2 for "Protein Structure Description with *ρ, θ* and *ϕ*: A Case Study with Caenopore-5"

### Protein Structure Description with $\rho$, $\theta$ and $\phi$: Supplementary Information

Wei Li *
Independent Researcher
Room 902, Building No. 41, Fourth Community, Yuhongjiayuan Village
No. 1168, Antong Road, Chenjia Town, Chongming Island
Shanghai, People's Republic of China

August 29, 2025

*

### 1 Supplementary Materials and Methods

#### 1.1 Geometric equivalence of Cartesian (CCS) and spherical (SCS) coordinate systems

By transforming the Cartesian coordinates into spherical coordinates (Equations 1, 2 and 3), this ABN-SCS approach decouples the spatial information of atoms into radial distance ($\rho$), polar angle ($\theta$), and azimuthal angle ($\phi$), where $\rho$ is defined as the equilibrium inter-atomic bond length (a physical constant), i.e., the inter-nuclear distance at which the system energy minimum occurs [1, 2],

$$\rho = \sqrt{x^2 + y^2 + z^2} \tag{1}$$

$$\theta = \arccos\left(\frac{z}{\rho}\right) \tag{2}$$

$$\phi = \arctan 2(y, x) \tag{3}$$

where:

- $\rho$ is the radial distance from the origin to the point, Figure 1.
- $\theta$ is the polar angle (angle from the positive $z$-axis, Figure 1).
- $\phi$ is the azimuthal angle (angle from the positive $x$-axis in the $xy$-plane, Figure 1).

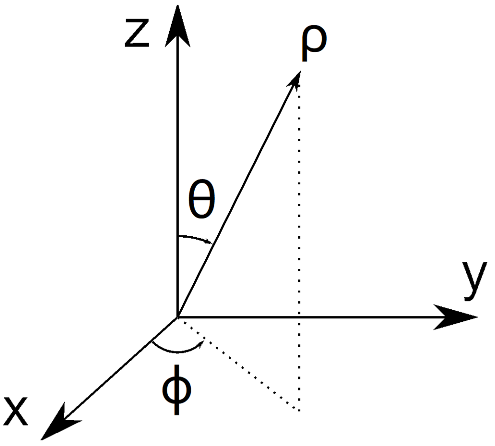


Figure 1: Geometrically, Cartesian and spherical coordinate systems are like two sides of one coin, and are both applicable in the specifications of atomic positions for protein structure description, either with [$x$, $y$, $z$] or [$\rho$, $\theta$, $\phi$].

An implementation of Equations 1, 2 and 3 is described with an in-house python script, whose source code is included in supplementary file **S3.pdf**.

#### 1.2 ABN-SCS: an alternative spherical coordinate system

As described in the main manuscript, in 2011, it was proposed for the first time that protein structure be represented in spherical coordinates ($\rho$, $\theta$ and $\phi$) [3,4], where a global spherical coordinate system (G-SCS) was employed for the separation of the protein outer layer from its inner core, and for the identification of protrusions and invaginations on the protein surface [3,4]. Thus, G-SCS also requires three parameters ($R$, $\theta$ and $\phi$), as it takes the geometric centroid (an imaginary point) of the protein molecule as origin, where $R$ is used to signify the distance between the imaginary centroid and any atom of the protein [3,4].

Here, in this manuscript, an alternative 3D spherical coordinate system, i.e., ABN-SCS, is defined to be based on the inter-atomic bonding network (ABN) for protein structure description with $\rho$, $\theta$ and $\phi$, where $\rho$ is defined as the equilibrium atomic bond length, i.e., the inter-nuclear distance at which the system energy minimum occurs [1,2].

#### 1.3 An ABN-SCS description of the chemical connectivity of twenty stand-alone natural amino acids

This section is described in detail in supplemtary file **S1.pdf**.

#### 1.4 An ABN-SCS description of protein peptide bond and backbone structures: an example of a model peptide with five amino acids

For an inter-atomic spherical coordinate system (ABN-SCS) approach to redefine protein structure with $\rho$, $\theta$ and $\phi$ [5], a short peptide with five amino acids (referred to below as **peptideX**) (Figure 2) is employed here as an example.

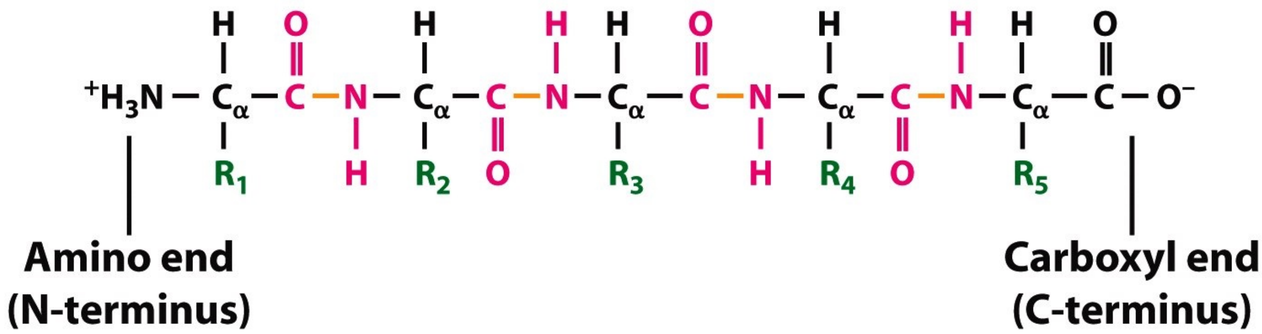


Figure 2: Chemical structure of a short model peptide with five amino acids (referred to below as **peptideX**).

Given an ABN-SCS description of the structures of twenty stand-alone natural amino acids, a logical next step is to use the ABN-SCS framework to define the formation of a peptide bond.

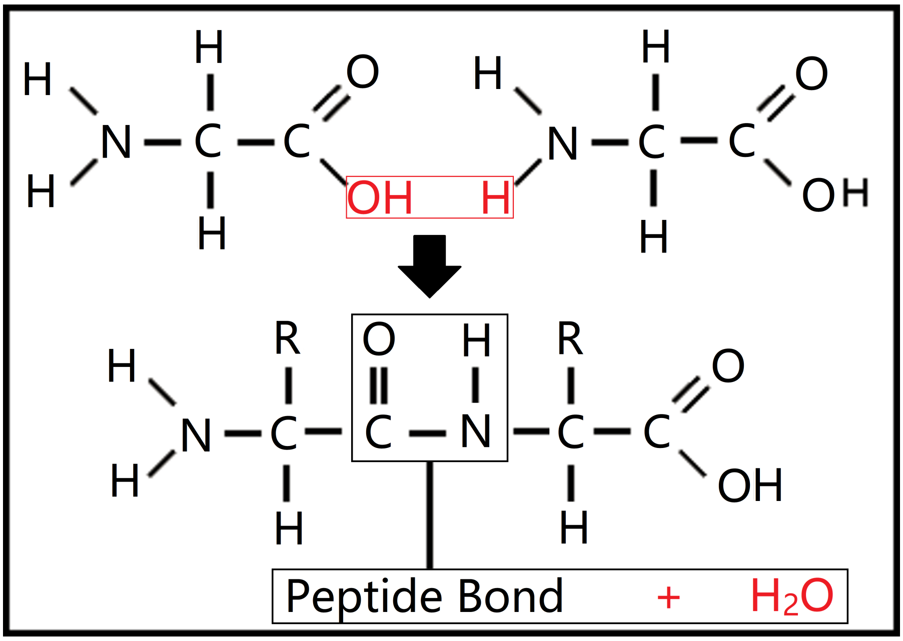


Figure 3: The formation of a peptide bond, which is the bond between the carbonyl carbon and the nitrogen in the amide fragment.

As shown in Figure 3, a peptide bond is formed by dehydration, which is the removal of the carboxyl hydroxyl on one amino acid and one hydrogen atom on the amine end of another amino acid. Specifically, the whole inter-atomic peptide bonding network of the backbone of the short peptide (Figure 2) is defined sequentially as below:

1. the peptide bond formed between $R_1$ and $R_2$ of **peptideX** (Figure 2);
2. the peptide bond formed between $R_2$ and $R_3$ of **peptideX** (Figure 2);

3. the peptide bond formed between $R_3$ and $R_4$ of **peptideX** (Figure 2);
4. the peptide bond formed between $R_4$ and $R_5$ of **peptideX** (Figure 2);

Afterwards, the whole inter-atomic covalent bonding network of the backbone of peptideX (Figure 2) is defined sequentially as below:

1. the covalent bond between N-terminal amide nitrogen atom (Atom_initial in ABN-SCS) of the first residue of peptideX (Figure 2) and three N-terminal amide hydrogen atoms (Atom_final in ABN-SCS) of the backbone of the first residue of peptideX (Figure 2);
2. the covalent bond between N-terminal amide nitrogen atom (Atom_initial in ABN-SCS) of the first residue of peptideX (Figure 2) and carbon $\alpha$ ($C_\alpha$, Atom_final in ABN-SCS) of the backbone of the first residue of peptideX (Figure 2);
3. the covalent bond between carbon $\alpha$ ($C_\alpha$, Atom_initial in ABN-SCS) of the first residue of peptideX (Figure 2) and hydrogen $\alpha$ ($H_\alpha$, Atom_final in ABN-SCS) of the backbone of the first residue of peptideX (Figure 2);
4. the covalent bond between carbon $\alpha$ ($C_\alpha$, Atom_initial in ABN-SCS) of the first residue of peptideX (Figure 2) and carbonyl carbon (Atom_final in ABN-SCS) of the backbone of the first residue of peptideX (Figure 2);
5. the covalent bond between carbonyl carbon (Atom_initial in ABN-SCS) of the first residue of peptideX (Figure 2) and carbonyl oxygen (the double-bonded oxygen atom, Atom_final in ABN-SCS) of the backbone of the first residue of peptideX (Figure 2);
6. the covalent bond between carbonyl carbon (Atom_initial in ABN-SCS) of the first residue of peptideX (Figure 2) and the amide nitrogen atom (Atom_final in ABN-SCS) of the second residue of peptideX (Figure 2);
7. the covalent bond between the amide nitrogen atom (Atom_initial in ABN-SCS) of the second residue of peptideX (Figure 2) and the amide hydrogen atom (Atom_final in ABN-SCS) of the backbone of the second residue of peptideX (Figure 2);
8. the covalent bond between N-terminal amide nitrogen atom (Atom_initial in ABN-SCS) of the second residue of peptideX (Figure 2) and carbon $\alpha$ ($C_\alpha$, Atom_final in ABN-SCS) of the backbone of the second residue of peptideX (Figure 2);
9. the covalent bond between carbon $\alpha$ ($C_\alpha$, Atom_initial in ABN-SCS) of the second residue of peptideX (Figure 2) and hydrogen $\alpha$ ($H_\alpha$, Atom_final in ABN-SCS) of the backbone of the second residue of peptideX (Figure 2);
10. the covalent bond between carbon $\alpha$ ($C_\alpha$, Atom_initial in ABN-SCS) of the second residue of peptideX (Figure 2) and carbonyl carbon (Atom_final in ABN-SCS) of the backbone of the second residue of peptideX (Figure 2);
11. the covalent bond between carbonyl carbon (Atom_initial in ABN-SCS) of the second residue of peptideX (Figure 2) and carbonyl oxygen (the double-bonded oxygen atom, Atom_final in ABN-SCS) of the backbone of the second residue of peptideX (Figure 2);
12. the covalent bond between carbonyl carbon (Atom_initial in ABN-SCS) of the second residue of peptideX (Figure 2) and the amide nitrogen atom (Atom_final in ABN-SCS) of the third residue of peptideX (Figure 2);

13. the covalent bond between the amide nitrogen atom (Atom_initial in ABN-SCS) of the third residue of peptideX (Figure 2) and the amide hydrogen atom (Atom_final in ABN-SCS) of the backbone of the third residue of peptideX (Figure 2);
14. the covalent bond between N-terminal amide nitrogen atom (Atom_initial in ABN-SCS) of the third residue of peptideX (Figure 2) and carbon α ($C_\alpha$, Atom_final in ABN-SCS) of the backbone of the third residue of peptideX (Figure 2);
15. the covalent bond between carbon α ($C_\alpha$, Atom_initial in ABN-SCS) of the third residue of peptideX (Figure 2) and hydrogen α ($H_\alpha$, Atom_final in ABN-SCS) of the backbone of the third residue of peptideX (Figure 2);
16. the covalent bond between carbon α ($C_\alpha$, Atom_initial in ABN-SCS) of the third residue of peptideX (Figure 2) and carbonyl carbon (Atom_final in ABN-SCS) of the backbone of the third residue of peptideX (Figure 2);
17. the covalent bond between carbonyl carbon (Atom_initial in ABN-SCS) of the third residue of peptideX (Figure 2) and carbonyl oxygen (the double-bonded oxygen atom, Atom_final in ABN-SCS) of the backbone of the third residue of peptideX (Figure 2);
18. the covalent bond between carbonyl carbon (Atom_initial in ABN-SCS) of the third residue of peptideX (Figure 2) and the amide nitrogen atom (Atom_final in ABN-SCS) of the fourth residue of peptideX (Figure 2);
19. the covalent bond between the amide nitrogen atom (Atom_initial in ABN-SCS) of the fourth residue of peptideX (Figure 2) and the amide hydrogen atom (Atom_final in ABN-SCS) of the backbone of the fourth residue of peptideX (Figure 2);
20. the covalent bond between N-terminal amide nitrogen atom (Atom_initial in ABN-SCS) of the fourth residue of peptideX (Figure 2) and carbon α ($C_\alpha$, Atom_final in ABN-SCS) of the backbone of the fourth residue of peptideX (Figure 2);
21. the covalent bond between carbon α ($C_\alpha$, Atom_initial in ABN-SCS) of the fourth residue of peptideX (Figure 2) and hydrogen α ($H_\alpha$, Atom_final in ABN-SCS) of the backbone of the fourth residue of peptideX (Figure 2);
22. the covalent bond between carbon α ($C_\alpha$, Atom_initial in ABN-SCS) of the fourth residue of peptideX (Figure 2) and carbonyl carbon (Atom_final in ABN-SCS) of the backbone of the fourth residue of peptideX (Figure 2);
23. the covalent bond between carbonyl carbon (Atom_initial in ABN-SCS) of the fourth residue of peptideX (Figure 2) and carbonyl oxygen (the double-bonded oxygen atom, Atom_final in ABN-SCS) of the backbone of the fourth residue of peptideX (Figure 2);
24. the covalent bond between carbonyl carbon (Atom_initial in ABN-SCS) of the fourth residue of peptideX (Figure 2) and the amide nitrogen atom (Atom_final in ABN-SCS) of the fifth residue of peptideX (Figure 2);
25. the covalent bond between the amide nitrogen atom (Atom_initial in ABN-SCS) of the fifth residue of peptideX (Figure 2) and the amide hydrogen atom (Atom_final in ABN-SCS) of the backbone of the fifth residue of peptideX (Figure 2);
26. the covalent bond between N-terminal amide nitrogen atom (Atom_initial in ABN-SCS) of the fifth residue of peptideX (Figure 2) and carbon α ($C_\alpha$, Atom_final in ABN-SCS) of the backbone of the fifth residue of peptideX (Figure 2);

27. the covalent bond between carbon α ($C_α$, Atom_initial in ABN-SCS) of the fifth residue of peptideX (Figure 2) and hydrogen α ($H_α$, Atom_final in ABN-SCS) of the backbone of the fifth residue of peptideX (Figure 2);

28. the covalent bond between carbon α ($C_α$, Atom_initial in ABN-SCS) of the fifth residue of peptideX (Figure 2) and carbonyl carbon (Atom_final in ABN-SCS) of the backbone of the fifth residue of peptideX (Figure 2);

29. the covalent double bond between carbonyl carbon (Atom_initial in ABN-SCS) of the fifth residue of peptideX (Figure 2) and carbonyl oxygen (the double-bonded oxygen atom, Atom_final in ABN-SCS) of the backbone of the fifth residue of peptideX (Figure 2);

30. the covalent single bond between carbonyl carbon (Atom_initial in ABN-SCS) of the fifth residue of peptideX (Figure 2) and the carboxyl hydroxyl oxygen atom (Atom_final in ABN-SCS) of the fifth residue of peptideX (Figure 2);

### 2 Supplementary Results

#### 2.1 Protein Structure Feature Extraction with the ABN-SCS Framework: A Case Study with Cp5

Overall, this article puts forward a reversible spherical geometric conversion method, which offers an approach for extracting essential structural features from protein backbones by transforming Cartesian coordinates ($x$, $y$, and $z$) into spherical coordinates $\rho$, $\theta$ and $\phi$. Specifically, the extraction of essential structural features from the backbone of 15 NMR structural models of Caenopore-5 [6–8] inculde:

1. a distribution pattern of the lengths ($\rho$) of 477 covalent bonds of the backbone of Caenopore-5 [6–8], as shown in Figure 4;
2. a distribution pattern of the azimuthal angle ($\theta$) of 477 covalent bonds of the backbone of Caenopore-5 [6–8], as shown in Figure 5;
3. a distribution pattern of the polar angle (also known as the zenith angle, $\phi$) of 477 covalent bonds of the backbone of Caenopore-5 [6–8], as shown in Figure 6;
4. a distribution pattern of the lengths ($\rho$) of 80 peptide bonds of Caenopore-5 [6–8], as shown in Figure 7;
5. a distribution pattern of the azimuthal angle ($\theta$) of 80 peptide bonds of Caenopore-5 [6–8], as shown in Figure 8;
6. a distribution pattern of the polar angle (also known as the zenith angle, $\phi$) of 80 peptide bonds of Caenopore-5 [6–8], as shown in Figure 9;

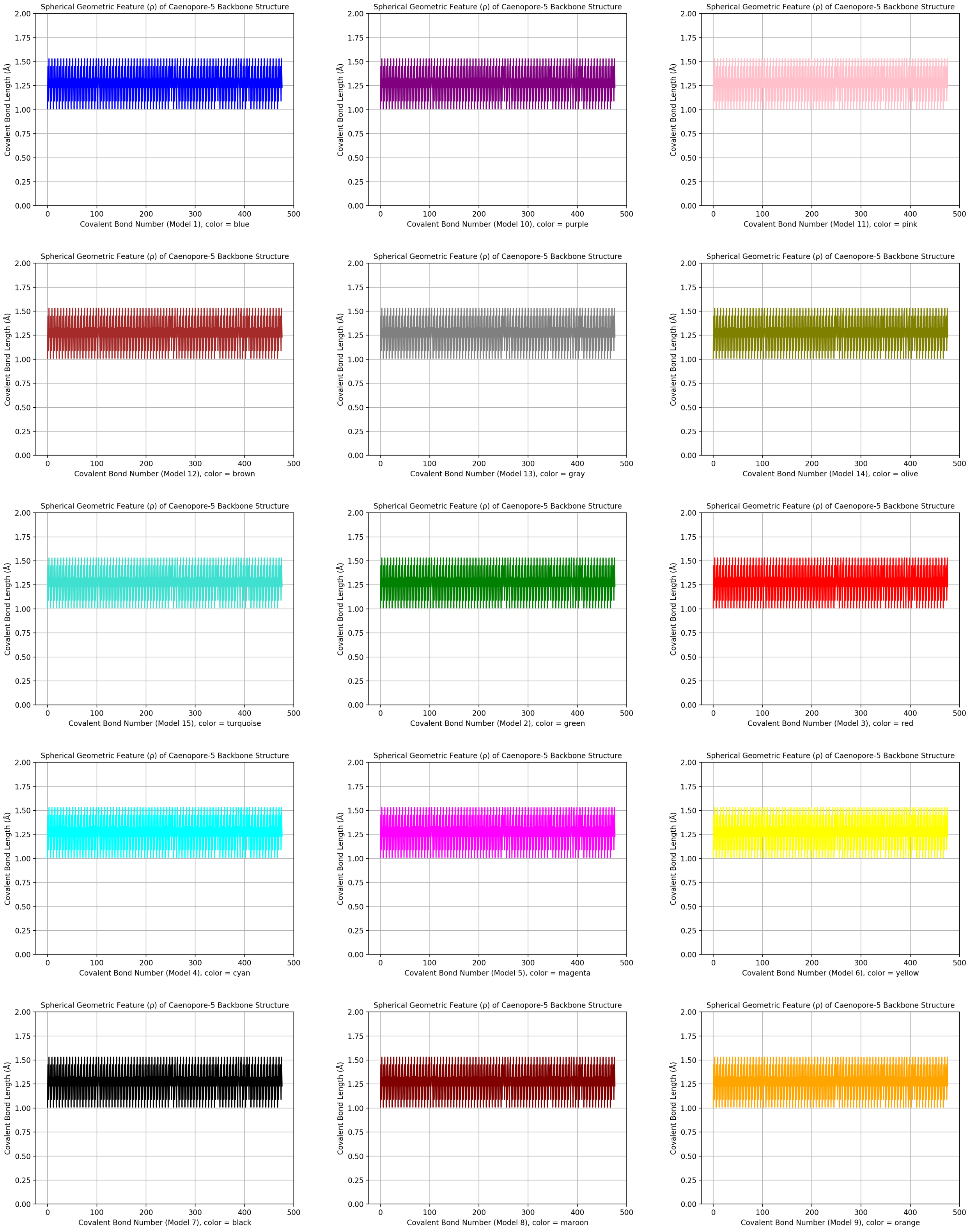


Figure 4: Distribution pattern of the lengths ($\rho$) of 477 covalent bonds of the backbone of Caenopore-5 [6–8] as reversibly extracted from the 15 NMR structural models of the three-dimensional NMR ensemble of Caenopore-5 (PDB ID: 2JSA) [6–8].

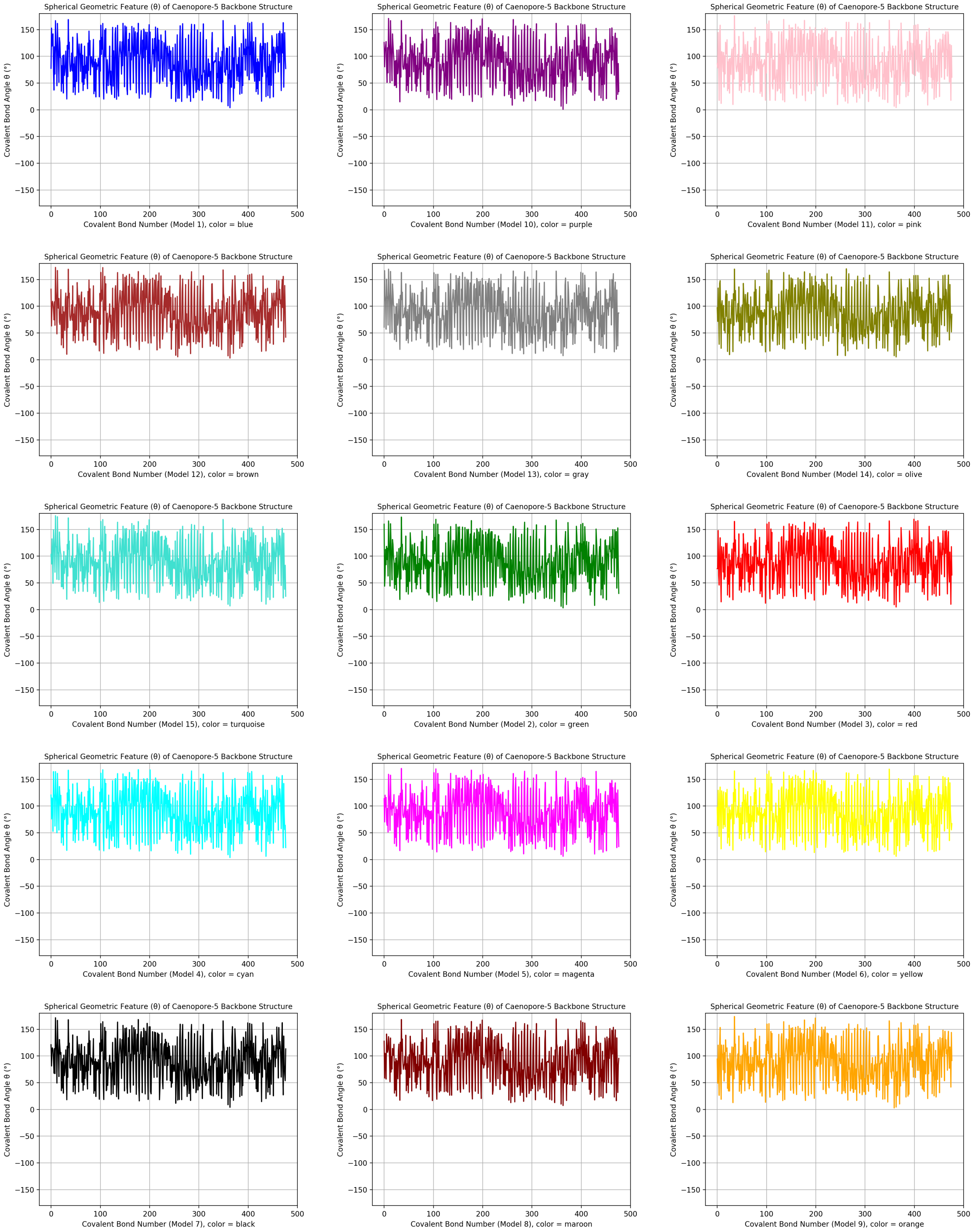


Figure 5: Distribution pattern of the azimuthal angle ($\theta$) of 477 covalent bonds of the backbone of Caenopore-5 [6–8] as reversibly extracted from the 15 NMR structural models of the three-dimensional NMR ensemble of Caenopore-5 (PDB ID: 2JSA) [6–8].

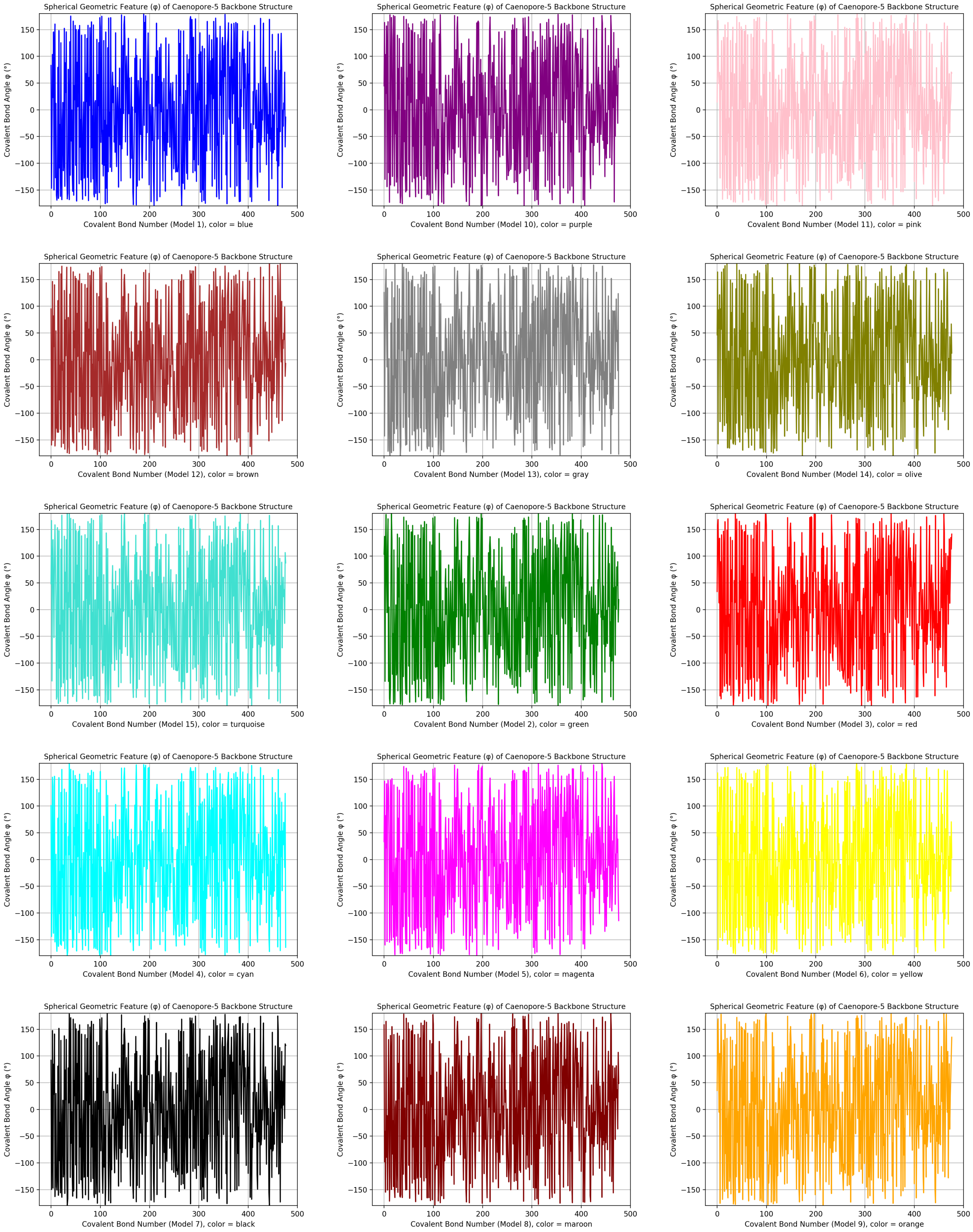


Figure 6: Distribution pattern of the polar angle (also known as the zenith angle, $\phi$) of 477 covalent bonds of the backbone of Caenopore-5 [6–8] as reversibly extracted from the 15 NMR structural models of the three-dimensional NMR ensemble of Caenopore-5 (PDB ID: 2JSA) [6–8].

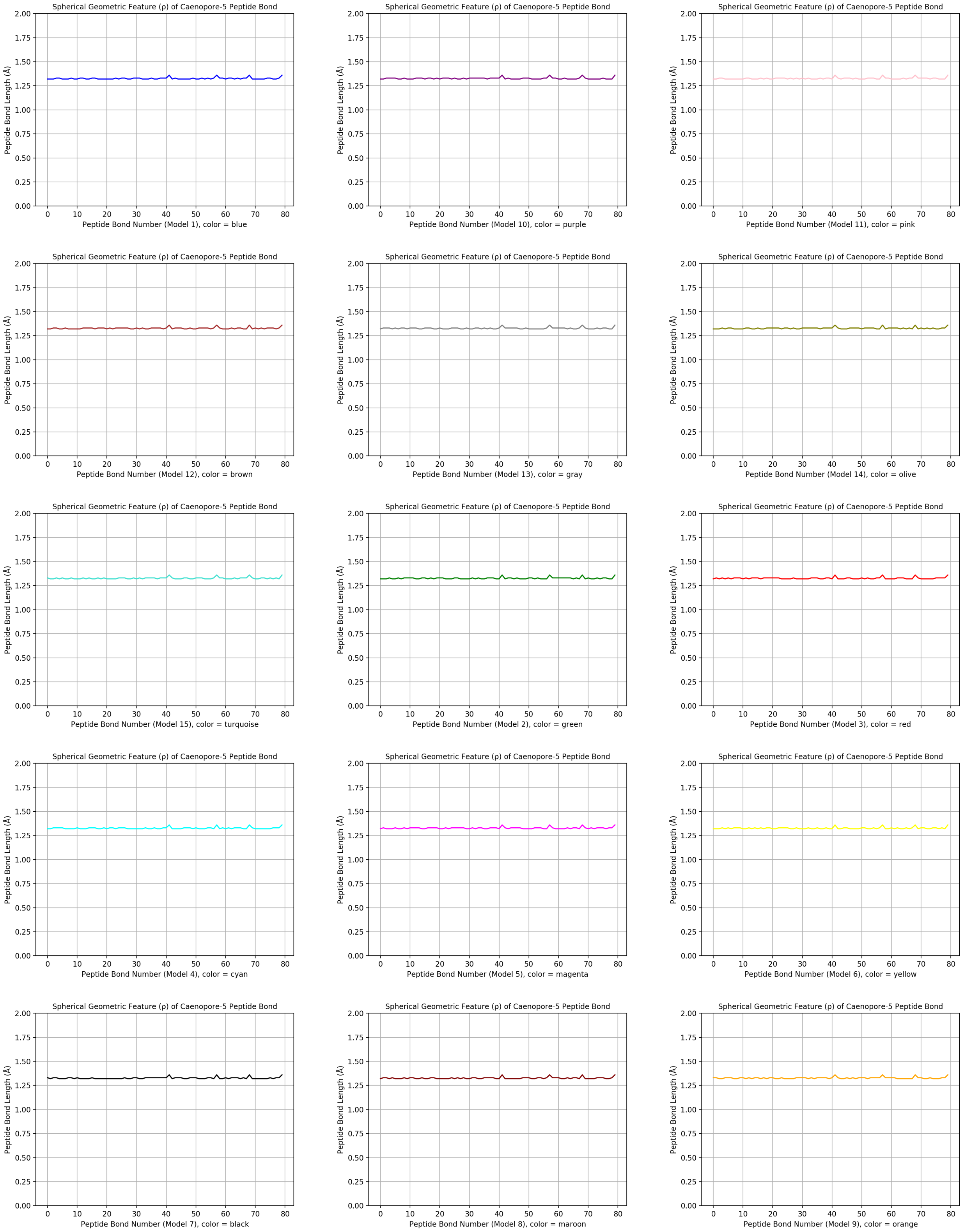


Figure 7: Distribution pattern of the lengths ($\rho$) of 80 peptide bonds of Caenopore-5 [6–8] as reversibly extracted from the 15 NMR structural models of the three-dimensional NMR ensemble of Caenopore-5 (PDB ID: 2JSA) [6–8].

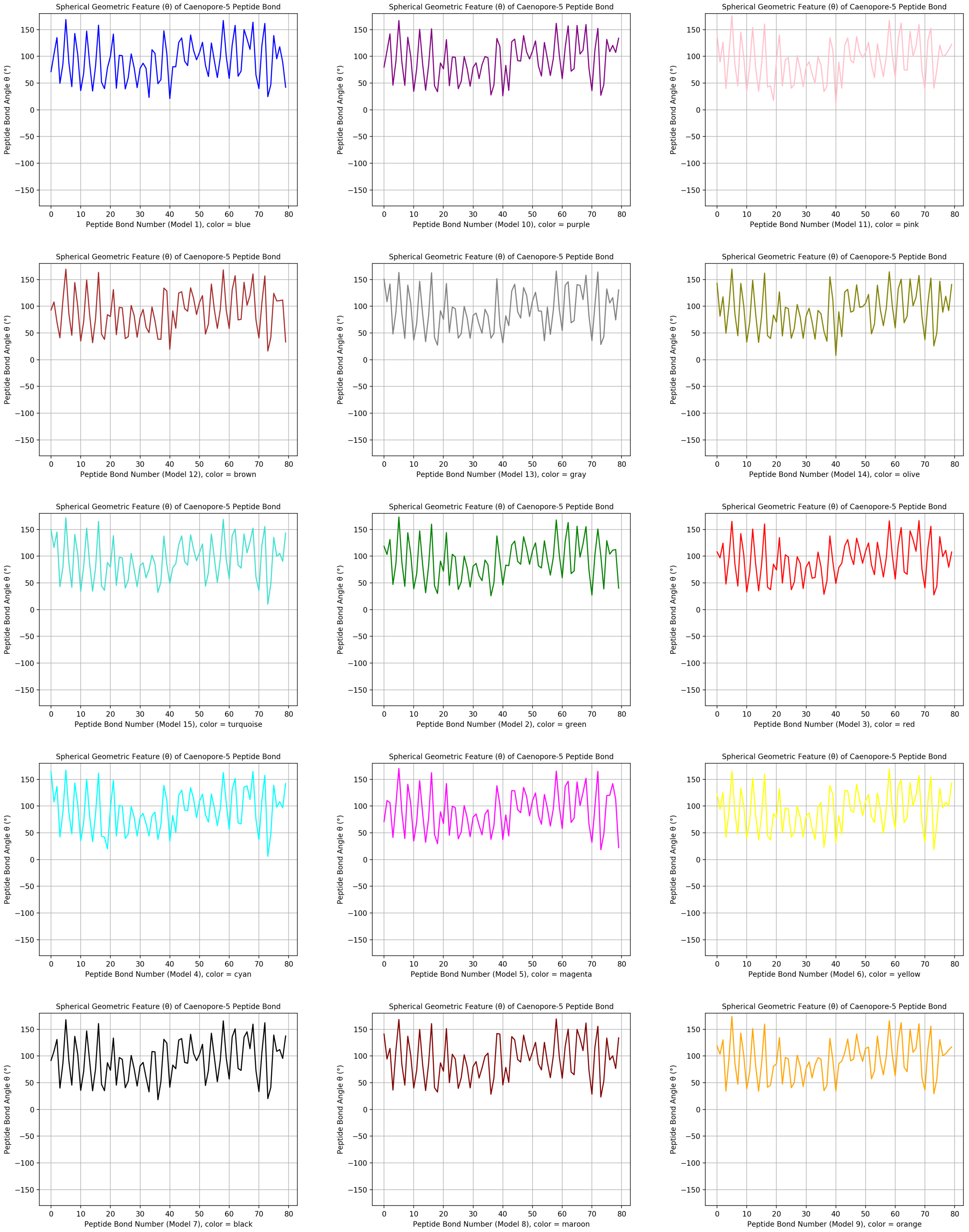


Figure 8: Distribution pattern of the azimuthal angle ($\theta$) of 80 peptide bonds of Caenopore-5 [6–8] as reversibly extracted from the 15 NMR structural models of the three-dimensional NMR ensemble of Caenopore-5 (PDB ID: 2JSA) [6–8].

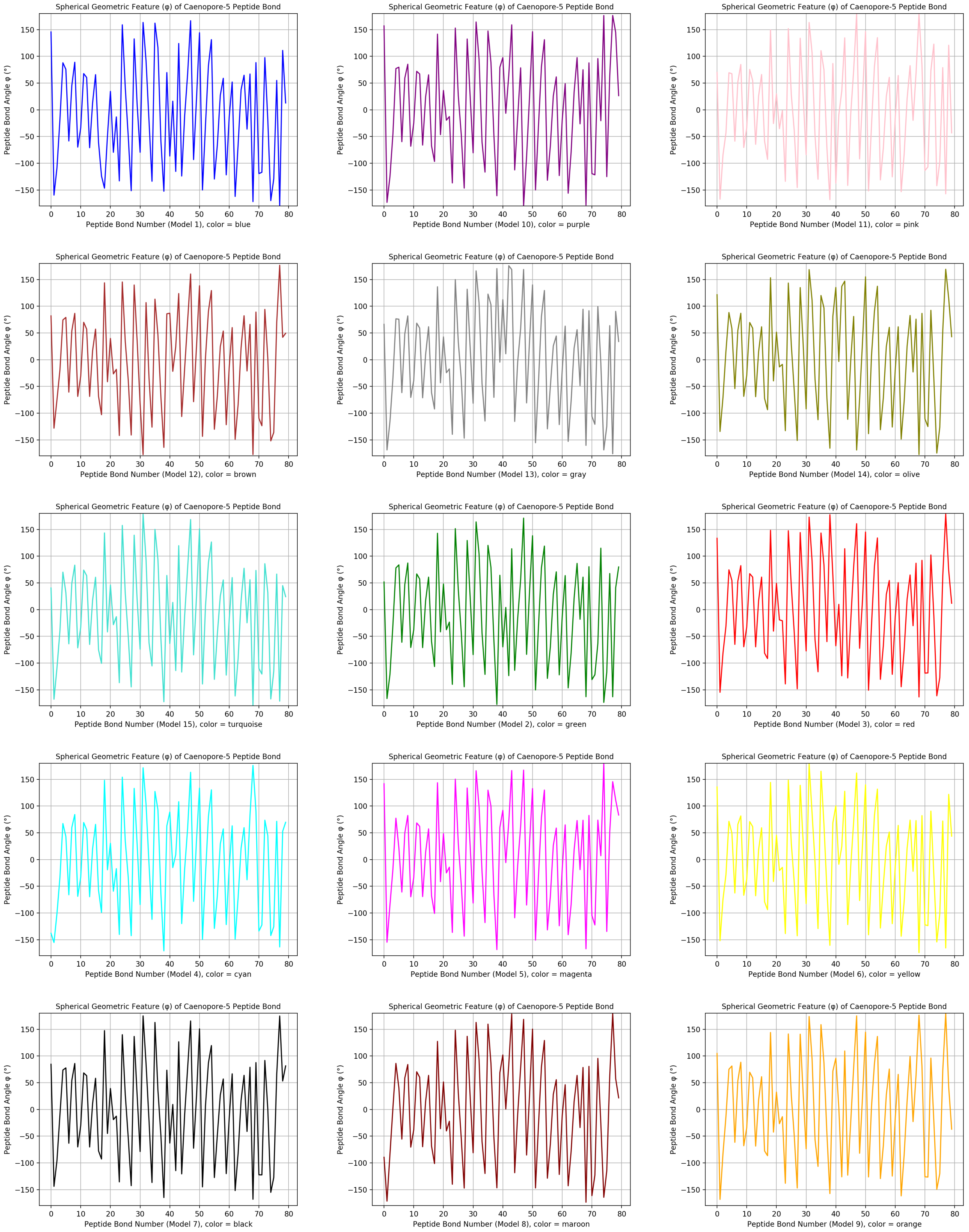


Figure 9: Distribution pattern of the polar angle (also known as the zenith angle, $\phi$) of 80 peptide bonds of Caenopore-5 [6–8] as reversibly extracted from the 15 NMR structural models of the three-dimensional NMR ensemble of Caenopore-5 (PDB ID: 2JSA) [6–8].

#### 2.2 Reversible extraction of $\rho$, $\theta$ and $\phi$ from protein structure: how?

Take Caenopore-5 [6–8] as example. The backbone atomic positions of Caenopore-5 are originally defined in the Cartesian coordinate system (CCS), where each atom's position is described by its [$x$, $y$, $z$] coordinates. These Cartesian coordinates are transformed into the SCS, where atomic positions are described in terms of radial distance ($\rho$), polar angle ($\theta$) and azimuthal angle ($\phi$), as described by Equations 1, 2 and 3. Afterwards, the spherical coordinates $\rho$, $\theta$ and $\phi$ can be used to fully describe the geometry of the covalent bonds, where covalent bond lengths and their spatial orientations are preserved, allowing for accurate representation of the protein structure. On the other hand, after the spherical coordinates $\rho$, $\theta$ and $\phi$ are extracted, they can be converted back into Cartesian coordinates using the geometric relationships between CCS and SCS:

$$x = \rho \sin\theta \cos\phi, \quad y = \rho \sin\theta \sin\phi, \quad z = \rho \cos\theta$$

#### 2.3 There is a total of 80 peptide bonds in Caenopore-5: why?

A peptide bond forms between the carboxyl group of one amino acid and the amino group of the next amino acid in the chain, as shown in Figure 3. For $N$ amino acids in a chain, there are $N - 1$ peptide bonds, since the first amino acid has no preceding amino acid to form a bond with, and the last amino acid has no subsequent amino acid to form a bond with.

Therefore, for a protein consisting of 81 amino acids, the number of peptide bonds is:

$$\text{Number of peptide bonds} = 81 - 1 = 80$$

Thus, Caenopore-5 (PDB ID: 2JSA) [6–8] with 81 amino acids contains 80 peptide bonds.

#### 2.4 There is 477 covalent bonds in Caenopore-5's backbone: why?

Caenopore-5 (PDB ID: 2JSA) consists of 81 amino acids [6–8]. Take Arginine 1 (Arg1) for example, whose six covalent bonds in the backbone of Caenopore-5 are included in Table 1. Specifically, the six covalent bonds consists of:

1. the covalent bond formed between N (Amine nitrogen) and first hydrogen attached to N (Amine nitrogen);
2. the covalent bond formed between N (Amine nitrogen) and C$\alpha$ (Alpha carbon);
3. the covalent bond formed between C$\alpha$ (Alpha carbon) and H$\alpha$ (Hydrogen attached to C$\alpha$);
4. the covalent bond formed between C$\alpha$ (Alpha carbon) and C (Carbonyl carbon);
5. the covalent bond formed between C (Carbonyl carbon) and O (Carbonyl oxygen);
6. the covalent bond formed between C (Carbonyl carbon) and N (Amine nitrogen) of the next amino acid residue.

| Bond ID | Model ID | Atom_Initial | Atom_Final | $X_i$ | $Y_i$ | $Z_i$ | $X_f$ | $Y_f$ | $Z_f$ | $\rho$ (Å) | $\theta$ (°) | $\phi$ (°) |
|---|---|---|---|---|---|---|---|---|---|---|---|---|
| 1 | 2jsa-1 | ARG_A_1_N | ARG_A_1_H | 14.733 | 4.945 | 1.144 | 14.853 | 5.924 | 1.361 | 1.01 | 77.59 | 83.01 |
| 2 | 2jsa-1 | ARG_A_1_N | ARG_A_1_CA | 14.733 | 4.945 | 1.144 | 14.178 | 4.583 | -0.15 | 1.45 | 152.88 | -146.89 |
| 3 | 2jsa-1 | ARG_A_1_CA | ARG_A_1_HA | 14.178 | 4.583 | -0.15 | 14.914 | 3.981 | -0.684 | 1.09 | 119.32 | -39.28 |
| 4 | 2jsa-1 | ARG_A_1_CA | ARG_A_1_C | 14.178 | 4.583 | -0.15 | 13.875 | 5.84 | -0.967 | 1.53 | 122.29 | 103.55 |
| 5 | 2jsa-1 | ARG_A_1_C | ARG_A_1_O | 13.875 | 5.84 | -0.967 | 14.57 | 6.135 | -1.939 | 1.23 | 142.16 | 23.00 |
| 6 | 2jsa-1 | ARG_A_1_C | SER_A_2_N | 13.875 | 5.84 | -0.967 | 12.838 | 6.548 | -0.544 | 1.32 | 71.38 | 145.68 |

Table 1: Six Arg1-specific covalent bonds in the backbone of Caenopore-5 [6–8]. This table is taken from Table 3, where $X_i$, $Y_i$ and $Z_i$ represent the Cartesian coordinates of **Atom_Initial**, and $X_f$, $Y_f$ and $Z_f$ represent the Cartesian coordinates of **Atom_Final**.

Thus, assume that there are a total of six covalent bonds in the backbone of Caenopore-5 for each amino acid residue, there are a total of $81 \times 6 = 486$ covalent bonds in the backbone of Caenopore-5. Nonetheless, there are a few exceptions.

As is known, glycine does not have a side chain, and its alpha carbon (C$\alpha$) is directly bonded to two hydrogens (H$\alpha_1$ and H$\alpha_2$). In PDB-format atom record, the two hydrogens (H$\alpha_1$ and H$\alpha_2$) are designated as **HA2** and **HA3**, respectively, as shown in Figure 10.

```
18266 ATOM    277  HE2 GLU A  17      -8.631   7.056   7.020  1.00  0.00           H
18267 ATOM    278  N   GLY A  18     -10.665   4.168   1.352  1.00  0.00           N
18268 ATOM    279  CA  GLY A  18     -11.832   4.218   0.486  1.00  0.00           C
18269 ATOM    280  C   GLY A  18     -12.197   2.823  -0.023  1.00  0.00           C
18270 ATOM    281  O   GLY A  18     -12.315   2.609  -1.228  1.00  0.00           O
18271 ATOM    282  H   GLY A  18      -9.780   3.878   0.960  1.00  0.00           H
18272 ATOM    283  HA2 GLY A  18     -12.674   4.626   1.045  1.00  0.00           H
18273 ATOM    284  HA3 GLY A  18     -11.617   4.864  -0.365  1.00  0.00           H
18274 ATOM    285  N   SER A  19     -12.366   1.908   0.922  1.00  0.00           N
```

Figure 10: Glycine 18 (Gly18) part of the PDB-format atom record of Caenopore-5 (PDB ID: 2jsa) [6–8].

Yet, in supplementary python script **S3.pdf** (Figure 11) for protein structure feature extraction, for all standard amino acids, the hydrogen attached to carbon (C$\alpha$) is designated as **HA** alone, excluding **HA2** and **HA3** (gray area of Figure 10). As a result, for the four glycines (Gly18, Gly45, Gly66 and Gly67), each glycine consists of five (Table 1), instead of six, covalent bonds in the backbone of Caenopore-5 (PDB ID: 2jsa) [6–8], because one covalent bond is not extracted, i.e., the covalent bond formed between C$\alpha$ (Alpha carbon) and H$\alpha$ (Hydrogen attached to C$\alpha$).

```python
for i in range(1,16):
    f='2jsa-' + str(i) + '.pdb'
    files.append(f)

backboneAtoms=[]
backboneAtoms.append('H')#N-terminal hydrogen, first atom
backboneAtoms.append('N')#N-terminal nitrogen
backboneAtoms.append('CA')#carbon alfa
backboneAtoms.append('HA')#hydrogen alfa
backboneAtoms.append('C')#carbonyl carbon
backboneAtoms.append('O')#carbonyl oxygen
```

Figure 11: Part of the supplementary python script **S3.pdf** for protein structure feature extraction with an alternative ABN-SCS approach of $\rho$, $\theta$ and $\phi$.

To address this issue, one straightforward solution is to add two lines of code (specifically for glycine) into supplementary file **S3.pdf**, as shown by the blue box in Figure 12 in comparison with Figure 10.

```python
for i in range(1,16):
    f='2jsa-' + str(i) + '.pdb'
    files.append(f)

backboneAtoms=[]
backboneAtoms.append('H')#N-terminal hydrogen, first atom
backboneAtoms.append('N')#N-terminal nitrogen
backboneAtoms.append('CA')#carbon alfa
backboneAtoms.append('HA')#hydrogen alfa
backboneAtoms.append('HA2')#hydrogen alfa for glycine
backboneAtoms.append('HA3')#hydrogen alfa for glycine
backboneAtoms.append('C')#carbonyl carbon
backboneAtoms.append('O')#carbonyl oxygen
```

Figure 12: Part of the newly edited python script for protein structure feature extraction with an alternative ABN-SCS approach of $\rho$, $\theta$ and $\phi$.

In addition, there are a total of four prolines in Caenopore-5 (PDB ID: 2jsa) [6–8], i.e., Pro43, Pro59, Pro70 and Pro81. Due to the chemical structure of proline, there is already one covalent bond between its N (Amine nitrogen) and its C$\delta$ (Delta carbon). Thus, in case a peptide bond (Figure 3) forms between proline and any other amino acid, the N (Amine nitrogen) of proline is covalently bonded to the N (Amine nitrogen) of the next amino acid residue. In light of the fact that the N (Amine nitrogen) of proline is already covalently bonded to its C$\alpha$ (Alpha carbon), for any proline, there is no hydrogen attached to N (Amine nitrogen) of the backbone of proline. As a result, for the four prolines (Pro43, Pro59, Pro70 and Pro81), each proline consists of five, instead of six, covalent bonds in the backbone of Caenopore-5 (PDB ID: 2jsa) [6–8], because one covalent bond does not exist, i.e., a covalent bond formed between N (Amine nitrogen) and first hydrogen attached to N (Amine nitrogen). Here, Pro81 is another exception, which is at the C-terminus of Caenopore-5. For Pro81, there is only four (Table 2), instead of five or six, covalent bonds in the backbone of Caenopore-5 (PDB ID:

2jsa) [6–8], because Pro81 does not form a peptide bond (Figure 3) with the next amino acid residue, i.e., there is no covalent bond for Pro81 formed between C (Carbonyl carbon) and N (Amine nitrogen) of the next amino acid residue.

| **Bond ID** | **Model ID** | **Atom_Initial** | **Atom_Final** | $\mathbf{X_i}$ | $\mathbf{Y_i}$ | $\mathbf{Z_i}$ | $\mathbf{X_f}$ | $\mathbf{Y_f}$ | $\mathbf{Z_f}$ | $\rho$ **(Å)** | $\theta$ **(°)** | $\phi$ **(°)** |
|---|---|---|---|---|---|---|---|---|---|---|---|---|
| 474 | 2jsa-15 | PRO_A_81_N | PRO_A_81_CA | 10.513 | 9.239 | 1.452 | 11.664 | 10.119 | 1.57 | 1.45 | 85.34 | 37.40 |
| 475 | 2jsa-15 | PRO_A_81_CA | PRO_A_81_HA | 11.664 | 10.119 | 1.57 | 12.271 | 9.827 | 2.427 | 1.09 | 38.17 | -25.69 |
| 476 | 2jsa-15 | PRO_A_81_CA | PRO_A_81_C | 11.664 | 10.119 | 1.57 | 11.225 | 11.574 | 1.758 | 1.53 | 82.95 | 106.79 |
| 477 | 2jsa-15 | PRO_A_81_C | PRO_A_81_O | 11.225 | 11.574 | 1.758 | 11.249 | 12.093 | 2.873 | 1.23 | 24.98 | 87.35 |

Table 2: Four Pro81-specific covalent bonds in the backbone of Caenopore-5 [6–8]. This table is taken from Table 3 in this file, where $X_i$, $Y_i$ and $Z_i$ represent the Cartesian coordinates of **Atom_Initial**, and $X_f$, $Y_f$ and $Z_f$ represent the Cartesian coordinates of **Atom_Final**.

Taken together, for Caenopore-5 (PDB ID: 2jsa) [6–8], a set of spherical coordinates ($\rho$, $\theta$ and $\phi$) are included in Table 3 (this file) for all covalent bonds in its backbone, and

$$\text{number of backbone covalent bonds} = 486 - 4 - 4 - 1 = 477$$

#### 2.5 An ABN-SCS Approach for Protein Backbone Structure Feature Extraction: A Case Study with Cp5

| Bond ID | Model ID | Atom_Initial | Atom_Final | $X_i$ | $Y_i$ | $Z_i$ | $X_f$ | $Y_f$ | $Z_f$ | ρ (Å) | θ (°) | ϕ (°) |
|---|---|---|---|---|---|---|---|---|---|---|---|---|
| 1 | 2jsa-1 | ARG_A_1_N | ARG_A_1_H | 14.733 | 4.945 | 1.144 | 14.853 | 5.924 | 1.361 | 1.01 | 77.59 | 83.01 |
| 2 | 2jsa-1 | ARG_A_1_N | ARG_A_1_CA | 14.733 | 4.945 | 1.144 | 14.178 | 4.583 | -0.15 | 1.45 | 152.88 | -146.89 |
| 3 | 2jsa-1 | ARG_A_1_CA | ARG_A_1_HA | 14.178 | 4.583 | -0.15 | 14.914 | 3.981 | -0.684 | 1.09 | 119.32 | -39.28 |
| 4 | 2jsa-1 | ARG_A_1_CA | ARG_A_1_C | 14.178 | 4.583 | -0.15 | 13.875 | 5.84 | -0.967 | 1.53 | 122.29 | 103.55 |
| 5 | 2jsa-1 | ARG_A_1_C | ARG_A_1_O | 13.875 | 5.84 | -0.967 | 14.57 | 6.135 | -1.939 | 1.23 | 142.16 | 23.00 |
| 6 | 2jsa-1 | ARG_A_1_C | SER_A_2_N | 13.875 | 5.84 | -0.967 | 12.838 | 6.548 | -0.544 | 1.32 | 71.38 | 145.68 |
| 7 | 2jsa-1 | SER_A_2_N | SER_A_2_H | 12.838 | 6.548 | -0.544 | 12.312 | 6.246 | 0.264 | 1.01 | 36.89 | -150.14 |
| 8 | 2jsa-1 | SER_A_2_N | SER_A_2_CA | 12.838 | 6.548 | -0.544 | 12.436 | 7.767 | -1.225 | 1.45 | 117.95 | 108.25 |
| 9 | 2jsa-1 | SER_A_2_CA | SER_A_2_HA | 12.436 | 7.767 | -1.225 | 11.492 | 8.102 | -0.796 | 1.09 | 66.82 | 160.46 |
| 10 | 2jsa-1 | SER_A_2_CA | SER_A_2_C | 12.436 | 7.767 | -1.225 | 12.226 | 7.489 | -2.715 | 1.53 | 166.84 | -127.07 |
| 11 | 2jsa-1 | SER_A_2_C | SER_A_2_O | 12.226 | 7.489 | -2.715 | 13.136 | 7.682 | -3.52 | 1.23 | 130.87 | 11.97 |
| 12 | 2jsa-1 | SER_A_2_C | ALA_A_3_N | 12.226 | 7.489 | -2.715 | 11.022 | 7.04 | -3.037 | 1.32 | 104.07 | -159.55 |
| 13 | 2jsa-1 | ALA_A_3_N | ALA_A_3_H | 11.022 | 7.04 | -3.037 | 10.322 | 6.904 | -2.322 | 1.01 | 44.92 | -169.01 |
| 14 | 2jsa-1 | ALA_A_3_N | ALA_A_3_CA | 11.022 | 7.04 | -3.037 | 10.682 | 6.734 | -4.416 | 1.45 | 161.65 | -138.01 |
| 15 | 2jsa-1 | ALA_A_3_CA | ALA_A_3_HA | 10.682 | 6.734 | -4.416 | 10.847 | 7.624 | -5.023 | 1.09 | 123.85 | 79.50 |
| 16 | 2jsa-1 | ALA_A_3_CA | ALA_A_3_C | 10.682 | 6.734 | -4.416 | 9.201 | 6.358 | -4.5 | 1.53 | 93.15 | -165.75 |
| 17 | 2jsa-1 | ALA_A_3_C | ALA_A_3_O | 9.201 | 6.358 | -4.5 | 8.384 | 6.865 | -3.733 | 1.23 | 51.42 | 148.18 |
| 18 | 2jsa-1 | ALA_A_3_C | LEU_A_4_N | 9.201 | 6.358 | -4.5 | 8.901 | 5.473 | -5.439 | 1.32 | 135.14 | -108.73 |
| 19 | 2jsa-1 | LEU_A_4_N | LEU_A_4_H | 8.901 | 5.473 | -5.439 | 9.626 | 5.098 | -6.034 | 1.01 | 126.09 | -27.35 |
| 20 | 2jsa-1 | LEU_A_4_N | LEU_A_4_CA | 8.901 | 5.473 | -5.439 | 7.533 | 5.024 | -5.634 | 1.45 | 97.71 | -161.83 |
| 21 | 2jsa-1 | LEU_A_4_CA | LEU_A_4_HA | 7.533 | 5.024 | -5.634 | 6.905 | 5.905 | -5.767 | 1.09 | 97.01 | 125.48 |
| 22 | 2jsa-1 | LEU_A_4_CA | LEU_A_4_C | 7.533 | 5.024 | -5.634 | 7.055 | 4.298 | -4.374 | 1.53 | 34.60 | -123.36 |
| 23 | 2jsa-1 | LEU_A_4_C | LEU_A_4_O | 7.055 | 4.298 | -4.374 | 5.862 | 4.047 | -4.213 | 1.23 | 82.48 | -168.12 |
| 24 | 2jsa-1 | LEU_A_4_C | SER_A_5_N | 7.055 | 4.298 | -4.374 | 8.012 | 3.981 | -3.514 | 1.33 | 49.53 | -18.33 |
| 25 | 2jsa-1 | SER_A_5_N | SER_A_5_H | 8.012 | 3.981 | -3.514 | 8.974 | 4.22 | -3.711 | 1.01 | 101.24 | 13.95 |
| 26 | 2jsa-1 | SER_A_5_N | SER_A_5_CA | 8.012 | 3.981 | -3.514 | 7.704 | 3.288 | -2.274 | 1.45 | 31.45 | -113.96 |
| 27 | 2jsa-1 | SER_A_5_CA | SER_A_5_HA | 7.704 | 3.288 | -2.274 | 7.472 | 2.25 | -2.511 | 1.09 | 102.56 | -102.60 |
| 28 | 2jsa-1 | SER_A_5_CA | SER_A_5_C | 7.704 | 3.288 | -2.274 | 6.482 | 3.925 | -1.609 | 1.53 | 64.24 | 152.47 |
| 29 | 2jsa-1 | SER_A_5_C | SER_A_5_O | 6.482 | 3.925 | -1.609 | 5.549 | 3.226 | -1.218 | 1.23 | 71.46 | -143.16 |
| 30 | 2jsa-1 | SER_A_5_C | CYS_A_6_N | 6.482 | 3.925 | -1.609 | 6.528 | 5.245 | -1.501 | 1.33 | 85.33 | 88.00 |
| 31 | 2jsa-1 | CYS_A_6_N | CYS_A_6_H | 6.528 | 5.245 | -1.501 | 7.328 | 5.756 | -1.845 | 1.01 | 109.92 | 32.57 |
| 32 | 2jsa-1 | CYS_A_6_N | CYS_A_6_CA | 6.528 | 5.245 | -1.501 | 5.436 | 5.984 | -0.89 | 1.45 | 65.14 | 145.91 |
| 33 | 2jsa-1 | CYS_A_6_CA | CYS_A_6_HA | 5.436 | 5.984 | -0.89 | 5.312 | 5.629 | 0.133 | 1.09 | 20.18 | -109.25 |
| 34 | 2jsa-1 | CYS_A_6_CA | CYS_A_6_C | 5.436 | 5.984 | -0.89 | 4.155 | 5.681 | -1.669 | 1.53 | 120.62 | -166.69 |
| 35 | 2jsa-1 | CYS_A_6_C | CYS_A_6_O | 4.155 | 5.681 | -1.669 | 3.163 | 5.24 | -1.091 | 1.23 | 61.97 | -156.03 |
| 36 | 2jsa-1 | CYS_A_6_C | GLN_A_7_N | 4.155 | 5.681 | -1.669 | 4.218 | 5.929 | -2.969 | 1.32 | 168.86 | 75.75 |
| 37 | 2jsa-1 | GLN_A_7_N | GLN_A_7_H | 4.218 | 5.929 | -2.969 | 5.068 | 6.292 | -3.377 | 1.01 | 113.82 | 23.13 |
| 38 | 2jsa-1 | GLN_A_7_N | GLN_A_7_CA | 4.218 | 5.929 | -2.969 | 3.075 | 5.688 | -3.833 | 1.45 | 126.49 | -168.09 |
| 39 | 2jsa-1 | GLN_A_7_CA | GLN_A_7_HA | 3.075 | 5.688 | -3.833 | 2.417 | 6.554 | -3.77 | 1.09 | 86.68 | 127.23 |
| 40 | 2jsa-1 | GLN_A_7_CA | GLN_A_7_C | 3.075 | 5.688 | -3.833 | 2.299 | 4.459 | -3.356 | 1.53 | 71.83 | -122.27 |
| 41 | 2jsa-1 | GLN_A_7_C | GLN_A_7_O | 2.299 | 4.459 | -3.356 | 1.117 | 4.555 | -3.029 | 1.23 | 74.58 | 175.36 |
| 42 | 2jsa-1 | GLN_A_7_C | MET_A_8_N | 2.299 | 4.459 | -3.356 | 2.995 | 3.332 | -3.33 | 1.32 | 88.88 | -58.30 |
| 43 | 2jsa-1 | MET_A_8_N | MET_A_8_H | 2.995 | 3.332 | -3.33 | 3.965 | 3.327 | -3.613 | 1.01 | 106.26 | -0.30 |
| 44 | 2jsa-1 | MET_A_8_N | MET_A_8_CA | 2.995 | 3.332 | -3.33 | 2.386 | 2.085 | -2.899 | 1.45 | 72.75 | -116.03 |
| 45 | 2jsa-1 | MET_A_8_CA | MET_A_8_HA | 2.386 | 2.085 | -2.899 | 1.716 | 1.734 | -3.683 | 1.09 | 136.03 | -152.35 |
| 46 | 2jsa-1 | MET_A_8_CA | MET_A_8_C | 2.386 | 2.085 | -2.899 | 1.584 | 2.284 | -1.611 | 1.53 | 32.68 | 166.06 |
| 47 | 2jsa-1 | MET_A_8_C | MET_A_8_O | 1.584 | 2.284 | -1.611 | 0.433 | 1.86 | -1.52 | 1.23 | 85.76 | -159.78 |
| 48 | 2jsa-1 | MET_A_8_C | CYS_A_9_N | 1.584 | 2.284 | -1.611 | 2.224 | 2.929 | -0.647 | 1.32 | 43.31 | 45.22 |
| 49 | 2jsa-1 | CYS_A_9_N | CYS_A_9_H | 2.224 | 2.929 | -0.647 | 3.171 | 3.251 | -0.791 | 1.01 | 98.19 | 18.78 |
| 50 | 2jsa-1 | CYS_A_9_N | CYS_A_9_CA | 2.224 | 2.929 | -0.647 | 1.585 | 3.189 | 0.632 | 1.45 | 28.34 | 157.86 |
| 51 | 2jsa-1 | CYS_A_9_CA | CYS_A_9_HA | 1.585 | 3.189 | 0.632 | 1.35 | 2.232 | 1.099 | 1.09 | 64.64 | -103.80 |
| 52 | 2jsa-1 | CYS_A_9_CA | CYS_A_9_C | 1.585 | 3.189 | 0.632 | 0.277 | 3.938 | 0.372 | 1.53 | 99.79 | 150.20 |
| 53 | 2jsa-1 | CYS_A_9_C | CYS_A_9_O | 0.277 | 3.938 | 0.372 | -0.691 | 3.78 | 1.113 | 1.23 | 52.93 | -170.73 |
| 54 | 2jsa-1 | CYS_A_9_C | GLU_A_10_N | 0.277 | 3.938 | 0.372 | 0.291 | 4.739 | -0.684 | 1.33 | 142.81 | 89.00 |
| 55 | 2jsa-1 | GLU_A_10_N | GLU_A_10_H | 0.291 | 4.739 | -0.684 | 1.124 | 4.82 | -1.25 | 1.01 | 124.07 | 5.55 |
| 56 | 2jsa-1 | GLU_A_10_N | GLU_A_10_CA | 0.291 | 4.739 | -0.684 | -0.882 | 5.513 | -1.051 | 1.45 | 104.64 | 146.58 |
| 57 | 2jsa-1 | GLU_A_10_CA | GLU_A_10_HA | -0.882 | 5.513 | -1.051 | -1.393 | 5.806 | -0.134 | 1.09 | 32.71 | 150.17 |
| 58 | 2jsa-1 | GLU_A_10_CA | GLU_A_10_C | -0.882 | 5.513 | -1.051 | -1.841 | 4.661 | -1.885 | 1.53 | 123.03 | -138.38 |
| 59 | 2jsa-1 | GLU_A_10_C | GLU_A_10_O | -1.841 | 4.661 | -1.885 | -2.935 | 5.107 | -2.228 | 1.23 | 106.19 | 157.82 |
| 60 | 2jsa-1 | GLU_A_10_C | LEU_A_11_N | -1.841 | 4.661 | -1.885 | -1.395 | 3.451 | -2.188 | 1.32 | 103.22 | -69.77 |
| 61 | 2jsa-1 | LEU_A_11_N | LEU_A_11_H | -1.395 | 3.451 | -2.188 | -0.483 | 3.151 | -1.873 | 1.01 | 71.84 | -18.21 |
| 62 | 2jsa-1 | LEU_A_11_N | LEU_A_11_CA | -1.395 | 3.451 | -2.188 | -2.2 | 2.532 | -2.975 | 1.45 | 122.79 | -131.22 |
| 63 | 2jsa-1 | LEU_A_11_CA | LEU_A_11_HA | -2.2 | 2.532 | -2.975 | -2.895 | 3.12 | -3.573 | 1.09 | 123.30 | 139.77 |
| 64 | 2jsa-1 | LEU_A_11_CA | LEU_A_11_C | -2.2 | 2.532 | -2.975 | -3.025 | 1.65 | -2.036 | 1.53 | 52.13 | -133.09 |
| 65 | 2jsa-1 | LEU_A_11_C | LEU_A_11_O | -3.025 | 1.65 | -2.036 | -4.199 | 1.39 | -2.296 | 1.23 | 102.20 | -167.51 |
| 66 | 2jsa-1 | LEU_A_11_C | VAL_A_12_N | -3.025 | 1.65 | -2.036 | -2.379 | 1.215 | -0.965 | 1.32 | 36.02 | -33.96 |
| 67 | 2jsa-1 | VAL_A_12_N | VAL_A_12_H | -2.379 | 1.215 | -0.965 | -1.413 | 1.469 | -0.815 | 1.01 | 81.46 | 14.73 |
| 68 | 2jsa-1 | VAL_A_12_N | VAL_A_12_CA | -2.379 | 1.215 | -0.965 | -3.039 | 0.367 | 0.014 | 1.45 | 47.66 | -127.89 |
| 69 | 2jsa-1 | VAL_A_12_CA | VAL_A_12_HA | -3.039 | 0.367 | 0.014 | -3.582 | -0.414 | -0.517 | 1.09 | 119.17 | -124.81 |
| 70 | 2jsa-1 | VAL_A_12_CA | VAL_A_12_C | -3.039 | 0.367 | 0.014 | -4.052 | 1.199 | 0.802 | 1.53 | 58.99 | 140.60 |
| 71 | 2jsa-1 | VAL_A_12_C | VAL_A_12_O | -4.052 | 1.199 | 0.802 | -5.206 | 0.8 | 0.952 | 1.23 | 83.00 | -160.93 |
| 72 | 2jsa-1 | VAL_A_12_C | VAL_A_13_N | -4.052 | 1.199 | 0.802 | -3.584 | 2.341 | 1.285 | 1.33 | 68.63 | 67.72 |
| 73 | 2jsa-1 | VAL_A_13_N | VAL_A_13_H | -3.584 | 2.341 | 1.285 | -2.623 | 2.606 | 1.122 | 1.01 | 99.29 | 15.42 |
| 74 | 2jsa-1 | VAL_A_13_N | VAL_A_13_CA | -3.584 | 2.341 | 1.285 | -4.435 | 3.233 | 2.055 | 1.45 | 58.01 | 133.65 |
| 75 | 2jsa-1 | VAL_A_13_CA | VAL_A_13_HA | -4.435 | 3.233 | 2.055 | -4.847 | 2.676 | 2.896 | 1.09 | 39.48 | -126.49 |
| 76 | 2jsa-1 | VAL_A_13_CA | VAL_A_13_C | -4.435 | 3.233 | 2.055 | -5.599 | 3.701 | 1.179 | 1.53 | 124.92 | 158.10 |
| 77 | 2jsa-1 | VAL_A_13_C | VAL_A_13_O | -5.599 | 3.701 | 1.179 | -6.762 | 3.494 | 1.523 | 1.23 | 73.76 | -169.91 |
| 78 | 2jsa-1 | VAL_A_13_C | LYS_A_14_N | -5.599 | 3.701 | 1.179 | -5.246 | 4.324 | 0.064 | 1.33 | 147.29 | 60.46 |
| 79 | 2jsa-1 | LYS_A_14_N | LYS_A_14_H | -5.246 | 4.324 | 0.064 | -4.269 | 4.46 | -0.155 | 1.01 | 102.52 | 7.92 |
| 80 | 2jsa-1 | LYS_A_14_N | LYS_A_14_CA | -5.246 | 4.324 | 0.064 | -6.247 | 4.823 | -0.863 | 1.45 | 129.65 | 153.50 |
| 81 | 2jsa-1 | LYS_A_14_CA | LYS_A_14_HA | -6.247 | 4.823 | -0.863 | -6.724 | 5.691 | -0.409 | 1.09 | 65.37 | 118.79 |
| 82 | 2jsa-1 | LYS_A_14_CA | LYS_A_14_C | -6.247 | 4.823 | -0.863 | -7.323 | 3.755 | -1.069 | 1.53 | 97.74 | -135.21 |
| 83 | 2jsa-1 | LYS_A_14_C | LYS_A_14_O | -7.323 | 3.755 | -1.069 | -8.493 | 4.078 | -1.272 | 1.23 | 99.49 | 164.57 |
| 84 | 2jsa-1 | LYS_A_14_C | LYS_A_15_N | -7.323 | 3.755 | -1.069 | -6.89 | 2.505 | -1.009 | 1.32 | 87.40 | -70.89 |
| 85 | 2jsa-1 | LYS_A_15_N | LYS_A_15_H | -6.89 | 2.505 | -1.009 | -5.913 | 2.313 | -0.838 | 1.01 | 80.26 | -11.12 |
| 86 | 2jsa-1 | LYS_A_15_N | LYS_A_15_CA | -6.89 | 2.505 | -1.009 | -7.802 | 1.387 | -1.186 | 1.45 | 96.99 | -129.21 |
| 87 | 2jsa-1 | LYS_A_15_CA | LYS_A_15_HA | -7.802 | 1.387 | -1.186 | -8.56 | 1.681 | -1.911 | 1.09 | 131.72 | 158.80 |
| 88 | 2jsa-1 | LYS_A_15_CA | LYS_A_15_C | -7.802 | 1.387 | -1.186 | -8.512 | 1.1 | 0.138 | 1.53 | 30.05 | -157.99 |
| 89 | 2jsa-1 | LYS_A_15_C | LYS_A_15_O | -8.512 | 1.1 | 0.138 | -9.688 | 0.737 | 0.15 | 1.23 | 89.44 | -162.85 |
| 90 | 2jsa-1 | LYS_A_15_C | TYR_A_16_N | -8.512 | 1.1 | 0.138 | -7.769 | 1.272 | 1.221 | 1.32 | 35.15 | 13.03 |
| 91 | 2jsa-1 | TYR_A_16_N | TYR_A_16_H | -7.769 | 1.272 | 1.221 | -6.808 | 1.571 | 1.136 | 1.01 | 94.83 | 17.28 |
| 92 | 2jsa-1 | TYR_A_16_N | TYR_A_16_CA | -7.769 | 1.272 | 1.221 | -8.313 | 1.035 | 2.548 | 1.45 | 24.09 | -156.46 |
| 93 | 2jsa-1 | TYR_A_16_CA | TYR_A_16_HA | -8.313 | 1.035 | 2.548 | -8.806 | 0.064 | 2.586 | 1.09 | 88.00 | -116.92 |
| 94 | 2jsa-1 | TYR_A_16_CA | TYR_A_16_C | -8.313 | 1.035 | 2.548 | -9.301 | 2.135 | 2.94 | 1.53 | 75.15 | 131.93 |
| 95 | 2jsa-1 | TYR_A_16_C | TYR_A_16_O | -9.301 | 2.135 | 2.94 | -10.499 | 1.884 | 3.063 | 1.23 | 84.26 | -168.17 |

| | | | | | | | | | | | | |
|---|---|---|---|---|---|---|---|---|---|---|---|---|
| 96 | 2jsa-1 | TYR_A_16_C | GLU_A_17_N | -9.301 | 2.135 | 2.94 | -8.762 | 3.332 | 3.124 | 1.33 | 82.02 | 65.76 |
| 97 | 2jsa-1 | GLU_A_17_N | GLU_A_17_H | -8.762 | 3.332 | 3.124 | -7.768 | 3.466 | 3.004 | 1.01 | 96.82 | 7.68 |
| 98 | 2jsa-1 | GLU_A_17_N | GLU_A_17_CA | -8.762 | 3.332 | 3.124 | -9.582 | 4.472 | 3.499 | 1.45 | 75.05 | 125.73 |
| 99 | 2jsa-1 | GLU_A_17_CA | GLU_A_17_HA | -9.582 | 4.472 | 3.499 | -9.988 | 4.283 | 4.493 | 1.09 | 24.25 | -155.04 |
| 100 | 2jsa-1 | GLU_A_17_CA | GLU_A_17_C | -9.582 | 4.472 | 3.499 | -10.75 | 4.629 | 2.524 | 1.53 | 129.60 | 172.34 |
| 101 | 2jsa-1 | GLU_A_17_C | GLU_A_17_O | -10.75 | 4.629 | 2.524 | -11.813 | 5.122 | 2.898 | 1.23 | 72.30 | 155.12 |
| 102 | 2jsa-1 | GLU_A_17_C | GLY_A_18_N | -10.75 | 4.629 | 2.524 | -10.513 | 4.202 | 1.292 | 1.33 | 158.38 | -60.97 |
| 103 | 2jsa-1 | GLY_A_18_N | GLY_A_18_H | -10.513 | 4.202 | 1.292 | -9.614 | 3.808 | 1.055 | 1.01 | 103.57 | -23.67 |
| 104 | 2jsa-1 | GLY_A_18_N | GLY_A_18_CA | -10.513 | 4.202 | 1.292 | -11.532 | 4.29 | 0.261 | 1.45 | 135.23 | 175.06 |
| 105 | 2jsa-1 | GLY_A_18_CA | GLY_A_18_C | -11.532 | 4.29 | 0.261 | -11.817 | 2.915 | -0.347 | 1.53 | 113.41 | -101.71 |
| 106 | 2jsa-1 | GLY_A_18_C | GLY_A_18_O | -11.817 | 2.915 | -0.347 | -11.513 | 2.673 | -1.514 | 1.23 | 161.58 | -38.52 |
| 107 | 2jsa-1 | GLY_A_18_C | SER_A_19_N | -11.817 | 2.915 | -0.347 | -12.397 | 2.05 | 0.472 | 1.32 | 51.82 | -123.84 |
| 108 | 2jsa-1 | SER_A_19_N | SER_A_19_H | -12.397 | 2.05 | 0.472 | -12.618 | 2.317 | 1.421 | 1.01 | 20.06 | 129.62 |
| 109 | 2jsa-1 | SER_A_19_N | SER_A_19_CA | -12.397 | 2.05 | 0.472 | -12.727 | 0.706 | 0.029 | 1.45 | 107.75 | -103.80 |
| 110 | 2jsa-1 | SER_A_19_CA | SER_A_19_HA | -12.727 | 0.706 | 0.029 | -12.201 | 0.516 | -0.907 | 1.09 | 149.14 | -19.86 |
| 111 | 2jsa-1 | SER_A_19_CA | SER_A_19_C | -12.727 | 0.706 | 0.029 | -14.232 | 0.591 | -0.222 | 1.53 | 99.44 | -175.63 |
| 112 | 2jsa-1 | SER_A_19_C | SER_A_19_O | -14.232 | 0.591 | -0.222 | -14.71 | 0.918 | -1.307 | 1.23 | 151.91 | 145.62 |
| 113 | 2jsa-1 | SER_A_19_C | ALA_A_20_N | -14.232 | 0.591 | -0.222 | -14.936 | 0.126 | 0.799 | 1.32 | 39.57 | -146.55 |
| 114 | 2jsa-1 | ALA_A_20_N | ALA_A_20_H | -14.936 | 0.126 | 0.799 | -14.475 | -0.125 | 1.662 | 1.01 | 31.31 | -28.57 |
| 115 | 2jsa-1 | ALA_A_20_N | ALA_A_20_CA | -14.936 | 0.126 | 0.799 | -16.377 | -0.035 | 0.703 | 1.45 | 93.79 | -173.62 |
| 116 | 2jsa-1 | ALA_A_20_CA | ALA_A_20_HA | -16.377 | -0.035 | 0.703 | -16.85 | 0.93 | 0.881 | 1.09 | 80.60 | 116.11 |
| 117 | 2jsa-1 | ALA_A_20_CA | ALA_A_20_C | -16.377 | -0.035 | 0.703 | -16.849 | -1.012 | 1.781 | 1.53 | 45.19 | -115.79 |
| 118 | 2jsa-1 | ALA_A_20_C | ALA_A_20_O | -16.849 | -1.012 | 1.781 | -17.938 | -0.855 | 2.333 | 1.23 | 63.36 | 171.80 |
| 119 | 2jsa-1 | ALA_A_20_C | ASP_A_21_N | -16.849 | -1.012 | 1.781 | -16.008 | -2.0 | 2.049 | 1.32 | 78.33 | -49.60 |
| 120 | 2jsa-1 | ASP_A_21_N | ASP_A_21_H | -16.008 | -2.0 | 2.049 | -15.129 | -2.068 | 1.555 | 1.01 | 119.26 | -4.42 |
| 121 | 2jsa-1 | ASP_A_21_N | ASP_A_21_CA | -16.008 | -2.0 | 2.049 | -16.326 | -3.003 | 3.052 | 1.45 | 46.37 | -107.59 |
| 122 | 2jsa-1 | ASP_A_21_CA | ASP_A_21_HA | -16.326 | -3.003 | 3.052 | -17.356 | -2.866 | 3.379 | 1.09 | 72.53 | 172.42 |
| 123 | 2jsa-1 | ASP_A_21_CA | ASP_A_21_C | -16.326 | -3.003 | 3.052 | -15.378 | -2.846 | 4.242 | 1.53 | 38.92 | 9.40 |
| 124 | 2jsa-1 | ASP_A_21_C | ASP_A_21_O | -15.378 | -2.846 | 4.242 | -15.634 | -3.383 | 5.319 | 1.23 | 28.91 | -115.49 |
| 125 | 2jsa-1 | ASP_A_21_C | LYS_A_22_N | -15.378 | -2.846 | 4.242 | -14.303 | -2.108 | 4.009 | 1.32 | 100.13 | 34.47 |
| 126 | 2jsa-1 | LYS_A_22_N | LYS_A_22_H | -14.303 | -2.108 | 4.009 | -14.156 | -1.696 | 3.099 | 1.01 | 154.33 | 70.36 |
| 127 | 2jsa-1 | LYS_A_22_N | LYS_A_22_CA | -14.303 | -2.108 | 4.009 | -13.316 | -1.874 | 5.049 | 1.45 | 44.28 | 13.34 |
| 128 | 2jsa-1 | LYS_A_22_CA | LYS_A_22_HA | -13.316 | -1.874 | 5.049 | -12.587 | -1.158 | 4.668 | 1.09 | 110.45 | 44.48 |
| 129 | 2jsa-1 | LYS_A_22_CA | LYS_A_22_C | -13.316 | -1.874 | 5.049 | -12.574 | -3.179 | 5.345 | 1.53 | 78.85 | -60.38 |
| 130 | 2jsa-1 | LYS_A_22_C | LYS_A_22_O | -12.574 | -3.179 | 5.345 | -12.164 | -3.42 | 6.479 | 1.23 | 22.75 | -30.45 |
| 131 | 2jsa-1 | LYS_A_22_C | ASP_A_23_N | -12.574 | -3.179 | 5.345 | -12.424 | -3.986 | 4.305 | 1.32 | 141.72 | -79.47 |
| 132 | 2jsa-1 | ASP_A_23_N | ASP_A_23_H | -12.424 | -3.986 | 4.305 | -12.788 | -3.723 | 3.4 | 1.01 | 153.61 | 144.15 |
| 133 | 2jsa-1 | ASP_A_23_N | ASP_A_23_CA | -12.424 | -3.986 | 4.305 | -11.738 | -5.26 | 4.44 | 1.45 | 84.67 | -61.70 |
| 134 | 2jsa-1 | ASP_A_23_CA | ASP_A_23_HA | -11.738 | -5.26 | 4.44 | -11.841 | -5.613 | 5.466 | 1.09 | 19.72 | -106.27 |
| 135 | 2jsa-1 | ASP_A_23_CA | ASP_A_23_C | -11.738 | -5.26 | 4.44 | -10.258 | -5.077 | 4.099 | 1.53 | 102.88 | 7.05 |
| 136 | 2jsa-1 | ASP_A_23_C | ASP_A_23_O | -10.258 | -5.077 | 4.099 | -9.914 | -4.769 | 2.959 | 1.23 | 157.95 | 41.84 |
| 137 | 2jsa-1 | ASP_A_23_C | ALA_A_24_N | -10.258 | -5.077 | 4.099 | -9.423 | -5.274 | 5.108 | 1.32 | 40.37 | -13.27 |
| 138 | 2jsa-1 | ALA_A_24_N | ALA_A_24_H | -9.423 | -5.274 | 5.108 | -9.777 | -5.525 | 6.02 | 1.01 | 25.45 | -144.66 |
| 139 | 2jsa-1 | ALA_A_24_N | ALA_A_24_CA | -9.423 | -5.274 | 5.108 | -7.987 | -5.134 | 4.929 | 1.45 | 97.07 | 5.57 |
| 140 | 2jsa-1 | ALA_A_24_CA | ALA_A_24_HA | -7.987 | -5.134 | 4.929 | -7.783 | -4.168 | 4.466 | 1.09 | 115.12 | 78.08 |
| 141 | 2jsa-1 | ALA_A_24_CA | ALA_A_24_C | -7.987 | -5.134 | 4.929 | -7.485 | -6.235 | 3.993 | 1.53 | 127.72 | -65.49 |
| 142 | 2jsa-1 | ALA_A_24_C | ALA_A_24_O | -7.485 | -6.235 | 3.993 | -6.337 | -6.204 | 3.552 | 1.23 | 111.01 | 1.55 |
| 143 | 2jsa-1 | ALA_A_24_C | ASN_A_25_N | -7.485 | -6.235 | 3.993 | -8.369 | -7.183 | 3.718 | 1.33 | 101.98 | -133.00 |
| 144 | 2jsa-1 | ASN_A_25_N | ASN_A_25_H | -8.369 | -7.183 | 3.718 | -9.296 | -7.145 | 4.117 | 1.01 | 66.73 | 177.65 |
| 145 | 2jsa-1 | ASN_A_25_N | ASN_A_25_CA | -8.369 | -7.183 | 3.718 | -8.029 | -8.292 | 2.842 | 1.45 | 127.06 | -72.96 |
| 146 | 2jsa-1 | ASN_A_25_CA | ASN_A_25_HA | -8.029 | -8.292 | 2.842 | -6.979 | -8.551 | 2.985 | 1.09 | 82.47 | -13.86 |
| 147 | 2jsa-1 | ASN_A_25_CA | ASN_A_25_C | -8.029 | -8.292 | 2.842 | -8.27 | -7.88 | 1.388 | 1.53 | 161.83 | 120.33 |
| 148 | 2jsa-1 | ASN_A_25_C | ASN_A_25_O | -8.27 | -7.88 | 1.388 | -7.374 | -7.99 | 0.552 | 1.23 | 132.80 | -7.00 |
| 149 | 2jsa-1 | ASN_A_25_C | VAL_A_26_N | -8.27 | -7.88 | 1.388 | -9.483 | -7.415 | 1.131 | 1.32 | 101.19 | 159.03 |
| 150 | 2jsa-1 | VAL_A_26_N | VAL_A_26_H | -9.483 | -7.415 | 1.131 | -10.17 | -7.353 | 1.868 | 1.01 | 43.10 | 174.84 |
| 151 | 2jsa-1 | VAL_A_26_N | VAL_A_26_CA | -9.483 | -7.415 | 1.131 | -9.852 | -6.986 | -0.207 | 1.45 | 157.08 | 130.70 |
| 152 | 2jsa-1 | VAL_A_26_CA | VAL_A_26_HA | -9.852 | -6.986 | -0.207 | -9.442 | -7.699 | -0.922 | 1.09 | 131.00 | -60.10 |
| 153 | 2jsa-1 | VAL_A_26_CA | VAL_A_26_C | -9.852 | -6.986 | -0.207 | -9.229 | -5.618 | -0.49 | 1.53 | 100.66 | 65.52 |
| 154 | 2jsa-1 | VAL_A_26_C | VAL_A_26_O | -9.229 | -5.618 | -0.49 | -9.323 | -5.108 | -1.605 | 1.23 | 155.06 | 100.44 |
| 155 | 2jsa-1 | VAL_A_26_C | ILE_A_27_N | -9.229 | -5.618 | -0.49 | -8.604 | -5.064 | 0.539 | 1.33 | 39.06 | 41.55 |
| 156 | 2jsa-1 | ILE_A_27_N | ILE_A_27_H | -8.604 | -5.064 | 0.539 | -8.563 | -5.543 | 1.428 | 1.01 | 28.40 | -85.11 |
| 157 | 2jsa-1 | ILE_A_27_N | ILE_A_27_CA | -8.604 | -5.064 | 0.539 | -7.965 | -3.765 | 0.415 | 1.45 | 94.90 | 63.81 |
| 158 | 2jsa-1 | ILE_A_27_CA | ILE_A_27_HA | -7.965 | -3.765 | 0.415 | -8.529 | -3.179 | -0.311 | 1.09 | 131.75 | 133.90 |
| 159 | 2jsa-1 | ILE_A_27_CA | ILE_A_27_C | -7.965 | -3.765 | 0.415 | -6.549 | -3.949 | -0.136 | 1.53 | 111.10 | -7.40 |
| 160 | 2jsa-1 | ILE_A_27_C | ILE_A_27_O | -6.549 | -3.949 | -0.136 | -6.175 | -3.315 | -1.121 | 1.23 | 143.23 | 59.46 |
| 161 | 2jsa-1 | ILE_A_27_C | LYS_A_28_N | -6.549 | -3.949 | -0.136 | -5.8 | -4.82 | 0.525 | 1.33 | 60.08 | -49.31 |
| 162 | 2jsa-1 | LYS_A_28_N | LYS_A_28_H | -5.8 | -4.82 | 0.525 | -6.173 | -5.306 | 1.328 | 1.01 | 37.34 | -127.51 |
| 163 | 2jsa-1 | LYS_A_28_N | LYS_A_28_CA | -5.8 | -4.82 | 0.525 | -4.434 | -5.095 | 0.114 | 1.45 | 106.43 | -11.38 |
| 164 | 2jsa-1 | LYS_A_28_CA | LYS_A_28_HA | -4.434 | -5.095 | 0.114 | -3.902 | -4.145 | 0.052 | 1.09 | 93.26 | 60.75 |
| 165 | 2jsa-1 | LYS_A_28_CA | LYS_A_28_C | -4.434 | -5.095 | 0.114 | -4.442 | -5.715 | -1.284 | 1.53 | 156.08 | -90.74 |
| 166 | 2jsa-1 | LYS_A_28_C | LYS_A_28_O | -4.442 | -5.715 | -1.284 | -3.454 | -5.625 | -2.012 | 1.23 | 126.27 | 5.20 |
| 167 | 2jsa-1 | LYS_A_28_C | LYS_A_29_N | -4.442 | -5.715 | -1.284 | -5.567 | -6.33 | -1.618 | 1.32 | 104.60 | -151.34 |
| 168 | 2jsa-1 | LYS_A_29_N | LYS_A_29_H | -5.567 | -6.33 | -1.618 | -6.339 | -6.364 | -0.968 | 1.01 | 49.93 | -177.48 |
| 169 | 2jsa-1 | LYS_A_29_N | LYS_A_29_CA | -5.567 | -6.33 | -1.618 | -5.716 | -6.965 | -2.916 | 1.45 | 153.32 | -103.21 |
| 170 | 2jsa-1 | LYS_A_29_CA | LYS_A_29_HA | -5.716 | -6.965 | -2.916 | -4.808 | -7.534 | -3.119 | 1.09 | 100.73 | -32.07 |
| 171 | 2jsa-1 | LYS_A_29_CA | LYS_A_29_C | -5.716 | -6.965 | -2.916 | -5.841 | -5.887 | -3.995 | 1.53 | 134.84 | 96.61 |
| 172 | 2jsa-1 | LYS_A_29_C | LYS_A_29_O | -5.841 | -5.887 | -3.995 | -5.156 | -5.942 | -5.015 | 1.23 | 146.03 | -4.59 |
| 173 | 2jsa-1 | LYS_A_29_C | ASP_A_30_N | -5.841 | -5.887 | -3.995 | -6.722 | -4.933 | -3.733 | 1.32 | 78.59 | 132.72 |
| 174 | 2jsa-1 | ASP_A_30_N | ASP_A_30_H | -6.722 | -4.933 | -3.733 | -7.252 | -4.952 | -2.874 | 1.01 | 31.69 | -177.95 |
| 175 | 2jsa-1 | ASP_A_30_N | ASP_A_30_CA | -6.722 | -4.933 | -3.733 | -6.946 | -3.844 | -4.668 | 1.45 | 130.06 | 101.62 |
| 176 | 2jsa-1 | ASP_A_30_CA | ASP_A_30_HA | -6.946 | -3.844 | -4.668 | -7.455 | -4.232 | -5.55 | 1.09 | 144.03 | -142.68 |
| 177 | 2jsa-1 | ASP_A_30_CA | ASP_A_30_C | -6.946 | -3.844 | -4.668 | -5.601 | -3.237 | -5.069 | 1.53 | 105.20 | 24.29 |
| 178 | 2jsa-1 | ASP_A_30_C | ASP_A_30_O | -5.601 | -3.237 | -5.069 | -5.381 | -2.924 | -6.238 | 1.23 | 161.88 | 54.90 |
| 179 | 2jsa-1 | ASP_A_30_C | PHE_A_31_N | -5.601 | -3.237 | -5.069 | -4.735 | -3.088 | -4.077 | 1.33 | 41.53 | 9.76 |
| 180 | 2jsa-1 | PHE_A_31_N | PHE_A_31_H | -4.735 | -3.088 | -4.077 | -4.983 | -3.367 | -3.139 | 1.01 | 21.70 | -131.63 |
| 181 | 2jsa-1 | PHE_A_31_N | PHE_A_31_CA | -4.735 | -3.088 | -4.077 | -3.417 | -2.524 | -4.312 | 1.45 | 99.31 | 23.17 |
| 182 | 2jsa-1 | PHE_A_31_CA | PHE_A_31_HA | -3.417 | -2.524 | -4.312 | -3.509 | -1.478 | -4.604 | 1.09 | 105.54 | 95.03 |
| 183 | 2jsa-1 | PHE_A_31_CA | PHE_A_31_C | -3.417 | -2.524 | -4.312 | -2.675 | -3.299 | -5.403 | 1.53 | 135.48 | -46.25 |
| 184 | 2jsa-1 | PHE_A_31_C | PHE_A_31_O | -2.675 | -3.299 | -5.403 | -2.343 | -2.742 | -6.448 | 1.23 | 148.18 | 59.20 |
| 185 | 2jsa-1 | PHE_A_31_C | ASP_A_32_N | -2.675 | -3.299 | -5.403 | -2.437 | -4.572 | -5.121 | 1.33 | 77.72 | -79.41 |
| 186 | 2jsa-1 | ASP_A_32_N | ASP_A_32_H | -2.437 | -4.572 | -5.121 | -2.738 | -4.963 | -4.24 | 1.01 | 29.25 | -127.59 |
| 187 | 2jsa-1 | ASP_A_32_N | ASP_A_32_CA | -2.437 | -4.572 | -5.121 | -1.741 | -5.429 | -6.065 | 1.45 | 130.53 | -50.92 |
| 188 | 2jsa-1 | ASP_A_32_CA | ASP_A_32_HA | -1.741 | -5.429 | -6.065 | -0.725 | -5.058 | -6.2 | 1.09 | 97.11 | 20.06 |
| 189 | 2jsa-1 | ASP_A_32_CA | ASP_A_32_C | -1.741 | -5.429 | -6.065 | -2.482 | -5.415 | -7.403 | 1.53 | 151.02 | 178.92 |
| 190 | 2jsa-1 | ASP_A_32_C | ASP_A_32_O | -2.482 | -5.415 | -7.403 | -1.905 | -5.737 | -8.441 | 1.23 | 147.52 | -29.16 |
| 191 | 2jsa-1 | ASP_A_32_C | ALA_A_33_N | -2.482 | -5.415 | -7.403 | -3.751 | -5.038 | -7.337 | 1.33 | 87.15 | 163.45 |

| | | | | | | | | | | | |
|---|---|---|---|---|---|---|---|---|---|---|---|
| 192 | 2jsa-1 | ALA_A_33_N | ALA_A_33_H | -3.751 | -5.038 | -7.337 | -4.161 | -4.787 | -6.449 | 1.01 | 28.43 | 148.53 |
| 193 | 2jsa-1 | ALA_A_33_N | ALA_A_33_CA | -3.751 | -5.038 | -7.337 | -4.577 | -4.978 | -8.53 | 1.45 | 145.23 | 175.85 |
| 194 | 2jsa-1 | ALA_A_33_CA | ALA_A_33_HA | -4.577 | -4.978 | -8.53 | -4.657 | -5.98 | -8.951 | 1.09 | 112.73 | -94.56 |
| 195 | 2jsa-1 | ALA_A_33_CA | ALA_A_33_C | -4.577 | -4.978 | -8.53 | -3.907 | -4.071 | -9.564 | 1.53 | 132.52 | 53.55 |
| 196 | 2jsa-1 | ALA_A_33_C | ALA_A_33_O | -3.907 | -4.071 | -9.564 | -3.39 | -4.549 | -10.573 | 1.23 | 145.09 | -42.76 |
| 197 | 2jsa-1 | ALA_A_33_C | GLU_A_34_N | -3.907 | -4.071 | -9.564 | -3.938 | -2.778 | -9.278 | 1.32 | 77.53 | 91.37 |
| 198 | 2jsa-1 | GLU_A_34_N | GLU_A_34_H | -3.938 | -2.778 | -9.278 | -4.38 | -2.455 | -8.429 | 1.01 | 32.81 | 143.84 |
| 199 | 2jsa-1 | GLU_A_34_N | GLU_A_34_CA | -3.938 | -2.778 | -9.278 | -3.34 | -1.799 | -10.171 | 1.45 | 127.90 | 58.58 |
| 200 | 2jsa-1 | GLU_A_34_CA | GLU_A_34_HA | -3.34 | -1.799 | -10.171 | -3.637 | -2.046 | -11.19 | 1.09 | 159.24 | -140.25 |
| 201 | 2jsa-1 | GLU_A_34_CA | GLU_A_34_C | -3.34 | -1.799 | -10.171 | -1.814 | -1.866 | -10.085 | 1.53 | 86.78 | -2.51 |
| 202 | 2jsa-1 | GLU_A_34_C | GLU_A_34_O | -1.814 | -1.866 | -10.085 | -1.126 | -1.747 | -11.098 | 1.23 | 145.42 | 9.81 |
| 203 | 2jsa-1 | GLU_A_34_C | CYS_A_35_N | -1.814 | -1.866 | -10.085 | -1.329 | -2.056 | -8.867 | 1.32 | 23.15 | -21.39 |
| 204 | 2jsa-1 | CYS_A_35_N | CYS_A_35_H | -1.329 | -2.056 | -8.867 | -1.954 | -2.147 | -8.079 | 1.01 | 38.71 | -171.72 |
| 205 | 2jsa-1 | CYS_A_35_N | CYS_A_35_CA | -1.329 | -2.056 | -8.867 | 0.103 | -2.139 | -8.636 | 1.45 | 80.85 | -3.32 |
| 206 | 2jsa-1 | CYS_A_35_CA | CYS_A_35_HA | 0.103 | -2.139 | -8.636 | 0.532 | -1.146 | -8.776 | 1.09 | 97.37 | 66.63 |
| 207 | 2jsa-1 | CYS_A_35_CA | CYS_A_35_C | 0.103 | -2.139 | -8.636 | 0.704 | -3.081 | -9.682 | 1.53 | 133.11 | -57.46 |
| 208 | 2jsa-1 | CYS_A_35_C | CYS_A_35_O | 0.704 | -3.081 | -9.682 | 1.888 | -2.986 | -9.999 | 1.23 | 104.94 | 4.59 |
| 209 | 2jsa-1 | CYS_A_35_C | LYS_A_36_N | 0.704 | -3.081 | -9.682 | -0.141 | -3.967 | -10.188 | 1.32 | 112.45 | -133.64 |
| 210 | 2jsa-1 | LYS_A_36_N | LYS_A_36_H | -0.141 | -3.967 | -10.188 | -1.102 | -3.987 | -9.879 | 1.01 | 72.18 | -178.81 |
| 211 | 2jsa-1 | LYS_A_36_N | LYS_A_36_CA | -0.141 | -3.967 | -10.188 | 0.292 | -4.925 | -11.192 | 1.45 | 133.68 | -65.68 |
| 212 | 2jsa-1 | LYS_A_36_CA | LYS_A_36_HA | 0.292 | -4.925 | -11.192 | 1.34 | -5.161 | -11.004 | 1.09 | 80.07 | -12.69 |
| 213 | 2jsa-1 | LYS_A_36_CA | LYS_A_36_C | 0.292 | -4.925 | -11.192 | 0.202 | -4.282 | -12.577 | 1.53 | 154.88 | 97.97 |
| 214 | 2jsa-1 | LYS_A_36_C | LYS_A_36_O | 0.202 | -4.282 | -12.577 | 1.207 | -4.156 | -13.274 | 1.23 | 124.53 | 7.15 |
| 215 | 2jsa-1 | LYS_A_36_C | LYS_A_37_N | 0.202 | -4.282 | -12.577 | -1.013 | -3.891 | -12.935 | 1.33 | 105.67 | 162.16 |
| 216 | 2jsa-1 | LYS_A_37_N | LYS_A_37_H | -1.013 | -3.891 | -12.935 | -1.795 | -4.026 | -12.31 | 1.01 | 51.78 | -170.21 |
| 217 | 2jsa-1 | LYS_A_37_N | LYS_A_37_CA | -1.013 | -3.891 | -12.935 | -1.248 | -3.264 | -14.224 | 1.45 | 152.55 | 110.55 |
| 218 | 2jsa-1 | LYS_A_37_CA | LYS_A_37_HA | -1.248 | -3.264 | -14.224 | -0.801 | -3.895 | -14.992 | 1.09 | 134.80 | -54.69 |
| 219 | 2jsa-1 | LYS_A_37_CA | LYS_A_37_C | -1.248 | -3.264 | -14.224 | -0.537 | -1.909 | -14.265 | 1.53 | 91.53 | 62.31 |
| 220 | 2jsa-1 | LYS_A_37_C | LYS_A_37_O | -0.537 | -1.909 | -14.265 | 0.376 | -1.707 | -15.064 | 1.23 | 130.51 | 12.48 |
| 221 | 2jsa-1 | LYS_A_37_C | LEU_A_38_N | -0.537 | -1.909 | -14.265 | -0.983 | -1.017 | -13.393 | 1.32 | 48.83 | 116.57 |
| 222 | 2jsa-1 | LEU_A_38_N | LEU_A_38_H | -0.983 | -1.017 | -13.393 | -1.738 | -1.252 | -12.765 | 1.01 | 51.54 | -162.71 |
| 223 | 2jsa-1 | LEU_A_38_N | LEU_A_38_CA | -0.983 | -1.017 | -13.393 | -0.401 | 0.312 | -13.319 | 1.45 | 87.08 | 66.35 |
| 224 | 2jsa-1 | LEU_A_38_CA | LEU_A_38_HA | -0.401 | 0.312 | -13.319 | -0.784 | 0.895 | -14.156 | 1.09 | 140.19 | 123.30 |
| 225 | 2jsa-1 | LEU_A_38_CA | LEU_A_38_C | -0.401 | 0.312 | -13.319 | 1.117 | 0.21 | -13.48 | 1.53 | 96.04 | -3.84 |
| 226 | 2jsa-1 | LEU_A_38_C | LEU_A_38_O | 1.117 | 0.21 | -13.48 | 1.716 | 0.967 | -14.243 | 1.23 | 128.32 | 51.65 |
| 227 | 2jsa-1 | LEU_A_38_C | PHE_A_39_N | 1.117 | 0.21 | -13.48 | 1.695 | -0.732 | -12.75 | 1.32 | 56.55 | -58.47 |
| 228 | 2jsa-1 | PHE_A_39_N | PHE_A_39_H | 1.695 | -0.732 | -12.75 | 1.14 | -1.319 | -12.144 | 1.01 | 53.12 | -133.39 |
| 229 | 2jsa-1 | PHE_A_39_N | PHE_A_39_CA | 1.695 | -0.732 | -12.75 | 3.132 | -0.943 | -12.803 | 1.45 | 92.09 | -8.35 |
| 230 | 2jsa-1 | PHE_A_39_CA | PHE_A_39_HA | 3.132 | -0.943 | -12.803 | 3.595 | -0.185 | -13.436 | 1.09 | 125.48 | 58.58 |
| 231 | 2jsa-1 | PHE_A_39_CA | PHE_A_39_C | 3.132 | -0.943 | -12.803 | 3.462 | -2.336 | -13.343 | 1.53 | 110.67 | -76.67 |
| 232 | 2jsa-1 | PHE_A_39_C | PHE_A_39_O | 3.462 | -2.336 | -13.343 | 4.256 | -3.064 | -12.749 | 1.23 | 61.13 | -42.52 |
| 233 | 2jsa-1 | PHE_A_39_C | HIS_A_40_N | 3.462 | -2.336 | -13.343 | 2.835 | -2.665 | -14.463 | 1.33 | 147.70 | -152.31 |
| 234 | 2jsa-1 | HIS_A_40_N | HIS_A_40_H | 2.835 | -2.665 | -14.463 | 2.194 | -2.016 | -14.896 | 1.01 | 115.39 | 134.64 |
| 235 | 2jsa-1 | HIS_A_40_N | HIS_A_40_CA | 2.835 | -2.665 | -14.463 | 3.053 | -3.958 | -15.09 | 1.45 | 115.56 | -80.43 |
| 236 | 2jsa-1 | HIS_A_40_CA | HIS_A_40_HA | 3.053 | -3.958 | -15.09 | 3.113 | -4.709 | -14.302 | 1.09 | 43.71 | -85.43 |
| 237 | 2jsa-1 | HIS_A_40_CA | HIS_A_40_C | 3.053 | -3.958 | -15.09 | 4.39 | -3.95 | -15.832 | 1.53 | 119.03 | 0.34 |
| 238 | 2jsa-1 | HIS_A_40_C | HIS_A_40_O | 4.39 | -3.95 | -15.832 | 4.976 | -5.004 | -16.074 | 1.23 | 101.35 | -60.93 |
| 239 | 2jsa-1 | HIS_A_40_C | THR_A_41_N | 4.39 | -3.95 | -15.832 | 4.835 | -2.749 | -16.172 | 1.33 | 104.87 | 69.67 |
| 240 | 2jsa-1 | THR_A_41_N | THR_A_41_H | 4.835 | -2.749 | -16.172 | 4.301 | -1.924 | -15.939 | 1.01 | 76.66 | 122.91 |
| 241 | 2jsa-1 | THR_A_41_N | THR_A_41_CA | 4.835 | -2.749 | -16.172 | 6.093 | -2.59 | -16.882 | 1.45 | 119.25 | 7.20 |
| 242 | 2jsa-1 | THR_A_41_CA | THR_A_41_HA | 6.093 | -2.59 | -16.882 | 6.457 | -3.575 | -17.175 | 1.09 | 105.59 | -69.72 |
| 243 | 2jsa-1 | THR_A_41_CA | THR_A_41_C | 6.093 | -2.59 | -16.882 | 7.143 | -1.953 | -15.97 | 1.53 | 53.40 | 31.24 |
| 244 | 2jsa-1 | THR_A_41_C | THR_A_41_O | 7.143 | -1.953 | -15.97 | 7.885 | -1.069 | -16.394 | 1.23 | 110.17 | 49.99 |
| 245 | 2jsa-1 | THR_A_41_C | ILE_A_42_N | 7.143 | -1.953 | -15.97 | 7.172 | -2.427 | -14.733 | 1.33 | 21.00 | -86.50 |
| 246 | 2jsa-1 | ILE_A_42_N | ILE_A_42_H | 7.172 | -2.427 | -14.733 | 6.532 | -3.156 | -14.452 | 1.01 | 73.85 | -131.28 |
| 247 | 2jsa-1 | ILE_A_42_N | ILE_A_42_CA | 7.172 | -2.427 | -14.733 | 8.119 | -1.915 | -13.757 | 1.45 | 47.80 | 28.40 |
| 248 | 2jsa-1 | ILE_A_42_CA | ILE_A_42_HA | 8.119 | -1.915 | -13.757 | 8.728 | -1.153 | -14.245 | 1.09 | 116.58 | 51.37 |
| 249 | 2jsa-1 | ILE_A_42_CA | ILE_A_42_C | 8.119 | -1.915 | -13.757 | 9.052 | -3.044 | -13.317 | 1.53 | 73.28 | -50.43 |
| 250 | 2jsa-1 | ILE_A_42_C | ILE_A_42_O | 9.052 | -3.044 | -13.317 | 8.626 | -4.191 | -13.185 | 1.23 | 83.84 | -110.38 |
| 251 | 2jsa-1 | ILE_A_42_C | PRO_A_43_N | 9.052 | -3.044 | -13.317 | 10.341 | -2.671 | -13.096 | 1.36 | 80.65 | 16.14 |
| 252 | 2jsa-1 | PRO_A_43_N | PRO_A_43_CA | 10.341 | -2.671 | -13.096 | 11.339 | -3.64 | -12.674 | 1.45 | 73.12 | -44.16 |
| 253 | 2jsa-1 | PRO_A_43_CA | PRO_A_43_HA | 11.339 | -3.64 | -12.674 | 11.253 | -4.549 | -13.27 | 1.09 | 123.13 | -95.40 |
| 254 | 2jsa-1 | PRO_A_43_CA | PRO_A_43_C | 11.339 | -3.64 | -12.674 | 11.156 | -4.007 | -11.2 | 1.53 | 15.55 | -116.50 |
| 255 | 2jsa-1 | PRO_A_43_C | PRO_A_43_O | 11.156 | -4.007 | -11.2 | 11.515 | -3.233 | -10.314 | 1.23 | 43.92 | 65.12 |
| 256 | 2jsa-1 | PRO_A_43_C | PHE_A_44_N | 11.156 | -4.007 | -11.2 | 10.599 | -5.189 | -10.982 | 1.32 | 80.53 | -115.23 |
| 257 | 2jsa-1 | PHE_A_44_N | PHE_A_44_H | 10.599 | -5.189 | -10.982 | 10.33 | -5.775 | -11.759 | 1.01 | 140.31 | -114.66 |
| 258 | 2jsa-1 | PHE_A_44_N | PHE_A_44_CA | 10.599 | -5.189 | -10.982 | 10.364 | -5.669 | -9.631 | 1.45 | 21.58 | -116.09 |
| 259 | 2jsa-1 | PHE_A_44_CA | PHE_A_44_HA | 10.364 | -5.669 | -9.631 | 10.203 | -6.747 | -9.644 | 1.09 | 90.68 | -98.49 |
| 260 | 2jsa-1 | PHE_A_44_CA | PHE_A_44_C | 10.364 | -5.669 | -9.631 | 9.153 | -4.975 | -9.005 | 1.53 | 65.84 | 150.18 |
| 261 | 2jsa-1 | PHE_A_44_C | PHE_A_44_O | 9.153 | -4.975 | -9.005 | 8.797 | -5.251 | -7.861 | 1.23 | 21.49 | -142.21 |
| 262 | 2jsa-1 | PHE_A_44_C | GLY_A_45_N | 9.153 | -4.975 | -9.005 | 8.552 | -4.087 | -9.784 | 1.33 | 126.00 | 124.09 |
| 263 | 2jsa-1 | GLY_A_45_N | GLY_A_45_H | 8.552 | -4.087 | -9.784 | 8.899 | -3.91 | -10.716 | 1.01 | 157.32 | 27.03 |
| 264 | 2jsa-1 | GLY_A_45_N | GLY_A_45_CA | 8.552 | -4.087 | -9.784 | 7.388 | -3.351 | -9.32 | 1.45 | 71.38 | 147.69 |
| 265 | 2jsa-1 | GLY_A_45_CA | GLY_A_45_C | 7.388 | -3.351 | -9.32 | 6.3 | -4.305 | -8.821 | 1.53 | 70.97 | -138.75 |
| 266 | 2jsa-1 | GLY_A_45_C | GLY_A_45_O | 6.3 | -4.305 | -8.821 | 5.969 | -4.309 | -7.637 | 1.23 | 15.62 | -179.31 |
| 267 | 2jsa-1 | GLY_A_45_C | THR_A_46_N | 6.3 | -4.305 | -8.821 | 5.774 | -5.089 | -9.75 | 1.32 | 134.54 | -123.86 |
| 268 | 2jsa-1 | THR_A_46_N | THR_A_46_H | 5.774 | -5.089 | -9.75 | 6.095 | -5.031 | -10.706 | 1.01 | 161.16 | 10.24 |
| 269 | 2jsa-1 | THR_A_46_N | THR_A_46_CA | 5.774 | -5.089 | -9.75 | 4.731 | -6.045 | -9.419 | 1.45 | 76.83 | -137.49 |
| 270 | 2jsa-1 | THR_A_46_CA | THR_A_46_HA | 4.731 | -6.045 | -9.419 | 3.875 | -5.503 | -9.019 | 1.09 | 68.46 | 147.66 |
| 271 | 2jsa-1 | THR_A_46_CA | THR_A_46_C | 4.731 | -6.045 | -9.419 | 5.218 | -7.016 | -8.342 | 1.53 | 45.25 | -63.36 |
| 272 | 2jsa-1 | THR_A_46_C | THR_A_46_O | 5.218 | -7.016 | -8.342 | 4.428 | -7.501 | -7.535 | 1.23 | 48.96 | -148.45 |
| 273 | 2jsa-1 | THR_A_46_C | ARG_A_47_N | 5.218 | -7.016 | -8.342 | 6.518 | -7.27 | -8.365 | 1.32 | 90.99 | -11.06 |
| 274 | 2jsa-1 | ARG_A_47_N | ARG_A_47_H | 6.518 | -7.27 | -8.365 | 7.109 | -6.835 | -9.059 | 1.01 | 133.40 | 36.35 |
| 275 | 2jsa-1 | ARG_A_47_N | ARG_A_47_CA | 6.518 | -7.27 | -8.365 | 7.121 | -8.175 | -7.401 | 1.45 | 48.44 | -56.32 |
| 276 | 2jsa-1 | ARG_A_47_CA | ARG_A_47_HA | 7.121 | -8.175 | -7.401 | 6.597 | -9.129 | -7.45 | 1.09 | 92.58 | -118.78 |
| 277 | 2jsa-1 | ARG_A_47_CA | ARG_A_47_C | 7.121 | -8.175 | -7.401 | 6.99 | -7.606 | -5.986 | 1.53 | 22.42 | 102.97 |
| 278 | 2jsa-1 | ARG_A_47_C | ARG_A_47_O | 6.99 | -7.606 | -5.986 | 6.443 | -8.259 | -5.099 | 1.23 | 43.84 | -129.95 |
| 279 | 2jsa-1 | ARG_A_47_C | GLU_A_48_N | 6.99 | -7.606 | -5.986 | 7.502 | -6.396 | -5.82 | 1.32 | 82.80 | 67.06 |
| 280 | 2jsa-1 | GLU_A_48_N | GLU_A_48_H | 7.502 | -6.396 | -5.82 | 7.938 | -5.917 | -6.595 | 1.01 | 140.11 | 47.69 |
| 281 | 2jsa-1 | GLU_A_48_N | GLU_A_48_CA | 7.502 | -6.396 | -5.82 | 7.449 | -5.732 | -4.529 | 1.45 | 27.29 | 94.56 |
| 282 | 2jsa-1 | GLU_A_48_CA | GLU_A_48_HA | 7.449 | -5.732 | -4.529 | 7.882 | -6.403 | -3.787 | 1.09 | 47.10 | -57.17 |
| 283 | 2jsa-1 | GLU_A_48_CA | GLU_A_48_C | 7.449 | -5.732 | -4.529 | 5.998 | -5.451 | -4.135 | 1.53 | 75.07 | 169.04 |
| 284 | 2jsa-1 | GLU_A_48_C | GLU_A_48_O | 5.998 | -5.451 | -4.135 | 5.667 | -5.419 | -2.95 | 1.23 | 15.68 | 174.48 |
| 285 | 2jsa-1 | GLU_A_48_C | CYS_A_49_N | 5.998 | -5.451 | -4.135 | 5.17 | -5.255 | -5.15 | 1.32 | 140.03 | 166.68 |
| 286 | 2jsa-1 | CYS_A_49_N | CYS_A_49_H | 5.17 | -5.255 | -5.15 | 5.512 | -5.296 | -6.1 | 1.01 | 160.07 | -6.84 |
| 287 | 2jsa-1 | CYS_A_49_N | CYS_A_49_CA | 5.17 | -5.255 | -5.15 | 3.761 | -4.979 | -4.925 | 1.45 | 81.09 | 168.92 |

| | | | | | | | | | | |
|---|---|---|---|---|---|---|---|---|---|---|
| 288 | 2jsa-1 | CYS_A_49_CA | CYS_A_49_HA | 3.761 | -4.979 | -4.925 | 3.674 | -3.993 | -4.468 | 1.09 | 65.22 | 95.04 |
| 289 | 2jsa-1 | CYS_A_49_CA | CYS_A_49_C | 3.761 | -4.979 | -4.925 | 3.218 | -6.02 | -3.944 | 1.53 | 50.12 | -117.55 |
| 290 | 2jsa-1 | CYS_A_49_C | CYS_A_49_O | 3.218 | -6.02 | -3.944 | 2.88 | -5.691 | -2.808 | 1.23 | 22.55 | 135.77 |
| 291 | 2jsa-1 | CYS_A_49_C | ASP_A_50_N | 3.218 | -6.02 | -3.944 | 3.151 | -7.255 | -4.419 | 1.32 | 111.01 | -93.11 |
| 292 | 2jsa-1 | ASP_A_50_N | ASP_A_50_H | 3.151 | -7.255 | -4.419 | 3.446 | -7.453 | -5.365 | 1.01 | 159.42 | -33.87 |
| 293 | 2jsa-1 | ASP_A_50_N | ASP_A_50_CA | 3.151 | -7.255 | -4.419 | 2.655 | -8.347 | -3.598 | 1.45 | 55.61 | -114.43 |
| 294 | 2jsa-1 | ASP_A_50_CA | ASP_A_50_HA | 2.655 | -8.347 | -3.598 | 1.607 | -8.166 | -3.36 | 1.09 | 77.39 | 170.20 |
| 295 | 2jsa-1 | ASP_A_50_CA | ASP_A_50_C | 2.655 | -8.347 | -3.598 | 3.474 | -8.423 | -2.309 | 1.53 | 32.54 | -5.30 |
| 296 | 2jsa-1 | ASP_A_50_C | ASP_A_50_O | 3.474 | -8.423 | -2.309 | 2.994 | -8.921 | -1.292 | 1.23 | 34.22 | -133.95 |
| 297 | 2jsa-1 | ASP_A_50_C | HIS_A_51_N | 3.474 | -8.423 | -2.309 | 4.698 | -7.922 | -2.393 | 1.33 | 93.63 | 22.26 |
| 298 | 2jsa-1 | HIS_A_51_N | HIS_A_51_H | 4.698 | -7.922 | -2.393 | 5.028 | -7.528 | -3.263 | 1.01 | 149.43 | 50.05 |
| 299 | 2jsa-1 | HIS_A_51_N | HIS_A_51_CA | 4.698 | -7.922 | -2.393 | 5.589 | -7.927 | -1.245 | 1.45 | 37.82 | -0.32 |
| 300 | 2jsa-1 | HIS_A_51_CA | HIS_A_51_HA | 5.589 | -7.927 | -1.245 | 5.382 | -8.824 | -0.662 | 1.09 | 57.65 | -102.99 |
| 301 | 2jsa-1 | HIS_A_51_CA | HIS_A_51_C | 5.589 | -7.927 | -1.245 | 5.294 | -6.713 | -0.361 | 1.53 | 54.72 | 103.66 |
| 302 | 2jsa-1 | HIS_A_51_C | HIS_A_51_O | 5.294 | -6.713 | -0.361 | 5.984 | -6.481 | 0.631 | 1.23 | 36.27 | 18.58 |
| 303 | 2jsa-1 | HIS_A_51_C | TYR_A_52_N | 5.294 | -6.713 | -0.361 | 4.268 | -5.972 | -0.752 | 1.32 | 107.17 | 144.16 |
| 304 | 2jsa-1 | TYR_A_52_N | TYR_A_52_H | 4.268 | -5.972 | -0.752 | 3.746 | -6.223 | -1.579 | 1.01 | 144.99 | -154.32 |
| 305 | 2jsa-1 | TYR_A_52_N | TYR_A_52_CA | 4.268 | -5.972 | -0.752 | 3.873 | -4.788 | -0.007 | 1.45 | 59.17 | 108.45 |
| 306 | 2jsa-1 | TYR_A_52_CA | TYR_A_52_HA | 3.873 | -4.788 | -0.007 | 4.618 | -4.554 | 0.754 | 1.09 | 45.74 | 17.44 |
| 307 | 2jsa-1 | TYR_A_52_CA | TYR_A_52_C | 3.873 | -4.788 | -0.007 | 2.505 | -4.985 | 0.649 | 1.53 | 64.61 | -171.81 |
| 308 | 2jsa-1 | TYR_A_52_C | TYR_A_52_O | 2.505 | -4.985 | 0.649 | 2.321 | -4.655 | 1.82 | 1.23 | 17.88 | 119.14 |
| 309 | 2jsa-1 | TYR_A_52_C | VAL_A_53_N | 2.505 | -4.985 | 0.649 | 1.58 | -5.52 | -0.134 | 1.32 | 126.23 | -149.96 |
| 310 | 2jsa-1 | VAL_A_53_N | VAL_A_53_H | 1.58 | -5.52 | -0.134 | 1.802 | -5.766 | -1.088 | 1.01 | 160.85 | -47.94 |
| 311 | 2jsa-1 | VAL_A_53_N | VAL_A_53_CA | 1.58 | -5.52 | -0.134 | 0.234 | -5.764 | 0.356 | 1.45 | 70.29 | -169.73 |
| 312 | 2jsa-1 | VAL_A_53_CA | VAL_A_53_HA | 0.234 | -5.764 | 0.356 | -0.201 | -4.811 | 0.657 | 1.09 | 73.97 | 114.53 |
| 313 | 2jsa-1 | VAL_A_53_CA | VAL_A_53_C | 0.234 | -5.764 | 0.356 | 0.301 | -6.666 | 1.59 | 1.53 | 36.24 | -85.75 |
| 314 | 2jsa-1 | VAL_A_53_C | VAL_A_53_O | 0.301 | -6.666 | 1.59 | -0.659 | -6.751 | 2.354 | 1.23 | 51.60 | -174.94 |
| 315 | 2jsa-1 | VAL_A_53_C | ASN_A_54_N | 0.301 | -6.666 | 1.59 | 1.445 | -7.317 | 1.747 | 1.33 | 83.20 | -29.64 |
| 316 | 2jsa-1 | ASN_A_54_N | ASN_A_54_H | 1.445 | -7.317 | 1.747 | 2.194 | -7.198 | 1.079 | 1.01 | 131.37 | 9.03 |
| 317 | 2jsa-1 | ASN_A_54_N | ASN_A_54_CA | 1.445 | -7.317 | 1.747 | 1.65 | -8.21 | 2.875 | 1.45 | 39.09 | -77.07 |
| 318 | 2jsa-1 | ASN_A_54_CA | ASN_A_54_HA | 1.65 | -8.21 | 2.875 | 0.889 | -8.012 | 3.63 | 1.09 | 46.16 | 165.42 |
| 319 | 2jsa-1 | ASN_A_54_CA | ASN_A_54_C | 1.65 | -8.21 | 2.875 | 3.041 | -7.968 | 3.465 | 1.53 | 67.32 | 9.87 |
| 320 | 2jsa-1 | ASN_A_54_C | ASN_A_54_O | 3.041 | -7.968 | 3.465 | 3.923 | -8.817 | 3.352 | 1.23 | 95.27 | -43.91 |
| 321 | 2jsa-1 | ASN_A_54_C | SER_A_55_N | 3.041 | -7.968 | 3.465 | 3.193 | -6.805 | 4.081 | 1.32 | 62.29 | 82.55 |
| 322 | 2jsa-1 | SER_A_55_N | SER_A_55_H | 3.193 | -6.805 | 4.081 | 2.423 | -6.154 | 4.136 | 1.01 | 86.88 | 139.79 |
| 323 | 2jsa-1 | SER_A_55_N | SER_A_55_CA | 3.193 | -6.805 | 4.081 | 4.461 | -6.44 | 4.689 | 1.45 | 65.26 | 16.06 |
| 324 | 2jsa-1 | SER_A_55_CA | SER_A_55_HA | 4.461 | -6.44 | 4.689 | 4.624 | -7.084 | 5.553 | 1.09 | 37.56 | -75.80 |
| 325 | 2jsa-1 | SER_A_55_CA | SER_A_55_C | 4.461 | -6.44 | 4.689 | 4.416 | -4.987 | 5.164 | 1.53 | 71.91 | 91.77 |
| 326 | 2jsa-1 | SER_A_55_C | SER_A_55_O | 4.416 | -4.987 | 5.164 | 5.014 | -4.644 | 6.183 | 1.23 | 34.08 | 29.84 |
| 327 | 2jsa-1 | SER_A_55_C | LYS_A_56_N | 4.416 | -4.987 | 5.164 | 3.7 | -4.171 | 4.404 | 1.33 | 125.00 | 131.27 |
| 328 | 2jsa-1 | LYS_A_56_N | LYS_A_56_H | 3.7 | -4.171 | 4.404 | 3.235 | -4.522 | 3.58 | 1.01 | 144.74 | -142.95 |
| 329 | 2jsa-1 | LYS_A_56_N | LYS_A_56_CA | 3.7 | -4.171 | 4.404 | 3.568 | -2.763 | 4.734 | 1.45 | 76.86 | 95.36 |
| 330 | 2jsa-1 | LYS_A_56_CA | LYS_A_56_HA | 3.568 | -2.763 | 4.734 | 3.953 | -2.616 | 5.743 | 1.09 | 22.22 | 20.90 |
| 331 | 2jsa-1 | LYS_A_56_CA | LYS_A_56_C | 3.568 | -2.763 | 4.734 | 2.087 | -2.381 | 4.741 | 1.53 | 89.74 | 165.54 |
| 332 | 2jsa-1 | LYS_A_56_C | LYS_A_56_O | 2.087 | -2.381 | 4.741 | 1.746 | -1.2 | 4.782 | 1.23 | 88.09 | 106.11 |
| 333 | 2jsa-1 | LYS_A_56_C | VAL_A_57_N | 2.087 | -2.381 | 4.741 | 1.245 | -3.403 | 4.699 | 1.32 | 91.82 | -129.48 |
| 334 | 2jsa-1 | VAL_A_57_N | VAL_A_57_H | 1.245 | -3.403 | 4.699 | 1.596 | -4.349 | 4.666 | 1.01 | 91.87 | -69.64 |
| 335 | 2jsa-1 | VAL_A_57_N | VAL_A_57_CA | 1.245 | -3.403 | 4.699 | -0.192 | -3.19 | 4.7 | 1.45 | 89.96 | 171.57 |
| 336 | 2jsa-1 | VAL_A_57_CA | VAL_A_57_HA | -0.192 | -3.19 | 4.7 | -0.388 | -2.166 | 4.381 | 1.09 | 107.01 | 100.84 |
| 337 | 2jsa-1 | VAL_A_57_CA | VAL_A_57_C | -0.192 | -3.19 | 4.7 | -0.728 | -3.355 | 6.123 | 1.53 | 21.51 | -162.89 |
| 338 | 2jsa-1 | VAL_A_57_C | VAL_A_57_O | -0.728 | -3.355 | 6.123 | -1.529 | -2.545 | 6.587 | 1.23 | 67.84 | 134.68 |
| 339 | 2jsa-1 | VAL_A_57_C | ASP_A_58_N | -0.728 | -3.355 | 6.123 | -0.263 | -4.409 | 6.778 | 1.33 | 60.38 | -66.19 |
| 340 | 2jsa-1 | ASP_A_58_N | ASP_A_58_H | -0.263 | -4.409 | 6.778 | 0.395 | -5.034 | 6.334 | 1.01 | 116.07 | -43.53 |
| 341 | 2jsa-1 | ASP_A_58_N | ASP_A_58_CA | -0.263 | -4.409 | 6.778 | -0.685 | -4.691 | 8.139 | 1.45 | 20.45 | -146.25 |
| 342 | 2jsa-1 | ASP_A_58_CA | ASP_A_58_HA | -0.685 | -4.691 | 8.139 | -1.745 | -4.942 | 8.138 | 1.09 | 90.05 | -166.68 |
| 343 | 2jsa-1 | ASP_A_58_CA | ASP_A_58_C | -0.685 | -4.691 | 8.139 | -0.444 | -3.455 | 9.009 | 1.53 | 55.36 | 78.97 |
| 344 | 2jsa-1 | ASP_A_58_C | ASP_A_58_O | -0.444 | -3.455 | 9.009 | -1.297 | -3.081 | 9.812 | 1.23 | 49.23 | 156.32 |
| 345 | 2jsa-1 | ASP_A_58_C | PRO_A_59_N | -0.444 | -3.455 | 9.009 | 0.754 | -2.841 | 8.815 | 1.36 | 98.20 | 27.14 |
| 346 | 2jsa-1 | PRO_A_59_N | PRO_A_59_CA | 0.754 | -2.841 | 8.815 | 1.118 | -1.655 | 9.572 | 1.45 | 58.61 | 72.94 |
| 347 | 2jsa-1 | PRO_A_59_CA | PRO_A_59_HA | 1.118 | -1.655 | 9.572 | 0.876 | -1.796 | 10.625 | 1.09 | 14.89 | -149.77 |
| 348 | 2jsa-1 | PRO_A_59_CA | PRO_A_59_C | 1.118 | -1.655 | 9.572 | 0.365 | -0.425 | 9.062 | 1.53 | 109.48 | 121.47 |
| 349 | 2jsa-1 | PRO_A_59_C | PRO_A_59_O | 0.365 | -0.425 | 9.062 | -0.321 | 0.249 | 9.828 | 1.23 | 51.46 | 135.51 |
| 350 | 2jsa-1 | PRO_A_59_C | ILE_A_60_N | 0.365 | -0.425 | 9.062 | 0.518 | -0.171 | 7.77 | 1.33 | 167.07 | 58.94 |
| 351 | 2jsa-1 | ILE_A_60_N | ILE_A_60_H | 0.518 | -0.171 | 7.77 | 1.097 | -0.77 | 7.199 | 1.01 | 124.43 | -45.97 |
| 352 | 2jsa-1 | ILE_A_60_N | ILE_A_60_CA | 0.518 | -0.171 | 7.77 | -0.139 | 0.966 | 7.149 | 1.45 | 115.31 | 120.02 |
| 353 | 2jsa-1 | ILE_A_60_CA | ILE_A_60_HA | -0.139 | 0.966 | 7.149 | 0.342 | 1.874 | 7.513 | 1.09 | 70.49 | 62.09 |
| 354 | 2jsa-1 | ILE_A_60_CA | ILE_A_60_C | -0.139 | 0.966 | 7.149 | -1.604 | 1.007 | 7.59 | 1.53 | 73.25 | 178.40 |
| 355 | 2jsa-1 | ILE_A_60_C | ILE_A_60_O | -1.604 | 1.007 | 7.59 | -2.069 | 2.017 | 8.115 | 1.23 | 64.73 | 114.72 |
| 356 | 2jsa-1 | ILE_A_60_C | ILE_A_61_N | -1.604 | 1.007 | 7.59 | -2.289 | -0.104 | 7.36 | 1.33 | 99.99 | -121.66 |
| 357 | 2jsa-1 | ILE_A_61_N | ILE_A_61_H | -2.289 | -0.104 | 7.36 | -1.84 | -0.898 | 6.925 | 1.01 | 115.50 | -60.51 |
| 358 | 2jsa-1 | ILE_A_61_N | ILE_A_61_CA | -2.289 | -0.104 | 7.36 | -3.691 | -0.207 | 7.727 | 1.45 | 75.37 | -175.80 |
| 359 | 2jsa-1 | ILE_A_61_CA | ILE_A_61_HA | -3.691 | -0.207 | 7.727 | -4.223 | 0.625 | 7.266 | 1.09 | 115.02 | 122.60 |
| 360 | 2jsa-1 | ILE_A_61_CA | ILE_A_61_C | -3.691 | -0.207 | 7.727 | -3.828 | -0.063 | 9.244 | 1.53 | 7.46 | 133.57 |
| 361 | 2jsa-1 | ILE_A_61_C | ILE_A_61_O | -3.828 | -0.063 | 9.244 | -4.901 | 0.271 | 9.744 | 1.23 | 66.01 | 162.71 |
| 362 | 2jsa-1 | ILE_A_61_C | HIS_A_62_N | -3.828 | -0.063 | 9.244 | -2.727 | -0.321 | 9.933 | 1.32 | 58.65 | -13.19 |
| 363 | 2jsa-1 | HIS_A_62_N | HIS_A_62_H | -2.727 | -0.321 | 9.933 | -1.879 | -0.591 | 9.455 | 1.01 | 118.24 | -17.66 |
| 364 | 2jsa-1 | HIS_A_62_N | HIS_A_62_CA | -2.727 | -0.321 | 9.933 | -2.711 | -0.224 | 11.383 | 1.45 | 3.88 | 80.63 |
| 365 | 2jsa-1 | HIS_A_62_CA | HIS_A_62_HA | -2.711 | -0.224 | 11.383 | -3.541 | -0.815 | 11.769 | 1.09 | 69.25 | -144.55 |
| 366 | 2jsa-1 | HIS_A_62_CA | HIS_A_62_C | -2.711 | -0.224 | 11.383 | -2.935 | 1.23 | 11.802 | 1.53 | 74.10 | 98.76 |
| 367 | 2jsa-1 | HIS_A_62_C | HIS_A_62_O | -2.935 | 1.23 | 11.802 | -3.714 | 1.504 | 12.713 | 1.23 | 42.19 | 160.62 |
| 368 | 2jsa-1 | HIS_A_62_C | GLU_A_63_N | -2.935 | 1.23 | 11.802 | -2.237 | 2.124 | 11.117 | 1.33 | 121.13 | 52.02 |
| 369 | 2jsa-1 | GLU_A_63_N | GLU_A_63_H | -2.237 | 2.124 | 11.117 | -1.614 | 1.826 | 10.379 | 1.01 | 136.90 | -25.56 |
| 370 | 2jsa-1 | GLU_A_63_N | GLU_A_63_CA | -2.237 | 2.124 | 11.117 | -2.349 | 3.543 | 11.407 | 1.45 | 78.48 | 94.51 |
| 371 | 2jsa-1 | GLU_A_63_CA | GLU_A_63_HA | -2.349 | 3.543 | 11.407 | -2.082 | 3.696 | 12.453 | 1.09 | 16.39 | 29.81 |
| 372 | 2jsa-1 | GLU_A_63_CA | GLU_A_63_C | -2.349 | 3.543 | 11.407 | -3.791 | 4.014 | 11.205 | 1.53 | 97.58 | 161.91 |
| 373 | 2jsa-1 | GLU_A_63_C | GLU_A_63_O | -3.791 | 4.014 | 11.205 | -4.429 | 4.493 | 12.141 | 1.23 | 40.44 | 143.10 |
| 374 | 2jsa-1 | GLU_A_63_C | LEU_A_64_N | -3.791 | 4.014 | 11.205 | -4.263 | 3.861 | 9.976 | 1.33 | 158.01 | -162.04 |
| 375 | 2jsa-1 | LEU_A_64_N | LEU_A_64_H | -4.263 | 3.861 | 9.976 | -3.681 | 3.459 | 9.255 | 1.01 | 135.55 | -34.63 |
| 376 | 2jsa-1 | LEU_A_64_N | LEU_A_64_CA | -4.263 | 3.861 | 9.976 | -5.617 | 4.264 | 9.638 | 1.45 | 103.46 | 163.43 |
| 377 | 2jsa-1 | LEU_A_64_CA | LEU_A_64_HA | -5.617 | 4.264 | 9.638 | -5.718 | 5.326 | 9.86 | 1.09 | 78.24 | 95.43 |
| 378 | 2jsa-1 | LEU_A_64_CA | LEU_A_64_C | -5.617 | 4.264 | 9.638 | -6.606 | 3.506 | 10.527 | 1.53 | 54.49 | -142.53 |
| 379 | 2jsa-1 | LEU_A_64_C | LEU_A_64_O | -6.606 | 3.506 | 10.527 | -7.767 | 3.896 | 10.641 | 1.23 | 84.68 | 161.43 |
| 380 | 2jsa-1 | LEU_A_64_C | GLU_A_65_N | -6.606 | 3.506 | 10.527 | -6.11 | 2.437 | 11.132 | 1.32 | 62.82 | -65.11 |
| 381 | 2jsa-1 | GLU_A_65_N | GLU_A_65_H | -6.11 | 2.437 | 11.132 | -5.144 | 2.176 | 10.992 | 1.01 | 97.96 | -15.12 |
| 382 | 2jsa-1 | GLU_A_65_N | GLU_A_65_CA | -6.11 | 2.437 | 11.132 | -6.936 | 1.621 | 12.006 | 1.45 | 53.03 | -135.35 |
| 383 | 2jsa-1 | GLU_A_65_CA | GLU_A_65_HA | -6.936 | 1.621 | 12.006 | -7.903 | 1.483 | 11.524 | 1.09 | 116.26 | -171.88 |

| | | | | | | | | | | | | |
|---|---|---|---|---|---|---|---|---|---|---|---|---|
| 384 | 2jsa-1 | GLU_A_65_CA | GLU_A_65_C | -6.936 | 1.621 | 12.006 | -7.161 | 2.334 | 13.341 | 1.53 | 29.25 | 107.51 |
| 385 | 2jsa-1 | GLU_A_65_C | GLU_A_65_O | -7.161 | 2.334 | 13.341 | -8.207 | 2.172 | 13.967 | 1.23 | 59.40 | -171.20 |
| 386 | 2jsa-1 | GLU_A_65_C | GLY_A_66_N | -7.161 | 2.334 | 13.341 | -6.161 | 3.109 | 13.738 | 1.33 | 72.58 | 37.78 |
| 387 | 2jsa-1 | GLY_A_66_N | GLY_A_66_H | -6.161 | 3.109 | 13.738 | -5.329 | 3.195 | 13.172 | 1.01 | 124.09 | 5.90 |
| 388 | 2jsa-1 | GLY_A_66_N | GLY_A_66_CA | -6.161 | 3.109 | 13.738 | -6.237 | 3.847 | 14.987 | 1.45 | 30.71 | 95.88 |
| 389 | 2jsa-1 | GLY_A_66_CA | GLY_A_66_C | -6.237 | 3.847 | 14.987 | -6.909 | 5.206 | 14.781 | 1.53 | 97.74 | 116.31 |
| 390 | 2jsa-1 | GLY_A_66_C | GLY_A_66_O | -6.909 | 5.206 | 14.781 | -7.65 | 5.673 | 15.645 | 1.23 | 45.39 | 147.78 |
| 391 | 2jsa-1 | GLY_A_66_C | GLY_A_67_N | -6.909 | 5.206 | 14.781 | -6.627 | 5.802 | 13.632 | 1.32 | 150.15 | 64.68 |
| 392 | 2jsa-1 | GLY_A_67_N | GLY_A_67_H | -6.627 | 5.802 | 13.632 | -6.008 | 5.359 | 12.968 | 1.01 | 131.10 | -35.59 |
| 393 | 2jsa-1 | GLY_A_67_N | GLY_A_67_CA | -6.627 | 5.802 | 13.632 | -7.195 | 7.098 | 13.301 | 1.45 | 103.17 | 113.67 |
| 394 | 2jsa-1 | GLY_A_67_CA | GLY_A_67_C | -7.195 | 7.098 | 13.301 | -6.14 | 8.017 | 12.683 | 1.53 | 113.83 | 41.06 |
| 395 | 2jsa-1 | GLY_A_67_C | GLY_A_67_O | -6.14 | 8.017 | 12.683 | -6.065 | 9.197 | 13.021 | 1.23 | 74.05 | 86.36 |
| 396 | 2jsa-1 | GLY_A_67_C | THR_A_68_N | -6.14 | 8.017 | 12.683 | -5.35 | 7.441 | 11.788 | 1.33 | 132.47 | -36.10 |
| 397 | 2jsa-1 | THR_A_68_N | THR_A_68_H | -5.35 | 7.441 | 11.788 | -5.468 | 6.465 | 11.558 | 1.01 | 103.17 | -96.89 |
| 398 | 2jsa-1 | THR_A_68_N | THR_A_68_CA | -5.35 | 7.441 | 11.788 | -4.303 | 8.195 | 11.12 | 1.45 | 117.37 | 35.76 |
| 399 | 2jsa-1 | THR_A_68_CA | THR_A_68_HA | -4.303 | 8.195 | 11.12 | -4.204 | 9.165 | 11.608 | 1.09 | 63.41 | 84.17 |
| 400 | 2jsa-1 | THR_A_68_CA | THR_A_68_C | -4.303 | 8.195 | 11.12 | -4.673 | 8.439 | 9.656 | 1.53 | 163.16 | 146.60 |
| 401 | 2jsa-1 | THR_A_68_C | THR_A_68_O | -4.673 | 8.439 | 9.656 | -5.365 | 7.628 | 9.043 | 1.23 | 119.90 | -130.47 |
| 402 | 2jsa-1 | THR_A_68_C | ALA_A_69_N | -4.673 | 8.439 | 9.656 | -4.196 | 9.561 | 9.137 | 1.33 | 113.06 | 66.97 |
| 403 | 2jsa-1 | ALA_A_69_N | ALA_A_69_H | -4.196 | 9.561 | 9.137 | -3.633 | 10.183 | 9.701 | 1.01 | 56.09 | 47.85 |
| 404 | 2jsa-1 | ALA_A_69_N | ALA_A_69_CA | -4.196 | 9.561 | 9.137 | -4.468 | 9.923 | 7.757 | 1.45 | 161.83 | 126.92 |
| 405 | 2jsa-1 | ALA_A_69_CA | ALA_A_69_HA | -4.468 | 9.923 | 7.757 | -5.483 | 9.616 | 7.508 | 1.09 | 103.21 | -163.17 |
| 406 | 2jsa-1 | ALA_A_69_CA | ALA_A_69_C | -4.468 | 9.923 | 7.757 | -3.498 | 9.178 | 6.838 | 1.53 | 126.92 | -37.53 |
| 407 | 2jsa-1 | ALA_A_69_C | ALA_A_69_O | -3.498 | 9.178 | 6.838 | -2.468 | 8.68 | 7.29 | 1.23 | 68.44 | -25.80 |
| 408 | 2jsa-1 | ALA_A_69_C | PRO_A_70_N | -3.498 | 9.178 | 6.838 | -3.871 | 9.124 | 5.531 | 1.36 | 163.91 | -171.76 |
| 409 | 2jsa-1 | PRO_A_70_N | PRO_A_70_CA | -3.871 | 9.124 | 5.531 | -3.045 | 8.448 | 4.545 | 1.45 | 132.73 | -39.30 |
| 410 | 2jsa-1 | PRO_A_70_CA | PRO_A_70_HA | -3.045 | 8.448 | 4.545 | -2.71 | 7.487 | 4.935 | 1.09 | 69.03 | -70.78 |
| 411 | 2jsa-1 | PRO_A_70_CA | PRO_A_70_C | -3.045 | 8.448 | 4.545 | -1.813 | 9.285 | 4.195 | 1.53 | 103.22 | 34.19 |
| 412 | 2jsa-1 | PRO_A_70_C | PRO_A_70_O | -1.813 | 9.285 | 4.195 | -0.944 | 8.835 | 3.45 | 1.23 | 127.28 | -27.38 |
| 413 | 2jsa-1 | PRO_A_70_C | LYS_A_71_N | -1.813 | 9.285 | 4.195 | -1.778 | 10.488 | 4.748 | 1.32 | 65.32 | 88.33 |
| 414 | 2jsa-1 | LYS_A_71_N | LYS_A_71_H | -1.778 | 10.488 | 4.748 | -2.529 | 10.792 | 5.351 | 1.01 | 53.34 | 157.96 |
| 415 | 2jsa-1 | LYS_A_71_N | LYS_A_71_CA | -1.778 | 10.488 | 4.748 | -0.668 | 11.393 | 4.504 | 1.45 | 99.67 | 39.19 |
| 416 | 2jsa-1 | LYS_A_71_CA | LYS_A_71_HA | -0.668 | 11.393 | 4.504 | -0.041 | 10.955 | 3.727 | 1.09 | 135.45 | -34.94 |
| 417 | 2jsa-1 | LYS_A_71_CA | LYS_A_71_C | -0.668 | 11.393 | 4.504 | 0.179 | 11.509 | 5.773 | 1.53 | 33.97 | 7.80 |
| 418 | 2jsa-1 | LYS_A_71_C | LYS_A_71_O | 0.179 | 11.509 | 5.773 | 1.163 | 12.246 | 5.802 | 1.23 | 88.65 | 36.83 |
| 419 | 2jsa-1 | LYS_A_71_C | ASP_A_72_N | 0.179 | 11.509 | 5.773 | -0.236 | 10.771 | 6.792 | 1.32 | 39.72 | -119.35 |
| 420 | 2jsa-1 | ASP_A_72_N | ASP_A_72_H | -0.236 | 10.771 | 6.792 | -1.055 | 10.189 | 6.698 | 1.01 | 95.34 | -144.60 |
| 421 | 2jsa-1 | ASP_A_72_N | ASP_A_72_CA | -0.236 | 10.771 | 6.792 | 0.472 | 10.782 | 8.061 | 1.45 | 29.16 | 0.89 |
| 422 | 2jsa-1 | ASP_A_72_CA | ASP_A_72_HA | 0.472 | 10.782 | 8.061 | 1.185 | 11.607 | 8.064 | 1.09 | 89.84 | 49.17 |
| 423 | 2jsa-1 | ASP_A_72_CA | ASP_A_72_C | 0.472 | 10.782 | 8.061 | 1.211 | 9.455 | 8.239 | 1.53 | 83.32 | -60.89 |
| 424 | 2jsa-1 | ASP_A_72_C | ASP_A_72_O | 1.211 | 9.455 | 8.239 | 2.208 | 9.387 | 8.957 | 1.23 | 54.30 | -3.90 |
| 425 | 2jsa-1 | ASP_A_72_C | VAL_A_73_N | 1.211 | 9.455 | 8.239 | 0.694 | 8.432 | 7.575 | 1.32 | 120.08 | -116.81 |
| 426 | 2jsa-1 | VAL_A_73_N | VAL_A_73_H | 0.694 | 8.432 | 7.575 | -0.129 | 8.56 | 7.004 | 1.01 | 124.43 | 171.16 |
| 427 | 2jsa-1 | VAL_A_73_N | VAL_A_73_CA | 0.694 | 8.432 | 7.575 | 1.293 | 7.11 | 7.651 | 1.45 | 87.00 | -65.62 |
| 428 | 2jsa-1 | VAL_A_73_CA | VAL_A_73_HA | 1.293 | 7.11 | 7.651 | 1.486 | 6.879 | 8.699 | 1.09 | 16.03 | -50.12 |
| 429 | 2jsa-1 | VAL_A_73_CA | VAL_A_73_C | 1.293 | 7.11 | 7.651 | 2.633 | 7.119 | 6.913 | 1.53 | 118.84 | 0.38 |
| 430 | 2jsa-1 | VAL_A_73_C | VAL_A_73_O | 2.633 | 7.119 | 6.913 | 3.663 | 6.76 | 7.483 | 1.23 | 62.41 | -19.22 |
| 431 | 2jsa-1 | VAL_A_73_C | CYS_A_74_N | 2.633 | 7.119 | 6.913 | 2.577 | 7.533 | 5.656 | 1.32 | 161.62 | 97.70 |
| 432 | 2jsa-1 | CYS_A_74_N | CYS_A_74_H | 2.577 | 7.533 | 5.656 | 1.696 | 7.815 | 5.252 | 1.01 | 113.59 | 162.25 |
| 433 | 2jsa-1 | CYS_A_74_N | CYS_A_74_CA | 2.577 | 7.533 | 5.656 | 3.774 | 7.593 | 4.834 | 1.45 | 124.44 | 2.87 |
| 434 | 2jsa-1 | CYS_A_74_CA | CYS_A_74_HA | 3.774 | 7.593 | 4.834 | 4.321 | 6.658 | 4.956 | 1.09 | 83.57 | -59.67 |
| 435 | 2jsa-1 | CYS_A_74_CA | CYS_A_74_C | 3.774 | 7.593 | 4.834 | 4.65 | 8.74 | 5.343 | 1.53 | 70.57 | 52.63 |
| 436 | 2jsa-1 | CYS_A_74_C | CYS_A_74_O | 4.65 | 8.74 | 5.343 | 4.83 | 9.742 | 4.653 | 1.23 | 124.13 | 79.82 |
| 437 | 2jsa-1 | CYS_A_74_C | THR_A_75_N | 4.65 | 8.74 | 5.343 | 5.17 | 8.554 | 6.547 | 1.32 | 24.64 | -19.68 |
| 438 | 2jsa-1 | THR_A_75_N | THR_A_75_H | 5.17 | 8.554 | 6.547 | 4.979 | 7.704 | 7.058 | 1.01 | 59.61 | -102.66 |
| 439 | 2jsa-1 | THR_A_75_N | THR_A_75_CA | 5.17 | 8.554 | 6.547 | 6.023 | 9.561 | 7.157 | 1.45 | 65.19 | 49.73 |
| 440 | 2jsa-1 | THR_A_75_CA | THR_A_75_HA | 6.023 | 9.561 | 7.157 | 6.897 | 9.711 | 6.522 | 1.09 | 125.61 | 9.74 |
| 441 | 2jsa-1 | THR_A_75_CA | THR_A_75_C | 6.023 | 9.561 | 7.157 | 6.512 | 9.086 | 8.527 | 1.53 | 26.46 | -44.17 |
| 442 | 2jsa-1 | THR_A_75_C | THR_A_75_O | 6.512 | 9.086 | 8.527 | 7.709 | 8.89 | 8.729 | 1.23 | 80.54 | -9.30 |
| 443 | 2jsa-1 | THR_A_75_C | LYS_A_76_N | 6.512 | 9.086 | 8.527 | 5.56 | 8.916 | 9.433 | 1.33 | 46.87 | -169.88 |
| 444 | 2jsa-1 | LYS_A_76_N | LYS_A_76_H | 5.56 | 8.916 | 9.433 | 4.595 | 9.097 | 9.197 | 1.01 | 103.52 | 169.38 |
| 445 | 2jsa-1 | LYS_A_76_N | LYS_A_76_CA | 5.56 | 8.916 | 9.433 | 5.879 | 8.468 | 10.778 | 1.45 | 22.24 | -54.55 |
| 446 | 2jsa-1 | LYS_A_76_CA | LYS_A_76_HA | 5.879 | 8.468 | 10.778 | 6.773 | 8.999 | 11.108 | 1.09 | 72.39 | 30.71 |
| 447 | 2jsa-1 | LYS_A_76_CA | LYS_A_76_C | 5.879 | 8.468 | 10.778 | 6.209 | 6.975 | 10.749 | 1.53 | 91.09 | -77.54 |
| 448 | 2jsa-1 | LYS_A_76_C | LYS_A_76_O | 6.209 | 6.975 | 10.749 | 6.936 | 6.479 | 11.609 | 1.23 | 45.66 | -34.30 |
| 449 | 2jsa-1 | LYS_A_76_C | LEU_A_77_N | 6.209 | 6.975 | 10.749 | 5.66 | 6.299 | 9.75 | 1.33 | 138.92 | -129.08 |
| 450 | 2jsa-1 | LEU_A_77_N | LEU_A_77_H | 5.66 | 6.299 | 9.75 | 5.073 | 6.772 | 9.078 | 1.01 | 131.71 | 141.14 |
| 451 | 2jsa-1 | LEU_A_77_N | LEU_A_77_CA | 5.66 | 6.299 | 9.75 | 5.888 | 4.872 | 9.598 | 1.45 | 96.00 | -80.92 |
| 452 | 2jsa-1 | LEU_A_77_CA | LEU_A_77_HA | 5.888 | 4.872 | 9.598 | 5.964 | 4.434 | 10.594 | 1.09 | 24.05 | -80.16 |
| 453 | 2jsa-1 | LEU_A_77_CA | LEU_A_77_C | 5.888 | 4.872 | 9.598 | 7.224 | 4.646 | 8.888 | 1.53 | 117.65 | -9.60 |
| 454 | 2jsa-1 | LEU_A_77_C | LEU_A_77_O | 7.224 | 4.646 | 8.888 | 7.534 | 3.527 | 8.482 | 1.23 | 109.27 | -74.52 |
| 455 | 2jsa-1 | LEU_A_77_C | ASN_A_78_N | 7.224 | 4.646 | 8.888 | 7.981 | 5.726 | 8.761 | 1.32 | 95.50 | 54.97 |
| 456 | 2jsa-1 | ASN_A_78_N | ASN_A_78_H | 7.981 | 5.726 | 8.761 | 7.663 | 6.615 | 9.12 | 1.01 | 69.18 | 109.68 |
| 457 | 2jsa-1 | ASN_A_78_N | ASN_A_78_CA | 7.981 | 5.726 | 8.761 | 9.277 | 5.66 | 8.108 | 1.45 | 116.71 | -2.92 |
| 458 | 2jsa-1 | ASN_A_78_CA | ASN_A_78_HA | 9.277 | 5.66 | 8.108 | 9.562 | 6.659 | 7.778 | 1.09 | 107.62 | 74.08 |
| 459 | 2jsa-1 | ASN_A_78_CA | ASN_A_78_C | 9.277 | 5.66 | 8.108 | 9.189 | 4.72 | 6.904 | 1.53 | 141.90 | -95.35 |
| 460 | 2jsa-1 | ASN_A_78_C | ASN_A_78_O | 9.189 | 4.72 | 6.904 | 10.158 | 4.038 | 6.573 | 1.23 | 105.61 | -35.14 |
| 461 | 2jsa-1 | ASN_A_78_C | GLU_A_79_N | 9.189 | 4.72 | 6.904 | 8.019 | 4.713 | 6.283 | 1.32 | 117.96 | -179.66 |
| 462 | 2jsa-1 | GLU_A_79_N | GLU_A_79_H | 8.019 | 4.713 | 6.283 | 7.266 | 5.301 | 6.611 | 1.01 | 71.05 | 142.01 |
| 463 | 2jsa-1 | GLU_A_79_N | GLU_A_79_CA | 8.019 | 4.713 | 6.283 | 7.792 | 3.868 | 5.123 | 1.45 | 142.97 | -105.04 |
| 464 | 2jsa-1 | GLU_A_79_CA | GLU_A_79_HA | 7.792 | 3.868 | 5.123 | 8.513 | 3.051 | 5.155 | 1.09 | 88.32 | -48.57 |
| 465 | 2jsa-1 | GLU_A_79_CA | GLU_A_79_C | 7.792 | 3.868 | 5.123 | 8.022 | 4.661 | 3.835 | 1.53 | 147.34 | 73.83 |
| 466 | 2jsa-1 | GLU_A_79_C | GLU_A_79_O | 8.022 | 4.661 | 3.835 | 8.612 | 4.151 | 2.885 | 1.23 | 140.62 | -40.84 |
| 467 | 2jsa-1 | GLU_A_79_C | CYS_A_80_N | 8.022 | 4.661 | 3.835 | 7.544 | 5.897 | 3.846 | 1.33 | 89.52 | 111.14 |
| 468 | 2jsa-1 | CYS_A_80_N | CYS_A_80_H | 7.544 | 5.897 | 3.846 | 7.07 | 6.252 | 4.664 | 1.01 | 35.90 | 143.17 |
| 469 | 2jsa-1 | CYS_A_80_N | CYS_A_80_CA | 7.544 | 5.897 | 3.846 | 7.69 | 6.766 | 2.69 | 1.45 | 142.68 | 80.46 |
| 470 | 2jsa-1 | CYS_A_80_CA | CYS_A_80_HA | 7.69 | 6.766 | 2.69 | 7.09 | 6.359 | 1.876 | 1.09 | 138.31 | -145.85 |
| 471 | 2jsa-1 | CYS_A_80_CA | CYS_A_80_C | 7.69 | 6.766 | 2.69 | 9.16 | 6.755 | 2.264 | 1.53 | 106.16 | -0.43 |
| 472 | 2jsa-1 | CYS_A_80_C | CYS_A_80_O | 9.16 | 6.755 | 2.264 | 9.467 | 6.572 | 1.087 | 1.23 | 163.11 | -30.80 |
| 473 | 2jsa-1 | CYS_A_80_C | PRO_A_81_N | 9.16 | 6.755 | 2.264 | 10.051 | 6.958 | 3.27 | 1.36 | 42.25 | 12.83 |
| 474 | 2jsa-1 | PRO_A_81_N | PRO_A_81_CA | 10.051 | 6.958 | 3.27 | 11.481 | 6.974 | 3.011 | 1.45 | 100.27 | 0.64 |
| 475 | 2jsa-1 | PRO_A_81_CA | PRO_A_81_HA | 11.481 | 6.974 | 3.011 | 11.693 | 7.571 | 2.124 | 1.09 | 144.46 | 70.45 |
| 476 | 2jsa-1 | PRO_A_81_CA | PRO_A_81_C | 11.481 | 6.974 | 3.011 | 12.012 | 5.557 | 2.781 | 1.53 | 98.64 | -69.46 |
| 477 | 2jsa-1 | PRO_A_81_C | PRO_A_81_O | 12.012 | 5.557 | 2.781 | 13.177 | 5.275 | 3.054 | 1.23 | 77.17 | -13.61 |
| 1 | 2jsa-2 | ARG_A_1_N | ARG_A_1_H | 14.924 | 2.313 | -0.565 | 14.842 | 2.65 | -1.513 | 1.01 | 159.90 | 103.68 |
| 2 | 2jsa-2 | ARG_A_1_N | ARG_A_1_CA | 14.924 | 2.313 | -0.565 | 14.187 | 3.0 | 0.482 | 1.45 | 43.90 | 137.01 |

| | | | | | | | | | | | | |
|---|---|---|---|---|---|---|---|---|---|---|---|---|
| 3 | 2jsa-2 | ARG_A_1_CA | ARG_A_1_HA | 14.187 | 3.0 | 0.482 | 13.499 | 2.29 | 0.941 | 1.09 | 65.10 | -134.10 |
| 4 | 2jsa-2 | ARG_A_1_CA | ARG_A_1_C | 14.187 | 3.0 | 0.482 | 13.384 | 4.161 | -0.108 | 1.53 | 112.68 | 124.67 |
| 5 | 2jsa-2 | ARG_A_1_C | ARG_A_1_O | 13.384 | 4.161 | -0.108 | 12.162 | 4.203 | 0.019 | 1.23 | 84.07 | 178.03 |
| 6 | 2jsa-2 | ARG_A_1_C | SER_A_2_N | 13.384 | 4.161 | -0.108 | 14.105 | 5.075 | -0.74 | 1.32 | 118.50 | 51.73 |
| 7 | 2jsa-2 | SER_A_2_N | SER_A_2_H | 14.105 | 5.075 | -0.74 | 15.108 | 4.976 | -0.806 | 1.01 | 93.75 | -5.64 |
| 8 | 2jsa-2 | SER_A_2_N | SER_A_2_CA | 14.105 | 5.075 | -0.74 | 13.475 | 6.234 | -1.35 | 1.45 | 114.82 | 118.53 |
| 9 | 2jsa-2 | SER_A_2_CA | SER_A_2_HA | 13.475 | 6.234 | -1.35 | 12.52 | 6.404 | -0.854 | 1.09 | 62.92 | 169.91 |
| 10 | 2jsa-2 | SER_A_2_CA | SER_A_2_C | 13.475 | 6.234 | -1.35 | 13.216 | 5.968 | -2.835 | 1.53 | 165.96 | -134.24 |
| 11 | 2jsa-2 | SER_A_2_C | SER_A_2_O | 13.216 | 5.968 | -2.835 | 14.133 | 6.044 | -3.651 | 1.23 | 131.57 | 4.74 |
| 12 | 2jsa-2 | SER_A_2_C | ALA_A_3_N | 13.216 | 5.968 | -2.835 | 11.964 | 5.663 | -3.139 | 1.32 | 103.27 | -166.31 |
| 13 | 2jsa-2 | ALA_A_3_N | ALA_A_3_H | 11.964 | 5.663 | -3.139 | 11.262 | 5.618 | -2.414 | 1.01 | 44.14 | -176.33 |
| 14 | 2jsa-2 | ALA_A_3_N | ALA_A_3_CA | 11.964 | 5.663 | -3.139 | 11.572 | 5.386 | -4.51 | 1.45 | 160.70 | -144.75 |
| 15 | 2jsa-2 | ALA_A_3_CA | ALA_A_3_HA | 11.572 | 5.386 | -4.51 | 11.971 | 6.17 | -5.153 | 1.09 | 126.16 | 63.03 |
| 16 | 2jsa-2 | ALA_A_3_CA | ALA_A_3_C | 11.572 | 5.386 | -4.51 | 10.047 | 5.404 | -4.616 | 1.53 | 93.98 | 179.32 |
| 17 | 2jsa-2 | ALA_A_3_C | ALA_A_3_O | 10.047 | 5.404 | -4.616 | 9.383 | 6.172 | -3.921 | 1.23 | 55.61 | 130.85 |
| 18 | 2jsa-2 | ALA_A_3_C | LEU_A_4_N | 10.047 | 5.404 | -4.616 | 9.535 | 4.549 | -5.489 | 1.32 | 131.22 | -120.91 |
| 19 | 2jsa-2 | LEU_A_4_N | LEU_A_4_H | 9.535 | 4.549 | -5.489 | 10.142 | 3.946 | -6.026 | 1.01 | 122.11 | -44.81 |
| 20 | 2jsa-2 | LEU_A_4_N | LEU_A_4_CA | 9.535 | 4.549 | -5.489 | 8.099 | 4.458 | -5.693 | 1.45 | 98.07 | -176.37 |
| 21 | 2jsa-2 | LEU_A_4_CA | LEU_A_4_HA | 8.099 | 4.458 | -5.693 | 7.695 | 5.47 | -5.7 | 1.09 | 90.37 | 111.76 |
| 22 | 2jsa-2 | LEU_A_4_CA | LEU_A_4_C | 8.099 | 4.458 | -5.693 | 7.467 | 3.713 | -4.516 | 1.53 | 39.69 | -130.31 |
| 23 | 2jsa-2 | LEU_A_4_C | LEU_A_4_O | 7.467 | 3.713 | -4.516 | 6.247 | 3.567 | -4.451 | 1.23 | 86.97 | -173.18 |
| 24 | 2jsa-2 | LEU_A_4_C | SER_A_5_N | 7.467 | 3.713 | -4.516 | 8.326 | 3.26 | -3.614 | 1.33 | 47.11 | -27.81 |
| 25 | 2jsa-2 | SER_A_5_N | SER_A_5_H | 8.326 | 3.26 | -3.614 | 9.317 | 3.416 | -3.731 | 1.01 | 96.65 | 8.95 |
| 26 | 2jsa-2 | SER_A_5_N | SER_A_5_CA | 8.326 | 3.26 | -3.614 | 7.866 | 2.534 | -2.442 | 1.45 | 36.25 | -122.36 |
| 27 | 2jsa-2 | SER_A_5_CA | SER_A_5_HA | 7.866 | 2.534 | -2.442 | 7.398 | 1.609 | -2.776 | 1.09 | 107.86 | -116.84 |
| 28 | 2jsa-2 | SER_A_5_CA | SER_A_5_C | 7.866 | 2.534 | -2.442 | 6.827 | 3.364 | -1.685 | 1.53 | 60.35 | 141.38 |
| 29 | 2jsa-2 | SER_A_5_C | SER_A_5_O | 6.827 | 3.364 | -1.685 | 5.828 | 2.83 | -1.21 | 1.23 | 67.25 | -151.87 |
| 30 | 2jsa-2 | SER_A_5_C | CYS_A_6_N | 6.827 | 3.364 | -1.685 | 7.101 | 4.657 | -1.597 | 1.32 | 86.19 | 78.04 |
| 31 | 2jsa-2 | CYS_A_6_N | CYS_A_6_H | 7.101 | 4.657 | -1.597 | 7.944 | 5.028 | -2.012 | 1.01 | 114.26 | 23.75 |
| 32 | 2jsa-2 | CYS_A_6_N | CYS_A_6_CA | 7.101 | 4.657 | -1.597 | 6.202 | 5.567 | -0.907 | 1.45 | 61.66 | 134.65 |
| 33 | 2jsa-2 | CYS_A_6_CA | CYS_A_6_HA | 6.202 | 5.567 | -0.907 | 6.097 | 5.229 | 0.124 | 1.09 | 18.95 | -107.26 |
| 34 | 2jsa-2 | CYS_A_6_CA | CYS_A_6_C | 6.202 | 5.567 | -0.907 | 4.833 | 5.495 | -1.585 | 1.53 | 116.32 | -176.99 |
| 35 | 2jsa-2 | CYS_A_6_C | CYS_A_6_O | 4.833 | 5.495 | -1.585 | 3.816 | 5.307 | -0.919 | 1.23 | 57.22 | -169.53 |
| 36 | 2jsa-2 | CYS_A_6_C | GLN_A_7_N | 4.833 | 5.495 | -1.585 | 4.85 | 5.65 | -2.9 | 1.32 | 173.24 | 83.74 |
| 37 | 2jsa-2 | GLN_A_7_N | GLN_A_7_H | 4.85 | 5.65 | -2.9 | 5.724 | 5.803 | -3.382 | 1.01 | 118.51 | 9.93 |
| 38 | 2jsa-2 | GLN_A_7_N | GLN_A_7_CA | 4.85 | 5.65 | -2.9 | 3.622 | 5.606 | -3.675 | 1.45 | 122.24 | -177.95 |
| 39 | 2jsa-2 | GLN_A_7_CA | GLN_A_7_HA | 3.622 | 5.606 | -3.675 | 3.058 | 6.514 | -3.464 | 1.09 | 78.83 | 121.85 |
| 40 | 2jsa-2 | GLN_A_7_CA | GLN_A_7_C | 3.622 | 5.606 | -3.675 | 2.773 | 4.404 | -3.256 | 1.53 | 74.11 | -125.23 |
| 41 | 2jsa-2 | GLN_A_7_C | GLN_A_7_O | 2.773 | 4.404 | -3.256 | 1.585 | 4.548 | -2.972 | 1.23 | 76.65 | 173.09 |
| 42 | 2jsa-2 | GLN_A_7_C | MET_A_8_N | 2.773 | 4.404 | -3.256 | 3.415 | 3.245 | -3.231 | 1.33 | 88.92 | -61.02 |
| 43 | 2jsa-2 | MET_A_8_N | MET_A_8_H | 3.415 | 3.245 | -3.231 | 4.393 | 3.2 | -3.48 | 1.01 | 104.27 | -2.63 |
| 44 | 2jsa-2 | MET_A_8_N | MET_A_8_CA | 3.415 | 3.245 | -3.231 | 2.734 | 2.02 | -2.85 | 1.45 | 74.79 | -119.07 |
| 45 | 2jsa-2 | MET_A_8_CA | MET_A_8_HA | 2.734 | 2.02 | -2.85 | 2.061 | 1.73 | -3.656 | 1.09 | 137.72 | -156.69 |
| 46 | 2jsa-2 | MET_A_8_CA | MET_A_8_C | 2.734 | 2.02 | -2.85 | 1.919 | 2.22 | -1.57 | 1.53 | 33.25 | 166.21 |
| 47 | 2jsa-2 | MET_A_8_C | MET_A_8_O | 1.919 | 2.22 | -1.57 | 0.776 | 1.774 | -1.483 | 1.23 | 85.94 | -158.68 |
| 48 | 2jsa-2 | MET_A_8_C | CYS_A_9_N | 1.919 | 2.22 | -1.57 | 2.539 | 2.888 | -0.61 | 1.32 | 43.51 | 47.13 |
| 49 | 2jsa-2 | CYS_A_9_N | CYS_A_9_H | 2.539 | 2.888 | -0.61 | 3.481 | 3.225 | -0.752 | 1.01 | 98.08 | 19.68 |
| 50 | 2jsa-2 | CYS_A_9_N | CYS_A_9_CA | 2.539 | 2.888 | -0.61 | 1.887 | 3.152 | 0.66 | 1.45 | 28.98 | 157.96 |
| 51 | 2jsa-2 | CYS_A_9_CA | CYS_A_9_HA | 1.887 | 3.152 | 0.66 | 1.621 | 2.197 | 1.114 | 1.09 | 65.39 | -105.56 |
| 52 | 2jsa-2 | CYS_A_9_CA | CYS_A_9_C | 1.887 | 3.152 | 0.66 | 0.601 | 3.937 | 0.387 | 1.53 | 100.27 | 148.60 |
| 53 | 2jsa-2 | CYS_A_9_C | CYS_A_9_O | 0.601 | 3.937 | 0.387 | -0.371 | 3.822 | 1.131 | 1.23 | 52.76 | -173.25 |
| 54 | 2jsa-2 | CYS_A_9_C | GLU_A_10_N | 0.601 | 3.937 | 0.387 | 0.639 | 4.717 | -0.685 | 1.33 | 143.93 | 87.21 |
| 55 | 2jsa-2 | GLU_A_10_N | GLU_A_10_H | 0.639 | 4.717 | -0.685 | 1.474 | 4.761 | -1.253 | 1.01 | 124.19 | 3.02 |
| 56 | 2jsa-2 | GLU_A_10_N | GLU_A_10_CA | 0.639 | 4.717 | -0.685 | -0.509 | 5.52 | -1.066 | 1.45 | 105.21 | 145.03 |
| 57 | 2jsa-2 | GLU_A_10_CA | GLU_A_10_HA | -0.509 | 5.52 | -1.066 | -1.0 | 5.859 | -0.154 | 1.09 | 33.19 | 145.38 |
| 58 | 2jsa-2 | GLU_A_10_CA | GLU_A_10_C | -0.509 | 5.52 | -1.066 | -1.506 | 4.676 | -1.865 | 1.53 | 121.45 | -139.75 |
| 59 | 2jsa-2 | GLU_A_10_C | GLU_A_10_O | -1.506 | 4.676 | -1.865 | -2.607 | 5.131 | -2.168 | 1.23 | 104.27 | 157.55 |
| 60 | 2jsa-2 | GLU_A_10_C | LEU_A_11_N | -1.506 | 4.676 | -1.865 | -1.082 | 3.461 | -2.181 | 1.33 | 103.80 | -70.76 |
| 61 | 2jsa-2 | LEU_A_11_N | LEU_A_11_H | -1.082 | 3.461 | -2.181 | -0.163 | 3.152 | -1.897 | 1.01 | 73.67 | -18.58 |
| 62 | 2jsa-2 | LEU_A_11_N | LEU_A_11_CA | -1.082 | 3.461 | -2.181 | -1.924 | 2.549 | -2.939 | 1.45 | 121.41 | -132.71 |
| 63 | 2jsa-2 | LEU_A_11_CA | LEU_A_11_HA | -1.924 | 2.549 | -2.939 | -2.601 | 3.144 | -3.552 | 1.09 | 124.22 | 138.69 |
| 64 | 2jsa-2 | LEU_A_11_CA | LEU_A_11_C | -1.924 | 2.549 | -2.939 | -2.775 | 1.726 | -1.971 | 1.53 | 50.73 | -135.96 |
| 65 | 2jsa-2 | LEU_A_11_C | LEU_A_11_O | -2.775 | 1.726 | -1.971 | -3.978 | 1.574 | -2.173 | 1.23 | 99.46 | -172.80 |
| 66 | 2jsa-2 | LEU_A_11_C | VAL_A_12_N | -2.775 | 1.726 | -1.971 | -2.117 | 1.216 | -0.94 | 1.33 | 38.92 | -37.78 |
| 67 | 2jsa-2 | VAL_A_12_N | VAL_A_12_H | -2.117 | 1.216 | -0.94 | -1.126 | 1.382 | -0.836 | 1.01 | 84.09 | 9.51 |
| 68 | 2jsa-2 | VAL_A_12_N | VAL_A_12_CA | -2.117 | 1.216 | -0.94 | -2.8 | 0.411 | 0.059 | 1.45 | 46.58 | -130.31 |
| 69 | 2jsa-2 | VAL_A_12_CA | VAL_A_12_HA | -2.8 | 0.411 | 0.059 | -3.336 | -0.39 | -0.45 | 1.09 | 117.84 | -123.79 |
| 70 | 2jsa-2 | VAL_A_12_CA | VAL_A_12_C | -2.8 | 0.411 | 0.059 | -3.825 | 1.278 | 0.793 | 1.53 | 61.33 | 139.77 |
| 71 | 2jsa-2 | VAL_A_12_C | VAL_A_12_O | -3.825 | 1.278 | 0.793 | -4.999 | 0.921 | 0.874 | 1.23 | 86.22 | -163.09 |
| 72 | 2jsa-2 | VAL_A_12_C | VAL_A_13_N | -3.825 | 1.278 | 0.793 | -3.344 | 2.401 | 1.307 | 1.33 | 67.18 | 66.81 |
| 73 | 2jsa-2 | VAL_A_13_N | VAL_A_13_H | -3.344 | 2.401 | 1.307 | -2.366 | 2.631 | 1.201 | 1.01 | 96.02 | 13.23 |
| 74 | 2jsa-2 | VAL_A_13_N | VAL_A_13_CA | -3.344 | 2.401 | 1.307 | -4.205 | 3.321 | 2.03 | 1.45 | 60.15 | 133.10 |
| 75 | 2jsa-2 | VAL_A_13_CA | VAL_A_13_HA | -4.205 | 3.321 | 2.03 | -4.641 | 2.791 | 2.877 | 1.09 | 39.02 | -129.44 |
| 76 | 2jsa-2 | VAL_A_13_CA | VAL_A_13_C | -4.205 | 3.321 | 2.03 | -5.345 | 3.773 | 1.114 | 1.53 | 126.76 | 158.37 |
| 77 | 2jsa-2 | VAL_A_13_C | VAL_A_13_O | -5.345 | 3.773 | 1.114 | -6.515 | 3.574 | 1.431 | 1.23 | 75.05 | -170.35 |
| 78 | 2jsa-2 | VAL_A_13_C | LYS_A_14_N | -5.345 | 3.773 | 1.114 | -4.96 | 4.374 | -0.002 | 1.32 | 147.40 | 57.36 |
| 79 | 2jsa-2 | LYS_A_14_N | LYS_A_14_H | -4.96 | 4.374 | -0.002 | -3.977 | 4.505 | -0.196 | 1.01 | 101.07 | 7.59 |
| 80 | 2jsa-2 | LYS_A_14_N | LYS_A_14_CA | -4.96 | 4.374 | -0.002 | -5.934 | 4.856 | -0.966 | 1.45 | 131.57 | 153.67 |
| 81 | 2jsa-2 | LYS_A_14_CA | LYS_A_14_HA | -5.934 | 4.856 | -0.966 | -6.43 | 5.727 | -0.537 | 1.09 | 66.83 | 119.66 |
| 82 | 2jsa-2 | LYS_A_14_CA | LYS_A_14_C | -5.934 | 4.856 | -0.966 | -6.998 | 3.779 | -1.192 | 1.53 | 98.49 | -134.65 |
| 83 | 2jsa-2 | LYS_A_14_C | LYS_A_14_O | -6.998 | 3.779 | -1.192 | -8.153 | 4.092 | -1.475 | 1.23 | 103.31 | 164.84 |
| 84 | 2jsa-2 | LYS_A_14_C | LYS_A_15_N | -6.998 | 3.779 | -1.192 | -6.569 | 2.533 | -1.058 | 1.32 | 84.19 | -71.00 |
| 85 | 2jsa-2 | LYS_A_15_N | LYS_A_15_H | -6.569 | 2.533 | -1.058 | -5.604 | 2.35 | -0.823 | 1.01 | 76.54 | -10.74 |
| 86 | 2jsa-2 | LYS_A_15_N | LYS_A_15_CA | -6.569 | 2.533 | -1.058 | -7.47 | 1.408 | -1.244 | 1.45 | 97.35 | -128.69 |
| 87 | 2jsa-2 | LYS_A_15_CA | LYS_A_15_HA | -7.47 | 1.408 | -1.244 | -8.167 | 1.663 | -2.042 | 1.09 | 137.08 | 159.90 |
| 88 | 2jsa-2 | LYS_A_15_CA | LYS_A_15_C | -7.47 | 1.408 | -1.244 | -8.283 | 1.194 | 0.033 | 1.53 | 33.36 | -165.25 |
| 89 | 2jsa-2 | LYS_A_15_C | LYS_A_15_O | -8.283 | 1.194 | 0.033 | -9.467 | 0.863 | -0.027 | 1.23 | 92.79 | -164.38 |
| 90 | 2jsa-2 | LYS_A_15_C | TYR_A_16_N | -8.283 | 1.194 | 0.033 | -7.617 | 1.393 | 1.162 | 1.33 | 31.62 | 16.64 |
| 91 | 2jsa-2 | TYR_A_16_N | TYR_A_16_H | -7.617 | 1.393 | 1.162 | -6.644 | 1.663 | 1.137 | 1.01 | 91.42 | 15.51 |
| 92 | 2jsa-2 | TYR_A_16_N | TYR_A_16_CA | -7.617 | 1.393 | 1.162 | -8.263 | 1.227 | 2.451 | 1.45 | 27.36 | -165.59 |
| 93 | 2jsa-2 | TYR_A_16_CA | TYR_A_16_HA | -8.263 | 1.227 | 2.451 | -8.849 | 0.308 | 2.464 | 1.09 | 89.32 | -122.52 |
| 94 | 2jsa-2 | TYR_A_16_CA | TYR_A_16_C | -8.263 | 1.227 | 2.451 | -9.163 | 2.421 | 2.773 | 1.53 | 77.85 | 127.01 |
| 95 | 2jsa-2 | TYR_A_16_C | TYR_A_16_O | -9.163 | 2.421 | 2.773 | -10.385 | 2.286 | 2.826 | 1.23 | 87.53 | -173.70 |
| 96 | 2jsa-2 | TYR_A_16_C | GLU_A_17_N | -9.163 | 2.421 | 2.773 | -8.524 | 3.564 | 2.979 | 1.33 | 81.06 | 60.79 |
| 97 | 2jsa-2 | GLU_A_17_N | GLU_A_17_H | -8.524 | 3.564 | 2.979 | -7.516 | 3.602 | 2.919 | 1.01 | 93.40 | 2.16 |
| 98 | 2jsa-2 | GLU_A_17_N | GLU_A_17_CA | -8.524 | 3.564 | 2.979 | -9.253 | 4.781 | 3.293 | 1.45 | 77.52 | 120.92 |

| | | | | | | | | | | | | |
|---|---|---|---|---|---|---|---|---|---|---|---|---|
| 99 | 2jsa-2 | GLU_A_17_CA | GLU_A_17_HA | -9.253 | 4.781 | 3.293 | -9.649 | 4.686 | 4.304 | 1.09 | 21.94 | -166.51 |
| 100 | 2jsa-2 | GLU_A_17_CA | GLU_A_17_C | -9.253 | 4.781 | 3.293 | -10.429 | 4.96 | 2.33 | 1.53 | 128.99 | 171.35 |
| 101 | 2jsa-2 | GLU_A_17_C | GLU_A_17_O | -10.429 | 4.96 | 2.33 | -11.492 | 5.437 | 2.726 | 1.23 | 71.23 | 155.83 |
| 102 | 2jsa-2 | GLU_A_17_C | GLY_A_18_N | -10.429 | 4.96 | 2.33 | -10.199 | 4.567 | 1.086 | 1.32 | 159.90 | -59.66 |
| 103 | 2jsa-2 | GLY_A_18_N | GLY_A_18_H | -10.199 | 4.567 | 1.086 | -9.3 | 4.183 | 0.832 | 1.01 | 104.56 | -23.13 |
| 104 | 2jsa-2 | GLY_A_18_N | GLY_A_18_CA | -10.199 | 4.567 | 1.086 | -11.226 | 4.678 | 0.065 | 1.45 | 134.67 | 173.83 |
| 105 | 2jsa-2 | GLY_A_18_CA | GLY_A_18_C | -11.226 | 4.678 | 0.065 | -11.643 | 3.298 | -0.446 | 1.53 | 109.52 | -106.81 |
| 106 | 2jsa-2 | GLY_A_18_C | GLY_A_18_O | -11.643 | 3.298 | -0.446 | -11.715 | 3.074 | -1.653 | 1.23 | 168.97 | -107.82 |
| 107 | 2jsa-2 | GLY_A_18_C | SER_A_19_N | -11.643 | 3.298 | -0.446 | -11.906 | 2.407 | 0.499 | 1.33 | 44.51 | -106.45 |
| 108 | 2jsa-2 | SER_A_19_N | SER_A_19_H | -11.906 | 2.407 | 0.499 | -11.826 | 2.661 | 1.474 | 1.01 | 15.28 | 72.52 |
| 109 | 2jsa-2 | SER_A_19_N | SER_A_19_CA | -11.906 | 2.407 | 0.499 | -12.314 | 1.054 | 0.16 | 1.45 | 103.49 | -106.78 |
| 110 | 2jsa-2 | SER_A_19_CA | SER_A_19_HA | -12.314 | 1.054 | 0.16 | -12.005 | 0.851 | -0.865 | 1.09 | 160.17 | -33.30 |
| 111 | 2jsa-2 | SER_A_19_CA | SER_A_19_C | -12.314 | 1.054 | 0.16 | -13.836 | 0.929 | 0.247 | 1.53 | 86.74 | -175.30 |
| 112 | 2jsa-2 | SER_A_19_C | SER_A_19_O | -13.836 | 0.929 | 0.247 | -14.481 | 0.482 | -0.701 | 1.23 | 140.38 | -145.28 |
| 113 | 2jsa-2 | SER_A_19_C | ALA_A_20_N | -13.836 | 0.929 | 0.247 | -14.367 | 1.331 | 1.392 | 1.32 | 30.19 | 142.87 |
| 114 | 2jsa-2 | ALA_A_20_N | ALA_A_20_H | -14.367 | 1.331 | 1.392 | -13.775 | 1.687 | 2.129 | 1.01 | 43.15 | 31.02 |
| 115 | 2jsa-2 | ALA_A_20_N | ALA_A_20_CA | -14.367 | 1.331 | 1.392 | -15.801 | 1.269 | 1.615 | 1.45 | 81.17 | -177.52 |
| 116 | 2jsa-2 | ALA_A_20_CA | ALA_A_20_HA | -15.801 | 1.269 | 1.615 | -16.06 | 1.956 | 2.421 | 1.09 | 42.33 | 110.66 |
| 117 | 2jsa-2 | ALA_A_20_CA | ALA_A_20_C | -15.801 | 1.269 | 1.615 | -16.186 | -0.148 | 2.044 | 1.53 | 73.71 | -105.20 |
| 118 | 2jsa-2 | ALA_A_20_C | ALA_A_20_O | -16.186 | -0.148 | 2.044 | -17.342 | -0.409 | 2.375 | 1.23 | 74.39 | -167.28 |
| 119 | 2jsa-2 | ALA_A_20_C | ASP_A_21_N | -16.186 | -0.148 | 2.044 | -15.194 | -1.027 | 2.025 | 1.33 | 90.82 | -41.54 |
| 120 | 2jsa-2 | ASP_A_21_N | ASP_A_21_H | -15.194 | -1.027 | 2.025 | -14.267 | -0.742 | 1.742 | 1.01 | 106.27 | 17.09 |
| 121 | 2jsa-2 | ASP_A_21_N | ASP_A_21_CA | -15.194 | -1.027 | 2.025 | -15.414 | -2.412 | 2.408 | 1.45 | 74.72 | -99.03 |
| 122 | 2jsa-2 | ASP_A_21_CA | ASP_A_21_HA | -15.414 | -2.412 | 2.408 | -16.486 | -2.605 | 2.446 | 1.09 | 88.00 | -169.79 |
| 123 | 2jsa-2 | ASP_A_21_CA | ASP_A_21_C | -15.414 | -2.412 | 2.408 | -14.792 | -2.661 | 3.783 | 1.53 | 25.98 | -21.82 |
| 124 | 2jsa-2 | ASP_A_21_C | ASP_A_21_O | -14.792 | -2.661 | 3.783 | -15.082 | -3.67 | 4.426 | 1.23 | 58.51 | -106.04 |
| 125 | 2jsa-2 | ASP_A_21_C | LYS_A_22_N | -14.792 | -2.661 | 3.783 | -13.949 | -1.725 | 4.195 | 1.33 | 71.89 | 47.99 |
| 126 | 2jsa-2 | LYS_A_22_N | LYS_A_22_H | -13.949 | -1.725 | 4.195 | -13.758 | -0.921 | 3.614 | 1.01 | 125.11 | 76.64 |
| 127 | 2jsa-2 | LYS_A_22_N | LYS_A_22_CA | -13.949 | -1.725 | 4.195 | -13.283 | -1.83 | 5.482 | 1.45 | 27.65 | -8.96 |
| 128 | 2jsa-2 | LYS_A_22_CA | LYS_A_22_HA | -13.283 | -1.83 | 5.482 | -12.45 | -1.126 | 5.491 | 1.09 | 89.53 | 40.20 |
| 129 | 2jsa-2 | LYS_A_22_CA | LYS_A_22_C | -13.283 | -1.83 | 5.482 | -12.705 | -3.238 | 5.638 | 1.53 | 84.15 | -67.68 |
| 130 | 2jsa-2 | LYS_A_22_C | LYS_A_22_O | -12.705 | -3.238 | 5.638 | -12.825 | -3.849 | 6.699 | 1.23 | 30.41 | -101.11 |
| 131 | 2jsa-2 | LYS_A_22_C | ASP_A_23_N | -12.705 | -3.238 | 5.638 | -12.088 | -3.713 | 4.565 | 1.33 | 144.03 | -37.59 |
| 132 | 2jsa-2 | ASP_A_23_N | ASP_A_23_H | -12.088 | -3.713 | 4.565 | -12.026 | -3.155 | 3.725 | 1.01 | 146.24 | 83.66 |
| 133 | 2jsa-2 | ASP_A_23_N | ASP_A_23_CA | -12.088 | -3.713 | 4.565 | -11.49 | -5.037 | 4.569 | 1.45 | 89.84 | -65.69 |
| 134 | 2jsa-2 | ASP_A_23_CA | ASP_A_23_HA | -11.49 | -5.037 | 4.569 | -11.565 | -5.459 | 5.571 | 1.09 | 23.16 | -100.08 |
| 135 | 2jsa-2 | ASP_A_23_CA | ASP_A_23_C | -11.49 | -5.037 | 4.569 | -10.02 | -4.929 | 4.16 | 1.53 | 105.51 | 4.20 |
| 136 | 2jsa-2 | ASP_A_23_C | ASP_A_23_O | -10.02 | -4.929 | 4.16 | -9.712 | -4.52 | 3.041 | 1.23 | 155.41 | 53.02 |
| 137 | 2jsa-2 | ASP_A_23_C | ALA_A_24_N | -10.02 | -4.929 | 4.16 | -9.151 | -5.303 | 5.086 | 1.32 | 45.61 | -23.29 |
| 138 | 2jsa-2 | ALA_A_24_N | ALA_A_24_H | -9.151 | -5.303 | 5.086 | -9.475 | -5.629 | 5.985 | 1.01 | 27.08 | -134.82 |
| 139 | 2jsa-2 | ALA_A_24_N | ALA_A_24_CA | -9.151 | -5.303 | 5.086 | -7.721 | -5.253 | 4.836 | 1.45 | 99.91 | 2.00 |
| 140 | 2jsa-2 | ALA_A_24_CA | ALA_A_24_HA | -7.721 | -5.253 | 4.836 | -7.478 | -4.295 | 4.377 | 1.09 | 114.91 | 75.77 |
| 141 | 2jsa-2 | ALA_A_24_CA | ALA_A_24_C | -7.721 | -5.253 | 4.836 | -7.339 | -6.369 | 3.86 | 1.53 | 129.61 | -71.10 |
| 142 | 2jsa-2 | ALA_A_24_C | ALA_A_24_O | -7.339 | -6.369 | 3.86 | -6.194 | -6.449 | 3.421 | 1.23 | 110.93 | -4.00 |
| 143 | 2jsa-2 | ALA_A_24_C | ASN_A_25_N | -7.339 | -6.369 | 3.86 | -8.322 | -7.201 | 3.55 | 1.32 | 103.53 | -139.76 |
| 144 | 2jsa-2 | ASN_A_25_N | ASN_A_25_H | -8.322 | -7.201 | 3.55 | -9.241 | -7.074 | 3.949 | 1.01 | 66.73 | 172.13 |
| 145 | 2jsa-2 | ASN_A_25_N | ASN_A_25_CA | -8.322 | -7.201 | 3.55 | -8.104 | -8.308 | 2.635 | 1.45 | 129.04 | -78.86 |
| 146 | 2jsa-2 | ASN_A_25_CA | ASN_A_25_HA | -8.104 | -8.308 | 2.635 | -7.078 | -8.661 | 2.74 | 1.09 | 84.47 | -18.99 |
| 147 | 2jsa-2 | ASN_A_25_CA | ASN_A_25_C | -8.104 | -8.308 | 2.635 | -8.346 | -7.834 | 1.201 | 1.53 | 159.64 | 117.05 |
| 148 | 2jsa-2 | ASN_A_25_C | ASN_A_25_O | -8.346 | -7.834 | 1.201 | -7.507 | -8.038 | 0.326 | 1.23 | 135.38 | -13.67 |
| 149 | 2jsa-2 | ASN_A_25_C | VAL_A_26_N | -8.346 | -7.834 | 1.201 | -9.498 | -7.21 | 1.004 | 1.32 | 98.55 | 151.56 |
| 150 | 2jsa-2 | VAL_A_26_N | VAL_A_26_H | -9.498 | -7.21 | 1.004 | -10.141 | -7.078 | 1.771 | 1.01 | 40.56 | 168.40 |
| 151 | 2jsa-2 | VAL_A_26_N | VAL_A_26_CA | -9.498 | -7.21 | 1.004 | -9.862 | -6.704 | -0.309 | 1.45 | 154.60 | 125.73 |
| 152 | 2jsa-2 | VAL_A_26_CA | VAL_A_26_HA | -9.862 | -6.704 | -0.309 | -9.54 | -7.428 | -1.058 | 1.09 | 133.39 | -66.02 |
| 153 | 2jsa-2 | VAL_A_26_CA | VAL_A_26_C | -9.862 | -6.704 | -0.309 | -9.121 | -5.391 | -0.569 | 1.53 | 99.78 | 60.56 |
| 154 | 2jsa-2 | VAL_A_26_C | VAL_A_26_O | -9.121 | -5.391 | -0.569 | -9.145 | -4.87 | -1.682 | 1.23 | 154.89 | 92.64 |
| 155 | 2jsa-2 | VAL_A_26_C | ILE_A_27_N | -9.121 | -5.391 | -0.569 | -8.48 | -4.893 | 0.479 | 1.33 | 37.76 | 37.84 |
| 156 | 2jsa-2 | ILE_A_27_N | ILE_A_27_H | -8.48 | -4.893 | 0.479 | -8.503 | -5.376 | 1.366 | 1.01 | 28.60 | -92.73 |
| 157 | 2jsa-2 | ILE_A_27_N | ILE_A_27_CA | -8.48 | -4.893 | 0.479 | -7.733 | -3.651 | 0.379 | 1.45 | 93.95 | 58.98 |
| 158 | 2jsa-2 | ILE_A_27_CA | ILE_A_27_HA | -7.733 | -3.651 | 0.379 | -8.237 | -3.012 | -0.346 | 1.09 | 131.70 | 128.26 |
| 159 | 2jsa-2 | ILE_A_27_CA | ILE_A_27_C | -7.733 | -3.651 | 0.379 | -6.332 | -3.946 | -0.158 | 1.53 | 110.56 | -11.89 |
| 160 | 2jsa-2 | ILE_A_27_C | ILE_A_27_O | -6.332 | -3.946 | -0.158 | -5.99 | -3.541 | -1.269 | 1.23 | 154.49 | 49.82 |
| 161 | 2jsa-2 | ILE_A_27_C | LYS_A_28_N | -6.332 | -3.946 | -0.158 | -5.556 | -4.647 | 0.656 | 1.33 | 52.10 | -42.09 |
| 162 | 2jsa-2 | LYS_A_28_N | LYS_A_28_H | -5.556 | -4.647 | 0.656 | -5.902 | -4.946 | 1.557 | 1.01 | 26.91 | -139.17 |
| 163 | 2jsa-2 | LYS_A_28_N | LYS_A_28_CA | -5.556 | -4.647 | 0.656 | -4.199 | -5.0 | 0.276 | 1.45 | 105.16 | -14.58 |
| 164 | 2jsa-2 | LYS_A_28_CA | LYS_A_28_HA | -4.199 | -5.0 | 0.276 | -3.624 | -4.079 | 0.181 | 1.09 | 95.00 | 58.02 |
| 165 | 2jsa-2 | LYS_A_28_CA | LYS_A_28_C | -4.199 | -5.0 | 0.276 | -4.218 | -5.679 | -1.095 | 1.53 | 153.64 | -91.60 |
| 166 | 2jsa-2 | LYS_A_28_C | LYS_A_28_O | -4.218 | -5.679 | -1.095 | -3.295 | -5.504 | -1.89 | 1.23 | 130.24 | 10.74 |
| 167 | 2jsa-2 | LYS_A_28_C | LYS_A_29_N | -4.218 | -5.679 | -1.095 | -5.276 | -6.439 | -1.328 | 1.32 | 100.14 | -144.31 |
| 168 | 2jsa-2 | LYS_A_29_N | LYS_A_29_H | -5.276 | -6.439 | -1.328 | -5.995 | -6.538 | -0.626 | 1.01 | 45.95 | -172.16 |
| 169 | 2jsa-2 | LYS_A_29_N | LYS_A_29_CA | -5.276 | -6.439 | -1.328 | -5.427 | -7.145 | -2.59 | 1.45 | 150.23 | -102.07 |
| 170 | 2jsa-2 | LYS_A_29_CA | LYS_A_29_HA | -5.427 | -7.145 | -2.59 | -4.504 | -7.692 | -2.783 | 1.09 | 100.20 | -30.65 |
| 171 | 2jsa-2 | LYS_A_29_CA | LYS_A_29_C | -5.427 | -7.145 | -2.59 | -5.611 | -6.129 | -3.719 | 1.53 | 137.56 | 100.27 |
| 172 | 2jsa-2 | LYS_A_29_C | LYS_A_29_O | -5.611 | -6.129 | -3.719 | -5.02 | -6.271 | -4.788 | 1.23 | 150.38 | -13.51 |
| 173 | 2jsa-2 | LYS_A_29_C | ASP_A_30_N | -5.611 | -6.129 | -3.719 | -6.432 | -5.127 | -3.443 | 1.32 | 77.97 | 129.33 |
| 174 | 2jsa-2 | ASP_A_30_N | ASP_A_30_H | -6.432 | -5.127 | -3.443 | -6.885 | -5.074 | -2.542 | 1.01 | 26.85 | 173.33 |
| 175 | 2jsa-2 | ASP_A_30_N | ASP_A_30_CA | -6.432 | -5.127 | -3.443 | -6.7 | -4.088 | -4.423 | 1.45 | 132.41 | 104.46 |
| 176 | 2jsa-2 | ASP_A_30_CA | ASP_A_30_HA | -6.7 | -4.088 | -4.423 | -7.214 | -4.528 | -5.277 | 1.09 | 141.61 | -139.44 |
| 177 | 2jsa-2 | ASP_A_30_CA | ASP_A_30_C | -6.7 | -4.088 | -4.423 | -5.378 | -3.466 | -4.875 | 1.53 | 107.19 | 25.20 |
| 178 | 2jsa-2 | ASP_A_30_C | ASP_A_30_O | -5.378 | -3.466 | -4.875 | -5.133 | -3.322 | -6.071 | 1.23 | 166.63 | 30.45 |
| 179 | 2jsa-2 | ASP_A_30_C | PHE_A_31_N | -5.378 | -3.466 | -4.875 | -4.561 | -3.112 | -3.894 | 1.32 | 42.23 | 23.43 |
| 180 | 2jsa-2 | PHE_A_31_N | PHE_A_31_H | -4.561 | -3.112 | -3.894 | -4.829 | -3.259 | -2.931 | 1.01 | 17.61 | -151.25 |
| 181 | 2jsa-2 | PHE_A_31_N | PHE_A_31_CA | -4.561 | -3.112 | -3.894 | -3.269 | -2.509 | -4.175 | 1.45 | 101.15 | 25.02 |
| 182 | 2jsa-2 | PHE_A_31_CA | PHE_A_31_HA | -3.269 | -2.509 | -4.175 | -3.402 | -1.467 | -4.466 | 1.09 | 105.48 | 97.27 |
| 183 | 2jsa-2 | PHE_A_31_CA | PHE_A_31_C | -3.269 | -2.509 | -4.175 | -2.542 | -3.265 | -5.289 | 1.53 | 136.73 | -46.12 |
| 184 | 2jsa-2 | PHE_A_31_C | PHE_A_31_O | -2.542 | -3.265 | -5.289 | -2.128 | -2.669 | -6.282 | 1.23 | 143.84 | 55.21 |
| 185 | 2jsa-2 | PHE_A_31_C | ASP_A_32_N | -2.542 | -3.265 | -5.289 | -2.409 | -4.567 | -5.087 | 1.32 | 81.23 | -84.17 |
| 186 | 2jsa-2 | ASP_A_32_N | ASP_A_32_H | -2.409 | -4.567 | -5.087 | -2.774 | -4.991 | -4.246 | 1.01 | 33.63 | -130.72 |
| 187 | 2jsa-2 | ASP_A_32_N | ASP_A_32_CA | -2.409 | -4.567 | -5.087 | -1.739 | -5.412 | -6.06 | 1.45 | 132.06 | -51.59 |
| 188 | 2jsa-2 | ASP_A_32_CA | ASP_A_32_HA | -1.739 | -5.412 | -6.06 | -0.71 | -5.072 | -6.177 | 1.09 | 96.16 | 18.28 |
| 189 | 2jsa-2 | ASP_A_32_CA | ASP_A_32_C | -1.739 | -5.412 | -6.06 | -2.473 | -5.321 | -7.399 | 1.53 | 151.09 | 172.93 |
| 190 | 2jsa-2 | ASP_A_32_C | ASP_A_32_O | -2.473 | -5.321 | -7.399 | -1.887 | -5.573 | -8.451 | 1.23 | 148.77 | -23.27 |
| 191 | 2jsa-2 | ASP_A_32_C | ALA_A_33_N | -2.473 | -5.321 | -7.399 | -3.746 | -4.959 | -7.316 | 1.33 | 86.41 | 164.13 |
| 192 | 2jsa-2 | ALA_A_33_N | ALA_A_33_H | -3.746 | -4.959 | -7.316 | -4.163 | -4.767 | -6.416 | 1.01 | 27.03 | 155.28 |
| 193 | 2jsa-2 | ALA_A_33_N | ALA_A_33_CA | -3.746 | -4.959 | -7.316 | -4.566 | -4.832 | -8.509 | 1.45 | 145.18 | 171.20 |
| 194 | 2jsa-2 | ALA_A_33_CA | ALA_A_33_HA | -4.566 | -4.832 | -8.509 | -4.401 | -5.703 | -9.143 | 1.09 | 125.57 | -79.27 |

| | | | | | | | | | | | |
|---|---|---|---|---|---|---|---|---|---|---|---|
| 195 | 2jsa-2 | ALA_A_33_CA | ALA_A_33_C | -4.566 | -4.832 | -8.509 | -4.141 | -3.582 | -9.283 | 1.53 | 120.38 | 71.22 |
| 196 | 2jsa-2 | ALA_A_33_C | ALA_A_33_O | -4.141 | -3.582 | -9.283 | -3.575 | -3.684 | -10.371 | 1.23 | 152.14 | -10.22 |
| 197 | 2jsa-2 | ALA_A_33_C | GLU_A_34_N | -4.141 | -3.582 | -9.283 | -4.432 | -2.432 | -8.694 | 1.32 | 63.59 | 104.20 |
| 198 | 2jsa-2 | GLU_A_34_N | GLU_A_34_H | -4.432 | -2.432 | -8.694 | -4.9 | -2.423 | -7.799 | 1.01 | 27.61 | 178.90 |
| 199 | 2jsa-2 | GLU_A_34_N | GLU_A_34_CA | -4.432 | -2.432 | -8.694 | -4.087 | -1.165 | -9.315 | 1.45 | 115.31 | 74.77 |
| 200 | 2jsa-2 | GLU_A_34_CA | GLU_A_34_HA | -4.087 | -1.165 | -9.315 | -4.659 | -1.075 | -10.238 | 1.09 | 147.90 | 171.06 |
| 201 | 2jsa-2 | GLU_A_34_CA | GLU_A_34_C | -4.087 | -1.165 | -9.315 | -2.599 | -1.133 | -9.668 | 1.53 | 103.34 | 1.23 |
| 202 | 2jsa-2 | GLU_A_34_C | GLU_A_34_O | -2.599 | -1.133 | -9.668 | -2.195 | -0.454 | -10.611 | 1.23 | 140.04 | 59.25 |
| 203 | 2jsa-2 | GLU_A_34_C | CYS_A_35_N | -2.599 | -1.133 | -9.668 | -1.824 | -1.878 | -8.893 | 1.33 | 54.21 | -43.87 |
| 204 | 2jsa-2 | CYS_A_35_N | CYS_A_35_H | -1.824 | -1.878 | -8.893 | -2.225 | -2.412 | -8.136 | 1.01 | 41.42 | -126.90 |
| 205 | 2jsa-2 | CYS_A_35_N | CYS_A_35_CA | -1.824 | -1.878 | -8.893 | -0.389 | -1.945 | -9.112 | 1.45 | 98.67 | -2.67 |
| 206 | 2jsa-2 | CYS_A_35_CA | CYS_A_35_HA | -0.389 | -1.945 | -9.112 | -0.021 | -0.934 | -9.29 | 1.09 | 99.39 | 70.00 |
| 207 | 2jsa-2 | CYS_A_35_CA | CYS_A_35_C | -0.389 | -1.945 | -9.112 | -0.133 | -2.784 | -10.365 | 1.53 | 145.01 | -73.03 |
| 208 | 2jsa-2 | CYS_A_35_C | CYS_A_35_O | -0.133 | -2.784 | -10.365 | 0.658 | -2.397 | -11.224 | 1.23 | 134.29 | 26.07 |
| 209 | 2jsa-2 | CYS_A_35_C | LYS_A_36_N | -0.133 | -2.784 | -10.365 | -0.817 | -3.916 | -10.43 | 1.32 | 92.81 | -121.14 |
| 210 | 2jsa-2 | LYS_A_36_N | LYS_A_36_H | -0.817 | -3.916 | -10.43 | -1.45 | -4.171 | -9.686 | 1.01 | 42.53 | -158.06 |
| 211 | 2jsa-2 | LYS_A_36_N | LYS_A_36_CA | -0.817 | -3.916 | -10.43 | -0.674 | -4.813 | -11.565 | 1.45 | 141.33 | -80.94 |
| 212 | 2jsa-2 | LYS_A_36_CA | LYS_A_36_HA | -0.674 | -4.813 | -11.565 | 0.353 | -5.178 | -11.578 | 1.09 | 90.68 | -19.57 |
| 213 | 2jsa-2 | LYS_A_36_CA | LYS_A_36_C | -0.674 | -4.813 | -11.565 | -0.906 | -4.033 | -12.86 | 1.53 | 147.85 | 106.56 |
| 214 | 2jsa-2 | LYS_A_36_C | LYS_A_36_O | -0.906 | -4.033 | -12.86 | -0.49 | -4.465 | -13.933 | 1.23 | 150.80 | -46.08 |
| 215 | 2jsa-2 | LYS_A_36_C | LYS_A_37_N | -0.906 | -4.033 | -12.86 | -1.569 | -2.895 | -12.716 | 1.32 | 83.76 | 120.23 |
| 216 | 2jsa-2 | LYS_A_37_N | LYS_A_37_H | -1.569 | -2.895 | -12.716 | -1.88 | -2.6 | -11.802 | 1.01 | 25.13 | 136.51 |
| 217 | 2jsa-2 | LYS_A_37_N | LYS_A_37_CA | -1.569 | -2.895 | -12.716 | -1.862 | -2.049 | -13.861 | 1.45 | 141.98 | 109.10 |
| 218 | 2jsa-2 | LYS_A_37_CA | LYS_A_37_HA | -1.862 | -2.049 | -13.861 | -1.611 | -2.605 | -14.764 | 1.09 | 145.96 | -65.70 |
| 219 | 2jsa-2 | LYS_A_37_CA | LYS_A_37_C | -1.862 | -2.049 | -13.861 | -0.968 | -0.809 | -13.815 | 1.53 | 88.28 | 54.21 |
| 220 | 2jsa-2 | LYS_A_37_C | LYS_A_37_O | -0.968 | -0.809 | -13.815 | -0.412 | -0.401 | -14.834 | 1.23 | 145.91 | 36.27 |
| 221 | 2jsa-2 | LYS_A_37_C | LEU_A_38_N | -0.968 | -0.809 | -13.815 | -0.858 | -0.241 | -12.622 | 1.33 | 25.87 | 79.04 |
| 222 | 2jsa-2 | LEU_A_38_N | LEU_A_38_H | -0.858 | -0.241 | -12.622 | -1.344 | -0.629 | -11.827 | 1.01 | 38.03 | -141.40 |
| 223 | 2jsa-2 | LEU_A_38_N | LEU_A_38_CA | -0.858 | -0.241 | -12.622 | -0.041 | 0.945 | -12.43 | 1.45 | 82.41 | 55.44 |
| 224 | 2jsa-2 | LEU_A_38_CA | LEU_A_38_HA | -0.041 | 0.945 | -12.43 | -0.349 | 1.689 | -13.164 | 1.09 | 132.35 | 112.49 |
| 225 | 2jsa-2 | LEU_A_38_CA | LEU_A_38_C | -0.041 | 0.945 | -12.43 | 1.425 | 0.6 | -12.704 | 1.53 | 100.31 | -13.24 |
| 226 | 2jsa-2 | LEU_A_38_C | LEU_A_38_O | 1.425 | 0.6 | -12.704 | 2.021 | 1.114 | -13.65 | 1.23 | 140.24 | 40.78 |
| 227 | 2jsa-2 | LEU_A_38_C | PHE_A_39_N | 1.425 | 0.6 | -12.704 | 1.962 | -0.269 | -11.86 | 1.33 | 50.44 | -58.29 |
| 228 | 2jsa-2 | PHE_A_39_N | PHE_A_39_H | 1.962 | -0.269 | -11.86 | 1.412 | -0.652 | -11.105 | 1.01 | 41.60 | -145.15 |
| 229 | 2jsa-2 | PHE_A_39_N | PHE_A_39_CA | 1.962 | -0.269 | -11.86 | 3.346 | -0.689 | -12.0 | 1.45 | 95.53 | -16.88 |
| 230 | 2jsa-2 | PHE_A_39_CA | PHE_A_39_HA | 3.346 | -0.689 | -12.0 | 3.943 | 0.132 | -12.398 | 1.09 | 111.41 | 53.98 |
| 231 | 2jsa-2 | PHE_A_39_CA | PHE_A_39_C | 3.346 | -0.689 | -12.0 | 3.461 | -1.903 | -12.923 | 1.53 | 127.12 | -84.59 |
| 232 | 2jsa-2 | PHE_A_39_C | PHE_A_39_O | 3.461 | -1.903 | -12.923 | 4.327 | -2.755 | -12.731 | 1.23 | 81.02 | -44.53 |
| 233 | 2jsa-2 | PHE_A_39_C | HIS_A_40_N | 3.461 | -1.903 | -12.923 | 2.574 | -1.944 | -13.907 | 1.33 | 137.94 | -177.35 |
| 234 | 2jsa-2 | HIS_A_40_N | HIS_A_40_H | 2.574 | -1.944 | -13.907 | 1.889 | -1.208 | -14.005 | 1.01 | 95.57 | 132.94 |
| 235 | 2jsa-2 | HIS_A_40_N | HIS_A_40_CA | 2.574 | -1.944 | -13.907 | 2.564 | -3.039 | -14.861 | 1.45 | 131.06 | -90.52 |
| 236 | 2jsa-2 | HIS_A_40_CA | HIS_A_40_HA | 2.564 | -3.039 | -14.861 | 2.587 | -3.973 | -14.3 | 1.09 | 59.02 | -88.59 |
| 237 | 2jsa-2 | HIS_A_40_CA | HIS_A_40_C | 2.564 | -3.039 | -14.861 | 3.825 | -2.979 | -15.725 | 1.53 | 124.39 | 2.72 |
| 238 | 2jsa-2 | HIS_A_40_C | HIS_A_40_O | 3.825 | -2.979 | -15.725 | 4.24 | -3.988 | -16.294 | 1.23 | 117.54 | -67.64 |
| 239 | 2jsa-2 | HIS_A_40_C | THR_A_41_N | 3.825 | -2.979 | -15.725 | 4.399 | -1.787 | -15.794 | 1.32 | 92.99 | 64.29 |
| 240 | 2jsa-2 | THR_A_41_N | THR_A_41_H | 4.399 | -1.787 | -15.794 | 4.003 | -1.001 | -15.298 | 1.01 | 60.60 | 116.74 |
| 241 | 2jsa-2 | THR_A_41_N | THR_A_41_CA | 4.399 | -1.787 | -15.794 | 5.606 | -1.582 | -16.579 | 1.45 | 122.67 | 9.64 |
| 242 | 2jsa-2 | THR_A_41_CA | THR_A_41_HA | 5.606 | -1.582 | -16.579 | 5.871 | -2.52 | -17.066 | 1.09 | 116.55 | -74.22 |
| 243 | 2jsa-2 | THR_A_41_CA | THR_A_41_C | 5.606 | -1.582 | -16.579 | 6.769 | -1.178 | -15.672 | 1.53 | 53.62 | 19.16 |
| 244 | 2jsa-2 | THR_A_41_C | THR_A_41_O | 6.769 | -1.178 | -15.672 | 7.328 | -0.092 | -15.819 | 1.23 | 96.86 | 62.76 |
| 245 | 2jsa-2 | THR_A_41_C | ILE_A_42_N | 6.769 | -1.178 | -15.672 | 7.101 | -2.074 | -14.755 | 1.32 | 46.18 | -69.67 |
| 246 | 2jsa-2 | ILE_A_42_N | ILE_A_42_H | 7.101 | -2.074 | -14.755 | 6.599 | -2.948 | -14.692 | 1.01 | 86.42 | -119.87 |
| 247 | 2jsa-2 | ILE_A_42_N | ILE_A_42_CA | 7.101 | -2.074 | -14.755 | 8.189 | -1.824 | -13.824 | 1.45 | 50.17 | 12.94 |
| 248 | 2jsa-2 | ILE_A_42_CA | ILE_A_42_HA | 8.189 | -1.824 | -13.824 | 8.909 | -1.168 | -14.314 | 1.09 | 116.71 | 42.34 |
| 249 | 2jsa-2 | ILE_A_42_CA | ILE_A_42_C | 8.189 | -1.824 | -13.824 | 8.9 | -3.143 | -13.511 | 1.53 | 78.20 | -61.67 |
| 250 | 2jsa-2 | ILE_A_42_C | ILE_A_42_O | 8.9 | -3.143 | -13.511 | 8.261 | -4.191 | -13.42 | 1.23 | 85.76 | -121.37 |
| 251 | 2jsa-2 | ILE_A_42_C | PRO_A_43_N | 8.9 | -3.143 | -13.511 | 10.247 | -3.046 | -13.352 | 1.36 | 83.29 | 4.12 |
| 252 | 2jsa-2 | PRO_A_43_N | PRO_A_43_CA | 10.247 | -3.046 | -13.352 | 11.051 | -4.218 | -13.052 | 1.45 | 78.08 | -55.55 |
| 253 | 2jsa-2 | PRO_A_43_CA | PRO_A_43_HA | 11.051 | -4.218 | -13.052 | 10.756 | -5.048 | -13.694 | 1.09 | 126.09 | -109.57 |
| 254 | 2jsa-2 | PRO_A_43_CA | PRO_A_43_C | 11.051 | -4.218 | -13.052 | 10.874 | -4.646 | -11.593 | 1.53 | 17.61 | -112.47 |
| 255 | 2jsa-2 | PRO_A_43_C | PRO_A_43_O | 10.874 | -4.646 | -11.593 | 11.385 | -3.991 | -10.685 | 1.23 | 42.46 | 52.04 |
| 256 | 2jsa-2 | PRO_A_43_C | PHE_A_44_N | 10.874 | -4.646 | -11.593 | 10.151 | -5.741 | -11.414 | 1.32 | 82.23 | -123.44 |
| 257 | 2jsa-2 | PHE_A_44_N | PHE_A_44_H | 10.151 | -5.741 | -11.414 | 9.763 | -6.229 | -12.208 | 1.01 | 141.86 | -128.49 |
| 258 | 2jsa-2 | PHE_A_44_N | PHE_A_44_CA | 10.151 | -5.741 | -11.414 | 9.901 | -6.263 | -10.082 | 1.45 | 23.49 | -115.59 |
| 259 | 2jsa-2 | PHE_A_44_CA | PHE_A_44_HA | 9.901 | -6.263 | -10.082 | 9.588 | -7.305 | -10.146 | 1.09 | 93.37 | -106.72 |
| 260 | 2jsa-2 | PHE_A_44_CA | PHE_A_44_C | 9.901 | -6.263 | -10.082 | 8.825 | -5.445 | -9.365 | 1.53 | 62.06 | 142.76 |
| 261 | 2jsa-2 | PHE_A_44_C | PHE_A_44_O | 8.825 | -5.445 | -9.365 | 8.445 | -5.762 | -8.24 | 1.23 | 23.74 | -140.16 |
| 262 | 2jsa-2 | PHE_A_44_C | GLY_A_45_N | 8.825 | -5.445 | -9.365 | 8.364 | -4.406 | -10.047 | 1.33 | 120.96 | 113.93 |
| 263 | 2jsa-2 | GLY_A_45_N | GLY_A_45_H | 8.364 | -4.406 | -10.047 | 8.724 | -4.203 | -10.969 | 1.01 | 155.86 | 29.42 |
| 264 | 2jsa-2 | GLY_A_45_N | GLY_A_45_CA | 8.364 | -4.406 | -10.047 | 7.339 | -3.54 | -9.49 | 1.45 | 67.46 | 139.81 |
| 265 | 2jsa-2 | GLY_A_45_CA | GLY_A_45_C | 7.339 | -3.54 | -9.49 | 6.149 | -4.356 | -8.981 | 1.53 | 70.57 | -145.56 |
| 266 | 2jsa-2 | GLY_A_45_C | GLY_A_45_O | 6.149 | -4.356 | -8.981 | 5.645 | -4.109 | -7.886 | 1.23 | 27.14 | 153.89 |
| 267 | 2jsa-2 | GLY_A_45_C | THR_A_46_N | 6.149 | -4.356 | -8.981 | 5.733 | -5.311 | -9.8 | 1.33 | 128.18 | -113.54 |
| 268 | 2jsa-2 | THR_A_46_N | THR_A_46_H | 5.733 | -5.311 | -9.8 | 6.195 | -5.457 | -10.686 | 1.01 | 151.33 | -17.54 |
| 269 | 2jsa-2 | THR_A_46_N | THR_A_46_CA | 5.733 | -5.311 | -9.8 | 4.612 | -6.165 | -9.445 | 1.45 | 75.86 | -142.70 |
| 270 | 2jsa-2 | THR_A_46_CA | THR_A_46_HA | 4.612 | -6.165 | -9.445 | 3.798 | -5.539 | -9.08 | 1.09 | 70.43 | 142.44 |
| 271 | 2jsa-2 | THR_A_46_CA | THR_A_46_C | 4.612 | -6.165 | -9.445 | 5.007 | -7.125 | -8.322 | 1.53 | 42.75 | -67.63 |
| 272 | 2jsa-2 | THR_A_46_C | THR_A_46_O | 5.007 | -7.125 | -8.322 | 4.175 | -7.494 | -7.496 | 1.23 | 47.78 | -156.08 |
| 273 | 2jsa-2 | THR_A_46_C | ARG_A_47_N | 5.007 | -7.125 | -8.322 | 6.277 | -7.502 | -8.329 | 1.32 | 90.30 | -16.53 |
| 274 | 2jsa-2 | ARG_A_47_N | ARG_A_47_H | 6.277 | -7.502 | -8.329 | 6.906 | -7.158 | -9.041 | 1.01 | 134.80 | 28.67 |
| 275 | 2jsa-2 | ARG_A_47_N | ARG_A_47_CA | 6.277 | -7.502 | -8.329 | 6.792 | -8.413 | -7.321 | 1.45 | 46.07 | -60.52 |
| 276 | 2jsa-2 | ARG_A_47_CA | ARG_A_47_HA | 6.792 | -8.413 | -7.321 | 6.164 | -9.304 | -7.31 | 1.09 | 89.42 | -125.18 |
| 277 | 2jsa-2 | ARG_A_47_CA | ARG_A_47_C | 6.792 | -8.413 | -7.321 | 6.748 | -7.756 | -5.939 | 1.53 | 25.48 | 93.83 |
| 278 | 2jsa-2 | ARG_A_47_C | ARG_A_47_O | 6.748 | -7.756 | -5.939 | 6.075 | -8.247 | -5.035 | 1.23 | 42.66 | -143.89 |
| 279 | 2jsa-2 | ARG_A_47_C | GLU_A_48_N | 6.748 | -7.756 | -5.939 | 7.476 | -6.655 | -5.821 | 1.33 | 84.89 | 56.53 |
| 280 | 2jsa-2 | GLU_A_48_N | GLU_A_48_H | 7.476 | -6.655 | -5.821 | 8.006 | -6.308 | -6.608 | 1.01 | 141.17 | 33.21 |
| 281 | 2jsa-2 | GLU_A_48_N | GLU_A_48_CA | 7.476 | -6.655 | -5.821 | 7.529 | -5.925 | -4.565 | 1.45 | 30.23 | 85.85 |
| 282 | 2jsa-2 | GLU_A_48_CA | GLU_A_48_HA | 7.529 | -5.925 | -4.565 | 8.009 | -6.564 | -3.823 | 1.09 | 47.13 | -53.09 |
| 283 | 2jsa-2 | GLU_A_48_CA | GLU_A_48_C | 7.529 | -5.925 | -4.565 | 6.115 | -5.603 | -4.079 | 1.53 | 71.47 | 167.17 |
| 284 | 2jsa-2 | GLU_A_48_C | GLU_A_48_O | 6.115 | -5.603 | -4.079 | 5.879 | -5.491 | -2.876 | 1.23 | 12.25 | 154.61 |
| 285 | 2jsa-2 | GLU_A_48_C | CYS_A_49_N | 6.115 | -5.603 | -4.079 | 5.211 | -5.463 | -5.037 | 1.32 | 136.32 | 171.20 |
| 286 | 2jsa-2 | CYS_A_49_N | CYS_A_49_H | 5.211 | -5.463 | -5.037 | 5.476 | -5.57 | -6.006 | 1.01 | 163.57 | -21.99 |
| 287 | 2jsa-2 | CYS_A_49_N | CYS_A_49_CA | 5.211 | -5.463 | -5.037 | 3.827 | -5.156 | -4.721 | 1.45 | 77.43 | 167.49 |
| 288 | 2jsa-2 | CYS_A_49_CA | CYS_A_49_HA | 3.827 | -5.156 | -4.721 | 3.806 | -4.243 | -4.125 | 1.09 | 56.87 | 91.32 |
| 289 | 2jsa-2 | CYS_A_49_CA | CYS_A_49_C | 3.827 | -5.156 | -4.721 | 3.261 | -6.3 | -3.877 | 1.53 | 56.53 | -116.32 |
| 290 | 2jsa-2 | CYS_A_49_C | CYS_A_49_O | 3.261 | -6.3 | -3.877 | 2.73 | -6.072 | -2.791 | 1.23 | 28.02 | 156.76 |

| | | | | | | | | | | | | |
|---|---|---|---|---|---|---|---|---|---|---|---|---|
| 291 | 2jsa-2 | CYS_A_49_C | ASP_A_50_N | 3.261 | -6.3 | -3.877 | 3.395 | -7.506 | -4.409 | 1.32 | 113.67 | -83.66 |
| 292 | 2jsa-2 | ASP_A_50_N | ASP_A_50_H | 3.395 | -7.506 | -4.409 | 3.843 | -7.619 | -5.307 | 1.01 | 152.77 | -14.16 |
| 293 | 2jsa-2 | ASP_A_50_N | ASP_A_50_CA | 3.395 | -7.506 | -4.409 | 2.904 | -8.687 | -3.719 | 1.45 | 61.65 | -112.58 |
| 294 | 2jsa-2 | ASP_A_50_CA | ASP_A_50_HA | 2.904 | -8.687 | -3.719 | 1.842 | -8.561 | -3.51 | 1.09 | 78.94 | 173.23 |
| 295 | 2jsa-2 | ASP_A_50_CA | ASP_A_50_C | 2.904 | -8.687 | -3.719 | 3.674 | -8.863 | -2.409 | 1.53 | 31.09 | -12.88 |
| 296 | 2jsa-2 | ASP_A_50_C | ASP_A_50_O | 3.674 | -8.863 | -2.409 | 3.283 | -9.661 | -1.558 | 1.23 | 46.24 | -116.10 |
| 297 | 2jsa-2 | ASP_A_50_C | HIS_A_51_N | 3.674 | -8.863 | -2.409 | 4.756 | -8.109 | -2.289 | 1.32 | 84.80 | 34.87 |
| 298 | 2jsa-2 | HIS_A_51_N | HIS_A_51_H | 4.756 | -8.109 | -2.289 | 5.02 | -7.476 | -3.031 | 1.01 | 137.25 | 67.36 |
| 299 | 2jsa-2 | HIS_A_51_N | HIS_A_51_CA | 4.756 | -8.109 | -2.289 | 5.585 | -8.173 | -1.097 | 1.45 | 34.90 | -4.41 |
| 300 | 2jsa-2 | HIS_A_51_CA | HIS_A_51_HA | 5.585 | -8.173 | -1.097 | 5.282 | -9.048 | -0.522 | 1.09 | 58.16 | -109.10 |
| 301 | 2jsa-2 | HIS_A_51_CA | HIS_A_51_C | 5.585 | -8.173 | -1.097 | 5.338 | -6.934 | -0.234 | 1.53 | 55.66 | 101.27 |
| 302 | 2jsa-2 | HIS_A_51_C | HIS_A_51_O | 5.338 | -6.934 | -0.234 | 5.975 | -6.759 | 0.804 | 1.23 | 32.47 | 15.36 |
| 303 | 2jsa-2 | HIS_A_51_C | TYR_A_52_N | 5.338 | -6.934 | -0.234 | 4.411 | -6.106 | -0.694 | 1.33 | 110.31 | 138.23 |
| 304 | 2jsa-2 | TYR_A_52_N | TYR_A_52_H | 4.411 | -6.106 | -0.694 | 3.927 | -6.313 | -1.556 | 1.01 | 148.59 | -156.84 |
| 305 | 2jsa-2 | TYR_A_52_N | TYR_A_52_CA | 4.411 | -6.106 | -0.694 | 4.072 | -4.889 | 0.024 | 1.45 | 60.39 | 105.57 |
| 306 | 2jsa-2 | TYR_A_52_CA | TYR_A_52_HA | 4.072 | -4.889 | 0.024 | 4.83 | -4.672 | 0.778 | 1.09 | 46.28 | 15.98 |
| 307 | 2jsa-2 | TYR_A_52_CA | TYR_A_52_C | 4.072 | -4.889 | 0.024 | 2.698 | -5.008 | 0.687 | 1.53 | 64.32 | -175.05 |
| 308 | 2jsa-2 | TYR_A_52_C | TYR_A_52_O | 2.698 | -5.008 | 0.687 | 2.527 | -4.621 | 1.842 | 1.23 | 20.12 | 113.84 |
| 309 | 2jsa-2 | TYR_A_52_C | VAL_A_53_N | 2.698 | -5.008 | 0.687 | 1.755 | -5.547 | -0.072 | 1.33 | 124.95 | -150.25 |
| 310 | 2jsa-2 | VAL_A_53_N | VAL_A_53_H | 1.755 | -5.547 | -0.072 | 1.968 | -5.844 | -1.013 | 1.01 | 158.77 | -54.35 |
| 311 | 2jsa-2 | VAL_A_53_N | VAL_A_53_CA | 1.755 | -5.547 | -0.072 | 0.401 | -5.722 | 0.429 | 1.45 | 69.85 | -172.64 |
| 312 | 2jsa-2 | VAL_A_53_CA | VAL_A_53_HA | 0.401 | -5.722 | 0.429 | -0.005 | -4.741 | 0.675 | 1.09 | 76.95 | 112.48 |
| 313 | 2jsa-2 | VAL_A_53_CA | VAL_A_53_C | 0.401 | -5.722 | 0.429 | 0.441 | -6.554 | 1.711 | 1.53 | 33.01 | -87.25 |
| 314 | 2jsa-2 | VAL_A_53_C | VAL_A_53_O | 0.441 | -6.554 | 1.711 | -0.516 | -6.552 | 2.486 | 1.23 | 51.00 | 179.88 |
| 315 | 2jsa-2 | VAL_A_53_C | ASN_A_54_N | 0.441 | -6.554 | 1.711 | 1.555 | -7.246 | 1.897 | 1.32 | 81.93 | -31.85 |
| 316 | 2jsa-2 | ASN_A_54_N | ASN_A_54_H | 1.555 | -7.246 | 1.897 | 2.301 | -7.2 | 1.218 | 1.01 | 132.25 | 3.53 |
| 317 | 2jsa-2 | ASN_A_54_N | ASN_A_54_CA | 1.555 | -7.246 | 1.897 | 1.731 | -8.081 | 3.073 | 1.45 | 35.97 | -78.10 |
| 318 | 2jsa-2 | ASN_A_54_CA | ASN_A_54_HA | 1.731 | -8.081 | 3.073 | 0.961 | -7.835 | 3.804 | 1.09 | 47.88 | 162.28 |
| 319 | 2jsa-2 | ASN_A_54_CA | ASN_A_54_C | 1.731 | -8.081 | 3.073 | 3.116 | -7.825 | 3.672 | 1.53 | 66.96 | 10.47 |
| 320 | 2jsa-2 | ASN_A_54_C | ASN_A_54_O | 3.116 | -7.825 | 3.672 | 3.894 | -8.757 | 3.867 | 1.23 | 80.88 | -50.15 |
| 321 | 2jsa-2 | ASN_A_54_C | SER_A_55_N | 3.116 | -7.825 | 3.672 | 3.381 | -6.556 | 3.948 | 1.33 | 77.98 | 78.20 |
| 322 | 2jsa-2 | SER_A_55_N | SER_A_55_H | 3.381 | -6.556 | 3.948 | 2.693 | -5.84 | 3.762 | 1.01 | 100.61 | 133.86 |
| 323 | 2jsa-2 | SER_A_55_N | SER_A_55_CA | 3.381 | -6.556 | 3.948 | 4.657 | -6.166 | 4.521 | 1.45 | 66.76 | 17.00 |
| 324 | 2jsa-2 | SER_A_55_CA | SER_A_55_HA | 4.657 | -6.166 | 4.521 | 4.89 | -6.853 | 5.335 | 1.09 | 41.71 | -71.27 |
| 325 | 2jsa-2 | SER_A_55_CA | SER_A_55_C | 4.657 | -6.166 | 4.521 | 4.565 | -4.749 | 5.088 | 1.53 | 68.23 | 93.71 |
| 326 | 2jsa-2 | SER_A_55_C | SER_A_55_O | 4.565 | -4.749 | 5.088 | 4.938 | -4.51 | 6.237 | 1.23 | 21.08 | 32.65 |
| 327 | 2jsa-2 | SER_A_55_C | LYS_A_56_N | 4.565 | -4.749 | 5.088 | 4.066 | -3.845 | 4.258 | 1.32 | 128.79 | 118.90 |
| 328 | 2jsa-2 | LYS_A_56_N | LYS_A_56_H | 4.066 | -3.845 | 4.258 | 3.78 | -4.112 | 3.327 | 1.01 | 157.20 | -136.97 |
| 329 | 2jsa-2 | LYS_A_56_N | LYS_A_56_CA | 4.066 | -3.845 | 4.258 | 3.92 | -2.456 | 4.663 | 1.45 | 73.83 | 96.00 |
| 330 | 2jsa-2 | LYS_A_56_CA | LYS_A_56_HA | 3.92 | -2.456 | 4.663 | 4.331 | -2.352 | 5.667 | 1.09 | 22.89 | 14.20 |
| 331 | 2jsa-2 | LYS_A_56_CA | LYS_A_56_C | 3.92 | -2.456 | 4.663 | 2.433 | -2.104 | 4.73 | 1.53 | 87.49 | 166.68 |
| 332 | 2jsa-2 | LYS_A_56_C | LYS_A_56_O | 2.433 | -2.104 | 4.73 | 2.074 | -0.933 | 4.853 | 1.23 | 84.27 | 107.04 |
| 333 | 2jsa-2 | LYS_A_56_C | VAL_A_57_N | 2.433 | -2.104 | 4.73 | 1.608 | -3.137 | 4.646 | 1.32 | 93.64 | -128.61 |
| 334 | 2jsa-2 | VAL_A_57_N | VAL_A_57_H | 1.608 | -3.137 | 4.646 | 1.974 | -4.073 | 4.546 | 1.01 | 95.68 | -68.64 |
| 335 | 2jsa-2 | VAL_A_57_N | VAL_A_57_CA | 1.608 | -3.137 | 4.646 | 0.168 | -2.949 | 4.695 | 1.45 | 88.07 | 172.56 |
| 336 | 2jsa-2 | VAL_A_57_CA | VAL_A_57_HA | 0.168 | -2.949 | 4.695 | -0.055 | -1.919 | 4.415 | 1.09 | 104.88 | 102.22 |
| 337 | 2jsa-2 | VAL_A_57_CA | VAL_A_57_C | 0.168 | -2.949 | 4.695 | -0.322 | -3.17 | 6.129 | 1.53 | 20.55 | -155.72 |
| 338 | 2jsa-2 | VAL_A_57_C | VAL_A_57_O | -0.322 | -3.17 | 6.129 | -1.045 | -2.339 | 6.675 | 1.23 | 63.63 | 131.02 |
| 339 | 2jsa-2 | VAL_A_57_C | ASP_A_58_N | -0.322 | -3.17 | 6.129 | 0.092 | -4.293 | 6.696 | 1.32 | 64.65 | -69.76 |
| 340 | 2jsa-2 | ASP_A_58_N | ASP_A_58_H | 0.092 | -4.293 | 6.696 | 0.686 | -4.932 | 6.186 | 1.01 | 120.31 | -47.09 |
| 341 | 2jsa-2 | ASP_A_58_N | ASP_A_58_CA | 0.092 | -4.293 | 6.696 | -0.295 | -4.632 | 8.054 | 1.45 | 20.75 | -138.78 |
| 342 | 2jsa-2 | ASP_A_58_CA | ASP_A_58_HA | -0.295 | -4.632 | 8.054 | -1.347 | -4.916 | 8.064 | 1.09 | 89.47 | -164.89 |
| 343 | 2jsa-2 | ASP_A_58_CA | ASP_A_58_C | -0.295 | -4.632 | 8.054 | -0.075 | -3.419 | 8.961 | 1.53 | 53.66 | 79.72 |
| 344 | 2jsa-2 | ASP_A_58_C | ASP_A_58_O | -0.075 | -3.419 | 8.961 | -0.929 | -3.09 | 9.781 | 1.23 | 48.14 | 158.93 |
| 345 | 2jsa-2 | ASP_A_58_C | PRO_A_59_N | -0.075 | -3.419 | 8.961 | 1.106 | -2.771 | 8.777 | 1.36 | 97.78 | 28.75 |
| 346 | 2jsa-2 | PRO_A_59_N | PRO_A_59_CA | 1.106 | -2.771 | 8.777 | 1.45 | -1.601 | 9.568 | 1.45 | 57.03 | 73.62 |
| 347 | 2jsa-2 | PRO_A_59_CA | PRO_A_59_HA | 1.45 | -1.601 | 9.568 | 1.221 | -1.78 | 10.619 | 1.09 | 15.46 | -141.99 |
| 348 | 2jsa-2 | PRO_A_59_CA | PRO_A_59_C | 1.45 | -1.601 | 9.568 | 0.663 | -0.374 | 9.102 | 1.53 | 107.73 | 122.68 |
| 349 | 2jsa-2 | PRO_A_59_C | PRO_A_59_O | 0.663 | -0.374 | 9.102 | 0.006 | 0.288 | 9.905 | 1.23 | 49.27 | 134.78 |
| 350 | 2jsa-2 | PRO_A_59_C | ILE_A_60_N | 0.663 | -0.374 | 9.102 | 0.756 | -0.108 | 7.807 | 1.33 | 167.72 | 70.73 |
| 351 | 2jsa-2 | ILE_A_60_N | ILE_A_60_H | 0.756 | -0.108 | 7.807 | 1.314 | -0.696 | 7.205 | 1.01 | 126.60 | -46.50 |
| 352 | 2jsa-2 | ILE_A_60_N | ILE_A_60_CA | 0.756 | -0.108 | 7.807 | 0.061 | 1.028 | 7.226 | 1.45 | 113.57 | 121.46 |
| 353 | 2jsa-2 | ILE_A_60_CA | ILE_A_60_HA | 0.061 | 1.028 | 7.226 | 0.53 | 1.938 | 7.601 | 1.09 | 69.88 | 62.73 |
| 354 | 2jsa-2 | ILE_A_60_CA | ILE_A_60_C | 0.061 | 1.028 | 7.226 | -1.395 | 1.027 | 7.697 | 1.53 | 72.07 | -179.96 |
| 355 | 2jsa-2 | ILE_A_60_C | ILE_A_60_O | -1.395 | 1.027 | 7.697 | -1.854 | 1.991 | 8.306 | 1.23 | 60.30 | 115.46 |
| 356 | 2jsa-2 | ILE_A_60_C | ILE_A_61_N | -1.395 | 1.027 | 7.697 | -2.079 | -0.068 | 7.397 | 1.33 | 103.08 | -121.99 |
| 357 | 2jsa-2 | ILE_A_61_N | ILE_A_61_H | -2.079 | -0.068 | 7.397 | -1.635 | -0.824 | 6.895 | 1.01 | 119.79 | -59.57 |
| 358 | 2jsa-2 | ILE_A_61_N | ILE_A_61_CA | -2.079 | -0.068 | 7.397 | -3.473 | -0.207 | 7.782 | 1.45 | 74.63 | -174.31 |
| 359 | 2jsa-2 | ILE_A_61_CA | ILE_A_61_HA | -3.473 | -0.207 | 7.782 | -4.029 | 0.62 | 7.34 | 1.09 | 113.92 | 123.91 |
| 360 | 2jsa-2 | ILE_A_61_CA | ILE_A_61_C | -3.473 | -0.207 | 7.782 | -3.592 | -0.088 | 9.303 | 1.53 | 6.31 | 135.00 |
| 361 | 2jsa-2 | ILE_A_61_C | ILE_A_61_O | -3.592 | -0.088 | 9.303 | -4.663 | 0.221 | 9.823 | 1.23 | 64.99 | 163.91 |
| 362 | 2jsa-2 | ILE_A_61_C | HIS_A_62_N | -3.592 | -0.088 | 9.303 | -2.476 | -0.338 | 9.973 | 1.33 | 59.64 | -12.63 |
| 363 | 2jsa-2 | HIS_A_62_N | HIS_A_62_H | -2.476 | -0.338 | 9.973 | -1.63 | -0.586 | 9.479 | 1.01 | 119.26 | -16.34 |
| 364 | 2jsa-2 | HIS_A_62_N | HIS_A_62_CA | -2.476 | -0.338 | 9.973 | -2.442 | -0.263 | 11.424 | 1.45 | 3.25 | 65.61 |
| 365 | 2jsa-2 | HIS_A_62_CA | HIS_A_62_HA | -2.442 | -0.263 | 11.424 | -3.236 | -0.9 | 11.813 | 1.09 | 69.09 | -141.26 |
| 366 | 2jsa-2 | HIS_A_62_CA | HIS_A_62_C | -2.442 | -0.263 | 11.424 | -2.73 | 1.172 | 11.87 | 1.53 | 73.05 | 101.35 |
| 367 | 2jsa-2 | HIS_A_62_C | HIS_A_62_O | -2.73 | 1.172 | 11.87 | -3.35 | 1.39 | 12.909 | 1.23 | 32.31 | 160.63 |
| 368 | 2jsa-2 | HIS_A_62_C | GLU_A_63_N | -2.73 | 1.172 | 11.87 | -2.265 | 2.113 | 11.061 | 1.33 | 127.62 | 63.70 |
| 369 | 2jsa-2 | GLU_A_63_N | GLU_A_63_H | -2.265 | 2.113 | 11.061 | -1.763 | 1.86 | 10.222 | 1.01 | 146.18 | -26.75 |
| 370 | 2jsa-2 | GLU_A_63_N | GLU_A_63_CA | -2.265 | 2.113 | 11.061 | -2.465 | 3.521 | 11.36 | 1.45 | 78.13 | 98.08 |
| 371 | 2jsa-2 | GLU_A_63_CA | GLU_A_63_HA | -2.465 | 3.521 | 11.36 | -2.348 | 3.658 | 12.435 | 1.09 | 9.51 | 49.50 |
| 372 | 2jsa-2 | GLU_A_63_CA | GLU_A_63_C | -2.465 | 3.521 | 11.36 | -3.881 | 3.951 | 10.972 | 1.53 | 104.69 | 163.11 |
| 373 | 2jsa-2 | GLU_A_63_C | GLU_A_63_O | -3.881 | 3.951 | 10.972 | -4.631 | 4.453 | 11.808 | 1.23 | 47.19 | 146.20 |
| 374 | 2jsa-2 | GLU_A_63_C | LEU_A_64_N | -3.881 | 3.951 | 10.972 | -4.204 | 3.736 | 9.705 | 1.33 | 162.97 | -146.35 |
| 375 | 2jsa-2 | LEU_A_64_N | LEU_A_64_H | -4.204 | 3.736 | 9.705 | -3.536 | 3.317 | 9.074 | 1.01 | 128.67 | -32.10 |
| 376 | 2jsa-2 | LEU_A_64_N | LEU_A_64_CA | -4.204 | 3.736 | 9.705 | -5.517 | 4.095 | 9.197 | 1.45 | 110.47 | 164.71 |
| 377 | 2jsa-2 | LEU_A_64_CA | LEU_A_64_HA | -5.517 | 4.095 | 9.197 | -5.568 | 5.182 | 9.134 | 1.09 | 93.31 | 92.69 |
| 378 | 2jsa-2 | LEU_A_64_CA | LEU_A_64_C | -5.517 | 4.095 | 9.197 | -6.586 | 3.636 | 10.189 | 1.53 | 49.55 | -156.76 |
| 379 | 2jsa-2 | LEU_A_64_C | LEU_A_64_O | -6.586 | 3.636 | 10.189 | -7.537 | 4.366 | 10.466 | 1.23 | 76.99 | 142.49 |
| 380 | 2jsa-2 | LEU_A_64_C | GLU_A_65_N | -6.586 | 3.636 | 10.189 | -6.396 | 2.427 | 10.697 | 1.33 | 67.46 | -81.07 |
| 381 | 2jsa-2 | GLU_A_65_N | GLU_A_65_H | -6.396 | 2.427 | 10.697 | -5.59 | 1.882 | 10.424 | 1.01 | 105.67 | -34.07 |
| 382 | 2jsa-2 | GLU_A_65_N | GLU_A_65_CA | -6.396 | 2.427 | 10.697 | -7.333 | 1.861 | 11.652 | 1.45 | 48.90 | -148.87 |
| 383 | 2jsa-2 | GLU_A_65_CA | GLU_A_65_HA | -7.333 | 1.861 | 11.652 | -8.325 | 1.879 | 11.203 | 1.09 | 114.35 | 178.96 |
| 384 | 2jsa-2 | GLU_A_65_CA | GLU_A_65_C | -7.333 | 1.861 | 11.652 | -7.366 | 2.706 | 12.928 | 1.53 | 33.53 | 92.24 |
| 385 | 2jsa-2 | GLU_A_65_C | GLU_A_65_O | -7.366 | 2.706 | 12.928 | -8.43 | 2.925 | 13.505 | 1.23 | 62.02 | 168.37 |
| 386 | 2jsa-2 | GLU_A_65_C | GLY_A_66_N | -7.366 | 2.706 | 12.928 | -6.187 | 3.16 | 13.33 | 1.33 | 72.35 | 21.06 |

| | | | | | | | | | | | | |
|---|---|---|---|---|---|---|---|---|---|---|---|---|
| 387 | 2jsa-2 | GLY_A_66_N | GLY_A_66_H | -6.187 | 3.16 | 13.33 | -5.352 | 2.942 | 12.805 | 1.01 | 121.31 | -14.63 |
| 388 | 2jsa-2 | GLY_A_66_N | GLY_A_66_CA | -6.187 | 3.16 | 13.33 | -6.068 | 3.976 | 14.526 | 1.45 | 34.59 | 81.70 |
| 389 | 2jsa-2 | GLY_A_66_CA | GLY_A_66_C | -6.068 | 3.976 | 14.526 | -6.851 | 5.283 | 14.378 | 1.53 | 95.55 | 120.93 |
| 390 | 2jsa-2 | GLY_A_66_C | GLY_A_66_O | -6.851 | 5.283 | 14.378 | -7.452 | 5.763 | 15.338 | 1.23 | 38.70 | 141.39 |
| 391 | 2jsa-2 | GLY_A_66_C | GLY_A_67_N | -6.851 | 5.283 | 14.378 | -6.818 | 5.821 | 13.168 | 1.32 | 155.99 | 86.49 |
| 392 | 2jsa-2 | GLY_A_67_N | GLY_A_67_H | -6.818 | 5.821 | 13.168 | -6.302 | 5.371 | 12.425 | 1.01 | 137.34 | -41.09 |
| 393 | 2jsa-2 | GLY_A_67_N | GLY_A_67_CA | -6.818 | 5.821 | 13.168 | -7.517 | 7.063 | 12.881 | 1.45 | 101.39 | 119.37 |
| 394 | 2jsa-2 | GLY_A_67_CA | GLY_A_67_C | -7.517 | 7.063 | 12.881 | -6.668 | 7.979 | 11.997 | 1.53 | 125.29 | 47.17 |
| 395 | 2jsa-2 | GLY_A_67_C | GLY_A_67_O | -6.668 | 7.979 | 11.997 | -7.148 | 9.005 | 11.519 | 1.23 | 112.88 | 115.07 |
| 396 | 2jsa-2 | GLY_A_67_C | THR_A_68_N | -6.668 | 7.979 | 11.997 | -5.421 | 7.573 | 11.805 | 1.33 | 98.33 | -18.03 |
| 397 | 2jsa-2 | THR_A_68_N | THR_A_68_H | -5.421 | 7.573 | 11.805 | -5.093 | 6.717 | 12.229 | 1.01 | 65.18 | -69.03 |
| 398 | 2jsa-2 | THR_A_68_N | THR_A_68_CA | -5.421 | 7.573 | 11.805 | -4.502 | 8.343 | 10.986 | 1.45 | 124.34 | 39.96 |
| 399 | 2jsa-2 | THR_A_68_CA | THR_A_68_HA | -4.502 | 8.343 | 10.986 | -4.426 | 9.35 | 11.398 | 1.09 | 67.81 | 85.68 |
| 400 | 2jsa-2 | THR_A_68_CA | THR_A_68_C | -4.502 | 8.343 | 10.986 | -5.03 | 8.459 | 9.555 | 1.53 | 159.30 | 167.61 |
| 401 | 2jsa-2 | THR_A_68_C | THR_A_68_O | -5.03 | 8.459 | 9.555 | -5.896 | 7.688 | 9.145 | 1.23 | 109.47 | -138.32 |
| 402 | 2jsa-2 | THR_A_68_C | ALA_A_69_N | -5.03 | 8.459 | 9.555 | -4.487 | 9.429 | 8.834 | 1.32 | 122.97 | 60.76 |
| 403 | 2jsa-2 | ALA_A_69_N | ALA_A_69_H | -4.487 | 9.429 | 8.834 | -3.78 | 10.028 | 9.237 | 1.01 | 66.50 | 40.27 |
| 404 | 2jsa-2 | ALA_A_69_N | ALA_A_69_CA | -4.487 | 9.429 | 8.834 | -4.892 | 9.655 | 7.457 | 1.45 | 161.39 | 150.84 |
| 405 | 2jsa-2 | ALA_A_69_CA | ALA_A_69_HA | -4.892 | 9.655 | 7.457 | -5.957 | 9.446 | 7.367 | 1.09 | 94.74 | -168.90 |
| 406 | 2jsa-2 | ALA_A_69_CA | ALA_A_69_C | -4.892 | 9.655 | 7.457 | -4.131 | 8.695 | 6.541 | 1.53 | 126.79 | -51.60 |
| 407 | 2jsa-2 | ALA_A_69_C | ALA_A_69_O | -4.131 | 8.695 | 6.541 | -3.109 | 8.135 | 6.934 | 1.23 | 71.36 | -28.72 |
| 408 | 2jsa-2 | ALA_A_69_C | PRO_A_70_N | -4.131 | 8.695 | 6.541 | -4.674 | 8.528 | 5.305 | 1.36 | 155.32 | -162.90 |
| 409 | 2jsa-2 | PRO_A_70_N | PRO_A_70_CA | -4.674 | 8.528 | 5.305 | -4.058 | 7.644 | 4.331 | 1.45 | 132.11 | -55.13 |
| 410 | 2jsa-2 | PRO_A_70_CA | PRO_A_70_HA | -4.058 | 7.644 | 4.331 | -3.776 | 6.704 | 4.805 | 1.09 | 64.22 | -73.30 |
| 411 | 2jsa-2 | PRO_A_70_CA | PRO_A_70_C | -4.058 | 7.644 | 4.331 | -2.802 | 8.28 | 3.73 | 1.53 | 113.12 | 26.86 |
| 412 | 2jsa-2 | PRO_A_70_C | PRO_A_70_O | -2.802 | 8.28 | 3.73 | -2.072 | 7.634 | 2.979 | 1.23 | 127.61 | -41.51 |
| 413 | 2jsa-2 | PRO_A_70_C | LYS_A_71_N | -2.802 | 8.28 | 3.73 | -2.588 | 9.538 | 4.085 | 1.32 | 74.45 | 80.35 |
| 414 | 2jsa-2 | LYS_A_71_N | LYS_A_71_H | -2.588 | 9.538 | 4.085 | -3.23 | 10.007 | 4.708 | 1.01 | 51.92 | 143.85 |
| 415 | 2jsa-2 | LYS_A_71_N | LYS_A_71_CA | -2.588 | 9.538 | 4.085 | -1.433 | 10.27 | 3.592 | 1.45 | 109.83 | 32.37 |
| 416 | 2jsa-2 | LYS_A_71_CA | LYS_A_71_HA | -1.433 | 10.27 | 3.592 | -0.86 | 9.602 | 2.948 | 1.09 | 126.19 | -49.38 |
| 417 | 2jsa-2 | LYS_A_71_CA | LYS_A_71_C | -1.433 | 10.27 | 3.592 | -0.538 | 10.659 | 4.77 | 1.53 | 39.64 | 23.49 |
| 418 | 2jsa-2 | LYS_A_71_C | LYS_A_71_O | -0.538 | 10.659 | 4.77 | 0.464 | 11.347 | 4.592 | 1.23 | 98.33 | 34.47 |
| 419 | 2jsa-2 | LYS_A_71_C | ASP_A_72_N | -0.538 | 10.659 | 4.77 | -0.935 | 10.2 | 5.949 | 1.33 | 27.24 | -130.86 |
| 420 | 2jsa-2 | ASP_A_72_N | ASP_A_72_H | -0.935 | 10.2 | 5.949 | -1.772 | 9.639 | 6.022 | 1.01 | 85.86 | -146.17 |
| 421 | 2jsa-2 | ASP_A_72_N | ASP_A_72_CA | -0.935 | 10.2 | 5.949 | -0.182 | 10.491 | 7.156 | 1.45 | 33.78 | 21.13 |
| 422 | 2jsa-2 | ASP_A_72_CA | ASP_A_72_HA | -0.182 | 10.491 | 7.156 | 0.602 | 11.211 | 6.922 | 1.09 | 102.40 | 42.56 |
| 423 | 2jsa-2 | ASP_A_72_CA | ASP_A_72_C | -0.182 | 10.491 | 7.156 | 0.441 | 9.199 | 7.688 | 1.53 | 69.65 | -64.26 |
| 424 | 2jsa-2 | ASP_A_72_C | ASP_A_72_O | 0.441 | 9.199 | 7.688 | 1.499 | 9.228 | 8.316 | 1.23 | 59.32 | 1.57 |
| 425 | 2jsa-2 | ASP_A_72_C | VAL_A_73_N | 0.441 | 9.199 | 7.688 | -0.241 | 8.096 | 7.417 | 1.32 | 101.80 | -121.73 |
| 426 | 2jsa-2 | VAL_A_73_N | VAL_A_73_H | -0.241 | 8.096 | 7.417 | -1.104 | 8.147 | 6.896 | 1.01 | 121.08 | 176.62 |
| 427 | 2jsa-2 | VAL_A_73_N | VAL_A_73_CA | -0.241 | 8.096 | 7.417 | 0.233 | 6.796 | 7.86 | 1.45 | 72.25 | -69.97 |
| 428 | 2jsa-2 | VAL_A_73_CA | VAL_A_73_HA | 0.233 | 6.796 | 7.86 | 0.099 | 6.731 | 8.94 | 1.09 | 7.85 | -154.12 |
| 429 | 2jsa-2 | VAL_A_73_CA | VAL_A_73_C | 0.233 | 6.796 | 7.86 | 1.728 | 6.672 | 7.559 | 1.53 | 101.35 | -4.74 |
| 430 | 2jsa-2 | VAL_A_73_C | VAL_A_73_O | 1.728 | 6.672 | 7.559 | 2.562 | 7.092 | 8.359 | 1.23 | 49.41 | 26.73 |
| 431 | 2jsa-2 | VAL_A_73_C | CYS_A_74_N | 1.728 | 6.672 | 7.559 | 2.02 | 6.095 | 6.403 | 1.32 | 150.78 | -63.16 |
| 432 | 2jsa-2 | CYS_A_74_N | CYS_A_74_H | 2.02 | 6.095 | 6.403 | 1.282 | 5.775 | 5.793 | 1.01 | 127.17 | -156.56 |
| 433 | 2jsa-2 | CYS_A_74_N | CYS_A_74_CA | 2.02 | 6.095 | 6.403 | 3.4 | 5.911 | 5.987 | 1.45 | 106.64 | -7.59 |
| 434 | 2jsa-2 | CYS_A_74_CA | CYS_A_74_HA | 3.4 | 5.911 | 5.987 | 3.789 | 5.016 | 6.473 | 1.09 | 63.53 | -66.51 |
| 435 | 2jsa-2 | CYS_A_74_CA | CYS_A_74_C | 3.4 | 5.911 | 5.987 | 4.212 | 7.116 | 6.466 | 1.53 | 71.76 | 56.03 |
| 436 | 2jsa-2 | CYS_A_74_C | CYS_A_74_O | 4.212 | 7.116 | 6.466 | 5.289 | 6.956 | 7.037 | 1.23 | 62.33 | -8.45 |
| 437 | 2jsa-2 | CYS_A_74_C | THR_A_75_N | 4.212 | 7.116 | 6.466 | 3.662 | 8.296 | 6.216 | 1.33 | 100.87 | 114.99 |
| 438 | 2jsa-2 | THR_A_75_N | THR_A_75_H | 3.662 | 8.296 | 6.216 | 2.772 | 8.352 | 5.743 | 1.01 | 117.94 | 176.40 |
| 439 | 2jsa-2 | THR_A_75_N | THR_A_75_CA | 3.662 | 8.296 | 6.216 | 4.321 | 9.527 | 6.614 | 1.45 | 74.09 | 61.84 |
| 440 | 2jsa-2 | THR_A_75_CA | THR_A_75_HA | 4.321 | 9.527 | 6.614 | 5.092 | 9.764 | 5.88 | 1.09 | 132.30 | 17.09 |
| 441 | 2jsa-2 | THR_A_75_CA | THR_A_75_C | 4.321 | 9.527 | 6.614 | 5.002 | 9.353 | 7.973 | 1.53 | 27.35 | -14.33 |
| 442 | 2jsa-2 | THR_A_75_C | THR_A_75_O | 5.002 | 9.353 | 7.973 | 6.23 | 9.306 | 8.055 | 1.23 | 86.18 | -2.19 |
| 443 | 2jsa-2 | THR_A_75_C | LYS_A_76_N | 5.002 | 9.353 | 7.973 | 4.177 | 9.259 | 9.005 | 1.32 | 38.82 | -173.50 |
| 444 | 2jsa-2 | LYS_A_76_N | LYS_A_76_H | 4.177 | 9.259 | 9.005 | 3.178 | 9.304 | 8.863 | 1.01 | 98.08 | 177.42 |
| 445 | 2jsa-2 | LYS_A_76_N | LYS_A_76_CA | 4.177 | 9.259 | 9.005 | 4.684 | 9.091 | 10.356 | 1.45 | 21.57 | -18.33 |
| 446 | 2jsa-2 | LYS_A_76_CA | LYS_A_76_HA | 4.684 | 9.091 | 10.356 | 5.459 | 9.84 | 10.522 | 1.09 | 81.24 | 44.02 |
| 447 | 2jsa-2 | LYS_A_76_CA | LYS_A_76_C | 4.684 | 9.091 | 10.356 | 5.336 | 7.712 | 10.483 | 1.53 | 85.24 | -64.69 |
| 448 | 2jsa-2 | LYS_A_76_C | LYS_A_76_O | 5.336 | 7.712 | 10.483 | 6.222 | 7.514 | 11.312 | 1.23 | 47.60 | -12.60 |
| 449 | 2jsa-2 | LYS_A_76_C | LEU_A_77_N | 5.336 | 7.712 | 10.483 | 4.87 | 6.793 | 9.649 | 1.33 | 128.99 | -116.89 |
| 450 | 2jsa-2 | LEU_A_77_N | LEU_A_77_H | 4.87 | 6.793 | 9.649 | 4.139 | 7.026 | 8.993 | 1.01 | 130.53 | 162.32 |
| 451 | 2jsa-2 | LEU_A_77_N | LEU_A_77_CA | 4.87 | 6.793 | 9.649 | 5.395 | 5.439 | 9.657 | 1.45 | 89.68 | -68.81 |
| 452 | 2jsa-2 | LEU_A_77_CA | LEU_A_77_HA | 5.395 | 5.439 | 9.657 | 5.6 | 5.163 | 10.692 | 1.09 | 18.38 | -53.40 |
| 453 | 2jsa-2 | LEU_A_77_CA | LEU_A_77_C | 5.395 | 5.439 | 9.657 | 6.722 | 5.407 | 8.896 | 1.53 | 119.83 | -1.38 |
| 454 | 2jsa-2 | LEU_A_77_C | LEU_A_77_O | 6.722 | 5.407 | 8.896 | 7.258 | 4.334 | 8.621 | 1.23 | 102.91 | -63.46 |
| 455 | 2jsa-2 | LEU_A_77_C | ASN_A_78_N | 6.722 | 5.407 | 8.896 | 7.213 | 6.596 | 8.577 | 1.33 | 103.93 | 67.56 |
| 456 | 2jsa-2 | ASN_A_78_N | ASN_A_78_H | 7.213 | 6.596 | 8.577 | 6.718 | 7.437 | 8.836 | 1.01 | 75.14 | 120.48 |
| 457 | 2jsa-2 | ASN_A_78_N | ASN_A_78_CA | 7.213 | 6.596 | 8.577 | 8.468 | 6.717 | 7.853 | 1.45 | 119.87 | 5.51 |
| 458 | 2jsa-2 | ASN_A_78_CA | ASN_A_78_HA | 8.468 | 6.717 | 7.853 | 8.535 | 7.715 | 7.42 | 1.09 | 113.41 | 86.16 |
| 459 | 2jsa-2 | ASN_A_78_CA | ASN_A_78_C | 8.468 | 6.717 | 7.853 | 8.515 | 5.664 | 6.743 | 1.53 | 136.48 | -87.44 |
| 460 | 2jsa-2 | ASN_A_78_C | ASN_A_78_O | 8.515 | 5.664 | 6.743 | 9.591 | 5.212 | 6.357 | 1.23 | 108.30 | -22.79 |
| 461 | 2jsa-2 | ASN_A_78_C | GLU_A_79_N | 8.515 | 5.664 | 6.743 | 7.334 | 5.303 | 6.263 | 1.32 | 111.24 | -163.00 |
| 462 | 2jsa-2 | GLU_A_79_N | GLU_A_79_H | 7.334 | 5.303 | 6.263 | 6.489 | 5.716 | 6.63 | 1.01 | 68.68 | 153.95 |
| 463 | 2jsa-2 | GLU_A_79_N | GLU_A_79_CA | 7.334 | 5.303 | 6.263 | 7.227 | 4.313 | 5.205 | 1.45 | 136.74 | -96.17 |
| 464 | 2jsa-2 | GLU_A_79_CA | GLU_A_79_HA | 7.227 | 4.313 | 5.205 | 8.203 | 3.843 | 5.084 | 1.09 | 96.37 | -25.71 |
| 465 | 2jsa-2 | GLU_A_79_CA | GLU_A_79_C | 7.227 | 4.313 | 5.205 | 6.85 | 4.985 | 3.884 | 1.53 | 149.75 | 119.29 |
| 466 | 2jsa-2 | GLU_A_79_C | GLU_A_79_O | 6.85 | 4.985 | 3.884 | 5.756 | 4.768 | 3.362 | 1.23 | 115.08 | -168.78 |
| 467 | 2jsa-2 | GLU_A_79_C | CYS_A_80_N | 6.85 | 4.985 | 3.884 | 7.775 | 5.787 | 3.379 | 1.32 | 112.42 | 40.93 |
| 468 | 2jsa-2 | CYS_A_80_N | CYS_A_80_H | 7.775 | 5.787 | 3.379 | 8.653 | 5.918 | 3.861 | 1.01 | 61.50 | 8.49 |
| 469 | 2jsa-2 | CYS_A_80_N | CYS_A_80_CA | 7.775 | 5.787 | 3.379 | 7.553 | 6.492 | 2.128 | 1.45 | 149.42 | 107.48 |
| 470 | 2jsa-2 | CYS_A_80_CA | CYS_A_80_HA | 7.553 | 6.492 | 2.128 | 7.615 | 5.771 | 1.313 | 1.09 | 138.40 | -85.09 |
| 471 | 2jsa-2 | CYS_A_80_CA | CYS_A_80_C | 7.553 | 6.492 | 2.128 | 8.67 | 7.523 | 1.952 | 1.53 | 96.60 | 42.71 |
| 472 | 2jsa-2 | CYS_A_80_C | CYS_A_80_O | 8.67 | 7.523 | 1.952 | 9.354 | 7.534 | 0.93 | 1.23 | 146.20 | 0.92 |
| 473 | 2jsa-2 | CYS_A_80_C | PRO_A_81_N | 8.67 | 7.523 | 1.952 | 8.824 | 8.387 | 2.992 | 1.36 | 40.16 | 79.89 |
| 474 | 2jsa-2 | PRO_A_81_N | PRO_A_81_CA | 8.824 | 8.387 | 2.992 | 9.845 | 9.419 | 2.963 | 1.45 | 91.14 | 45.31 |
| 475 | 2jsa-2 | PRO_A_81_CA | PRO_A_81_HA | 9.845 | 9.419 | 2.963 | 9.848 | 9.914 | 1.992 | 1.09 | 152.99 | 89.65 |
| 476 | 2jsa-2 | PRO_A_81_CA | PRO_A_81_C | 9.845 | 9.419 | 2.963 | 11.233 | 8.823 | 3.207 | 1.53 | 80.82 | -23.24 |
| 477 | 2jsa-2 | PRO_A_81_C | PRO_A_81_O | 11.233 | 8.823 | 3.207 | 11.821 | 9.025 | 4.269 | 1.23 | 30.35 | 18.96 |
| 1 | 2jsa-3 | ARG_A_1_N | ARG_A_1_H | 14.994 | 0.726 | -3.438 | 15.81 | 1.274 | -3.203 | 1.01 | 76.55 | 33.88 |
| 2 | 2jsa-3 | ARG_A_1_N | ARG_A_1_CA | 14.994 | 0.726 | -3.438 | 13.682 | 1.257 | -3.109 | 1.45 | 76.91 | 157.97 |
| 3 | 2jsa-3 | ARG_A_1_CA | ARG_A_1_HA | 13.682 | 1.257 | -3.109 | 13.116 | 1.371 | -4.033 | 1.09 | 148.00 | 168.61 |
| 4 | 2jsa-3 | ARG_A_1_CA | ARG_A_1_C | 13.682 | 1.257 | -3.109 | 13.816 | 2.625 | -2.435 | 1.53 | 63.88 | 84.41 |
| 5 | 2jsa-3 | ARG_A_1_C | ARG_A_1_O | 13.816 | 2.625 | -2.435 | 14.679 | 2.818 | -1.58 | 1.23 | 45.97 | 12.61 |

| | | | | | | | | | | | |
|---|---|---|---|---|---|---|---|---|---|---|---|
| 6 | 2jsa-3 | ARG_A_1_C | SER_A_2_N | 13.816 | 2.625 | -2.435 | 12.949 | 3.538 | -2.847 | 1.32 | 108.12 | 133.52 |
| 7 | 2jsa-3 | SER_A_2_N | SER_A_2_H | 12.949 | 3.538 | -2.847 | 12.266 | 3.309 | -3.555 | 1.01 | 134.50 | -161.46 |
| 8 | 2jsa-3 | SER_A_2_N | SER_A_2_CA | 12.949 | 3.538 | -2.847 | 12.959 | 4.882 | -2.294 | 1.45 | 67.64 | 89.57 |
| 9 | 2jsa-3 | SER_A_2_CA | SER_A_2_HA | 12.959 | 4.882 | -2.294 | 12.692 | 4.82 | -1.239 | 1.09 | 14.56 | -166.93 |
| 10 | 2jsa-3 | SER_A_2_CA | SER_A_2_C | 12.959 | 4.882 | -2.294 | 11.925 | 5.751 | -3.011 | 1.53 | 117.96 | 139.96 |
| 11 | 2jsa-3 | SER_A_2_C | SER_A_2_O | 11.925 | 5.751 | -3.011 | 12.216 | 6.883 | -3.391 | 1.23 | 108.01 | 75.58 |
| 12 | 2jsa-3 | SER_A_2_C | ALA_A_3_N | 11.925 | 5.751 | -3.011 | 10.736 | 5.187 | -3.172 | 1.33 | 96.98 | -154.62 |
| 13 | 2jsa-3 | ALA_A_3_N | ALA_A_3_H | 10.736 | 5.187 | -3.172 | 10.568 | 4.251 | -2.833 | 1.01 | 70.38 | -100.18 |
| 14 | 2jsa-3 | ALA_A_3_N | ALA_A_3_CA | 10.736 | 5.187 | -3.172 | 9.655 | 5.896 | -3.836 | 1.45 | 117.19 | 146.74 |
| 15 | 2jsa-3 | ALA_A_3_CA | ALA_A_3_HA | 9.655 | 5.896 | -3.836 | 9.388 | 6.767 | -3.238 | 1.09 | 56.72 | 107.04 |
| 16 | 2jsa-3 | ALA_A_3_CA | ALA_A_3_C | 9.655 | 5.896 | -3.836 | 8.433 | 4.981 | -3.93 | 1.53 | 93.52 | -143.18 |
| 17 | 2jsa-3 | ALA_A_3_C | ALA_A_3_O | 8.433 | 4.981 | -3.93 | 7.393 | 5.267 | -3.339 | 1.23 | 61.28 | 164.62 |
| 18 | 2jsa-3 | ALA_A_3_C | LEU_A_4_N | 8.433 | 4.981 | -3.93 | 8.599 | 3.9 | -4.678 | 1.32 | 124.37 | -81.27 |
| 19 | 2jsa-3 | LEU_A_4_N | LEU_A_4_H | 8.599 | 3.9 | -4.678 | 9.483 | 3.728 | -5.135 | 1.01 | 116.91 | -11.01 |
| 20 | 2jsa-3 | LEU_A_4_N | LEU_A_4_CA | 8.599 | 3.9 | -4.678 | 7.521 | 2.942 | -4.857 | 1.45 | 97.08 | -138.37 |
| 21 | 2jsa-3 | LEU_A_4_CA | LEU_A_4_HA | 7.521 | 2.942 | -4.857 | 6.75 | 3.412 | -5.467 | 1.09 | 124.04 | 148.63 |
| 22 | 2jsa-3 | LEU_A_4_CA | LEU_A_4_C | 7.521 | 2.942 | -4.857 | 6.904 | 2.615 | -3.496 | 1.53 | 27.16 | -152.08 |
| 23 | 2jsa-3 | LEU_A_4_C | LEU_A_4_O | 6.904 | 2.615 | -3.496 | 5.71 | 2.825 | -3.285 | 1.23 | 80.13 | 170.02 |
| 24 | 2jsa-3 | LEU_A_4_C | SER_A_5_N | 6.904 | 2.615 | -3.496 | 7.745 | 2.106 | -2.607 | 1.33 | 47.88 | -31.18 |
| 25 | 2jsa-3 | SER_A_5_N | SER_A_5_H | 7.745 | 2.106 | -2.607 | 8.715 | 1.961 | -2.849 | 1.01 | 103.86 | -8.50 |
| 26 | 2jsa-3 | SER_A_5_N | SER_A_5_CA | 7.745 | 2.106 | -2.607 | 7.296 | 1.748 | -1.273 | 1.45 | 23.29 | -141.43 |
| 27 | 2jsa-3 | SER_A_5_CA | SER_A_5_HA | 7.296 | 1.748 | -1.273 | 6.827 | 0.766 | -1.323 | 1.09 | 92.63 | -115.53 |
| 28 | 2jsa-3 | SER_A_5_CA | SER_A_5_C | 7.296 | 1.748 | -1.273 | 6.262 | 2.762 | -0.778 | 1.53 | 71.13 | 135.56 |
| 29 | 2jsa-3 | SER_A_5_C | SER_A_5_O | 6.262 | 2.762 | -0.778 | 5.193 | 2.381 | -0.301 | 1.23 | 67.20 | -160.38 |
| 30 | 2jsa-3 | SER_A_5_C | CYS_A_6_N | 6.262 | 2.762 | -0.778 | 6.615 | 4.032 | -0.91 | 1.32 | 95.72 | 74.47 |
| 31 | 2jsa-3 | CYS_A_6_N | CYS_A_6_H | 6.615 | 4.032 | -0.91 | 7.51 | 4.272 | -1.313 | 1.01 | 113.50 | 15.01 |
| 32 | 2jsa-3 | CYS_A_6_N | CYS_A_6_CA | 6.615 | 4.032 | -0.91 | 5.731 | 5.103 | -0.483 | 1.45 | 72.91 | 129.54 |
| 33 | 2jsa-3 | CYS_A_6_CA | CYS_A_6_HA | 5.731 | 5.103 | -0.483 | 5.443 | 4.918 | 0.552 | 1.09 | 18.30 | -147.28 |
| 34 | 2jsa-3 | CYS_A_6_CA | CYS_A_6_C | 5.731 | 5.103 | -0.483 | 4.473 | 5.062 | -1.354 | 1.53 | 124.68 | -178.13 |
| 35 | 2jsa-3 | CYS_A_6_C | CYS_A_6_O | 4.473 | 5.062 | -1.354 | 3.38 | 4.791 | -0.861 | 1.23 | 66.36 | -166.07 |
| 36 | 2jsa-3 | CYS_A_6_C | GLN_A_7_N | 4.473 | 5.062 | -1.354 | 4.67 | 5.339 | -2.635 | 1.33 | 165.14 | 54.58 |
| 37 | 2jsa-3 | GLN_A_7_N | GLN_A_7_H | 4.67 | 5.339 | -2.635 | 5.598 | 5.556 | -2.971 | 1.01 | 109.42 | 13.16 |
| 38 | 2jsa-3 | GLN_A_7_N | GLN_A_7_CA | 4.67 | 5.339 | -2.635 | 3.566 | 5.339 | -3.579 | 1.45 | 130.53 | 180.00 |
| 39 | 2jsa-3 | GLN_A_7_CA | GLN_A_7_HA | 3.566 | 5.339 | -3.579 | 3.031 | 6.282 | -3.471 | 1.09 | 84.31 | 119.57 |
| 40 | 2jsa-3 | GLN_A_7_CA | GLN_A_7_C | 3.566 | 5.339 | -3.579 | 2.598 | 4.197 | -3.263 | 1.53 | 78.08 | -130.29 |
| 41 | 2jsa-3 | GLN_A_7_C | GLN_A_7_O | 2.598 | 4.197 | -3.263 | 1.398 | 4.42 | -3.114 | 1.23 | 83.04 | 169.47 |
| 42 | 2jsa-3 | GLN_A_7_C | MET_A_8_N | 2.598 | 4.197 | -3.263 | 3.156 | 2.999 | -3.171 | 1.32 | 86.02 | -65.03 |
| 43 | 2jsa-3 | MET_A_8_N | MET_A_8_H | 3.156 | 2.999 | -3.171 | 4.151 | 2.89 | -3.308 | 1.01 | 97.79 | -6.25 |
| 44 | 2jsa-3 | MET_A_8_N | MET_A_8_CA | 3.156 | 2.999 | -3.171 | 2.357 | 1.822 | -2.875 | 1.45 | 78.25 | -124.17 |
| 45 | 2jsa-3 | MET_A_8_CA | MET_A_8_HA | 2.357 | 1.822 | -2.875 | 1.717 | 1.61 | -3.731 | 1.09 | 141.78 | -161.67 |
| 46 | 2jsa-3 | MET_A_8_CA | MET_A_8_C | 2.357 | 1.822 | -2.875 | 1.479 | 2.05 | -1.643 | 1.53 | 36.36 | 165.44 |
| 47 | 2jsa-3 | MET_A_8_C | MET_A_8_O | 1.479 | 2.05 | -1.643 | 0.35 | 1.565 | -1.581 | 1.23 | 87.11 | -156.75 |
| 48 | 2jsa-3 | MET_A_8_C | CYS_A_9_N | 1.479 | 2.05 | -1.643 | 2.031 | 2.79 | -0.692 | 1.33 | 44.15 | 53.28 |
| 49 | 2jsa-3 | CYS_A_9_N | CYS_A_9_H | 2.031 | 2.79 | -0.692 | 2.965 | 3.158 | -0.81 | 1.01 | 96.70 | 21.50 |
| 50 | 2jsa-3 | CYS_A_9_N | CYS_A_9_CA | 2.031 | 2.79 | -0.692 | 1.312 | 3.089 | 0.535 | 1.45 | 32.40 | 157.42 |
| 51 | 2jsa-3 | CYS_A_9_CA | CYS_A_9_HA | 1.312 | 3.089 | 0.535 | 1.03 | 2.147 | 1.005 | 1.09 | 64.45 | -106.67 |
| 52 | 2jsa-3 | CYS_A_9_CA | CYS_A_9_C | 1.312 | 3.089 | 0.535 | 0.038 | 3.853 | 0.172 | 1.53 | 103.73 | 149.05 |
| 53 | 2jsa-3 | CYS_A_9_C | CYS_A_9_O | 0.038 | 3.853 | 0.172 | -0.99 | 3.703 | 0.83 | 1.23 | 57.65 | -171.70 |
| 54 | 2jsa-3 | CYS_A_9_C | GLU_A_10_N | 0.038 | 3.853 | 0.172 | 0.147 | 4.657 | -0.876 | 1.33 | 142.25 | 82.28 |
| 55 | 2jsa-3 | GLU_A_10_N | GLU_A_10_H | 0.147 | 4.657 | -0.876 | 1.024 | 4.73 | -1.371 | 1.01 | 119.36 | 4.76 |
| 56 | 2jsa-3 | GLU_A_10_N | GLU_A_10_CA | 0.147 | 4.657 | -0.876 | -0.984 | 5.446 | -1.335 | 1.45 | 108.41 | 145.10 |
| 57 | 2jsa-3 | GLU_A_10_CA | GLU_A_10_HA | -0.984 | 5.446 | -1.335 | -1.522 | 5.806 | -0.458 | 1.09 | 36.43 | 146.21 |
| 58 | 2jsa-3 | GLU_A_10_CA | GLU_A_10_C | -0.984 | 5.446 | -1.335 | -1.936 | 4.579 | -2.162 | 1.53 | 122.71 | -137.68 |
| 59 | 2jsa-3 | GLU_A_10_C | GLU_A_10_O | -1.936 | 4.579 | -2.162 | -3.031 | 5.014 | -2.513 | 1.23 | 106.59 | 158.33 |
| 60 | 2jsa-3 | GLU_A_10_C | LEU_A_11_N | -1.936 | 4.579 | -2.162 | -1.481 | 3.367 | -2.449 | 1.33 | 102.50 | -69.42 |
| 61 | 2jsa-3 | LEU_A_11_N | LEU_A_11_H | -1.481 | 3.367 | -2.449 | -0.568 | 3.076 | -2.128 | 1.01 | 71.48 | -17.68 |
| 62 | 2jsa-3 | LEU_A_11_N | LEU_A_11_CA | -1.481 | 3.367 | -2.449 | -2.279 | 2.435 | -3.227 | 1.45 | 122.38 | -130.57 |
| 63 | 2jsa-3 | LEU_A_11_CA | LEU_A_11_HA | -2.279 | 2.435 | -3.227 | -2.954 | 3.014 | -3.857 | 1.09 | 125.31 | 139.38 |
| 64 | 2jsa-3 | LEU_A_11_CA | LEU_A_11_C | -2.279 | 2.435 | -3.227 | -3.136 | 1.589 | -2.282 | 1.53 | 51.88 | -135.37 |
| 65 | 2jsa-3 | LEU_A_11_C | LEU_A_11_O | -3.136 | 1.589 | -2.282 | -4.298 | 1.313 | -2.572 | 1.23 | 103.65 | -166.64 |
| 66 | 2jsa-3 | LEU_A_11_C | VAL_A_12_N | -3.136 | 1.589 | -2.282 | -2.526 | 1.202 | -1.172 | 1.32 | 33.06 | -32.39 |
| 67 | 2jsa-3 | VAL_A_12_N | VAL_A_12_H | -2.526 | 1.202 | -1.172 | -1.566 | 1.466 | -1.0 | 1.01 | 80.20 | 15.38 |
| 68 | 2jsa-3 | VAL_A_12_N | VAL_A_12_CA | -2.526 | 1.202 | -1.172 | -3.218 | 0.393 | -0.182 | 1.45 | 47.08 | -130.54 |
| 69 | 2jsa-3 | VAL_A_12_CA | VAL_A_12_HA | -3.218 | 0.393 | -0.182 | -3.766 | -0.392 | -0.702 | 1.09 | 118.51 | -124.92 |
| 70 | 2jsa-3 | VAL_A_12_CA | VAL_A_12_C | -3.218 | 0.393 | -0.182 | -4.23 | 1.265 | 0.566 | 1.53 | 60.75 | 139.25 |
| 71 | 2jsa-3 | VAL_A_12_C | VAL_A_12_O | -4.23 | 1.265 | 0.566 | -5.393 | 0.89 | 0.707 | 1.23 | 83.42 | -162.13 |
| 72 | 2jsa-3 | VAL_A_12_C | VAL_A_13_N | -4.23 | 1.265 | 0.566 | -3.75 | 2.412 | 1.024 | 1.33 | 69.78 | 67.29 |
| 73 | 2jsa-3 | VAL_A_13_N | VAL_A_13_H | -3.75 | 2.412 | 1.024 | -2.782 | 2.656 | 0.871 | 1.01 | 98.71 | 14.15 |
| 74 | 2jsa-3 | VAL_A_13_N | VAL_A_13_CA | -3.75 | 2.412 | 1.024 | -4.598 | 3.34 | 1.753 | 1.45 | 59.89 | 132.42 |
| 75 | 2jsa-3 | VAL_A_13_CA | VAL_A_13_HA | -4.598 | 3.34 | 1.753 | -4.98 | 2.833 | 2.639 | 1.09 | 35.62 | -127.00 |
| 76 | 2jsa-3 | VAL_A_13_CA | VAL_A_13_C | -4.598 | 3.34 | 1.753 | -5.791 | 3.73 | 0.877 | 1.53 | 124.91 | 161.90 |
| 77 | 2jsa-3 | VAL_A_13_C | VAL_A_13_O | -5.791 | 3.73 | 0.877 | -6.941 | 3.517 | 1.257 | 1.23 | 72.00 | -169.51 |
| 78 | 2jsa-3 | VAL_A_13_C | LYS_A_14_N | -5.791 | 3.73 | 0.877 | -5.475 | 4.296 | -0.278 | 1.32 | 150.70 | 60.83 |
| 79 | 2jsa-3 | LYS_A_14_N | LYS_A_14_H | -5.475 | 4.296 | -0.278 | -4.506 | 4.442 | -0.525 | 1.01 | 104.15 | 8.57 |
| 80 | 2jsa-3 | LYS_A_14_N | LYS_A_14_CA | -5.475 | 4.296 | -0.278 | -6.506 | 4.718 | -1.212 | 1.45 | 129.98 | 157.74 |
| 81 | 2jsa-3 | LYS_A_14_CA | LYS_A_14_HA | -6.506 | 4.718 | -1.212 | -6.988 | 5.606 | -0.803 | 1.09 | 67.96 | 118.49 |
| 82 | 2jsa-3 | LYS_A_14_CA | LYS_A_14_C | -6.506 | 4.718 | -1.212 | -7.567 | 3.622 | -1.323 | 1.53 | 94.16 | -134.07 |
| 83 | 2jsa-3 | LYS_A_14_C | LYS_A_14_O | -7.567 | 3.622 | -1.323 | -8.75 | 3.913 | -1.492 | 1.23 | 97.90 | 166.18 |
| 84 | 2jsa-3 | LYS_A_14_C | LYS_A_15_N | -7.567 | 3.622 | -1.323 | -7.106 | 2.383 | -1.221 | 1.33 | 85.59 | -69.59 |
| 85 | 2jsa-3 | LYS_A_15_N | LYS_A_15_H | -7.106 | 2.383 | -1.221 | -6.119 | 2.217 | -1.08 | 1.01 | 81.98 | -9.55 |
| 86 | 2jsa-3 | LYS_A_15_N | LYS_A_15_CA | -7.106 | 2.383 | -1.221 | -8.001 | 1.242 | -1.308 | 1.45 | 93.43 | -128.11 |
| 87 | 2jsa-3 | LYS_A_15_CA | LYS_A_15_HA | -8.001 | 1.242 | -1.308 | -8.74 | 1.451 | -2.081 | 1.09 | 135.19 | 164.21 |
| 88 | 2jsa-3 | LYS_A_15_CA | LYS_A_15_C | -8.001 | 1.242 | -1.308 | -8.748 | 1.083 | 0.017 | 1.53 | 29.96 | -167.98 |
| 89 | 2jsa-3 | LYS_A_15_C | LYS_A_15_O | -8.748 | 1.083 | 0.017 | -9.937 | 0.769 | 0.031 | 1.23 | 89.35 | -165.21 |
| 90 | 2jsa-3 | LYS_A_15_C | TYR_A_16_N | -8.748 | 1.083 | 0.017 | -8.019 | 1.306 | 1.101 | 1.33 | 35.12 | 17.01 |
| 91 | 2jsa-3 | TYR_A_16_N | TYR_A_16_H | -8.019 | 1.306 | 1.101 | -7.045 | 1.56 | 1.015 | 1.01 | 94.88 | 14.62 |
| 92 | 2jsa-3 | TYR_A_16_N | TYR_A_16_CA | -8.019 | 1.306 | 1.101 | -8.596 | 1.191 | 2.429 | 1.45 | 23.89 | -168.73 |
| 93 | 2jsa-3 | TYR_A_16_CA | TYR_A_16_HA | -8.596 | 1.191 | 2.429 | -9.175 | 0.271 | 2.51 | 1.09 | 85.74 | -122.18 |
| 94 | 2jsa-3 | TYR_A_16_CA | TYR_A_16_C | -8.596 | 1.191 | 2.429 | -9.486 | 2.395 | 2.746 | 1.53 | 78.05 | 126.47 |
| 95 | 2jsa-3 | TYR_A_16_C | TYR_A_16_O | -9.486 | 2.395 | 2.746 | -10.704 | 2.259 | 2.863 | 1.23 | 84.55 | -173.63 |
| 96 | 2jsa-3 | TYR_A_16_C | GLU_A_17_N | -9.486 | 2.395 | 2.746 | -8.844 | 3.547 | 2.876 | 1.33 | 84.37 | 60.87 |
| 97 | 2jsa-3 | GLU_A_17_N | GLU_A_17_H | -8.844 | 3.547 | 2.876 | -7.84 | 3.585 | 2.766 | 1.01 | 96.25 | 2.17 |
| 98 | 2jsa-3 | GLU_A_17_N | GLU_A_17_CA | -8.844 | 3.547 | 2.876 | -9.561 | 4.774 | 3.176 | 1.45 | 78.08 | 120.30 |
| 99 | 2jsa-3 | GLU_A_17_CA | GLU_A_17_HA | -9.561 | 4.774 | 3.176 | -9.788 | 4.779 | 4.242 | 1.09 | 12.02 | 178.74 |
| 100 | 2jsa-3 | GLU_A_17_CA | GLU_A_17_C | -9.561 | 4.774 | 3.176 | -10.881 | 4.821 | 2.402 | 1.53 | 120.37 | 177.96 |
| 101 | 2jsa-3 | GLU_A_17_C | GLU_A_17_O | -10.881 | 4.821 | 2.402 | -11.904 | 5.243 | 2.939 | 1.23 | 64.11 | 157.58 |

| | | | | | | | | | | | | |
|---|---|---|---|---|---|---|---|---|---|---|---|---|
| 102 | 2jsa-3 | GLU_A_17_C | GLY_A_18_N | -10.881 | 4.821 | 2.402 | -10.814 | 4.38 | 1.155 | 1.32 | 160.32 | -81.36 |
| 103 | 2jsa-3 | GLY_A_18_N | GLY_A_18_H | -10.814 | 4.38 | 1.155 | -9.937 | 4.046 | 0.78 | 1.01 | 111.78 | -20.85 |
| 104 | 2jsa-3 | GLY_A_18_N | GLY_A_18_CA | -10.814 | 4.38 | 1.155 | -11.992 | 4.366 | 0.303 | 1.45 | 125.87 | -179.32 |
| 105 | 2jsa-3 | GLY_A_18_CA | GLY_A_18_C | -11.992 | 4.366 | 0.303 | -12.343 | 2.94 | -0.124 | 1.53 | 106.21 | -103.83 |
| 106 | 2jsa-3 | GLY_A_18_C | GLY_A_18_O | -12.343 | 2.94 | -0.124 | -12.578 | 2.681 | -1.303 | 1.23 | 163.48 | -132.22 |
| 107 | 2jsa-3 | GLY_A_18_C | SER_A_19_N | -12.343 | 2.94 | -0.124 | -12.367 | 2.051 | 0.859 | 1.33 | 42.14 | -91.55 |
| 108 | 2jsa-3 | SER_A_19_N | SER_A_19_H | -12.367 | 2.051 | 0.859 | -12.163 | 2.335 | 1.807 | 1.01 | 20.25 | 54.31 |
| 109 | 2jsa-3 | SER_A_19_N | SER_A_19_CA | -12.367 | 2.051 | 0.859 | -12.686 | 0.658 | 0.599 | 1.45 | 100.31 | -102.90 |
| 110 | 2jsa-3 | SER_A_19_CA | SER_A_19_HA | -12.686 | 0.658 | 0.599 | -12.237 | 0.376 | -0.354 | 1.09 | 150.91 | -32.13 |
| 111 | 2jsa-3 | SER_A_19_CA | SER_A_19_C | -12.686 | 0.658 | 0.599 | -14.202 | 0.477 | 0.501 | 1.53 | 93.67 | -173.19 |
| 112 | 2jsa-3 | SER_A_19_C | SER_A_19_O | -14.202 | 0.477 | 0.501 | -14.707 | -0.026 | -0.502 | 1.23 | 144.60 | -135.11 |
| 113 | 2jsa-3 | SER_A_19_C | ALA_A_20_N | -14.202 | 0.477 | 0.501 | -14.886 | 0.898 | 1.555 | 1.33 | 37.31 | 148.39 |
| 114 | 2jsa-3 | ALA_A_20_N | ALA_A_20_H | -14.886 | 0.898 | 1.555 | -14.404 | 1.302 | 2.346 | 1.01 | 38.49 | 39.97 |
| 115 | 2jsa-3 | ALA_A_20_N | ALA_A_20_CA | -14.886 | 0.898 | 1.555 | -16.335 | 0.789 | 1.599 | 1.45 | 88.27 | -175.70 |
| 116 | 2jsa-3 | ALA_A_20_CA | ALA_A_20_HA | -16.335 | 0.789 | 1.599 | -16.719 | 1.515 | 2.316 | 1.09 | 48.88 | 117.88 |
| 117 | 2jsa-3 | ALA_A_20_CA | ALA_A_20_C | -16.335 | 0.789 | 1.599 | -16.722 | -0.613 | 2.074 | 1.53 | 71.91 | -105.43 |
| 118 | 2jsa-3 | ALA_A_20_C | ALA_A_20_O | -16.722 | -0.613 | 2.074 | -17.895 | -0.886 | 2.323 | 1.23 | 78.32 | -166.90 |
| 119 | 2jsa-3 | ALA_A_20_C | ASP_A_21_N | -16.722 | -0.613 | 2.074 | -15.713 | -1.465 | 2.186 | 1.33 | 85.15 | -40.18 |
| 120 | 2jsa-3 | ASP_A_21_N | ASP_A_21_H | -15.713 | -1.465 | 2.186 | -14.772 | -1.172 | 1.963 | 1.01 | 102.75 | 17.30 |
| 121 | 2jsa-3 | ASP_A_21_N | ASP_A_21_CA | -15.713 | -1.465 | 2.186 | -15.934 | -2.832 | 2.628 | 1.45 | 72.30 | -99.18 |
| 122 | 2jsa-3 | ASP_A_21_CA | ASP_A_21_HA | -15.934 | -2.832 | 2.628 | -17.005 | -3.004 | 2.733 | 1.09 | 84.47 | -170.88 |
| 123 | 2jsa-3 | ASP_A_21_CA | ASP_A_21_C | -15.934 | -2.832 | 2.628 | -15.238 | -3.048 | 3.972 | 1.53 | 28.47 | -17.24 |
| 124 | 2jsa-3 | ASP_A_21_C | ASP_A_21_O | -15.238 | -3.048 | 3.972 | -15.465 | -4.057 | 4.639 | 1.23 | 57.18 | -102.68 |
| 125 | 2jsa-3 | ASP_A_21_C | LYS_A_22_N | -15.238 | -3.048 | 3.972 | -14.405 | -2.082 | 4.333 | 1.33 | 74.20 | 49.23 |
| 126 | 2jsa-3 | LYS_A_22_N | LYS_A_22_H | -14.405 | -2.082 | 4.333 | -14.267 | -1.279 | 3.736 | 1.01 | 126.23 | 80.25 |
| 127 | 2jsa-3 | LYS_A_22_N | LYS_A_22_CA | -14.405 | -2.082 | 4.333 | -13.675 | -2.154 | 5.588 | 1.45 | 30.31 | -5.63 |
| 128 | 2jsa-3 | LYS_A_22_CA | LYS_A_22_HA | -13.675 | -2.154 | 5.588 | -12.866 | -1.424 | 5.55 | 1.09 | 92.00 | 42.06 |
| 129 | 2jsa-3 | LYS_A_22_CA | LYS_A_22_C | -13.675 | -2.154 | 5.588 | -13.044 | -3.54 | 5.727 | 1.53 | 84.78 | -65.52 |
| 130 | 2jsa-3 | LYS_A_22_C | LYS_A_22_O | -13.044 | -3.54 | 5.727 | -13.365 | -4.282 | 6.654 | 1.23 | 41.09 | -113.39 |
| 131 | 2jsa-3 | LYS_A_22_C | ASP_A_23_N | -13.044 | -3.54 | 5.727 | -12.156 | -3.848 | 4.792 | 1.33 | 134.85 | -19.13 |
| 132 | 2jsa-3 | ASP_A_23_N | ASP_A_23_H | -12.156 | -3.848 | 4.792 | -11.94 | -3.188 | 4.058 | 1.01 | 136.59 | 71.88 |
| 133 | 2jsa-3 | ASP_A_23_N | ASP_A_23_CA | -12.156 | -3.848 | 4.792 | -11.477 | -5.133 | 4.799 | 1.45 | 89.72 | -62.15 |
| 134 | 2jsa-3 | ASP_A_23_CA | ASP_A_23_HA | -11.477 | -5.133 | 4.799 | -11.466 | -5.525 | 5.816 | 1.09 | 21.09 | -88.39 |
| 135 | 2jsa-3 | ASP_A_23_CA | ASP_A_23_C | -11.477 | -5.133 | 4.799 | -10.043 | -4.949 | 4.298 | 1.53 | 109.11 | 7.31 |
| 136 | 2jsa-3 | ASP_A_23_C | ASP_A_23_O | -10.043 | -4.949 | 4.298 | -9.827 | -4.497 | 3.174 | 1.23 | 155.98 | 64.46 |
| 137 | 2jsa-3 | ASP_A_23_C | ALA_A_24_N | -10.043 | -4.949 | 4.298 | -9.099 | -5.308 | 5.156 | 1.33 | 49.65 | -20.82 |
| 138 | 2jsa-3 | ALA_A_24_N | ALA_A_24_H | -9.099 | -5.308 | 5.156 | -9.349 | -5.67 | 6.065 | 1.01 | 25.83 | -124.63 |
| 139 | 2jsa-3 | ALA_A_24_N | ALA_A_24_CA | -9.099 | -5.308 | 5.156 | -7.693 | -5.189 | 4.814 | 1.45 | 103.62 | 4.84 |
| 140 | 2jsa-3 | ALA_A_24_CA | ALA_A_24_HA | -7.693 | -5.189 | 4.814 | -7.535 | -4.238 | 4.306 | 1.09 | 117.79 | 80.57 |
| 141 | 2jsa-3 | ALA_A_24_CA | ALA_A_24_C | -7.693 | -5.189 | 4.814 | -7.307 | -6.319 | 3.856 | 1.53 | 128.74 | -71.14 |
| 142 | 2jsa-3 | ALA_A_24_C | ALA_A_24_O | -7.307 | -6.319 | 3.856 | -6.164 | -6.395 | 3.41 | 1.23 | 111.27 | -3.80 |
| 143 | 2jsa-3 | ALA_A_24_C | ASN_A_25_N | -7.307 | -6.319 | 3.856 | -8.283 | -7.168 | 3.57 | 1.32 | 102.47 | -138.98 |
| 144 | 2jsa-3 | ASN_A_25_N | ASN_A_25_H | -8.283 | -7.168 | 3.57 | -9.2 | -7.043 | 3.974 | 1.01 | 66.42 | 172.24 |
| 145 | 2jsa-3 | ASN_A_25_N | ASN_A_25_CA | -8.283 | -7.168 | 3.57 | -8.06 | -8.29 | 2.675 | 1.45 | 128.04 | -78.76 |
| 146 | 2jsa-3 | ASN_A_25_CA | ASN_A_25_HA | -8.06 | -8.29 | 2.675 | -7.035 | -8.643 | 2.794 | 1.09 | 83.74 | -19.00 |
| 147 | 2jsa-3 | ASN_A_25_CA | ASN_A_25_C | -8.06 | -8.29 | 2.675 | -8.29 | -7.838 | 1.231 | 1.53 | 160.65 | 116.97 |
| 148 | 2jsa-3 | ASN_A_25_C | ASN_A_25_O | -8.29 | -7.838 | 1.231 | -7.481 | -8.127 | 0.35 | 1.23 | 135.72 | -19.66 |
| 149 | 2jsa-3 | ASN_A_25_C | VAL_A_26_N | -8.29 | -7.838 | 1.231 | -9.396 | -7.136 | 1.034 | 1.32 | 98.55 | 147.60 |
| 150 | 2jsa-3 | VAL_A_26_N | VAL_A_26_H | -9.396 | -7.136 | 1.034 | -10.015 | -6.935 | 1.806 | 1.01 | 40.13 | 162.01 |
| 151 | 2jsa-3 | VAL_A_26_N | VAL_A_26_CA | -9.396 | -7.136 | 1.034 | -9.742 | -6.641 | -0.288 | 1.45 | 155.45 | 124.95 |
| 152 | 2jsa-3 | VAL_A_26_CA | VAL_A_26_HA | -9.742 | -6.641 | -0.288 | -9.481 | -7.405 | -1.021 | 1.09 | 132.24 | -71.14 |
| 153 | 2jsa-3 | VAL_A_26_CA | VAL_A_26_C | -9.742 | -6.641 | -0.288 | -8.916 | -5.39 | -0.592 | 1.53 | 101.46 | 56.56 |
| 154 | 2jsa-3 | VAL_A_26_C | VAL_A_26_O | -8.916 | -5.39 | -0.592 | -8.815 | -4.974 | -1.745 | 1.23 | 159.63 | 76.35 |
| 155 | 2jsa-3 | VAL_A_26_C | ILE_A_27_N | -8.916 | -5.39 | -0.592 | -8.347 | -4.826 | 0.462 | 1.32 | 37.24 | 44.75 |
| 156 | 2jsa-3 | ILE_A_27_N | ILE_A_27_H | -8.347 | -4.826 | 0.462 | -8.472 | -5.224 | 1.382 | 1.01 | 24.39 | -107.44 |
| 157 | 2jsa-3 | ILE_A_27_N | ILE_A_27_CA | -8.347 | -4.826 | 0.462 | -7.532 | -3.629 | 0.323 | 1.45 | 95.48 | 55.75 |
| 158 | 2jsa-3 | ILE_A_27_CA | ILE_A_27_HA | -7.532 | -3.629 | 0.323 | -8.011 | -2.977 | -0.407 | 1.09 | 132.06 | 126.30 |
| 159 | 2jsa-3 | ILE_A_27_CA | ILE_A_27_C | -7.532 | -3.629 | 0.323 | -6.158 | -4.016 | -0.23 | 1.53 | 111.18 | -15.73 |
| 160 | 2jsa-3 | ILE_A_27_C | ILE_A_27_O | -6.158 | -4.016 | -0.23 | -5.823 | -3.672 | -1.363 | 1.23 | 157.03 | 45.76 |
| 161 | 2jsa-3 | ILE_A_27_C | LYS_A_28_N | -6.158 | -4.016 | -0.23 | -5.401 | -4.724 | 0.594 | 1.32 | 51.52 | -43.08 |
| 162 | 2jsa-3 | LYS_A_28_N | LYS_A_28_H | -5.401 | -4.724 | 0.594 | -5.742 | -4.971 | 1.512 | 1.01 | 24.64 | -144.08 |
| 163 | 2jsa-3 | LYS_A_28_N | LYS_A_28_CA | -5.401 | -4.724 | 0.594 | -4.073 | -5.16 | 0.203 | 1.45 | 105.63 | -18.18 |
| 164 | 2jsa-3 | LYS_A_28_CA | LYS_A_28_HA | -4.073 | -5.16 | 0.203 | -3.449 | -4.274 | 0.08 | 1.09 | 96.48 | 54.84 |
| 165 | 2jsa-3 | LYS_A_28_CA | LYS_A_28_C | -4.073 | -5.16 | 0.203 | -4.151 | -5.866 | -1.152 | 1.53 | 152.34 | -96.30 |
| 166 | 2jsa-3 | LYS_A_28_C | LYS_A_28_O | -4.151 | -5.866 | -1.152 | -3.228 | -5.774 | -1.959 | 1.23 | 131.02 | 5.69 |
| 167 | 2jsa-3 | LYS_A_28_C | LYS_A_29_N | -4.151 | -5.866 | -1.152 | -5.263 | -6.557 | -1.36 | 1.33 | 99.03 | -148.14 |
| 168 | 2jsa-3 | LYS_A_29_N | LYS_A_29_H | -5.263 | -6.557 | -1.36 | -5.981 | -6.589 | -0.651 | 1.01 | 45.39 | -177.45 |
| 169 | 2jsa-3 | LYS_A_29_N | LYS_A_29_CA | -5.263 | -6.557 | -1.36 | -5.474 | -7.279 | -2.603 | 1.45 | 148.82 | -106.29 |
| 170 | 2jsa-3 | LYS_A_29_CA | LYS_A_29_HA | -5.474 | -7.279 | -2.603 | -4.582 | -7.872 | -2.804 | 1.09 | 100.63 | -33.62 |
| 171 | 2jsa-3 | LYS_A_29_CA | LYS_A_29_C | -5.474 | -7.279 | -2.603 | -5.635 | -6.277 | -3.749 | 1.53 | 138.47 | 99.13 |
| 172 | 2jsa-3 | LYS_A_29_C | LYS_A_29_O | -5.635 | -6.277 | -3.749 | -4.806 | -6.227 | -4.655 | 1.23 | 137.49 | 3.45 |
| 173 | 2jsa-3 | LYS_A_29_C | ASP_A_30_N | -5.635 | -6.277 | -3.749 | -6.707 | -5.504 | -3.67 | 1.32 | 86.58 | 144.21 |
| 174 | 2jsa-3 | ASP_A_30_N | ASP_A_30_H | -6.707 | -5.504 | -3.67 | -7.347 | -5.601 | -2.895 | 1.01 | 39.87 | -171.38 |
| 175 | 2jsa-3 | ASP_A_30_N | ASP_A_30_CA | -6.707 | -5.504 | -3.67 | -6.988 | -4.506 | -4.688 | 1.45 | 134.48 | 105.73 |
| 176 | 2jsa-3 | ASP_A_30_CA | ASP_A_30_HA | -6.988 | -4.506 | -4.688 | -7.411 | -5.0 | -5.563 | 1.09 | 143.38 | -130.57 |
| 177 | 2jsa-3 | ASP_A_30_CA | ASP_A_30_C | -6.988 | -4.506 | -4.688 | -5.689 | -3.794 | -5.07 | 1.53 | 104.46 | 28.73 |
| 178 | 2jsa-3 | ASP_A_30_C | ASP_A_30_O | -5.689 | -3.794 | -5.07 | -5.369 | -3.67 | -6.251 | 1.23 | 163.80 | 21.18 |
| 179 | 2jsa-3 | ASP_A_30_C | PHE_A_31_N | -5.689 | -3.794 | -5.07 | -4.974 | -3.346 | -4.049 | 1.32 | 39.57 | 32.07 |
| 180 | 2jsa-3 | PHE_A_31_N | PHE_A_31_H | -4.974 | -3.346 | -4.049 | -5.297 | -3.485 | -3.102 | 1.01 | 20.37 | -156.72 |
| 181 | 2jsa-3 | PHE_A_31_N | PHE_A_31_CA | -4.974 | -3.346 | -4.049 | -3.717 | -2.65 | -4.263 | 1.45 | 98.47 | 28.97 |
| 182 | 2jsa-3 | PHE_A_31_CA | PHE_A_31_HA | -3.717 | -2.65 | -4.263 | -3.91 | -1.617 | -4.551 | 1.09 | 105.33 | 100.58 |
| 183 | 2jsa-3 | PHE_A_31_CA | PHE_A_31_C | -3.717 | -2.65 | -4.263 | -2.886 | -3.34 | -5.347 | 1.53 | 135.10 | -39.70 |
| 184 | 2jsa-3 | PHE_A_31_C | PHE_A_31_O | -2.886 | -3.34 | -5.347 | -2.531 | -2.721 | -6.349 | 1.23 | 144.54 | 60.17 |
| 185 | 2jsa-3 | PHE_A_31_C | ASP_A_32_N | -2.886 | -3.34 | -5.347 | -2.6 | -4.611 | -5.11 | 1.32 | 79.69 | -77.32 |
| 186 | 2jsa-3 | ASP_A_32_N | ASP_A_32_H | -2.6 | -4.611 | -5.11 | -2.923 | -5.055 | -4.263 | 1.01 | 32.95 | -126.04 |
| 187 | 2jsa-3 | ASP_A_32_N | ASP_A_32_CA | -2.6 | -4.611 | -5.11 | -1.819 | -5.392 | -6.054 | 1.45 | 130.52 | -45.00 |
| 188 | 2jsa-3 | ASP_A_32_CA | ASP_A_32_HA | -1.819 | -5.392 | -6.054 | -0.805 | -4.994 | -6.092 | 1.09 | 92.00 | 21.43 |
| 189 | 2jsa-3 | ASP_A_32_CA | ASP_A_32_C | -1.819 | -5.392 | -6.054 | -2.464 | -5.305 | -7.439 | 1.53 | 154.83 | 172.32 |
| 190 | 2jsa-3 | ASP_A_32_C | ASP_A_32_O | -2.464 | -5.305 | -7.439 | -1.775 | -5.382 | -8.454 | 1.23 | 145.67 | -6.38 |
| 191 | 2jsa-3 | ASP_A_32_C | ALA_A_33_N | -2.464 | -5.305 | -7.439 | -3.779 | -5.145 | -7.433 | 1.32 | 89.74 | 173.06 |
| 192 | 2jsa-3 | ALA_A_33_N | ALA_A_33_H | -3.779 | -5.145 | -7.433 | -4.281 | -5.09 | -6.558 | 1.01 | 29.99 | 173.75 |
| 193 | 2jsa-3 | ALA_A_33_N | ALA_A_33_CA | -3.779 | -5.145 | -7.433 | -4.525 | -5.046 | -8.678 | 1.45 | 148.85 | 172.44 |
| 194 | 2jsa-3 | ALA_A_33_CA | ALA_A_33_HA | -4.525 | -5.046 | -8.678 | -4.32 | -5.93 | -9.281 | 1.09 | 123.60 | -76.94 |
| 195 | 2jsa-3 | ALA_A_33_CA | ALA_A_33_C | -4.525 | -5.046 | -8.678 | -4.057 | -3.814 | -9.452 | 1.53 | 120.43 | 69.20 |
| 196 | 2jsa-3 | ALA_A_33_C | ALA_A_33_O | -4.057 | -3.814 | -9.452 | -3.694 | -3.913 | -10.624 | 1.23 | 162.20 | -15.26 |
| 197 | 2jsa-3 | ALA_A_33_C | GLU_A_34_N | -4.057 | -3.814 | -9.452 | -4.078 | -2.68 | -8.767 | 1.32 | 58.87 | 91.06 |

| | | | | | | | | | | | | |
|---|---|---|---|---|---|---|---|---|---|---|---|---|
| 198 | 2jsa-3 | GLU_A_34_N | GLU_A_34_H | -4.078 | -2.68 | -8.767 | -4.386 | -2.673 | -7.805 | 1.01 | 17.76 | 178.70 |
| 199 | 2jsa-3 | GLU_A_34_N | GLU_A_34_CA | -4.078 | -2.68 | -8.767 | -3.66 | -1.429 | -9.376 | 1.45 | 114.78 | 71.52 |
| 200 | 2jsa-3 | GLU_A_34_CA | GLU_A_34_HA | -3.66 | -1.429 | -9.376 | -4.02 | -1.418 | -10.404 | 1.09 | 160.69 | 178.25 |
| 201 | 2jsa-3 | GLU_A_34_CA | GLU_A_34_C | -3.66 | -1.429 | -9.376 | -2.133 | -1.335 | -9.401 | 1.53 | 90.94 | 3.52 |
| 202 | 2jsa-3 | GLU_A_34_C | GLU_A_34_O | -2.133 | -1.335 | -9.401 | -1.573 | -0.418 | -10.001 | 1.23 | 119.18 | 58.59 |
| 203 | 2jsa-3 | GLU_A_34_C | CYS_A_35_N | -2.133 | -1.335 | -9.401 | -1.502 | -2.296 | -8.742 | 1.33 | 60.18 | -56.71 |
| 204 | 2jsa-3 | CYS_A_35_N | CYS_A_35_H | -1.502 | -2.296 | -8.742 | -2.026 | -3.019 | -8.271 | 1.01 | 62.19 | -125.93 |
| 205 | 2jsa-3 | CYS_A_35_N | CYS_A_35_CA | -1.502 | -2.296 | -8.742 | -0.051 | -2.334 | -8.682 | 1.45 | 87.63 | -1.50 |
| 206 | 2jsa-3 | CYS_A_35_CA | CYS_A_35_HA | -0.051 | -2.334 | -8.682 | 0.314 | -1.314 | -8.556 | 1.09 | 83.37 | 70.31 |
| 207 | 2jsa-3 | CYS_A_35_CA | CYS_A_35_C | -0.051 | -2.334 | -8.682 | 0.472 | -2.876 | -10.013 | 1.53 | 150.50 | -46.02 |
| 208 | 2jsa-3 | CYS_A_35_C | CYS_A_35_O | 0.472 | -2.876 | -10.013 | 1.347 | -2.273 | -10.633 | 1.23 | 120.26 | 34.57 |
| 209 | 2jsa-3 | CYS_A_35_C | LYS_A_36_N | 0.472 | -2.876 | -10.013 | -0.086 | -4.009 | -10.415 | 1.33 | 107.66 | -116.22 |
| 210 | 2jsa-3 | LYS_A_36_N | LYS_A_36_H | -0.086 | -4.009 | -10.415 | -0.801 | -4.45 | -9.854 | 1.01 | 56.26 | -148.33 |
| 211 | 2jsa-3 | LYS_A_36_N | LYS_A_36_CA | -0.086 | -4.009 | -10.415 | 0.313 | -4.64 | -11.661 | 1.45 | 149.07 | -57.69 |
| 212 | 2jsa-3 | LYS_A_36_CA | LYS_A_36_HA | 0.313 | -4.64 | -11.661 | 1.401 | -4.718 | -11.667 | 1.09 | 90.32 | -4.10 |
| 213 | 2jsa-3 | LYS_A_36_CA | LYS_A_36_C | 0.313 | -4.64 | -11.661 | -0.095 | -3.746 | -12.833 | 1.53 | 140.02 | 114.53 |
| 214 | 2jsa-3 | LYS_A_36_C | LYS_A_36_O | -0.095 | -3.746 | -12.833 | 0.531 | -3.779 | -13.891 | 1.23 | 149.35 | -3.02 |
| 215 | 2jsa-3 | LYS_A_36_C | LYS_A_37_N | -0.095 | -3.746 | -12.833 | -1.143 | -2.967 | -12.606 | 1.33 | 80.14 | 143.38 |
| 216 | 2jsa-3 | LYS_A_37_N | LYS_A_37_H | -1.143 | -2.967 | -12.606 | -1.61 | -2.991 | -11.711 | 1.01 | 27.59 | -177.06 |
| 217 | 2jsa-3 | LYS_A_37_N | LYS_A_37_CA | -1.143 | -2.967 | -12.606 | -1.642 | -2.066 | -13.63 | 1.45 | 134.83 | 118.98 |
| 218 | 2jsa-3 | LYS_A_37_CA | LYS_A_37_HA | -1.642 | -2.066 | -13.63 | -1.517 | -2.552 | -14.597 | 1.09 | 152.57 | -75.58 |
| 219 | 2jsa-3 | LYS_A_37_CA | LYS_A_37_C | -1.642 | -2.066 | -13.63 | -0.796 | -0.79 | -13.637 | 1.53 | 90.26 | 56.46 |
| 220 | 2jsa-3 | LYS_A_37_C | LYS_A_37_O | -0.796 | -0.79 | -13.637 | -0.238 | -0.417 | -14.668 | 1.23 | 146.94 | 33.76 |
| 221 | 2jsa-3 | LYS_A_37_C | LEU_A_38_N | -0.796 | -0.79 | -13.637 | -0.728 | -0.157 | -12.475 | 1.32 | 28.72 | 83.87 |
| 222 | 2jsa-3 | LEU_A_38_N | LEU_A_38_H | -0.728 | -0.157 | -12.475 | -1.212 | -0.522 | -11.667 | 1.01 | 36.88 | -142.98 |
| 223 | 2jsa-3 | LEU_A_38_N | LEU_A_38_CA | -0.728 | -0.157 | -12.475 | 0.04 | 1.068 | -12.335 | 1.45 | 84.47 | 57.91 |
| 224 | 2jsa-3 | LEU_A_38_CA | LEU_A_38_HA | 0.04 | 1.068 | -12.335 | -0.3 | 1.769 | -13.097 | 1.09 | 134.36 | 115.87 |
| 225 | 2jsa-3 | LEU_A_38_CA | LEU_A_38_C | 0.04 | 1.068 | -12.335 | 1.517 | 0.772 | -12.602 | 1.53 | 100.05 | -11.33 |
| 226 | 2jsa-3 | LEU_A_38_C | LEU_A_38_O | 1.517 | 0.772 | -12.602 | 2.123 | 1.371 | -13.489 | 1.23 | 136.15 | 44.67 |
| 227 | 2jsa-3 | LEU_A_38_C | PHE_A_39_N | 1.517 | 0.772 | -12.602 | 2.054 | -0.152 | -11.82 | 1.32 | 53.81 | -59.84 |
| 228 | 2jsa-3 | PHE_A_39_N | PHE_A_39_H | 2.054 | -0.152 | -11.82 | 1.495 | -0.605 | -11.111 | 1.01 | 45.42 | -140.98 |
| 229 | 2jsa-3 | PHE_A_39_N | PHE_A_39_CA | 2.054 | -0.152 | -11.82 | 3.449 | -0.535 | -11.96 | 1.45 | 95.53 | -15.35 |
| 230 | 2jsa-3 | PHE_A_39_CA | PHE_A_39_HA | 3.449 | -0.535 | -11.96 | 4.047 | 0.336 | -12.228 | 1.09 | 104.23 | 55.53 |
| 231 | 2jsa-3 | PHE_A_39_CA | PHE_A_39_C | 3.449 | -0.535 | -11.96 | 3.614 | -1.62 | -13.025 | 1.53 | 134.14 | -81.35 |
| 232 | 2jsa-3 | PHE_A_39_C | PHE_A_39_O | 3.614 | -1.62 | -13.025 | 4.52 | -2.448 | -12.938 | 1.23 | 85.95 | -42.42 |
| 233 | 2jsa-3 | PHE_A_39_C | HIS_A_40_N | 3.614 | -1.62 | -13.025 | 2.725 | -1.583 | -14.007 | 1.33 | 137.82 | 177.62 |
| 234 | 2jsa-3 | HIS_A_40_N | HIS_A_40_H | 2.725 | -1.583 | -14.007 | 2.006 | -0.874 | -14.015 | 1.01 | 90.45 | 135.40 |
| 235 | 2jsa-3 | HIS_A_40_N | HIS_A_40_CA | 2.725 | -1.583 | -14.007 | 2.761 | -2.553 | -15.087 | 1.45 | 138.05 | -87.87 |
| 236 | 2jsa-3 | HIS_A_40_CA | HIS_A_40_HA | 2.761 | -2.553 | -15.087 | 2.663 | -3.547 | -14.65 | 1.09 | 66.37 | -95.63 |
| 237 | 2jsa-3 | HIS_A_40_CA | HIS_A_40_C | 2.761 | -2.553 | -15.087 | 4.115 | -2.483 | -15.795 | 1.53 | 117.57 | 2.96 |
| 238 | 2jsa-3 | HIS_A_40_C | HIS_A_40_O | 4.115 | -2.483 | -15.795 | 4.596 | -3.484 | -16.325 | 1.23 | 115.51 | -64.33 |
| 239 | 2jsa-3 | HIS_A_40_C | THR_A_41_N | 4.115 | -2.483 | -15.795 | 4.694 | -1.291 | -15.78 | 1.33 | 89.35 | 64.09 |
| 240 | 2jsa-3 | THR_A_41_N | THR_A_41_H | 4.694 | -1.291 | -15.78 | 4.243 | -0.512 | -15.323 | 1.01 | 63.08 | 120.07 |
| 241 | 2jsa-3 | THR_A_41_N | THR_A_41_CA | 4.694 | -1.291 | -15.78 | 5.983 | -1.078 | -16.415 | 1.45 | 115.92 | 9.38 |
| 242 | 2jsa-3 | THR_A_41_CA | THR_A_41_HA | 5.983 | -1.078 | -16.415 | 6.33 | -2.024 | -16.83 | 1.09 | 112.38 | -69.86 |
| 243 | 2jsa-3 | THR_A_41_CA | THR_A_41_C | 5.983 | -1.078 | -16.415 | 7.013 | -0.609 | -15.384 | 1.53 | 47.67 | 24.48 |
| 244 | 2jsa-3 | THR_A_41_C | THR_A_41_O | 7.013 | -0.609 | -15.384 | 7.431 | 0.548 | -15.4 | 1.23 | 90.75 | 70.14 |
| 245 | 2jsa-3 | THR_A_41_C | ILE_A_42_N | 7.013 | -0.609 | -15.384 | 7.392 | -1.531 | -14.512 | 1.32 | 48.82 | -67.65 |
| 246 | 2jsa-3 | ILE_A_42_N | ILE_A_42_H | 7.392 | -1.531 | -14.512 | 7.007 | -2.463 | -14.558 | 1.01 | 92.61 | -112.45 |
| 247 | 2jsa-3 | ILE_A_42_N | ILE_A_42_CA | 7.392 | -1.531 | -14.512 | 8.364 | -1.228 | -13.476 | 1.45 | 44.50 | 17.31 |
| 248 | 2jsa-3 | ILE_A_42_CA | ILE_A_42_HA | 8.364 | -1.228 | -13.476 | 9.043 | -0.468 | -13.863 | 1.09 | 110.79 | 48.22 |
| 249 | 2jsa-3 | ILE_A_42_CA | ILE_A_42_C | 8.364 | -1.228 | -13.476 | 9.191 | -2.479 | -13.176 | 1.53 | 78.69 | -56.53 |
| 250 | 2jsa-3 | ILE_A_42_C | ILE_A_42_O | 9.191 | -2.479 | -13.176 | 8.667 | -3.592 | -13.187 | 1.23 | 90.51 | -115.21 |
| 251 | 2jsa-3 | ILE_A_42_C | PRO_A_43_N | 9.191 | -2.479 | -13.176 | 10.505 | -2.25 | -12.907 | 1.36 | 78.60 | 9.89 |
| 252 | 2jsa-3 | PRO_A_43_N | PRO_A_43_CA | 10.505 | -2.25 | -12.907 | 11.409 | -3.347 | -12.604 | 1.45 | 77.97 | -50.51 |
| 253 | 2jsa-3 | PRO_A_43_CA | PRO_A_43_HA | 11.409 | -3.347 | -12.604 | 11.253 | -4.165 | -13.307 | 1.09 | 130.17 | -100.80 |
| 254 | 2jsa-3 | PRO_A_43_CA | PRO_A_43_C | 11.409 | -3.347 | -12.604 | 11.175 | -3.874 | -11.187 | 1.53 | 22.14 | -113.94 |
| 255 | 2jsa-3 | PRO_A_43_C | PRO_A_43_O | 11.175 | -3.874 | -11.187 | 11.654 | -3.289 | -10.216 | 1.23 | 37.91 | 50.69 |
| 256 | 2jsa-3 | PRO_A_43_C | PHE_A_44_N | 11.175 | -3.874 | -11.187 | 10.441 | -4.974 | -11.113 | 1.32 | 86.80 | -123.71 |
| 257 | 2jsa-3 | PHE_A_44_N | PHE_A_44_H | 10.441 | -4.974 | -11.113 | 10.081 | -5.402 | -11.953 | 1.01 | 146.34 | -130.07 |
| 258 | 2jsa-3 | PHE_A_44_N | PHE_A_44_CA | 10.441 | -4.974 | -11.113 | 10.138 | -5.587 | -9.83 | 1.45 | 28.06 | -116.30 |
| 259 | 2jsa-3 | PHE_A_44_CA | PHE_A_44_HA | 10.138 | -5.587 | -9.83 | 9.811 | -6.616 | -9.979 | 1.09 | 97.86 | -107.63 |
| 260 | 2jsa-3 | PHE_A_44_CA | PHE_A_44_C | 10.138 | -5.587 | -9.83 | 9.051 | -4.803 | -9.09 | 1.53 | 61.10 | 144.20 |
| 261 | 2jsa-3 | PHE_A_44_C | PHE_A_44_O | 9.051 | -4.803 | -9.09 | 8.672 | -5.162 | -7.977 | 1.23 | 25.13 | -136.55 |
| 262 | 2jsa-3 | PHE_A_44_C | GLY_A_45_N | 9.051 | -4.803 | -9.09 | 8.582 | -3.748 | -9.74 | 1.32 | 119.38 | 113.97 |
| 263 | 2jsa-3 | GLY_A_45_N | GLY_A_45_H | 8.582 | -3.748 | -9.74 | 8.942 | -3.513 | -10.654 | 1.01 | 154.81 | 33.14 |
| 264 | 2jsa-3 | GLY_A_45_N | GLY_A_45_CA | 8.582 | -3.748 | -9.74 | 7.547 | -2.91 | -9.159 | 1.45 | 66.43 | 141.00 |
| 265 | 2jsa-3 | GLY_A_45_CA | GLY_A_45_C | 7.547 | -2.91 | -9.159 | 6.416 | -3.762 | -8.577 | 1.53 | 67.66 | -143.01 |
| 266 | 2jsa-3 | GLY_A_45_C | GLY_A_45_O | 6.416 | -3.762 | -8.577 | 6.128 | -3.687 | -7.383 | 1.23 | 14.00 | 165.40 |
| 267 | 2jsa-3 | GLY_A_45_C | THR_A_46_N | 6.416 | -3.762 | -8.577 | 5.805 | -4.55 | -9.447 | 1.32 | 131.10 | -127.79 |
| 268 | 2jsa-3 | THR_A_46_N | THR_A_46_H | 5.805 | -4.55 | -9.447 | 6.095 | -4.559 | -10.415 | 1.01 | 163.31 | -1.78 |
| 269 | 2jsa-3 | THR_A_46_N | THR_A_46_CA | 5.805 | -4.55 | -9.447 | 4.712 | -5.416 | -9.036 | 1.45 | 73.58 | -141.61 |
| 270 | 2jsa-3 | THR_A_46_CA | THR_A_46_HA | 4.712 | -5.416 | -9.036 | 3.949 | -4.808 | -8.55 | 1.09 | 63.52 | 141.45 |
| 271 | 2jsa-3 | THR_A_46_CA | THR_A_46_C | 4.712 | -5.416 | -9.036 | 5.203 | -6.451 | -8.024 | 1.53 | 48.54 | -64.62 |
| 272 | 2jsa-3 | THR_A_46_C | THR_A_46_O | 5.203 | -6.451 | -8.024 | 4.508 | -6.757 | -7.056 | 1.23 | 38.11 | -156.24 |
| 273 | 2jsa-3 | THR_A_46_C | ARG_A_47_N | 5.203 | -6.451 | -8.024 | 6.399 | -6.962 | -8.281 | 1.33 | 101.18 | -23.13 |
| 274 | 2jsa-3 | ARG_A_47_N | ARG_A_47_H | 6.399 | -6.962 | -8.281 | 6.916 | -6.663 | -9.096 | 1.01 | 143.77 | 30.04 |
| 275 | 2jsa-3 | ARG_A_47_N | ARG_A_47_CA | 6.399 | -6.962 | -8.281 | 6.992 | -7.958 | -7.403 | 1.45 | 52.86 | -59.23 |
| 276 | 2jsa-3 | ARG_A_47_CA | ARG_A_47_HA | 6.992 | -7.958 | -7.403 | 6.386 | -8.863 | -7.45 | 1.09 | 92.47 | -123.81 |
| 277 | 2jsa-3 | ARG_A_47_CA | ARG_A_47_C | 6.992 | -7.958 | -7.403 | 7.015 | -7.448 | -5.962 | 1.53 | 19.51 | 87.42 |
| 278 | 2jsa-3 | ARG_A_47_C | ARG_A_47_O | 7.015 | -7.448 | -5.962 | 6.829 | -8.22 | -5.023 | 1.23 | 40.22 | -103.55 |
| 279 | 2jsa-3 | ARG_A_47_C | GLU_A_48_N | 7.015 | -7.448 | -5.962 | 7.245 | -6.149 | -5.83 | 1.33 | 84.29 | 79.96 |
| 280 | 2jsa-3 | GLU_A_48_N | GLU_A_48_H | 7.245 | -6.149 | -5.83 | 7.388 | -5.573 | -6.647 | 1.01 | 144.00 | 76.06 |
| 281 | 2jsa-3 | GLU_A_48_N | GLU_A_48_CA | 7.245 | -6.149 | -5.83 | 7.296 | -5.527 | -4.519 | 1.45 | 25.46 | 85.31 |
| 282 | 2jsa-3 | GLU_A_48_CA | GLU_A_48_HA | 7.296 | -5.527 | -4.519 | 7.818 | -6.205 | -3.844 | 1.09 | 51.73 | -52.41 |
| 283 | 2jsa-3 | GLU_A_48_CA | GLU_A_48_C | 7.296 | -5.527 | -4.519 | 5.881 | -5.307 | -3.98 | 1.53 | 69.37 | 171.16 |
| 284 | 2jsa-3 | GLU_A_48_C | GLU_A_48_O | 5.881 | -5.307 | -3.98 | 5.645 | -5.426 | -2.778 | 1.23 | 12.40 | -153.24 |
| 285 | 2jsa-3 | GLU_A_48_C | CYS_A_49_N | 5.881 | -5.307 | -3.98 | 4.976 | -4.992 | -4.895 | 1.32 | 133.68 | 160.81 |
| 286 | 2jsa-3 | CYS_A_49_N | CYS_A_49_H | 4.976 | -4.992 | -4.895 | 5.241 | -4.913 | -5.866 | 1.01 | 164.10 | 16.60 |
| 287 | 2jsa-3 | CYS_A_49_N | CYS_A_49_CA | 4.976 | -4.992 | -4.895 | 3.59 | -4.755 | -4.527 | 1.45 | 75.33 | 170.30 |
| 288 | 2jsa-3 | CYS_A_49_CA | CYS_A_49_HA | 3.59 | -4.755 | -4.527 | 3.555 | -3.893 | -3.86 | 1.09 | 52.29 | 92.33 |
| 289 | 2jsa-3 | CYS_A_49_CA | CYS_A_49_C | 3.59 | -4.755 | -4.527 | 3.079 | -5.981 | -3.767 | 1.53 | 60.22 | -112.63 |
| 290 | 2jsa-3 | CYS_A_49_C | CYS_A_49_O | 3.079 | -5.981 | -3.767 | 2.377 | -5.846 | -2.765 | 1.23 | 35.51 | 169.11 |
| 291 | 2jsa-3 | CYS_A_49_C | ASP_A_50_N | 3.079 | -5.981 | -3.767 | 3.452 | -7.148 | -4.27 | 1.32 | 112.32 | -72.27 |
| 292 | 2jsa-3 | ASP_A_50_N | ASP_A_50_H | 3.452 | -7.148 | -4.27 | 4.032 | -7.184 | -5.097 | 1.01 | 144.91 | -3.55 |
| 293 | 2jsa-3 | ASP_A_50_N | ASP_A_50_CA | 3.452 | -7.148 | -4.27 | 3.04 | -8.396 | -3.651 | 1.45 | 64.78 | -108.27 |

| | | | | | | | | | | | |
|---|---|---|---|---|---|---|---|---|---|---|---|
| 294 | 2jsa-3 | ASP_A_50_CA | ASP_A_50_HA | 3.04 | -8.396 | -3.651 | 1.955 | -8.403 | -3.553 | 1.09 | 84.84 | -179.63 |
| 295 | 2jsa-3 | ASP_A_50_CA | ASP_A_50_C | 3.04 | -8.396 | -3.651 | 3.688 | -8.513 | -2.27 | 1.53 | 25.49 | -10.23 |
| 296 | 2jsa-3 | ASP_A_50_C | ASP_A_50_O | 3.688 | -8.513 | -2.27 | 3.063 | -8.991 | -1.325 | 1.23 | 39.78 | -142.59 |
| 297 | 2jsa-3 | ASP_A_50_C | HIS_A_51_N | 3.688 | -8.513 | -2.27 | 4.934 | -8.069 | -2.196 | 1.32 | 86.80 | 19.61 |
| 298 | 2jsa-3 | HIS_A_51_N | HIS_A_51_H | 4.934 | -8.069 | -2.196 | 5.388 | -7.689 | -3.014 | 1.01 | 144.10 | 39.93 |
| 299 | 2jsa-3 | HIS_A_51_N | HIS_A_51_CA | 4.934 | -8.069 | -2.196 | 5.674 | -8.118 | -0.947 | 1.45 | 30.70 | -3.79 |
| 300 | 2jsa-3 | HIS_A_51_CA | HIS_A_51_HA | 5.674 | -8.118 | -0.947 | 5.329 | -8.986 | -0.385 | 1.09 | 58.97 | -111.68 |
| 301 | 2jsa-3 | HIS_A_51_CA | HIS_A_51_C | 5.674 | -8.118 | -0.947 | 5.367 | -6.869 | -0.119 | 1.53 | 57.23 | 103.81 |
| 302 | 2jsa-3 | HIS_A_51_C | HIS_A_51_O | 5.367 | -6.869 | -0.119 | 6.05 | -6.588 | 0.865 | 1.23 | 36.89 | 22.36 |
| 303 | 2jsa-3 | HIS_A_51_C | TYR_A_52_N | 5.367 | -6.869 | -0.119 | 4.338 | -6.152 | -0.548 | 1.33 | 108.88 | 145.13 |
| 304 | 2jsa-3 | TYR_A_52_N | TYR_A_52_H | 4.338 | -6.152 | -0.548 | 3.822 | -6.442 | -1.366 | 1.01 | 144.11 | -150.66 |
| 305 | 2jsa-3 | TYR_A_52_N | TYR_A_52_CA | 4.338 | -6.152 | -0.548 | 3.932 | -4.939 | 0.142 | 1.45 | 61.66 | 108.51 |
| 306 | 2jsa-3 | TYR_A_52_CA | TYR_A_52_HA | 3.932 | -4.939 | 0.142 | 4.686 | -4.651 | 0.875 | 1.09 | 47.76 | 20.91 |
| 307 | 2jsa-3 | TYR_A_52_CA | TYR_A_52_C | 3.932 | -4.939 | 0.142 | 2.581 | -5.129 | 0.834 | 1.53 | 63.10 | -171.99 |
| 308 | 2jsa-3 | TYR_A_52_C | TYR_A_52_O | 2.581 | -5.129 | 0.834 | 2.429 | -4.796 | 2.009 | 1.23 | 17.30 | 114.53 |
| 309 | 2jsa-3 | TYR_A_52_C | VAL_A_53_N | 2.581 | -5.129 | 0.834 | 1.633 | -5.66 | 0.077 | 1.32 | 124.86 | -150.75 |
| 310 | 2jsa-3 | VAL_A_53_N | VAL_A_53_H | 1.633 | -5.66 | 0.077 | 1.829 | -5.907 | -0.882 | 1.01 | 161.80 | -51.57 |
| 311 | 2jsa-3 | VAL_A_53_N | VAL_A_53_CA | 1.633 | -5.66 | 0.077 | 0.3 | -5.898 | 0.602 | 1.45 | 68.81 | -169.88 |
| 312 | 2jsa-3 | VAL_A_53_CA | VAL_A_53_HA | 0.3 | -5.898 | 0.602 | -0.146 | -4.936 | 0.854 | 1.09 | 76.63 | 114.87 |
| 313 | 2jsa-3 | VAL_A_53_CA | VAL_A_53_C | 0.3 | -5.898 | 0.602 | 0.401 | -6.725 | 1.885 | 1.53 | 33.00 | -83.04 |
| 314 | 2jsa-3 | VAL_A_53_C | VAL_A_53_O | 0.401 | -6.725 | 1.885 | -0.528 | -6.746 | 2.69 | 1.23 | 49.10 | -178.71 |
| 315 | 2jsa-3 | VAL_A_53_C | ASN_A_54_N | 0.401 | -6.725 | 1.885 | 1.54 | -7.388 | 2.035 | 1.33 | 83.51 | -30.20 |
| 316 | 2jsa-3 | ASN_A_54_N | ASN_A_54_H | 1.54 | -7.388 | 2.035 | 2.262 | -7.324 | 1.332 | 1.01 | 134.12 | 5.07 |
| 317 | 2jsa-3 | ASN_A_54_N | ASN_A_54_CA | 1.54 | -7.388 | 2.035 | 1.775 | -8.215 | 3.205 | 1.45 | 36.31 | -74.14 |
| 318 | 2jsa-3 | ASN_A_54_CA | ASN_A_54_HA | 1.775 | -8.215 | 3.205 | 1.042 | -7.965 | 3.972 | 1.09 | 45.28 | 161.17 |
| 319 | 2jsa-3 | ASN_A_54_CA | ASN_A_54_C | 1.775 | -8.215 | 3.205 | 3.188 | -7.954 | 3.733 | 1.53 | 69.82 | 10.47 |
| 320 | 2jsa-3 | ASN_A_54_C | ASN_A_54_O | 3.188 | -7.954 | 3.733 | 4.061 | -8.816 | 3.634 | 1.23 | 94.61 | -44.64 |
| 321 | 2jsa-3 | ASN_A_54_C | SER_A_55_N | 3.188 | -7.954 | 3.733 | 3.369 | -6.763 | 4.283 | 1.32 | 65.46 | 81.36 |
| 322 | 2jsa-3 | SER_A_55_N | SER_A_55_H | 3.369 | -6.763 | 4.283 | 2.605 | -6.104 | 4.333 | 1.01 | 87.16 | 139.22 |
| 323 | 2jsa-3 | SER_A_55_N | SER_A_55_CA | 3.369 | -6.763 | 4.283 | 4.661 | -6.377 | 4.826 | 1.45 | 68.07 | 16.63 |
| 324 | 2jsa-3 | SER_A_55_CA | SER_A_55_HA | 4.661 | -6.377 | 4.826 | 4.843 | -6.965 | 5.726 | 1.09 | 34.37 | -72.80 |
| 325 | 2jsa-3 | SER_A_55_CA | SER_A_55_C | 4.661 | -6.377 | 4.826 | 4.649 | -4.895 | 5.204 | 1.53 | 75.69 | 90.46 |
| 326 | 2jsa-3 | SER_A_55_C | SER_A_55_O | 4.649 | -4.895 | 5.204 | 5.299 | -4.492 | 6.167 | 1.23 | 38.46 | 31.80 |
| 327 | 2jsa-3 | SER_A_55_C | LYS_A_56_N | 4.649 | -4.895 | 5.204 | 3.903 | -4.126 | 4.426 | 1.32 | 125.99 | 134.13 |
| 328 | 2jsa-3 | LYS_A_56_N | LYS_A_56_H | 3.903 | -4.126 | 4.426 | 3.395 | -4.527 | 3.651 | 1.01 | 140.13 | -141.71 |
| 329 | 2jsa-3 | LYS_A_56_N | LYS_A_56_CA | 3.903 | -4.126 | 4.426 | 3.798 | -2.696 | 4.667 | 1.45 | 80.46 | 94.20 |
| 330 | 2jsa-3 | LYS_A_56_CA | LYS_A_56_HA | 3.798 | -2.696 | 4.667 | 4.214 | -2.489 | 5.654 | 1.09 | 25.21 | 26.45 |
| 331 | 2jsa-3 | LYS_A_56_CA | LYS_A_56_C | 3.798 | -2.696 | 4.667 | 2.322 | -2.294 | 4.691 | 1.53 | 89.10 | 164.76 |
| 332 | 2jsa-3 | LYS_A_56_C | LYS_A_56_O | 2.322 | -2.294 | 4.691 | 1.998 | -1.112 | 4.783 | 1.23 | 85.71 | 105.33 |
| 333 | 2jsa-3 | LYS_A_56_C | VAL_A_57_N | 2.322 | -2.294 | 4.691 | 1.466 | -3.302 | 4.605 | 1.33 | 93.72 | -130.34 |
| 334 | 2jsa-3 | VAL_A_57_N | VAL_A_57_H | 1.466 | -3.302 | 4.605 | 1.804 | -4.251 | 4.53 | 1.01 | 94.26 | -70.40 |
| 335 | 2jsa-3 | VAL_A_57_N | VAL_A_57_CA | 1.466 | -3.302 | 4.605 | 0.032 | -3.069 | 4.616 | 1.45 | 89.57 | 170.77 |
| 336 | 2jsa-3 | VAL_A_57_CA | VAL_A_57_HA | 0.032 | -3.069 | 4.616 | -0.151 | -2.039 | 4.309 | 1.09 | 106.35 | 100.07 |
| 337 | 2jsa-3 | VAL_A_57_CA | VAL_A_57_C | 0.032 | -3.069 | 4.616 | -0.499 | -3.241 | 6.04 | 1.53 | 21.40 | -162.05 |
| 338 | 2jsa-3 | VAL_A_57_C | VAL_A_57_O | -0.499 | -3.241 | 6.04 | -1.285 | -2.423 | 6.516 | 1.23 | 67.24 | 133.86 |
| 339 | 2jsa-3 | VAL_A_57_C | ASP_A_58_N | -0.499 | -3.241 | 6.04 | -0.047 | -4.309 | 6.681 | 1.33 | 61.07 | -67.06 |
| 340 | 2jsa-3 | ASP_A_58_N | ASP_A_58_H | -0.047 | -4.309 | 6.681 | 0.599 | -4.939 | 6.227 | 1.01 | 116.71 | -44.28 |
| 341 | 2jsa-3 | ASP_A_58_N | ASP_A_58_CA | -0.047 | -4.309 | 6.681 | -0.466 | -4.599 | 8.041 | 1.45 | 20.54 | -145.31 |
| 342 | 2jsa-3 | ASP_A_58_CA | ASP_A_58_HA | -0.466 | -4.599 | 8.041 | -1.516 | -4.891 | 8.035 | 1.09 | 90.32 | -164.46 |
| 343 | 2jsa-3 | ASP_A_58_CA | ASP_A_58_C | -0.466 | -4.599 | 8.041 | -0.279 | -3.35 | 8.905 | 1.53 | 55.62 | 81.48 |
| 344 | 2jsa-3 | ASP_A_58_C | ASP_A_58_O | -0.279 | -3.35 | 8.905 | -1.158 | -2.994 | 9.688 | 1.23 | 50.46 | 157.95 |
| 345 | 2jsa-3 | ASP_A_58_C | PRO_A_59_N | -0.279 | -3.35 | 8.905 | 0.904 | -2.702 | 8.729 | 1.36 | 97.43 | 28.71 |
| 346 | 2jsa-3 | PRO_A_59_N | PRO_A_59_CA | 0.904 | -2.702 | 8.729 | 1.218 | -1.501 | 9.483 | 1.45 | 58.73 | 75.35 |
| 347 | 2jsa-3 | PRO_A_59_CA | PRO_A_59_HA | 1.218 | -1.501 | 9.483 | 0.959 | -1.641 | 10.533 | 1.09 | 15.66 | -151.61 |
| 348 | 2jsa-3 | PRO_A_59_CA | PRO_A_59_C | 1.218 | -1.501 | 9.483 | 0.44 | -0.298 | 8.948 | 1.53 | 110.48 | 122.89 |
| 349 | 2jsa-3 | PRO_A_59_C | PRO_A_59_O | 0.44 | -0.298 | 8.948 | -0.292 | 0.353 | 9.69 | 1.23 | 52.86 | 138.35 |
| 350 | 2jsa-3 | PRO_A_59_C | ILE_A_60_N | 0.44 | -0.298 | 8.948 | 0.625 | -0.038 | 7.662 | 1.32 | 166.06 | 54.57 |
| 351 | 2jsa-3 | ILE_A_60_N | ILE_A_60_H | 0.625 | -0.038 | 7.662 | 1.243 | -0.616 | 7.11 | 1.01 | 123.12 | -43.08 |
| 352 | 2jsa-3 | ILE_A_60_N | ILE_A_60_CA | 0.625 | -0.038 | 7.662 | -0.051 | 1.076 | 7.018 | 1.45 | 116.30 | 121.25 |
| 353 | 2jsa-3 | ILE_A_60_CA | ILE_A_60_HA | -0.051 | 1.076 | 7.018 | 0.401 | 2.0 | 7.38 | 1.09 | 70.61 | 63.93 |
| 354 | 2jsa-3 | ILE_A_60_CA | ILE_A_60_C | -0.051 | 1.076 | 7.018 | -1.523 | 1.084 | 7.436 | 1.53 | 74.15 | 179.69 |
| 355 | 2jsa-3 | ILE_A_60_C | ILE_A_60_O | -1.523 | 1.084 | 7.436 | -2.008 | 2.069 | 7.989 | 1.23 | 63.27 | 116.21 |
| 356 | 2jsa-3 | ILE_A_60_C | ILE_A_61_N | -1.523 | 1.084 | 7.436 | -2.19 | -0.025 | 7.156 | 1.32 | 102.21 | -121.02 |
| 357 | 2jsa-3 | ILE_A_61_N | ILE_A_61_H | -2.19 | -0.025 | 7.156 | -1.725 | -0.797 | 6.701 | 1.01 | 116.79 | -58.94 |
| 358 | 2jsa-3 | ILE_A_61_N | ILE_A_61_CA | -2.19 | -0.025 | 7.156 | -3.597 | -0.158 | 7.496 | 1.45 | 76.47 | -174.60 |
| 359 | 2jsa-3 | ILE_A_61_CA | ILE_A_61_HA | -3.597 | -0.158 | 7.496 | -4.14 | 0.65 | 7.006 | 1.09 | 116.72 | 123.90 |
| 360 | 2jsa-3 | ILE_A_61_CA | ILE_A_61_C | -3.597 | -0.158 | 7.496 | -3.77 | 0.014 | 9.007 | 1.53 | 9.17 | 135.17 |
| 361 | 2jsa-3 | ILE_A_61_C | ILE_A_61_O | -3.77 | 0.014 | 9.007 | -4.853 | 0.364 | 9.475 | 1.23 | 67.65 | 162.09 |
| 362 | 2jsa-3 | ILE_A_61_C | HIS_A_62_N | -3.77 | 0.014 | 9.007 | -2.687 | -0.238 | 9.727 | 1.32 | 57.08 | -13.10 |
| 363 | 2jsa-3 | HIS_A_62_N | HIS_A_62_H | -2.687 | -0.238 | 9.727 | -1.829 | -0.521 | 9.274 | 1.01 | 116.63 | -18.25 |
| 364 | 2jsa-3 | HIS_A_62_N | HIS_A_62_CA | -2.687 | -0.238 | 9.727 | -2.706 | -0.115 | 11.174 | 1.45 | 4.92 | 98.78 |
| 365 | 2jsa-3 | HIS_A_62_CA | HIS_A_62_HA | -2.706 | -0.115 | 11.174 | -3.541 | -0.705 | 11.551 | 1.09 | 69.76 | -144.76 |
| 366 | 2jsa-3 | HIS_A_62_CA | HIS_A_62_C | -2.706 | -0.115 | 11.174 | -2.949 | 1.346 | 11.562 | 1.53 | 75.32 | 99.44 |
| 367 | 2jsa-3 | HIS_A_62_C | HIS_A_62_O | -2.949 | 1.346 | 11.562 | -3.769 | 1.632 | 12.433 | 1.23 | 44.92 | 160.77 |
| 368 | 2jsa-3 | HIS_A_62_C | GLU_A_63_N | -2.949 | 1.346 | 11.562 | -2.221 | 2.23 | 10.897 | 1.32 | 120.14 | 50.53 |
| 369 | 2jsa-3 | GLU_A_63_N | GLU_A_63_H | -2.221 | 2.23 | 10.897 | -1.565 | 1.922 | 10.193 | 1.01 | 134.17 | -25.15 |
| 370 | 2jsa-3 | GLU_A_63_N | GLU_A_63_CA | -2.221 | 2.23 | 10.897 | -2.346 | 3.653 | 11.16 | 1.45 | 79.57 | 95.02 |
| 371 | 2jsa-3 | GLU_A_63_CA | GLU_A_63_HA | -2.346 | 3.653 | 11.16 | -2.131 | 3.821 | 12.216 | 1.09 | 14.49 | 38.00 |
| 372 | 2jsa-3 | GLU_A_63_CA | GLU_A_63_C | -2.346 | 3.653 | 11.16 | -3.776 | 4.12 | 10.88 | 1.53 | 100.54 | 161.91 |
| 373 | 2jsa-3 | GLU_A_63_C | GLU_A_63_O | -3.776 | 4.12 | 10.88 | -4.399 | 4.767 | 11.72 | 1.23 | 46.92 | 133.92 |
| 374 | 2jsa-3 | GLU_A_63_C | LEU_A_64_N | -3.776 | 4.12 | 10.88 | -4.255 | 3.775 | 9.693 | 1.33 | 153.56 | -144.24 |
| 375 | 2jsa-3 | LEU_A_64_N | LEU_A_64_H | -4.255 | 3.775 | 9.693 | -3.686 | 3.243 | 9.05 | 1.01 | 129.54 | -43.08 |
| 376 | 2jsa-3 | LEU_A_64_N | LEU_A_64_CA | -4.255 | 3.775 | 9.693 | -5.599 | 4.151 | 9.291 | 1.45 | 106.07 | 164.37 |
| 377 | 2jsa-3 | LEU_A_64_CA | LEU_A_64_HA | -5.599 | 4.151 | 9.291 | -5.678 | 5.236 | 9.354 | 1.09 | 86.69 | 94.16 |
| 378 | 2jsa-3 | LEU_A_64_CA | LEU_A_64_C | -5.599 | 4.151 | 9.291 | -6.608 | 3.554 | 10.274 | 1.53 | 50.02 | -149.39 |
| 379 | 2jsa-3 | LEU_A_64_C | LEU_A_64_O | -6.608 | 3.554 | 10.274 | -7.609 | 4.188 | 10.604 | 1.23 | 74.44 | 147.65 |
| 380 | 2jsa-3 | LEU_A_64_C | GLU_A_65_N | -6.608 | 3.554 | 10.274 | -6.309 | 2.34 | 10.714 | 1.33 | 70.61 | -76.16 |
| 381 | 2jsa-3 | GLU_A_65_N | GLU_A_65_H | -6.309 | 2.34 | 10.714 | -5.467 | 1.878 | 10.4 | 1.01 | 108.10 | -28.75 |
| 382 | 2jsa-3 | GLU_A_65_N | GLU_A_65_CA | -6.309 | 2.34 | 10.714 | -7.177 | 1.65 | 11.653 | 1.45 | 49.74 | -141.52 |
| 383 | 2jsa-3 | GLU_A_65_CA | GLU_A_65_HA | -7.177 | 1.65 | 11.653 | -8.161 | 1.556 | 11.196 | 1.09 | 114.81 | -174.54 |
| 384 | 2jsa-3 | GLU_A_65_CA | GLU_A_65_C | -7.177 | 1.65 | 11.653 | -7.321 | 2.464 | 12.94 | 1.53 | 32.71 | 100.03 |
| 385 | 2jsa-3 | GLU_A_65_C | GLU_A_65_O | -7.321 | 2.464 | 12.94 | -8.433 | 2.7 | 13.408 | 1.23 | 67.62 | 168.02 |
| 386 | 2jsa-3 | GLU_A_65_C | GLY_A_66_N | -7.321 | 2.464 | 12.94 | -6.179 | 2.871 | 13.475 | 1.33 | 66.19 | 19.62 |
| 387 | 2jsa-3 | GLY_A_66_N | GLY_A_66_H | -6.179 | 2.871 | 13.475 | -5.3 | 2.64 | 13.035 | 1.01 | 115.83 | -14.72 |
| 388 | 2jsa-3 | GLY_A_66_N | GLY_A_66_CA | -6.179 | 2.871 | 13.475 | -6.164 | 3.655 | 14.7 | 1.45 | 32.62 | 88.90 |
| 389 | 2jsa-3 | GLY_A_66_CA | GLY_A_66_C | -6.164 | 3.655 | 14.7 | -6.888 | 4.988 | 14.506 | 1.53 | 97.29 | 118.51 |

| | | | | | | | | | | |
|---|---|---|---|---|---|---|---|---|---|---|
| 390 | 2jsa-3 | GLY_A_66_C | GLY_A_66_O | -6.888 | 4.988 | 14.506 | -7.685 | 5.393 | 15.35 | 1.23 | 46.65 | 153.06 |
| 391 | 2jsa-3 | GLY_A_66_C | GLY_A_67_N | -6.888 | 4.988 | 14.506 | -6.585 | 5.633 | 13.39 | 1.32 | 147.44 | 64.84 |
| 392 | 2jsa-3 | GLY_A_67_N | GLY_A_67_H | -6.585 | 5.633 | 13.39 | -5.919 | 5.24 | 12.741 | 1.01 | 130.01 | -30.54 |
| 393 | 2jsa-3 | GLY_A_67_N | GLY_A_67_CA | -6.585 | 5.633 | 13.39 | -7.198 | 6.913 | 13.074 | 1.45 | 102.55 | 115.59 |
| 394 | 2jsa-3 | GLY_A_67_CA | GLY_A_67_C | -7.198 | 6.913 | 13.074 | -6.157 | 7.906 | 12.554 | 1.53 | 109.87 | 43.65 |
| 395 | 2jsa-3 | GLY_A_67_C | GLY_A_67_O | -6.157 | 7.906 | 12.554 | -6.159 | 9.073 | 12.941 | 1.23 | 71.65 | 90.10 |
| 396 | 2jsa-3 | GLY_A_67_C | THR_A_68_N | -6.157 | 7.906 | 12.554 | -5.291 | 7.405 | 11.686 | 1.32 | 130.94 | -30.05 |
| 397 | 2jsa-3 | THR_A_68_N | THR_A_68_H | -5.291 | 7.405 | 11.686 | -5.348 | 6.434 | 11.413 | 1.01 | 105.68 | -93.36 |
| 398 | 2jsa-3 | THR_A_68_N | THR_A_68_CA | -5.291 | 7.405 | 11.686 | -4.245 | 8.234 | 11.108 | 1.45 | 113.42 | 38.40 |
| 399 | 2jsa-3 | THR_A_68_CA | THR_A_68_HA | -4.245 | 8.234 | 11.108 | -4.284 | 9.22 | 11.571 | 1.09 | 64.86 | 92.27 |
| 400 | 2jsa-3 | THR_A_68_CA | THR_A_68_C | -4.245 | 8.234 | 11.108 | -4.473 | 8.409 | 9.606 | 1.53 | 169.17 | 142.49 |
| 401 | 2jsa-3 | THR_A_68_C | THR_A_68_O | -4.473 | 8.409 | 9.606 | -4.7 | 7.434 | 8.891 | 1.23 | 125.54 | -103.11 |
| 402 | 2jsa-3 | THR_A_68_C | ALA_A_69_N | -4.473 | 8.409 | 9.606 | -4.404 | 9.658 | 9.171 | 1.32 | 109.18 | 86.84 |
| 403 | 2jsa-3 | ALA_A_69_N | ALA_A_69_H | -4.404 | 9.658 | 9.171 | -4.214 | 10.411 | 9.817 | 1.01 | 50.25 | 75.84 |
| 404 | 2jsa-3 | ALA_A_69_N | ALA_A_69_CA | -4.404 | 9.658 | 9.171 | -4.599 | 9.974 | 7.766 | 1.45 | 165.20 | 121.68 |
| 405 | 2jsa-3 | ALA_A_69_CA | ALA_A_69_HA | -4.599 | 9.974 | 7.766 | -5.607 | 9.677 | 7.477 | 1.09 | 105.38 | -163.58 |
| 406 | 2jsa-3 | ALA_A_69_CA | ALA_A_69_C | -4.599 | 9.974 | 7.766 | -3.6 | 9.178 | 6.925 | 1.53 | 123.36 | -38.55 |
| 407 | 2jsa-3 | ALA_A_69_C | ALA_A_69_O | -3.6 | 9.178 | 6.925 | -2.606 | 8.674 | 7.448 | 1.23 | 64.86 | -26.89 |
| 408 | 2jsa-3 | ALA_A_69_C | PRO_A_70_N | -3.6 | 9.178 | 6.925 | -3.904 | 9.087 | 5.603 | 1.36 | 166.50 | -163.34 |
| 409 | 2jsa-3 | PRO_A_70_N | PRO_A_70_CA | -3.904 | 9.087 | 5.603 | -3.044 | 8.361 | 4.684 | 1.45 | 129.23 | -40.17 |
| 410 | 2jsa-3 | PRO_A_70_CA | PRO_A_70_HA | -3.044 | 8.361 | 4.684 | -2.752 | 7.406 | 5.122 | 1.09 | 66.32 | -73.00 |
| 411 | 2jsa-3 | PRO_A_70_CA | PRO_A_70_C | -3.044 | 8.361 | 4.684 | -1.776 | 9.159 | 4.373 | 1.53 | 101.73 | 32.18 |
| 412 | 2jsa-3 | PRO_A_70_C | PRO_A_70_O | -1.776 | 9.159 | 4.373 | -0.803 | 8.611 | 3.857 | 1.23 | 114.80 | -29.39 |
| 413 | 2jsa-3 | PRO_A_70_C | LYS_A_71_N | -1.776 | 9.159 | 4.373 | -1.827 | 10.442 | 4.702 | 1.33 | 75.63 | 92.28 |
| 414 | 2jsa-3 | LYS_A_71_N | LYS_A_71_H | -1.827 | 10.442 | 4.702 | -2.659 | 10.826 | 5.125 | 1.01 | 65.22 | 155.22 |
| 415 | 2jsa-3 | LYS_A_71_N | LYS_A_71_CA | -1.827 | 10.442 | 4.702 | -0.695 | 11.321 | 4.464 | 1.45 | 99.43 | 37.83 |
| 416 | 2jsa-3 | LYS_A_71_CA | LYS_A_71_HA | -0.695 | 11.321 | 4.464 | -0.067 | 10.864 | 3.699 | 1.09 | 134.57 | -36.04 |
| 417 | 2jsa-3 | LYS_A_71_CA | LYS_A_71_C | -0.695 | 11.321 | 4.464 | 0.138 | 11.431 | 5.742 | 1.53 | 33.32 | 7.52 |
| 418 | 2jsa-3 | LYS_A_71_C | LYS_A_71_O | 0.138 | 11.431 | 5.742 | 1.112 | 12.181 | 5.791 | 1.23 | 87.72 | 37.60 |
| 419 | 2jsa-3 | LYS_A_71_C | ASP_A_72_N | 0.138 | 11.431 | 5.742 | -0.277 | 10.674 | 6.747 | 1.32 | 40.66 | -118.73 |
| 420 | 2jsa-3 | ASP_A_72_N | ASP_A_72_H | -0.277 | 10.674 | 6.747 | -1.088 | 10.082 | 6.637 | 1.01 | 96.25 | -143.87 |
| 421 | 2jsa-3 | ASP_A_72_N | ASP_A_72_CA | -0.277 | 10.674 | 6.747 | 0.418 | 10.677 | 8.024 | 1.45 | 28.56 | 0.25 |
| 422 | 2jsa-3 | ASP_A_72_CA | ASP_A_72_HA | 0.418 | 10.677 | 8.024 | 1.113 | 11.517 | 8.05 | 1.09 | 88.63 | 50.40 |
| 423 | 2jsa-3 | ASP_A_72_CA | ASP_A_72_C | 0.418 | 10.677 | 8.024 | 1.184 | 9.362 | 8.185 | 1.53 | 83.96 | -59.78 |
| 424 | 2jsa-3 | ASP_A_72_C | ASP_A_72_O | 1.184 | 9.362 | 8.185 | 2.246 | 9.331 | 8.805 | 1.23 | 59.73 | -1.67 |
| 425 | 2jsa-3 | ASP_A_72_C | VAL_A_73_N | 1.184 | 9.362 | 8.185 | 0.616 | 8.309 | 7.616 | 1.32 | 115.43 | -118.34 |
| 426 | 2jsa-3 | VAL_A_73_N | VAL_A_73_H | 0.616 | 8.309 | 7.616 | -0.258 | 8.409 | 7.121 | 1.01 | 119.37 | 173.47 |
| 427 | 2jsa-3 | VAL_A_73_N | VAL_A_73_CA | 0.616 | 8.309 | 7.616 | 1.232 | 6.995 | 7.69 | 1.45 | 87.08 | -64.88 |
| 428 | 2jsa-3 | VAL_A_73_CA | VAL_A_73_HA | 1.232 | 6.995 | 7.69 | 1.501 | 6.798 | 8.728 | 1.09 | 17.81 | -36.22 |
| 429 | 2jsa-3 | VAL_A_73_CA | VAL_A_73_C | 1.232 | 6.995 | 7.69 | 2.516 | 6.993 | 6.858 | 1.53 | 122.94 | -0.09 |
| 430 | 2jsa-3 | VAL_A_73_C | VAL_A_73_O | 2.516 | 6.993 | 6.858 | 3.556 | 6.522 | 7.317 | 1.23 | 68.10 | -24.37 |
| 431 | 2jsa-3 | VAL_A_73_C | CYS_A_74_N | 2.516 | 6.993 | 6.858 | 2.401 | 7.522 | 5.649 | 1.32 | 155.88 | 102.26 |
| 432 | 2jsa-3 | CYS_A_74_N | CYS_A_74_H | 2.401 | 7.522 | 5.649 | 1.514 | 7.89 | 5.337 | 1.01 | 108.00 | 157.47 |
| 433 | 2jsa-3 | CYS_A_74_N | CYS_A_74_CA | 2.401 | 7.522 | 5.649 | 3.54 | 7.586 | 4.749 | 1.45 | 128.27 | 3.22 |
| 434 | 2jsa-3 | CYS_A_74_CA | CYS_A_74_HA | 3.54 | 7.586 | 4.749 | 4.122 | 6.671 | 4.866 | 1.09 | 83.84 | -57.54 |
| 435 | 2jsa-3 | CYS_A_74_CA | CYS_A_74_C | 3.54 | 7.586 | 4.749 | 4.412 | 8.773 | 5.161 | 1.53 | 74.37 | 53.70 |
| 436 | 2jsa-3 | CYS_A_74_C | CYS_A_74_O | 4.412 | 8.773 | 5.161 | 4.52 | 9.754 | 4.426 | 1.23 | 126.68 | 83.72 |
| 437 | 2jsa-3 | CYS_A_74_C | THR_A_75_N | 4.412 | 8.773 | 5.161 | 5.013 | 8.647 | 6.335 | 1.32 | 27.61 | -11.84 |
| 438 | 2jsa-3 | THR_A_75_N | THR_A_75_H | 5.013 | 8.647 | 6.335 | 4.88 | 7.812 | 6.887 | 1.01 | 56.86 | -99.05 |
| 439 | 2jsa-3 | THR_A_75_N | THR_A_75_CA | 5.013 | 8.647 | 6.335 | 5.873 | 9.697 | 6.854 | 1.45 | 69.07 | 50.68 |
| 440 | 2jsa-3 | THR_A_75_CA | THR_A_75_HA | 5.873 | 9.697 | 6.854 | 6.683 | 9.87 | 6.145 | 1.09 | 130.56 | 12.06 |
| 441 | 2jsa-3 | THR_A_75_CA | THR_A_75_C | 5.873 | 9.697 | 6.854 | 6.496 | 9.268 | 8.184 | 1.53 | 29.63 | -34.55 |
| 442 | 2jsa-3 | THR_A_75_C | THR_A_75_O | 6.496 | 9.268 | 8.184 | 7.718 | 9.215 | 8.311 | 1.23 | 84.07 | -2.48 |
| 443 | 2jsa-3 | THR_A_75_C | LYS_A_76_N | 6.496 | 9.268 | 8.184 | 5.628 | 8.972 | 9.14 | 1.32 | 43.81 | -161.17 |
| 444 | 2jsa-3 | LYS_A_76_N | LYS_A_76_H | 5.628 | 8.972 | 9.14 | 4.636 | 9.038 | 8.964 | 1.01 | 100.04 | 176.19 |
| 445 | 2jsa-3 | LYS_A_76_N | LYS_A_76_CA | 5.628 | 8.972 | 9.14 | 6.079 | 8.549 | 10.455 | 1.45 | 25.18 | -43.17 |
| 446 | 2jsa-3 | LYS_A_76_CA | LYS_A_76_HA | 6.079 | 8.549 | 10.455 | 6.93 | 9.17 | 10.736 | 1.09 | 75.07 | 36.12 |
| 447 | 2jsa-3 | LYS_A_76_CA | LYS_A_76_C | 6.079 | 8.549 | 10.455 | 6.565 | 7.1 | 10.381 | 1.53 | 92.77 | -71.46 |
| 448 | 2jsa-3 | LYS_A_76_C | LYS_A_76_O | 6.565 | 7.1 | 10.381 | 7.406 | 6.682 | 11.175 | 1.23 | 49.79 | -26.43 |
| 449 | 2jsa-3 | LYS_A_76_C | LEU_A_77_N | 6.565 | 7.1 | 10.381 | 6.014 | 6.373 | 9.42 | 1.33 | 136.49 | -127.16 |
| 450 | 2jsa-3 | LEU_A_77_N | LEU_A_77_H | 6.014 | 6.373 | 9.42 | 5.328 | 6.783 | 8.802 | 1.01 | 127.71 | 149.13 |
| 451 | 2jsa-3 | LEU_A_77_N | LEU_A_77_CA | 6.014 | 6.373 | 9.42 | 6.38 | 4.979 | 9.232 | 1.45 | 97.43 | -75.29 |
| 452 | 2jsa-3 | LEU_A_77_CA | LEU_A_77_HA | 6.38 | 4.979 | 9.232 | 6.633 | 4.562 | 10.207 | 1.09 | 26.58 | -58.75 |
| 453 | 2jsa-3 | LEU_A_77_CA | LEU_A_77_C | 6.38 | 4.979 | 9.232 | 7.631 | 4.9 | 8.354 | 1.53 | 125.01 | -3.61 |
| 454 | 2jsa-3 | LEU_A_77_C | LEU_A_77_O | 7.631 | 4.9 | 8.354 | 8.007 | 3.821 | 7.899 | 1.23 | 111.71 | -70.79 |
| 455 | 2jsa-3 | LEU_A_77_C | ASN_A_78_N | 7.631 | 4.9 | 8.354 | 8.242 | 6.057 | 8.142 | 1.33 | 99.20 | 62.16 |
| 456 | 2jsa-3 | ASN_A_78_N | ASN_A_78_H | 8.242 | 6.057 | 8.142 | 7.878 | 6.907 | 8.549 | 1.01 | 66.24 | 113.18 |
| 457 | 2jsa-3 | ASN_A_78_N | ASN_A_78_CA | 8.242 | 6.057 | 8.142 | 9.442 | 6.131 | 7.327 | 1.45 | 124.13 | 3.53 |
| 458 | 2jsa-3 | ASN_A_78_CA | ASN_A_78_HA | 9.442 | 6.131 | 7.327 | 9.647 | 7.175 | 7.088 | 1.09 | 102.66 | 78.89 |
| 459 | 2jsa-3 | ASN_A_78_CA | ASN_A_78_C | 9.442 | 6.131 | 7.327 | 9.229 | 5.336 | 6.038 | 1.53 | 147.44 | -105.00 |
| 460 | 2jsa-3 | ASN_A_78_C | ASN_A_78_O | 9.229 | 5.336 | 6.038 | 10.163 | 4.732 | 5.515 | 1.23 | 115.18 | -32.89 |
| 461 | 2jsa-3 | ASN_A_78_C | GLU_A_79_N | 9.229 | 5.336 | 6.038 | 7.992 | 5.362 | 5.563 | 1.33 | 111.00 | 178.80 |
| 462 | 2jsa-3 | GLU_A_79_N | GLU_A_79_H | 7.992 | 5.362 | 5.563 | 7.273 | 5.881 | 6.046 | 1.01 | 61.42 | 144.18 |
| 463 | 2jsa-3 | GLU_A_79_N | GLU_A_79_CA | 7.992 | 5.362 | 5.563 | 7.642 | 4.651 | 4.344 | 1.45 | 146.97 | -116.21 |
| 464 | 2jsa-3 | GLU_A_79_CA | GLU_A_79_HA | 7.642 | 4.651 | 4.344 | 8.492 | 4.03 | 4.061 | 1.09 | 105.05 | -36.15 |
| 465 | 2jsa-3 | GLU_A_79_CA | GLU_A_79_C | 7.642 | 4.651 | 4.344 | 7.373 | 5.642 | 3.211 | 1.53 | 137.81 | 105.19 |
| 466 | 2jsa-3 | GLU_A_79_C | GLU_A_79_O | 7.373 | 5.642 | 3.211 | 6.839 | 5.268 | 2.168 | 1.23 | 147.99 | -144.99 |
| 467 | 2jsa-3 | GLU_A_79_C | CYS_A_80_N | 7.373 | 5.642 | 3.211 | 7.754 | 6.888 | 3.453 | 1.33 | 79.48 | 73.00 |
| 468 | 2jsa-3 | CYS_A_80_N | CYS_A_80_H | 7.754 | 6.888 | 3.453 | 8.189 | 7.126 | 4.334 | 1.01 | 29.37 | 28.68 |
| 469 | 2jsa-3 | CYS_A_80_N | CYS_A_80_CA | 7.754 | 6.888 | 3.453 | 7.559 | 7.935 | 2.465 | 1.45 | 132.85 | 100.55 |
| 470 | 2jsa-3 | CYS_A_80_CA | CYS_A_80_HA | 7.559 | 7.935 | 2.465 | 8.101 | 7.658 | 1.56 | 1.09 | 146.08 | -27.07 |
| 471 | 2jsa-3 | CYS_A_80_CA | CYS_A_80_C | 7.559 | 7.935 | 2.465 | 8.159 | 9.232 | 3.014 | 1.53 | 68.98 | 65.17 |
| 472 | 2jsa-3 | CYS_A_80_C | CYS_A_80_O | 8.159 | 9.232 | 3.014 | 7.509 | 9.947 | 3.775 | 1.23 | 51.78 | 132.27 |
| 473 | 2jsa-3 | CYS_A_80_C | PRO_A_81_N | 8.159 | 9.232 | 3.014 | 9.426 | 9.5 | 2.598 | 1.36 | 107.81 | 11.94 |
| 474 | 2jsa-3 | PRO_A_81_N | PRO_A_81_CA | 9.426 | 9.5 | 2.598 | 10.12 | 10.697 | 3.04 | 1.45 | 72.28 | 59.90 |
| 475 | 2jsa-3 | PRO_A_81_CA | PRO_A_81_HA | 10.12 | 10.697 | 3.04 | 9.986 | 10.832 | 4.113 | 1.09 | 10.05 | 134.79 |
| 476 | 2jsa-3 | PRO_A_81_CA | PRO_A_81_C | 10.12 | 10.697 | 3.04 | 9.584 | 11.939 | 2.324 | 1.53 | 117.89 | 113.34 |
| 477 | 2jsa-3 | PRO_A_81_C | PRO_A_81_O | 9.584 | 11.939 | 2.324 | 8.715 | 12.635 | 2.848 | 1.23 | 64.80 | 141.31 |
| 1 | 2jsa-4 | ARG_A_1_N | ARG_A_1_H | 16.266 | 6.009 | 2.145 | 16.103 | 6.862 | 1.629 | 1.01 | 120.72 | 100.82 |
| 2 | 2jsa-4 | ARG_A_1_N | ARG_A_1_CA | 16.266 | 6.009 | 2.145 | 15.125 | 5.18 | 2.495 | 1.45 | 76.06 | -144.00 |
| 3 | 2jsa-4 | ARG_A_1_CA | ARG_A_1_HA | 15.125 | 5.18 | 2.495 | 15.256 | 4.201 | 2.033 | 1.09 | 115.07 | -82.38 |
| 4 | 2jsa-4 | ARG_A_1_CA | ARG_A_1_C | 15.125 | 5.18 | 2.495 | 13.831 | 5.809 | 1.973 | 1.53 | 109.94 | 154.08 |
| 5 | 2jsa-4 | ARG_A_1_C | ARG_A_1_O | 13.831 | 5.809 | 1.973 | 13.117 | 6.48 | 2.718 | 1.23 | 52.75 | 136.78 |
| 6 | 2jsa-4 | ARG_A_1_C | SER_A_2_N | 13.831 | 5.809 | 1.973 | 13.569 | 5.572 | 0.696 | 1.32 | 164.54 | -137.87 |
| 7 | 2jsa-4 | SER_A_2_N | SER_A_2_H | 13.569 | 5.572 | 0.696 | 14.201 | 5.011 | 0.143 | 1.01 | 123.20 | -41.59 |
| 8 | 2jsa-4 | SER_A_2_N | SER_A_2_CA | 13.569 | 5.572 | 0.696 | 12.375 | 6.108 | 0.065 | 1.45 | 115.74 | 155.82 |

| | | | | | | | | | | | |
|---|---|---|---|---|---|---|---|---|---|---|---|
| 9 | 2jsa-4 | SER_A_2_CA | SER_A_2_HA | 12.375 | 6.108 | 0.065 | 11.505 | 5.686 | 0.567 | 1.09 | 62.56 | -154.12 |
| 10 | 2jsa-4 | SER_A_2_CA | SER_A_2_C | 12.375 | 6.108 | 0.065 | 12.335 | 5.701 | -1.41 | 1.53 | 164.50 | -95.61 |
| 11 | 2jsa-4 | SER_A_2_C | SER_A_2_O | 12.335 | 5.701 | -1.41 | 13.316 | 5.872 | -2.131 | 1.23 | 125.91 | 9.89 |
| 12 | 2jsa-4 | SER_A_2_C | ALA_A_3_N | 12.335 | 5.701 | -1.41 | 11.19 | 5.171 | -1.813 | 1.32 | 107.71 | -155.16 |
| 13 | 2jsa-4 | ALA_A_3_N | ALA_A_3_H | 11.19 | 5.171 | -1.813 | 10.425 | 5.059 | -1.163 | 1.01 | 49.95 | -171.67 |
| 14 | 2jsa-4 | ALA_A_3_N | ALA_A_3_CA | 11.19 | 5.171 | -1.813 | 11.009 | 4.74 | -3.189 | 1.45 | 161.24 | -112.78 |
| 15 | 2jsa-4 | ALA_A_3_CA | ALA_A_3_HA | 11.009 | 4.74 | -3.189 | 11.197 | 5.586 | -3.85 | 1.09 | 127.33 | 77.47 |
| 16 | 2jsa-4 | ALA_A_3_CA | ALA_A_3_C | 11.009 | 4.74 | -3.189 | 9.563 | 4.281 | -3.389 | 1.53 | 97.51 | -162.39 |
| 17 | 2jsa-4 | ALA_A_3_C | ALA_A_3_O | 9.563 | 4.281 | -3.389 | 8.662 | 4.733 | -2.685 | 1.23 | 55.07 | 153.36 |
| 18 | 2jsa-4 | ALA_A_3_C | LEU_A_4_N | 9.563 | 4.281 | -3.389 | 9.387 | 3.389 | -4.353 | 1.33 | 136.68 | -101.16 |
| 19 | 2jsa-4 | LEU_A_4_N | LEU_A_4_H | 9.387 | 3.389 | -4.353 | 10.175 | 3.062 | -4.893 | 1.01 | 122.33 | -22.54 |
| 20 | 2jsa-4 | LEU_A_4_N | LEU_A_4_CA | 9.387 | 3.389 | -4.353 | 8.066 | 2.864 | -4.654 | 1.45 | 101.96 | -158.33 |
| 21 | 2jsa-4 | LEU_A_4_CA | LEU_A_4_HA | 8.066 | 2.864 | -4.654 | 7.516 | 3.63 | -5.201 | 1.09 | 120.12 | 125.68 |
| 22 | 2jsa-4 | LEU_A_4_CA | LEU_A_4_C | 8.066 | 2.864 | -4.654 | 7.318 | 2.598 | -3.348 | 1.53 | 31.29 | -160.42 |
| 23 | 2jsa-4 | LEU_A_4_C | LEU_A_4_O | 7.318 | 2.598 | -3.348 | 6.115 | 2.84 | -3.257 | 1.23 | 85.76 | 168.63 |
| 24 | 2jsa-4 | LEU_A_4_C | SER_A_5_N | 7.318 | 2.598 | -3.348 | 8.059 | 2.103 | -2.367 | 1.33 | 42.25 | -33.74 |
| 25 | 2jsa-4 | SER_A_5_N | SER_A_5_H | 8.059 | 2.103 | -2.367 | 9.044 | 1.93 | -2.512 | 1.01 | 98.25 | -9.96 |
| 26 | 2jsa-4 | SER_A_5_N | SER_A_5_CA | 8.059 | 2.103 | -2.367 | 7.481 | 1.801 | -1.07 | 1.45 | 26.69 | -152.41 |
| 27 | 2jsa-4 | SER_A_5_CA | SER_A_5_HA | 7.481 | 1.801 | -1.07 | 6.94 | 0.859 | -1.149 | 1.09 | 94.16 | -119.87 |
| 28 | 2jsa-4 | SER_A_5_CA | SER_A_5_C | 7.481 | 1.801 | -1.07 | 6.496 | 2.9 | -0.667 | 1.53 | 74.73 | 131.87 |
| 29 | 2jsa-4 | SER_A_5_C | SER_A_5_O | 6.496 | 2.9 | -0.667 | 5.327 | 2.626 | -0.404 | 1.23 | 77.64 | -166.81 |
| 30 | 2jsa-4 | SER_A_5_C | CYS_A_6_N | 6.496 | 2.9 | -0.667 | 7.006 | 4.123 | -0.632 | 1.33 | 88.49 | 67.36 |
| 31 | 2jsa-4 | CYS_A_6_N | CYS_A_6_H | 7.006 | 4.123 | -0.632 | 7.978 | 4.276 | -0.863 | 1.01 | 103.21 | 8.95 |
| 32 | 2jsa-4 | CYS_A_6_N | CYS_A_6_CA | 7.006 | 4.123 | -0.632 | 6.187 | 5.265 | -0.265 | 1.45 | 75.36 | 125.65 |
| 33 | 2jsa-4 | CYS_A_6_CA | CYS_A_6_HA | 6.187 | 5.265 | -0.265 | 5.887 | 5.152 | 0.777 | 1.09 | 17.10 | -159.36 |
| 34 | 2jsa-4 | CYS_A_6_CA | CYS_A_6_C | 6.187 | 5.265 | -0.265 | 4.93 | 5.255 | -1.138 | 1.53 | 124.78 | -179.54 |
| 35 | 2jsa-4 | CYS_A_6_C | CYS_A_6_O | 4.93 | 5.255 | -1.138 | 3.823 | 5.07 | -0.636 | 1.23 | 65.90 | -170.51 |
| 36 | 2jsa-4 | CYS_A_6_C | GLN_A_7_N | 4.93 | 5.255 | -1.138 | 5.145 | 5.456 | -2.43 | 1.33 | 167.17 | 43.07 |
| 37 | 2jsa-4 | GLN_A_7_N | GLN_A_7_H | 5.145 | 5.456 | -2.43 | 6.084 | 5.602 | -2.773 | 1.01 | 109.85 | 8.84 |
| 38 | 2jsa-4 | GLN_A_7_N | GLN_A_7_CA | 5.145 | 5.456 | -2.43 | 4.043 | 5.473 | -3.378 | 1.45 | 130.70 | 179.12 |
| 39 | 2jsa-4 | GLN_A_7_CA | GLN_A_7_HA | 4.043 | 5.473 | -3.378 | 3.515 | 6.419 | -3.264 | 1.09 | 83.99 | 119.17 |
| 40 | 2jsa-4 | GLN_A_7_CA | GLN_A_7_C | 4.043 | 5.473 | -3.378 | 3.066 | 4.336 | -3.075 | 1.53 | 78.57 | -130.67 |
| 41 | 2jsa-4 | GLN_A_7_C | GLN_A_7_O | 3.066 | 4.336 | -3.075 | 1.873 | 4.572 | -2.885 | 1.23 | 81.12 | 168.81 |
| 42 | 2jsa-4 | GLN_A_7_C | MET_A_8_N | 3.066 | 4.336 | -3.075 | 3.606 | 3.127 | -3.038 | 1.32 | 88.40 | -65.93 |
| 43 | 2jsa-4 | MET_A_8_N | MET_A_8_H | 3.606 | 3.127 | -3.038 | 4.595 | 3.007 | -3.204 | 1.01 | 99.46 | -6.92 |
| 44 | 2jsa-4 | MET_A_8_N | MET_A_8_CA | 3.606 | 3.127 | -3.038 | 2.796 | 1.953 | -2.761 | 1.45 | 79.01 | -124.60 |
| 45 | 2jsa-4 | MET_A_8_CA | MET_A_8_HA | 2.796 | 1.953 | -2.761 | 2.143 | 1.77 | -3.614 | 1.09 | 141.51 | -164.34 |
| 46 | 2jsa-4 | MET_A_8_CA | MET_A_8_C | 2.796 | 1.953 | -2.761 | 1.936 | 2.163 | -1.513 | 1.53 | 35.35 | 166.28 |
| 47 | 2jsa-4 | MET_A_8_C | MET_A_8_O | 1.936 | 2.163 | -1.513 | 0.869 | 1.563 | -1.381 | 1.23 | 83.85 | -150.65 |
| 48 | 2jsa-4 | MET_A_8_C | CYS_A_9_N | 1.936 | 2.163 | -1.513 | 2.43 | 3.015 | -0.628 | 1.32 | 48.06 | 59.89 |
| 49 | 2jsa-4 | CYS_A_9_N | CYS_A_9_H | 2.43 | 3.015 | -0.628 | 3.314 | 3.473 | -0.802 | 1.01 | 99.91 | 27.39 |
| 50 | 2jsa-4 | CYS_A_9_N | CYS_A_9_CA | 2.43 | 3.015 | -0.628 | 1.721 | 3.312 | 0.604 | 1.45 | 31.96 | 157.27 |
| 51 | 2jsa-4 | CYS_A_9_CA | CYS_A_9_HA | 1.721 | 3.312 | 0.604 | 1.461 | 2.369 | 1.086 | 1.09 | 63.77 | -105.41 |
| 52 | 2jsa-4 | CYS_A_9_CA | CYS_A_9_C | 1.721 | 3.312 | 0.604 | 0.429 | 4.05 | 0.249 | 1.53 | 103.42 | 150.26 |
| 53 | 2jsa-4 | CYS_A_9_C | CYS_A_9_O | 0.429 | 4.05 | 0.249 | -0.587 | 3.891 | 0.925 | 1.23 | 56.68 | -171.11 |
| 54 | 2jsa-4 | CYS_A_9_C | GLU_A_10_N | 0.429 | 4.05 | 0.249 | 0.509 | 4.84 | -0.811 | 1.32 | 143.16 | 84.22 |
| 55 | 2jsa-4 | GLU_A_10_N | GLU_A_10_H | 0.509 | 4.84 | -0.811 | 1.378 | 4.921 | -1.32 | 1.01 | 120.25 | 5.33 |
| 56 | 2jsa-4 | GLU_A_10_N | GLU_A_10_CA | 0.509 | 4.84 | -0.811 | -0.641 | 5.603 | -1.265 | 1.45 | 108.21 | 146.44 |
| 57 | 2jsa-4 | GLU_A_10_CA | GLU_A_10_HA | -0.641 | 5.603 | -1.265 | -1.17 | 5.97 | -0.385 | 1.09 | 36.19 | 145.25 |
| 58 | 2jsa-4 | GLU_A_10_CA | GLU_A_10_C | -0.641 | 5.603 | -1.265 | -1.593 | 4.707 | -2.06 | 1.53 | 121.30 | -136.74 |
| 59 | 2jsa-4 | GLU_A_10_C | GLU_A_10_O | -1.593 | 4.707 | -2.06 | -2.704 | 5.117 | -2.395 | 1.23 | 105.80 | 159.74 |
| 60 | 2jsa-4 | GLU_A_10_C | LEU_A_11_N | -1.593 | 4.707 | -2.06 | -1.125 | 3.499 | -2.338 | 1.32 | 102.11 | -68.82 |
| 61 | 2jsa-4 | LEU_A_11_N | LEU_A_11_H | -1.125 | 3.499 | -2.338 | -0.202 | 3.228 | -2.029 | 1.01 | 72.19 | -16.36 |
| 62 | 2jsa-4 | LEU_A_11_N | LEU_A_11_CA | -1.125 | 3.499 | -2.338 | -1.919 | 2.541 | -3.087 | 1.45 | 121.05 | -129.65 |
| 63 | 2jsa-4 | LEU_A_11_CA | LEU_A_11_HA | -1.919 | 2.541 | -3.087 | -2.602 | 3.098 | -3.728 | 1.09 | 126.03 | 140.80 |
| 64 | 2jsa-4 | LEU_A_11_CA | LEU_A_11_C | -1.919 | 2.541 | -3.087 | -2.764 | 1.715 | -2.115 | 1.53 | 50.56 | -135.65 |
| 65 | 2jsa-4 | LEU_A_11_C | LEU_A_11_O | -2.764 | 1.715 | -2.115 | -3.947 | 1.48 | -2.358 | 1.23 | 101.39 | -168.76 |
| 66 | 2jsa-4 | LEU_A_11_C | VAL_A_12_N | -2.764 | 1.715 | -2.115 | -2.123 | 1.295 | -1.033 | 1.33 | 35.31 | -33.23 |
| 67 | 2jsa-4 | VAL_A_12_N | VAL_A_12_H | -2.123 | 1.295 | -1.033 | -1.148 | 1.525 | -0.898 | 1.01 | 82.32 | 13.27 |
| 68 | 2jsa-4 | VAL_A_12_N | VAL_A_12_CA | -2.123 | 1.295 | -1.033 | -2.8 | 0.501 | -0.023 | 1.45 | 45.93 | -130.45 |
| 69 | 2jsa-4 | VAL_A_12_CA | VAL_A_12_HA | -2.8 | 0.501 | -0.023 | -3.308 | -0.325 | -0.519 | 1.09 | 117.09 | -121.59 |
| 70 | 2jsa-4 | VAL_A_12_CA | VAL_A_12_C | -2.8 | 0.501 | -0.023 | -3.858 | 1.361 | 0.672 | 1.53 | 62.99 | 140.89 |
| 71 | 2jsa-4 | VAL_A_12_C | VAL_A_12_O | -3.858 | 1.361 | 0.672 | -5.019 | 0.966 | 0.768 | 1.23 | 85.52 | -161.21 |
| 72 | 2jsa-4 | VAL_A_12_C | VAL_A_13_N | -3.858 | 1.361 | 0.672 | -3.419 | 2.521 | 1.138 | 1.32 | 69.41 | 69.27 |
| 73 | 2jsa-4 | VAL_A_13_N | VAL_A_13_H | -3.419 | 2.521 | 1.138 | -2.45 | 2.782 | 1.022 | 1.01 | 96.59 | 15.07 |
| 74 | 2jsa-4 | VAL_A_13_N | VAL_A_13_CA | -3.419 | 2.521 | 1.138 | -4.314 | 3.439 | 1.822 | 1.45 | 61.92 | 134.27 |
| 75 | 2jsa-4 | VAL_A_13_CA | VAL_A_13_HA | -4.314 | 3.439 | 1.822 | -4.739 | 2.927 | 2.685 | 1.09 | 37.63 | -129.70 |
| 76 | 2jsa-4 | VAL_A_13_CA | VAL_A_13_C | -4.314 | 3.439 | 1.822 | -5.46 | 3.821 | 0.883 | 1.53 | 127.86 | 161.57 |
| 77 | 2jsa-4 | VAL_A_13_C | VAL_A_13_O | -5.46 | 3.821 | 0.883 | -6.629 | 3.616 | 1.21 | 1.23 | 74.60 | -170.05 |
| 78 | 2jsa-4 | VAL_A_13_C | LYS_A_14_N | -5.46 | 3.821 | 0.883 | -5.087 | 4.367 | -0.264 | 1.32 | 150.04 | 55.66 |
| 79 | 2jsa-4 | LYS_A_14_N | LYS_A_14_H | -5.087 | 4.367 | -0.264 | -4.107 | 4.504 | -0.466 | 1.01 | 101.54 | 7.96 |
| 80 | 2jsa-4 | LYS_A_14_N | LYS_A_14_CA | -5.087 | 4.367 | -0.264 | -6.069 | 4.779 | -1.252 | 1.45 | 132.85 | 157.24 |
| 81 | 2jsa-4 | LYS_A_14_CA | LYS_A_14_HA | -6.069 | 4.779 | -1.252 | -6.586 | 5.658 | -0.867 | 1.09 | 69.32 | 120.46 |
| 82 | 2jsa-4 | LYS_A_14_CA | LYS_A_14_C | -6.069 | 4.779 | -1.252 | -7.106 | 3.668 | -1.428 | 1.53 | 96.61 | -133.03 |
| 83 | 2jsa-4 | LYS_A_14_C | LYS_A_14_O | -7.106 | 3.668 | -1.428 | -8.27 | 3.94 | -1.719 | 1.23 | 103.68 | 166.85 |
| 84 | 2jsa-4 | LYS_A_14_C | LYS_A_15_N | -7.106 | 3.668 | -1.428 | -6.648 | 2.439 | -1.242 | 1.32 | 81.93 | -69.56 |
| 85 | 2jsa-4 | LYS_A_15_N | LYS_A_15_H | -6.648 | 2.439 | -1.242 | -5.678 | 2.289 | -1.003 | 1.01 | 76.31 | -8.79 |
| 86 | 2jsa-4 | LYS_A_15_N | LYS_A_15_CA | -6.648 | 2.439 | -1.242 | -7.521 | 1.285 | -1.376 | 1.45 | 95.29 | -127.11 |
| 87 | 2jsa-4 | LYS_A_15_CA | LYS_A_15_HA | -7.521 | 1.285 | -1.376 | -8.242 | 1.498 | -2.165 | 1.09 | 136.38 | 163.54 |
| 88 | 2jsa-4 | LYS_A_15_CA | LYS_A_15_C | -7.521 | 1.285 | -1.376 | -8.301 | 1.088 | -0.075 | 1.53 | 31.73 | -165.83 |
| 89 | 2jsa-4 | LYS_A_15_C | LYS_A_15_O | -8.301 | 1.088 | -0.075 | -9.475 | 0.723 | -0.099 | 1.23 | 91.12 | -162.73 |
| 90 | 2jsa-4 | LYS_A_15_C | TYR_A_16_N | -8.301 | 1.088 | -0.075 | -7.616 | 1.339 | 1.032 | 1.33 | 33.39 | 20.12 |
| 91 | 2jsa-4 | TYR_A_16_N | TYR_A_16_H | -7.616 | 1.339 | 1.032 | -6.652 | 1.636 | 0.977 | 1.01 | 93.12 | 17.12 |
| 92 | 2jsa-4 | TYR_A_16_N | TYR_A_16_CA | -7.616 | 1.339 | 1.032 | -8.229 | 1.192 | 2.341 | 1.45 | 25.71 | -166.51 |
| 93 | 2jsa-4 | TYR_A_16_CA | TYR_A_16_HA | -8.229 | 1.192 | 2.341 | -8.736 | 0.23 | 2.415 | 1.09 | 86.11 | -117.79 |
| 94 | 2jsa-4 | TYR_A_16_CA | TYR_A_16_C | -8.229 | 1.192 | 2.341 | -9.222 | 2.325 | 2.61 | 1.53 | 79.88 | 131.23 |
| 95 | 2jsa-4 | TYR_A_16_C | TYR_A_16_O | -9.222 | 2.325 | 2.61 | -10.425 | 2.091 | 2.71 | 1.23 | 85.34 | -168.99 |
| 96 | 2jsa-4 | TYR_A_16_C | GLU_A_17_N | -9.222 | 2.325 | 2.61 | -8.68 | 3.53 | 2.72 | 1.33 | 85.24 | 65.78 |
| 97 | 2jsa-4 | GLU_A_17_N | GLU_A_17_H | -8.68 | 3.53 | 2.72 | -7.681 | 3.649 | 2.626 | 1.01 | 95.34 | 6.79 |
| 98 | 2jsa-4 | GLU_A_17_N | GLU_A_17_CA | -8.68 | 3.53 | 2.72 | -9.503 | 4.7 | 2.976 | 1.45 | 79.85 | 125.12 |
| 99 | 2jsa-4 | GLU_A_17_CA | GLU_A_17_HA | -9.503 | 4.7 | 2.976 | -9.833 | 4.66 | 4.014 | 1.09 | 17.76 | -173.09 |
| 100 | 2jsa-4 | GLU_A_17_CA | GLU_A_17_C | -9.503 | 4.7 | 2.976 | -10.74 | 4.686 | 2.077 | 1.53 | 126.01 | -179.35 |
| 101 | 2jsa-4 | GLU_A_17_C | GLU_A_17_O | -10.74 | 4.686 | 2.077 | -11.844 | 4.985 | 2.529 | 1.23 | 68.44 | 164.85 |
| 102 | 2jsa-4 | GLU_A_17_C | GLY_A_18_N | -10.74 | 4.686 | 2.077 | -10.515 | 4.334 | 0.819 | 1.33 | 161.63 | -57.41 |
| 103 | 2jsa-4 | GLY_A_18_N | GLY_A_18_H | -10.515 | 4.334 | 0.819 | -9.58 | 4.101 | 0.516 | 1.01 | 107.46 | -13.99 |
| 104 | 2jsa-4 | GLY_A_18_N | GLY_A_18_CA | -10.515 | 4.334 | 0.819 | -11.598 | 4.276 | -0.147 | 1.45 | 131.69 | -176.93 |

| | | | | | | | | | | | | |
|---|---|---|---|---|---|---|---|---|---|---|---|---|
| 105 | 2jsa-4 | GLY_A_18_CA | GLY_A_18_C | -11.598 | 4.276 | -0.147 | -11.824 | 2.843 | -0.633 | 1.53 | 108.52 | -98.96 |
| 106 | 2jsa-4 | GLY_A_18_C | GLY_A_18_O | -11.824 | 2.843 | -0.633 | -11.864 | 2.59 | -1.836 | 1.23 | 167.98 | -98.98 |
| 107 | 2jsa-4 | GLY_A_18_C | SER_A_19_N | -11.824 | 2.843 | -0.633 | -11.966 | 1.942 | 0.328 | 1.32 | 43.51 | -98.96 |
| 108 | 2jsa-4 | SER_A_19_N | SER_A_19_H | -11.966 | 1.942 | 0.328 | -11.922 | 2.221 | 1.298 | 1.01 | 16.23 | 81.04 |
| 109 | 2jsa-4 | SER_A_19_N | SER_A_19_CA | -11.966 | 1.942 | 0.328 | -12.188 | 0.541 | 0.012 | 1.45 | 102.56 | -99.00 |
| 110 | 2jsa-4 | SER_A_19_CA | SER_A_19_HA | -12.188 | 0.541 | 0.012 | -11.7 | 0.324 | -0.938 | 1.09 | 150.66 | -23.97 |
| 111 | 2jsa-4 | SER_A_19_CA | SER_A_19_C | -12.188 | 0.541 | 0.012 | -13.686 | 0.267 | -0.132 | 1.53 | 95.40 | -169.63 |
| 112 | 2jsa-4 | SER_A_19_C | SER_A_19_O | -13.686 | 0.267 | -0.132 | -14.116 | -0.346 | -1.109 | 1.23 | 142.53 | -125.05 |
| 113 | 2jsa-4 | SER_A_19_C | ALA_A_20_N | -13.686 | 0.267 | -0.132 | -14.438 | 0.733 | 0.853 | 1.32 | 41.93 | 148.21 |
| 114 | 2jsa-4 | ALA_A_20_N | ALA_A_20_H | -14.438 | 0.733 | 0.853 | -14.015 | 1.227 | 1.626 | 1.01 | 40.08 | 49.43 |
| 115 | 2jsa-4 | ALA_A_20_N | ALA_A_20_CA | -14.438 | 0.733 | 0.853 | -15.879 | 0.546 | 0.847 | 1.45 | 90.24 | -172.61 |
| 116 | 2jsa-4 | ALA_A_20_CA | ALA_A_20_HA | -15.879 | 0.546 | 0.847 | -16.337 | 1.337 | 1.441 | 1.09 | 56.98 | 120.07 |
| 117 | 2jsa-4 | ALA_A_20_CA | ALA_A_20_C | -15.879 | 0.546 | 0.847 | -16.217 | -0.799 | 1.491 | 1.53 | 65.09 | -104.11 |
| 118 | 2jsa-4 | ALA_A_20_C | ALA_A_20_O | -16.217 | -0.799 | 1.491 | -16.842 | -1.653 | 0.864 | 1.23 | 120.65 | -126.20 |
| 119 | 2jsa-4 | ALA_A_20_C | ASP_A_21_N | -16.217 | -0.799 | 1.491 | -15.788 | -0.948 | 2.736 | 1.33 | 20.04 | -19.15 |
| 120 | 2jsa-4 | ASP_A_21_N | ASP_A_21_H | -15.788 | -0.948 | 2.736 | -15.278 | -0.204 | 3.191 | 1.01 | 63.23 | 55.57 |
| 121 | 2jsa-4 | ASP_A_21_N | ASP_A_21_CA | -15.788 | -0.948 | 2.736 | -16.037 | -2.176 | 3.472 | 1.45 | 59.57 | -101.46 |
| 122 | 2jsa-4 | ASP_A_21_CA | ASP_A_21_HA | -16.037 | -2.176 | 3.472 | -17.069 | -2.177 | 3.823 | 1.09 | 71.22 | -179.94 |
| 123 | 2jsa-4 | ASP_A_21_CA | ASP_A_21_C | -16.037 | -2.176 | 3.472 | -15.082 | -2.253 | 4.664 | 1.53 | 38.79 | -4.61 |
| 124 | 2jsa-4 | ASP_A_21_C | ASP_A_21_O | -15.082 | -2.253 | 4.664 | -15.386 | -2.899 | 5.667 | 1.23 | 35.44 | -115.20 |
| 125 | 2jsa-4 | ASP_A_21_C | LYS_A_22_N | -15.082 | -2.253 | 4.664 | -13.947 | -1.587 | 4.517 | 1.32 | 96.37 | 30.40 |
| 126 | 2jsa-4 | LYS_A_22_N | LYS_A_22_H | -13.947 | -1.587 | 4.517 | -13.763 | -1.078 | 3.665 | 1.01 | 147.57 | 70.13 |
| 127 | 2jsa-4 | LYS_A_22_N | LYS_A_22_CA | -13.947 | -1.587 | 4.517 | -12.946 | -1.572 | 5.57 | 1.45 | 43.55 | 0.86 |
| 128 | 2jsa-4 | LYS_A_22_CA | LYS_A_22_HA | -12.946 | -1.572 | 5.57 | -12.13 | -0.924 | 5.251 | 1.09 | 107.02 | 38.45 |
| 129 | 2jsa-4 | LYS_A_22_CA | LYS_A_22_C | -12.946 | -1.572 | 5.57 | -12.378 | -2.982 | 5.747 | 1.53 | 83.36 | -68.06 |
| 130 | 2jsa-4 | LYS_A_22_C | LYS_A_22_O | -12.378 | -2.982 | 5.747 | -12.278 | -3.478 | 6.868 | 1.23 | 24.29 | -78.60 |
| 131 | 2jsa-4 | LYS_A_22_C | ASP_A_23_N | -12.378 | -2.982 | 5.747 | -12.018 | -3.586 | 4.623 | 1.33 | 147.97 | -59.20 |
| 132 | 2jsa-4 | ASP_A_23_N | ASP_A_23_H | -12.018 | -3.586 | 4.623 | -12.127 | -3.117 | 3.735 | 1.01 | 151.53 | 103.08 |
| 133 | 2jsa-4 | ASP_A_23_N | ASP_A_23_CA | -12.018 | -3.586 | 4.623 | -11.463 | -4.928 | 4.641 | 1.45 | 89.29 | -67.53 |
| 134 | 2jsa-4 | ASP_A_23_CA | ASP_A_23_HA | -11.463 | -4.928 | 4.641 | -11.531 | -5.328 | 5.653 | 1.09 | 21.85 | -99.65 |
| 135 | 2jsa-4 | ASP_A_23_CA | ASP_A_23_C | -11.463 | -4.928 | 4.641 | -9.999 | -4.875 | 4.2 | 1.53 | 106.75 | 2.07 |
| 136 | 2jsa-4 | ASP_A_23_C | ASP_A_23_O | -9.999 | -4.875 | 4.2 | -9.706 | -4.598 | 3.038 | 1.23 | 160.86 | 43.39 |
| 137 | 2jsa-4 | ASP_A_23_C | ALA_A_24_N | -9.999 | -4.875 | 4.2 | -9.118 | -5.145 | 5.153 | 1.33 | 44.04 | -17.04 |
| 138 | 2jsa-4 | ALA_A_24_N | ALA_A_24_H | -9.118 | -5.145 | 5.153 | -9.431 | -5.364 | 6.088 | 1.01 | 22.22 | -145.02 |
| 139 | 2jsa-4 | ALA_A_24_N | ALA_A_24_CA | -9.118 | -5.145 | 5.153 | -7.691 | -5.131 | 4.878 | 1.45 | 100.91 | 0.56 |
| 140 | 2jsa-4 | ALA_A_24_CA | ALA_A_24_HA | -7.691 | -5.131 | 4.878 | -7.434 | -4.185 | 4.401 | 1.09 | 115.95 | 74.80 |
| 141 | 2jsa-4 | ALA_A_24_CA | ALA_A_24_C | -7.691 | -5.131 | 4.878 | -7.352 | -6.268 | 3.912 | 1.53 | 129.15 | -73.40 |
| 142 | 2jsa-4 | ALA_A_24_C | ALA_A_24_O | -7.352 | -6.268 | 3.912 | -6.222 | -6.363 | 3.433 | 1.23 | 112.90 | -4.81 |
| 143 | 2jsa-4 | ALA_A_24_C | ASN_A_25_N | -7.352 | -6.268 | 3.912 | -8.349 | -7.1 | 3.654 | 1.32 | 101.24 | -140.15 |
| 144 | 2jsa-4 | ASN_A_25_N | ASN_A_25_H | -8.349 | -7.1 | 3.654 | -9.252 | -6.959 | 4.083 | 1.01 | 64.85 | 171.13 |
| 145 | 2jsa-4 | ASN_A_25_N | ASN_A_25_CA | -8.349 | -7.1 | 3.654 | -8.17 | -8.227 | 2.755 | 1.45 | 128.23 | -80.98 |
| 146 | 2jsa-4 | ASN_A_25_CA | ASN_A_25_HA | -8.17 | -8.227 | 2.755 | -7.155 | -8.611 | 2.863 | 1.09 | 84.32 | -20.72 |
| 147 | 2jsa-4 | ASN_A_25_CA | ASN_A_25_C | -8.17 | -8.227 | 2.755 | -8.401 | -7.765 | 1.314 | 1.53 | 160.28 | 116.57 |
| 148 | 2jsa-4 | ASN_A_25_C | ASN_A_25_O | -8.401 | -7.765 | 1.314 | -7.533 | -7.93 | 0.458 | 1.23 | 134.09 | -10.76 |
| 149 | 2jsa-4 | ASN_A_25_C | VAL_A_26_N | -8.401 | -7.765 | 1.314 | -9.577 | -7.196 | 1.091 | 1.33 | 99.69 | 154.18 |
| 150 | 2jsa-4 | VAL_A_26_N | VAL_A_26_H | -9.577 | -7.196 | 1.091 | -10.244 | -7.094 | 1.843 | 1.01 | 41.90 | 171.31 |
| 151 | 2jsa-4 | VAL_A_26_N | VAL_A_26_CA | -9.577 | -7.196 | 1.091 | -9.932 | -6.709 | -0.231 | 1.45 | 155.49 | 126.09 |
| 152 | 2jsa-4 | VAL_A_26_CA | VAL_A_26_HA | -9.932 | -6.709 | -0.231 | -9.634 | -7.458 | -0.965 | 1.09 | 132.32 | -68.30 |
| 153 | 2jsa-4 | VAL_A_26_CA | VAL_A_26_C | -9.932 | -6.709 | -0.231 | -9.156 | -5.423 | -0.523 | 1.53 | 101.00 | 58.89 |
| 154 | 2jsa-4 | VAL_A_26_C | VAL_A_26_O | -9.156 | -5.423 | -0.523 | -9.202 | -4.906 | -1.638 | 1.23 | 155.04 | 95.08 |
| 155 | 2jsa-4 | VAL_A_26_C | ILE_A_27_N | -9.156 | -5.423 | -0.523 | -8.461 | -4.945 | 0.499 | 1.33 | 39.53 | 34.52 |
| 156 | 2jsa-4 | ILE_A_27_N | ILE_A_27_H | -8.461 | -4.945 | 0.499 | -8.468 | -5.424 | 1.388 | 1.01 | 28.32 | -90.84 |
| 157 | 2jsa-4 | ILE_A_27_N | ILE_A_27_CA | -8.461 | -4.945 | 0.499 | -7.676 | -3.73 | 0.366 | 1.45 | 95.25 | 57.13 |
| 158 | 2jsa-4 | ILE_A_27_CA | ILE_A_27_HA | -7.676 | -3.73 | 0.366 | -8.162 | -3.092 | -0.372 | 1.09 | 132.62 | 127.30 |
| 159 | 2jsa-4 | ILE_A_27_CA | ILE_A_27_C | -7.676 | -3.73 | 0.366 | -6.286 | -4.082 | -0.168 | 1.53 | 110.43 | -14.21 |
| 160 | 2jsa-4 | ILE_A_27_C | ILE_A_27_O | -6.286 | -4.082 | -0.168 | -5.997 | -3.869 | -1.344 | 1.23 | 163.02 | 36.39 |
| 161 | 2jsa-4 | ILE_A_27_C | LYS_A_28_N | -6.286 | -4.082 | -0.168 | -5.462 | -4.614 | 0.723 | 1.33 | 47.75 | -32.85 |
| 162 | 2jsa-4 | LYS_A_28_N | LYS_A_28_H | -5.462 | -4.614 | 0.723 | -5.767 | -4.76 | 1.674 | 1.01 | 19.57 | -154.42 |
| 163 | 2jsa-4 | LYS_A_28_N | LYS_A_28_CA | -5.462 | -4.614 | 0.723 | -4.109 | -4.997 | 0.357 | 1.45 | 104.59 | -15.81 |
| 164 | 2jsa-4 | LYS_A_28_CA | LYS_A_28_HA | -4.109 | -4.997 | 0.357 | -3.514 | -4.088 | 0.262 | 1.09 | 95.00 | 56.79 |
| 165 | 2jsa-4 | LYS_A_28_CA | LYS_A_28_C | -4.109 | -4.997 | 0.357 | -4.131 | -5.684 | -1.011 | 1.53 | 153.32 | -91.83 |
| 166 | 2jsa-4 | LYS_A_28_C | LYS_A_28_O | -4.131 | -5.684 | -1.011 | -3.233 | -5.48 | -1.826 | 1.23 | 131.51 | 12.80 |
| 167 | 2jsa-4 | LYS_A_28_C | LYS_A_29_N | -4.131 | -5.684 | -1.011 | -5.167 | -6.483 | -1.22 | 1.32 | 99.08 | -142.36 |
| 168 | 2jsa-4 | LYS_A_29_N | LYS_A_29_H | -5.167 | -6.483 | -1.22 | -5.869 | -6.603 | -0.504 | 1.01 | 44.85 | -170.30 |
| 169 | 2jsa-4 | LYS_A_29_N | LYS_A_29_CA | -5.167 | -6.483 | -1.22 | -5.318 | -7.201 | -2.474 | 1.45 | 149.67 | -101.88 |
| 170 | 2jsa-4 | LYS_A_29_CA | LYS_A_29_HA | -5.318 | -7.201 | -2.474 | -4.388 | -7.736 | -2.671 | 1.09 | 100.40 | -29.91 |
| 171 | 2jsa-4 | LYS_A_29_CA | LYS_A_29_C | -5.318 | -7.201 | -2.474 | -5.527 | -6.196 | -3.609 | 1.53 | 137.87 | 101.75 |
| 172 | 2jsa-4 | LYS_A_29_C | LYS_A_29_O | -5.527 | -6.196 | -3.609 | -4.894 | -6.3 | -4.658 | 1.23 | 148.55 | -9.33 |
| 173 | 2jsa-4 | LYS_A_29_C | ASP_A_30_N | -5.527 | -6.196 | -3.609 | -6.417 | -5.247 | -3.36 | 1.32 | 79.17 | 133.16 |
| 174 | 2jsa-4 | ASP_A_30_N | ASP_A_30_H | -6.417 | -5.247 | -3.36 | -6.9 | -5.223 | -2.473 | 1.01 | 28.60 | 177.16 |
| 175 | 2jsa-4 | ASP_A_30_N | ASP_A_30_CA | -6.417 | -5.247 | -3.36 | -6.718 | -4.226 | -4.349 | 1.45 | 132.90 | 106.43 |
| 176 | 2jsa-4 | ASP_A_30_CA | ASP_A_30_HA | -6.718 | -4.226 | -4.349 | -7.25 | -4.683 | -5.183 | 1.09 | 139.94 | -139.34 |
| 177 | 2jsa-4 | ASP_A_30_CA | ASP_A_30_C | -6.718 | -4.226 | -4.349 | -5.412 | -3.602 | -4.844 | 1.53 | 108.88 | 25.54 |
| 178 | 2jsa-4 | ASP_A_30_C | ASP_A_30_O | -5.412 | -3.602 | -4.844 | -5.171 | -3.531 | -6.049 | 1.23 | 168.22 | 16.42 |
| 179 | 2jsa-4 | ASP_A_30_C | PHE_A_31_N | -5.412 | -3.602 | -4.844 | -4.602 | -3.166 | -3.891 | 1.32 | 43.99 | 28.29 |
| 180 | 2jsa-4 | PHE_A_31_N | PHE_A_31_H | -4.602 | -3.166 | -3.891 | -4.864 | -3.257 | -2.92 | 1.01 | 15.94 | -160.85 |
| 181 | 2jsa-4 | PHE_A_31_N | PHE_A_31_CA | -4.602 | -3.166 | -3.891 | -3.326 | -2.551 | -4.215 | 1.45 | 102.88 | 25.73 |
| 182 | 2jsa-4 | PHE_A_31_CA | PHE_A_31_HA | -3.326 | -2.551 | -4.215 | -3.484 | -1.528 | -4.556 | 1.09 | 108.23 | 98.78 |
| 183 | 2jsa-4 | PHE_A_31_CA | PHE_A_31_C | -3.326 | -2.551 | -4.215 | -2.593 | -3.346 | -5.298 | 1.53 | 135.04 | -47.32 |
| 184 | 2jsa-4 | PHE_A_31_C | PHE_A_31_O | -2.593 | -3.346 | -5.298 | -2.175 | -2.785 | -6.309 | 1.23 | 145.32 | 53.31 |
| 185 | 2jsa-4 | PHE_A_31_C | ASP_A_32_N | -2.593 | -3.346 | -5.298 | -2.461 | -4.64 | -5.049 | 1.32 | 79.16 | -84.18 |
| 186 | 2jsa-4 | ASP_A_32_N | ASP_A_32_H | -2.461 | -4.64 | -5.049 | -2.829 | -5.034 | -4.195 | 1.01 | 32.26 | -133.05 |
| 187 | 2jsa-4 | ASP_A_32_N | ASP_A_32_CA | -2.461 | -4.64 | -5.049 | -1.787 | -5.519 | -5.99 | 1.45 | 130.35 | -52.52 |
| 188 | 2jsa-4 | ASP_A_32_CA | ASP_A_32_HA | -1.787 | -5.519 | -5.99 | -0.761 | -5.176 | -6.125 | 1.09 | 97.11 | 18.49 |
| 189 | 2jsa-4 | ASP_A_32_CA | ASP_A_32_C | -1.787 | -5.519 | -5.99 | -2.526 | -5.488 | -7.33 | 1.53 | 151.10 | 177.60 |
| 190 | 2jsa-4 | ASP_A_32_C | ASP_A_32_O | -2.526 | -5.488 | -7.33 | -1.911 | -5.634 | -8.385 | 1.23 | 149.07 | -13.35 |
| 191 | 2jsa-4 | ASP_A_32_C | ALA_A_33_N | -2.526 | -5.488 | -7.33 | -3.834 | -5.297 | -7.242 | 1.32 | 86.19 | 171.69 |
| 192 | 2jsa-4 | ALA_A_33_N | ALA_A_33_H | -3.834 | -5.297 | -7.242 | -4.272 | -5.185 | -6.339 | 1.01 | 26.60 | 165.66 |
| 193 | 2jsa-4 | ALA_A_33_N | ALA_A_33_CA | -3.834 | -5.297 | -7.242 | -4.664 | -5.245 | -8.435 | 1.45 | 145.12 | 176.42 |
| 194 | 2jsa-4 | ALA_A_33_CA | ALA_A_33_HA | -4.664 | -5.245 | -8.435 | -4.508 | -6.157 | -9.01 | 1.09 | 121.86 | -80.29 |
| 195 | 2jsa-4 | ALA_A_33_CA | ALA_A_33_C | -4.664 | -5.245 | -8.435 | -4.24 | -4.053 | -9.294 | 1.53 | 124.18 | 70.42 |
| 196 | 2jsa-4 | ALA_A_33_C | ALA_A_33_O | -4.24 | -4.053 | -9.294 | -3.682 | -4.23 | -10.376 | 1.23 | 151.59 | -17.60 |
| 197 | 2jsa-4 | ALA_A_33_C | GLU_A_34_N | -4.24 | -4.053 | -9.294 | -4.52 | -2.864 | -8.781 | 1.32 | 67.22 | 103.25 |
| 198 | 2jsa-4 | GLU_A_34_N | GLU_A_34_H | -4.52 | -2.864 | -8.781 | -4.98 | -2.793 | -7.885 | 1.01 | 27.45 | 171.23 |
| 199 | 2jsa-4 | GLU_A_34_N | GLU_A_34_CA | -4.52 | -2.864 | -8.781 | -4.174 | -1.643 | -9.488 | 1.45 | 119.12 | 74.18 |
| 200 | 2jsa-4 | GLU_A_34_CA | GLU_A_34_HA | -4.174 | -1.643 | -9.488 | -4.832 | -1.556 | -10.352 | 1.09 | 142.47 | 172.47 |

| | | | | | | | | | | | |
|---|---|---|---|---|---|---|---|---|---|---|---|
| 201 | 2jsa-4 | GLU_A_34_CA | GLU_A_34_C | -4.174 | -1.643 | -9.488 | -2.73 | -1.709 | -9.989 | 1.53 | 109.12 | -2.62 |
| 202 | 2jsa-4 | GLU_A_34_C | GLU_A_34_O | -2.73 | -1.709 | -9.989 | -2.482 | -1.629 | -11.191 | 1.23 | 167.77 | 17.88 |
| 203 | 2jsa-4 | GLU_A_34_C | CYS_A_35_N | -2.73 | -1.709 | -9.989 | -1.815 | -1.856 | -9.042 | 1.33 | 44.38 | -9.13 |
| 204 | 2jsa-4 | CYS_A_35_N | CYS_A_35_H | -1.815 | -1.856 | -9.042 | -2.091 | -1.916 | -8.072 | 1.01 | 16.23 | -167.74 |
| 205 | 2jsa-4 | CYS_A_35_N | CYS_A_35_CA | -1.815 | -1.856 | -9.042 | -0.403 | -1.935 | -9.371 | 1.45 | 103.10 | -3.20 |
| 206 | 2jsa-4 | CYS_A_35_CA | CYS_A_35_HA | -0.403 | -1.935 | -9.371 | -0.065 | -0.944 | -9.675 | 1.09 | 106.19 | 71.17 |
| 207 | 2jsa-4 | CYS_A_35_CA | CYS_A_35_C | -0.403 | -1.935 | -9.371 | -0.238 | -2.89 | -10.555 | 1.53 | 140.70 | -80.20 |
| 208 | 2jsa-4 | CYS_A_35_C | CYS_A_35_O | -0.238 | -2.89 | -10.555 | 0.312 | -2.514 | -11.589 | 1.23 | 147.20 | 34.36 |
| 209 | 2jsa-4 | CYS_A_35_C | LYS_A_36_N | -0.238 | -2.89 | -10.555 | -0.726 | -4.107 | -10.365 | 1.32 | 81.75 | -111.85 |
| 210 | 2jsa-4 | LYS_A_36_N | LYS_A_36_H | -0.726 | -4.107 | -10.365 | -1.166 | -4.347 | -9.488 | 1.01 | 29.75 | -151.39 |
| 211 | 2jsa-4 | LYS_A_36_N | LYS_A_36_CA | -0.726 | -4.107 | -10.365 | -0.64 | -5.119 | -11.405 | 1.45 | 135.68 | -85.14 |
| 212 | 2jsa-4 | LYS_A_36_CA | LYS_A_36_HA | -0.64 | -5.119 | -11.405 | 0.379 | -5.506 | -11.416 | 1.09 | 90.58 | -20.80 |
| 213 | 2jsa-4 | LYS_A_36_CA | LYS_A_36_C | -0.64 | -5.119 | -11.405 | -0.906 | -4.47 | -12.765 | 1.53 | 152.72 | 112.29 |
| 214 | 2jsa-4 | LYS_A_36_C | LYS_A_36_O | -0.906 | -4.47 | -12.765 | -0.389 | -4.924 | -13.785 | 1.23 | 146.00 | -41.29 |
| 215 | 2jsa-4 | LYS_A_36_C | LYS_A_37_N | -0.906 | -4.47 | -12.765 | -1.71 | -3.418 | -12.735 | 1.32 | 88.70 | 127.39 |
| 216 | 2jsa-4 | LYS_A_37_N | LYS_A_37_H | -1.71 | -3.418 | -12.735 | -2.099 | -3.099 | -11.859 | 1.01 | 29.87 | 140.65 |
| 217 | 2jsa-4 | LYS_A_37_N | LYS_A_37_CA | -1.71 | -3.418 | -12.735 | -2.049 | -2.702 | -13.954 | 1.45 | 146.98 | 115.34 |
| 218 | 2jsa-4 | LYS_A_37_CA | LYS_A_37_HA | -2.049 | -2.702 | -13.954 | -1.817 | -3.348 | -14.801 | 1.09 | 140.98 | -70.25 |
| 219 | 2jsa-4 | LYS_A_37_CA | LYS_A_37_C | -2.049 | -2.702 | -13.954 | -1.173 | -1.453 | -14.068 | 1.53 | 94.27 | 54.96 |
| 220 | 2jsa-4 | LYS_A_37_C | LYS_A_37_O | -1.173 | -1.453 | -14.068 | -0.493 | -1.258 | -15.074 | 1.23 | 144.89 | 16.00 |
| 221 | 2jsa-4 | LYS_A_37_C | LEU_A_38_N | -1.173 | -1.453 | -14.068 | -1.217 | -0.64 | -13.022 | 1.33 | 37.90 | 93.10 |
| 222 | 2jsa-4 | LEU_A_38_N | LEU_A_38_H | -1.217 | -0.64 | -13.022 | -1.799 | -0.863 | -12.228 | 1.01 | 38.13 | -159.04 |
| 223 | 2jsa-4 | LEU_A_38_N | LEU_A_38_CA | -1.217 | -0.64 | -13.022 | -0.434 | 0.583 | -12.993 | 1.45 | 88.86 | 57.37 |
| 224 | 2jsa-4 | LEU_A_38_CA | LEU_A_38_HA | -0.434 | 0.583 | -12.993 | -0.846 | 1.265 | -13.737 | 1.09 | 133.04 | 121.14 |
| 225 | 2jsa-4 | LEU_A_38_CA | LEU_A_38_C | -0.434 | 0.583 | -12.993 | 1.008 | 0.272 | -13.396 | 1.53 | 105.28 | -12.17 |
| 226 | 2jsa-4 | LEU_A_38_C | LEU_A_38_O | 1.008 | 0.272 | -13.396 | 1.608 | 1.001 | -14.186 | 1.23 | 129.92 | 50.54 |
| 227 | 2jsa-4 | LEU_A_38_C | PHE_A_39_N | 1.008 | 0.272 | -13.396 | 1.525 | -0.812 | -12.837 | 1.32 | 65.04 | -64.50 |
| 228 | 2jsa-4 | PHE_A_39_N | PHE_A_39_H | 1.525 | -0.812 | -12.837 | 0.974 | -1.365 | -12.196 | 1.01 | 50.61 | -134.90 |
| 229 | 2jsa-4 | PHE_A_39_N | PHE_A_39_CA | 1.525 | -0.812 | -12.837 | 2.886 | -1.23 | -13.128 | 1.45 | 101.55 | -17.07 |
| 230 | 2jsa-4 | PHE_A_39_CA | PHE_A_39_HA | 2.886 | -1.23 | -13.128 | 3.371 | -0.492 | -13.767 | 1.09 | 125.89 | 56.69 |
| 231 | 2jsa-4 | PHE_A_39_CA | PHE_A_39_C | 2.886 | -1.23 | -13.128 | 2.905 | -2.596 | -13.815 | 1.53 | 116.70 | -89.20 |
| 232 | 2jsa-4 | PHE_A_39_C | PHE_A_39_O | 2.905 | -2.596 | -13.815 | 3.681 | -3.473 | -13.437 | 1.23 | 72.11 | -48.50 |
| 233 | 2jsa-4 | PHE_A_39_C | HIS_A_40_N | 2.905 | -2.596 | -13.815 | 2.042 | -2.736 | -14.81 | 1.32 | 138.70 | -170.79 |
| 234 | 2jsa-4 | HIS_A_40_N | HIS_A_40_H | 2.042 | -2.736 | -14.81 | 1.433 | -1.972 | -15.065 | 1.01 | 104.63 | 128.56 |
| 235 | 2jsa-4 | HIS_A_40_N | HIS_A_40_CA | 2.042 | -2.736 | -14.81 | 1.95 | -3.981 | -15.553 | 1.45 | 120.76 | -94.23 |
| 236 | 2jsa-4 | HIS_A_40_CA | HIS_A_40_HA | 1.95 | -3.981 | -15.553 | 1.865 | -4.796 | -14.834 | 1.09 | 48.73 | -95.95 |
| 237 | 2jsa-4 | HIS_A_40_CA | HIS_A_40_C | 1.95 | -3.981 | -15.553 | 3.234 | -4.196 | -16.355 | 1.53 | 121.63 | -9.51 |
| 238 | 2jsa-4 | HIS_A_40_C | HIS_A_40_O | 3.234 | -4.196 | -16.355 | 3.678 | -5.329 | -16.532 | 1.23 | 98.28 | -68.60 |
| 239 | 2jsa-4 | HIS_A_40_C | THR_A_41_N | 3.234 | -4.196 | -16.355 | 3.797 | -3.089 | -16.818 | 1.33 | 110.45 | 63.04 |
| 240 | 2jsa-4 | THR_A_41_N | THR_A_41_H | 3.797 | -3.089 | -16.818 | 3.378 | -2.189 | -16.632 | 1.01 | 79.39 | 114.96 |
| 241 | 2jsa-4 | THR_A_41_N | THR_A_41_CA | 3.797 | -3.089 | -16.818 | 5.022 | -3.143 | -17.597 | 1.45 | 122.43 | -2.52 |
| 242 | 2jsa-4 | THR_A_41_CA | THR_A_41_HA | 5.022 | -3.143 | -17.597 | 4.965 | -3.99 | -18.28 | 1.09 | 128.82 | -93.85 |
| 243 | 2jsa-4 | THR_A_41_CA | THR_A_41_C | 5.022 | -3.143 | -17.597 | 6.229 | -3.357 | -16.681 | 1.53 | 53.23 | -10.05 |
| 244 | 2jsa-4 | THR_A_41_C | THR_A_41_O | 6.229 | -3.357 | -16.681 | 7.098 | -4.176 | -16.975 | 1.23 | 103.83 | -43.30 |
| 245 | 2jsa-4 | THR_A_41_C | ILE_A_42_N | 6.229 | -3.357 | -16.681 | 6.243 | -2.606 | -15.589 | 1.33 | 34.52 | 88.93 |
| 246 | 2jsa-4 | ILE_A_42_N | ILE_A_42_H | 6.243 | -2.606 | -15.589 | 5.494 | -1.953 | -15.411 | 1.01 | 79.84 | 138.92 |
| 247 | 2jsa-4 | ILE_A_42_N | ILE_A_42_CA | 6.243 | -2.606 | -15.589 | 7.327 | -2.704 | -14.628 | 1.45 | 48.56 | -5.17 |
| 248 | 2jsa-4 | ILE_A_42_CA | ILE_A_42_HA | 7.327 | -2.704 | -14.628 | 8.234 | -2.33 | -15.104 | 1.09 | 115.88 | 22.41 |
| 249 | 2jsa-4 | ILE_A_42_CA | ILE_A_42_C | 7.327 | -2.704 | -14.628 | 7.56 | -4.175 | -14.276 | 1.53 | 76.70 | -81.00 |
| 250 | 2jsa-4 | ILE_A_42_C | ILE_A_42_O | 7.56 | -4.175 | -14.276 | 6.61 | -4.948 | -14.168 | 1.23 | 84.96 | -140.87 |
| 251 | 2jsa-4 | ILE_A_42_C | PRO_A_43_N | 7.56 | -4.175 | -14.276 | 8.863 | -4.525 | -14.101 | 1.36 | 82.61 | -15.04 |
| 252 | 2jsa-4 | PRO_A_43_N | PRO_A_43_CA | 8.863 | -4.525 | -14.101 | 9.232 | -5.889 | -13.763 | 1.45 | 76.55 | -74.86 |
| 253 | 2jsa-4 | PRO_A_43_CA | PRO_A_43_HA | 9.232 | -5.889 | -13.763 | 8.681 | -6.591 | -14.388 | 1.09 | 125.01 | -128.13 |
| 254 | 2jsa-4 | PRO_A_43_CA | PRO_A_43_C | 9.232 | -5.889 | -13.763 | 8.919 | -6.195 | -12.298 | 1.53 | 16.64 | -135.65 |
| 255 | 2jsa-4 | PRO_A_43_C | PRO_A_43_O | 8.919 | -6.195 | -12.298 | 7.793 | -6.564 | -11.964 | 1.23 | 74.26 | -161.86 |
| 256 | 2jsa-4 | PRO_A_43_C | PHE_A_44_N | 8.919 | -6.195 | -12.298 | 9.932 | -6.03 | -11.461 | 1.32 | 50.80 | 9.25 |
| 257 | 2jsa-4 | PHE_A_44_N | PHE_A_44_H | 9.932 | -6.03 | -11.461 | 10.831 | -5.724 | -11.805 | 1.01 | 109.91 | 18.80 |
| 258 | 2jsa-4 | PHE_A_44_N | PHE_A_44_CA | 9.932 | -6.03 | -11.461 | 9.778 | -6.283 | -10.038 | 1.45 | 11.76 | -121.33 |
| 259 | 2jsa-4 | PHE_A_44_CA | PHE_A_44_HA | 9.778 | -6.283 | -10.038 | 9.679 | -7.354 | -9.861 | 1.09 | 80.65 | -95.28 |
| 260 | 2jsa-4 | PHE_A_44_CA | PHE_A_44_C | 9.778 | -6.283 | -10.038 | 8.554 | -5.554 | -9.481 | 1.53 | 68.65 | 149.22 |
| 261 | 2jsa-4 | PHE_A_44_C | PHE_A_44_O | 8.554 | -5.554 | -9.481 | 7.974 | -5.981 | -8.483 | 1.23 | 35.82 | -143.64 |
| 262 | 2jsa-4 | PHE_A_44_C | GLY_A_45_N | 8.554 | -5.554 | -9.481 | 8.196 | -4.468 | -10.15 | 1.32 | 120.33 | 108.24 |
| 263 | 2jsa-4 | GLY_A_45_N | GLY_A_45_H | 8.196 | -4.468 | -10.15 | 8.719 | -4.177 | -10.964 | 1.01 | 143.67 | 29.09 |
| 264 | 2jsa-4 | GLY_A_45_N | GLY_A_45_CA | 8.196 | -4.468 | -10.15 | 7.052 | -3.675 | -9.733 | 1.45 | 73.32 | 145.27 |
| 265 | 2jsa-4 | GLY_A_45_CA | GLY_A_45_C | 7.052 | -3.675 | -9.733 | 5.947 | -4.567 | -9.162 | 1.53 | 68.10 | -141.09 |
| 266 | 2jsa-4 | GLY_A_45_C | GLY_A_45_O | 5.947 | -4.567 | -9.162 | 5.585 | -4.441 | -7.994 | 1.23 | 18.17 | 160.81 |
| 267 | 2jsa-4 | GLY_A_45_C | THR_A_46_N | 5.947 | -4.567 | -9.162 | 5.442 | -5.448 | -10.013 | 1.32 | 129.96 | -119.82 |
| 268 | 2jsa-4 | THR_A_46_N | THR_A_46_H | 5.442 | -5.448 | -10.013 | 5.79 | -5.496 | -10.96 | 1.01 | 159.65 | -7.85 |
| 269 | 2jsa-4 | THR_A_46_N | THR_A_46_CA | 5.442 | -5.448 | -10.013 | 4.386 | -6.359 | -9.609 | 1.45 | 73.84 | -139.22 |
| 270 | 2jsa-4 | THR_A_46_CA | THR_A_46_HA | 4.386 | -6.359 | -9.609 | 3.527 | -5.774 | -9.279 | 1.09 | 72.38 | 145.74 |
| 271 | 2jsa-4 | THR_A_46_CA | THR_A_46_C | 4.386 | -6.359 | -9.609 | 4.848 | -7.222 | -8.433 | 1.53 | 39.77 | -61.84 |
| 272 | 2jsa-4 | THR_A_46_C | THR_A_46_O | 4.848 | -7.222 | -8.433 | 4.072 | -7.502 | -7.52 | 1.23 | 42.10 | -160.16 |
| 273 | 2jsa-4 | THR_A_46_C | ARG_A_47_N | 4.848 | -7.222 | -8.433 | 6.11 | -7.621 | -8.494 | 1.32 | 92.64 | -17.55 |
| 274 | 2jsa-4 | ARG_A_47_N | ARG_A_47_H | 6.11 | -7.621 | -8.494 | 6.691 | -7.354 | -9.276 | 1.01 | 140.73 | 24.68 |
| 275 | 2jsa-4 | ARG_A_47_N | ARG_A_47_CA | 6.11 | -7.621 | -8.494 | 6.686 | -8.447 | -7.446 | 1.45 | 43.86 | -55.11 |
| 276 | 2jsa-4 | ARG_A_47_CA | ARG_A_47_HA | 6.686 | -8.447 | -7.446 | 6.061 | -9.331 | -7.322 | 1.09 | 83.47 | -125.26 |
| 277 | 2jsa-4 | ARG_A_47_CA | ARG_A_47_C | 6.686 | -8.447 | -7.446 | 6.726 | -7.676 | -6.125 | 1.53 | 30.30 | 87.03 |
| 278 | 2jsa-4 | ARG_A_47_C | ARG_A_47_O | 6.726 | -7.676 | -6.125 | 6.142 | -8.108 | -5.132 | 1.23 | 36.19 | -143.51 |
| 279 | 2jsa-4 | ARG_A_47_C | GLU_A_48_N | 6.726 | -7.676 | -6.125 | 7.421 | -6.548 | -6.154 | 1.33 | 91.25 | 58.36 |
| 280 | 2jsa-4 | GLU_A_48_N | GLU_A_48_H | 7.421 | -6.548 | -6.154 | 7.876 | -6.253 | -7.006 | 1.01 | 147.52 | 32.96 |
| 281 | 2jsa-4 | GLU_A_48_N | GLU_A_48_CA | 7.421 | -6.548 | -6.154 | 7.544 | -5.714 | -4.971 | 1.45 | 35.47 | 81.61 |
| 282 | 2jsa-4 | GLU_A_48_CA | GLU_A_48_HA | 7.544 | -5.714 | -4.971 | 8.066 | -6.289 | -4.205 | 1.09 | 45.39 | -47.77 |
| 283 | 2jsa-4 | GLU_A_48_CA | GLU_A_48_C | 7.544 | -5.714 | -4.971 | 6.16 | -5.345 | -4.435 | 1.53 | 69.48 | 165.07 |
| 284 | 2jsa-4 | GLU_A_48_C | GLU_A_48_O | 6.16 | -5.345 | -4.435 | 5.948 | -5.315 | -3.224 | 1.23 | 10.03 | 171.95 |
| 285 | 2jsa-4 | GLU_A_48_C | CYS_A_49_N | 6.16 | -5.345 | -4.435 | 5.254 | -5.073 | -5.363 | 1.33 | 134.45 | 163.29 |
| 286 | 2jsa-4 | CYS_A_49_N | CYS_A_49_H | 5.254 | -5.073 | -5.363 | 5.499 | -5.118 | -6.342 | 1.01 | 165.72 | -10.41 |
| 287 | 2jsa-4 | CYS_A_49_N | CYS_A_49_CA | 5.254 | -5.073 | -5.363 | 3.896 | -4.707 | -4.999 | 1.45 | 75.49 | 164.92 |
| 288 | 2jsa-4 | CYS_A_49_CA | CYS_A_49_HA | 3.896 | -4.707 | -4.999 | 3.942 | -3.863 | -4.31 | 1.09 | 50.82 | 86.88 |
| 289 | 2jsa-4 | CYS_A_49_CA | CYS_A_49_C | 3.896 | -4.707 | -4.999 | 3.257 | -5.894 | -4.275 | 1.53 | 61.76 | -118.29 |
| 290 | 2jsa-4 | CYS_A_49_C | CYS_A_49_O | 3.257 | -5.894 | -4.275 | 2.57 | -5.716 | -3.27 | 1.23 | 35.23 | 165.47 |
| 291 | 2jsa-4 | CYS_A_49_C | ASP_A_50_N | 3.257 | -5.894 | -4.275 | 3.505 | -7.079 | -4.814 | 1.33 | 114.00 | -78.18 |
| 292 | 2jsa-4 | ASP_A_50_N | ASP_A_50_H | 3.505 | -7.079 | -4.814 | 4.078 | -7.151 | -5.643 | 1.01 | 145.14 | -7.16 |
| 293 | 2jsa-4 | ASP_A_50_N | ASP_A_50_CA | 3.505 | -7.079 | -4.814 | 2.962 | -8.294 | -4.231 | 1.45 | 66.34 | -114.08 |
| 294 | 2jsa-4 | ASP_A_50_CA | ASP_A_50_HA | 2.962 | -8.294 | -4.231 | 1.902 | -8.148 | -4.027 | 1.09 | 79.21 | 172.16 |
| 295 | 2jsa-4 | ASP_A_50_CA | ASP_A_50_C | 2.962 | -8.294 | -4.231 | 3.704 | -8.607 | -2.931 | 1.53 | 31.78 | -22.87 |
| 296 | 2jsa-4 | ASP_A_50_C | ASP_A_50_O | 3.704 | -8.607 | -2.931 | 3.344 | -9.541 | -2.216 | 1.23 | 54.46 | -111.08 |

| | | | | | | | | | | | |
|---|---|---|---|---|---|---|---|---|---|---|---|
| 297 | 2jsa-4 | ASP_A_50_C | HIS_A_51_N | 3.704 | -8.607 | -2.931 | 4.726 | -7.808 | -2.664 | 1.32 | 78.37 | 38.02 |
| 298 | 2jsa-4 | HIS_A_51_N | HIS_A_51_H | 4.726 | -7.808 | -2.664 | 4.966 | -7.059 | -3.298 | 1.01 | 128.87 | 72.23 |
| 299 | 2jsa-4 | HIS_A_51_N | HIS_A_51_CA | 4.726 | -7.808 | -2.664 | 5.522 | -7.988 | -1.461 | 1.45 | 34.15 | -12.74 |
| 300 | 2jsa-4 | HIS_A_51_CA | HIS_A_51_HA | 5.522 | -7.988 | -1.461 | 5.165 | -8.883 | -0.952 | 1.09 | 62.16 | -111.75 |
| 301 | 2jsa-4 | HIS_A_51_CA | HIS_A_51_C | 5.522 | -7.988 | -1.461 | 5.311 | -6.799 | -0.522 | 1.53 | 52.13 | 100.06 |
| 302 | 2jsa-4 | HIS_A_51_C | HIS_A_51_O | 5.311 | -6.799 | -0.522 | 5.92 | -6.728 | 0.544 | 1.23 | 29.91 | 6.65 |
| 303 | 2jsa-4 | HIS_A_51_C | TYR_A_52_N | 5.311 | -6.799 | -0.522 | 4.447 | -5.892 | -0.954 | 1.33 | 109.03 | 133.61 |
| 304 | 2jsa-4 | TYR_A_52_N | TYR_A_52_H | 4.447 | -5.892 | -0.954 | 3.985 | -6.013 | -1.843 | 1.01 | 151.75 | -165.32 |
| 305 | 2jsa-4 | TYR_A_52_N | TYR_A_52_CA | 4.447 | -5.892 | -0.954 | 4.148 | -4.709 | -0.166 | 1.45 | 57.15 | 104.18 |
| 306 | 2jsa-4 | TYR_A_52_CA | TYR_A_52_HA | 4.148 | -4.709 | -0.166 | 4.944 | -4.525 | 0.556 | 1.09 | 48.53 | 13.02 |
| 307 | 2jsa-4 | TYR_A_52_CA | TYR_A_52_C | 4.148 | -4.709 | -0.166 | 2.809 | -4.858 | 0.562 | 1.53 | 61.62 | -173.65 |
| 308 | 2jsa-4 | TYR_A_52_C | TYR_A_52_O | 2.809 | -4.858 | 0.562 | 2.682 | -4.466 | 1.72 | 1.23 | 19.59 | 107.95 |
| 309 | 2jsa-4 | TYR_A_52_C | VAL_A_53_N | 2.809 | -4.858 | 0.562 | 1.846 | -5.426 | -0.148 | 1.32 | 122.42 | -149.47 |
| 310 | 2jsa-4 | VAL_A_53_N | VAL_A_53_H | 1.846 | -5.426 | -0.148 | 2.023 | -5.724 | -1.096 | 1.01 | 159.92 | -59.29 |
| 311 | 2jsa-4 | VAL_A_53_N | VAL_A_53_CA | 1.846 | -5.426 | -0.148 | 0.523 | -5.632 | 0.415 | 1.45 | 67.19 | -171.15 |
| 312 | 2jsa-4 | VAL_A_53_CA | VAL_A_53_HA | 0.523 | -5.632 | 0.415 | 0.105 | -4.66 | 0.677 | 1.09 | 76.09 | 113.27 |
| 313 | 2jsa-4 | VAL_A_53_CA | VAL_A_53_C | 0.523 | -5.632 | 0.415 | 0.643 | -6.459 | 1.697 | 1.53 | 33.10 | -81.74 |
| 314 | 2jsa-4 | VAL_A_53_C | VAL_A_53_O | 0.643 | -6.459 | 1.697 | -0.283 | -6.493 | 2.507 | 1.23 | 48.84 | -177.90 |
| 315 | 2jsa-4 | VAL_A_53_C | ASN_A_54_N | 0.643 | -6.459 | 1.697 | 1.79 | -7.106 | 1.841 | 1.32 | 83.76 | -29.43 |
| 316 | 2jsa-4 | ASN_A_54_N | ASN_A_54_H | 1.79 | -7.106 | 1.841 | 2.508 | -7.033 | 1.134 | 1.01 | 134.41 | 5.81 |
| 317 | 2jsa-4 | ASN_A_54_N | ASN_A_54_CA | 1.79 | -7.106 | 1.841 | 2.043 | -7.93 | 3.011 | 1.45 | 36.38 | -72.93 |
| 318 | 2jsa-4 | ASN_A_54_CA | ASN_A_54_HA | 2.043 | -7.93 | 3.011 | 1.318 | -7.682 | 3.786 | 1.09 | 44.67 | 161.12 |
| 319 | 2jsa-4 | ASN_A_54_CA | ASN_A_54_C | 2.043 | -7.93 | 3.011 | 3.46 | -7.662 | 3.522 | 1.53 | 70.49 | 10.71 |
| 320 | 2jsa-4 | ASN_A_54_C | ASN_A_54_O | 3.46 | -7.662 | 3.522 | 4.31 | -8.551 | 3.498 | 1.23 | 91.12 | -46.28 |
| 321 | 2jsa-4 | ASN_A_54_C | SER_A_55_N | 3.46 | -7.662 | 3.522 | 3.669 | -6.434 | 3.973 | 1.32 | 70.10 | 80.34 |
| 322 | 2jsa-4 | SER_A_55_N | SER_A_55_H | 3.669 | -6.434 | 3.973 | 2.922 | -5.754 | 3.964 | 1.01 | 90.51 | 137.69 |
| 323 | 2jsa-4 | SER_A_55_N | SER_A_55_CA | 3.669 | -6.434 | 3.973 | 4.968 | -6.038 | 4.489 | 1.45 | 69.19 | 16.95 |
| 324 | 2jsa-4 | SER_A_55_CA | SER_A_55_HA | 4.968 | -6.038 | 4.489 | 5.238 | -6.72 | 5.296 | 1.09 | 42.27 | -68.40 |
| 325 | 2jsa-4 | SER_A_55_CA | SER_A_55_C | 4.968 | -6.038 | 4.489 | 4.897 | -4.617 | 5.052 | 1.53 | 68.41 | 92.86 |
| 326 | 2jsa-4 | SER_A_55_C | SER_A_55_O | 4.897 | -4.617 | 5.052 | 5.482 | -4.327 | 6.093 | 1.23 | 32.10 | 26.37 |
| 327 | 2jsa-4 | SER_A_55_C | LYS_A_56_N | 4.897 | -4.617 | 5.052 | 4.173 | -3.767 | 4.337 | 1.33 | 122.63 | 130.42 |
| 328 | 2jsa-4 | LYS_A_56_N | LYS_A_56_H | 4.173 | -3.767 | 4.337 | 3.718 | -4.074 | 3.49 | 1.01 | 147.06 | -145.99 |
| 329 | 2jsa-4 | LYS_A_56_N | LYS_A_56_CA | 4.173 | -3.767 | 4.337 | 4.018 | -2.384 | 4.751 | 1.45 | 73.43 | 96.39 |
| 330 | 2jsa-4 | LYS_A_56_CA | LYS_A_56_HA | 4.018 | -2.384 | 4.751 | 4.39 | -2.294 | 5.772 | 1.09 | 20.55 | 13.60 |
| 331 | 2jsa-4 | LYS_A_56_CA | LYS_A_56_C | 4.018 | -2.384 | 4.751 | 2.53 | -2.023 | 4.764 | 1.53 | 89.51 | 166.36 |
| 332 | 2jsa-4 | LYS_A_56_C | LYS_A_56_O | 2.53 | -2.023 | 4.764 | 2.172 | -0.856 | 4.904 | 1.23 | 83.46 | 107.05 |
| 333 | 2jsa-4 | LYS_A_56_C | VAL_A_57_N | 2.53 | -2.023 | 4.764 | 1.704 | -3.049 | 4.615 | 1.33 | 96.45 | -128.84 |
| 334 | 2jsa-4 | VAL_A_57_N | VAL_A_57_H | 1.704 | -3.049 | 4.615 | 2.07 | -3.984 | 4.504 | 1.01 | 96.31 | -68.62 |
| 335 | 2jsa-4 | VAL_A_57_N | VAL_A_57_CA | 1.704 | -3.049 | 4.615 | 0.265 | -2.856 | 4.608 | 1.45 | 90.28 | 172.36 |
| 336 | 2jsa-4 | VAL_A_57_CA | VAL_A_57_HA | 0.265 | -2.856 | 4.608 | 0.058 | -1.821 | 4.336 | 1.09 | 104.45 | 101.31 |
| 337 | 2jsa-4 | VAL_A_57_CA | VAL_A_57_C | 0.265 | -2.856 | 4.608 | -0.285 | -3.096 | 6.015 | 1.53 | 23.10 | -156.43 |
| 338 | 2jsa-4 | VAL_A_57_C | VAL_A_57_O | -0.285 | -3.096 | 6.015 | -1.075 | -2.3 | 6.521 | 1.23 | 65.72 | 134.78 |
| 339 | 2jsa-4 | VAL_A_57_C | ASP_A_58_N | -0.285 | -3.096 | 6.015 | 0.154 | -4.197 | 6.607 | 1.32 | 63.46 | -68.26 |
| 340 | 2jsa-4 | ASP_A_58_N | ASP_A_58_H | 0.154 | -4.197 | 6.607 | 0.802 | -4.808 | 6.13 | 1.01 | 118.17 | -43.32 |
| 341 | 2jsa-4 | ASP_A_58_N | ASP_A_58_CA | 0.154 | -4.197 | 6.607 | -0.284 | -4.551 | 7.947 | 1.45 | 22.80 | -141.05 |
| 342 | 2jsa-4 | ASP_A_58_CA | ASP_A_58_HA | -0.284 | -4.551 | 7.947 | -1.333 | -4.845 | 7.912 | 1.09 | 91.84 | -164.34 |
| 343 | 2jsa-4 | ASP_A_58_CA | ASP_A_58_C | -0.284 | -4.551 | 7.947 | -0.113 | -3.344 | 8.872 | 1.53 | 52.81 | 81.94 |
| 344 | 2jsa-4 | ASP_A_58_C | ASP_A_58_O | -0.113 | -3.344 | 8.872 | -1.004 | -3.03 | 9.66 | 1.23 | 50.17 | 160.59 |
| 345 | 2jsa-4 | ASP_A_58_C | PRO_A_59_N | -0.113 | -3.344 | 8.872 | 1.067 | -2.682 | 8.741 | 1.36 | 95.53 | 29.29 |
| 346 | 2jsa-4 | PRO_A_59_N | PRO_A_59_CA | 1.067 | -2.682 | 8.741 | 1.366 | -1.516 | 9.554 | 1.45 | 55.96 | 75.62 |
| 347 | 2jsa-4 | PRO_A_59_CA | PRO_A_59_HA | 1.366 | -1.516 | 9.554 | 1.096 | -1.706 | 10.593 | 1.09 | 17.63 | -144.87 |
| 348 | 2jsa-4 | PRO_A_59_CA | PRO_A_59_C | 1.366 | -1.516 | 9.554 | 0.586 | -0.293 | 9.067 | 1.53 | 108.56 | 122.53 |
| 349 | 2jsa-4 | PRO_A_59_C | PRO_A_59_O | 0.586 | -0.293 | 9.067 | -0.17 | 0.309 | 9.828 | 1.23 | 51.78 | 141.47 |
| 350 | 2jsa-4 | PRO_A_59_C | ILE_A_60_N | 0.586 | -0.293 | 9.067 | 0.798 | 0.038 | 7.802 | 1.32 | 162.74 | 57.36 |
| 351 | 2jsa-4 | ILE_A_60_N | ILE_A_60_H | 0.798 | 0.038 | 7.802 | 1.435 | -0.503 | 7.234 | 1.01 | 124.20 | -40.34 |
| 352 | 2jsa-4 | ILE_A_60_N | ILE_A_60_CA | 0.798 | 0.038 | 7.802 | 0.124 | 1.178 | 7.204 | 1.45 | 114.30 | 120.59 |
| 353 | 2jsa-4 | ILE_A_60_CA | ILE_A_60_HA | 0.124 | 1.178 | 7.204 | 0.566 | 2.085 | 7.617 | 1.09 | 67.74 | 64.02 |
| 354 | 2jsa-4 | ILE_A_60_CA | ILE_A_60_C | 0.124 | 1.178 | 7.204 | -1.353 | 1.158 | 7.601 | 1.53 | 74.96 | -179.22 |
| 355 | 2jsa-4 | ILE_A_60_C | ILE_A_60_O | -1.353 | 1.158 | 7.601 | -1.845 | 2.099 | 8.223 | 1.23 | 59.64 | 117.60 |
| 356 | 2jsa-4 | ILE_A_60_C | ILE_A_61_N | -1.353 | 1.158 | 7.601 | -2.019 | 0.076 | 7.225 | 1.33 | 106.49 | -121.61 |
| 357 | 2jsa-4 | ILE_A_61_N | ILE_A_61_H | -2.019 | 0.076 | 7.225 | -1.549 | -0.659 | 6.716 | 1.01 | 120.26 | -57.40 |
| 358 | 2jsa-4 | ILE_A_61_N | ILE_A_61_CA | -2.019 | 0.076 | 7.225 | -3.43 | -0.079 | 7.534 | 1.45 | 77.72 | -173.73 |
| 359 | 2jsa-4 | ILE_A_61_CA | ILE_A_61_HA | -3.43 | -0.079 | 7.534 | -3.971 | 0.739 | 7.058 | 1.09 | 115.89 | 123.48 |
| 360 | 2jsa-4 | ILE_A_61_CA | ILE_A_61_C | -3.43 | -0.079 | 7.534 | -3.634 | 0.047 | 9.046 | 1.53 | 9.01 | 148.30 |
| 361 | 2jsa-4 | ILE_A_61_C | ILE_A_61_O | -3.634 | 0.047 | 9.046 | -4.727 | 0.377 | 9.503 | 1.23 | 68.19 | 163.20 |
| 362 | 2jsa-4 | ILE_A_61_C | HIS_A_62_N | -3.634 | 0.047 | 9.046 | -2.564 | -0.223 | 9.779 | 1.32 | 56.41 | -14.16 |
| 363 | 2jsa-4 | HIS_A_62_N | HIS_A_62_H | -2.564 | -0.223 | 9.779 | -1.697 | -0.49 | 9.335 | 1.01 | 116.08 | -17.12 |
| 364 | 2jsa-4 | HIS_A_62_N | HIS_A_62_CA | -2.564 | -0.223 | 9.779 | -2.612 | -0.143 | 11.229 | 1.45 | 3.68 | 120.96 |
| 365 | 2jsa-4 | HIS_A_62_CA | HIS_A_62_HA | -2.612 | -0.143 | 11.229 | -3.493 | -0.688 | 11.567 | 1.09 | 71.93 | -148.26 |
| 366 | 2jsa-4 | HIS_A_62_CA | HIS_A_62_C | -2.612 | -0.143 | 11.229 | -2.769 | 1.316 | 11.658 | 1.53 | 73.70 | 96.14 |
| 367 | 2jsa-4 | HIS_A_62_C | HIS_A_62_O | -2.769 | 1.316 | 11.658 | -3.284 | 1.597 | 12.74 | 1.23 | 28.47 | 151.38 |
| 368 | 2jsa-4 | HIS_A_62_C | GLU_A_63_N | -2.769 | 1.316 | 11.658 | -2.316 | 2.208 | 10.789 | 1.33 | 130.98 | 63.08 |
| 369 | 2jsa-4 | GLU_A_63_N | GLU_A_63_H | -2.316 | 2.208 | 10.789 | -1.904 | 1.905 | 9.918 | 1.01 | 149.58 | -36.33 |
| 370 | 2jsa-4 | GLU_A_63_N | GLU_A_63_CA | -2.316 | 2.208 | 10.789 | -2.401 | 3.632 | 11.065 | 1.45 | 79.05 | 93.42 |
| 371 | 2jsa-4 | GLU_A_63_CA | GLU_A_63_HA | -2.401 | 3.632 | 11.065 | -2.137 | 3.789 | 12.111 | 1.09 | 16.36 | 30.74 |
| 372 | 2jsa-4 | GLU_A_63_CA | GLU_A_63_C | -2.401 | 3.632 | 11.065 | -3.831 | 4.131 | 10.85 | 1.53 | 98.08 | 160.76 |
| 373 | 2jsa-4 | GLU_A_63_C | GLU_A_63_O | -3.831 | 4.131 | 10.85 | -4.403 | 4.781 | 11.723 | 1.23 | 44.76 | 131.35 |
| 374 | 2jsa-4 | GLU_A_63_C | LEU_A_64_N | -3.831 | 4.131 | 10.85 | -4.367 | 3.806 | 9.683 | 1.32 | 151.76 | -148.77 |
| 375 | 2jsa-4 | LEU_A_64_N | LEU_A_64_H | -4.367 | 3.806 | 9.683 | -3.837 | 3.268 | 9.012 | 1.01 | 131.62 | -45.43 |
| 376 | 2jsa-4 | LEU_A_64_N | LEU_A_64_CA | -4.367 | 3.806 | 9.683 | -5.72 | 4.213 | 9.342 | 1.45 | 103.57 | 163.26 |
| 377 | 2jsa-4 | LEU_A_64_CA | LEU_A_64_HA | -5.72 | 4.213 | 9.342 | -5.799 | 5.288 | 9.503 | 1.09 | 81.50 | 94.20 |
| 378 | 2jsa-4 | LEU_A_64_CA | LEU_A_64_C | -5.72 | 4.213 | 9.342 | -6.708 | 3.527 | 10.287 | 1.53 | 51.84 | -145.23 |
| 379 | 2jsa-4 | LEU_A_64_C | LEU_A_64_O | -6.708 | 3.527 | 10.287 | -7.788 | 4.054 | 10.552 | 1.23 | 77.56 | 153.99 |
| 380 | 2jsa-4 | LEU_A_64_C | GLU_A_65_N | -6.708 | 3.527 | 10.287 | -6.303 | 2.361 | 10.77 | 1.33 | 68.63 | -70.85 |
| 381 | 2jsa-4 | GLU_A_65_N | GLU_A_65_H | -6.303 | 2.361 | 10.77 | -5.401 | 1.988 | 10.509 | 1.01 | 104.97 | -22.47 |
| 382 | 2jsa-4 | GLU_A_65_N | GLU_A_65_CA | -6.303 | 2.361 | 10.77 | -7.139 | 1.597 | 11.68 | 1.45 | 51.22 | -137.58 |
| 383 | 2jsa-4 | GLU_A_65_CA | GLU_A_65_HA | -7.139 | 1.597 | 11.68 | -8.126 | 1.499 | 11.229 | 1.09 | 114.45 | -174.33 |
| 384 | 2jsa-4 | GLU_A_65_CA | GLU_A_65_C | -7.139 | 1.597 | 11.68 | -7.29 | 2.337 | 13.011 | 1.53 | 29.57 | 101.53 |
| 385 | 2jsa-4 | GLU_A_65_C | GLU_A_65_O | -7.29 | 2.337 | 13.011 | -8.399 | 2.484 | 13.522 | 1.23 | 65.45 | 172.45 |
| 386 | 2jsa-4 | GLU_A_65_C | GLY_A_66_N | -7.29 | 2.337 | 13.011 | -6.157 | 2.784 | 13.534 | 1.33 | 66.76 | 21.53 |
| 387 | 2jsa-4 | GLY_A_66_N | GLY_A_66_H | -6.157 | 2.784 | 13.534 | -5.28 | 2.627 | 13.058 | 1.01 | 118.11 | -10.15 |
| 388 | 2jsa-4 | GLY_A_66_N | GLY_A_66_CA | -6.157 | 2.784 | 13.534 | -6.149 | 3.505 | 14.795 | 1.45 | 29.76 | 89.36 |
| 389 | 2jsa-4 | GLY_A_66_CA | GLY_A_66_C | -6.149 | 3.505 | 14.795 | -6.874 | 4.847 | 14.665 | 1.53 | 94.87 | 118.38 |
| 390 | 2jsa-4 | GLY_A_66_C | GLY_A_66_O | -6.874 | 4.847 | 14.665 | -7.818 | 5.121 | 15.404 | 1.23 | 53.06 | 163.81 |
| 391 | 2jsa-4 | GLY_A_66_C | GLY_A_67_N | -6.874 | 4.847 | 14.665 | -6.406 | 5.647 | 13.718 | 1.33 | 135.62 | 59.67 |
| 392 | 2jsa-4 | GLY_A_67_N | GLY_A_67_H | -6.406 | 5.647 | 13.718 | -5.626 | 5.355 | 13.146 | 1.01 | 124.48 | -20.52 |

| | | | | | | | | | | | | |
|---|---|---|---|---|---|---|---|---|---|---|---|---|
| 393 | 2jsa-4 | GLY_A_67_N | GLY_A_67_CA | -6.406 | 5.647 | 13.718 | -6.997 | 6.953 | 13.481 | 1.45 | 99.39 | 114.35 |
| 394 | 2jsa-4 | GLY_A_67_CA | GLY_A_67_C | -6.997 | 6.953 | 13.481 | -5.999 | 7.892 | 12.801 | 1.53 | 116.39 | 43.26 |
| 395 | 2jsa-4 | GLY_A_67_C | GLY_A_67_O | -5.999 | 7.892 | 12.801 | -5.887 | 9.059 | 13.174 | 1.23 | 72.35 | 84.52 |
| 396 | 2jsa-4 | GLY_A_67_C | THR_A_68_N | -5.999 | 7.892 | 12.801 | -5.298 | 7.347 | 11.818 | 1.32 | 137.91 | -37.86 |
| 397 | 2jsa-4 | THR_A_68_N | THR_A_68_H | -5.298 | 7.347 | 11.818 | -5.441 | 6.38 | 11.566 | 1.01 | 104.46 | -98.41 |
| 398 | 2jsa-4 | THR_A_68_N | THR_A_68_CA | -5.298 | 7.347 | 11.818 | -4.313 | 8.122 | 11.083 | 1.45 | 120.39 | 38.20 |
| 399 | 2jsa-4 | THR_A_68_CA | THR_A_68_HA | -4.313 | 8.122 | 11.083 | -4.256 | 9.12 | 11.519 | 1.09 | 66.43 | 86.73 |
| 400 | 2jsa-4 | THR_A_68_CA | THR_A_68_C | -4.313 | 8.122 | 11.083 | -4.734 | 8.271 | 9.62 | 1.53 | 163.02 | 160.51 |
| 401 | 2jsa-4 | THR_A_68_C | THR_A_68_O | -4.734 | 8.271 | 9.62 | -5.17 | 7.305 | 8.994 | 1.23 | 120.57 | -114.29 |
| 402 | 2jsa-4 | THR_A_68_C | ALA_A_69_N | -4.734 | 8.271 | 9.62 | -4.589 | 9.487 | 9.116 | 1.32 | 112.37 | 83.20 |
| 403 | 2jsa-4 | ALA_A_69_N | ALA_A_69_H | -4.589 | 9.487 | 9.116 | -4.224 | 10.234 | 9.689 | 1.01 | 55.43 | 63.96 |
| 404 | 2jsa-4 | ALA_A_69_N | ALA_A_69_CA | -4.589 | 9.487 | 9.116 | -4.949 | 9.775 | 7.738 | 1.45 | 161.50 | 141.34 |
| 405 | 2jsa-4 | ALA_A_69_CA | ALA_A_69_HA | -4.949 | 9.775 | 7.738 | -5.968 | 9.432 | 7.564 | 1.09 | 99.19 | -161.40 |
| 406 | 2jsa-4 | ALA_A_69_CA | ALA_A_69_C | -4.949 | 9.775 | 7.738 | -4.014 | 9.008 | 6.8 | 1.53 | 127.80 | -39.36 |
| 407 | 2jsa-4 | ALA_A_69_C | ALA_A_69_O | -4.014 | 9.008 | 6.8 | -3.023 | 8.429 | 7.243 | 1.23 | 68.89 | -30.30 |
| 408 | 2jsa-4 | ALA_A_69_C | PRO_A_70_N | -4.014 | 9.008 | 6.8 | -4.369 | 9.032 | 5.488 | 1.36 | 164.83 | 176.13 |
| 409 | 2jsa-4 | PRO_A_70_N | PRO_A_70_CA | -4.369 | 9.032 | 5.488 | -3.573 | 8.347 | 4.484 | 1.45 | 133.71 | -40.71 |
| 410 | 2jsa-4 | PRO_A_70_CA | PRO_A_70_HA | -3.573 | 8.347 | 4.484 | -3.304 | 7.351 | 4.836 | 1.09 | 71.16 | -74.89 |
| 411 | 2jsa-4 | PRO_A_70_CA | PRO_A_70_C | -3.573 | 8.347 | 4.484 | -2.286 | 9.119 | 4.185 | 1.53 | 101.27 | 30.96 |
| 412 | 2jsa-4 | PRO_A_70_C | PRO_A_70_O | -2.286 | 9.119 | 4.185 | -1.323 | 8.551 | 3.671 | 1.23 | 114.69 | -30.53 |
| 413 | 2jsa-4 | PRO_A_70_C | LYS_A_71_N | -2.286 | 9.119 | 4.185 | -2.311 | 10.401 | 4.52 | 1.33 | 75.36 | 91.12 |
| 414 | 2jsa-4 | LYS_A_71_N | LYS_A_71_H | -2.311 | 10.401 | 4.52 | -3.137 | 10.801 | 4.94 | 1.01 | 65.41 | 154.16 |
| 415 | 2jsa-4 | LYS_A_71_N | LYS_A_71_CA | -2.311 | 10.401 | 4.52 | -1.158 | 11.255 | 4.293 | 1.45 | 98.99 | 36.53 |
| 416 | 2jsa-4 | LYS_A_71_CA | LYS_A_71_HA | -1.158 | 11.255 | 4.293 | -0.547 | 10.799 | 3.514 | 1.09 | 135.62 | -36.73 |
| 417 | 2jsa-4 | LYS_A_71_CA | LYS_A_71_C | -1.158 | 11.255 | 4.293 | -0.315 | 11.317 | 5.569 | 1.53 | 33.52 | 4.21 |
| 418 | 2jsa-4 | LYS_A_71_C | LYS_A_71_O | -0.315 | 11.317 | 5.569 | 0.785 | 11.867 | 5.565 | 1.23 | 90.19 | 26.57 |
| 419 | 2jsa-4 | LYS_A_71_C | ASP_A_72_N | -0.315 | 11.317 | 5.569 | -0.864 | 10.744 | 6.63 | 1.32 | 36.79 | -133.77 |
| 420 | 2jsa-4 | ASP_A_72_N | ASP_A_72_H | -0.864 | 10.744 | 6.63 | -1.773 | 10.31 | 6.561 | 1.01 | 93.92 | -154.48 |
| 421 | 2jsa-4 | ASP_A_72_N | ASP_A_72_CA | -0.864 | 10.744 | 6.63 | -0.176 | 10.727 | 7.91 | 1.45 | 28.27 | -1.42 |
| 422 | 2jsa-4 | ASP_A_72_CA | ASP_A_72_HA | -0.176 | 10.727 | 7.91 | 0.556 | 11.535 | 7.933 | 1.09 | 88.79 | 47.83 |
| 423 | 2jsa-4 | ASP_A_72_CA | ASP_A_72_C | -0.176 | 10.727 | 7.91 | 0.528 | 9.381 | 8.087 | 1.53 | 83.35 | -62.39 |
| 424 | 2jsa-4 | ASP_A_72_C | ASP_A_72_O | 0.528 | 9.381 | 8.087 | 1.631 | 9.32 | 8.628 | 1.23 | 63.91 | -3.17 |
| 425 | 2jsa-4 | ASP_A_72_C | VAL_A_73_N | 0.528 | 9.381 | 8.087 | -0.137 | 8.334 | 7.621 | 1.32 | 110.59 | -122.42 |
| 426 | 2jsa-4 | VAL_A_73_N | VAL_A_73_H | -0.137 | 8.334 | 7.621 | -1.04 | 8.458 | 7.188 | 1.01 | 115.41 | 172.18 |
| 427 | 2jsa-4 | VAL_A_73_N | VAL_A_73_CA | -0.137 | 8.334 | 7.621 | 0.412 | 6.992 | 7.721 | 1.45 | 86.05 | -67.75 |
| 428 | 2jsa-4 | VAL_A_73_CA | VAL_A_73_HA | 0.412 | 6.992 | 7.721 | 0.442 | 6.71 | 8.774 | 1.09 | 15.07 | -83.93 |
| 429 | 2jsa-4 | VAL_A_73_CA | VAL_A_73_C | 0.412 | 6.992 | 7.721 | 1.847 | 6.993 | 7.189 | 1.53 | 110.34 | 0.04 |
| 430 | 2jsa-4 | VAL_A_73_C | VAL_A_73_O | 1.847 | 6.993 | 7.189 | 2.769 | 6.566 | 7.882 | 1.23 | 55.70 | -24.85 |
| 431 | 2jsa-4 | VAL_A_73_C | CYS_A_74_N | 1.847 | 6.993 | 7.189 | 1.989 | 7.476 | 5.964 | 1.32 | 157.66 | 73.62 |
| 432 | 2jsa-4 | CYS_A_74_N | CYS_A_74_H | 1.989 | 7.476 | 5.964 | 1.185 | 7.811 | 5.453 | 1.01 | 120.40 | 157.38 |
| 433 | 2jsa-4 | CYS_A_74_N | CYS_A_74_CA | 1.989 | 7.476 | 5.964 | 3.296 | 7.537 | 5.33 | 1.45 | 115.85 | 2.67 |
| 434 | 2jsa-4 | CYS_A_74_CA | CYS_A_74_HA | 3.296 | 7.537 | 5.33 | 3.868 | 6.662 | 5.638 | 1.09 | 73.58 | -56.83 |
| 435 | 2jsa-4 | CYS_A_74_CA | CYS_A_74_C | 3.296 | 7.537 | 5.33 | 4.015 | 8.791 | 5.834 | 1.53 | 70.78 | 60.17 |
| 436 | 2jsa-4 | CYS_A_74_C | CYS_A_74_O | 4.015 | 8.791 | 5.834 | 4.454 | 9.62 | 5.039 | 1.23 | 130.28 | 62.10 |
| 437 | 2jsa-4 | CYS_A_74_C | THR_A_75_N | 4.015 | 8.791 | 5.834 | 4.114 | 8.889 | 7.151 | 1.32 | 6.04 | 44.71 |
| 438 | 2jsa-4 | THR_A_75_N | THR_A_75_H | 4.114 | 8.889 | 7.151 | 3.731 | 8.168 | 7.745 | 1.01 | 53.96 | -117.98 |
| 439 | 2jsa-4 | THR_A_75_N | THR_A_75_CA | 4.114 | 8.889 | 7.151 | 4.772 | 10.026 | 7.77 | 1.45 | 64.77 | 59.94 |
| 440 | 2jsa-4 | THR_A_75_CA | THR_A_75_HA | 4.772 | 10.026 | 7.77 | 5.513 | 10.423 | 7.075 | 1.09 | 129.58 | 28.18 |
| 441 | 2jsa-4 | THR_A_75_CA | THR_A_75_C | 4.772 | 10.026 | 7.77 | 5.504 | 9.592 | 9.041 | 1.53 | 33.80 | -30.66 |
| 442 | 2jsa-4 | THR_A_75_C | THR_A_75_O | 5.504 | 9.592 | 9.041 | 6.711 | 9.798 | 9.167 | 1.23 | 84.12 | 9.69 |
| 443 | 2jsa-4 | THR_A_75_C | LYS_A_76_N | 5.504 | 9.592 | 9.041 | 4.744 | 9.001 | 9.95 | 1.32 | 46.64 | -142.13 |
| 444 | 2jsa-4 | LYS_A_76_N | LYS_A_76_H | 4.744 | 9.001 | 9.95 | 3.758 | 8.865 | 9.779 | 1.01 | 99.75 | -172.15 |
| 445 | 2jsa-4 | LYS_A_76_N | LYS_A_76_CA | 4.744 | 9.001 | 9.95 | 5.307 | 8.537 | 11.208 | 1.45 | 30.11 | -39.49 |
| 446 | 2jsa-4 | LYS_A_76_CA | LYS_A_76_HA | 5.307 | 8.537 | 11.208 | 6.098 | 9.228 | 11.501 | 1.09 | 74.41 | 41.14 |
| 447 | 2jsa-4 | LYS_A_76_CA | LYS_A_76_C | 5.307 | 8.537 | 11.208 | 5.944 | 7.162 | 10.999 | 1.53 | 97.85 | -65.14 |
| 448 | 2jsa-4 | LYS_A_76_C | LYS_A_76_O | 5.944 | 7.162 | 10.999 | 6.858 | 6.781 | 11.728 | 1.23 | 53.64 | -22.63 |
| 449 | 2jsa-4 | LYS_A_76_C | LEU_A_77_N | 5.944 | 7.162 | 10.999 | 5.438 | 6.455 | 10.0 | 1.32 | 138.97 | -125.59 |
| 450 | 2jsa-4 | LEU_A_77_N | LEU_A_77_H | 5.438 | 6.455 | 10.0 | 4.686 | 6.832 | 9.441 | 1.01 | 123.60 | 153.37 |
| 451 | 2jsa-4 | LEU_A_77_N | LEU_A_77_CA | 5.438 | 6.455 | 10.0 | 5.947 | 5.131 | 9.685 | 1.45 | 102.52 | -68.97 |
| 452 | 2jsa-4 | LEU_A_77_CA | LEU_A_77_HA | 5.947 | 5.131 | 9.685 | 6.15 | 4.616 | 10.624 | 1.09 | 30.52 | -68.49 |
| 453 | 2jsa-4 | LEU_A_77_CA | LEU_A_77_C | 5.947 | 5.131 | 9.685 | 7.272 | 5.264 | 8.933 | 1.53 | 119.45 | 5.73 |
| 454 | 2jsa-4 | LEU_A_77_C | LEU_A_77_O | 7.272 | 5.264 | 8.933 | 7.875 | 4.264 | 8.548 | 1.23 | 108.25 | -58.91 |
| 455 | 2jsa-4 | LEU_A_77_C | ASN_A_78_N | 7.272 | 5.264 | 8.933 | 7.688 | 6.509 | 8.749 | 1.33 | 97.98 | 71.52 |
| 456 | 2jsa-4 | ASN_A_78_N | ASN_A_78_H | 7.688 | 6.509 | 8.749 | 7.142 | 7.285 | 9.096 | 1.01 | 69.91 | 125.13 |
| 457 | 2jsa-4 | ASN_A_78_N | ASN_A_78_CA | 7.688 | 6.509 | 8.749 | 8.931 | 6.786 | 8.049 | 1.45 | 118.80 | 12.56 |
| 458 | 2jsa-4 | ASN_A_78_CA | ASN_A_78_HA | 8.931 | 6.786 | 8.049 | 9.048 | 7.865 | 7.944 | 1.09 | 95.53 | 83.81 |
| 459 | 2jsa-4 | ASN_A_78_CA | ASN_A_78_C | 8.931 | 6.786 | 8.049 | 8.893 | 6.13 | 6.667 | 1.53 | 154.57 | -93.32 |
| 460 | 2jsa-4 | ASN_A_78_C | ASN_A_78_O | 8.893 | 6.13 | 6.667 | 9.928 | 5.965 | 6.025 | 1.23 | 121.49 | -9.06 |
| 461 | 2jsa-4 | ASN_A_78_C | GLU_A_79_N | 8.893 | 6.13 | 6.667 | 7.686 | 5.772 | 6.251 | 1.33 | 108.29 | -163.48 |
| 462 | 2jsa-4 | GLU_A_79_N | GLU_A_79_H | 7.686 | 5.772 | 6.251 | 6.878 | 5.938 | 6.833 | 1.01 | 54.79 | 168.39 |
| 463 | 2jsa-4 | GLU_A_79_N | GLU_A_79_CA | 7.686 | 5.772 | 6.251 | 7.498 | 5.138 | 4.958 | 1.45 | 152.91 | -106.52 |
| 464 | 2jsa-4 | GLU_A_79_CA | GLU_A_79_HA | 7.498 | 5.138 | 4.958 | 8.462 | 4.749 | 4.63 | 1.09 | 107.51 | -21.98 |
| 465 | 2jsa-4 | GLU_A_79_CA | GLU_A_79_C | 7.498 | 5.138 | 4.958 | 7.018 | 6.162 | 3.928 | 1.53 | 132.33 | 115.11 |
| 466 | 2jsa-4 | GLU_A_79_C | GLU_A_79_O | 7.018 | 6.162 | 3.928 | 5.967 | 5.986 | 3.316 | 1.23 | 119.87 | -170.49 |
| 467 | 2jsa-4 | GLU_A_79_C | CYS_A_80_N | 7.018 | 6.162 | 3.928 | 7.812 | 7.212 | 3.771 | 1.33 | 96.80 | 52.90 |
| 468 | 2jsa-4 | CYS_A_80_N | CYS_A_80_H | 7.812 | 7.212 | 3.771 | 8.663 | 7.292 | 4.31 | 1.01 | 57.76 | 5.37 |
| 469 | 2jsa-4 | CYS_A_80_N | CYS_A_80_CA | 7.812 | 7.212 | 3.771 | 7.481 | 8.265 | 2.826 | 1.45 | 130.57 | 107.45 |
| 470 | 2jsa-4 | CYS_A_80_CA | CYS_A_80_HA | 7.481 | 8.265 | 2.826 | 7.702 | 7.906 | 1.821 | 1.09 | 157.24 | -58.38 |
| 471 | 2jsa-4 | CYS_A_80_CA | CYS_A_80_C | 7.481 | 8.265 | 2.826 | 8.375 | 9.472 | 3.119 | 1.53 | 78.96 | 53.47 |
| 472 | 2jsa-4 | CYS_A_80_C | CYS_A_80_O | 8.375 | 9.472 | 3.119 | 8.782 | 9.685 | 4.26 | 1.23 | 21.93 | 27.62 |
| 473 | 2jsa-4 | CYS_A_80_C | PRO_A_81_N | 8.375 | 9.472 | 3.119 | 8.663 | 10.25 | 2.042 | 1.36 | 142.39 | 69.69 |
| 474 | 2jsa-4 | PRO_A_81_N | PRO_A_81_CA | 8.663 | 10.25 | 2.042 | 9.5 | 11.429 | 2.171 | 1.45 | 84.90 | 54.63 |
| 475 | 2jsa-4 | PRO_A_81_CA | PRO_A_81_HA | 9.5 | 11.429 | 2.171 | 10.376 | 11.199 | 2.778 | 1.09 | 56.17 | -14.71 |
| 476 | 2jsa-4 | PRO_A_81_CA | PRO_A_81_C | 9.5 | 11.429 | 2.171 | 8.736 | 12.575 | 2.838 | 1.53 | 64.16 | 123.69 |
| 477 | 2jsa-4 | PRO_A_81_C | PRO_A_81_O | 8.736 | 12.575 | 2.838 | 8.285 | 12.447 | 3.975 | 1.23 | 22.41 | -164.16 |
| 1 | 2jsa-5 | ARG_A_1_N | ARG_A_1_H | 14.073 | 1.616 | -0.782 | 14.844 | 2.125 | -1.191 | 1.01 | 113.88 | 33.43 |
| 2 | 2jsa-5 | ARG_A_1_N | ARG_A_1_CA | 14.073 | 1.616 | -0.782 | 12.928 | 2.365 | -0.295 | 1.45 | 70.41 | 146.81 |
| 3 | 2jsa-5 | ARG_A_1_CA | ARG_A_1_HA | 12.928 | 2.365 | -0.295 | 12.046 | 2.042 | -0.848 | 1.09 | 120.49 | -159.89 |
| 4 | 2jsa-5 | ARG_A_1_CA | ARG_A_1_C | 12.928 | 2.365 | -0.295 | 13.136 | 3.863 | -0.521 | 1.53 | 98.50 | 82.09 |
| 5 | 2jsa-5 | ARG_A_1_C | ARG_A_1_O | 13.136 | 3.863 | -0.521 | 14.159 | 4.275 | -1.066 | 1.23 | 116.30 | 21.94 |
| 6 | 2jsa-5 | ARG_A_1_C | SER_A_2_N | 13.136 | 3.863 | -0.521 | 12.151 | 4.637 | -0.092 | 1.32 | 71.10 | 141.84 |
| 7 | 2jsa-5 | SER_A_2_N | SER_A_2_H | 12.151 | 4.637 | -0.092 | 11.339 | 4.229 | 0.348 | 1.01 | 64.16 | -153.32 |
| 8 | 2jsa-5 | SER_A_2_N | SER_A_2_CA | 12.151 | 4.637 | -0.092 | 12.213 | 6.081 | -0.24 | 1.45 | 95.85 | 87.54 |
| 9 | 2jsa-5 | SER_A_2_CA | SER_A_2_HA | 12.213 | 6.081 | -0.24 | 11.486 | 6.526 | 0.439 | 1.09 | 51.46 | 148.53 |
| 10 | 2jsa-5 | SER_A_2_CA | SER_A_2_C | 12.213 | 6.081 | -0.24 | 11.851 | 6.475 | -1.674 | 1.53 | 159.54 | 132.58 |
| 11 | 2jsa-5 | SER_A_2_C | SER_A_2_O | 11.851 | 6.475 | -1.674 | 12.578 | 7.232 | -2.316 | 1.23 | 121.45 | 46.16 |

| | | | | | | | | | | | | |
|---|---|---|---|---|---|---|---|---|---|---|---|---|
| 12 | 2jsa-5 | SER_A_2_C | ALA_A_3_N | 11.851 | 6.475 | -1.674 | 10.728 | 5.943 | -2.134 | 1.33 | 110.31 | -154.65 |
| 13 | 2jsa-5 | ALA_A_3_N | ALA_A_3_H | 10.728 | 5.943 | -2.134 | 10.181 | 5.327 | -1.55 | 1.01 | 54.67 | -131.60 |
| 14 | 2jsa-5 | ALA_A_3_N | ALA_A_3_CA | 10.728 | 5.943 | -2.134 | 10.261 | 6.23 | -3.48 | 1.45 | 157.84 | 148.43 |
| 15 | 2jsa-5 | ALA_A_3_CA | ALA_A_3_HA | 10.261 | 6.23 | -3.48 | 10.086 | 7.302 | -3.569 | 1.09 | 94.68 | 99.27 |
| 16 | 2jsa-5 | ALA_A_3_CA | ALA_A_3_C | 10.261 | 6.23 | -3.48 | 8.937 | 5.501 | -3.722 | 1.53 | 99.10 | -151.16 |
| 17 | 2jsa-5 | ALA_A_3_C | ALA_A_3_O | 8.937 | 5.501 | -3.722 | 7.865 | 6.087 | -3.569 | 1.23 | 82.86 | 151.34 |
| 18 | 2jsa-5 | ALA_A_3_C | LEU_A_4_N | 8.937 | 5.501 | -3.722 | 9.054 | 4.236 | -4.095 | 1.32 | 106.36 | -84.72 |
| 19 | 2jsa-5 | LEU_A_4_N | LEU_A_4_H | 9.054 | 4.236 | -4.095 | 9.968 | 3.819 | -4.201 | 1.01 | 96.02 | -24.52 |
| 20 | 2jsa-5 | LEU_A_4_N | LEU_A_4_CA | 9.054 | 4.236 | -4.095 | 7.88 | 3.422 | -4.359 | 1.45 | 100.47 | -145.26 |
| 21 | 2jsa-5 | LEU_A_4_CA | LEU_A_4_HA | 7.88 | 3.422 | -4.359 | 7.173 | 4.025 | -4.929 | 1.09 | 121.53 | 139.54 |
| 22 | 2jsa-5 | LEU_A_4_CA | LEU_A_4_C | 7.88 | 3.422 | -4.359 | 7.213 | 3.052 | -3.033 | 1.53 | 29.91 | -150.98 |
| 23 | 2jsa-5 | LEU_A_4_C | LEU_A_4_O | 7.213 | 3.052 | -3.033 | 5.987 | 3.011 | -2.941 | 1.23 | 85.71 | -178.08 |
| 24 | 2jsa-5 | LEU_A_4_C | SER_A_5_N | 7.213 | 3.052 | -3.033 | 8.049 | 2.789 | -2.04 | 1.32 | 41.43 | -17.46 |
| 25 | 2jsa-5 | SER_A_5_N | SER_A_5_H | 8.049 | 2.789 | -2.04 | 9.047 | 2.841 | -2.188 | 1.01 | 98.42 | 2.98 |
| 26 | 2jsa-5 | SER_A_5_N | SER_A_5_CA | 8.049 | 2.789 | -2.04 | 7.556 | 2.423 | -0.723 | 1.45 | 25.00 | -143.41 |
| 27 | 2jsa-5 | SER_A_5_CA | SER_A_5_HA | 7.556 | 2.423 | -0.723 | 7.079 | 1.446 | -0.796 | 1.09 | 93.84 | -116.02 |
| 28 | 2jsa-5 | SER_A_5_CA | SER_A_5_C | 7.556 | 2.423 | -0.723 | 6.516 | 3.442 | -0.252 | 1.53 | 72.07 | 135.58 |
| 29 | 2jsa-5 | SER_A_5_C | SER_A_5_O | 6.516 | 3.442 | -0.252 | 5.507 | 3.073 | 0.346 | 1.23 | 60.90 | -159.91 |
| 30 | 2jsa-5 | SER_A_5_C | CYS_A_6_N | 6.516 | 3.442 | -0.252 | 6.799 | 4.704 | -0.539 | 1.32 | 102.51 | 77.36 |
| 31 | 2jsa-5 | CYS_A_6_N | CYS_A_6_H | 6.799 | 4.704 | -0.539 | 7.649 | 4.935 | -1.034 | 1.01 | 119.33 | 15.20 |
| 32 | 2jsa-5 | CYS_A_6_N | CYS_A_6_CA | 6.799 | 4.704 | -0.539 | 5.901 | 5.779 | -0.152 | 1.45 | 74.56 | 129.87 |
| 33 | 2jsa-5 | CYS_A_6_CA | CYS_A_6_HA | 5.901 | 5.779 | -0.152 | 5.624 | 5.635 | 0.892 | 1.09 | 16.65 | -152.53 |
| 34 | 2jsa-5 | CYS_A_6_CA | CYS_A_6_C | 5.901 | 5.779 | -0.152 | 4.638 | 5.682 | -1.009 | 1.53 | 124.08 | -175.61 |
| 35 | 2jsa-5 | CYS_A_6_C | CYS_A_6_O | 4.638 | 5.682 | -1.009 | 3.535 | 5.539 | -0.484 | 1.23 | 64.73 | -172.61 |
| 36 | 2jsa-5 | CYS_A_6_C | GLN_A_7_N | 4.638 | 5.682 | -1.009 | 4.841 | 5.763 | -2.317 | 1.33 | 170.51 | 21.75 |
| 37 | 2jsa-5 | GLN_A_7_N | GLN_A_7_H | 4.841 | 5.763 | -2.317 | 5.776 | 5.881 | -2.68 | 1.01 | 111.07 | 7.19 |
| 38 | 2jsa-5 | GLN_A_7_N | GLN_A_7_CA | 4.841 | 5.763 | -2.317 | 3.732 | 5.686 | -3.252 | 1.45 | 130.07 | -176.03 |
| 39 | 2jsa-5 | GLN_A_7_CA | GLN_A_7_HA | 3.732 | 5.686 | -3.252 | 3.123 | 6.581 | -3.126 | 1.09 | 83.36 | 124.23 |
| 40 | 2jsa-5 | GLN_A_7_CA | GLN_A_7_C | 3.732 | 5.686 | -3.252 | 2.863 | 4.465 | -2.946 | 1.53 | 78.46 | -125.44 |
| 41 | 2jsa-5 | GLN_A_7_C | GLN_A_7_O | 2.863 | 4.465 | -2.946 | 1.656 | 4.592 | -2.746 | 1.23 | 80.64 | 173.99 |
| 42 | 2jsa-5 | GLN_A_7_C | MET_A_8_N | 2.863 | 4.465 | -2.946 | 3.509 | 3.309 | -2.918 | 1.32 | 88.79 | -60.80 |
| 43 | 2jsa-5 | MET_A_8_N | MET_A_8_H | 3.509 | 3.309 | -2.918 | 4.504 | 3.279 | -3.093 | 1.01 | 99.97 | -1.73 |
| 44 | 2jsa-5 | MET_A_8_N | MET_A_8_CA | 3.509 | 3.309 | -2.918 | 2.811 | 2.066 | -2.639 | 1.45 | 78.93 | -119.32 |
| 45 | 2jsa-5 | MET_A_8_CA | MET_A_8_HA | 2.811 | 2.066 | -2.639 | 2.2 | 1.807 | -3.503 | 1.09 | 142.47 | -157.03 |
| 46 | 2jsa-5 | MET_A_8_CA | MET_A_8_C | 2.811 | 2.066 | -2.639 | 1.905 | 2.21 | -1.414 | 1.53 | 36.83 | 170.97 |
| 47 | 2jsa-5 | MET_A_8_C | MET_A_8_O | 1.905 | 2.21 | -1.414 | 0.766 | 1.747 | -1.423 | 1.23 | 90.42 | -157.88 |
| 48 | 2jsa-5 | MET_A_8_C | CYS_A_9_N | 1.905 | 2.21 | -1.414 | 2.446 | 2.853 | -0.39 | 1.32 | 39.37 | 49.92 |
| 49 | 2jsa-5 | CYS_A_9_N | CYS_A_9_H | 2.446 | 2.853 | -0.39 | 3.39 | 3.208 | -0.453 | 1.01 | 93.57 | 20.61 |
| 50 | 2jsa-5 | CYS_A_9_N | CYS_A_9_CA | 2.446 | 2.853 | -0.39 | 1.7 | 3.063 | 0.84 | 1.45 | 32.21 | 164.28 |
| 51 | 2jsa-5 | CYS_A_9_CA | CYS_A_9_HA | 1.7 | 3.063 | 0.84 | 1.414 | 2.09 | 1.239 | 1.09 | 68.52 | -106.38 |
| 52 | 2jsa-5 | CYS_A_9_CA | CYS_A_9_C | 1.7 | 3.063 | 0.84 | 0.428 | 3.845 | 0.503 | 1.53 | 102.72 | 148.42 |
| 53 | 2jsa-5 | CYS_A_9_C | CYS_A_9_O | 0.428 | 3.845 | 0.503 | -0.597 | 3.683 | 1.163 | 1.23 | 57.54 | -171.02 |
| 54 | 2jsa-5 | CYS_A_9_C | GLU_A_10_N | 0.428 | 3.845 | 0.503 | 0.538 | 4.675 | -0.524 | 1.33 | 140.81 | 82.45 |
| 55 | 2jsa-5 | GLU_A_10_N | GLU_A_10_H | 0.538 | 4.675 | -0.524 | 1.413 | 4.756 | -1.022 | 1.01 | 119.54 | 5.29 |
| 56 | 2jsa-5 | GLU_A_10_N | GLU_A_10_CA | 0.538 | 4.675 | -0.524 | -0.591 | 5.482 | -0.955 | 1.45 | 107.25 | 144.44 |
| 57 | 2jsa-5 | GLU_A_10_CA | GLU_A_10_HA | -0.591 | 5.482 | -0.955 | -1.122 | 5.823 | -0.066 | 1.09 | 35.37 | 147.29 |
| 58 | 2jsa-5 | GLU_A_10_CA | GLU_A_10_C | -0.591 | 5.482 | -0.955 | -1.553 | 4.641 | -1.798 | 1.53 | 123.41 | -138.84 |
| 59 | 2jsa-5 | GLU_A_10_C | GLU_A_10_O | -1.553 | 4.641 | -1.798 | -2.652 | 5.09 | -2.125 | 1.23 | 105.40 | 157.78 |
| 60 | 2jsa-5 | GLU_A_10_C | LEU_A_11_N | -1.553 | 4.641 | -1.798 | -1.106 | 3.438 | -2.125 | 1.32 | 104.29 | -69.62 |
| 61 | 2jsa-5 | LEU_A_11_N | LEU_A_11_H | -1.106 | 3.438 | -2.125 | -0.191 | 3.135 | -1.821 | 1.01 | 72.49 | -18.32 |
| 62 | 2jsa-5 | LEU_A_11_N | LEU_A_11_CA | -1.106 | 3.438 | -2.125 | -1.913 | 2.531 | -2.923 | 1.45 | 123.32 | -131.66 |
| 63 | 2jsa-5 | LEU_A_11_CA | LEU_A_11_HA | -1.913 | 2.531 | -2.923 | -2.597 | 3.129 | -3.525 | 1.09 | 123.53 | 138.84 |
| 64 | 2jsa-5 | LEU_A_11_CA | LEU_A_11_C | -1.913 | 2.531 | -2.923 | -2.755 | 1.654 | -1.995 | 1.53 | 52.65 | -133.83 |
| 65 | 2jsa-5 | LEU_A_11_C | LEU_A_11_O | -2.755 | 1.654 | -1.995 | -3.92 | 1.384 | -2.278 | 1.23 | 103.31 | -166.95 |
| 66 | 2jsa-5 | LEU_A_11_C | VAL_A_12_N | -2.755 | 1.654 | -1.995 | -2.131 | 1.233 | -0.904 | 1.33 | 34.60 | -34.01 |
| 67 | 2jsa-5 | VAL_A_12_N | VAL_A_12_H | -2.131 | 1.233 | -0.904 | -1.17 | 1.494 | -0.735 | 1.01 | 80.37 | 15.19 |
| 68 | 2jsa-5 | VAL_A_12_N | VAL_A_12_CA | -2.131 | 1.233 | -0.904 | -2.809 | 0.392 | 0.068 | 1.45 | 48.02 | -128.88 |
| 69 | 2jsa-5 | VAL_A_12_CA | VAL_A_12_HA | -2.809 | 0.392 | 0.068 | -3.352 | -0.386 | -0.469 | 1.09 | 119.51 | -124.91 |
| 70 | 2jsa-5 | VAL_A_12_CA | VAL_A_12_C | -2.809 | 0.392 | 0.068 | -3.825 | 1.232 | 0.844 | 1.53 | 59.52 | 140.42 |
| 71 | 2jsa-5 | VAL_A_12_C | VAL_A_12_O | -3.825 | 1.232 | 0.844 | -4.972 | 0.823 | 1.016 | 1.23 | 81.96 | -160.37 |
| 72 | 2jsa-5 | VAL_A_12_C | VAL_A_13_N | -3.825 | 1.232 | 0.844 | -3.368 | 2.394 | 1.289 | 1.33 | 70.38 | 68.53 |
| 73 | 2jsa-5 | VAL_A_13_N | VAL_A_13_H | -3.368 | 2.394 | 1.289 | -2.412 | 2.668 | 1.109 | 1.01 | 100.26 | 15.99 |
| 74 | 2jsa-5 | VAL_A_13_N | VAL_A_13_CA | -3.368 | 2.394 | 1.289 | -4.224 | 3.295 | 2.041 | 1.45 | 58.82 | 133.53 |
| 75 | 2jsa-5 | VAL_A_13_CA | VAL_A_13_HA | -4.224 | 3.295 | 2.041 | -4.621 | 2.755 | 2.901 | 1.09 | 37.93 | -126.32 |
| 76 | 2jsa-5 | VAL_A_13_CA | VAL_A_13_C | -4.224 | 3.295 | 2.041 | -5.401 | 3.724 | 1.166 | 1.53 | 124.93 | 159.97 |
| 77 | 2jsa-5 | VAL_A_13_C | VAL_A_13_O | -5.401 | 3.724 | 1.166 | -6.557 | 3.486 | 1.511 | 1.23 | 73.71 | -168.37 |
| 78 | 2jsa-5 | VAL_A_13_C | LYS_A_14_N | -5.401 | 3.724 | 1.166 | -5.068 | 4.35 | 0.046 | 1.33 | 147.66 | 61.99 |
| 79 | 2jsa-5 | LYS_A_14_N | LYS_A_14_H | -5.068 | 4.35 | 0.046 | -4.096 | 4.512 | -0.176 | 1.01 | 102.70 | 9.46 |
| 80 | 2jsa-5 | LYS_A_14_N | LYS_A_14_CA | -5.068 | 4.35 | 0.046 | -6.084 | 4.815 | -0.883 | 1.45 | 129.74 | 155.41 |
| 81 | 2jsa-5 | LYS_A_14_CA | LYS_A_14_HA | -6.084 | 4.815 | -0.883 | -6.587 | 5.671 | -0.432 | 1.09 | 65.57 | 120.44 |
| 82 | 2jsa-5 | LYS_A_14_CA | LYS_A_14_C | -6.084 | 4.815 | -0.883 | -7.128 | 3.714 | -1.084 | 1.53 | 97.55 | -133.48 |
| 83 | 2jsa-5 | LYS_A_14_C | LYS_A_14_O | -7.128 | 3.714 | -1.084 | -8.303 | 4.001 | -1.302 | 1.23 | 100.22 | 166.27 |
| 84 | 2jsa-5 | LYS_A_14_C | LYS_A_15_N | -7.128 | 3.714 | -1.084 | -6.659 | 2.477 | -1.002 | 1.33 | 86.45 | -69.24 |
| 85 | 2jsa-5 | LYS_A_15_N | LYS_A_15_H | -6.659 | 2.477 | -1.002 | -5.679 | 2.316 | -0.819 | 1.01 | 79.56 | -9.33 |
| 86 | 2jsa-5 | LYS_A_15_N | LYS_A_15_CA | -6.659 | 2.477 | -1.002 | -7.537 | 1.332 | -1.172 | 1.45 | 96.72 | -127.48 |
| 87 | 2jsa-5 | LYS_A_15_CA | LYS_A_15_HA | -7.537 | 1.332 | -1.172 | -8.247 | 1.566 | -1.964 | 1.09 | 136.65 | 161.76 |
| 88 | 2jsa-5 | LYS_A_15_CA | LYS_A_15_C | -7.537 | 1.332 | -1.172 | -8.333 | 1.11 | 0.116 | 1.53 | 32.68 | -164.42 |
| 89 | 2jsa-5 | LYS_A_15_C | LYS_A_15_O | -8.333 | 1.11 | 0.116 | -9.523 | 0.802 | 0.068 | 1.23 | 92.24 | -165.49 |
| 90 | 2jsa-5 | LYS_A_15_C | TYR_A_16_N | -8.333 | 1.11 | 0.116 | -7.644 | 1.275 | 1.235 | 1.32 | 32.34 | 13.47 |
| 91 | 2jsa-5 | TYR_A_16_N | TYR_A_16_H | -7.644 | 1.275 | 1.235 | -6.666 | 1.528 | 1.198 | 1.01 | 92.10 | 14.50 |
| 92 | 2jsa-5 | TYR_A_16_N | TYR_A_16_CA | -7.644 | 1.275 | 1.235 | -8.272 | 1.096 | 2.534 | 1.45 | 26.69 | -164.09 |
| 93 | 2jsa-5 | TYR_A_16_CA | TYR_A_16_HA | -8.272 | 1.096 | 2.534 | -8.897 | 0.203 | 2.532 | 1.09 | 90.11 | -124.99 |
| 94 | 2jsa-5 | TYR_A_16_CA | TYR_A_16_C | -8.272 | 1.096 | 2.534 | -9.113 | 2.318 | 2.908 | 1.53 | 75.85 | 124.54 |
| 95 | 2jsa-5 | TYR_A_16_C | TYR_A_16_O | -9.113 | 2.318 | 2.908 | -10.341 | 2.253 | 2.909 | 1.23 | 89.95 | -176.97 |
| 96 | 2jsa-5 | TYR_A_16_C | GLU_A_17_N | -9.113 | 2.318 | 2.908 | -8.419 | 3.403 | 3.219 | 1.32 | 76.42 | 57.40 |
| 97 | 2jsa-5 | GLU_A_17_N | GLU_A_17_H | -8.419 | 3.403 | 3.219 | -7.409 | 3.384 | 3.197 | 1.01 | 91.25 | -1.08 |
| 98 | 2jsa-5 | GLU_A_17_N | GLU_A_17_CA | -8.419 | 3.403 | 3.219 | -9.086 | 4.637 | 3.595 | 1.45 | 74.99 | 118.39 |
| 99 | 2jsa-5 | GLU_A_17_CA | GLU_A_17_HA | -9.086 | 4.637 | 3.595 | -9.427 | 4.536 | 4.625 | 1.09 | 19.05 | -163.50 |
| 100 | 2jsa-5 | GLU_A_17_CA | GLU_A_17_C | -9.086 | 4.637 | 3.595 | -10.309 | 4.872 | 2.704 | 1.53 | 125.58 | 169.12 |
| 101 | 2jsa-5 | GLU_A_17_C | GLU_A_17_O | -10.309 | 4.872 | 2.704 | -11.333 | 5.37 | 3.168 | 1.23 | 67.83 | 154.07 |
| 102 | 2jsa-5 | GLU_A_17_C | GLY_A_18_N | -10.309 | 4.872 | 2.704 | -10.16 | 4.502 | 1.44 | 1.33 | 162.49 | -68.07 |
| 103 | 2jsa-5 | GLY_A_18_N | GLY_A_18_H | -10.16 | 4.502 | 1.44 | -9.288 | 4.099 | 1.127 | 1.01 | 108.05 | -24.80 |
| 104 | 2jsa-5 | GLY_A_18_N | GLY_A_18_CA | -10.16 | 4.502 | 1.44 | -11.239 | 4.666 | 0.481 | 1.45 | 131.31 | 171.36 |
| 105 | 2jsa-5 | GLY_A_18_CA | GLY_A_18_C | -11.239 | 4.666 | 0.481 | -11.682 | 3.314 | -0.082 | 1.53 | 111.59 | -108.14 |
| 106 | 2jsa-5 | GLY_A_18_C | GLY_A_18_O | -11.682 | 3.314 | -0.082 | -11.847 | 3.165 | -1.291 | 1.23 | 169.58 | -137.92 |
| 107 | 2jsa-5 | GLY_A_18_C | SER_A_19_N | -11.682 | 3.314 | -0.082 | -11.863 | 2.363 | 0.823 | 1.33 | 46.93 | -100.78 |

| | | | | | | | | | | | | |
|---|---|---|---|---|---|---|---|---|---|---|---|---|
| 108 | 2jsa-5 | SER_A_19_N | SER_A_19_H | -11.863 | 2.363 | 0.823 | -11.709 | 2.557 | 1.802 | 1.01 | 14.20 | 51.56 |
| 109 | 2jsa-5 | SER_A_19_N | SER_A_19_CA | -11.863 | 2.363 | 0.823 | -12.285 | 1.028 | 0.432 | 1.45 | 105.60 | -107.54 |
| 110 | 2jsa-5 | SER_A_19_CA | SER_A_19_HA | -12.285 | 1.028 | 0.432 | -11.978 | 0.861 | -0.6 | 1.09 | 161.29 | -28.55 |
| 111 | 2jsa-5 | SER_A_19_CA | SER_A_19_C | -12.285 | 1.028 | 0.432 | -13.808 | 0.916 | 0.515 | 1.53 | 86.89 | -175.79 |
| 112 | 2jsa-5 | SER_A_19_C | SER_A_19_O | -13.808 | 0.916 | 0.515 | -14.455 | 0.489 | -0.44 | 1.23 | 140.93 | -146.58 |
| 113 | 2jsa-5 | SER_A_19_C | ALA_A_20_N | -13.808 | 0.916 | 0.515 | -14.337 | 1.305 | 1.666 | 1.33 | 29.70 | 143.67 |
| 114 | 2jsa-5 | ALA_A_20_N | ALA_A_20_H | -14.337 | 1.305 | 1.666 | -13.743 | 1.644 | 2.41 | 1.01 | 42.59 | 29.71 |
| 115 | 2jsa-5 | ALA_A_20_N | ALA_A_20_CA | -14.337 | 1.305 | 1.666 | -15.773 | 1.253 | 1.886 | 1.45 | 81.30 | -177.93 |
| 116 | 2jsa-5 | ALA_A_20_CA | ALA_A_20_HA | -15.773 | 1.253 | 1.886 | -16.018 | 1.871 | 2.75 | 1.09 | 37.58 | 111.63 |
| 117 | 2jsa-5 | ALA_A_20_CA | ALA_A_20_C | -15.773 | 1.253 | 1.886 | -16.185 | -0.189 | 2.189 | 1.53 | 78.58 | -105.95 |
| 118 | 2jsa-5 | ALA_A_20_C | ALA_A_20_O | -16.185 | -0.189 | 2.189 | -17.364 | -0.47 | 2.398 | 1.23 | 80.22 | -166.59 |
| 119 | 2jsa-5 | ALA_A_20_C | ASP_A_21_N | -16.185 | -0.189 | 2.189 | -15.191 | -1.065 | 2.204 | 1.33 | 89.35 | -41.39 |
| 120 | 2jsa-5 | ASP_A_21_N | ASP_A_21_H | -15.191 | -1.065 | 2.204 | -14.244 | -0.763 | 2.023 | 1.01 | 100.32 | 17.69 |
| 121 | 2jsa-5 | ASP_A_21_N | ASP_A_21_CA | -15.191 | -1.065 | 2.204 | -15.436 | -2.47 | 2.478 | 1.45 | 79.12 | -99.89 |
| 122 | 2jsa-5 | ASP_A_21_CA | ASP_A_21_HA | -15.436 | -2.47 | 2.478 | -16.511 | -2.645 | 2.505 | 1.09 | 88.58 | -170.75 |
| 123 | 2jsa-5 | ASP_A_21_CA | ASP_A_21_C | -15.436 | -2.47 | 2.478 | -14.816 | -2.838 | 3.828 | 1.53 | 28.11 | -30.69 |
| 124 | 2jsa-5 | ASP_A_21_C | ASP_A_21_O | -14.816 | -2.838 | 3.828 | -15.078 | -3.915 | 4.362 | 1.23 | 64.28 | -103.67 |
| 125 | 2jsa-5 | ASP_A_21_C | LYS_A_22_N | -14.816 | -2.838 | 3.828 | -14.006 | -1.923 | 4.339 | 1.32 | 67.31 | 48.48 |
| 126 | 2jsa-5 | LYS_A_22_N | LYS_A_22_H | -14.006 | -1.923 | 4.339 | -13.836 | -1.059 | 3.844 | 1.01 | 119.34 | 78.87 |
| 127 | 2jsa-5 | LYS_A_22_N | LYS_A_22_CA | -14.006 | -1.923 | 4.339 | -13.347 | -2.139 | 5.617 | 1.45 | 28.49 | -18.15 |
| 128 | 2jsa-5 | LYS_A_22_CA | LYS_A_22_HA | -13.347 | -2.139 | 5.617 | -12.507 | -1.447 | 5.685 | 1.09 | 86.42 | 39.48 |
| 129 | 2jsa-5 | LYS_A_22_CA | LYS_A_22_C | -13.347 | -2.139 | 5.617 | -12.782 | -3.559 | 5.661 | 1.53 | 88.35 | -68.30 |
| 130 | 2jsa-5 | LYS_A_22_C | LYS_A_22_O | -12.782 | -3.559 | 5.661 | -13.02 | -4.295 | 6.617 | 1.23 | 38.98 | -107.92 |
| 131 | 2jsa-5 | LYS_A_22_C | ASP_A_23_N | -12.782 | -3.559 | 5.661 | -12.046 | -3.902 | 4.615 | 1.32 | 142.18 | -24.99 |
| 132 | 2jsa-5 | ASP_A_23_N | ASP_A_23_H | -12.046 | -3.902 | 4.615 | -11.892 | -3.245 | 3.863 | 1.01 | 138.10 | 76.81 |
| 133 | 2jsa-5 | ASP_A_23_N | ASP_A_23_CA | -12.046 | -3.902 | 4.615 | -11.446 | -5.223 | 4.523 | 1.45 | 93.63 | -65.57 |
| 134 | 2jsa-5 | ASP_A_23_CA | ASP_A_23_HA | -11.446 | -5.223 | 4.523 | -11.474 | -5.696 | 5.505 | 1.09 | 25.76 | -93.39 |
| 135 | 2jsa-5 | ASP_A_23_CA | ASP_A_23_C | -11.446 | -5.223 | 4.523 | -9.997 | -5.088 | 4.052 | 1.53 | 107.93 | 5.32 |
| 136 | 2jsa-5 | ASP_A_23_C | ASP_A_23_O | -9.997 | -5.088 | 4.052 | -9.744 | -4.784 | 2.887 | 1.23 | 161.25 | 50.23 |
| 137 | 2jsa-5 | ASP_A_23_C | ALA_A_24_N | -9.997 | -5.088 | 4.052 | -9.081 | -5.32 | 4.982 | 1.33 | 45.46 | -14.21 |
| 138 | 2jsa-5 | ALA_A_24_N | ALA_A_24_H | -9.081 | -5.32 | 4.982 | -9.36 | -5.565 | 5.921 | 1.01 | 21.58 | -138.71 |
| 139 | 2jsa-5 | ALA_A_24_N | ALA_A_24_CA | -9.081 | -5.32 | 4.982 | -7.663 | -5.227 | 4.676 | 1.45 | 102.15 | 3.75 |
| 140 | 2jsa-5 | ALA_A_24_CA | ALA_A_24_HA | -7.663 | -5.227 | 4.676 | -7.46 | -4.246 | 4.245 | 1.09 | 113.28 | 78.31 |
| 141 | 2jsa-5 | ALA_A_24_CA | ALA_A_24_C | -7.663 | -5.227 | 4.676 | -7.299 | -6.296 | 3.642 | 1.53 | 132.48 | -71.20 |
| 142 | 2jsa-5 | ALA_A_24_C | ALA_A_24_O | -7.299 | -6.296 | 3.642 | -6.2 | -6.279 | 3.09 | 1.23 | 116.67 | 0.89 |
| 143 | 2jsa-5 | ALA_A_24_C | ASN_A_25_N | -7.299 | -6.296 | 3.642 | -8.242 | -7.198 | 3.413 | 1.32 | 99.95 | -136.27 |
| 144 | 2jsa-5 | ASN_A_25_N | ASN_A_25_H | -8.242 | -7.198 | 3.413 | -9.123 | -7.149 | 3.905 | 1.01 | 60.86 | 176.82 |
| 145 | 2jsa-5 | ASN_A_25_N | ASN_A_25_CA | -8.242 | -7.198 | 3.413 | -8.033 | -8.271 | 2.456 | 1.45 | 131.20 | -78.98 |
| 146 | 2jsa-5 | ASN_A_25_CA | ASN_A_25_HA | -8.033 | -8.271 | 2.456 | -7.006 | -8.627 | 2.538 | 1.09 | 85.69 | -19.12 |
| 147 | 2jsa-5 | ASN_A_25_CA | ASN_A_25_C | -8.033 | -8.271 | 2.456 | -8.289 | -7.745 | 1.043 | 1.53 | 157.51 | 115.95 |
| 148 | 2jsa-5 | ASN_A_25_C | ASN_A_25_O | -8.289 | -7.745 | 1.043 | -7.467 | -7.932 | 0.148 | 1.23 | 136.71 | -12.82 |
| 149 | 2jsa-5 | ASN_A_25_C | VAL_A_26_N | -8.289 | -7.745 | 1.043 | -9.434 | -7.096 | 0.885 | 1.33 | 96.85 | 150.45 |
| 150 | 2jsa-5 | VAL_A_26_N | VAL_A_26_H | -9.434 | -7.096 | 0.885 | -10.064 | -6.979 | 1.666 | 1.01 | 39.37 | 169.48 |
| 151 | 2jsa-5 | VAL_A_26_N | VAL_A_26_CA | -9.434 | -7.096 | 0.885 | -9.808 | -6.541 | -0.405 | 1.45 | 152.58 | 123.98 |
| 152 | 2jsa-5 | VAL_A_26_CA | VAL_A_26_HA | -9.808 | -6.541 | -0.405 | -9.537 | -7.258 | -1.18 | 1.09 | 135.32 | -69.30 |
| 153 | 2jsa-5 | VAL_A_26_CA | VAL_A_26_C | -9.808 | -6.541 | -0.405 | -9.018 | -5.255 | -0.651 | 1.53 | 99.26 | 58.44 |
| 154 | 2jsa-5 | VAL_A_26_C | VAL_A_26_O | -9.018 | -5.255 | -0.651 | -9.067 | -4.691 | -1.743 | 1.23 | 152.60 | 94.97 |
| 155 | 2jsa-5 | VAL_A_26_C | ILE_A_27_N | -9.018 | -5.255 | -0.651 | -8.305 | -4.83 | 0.383 | 1.33 | 38.76 | 30.80 |
| 156 | 2jsa-5 | ILE_A_27_N | ILE_A_27_H | -8.305 | -4.83 | 0.383 | -8.309 | -5.346 | 1.251 | 1.01 | 30.73 | -90.44 |
| 157 | 2jsa-5 | ILE_A_27_N | ILE_A_27_CA | -8.305 | -4.83 | 0.383 | -7.505 | -3.621 | 0.292 | 1.45 | 93.59 | 56.51 |
| 158 | 2jsa-5 | ILE_A_27_CA | ILE_A_27_HA | -7.505 | -3.621 | 0.292 | -7.969 | -2.964 | -0.443 | 1.09 | 132.42 | 125.23 |
| 159 | 2jsa-5 | ILE_A_27_CA | ILE_A_27_C | -7.505 | -3.621 | 0.292 | -6.108 | -3.976 | -0.22 | 1.53 | 109.56 | -14.26 |
| 160 | 2jsa-5 | ILE_A_27_C | ILE_A_27_O | -6.108 | -3.976 | -0.22 | -5.72 | -3.567 | -1.313 | 1.23 | 152.72 | 46.51 |
| 161 | 2jsa-5 | ILE_A_27_C | LYS_A_28_N | -6.108 | -3.976 | -0.22 | -5.389 | -4.735 | 0.596 | 1.33 | 52.03 | -46.55 |
| 162 | 2jsa-5 | LYS_A_28_N | LYS_A_28_H | -5.389 | -4.735 | 0.596 | -5.771 | -5.035 | 1.481 | 1.01 | 28.76 | -141.86 |
| 163 | 2jsa-5 | LYS_A_28_N | LYS_A_28_CA | -5.389 | -4.735 | 0.596 | -4.044 | -5.15 | 0.239 | 1.45 | 104.23 | -17.15 |
| 164 | 2jsa-5 | LYS_A_28_CA | LYS_A_28_HA | -4.044 | -5.15 | 0.239 | -3.416 | -4.26 | 0.197 | 1.09 | 92.21 | 54.79 |
| 165 | 2jsa-5 | LYS_A_28_CA | LYS_A_28_C | -4.044 | -5.15 | 0.239 | -4.06 | -5.774 | -1.158 | 1.53 | 155.92 | -91.47 |
| 166 | 2jsa-5 | LYS_A_28_C | LYS_A_28_O | -4.06 | -5.774 | -1.158 | -3.152 | -5.543 | -1.955 | 1.23 | 130.39 | 14.27 |
| 167 | 2jsa-5 | LYS_A_28_C | LYS_A_29_N | -4.06 | -5.774 | -1.158 | -5.103 | -6.552 | -1.412 | 1.33 | 101.05 | -143.28 |
| 168 | 2jsa-5 | LYS_A_29_N | LYS_A_29_H | -5.103 | -6.552 | -1.412 | -5.812 | -6.697 | -0.708 | 1.01 | 45.79 | -168.44 |
| 169 | 2jsa-5 | LYS_A_29_N | LYS_A_29_CA | -5.103 | -6.552 | -1.412 | -5.249 | -7.21 | -2.699 | 1.45 | 152.36 | -102.51 |
| 170 | 2jsa-5 | LYS_A_29_CA | LYS_A_29_HA | -5.249 | -7.21 | -2.699 | -4.321 | -7.741 | -2.914 | 1.09 | 101.37 | -29.78 |
| 171 | 2jsa-5 | LYS_A_29_CA | LYS_A_29_C | -5.249 | -7.21 | -2.699 | -5.443 | -6.154 | -3.789 | 1.53 | 135.43 | 100.41 |
| 172 | 2jsa-5 | LYS_A_29_C | LYS_A_29_O | -5.443 | -6.154 | -3.789 | -4.793 | -6.206 | -4.831 | 1.23 | 147.96 | -4.57 |
| 173 | 2jsa-5 | LYS_A_29_C | ASP_A_30_N | -5.443 | -6.154 | -3.789 | -6.341 | -5.22 | -3.509 | 1.33 | 77.81 | 133.87 |
| 174 | 2jsa-5 | ASP_A_30_N | ASP_A_30_H | -6.341 | -5.22 | -3.509 | -6.84 | -5.24 | -2.631 | 1.01 | 29.63 | -177.70 |
| 175 | 2jsa-5 | ASP_A_30_N | ASP_A_30_CA | -6.341 | -5.22 | -3.509 | -6.628 | -4.153 | -4.452 | 1.45 | 130.48 | 105.06 |
| 176 | 2jsa-5 | ASP_A_30_CA | ASP_A_30_HA | -6.628 | -4.153 | -4.452 | -7.164 | -4.567 | -5.306 | 1.09 | 141.58 | -142.32 |
| 177 | 2jsa-5 | ASP_A_30_CA | ASP_A_30_C | -6.628 | -4.153 | -4.452 | -5.314 | -3.523 | -4.919 | 1.53 | 107.77 | 25.62 |
| 178 | 2jsa-5 | ASP_A_30_C | ASP_A_30_O | -5.314 | -3.523 | -4.919 | -5.071 | -3.404 | -6.119 | 1.23 | 167.29 | 26.09 |
| 179 | 2jsa-5 | ASP_A_30_C | PHE_A_31_N | -5.314 | -3.523 | -4.919 | -4.502 | -3.135 | -3.947 | 1.32 | 42.80 | 25.54 |
| 180 | 2jsa-5 | PHE_A_31_N | PHE_A_31_H | -4.502 | -3.135 | -3.947 | -4.768 | -3.263 | -2.981 | 1.01 | 16.99 | -154.30 |
| 181 | 2jsa-5 | PHE_A_31_N | PHE_A_31_CA | -4.502 | -3.135 | -3.947 | -3.219 | -2.521 | -4.243 | 1.45 | 101.76 | 25.57 |
| 182 | 2jsa-5 | PHE_A_31_CA | PHE_A_31_HA | -3.219 | -2.521 | -4.243 | -3.366 | -1.485 | -4.549 | 1.09 | 106.30 | 98.08 |
| 183 | 2jsa-5 | PHE_A_31_CA | PHE_A_31_C | -3.219 | -2.521 | -4.243 | -2.487 | -3.285 | -5.349 | 1.53 | 136.27 | -46.23 |
| 184 | 2jsa-5 | PHE_A_31_C | PHE_A_31_O | -2.487 | -3.285 | -5.349 | -2.126 | -2.707 | -6.374 | 1.23 | 146.38 | 58.01 |
| 185 | 2jsa-5 | PHE_A_31_C | ASP_A_32_N | -2.487 | -3.285 | -5.349 | -2.291 | -4.572 | -5.104 | 1.32 | 79.34 | -81.34 |
| 186 | 2jsa-5 | ASP_A_32_N | ASP_A_32_H | -2.291 | -4.572 | -5.104 | -2.614 | -4.98 | -4.239 | 1.01 | 31.03 | -128.37 |
| 187 | 2jsa-5 | ASP_A_32_N | ASP_A_32_CA | -2.291 | -4.572 | -5.104 | -1.61 | -5.421 | -6.066 | 1.45 | 131.47 | -51.27 |
| 188 | 2jsa-5 | ASP_A_32_CA | ASP_A_32_HA | -1.61 | -5.421 | -6.066 | -0.604 | -5.035 | -6.232 | 1.09 | 98.76 | 20.99 |
| 189 | 2jsa-5 | ASP_A_32_CA | ASP_A_32_C | -1.61 | -5.421 | -6.066 | -2.39 | -5.423 | -7.383 | 1.53 | 149.36 | -179.85 |
| 190 | 2jsa-5 | ASP_A_32_C | ASP_A_32_O | -2.39 | -5.423 | -7.383 | -1.828 | -5.703 | -8.44 | 1.23 | 149.29 | -26.48 |
| 191 | 2jsa-5 | ASP_A_32_C | ALA_A_33_N | -2.39 | -5.423 | -7.383 | -3.672 | -5.106 | -7.274 | 1.33 | 85.28 | 166.11 |
| 192 | 2jsa-5 | ALA_A_33_N | ALA_A_33_H | -3.672 | -5.106 | -7.274 | -4.067 | -4.888 | -6.37 | 1.01 | 26.52 | 151.11 |
| 193 | 2jsa-5 | ALA_A_33_N | ALA_A_33_CA | -3.672 | -5.106 | -7.274 | -4.535 | -5.066 | -8.443 | 1.45 | 143.53 | 177.35 |
| 194 | 2jsa-5 | ALA_A_33_CA | ALA_A_33_HA | -4.535 | -5.066 | -8.443 | -4.374 | -5.971 | -9.029 | 1.09 | 122.52 | -79.91 |
| 195 | 2jsa-5 | ALA_A_33_CA | ALA_A_33_C | -4.535 | -5.066 | -8.443 | -4.162 | -3.858 | -9.306 | 1.53 | 124.32 | 72.84 |
| 196 | 2jsa-5 | ALA_A_33_C | ALA_A_33_O | -4.162 | -3.858 | -9.306 | -3.727 | -4.017 | -10.446 | 1.23 | 157.89 | -20.08 |
| 197 | 2jsa-5 | ALA_A_33_C | GLU_A_34_N | -4.162 | -3.858 | -9.306 | -4.346 | -2.68 | -8.73 | 1.32 | 64.21 | 98.88 |
| 198 | 2jsa-5 | GLU_A_34_N | GLU_A_34_H | -4.346 | -2.68 | -8.73 | -4.707 | -2.626 | -7.788 | 1.01 | 21.18 | 171.49 |
| 199 | 2jsa-5 | GLU_A_34_N | GLU_A_34_CA | -4.346 | -2.68 | -8.73 | -4.035 | -1.447 | -9.431 | 1.45 | 118.87 | 75.84 |
| 200 | 2jsa-5 | GLU_A_34_CA | GLU_A_34_HA | -4.035 | -1.447 | -9.431 | -4.676 | -1.388 | -10.31 | 1.09 | 143.78 | 174.74 |
| 201 | 2jsa-5 | GLU_A_34_CA | GLU_A_34_C | -4.035 | -1.447 | -9.431 | -2.579 | -1.456 | -9.899 | 1.53 | 107.82 | -0.35 |
| 202 | 2jsa-5 | GLU_A_34_C | GLU_A_34_O | -2.579 | -1.456 | -9.899 | -2.29 | -1.13 | -11.051 | 1.23 | 159.28 | 48.44 |
| 203 | 2jsa-5 | GLU_A_34_C | CYS_A_35_N | -2.579 | -1.456 | -9.899 | -1.698 | -1.832 | -8.983 | 1.33 | 46.28 | -23.11 |

| | | | | | | | | | | | |
|---|---|---|---|---|---|---|---|---|---|---|
| 204 | 2jsa-5 | CYS_A_35_N | CYS_A_35_H | -1.698 | -1.832 | -8.983 | -2.005 | -2.087 | -8.055 | 1.01 | 23.27 | -140.29 |
| 205 | 2jsa-5 | CYS_A_35_N | CYS_A_35_CA | -1.698 | -1.832 | -8.983 | -0.278 | -1.887 | -9.289 | 1.45 | 102.15 | -2.22 |
| 206 | 2jsa-5 | CYS_A_35_CA | CYS_A_35_HA | -0.278 | -1.887 | -9.289 | 0.046 | -0.892 | -9.596 | 1.09 | 106.35 | 71.96 |
| 207 | 2jsa-5 | CYS_A_35_CA | CYS_A_35_C | -0.278 | -1.887 | -9.289 | -0.076 | -2.849 | -10.461 | 1.53 | 140.01 | -78.14 |
| 208 | 2jsa-5 | CYS_A_35_C | CYS_A_35_O | -0.076 | -2.849 | -10.461 | 0.617 | -2.523 | -11.424 | 1.23 | 141.51 | 25.19 |
| 209 | 2jsa-5 | CYS_A_35_C | LYS_A_36_N | -0.076 | -2.849 | -10.461 | -0.694 | -4.016 | -10.342 | 1.33 | 84.85 | -117.90 |
| 210 | 2jsa-5 | LYS_A_36_N | LYS_A_36_H | -0.694 | -4.016 | -10.342 | -1.248 | -4.217 | -9.522 | 1.01 | 35.70 | -160.06 |
| 211 | 2jsa-5 | LYS_A_36_N | LYS_A_36_CA | -0.694 | -4.016 | -10.342 | -0.59 | -5.026 | -11.38 | 1.45 | 135.63 | -84.12 |
| 212 | 2jsa-5 | LYS_A_36_CA | LYS_A_36_HA | -0.59 | -5.026 | -11.38 | 0.392 | -5.493 | -11.3 | 1.09 | 85.79 | -25.43 |
| 213 | 2jsa-5 | LYS_A_36_CA | LYS_A_36_C | -0.59 | -5.026 | -11.38 | -0.674 | -4.353 | -12.752 | 1.53 | 153.70 | 97.11 |
| 214 | 2jsa-5 | LYS_A_36_C | LYS_A_36_O | -0.674 | -4.353 | -12.752 | 0.014 | -4.759 | -13.688 | 1.23 | 139.52 | -30.55 |
| 215 | 2jsa-5 | LYS_A_36_C | LYS_A_37_N | -0.674 | -4.353 | -12.752 | -1.521 | -3.338 | -12.826 | 1.32 | 93.20 | 129.84 |
| 216 | 2jsa-5 | LYS_A_37_N | LYS_A_37_H | -1.521 | -3.338 | -12.826 | -2.054 | -3.06 | -12.014 | 1.01 | 36.51 | 152.45 |
| 217 | 2jsa-5 | LYS_A_37_N | LYS_A_37_CA | -1.521 | -3.338 | -12.826 | -1.703 | -2.605 | -14.068 | 1.45 | 148.70 | 103.94 |
| 218 | 2jsa-5 | LYS_A_37_CA | LYS_A_37_HA | -1.703 | -2.605 | -14.068 | -1.39 | -3.25 | -14.889 | 1.09 | 138.87 | -64.11 |
| 219 | 2jsa-5 | LYS_A_37_CA | LYS_A_37_C | -1.703 | -2.605 | -14.068 | -0.791 | -1.376 | -14.069 | 1.53 | 90.04 | 53.42 |
| 220 | 2jsa-5 | LYS_A_37_C | LYS_A_37_O | -0.791 | -1.376 | -14.069 | 0.001 | -1.186 | -14.991 | 1.23 | 138.54 | 13.49 |
| 221 | 2jsa-5 | LYS_A_37_C | LEU_A_38_N | -0.791 | -1.376 | -14.069 | -0.933 | -0.574 | -13.024 | 1.32 | 37.93 | 100.04 |
| 222 | 2jsa-5 | LEU_A_38_N | LEU_A_38_H | -0.933 | -0.574 | -13.024 | -1.604 | -0.792 | -12.302 | 1.01 | 44.34 | -162.00 |
| 223 | 2jsa-5 | LEU_A_38_N | LEU_A_38_CA | -0.933 | -0.574 | -13.024 | -0.131 | 0.631 | -12.892 | 1.45 | 84.79 | 56.35 |
| 224 | 2jsa-5 | LEU_A_38_CA | LEU_A_38_HA | -0.131 | 0.631 | -12.892 | -0.478 | 1.35 | -13.634 | 1.09 | 132.90 | 115.76 |
| 225 | 2jsa-5 | LEU_A_38_CA | LEU_A_38_C | -0.131 | 0.631 | -12.892 | 1.329 | 0.305 | -13.209 | 1.53 | 101.96 | -12.59 |
| 226 | 2jsa-5 | LEU_A_38_C | LEU_A_38_O | 1.329 | 0.305 | -13.209 | 1.98 | 1.025 | -13.965 | 1.23 | 127.91 | 47.88 |
| 227 | 2jsa-5 | LEU_A_38_C | PHE_A_39_N | 1.329 | 0.305 | -13.209 | 1.802 | -0.782 | -12.617 | 1.33 | 63.46 | -66.48 |
| 228 | 2jsa-5 | PHE_A_39_N | PHE_A_39_H | 1.802 | -0.782 | -12.617 | 1.21 | -1.328 | -12.008 | 1.01 | 52.90 | -137.31 |
| 229 | 2jsa-5 | PHE_A_39_N | PHE_A_39_CA | 1.802 | -0.782 | -12.617 | 3.174 | -1.212 | -12.828 | 1.45 | 98.35 | -17.40 |
| 230 | 2jsa-5 | PHE_A_39_CA | PHE_A_39_HA | 3.174 | -1.212 | -12.828 | 3.698 | -0.488 | -13.452 | 1.09 | 124.92 | 54.10 |
| 231 | 2jsa-5 | PHE_A_39_CA | PHE_A_39_C | 3.174 | -1.212 | -12.828 | 3.22 | -2.591 | -13.49 | 1.53 | 115.63 | -88.09 |
| 232 | 2jsa-5 | PHE_A_39_C | PHE_A_39_O | 3.22 | -2.591 | -13.49 | 4.019 | -3.443 | -13.102 | 1.23 | 71.62 | -46.84 |
| 233 | 2jsa-5 | PHE_A_39_C | HIS_A_40_N | 3.22 | -2.591 | -13.49 | 2.353 | -2.768 | -14.477 | 1.33 | 138.12 | -168.46 |
| 234 | 2jsa-5 | HIS_A_40_N | HIS_A_40_H | 2.353 | -2.768 | -14.477 | 1.725 | -2.023 | -14.742 | 1.01 | 105.21 | 130.13 |
| 235 | 2jsa-5 | HIS_A_40_N | HIS_A_40_CA | 2.353 | -2.768 | -14.477 | 2.286 | -4.029 | -15.195 | 1.45 | 119.62 | -93.04 |
| 236 | 2jsa-5 | HIS_A_40_CA | HIS_A_40_HA | 2.286 | -4.029 | -15.195 | 2.237 | -4.833 | -14.46 | 1.09 | 47.62 | -93.49 |
| 237 | 2jsa-5 | HIS_A_40_CA | HIS_A_40_C | 2.286 | -4.029 | -15.195 | 3.562 | -4.221 | -16.016 | 1.53 | 122.47 | -8.56 |
| 238 | 2jsa-5 | HIS_A_40_C | HIS_A_40_O | 3.562 | -4.221 | -16.016 | 3.925 | -5.347 | -16.352 | 1.23 | 105.85 | -72.13 |
| 239 | 2jsa-5 | HIS_A_40_C | THR_A_41_N | 3.562 | -4.221 | -16.016 | 4.209 | -3.104 | -16.315 | 1.33 | 103.04 | 59.92 |
| 240 | 2jsa-5 | THR_A_41_N | THR_A_41_H | 4.209 | -3.104 | -16.315 | 3.853 | -2.211 | -16.004 | 1.01 | 72.07 | 111.74 |
| 241 | 2jsa-5 | THR_A_41_N | THR_A_41_CA | 4.209 | -3.104 | -16.315 | 5.437 | -3.135 | -17.09 | 1.45 | 122.25 | -1.45 |
| 242 | 2jsa-5 | THR_A_41_CA | THR_A_41_HA | 5.437 | -3.135 | -17.09 | 5.424 | -4.017 | -17.73 | 1.09 | 125.96 | -90.84 |
| 243 | 2jsa-5 | THR_A_41_CA | THR_A_41_C | 5.437 | -3.135 | -17.09 | 6.65 | -3.243 | -16.164 | 1.53 | 52.75 | -5.09 |
| 244 | 2jsa-5 | THR_A_41_C | THR_A_41_O | 6.65 | -3.243 | -16.164 | 7.562 | -4.03 | -16.418 | 1.23 | 101.91 | -40.79 |
| 245 | 2jsa-5 | THR_A_41_C | ILE_A_42_N | 6.65 | -3.243 | -16.164 | 6.622 | -2.442 | -15.109 | 1.32 | 37.22 | 92.00 |
| 246 | 2jsa-5 | ILE_A_42_N | ILE_A_42_H | 6.622 | -2.442 | -15.109 | 5.839 | -1.821 | -14.962 | 1.01 | 81.63 | 141.58 |
| 247 | 2jsa-5 | ILE_A_42_N | ILE_A_42_CA | 6.622 | -2.442 | -15.109 | 7.708 | -2.438 | -14.143 | 1.45 | 48.35 | 0.21 |
| 248 | 2jsa-5 | ILE_A_42_CA | ILE_A_42_HA | 7.708 | -2.438 | -14.143 | 8.535 | -1.869 | -14.567 | 1.09 | 112.90 | 34.53 |
| 249 | 2jsa-5 | ILE_A_42_CA | ILE_A_42_C | 7.708 | -2.438 | -14.143 | 8.195 | -3.871 | -13.924 | 1.53 | 81.77 | -71.23 |
| 250 | 2jsa-5 | ILE_A_42_C | ILE_A_42_O | 8.195 | -3.871 | -13.924 | 7.395 | -4.805 | -13.893 | 1.23 | 88.56 | -130.58 |
| 251 | 2jsa-5 | ILE_A_42_C | PRO_A_43_N | 8.195 | -3.871 | -13.924 | 9.54 | -4.005 | -13.772 | 1.36 | 83.58 | -5.69 |
| 252 | 2jsa-5 | PRO_A_43_N | PRO_A_43_CA | 9.54 | -4.005 | -13.772 | 10.143 | -5.309 | -13.557 | 1.45 | 81.49 | -65.18 |
| 253 | 2jsa-5 | PRO_A_43_CA | PRO_A_43_HA | 10.143 | -5.309 | -13.557 | 9.716 | -6.036 | -14.247 | 1.09 | 129.30 | -120.43 |
| 254 | 2jsa-5 | PRO_A_43_CA | PRO_A_43_C | 10.143 | -5.309 | -13.557 | 9.9 | -5.796 | -12.127 | 1.53 | 20.84 | -116.52 |
| 255 | 2jsa-5 | PRO_A_43_C | PRO_A_43_O | 9.9 | -5.796 | -12.127 | 9.437 | -6.917 | -11.919 | 1.23 | 80.27 | -112.44 |
| 256 | 2jsa-5 | PRO_A_43_C | PHE_A_44_N | 9.9 | -5.796 | -12.127 | 10.222 | -4.929 | -11.178 | 1.33 | 44.26 | 69.63 |
| 257 | 2jsa-5 | PHE_A_44_N | PHE_A_44_H | 10.222 | -4.929 | -11.178 | 10.598 | -4.024 | -11.422 | 1.01 | 103.98 | 67.44 |
| 258 | 2jsa-5 | PHE_A_44_N | PHE_A_44_CA | 10.222 | -4.929 | -11.178 | 10.044 | -5.257 | -9.773 | 1.45 | 14.88 | -118.49 |
| 259 | 2jsa-5 | PHE_A_44_CA | PHE_A_44_HA | 10.044 | -5.257 | -9.773 | 9.866 | -6.326 | -9.662 | 1.09 | 84.15 | -99.45 |
| 260 | 2jsa-5 | PHE_A_44_CA | PHE_A_44_C | 10.044 | -5.257 | -9.773 | 8.874 | -4.475 | -9.172 | 1.53 | 66.87 | 146.24 |
| 261 | 2jsa-5 | PHE_A_44_C | PHE_A_44_O | 8.874 | -4.475 | -9.172 | 8.905 | -4.113 | -7.996 | 1.23 | 17.17 | 85.11 |
| 262 | 2jsa-5 | PHE_A_44_C | GLY_A_45_N | 8.874 | -4.475 | -9.172 | 7.874 | -4.235 | -10.005 | 1.32 | 129.01 | 166.50 |
| 263 | 2jsa-5 | GLY_A_45_N | GLY_A_45_H | 7.874 | -4.235 | -10.005 | 7.919 | -4.561 | -10.96 | 1.01 | 160.99 | -82.14 |
| 264 | 2jsa-5 | GLY_A_45_N | GLY_A_45_CA | 7.874 | -4.235 | -10.005 | 6.696 | -3.502 | -9.57 | 1.45 | 72.59 | 148.11 |
| 265 | 2jsa-5 | GLY_A_45_CA | GLY_A_45_C | 6.696 | -3.502 | -9.57 | 5.622 | -4.454 | -9.04 | 1.53 | 69.73 | -138.45 |
| 266 | 2jsa-5 | GLY_A_45_C | GLY_A_45_O | 5.622 | -4.454 | -9.04 | 5.132 | -4.285 | -7.925 | 1.23 | 24.93 | 160.97 |
| 267 | 2jsa-5 | GLY_A_45_C | THR_A_46_N | 5.622 | -4.454 | -9.04 | 5.286 | -5.434 | -9.867 | 1.33 | 128.60 | -108.92 |
| 268 | 2jsa-5 | THR_A_46_N | THR_A_46_H | 5.286 | -5.434 | -9.867 | 5.731 | -5.512 | -10.771 | 1.01 | 153.45 | -9.94 |
| 269 | 2jsa-5 | THR_A_46_N | THR_A_46_CA | 5.286 | -5.434 | -9.867 | 4.278 | -6.413 | -9.496 | 1.45 | 75.21 | -135.84 |
| 270 | 2jsa-5 | THR_A_46_CA | THR_A_46_HA | 4.278 | -6.413 | -9.496 | 3.418 | -5.889 | -9.078 | 1.09 | 67.46 | 148.65 |
| 271 | 2jsa-5 | THR_A_46_CA | THR_A_46_C | 4.278 | -6.413 | -9.496 | 4.822 | -7.357 | -8.422 | 1.53 | 45.41 | -60.05 |
| 272 | 2jsa-5 | THR_A_46_C | THR_A_46_O | 4.822 | -7.357 | -8.422 | 4.073 | -7.831 | -7.57 | 1.23 | 46.13 | -147.67 |
| 273 | 2jsa-5 | THR_A_46_C | ARG_A_47_N | 4.822 | -7.357 | -8.422 | 6.122 | -7.602 | -8.498 | 1.33 | 93.29 | -10.67 |
| 274 | 2jsa-5 | ARG_A_47_N | ARG_A_47_H | 6.122 | -7.602 | -8.498 | 6.678 | -7.177 | -9.226 | 1.01 | 136.13 | 37.39 |
| 275 | 2jsa-5 | ARG_A_47_N | ARG_A_47_CA | 6.122 | -7.602 | -8.498 | 6.775 | -8.482 | -7.544 | 1.45 | 48.96 | -53.42 |
| 276 | 2jsa-5 | ARG_A_47_CA | ARG_A_47_HA | 6.775 | -8.482 | -7.544 | 6.207 | -9.41 | -7.487 | 1.09 | 87.00 | -121.47 |
| 277 | 2jsa-5 | ARG_A_47_CA | ARG_A_47_C | 6.775 | -8.482 | -7.544 | 6.807 | -7.831 | -6.159 | 1.53 | 25.20 | 87.19 |
| 278 | 2jsa-5 | ARG_A_47_C | ARG_A_47_O | 6.807 | -7.831 | -6.159 | 6.289 | -8.391 | -5.193 | 1.23 | 38.30 | -132.77 |
| 279 | 2jsa-5 | ARG_A_47_C | GLU_A_48_N | 6.807 | -7.831 | -6.159 | 7.421 | -6.658 | -6.105 | 1.33 | 87.66 | 62.37 |
| 280 | 2jsa-5 | GLU_A_48_N | GLU_A_48_H | 7.421 | -6.658 | -6.105 | 7.824 | -6.256 | -6.939 | 1.01 | 145.69 | 44.93 |
| 281 | 2jsa-5 | GLU_A_48_N | GLU_A_48_CA | 7.421 | -6.658 | -6.105 | 7.528 | -5.926 | -4.856 | 1.45 | 30.64 | 81.68 |
| 282 | 2jsa-5 | GLU_A_48_CA | GLU_A_48_HA | 7.528 | -5.926 | -4.856 | 8.012 | -6.574 | -4.125 | 1.09 | 47.89 | -53.24 |
| 283 | 2jsa-5 | GLU_A_48_CA | GLU_A_48_C | 7.528 | -5.926 | -4.856 | 6.137 | -5.565 | -4.332 | 1.53 | 69.97 | 165.45 |
| 284 | 2jsa-5 | GLU_A_48_C | GLU_A_48_O | 6.137 | -5.565 | -4.332 | 5.927 | -5.481 | -3.122 | 1.23 | 10.59 | 158.20 |
| 285 | 2jsa-5 | GLU_A_48_C | CYS_A_49_N | 6.137 | -5.565 | -4.332 | 5.222 | -5.359 | -5.268 | 1.33 | 134.94 | 167.31 |
| 286 | 2jsa-5 | CYS_A_49_N | CYS_A_49_H | 5.222 | -5.359 | -5.268 | 5.465 | -5.444 | -6.244 | 1.01 | 165.22 | -19.28 |
| 287 | 2jsa-5 | CYS_A_49_N | CYS_A_49_CA | 5.222 | -5.359 | -5.268 | 3.856 | -5.009 | -4.915 | 1.45 | 75.95 | 165.63 |
| 288 | 2jsa-5 | CYS_A_49_CA | CYS_A_49_HA | 3.856 | -5.009 | -4.915 | 3.885 | -4.153 | -4.24 | 1.09 | 51.76 | 88.06 |
| 289 | 2jsa-5 | CYS_A_49_CA | CYS_A_49_C | 3.856 | -5.009 | -4.915 | 3.233 | -6.193 | -4.173 | 1.53 | 60.99 | -117.75 |
| 290 | 2jsa-5 | CYS_A_49_C | CYS_A_49_O | 3.233 | -6.193 | -4.173 | 2.688 | -6.029 | -3.082 | 1.23 | 27.55 | 163.25 |
| 291 | 2jsa-5 | CYS_A_49_C | ASP_A_50_N | 3.233 | -6.193 | -4.173 | 3.334 | -7.358 | -4.795 | 1.32 | 118.01 | -85.05 |
| 292 | 2jsa-5 | ASP_A_50_N | ASP_A_50_H | 3.334 | -7.358 | -4.795 | 3.796 | -7.418 | -5.691 | 1.01 | 152.53 | -7.40 |
| 293 | 2jsa-5 | ASP_A_50_N | ASP_A_50_CA | 3.334 | -7.358 | -4.795 | 2.786 | -8.57 | -4.207 | 1.45 | 66.15 | -114.33 |
| 294 | 2jsa-5 | ASP_A_50_CA | ASP_A_50_HA | 2.786 | -8.57 | -4.207 | 1.72 | -8.429 | -4.029 | 1.09 | 80.60 | 172.47 |
| 295 | 2jsa-5 | ASP_A_50_CA | ASP_A_50_C | 2.786 | -8.57 | -4.207 | 3.5 | -8.858 | -2.884 | 1.53 | 30.20 | -21.97 |
| 296 | 2jsa-5 | ASP_A_50_C | ASP_A_50_O | 3.5 | -8.858 | -2.884 | 3.034 | -9.67 | -2.088 | 1.23 | 49.63 | -119.85 |
| 297 | 2jsa-5 | ASP_A_50_C | HIS_A_51_N | 3.5 | -8.858 | -2.884 | 4.619 | -8.176 | -2.693 | 1.32 | 81.71 | 31.36 |
| 298 | 2jsa-5 | HIS_A_51_N | HIS_A_51_H | 4.619 | -8.176 | -2.693 | 4.944 | -7.524 | -3.393 | 1.01 | 133.86 | 63.51 |
| 299 | 2jsa-5 | HIS_A_51_N | HIS_A_51_CA | 4.619 | -8.176 | -2.693 | 5.403 | -8.348 | -1.48 | 1.45 | 33.49 | -12.37 |

| | | | | | | | | | | | |
|---|---|---|---|---|---|---|---|---|---|---|---|
| 300 | 2jsa-5 | HIS_A_51_CA | HIS_A_51_HA | 5.403 | -8.348 | -1.48 | 5.035 | -9.234 | -0.964 | 1.09 | 61.73 | -112.56 |
| 301 | 2jsa-5 | HIS_A_51_CA | HIS_A_51_C | 5.403 | -8.348 | -1.48 | 5.191 | -7.146 | -0.558 | 1.53 | 52.93 | 100.00 |
| 302 | 2jsa-5 | HIS_A_51_C | HIS_A_51_O | 5.191 | -7.146 | -0.558 | 5.783 | -7.073 | 0.517 | 1.23 | 29.02 | 7.03 |
| 303 | 2jsa-5 | HIS_A_51_C | TYR_A_52_N | 5.191 | -7.146 | -0.558 | 4.346 | -6.233 | -1.014 | 1.32 | 110.13 | 132.78 |
| 304 | 2jsa-5 | TYR_A_52_N | TYR_A_52_H | 4.346 | -6.233 | -1.014 | 3.897 | -6.358 | -1.91 | 1.01 | 152.52 | -164.44 |
| 305 | 2jsa-5 | TYR_A_52_N | TYR_A_52_CA | 4.346 | -6.233 | -1.014 | 4.049 | -5.037 | -0.242 | 1.45 | 57.93 | 103.95 |
| 306 | 2jsa-5 | TYR_A_52_CA | TYR_A_52_HA | 4.049 | -5.037 | -0.242 | 4.819 | -4.874 | 0.513 | 1.09 | 46.19 | 11.95 |
| 307 | 2jsa-5 | TYR_A_52_CA | TYR_A_52_C | 4.049 | -5.037 | -0.242 | 2.679 | -5.144 | 0.429 | 1.53 | 63.97 | -175.53 |
| 308 | 2jsa-5 | TYR_A_52_C | TYR_A_52_O | 2.679 | -5.144 | 0.429 | 2.518 | -4.752 | 1.585 | 1.23 | 20.13 | 112.33 |
| 309 | 2jsa-5 | TYR_A_52_C | VAL_A_53_N | 2.679 | -5.144 | 0.429 | 1.727 | -5.677 | -0.322 | 1.32 | 124.54 | -150.76 |
| 310 | 2jsa-5 | VAL_A_53_N | VAL_A_53_H | 1.727 | -5.677 | -0.322 | 1.932 | -5.978 | -1.264 | 1.01 | 158.86 | -55.74 |
| 311 | 2jsa-5 | VAL_A_53_N | VAL_A_53_CA | 1.727 | -5.677 | -0.322 | 0.376 | -5.84 | 0.187 | 1.45 | 69.49 | -173.12 |
| 312 | 2jsa-5 | VAL_A_53_CA | VAL_A_53_HA | 0.376 | -5.84 | 0.187 | -0.021 | -4.855 | 0.433 | 1.09 | 76.96 | 111.95 |
| 313 | 2jsa-5 | VAL_A_53_CA | VAL_A_53_C | 0.376 | -5.84 | 0.187 | 0.416 | -6.669 | 1.471 | 1.53 | 32.88 | -87.24 |
| 314 | 2jsa-5 | VAL_A_53_C | VAL_A_53_O | 0.416 | -6.669 | 1.471 | -0.537 | -6.662 | 2.249 | 1.23 | 50.77 | 179.58 |
| 315 | 2jsa-5 | VAL_A_53_C | ASN_A_54_N | 0.416 | -6.669 | 1.471 | 1.529 | -7.364 | 1.655 | 1.33 | 82.02 | -31.98 |
| 316 | 2jsa-5 | ASN_A_54_N | ASN_A_54_H | 1.529 | -7.364 | 1.655 | 2.274 | -7.322 | 0.974 | 1.01 | 132.38 | 3.23 |
| 317 | 2jsa-5 | ASN_A_54_N | ASN_A_54_CA | 1.529 | -7.364 | 1.655 | 1.705 | -8.197 | 2.833 | 1.45 | 35.86 | -78.07 |
| 318 | 2jsa-5 | ASN_A_54_CA | ASN_A_54_HA | 1.705 | -8.197 | 2.833 | 0.976 | -7.905 | 3.589 | 1.09 | 46.09 | 158.17 |
| 319 | 2jsa-5 | ASN_A_54_CA | ASN_A_54_C | 1.705 | -8.197 | 2.833 | 3.123 | -8.009 | 3.377 | 1.53 | 69.18 | 7.55 |
| 320 | 2jsa-5 | ASN_A_54_C | ASN_A_54_O | 3.123 | -8.009 | 3.377 | 3.947 | -8.919 | 3.3 | 1.23 | 93.59 | -47.84 |
| 321 | 2jsa-5 | ASN_A_54_C | SER_A_55_N | 3.123 | -8.009 | 3.377 | 3.363 | -6.822 | 3.915 | 1.33 | 66.05 | 78.57 |
| 322 | 2jsa-5 | SER_A_55_N | SER_A_55_H | 3.363 | -6.822 | 3.915 | 2.637 | -6.12 | 3.945 | 1.01 | 88.30 | 135.96 |
| 323 | 2jsa-5 | SER_A_55_N | SER_A_55_CA | 3.363 | -6.822 | 3.915 | 4.667 | -6.504 | 4.472 | 1.45 | 67.46 | 13.70 |
| 324 | 2jsa-5 | SER_A_55_CA | SER_A_55_HA | 4.667 | -6.504 | 4.472 | 4.863 | -7.189 | 5.297 | 1.09 | 40.81 | -74.03 |
| 325 | 2jsa-5 | SER_A_55_CA | SER_A_55_C | 4.667 | -6.504 | 4.472 | 4.669 | -5.072 | 5.012 | 1.53 | 69.34 | 89.92 |
| 326 | 2jsa-5 | SER_A_55_C | SER_A_55_O | 4.669 | -5.072 | 5.012 | 5.311 | -4.786 | 6.022 | 1.23 | 34.83 | 24.01 |
| 327 | 2jsa-5 | SER_A_55_C | LYS_A_56_N | 4.669 | -5.072 | 5.012 | 3.945 | -4.209 | 4.314 | 1.33 | 121.78 | 129.99 |
| 328 | 2jsa-5 | LYS_A_56_N | LYS_A_56_H | 3.945 | -4.209 | 4.314 | 3.444 | -4.514 | 3.492 | 1.01 | 144.49 | -148.67 |
| 329 | 2jsa-5 | LYS_A_56_N | LYS_A_56_CA | 3.945 | -4.209 | 4.314 | 3.855 | -2.814 | 4.71 | 1.45 | 74.18 | 93.69 |
| 330 | 2jsa-5 | LYS_A_56_CA | LYS_A_56_HA | 3.855 | -2.814 | 4.71 | 4.222 | -2.73 | 5.733 | 1.09 | 20.20 | 12.89 |
| 331 | 2jsa-5 | LYS_A_56_CA | LYS_A_56_C | 3.855 | -2.814 | 4.71 | 2.389 | -2.38 | 4.706 | 1.53 | 90.15 | 163.51 |
| 332 | 2jsa-5 | LYS_A_56_C | LYS_A_56_O | 2.389 | -2.38 | 4.706 | 2.089 | -1.193 | 4.823 | 1.23 | 84.54 | 104.18 |
| 333 | 2jsa-5 | LYS_A_56_C | VAL_A_57_N | 2.389 | -2.38 | 4.706 | 1.514 | -3.365 | 4.567 | 1.32 | 96.02 | -131.62 |
| 334 | 2jsa-5 | VAL_A_57_N | VAL_A_57_H | 1.514 | -3.365 | 4.567 | 1.833 | -4.319 | 4.474 | 1.01 | 95.28 | -71.51 |
| 335 | 2jsa-5 | VAL_A_57_N | VAL_A_57_CA | 1.514 | -3.365 | 4.567 | 0.085 | -3.1 | 4.545 | 1.45 | 90.87 | 169.49 |
| 336 | 2jsa-5 | VAL_A_57_CA | VAL_A_57_HA | 0.085 | -3.1 | 4.545 | -0.067 | -2.052 | 4.285 | 1.09 | 103.79 | 98.25 |
| 337 | 2jsa-5 | VAL_A_57_CA | VAL_A_57_C | 0.085 | -3.1 | 4.545 | -0.493 | -3.331 | 5.942 | 1.53 | 24.02 | -158.22 |
| 338 | 2jsa-5 | VAL_A_57_C | VAL_A_57_O | -0.493 | -3.331 | 5.942 | -1.29 | -2.528 | 6.429 | 1.23 | 66.71 | 134.79 |
| 339 | 2jsa-5 | VAL_A_57_C | ASP_A_58_N | -0.493 | -3.331 | 5.942 | -0.072 | -4.431 | 6.548 | 1.32 | 62.77 | -69.06 |
| 340 | 2jsa-5 | ASP_A_58_N | ASP_A_58_H | -0.072 | -4.431 | 6.548 | 0.583 | -5.047 | 6.087 | 1.01 | 117.14 | -43.24 |
| 341 | 2jsa-5 | ASP_A_58_N | ASP_A_58_CA | -0.072 | -4.431 | 6.548 | -0.538 | -4.777 | 7.88 | 1.45 | 23.54 | -143.41 |
| 342 | 2jsa-5 | ASP_A_58_CA | ASP_A_58_HA | -0.538 | -4.777 | 7.88 | -1.599 | -5.023 | 7.833 | 1.09 | 92.47 | -166.95 |
| 343 | 2jsa-5 | ASP_A_58_CA | ASP_A_58_C | -0.538 | -4.777 | 7.88 | -0.321 | -3.587 | 8.817 | 1.53 | 52.24 | 79.67 |
| 344 | 2jsa-5 | ASP_A_58_C | ASP_A_58_O | -0.321 | -3.587 | 8.817 | -1.204 | -3.24 | 9.6 | 1.23 | 50.47 | 158.55 |
| 345 | 2jsa-5 | ASP_A_58_C | PRO_A_59_N | -0.321 | -3.587 | 8.817 | 0.891 | -2.981 | 8.705 | 1.36 | 94.72 | 26.57 |
| 346 | 2jsa-5 | PRO_A_59_N | PRO_A_59_CA | 0.891 | -2.981 | 8.705 | 1.236 | -1.837 | 9.533 | 1.45 | 55.28 | 73.22 |
| 347 | 2jsa-5 | PRO_A_59_CA | PRO_A_59_HA | 1.236 | -1.837 | 9.533 | 0.947 | -2.024 | 10.567 | 1.09 | 18.41 | -147.09 |
| 348 | 2jsa-5 | PRO_A_59_CA | PRO_A_59_C | 1.236 | -1.837 | 9.533 | 0.52 | -0.575 | 9.049 | 1.53 | 108.45 | 119.57 |
| 349 | 2jsa-5 | PRO_A_59_C | PRO_A_59_O | 0.52 | -0.575 | 9.049 | -0.157 | 0.096 | 9.828 | 1.23 | 50.74 | 135.26 |
| 350 | 2jsa-5 | PRO_A_59_C | ILE_A_60_N | 0.52 | -0.575 | 9.049 | 0.693 | -0.288 | 7.767 | 1.33 | 165.35 | 58.92 |
| 351 | 2jsa-5 | ILE_A_60_N | ILE_A_60_H | 0.693 | -0.288 | 7.767 | 1.263 | -0.885 | 7.185 | 1.01 | 125.19 | -46.33 |
| 352 | 2jsa-5 | ILE_A_60_N | ILE_A_60_CA | 0.693 | -0.288 | 7.767 | 0.072 | 0.882 | 7.171 | 1.45 | 114.23 | 117.96 |
| 353 | 2jsa-5 | ILE_A_60_CA | ILE_A_60_HA | 0.072 | 0.882 | 7.171 | 0.592 | 1.766 | 7.542 | 1.09 | 70.11 | 59.53 |
| 354 | 2jsa-5 | ILE_A_60_CA | ILE_A_60_C | 0.072 | 0.882 | 7.171 | -1.385 | 0.971 | 7.631 | 1.53 | 72.51 | 176.50 |
| 355 | 2jsa-5 | ILE_A_60_C | ILE_A_60_O | -1.385 | 0.971 | 7.631 | -1.807 | 1.993 | 8.172 | 1.23 | 63.93 | 112.44 |
| 356 | 2jsa-5 | ILE_A_60_C | ILE_A_61_N | -1.385 | 0.971 | 7.631 | -2.112 | -0.111 | 7.401 | 1.32 | 100.01 | -123.90 |
| 357 | 2jsa-5 | ILE_A_61_N | ILE_A_61_H | -2.112 | -0.111 | 7.401 | -1.698 | -0.916 | 6.953 | 1.01 | 116.33 | -62.78 |
| 358 | 2jsa-5 | ILE_A_61_N | ILE_A_61_CA | -2.112 | -0.111 | 7.401 | -3.512 | -0.168 | 7.785 | 1.45 | 74.67 | -177.67 |
| 359 | 2jsa-5 | ILE_A_61_CA | ILE_A_61_HA | -3.512 | -0.168 | 7.785 | -4.031 | 0.653 | 7.291 | 1.09 | 116.96 | 122.30 |
| 360 | 2jsa-5 | ILE_A_61_CA | ILE_A_61_C | -3.512 | -0.168 | 7.785 | -3.631 | 0.048 | 9.295 | 1.53 | 9.28 | 118.85 |
| 361 | 2jsa-5 | ILE_A_61_C | ILE_A_61_O | -3.631 | 0.048 | 9.295 | -4.667 | 0.5 | 9.781 | 1.23 | 66.73 | 156.43 |
| 362 | 2jsa-5 | ILE_A_61_C | HIS_A_62_N | -3.631 | 0.048 | 9.295 | -2.556 | -0.285 | 9.994 | 1.32 | 58.15 | -17.21 |
| 363 | 2jsa-5 | HIS_A_62_N | HIS_A_62_H | -2.556 | -0.285 | 9.994 | -1.738 | -0.651 | 9.527 | 1.01 | 117.52 | -24.11 |
| 364 | 2jsa-5 | HIS_A_62_N | HIS_A_62_CA | -2.556 | -0.285 | 9.994 | -2.527 | -0.133 | 11.439 | 1.45 | 6.11 | 79.20 |
| 365 | 2jsa-5 | HIS_A_62_CA | HIS_A_62_HA | -2.527 | -0.133 | 11.439 | -3.427 | -0.595 | 11.844 | 1.09 | 68.18 | -152.83 |
| 366 | 2jsa-5 | HIS_A_62_CA | HIS_A_62_C | -2.527 | -0.133 | 11.439 | -2.559 | 1.354 | 11.8 | 1.53 | 76.36 | 91.23 |
| 367 | 2jsa-5 | HIS_A_62_C | HIS_A_62_O | -2.559 | 1.354 | 11.8 | -2.922 | 1.718 | 12.916 | 1.23 | 24.73 | 134.92 |
| 368 | 2jsa-5 | HIS_A_62_C | GLU_A_63_N | -2.559 | 1.354 | 11.8 | -2.175 | 2.173 | 10.832 | 1.32 | 136.94 | 64.88 |
| 369 | 2jsa-5 | GLU_A_63_N | GLU_A_63_H | -2.175 | 2.173 | 10.832 | -1.89 | 1.803 | 9.936 | 1.01 | 152.47 | -52.39 |
| 370 | 2jsa-5 | GLU_A_63_N | GLU_A_63_CA | -2.175 | 2.173 | 10.832 | -2.155 | 3.612 | 11.033 | 1.45 | 82.05 | 89.20 |
| 371 | 2jsa-5 | GLU_A_63_CA | GLU_A_63_HA | -2.155 | 3.612 | 11.033 | -1.941 | 3.803 | 12.085 | 1.09 | 15.25 | 41.75 |
| 372 | 2jsa-5 | GLU_A_63_CA | GLU_A_63_C | -2.155 | 3.612 | 11.033 | -3.522 | 4.213 | 10.703 | 1.53 | 102.46 | 156.27 |
| 373 | 2jsa-5 | GLU_A_63_C | GLU_A_63_O | -3.522 | 4.213 | 10.703 | -4.024 | 5.064 | 11.436 | 1.23 | 53.43 | 120.54 |
| 374 | 2jsa-5 | GLU_A_63_C | LEU_A_64_N | -3.522 | 4.213 | 10.703 | -4.087 | 3.748 | 9.599 | 1.32 | 146.46 | -140.55 |
| 375 | 2jsa-5 | LEU_A_64_N | LEU_A_64_H | -4.087 | 3.748 | 9.599 | -3.616 | 3.048 | 9.043 | 1.01 | 123.38 | -56.07 |
| 376 | 2jsa-5 | LEU_A_64_N | LEU_A_64_CA | -4.087 | 3.748 | 9.599 | -5.387 | 4.228 | 9.163 | 1.45 | 107.46 | 159.73 |
| 377 | 2jsa-5 | LEU_A_64_CA | LEU_A_64_HA | -5.387 | 4.228 | 9.163 | -5.357 | 5.318 | 9.151 | 1.09 | 90.63 | 88.42 |
| 378 | 2jsa-5 | LEU_A_64_CA | LEU_A_64_C | -5.387 | 4.228 | 9.163 | -6.45 | 3.804 | 10.178 | 1.53 | 48.43 | -158.25 |
| 379 | 2jsa-5 | LEU_A_64_C | LEU_A_64_O | -6.45 | 3.804 | 10.178 | -7.342 | 4.584 | 10.511 | 1.23 | 74.30 | 138.83 |
| 380 | 2jsa-5 | LEU_A_64_C | GLU_A_65_N | -6.45 | 3.804 | 10.178 | -6.319 | 2.57 | 10.643 | 1.33 | 69.46 | -83.94 |
| 381 | 2jsa-5 | GLU_A_65_N | GLU_A_65_H | -6.319 | 2.57 | 10.643 | -5.559 | 1.985 | 10.326 | 1.01 | 108.29 | -37.59 |
| 382 | 2jsa-5 | GLU_A_65_N | GLU_A_65_CA | -6.319 | 2.57 | 10.643 | -7.258 | 2.033 | 11.614 | 1.45 | 48.09 | -150.24 |
| 383 | 2jsa-5 | GLU_A_65_CA | GLU_A_65_HA | -7.258 | 2.033 | 11.614 | -8.267 | 2.215 | 11.245 | 1.09 | 109.79 | 169.78 |
| 384 | 2jsa-5 | GLU_A_65_CA | GLU_A_65_C | -7.258 | 2.033 | 11.614 | -7.101 | 2.75 | 12.957 | 1.53 | 28.66 | 77.65 |
| 385 | 2jsa-5 | GLU_A_65_C | GLU_A_65_O | -7.101 | 2.75 | 12.957 | -8.074 | 2.927 | 13.687 | 1.23 | 53.57 | 169.69 |
| 386 | 2jsa-5 | GLU_A_65_C | GLY_A_66_N | -7.101 | 2.75 | 12.957 | -5.868 | 3.142 | 13.241 | 1.32 | 77.62 | 17.64 |
| 387 | 2jsa-5 | GLY_A_66_N | GLY_A_66_H | -5.868 | 3.142 | 13.241 | -5.114 | 2.961 | 12.593 | 1.01 | 129.88 | -13.50 |
| 388 | 2jsa-5 | GLY_A_66_N | GLY_A_66_CA | -5.868 | 3.142 | 13.241 | -5.571 | 3.836 | 14.482 | 1.45 | 31.31 | 66.83 |
| 389 | 2jsa-5 | GLY_A_66_CA | GLY_A_66_C | -5.571 | 3.836 | 14.482 | -6.2 | 5.231 | 14.495 | 1.53 | 89.51 | 114.27 |
| 390 | 2jsa-5 | GLY_A_66_C | GLY_A_66_O | -6.2 | 5.231 | 14.495 | -6.854 | 5.612 | 15.464 | 1.23 | 37.99 | 149.78 |
| 391 | 2jsa-5 | GLY_A_66_C | GLY_A_67_N | -6.2 | 5.231 | 14.495 | -5.978 | 5.955 | 13.407 | 1.33 | 145.16 | 72.95 |
| 392 | 2jsa-5 | GLY_A_67_N | GLY_A_67_H | -5.978 | 5.955 | 13.407 | -5.428 | 5.578 | 12.648 | 1.01 | 138.70 | -34.43 |
| 393 | 2jsa-5 | GLY_A_67_N | GLY_A_67_CA | -5.978 | 5.955 | 13.407 | -6.516 | 7.299 | 13.281 | 1.45 | 94.97 | 111.82 |
| 394 | 2jsa-5 | GLY_A_67_CA | GLY_A_67_C | -6.516 | 7.299 | 13.281 | -5.772 | 8.087 | 12.201 | 1.53 | 134.90 | 46.65 |
| 395 | 2jsa-5 | GLY_A_67_C | GLY_A_67_O | -5.772 | 8.087 | 12.201 | -6.316 | 9.033 | 11.632 | 1.23 | 117.54 | 119.90 |

| | | | | | | | | | | | | |
|---|---|---|---|---|---|---|---|---|---|---|---|---|
| 396 | 2jsa-5 | GLY_A_67_C | THR_A_68_N | -5.772 | 8.087 | 12.201 | -4.54 | 7.668 | 11.95 | 1.33 | 100.92 | -18.78 |
| 397 | 2jsa-5 | THR_A_68_N | THR_A_68_H | -4.54 | 7.668 | 11.95 | -4.157 | 6.882 | 12.456 | 1.01 | 59.94 | -64.02 |
| 398 | 2jsa-5 | THR_A_68_N | THR_A_68_CA | -4.54 | 7.668 | 11.95 | -3.716 | 8.324 | 10.948 | 1.45 | 133.57 | 38.52 |
| 399 | 2jsa-5 | THR_A_68_CA | THR_A_68_HA | -3.716 | 8.324 | 10.948 | -3.524 | 9.348 | 11.268 | 1.09 | 72.93 | 79.38 |
| 400 | 2jsa-5 | THR_A_68_CA | THR_A_68_C | -3.716 | 8.324 | 10.948 | -4.451 | 8.379 | 9.607 | 1.53 | 151.21 | 175.72 |
| 401 | 2jsa-5 | THR_A_68_C | THR_A_68_O | -4.451 | 8.379 | 9.607 | -5.257 | 7.503 | 9.301 | 1.23 | 104.42 | -132.62 |
| 402 | 2jsa-5 | THR_A_68_C | ALA_A_69_N | -4.451 | 8.379 | 9.607 | -4.144 | 9.417 | 8.844 | 1.32 | 125.18 | 73.52 |
| 403 | 2jsa-5 | ALA_A_69_N | ALA_A_69_H | -4.144 | 9.417 | 8.844 | -3.471 | 10.1 | 9.161 | 1.01 | 71.71 | 45.42 |
| 404 | 2jsa-5 | ALA_A_69_N | ALA_A_69_CA | -4.144 | 9.417 | 8.844 | -4.765 | 9.597 | 7.542 | 1.45 | 153.59 | 163.84 |
| 405 | 2jsa-5 | ALA_A_69_CA | ALA_A_69_HA | -4.765 | 9.597 | 7.542 | -5.776 | 9.191 | 7.576 | 1.09 | 88.21 | -158.12 |
| 406 | 2jsa-5 | ALA_A_69_CA | ALA_A_69_C | -4.765 | 9.597 | 7.542 | -3.968 | 8.823 | 6.49 | 1.53 | 133.44 | -44.16 |
| 407 | 2jsa-5 | ALA_A_69_C | ALA_A_69_O | -3.968 | 8.823 | 6.49 | -2.851 | 8.381 | 6.753 | 1.23 | 77.65 | -21.59 |
| 408 | 2jsa-5 | ALA_A_69_C | PRO_A_70_N | -3.968 | 8.823 | 6.49 | -4.59 | 8.679 | 5.289 | 1.36 | 152.00 | -166.97 |
| 409 | 2jsa-5 | PRO_A_70_N | PRO_A_70_CA | -4.59 | 8.679 | 5.289 | -3.952 | 7.965 | 4.197 | 1.45 | 138.75 | -48.22 |
| 410 | 2jsa-5 | PRO_A_70_CA | PRO_A_70_HA | -3.952 | 7.965 | 4.197 | -3.51 | 7.039 | 4.564 | 1.09 | 70.32 | -64.48 |
| 411 | 2jsa-5 | PRO_A_70_CA | PRO_A_70_C | -3.952 | 7.965 | 4.197 | -2.847 | 8.811 | 3.562 | 1.53 | 114.53 | 37.44 |
| 412 | 2jsa-5 | PRO_A_70_C | PRO_A_70_O | -2.847 | 8.811 | 3.562 | -2.177 | 8.367 | 2.63 | 1.23 | 139.23 | -33.53 |
| 413 | 2jsa-5 | PRO_A_70_C | LYS_A_71_N | -2.847 | 8.811 | 3.562 | -2.689 | 10.017 | 4.091 | 1.33 | 66.49 | 82.54 |
| 414 | 2jsa-5 | LYS_A_71_N | LYS_A_71_H | -2.689 | 10.017 | 4.091 | -3.275 | 10.318 | 4.856 | 1.01 | 40.73 | 152.81 |
| 415 | 2jsa-5 | LYS_A_71_N | LYS_A_71_CA | -2.689 | 10.017 | 4.091 | -1.677 | 10.928 | 3.587 | 1.45 | 110.31 | 41.99 |
| 416 | 2jsa-5 | LYS_A_71_CA | LYS_A_71_HA | -1.677 | 10.928 | 3.587 | -1.143 | 10.427 | 2.779 | 1.09 | 137.82 | -43.17 |
| 417 | 2jsa-5 | LYS_A_71_CA | LYS_A_71_C | -1.677 | 10.928 | 3.587 | -0.667 | 11.225 | 4.697 | 1.53 | 43.48 | 16.39 |
| 418 | 2jsa-5 | LYS_A_71_C | LYS_A_71_O | -0.667 | 11.225 | 4.697 | 0.231 | 12.047 | 4.52 | 1.23 | 98.27 | 42.47 |
| 419 | 2jsa-5 | LYS_A_71_C | ASP_A_72_N | -0.667 | 11.225 | 4.697 | -0.848 | 10.541 | 5.817 | 1.32 | 32.28 | -104.82 |
| 420 | 2jsa-5 | ASP_A_72_N | ASP_A_72_H | -0.848 | 10.541 | 5.817 | -1.609 | 9.882 | 5.893 | 1.01 | 85.68 | -139.11 |
| 421 | 2jsa-5 | ASP_A_72_N | ASP_A_72_CA | -0.848 | 10.541 | 5.817 | 0.036 | 10.721 | 6.956 | 1.45 | 38.38 | 11.51 |
| 422 | 2jsa-5 | ASP_A_72_CA | ASP_A_72_HA | 0.036 | 10.721 | 6.956 | 0.736 | 11.528 | 6.739 | 1.09 | 101.48 | 49.06 |
| 423 | 2jsa-5 | ASP_A_72_CA | ASP_A_72_C | 0.036 | 10.721 | 6.956 | 0.804 | 9.423 | 7.211 | 1.53 | 80.40 | -59.39 |
| 424 | 2jsa-5 | ASP_A_72_C | ASP_A_72_O | 0.804 | 9.423 | 7.211 | 1.983 | 9.452 | 7.559 | 1.23 | 73.56 | 1.41 |
| 425 | 2jsa-5 | ASP_A_72_C | VAL_A_73_N | 0.804 | 9.423 | 7.211 | 0.102 | 8.313 | 7.028 | 1.33 | 97.93 | -122.31 |
| 426 | 2jsa-5 | VAL_A_73_N | VAL_A_73_H | 0.102 | 8.313 | 7.028 | -0.865 | 8.363 | 6.741 | 1.01 | 106.51 | 177.04 |
| 427 | 2jsa-5 | VAL_A_73_N | VAL_A_73_CA | 0.102 | 8.313 | 7.028 | 0.704 | 7.007 | 7.235 | 1.45 | 81.81 | -65.25 |
| 428 | 2jsa-5 | VAL_A_73_CA | VAL_A_73_HA | 0.704 | 7.007 | 7.235 | 0.766 | 6.822 | 8.308 | 1.09 | 10.31 | -71.47 |
| 429 | 2jsa-5 | VAL_A_73_CA | VAL_A_73_C | 0.704 | 7.007 | 7.235 | 2.126 | 7.009 | 6.671 | 1.53 | 111.63 | 0.08 |
| 430 | 2jsa-5 | VAL_A_73_C | VAL_A_73_O | 2.126 | 7.009 | 6.671 | 3.082 | 6.719 | 7.389 | 1.23 | 54.29 | -16.88 |
| 431 | 2jsa-5 | VAL_A_73_C | CYS_A_74_N | 2.126 | 7.009 | 6.671 | 2.222 | 7.339 | 5.392 | 1.32 | 164.96 | 73.78 |
| 432 | 2jsa-5 | CYS_A_74_N | CYS_A_74_H | 2.222 | 7.339 | 5.392 | 1.393 | 7.567 | 4.862 | 1.01 | 121.65 | 164.62 |
| 433 | 2jsa-5 | CYS_A_74_N | CYS_A_74_CA | 2.222 | 7.339 | 5.392 | 3.511 | 7.383 | 4.724 | 1.45 | 117.38 | 1.96 |
| 434 | 2jsa-5 | CYS_A_74_CA | CYS_A_74_HA | 3.511 | 7.383 | 4.724 | 4.111 | 6.544 | 5.079 | 1.09 | 71.01 | -54.43 |
| 435 | 2jsa-5 | CYS_A_74_CA | CYS_A_74_C | 3.511 | 7.383 | 4.724 | 4.212 | 8.684 | 5.119 | 1.53 | 75.04 | 61.68 |
| 436 | 2jsa-5 | CYS_A_74_C | CYS_A_74_O | 4.212 | 8.684 | 5.119 | 4.358 | 9.589 | 4.3 | 1.23 | 131.78 | 80.84 |
| 437 | 2jsa-5 | CYS_A_74_C | THR_A_75_N | 4.212 | 8.684 | 5.119 | 4.627 | 8.736 | 6.377 | 1.33 | 18.39 | 7.14 |
| 438 | 2jsa-5 | THR_A_75_N | THR_A_75_H | 4.627 | 8.736 | 6.377 | 4.473 | 7.954 | 6.997 | 1.01 | 52.12 | -101.14 |
| 439 | 2jsa-5 | THR_A_75_N | THR_A_75_CA | 4.627 | 8.736 | 6.377 | 5.31 | 9.911 | 6.891 | 1.45 | 69.28 | 59.83 |
| 440 | 2jsa-5 | THR_A_75_CA | THR_A_75_HA | 5.31 | 9.911 | 6.891 | 6.202 | 10.088 | 6.29 | 1.09 | 123.46 | 11.22 |
| 441 | 2jsa-5 | THR_A_75_CA | THR_A_75_C | 5.31 | 9.911 | 6.891 | 5.755 | 9.68 | 8.337 | 1.53 | 19.12 | -27.43 |
| 442 | 2jsa-5 | THR_A_75_C | THR_A_75_O | 5.755 | 9.68 | 8.337 | 6.943 | 9.509 | 8.604 | 1.23 | 77.46 | -8.19 |
| 443 | 2jsa-5 | THR_A_75_C | LYS_A_76_N | 5.755 | 9.68 | 8.337 | 4.777 | 9.682 | 9.231 | 1.33 | 47.57 | 179.88 |
| 444 | 2jsa-5 | LYS_A_76_N | LYS_A_76_H | 4.777 | 9.682 | 9.231 | 3.822 | 9.83 | 8.94 | 1.01 | 106.76 | 171.19 |
| 445 | 2jsa-5 | LYS_A_76_N | LYS_A_76_CA | 4.777 | 9.682 | 9.231 | 5.053 | 9.473 | 10.641 | 1.45 | 13.80 | -37.13 |
| 446 | 2jsa-5 | LYS_A_76_CA | LYS_A_76_HA | 5.053 | 9.473 | 10.641 | 5.971 | 10.008 | 10.888 | 1.09 | 76.91 | 30.23 |
| 447 | 2jsa-5 | LYS_A_76_CA | LYS_A_76_C | 5.053 | 9.473 | 10.641 | 5.301 | 7.985 | 10.895 | 1.53 | 80.44 | -80.54 |
| 448 | 2jsa-5 | LYS_A_76_C | LYS_A_76_O | 5.301 | 7.985 | 10.895 | 6.192 | 7.622 | 11.66 | 1.23 | 51.51 | -22.17 |
| 449 | 2jsa-5 | LYS_A_76_C | LEU_A_77_N | 5.301 | 7.985 | 10.895 | 4.495 | 7.164 | 10.237 | 1.33 | 119.77 | -134.47 |
| 450 | 2jsa-5 | LEU_A_77_N | LEU_A_77_H | 4.495 | 7.164 | 10.237 | 3.784 | 7.533 | 9.623 | 1.01 | 127.47 | 152.57 |
| 451 | 2jsa-5 | LEU_A_77_N | LEU_A_77_CA | 4.495 | 7.164 | 10.237 | 4.615 | 5.723 | 10.382 | 1.45 | 84.27 | -85.24 |
| 452 | 2jsa-5 | LEU_A_77_CA | LEU_A_77_HA | 4.615 | 5.723 | 10.382 | 4.44 | 5.473 | 11.429 | 1.09 | 16.25 | -124.99 |
| 453 | 2jsa-5 | LEU_A_77_CA | LEU_A_77_C | 4.615 | 5.723 | 10.382 | 6.043 | 5.298 | 10.035 | 1.53 | 103.11 | -16.57 |
| 454 | 2jsa-5 | LEU_A_77_C | LEU_A_77_O | 6.043 | 5.298 | 10.035 | 6.464 | 4.192 | 10.371 | 1.23 | 74.15 | -69.16 |
| 455 | 2jsa-5 | LEU_A_77_C | ASN_A_78_N | 6.043 | 5.298 | 10.035 | 6.749 | 6.198 | 9.367 | 1.32 | 120.28 | 51.89 |
| 456 | 2jsa-5 | ASN_A_78_N | ASN_A_78_H | 6.749 | 6.198 | 9.367 | 6.338 | 7.088 | 9.124 | 1.01 | 103.92 | 114.79 |
| 457 | 2jsa-5 | ASN_A_78_N | ASN_A_78_CA | 6.749 | 6.198 | 9.367 | 8.122 | 5.93 | 8.971 | 1.45 | 105.81 | -11.04 |
| 458 | 2jsa-5 | ASN_A_78_CA | ASN_A_78_HA | 8.122 | 5.93 | 8.971 | 8.615 | 6.872 | 8.731 | 1.09 | 102.72 | 62.37 |
| 459 | 2jsa-5 | ASN_A_78_CA | ASN_A_78_C | 8.122 | 5.93 | 8.971 | 8.124 | 5.015 | 7.745 | 1.53 | 143.26 | -89.87 |
| 460 | 2jsa-5 | ASN_A_78_C | ASN_A_78_O | 8.124 | 5.015 | 7.745 | 8.72 | 3.938 | 7.771 | 1.23 | 88.79 | -61.04 |
| 461 | 2jsa-5 | ASN_A_78_C | GLU_A_79_N | 8.124 | 5.015 | 7.745 | 7.451 | 5.475 | 6.7 | 1.33 | 142.04 | 145.65 |
| 462 | 2jsa-5 | GLU_A_79_N | GLU_A_79_H | 7.451 | 5.475 | 6.7 | 6.985 | 6.37 | 6.749 | 1.01 | 87.22 | 117.50 |
| 463 | 2jsa-5 | GLU_A_79_N | GLU_A_79_CA | 7.451 | 5.475 | 6.7 | 7.369 | 4.711 | 5.468 | 1.45 | 148.05 | -96.13 |
| 464 | 2jsa-5 | GLU_A_79_CA | GLU_A_79_HA | 7.369 | 4.711 | 5.468 | 7.73 | 3.703 | 5.672 | 1.09 | 79.21 | -70.30 |
| 465 | 2jsa-5 | GLU_A_79_CA | GLU_A_79_C | 7.369 | 4.711 | 5.468 | 8.265 | 5.335 | 4.396 | 1.53 | 134.47 | 34.85 |
| 466 | 2jsa-5 | GLU_A_79_C | GLU_A_79_O | 8.265 | 5.335 | 4.396 | 9.313 | 4.784 | 4.06 | 1.23 | 105.84 | -27.73 |
| 467 | 2jsa-5 | GLU_A_79_C | CYS_A_80_N | 8.265 | 5.335 | 4.396 | 7.82 | 6.475 | 3.888 | 1.33 | 112.54 | 111.32 |
| 468 | 2jsa-5 | CYS_A_80_N | CYS_A_80_H | 7.82 | 6.475 | 3.888 | 6.949 | 6.871 | 4.211 | 1.01 | 71.35 | 155.55 |
| 469 | 2jsa-5 | CYS_A_80_N | CYS_A_80_CA | 7.82 | 6.475 | 3.888 | 8.57 | 7.179 | 2.862 | 1.45 | 134.93 | 43.19 |
| 470 | 2jsa-5 | CYS_A_80_CA | CYS_A_80_HA | 8.57 | 7.179 | 2.862 | 8.639 | 6.535 | 1.985 | 1.09 | 143.55 | -83.88 |
| 471 | 2jsa-5 | CYS_A_80_CA | CYS_A_80_C | 8.57 | 7.179 | 2.862 | 9.983 | 7.436 | 3.388 | 1.53 | 69.88 | 10.31 |
| 472 | 2jsa-5 | CYS_A_80_C | CYS_A_80_O | 9.983 | 7.436 | 3.388 | 10.964 | 7.183 | 2.691 | 1.23 | 124.53 | -14.46 |
| 473 | 2jsa-5 | CYS_A_80_C | PRO_A_81_N | 9.983 | 7.436 | 3.388 | 10.043 | 7.948 | 4.647 | 1.36 | 22.27 | 83.32 |
| 474 | 2jsa-5 | PRO_A_81_N | PRO_A_81_CA | 10.043 | 7.948 | 4.647 | 11.32 | 8.241 | 5.275 | 1.45 | 64.39 | 12.92 |
| 475 | 2jsa-5 | PRO_A_81_CA | PRO_A_81_HA | 11.32 | 8.241 | 5.275 | 11.972 | 8.758 | 4.571 | 1.09 | 130.23 | 38.41 |
| 476 | 2jsa-5 | PRO_A_81_CA | PRO_A_81_C | 11.32 | 8.241 | 5.275 | 12.016 | 6.957 | 5.727 | 1.53 | 72.80 | -61.54 |
| 477 | 2jsa-5 | PRO_A_81_C | PRO_A_81_O | 12.016 | 6.957 | 5.727 | 11.808 | 6.496 | 6.849 | 1.23 | 24.26 | -114.28 |
| 1 | 2jsa-6 | ARG_A_1_N | ARG_A_1_H | 14.712 | 1.733 | -2.113 | 15.435 | 2.427 | -2.244 | 1.01 | 97.45 | 43.83 |
| 2 | 2jsa-6 | ARG_A_1_N | ARG_A_1_CA | 14.712 | 1.733 | -2.113 | 13.475 | 2.133 | -1.465 | 1.45 | 63.51 | 162.08 |
| 3 | 2jsa-6 | ARG_A_1_CA | ARG_A_1_HA | 13.475 | 2.133 | -1.465 | 12.653 | 1.967 | -2.161 | 1.09 | 129.69 | -168.58 |
| 4 | 2jsa-6 | ARG_A_1_CA | ARG_A_1_C | 13.475 | 2.133 | -1.465 | 13.524 | 3.618 | -1.098 | 1.53 | 76.13 | 88.11 |
| 5 | 2jsa-6 | ARG_A_1_C | ARG_A_1_O | 13.524 | 3.618 | -1.098 | 14.266 | 4.014 | -0.201 | 1.23 | 43.16 | 28.09 |
| 6 | 2jsa-6 | ARG_A_1_C | SER_A_2_N | 13.524 | 3.618 | -1.098 | 12.724 | 4.398 | -1.81 | 1.32 | 122.51 | 135.73 |
| 7 | 2jsa-6 | SER_A_2_N | SER_A_2_H | 12.724 | 4.398 | -1.81 | 12.139 | 4.003 | -2.532 | 1.01 | 135.65 | -145.97 |
| 8 | 2jsa-6 | SER_A_2_N | SER_A_2_CA | 12.724 | 4.398 | -1.81 | 12.668 | 5.829 | -1.571 | 1.45 | 80.53 | 92.24 |
| 9 | 2jsa-6 | SER_A_2_CA | SER_A_2_HA | 12.668 | 5.829 | -1.571 | 12.331 | 5.991 | -0.547 | 1.09 | 20.06 | 154.33 |
| 10 | 2jsa-6 | SER_A_2_CA | SER_A_2_C | 12.668 | 5.829 | -1.571 | 11.665 | 6.481 | -2.525 | 1.53 | 128.57 | 146.97 |
| 11 | 2jsa-6 | SER_A_2_C | SER_A_2_O | 11.665 | 6.481 | -2.525 | 11.961 | 7.507 | -3.135 | 1.23 | 119.74 | 73.91 |
| 12 | 2jsa-6 | SER_A_2_C | ALA_A_3_N | 11.665 | 6.481 | -2.525 | 10.501 | 5.857 | -2.625 | 1.32 | 94.33 | -151.81 |
| 13 | 2jsa-6 | ALA_A_3_N | ALA_A_3_H | 10.501 | 5.857 | -2.625 | 10.328 | 5.016 | -2.094 | 1.01 | 58.27 | -101.62 |
| 14 | 2jsa-6 | ALA_A_3_N | ALA_A_3_CA | 10.501 | 5.857 | -2.625 | 9.453 | 6.363 | -3.496 | 1.45 | 126.81 | 154.23 |

| | | | | | | | | | | | |
|---|---|---|---|---|---|---|---|---|---|---|---|
| 15 | 2jsa-6 | ALA_A_3_CA | ALA_A_3_HA | 9.453 | 6.363 | -3.496 | 9.03 | 7.263 | -3.049 | 1.09 | 65.80 | 115.17 |
| 16 | 2jsa-6 | ALA_A_3_CA | ALA_A_3_C | 9.453 | 6.363 | -3.496 | 8.345 | 5.315 | -3.616 | 1.53 | 94.50 | -136.59 |
| 17 | 2jsa-6 | ALA_A_3_C | ALA_A_3_O | 8.345 | 5.315 | -3.616 | 7.275 | 5.467 | -3.027 | 1.23 | 61.41 | 171.91 |
| 18 | 2jsa-6 | ALA_A_3_C | LEU_A_4_N | 8.345 | 5.315 | -3.616 | 8.638 | 4.275 | -4.382 | 1.32 | 125.33 | -74.27 |
| 19 | 2jsa-6 | LEU_A_4_N | LEU_A_4_H | 8.638 | 4.275 | -4.382 | 9.538 | 4.216 | -4.836 | 1.01 | 116.72 | -3.75 |
| 20 | 2jsa-6 | LEU_A_4_N | LEU_A_4_CA | 8.638 | 4.275 | -4.382 | 7.68 | 3.202 | -4.586 | 1.45 | 98.07 | -131.76 |
| 21 | 2jsa-6 | LEU_A_4_CA | LEU_A_4_HA | 7.68 | 3.202 | -4.586 | 6.938 | 3.546 | -5.306 | 1.09 | 131.36 | 155.13 |
| 22 | 2jsa-6 | LEU_A_4_CA | LEU_A_4_C | 7.68 | 3.202 | -4.586 | 6.953 | 2.915 | -3.27 | 1.53 | 30.71 | -158.46 |
| 23 | 2jsa-6 | LEU_A_4_C | LEU_A_4_O | 6.953 | 2.915 | -3.27 | 5.738 | 3.078 | -3.18 | 1.23 | 85.80 | 172.36 |
| 24 | 2jsa-6 | LEU_A_4_C | SER_A_5_N | 6.953 | 2.915 | -3.27 | 7.73 | 2.492 | -2.283 | 1.33 | 41.87 | -28.56 |
| 25 | 2jsa-6 | SER_A_5_N | SER_A_5_H | 7.73 | 2.492 | -2.283 | 8.724 | 2.381 | -2.429 | 1.01 | 98.31 | -6.37 |
| 26 | 2jsa-6 | SER_A_5_N | SER_A_5_CA | 7.73 | 2.492 | -2.283 | 7.176 | 2.181 | -0.976 | 1.45 | 25.92 | -150.69 |
| 27 | 2jsa-6 | SER_A_5_CA | SER_A_5_HA | 7.176 | 2.181 | -0.976 | 6.705 | 1.2 | -1.029 | 1.09 | 92.79 | -115.65 |
| 28 | 2jsa-6 | SER_A_5_CA | SER_A_5_C | 7.176 | 2.181 | -0.976 | 6.113 | 3.216 | -0.599 | 1.53 | 75.74 | 135.76 |
| 29 | 2jsa-6 | SER_A_5_C | SER_A_5_O | 6.113 | 3.216 | -0.599 | 4.966 | 2.862 | -0.332 | 1.23 | 77.46 | -162.85 |
| 30 | 2jsa-6 | SER_A_5_C | CYS_A_6_N | 6.113 | 3.216 | -0.599 | 6.533 | 4.471 | -0.589 | 1.32 | 89.57 | 71.50 |
| 31 | 2jsa-6 | CYS_A_6_N | CYS_A_6_H | 6.533 | 4.471 | -0.589 | 7.492 | 4.689 | -0.821 | 1.01 | 103.27 | 12.81 |
| 32 | 2jsa-6 | CYS_A_6_N | CYS_A_6_CA | 6.533 | 4.471 | -0.589 | 5.632 | 5.559 | -0.248 | 1.45 | 76.43 | 129.63 |
| 33 | 2jsa-6 | CYS_A_6_CA | CYS_A_6_HA | 5.632 | 5.559 | -0.248 | 5.326 | 5.438 | 0.791 | 1.09 | 17.57 | -158.42 |
| 34 | 2jsa-6 | CYS_A_6_CA | CYS_A_6_C | 5.632 | 5.559 | -0.248 | 4.391 | 5.451 | -1.137 | 1.53 | 125.51 | -175.03 |
| 35 | 2jsa-6 | CYS_A_6_C | CYS_A_6_O | 4.391 | 5.451 | -1.137 | 3.301 | 5.156 | -0.654 | 1.23 | 66.84 | -164.86 |
| 36 | 2jsa-6 | CYS_A_6_C | GLN_A_7_N | 4.391 | 5.451 | -1.137 | 4.601 | 5.698 | -2.423 | 1.33 | 165.85 | 49.63 |
| 37 | 2jsa-6 | GLN_A_7_N | GLN_A_7_H | 4.601 | 5.698 | -2.423 | 5.526 | 5.938 | -2.751 | 1.01 | 108.94 | 14.55 |
| 38 | 2jsa-6 | GLN_A_7_N | GLN_A_7_CA | 4.601 | 5.698 | -2.423 | 3.514 | 5.631 | -3.384 | 1.45 | 131.43 | -176.47 |
| 39 | 2jsa-6 | GLN_A_7_CA | GLN_A_7_HA | 3.514 | 5.631 | -3.384 | 2.926 | 6.544 | -3.29 | 1.09 | 85.05 | 122.78 |
| 40 | 2jsa-6 | GLN_A_7_CA | GLN_A_7_C | 3.514 | 5.631 | -3.384 | 2.605 | 4.44 | -3.076 | 1.53 | 78.38 | -127.35 |
| 41 | 2jsa-6 | GLN_A_7_C | GLN_A_7_O | 2.605 | 4.44 | -3.076 | 1.401 | 4.606 | -2.883 | 1.23 | 80.98 | 172.15 |
| 42 | 2jsa-6 | GLN_A_7_C | MET_A_8_N | 2.605 | 4.44 | -3.076 | 3.215 | 3.265 | -3.041 | 1.32 | 88.49 | -62.56 |
| 43 | 2jsa-6 | MET_A_8_N | MET_A_8_H | 3.215 | 3.265 | -3.041 | 4.209 | 3.203 | -3.212 | 1.01 | 99.74 | -3.57 |
| 44 | 2jsa-6 | MET_A_8_N | MET_A_8_CA | 3.215 | 3.265 | -3.041 | 2.476 | 2.045 | -2.76 | 1.45 | 78.86 | -121.20 |
| 45 | 2jsa-6 | MET_A_8_CA | MET_A_8_HA | 2.476 | 2.045 | -2.76 | 1.889 | 1.782 | -3.639 | 1.09 | 143.80 | -155.87 |
| 46 | 2jsa-6 | MET_A_8_CA | MET_A_8_C | 2.476 | 2.045 | -2.76 | 1.532 | 2.238 | -1.572 | 1.53 | 39.04 | 168.45 |
| 47 | 2jsa-6 | MET_A_8_C | MET_A_8_O | 1.532 | 2.238 | -1.572 | 0.488 | 1.593 | -1.493 | 1.23 | 86.32 | -148.29 |
| 48 | 2jsa-6 | MET_A_8_C | CYS_A_9_N | 1.532 | 2.238 | -1.572 | 1.933 | 3.129 | -0.677 | 1.33 | 47.51 | 65.77 |
| 49 | 2jsa-6 | CYS_A_9_N | CYS_A_9_H | 1.933 | 3.129 | -0.677 | 2.804 | 3.625 | -0.807 | 1.01 | 97.39 | 29.66 |
| 50 | 2jsa-6 | CYS_A_9_N | CYS_A_9_CA | 1.933 | 3.129 | -0.677 | 1.137 | 3.414 | 0.505 | 1.45 | 35.58 | 160.30 |
| 51 | 2jsa-6 | CYS_A_9_CA | CYS_A_9_HA | 1.137 | 3.414 | 0.505 | 0.897 | 2.469 | 0.993 | 1.09 | 63.41 | -104.25 |
| 52 | 2jsa-6 | CYS_A_9_CA | CYS_A_9_C | 1.137 | 3.414 | 0.505 | -0.168 | 4.076 | 0.055 | 1.53 | 107.09 | 153.10 |
| 53 | 2jsa-6 | CYS_A_9_C | CYS_A_9_O | -0.168 | 4.076 | 0.055 | -1.245 | 3.726 | 0.537 | 1.23 | 66.94 | -162.00 |
| 54 | 2jsa-6 | CYS_A_9_C | GLU_A_10_N | -0.168 | 4.076 | 0.055 | -0.029 | 5.022 | -0.863 | 1.33 | 133.83 | 81.64 |
| 55 | 2jsa-6 | GLU_A_10_N | GLU_A_10_H | -0.029 | 5.022 | -0.863 | 0.888 | 5.258 | -1.215 | 1.01 | 110.39 | 14.43 |
| 56 | 2jsa-6 | GLU_A_10_N | GLU_A_10_CA | -0.029 | 5.022 | -0.863 | -1.183 | 5.736 | -1.382 | 1.45 | 110.93 | 148.25 |
| 57 | 2jsa-6 | GLU_A_10_CA | GLU_A_10_HA | -1.183 | 5.736 | -1.382 | -1.755 | 6.113 | -0.535 | 1.09 | 38.97 | 146.61 |
| 58 | 2jsa-6 | GLU_A_10_CA | GLU_A_10_C | -1.183 | 5.736 | -1.382 | -2.077 | 4.788 | -2.183 | 1.53 | 121.58 | -133.32 |
| 59 | 2jsa-6 | GLU_A_10_C | GLU_A_10_O | -2.077 | 4.788 | -2.183 | -3.185 | 5.154 | -2.572 | 1.23 | 108.44 | 161.72 |
| 60 | 2jsa-6 | GLU_A_10_C | LEU_A_11_N | -2.077 | 4.788 | -2.183 | -1.563 | 3.587 | -2.405 | 1.33 | 99.64 | -66.83 |
| 61 | 2jsa-6 | LEU_A_11_N | LEU_A_11_H | -1.563 | 3.587 | -2.405 | -0.645 | 3.354 | -2.054 | 1.01 | 69.67 | -14.24 |
| 62 | 2jsa-6 | LEU_A_11_N | LEU_A_11_CA | -1.563 | 3.587 | -2.405 | -2.302 | 2.584 | -3.153 | 1.45 | 120.98 | -126.38 |
| 63 | 2jsa-6 | LEU_A_11_CA | LEU_A_11_HA | -2.302 | 2.584 | -3.153 | -2.998 | 3.1 | -3.813 | 1.09 | 127.30 | 143.45 |
| 64 | 2jsa-6 | LEU_A_11_CA | LEU_A_11_C | -2.302 | 2.584 | -3.153 | -3.124 | 1.734 | -2.182 | 1.53 | 50.61 | -134.04 |
| 65 | 2jsa-6 | LEU_A_11_C | LEU_A_11_O | -3.124 | 1.734 | -2.182 | -4.329 | 1.564 | -2.368 | 1.23 | 98.69 | -171.97 |
| 66 | 2jsa-6 | LEU_A_11_C | VAL_A_12_N | -3.124 | 1.734 | -2.182 | -2.441 | 1.223 | -1.169 | 1.32 | 40.10 | -36.80 |
| 67 | 2jsa-6 | VAL_A_12_N | VAL_A_12_H | -2.441 | 1.223 | -1.169 | -1.451 | 1.405 | -1.079 | 1.01 | 84.89 | 10.42 |
| 68 | 2jsa-6 | VAL_A_12_N | VAL_A_12_CA | -2.441 | 1.223 | -1.169 | -3.093 | 0.394 | -0.169 | 1.45 | 46.52 | -128.18 |
| 69 | 2jsa-6 | VAL_A_12_CA | VAL_A_12_HA | -3.093 | 0.394 | -0.169 | -3.6 | -0.426 | -0.678 | 1.09 | 117.83 | -121.73 |
| 70 | 2jsa-6 | VAL_A_12_CA | VAL_A_12_C | -3.093 | 0.394 | -0.169 | -4.149 | 1.225 | 0.565 | 1.53 | 61.36 | 141.80 |
| 71 | 2jsa-6 | VAL_A_12_C | VAL_A_12_O | -4.149 | 1.225 | 0.565 | -5.282 | 0.78 | 0.739 | 1.23 | 81.86 | -158.56 |
| 72 | 2jsa-6 | VAL_A_12_C | VAL_A_13_N | -4.149 | 1.225 | 0.565 | -3.738 | 2.415 | 0.976 | 1.32 | 71.92 | 70.95 |
| 73 | 2jsa-6 | VAL_A_13_N | VAL_A_13_H | -3.738 | 2.415 | 0.976 | -2.79 | 2.715 | 0.799 | 1.01 | 100.09 | 17.56 |
| 74 | 2jsa-6 | VAL_A_13_N | VAL_A_13_CA | -3.738 | 2.415 | 0.976 | -4.634 | 3.311 | 1.687 | 1.45 | 60.70 | 135.00 |
| 75 | 2jsa-6 | VAL_A_13_CA | VAL_A_13_HA | -4.634 | 3.311 | 1.687 | -4.992 | 2.801 | 2.581 | 1.09 | 34.88 | -125.07 |
| 76 | 2jsa-6 | VAL_A_13_CA | VAL_A_13_C | -4.634 | 3.311 | 1.687 | -5.842 | 3.625 | 0.803 | 1.53 | 125.31 | 165.43 |
| 77 | 2jsa-6 | VAL_A_13_C | VAL_A_13_O | -5.842 | 3.625 | 0.803 | -6.983 | 3.38 | 1.193 | 1.23 | 71.52 | -167.88 |
| 78 | 2jsa-6 | VAL_A_13_C | LYS_A_14_N | -5.842 | 3.625 | 0.803 | -5.551 | 4.162 | -0.373 | 1.33 | 152.55 | 61.55 |
| 79 | 2jsa-6 | LYS_A_14_N | LYS_A_14_H | -5.551 | 4.162 | -0.373 | -4.589 | 4.336 | -0.629 | 1.01 | 104.67 | 10.25 |
| 80 | 2jsa-6 | LYS_A_14_N | LYS_A_14_CA | -5.551 | 4.162 | -0.373 | -6.6 | 4.512 | -1.316 | 1.45 | 130.46 | 161.55 |
| 81 | 2jsa-6 | LYS_A_14_CA | LYS_A_14_HA | -6.6 | 4.512 | -1.316 | -7.104 | 5.404 | -0.944 | 1.09 | 70.04 | 119.47 |
| 82 | 2jsa-6 | LYS_A_14_CA | LYS_A_14_C | -6.6 | 4.512 | -1.316 | -7.631 | 3.382 | -1.368 | 1.53 | 91.95 | -132.38 |
| 83 | 2jsa-6 | LYS_A_14_C | LYS_A_14_O | -7.631 | 3.382 | -1.368 | -8.824 | 3.633 | -1.528 | 1.23 | 97.48 | 168.12 |
| 84 | 2jsa-6 | LYS_A_14_C | LYS_A_15_N | -7.631 | 3.382 | -1.368 | -7.132 | 2.163 | -1.232 | 1.32 | 84.11 | -67.74 |
| 85 | 2jsa-6 | LYS_A_15_N | LYS_A_15_H | -7.132 | 2.163 | -1.232 | -6.138 | 2.031 | -1.105 | 1.01 | 82.78 | -7.56 |
| 86 | 2jsa-6 | LYS_A_15_N | LYS_A_15_CA | -7.132 | 2.163 | -1.232 | -7.995 | 0.994 | -1.262 | 1.45 | 91.18 | -126.44 |
| 87 | 2jsa-6 | LYS_A_15_CA | LYS_A_15_HA | -7.995 | 0.994 | -1.262 | -8.76 | 1.158 | -2.02 | 1.09 | 134.09 | 167.90 |
| 88 | 2jsa-6 | LYS_A_15_CA | LYS_A_15_C | -7.995 | 0.994 | -1.262 | -8.7 | 0.852 | 0.088 | 1.53 | 28.04 | -168.61 |
| 89 | 2jsa-6 | LYS_A_15_C | LYS_A_15_O | -8.7 | 0.852 | 0.088 | -9.884 | 0.522 | 0.142 | 1.23 | 87.48 | -164.43 |
| 90 | 2jsa-6 | LYS_A_15_C | TYR_A_16_N | -8.7 | 0.852 | 0.088 | -7.943 | 1.107 | 1.146 | 1.33 | 37.05 | 18.62 |
| 91 | 2jsa-6 | TYR_A_16_N | TYR_A_16_H | -7.943 | 1.107 | 1.146 | -6.975 | 1.371 | 1.027 | 1.01 | 96.76 | 15.26 |
| 92 | 2jsa-6 | TYR_A_16_N | TYR_A_16_CA | -7.943 | 1.107 | 1.146 | -8.482 | 1.012 | 2.492 | 1.45 | 22.13 | -170.00 |
| 93 | 2jsa-6 | TYR_A_16_CA | TYR_A_16_HA | -8.482 | 1.012 | 2.492 | -9.068 | 0.1 | 2.6 | 1.09 | 84.31 | -122.72 |
| 94 | 2jsa-6 | TYR_A_16_CA | TYR_A_16_C | -8.482 | 1.012 | 2.492 | -9.348 | 2.228 | 2.823 | 1.53 | 77.50 | 125.46 |
| 95 | 2jsa-6 | TYR_A_16_C | TYR_A_16_O | -9.348 | 2.228 | 2.823 | -10.571 | 2.121 | 2.904 | 1.23 | 86.23 | -175.00 |
| 96 | 2jsa-6 | TYR_A_16_C | GLU_A_17_N | -9.348 | 2.228 | 2.823 | -8.681 | 3.358 | 3.006 | 1.32 | 82.06 | 59.45 |
| 97 | 2jsa-6 | GLU_A_17_N | GLU_A_17_H | -8.681 | 3.358 | 3.006 | -7.674 | 3.373 | 2.924 | 1.01 | 94.65 | 0.85 |
| 98 | 2jsa-6 | GLU_A_17_N | GLU_A_17_CA | -8.681 | 3.358 | 3.006 | -9.376 | 4.595 | 3.326 | 1.45 | 77.29 | 119.33 |
| 99 | 2jsa-6 | GLU_A_17_CA | GLU_A_17_HA | -9.376 | 4.595 | 3.326 | -9.646 | 4.566 | 4.382 | 1.09 | 14.42 | -173.87 |
| 100 | 2jsa-6 | GLU_A_17_CA | GLU_A_17_C | -9.376 | 4.595 | 3.326 | -10.661 | 4.71 | 2.504 | 1.53 | 122.50 | 174.89 |
| 101 | 2jsa-6 | GLU_A_17_C | GLU_A_17_O | -10.661 | 4.71 | 2.504 | -11.678 | 5.192 | 2.999 | 1.23 | 66.26 | 154.64 |
| 102 | 2jsa-6 | GLU_A_17_C | GLY_A_18_N | -10.661 | 4.71 | 2.504 | -10.573 | 4.259 | 1.261 | 1.33 | 159.71 | -78.96 |
| 103 | 2jsa-6 | GLY_A_18_N | GLY_A_18_H | -10.573 | 4.259 | 1.261 | -9.703 | 3.875 | 0.92 | 1.01 | 109.73 | -23.82 |
| 104 | 2jsa-6 | GLY_A_18_N | GLY_A_18_CA | -10.573 | 4.259 | 1.261 | -11.716 | 4.305 | 0.365 | 1.45 | 128.07 | 177.70 |
| 105 | 2jsa-6 | GLY_A_18_CA | GLY_A_18_C | -11.716 | 4.305 | 0.365 | -12.088 | 2.904 | -0.122 | 1.53 | 108.57 | -104.87 |
| 106 | 2jsa-6 | GLY_A_18_C | GLY_A_18_O | -12.088 | 2.904 | -0.122 | -12.307 | 2.696 | -1.314 | 1.23 | 165.78 | -136.48 |
| 107 | 2jsa-6 | GLY_A_18_C | SER_A_19_N | -12.088 | 2.904 | -0.122 | -12.149 | 1.98 | 0.825 | 1.32 | 44.36 | -93.78 |
| 108 | 2jsa-6 | SER_A_19_N | SER_A_19_H | -12.149 | 1.98 | 0.825 | -11.957 | 2.224 | 1.787 | 1.01 | 17.89 | 51.80 |
| 109 | 2jsa-6 | SER_A_19_N | SER_A_19_CA | -12.149 | 1.98 | 0.825 | -12.491 | 0.603 | 0.507 | 1.45 | 102.63 | -103.95 |
| 110 | 2jsa-6 | SER_A_19_CA | SER_A_19_HA | -12.491 | 0.603 | 0.507 | -12.083 | 0.368 | -0.476 | 1.09 | 154.41 | -29.94 |

| | | | | | | | | | | |
|---|---|---|---|---|---|---|---|---|---|---|
| 111 | 2jsa-6 | SER_A_19_CA | SER_A_19_C | -12.491 | 0.603 | 0.507 | -14.011 | 0.44 | 0.459 | 1.53 | 91.80 | -173.88 |
| 112 | 2jsa-6 | SER_A_19_C | SER_A_19_O | -14.011 | 0.44 | 0.459 | -14.552 | -0.089 | -0.51 | 1.23 | 142.02 | -135.64 |
| 113 | 2jsa-6 | SER_A_19_C | ALA_A_20_N | -14.011 | 0.44 | 0.459 | -14.657 | 0.904 | 1.519 | 1.33 | 36.88 | 144.31 |
| 114 | 2jsa-6 | ALA_A_20_N | ALA_A_20_H | -14.657 | 0.904 | 1.519 | -14.146 | 1.327 | 2.281 | 1.01 | 41.04 | 39.62 |
| 115 | 2jsa-6 | ALA_A_20_N | ALA_A_20_CA | -14.657 | 0.904 | 1.519 | -16.105 | 0.815 | 1.61 | 1.45 | 86.41 | -176.48 |
| 116 | 2jsa-6 | ALA_A_20_CA | ALA_A_20_HA | -16.105 | 0.815 | 1.61 | -16.458 | 1.57 | 2.313 | 1.09 | 49.85 | 115.06 |
| 117 | 2jsa-6 | ALA_A_20_CA | ALA_A_20_C | -16.105 | 0.815 | 1.61 | -16.495 | -0.564 | 2.145 | 1.53 | 69.53 | -105.79 |
| 118 | 2jsa-6 | ALA_A_20_C | ALA_A_20_O | -16.495 | -0.564 | 2.145 | -17.66 | -0.808 | 2.455 | 1.23 | 75.40 | -168.17 |
| 119 | 2jsa-6 | ALA_A_20_C | ASP_A_21_N | -16.495 | -0.564 | 2.145 | -15.497 | -1.432 | 2.237 | 1.33 | 86.02 | -41.01 |
| 120 | 2jsa-6 | ASP_A_21_N | ASP_A_21_H | -15.497 | -1.432 | 2.237 | -14.562 | -1.164 | 1.963 | 1.01 | 105.73 | 15.99 |
| 121 | 2jsa-6 | ASP_A_21_N | ASP_A_21_CA | -15.497 | -1.432 | 2.237 | -15.721 | -2.78 | 2.73 | 1.45 | 70.16 | -99.43 |
| 122 | 2jsa-6 | ASP_A_21_CA | ASP_A_21_HA | -15.721 | -2.78 | 2.73 | -16.789 | -2.928 | 2.891 | 1.09 | 81.51 | -172.11 |
| 123 | 2jsa-6 | ASP_A_21_CA | ASP_A_21_C | -15.721 | -2.78 | 2.73 | -14.966 | -2.969 | 4.047 | 1.53 | 30.58 | -14.05 |
| 124 | 2jsa-6 | ASP_A_21_C | ASP_A_21_O | -14.966 | -2.969 | 4.047 | -15.217 | -3.926 | 4.777 | 1.23 | 53.58 | -104.70 |
| 125 | 2jsa-6 | ASP_A_21_C | LYS_A_22_N | -14.966 | -2.969 | 4.047 | -14.058 | -2.042 | 4.31 | 1.32 | 78.54 | 45.59 |
| 126 | 2jsa-6 | LYS_A_22_N | LYS_A_22_H | -14.058 | -2.042 | 4.31 | -13.904 | -1.282 | 3.663 | 1.01 | 129.84 | 78.55 |
| 127 | 2jsa-6 | LYS_A_22_N | LYS_A_22_CA | -14.058 | -2.042 | 4.31 | -13.266 | -2.094 | 5.527 | 1.45 | 33.11 | -3.76 |
| 128 | 2jsa-6 | LYS_A_22_CA | LYS_A_22_HA | -13.266 | -2.094 | 5.527 | -12.429 | -1.404 | 5.415 | 1.09 | 95.89 | 39.50 |
| 129 | 2jsa-6 | LYS_A_22_CA | LYS_A_22_C | -13.266 | -2.094 | 5.527 | -12.689 | -3.501 | 5.695 | 1.53 | 83.70 | -67.70 |
| 130 | 2jsa-6 | LYS_A_22_C | LYS_A_22_O | -12.689 | -3.501 | 5.695 | -13.075 | -4.229 | 6.608 | 1.23 | 42.07 | -117.93 |
| 131 | 2jsa-6 | LYS_A_22_C | ASP_A_23_N | -12.689 | -3.501 | 5.695 | -11.776 | -3.843 | 4.798 | 1.32 | 132.62 | -20.54 |
| 132 | 2jsa-6 | ASP_A_23_N | ASP_A_23_H | -11.776 | -3.843 | 4.798 | -11.508 | -3.194 | 4.072 | 1.01 | 135.96 | 67.56 |
| 133 | 2jsa-6 | ASP_A_23_N | ASP_A_23_CA | -11.776 | -3.843 | 4.798 | -11.143 | -5.15 | 4.836 | 1.45 | 88.50 | -64.16 |
| 134 | 2jsa-6 | ASP_A_23_CA | ASP_A_23_HA | -11.143 | -5.15 | 4.836 | -11.09 | -5.491 | 5.87 | 1.09 | 18.46 | -81.17 |
| 135 | 2jsa-6 | ASP_A_23_CA | ASP_A_23_C | -11.143 | -5.15 | 4.836 | -9.733 | -5.046 | 4.25 | 1.53 | 112.51 | 4.22 |
| 136 | 2jsa-6 | ASP_A_23_C | ASP_A_23_O | -9.733 | -5.046 | 4.25 | -9.568 | -4.742 | 3.069 | 1.23 | 163.68 | 61.51 |
| 137 | 2jsa-6 | ASP_A_23_C | ALA_A_24_N | -9.733 | -5.046 | 4.25 | -8.752 | -5.304 | 5.103 | 1.33 | 49.94 | -14.73 |
| 138 | 2jsa-6 | ALA_A_24_N | ALA_A_24_H | -8.752 | -5.304 | 5.103 | -8.961 | -5.547 | 6.061 | 1.01 | 18.50 | -130.70 |
| 139 | 2jsa-6 | ALA_A_24_N | ALA_A_24_CA | -8.752 | -5.304 | 5.103 | -7.362 | -5.243 | 4.684 | 1.45 | 106.76 | 2.51 |
| 140 | 2jsa-6 | ALA_A_24_CA | ALA_A_24_HA | -7.362 | -5.243 | 4.684 | -7.181 | -4.279 | 4.208 | 1.09 | 115.89 | 79.37 |
| 141 | 2jsa-6 | ALA_A_24_CA | ALA_A_24_C | -7.362 | -5.243 | 4.684 | -7.093 | -6.347 | 3.66 | 1.53 | 132.02 | -76.31 |
| 142 | 2jsa-6 | ALA_A_24_C | ALA_A_24_O | -7.093 | -6.347 | 3.66 | -6.026 | -6.385 | 3.048 | 1.23 | 119.82 | -2.04 |
| 143 | 2jsa-6 | ALA_A_24_C | ASN_A_25_N | -7.093 | -6.347 | 3.66 | -8.078 | -7.22 | 3.505 | 1.33 | 96.72 | -138.45 |
| 144 | 2jsa-6 | ASN_A_25_N | ASN_A_25_H | -8.078 | -7.22 | 3.505 | -8.927 | -7.128 | 4.043 | 1.01 | 57.79 | 173.82 |
| 145 | 2jsa-6 | ASN_A_25_N | ASN_A_25_CA | -8.078 | -7.22 | 3.505 | -7.962 | -8.322 | 2.565 | 1.45 | 130.31 | -83.99 |
| 146 | 2jsa-6 | ASN_A_25_CA | ASN_A_25_HA | -7.962 | -8.322 | 2.565 | -6.943 | -8.71 | 2.595 | 1.09 | 88.42 | -20.85 |
| 147 | 2jsa-6 | ASN_A_25_CA | ASN_A_25_C | -7.962 | -8.322 | 2.565 | -8.286 | -7.821 | 1.156 | 1.53 | 157.05 | 122.89 |
| 148 | 2jsa-6 | ASN_A_25_C | ASN_A_25_O | -8.286 | -7.821 | 1.156 | -7.528 | -8.061 | 0.219 | 1.23 | 139.68 | -17.57 |
| 149 | 2jsa-6 | ASN_A_25_C | VAL_A_26_N | -8.286 | -7.821 | 1.156 | -9.415 | -7.135 | 1.051 | 1.33 | 94.54 | 148.72 |
| 150 | 2jsa-6 | VAL_A_26_N | VAL_A_26_H | -9.415 | -7.135 | 1.051 | -9.99 | -6.976 | 1.865 | 1.01 | 36.24 | 164.54 |
| 151 | 2jsa-6 | VAL_A_26_N | VAL_A_26_CA | -9.415 | -7.135 | 1.051 | -9.849 | -6.599 | -0.226 | 1.45 | 151.63 | 129.00 |
| 152 | 2jsa-6 | VAL_A_26_CA | VAL_A_26_HA | -9.849 | -6.599 | -0.226 | -9.652 | -7.345 | -0.996 | 1.09 | 134.94 | -75.21 |
| 153 | 2jsa-6 | VAL_A_26_CA | VAL_A_26_C | -9.849 | -6.599 | -0.226 | -9.03 | -5.35 | -0.556 | 1.53 | 102.46 | 56.75 |
| 154 | 2jsa-6 | VAL_A_26_C | VAL_A_26_O | -9.03 | -5.35 | -0.556 | -9.104 | -4.83 | -1.668 | 1.23 | 154.72 | 98.10 |
| 155 | 2jsa-6 | VAL_A_26_C | ILE_A_27_N | -9.03 | -5.35 | -0.556 | -8.267 | -4.904 | 0.432 | 1.33 | 41.81 | 30.31 |
| 156 | 2jsa-6 | ILE_A_27_N | ILE_A_27_H | -8.267 | -4.904 | 0.432 | -8.254 | -5.383 | 1.321 | 1.01 | 28.32 | -88.45 |
| 157 | 2jsa-6 | ILE_A_27_N | ILE_A_27_CA | -8.267 | -4.904 | 0.432 | -7.436 | -3.725 | 0.262 | 1.45 | 96.72 | 54.82 |
| 158 | 2jsa-6 | ILE_A_27_CA | ILE_A_27_HA | -7.436 | -3.725 | 0.262 | -7.9 | -3.091 | -0.494 | 1.09 | 133.90 | 126.20 |
| 159 | 2jsa-6 | ILE_A_27_CA | ILE_A_27_C | -7.436 | -3.725 | 0.262 | -6.062 | -4.147 | -0.264 | 1.53 | 110.10 | -17.07 |
| 160 | 2jsa-6 | ILE_A_27_C | ILE_A_27_O | -6.062 | -4.147 | -0.264 | -5.711 | -3.846 | -1.405 | 1.23 | 157.94 | 40.61 |
| 161 | 2jsa-6 | ILE_A_27_C | LYS_A_28_N | -6.062 | -4.147 | -0.264 | -5.323 | -4.838 | 0.591 | 1.32 | 49.80 | -43.08 |
| 162 | 2jsa-6 | LYS_A_28_N | LYS_A_28_H | -5.323 | -4.838 | 0.591 | -5.676 | -5.05 | 1.513 | 1.01 | 24.07 | -149.01 |
| 163 | 2jsa-6 | LYS_A_28_N | LYS_A_28_CA | -5.323 | -4.838 | 0.591 | -3.996 | -5.304 | 0.226 | 1.45 | 104.55 | -19.35 |
| 164 | 2jsa-6 | LYS_A_28_CA | LYS_A_28_HA | -3.996 | -5.304 | 0.226 | -3.334 | -4.439 | 0.181 | 1.09 | 92.37 | 52.57 |
| 165 | 2jsa-6 | LYS_A_28_CA | LYS_A_28_C | -3.996 | -5.304 | 0.226 | -4.045 | -5.925 | -1.171 | 1.53 | 155.97 | -94.51 |
| 166 | 2jsa-6 | LYS_A_28_C | LYS_A_28_O | -4.045 | -5.925 | -1.171 | -3.163 | -5.682 | -1.994 | 1.23 | 131.97 | 15.40 |
| 167 | 2jsa-6 | LYS_A_28_C | LYS_A_29_N | -4.045 | -5.925 | -1.171 | -5.082 | -6.717 | -1.396 | 1.32 | 99.78 | -142.63 |
| 168 | 2jsa-6 | LYS_A_29_N | LYS_A_29_H | -5.082 | -6.717 | -1.396 | -5.768 | -6.874 | -0.672 | 1.01 | 44.19 | -167.11 |
| 169 | 2jsa-6 | LYS_A_29_N | LYS_A_29_CA | -5.082 | -6.717 | -1.396 | -5.258 | -7.375 | -2.68 | 1.45 | 152.06 | -104.97 |
| 170 | 2jsa-6 | LYS_A_29_CA | LYS_A_29_HA | -5.258 | -7.375 | -2.68 | -4.334 | -7.904 | -2.917 | 1.09 | 102.55 | -29.79 |
| 171 | 2jsa-6 | LYS_A_29_CA | LYS_A_29_C | -5.258 | -7.375 | -2.68 | -5.479 | -6.318 | -3.764 | 1.53 | 135.11 | 101.81 |
| 172 | 2jsa-6 | LYS_A_29_C | LYS_A_29_O | -5.479 | -6.318 | -3.764 | -4.777 | -6.303 | -4.773 | 1.23 | 145.17 | 1.22 |
| 173 | 2jsa-6 | LYS_A_29_C | ASP_A_30_N | -5.479 | -6.318 | -3.764 | -6.46 | -5.461 | -3.518 | 1.33 | 79.31 | 138.86 |
| 174 | 2jsa-6 | ASP_A_30_N | ASP_A_30_H | -6.46 | -5.461 | -3.518 | -6.998 | -5.534 | -2.666 | 1.01 | 32.51 | -172.27 |
| 175 | 2jsa-6 | ASP_A_30_N | ASP_A_30_CA | -6.46 | -5.461 | -3.518 | -6.783 | -4.404 | -4.461 | 1.45 | 130.47 | 106.99 |
| 176 | 2jsa-6 | ASP_A_30_CA | ASP_A_30_HA | -6.783 | -4.404 | -4.461 | -7.295 | -4.837 | -5.32 | 1.09 | 142.02 | -139.78 |
| 177 | 2jsa-6 | ASP_A_30_CA | ASP_A_30_C | -6.783 | -4.404 | -4.461 | -5.492 | -3.72 | -4.915 | 1.53 | 107.26 | 27.92 |
| 178 | 2jsa-6 | ASP_A_30_C | ASP_A_30_O | -5.492 | -3.72 | -4.915 | -5.218 | -3.635 | -6.111 | 1.23 | 166.51 | 17.23 |
| 179 | 2jsa-6 | ASP_A_30_C | PHE_A_31_N | -5.492 | -3.72 | -4.915 | -4.733 | -3.249 | -3.937 | 1.32 | 42.41 | 31.82 |
| 180 | 2jsa-6 | PHE_A_31_N | PHE_A_31_H | -4.733 | -3.249 | -3.937 | -5.02 | -3.354 | -2.975 | 1.01 | 17.62 | -159.90 |
| 181 | 2jsa-6 | PHE_A_31_N | PHE_A_31_CA | -4.733 | -3.249 | -3.937 | -3.478 | -2.575 | -4.221 | 1.45 | 101.27 | 28.24 |
| 182 | 2jsa-6 | PHE_A_31_CA | PHE_A_31_HA | -3.478 | -2.575 | -4.221 | -3.67 | -1.542 | -4.512 | 1.09 | 105.48 | 100.53 |
| 183 | 2jsa-6 | PHE_A_31_CA | PHE_A_31_C | -3.478 | -2.575 | -4.221 | -2.712 | -3.29 | -5.337 | 1.53 | 136.80 | -43.03 |
| 184 | 2jsa-6 | PHE_A_31_C | PHE_A_31_O | -2.712 | -3.29 | -5.337 | -2.307 | -2.665 | -6.315 | 1.23 | 142.71 | 57.06 |
| 185 | 2jsa-6 | PHE_A_31_C | ASP_A_32_N | -2.712 | -3.29 | -5.337 | -2.538 | -4.59 | -5.151 | 1.32 | 81.93 | -82.38 |
| 186 | 2jsa-6 | ASP_A_32_N | ASP_A_32_H | -2.538 | -4.59 | -5.151 | -2.898 | -5.038 | -4.321 | 1.01 | 34.70 | -128.78 |
| 187 | 2jsa-6 | ASP_A_32_N | ASP_A_32_CA | -2.538 | -4.59 | -5.151 | -1.828 | -5.396 | -6.129 | 1.45 | 132.32 | -48.62 |
| 188 | 2jsa-6 | ASP_A_32_CA | ASP_A_32_HA | -1.828 | -5.396 | -6.129 | -0.806 | -5.026 | -6.221 | 1.09 | 94.84 | 19.90 |
| 189 | 2jsa-6 | ASP_A_32_CA | ASP_A_32_C | -1.828 | -5.396 | -6.129 | -2.541 | -5.299 | -7.479 | 1.53 | 151.94 | 172.25 |
| 190 | 2jsa-6 | ASP_A_32_C | ASP_A_32_O | -2.541 | -5.299 | -7.479 | -1.895 | -5.252 | -8.524 | 1.23 | 148.21 | 4.16 |
| 191 | 2jsa-6 | ASP_A_32_C | ALA_A_33_N | -2.541 | -5.299 | -7.479 | -3.864 | -5.271 | -7.412 | 1.32 | 87.10 | 178.79 |
| 192 | 2jsa-6 | ALA_A_33_N | ALA_A_33_H | -3.864 | -5.271 | -7.412 | -4.329 | -5.314 | -6.516 | 1.01 | 27.53 | -174.72 |
| 193 | 2jsa-6 | ALA_A_33_N | ALA_A_33_CA | -3.864 | -5.271 | -7.412 | -4.672 | -5.179 | -8.616 | 1.45 | 145.96 | 173.50 |
| 194 | 2jsa-6 | ALA_A_33_CA | ALA_A_33_HA | -4.672 | -5.179 | -8.616 | -4.503 | -6.07 | -9.22 | 1.09 | 123.66 | -79.26 |
| 195 | 2jsa-6 | ALA_A_33_CA | ALA_A_33_C | -4.672 | -5.179 | -8.616 | -4.237 | -3.956 | -9.426 | 1.53 | 121.96 | 70.42 |
| 196 | 2jsa-6 | ALA_A_33_C | ALA_A_33_O | -4.237 | -3.956 | -9.426 | -3.975 | -4.062 | -10.624 | 1.23 | 166.73 | -22.03 |
| 197 | 2jsa-6 | ALA_A_33_C | GLU_A_34_N | -4.237 | -3.956 | -9.426 | -4.171 | -2.824 | -8.74 | 1.33 | 58.83 | 86.66 |
| 198 | 2jsa-6 | GLU_A_34_N | GLU_A_34_H | -4.171 | -2.824 | -8.74 | -4.398 | -2.811 | -7.756 | 1.01 | 13.01 | 176.72 |
| 199 | 2jsa-6 | GLU_A_34_N | GLU_A_34_CA | -4.171 | -2.824 | -8.74 | -3.772 | -1.584 | -9.382 | 1.45 | 116.24 | 72.16 |
| 200 | 2jsa-6 | GLU_A_34_CA | GLU_A_34_HA | -3.772 | -1.584 | -9.382 | -4.33 | -1.491 | -10.313 | 1.09 | 148.72 | 170.54 |
| 201 | 2jsa-6 | GLU_A_34_CA | GLU_A_34_C | -3.772 | -1.584 | -9.382 | -2.279 | -1.616 | -9.717 | 1.53 | 102.64 | -1.23 |
| 202 | 2jsa-6 | GLU_A_34_C | GLU_A_34_O | -2.279 | -1.616 | -9.717 | -1.901 | -1.567 | -10.887 | 1.23 | 161.96 | 7.39 |
| 203 | 2jsa-6 | GLU_A_34_C | CYS_A_35_N | -2.279 | -1.616 | -9.717 | -1.472 | -1.699 | -8.671 | 1.32 | 37.80 | -5.87 |
| 204 | 2jsa-6 | CYS_A_35_N | CYS_A_35_H | -1.472 | -1.699 | -8.671 | -1.853 | -1.736 | -7.736 | 1.01 | 22.26 | -174.45 |
| 205 | 2jsa-6 | CYS_A_35_N | CYS_A_35_CA | -1.472 | -1.699 | -8.671 | -0.028 | -1.739 | -8.84 | 1.45 | 96.67 | -1.59 |
| 206 | 2jsa-6 | CYS_A_35_CA | CYS_A_35_HA | -0.028 | -1.739 | -8.84 | 0.323 | -0.726 | -9.039 | 1.09 | 100.52 | 70.89 |

| | | | | | | | | | | | |
|---|---|---|---|---|---|---|---|---|---|---|---|
| 207 | 2jsa-6 | CYS_A_35_CA | CYS_A_35_C | -0.028 | -1.739 | -8.84 | 0.288 | -2.611 | -10.056 | 1.53 | 142.67 | -70.08 |
| 208 | 2jsa-6 | CYS_A_35_C | CYS_A_35_O | 0.288 | -2.611 | -10.056 | 1.251 | -2.351 | -10.777 | 1.23 | 125.86 | 15.11 |
| 209 | 2jsa-6 | CYS_A_35_C | LYS_A_36_N | 0.288 | -2.611 | -10.056 | -0.539 | -3.628 | -10.247 | 1.32 | 98.29 | -129.12 |
| 210 | 2jsa-6 | LYS_A_36_N | LYS_A_36_H | -0.539 | -3.628 | -10.247 | -1.31 | -3.782 | -9.613 | 1.01 | 51.12 | -168.70 |
| 211 | 2jsa-6 | LYS_A_36_N | LYS_A_36_CA | -0.539 | -3.628 | -10.247 | -0.359 | -4.54 | -11.364 | 1.45 | 140.23 | -78.84 |
| 212 | 2jsa-6 | LYS_A_36_CA | LYS_A_36_HA | -0.359 | -4.54 | -11.364 | 0.642 | -4.967 | -11.293 | 1.09 | 86.27 | -23.10 |
| 213 | 2jsa-6 | LYS_A_36_CA | LYS_A_36_C | -0.359 | -4.54 | -11.364 | -0.438 | -3.755 | -12.676 | 1.53 | 148.98 | 95.75 |
| 214 | 2jsa-6 | LYS_A_36_C | LYS_A_36_O | -0.438 | -3.755 | -12.676 | 0.586 | -3.466 | -13.293 | 1.23 | 120.11 | 15.76 |
| 215 | 2jsa-6 | LYS_A_36_C | LYS_A_37_N | -0.438 | -3.755 | -12.676 | -1.664 | -3.431 | -13.061 | 1.33 | 106.89 | 165.20 |
| 216 | 2jsa-6 | LYS_A_37_N | LYS_A_37_H | -1.664 | -3.431 | -13.061 | -2.46 | -3.701 | -12.502 | 1.01 | 56.37 | -161.26 |
| 217 | 2jsa-6 | LYS_A_37_N | LYS_A_37_CA | -1.664 | -3.431 | -13.061 | -1.89 | -2.686 | -14.287 | 1.45 | 147.58 | 106.88 |
| 218 | 2jsa-6 | LYS_A_37_CA | LYS_A_37_HA | -1.89 | -2.686 | -14.287 | -1.49 | -3.271 | -15.115 | 1.09 | 139.44 | -55.64 |
| 219 | 2jsa-6 | LYS_A_37_CA | LYS_A_37_C | -1.89 | -2.686 | -14.287 | -1.112 | -1.37 | -14.23 | 1.53 | 87.86 | 59.41 |
| 220 | 2jsa-6 | LYS_A_37_C | LYS_A_37_O | -1.112 | -1.37 | -14.23 | -0.735 | -0.822 | -15.264 | 1.23 | 147.25 | 55.47 |
| 221 | 2jsa-6 | LYS_A_37_C | LEU_A_38_N | -1.112 | -1.37 | -14.23 | -0.897 | -0.9 | -13.01 | 1.32 | 22.96 | 65.42 |
| 222 | 2jsa-6 | LEU_A_38_N | LEU_A_38_H | -0.897 | -0.9 | -13.01 | -1.237 | -1.403 | -12.203 | 1.01 | 36.96 | -124.06 |
| 223 | 2jsa-6 | LEU_A_38_N | LEU_A_38_CA | -0.897 | -0.9 | -13.01 | -0.171 | 0.342 | -12.804 | 1.45 | 81.85 | 59.69 |
| 224 | 2jsa-6 | LEU_A_38_CA | LEU_A_38_HA | -0.171 | 0.342 | -12.804 | -0.616 | 1.103 | -13.445 | 1.09 | 126.02 | 120.32 |
| 225 | 2jsa-6 | LEU_A_38_CA | LEU_A_38_C | -0.171 | 0.342 | -12.804 | 1.283 | 0.157 | -13.243 | 1.53 | 106.67 | -7.25 |
| 226 | 2jsa-6 | LEU_A_38_C | LEU_A_38_O | 1.283 | 0.157 | -13.243 | 1.778 | 0.9 | -14.089 | 1.23 | 133.46 | 56.33 |
| 227 | 2jsa-6 | LEU_A_38_C | PHE_A_39_N | 1.283 | 0.157 | -13.243 | 1.925 | -0.837 | -12.648 | 1.32 | 63.31 | -57.14 |
| 228 | 2jsa-6 | PHE_A_39_N | PHE_A_39_H | 1.925 | -0.837 | -12.648 | 1.454 | -1.407 | -11.961 | 1.01 | 47.10 | -129.57 |
| 229 | 2jsa-6 | PHE_A_39_N | PHE_A_39_CA | 1.925 | -0.837 | -12.648 | 3.312 | -1.129 | -12.967 | 1.45 | 102.68 | -11.89 |
| 230 | 2jsa-6 | PHE_A_39_CA | PHE_A_39_HA | 3.312 | -1.129 | -12.967 | 3.686 | -0.403 | -13.689 | 1.09 | 131.48 | 62.74 |
| 231 | 2jsa-6 | PHE_A_39_CA | PHE_A_39_C | 3.312 | -1.129 | -12.967 | 3.459 | -2.542 | -13.535 | 1.53 | 111.79 | -84.06 |
| 232 | 2jsa-6 | PHE_A_39_C | PHE_A_39_O | 3.459 | -2.542 | -13.535 | 4.294 | -3.317 | -13.073 | 1.23 | 67.93 | -42.87 |
| 233 | 2jsa-6 | PHE_A_39_C | HIS_A_40_N | 3.459 | -2.542 | -13.535 | 2.633 | -2.835 | -14.529 | 1.33 | 138.60 | -160.47 |
| 234 | 2jsa-6 | HIS_A_40_N | HIS_A_40_H | 2.633 | -2.835 | -14.529 | 1.97 | -2.148 | -14.858 | 1.01 | 109.01 | 133.98 |
| 235 | 2jsa-6 | HIS_A_40_N | HIS_A_40_CA | 2.633 | -2.835 | -14.529 | 2.66 | -4.141 | -15.165 | 1.45 | 115.96 | -88.82 |
| 236 | 2jsa-6 | HIS_A_40_CA | HIS_A_40_HA | 2.66 | -4.141 | -15.165 | 2.598 | -4.897 | -14.383 | 1.09 | 44.13 | -94.69 |
| 237 | 2jsa-6 | HIS_A_40_CA | HIS_A_40_C | 2.66 | -4.141 | -15.165 | 3.992 | -4.329 | -15.894 | 1.53 | 118.45 | -8.03 |
| 238 | 2jsa-6 | HIS_A_40_C | HIS_A_40_O | 3.992 | -4.329 | -15.894 | 4.603 | -5.393 | -15.811 | 1.23 | 86.13 | -60.13 |
| 239 | 2jsa-6 | HIS_A_40_C | THR_A_41_N | 3.992 | -4.329 | -15.894 | 4.401 | -3.28 | -16.591 | 1.32 | 121.76 | 68.70 |
| 240 | 2jsa-6 | THR_A_41_N | THR_A_41_H | 4.401 | -3.28 | -16.591 | 3.847 | -2.436 | -16.613 | 1.01 | 91.25 | 123.28 |
| 241 | 2jsa-6 | THR_A_41_N | THR_A_41_CA | 4.401 | -3.28 | -16.591 | 5.649 | -3.316 | -17.335 | 1.45 | 120.79 | -1.65 |
| 242 | 2jsa-6 | THR_A_41_CA | THR_A_41_HA | 5.649 | -3.316 | -17.335 | 5.651 | -4.199 | -17.974 | 1.09 | 125.89 | -89.87 |
| 243 | 2jsa-6 | THR_A_41_CA | THR_A_41_C | 5.649 | -3.316 | -17.335 | 6.837 | -3.426 | -16.377 | 1.53 | 51.24 | -5.29 |
| 244 | 2jsa-6 | THR_A_41_C | THR_A_41_O | 6.837 | -3.426 | -16.377 | 7.816 | -4.11 | -16.673 | 1.23 | 103.92 | -34.94 |
| 245 | 2jsa-6 | THR_A_41_C | ILE_A_42_N | 6.837 | -3.426 | -16.377 | 6.712 | -2.744 | -15.248 | 1.32 | 31.56 | 100.39 |
| 246 | 2jsa-6 | ILE_A_42_N | ILE_A_42_H | 6.712 | -2.744 | -15.248 | 5.877 | -2.204 | -15.069 | 1.01 | 79.80 | 147.11 |
| 247 | 2jsa-6 | ILE_A_42_N | ILE_A_42_CA | 6.712 | -2.744 | -15.248 | 7.764 | -2.756 | -14.245 | 1.45 | 46.37 | -0.65 |
| 248 | 2jsa-6 | ILE_A_42_CA | ILE_A_42_HA | 7.764 | -2.756 | -14.245 | 8.638 | -2.257 | -14.665 | 1.09 | 112.65 | 29.72 |
| 249 | 2jsa-6 | ILE_A_42_CA | ILE_A_42_C | 7.764 | -2.756 | -14.245 | 8.156 | -4.204 | -13.943 | 1.53 | 78.62 | -74.85 |
| 250 | 2jsa-6 | ILE_A_42_C | ILE_A_42_O | 8.156 | -4.204 | -13.943 | 7.298 | -5.084 | -13.883 | 1.23 | 87.21 | -134.27 |
| 251 | 2jsa-6 | ILE_A_42_C | PRO_A_43_N | 8.156 | -4.204 | -13.943 | 9.487 | -4.411 | -13.755 | 1.36 | 82.05 | -8.84 |
| 252 | 2jsa-6 | PRO_A_43_N | PRO_A_43_CA | 9.487 | -4.411 | -13.755 | 10.004 | -5.737 | -13.459 | 1.45 | 78.25 | -68.70 |
| 253 | 2jsa-6 | PRO_A_43_CA | PRO_A_43_HA | 10.004 | -5.737 | -13.459 | 9.547 | -6.472 | -14.121 | 1.09 | 127.41 | -121.87 |
| 254 | 2jsa-6 | PRO_A_43_CA | PRO_A_43_C | 10.004 | -5.737 | -13.459 | 9.703 | -6.133 | -12.013 | 1.53 | 18.98 | -127.24 |
| 255 | 2jsa-6 | PRO_A_43_C | PRO_A_43_O | 9.703 | -6.133 | -12.013 | 8.692 | -6.779 | -11.74 | 1.23 | 77.18 | -147.42 |
| 256 | 2jsa-6 | PRO_A_43_C | PHE_A_44_N | 9.703 | -6.133 | -12.013 | 10.598 | -5.728 | -11.124 | 1.32 | 47.86 | 24.35 |
| 257 | 2jsa-6 | PHE_A_44_N | PHE_A_44_H | 10.598 | -5.728 | -11.124 | 11.406 | -5.199 | -11.42 | 1.01 | 107.04 | 33.21 |
| 258 | 2jsa-6 | PHE_A_44_N | PHE_A_44_CA | 10.598 | -5.728 | -11.124 | 10.441 | -6.033 | -9.711 | 1.45 | 13.65 | -117.24 |
| 259 | 2jsa-6 | PHE_A_44_CA | PHE_A_44_HA | 10.441 | -6.033 | -9.711 | 10.268 | -7.101 | -9.58 | 1.09 | 83.10 | -99.20 |
| 260 | 2jsa-6 | PHE_A_44_CA | PHE_A_44_C | 10.441 | -6.033 | -9.711 | 9.277 | -5.244 | -9.107 | 1.53 | 66.76 | 145.87 |
| 261 | 2jsa-6 | PHE_A_44_C | PHE_A_44_O | 9.277 | -5.244 | -9.107 | 8.979 | -5.382 | -7.921 | 1.23 | 15.48 | -155.15 |
| 262 | 2jsa-6 | PHE_A_44_C | GLY_A_45_N | 9.277 | -5.244 | -9.107 | 8.651 | -4.435 | -9.949 | 1.32 | 129.46 | 127.73 |
| 263 | 2jsa-6 | GLY_A_45_N | GLY_A_45_H | 8.651 | -4.435 | -9.949 | 8.951 | -4.373 | -10.911 | 1.01 | 162.34 | 11.68 |
| 264 | 2jsa-6 | GLY_A_45_N | GLY_A_45_CA | 8.651 | -4.435 | -9.949 | 7.528 | -3.624 | -9.512 | 1.45 | 72.49 | 144.16 |
| 265 | 2jsa-6 | GLY_A_45_CA | GLY_A_45_C | 7.528 | -3.624 | -9.512 | 6.434 | -4.493 | -8.889 | 1.53 | 65.97 | -141.54 |
| 266 | 2jsa-6 | GLY_A_45_C | GLY_A_45_O | 6.434 | -4.493 | -8.889 | 6.117 | -4.348 | -7.709 | 1.23 | 16.46 | 155.42 |
| 267 | 2jsa-6 | GLY_A_45_C | THR_A_46_N | 6.434 | -4.493 | -8.889 | 5.887 | -5.379 | -9.709 | 1.33 | 128.22 | -121.69 |
| 268 | 2jsa-6 | THR_A_46_N | THR_A_46_H | 5.887 | -5.379 | -9.709 | 6.198 | -5.444 | -10.668 | 1.01 | 161.67 | -11.81 |
| 269 | 2jsa-6 | THR_A_46_N | THR_A_46_CA | 5.887 | -5.379 | -9.709 | 4.835 | -6.272 | -9.252 | 1.45 | 71.68 | -139.67 |
| 270 | 2jsa-6 | THR_A_46_CA | THR_A_46_HA | 4.835 | -6.272 | -9.252 | 4.006 | -5.673 | -8.876 | 1.09 | 69.81 | 144.15 |
| 271 | 2jsa-6 | THR_A_46_CA | THR_A_46_C | 4.835 | -6.272 | -9.252 | 5.341 | -7.149 | -8.104 | 1.53 | 41.41 | -60.02 |
| 272 | 2jsa-6 | THR_A_46_C | THR_A_46_O | 5.341 | -7.149 | -8.104 | 4.574 | -7.521 | -7.218 | 1.23 | 43.89 | -154.13 |
| 273 | 2jsa-6 | THR_A_46_C | ARG_A_47_N | 5.341 | -7.149 | -8.104 | 6.63 | -7.454 | -8.159 | 1.33 | 92.38 | -13.31 |
| 274 | 2jsa-6 | ARG_A_47_N | ARG_A_47_H | 6.63 | -7.454 | -8.159 | 7.202 | -7.114 | -8.919 | 1.01 | 138.80 | 30.73 |
| 275 | 2jsa-6 | ARG_A_47_N | ARG_A_47_CA | 6.63 | -7.454 | -8.159 | 7.247 | -8.279 | -7.135 | 1.45 | 45.17 | -53.21 |
| 276 | 2jsa-6 | ARG_A_47_CA | ARG_A_47_HA | 7.247 | -8.279 | -7.135 | 6.723 | -9.234 | -7.101 | 1.09 | 88.21 | -118.75 |
| 277 | 2jsa-6 | ARG_A_47_CA | ARG_A_47_C | 7.247 | -8.279 | -7.135 | 7.136 | -7.602 | -5.768 | 1.53 | 26.65 | 99.31 |
| 278 | 2jsa-6 | ARG_A_47_C | ARG_A_47_O | 7.136 | -7.602 | -5.768 | 6.667 | -8.211 | -4.806 | 1.23 | 38.63 | -127.60 |
| 279 | 2jsa-6 | ARG_A_47_C | GLU_A_48_N | 7.136 | -7.602 | -5.768 | 7.573 | -6.352 | -5.724 | 1.32 | 88.10 | 70.73 |
| 280 | 2jsa-6 | GLU_A_48_N | GLU_A_48_H | 7.573 | -6.352 | -5.724 | 7.945 | -5.914 | -6.555 | 1.01 | 145.34 | 49.66 |
| 281 | 2jsa-6 | GLU_A_48_N | GLU_A_48_CA | 7.573 | -6.352 | -5.724 | 7.528 | -5.586 | -4.49 | 1.45 | 31.87 | 93.36 |
| 282 | 2jsa-6 | GLU_A_48_CA | GLU_A_48_HA | 7.528 | -5.586 | -4.49 | 7.959 | -6.197 | -3.697 | 1.09 | 43.32 | -54.80 |
| 283 | 2jsa-6 | GLU_A_48_CA | GLU_A_48_C | 7.528 | -5.586 | -4.49 | 6.08 | -5.265 | -4.118 | 1.53 | 75.92 | 167.50 |
| 284 | 2jsa-6 | GLU_A_48_C | GLU_A_48_O | 6.08 | -5.265 | -4.118 | 5.724 | -5.26 | -2.94 | 1.23 | 16.82 | 179.20 |
| 285 | 2jsa-6 | GLU_A_48_C | CYS_A_49_N | 6.08 | -5.265 | -4.118 | 5.282 | -5.004 | -5.143 | 1.32 | 140.68 | 161.89 |
| 286 | 2jsa-6 | CYS_A_49_N | CYS_A_49_H | 5.282 | -5.004 | -5.143 | 5.643 | -5.026 | -6.086 | 1.01 | 159.02 | -3.49 |
| 287 | 2jsa-6 | CYS_A_49_N | CYS_A_49_CA | 5.282 | -5.004 | -5.143 | 3.88 | -4.683 | -4.938 | 1.45 | 81.89 | 167.10 |
| 288 | 2jsa-6 | CYS_A_49_CA | CYS_A_49_HA | 3.88 | -4.683 | -4.938 | 3.82 | -3.796 | -4.306 | 1.09 | 54.59 | 93.87 |
| 289 | 2jsa-6 | CYS_A_49_CA | CYS_A_49_C | 3.88 | -4.683 | -4.938 | 3.223 | -5.852 | -4.2 | 1.53 | 61.17 | -119.34 |
| 290 | 2jsa-6 | CYS_A_49_C | CYS_A_49_O | 3.223 | -5.852 | -4.2 | 2.491 | -5.647 | -3.232 | 1.23 | 38.14 | 164.35 |
| 291 | 2jsa-6 | CYS_A_49_C | ASP_A_50_N | 3.223 | -5.852 | -4.2 | 3.506 | -7.051 | -4.685 | 1.32 | 111.49 | -76.72 |
| 292 | 2jsa-6 | ASP_A_50_N | ASP_A_50_H | 3.506 | -7.051 | -4.685 | 4.116 | -7.145 | -5.485 | 1.01 | 142.35 | -8.76 |
| 293 | 2jsa-6 | ASP_A_50_N | ASP_A_50_CA | 3.506 | -7.051 | -4.685 | 2.952 | -8.252 | -4.085 | 1.45 | 65.60 | -114.76 |
| 294 | 2jsa-6 | ASP_A_50_CA | ASP_A_50_HA | 2.952 | -8.252 | -4.085 | 1.878 | -8.12 | -3.953 | 1.09 | 83.05 | 172.99 |
| 295 | 2jsa-6 | ASP_A_50_CA | ASP_A_50_C | 2.952 | -8.252 | -4.085 | 3.617 | -8.492 | -2.729 | 1.53 | 27.54 | -19.84 |
| 296 | 2jsa-6 | ASP_A_50_C | ASP_A_50_O | 3.617 | -8.492 | -2.729 | 3.101 | -9.246 | -1.905 | 1.23 | 47.95 | -124.39 |
| 297 | 2jsa-6 | ASP_A_50_C | HIS_A_51_N | 3.617 | -8.492 | -2.729 | 4.753 | -7.838 | -2.538 | 1.32 | 81.71 | 29.93 |
| 298 | 2jsa-6 | HIS_A_51_N | HIS_A_51_H | 4.753 | -7.838 | -2.538 | 5.12 | -7.235 | -3.261 | 1.01 | 135.69 | 58.67 |
| 299 | 2jsa-6 | HIS_A_51_N | HIS_A_51_CA | 4.753 | -7.838 | -2.538 | 5.495 | -7.97 | -1.296 | 1.45 | 31.25 | -10.09 |
| 300 | 2jsa-6 | HIS_A_51_CA | HIS_A_51_HA | 5.495 | -7.97 | -1.296 | 5.133 | -8.86 | -0.781 | 1.09 | 61.81 | -112.13 |
| 301 | 2jsa-6 | HIS_A_51_CA | HIS_A_51_C | 5.495 | -7.97 | -1.296 | 5.215 | -6.763 | -0.399 | 1.53 | 54.10 | 103.06 |
| 302 | 2jsa-6 | HIS_A_51_C | HIS_A_51_O | 5.215 | -6.763 | -0.399 | 5.855 | -6.593 | 0.638 | 1.23 | 32.56 | 14.88 |

| | | | | | | | | | | | | |
|---|---|---|---|---|---|---|---|---|---|---|---|---|
| 303 | 2jsa-6 | HIS_A_51_C | TYR_A_52_N | 5.215 | -6.763 | -0.399 | 4.259 | -5.954 | -0.832 | 1.33 | 109.07 | 139.76 |
| 304 | 2jsa-6 | TYR_A_52_N | TYR_A_52_H | 4.259 | -5.954 | -0.832 | 3.776 | -6.154 | -1.696 | 1.01 | 148.82 | -157.51 |
| 305 | 2jsa-6 | TYR_A_52_N | TYR_A_52_CA | 4.259 | -5.954 | -0.832 | 3.887 | -4.767 | -0.081 | 1.45 | 58.88 | 107.40 |
| 306 | 2jsa-6 | TYR_A_52_CA | TYR_A_52_HA | 3.887 | -4.767 | -0.081 | 4.64 | -4.549 | 0.677 | 1.09 | 45.96 | 16.15 |
| 307 | 2jsa-6 | TYR_A_52_CA | TYR_A_52_C | 3.887 | -4.767 | -0.081 | 2.519 | -4.942 | 0.581 | 1.53 | 64.36 | -172.71 |
| 308 | 2jsa-6 | TYR_A_52_C | TYR_A_52_O | 2.519 | -4.942 | 0.581 | 2.289 | -4.444 | 1.681 | 1.23 | 26.50 | 114.79 |
| 309 | 2jsa-6 | TYR_A_52_C | VAL_A_53_N | 2.519 | -4.942 | 0.581 | 1.646 | -5.654 | -0.118 | 1.33 | 121.82 | -140.80 |
| 310 | 2jsa-6 | VAL_A_53_N | VAL_A_53_H | 1.646 | -5.654 | -0.118 | 1.904 | -6.035 | -1.017 | 1.01 | 152.90 | -55.90 |
| 311 | 2jsa-6 | VAL_A_53_N | VAL_A_53_CA | 1.646 | -5.654 | -0.118 | 0.307 | -5.901 | 0.389 | 1.45 | 69.58 | -169.55 |
| 312 | 2jsa-6 | VAL_A_53_CA | VAL_A_53_HA | 0.307 | -5.901 | 0.389 | -0.136 | -4.946 | 0.672 | 1.09 | 74.95 | 114.89 |
| 313 | 2jsa-6 | VAL_A_53_CA | VAL_A_53_C | 0.307 | -5.901 | 0.389 | 0.394 | -6.775 | 1.642 | 1.53 | 35.03 | -84.32 |
| 314 | 2jsa-6 | VAL_A_53_C | VAL_A_53_O | 0.394 | -6.775 | 1.642 | -0.595 | -6.945 | 2.354 | 1.23 | 54.64 | -170.25 |
| 315 | 2jsa-6 | VAL_A_53_C | ASN_A_54_N | 0.394 | -6.775 | 1.642 | 1.585 | -7.305 | 1.873 | 1.32 | 79.95 | -23.99 |
| 316 | 2jsa-6 | ASN_A_54_N | ASN_A_54_H | 1.585 | -7.305 | 1.873 | 2.354 | -7.121 | 1.245 | 1.01 | 128.46 | 13.46 |
| 317 | 2jsa-6 | ASN_A_54_N | ASN_A_54_CA | 1.585 | -7.305 | 1.873 | 1.814 | -8.157 | 3.028 | 1.45 | 37.37 | -74.96 |
| 318 | 2jsa-6 | ASN_A_54_CA | ASN_A_54_HA | 1.814 | -8.157 | 3.028 | 1.046 | -7.959 | 3.775 | 1.09 | 46.71 | 165.54 |
| 319 | 2jsa-6 | ASN_A_54_CA | ASN_A_54_C | 1.814 | -8.157 | 3.028 | 3.196 | -7.857 | 3.612 | 1.53 | 67.56 | 12.25 |
| 320 | 2jsa-6 | ASN_A_54_C | ASN_A_54_O | 3.196 | -7.857 | 3.612 | 4.069 | -8.723 | 3.628 | 1.23 | 89.25 | -44.77 |
| 321 | 2jsa-6 | ASN_A_54_C | SER_A_55_N | 3.196 | -7.857 | 3.612 | 3.352 | -6.627 | 4.077 | 1.32 | 69.44 | 82.77 |
| 322 | 2jsa-6 | SER_A_55_N | SER_A_55_H | 3.352 | -6.627 | 4.077 | 2.59 | -5.966 | 4.032 | 1.01 | 92.55 | 139.06 |
| 323 | 2jsa-6 | SER_A_55_N | SER_A_55_CA | 3.352 | -6.627 | 4.077 | 4.613 | -6.201 | 4.661 | 1.45 | 66.31 | 18.67 |
| 324 | 2jsa-6 | SER_A_55_CA | SER_A_55_HA | 4.613 | -6.201 | 4.661 | 4.849 | -6.869 | 5.489 | 1.09 | 40.55 | -70.54 |
| 325 | 2jsa-6 | SER_A_55_CA | SER_A_55_C | 4.613 | -6.201 | 4.661 | 4.485 | -4.776 | 5.203 | 1.53 | 69.25 | 95.13 |
| 326 | 2jsa-6 | SER_A_55_C | SER_A_55_O | 4.485 | -4.776 | 5.203 | 5.036 | -4.456 | 6.254 | 1.23 | 31.23 | 30.15 |
| 327 | 2jsa-6 | SER_A_55_C | LYS_A_56_N | 4.485 | -4.776 | 5.203 | 3.754 | -3.958 | 4.459 | 1.33 | 124.14 | 131.79 |
| 328 | 2jsa-6 | LYS_A_56_N | LYS_A_56_H | 3.754 | -3.958 | 4.459 | 3.33 | -4.289 | 3.605 | 1.01 | 147.79 | -142.02 |
| 329 | 2jsa-6 | LYS_A_56_N | LYS_A_56_CA | 3.754 | -3.958 | 4.459 | 3.547 | -2.575 | 4.852 | 1.45 | 74.30 | 98.51 |
| 330 | 2jsa-6 | LYS_A_56_CA | LYS_A_56_HA | 3.547 | -2.575 | 4.852 | 3.805 | -2.481 | 5.907 | 1.09 | 14.59 | 20.02 |
| 331 | 2jsa-6 | LYS_A_56_CA | LYS_A_56_C | 3.547 | -2.575 | 4.852 | 2.065 | -2.223 | 4.701 | 1.53 | 95.66 | 166.64 |
| 332 | 2jsa-6 | LYS_A_56_C | LYS_A_56_O | 2.065 | -2.223 | 4.701 | 1.703 | -1.048 | 4.666 | 1.23 | 91.63 | 107.12 |
| 333 | 2jsa-6 | LYS_A_56_C | VAL_A_57_N | 2.065 | -2.223 | 4.701 | 1.249 | -3.263 | 4.618 | 1.32 | 93.59 | -128.12 |
| 334 | 2jsa-6 | VAL_A_57_N | VAL_A_57_H | 1.249 | -3.263 | 4.618 | 1.618 | -4.203 | 4.655 | 1.01 | 87.90 | -68.57 |
| 335 | 2jsa-6 | VAL_A_57_N | VAL_A_57_CA | 1.249 | -3.263 | 4.618 | -0.186 | -3.079 | 4.473 | 1.45 | 95.72 | 172.69 |
| 336 | 2jsa-6 | VAL_A_57_CA | VAL_A_57_HA | -0.186 | -3.079 | 4.473 | -0.37 | -2.059 | 4.135 | 1.09 | 108.06 | 100.23 |
| 337 | 2jsa-6 | VAL_A_57_CA | VAL_A_57_C | -0.186 | -3.079 | 4.473 | -0.858 | -3.256 | 5.836 | 1.53 | 27.01 | -165.24 |
| 338 | 2jsa-6 | VAL_A_57_C | VAL_A_57_O | -0.858 | -3.256 | 5.836 | -1.949 | -2.734 | 6.066 | 1.23 | 79.23 | 154.43 |
| 339 | 2jsa-6 | VAL_A_57_C | ASP_A_58_N | -0.858 | -3.256 | 5.836 | -0.181 | -3.994 | 6.704 | 1.33 | 49.08 | -47.47 |
| 340 | 2jsa-6 | ASP_A_58_N | ASP_A_58_H | -0.181 | -3.994 | 6.704 | 0.711 | -4.393 | 6.446 | 1.01 | 104.79 | -24.10 |
| 341 | 2jsa-6 | ASP_A_58_N | ASP_A_58_CA | -0.181 | -3.994 | 6.704 | -0.7 | -4.246 | 8.037 | 1.45 | 23.40 | -154.10 |
| 342 | 2jsa-6 | ASP_A_58_CA | ASP_A_58_HA | -0.7 | -4.246 | 8.037 | -1.768 | -4.452 | 7.971 | 1.09 | 93.47 | -169.08 |
| 343 | 2jsa-6 | ASP_A_58_CA | ASP_A_58_C | -0.7 | -4.246 | 8.037 | -0.46 | -3.014 | 8.914 | 1.53 | 55.06 | 78.98 |
| 344 | 2jsa-6 | ASP_A_58_C | ASP_A_58_O | -0.46 | -3.014 | 8.914 | -1.34 | -2.606 | 9.671 | 1.23 | 52.03 | 155.13 |
| 345 | 2jsa-6 | ASP_A_58_C | PRO_A_59_N | -0.46 | -3.014 | 8.914 | 0.765 | -2.442 | 8.778 | 1.36 | 95.74 | 25.03 |
| 346 | 2jsa-6 | PRO_A_59_N | PRO_A_59_CA | 0.765 | -2.442 | 8.778 | 1.132 | -1.265 | 9.548 | 1.45 | 58.01 | 72.68 |
| 347 | 2jsa-6 | PRO_A_59_CA | PRO_A_59_HA | 1.132 | -1.265 | 9.548 | 0.832 | -1.392 | 10.588 | 1.09 | 17.39 | -157.06 |
| 348 | 2jsa-6 | PRO_A_59_CA | PRO_A_59_C | 1.132 | -1.265 | 9.548 | 0.448 | -0.013 | 8.996 | 1.53 | 111.15 | 118.65 |
| 349 | 2jsa-6 | PRO_A_59_C | PRO_A_59_O | 0.448 | -0.013 | 8.996 | -0.189 | 0.731 | 9.741 | 1.23 | 52.74 | 130.57 |
| 350 | 2jsa-6 | PRO_A_59_C | ILE_A_60_N | 0.448 | -0.013 | 8.996 | 0.602 | 0.181 | 7.695 | 1.32 | 169.22 | 51.56 |
| 351 | 2jsa-6 | ILE_A_60_N | ILE_A_60_H | 0.602 | 0.181 | 7.695 | 1.138 | -0.473 | 7.143 | 1.01 | 123.14 | -50.66 |
| 352 | 2jsa-6 | ILE_A_60_N | ILE_A_60_CA | 0.602 | 0.181 | 7.695 | 0.007 | 1.331 | 7.034 | 1.45 | 117.04 | 117.36 |
| 353 | 2jsa-6 | ILE_A_60_CA | ILE_A_60_HA | 0.007 | 1.331 | 7.034 | 0.51 | 2.227 | 7.398 | 1.09 | 70.49 | 60.69 |
| 354 | 2jsa-6 | ILE_A_60_CA | ILE_A_60_C | 0.007 | 1.331 | 7.034 | -1.468 | 1.432 | 7.427 | 1.53 | 75.11 | 176.08 |
| 355 | 2jsa-6 | ILE_A_60_C | ILE_A_60_O | -1.468 | 1.432 | 7.427 | -1.962 | 2.52 | 7.719 | 1.23 | 76.27 | 114.42 |
| 356 | 2jsa-6 | ILE_A_60_C | ILE_A_61_N | -1.468 | 1.432 | 7.427 | -2.129 | 0.284 | 7.421 | 1.32 | 90.26 | -119.93 |
| 357 | 2jsa-6 | ILE_A_61_N | ILE_A_61_H | -2.129 | 0.284 | 7.421 | -1.657 | -0.574 | 7.171 | 1.01 | 104.32 | -61.18 |
| 358 | 2jsa-6 | ILE_A_61_N | ILE_A_61_CA | -2.129 | 0.284 | 7.421 | -3.538 | 0.231 | 7.773 | 1.45 | 75.98 | -177.85 |
| 359 | 2jsa-6 | ILE_A_61_CA | ILE_A_61_HA | -3.538 | 0.231 | 7.773 | -4.042 | 1.059 | 7.274 | 1.09 | 117.24 | 121.33 |
| 360 | 2jsa-6 | ILE_A_61_CA | ILE_A_61_C | -3.538 | 0.231 | 7.773 | -3.69 | 0.436 | 9.282 | 1.53 | 9.60 | 126.56 |
| 361 | 2jsa-6 | ILE_A_61_C | ILE_A_61_O | -3.69 | 0.436 | 9.282 | -4.764 | 0.803 | 9.757 | 1.23 | 67.29 | 161.13 |
| 362 | 2jsa-6 | ILE_A_61_C | HIS_A_62_N | -3.69 | 0.436 | 9.282 | -2.599 | 0.191 | 9.994 | 1.33 | 57.51 | -12.66 |
| 363 | 2jsa-6 | HIS_A_62_N | HIS_A_62_H | -2.599 | 0.191 | 9.994 | -1.749 | -0.108 | 9.536 | 1.01 | 116.94 | -19.38 |
| 364 | 2jsa-6 | HIS_A_62_N | HIS_A_62_CA | -2.599 | 0.191 | 9.994 | -2.598 | 0.344 | 11.438 | 1.45 | 6.05 | 89.63 |
| 365 | 2jsa-6 | HIS_A_62_CA | HIS_A_62_HA | -2.598 | 0.344 | 11.438 | -3.5 | -0.127 | 11.827 | 1.09 | 69.08 | -152.43 |
| 366 | 2jsa-6 | HIS_A_62_CA | HIS_A_62_C | -2.598 | 0.344 | 11.438 | -2.652 | 1.83 | 11.796 | 1.53 | 76.46 | 92.08 |
| 367 | 2jsa-6 | HIS_A_62_C | HIS_A_62_O | -2.652 | 1.83 | 11.796 | -3.049 | 2.192 | 12.904 | 1.23 | 25.87 | 137.64 |
| 368 | 2jsa-6 | HIS_A_62_C | GLU_A_63_N | -2.652 | 1.83 | 11.796 | -2.249 | 2.652 | 10.839 | 1.32 | 136.27 | 63.88 |
| 369 | 2jsa-6 | GLU_A_63_N | GLU_A_63_H | -2.249 | 2.652 | 10.839 | -1.936 | 2.285 | 9.952 | 1.01 | 151.46 | -49.54 |
| 370 | 2jsa-6 | GLU_A_63_N | GLU_A_63_CA | -2.249 | 2.652 | 10.839 | -2.247 | 4.092 | 11.04 | 1.45 | 82.05 | 89.92 |
| 371 | 2jsa-6 | GLU_A_63_CA | GLU_A_63_HA | -2.247 | 4.092 | 11.04 | -2.01 | 4.286 | 12.086 | 1.09 | 16.32 | 39.30 |
| 372 | 2jsa-6 | GLU_A_63_CA | GLU_A_63_C | -2.247 | 4.092 | 11.04 | -3.632 | 4.671 | 10.745 | 1.53 | 101.12 | 157.31 |
| 373 | 2jsa-6 | GLU_A_63_C | GLU_A_63_O | -3.632 | 4.671 | 10.745 | -4.164 | 5.451 | 11.533 | 1.23 | 50.15 | 124.30 |
| 374 | 2jsa-6 | GLU_A_63_C | LEU_A_64_N | -3.632 | 4.671 | 10.745 | -4.177 | 4.266 | 9.606 | 1.33 | 149.20 | -143.38 |
| 375 | 2jsa-6 | LEU_A_64_N | LEU_A_64_H | -4.177 | 4.266 | 9.606 | -3.682 | 3.622 | 9.006 | 1.01 | 126.45 | -52.45 |
| 376 | 2jsa-6 | LEU_A_64_N | LEU_A_64_CA | -4.177 | 4.266 | 9.606 | -5.49 | 4.735 | 9.197 | 1.45 | 106.35 | 160.34 |
| 377 | 2jsa-6 | LEU_A_64_CA | LEU_A_64_HA | -5.49 | 4.735 | 9.197 | -5.495 | 5.823 | 9.264 | 1.09 | 86.48 | 90.26 |
| 378 | 2jsa-6 | LEU_A_64_CA | LEU_A_64_C | -5.49 | 4.735 | 9.197 | -6.543 | 4.204 | 10.172 | 1.53 | 50.42 | -153.24 |
| 379 | 2jsa-6 | LEU_A_64_C | LEU_A_64_O | -6.543 | 4.204 | 10.172 | -7.52 | 4.89 | 10.469 | 1.23 | 76.03 | 144.93 |
| 380 | 2jsa-6 | LEU_A_64_C | GLU_A_65_N | -6.543 | 4.204 | 10.172 | -6.308 | 2.988 | 10.642 | 1.32 | 69.22 | -79.06 |
| 381 | 2jsa-6 | GLU_A_65_N | GLU_A_65_H | -6.308 | 2.988 | 10.642 | -5.482 | 2.482 | 10.354 | 1.01 | 106.56 | -31.49 |
| 382 | 2jsa-6 | GLU_A_65_N | GLU_A_65_CA | -6.308 | 2.988 | 10.642 | -7.224 | 2.358 | 11.576 | 1.45 | 49.97 | -145.48 |
| 383 | 2jsa-6 | GLU_A_65_CA | GLU_A_65_HA | -7.224 | 2.358 | 11.576 | -8.24 | 2.549 | 11.232 | 1.09 | 108.41 | 169.35 |
| 384 | 2jsa-6 | GLU_A_65_CA | GLU_A_65_C | -7.224 | 2.358 | 11.576 | -7.065 | 2.969 | 12.97 | 1.53 | 24.37 | 75.41 |
| 385 | 2jsa-6 | GLU_A_65_C | GLU_A_65_O | -7.065 | 2.969 | 12.97 | -8.016 | 2.998 | 13.751 | 1.23 | 50.62 | 178.25 |
| 386 | 2jsa-6 | GLU_A_65_C | GLY_A_66_N | -7.065 | 2.969 | 12.97 | -5.858 | 3.443 | 13.24 | 1.32 | 78.24 | 21.44 |
| 387 | 2jsa-6 | GLY_A_66_N | GLY_A_66_H | -5.858 | 3.443 | 13.24 | -5.123 | 3.386 | 12.549 | 1.01 | 133.15 | -4.43 |
| 388 | 2jsa-6 | GLY_A_66_N | GLY_A_66_CA | -5.858 | 3.443 | 13.24 | -5.562 | 4.052 | 14.525 | 1.45 | 27.79 | 64.08 |
| 389 | 2jsa-6 | GLY_A_66_CA | GLY_A_66_C | -5.562 | 4.052 | 14.525 | -6.125 | 5.472 | 14.601 | 1.53 | 87.15 | 111.63 |
| 390 | 2jsa-6 | GLY_A_66_C | GLY_A_66_O | -6.125 | 5.472 | 14.601 | -6.73 | 5.852 | 15.603 | 1.23 | 35.49 | 147.87 |
| 391 | 2jsa-6 | GLY_A_66_C | GLY_A_67_N | -6.125 | 5.472 | 14.601 | -5.907 | 6.219 | 13.528 | 1.33 | 144.05 | 73.73 |
| 392 | 2jsa-6 | GLY_A_67_N | GLY_A_67_H | -5.907 | 6.219 | 13.528 | -5.401 | 5.842 | 12.739 | 1.01 | 141.35 | -36.69 |
| 393 | 2jsa-6 | GLY_A_67_N | GLY_A_67_CA | -5.907 | 6.219 | 13.528 | -6.386 | 7.589 | 13.461 | 1.45 | 92.64 | 109.27 |
| 394 | 2jsa-6 | GLY_A_67_CA | GLY_A_67_C | -6.386 | 7.589 | 13.461 | -5.53 | 8.422 | 12.504 | 1.53 | 128.70 | 44.22 |
| 395 | 2jsa-6 | GLY_A_67_C | GLY_A_67_O | -5.53 | 8.422 | 12.504 | -5.97 | 9.462 | 12.016 | 1.23 | 113.37 | 112.93 |
| 396 | 2jsa-6 | GLY_A_67_C | THR_A_68_N | -5.53 | 8.422 | 12.504 | -4.322 | 7.934 | 12.265 | 1.32 | 100.40 | -22.00 |
| 397 | 2jsa-6 | THR_A_68_N | THR_A_68_H | -4.322 | 7.934 | 12.265 | -4.026 | 7.072 | 12.701 | 1.01 | 64.43 | -71.05 |
| 398 | 2jsa-6 | THR_A_68_N | THR_A_68_CA | -4.322 | 7.934 | 12.265 | -3.399 | 8.62 | 11.376 | 1.45 | 127.71 | 36.62 |

| | | | | | | | | | | | |
|---|---|---|---|---|---|---|---|---|---|---|---|
| 399 | 2jsa-6 | THR_A_68_CA | THR_A_68_HA | -3.399 | 8.62 | 11.376 | -3.225 | 9.626 | 11.76 | 1.09 | 69.39 | 80.19 |
| 400 | 2jsa-6 | THR_A_68_CA | THR_A_68_C | -3.399 | 8.62 | 11.376 | -4.004 | 8.747 | 9.976 | 1.53 | 156.18 | 168.14 |
| 401 | 2jsa-6 | THR_A_68_C | THR_A_68_O | -4.004 | 8.747 | 9.976 | -4.745 | 7.87 | 9.534 | 1.23 | 111.06 | -130.20 |
| 402 | 2jsa-6 | THR_A_68_C | ALA_A_69_N | -4.004 | 8.747 | 9.976 | -3.665 | 9.846 | 9.317 | 1.33 | 119.81 | 72.86 |
| 403 | 2jsa-6 | ALA_A_69_N | ALA_A_69_H | -3.665 | 9.846 | 9.317 | -3.051 | 10.526 | 9.743 | 1.01 | 65.06 | 47.92 |
| 404 | 2jsa-6 | ALA_A_69_N | ALA_A_69_CA | -3.665 | 9.846 | 9.317 | -4.165 | 10.099 | 7.977 | 1.45 | 157.31 | 153.16 |
| 405 | 2jsa-6 | ALA_A_69_CA | ALA_A_69_HA | -4.165 | 10.099 | 7.977 | -5.208 | 9.787 | 7.927 | 1.09 | 92.63 | -163.35 |
| 406 | 2jsa-6 | ALA_A_69_CA | ALA_A_69_C | -4.165 | 10.099 | 7.977 | -3.362 | 9.269 | 6.973 | 1.53 | 131.00 | -45.95 |
| 407 | 2jsa-6 | ALA_A_69_C | ALA_A_69_O | -3.362 | 9.269 | 6.973 | -2.323 | 8.707 | 7.316 | 1.23 | 73.81 | -28.41 |
| 408 | 2jsa-6 | ALA_A_69_C | PRO_A_70_N | -3.362 | 9.269 | 6.973 | -3.889 | 9.215 | 5.721 | 1.36 | 157.07 | -174.15 |
| 409 | 2jsa-6 | PRO_A_70_N | PRO_A_70_CA | -3.889 | 9.215 | 5.721 | -3.234 | 8.463 | 4.664 | 1.45 | 136.67 | -48.94 |
| 410 | 2jsa-6 | PRO_A_70_CA | PRO_A_70_HA | -3.234 | 8.463 | 4.664 | -2.921 | 7.488 | 5.038 | 1.09 | 69.94 | -72.20 |
| 411 | 2jsa-6 | PRO_A_70_CA | PRO_A_70_C | -3.234 | 8.463 | 4.664 | -1.997 | 9.204 | 4.15 | 1.53 | 109.62 | 30.92 |
| 412 | 2jsa-6 | PRO_A_70_C | PRO_A_70_O | -1.997 | 9.204 | 4.15 | -1.242 | 8.67 | 3.339 | 1.23 | 131.25 | -35.27 |
| 413 | 2jsa-6 | PRO_A_70_C | LYS_A_71_N | -1.997 | 9.204 | 4.15 | -1.83 | 10.422 | 4.643 | 1.32 | 68.15 | 82.19 |
| 414 | 2jsa-6 | LYS_A_71_N | LYS_A_71_H | -1.83 | 10.422 | 4.643 | -2.492 | 10.798 | 5.307 | 1.01 | 48.91 | 150.40 |
| 415 | 2jsa-6 | LYS_A_71_N | LYS_A_71_CA | -1.83 | 10.422 | 4.643 | -0.698 | 11.241 | 4.243 | 1.45 | 105.98 | 35.89 |
| 416 | 2jsa-6 | LYS_A_71_CA | LYS_A_71_HA | -0.698 | 11.241 | 4.243 | -0.186 | 10.734 | 3.425 | 1.09 | 138.62 | -44.72 |
| 417 | 2jsa-6 | LYS_A_71_CA | LYS_A_71_C | -0.698 | 11.241 | 4.243 | 0.285 | 11.349 | 5.41 | 1.53 | 40.28 | 6.27 |
| 418 | 2jsa-6 | LYS_A_71_C | LYS_A_71_O | 0.285 | 11.349 | 5.41 | 1.341 | 11.968 | 5.283 | 1.23 | 95.92 | 30.38 |
| 419 | 2jsa-6 | LYS_A_71_C | ASP_A_72_N | 0.285 | 11.349 | 5.41 | -0.097 | 10.738 | 6.523 | 1.33 | 32.92 | -122.01 |
| 420 | 2jsa-6 | ASP_A_72_N | ASP_A_72_H | -0.097 | 10.738 | 6.523 | -0.979 | 10.248 | 6.558 | 1.01 | 88.01 | -150.95 |
| 421 | 2jsa-6 | ASP_A_72_N | ASP_A_72_CA | -0.097 | 10.738 | 6.523 | 0.738 | 10.759 | 7.712 | 1.45 | 35.09 | 1.44 |
| 422 | 2jsa-6 | ASP_A_72_CA | ASP_A_72_HA | 0.738 | 10.759 | 7.712 | 1.48 | 11.552 | 7.613 | 1.09 | 95.21 | 46.90 |
| 423 | 2jsa-6 | ASP_A_72_CA | ASP_A_72_C | 0.738 | 10.759 | 7.712 | 1.438 | 9.406 | 7.861 | 1.53 | 84.41 | -62.64 |
| 424 | 2jsa-6 | ASP_A_72_C | ASP_A_72_O | 1.438 | 9.406 | 7.861 | 2.561 | 9.337 | 8.358 | 1.23 | 66.17 | -3.52 |
| 425 | 2jsa-6 | ASP_A_72_C | VAL_A_73_N | 1.438 | 9.406 | 7.861 | 0.745 | 8.365 | 7.421 | 1.33 | 109.38 | -123.65 |
| 426 | 2jsa-6 | VAL_A_73_N | VAL_A_73_H | 0.745 | 8.365 | 7.421 | -0.173 | 8.497 | 7.023 | 1.01 | 113.23 | 171.82 |
| 427 | 2jsa-6 | VAL_A_73_N | VAL_A_73_CA | 0.745 | 8.365 | 7.421 | 1.287 | 7.019 | 7.5 | 1.45 | 86.88 | -68.07 |
| 428 | 2jsa-6 | VAL_A_73_CA | VAL_A_73_HA | 1.287 | 7.019 | 7.5 | 1.474 | 6.784 | 8.548 | 1.09 | 15.99 | -51.49 |
| 429 | 2jsa-6 | VAL_A_73_CA | VAL_A_73_C | 1.287 | 7.019 | 7.5 | 2.622 | 6.967 | 6.757 | 1.53 | 119.08 | -2.23 |
| 430 | 2jsa-6 | VAL_A_73_C | VAL_A_73_O | 2.622 | 6.967 | 6.757 | 3.605 | 6.439 | 7.274 | 1.23 | 65.14 | -28.24 |
| 431 | 2jsa-6 | VAL_A_73_C | CYS_A_74_N | 2.622 | 6.967 | 6.757 | 2.616 | 7.524 | 5.555 | 1.32 | 155.14 | 90.62 |
| 432 | 2jsa-6 | CYS_A_74_N | CYS_A_74_H | 2.616 | 7.524 | 5.555 | 1.772 | 7.943 | 5.193 | 1.01 | 111.02 | 153.60 |
| 433 | 2jsa-6 | CYS_A_74_N | CYS_A_74_CA | 2.616 | 7.524 | 5.555 | 3.816 | 7.547 | 4.734 | 1.45 | 124.37 | 1.10 |
| 434 | 2jsa-6 | CYS_A_74_CA | CYS_A_74_HA | 3.816 | 7.547 | 4.734 | 4.298 | 6.572 | 4.803 | 1.09 | 86.37 | -63.69 |
| 435 | 2jsa-6 | CYS_A_74_CA | CYS_A_74_C | 3.816 | 7.547 | 4.734 | 4.767 | 8.602 | 5.304 | 1.53 | 68.13 | 47.97 |
| 436 | 2jsa-6 | CYS_A_74_C | CYS_A_74_O | 4.767 | 8.602 | 5.304 | 5.13 | 9.553 | 4.615 | 1.23 | 124.09 | 69.11 |
| 437 | 2jsa-6 | CYS_A_74_C | THR_A_75_N | 4.767 | 8.602 | 5.304 | 5.143 | 8.398 | 6.558 | 1.32 | 18.84 | -28.48 |
| 438 | 2jsa-6 | THR_A_75_N | THR_A_75_H | 5.143 | 8.398 | 6.558 | 4.805 | 7.593 | 7.066 | 1.01 | 59.81 | -112.78 |
| 439 | 2jsa-6 | THR_A_75_N | THR_A_75_CA | 5.143 | 8.398 | 6.558 | 6.044 | 9.319 | 7.229 | 1.45 | 62.49 | 45.63 |
| 440 | 2jsa-6 | THR_A_75_CA | THR_A_75_HA | 6.044 | 9.319 | 7.229 | 7.03 | 9.241 | 6.77 | 1.09 | 114.89 | -4.52 |
| 441 | 2jsa-6 | THR_A_75_CA | THR_A_75_C | 6.044 | 9.319 | 7.229 | 6.183 | 8.949 | 8.706 | 1.53 | 14.98 | -69.41 |
| 442 | 2jsa-6 | THR_A_75_C | THR_A_75_O | 6.183 | 8.949 | 8.706 | 7.242 | 9.143 | 9.302 | 1.23 | 61.03 | 10.38 |
| 443 | 2jsa-6 | THR_A_75_C | LYS_A_76_N | 6.183 | 8.949 | 8.706 | 5.098 | 8.423 | 9.256 | 1.33 | 65.48 | -154.14 |
| 444 | 2jsa-6 | LYS_A_76_N | LYS_A_76_H | 5.098 | 8.423 | 9.256 | 4.261 | 8.293 | 8.706 | 1.01 | 123.00 | -171.17 |
| 445 | 2jsa-6 | LYS_A_76_N | LYS_A_76_CA | 5.098 | 8.423 | 9.256 | 5.086 | 8.024 | 10.654 | 1.45 | 15.94 | -91.72 |
| 446 | 2jsa-6 | LYS_A_76_CA | LYS_A_76_HA | 5.086 | 8.024 | 10.654 | 5.536 | 8.827 | 11.239 | 1.09 | 57.56 | 60.73 |
| 447 | 2jsa-6 | LYS_A_76_CA | LYS_A_76_C | 5.086 | 8.024 | 10.654 | 5.955 | 6.778 | 10.831 | 1.53 | 83.35 | -55.11 |
| 448 | 2jsa-6 | LYS_A_76_C | LYS_A_76_O | 5.955 | 6.778 | 10.831 | 6.76 | 6.704 | 11.758 | 1.23 | 41.09 | -5.25 |
| 449 | 2jsa-6 | LYS_A_76_C | LEU_A_77_N | 5.955 | 6.778 | 10.831 | 5.764 | 5.827 | 9.927 | 1.33 | 132.98 | -101.36 |
| 450 | 2jsa-6 | LEU_A_77_N | LEU_A_77_H | 5.764 | 5.827 | 9.927 | 5.086 | 5.956 | 9.19 | 1.01 | 136.88 | 169.23 |
| 451 | 2jsa-6 | LEU_A_77_N | LEU_A_77_CA | 5.764 | 5.827 | 9.927 | 6.519 | 4.587 | 9.972 | 1.45 | 88.22 | -58.66 |
| 452 | 2jsa-6 | LEU_A_77_CA | LEU_A_77_HA | 6.519 | 4.587 | 9.972 | 6.799 | 4.4 | 11.009 | 1.09 | 17.99 | -33.74 |
| 453 | 2jsa-6 | LEU_A_77_CA | LEU_A_77_C | 6.519 | 4.587 | 9.972 | 7.809 | 4.752 | 9.164 | 1.53 | 121.85 | 7.29 |
| 454 | 2jsa-6 | LEU_A_77_C | LEU_A_77_O | 7.809 | 4.752 | 9.164 | 8.394 | 3.767 | 8.716 | 1.23 | 111.36 | -59.29 |
| 455 | 2jsa-6 | LEU_A_77_C | ASN_A_78_N | 7.809 | 4.752 | 9.164 | 8.213 | 6.003 | 9.003 | 1.32 | 96.98 | 72.10 |
| 456 | 2jsa-6 | ASN_A_78_N | ASN_A_78_H | 8.213 | 6.003 | 9.003 | 7.682 | 6.764 | 9.401 | 1.01 | 66.79 | 124.91 |
| 457 | 2jsa-6 | ASN_A_78_N | ASN_A_78_CA | 8.213 | 6.003 | 9.003 | 9.422 | 6.309 | 8.256 | 1.45 | 120.92 | 14.20 |
| 458 | 2jsa-6 | ASN_A_78_CA | ASN_A_78_HA | 9.422 | 6.309 | 8.256 | 9.522 | 7.391 | 8.165 | 1.09 | 94.79 | 84.72 |
| 459 | 2jsa-6 | ASN_A_78_CA | ASN_A_78_C | 9.422 | 6.309 | 8.256 | 9.331 | 5.677 | 6.866 | 1.53 | 155.33 | -98.19 |
| 460 | 2jsa-6 | ASN_A_78_C | ASN_A_78_O | 9.331 | 5.677 | 6.866 | 10.346 | 5.496 | 6.196 | 1.23 | 123.02 | -10.11 |
| 461 | 2jsa-6 | ASN_A_78_C | GLU_A_79_N | 9.331 | 5.677 | 6.866 | 8.106 | 5.358 | 6.474 | 1.33 | 107.21 | -165.40 |
| 462 | 2jsa-6 | GLU_A_79_N | GLU_A_79_H | 8.106 | 5.358 | 6.474 | 7.317 | 5.536 | 7.078 | 1.01 | 53.25 | 167.29 |
| 463 | 2jsa-6 | GLU_A_79_N | GLU_A_79_CA | 8.106 | 5.358 | 6.474 | 7.87 | 4.75 | 5.176 | 1.45 | 153.32 | -111.21 |
| 464 | 2jsa-6 | GLU_A_79_CA | GLU_A_79_HA | 7.87 | 4.75 | 5.176 | 8.396 | 3.795 | 5.15 | 1.09 | 91.37 | -61.15 |
| 465 | 2jsa-6 | GLU_A_79_CA | GLU_A_79_C | 7.87 | 4.75 | 5.176 | 8.432 | 5.638 | 4.064 | 1.53 | 136.62 | 57.67 |
| 466 | 2jsa-6 | GLU_A_79_C | GLU_A_79_O | 8.432 | 5.638 | 4.064 | 9.452 | 5.311 | 3.459 | 1.23 | 119.46 | -17.78 |
| 467 | 2jsa-6 | GLU_A_79_C | CYS_A_80_N | 8.432 | 5.638 | 4.064 | 7.742 | 6.744 | 3.829 | 1.32 | 100.22 | 121.96 |
| 468 | 2jsa-6 | CYS_A_80_N | CYS_A_80_H | 7.742 | 6.744 | 3.829 | 6.912 | 6.95 | 4.367 | 1.01 | 57.83 | 166.06 |
| 469 | 2jsa-6 | CYS_A_80_N | CYS_A_80_CA | 7.742 | 6.744 | 3.829 | 8.16 | 7.682 | 2.8 | 1.45 | 135.06 | 65.98 |
| 470 | 2jsa-6 | CYS_A_80_CA | CYS_A_80_HA | 8.16 | 7.682 | 2.8 | 7.906 | 7.26 | 1.828 | 1.09 | 153.13 | -121.04 |
| 471 | 2jsa-6 | CYS_A_80_CA | CYS_A_80_C | 8.16 | 7.682 | 2.8 | 9.679 | 7.839 | 2.88 | 1.53 | 87.00 | 5.90 |
| 472 | 2jsa-6 | CYS_A_80_C | CYS_A_80_O | 9.679 | 7.839 | 2.88 | 10.292 | 7.476 | 3.884 | 1.23 | 35.36 | -30.63 |
| 473 | 2jsa-6 | CYS_A_80_C | PRO_A_81_N | 9.679 | 7.839 | 2.88 | 10.259 | 8.395 | 1.784 | 1.36 | 143.76 | 43.79 |
| 474 | 2jsa-6 | PRO_A_81_N | PRO_A_81_CA | 10.259 | 8.395 | 1.784 | 11.695 | 8.605 | 1.721 | 1.45 | 92.49 | 8.32 |
| 475 | 2jsa-6 | PRO_A_81_CA | PRO_A_81_HA | 11.695 | 8.605 | 1.721 | 12.218 | 7.717 | 2.077 | 1.09 | 70.94 | -59.50 |
| 476 | 2jsa-6 | PRO_A_81_CA | PRO_A_81_C | 11.695 | 8.605 | 1.721 | 12.114 | 9.794 | 2.588 | 1.53 | 55.48 | 70.59 |
| 477 | 2jsa-6 | PRO_A_81_C | PRO_A_81_O | 12.114 | 9.794 | 2.588 | 13.246 | 9.85 | 3.065 | 1.23 | 67.18 | 2.83 |
| 1 | 2jsa-7 | ARG_A_1_N | ARG_A_1_H | 15.033 | 4.568 | 0.099 | 15.002 | 5.432 | -0.423 | 1.01 | 121.12 | 92.05 |
| 2 | 2jsa-7 | ARG_A_1_N | ARG_A_1_CA | 15.033 | 4.568 | 0.099 | 13.793 | 3.844 | 0.318 | 1.45 | 81.33 | -149.72 |
| 3 | 2jsa-7 | ARG_A_1_CA | ARG_A_1_HA | 13.793 | 3.844 | 0.318 | 13.886 | 2.858 | -0.136 | 1.09 | 114.63 | -84.61 |
| 4 | 2jsa-7 | ARG_A_1_CA | ARG_A_1_C | 13.793 | 3.844 | 0.318 | 12.624 | 4.584 | -0.334 | 1.53 | 115.23 | 147.67 |
| 5 | 2jsa-7 | ARG_A_1_C | ARG_A_1_O | 12.624 | 4.584 | -0.334 | 11.659 | 3.962 | -0.775 | 1.23 | 111.01 | -147.20 |
| 6 | 2jsa-7 | ARG_A_1_C | SER_A_2_N | 12.624 | 4.584 | -0.334 | 12.748 | 5.903 | -0.375 | 1.33 | 91.77 | 84.63 |
| 7 | 2jsa-7 | SER_A_2_N | SER_A_2_H | 12.748 | 5.903 | -0.375 | 13.571 | 6.348 | 0.006 | 1.01 | 67.84 | 28.40 |
| 8 | 2jsa-7 | SER_A_2_N | SER_A_2_CA | 12.748 | 5.903 | -0.375 | 11.714 | 6.735 | -0.965 | 1.45 | 113.97 | 141.18 |
| 9 | 2jsa-7 | SER_A_2_CA | SER_A_2_HA | 11.714 | 6.735 | -0.965 | 10.755 | 6.441 | -0.54 | 1.09 | 67.04 | -162.96 |
| 10 | 2jsa-7 | SER_A_2_CA | SER_A_2_C | 11.714 | 6.735 | -0.965 | 11.668 | 6.516 | -2.48 | 1.53 | 171.60 | -101.86 |
| 11 | 2jsa-7 | SER_A_2_C | SER_A_2_O | 11.668 | 6.516 | -2.48 | 12.527 | 7.011 | -3.208 | 1.23 | 126.29 | 29.95 |
| 12 | 2jsa-7 | SER_A_2_C | ALA_A_3_N | 11.668 | 6.516 | -2.48 | 10.656 | 5.776 | -2.907 | 1.32 | 108.81 | -143.82 |
| 13 | 2jsa-7 | ALA_A_3_N | ALA_A_3_H | 10.656 | 5.776 | -2.907 | 9.987 | 5.402 | -2.25 | 1.01 | 49.40 | -150.79 |
| 14 | 2jsa-7 | ALA_A_3_N | ALA_A_3_CA | 10.656 | 5.776 | -2.907 | 10.487 | 5.486 | -4.322 | 1.45 | 166.66 | -120.23 |
| 15 | 2jsa-7 | ALA_A_3_CA | ALA_A_3_HA | 10.487 | 5.486 | -4.322 | 10.631 | 6.406 | -4.888 | 1.09 | 121.29 | 81.10 |
| 16 | 2jsa-7 | ALA_A_3_CA | ALA_A_3_C | 10.487 | 5.486 | -4.322 | 9.063 | 4.984 | -4.568 | 1.53 | 99.25 | -160.58 |
| 17 | 2jsa-7 | ALA_A_3_C | ALA_A_3_O | 9.063 | 4.984 | -4.568 | 8.11 | 5.495 | -3.981 | 1.23 | 61.51 | 151.80 |

| | | | | | | | | | | | |
|---|---|---|---|---|---|---|---|---|---|---|---|
| 18 | 2jsa-7 | ALA_A_3_C | LEU_A_4_N | 9.063 | 4.984 | -4.568 | 8.962 | 3.988 | -5.436 | 1.33 | 130.93 | -95.79 |
| 19 | 2jsa-7 | LEU_A_4_N | LEU_A_4_H | 8.962 | 3.988 | -5.436 | 9.789 | 3.616 | -5.881 | 1.01 | 116.14 | -24.22 |
| 20 | 2jsa-7 | LEU_A_4_N | LEU_A_4_CA | 8.962 | 3.988 | -5.436 | 7.671 | 3.411 | -5.767 | 1.45 | 103.17 | -155.92 |
| 21 | 2jsa-7 | LEU_A_4_CA | LEU_A_4_HA | 7.671 | 3.411 | -5.767 | 7.01 | 4.22 | -6.078 | 1.09 | 106.58 | 129.25 |
| 22 | 2jsa-7 | LEU_A_4_CA | LEU_A_4_C | 7.671 | 3.411 | -5.767 | 7.067 | 2.772 | -4.515 | 1.53 | 35.08 | -133.39 |
| 23 | 2jsa-7 | LEU_A_4_C | LEU_A_4_O | 7.067 | 2.772 | -4.515 | 5.883 | 2.438 | -4.494 | 1.23 | 89.02 | -164.25 |
| 24 | 2jsa-7 | LEU_A_4_C | SER_A_5_N | 7.067 | 2.772 | -4.515 | 7.907 | 2.623 | -3.501 | 1.33 | 40.08 | -10.06 |
| 25 | 2jsa-7 | SER_A_5_N | SER_A_5_H | 7.907 | 2.623 | -3.501 | 8.868 | 2.923 | -3.588 | 1.01 | 94.94 | 17.34 |
| 26 | 2jsa-7 | SER_A_5_N | SER_A_5_CA | 7.907 | 2.623 | -3.501 | 7.471 | 2.03 | -2.248 | 1.45 | 30.43 | -126.32 |
| 27 | 2jsa-7 | SER_A_5_CA | SER_A_5_HA | 7.471 | 2.03 | -2.248 | 6.985 | 1.08 | -2.47 | 1.09 | 101.75 | -117.09 |
| 28 | 2jsa-7 | SER_A_5_CA | SER_A_5_C | 7.471 | 2.03 | -2.248 | 6.459 | 2.949 | -1.559 | 1.53 | 63.25 | 137.76 |
| 29 | 2jsa-7 | SER_A_5_C | SER_A_5_O | 6.459 | 2.949 | -1.559 | 5.395 | 2.499 | -1.136 | 1.23 | 69.89 | -157.07 |
| 30 | 2jsa-7 | SER_A_5_C | CYS_A_6_N | 6.459 | 2.949 | -1.559 | 6.828 | 4.218 | -1.467 | 1.32 | 86.02 | 73.79 |
| 31 | 2jsa-7 | CYS_A_6_N | CYS_A_6_H | 6.828 | 4.218 | -1.467 | 7.719 | 4.517 | -1.837 | 1.01 | 111.49 | 18.55 |
| 32 | 2jsa-7 | CYS_A_6_N | CYS_A_6_CA | 6.828 | 4.218 | -1.467 | 5.966 | 5.204 | -0.837 | 1.45 | 64.31 | 131.16 |
| 33 | 2jsa-7 | CYS_A_6_CA | CYS_A_6_HA | 5.966 | 5.204 | -0.837 | 5.804 | 4.906 | 0.199 | 1.09 | 18.13 | -118.53 |
| 34 | 2jsa-7 | CYS_A_6_CA | CYS_A_6_C | 5.966 | 5.204 | -0.837 | 4.619 | 5.197 | -1.562 | 1.53 | 118.29 | -179.70 |
| 35 | 2jsa-7 | CYS_A_6_C | CYS_A_6_O | 4.619 | 5.197 | -1.562 | 3.581 | 4.953 | -0.948 | 1.23 | 60.07 | -166.77 |
| 36 | 2jsa-7 | CYS_A_6_C | GLN_A_7_N | 4.619 | 5.197 | -1.562 | 4.678 | 5.468 | -2.857 | 1.32 | 167.91 | 77.72 |
| 37 | 2jsa-7 | GLN_A_7_N | GLN_A_7_H | 4.678 | 5.468 | -2.857 | 5.567 | 5.66 | -3.296 | 1.01 | 115.77 | 12.19 |
| 38 | 2jsa-7 | GLN_A_7_N | GLN_A_7_CA | 4.678 | 5.468 | -2.857 | 3.474 | 5.497 | -3.671 | 1.45 | 124.05 | 178.62 |
| 39 | 2jsa-7 | GLN_A_7_CA | GLN_A_7_HA | 3.474 | 5.497 | -3.671 | 2.933 | 6.415 | -3.441 | 1.09 | 77.82 | 120.51 |
| 40 | 2jsa-7 | GLN_A_7_CA | GLN_A_7_C | 3.474 | 5.497 | -3.671 | 2.573 | 4.308 | -3.329 | 1.53 | 77.09 | -127.15 |
| 41 | 2jsa-7 | GLN_A_7_C | GLN_A_7_O | 2.573 | 4.308 | -3.329 | 1.389 | 4.483 | -3.049 | 1.23 | 76.83 | 171.59 |
| 42 | 2jsa-7 | GLN_A_7_C | MET_A_8_N | 2.573 | 4.308 | -3.329 | 3.17 | 3.126 | -3.362 | 1.32 | 91.43 | -63.20 |
| 43 | 2jsa-7 | MET_A_8_N | MET_A_8_H | 3.17 | 3.126 | -3.362 | 4.149 | 3.057 | -3.601 | 1.01 | 103.69 | -4.03 |
| 44 | 2jsa-7 | MET_A_8_N | MET_A_8_CA | 3.17 | 3.126 | -3.362 | 2.436 | 1.909 | -3.059 | 1.45 | 77.96 | -121.10 |
| 45 | 2jsa-7 | MET_A_8_CA | MET_A_8_HA | 2.436 | 1.909 | -3.059 | 1.761 | 1.693 | -3.887 | 1.09 | 139.44 | -162.26 |
| 46 | 2jsa-7 | MET_A_8_CA | MET_A_8_C | 2.436 | 1.909 | -3.059 | 1.616 | 2.068 | -1.778 | 1.53 | 33.11 | 169.03 |
| 47 | 2jsa-7 | MET_A_8_C | MET_A_8_O | 1.616 | 2.068 | -1.778 | 0.528 | 1.506 | -1.659 | 1.23 | 84.45 | -152.68 |
| 48 | 2jsa-7 | MET_A_8_C | CYS_A_9_N | 1.616 | 2.068 | -1.778 | 2.169 | 2.837 | -0.851 | 1.33 | 45.62 | 54.28 |
| 49 | 2jsa-7 | CYS_A_9_N | CYS_A_9_H | 2.169 | 2.837 | -0.851 | 3.067 | 3.269 | -1.017 | 1.01 | 99.46 | 25.69 |
| 50 | 2jsa-7 | CYS_A_9_N | CYS_A_9_CA | 2.169 | 2.837 | -0.851 | 1.502 | 3.077 | 0.417 | 1.45 | 29.21 | 160.21 |
| 51 | 2jsa-7 | CYS_A_9_CA | CYS_A_9_HA | 1.502 | 3.077 | 0.417 | 1.268 | 2.113 | 0.869 | 1.09 | 65.50 | -103.64 |
| 52 | 2jsa-7 | CYS_A_9_CA | CYS_A_9_C | 1.502 | 3.077 | 0.417 | 0.191 | 3.816 | 0.138 | 1.53 | 100.50 | 150.59 |
| 53 | 2jsa-7 | CYS_A_9_C | CYS_A_9_O | 0.191 | 3.816 | 0.138 | -0.821 | 3.559 | 0.788 | 1.23 | 58.10 | -165.75 |
| 54 | 2jsa-7 | CYS_A_9_C | GLU_A_10_N | 0.191 | 3.816 | 0.138 | 0.254 | 4.722 | -0.828 | 1.33 | 136.77 | 86.02 |
| 55 | 2jsa-7 | GLU_A_10_N | GLU_A_10_H | 0.254 | 4.722 | -0.828 | 1.122 | 4.883 | -1.319 | 1.01 | 119.08 | 10.51 |
| 56 | 2jsa-7 | GLU_A_10_N | GLU_A_10_CA | 0.254 | 4.722 | -0.828 | -0.915 | 5.499 | -1.201 | 1.45 | 104.88 | 146.39 |
| 57 | 2jsa-7 | GLU_A_10_CA | GLU_A_10_HA | -0.915 | 5.499 | -1.201 | -1.411 | 5.822 | -0.286 | 1.09 | 32.90 | 146.93 |
| 58 | 2jsa-7 | GLU_A_10_CA | GLU_A_10_C | -0.915 | 5.499 | -1.201 | -1.894 | 4.637 | -2.0 | 1.53 | 121.49 | -138.64 |
| 59 | 2jsa-7 | GLU_A_10_C | GLU_A_10_O | -1.894 | 4.637 | -2.0 | -2.998 | 5.078 | -2.317 | 1.23 | 104.93 | 158.23 |
| 60 | 2jsa-7 | GLU_A_10_C | LEU_A_11_N | -1.894 | 4.637 | -2.0 | -1.455 | 3.425 | -2.302 | 1.32 | 103.19 | -70.09 |
| 61 | 2jsa-7 | LEU_A_11_N | LEU_A_11_H | -1.455 | 3.425 | -2.302 | -0.535 | 3.129 | -2.008 | 1.01 | 73.08 | -17.84 |
| 62 | 2jsa-7 | LEU_A_11_N | LEU_A_11_CA | -1.455 | 3.425 | -2.302 | -2.28 | 2.497 | -3.058 | 1.45 | 121.33 | -131.64 |
| 63 | 2jsa-7 | LEU_A_11_CA | LEU_A_11_HA | -2.28 | 2.497 | -3.058 | -2.997 | 3.078 | -3.637 | 1.09 | 122.10 | 140.98 |
| 64 | 2jsa-7 | LEU_A_11_CA | LEU_A_11_C | -2.28 | 2.497 | -3.058 | -3.07 | 1.616 | -2.088 | 1.53 | 50.66 | -131.88 |
| 65 | 2jsa-7 | LEU_A_11_C | LEU_A_11_O | -3.07 | 1.616 | -2.088 | -4.238 | 1.315 | -2.327 | 1.23 | 101.21 | -165.55 |
| 66 | 2jsa-7 | LEU_A_11_C | VAL_A_12_N | -3.07 | 1.616 | -2.088 | -2.4 | 1.227 | -1.012 | 1.33 | 35.75 | -30.14 |
| 67 | 2jsa-7 | VAL_A_12_N | VAL_A_12_H | -2.4 | 1.227 | -1.012 | -1.44 | 1.513 | -0.879 | 1.01 | 82.44 | 16.59 |
| 68 | 2jsa-7 | VAL_A_12_N | VAL_A_12_CA | -2.4 | 1.227 | -1.012 | -3.025 | 0.387 | -0.005 | 1.45 | 46.12 | -126.65 |
| 69 | 2jsa-7 | VAL_A_12_CA | VAL_A_12_HA | -3.025 | 0.387 | -0.005 | -3.581 | -0.402 | -0.511 | 1.09 | 117.66 | -125.17 |
| 70 | 2jsa-7 | VAL_A_12_CA | VAL_A_12_C | -3.025 | 0.387 | -0.005 | -4.017 | 1.223 | 0.806 | 1.53 | 57.99 | 139.88 |
| 71 | 2jsa-7 | VAL_A_12_C | VAL_A_12_O | -4.017 | 1.223 | 0.806 | -5.153 | 0.804 | 1.026 | 1.23 | 79.70 | -159.75 |
| 72 | 2jsa-7 | VAL_A_12_C | VAL_A_13_N | -4.017 | 1.223 | 0.806 | -3.553 | 2.39 | 1.228 | 1.32 | 71.43 | 68.32 |
| 73 | 2jsa-7 | VAL_A_13_N | VAL_A_13_H | -3.553 | 2.39 | 1.228 | -2.607 | 2.67 | 1.01 | 1.01 | 102.46 | 16.49 |
| 74 | 2jsa-7 | VAL_A_13_N | VAL_A_13_CA | -3.553 | 2.39 | 1.228 | -4.386 | 3.289 | 2.01 | 1.45 | 57.46 | 132.82 |
| 75 | 2jsa-7 | VAL_A_13_CA | VAL_A_13_HA | -4.386 | 3.289 | 2.01 | -4.778 | 2.739 | 2.866 | 1.09 | 38.27 | -125.48 |
| 76 | 2jsa-7 | VAL_A_13_CA | VAL_A_13_C | -4.386 | 3.289 | 2.01 | -5.57 | 3.749 | 1.158 | 1.53 | 123.85 | 158.77 |
| 77 | 2jsa-7 | VAL_A_13_C | VAL_A_13_O | -5.57 | 3.749 | 1.158 | -6.724 | 3.519 | 1.516 | 1.23 | 73.08 | -168.73 |
| 78 | 2jsa-7 | VAL_A_13_C | LYS_A_14_N | -5.57 | 3.749 | 1.158 | -5.244 | 4.393 | 0.047 | 1.32 | 146.99 | 63.15 |
| 79 | 2jsa-7 | LYS_A_14_N | LYS_A_14_H | -5.244 | 4.393 | 0.047 | -4.273 | 4.548 | -0.184 | 1.01 | 103.22 | 9.07 |
| 80 | 2jsa-7 | LYS_A_14_N | LYS_A_14_CA | -5.244 | 4.393 | 0.047 | -6.266 | 4.888 | -0.858 | 1.45 | 128.55 | 154.16 |
| 81 | 2jsa-7 | LYS_A_14_CA | LYS_A_14_HA | -6.266 | 4.888 | -0.858 | -6.738 | 5.754 | -0.393 | 1.09 | 64.76 | 118.59 |
| 82 | 2jsa-7 | LYS_A_14_CA | LYS_A_14_C | -6.266 | 4.888 | -0.858 | -7.342 | 3.815 | -1.043 | 1.53 | 96.94 | -135.08 |
| 83 | 2jsa-7 | LYS_A_14_C | LYS_A_14_O | -7.342 | 3.815 | -1.043 | -8.525 | 4.132 | -1.16 | 1.23 | 95.46 | 165.00 |
| 84 | 2jsa-7 | LYS_A_14_C | LYS_A_15_N | -7.342 | 3.815 | -1.043 | -6.893 | 2.569 | -1.062 | 1.32 | 90.82 | -70.18 |
| 85 | 2jsa-7 | LYS_A_15_N | LYS_A_15_H | -6.893 | 2.569 | -1.062 | -5.905 | 2.383 | -0.96 | 1.01 | 84.21 | -10.66 |
| 86 | 2jsa-7 | LYS_A_15_N | LYS_A_15_CA | -6.893 | 2.569 | -1.062 | -7.802 | 1.448 | -1.229 | 1.45 | 96.60 | -129.04 |
| 87 | 2jsa-7 | LYS_A_15_CA | LYS_A_15_HA | -7.802 | 1.448 | -1.229 | -8.555 | 1.731 | -1.965 | 1.09 | 132.46 | 159.40 |
| 88 | 2jsa-7 | LYS_A_15_CA | LYS_A_15_C | -7.802 | 1.448 | -1.229 | -8.522 | 1.18 | 0.093 | 1.53 | 30.16 | -159.58 |
| 89 | 2jsa-7 | LYS_A_15_C | LYS_A_15_O | -8.522 | 1.18 | 0.093 | -9.71 | 0.856 | 0.103 | 1.23 | 89.53 | -164.74 |
| 90 | 2jsa-7 | LYS_A_15_C | TYR_A_16_N | -8.522 | 1.18 | 0.093 | -7.776 | 1.324 | 1.178 | 1.32 | 35.00 | 10.93 |
| 91 | 2jsa-7 | TYR_A_16_N | TYR_A_16_H | -7.776 | 1.324 | 1.178 | -6.805 | 1.592 | 1.096 | 1.01 | 94.65 | 15.43 |
| 92 | 2jsa-7 | TYR_A_16_N | TYR_A_16_CA | -7.776 | 1.324 | 1.178 | -8.329 | 1.101 | 2.503 | 1.45 | 24.23 | -158.04 |
| 93 | 2jsa-7 | TYR_A_16_CA | TYR_A_16_HA | -8.329 | 1.101 | 2.503 | -8.922 | 0.187 | 2.516 | 1.09 | 89.32 | -122.98 |
| 94 | 2jsa-7 | TYR_A_16_CA | TYR_A_16_C | -8.329 | 1.101 | 2.503 | -9.191 | 2.286 | 2.943 | 1.53 | 73.29 | 126.03 |
| 95 | 2jsa-7 | TYR_A_16_C | TYR_A_16_O | -9.191 | 2.286 | 2.943 | -10.417 | 2.187 | 2.979 | 1.23 | 88.32 | -175.38 |
| 96 | 2jsa-7 | TYR_A_16_C | GLU_A_17_N | -9.191 | 2.286 | 2.943 | -8.516 | 3.38 | 3.267 | 1.33 | 75.85 | 58.33 |
| 97 | 2jsa-7 | GLU_A_17_N | GLU_A_17_H | -8.516 | 3.38 | 3.267 | -7.507 | 3.391 | 3.213 | 1.01 | 93.06 | 0.62 |
| 98 | 2jsa-7 | GLU_A_17_N | GLU_A_17_CA | -8.516 | 3.38 | 3.267 | -9.206 | 4.581 | 3.703 | 1.45 | 72.53 | 119.88 |
| 99 | 2jsa-7 | GLU_A_17_CA | GLU_A_17_HA | -9.206 | 4.581 | 3.703 | -9.529 | 4.429 | 4.733 | 1.09 | 19.12 | -154.80 |
| 100 | 2jsa-7 | GLU_A_17_CA | GLU_A_17_C | -9.206 | 4.581 | 3.703 | -10.446 | 4.825 | 2.841 | 1.53 | 124.30 | 168.87 |
| 101 | 2jsa-7 | GLU_A_17_C | GLU_A_17_O | -10.446 | 4.825 | 2.841 | -11.436 | 5.383 | 3.313 | 1.23 | 67.45 | 150.59 |
| 102 | 2jsa-7 | GLU_A_17_C | GLY_A_18_N | -10.446 | 4.825 | 2.841 | -10.351 | 4.397 | 1.591 | 1.32 | 160.67 | -77.49 |
| 103 | 2jsa-7 | GLY_A_18_N | GLY_A_18_H | -10.351 | 4.397 | 1.591 | -9.505 | 3.948 | 1.27 | 1.01 | 108.53 | -27.96 |
| 104 | 2jsa-7 | GLY_A_18_N | GLY_A_18_CA | -10.351 | 4.397 | 1.591 | -11.453 | 4.562 | 0.658 | 1.45 | 129.94 | 171.48 |
| 105 | 2jsa-7 | GLY_A_18_CA | GLY_A_18_C | -11.453 | 4.562 | 0.658 | -11.927 | 3.208 | 0.125 | 1.53 | 110.38 | -109.29 |
| 106 | 2jsa-7 | GLY_A_18_C | GLY_A_18_O | -11.927 | 3.208 | 0.125 | -12.236 | 3.076 | -1.059 | 1.23 | 164.16 | -156.87 |
| 107 | 2jsa-7 | GLY_A_18_C | SER_A_19_N | -11.927 | 3.208 | 0.125 | -11.972 | 2.237 | 1.025 | 1.32 | 47.20 | -92.65 |
| 108 | 2jsa-7 | SER_A_19_N | SER_A_19_H | -11.972 | 2.237 | 1.025 | -11.706 | 2.418 | 1.983 | 1.01 | 18.56 | 34.23 |
| 109 | 2jsa-7 | SER_A_19_N | SER_A_19_CA | -11.972 | 2.237 | 1.025 | -12.403 | 0.898 | 0.66 | 1.45 | 104.55 | -107.84 |
| 110 | 2jsa-7 | SER_A_19_CA | SER_A_19_HA | -12.403 | 0.898 | 0.66 | -11.984 | 0.659 | -0.317 | 1.09 | 153.72 | -29.70 |
| 111 | 2jsa-7 | SER_A_19_CA | SER_A_19_C | -12.403 | 0.898 | 0.66 | -13.93 | 0.849 | 0.566 | 1.53 | 93.52 | -178.16 |
| 112 | 2jsa-7 | SER_A_19_C | SER_A_19_O | -13.93 | 0.849 | 0.566 | -14.479 | 0.45 | -0.46 | 1.23 | 146.52 | -143.99 |
| 113 | 2jsa-7 | SER_A_19_C | ALA_A_20_N | -13.93 | 0.849 | 0.566 | -14.572 | 1.258 | 1.649 | 1.32 | 35.10 | 147.50 |

| | | | | | | | | | | | | |
|---|---|---|---|---|---|---|---|---|---|---|---|---|
| 114 | 2jsa-7 | ALA_A_20_N | ALA_A_20_H | -14.572 | 1.258 | 1.649 | -14.054 | 1.571 | 2.458 | 1.01 | 36.80 | 31.14 |
| 115 | 2jsa-7 | ALA_A_20_N | ALA_A_20_CA | -14.572 | 1.258 | 1.649 | -16.024 | 1.266 | 1.702 | 1.45 | 87.91 | 179.68 |
| 116 | 2jsa-7 | ALA_A_20_CA | ALA_A_20_HA | -16.024 | 1.266 | 1.702 | -16.343 | 1.968 | 2.473 | 1.09 | 45.00 | 114.44 |
| 117 | 2jsa-7 | ALA_A_20_CA | ALA_A_20_C | -16.024 | 1.266 | 1.702 | -16.523 | -0.13 | 2.081 | 1.53 | 75.66 | -109.67 |
| 118 | 2jsa-7 | ALA_A_20_C | ALA_A_20_O | -16.523 | -0.13 | 2.081 | -17.716 | -0.326 | 2.307 | 1.23 | 79.41 | -170.67 |
| 119 | 2jsa-7 | ALA_A_20_C | ASP_A_21_N | -16.523 | -0.13 | 2.081 | -15.585 | -1.064 | 2.138 | 1.32 | 87.53 | -44.88 |
| 120 | 2jsa-7 | ASP_A_21_N | ASP_A_21_H | -15.585 | -1.064 | 2.138 | -14.622 | -0.831 | 1.939 | 1.01 | 101.36 | 13.60 |
| 121 | 2jsa-7 | ASP_A_21_N | ASP_A_21_CA | -15.585 | -1.064 | 2.138 | -15.914 | -2.436 | 2.485 | 1.45 | 76.18 | -103.48 |
| 122 | 2jsa-7 | ASP_A_21_CA | ASP_A_21_HA | -15.914 | -2.436 | 2.485 | -16.997 | -2.537 | 2.545 | 1.09 | 86.84 | -174.67 |
| 123 | 2jsa-7 | ASP_A_21_CA | ASP_A_21_C | -15.914 | -2.436 | 2.485 | -15.285 | -2.78 | 3.837 | 1.53 | 27.94 | -28.67 |
| 124 | 2jsa-7 | ASP_A_21_C | ASP_A_21_O | -15.285 | -2.78 | 3.837 | -15.691 | -3.742 | 4.488 | 1.23 | 58.06 | -112.88 |
| 125 | 2jsa-7 | ASP_A_21_C | LYS_A_22_N | -15.285 | -2.78 | 3.837 | -14.302 | -1.977 | 4.217 | 1.32 | 73.33 | 39.24 |
| 126 | 2jsa-7 | LYS_A_22_N | LYS_A_22_H | -14.302 | -1.977 | 4.217 | -14.019 | -1.205 | 3.63 | 1.01 | 125.52 | 69.87 |
| 127 | 2jsa-7 | LYS_A_22_N | LYS_A_22_CA | -14.302 | -1.977 | 4.217 | -13.612 | -2.186 | 5.479 | 1.45 | 29.74 | -16.85 |
| 128 | 2jsa-7 | LYS_A_22_CA | LYS_A_22_HA | -13.612 | -2.186 | 5.479 | -12.747 | -1.523 | 5.504 | 1.09 | 88.69 | 37.47 |
| 129 | 2jsa-7 | LYS_A_22_CA | LYS_A_22_C | -13.612 | -2.186 | 5.479 | -13.096 | -3.625 | 5.541 | 1.53 | 87.68 | -70.27 |
| 130 | 2jsa-7 | LYS_A_22_C | LYS_A_22_O | -13.096 | -3.625 | 5.541 | -13.52 | -4.403 | 6.395 | 1.23 | 46.05 | -118.59 |
| 131 | 2jsa-7 | LYS_A_22_C | ASP_A_23_N | -13.096 | -3.625 | 5.541 | -12.189 | -3.936 | 4.627 | 1.32 | 133.63 | -18.93 |
| 132 | 2jsa-7 | ASP_A_23_N | ASP_A_23_H | -12.189 | -3.936 | 4.627 | -11.887 | -3.246 | 3.954 | 1.01 | 131.78 | 66.36 |
| 133 | 2jsa-7 | ASP_A_23_N | ASP_A_23_CA | -12.189 | -3.936 | 4.627 | -11.611 | -5.267 | 4.569 | 1.45 | 92.29 | -66.53 |
| 134 | 2jsa-7 | ASP_A_23_CA | ASP_A_23_HA | -11.611 | -5.267 | 4.569 | -11.635 | -5.71 | 5.565 | 1.09 | 24.01 | -93.10 |
| 135 | 2jsa-7 | ASP_A_23_CA | ASP_A_23_C | -11.611 | -5.267 | 4.569 | -10.166 | -5.17 | 4.077 | 1.53 | 108.76 | 3.84 |
| 136 | 2jsa-7 | ASP_A_23_C | ASP_A_23_O | -10.166 | -5.17 | 4.077 | -9.924 | -4.918 | 2.897 | 1.23 | 163.51 | 46.16 |
| 137 | 2jsa-7 | ASP_A_23_C | ALA_A_24_N | -10.166 | -5.17 | 4.077 | -9.243 | -5.375 | 5.005 | 1.32 | 45.53 | -12.52 |
| 138 | 2jsa-7 | ALA_A_24_N | ALA_A_24_H | -9.243 | -5.375 | 5.005 | -9.515 | -5.577 | 5.956 | 1.01 | 19.61 | -143.40 |
| 139 | 2jsa-7 | ALA_A_24_N | ALA_A_24_CA | -9.243 | -5.375 | 5.005 | -7.828 | -5.313 | 4.681 | 1.45 | 102.89 | 2.51 |
| 140 | 2jsa-7 | ALA_A_24_CA | ALA_A_24_HA | -7.828 | -5.313 | 4.681 | -7.611 | -4.339 | 4.242 | 1.09 | 113.75 | 77.44 |
| 141 | 2jsa-7 | ALA_A_24_CA | ALA_A_24_C | -7.828 | -5.313 | 4.681 | -7.499 | -6.395 | 3.649 | 1.53 | 132.38 | -73.09 |
| 142 | 2jsa-7 | ALA_A_24_C | ALA_A_24_O | -7.499 | -6.395 | 3.649 | -6.43 | -6.371 | 3.041 | 1.23 | 119.62 | 1.29 |
| 143 | 2jsa-7 | ALA_A_24_C | ASN_A_25_N | -7.499 | -6.395 | 3.649 | -8.436 | -7.316 | 3.484 | 1.32 | 97.16 | -135.49 |
| 144 | 2jsa-7 | ASN_A_25_N | ASN_A_25_H | -8.436 | -7.316 | 3.484 | -9.291 | -7.271 | 4.019 | 1.01 | 58.00 | 176.99 |
| 145 | 2jsa-7 | ASN_A_25_N | ASN_A_25_CA | -8.436 | -7.316 | 3.484 | -8.259 | -8.404 | 2.538 | 1.45 | 130.64 | -80.76 |
| 146 | 2jsa-7 | ASN_A_25_CA | ASN_A_25_HA | -8.259 | -8.404 | 2.538 | -7.23 | -8.759 | 2.592 | 1.09 | 87.16 | -19.03 |
| 147 | 2jsa-7 | ASN_A_25_CA | ASN_A_25_C | -8.259 | -8.404 | 2.538 | -8.562 | -7.898 | 1.125 | 1.53 | 157.34 | 120.91 |
| 148 | 2jsa-7 | ASN_A_25_C | ASN_A_25_O | -8.562 | -7.898 | 1.125 | -7.89 | -8.281 | 0.168 | 1.23 | 141.05 | -29.68 |
| 149 | 2jsa-7 | ASN_A_25_C | VAL_A_26_N | -8.562 | -7.898 | 1.125 | -9.573 | -7.046 | 1.04 | 1.32 | 93.68 | 139.88 |
| 150 | 2jsa-7 | VAL_A_26_N | VAL_A_26_H | -9.573 | -7.046 | 1.04 | -10.08 | -6.775 | 1.87 | 1.01 | 34.71 | 151.87 |
| 151 | 2jsa-7 | VAL_A_26_N | VAL_A_26_CA | -9.573 | -7.046 | 1.04 | -9.973 | -6.485 | -0.239 | 1.45 | 151.69 | 125.49 |
| 152 | 2jsa-7 | VAL_A_26_CA | VAL_A_26_HA | -9.973 | -6.485 | -0.239 | -9.754 | -7.215 | -1.019 | 1.09 | 135.66 | -73.30 |
| 153 | 2jsa-7 | VAL_A_26_CA | VAL_A_26_C | -9.973 | -6.485 | -0.239 | -9.147 | -5.228 | -0.52 | 1.53 | 100.58 | 56.69 |
| 154 | 2jsa-7 | VAL_A_26_C | VAL_A_26_O | -9.147 | -5.228 | -0.52 | -9.217 | -4.667 | -1.612 | 1.23 | 152.63 | 97.11 |
| 155 | 2jsa-7 | VAL_A_26_C | ILE_A_27_N | -9.147 | -5.228 | -0.52 | -8.386 | -4.822 | 0.485 | 1.32 | 40.64 | 28.08 |
| 156 | 2jsa-7 | ILE_A_27_N | ILE_A_27_H | -8.386 | -4.822 | 0.485 | -8.378 | -5.332 | 1.357 | 1.01 | 30.32 | -89.10 |
| 157 | 2jsa-7 | ILE_A_27_N | ILE_A_27_CA | -8.386 | -4.822 | 0.485 | -7.548 | -3.642 | 0.359 | 1.45 | 94.98 | 54.62 |
| 158 | 2jsa-7 | ILE_A_27_CA | ILE_A_27_HA | -7.548 | -3.642 | 0.359 | -7.986 | -2.996 | -0.402 | 1.09 | 134.28 | 124.14 |
| 159 | 2jsa-7 | ILE_A_27_CA | ILE_A_27_C | -7.548 | -3.642 | 0.359 | -6.159 | -4.057 | -0.128 | 1.53 | 108.57 | -16.63 |
| 160 | 2jsa-7 | ILE_A_27_C | ILE_A_27_O | -6.159 | -4.057 | -0.128 | -5.711 | -3.616 | -1.184 | 1.23 | 149.23 | 44.55 |
| 161 | 2jsa-7 | ILE_A_27_C | LYS_A_28_N | -6.159 | -4.057 | -0.128 | -5.515 | -4.901 | 0.666 | 1.33 | 53.21 | -52.66 |
| 162 | 2jsa-7 | LYS_A_28_N | LYS_A_28_H | -5.515 | -4.901 | 0.666 | -5.944 | -5.222 | 1.522 | 1.01 | 32.04 | -143.19 |
| 163 | 2jsa-7 | LYS_A_28_N | LYS_A_28_CA | -5.515 | -4.901 | 0.666 | -4.186 | -5.381 | 0.328 | 1.45 | 103.45 | -19.86 |
| 164 | 2jsa-7 | LYS_A_28_CA | LYS_A_28_HA | -4.186 | -5.381 | 0.328 | -3.511 | -4.525 | 0.312 | 1.09 | 90.84 | 51.74 |
| 165 | 2jsa-7 | LYS_A_28_CA | LYS_A_28_C | -4.186 | -5.381 | 0.328 | -4.208 | -5.982 | -1.079 | 1.53 | 156.86 | -92.10 |
| 166 | 2jsa-7 | LYS_A_28_C | LYS_A_28_O | -4.208 | -5.982 | -1.079 | -3.314 | -5.722 | -1.882 | 1.23 | 130.78 | 16.22 |
| 167 | 2jsa-7 | LYS_A_28_C | LYS_A_29_N | -4.208 | -5.982 | -1.079 | -5.239 | -6.773 | -1.334 | 1.32 | 101.10 | -142.50 |
| 168 | 2jsa-7 | LYS_A_29_N | LYS_A_29_H | -5.239 | -6.773 | -1.334 | -5.938 | -6.942 | -0.625 | 1.01 | 45.41 | -166.41 |
| 169 | 2jsa-7 | LYS_A_29_N | LYS_A_29_CA | -5.239 | -6.773 | -1.334 | -5.388 | -7.412 | -2.63 | 1.45 | 153.15 | -103.13 |
| 170 | 2jsa-7 | LYS_A_29_CA | LYS_A_29_HA | -5.388 | -7.412 | -2.63 | -4.443 | -7.895 | -2.879 | 1.09 | 103.20 | -27.07 |
| 171 | 2jsa-7 | LYS_A_29_CA | LYS_A_29_C | -5.388 | -7.412 | -2.63 | -5.659 | -6.346 | -3.693 | 1.53 | 134.02 | 104.26 |
| 172 | 2jsa-7 | LYS_A_29_C | LYS_A_29_O | -5.659 | -6.346 | -3.693 | -5.031 | -6.345 | -4.75 | 1.23 | 149.28 | 0.09 |
| 173 | 2jsa-7 | LYS_A_29_C | ASP_A_30_N | -5.659 | -6.346 | -3.693 | -6.593 | -5.462 | -3.374 | 1.32 | 76.07 | 136.58 |
| 174 | 2jsa-7 | ASP_A_30_N | ASP_A_30_H | -6.593 | -5.462 | -3.374 | -7.068 | -5.523 | -2.485 | 1.01 | 28.31 | -172.68 |
| 175 | 2jsa-7 | ASP_A_30_N | ASP_A_30_CA | -6.593 | -5.462 | -3.374 | -6.954 | -4.392 | -4.287 | 1.45 | 128.96 | 108.64 |
| 176 | 2jsa-7 | ASP_A_30_CA | ASP_A_30_HA | -6.954 | -4.392 | -4.287 | -7.497 | -4.814 | -5.132 | 1.09 | 140.86 | -142.15 |
| 177 | 2jsa-7 | ASP_A_30_CA | ASP_A_30_C | -6.954 | -4.392 | -4.287 | -5.684 | -3.698 | -4.781 | 1.53 | 108.85 | 28.65 |
| 178 | 2jsa-7 | ASP_A_30_C | ASP_A_30_O | -5.684 | -3.698 | -4.781 | -5.48 | -3.552 | -5.985 | 1.23 | 168.23 | 35.59 |
| 179 | 2jsa-7 | ASP_A_30_C | PHE_A_31_N | -5.684 | -3.698 | -4.781 | -4.862 | -3.289 | -3.825 | 1.33 | 43.84 | 26.45 |
| 180 | 2jsa-7 | PHE_A_31_N | PHE_A_31_H | -4.862 | -3.289 | -3.825 | -5.094 | -3.443 | -2.854 | 1.01 | 16.00 | -146.42 |
| 181 | 2jsa-7 | PHE_A_31_N | PHE_A_31_CA | -4.862 | -3.289 | -3.825 | -3.616 | -2.614 | -4.147 | 1.45 | 102.80 | 28.45 |
| 182 | 2jsa-7 | PHE_A_31_CA | PHE_A_31_HA | -3.616 | -2.614 | -4.147 | -3.815 | -1.576 | -4.414 | 1.09 | 104.18 | 100.85 |
| 183 | 2jsa-7 | PHE_A_31_CA | PHE_A_31_C | -3.616 | -2.614 | -4.147 | -2.897 | -3.313 | -5.302 | 1.53 | 139.04 | -44.19 |
| 184 | 2jsa-7 | PHE_A_31_C | PHE_A_31_O | -2.897 | -3.313 | -5.302 | -2.597 | -2.689 | -6.32 | 1.23 | 145.78 | 64.32 |
| 185 | 2jsa-7 | PHE_A_31_C | ASP_A_32_N | -2.897 | -3.313 | -5.302 | -2.64 | -4.598 | -5.106 | 1.33 | 81.49 | -78.69 |
| 186 | 2jsa-7 | ASP_A_32_N | ASP_A_32_H | -2.64 | -4.598 | -5.106 | -2.914 | -5.046 | -4.243 | 1.01 | 31.32 | -121.45 |
| 187 | 2jsa-7 | ASP_A_32_N | ASP_A_32_CA | -2.64 | -4.598 | -5.106 | -1.961 | -5.388 | -6.119 | 1.45 | 134.20 | -49.32 |
| 188 | 2jsa-7 | ASP_A_32_CA | ASP_A_32_HA | -1.961 | -5.388 | -6.119 | -0.917 | -5.079 | -6.171 | 1.09 | 92.73 | 16.49 |
| 189 | 2jsa-7 | ASP_A_32_CA | ASP_A_32_C | -1.961 | -5.388 | -6.119 | -2.638 | -5.164 | -7.473 | 1.53 | 152.23 | 161.69 |
| 190 | 2jsa-7 | ASP_A_32_C | ASP_A_32_O | -2.638 | -5.164 | -7.473 | -1.967 | -5.1 | -8.502 | 1.23 | 146.77 | 5.45 |
| 191 | 2jsa-7 | ASP_A_32_C | ALA_A_33_N | -2.638 | -5.164 | -7.473 | -3.957 | -5.05 | -7.428 | 1.32 | 88.05 | 175.06 |
| 192 | 2jsa-7 | ALA_A_33_N | ALA_A_33_H | -3.957 | -5.05 | -7.428 | -4.443 | -5.113 | -6.545 | 1.01 | 29.03 | -172.61 |
| 193 | 2jsa-7 | ALA_A_33_N | ALA_A_33_CA | -3.957 | -5.05 | -7.428 | -4.731 | -4.835 | -8.638 | 1.45 | 146.42 | 164.48 |
| 194 | 2jsa-7 | ALA_A_33_CA | ALA_A_33_HA | -4.731 | -4.835 | -8.638 | -4.674 | -5.732 | -9.254 | 1.09 | 124.42 | -86.36 |
| 195 | 2jsa-7 | ALA_A_33_CA | ALA_A_33_C | -4.731 | -4.835 | -8.638 | -4.127 | -3.669 | -9.423 | 1.53 | 120.87 | 62.62 |
| 196 | 2jsa-7 | ALA_A_33_C | ALA_A_33_O | -4.127 | -3.669 | -9.423 | -3.761 | -3.824 | -10.588 | 1.23 | 161.16 | -22.95 |
| 197 | 2jsa-7 | ALA_A_33_C | GLU_A_34_N | -4.127 | -3.669 | -9.423 | -4.042 | -2.528 | -8.756 | 1.32 | 59.76 | 85.74 |
| 198 | 2jsa-7 | GLU_A_34_N | GLU_A_34_H | -4.042 | -2.528 | -8.756 | -4.362 | -2.473 | -7.8 | 1.01 | 18.76 | 170.25 |
| 199 | 2jsa-7 | GLU_A_34_N | GLU_A_34_CA | -4.042 | -2.528 | -8.756 | -3.489 | -1.337 | -9.377 | 1.45 | 115.31 | 65.09 |
| 200 | 2jsa-7 | GLU_A_34_CA | GLU_A_34_HA | -3.489 | -1.337 | -9.377 | -3.943 | -1.228 | -10.362 | 1.09 | 154.64 | 166.50 |
| 201 | 2jsa-7 | GLU_A_34_CA | GLU_A_34_C | -3.489 | -1.337 | -9.377 | -1.977 | -1.486 | -9.559 | 1.53 | 96.83 | -5.63 |
| 202 | 2jsa-7 | GLU_A_34_C | GLU_A_34_O | -1.977 | -1.486 | -9.559 | -1.466 | -1.365 | -10.671 | 1.23 | 154.72 | 13.32 |
| 203 | 2jsa-7 | GLU_A_34_C | CYS_A_35_N | -1.977 | -1.486 | -9.559 | -1.303 | -1.747 | -8.447 | 1.33 | 33.02 | -21.17 |
| 204 | 2jsa-7 | CYS_A_35_N | CYS_A_35_H | -1.303 | -1.747 | -8.447 | -1.788 | -1.834 | -7.566 | 1.01 | 29.22 | -169.83 |
| 205 | 2jsa-7 | CYS_A_35_N | CYS_A_35_CA | -1.303 | -1.747 | -8.447 | 0.14 | -1.914 | -8.47 | 1.45 | 90.91 | -6.60 |
| 206 | 2jsa-7 | CYS_A_35_CA | CYS_A_35_HA | 0.14 | -1.914 | -8.47 | 0.599 | -0.926 | -8.522 | 1.09 | 92.73 | 65.08 |
| 207 | 2jsa-7 | CYS_A_35_CA | CYS_A_35_C | 0.14 | -1.914 | -8.47 | 0.515 | -2.692 | -9.733 | 1.53 | 145.64 | -64.27 |
| 208 | 2jsa-7 | CYS_A_35_C | CYS_A_35_O | 0.515 | -2.692 | -9.733 | 1.595 | -2.496 | -10.289 | 1.23 | 116.86 | 10.29 |
| 209 | 2jsa-7 | CYS_A_35_C | LYS_A_36_N | 0.515 | -2.692 | -9.733 | -0.399 | -3.556 | -10.151 | 1.33 | 108.38 | -136.61 |

| | | | | | | | | | | | | |
|---|---|---|---|---|---|---|---|---|---|---|---|---|
| 210 | 2jsa-7 | LYS_A_36_N | LYS_A_36_H | -0.399 | -3.556 | -10.151 | -1.266 | -3.663 | -9.645 | 1.01 | 59.92 | -172.96 |
| 211 | 2jsa-7 | LYS_A_36_N | LYS_A_36_CA | -0.399 | -3.556 | -10.151 | -0.178 | -4.364 | -11.338 | 1.45 | 144.79 | -74.70 |
| 212 | 2jsa-7 | LYS_A_36_CA | LYS_A_36_HA | -0.178 | -4.364 | -11.338 | 0.816 | -4.807 | -11.265 | 1.09 | 86.16 | -24.02 |
| 213 | 2jsa-7 | LYS_A_36_CA | LYS_A_36_C | -0.178 | -4.364 | -11.338 | -0.196 | -3.461 | -12.573 | 1.53 | 143.82 | 91.14 |
| 214 | 2jsa-7 | LYS_A_36_C | LYS_A_36_O | -0.196 | -3.461 | -12.573 | 0.855 | -3.132 | -13.121 | 1.23 | 116.45 | 17.38 |
| 215 | 2jsa-7 | LYS_A_36_C | LYS_A_37_N | -0.196 | -3.461 | -12.573 | -1.402 | -3.088 | -12.977 | 1.33 | 107.75 | 162.81 |
| 216 | 2jsa-7 | LYS_A_37_N | LYS_A_37_H | -1.402 | -3.088 | -12.977 | -2.223 | -3.398 | -12.479 | 1.01 | 60.43 | -159.31 |
| 217 | 2jsa-7 | LYS_A_37_N | LYS_A_37_CA | -1.402 | -3.088 | -12.977 | -1.57 | -2.23 | -14.138 | 1.45 | 143.02 | 101.08 |
| 218 | 2jsa-7 | LYS_A_37_CA | LYS_A_37_HA | -1.57 | -2.23 | -14.138 | -1.117 | -2.729 | -14.995 | 1.09 | 141.82 | -47.77 |
| 219 | 2jsa-7 | LYS_A_37_CA | LYS_A_37_C | -1.57 | -2.23 | -14.138 | -0.815 | -0.919 | -13.912 | 1.53 | 81.50 | 60.06 |
| 220 | 2jsa-7 | LYS_A_37_C | LYS_A_37_O | -0.815 | -0.919 | -13.912 | -0.56 | -0.175 | -14.856 | 1.23 | 140.20 | 71.08 |
| 221 | 2jsa-7 | LYS_A_37_C | LEU_A_38_N | -0.815 | -0.919 | -13.912 | -0.479 | -0.676 | -12.653 | 1.33 | 18.23 | 35.87 |
| 222 | 2jsa-7 | LEU_A_38_N | LEU_A_38_H | -0.479 | -0.676 | -12.653 | -0.722 | -1.333 | -11.926 | 1.01 | 43.94 | -110.30 |
| 223 | 2jsa-7 | LEU_A_38_N | LEU_A_38_CA | -0.479 | -0.676 | -12.653 | 0.242 | 0.532 | -12.29 | 1.45 | 75.53 | 59.17 |
| 224 | 2jsa-7 | LEU_A_38_CA | LEU_A_38_HA | 0.242 | 0.532 | -12.29 | -0.155 | 1.354 | -12.886 | 1.09 | 123.14 | 115.78 |
| 225 | 2jsa-7 | LEU_A_38_CA | LEU_A_38_C | 0.242 | 0.532 | -12.29 | 1.719 | 0.368 | -12.652 | 1.53 | 103.69 | -6.34 |
| 226 | 2jsa-7 | LEU_A_38_C | LEU_A_38_O | 1.719 | 0.368 | -12.652 | 2.189 | 0.941 | -13.634 | 1.23 | 142.96 | 50.64 |
| 227 | 2jsa-7 | LEU_A_38_C | PHE_A_39_N | 1.719 | 0.368 | -12.652 | 2.411 | -0.417 | -11.839 | 1.33 | 52.16 | -48.60 |
| 228 | 2jsa-7 | PHE_A_39_N | PHE_A_39_H | 2.411 | -0.417 | -11.839 | 1.959 | -0.856 | -11.05 | 1.01 | 38.61 | -135.84 |
| 229 | 2jsa-7 | PHE_A_39_N | PHE_A_39_CA | 2.411 | -0.417 | -11.839 | 3.825 | -0.664 | -12.06 | 1.45 | 98.75 | -9.91 |
| 230 | 2jsa-7 | PHE_A_39_CA | PHE_A_39_HA | 3.825 | -0.664 | -12.06 | 4.275 | 0.189 | -12.568 | 1.09 | 117.78 | 62.19 |
| 231 | 2jsa-7 | PHE_A_39_CA | PHE_A_39_C | 3.825 | -0.664 | -12.06 | 4.037 | -1.93 | -12.894 | 1.53 | 123.01 | -80.49 |
| 232 | 2jsa-7 | PHE_A_39_C | PHE_A_39_O | 4.037 | -1.93 | -12.894 | 5.039 | -2.624 | -12.73 | 1.23 | 82.34 | -34.71 |
| 233 | 2jsa-7 | PHE_A_39_C | HIS_A_40_N | 4.037 | -1.93 | -12.894 | 3.076 | -2.192 | -13.768 | 1.33 | 131.27 | -164.75 |
| 234 | 2jsa-7 | HIS_A_40_N | HIS_A_40_H | 3.076 | -2.192 | -13.768 | 2.279 | -1.577 | -13.847 | 1.01 | 94.49 | 142.34 |
| 235 | 2jsa-7 | HIS_A_40_N | HIS_A_40_CA | 3.076 | -2.192 | -13.768 | 3.146 | -3.362 | -14.626 | 1.45 | 126.21 | -86.58 |
| 236 | 2jsa-7 | HIS_A_40_CA | HIS_A_40_HA | 3.146 | -3.362 | -14.626 | 3.037 | -4.247 | -13.998 | 1.09 | 54.84 | -97.02 |
| 237 | 2jsa-7 | HIS_A_40_CA | HIS_A_40_C | 3.146 | -3.362 | -14.626 | 4.519 | -3.428 | -15.295 | 1.53 | 115.95 | -2.75 |
| 238 | 2jsa-7 | HIS_A_40_C | HIS_A_40_O | 4.519 | -3.428 | -15.295 | 5.248 | -4.406 | -15.131 | 1.23 | 82.34 | -53.30 |
| 239 | 2jsa-7 | HIS_A_40_C | THR_A_41_N | 4.519 | -3.428 | -15.295 | 4.832 | -2.375 | -16.036 | 1.33 | 124.00 | 73.45 |
| 240 | 2jsa-7 | THR_A_41_N | THR_A_41_H | 4.832 | -2.375 | -16.036 | 4.184 | -1.606 | -16.126 | 1.01 | 95.11 | 130.12 |
| 241 | 2jsa-7 | THR_A_41_N | THR_A_41_CA | 4.832 | -2.375 | -16.036 | 6.107 | -2.301 | -16.731 | 1.45 | 118.55 | 3.32 |
| 242 | 2jsa-7 | THR_A_41_CA | THR_A_41_HA | 6.107 | -2.301 | -16.731 | 6.423 | -3.311 | -16.993 | 1.09 | 103.91 | -72.63 |
| 243 | 2jsa-7 | THR_A_41_CA | THR_A_41_C | 6.107 | -2.301 | -16.731 | 7.176 | -1.694 | -15.82 | 1.53 | 53.46 | 29.59 |
| 244 | 2jsa-7 | THR_A_41_C | THR_A_41_O | 7.176 | -1.694 | -15.82 | 7.604 | -0.561 | -16.028 | 1.23 | 99.74 | 69.31 |
| 245 | 2jsa-7 | THR_A_41_C | ILE_A_42_N | 7.176 | -1.694 | -15.82 | 7.577 | -2.477 | -14.829 | 1.33 | 41.60 | -62.88 |
| 246 | 2jsa-7 | ILE_A_42_N | ILE_A_42_H | 7.577 | -2.477 | -14.829 | 7.183 | -3.4 | -14.718 | 1.01 | 83.69 | -113.12 |
| 247 | 2jsa-7 | ILE_A_42_N | ILE_A_42_CA | 7.577 | -2.477 | -14.829 | 8.588 | -2.031 | -13.885 | 1.45 | 49.49 | 23.80 |
| 248 | 2jsa-7 | ILE_A_42_CA | ILE_A_42_HA | 8.588 | -2.031 | -13.885 | 9.277 | -1.373 | -14.414 | 1.09 | 119.04 | 43.68 |
| 249 | 2jsa-7 | ILE_A_42_CA | ILE_A_42_C | 8.588 | -2.031 | -13.885 | 9.386 | -3.241 | -13.392 | 1.53 | 71.22 | -56.59 |
| 250 | 2jsa-7 | ILE_A_42_C | ILE_A_42_O | 9.386 | -3.241 | -13.392 | 8.824 | -4.312 | -13.173 | 1.23 | 79.74 | -117.69 |
| 251 | 2jsa-7 | ILE_A_42_C | PRO_A_43_N | 9.386 | -3.241 | -13.392 | 10.718 | -3.021 | -13.227 | 1.36 | 83.03 | 9.38 |
| 252 | 2jsa-7 | PRO_A_43_N | PRO_A_43_CA | 10.718 | -3.021 | -13.227 | 11.599 | -4.079 | -12.763 | 1.45 | 71.38 | -50.22 |
| 253 | 2jsa-7 | PRO_A_43_CA | PRO_A_43_HA | 11.599 | -4.079 | -12.763 | 11.381 | -5.005 | -13.294 | 1.09 | 119.17 | -103.25 |
| 254 | 2jsa-7 | PRO_A_43_CA | PRO_A_43_C | 11.599 | -4.079 | -12.763 | 11.418 | -4.326 | -11.263 | 1.53 | 11.54 | -126.23 |
| 255 | 2jsa-7 | PRO_A_43_C | PRO_A_43_O | 11.418 | -4.326 | -11.263 | 11.755 | -3.471 | -10.445 | 1.23 | 48.33 | 68.49 |
| 256 | 2jsa-7 | PRO_A_43_C | PHE_A_44_N | 11.418 | -4.326 | -11.263 | 10.886 | -5.497 | -10.948 | 1.32 | 76.24 | -114.43 |
| 257 | 2jsa-7 | PHE_A_44_N | PHE_A_44_H | 10.886 | -5.497 | -10.948 | 10.633 | -6.151 | -11.674 | 1.01 | 135.99 | -111.15 |
| 258 | 2jsa-7 | PHE_A_44_N | PHE_A_44_CA | 10.886 | -5.497 | -10.948 | 10.656 | -5.867 | -9.562 | 1.45 | 17.45 | -121.87 |
| 259 | 2jsa-7 | PHE_A_44_CA | PHE_A_44_HA | 10.656 | -5.867 | -9.562 | 10.49 | -6.942 | -9.488 | 1.09 | 86.11 | -98.78 |
| 260 | 2jsa-7 | PHE_A_44_CA | PHE_A_44_C | 10.656 | -5.867 | -9.562 | 9.452 | -5.119 | -8.988 | 1.53 | 67.95 | 148.15 |
| 261 | 2jsa-7 | PHE_A_44_C | PHE_A_44_O | 9.452 | -5.119 | -8.988 | 9.106 | -5.294 | -7.82 | 1.23 | 18.36 | -153.17 |
| 262 | 2jsa-7 | PHE_A_44_C | GLY_A_45_N | 9.452 | -5.119 | -8.988 | 8.844 | -4.301 | -9.835 | 1.33 | 129.73 | 126.62 |
| 263 | 2jsa-7 | GLY_A_45_N | GLY_A_45_H | 8.844 | -4.301 | -9.835 | 9.183 | -4.206 | -10.782 | 1.01 | 159.61 | 15.65 |
| 264 | 2jsa-7 | GLY_A_45_N | GLY_A_45_CA | 8.844 | -4.301 | -9.835 | 7.685 | -3.526 | -9.426 | 1.45 | 73.65 | 146.23 |
| 265 | 2jsa-7 | GLY_A_45_CA | GLY_A_45_C | 7.685 | -3.526 | -9.426 | 6.569 | -4.437 | -8.913 | 1.53 | 70.40 | -140.77 |
| 266 | 2jsa-7 | GLY_A_45_C | GLY_A_45_O | 6.569 | -4.437 | -8.913 | 6.19 | -4.363 | -7.745 | 1.23 | 18.29 | 168.95 |
| 267 | 2jsa-7 | GLY_A_45_C | THR_A_46_N | 6.569 | -4.437 | -8.913 | 6.074 | -5.277 | -9.811 | 1.33 | 132.65 | -120.51 |
| 268 | 2jsa-7 | THR_A_46_N | THR_A_46_H | 6.074 | -5.277 | -9.811 | 6.435 | -5.284 | -10.754 | 1.01 | 159.05 | -1.11 |
| 269 | 2jsa-7 | THR_A_46_N | THR_A_46_CA | 6.074 | -5.277 | -9.811 | 5.009 | -6.202 | -9.463 | 1.45 | 76.14 | -139.02 |
| 270 | 2jsa-7 | THR_A_46_CA | THR_A_46_HA | 5.009 | -6.202 | -9.463 | 4.161 | -5.632 | -9.084 | 1.09 | 69.65 | 146.09 |
| 271 | 2jsa-7 | THR_A_46_CA | THR_A_46_C | 5.009 | -6.202 | -9.463 | 5.468 | -7.153 | -8.357 | 1.53 | 43.67 | -64.24 |
| 272 | 2jsa-7 | THR_A_46_C | THR_A_46_O | 5.468 | -7.153 | -8.357 | 4.652 | -7.659 | -7.589 | 1.23 | 51.34 | -148.20 |
| 273 | 2jsa-7 | THR_A_46_C | ARG_A_47_N | 5.468 | -7.153 | -8.357 | 6.775 | -7.369 | -8.311 | 1.33 | 88.01 | -9.38 |
| 274 | 2jsa-7 | ARG_A_47_N | ARG_A_47_H | 6.775 | -7.369 | -8.311 | 7.388 | -6.92 | -8.976 | 1.01 | 131.19 | 36.22 |
| 275 | 2jsa-7 | ARG_A_47_N | ARG_A_47_CA | 6.775 | -7.369 | -8.311 | 7.354 | -8.25 | -7.312 | 1.45 | 46.54 | -56.69 |
| 276 | 2jsa-7 | ARG_A_47_CA | ARG_A_47_HA | 7.354 | -8.25 | -7.312 | 6.821 | -9.2 | -7.342 | 1.09 | 91.58 | -119.29 |
| 277 | 2jsa-7 | ARG_A_47_CA | ARG_A_47_C | 7.354 | -8.25 | -7.312 | 7.206 | -7.64 | -5.916 | 1.53 | 24.21 | 103.64 |
| 278 | 2jsa-7 | ARG_A_47_C | ARG_A_47_O | 7.206 | -7.64 | -5.916 | 6.841 | -8.333 | -4.968 | 1.23 | 39.56 | -117.78 |
| 279 | 2jsa-7 | ARG_A_47_C | GLU_A_48_N | 7.206 | -7.64 | -5.916 | 7.497 | -6.35 | -5.835 | 1.32 | 86.49 | 77.29 |
| 280 | 2jsa-7 | GLU_A_48_N | GLU_A_48_H | 7.497 | -6.35 | -5.835 | 7.791 | -5.842 | -6.657 | 1.01 | 144.47 | 59.94 |
| 281 | 2jsa-7 | GLU_A_48_N | GLU_A_48_CA | 7.497 | -6.35 | -5.835 | 7.4 | -5.639 | -4.571 | 1.45 | 29.58 | 97.77 |
| 282 | 2jsa-7 | GLU_A_48_CA | GLU_A_48_HA | 7.4 | -5.639 | -4.571 | 7.859 | -6.258 | -3.799 | 1.09 | 44.95 | -53.44 |
| 283 | 2jsa-7 | GLU_A_48_CA | GLU_A_48_C | 7.4 | -5.639 | -4.571 | 5.934 | -5.415 | -4.199 | 1.53 | 75.92 | 171.31 |
| 284 | 2jsa-7 | GLU_A_48_C | GLU_A_48_O | 5.934 | -5.415 | -4.199 | 5.579 | -5.437 | -3.021 | 1.23 | 16.80 | -176.45 |
| 285 | 2jsa-7 | GLU_A_48_C | CYS_A_49_N | 5.934 | -5.415 | -4.199 | 5.121 | -5.206 | -5.223 | 1.32 | 140.66 | 165.58 |
| 286 | 2jsa-7 | CYS_A_49_N | CYS_A_49_H | 5.121 | -5.206 | -5.223 | 5.482 | -5.203 | -6.166 | 1.01 | 159.05 | 0.48 |
| 287 | 2jsa-7 | CYS_A_49_N | CYS_A_49_CA | 5.121 | -5.206 | -5.223 | 3.7 | -4.978 | -5.018 | 1.45 | 81.89 | 170.88 |
| 288 | 2jsa-7 | CYS_A_49_CA | CYS_A_49_HA | 3.7 | -4.978 | -5.018 | 3.577 | -4.028 | -4.497 | 1.09 | 61.46 | 97.38 |
| 289 | 2jsa-7 | CYS_A_49_CA | CYS_A_49_C | 3.7 | -4.978 | -5.018 | 3.16 | -6.097 | -4.125 | 1.53 | 54.29 | -115.76 |
| 290 | 2jsa-7 | CYS_A_49_C | CYS_A_49_O | 3.16 | -6.097 | -4.125 | 2.422 | -5.836 | -3.175 | 1.23 | 39.49 | 160.52 |
| 291 | 2jsa-7 | CYS_A_49_C | ASP_A_50_N | 3.16 | -6.097 | -4.125 | 3.546 | -7.319 | -4.462 | 1.33 | 104.73 | -72.47 |
| 292 | 2jsa-7 | ASP_A_50_N | ASP_A_50_H | 3.546 | -7.319 | -4.462 | 4.153 | -7.46 | -5.257 | 1.01 | 141.91 | -13.08 |
| 293 | 2jsa-7 | ASP_A_50_N | ASP_A_50_CA | 3.546 | -7.319 | -4.462 | 3.109 | -8.478 | -3.702 | 1.45 | 58.47 | -110.66 |
| 294 | 2jsa-7 | ASP_A_50_CA | ASP_A_50_HA | 3.109 | -8.478 | -3.702 | 2.046 | -8.38 | -3.483 | 1.09 | 78.41 | 174.73 |
| 295 | 2jsa-7 | ASP_A_50_CA | ASP_A_50_C | 3.109 | -8.478 | -3.702 | 3.903 | -8.556 | -2.397 | 1.53 | 31.44 | -5.61 |
| 296 | 2jsa-7 | ASP_A_50_C | ASP_A_50_O | 3.903 | -8.556 | -2.397 | 3.435 | -9.13 | -1.414 | 1.23 | 36.99 | -129.19 |
| 297 | 2jsa-7 | ASP_A_50_C | HIS_A_51_N | 3.903 | -8.556 | -2.397 | 5.091 | -7.97 | -2.429 | 1.33 | 91.38 | 26.26 |
| 298 | 2jsa-7 | HIS_A_51_N | HIS_A_51_H | 5.091 | -7.97 | -2.429 | 5.413 | -7.517 | -3.272 | 1.01 | 146.60 | 54.59 |
| 299 | 2jsa-7 | HIS_A_51_N | HIS_A_51_CA | 5.091 | -7.97 | -2.429 | 5.954 | -7.966 | -1.26 | 1.45 | 36.44 | 0.27 |
| 300 | 2jsa-7 | HIS_A_51_CA | HIS_A_51_HA | 5.954 | -7.966 | -1.26 | 5.72 | -8.849 | -0.666 | 1.09 | 56.97 | -104.84 |
| 301 | 2jsa-7 | HIS_A_51_CA | HIS_A_51_C | 5.954 | -7.966 | -1.26 | 5.656 | -6.733 | -0.405 | 1.53 | 56.02 | 103.59 |
| 302 | 2jsa-7 | HIS_A_51_C | HIS_A_51_O | 5.656 | -6.733 | -0.405 | 6.43 | -6.388 | 0.486 | 1.23 | 43.56 | 24.02 |
| 303 | 2jsa-7 | HIS_A_51_C | TYR_A_52_N | 5.656 | -6.733 | -0.405 | 4.53 | -6.102 | -0.707 | 1.33 | 103.17 | 150.73 |
| 304 | 2jsa-7 | TYR_A_52_N | TYR_A_52_H | 4.53 | -6.102 | -0.707 | 3.941 | -6.441 | -1.454 | 1.01 | 137.71 | -150.08 |
| 305 | 2jsa-7 | TYR_A_52_N | TYR_A_52_CA | 4.53 | -6.102 | -0.707 | 4.12 | -4.914 | 0.024 | 1.45 | 59.82 | 109.04 |

| | | | | | | | | | | | |
|---|---|---|---|---|---|---|---|---|---|---|---|
| 306 | 2jsa-7 | TYR_A_52_CA | TYR_A_52_HA | 4.12 | -4.914 | 0.024 | 4.891 | -4.625 | 0.739 | 1.09 | 49.03 | 20.55 |
| 307 | 2jsa-7 | TYR_A_52_CA | TYR_A_52_C | 4.12 | -4.914 | 0.024 | 2.797 | -5.15 | 0.754 | 1.53 | 61.49 | -169.89 |
| 308 | 2jsa-7 | TYR_A_52_C | TYR_A_52_O | 2.797 | -5.15 | 0.754 | 2.642 | -4.751 | 1.908 | 1.23 | 20.35 | 111.23 |
| 309 | 2jsa-7 | TYR_A_52_C | VAL_A_53_N | 2.797 | -5.15 | 0.754 | 1.878 | -5.798 | 0.054 | 1.32 | 121.90 | -144.81 |
| 310 | 2jsa-7 | VAL_A_53_N | VAL_A_53_H | 1.878 | -5.798 | 0.054 | 2.076 | -6.097 | -0.89 | 1.01 | 159.20 | -56.49 |
| 311 | 2jsa-7 | VAL_A_53_N | VAL_A_53_CA | 1.878 | -5.798 | 0.054 | 0.573 | -6.092 | 0.622 | 1.45 | 66.99 | -167.30 |
| 312 | 2jsa-7 | VAL_A_53_CA | VAL_A_53_HA | 0.573 | -6.092 | 0.622 | 0.256 | -5.236 | 1.218 | 1.09 | 56.86 | 110.32 |
| 313 | 2jsa-7 | VAL_A_53_CA | VAL_A_53_C | 0.573 | -6.092 | 0.622 | 0.688 | -7.302 | 1.553 | 1.53 | 52.55 | -84.57 |
| 314 | 2jsa-7 | VAL_A_53_C | VAL_A_53_O | 0.688 | -7.302 | 1.553 | -0.047 | -8.276 | 1.404 | 1.23 | 96.96 | -127.04 |
| 315 | 2jsa-7 | VAL_A_53_C | ASN_A_54_N | 0.688 | -7.302 | 1.553 | 1.616 | -7.199 | 2.493 | 1.32 | 44.81 | 6.33 |
| 316 | 2jsa-7 | ASN_A_54_N | ASN_A_54_H | 1.616 | -7.199 | 2.493 | 2.185 | -6.367 | 2.557 | 1.01 | 86.37 | 55.63 |
| 317 | 2jsa-7 | ASN_A_54_N | ASN_A_54_CA | 1.616 | -7.199 | 2.493 | 1.836 | -8.272 | 3.447 | 1.45 | 48.94 | -78.41 |
| 318 | 2jsa-7 | ASN_A_54_CA | ASN_A_54_HA | 1.836 | -8.272 | 3.447 | 0.944 | -8.391 | 4.062 | 1.09 | 55.65 | -172.40 |
| 319 | 2jsa-7 | ASN_A_54_CA | ASN_A_54_C | 1.836 | -8.272 | 3.447 | 3.035 | -7.924 | 4.332 | 1.53 | 54.67 | 16.18 |
| 320 | 2jsa-7 | ASN_A_54_C | ASN_A_54_O | 3.035 | -7.924 | 4.332 | 3.854 | -8.787 | 4.642 | 1.23 | 75.40 | -46.50 |
| 321 | 2jsa-7 | ASN_A_54_C | SER_A_55_N | 3.035 | -7.924 | 4.332 | 3.1 | -6.657 | 4.714 | 1.32 | 73.24 | 87.06 |
| 322 | 2jsa-7 | SER_A_55_N | SER_A_55_H | 3.1 | -6.657 | 4.714 | 2.39 | -6.001 | 4.422 | 1.01 | 106.81 | 137.26 |
| 323 | 2jsa-7 | SER_A_55_N | SER_A_55_CA | 3.1 | -6.657 | 4.714 | 4.185 | -6.184 | 5.557 | 1.45 | 54.54 | 23.55 |
| 324 | 2jsa-7 | SER_A_55_CA | SER_A_55_HA | 4.185 | -6.184 | 5.557 | 4.121 | -6.696 | 6.517 | 1.09 | 28.26 | -97.13 |
| 325 | 2jsa-7 | SER_A_55_CA | SER_A_55_C | 4.185 | -6.184 | 5.557 | 4.045 | -4.679 | 5.794 | 1.53 | 81.09 | 95.31 |
| 326 | 2jsa-7 | SER_A_55_C | SER_A_55_O | 4.045 | -4.679 | 5.794 | 4.292 | -4.195 | 6.898 | 1.23 | 26.21 | 62.96 |
| 327 | 2jsa-7 | SER_A_55_C | LYS_A_56_N | 4.045 | -4.679 | 5.794 | 3.649 | -3.98 | 4.74 | 1.33 | 142.68 | 119.53 |
| 328 | 2jsa-7 | LYS_A_56_N | LYS_A_56_H | 3.649 | -3.98 | 4.74 | 3.464 | -4.444 | 3.863 | 1.01 | 150.34 | -111.74 |
| 329 | 2jsa-7 | LYS_A_56_N | LYS_A_56_CA | 3.649 | -3.98 | 4.74 | 3.473 | -2.54 | 4.82 | 1.45 | 86.84 | 96.97 |
| 330 | 2jsa-7 | LYS_A_56_CA | LYS_A_56_HA | 3.473 | -2.54 | 4.82 | 3.803 | -2.214 | 5.807 | 1.09 | 25.17 | 44.65 |
| 331 | 2jsa-7 | LYS_A_56_CA | LYS_A_56_C | 3.473 | -2.54 | 4.82 | 1.986 | -2.204 | 4.692 | 1.53 | 94.80 | 167.27 |
| 332 | 2jsa-7 | LYS_A_56_C | LYS_A_56_O | 1.986 | -2.204 | 4.692 | 1.602 | -1.037 | 4.755 | 1.23 | 87.06 | 108.21 |
| 333 | 2jsa-7 | LYS_A_56_C | VAL_A_57_N | 1.986 | -2.204 | 4.692 | 1.189 | -3.248 | 4.515 | 1.33 | 97.67 | -127.36 |
| 334 | 2jsa-7 | VAL_A_57_N | VAL_A_57_H | 1.189 | -3.248 | 4.515 | 1.575 | -4.18 | 4.472 | 1.01 | 92.44 | -67.50 |
| 335 | 2jsa-7 | VAL_A_57_N | VAL_A_57_CA | 1.189 | -3.248 | 4.515 | -0.248 | -3.079 | 4.379 | 1.45 | 95.37 | 173.29 |
| 336 | 2jsa-7 | VAL_A_57_CA | VAL_A_57_HA | -0.248 | -3.079 | 4.379 | -0.449 | -2.039 | 4.12 | 1.09 | 103.74 | 100.94 |
| 337 | 2jsa-7 | VAL_A_57_CA | VAL_A_57_C | -0.248 | -3.079 | 4.379 | -0.921 | -3.372 | 5.72 | 1.53 | 28.69 | -156.47 |
| 338 | 2jsa-7 | VAL_A_57_C | VAL_A_57_O | -0.921 | -3.372 | 5.72 | -2.028 | -2.901 | 5.979 | 1.23 | 77.85 | 156.95 |
| 339 | 2jsa-7 | VAL_A_57_C | ASP_A_58_N | -0.921 | -3.372 | 5.72 | -0.227 | -4.15 | 6.537 | 1.32 | 51.92 | -48.27 |
| 340 | 2jsa-7 | ASP_A_58_N | ASP_A_58_H | -0.227 | -4.15 | 6.537 | 0.677 | -4.503 | 6.257 | 1.01 | 106.09 | -21.33 |
| 341 | 2jsa-7 | ASP_A_58_N | ASP_A_58_CA | -0.227 | -4.15 | 6.537 | -0.745 | -4.513 | 7.846 | 1.45 | 25.79 | -144.98 |
| 342 | 2jsa-7 | ASP_A_58_CA | ASP_A_58_HA | -0.745 | -4.513 | 7.846 | -1.801 | -4.764 | 7.754 | 1.09 | 94.84 | -166.63 |
| 343 | 2jsa-7 | ASP_A_58_CA | ASP_A_58_C | -0.745 | -4.513 | 7.846 | -0.572 | -3.331 | 8.803 | 1.53 | 51.30 | 81.67 |
| 344 | 2jsa-7 | ASP_A_58_C | ASP_A_58_O | -0.572 | -3.331 | 8.803 | -1.486 | -2.999 | 9.556 | 1.23 | 52.25 | 160.04 |
| 345 | 2jsa-7 | ASP_A_58_C | PRO_A_59_N | -0.572 | -3.331 | 8.803 | 0.638 | -2.714 | 8.741 | 1.36 | 92.61 | 27.02 |
| 346 | 2jsa-7 | PRO_A_59_N | PRO_A_59_CA | 0.638 | -2.714 | 8.741 | 0.943 | -1.577 | 9.593 | 1.45 | 54.10 | 74.98 |
| 347 | 2jsa-7 | PRO_A_59_CA | PRO_A_59_HA | 0.943 | -1.577 | 9.593 | 0.617 | -1.777 | 10.614 | 1.09 | 20.54 | -148.47 |
| 348 | 2jsa-7 | PRO_A_59_CA | PRO_A_59_C | 0.943 | -1.577 | 9.593 | 0.235 | -0.315 | 9.097 | 1.53 | 108.92 | 119.29 |
| 349 | 2jsa-7 | PRO_A_59_C | PRO_A_59_O | 0.235 | -0.315 | 9.097 | -0.439 | 0.367 | 9.868 | 1.23 | 51.20 | 134.66 |
| 350 | 2jsa-7 | PRO_A_59_C | ILE_A_60_N | 0.235 | -0.315 | 9.097 | 0.412 | -0.042 | 7.813 | 1.32 | 165.78 | 57.04 |
| 351 | 2jsa-7 | ILE_A_60_N | ILE_A_60_H | 0.412 | -0.042 | 7.813 | 0.979 | -0.648 | 7.238 | 1.01 | 124.72 | -46.90 |
| 352 | 2jsa-7 | ILE_A_60_N | ILE_A_60_CA | 0.412 | -0.042 | 7.813 | -0.201 | 1.126 | 7.205 | 1.45 | 114.75 | 117.69 |
| 353 | 2jsa-7 | ILE_A_60_CA | ILE_A_60_HA | -0.201 | 1.126 | 7.205 | 0.283 | 2.013 | 7.615 | 1.09 | 67.91 | 61.38 |
| 354 | 2jsa-7 | ILE_A_60_CA | ILE_A_60_C | -0.201 | 1.126 | 7.205 | -1.68 | 1.182 | 7.594 | 1.53 | 75.27 | 177.83 |
| 355 | 2jsa-7 | ILE_A_60_C | ILE_A_60_O | -1.68 | 1.182 | 7.594 | -2.179 | 2.231 | 8.003 | 1.23 | 70.60 | 115.44 |
| 356 | 2jsa-7 | ILE_A_60_C | ILE_A_61_N | -1.68 | 1.182 | 7.594 | -2.339 | 0.042 | 7.454 | 1.32 | 96.07 | -120.03 |
| 357 | 2jsa-7 | ILE_A_61_N | ILE_A_61_H | -2.339 | 0.042 | 7.454 | -1.864 | -0.782 | 7.113 | 1.01 | 109.72 | -60.04 |
| 358 | 2jsa-7 | ILE_A_61_N | ILE_A_61_CA | -2.339 | 0.042 | 7.454 | -3.751 | -0.052 | 7.787 | 1.45 | 76.76 | -176.19 |
| 359 | 2jsa-7 | ILE_A_61_CA | ILE_A_61_HA | -3.751 | -0.052 | 7.787 | -4.267 | 0.78 | 7.307 | 1.09 | 116.12 | 121.81 |
| 360 | 2jsa-7 | ILE_A_61_CA | ILE_A_61_C | -3.751 | -0.052 | 7.787 | -3.926 | 0.103 | 9.299 | 1.53 | 8.79 | 138.47 |
| 361 | 2jsa-7 | ILE_A_61_C | ILE_A_61_O | -3.926 | 0.103 | 9.299 | -4.996 | 0.489 | 9.768 | 1.23 | 67.59 | 160.16 |
| 362 | 2jsa-7 | ILE_A_61_C | HIS_A_62_N | -3.926 | 0.103 | 9.299 | -2.858 | -0.204 | 10.021 | 1.33 | 56.99 | -16.04 |
| 363 | 2jsa-7 | HIS_A_62_N | HIS_A_62_H | -2.858 | -0.204 | 10.021 | -2.01 | -0.514 | 9.568 | 1.01 | 116.64 | -20.08 |
| 364 | 2jsa-7 | HIS_A_62_N | HIS_A_62_CA | -2.858 | -0.204 | 10.021 | -2.879 | -0.104 | 11.47 | 1.45 | 4.03 | 101.86 |
| 365 | 2jsa-7 | HIS_A_62_CA | HIS_A_62_HA | -2.879 | -0.104 | 11.47 | -3.804 | -0.558 | 11.825 | 1.09 | 70.99 | -153.86 |
| 366 | 2jsa-7 | HIS_A_62_CA | HIS_A_62_C | -2.879 | -0.104 | 11.47 | -2.891 | 1.37 | 11.882 | 1.53 | 74.38 | 90.47 |
| 367 | 2jsa-7 | HIS_A_62_C | HIS_A_62_O | -2.891 | 1.37 | 11.882 | -3.222 | 1.697 | 13.02 | 1.23 | 22.24 | 135.35 |
| 368 | 2jsa-7 | HIS_A_62_C | GLU_A_63_N | -2.891 | 1.37 | 11.882 | -2.525 | 2.219 | 10.934 | 1.32 | 135.72 | 66.68 |
| 369 | 2jsa-7 | GLU_A_63_N | GLU_A_63_H | -2.525 | 2.219 | 10.934 | -2.265 | 1.878 | 10.019 | 1.01 | 154.89 | -52.68 |
| 370 | 2jsa-7 | GLU_A_63_N | GLU_A_63_CA | -2.525 | 2.219 | 10.934 | -2.489 | 3.65 | 11.183 | 1.45 | 80.13 | 88.56 |
| 371 | 2jsa-7 | GLU_A_63_CA | GLU_A_63_HA | -2.489 | 3.65 | 11.183 | -2.209 | 3.804 | 12.225 | 1.09 | 17.05 | 28.81 |
| 372 | 2jsa-7 | GLU_A_63_CA | GLU_A_63_C | -2.489 | 3.65 | 11.183 | -3.873 | 4.262 | 10.962 | 1.53 | 98.31 | 156.15 |
| 373 | 2jsa-7 | GLU_A_63_C | GLU_A_63_O | -3.873 | 4.262 | 10.962 | -4.383 | 4.98 | 11.82 | 1.23 | 45.75 | 125.39 |
| 374 | 2jsa-7 | GLU_A_63_C | LEU_A_64_N | -3.873 | 4.262 | 10.962 | -4.442 | 3.956 | 9.804 | 1.33 | 150.84 | -151.73 |
| 375 | 2jsa-7 | LEU_A_64_N | LEU_A_64_H | -4.442 | 3.956 | 9.804 | -3.964 | 3.359 | 9.144 | 1.01 | 130.79 | -51.32 |
| 376 | 2jsa-7 | LEU_A_64_N | LEU_A_64_CA | -4.442 | 3.956 | 9.804 | -5.758 | 4.468 | 9.459 | 1.45 | 103.73 | 158.74 |
| 377 | 2jsa-7 | LEU_A_64_CA | LEU_A_64_HA | -5.758 | 4.468 | 9.459 | -5.747 | 5.549 | 9.601 | 1.09 | 82.52 | 89.42 |
| 378 | 2jsa-7 | LEU_A_64_CA | LEU_A_64_C | -5.758 | 4.468 | 9.459 | -6.796 | 3.883 | 10.418 | 1.53 | 51.17 | -150.60 |
| 379 | 2jsa-7 | LEU_A_64_C | LEU_A_64_O | -6.796 | 3.883 | 10.418 | -7.711 | 4.583 | 10.849 | 1.23 | 69.49 | 142.58 |
| 380 | 2jsa-7 | LEU_A_64_C | GLU_A_65_N | -6.796 | 3.883 | 10.418 | -6.619 | 2.606 | 10.724 | 1.33 | 76.65 | -82.11 |
| 381 | 2jsa-7 | GLU_A_65_N | GLU_A_65_H | -6.619 | 2.606 | 10.724 | -5.842 | 2.092 | 10.332 | 1.01 | 112.82 | -33.49 |
| 382 | 2jsa-7 | GLU_A_65_N | GLU_A_65_CA | -6.619 | 2.606 | 10.724 | -7.529 | 1.92 | 11.624 | 1.45 | 51.70 | -142.99 |
| 383 | 2jsa-7 | GLU_A_65_CA | GLU_A_65_HA | -7.529 | 1.92 | 11.624 | -8.542 | 2.05 | 11.243 | 1.09 | 110.46 | 172.69 |
| 384 | 2jsa-7 | GLU_A_65_CA | GLU_A_65_C | -7.529 | 1.92 | 11.624 | -7.459 | 2.537 | 13.022 | 1.53 | 23.95 | 83.53 |
| 385 | 2jsa-7 | GLU_A_65_C | GLU_A_65_O | -7.459 | 2.537 | 13.022 | -8.475 | 2.652 | 13.705 | 1.23 | 56.26 | 173.54 |
| 386 | 2jsa-7 | GLU_A_65_C | GLY_A_66_N | -7.459 | 2.537 | 13.022 | -6.249 | 2.917 | 13.407 | 1.33 | 73.11 | 17.43 |
| 387 | 2jsa-7 | GLY_A_66_N | GLY_A_66_H | -6.249 | 2.917 | 13.407 | -5.456 | 2.791 | 12.794 | 1.01 | 127.36 | -9.03 |
| 388 | 2jsa-7 | GLY_A_66_N | GLY_A_66_CA | -6.249 | 2.917 | 13.407 | -6.034 | 3.52 | 14.711 | 1.45 | 26.15 | 70.38 |
| 389 | 2jsa-7 | GLY_A_66_CA | GLY_A_66_C | -6.034 | 3.52 | 14.711 | -6.648 | 4.919 | 14.777 | 1.53 | 87.53 | 113.70 |
| 390 | 2jsa-7 | GLY_A_66_C | GLY_A_66_O | -6.648 | 4.919 | 14.777 | -7.462 | 5.202 | 15.655 | 1.23 | 44.47 | 160.83 |
| 391 | 2jsa-7 | GLY_A_66_C | GLY_A_67_N | -6.648 | 4.919 | 14.777 | -6.236 | 5.757 | 13.837 | 1.32 | 135.19 | 63.82 |
| 392 | 2jsa-7 | GLY_A_67_N | GLY_A_67_H | -6.236 | 5.757 | 13.837 | -5.564 | 5.456 | 13.146 | 1.01 | 133.18 | -24.13 |
| 393 | 2jsa-7 | GLY_A_67_N | GLY_A_67_CA | -6.236 | 5.757 | 13.837 | -6.736 | 7.119 | 13.778 | 1.45 | 92.33 | 110.16 |
| 394 | 2jsa-7 | GLY_A_67_CA | GLY_A_67_C | -6.736 | 7.119 | 13.778 | -5.728 | 8.045 | 13.095 | 1.53 | 116.52 | 42.57 |
| 395 | 2jsa-7 | GLY_A_67_C | GLY_A_67_O | -5.728 | 8.045 | 13.095 | -5.49 | 9.16 | 13.558 | 1.23 | 67.90 | 77.95 |
| 396 | 2jsa-7 | GLY_A_67_C | THR_A_68_N | -5.728 | 8.045 | 13.095 | -5.16 | 7.549 | 12.005 | 1.33 | 145.32 | -41.13 |
| 397 | 2jsa-7 | THR_A_68_N | THR_A_68_H | -5.16 | 7.549 | 12.005 | -5.403 | 6.624 | 11.682 | 1.01 | 108.66 | -104.72 |
| 398 | 2jsa-7 | THR_A_68_N | THR_A_68_CA | -5.16 | 7.549 | 12.005 | -4.183 | 8.317 | 11.254 | 1.45 | 121.15 | 38.17 |
| 399 | 2jsa-7 | THR_A_68_CA | THR_A_68_HA | -4.183 | 8.317 | 11.254 | -4.11 | 9.313 | 11.691 | 1.09 | 66.37 | 85.81 |
| 400 | 2jsa-7 | THR_A_68_CA | THR_A_68_C | -4.183 | 8.317 | 11.254 | -4.629 | 8.472 | 9.799 | 1.53 | 162.02 | 160.84 |
| 401 | 2jsa-7 | THR_A_68_C | THR_A_68_O | -4.629 | 8.472 | 9.799 | -5.161 | 7.534 | 9.206 | 1.23 | 118.81 | -119.56 |

| | | | | | | | | | | |
|---|---|---|---|---|---|---|---|---|---|---|
| 402 | 2jsa-7 | THR_A_68_C | ALA_A_69_N | -4.629 | 8.472 | 9.799 | -4.397 | 9.662 | 9.265 | 1.32 | 113.77 | 78.97 |
| 403 | 2jsa-7 | ALA_A_69_N | ALA_A_69_H | -4.397 | 9.662 | 9.265 | -3.956 | 10.388 | 9.812 | 1.01 | 57.22 | 58.72 |
| 404 | 2jsa-7 | ALA_A_69_N | ALA_A_69_CA | -4.397 | 9.662 | 9.265 | -4.769 | 9.951 | 7.891 | 1.45 | 161.08 | 142.16 |
| 405 | 2jsa-7 | ALA_A_69_CA | ALA_A_69_HA | -4.769 | 9.951 | 7.891 | -5.808 | 9.659 | 7.74 | 1.09 | 97.96 | -164.30 |
| 406 | 2jsa-7 | ALA_A_69_CA | ALA_A_69_C | -4.769 | 9.951 | 7.891 | -3.892 | 9.127 | 6.946 | 1.53 | 128.14 | -43.22 |
| 407 | 2jsa-7 | ALA_A_69_C | ALA_A_69_O | -3.892 | 9.127 | 6.946 | -2.852 | 8.61 | 7.35 | 1.23 | 70.82 | -26.43 |
| 408 | 2jsa-7 | ALA_A_69_C | PRO_A_70_N | -3.892 | 9.127 | 6.946 | -4.356 | 9.028 | 5.671 | 1.36 | 159.59 | -167.96 |
| 409 | 2jsa-7 | PRO_A_70_N | PRO_A_70_CA | -4.356 | 9.028 | 5.671 | -3.626 | 8.276 | 4.665 | 1.45 | 133.83 | -45.85 |
| 410 | 2jsa-7 | PRO_A_70_CA | PRO_A_70_HA | -3.626 | 8.276 | 4.665 | -3.3 | 7.321 | 5.077 | 1.09 | 67.79 | -71.15 |
| 411 | 2jsa-7 | PRO_A_70_CA | PRO_A_70_C | -3.626 | 8.276 | 4.665 | -2.392 | 9.047 | 4.193 | 1.53 | 107.97 | 32.00 |
| 412 | 2jsa-7 | PRO_A_70_C | PRO_A_70_O | -2.392 | 9.047 | 4.193 | -1.534 | 8.493 | 3.506 | 1.23 | 123.93 | -32.85 |
| 413 | 2jsa-7 | PRO_A_70_C | LYS_A_71_N | -2.392 | 9.047 | 4.193 | -2.341 | 10.313 | 4.579 | 1.32 | 73.06 | 87.69 |
| 414 | 2jsa-7 | LYS_A_71_N | LYS_A_71_H | -2.341 | 10.313 | 4.579 | -3.082 | 10.703 | 5.143 | 1.01 | 56.04 | 152.24 |
| 415 | 2jsa-7 | LYS_A_71_N | LYS_A_71_CA | -2.341 | 10.313 | 4.579 | -1.226 | 11.166 | 4.204 | 1.45 | 104.96 | 37.42 |
| 416 | 2jsa-7 | LYS_A_71_CA | LYS_A_71_HA | -1.226 | 11.166 | 4.204 | -0.679 | 10.672 | 3.401 | 1.09 | 137.45 | -42.09 |
| 417 | 2jsa-7 | LYS_A_71_CA | LYS_A_71_C | -1.226 | 11.166 | 4.204 | -0.275 | 11.309 | 5.394 | 1.53 | 38.94 | 8.55 |
| 418 | 2jsa-7 | LYS_A_71_C | LYS_A_71_O | -0.275 | 11.309 | 5.394 | 0.764 | 11.959 | 5.289 | 1.23 | 94.90 | 32.03 |
| 419 | 2jsa-7 | LYS_A_71_C | ASP_A_72_N | -0.275 | 11.309 | 5.394 | -0.665 | 10.691 | 6.499 | 1.32 | 33.48 | -122.25 |
| 420 | 2jsa-7 | ASP_A_72_N | ASP_A_72_H | -0.665 | 10.691 | 6.499 | -1.532 | 10.174 | 6.514 | 1.01 | 89.15 | -149.19 |
| 421 | 2jsa-7 | ASP_A_72_N | ASP_A_72_CA | -0.665 | 10.691 | 6.499 | 0.14 | 10.742 | 7.709 | 1.45 | 33.69 | 3.63 |
| 422 | 2jsa-7 | ASP_A_72_CA | ASP_A_72_HA | 0.14 | 10.742 | 7.709 | 0.855 | 11.561 | 7.628 | 1.09 | 94.26 | 48.88 |
| 423 | 2jsa-7 | ASP_A_72_CA | ASP_A_72_C | 0.14 | 10.742 | 7.709 | 0.885 | 9.416 | 7.875 | 1.53 | 83.77 | -60.67 |
| 424 | 2jsa-7 | ASP_A_72_C | ASP_A_72_O | 0.885 | 9.416 | 7.875 | 2.032 | 9.396 | 8.319 | 1.23 | 68.84 | -1.00 |
| 425 | 2jsa-7 | ASP_A_72_C | VAL_A_73_N | 0.885 | 9.416 | 7.875 | 0.203 | 8.341 | 7.51 | 1.32 | 106.00 | -122.39 |
| 426 | 2jsa-7 | VAL_A_73_N | VAL_A_73_H | 0.203 | 8.341 | 7.51 | -0.737 | 8.432 | 7.153 | 1.01 | 110.71 | 174.47 |
| 427 | 2jsa-7 | VAL_A_73_N | VAL_A_73_CA | 0.203 | 8.341 | 7.51 | 0.787 | 7.013 | 7.614 | 1.45 | 85.90 | -66.26 |
| 428 | 2jsa-7 | VAL_A_73_CA | VAL_A_73_HA | 0.787 | 7.013 | 7.614 | 0.945 | 6.787 | 8.669 | 1.09 | 14.65 | -55.04 |
| 429 | 2jsa-7 | VAL_A_73_CA | VAL_A_73_C | 0.787 | 7.013 | 7.614 | 2.15 | 7.005 | 6.917 | 1.53 | 117.08 | -0.34 |
| 430 | 2jsa-7 | VAL_A_73_C | VAL_A_73_O | 2.15 | 7.005 | 6.917 | 3.158 | 6.642 | 7.523 | 1.23 | 60.51 | -19.80 |
| 431 | 2jsa-7 | VAL_A_73_C | CYS_A_74_N | 2.15 | 7.005 | 6.917 | 2.138 | 7.409 | 5.656 | 1.32 | 162.23 | 91.70 |
| 432 | 2jsa-7 | CYS_A_74_N | CYS_A_74_H | 2.138 | 7.409 | 5.656 | 1.272 | 7.694 | 5.222 | 1.01 | 115.46 | 161.78 |
| 433 | 2jsa-7 | CYS_A_74_N | CYS_A_74_CA | 2.138 | 7.409 | 5.656 | 3.359 | 7.453 | 4.872 | 1.45 | 122.69 | 2.06 |
| 434 | 2jsa-7 | CYS_A_74_CA | CYS_A_74_HA | 3.359 | 7.453 | 4.872 | 3.957 | 6.575 | 5.119 | 1.09 | 76.91 | -55.74 |
| 435 | 2jsa-7 | CYS_A_74_CA | CYS_A_74_C | 3.359 | 7.453 | 4.872 | 4.143 | 8.703 | 5.273 | 1.53 | 74.80 | 57.90 |
| 436 | 2jsa-7 | CYS_A_74_C | CYS_A_74_O | 4.143 | 8.703 | 5.273 | 4.31 | 9.62 | 4.469 | 1.23 | 130.78 | 79.68 |
| 437 | 2jsa-7 | CYS_A_74_C | THR_A_75_N | 4.143 | 8.703 | 5.273 | 4.604 | 8.701 | 6.515 | 1.32 | 20.36 | -0.25 |
| 438 | 2jsa-7 | THR_A_75_N | THR_A_75_H | 4.604 | 8.701 | 6.515 | 4.43 | 7.912 | 7.121 | 1.01 | 53.13 | -102.44 |
| 439 | 2jsa-7 | THR_A_75_N | THR_A_75_CA | 4.604 | 8.701 | 6.515 | 5.367 | 9.824 | 7.033 | 1.45 | 69.12 | 55.81 |
| 440 | 2jsa-7 | THR_A_75_CA | THR_A_75_HA | 5.367 | 9.824 | 7.033 | 6.122 | 10.103 | 6.298 | 1.09 | 132.40 | 20.28 |
| 441 | 2jsa-7 | THR_A_75_CA | THR_A_75_C | 5.367 | 9.824 | 7.033 | 6.089 | 9.429 | 8.322 | 1.53 | 32.56 | -28.68 |
| 442 | 2jsa-7 | THR_A_75_C | THR_A_75_O | 6.089 | 9.429 | 8.322 | 7.318 | 9.465 | 8.386 | 1.23 | 87.02 | 1.68 |
| 443 | 2jsa-7 | THR_A_75_C | LYS_A_76_N | 6.089 | 9.429 | 8.322 | 5.297 | 9.06 | 9.317 | 1.32 | 41.29 | -155.02 |
| 444 | 2jsa-7 | LYS_A_76_N | LYS_A_76_H | 5.297 | 9.06 | 9.317 | 4.295 | 9.054 | 9.194 | 1.01 | 97.00 | -179.66 |
| 445 | 2jsa-7 | LYS_A_76_N | LYS_A_76_CA | 5.297 | 9.06 | 9.317 | 5.846 | 8.658 | 10.601 | 1.45 | 27.92 | -36.21 |
| 446 | 2jsa-7 | LYS_A_76_CA | LYS_A_76_HA | 5.846 | 8.658 | 10.601 | 6.742 | 9.252 | 10.786 | 1.09 | 80.24 | 33.54 |
| 447 | 2jsa-7 | LYS_A_76_CA | LYS_A_76_C | 5.846 | 8.658 | 10.601 | 6.268 | 7.19 | 10.534 | 1.53 | 92.51 | -73.96 |
| 448 | 2jsa-7 | LYS_A_76_C | LYS_A_76_O | 6.268 | 7.19 | 10.534 | 7.039 | 6.722 | 11.37 | 1.23 | 47.17 | -31.26 |
| 449 | 2jsa-7 | LYS_A_76_C | LEU_A_77_N | 6.268 | 7.19 | 10.534 | 5.744 | 6.501 | 9.53 | 1.33 | 139.23 | -127.25 |
| 450 | 2jsa-7 | LEU_A_77_N | LEU_A_77_H | 5.744 | 6.501 | 9.53 | 5.117 | 6.95 | 8.878 | 1.01 | 130.21 | 144.39 |
| 451 | 2jsa-7 | LEU_A_77_N | LEU_A_77_CA | 5.744 | 6.501 | 9.53 | 6.056 | 5.095 | 9.343 | 1.45 | 97.40 | -77.49 |
| 452 | 2jsa-7 | LEU_A_77_CA | LEU_A_77_HA | 6.056 | 5.095 | 9.343 | 6.188 | 4.645 | 10.327 | 1.09 | 25.48 | -73.65 |
| 453 | 2jsa-7 | LEU_A_77_CA | LEU_A_77_C | 6.056 | 5.095 | 9.343 | 7.383 | 4.968 | 8.591 | 1.53 | 119.43 | -5.47 |
| 454 | 2jsa-7 | LEU_A_77_C | LEU_A_77_O | 7.383 | 4.968 | 8.591 | 7.898 | 3.865 | 8.417 | 1.23 | 98.13 | -64.97 |
| 455 | 2jsa-7 | LEU_A_77_C | ASN_A_78_N | 7.383 | 4.968 | 8.591 | 7.899 | 6.112 | 8.167 | 1.32 | 108.67 | 65.72 |
| 456 | 2jsa-7 | ASN_A_78_N | ASN_A_78_H | 7.899 | 6.112 | 8.167 | 7.421 | 6.983 | 8.347 | 1.01 | 79.73 | 118.76 |
| 457 | 2jsa-7 | ASN_A_78_N | ASN_A_78_CA | 7.899 | 6.112 | 8.167 | 9.156 | 6.142 | 7.44 | 1.45 | 120.04 | 1.37 |
| 458 | 2jsa-7 | ASN_A_78_CA | ASN_A_78_HA | 9.156 | 6.142 | 7.44 | 9.478 | 7.177 | 7.327 | 1.09 | 95.95 | 72.72 |
| 459 | 2jsa-7 | ASN_A_78_CA | ASN_A_78_C | 9.156 | 6.142 | 7.44 | 8.958 | 5.506 | 6.062 | 1.53 | 154.20 | -107.29 |
| 460 | 2jsa-7 | ASN_A_78_C | ASN_A_78_O | 8.958 | 5.506 | 6.062 | 9.892 | 4.944 | 5.494 | 1.23 | 117.52 | -31.04 |
| 461 | 2jsa-7 | ASN_A_78_C | GLU_A_79_N | 8.958 | 5.506 | 6.062 | 7.734 | 5.616 | 5.565 | 1.33 | 112.02 | 174.86 |
| 462 | 2jsa-7 | GLU_A_79_N | GLU_A_79_H | 7.734 | 5.616 | 5.565 | 7.014 | 6.094 | 6.088 | 1.01 | 58.82 | 146.42 |
| 463 | 2jsa-7 | GLU_A_79_N | GLU_A_79_CA | 7.734 | 5.616 | 5.565 | 7.402 | 5.057 | 4.266 | 1.45 | 153.41 | -120.71 |
| 464 | 2jsa-7 | GLU_A_79_CA | GLU_A_79_HA | 7.402 | 5.057 | 4.266 | 8.315 | 4.65 | 3.831 | 1.09 | 113.52 | -24.03 |
| 465 | 2jsa-7 | GLU_A_79_CA | GLU_A_79_C | 7.402 | 5.057 | 4.266 | 6.872 | 6.152 | 3.337 | 1.53 | 127.37 | 115.83 |
| 466 | 2jsa-7 | GLU_A_79_C | GLU_A_79_O | 6.872 | 6.152 | 3.337 | 5.788 | 6.021 | 2.772 | 1.23 | 117.36 | -173.11 |
| 467 | 2jsa-7 | GLU_A_79_C | CYS_A_80_N | 6.872 | 6.152 | 3.337 | 7.662 | 7.209 | 3.21 | 1.33 | 95.50 | 53.23 |
| 468 | 2jsa-7 | CYS_A_80_N | CYS_A_80_H | 7.662 | 7.209 | 3.21 | 8.541 | 7.25 | 3.706 | 1.01 | 60.59 | 2.67 |
| 469 | 2jsa-7 | CYS_A_80_N | CYS_A_80_CA | 7.662 | 7.209 | 3.21 | 7.286 | 8.326 | 2.36 | 1.45 | 125.80 | 108.60 |
| 470 | 2jsa-7 | CYS_A_80_CA | CYS_A_80_HA | 7.286 | 8.326 | 2.36 | 7.441 | 8.034 | 1.321 | 1.09 | 162.35 | -62.04 |
| 471 | 2jsa-7 | CYS_A_80_CA | CYS_A_80_C | 7.286 | 8.326 | 2.36 | 8.211 | 9.503 | 2.678 | 1.53 | 78.01 | 51.84 |
| 472 | 2jsa-7 | CYS_A_80_C | CYS_A_80_O | 8.211 | 9.503 | 2.678 | 8.769 | 9.581 | 3.771 | 1.23 | 27.27 | 7.96 |
| 473 | 2jsa-7 | CYS_A_80_C | PRO_A_81_N | 8.211 | 9.503 | 2.678 | 8.347 | 10.413 | 1.677 | 1.36 | 137.41 | 81.50 |
| 474 | 2jsa-7 | PRO_A_81_N | PRO_A_81_CA | 8.347 | 10.413 | 1.677 | 9.194 | 11.583 | 1.839 | 1.45 | 83.60 | 54.10 |
| 475 | 2jsa-7 | PRO_A_81_CA | PRO_A_81_HA | 9.194 | 11.583 | 1.839 | 10.145 | 11.296 | 2.289 | 1.09 | 65.63 | -16.79 |
| 476 | 2jsa-7 | PRO_A_81_CA | PRO_A_81_C | 9.194 | 11.583 | 1.839 | 8.527 | 12.623 | 2.74 | 1.53 | 53.90 | 122.67 |
| 477 | 2jsa-7 | PRO_A_81_C | PRO_A_81_O | 8.527 | 12.623 | 2.74 | 7.968 | 13.604 | 2.253 | 1.23 | 113.33 | 119.68 |
| 1 | 2jsa-8 | ARG_A_1_N | ARG_A_1_H | 15.458 | 8.607 | -0.117 | 14.716 | 8.902 | -0.735 | 1.01 | 127.74 | 158.32 |
| 2 | 2jsa-8 | ARG_A_1_N | ARG_A_1_CA | 15.458 | 8.607 | -0.117 | 15.288 | 7.375 | 0.635 | 1.45 | 58.84 | -97.86 |
| 3 | 2jsa-8 | ARG_A_1_CA | ARG_A_1_HA | 15.288 | 7.375 | 0.635 | 16.084 | 6.686 | 0.351 | 1.09 | 105.10 | -40.88 |
| 4 | 2jsa-8 | ARG_A_1_CA | ARG_A_1_C | 15.288 | 7.375 | 0.635 | 13.936 | 6.737 | 0.31 | 1.53 | 102.26 | -154.74 |
| 5 | 2jsa-8 | ARG_A_1_C | ARG_A_1_O | 13.936 | 6.737 | 0.31 | 12.941 | 7.005 | 0.982 | 1.23 | 56.89 | 164.93 |
| 6 | 2jsa-8 | ARG_A_1_C | SER_A_2_N | 13.936 | 6.737 | 0.31 | 13.944 | 5.905 | -0.721 | 1.32 | 141.10 | -89.45 |
| 7 | 2jsa-8 | SER_A_2_N | SER_A_2_H | 13.944 | 5.905 | -0.721 | 14.8 | 5.731 | -1.228 | 1.01 | 120.13 | -11.49 |
| 8 | 2jsa-8 | SER_A_2_N | SER_A_2_CA | 13.944 | 5.905 | -0.721 | 12.73 | 5.227 | -1.143 | 1.45 | 106.88 | -150.82 |
| 9 | 2jsa-8 | SER_A_2_CA | SER_A_2_HA | 12.73 | 5.227 | -1.143 | 13.006 | 4.251 | -1.541 | 1.09 | 111.42 | -74.21 |
| 10 | 2jsa-8 | SER_A_2_CA | SER_A_2_C | 12.73 | 5.227 | -1.143 | 12.039 | 6.029 | -2.247 | 1.53 | 136.20 | 130.75 |
| 11 | 2jsa-8 | SER_A_2_C | SER_A_2_O | 12.039 | 6.029 | -2.247 | 12.687 | 6.789 | -2.966 | 1.23 | 125.75 | 49.55 |
| 12 | 2jsa-8 | SER_A_2_C | ALA_A_3_N | 12.039 | 6.029 | -2.247 | 10.732 | 5.833 | -2.349 | 1.33 | 94.41 | -171.47 |
| 13 | 2jsa-8 | ALA_A_3_N | ALA_A_3_H | 10.732 | 5.833 | -2.349 | 10.265 | 5.191 | -1.725 | 1.01 | 51.83 | -126.03 |
| 14 | 2jsa-8 | ALA_A_3_N | ALA_A_3_CA | 10.732 | 5.833 | -2.349 | 9.946 | 6.529 | -3.353 | 1.45 | 133.72 | 138.48 |
| 15 | 2jsa-8 | ALA_A_3_CA | ALA_A_3_HA | 9.946 | 6.529 | -3.353 | 9.704 | 7.524 | -2.98 | 1.09 | 69.99 | 103.67 |
| 16 | 2jsa-8 | ALA_A_3_CA | ALA_A_3_C | 9.946 | 6.529 | -3.353 | 8.64 | 5.77 | -3.589 | 1.53 | 98.88 | -149.84 |
| 17 | 2jsa-8 | ALA_A_3_C | ALA_A_3_O | 8.64 | 5.77 | -3.589 | 7.561 | 6.273 | -3.282 | 1.23 | 75.54 | 155.01 |
| 18 | 2jsa-8 | ALA_A_3_C | LEU_A_4_N | 8.64 | 5.77 | -3.589 | 8.78 | 4.569 | -4.131 | 1.33 | 114.14 | -83.35 |
| 19 | 2jsa-8 | LEU_A_4_N | LEU_A_4_H | 8.78 | 4.569 | -4.131 | 9.7 | 4.217 | -4.356 | 1.01 | 102.87 | -20.94 |
| 20 | 2jsa-8 | LEU_A_4_N | LEU_A_4_CA | 8.78 | 4.569 | -4.131 | 7.624 | 3.735 | -4.413 | 1.45 | 101.19 | -144.19 |

| | | | | | | | | | | | |
|---|---|---|---|---|---|---|---|---|---|---|
| 21 | 2jsa-8 | LEU_A_4_CA | LEU_A_4_HA | 7.624 | 3.735 | -4.413 | 6.914 | 4.326 | -4.99 | 1.09 | 121.99 | 140.23 |
| 22 | 2jsa-8 | LEU_A_4_CA | LEU_A_4_C | 7.624 | 3.735 | -4.413 | 6.948 | 3.349 | -3.096 | 1.53 | 30.59 | -150.27 |
| 23 | 2jsa-8 | LEU_A_4_C | LEU_A_4_O | 6.948 | 3.349 | -3.096 | 5.761 | 3.026 | -3.074 | 1.23 | 88.98 | -164.78 |
| 24 | 2jsa-8 | LEU_A_4_C | SER_A_5_N | 6.948 | 3.349 | -3.096 | 7.732 | 3.395 | -2.029 | 1.32 | 36.35 | 3.36 |
| 25 | 2jsa-8 | SER_A_5_N | SER_A_5_H | 7.732 | 3.395 | -2.029 | 8.7 | 3.67 | -2.12 | 1.01 | 95.17 | 15.86 |
| 26 | 2jsa-8 | SER_A_5_N | SER_A_5_CA | 7.732 | 3.395 | -2.029 | 7.223 | 3.054 | -0.71 | 1.45 | 24.91 | -146.18 |
| 27 | 2jsa-8 | SER_A_5_CA | SER_A_5_HA | 7.223 | 3.054 | -0.71 | 6.881 | 2.019 | -0.733 | 1.09 | 91.21 | -108.29 |
| 28 | 2jsa-8 | SER_A_5_CA | SER_A_5_C | 7.223 | 3.054 | -0.71 | 6.038 | 3.954 | -0.358 | 1.53 | 76.69 | 142.78 |
| 29 | 2jsa-8 | SER_A_5_C | SER_A_5_O | 6.038 | 3.954 | -0.358 | 5.099 | 3.518 | 0.306 | 1.23 | 57.33 | -155.09 |
| 30 | 2jsa-8 | SER_A_5_C | CYS_A_6_N | 6.038 | 3.954 | -0.358 | 6.121 | 5.194 | -0.818 | 1.33 | 110.31 | 86.17 |
| 31 | 2jsa-8 | CYS_A_6_N | CYS_A_6_H | 6.121 | 5.194 | -0.818 | 6.924 | 5.484 | -1.358 | 1.01 | 122.31 | 19.86 |
| 32 | 2jsa-8 | CYS_A_6_N | CYS_A_6_CA | 6.121 | 5.194 | -0.818 | 5.066 | 6.16 | -0.559 | 1.45 | 79.74 | 137.52 |
| 33 | 2jsa-8 | CYS_A_6_CA | CYS_A_6_HA | 5.066 | 6.16 | -0.559 | 4.758 | 6.061 | 0.482 | 1.09 | 17.26 | -162.18 |
| 34 | 2jsa-8 | CYS_A_6_CA | CYS_A_6_C | 5.066 | 6.16 | -0.559 | 3.873 | 5.82 | -1.455 | 1.53 | 125.84 | -164.09 |
| 35 | 2jsa-8 | CYS_A_6_C | CYS_A_6_O | 3.873 | 5.82 | -1.455 | 2.819 | 5.417 | -0.964 | 1.23 | 66.49 | -159.08 |
| 36 | 2jsa-8 | CYS_A_6_C | GLN_A_7_N | 3.873 | 5.82 | -1.455 | 4.078 | 5.994 | -2.752 | 1.32 | 168.29 | 40.32 |
| 37 | 2jsa-8 | GLN_A_7_N | GLN_A_7_H | 4.078 | 5.994 | -2.752 | 4.971 | 6.33 | -3.085 | 1.01 | 109.24 | 20.62 |
| 38 | 2jsa-8 | GLN_A_7_N | GLN_A_7_CA | 4.078 | 5.994 | -2.752 | 3.033 | 5.71 | -3.719 | 1.45 | 131.76 | -164.80 |
| 39 | 2jsa-8 | GLN_A_7_CA | GLN_A_7_HA | 3.033 | 5.71 | -3.719 | 2.334 | 6.546 | -3.714 | 1.09 | 89.74 | 129.90 |
| 40 | 2jsa-8 | GLN_A_7_CA | GLN_A_7_C | 3.033 | 5.71 | -3.719 | 2.27 | 4.444 | -3.322 | 1.53 | 74.97 | -121.08 |
| 41 | 2jsa-8 | GLN_A_7_C | GLN_A_7_O | 2.27 | 4.444 | -3.322 | 1.054 | 4.478 | -3.146 | 1.23 | 81.77 | 178.40 |
| 42 | 2jsa-8 | GLN_A_7_C | MET_A_8_N | 2.27 | 4.444 | -3.322 | 3.018 | 3.358 | -3.194 | 1.32 | 84.46 | -55.44 |
| 43 | 2jsa-8 | MET_A_8_N | MET_A_8_H | 3.018 | 3.358 | -3.194 | 4.014 | 3.404 | -3.357 | 1.01 | 99.28 | 2.64 |
| 44 | 2jsa-8 | MET_A_8_N | MET_A_8_CA | 3.018 | 3.358 | -3.194 | 2.428 | 2.084 | -2.82 | 1.45 | 75.08 | -114.85 |
| 45 | 2jsa-8 | MET_A_8_CA | MET_A_8_HA | 2.428 | 2.084 | -2.82 | 1.805 | 1.733 | -3.642 | 1.09 | 138.98 | -150.60 |
| 46 | 2jsa-8 | MET_A_8_CA | MET_A_8_C | 2.428 | 2.084 | -2.82 | 1.563 | 2.226 | -1.567 | 1.53 | 34.98 | 170.68 |
| 47 | 2jsa-8 | MET_A_8_C | MET_A_8_O | 1.563 | 2.226 | -1.567 | 0.5 | 1.614 | -1.469 | 1.23 | 85.43 | -150.07 |
| 48 | 2jsa-8 | MET_A_8_C | CYS_A_9_N | 1.563 | 2.226 | -1.567 | 2.05 | 3.037 | -0.64 | 1.32 | 45.58 | 59.02 |
| 49 | 2jsa-8 | CYS_A_9_N | CYS_A_9_H | 2.05 | 3.037 | -0.64 | 2.931 | 3.509 | -0.789 | 1.01 | 98.48 | 28.18 |
| 50 | 2jsa-8 | CYS_A_9_N | CYS_A_9_CA | 2.05 | 3.037 | -0.64 | 1.335 | 3.267 | 0.604 | 1.45 | 31.12 | 162.17 |
| 51 | 2jsa-8 | CYS_A_9_CA | CYS_A_9_HA | 1.335 | 3.267 | 0.604 | 1.086 | 2.3 | 1.041 | 1.09 | 66.36 | -104.44 |
| 52 | 2jsa-8 | CYS_A_9_CA | CYS_A_9_C | 1.335 | 3.267 | 0.604 | 0.034 | 4.006 | 0.28 | 1.53 | 102.22 | 150.40 |
| 53 | 2jsa-8 | CYS_A_9_C | CYS_A_9_O | 0.034 | 4.006 | 0.28 | -0.998 | 3.753 | 0.899 | 1.23 | 59.78 | -166.23 |
| 54 | 2jsa-8 | CYS_A_9_C | GLU_A_10_N | 0.034 | 4.006 | 0.28 | 0.127 | 4.904 | -0.69 | 1.33 | 137.05 | 84.09 |
| 55 | 2jsa-8 | GLU_A_10_N | GLU_A_10_H | 0.127 | 4.904 | -0.69 | 1.01 | 5.06 | -1.157 | 1.01 | 117.51 | 10.02 |
| 56 | 2jsa-8 | GLU_A_10_N | GLU_A_10_CA | 0.127 | 4.904 | -0.69 | -1.03 | 5.68 | -1.103 | 1.45 | 106.51 | 146.15 |
| 57 | 2jsa-8 | GLU_A_10_CA | GLU_A_10_HA | -1.03 | 5.68 | -1.103 | -1.55 | 6.013 | -0.205 | 1.09 | 34.51 | 147.37 |
| 58 | 2jsa-8 | GLU_A_10_CA | GLU_A_10_C | -1.03 | 5.68 | -1.103 | -1.988 | 4.811 | -1.919 | 1.53 | 122.25 | -137.79 |
| 59 | 2jsa-8 | GLU_A_10_C | GLU_A_10_O | -1.988 | 4.811 | -1.919 | -3.073 | 5.257 | -2.291 | 1.23 | 107.59 | 157.65 |
| 60 | 2jsa-8 | GLU_A_10_C | LEU_A_11_N | -1.988 | 4.811 | -1.919 | -1.553 | 3.586 | -2.175 | 1.32 | 101.14 | -70.45 |
| 61 | 2jsa-8 | LEU_A_11_N | LEU_A_11_H | -1.553 | 3.586 | -2.175 | -0.649 | 3.286 | -1.837 | 1.01 | 70.46 | -18.36 |
| 62 | 2jsa-8 | LEU_A_11_N | LEU_A_11_CA | -1.553 | 3.586 | -2.175 | -2.359 | 2.65 | -2.942 | 1.45 | 121.84 | -130.73 |
| 63 | 2jsa-8 | LEU_A_11_CA | LEU_A_11_HA | -2.359 | 2.65 | -2.942 | -3.047 | 3.226 | -3.561 | 1.09 | 124.60 | 140.06 |
| 64 | 2jsa-8 | LEU_A_11_CA | LEU_A_11_C | -2.359 | 2.65 | -2.942 | -3.195 | 1.8 | -1.983 | 1.53 | 51.19 | -134.52 |
| 65 | 2jsa-8 | LEU_A_11_C | LEU_A_11_O | -3.195 | 1.8 | -1.983 | -4.4 | 1.642 | -2.177 | 1.23 | 99.07 | -172.53 |
| 66 | 2jsa-8 | LEU_A_11_C | VAL_A_12_N | -3.195 | 1.8 | -1.983 | -2.523 | 1.274 | -0.969 | 1.33 | 40.08 | -38.05 |
| 67 | 2jsa-8 | VAL_A_12_N | VAL_A_12_H | -2.523 | 1.274 | -0.969 | -1.532 | 1.446 | -0.872 | 1.01 | 84.49 | 9.85 |
| 68 | 2jsa-8 | VAL_A_12_N | VAL_A_12_CA | -2.523 | 1.274 | -0.969 | -3.189 | 0.444 | 0.02 | 1.45 | 47.10 | -128.74 |
| 69 | 2jsa-8 | VAL_A_12_CA | VAL_A_12_HA | -3.189 | 0.444 | 0.02 | -3.718 | -0.354 | -0.5 | 1.09 | 118.51 | -123.54 |
| 70 | 2jsa-8 | VAL_A_12_CA | VAL_A_12_C | -3.189 | 0.444 | 0.02 | -4.221 | 1.285 | 0.774 | 1.53 | 60.47 | 140.82 |
| 71 | 2jsa-8 | VAL_A_12_C | VAL_A_12_O | -4.221 | 1.285 | 0.774 | -5.347 | 0.843 | 0.99 | 1.23 | 79.88 | -158.57 |
| 72 | 2jsa-8 | VAL_A_12_C | VAL_A_13_N | -4.221 | 1.285 | 0.774 | -3.797 | 2.482 | 1.154 | 1.33 | 73.34 | 70.49 |
| 73 | 2jsa-8 | VAL_A_13_N | VAL_A_13_H | -3.797 | 2.482 | 1.154 | -2.855 | 2.779 | 0.942 | 1.01 | 102.11 | 17.50 |
| 74 | 2jsa-8 | VAL_A_13_N | VAL_A_13_CA | -3.797 | 2.482 | 1.154 | -4.67 | 3.389 | 1.88 | 1.45 | 60.03 | 133.91 |
| 75 | 2jsa-8 | VAL_A_13_CA | VAL_A_13_HA | -4.67 | 3.389 | 1.88 | -5.064 | 2.864 | 2.75 | 1.09 | 37.03 | -126.89 |
| 76 | 2jsa-8 | VAL_A_13_CA | VAL_A_13_C | -4.67 | 3.389 | 1.88 | -5.848 | 3.779 | 0.986 | 1.53 | 125.77 | 161.68 |
| 77 | 2jsa-8 | VAL_A_13_C | VAL_A_13_O | -5.848 | 3.779 | 0.986 | -7.005 | 3.553 | 1.342 | 1.23 | 73.20 | -168.95 |
| 78 | 2jsa-8 | VAL_A_13_C | LYS_A_14_N | -5.848 | 3.779 | 0.986 | -5.516 | 4.359 | -0.158 | 1.32 | 149.71 | 60.21 |
| 79 | 2jsa-8 | LYS_A_14_N | LYS_A_14_H | -5.516 | 4.359 | -0.158 | -4.544 | 4.515 | -0.386 | 1.01 | 103.04 | 9.12 |
| 80 | 2jsa-8 | LYS_A_14_N | LYS_A_14_CA | -5.516 | 4.359 | -0.158 | -6.533 | 4.782 | -1.105 | 1.45 | 130.69 | 157.42 |
| 81 | 2jsa-8 | LYS_A_14_CA | LYS_A_14_HA | -6.533 | 4.782 | -1.105 | -7.025 | 5.666 | -0.699 | 1.09 | 68.13 | 119.10 |
| 82 | 2jsa-8 | LYS_A_14_CA | LYS_A_14_C | -6.533 | 4.782 | -1.105 | -7.589 | 3.683 | -1.24 | 1.53 | 95.06 | -133.86 |
| 83 | 2jsa-8 | LYS_A_14_C | LYS_A_14_O | -7.589 | 3.683 | -1.24 | -8.764 | 3.971 | -1.462 | 1.23 | 100.40 | 166.23 |
| 84 | 2jsa-8 | LYS_A_14_C | LYS_A_15_N | -7.589 | 3.683 | -1.24 | -7.132 | 2.448 | -1.098 | 1.32 | 83.85 | -69.69 |
| 85 | 2jsa-8 | LYS_A_15_N | LYS_A_15_H | -7.132 | 2.448 | -1.098 | -6.152 | 2.286 | -0.914 | 1.01 | 79.51 | -9.39 |
| 86 | 2jsa-8 | LYS_A_15_N | LYS_A_15_CA | -7.132 | 2.448 | -1.098 | -8.022 | 1.305 | -1.202 | 1.45 | 94.11 | -127.91 |
| 87 | 2jsa-8 | LYS_A_15_CA | LYS_A_15_HA | -8.022 | 1.305 | -1.202 | -8.771 | 1.529 | -1.961 | 1.09 | 134.15 | 163.35 |
| 88 | 2jsa-8 | LYS_A_15_CA | LYS_A_15_C | -8.022 | 1.305 | -1.202 | -8.752 | 1.111 | 0.129 | 1.53 | 29.57 | -165.12 |
| 89 | 2jsa-8 | LYS_A_15_C | LYS_A_15_O | -8.752 | 1.111 | 0.129 | -9.942 | 0.799 | 0.149 | 1.23 | 89.07 | -165.31 |
| 90 | 2jsa-8 | LYS_A_15_C | TYR_A_16_N | -8.752 | 1.111 | 0.129 | -8.009 | 1.304 | 1.209 | 1.33 | 35.40 | 14.56 |
| 91 | 2jsa-8 | TYR_A_16_N | TYR_A_16_H | -8.009 | 1.304 | 1.209 | -7.035 | 1.558 | 1.118 | 1.01 | 95.17 | 14.62 |
| 92 | 2jsa-8 | TYR_A_16_N | TYR_A_16_CA | -8.009 | 1.304 | 1.209 | -8.571 | 1.155 | 2.54 | 1.45 | 23.60 | -165.15 |
| 93 | 2jsa-8 | TYR_A_16_CA | TYR_A_16_HA | -8.571 | 1.155 | 2.54 | -9.105 | 0.208 | 2.62 | 1.09 | 85.79 | -119.42 |
| 94 | 2jsa-8 | TYR_A_16_CA | TYR_A_16_C | -8.571 | 1.155 | 2.54 | -9.514 | 2.313 | 2.871 | 1.53 | 77.50 | 129.16 |
| 95 | 2jsa-8 | TYR_A_16_C | TYR_A_16_O | -9.514 | 2.313 | 2.871 | -10.72 | 2.115 | 3.014 | 1.23 | 83.33 | -170.68 |
| 96 | 2jsa-8 | TYR_A_16_C | GLU_A_17_N | -9.514 | 2.313 | 2.871 | -8.93 | 3.497 | 2.98 | 1.32 | 85.28 | 63.75 |
| 97 | 2jsa-8 | GLU_A_17_N | GLU_A_17_H | -8.93 | 3.497 | 2.98 | -7.932 | 3.586 | 2.847 | 1.01 | 97.56 | 5.10 |
| 98 | 2jsa-8 | GLU_A_17_N | GLU_A_17_CA | -8.93 | 3.497 | 2.98 | -9.704 | 4.688 | 3.291 | 1.45 | 77.65 | 123.02 |
| 99 | 2jsa-8 | GLU_A_17_CA | GLU_A_17_HA | -9.704 | 4.688 | 3.291 | -10.01 | 4.628 | 4.336 | 1.09 | 16.62 | -168.91 |
| 100 | 2jsa-8 | GLU_A_17_CA | GLU_A_17_C | -9.704 | 4.688 | 3.291 | -10.963 | 4.743 | 2.424 | 1.53 | 124.53 | 177.50 |
| 101 | 2jsa-8 | GLU_A_17_C | GLU_A_17_O | -10.963 | 4.743 | 2.424 | -12.026 | 5.148 | 2.892 | 1.23 | 67.64 | 159.14 |
| 102 | 2jsa-8 | GLU_A_17_C | GLY_A_18_N | -10.963 | 4.743 | 2.424 | -10.802 | 4.332 | 1.175 | 1.32 | 160.54 | -68.61 |
| 103 | 2jsa-8 | GLY_A_18_N | GLY_A_18_H | -10.802 | 4.332 | 1.175 | -9.897 | 4.013 | 0.858 | 1.01 | 108.28 | -19.42 |
| 104 | 2jsa-8 | GLY_A_18_N | GLY_A_18_CA | -10.802 | 4.332 | 1.175 | -11.912 | 4.33 | 0.238 | 1.45 | 130.17 | -179.90 |
| 105 | 2jsa-8 | GLY_A_18_CA | GLY_A_18_C | -11.912 | 4.33 | 0.238 | -12.277 | 2.903 | -0.178 | 1.53 | 105.77 | -104.35 |
| 106 | 2jsa-8 | GLY_A_18_C | GLY_A_18_O | -12.277 | 2.903 | -0.178 | -12.394 | 2.61 | -1.366 | 1.23 | 165.13 | -111.77 |
| 107 | 2jsa-8 | GLY_A_18_C | SER_A_19_N | -12.277 | 2.903 | -0.178 | -12.445 | 2.054 | 0.826 | 1.33 | 40.76 | -101.19 |
| 108 | 2jsa-8 | SER_A_19_N | SER_A_19_H | -12.445 | 2.054 | 0.826 | -12.332 | 2.365 | 1.781 | 1.01 | 19.11 | 70.03 |
| 109 | 2jsa-8 | SER_A_19_N | SER_A_19_CA | -12.445 | 2.054 | 0.826 | -12.794 | 0.664 | 0.578 | 1.45 | 99.82 | -104.09 |
| 110 | 2jsa-8 | SER_A_19_CA | SER_A_19_HA | -12.794 | 0.664 | 0.578 | -12.62 | 0.45 | -0.476 | 1.09 | 165.34 | -50.89 |
| 111 | 2jsa-8 | SER_A_19_CA | SER_A_19_C | -12.794 | 0.664 | 0.578 | -14.275 | 0.434 | 0.888 | 1.53 | 78.31 | -171.17 |
| 112 | 2jsa-8 | SER_A_19_C | SER_A_19_O | -14.275 | 0.434 | 0.888 | -14.974 | -0.236 | 0.13 | 1.23 | 128.06 | -136.21 |
| 113 | 2jsa-8 | SER_A_19_C | ALA_A_20_N | -14.275 | 0.434 | 0.888 | -14.708 | 1.001 | 2.005 | 1.33 | 32.57 | 127.37 |
| 114 | 2jsa-8 | ALA_A_20_N | ALA_A_20_H | -14.708 | 1.001 | 2.005 | -14.075 | 1.536 | 2.583 | 1.01 | 55.11 | 40.20 |
| 115 | 2jsa-8 | ALA_A_20_N | ALA_A_20_CA | -14.708 | 1.001 | 2.005 | -16.093 | 0.866 | 2.423 | 1.45 | 73.28 | -174.43 |
| 116 | 2jsa-8 | ALA_A_20_CA | ALA_A_20_HA | -16.093 | 0.866 | 2.423 | -16.248 | 1.467 | 3.319 | 1.09 | 34.71 | 104.46 |

| | | | | | | | | | | | |
|---|---|---|---|---|---|---|---|---|---|---|---|
| 117 | 2jsa-8 | ALA_A_20_CA | ALA_A_20_C | -16.093 | 0.866 | 2.423 | -16.376 | -0.598 | 2.767 | 1.53 | 77.01 | -100.94 |
| 118 | 2jsa-8 | ALA_A_20_C | ALA_A_20_O | -16.376 | -0.598 | 2.767 | -17.528 | -0.982 | 2.963 | 1.23 | 80.83 | -161.57 |
| 119 | 2jsa-8 | ALA_A_20_C | ASP_A_21_N | -16.376 | -0.598 | 2.767 | -15.305 | -1.375 | 2.83 | 1.32 | 87.27 | -35.96 |
| 120 | 2jsa-8 | ASP_A_21_N | ASP_A_21_H | -15.305 | -1.375 | 2.83 | -14.386 | -0.992 | 2.657 | 1.01 | 99.86 | 22.62 |
| 121 | 2jsa-8 | ASP_A_21_N | ASP_A_21_CA | -15.305 | -1.375 | 2.83 | -15.425 | -2.788 | 3.147 | 1.45 | 77.40 | -94.85 |
| 122 | 2jsa-8 | ASP_A_21_CA | ASP_A_21_HA | -15.425 | -2.788 | 3.147 | -16.477 | -3.033 | 3.292 | 1.09 | 82.35 | -166.89 |
| 123 | 2jsa-8 | ASP_A_21_CA | ASP_A_21_C | -15.425 | -2.788 | 3.147 | -14.639 | -3.088 | 4.426 | 1.53 | 33.34 | -20.89 |
| 124 | 2jsa-8 | ASP_A_21_C | ASP_A_21_O | -14.639 | -3.088 | 4.426 | -14.757 | -4.175 | 4.99 | 1.23 | 62.71 | -96.20 |
| 125 | 2jsa-8 | ASP_A_21_C | LYS_A_22_N | -14.639 | -3.088 | 4.426 | -13.856 | -2.105 | 4.845 | 1.32 | 71.56 | 51.46 |
| 126 | 2jsa-8 | LYS_A_22_N | LYS_A_22_H | -13.856 | -2.105 | 4.845 | -13.808 | -1.238 | 4.33 | 1.01 | 120.67 | 86.83 |
| 127 | 2jsa-8 | LYS_A_22_N | LYS_A_22_CA | -13.856 | -2.105 | 4.845 | -13.053 | -2.25 | 6.047 | 1.45 | 34.17 | -10.24 |
| 128 | 2jsa-8 | LYS_A_22_CA | LYS_A_22_HA | -13.053 | -2.25 | 6.047 | -12.244 | -1.52 | 6.003 | 1.09 | 92.31 | 42.06 |
| 129 | 2jsa-8 | LYS_A_22_CA | LYS_A_22_C | -13.053 | -2.25 | 6.047 | -12.42 | -3.643 | 6.067 | 1.53 | 89.25 | -65.56 |
| 130 | 2jsa-8 | LYS_A_22_C | LYS_A_22_O | -12.42 | -3.643 | 6.067 | -12.387 | -4.299 | 7.108 | 1.23 | 32.25 | -87.12 |
| 131 | 2jsa-8 | LYS_A_22_C | ASP_A_23_N | -12.42 | -3.643 | 6.067 | -11.933 | -4.054 | 4.906 | 1.32 | 151.24 | -40.16 |
| 132 | 2jsa-8 | ASP_A_23_N | ASP_A_23_H | -11.933 | -4.054 | 4.906 | -11.996 | -3.462 | 4.09 | 1.01 | 143.89 | 96.07 |
| 133 | 2jsa-8 | ASP_A_23_N | ASP_A_23_CA | -11.933 | -4.054 | 4.906 | -11.302 | -5.356 | 4.777 | 1.45 | 95.09 | -64.14 |
| 134 | 2jsa-8 | ASP_A_23_CA | ASP_A_23_HA | -11.302 | -5.356 | 4.777 | -11.294 | -5.845 | 5.751 | 1.09 | 26.66 | -89.06 |
| 135 | 2jsa-8 | ASP_A_23_CA | ASP_A_23_C | -11.302 | -5.356 | 4.777 | -9.869 | -5.178 | 4.274 | 1.53 | 109.21 | 7.08 |
| 136 | 2jsa-8 | ASP_A_23_C | ASP_A_23_O | -9.869 | -5.178 | 4.274 | -9.654 | -4.71 | 3.156 | 1.23 | 155.27 | 65.33 |
| 137 | 2jsa-8 | ASP_A_23_C | ALA_A_24_N | -9.869 | -5.178 | 4.274 | -8.926 | -5.56 | 5.122 | 1.32 | 50.19 | -22.05 |
| 138 | 2jsa-8 | ALA_A_24_N | ALA_A_24_H | -8.926 | -5.56 | 5.122 | -9.176 | -5.936 | 6.025 | 1.01 | 26.57 | -123.62 |
| 139 | 2jsa-8 | ALA_A_24_N | ALA_A_24_CA | -8.926 | -5.56 | 5.122 | -7.519 | -5.447 | 4.777 | 1.45 | 103.73 | 4.59 |
| 140 | 2jsa-8 | ALA_A_24_CA | ALA_A_24_HA | -7.519 | -5.447 | 4.777 | -7.345 | -4.469 | 4.327 | 1.09 | 114.37 | 79.91 |
| 141 | 2jsa-8 | ALA_A_24_CA | ALA_A_24_C | -7.519 | -5.447 | 4.777 | -7.163 | -6.523 | 3.749 | 1.53 | 132.21 | -71.69 |
| 142 | 2jsa-8 | ALA_A_24_C | ALA_A_24_O | -7.163 | -6.523 | 3.749 | -6.032 | -6.577 | 3.268 | 1.23 | 113.02 | -2.73 |
| 143 | 2jsa-8 | ALA_A_24_C | ASN_A_25_N | -7.163 | -6.523 | 3.749 | -8.149 | -7.353 | 3.443 | 1.32 | 103.36 | -139.91 |
| 144 | 2jsa-8 | ASN_A_25_N | ASN_A_25_H | -8.149 | -7.353 | 3.443 | -9.054 | -7.249 | 3.878 | 1.01 | 64.47 | 173.44 |
| 145 | 2jsa-8 | ASN_A_25_N | ASN_A_25_CA | -8.149 | -7.353 | 3.443 | -7.954 | -8.425 | 2.481 | 1.45 | 131.44 | -79.69 |
| 146 | 2jsa-8 | ASN_A_25_CA | ASN_A_25_HA | -7.954 | -8.425 | 2.481 | -6.917 | -8.76 | 2.526 | 1.09 | 87.64 | -17.90 |
| 147 | 2jsa-8 | ASN_A_25_CA | ASN_A_25_C | -7.954 | -8.425 | 2.481 | -8.272 | -7.908 | 1.077 | 1.53 | 156.62 | 121.60 |
| 148 | 2jsa-8 | ASN_A_25_C | ASN_A_25_O | -8.272 | -7.908 | 1.077 | -7.513 | -8.143 | 0.139 | 1.23 | 139.73 | -17.20 |
| 149 | 2jsa-8 | ASN_A_25_C | VAL_A_26_N | -8.272 | -7.908 | 1.077 | -9.396 | -7.213 | 0.978 | 1.33 | 94.28 | 148.27 |
| 150 | 2jsa-8 | VAL_A_26_N | VAL_A_26_H | -9.396 | -7.213 | 0.978 | -9.972 | -7.058 | 1.793 | 1.01 | 36.20 | 164.94 |
| 151 | 2jsa-8 | VAL_A_26_N | VAL_A_26_CA | -9.396 | -7.213 | 0.978 | -9.824 | -6.661 | -0.296 | 1.45 | 151.27 | 127.79 |
| 152 | 2jsa-8 | VAL_A_26_CA | VAL_A_26_HA | -9.824 | -6.661 | -0.296 | -9.524 | -7.348 | -1.087 | 1.09 | 136.54 | -66.41 |
| 153 | 2jsa-8 | VAL_A_26_CA | VAL_A_26_C | -9.824 | -6.661 | -0.296 | -9.114 | -5.326 | -0.533 | 1.53 | 98.91 | 61.99 |
| 154 | 2jsa-8 | VAL_A_26_C | VAL_A_26_O | -9.114 | -5.326 | -0.533 | -9.225 | -4.743 | -1.61 | 1.23 | 151.14 | 100.78 |
| 155 | 2jsa-8 | VAL_A_26_C | ILE_A_27_N | -9.114 | -5.326 | -0.533 | -8.399 | -4.883 | 0.49 | 1.32 | 39.43 | 31.78 |
| 156 | 2jsa-8 | ILE_A_27_N | ILE_A_27_H | -8.399 | -4.883 | 0.49 | -8.35 | -5.417 | 1.346 | 1.01 | 32.07 | -84.76 |
| 157 | 2jsa-8 | ILE_A_27_N | ILE_A_27_CA | -8.399 | -4.883 | 0.49 | -7.671 | -3.628 | 0.407 | 1.45 | 93.27 | 59.88 |
| 158 | 2jsa-8 | ILE_A_27_CA | ILE_A_27_HA | -7.671 | -3.628 | 0.407 | -8.175 | -2.992 | -0.321 | 1.09 | 131.90 | 128.40 |
| 159 | 2jsa-8 | ILE_A_27_CA | ILE_A_27_C | -7.671 | -3.628 | 0.407 | -6.257 | -3.898 | -0.112 | 1.53 | 109.83 | -10.81 |
| 160 | 2jsa-8 | ILE_A_27_C | ILE_A_27_O | -6.257 | -3.898 | -0.112 | -5.821 | -3.283 | -1.083 | 1.23 | 142.17 | 54.67 |
| 161 | 2jsa-8 | ILE_A_27_C | LYS_A_28_N | -6.257 | -3.898 | -0.112 | -5.58 | -4.818 | 0.56 | 1.33 | 59.53 | -53.65 |
| 162 | 2jsa-8 | LYS_A_28_N | LYS_A_28_H | -5.58 | -4.818 | 0.56 | -6.002 | -5.283 | 1.351 | 1.01 | 38.44 | -132.22 |
| 163 | 2jsa-8 | LYS_A_28_N | LYS_A_28_CA | -5.58 | -4.818 | 0.56 | -4.225 | -5.177 | 0.179 | 1.45 | 105.21 | -14.84 |
| 164 | 2jsa-8 | LYS_A_28_CA | LYS_A_28_HA | -4.225 | -5.177 | 0.179 | -3.637 | -4.261 | 0.122 | 1.09 | 93.00 | 57.30 |
| 165 | 2jsa-8 | LYS_A_28_CA | LYS_A_28_C | -4.225 | -5.177 | 0.179 | -4.241 | -5.807 | -1.215 | 1.53 | 155.67 | -91.45 |
| 166 | 2jsa-8 | LYS_A_28_C | LYS_A_28_O | -4.241 | -5.807 | -1.215 | -3.294 | -5.647 | -1.984 | 1.23 | 128.68 | 9.59 |
| 167 | 2jsa-8 | LYS_A_28_C | LYS_A_29_N | -4.241 | -5.807 | -1.215 | -5.327 | -6.51 | -1.499 | 1.32 | 102.38 | -147.08 |
| 168 | 2jsa-8 | LYS_A_29_N | LYS_A_29_H | -5.327 | -6.51 | -1.499 | -6.067 | -6.6 | -0.818 | 1.01 | 47.59 | -173.07 |
| 169 | 2jsa-8 | LYS_A_29_N | LYS_A_29_CA | -5.327 | -6.51 | -1.499 | -5.479 | -7.164 | -2.787 | 1.45 | 152.47 | -103.08 |
| 170 | 2jsa-8 | LYS_A_29_CA | LYS_A_29_HA | -5.479 | -7.164 | -2.787 | -4.556 | -7.702 | -3.003 | 1.09 | 101.43 | -30.24 |
| 171 | 2jsa-8 | LYS_A_29_CA | LYS_A_29_C | -5.479 | -7.164 | -2.787 | -5.664 | -6.105 | -3.874 | 1.53 | 135.32 | 99.91 |
| 172 | 2jsa-8 | LYS_A_29_C | LYS_A_29_O | -5.664 | -6.105 | -3.874 | -4.961 | -6.117 | -4.884 | 1.23 | 145.16 | -0.98 |
| 173 | 2jsa-8 | LYS_A_29_C | ASP_A_30_N | -5.664 | -6.105 | -3.874 | -6.614 | -5.213 | -3.632 | 1.33 | 79.48 | 136.80 |
| 174 | 2jsa-8 | ASP_A_30_N | ASP_A_30_H | -6.614 | -5.213 | -3.632 | -7.154 | -5.264 | -2.78 | 1.01 | 32.48 | -174.60 |
| 175 | 2jsa-8 | ASP_A_30_N | ASP_A_30_CA | -6.614 | -5.213 | -3.632 | -6.9 | -4.149 | -4.579 | 1.45 | 130.68 | 105.05 |
| 176 | 2jsa-8 | ASP_A_30_CA | ASP_A_30_HA | -6.9 | -4.149 | -4.579 | -7.382 | -4.575 | -5.459 | 1.09 | 143.83 | -138.53 |
| 177 | 2jsa-8 | ASP_A_30_CA | ASP_A_30_C | -6.9 | -4.149 | -4.579 | -5.591 | -3.466 | -4.982 | 1.53 | 105.27 | 27.55 |
| 178 | 2jsa-8 | ASP_A_30_C | ASP_A_30_O | -5.591 | -3.466 | -4.982 | -5.327 | -3.276 | -6.168 | 1.23 | 164.66 | 35.74 |
| 179 | 2jsa-8 | ASP_A_30_C | PHE_A_31_N | -5.591 | -3.466 | -4.982 | -4.807 | -3.117 | -3.973 | 1.32 | 40.38 | 24.00 |
| 180 | 2jsa-8 | PHE_A_31_N | PHE_A_31_H | -4.807 | -3.117 | -3.973 | -5.088 | -3.306 | -3.021 | 1.01 | 19.58 | -146.08 |
| 181 | 2jsa-8 | PHE_A_31_N | PHE_A_31_CA | -4.807 | -3.117 | -3.973 | -3.534 | -2.459 | -4.208 | 1.45 | 99.31 | 27.33 |
| 182 | 2jsa-8 | PHE_A_31_CA | PHE_A_31_HA | -3.534 | -2.459 | -4.208 | -3.701 | -1.423 | -4.503 | 1.09 | 105.70 | 99.16 |
| 183 | 2jsa-8 | PHE_A_31_CA | PHE_A_31_C | -3.534 | -2.459 | -4.208 | -2.736 | -3.183 | -5.295 | 1.53 | 135.25 | -42.22 |
| 184 | 2jsa-8 | PHE_A_31_C | PHE_A_31_O | -2.736 | -3.183 | -5.295 | -2.334 | -2.571 | -6.285 | 1.23 | 143.51 | 56.70 |
| 185 | 2jsa-8 | PHE_A_31_C | ASP_A_32_N | -2.736 | -3.183 | -5.295 | -2.532 | -4.473 | -5.076 | 1.32 | 80.48 | -81.01 |
| 186 | 2jsa-8 | ASP_A_32_N | ASP_A_32_H | -2.532 | -4.473 | -5.076 | -2.892 | -4.91 | -4.24 | 1.01 | 34.11 | -129.48 |
| 187 | 2jsa-8 | ASP_A_32_N | ASP_A_32_CA | -2.532 | -4.473 | -5.076 | -1.789 | -5.285 | -6.025 | 1.45 | 130.77 | -47.54 |
| 188 | 2jsa-8 | ASP_A_32_CA | ASP_A_32_HA | -1.789 | -5.285 | -6.025 | -0.76 | -4.926 | -6.073 | 1.09 | 92.52 | 19.23 |
| 189 | 2jsa-8 | ASP_A_32_CA | ASP_A_32_C | -1.789 | -5.285 | -6.025 | -2.443 | -5.176 | -7.404 | 1.53 | 154.32 | 170.54 |
| 190 | 2jsa-8 | ASP_A_32_C | ASP_A_32_O | -2.443 | -5.176 | -7.404 | -1.804 | -5.442 | -8.421 | 1.23 | 145.76 | -22.60 |
| 191 | 2jsa-8 | ASP_A_32_C | ALA_A_33_N | -2.443 | -5.176 | -7.404 | -3.709 | -4.785 | -7.394 | 1.33 | 89.57 | 162.84 |
| 192 | 2jsa-8 | ALA_A_33_N | ALA_A_33_H | -3.709 | -4.785 | -7.394 | -4.172 | -4.584 | -6.519 | 1.01 | 29.98 | 156.53 |
| 193 | 2jsa-8 | ALA_A_33_N | ALA_A_33_CA | -3.709 | -4.785 | -7.394 | -4.456 | -4.638 | -8.631 | 1.45 | 148.39 | 168.87 |
| 194 | 2jsa-8 | ALA_A_33_CA | ALA_A_33_HA | -4.456 | -4.638 | -8.631 | -4.275 | -5.513 | -9.255 | 1.09 | 124.93 | -78.31 |
| 195 | 2jsa-8 | ALA_A_33_CA | ALA_A_33_C | -4.456 | -4.638 | -8.631 | -3.959 | -3.399 | -9.378 | 1.53 | 119.23 | 68.14 |
| 196 | 2jsa-8 | ALA_A_33_C | ALA_A_33_O | -3.959 | -3.399 | -9.378 | -3.541 | -3.491 | -10.531 | 1.23 | 159.63 | -12.41 |
| 197 | 2jsa-8 | ALA_A_33_C | GLU_A_34_N | -3.959 | -3.399 | -9.378 | -4.021 | -2.268 | -8.69 | 1.33 | 58.73 | 93.14 |
| 198 | 2jsa-8 | GLU_A_34_N | GLU_A_34_H | -4.021 | -2.268 | -8.69 | -4.375 | -2.267 | -7.744 | 1.01 | 20.52 | 179.84 |
| 199 | 2jsa-8 | GLU_A_34_N | GLU_A_34_CA | -4.021 | -2.268 | -8.69 | -3.583 | -1.012 | -9.273 | 1.45 | 113.67 | 70.78 |
| 200 | 2jsa-8 | GLU_A_34_CA | GLU_A_34_HA | -3.583 | -1.012 | -9.273 | -3.824 | -1.031 | -10.336 | 1.09 | 167.19 | -175.49 |
| 201 | 2jsa-8 | GLU_A_34_CA | GLU_A_34_C | -3.583 | -1.012 | -9.273 | -2.067 | -0.86 | -9.127 | 1.53 | 84.53 | 5.73 |
| 202 | 2jsa-8 | GLU_A_34_C | GLU_A_34_O | -2.067 | -0.86 | -9.127 | -1.533 | 0.239 | -9.274 | 1.23 | 96.86 | 64.09 |
| 203 | 2jsa-8 | GLU_A_34_C | CYS_A_35_N | -2.067 | -0.86 | -9.127 | -1.417 | -1.978 | -8.84 | 1.32 | 77.49 | -59.83 |
| 204 | 2jsa-8 | CYS_A_35_N | CYS_A_35_H | -1.417 | -1.978 | -8.84 | -1.921 | -2.847 | -8.734 | 1.01 | 83.98 | -120.11 |
| 205 | 2jsa-8 | CYS_A_35_N | CYS_A_35_CA | -1.417 | -1.978 | -8.84 | 0.027 | -1.982 | -8.673 | 1.45 | 83.40 | -0.16 |
| 206 | 2jsa-8 | CYS_A_35_CA | CYS_A_35_HA | 0.027 | -1.982 | -8.673 | 0.393 | -0.965 | -8.82 | 1.09 | 97.74 | 70.21 |
| 207 | 2jsa-8 | CYS_A_35_CA | CYS_A_35_C | 0.027 | -1.982 | -8.673 | 0.633 | -2.879 | -9.754 | 1.53 | 134.96 | -55.96 |
| 208 | 2jsa-8 | CYS_A_35_C | CYS_A_35_O | 0.633 | -2.879 | -9.754 | 1.649 | -2.533 | -10.355 | 1.23 | 119.25 | 18.81 |
| 209 | 2jsa-8 | CYS_A_35_C | LYS_A_36_N | 0.633 | -2.879 | -9.754 | -0.016 | -4.013 | -9.969 | 1.32 | 99.34 | -119.78 |
| 210 | 2jsa-8 | LYS_A_36_N | LYS_A_36_H | -0.016 | -4.013 | -9.969 | -0.846 | -4.232 | -9.437 | 1.01 | 58.21 | -165.22 |
| 211 | 2jsa-8 | LYS_A_36_N | LYS_A_36_CA | -0.016 | -4.013 | -9.969 | 0.446 | -4.963 | -10.968 | 1.45 | 133.40 | -64.07 |
| 212 | 2jsa-8 | LYS_A_36_CA | LYS_A_36_HA | 0.446 | -4.963 | -10.968 | 1.493 | -5.188 | -10.761 | 1.09 | 79.06 | -12.13 |

| | | | | | | | | | | | |
|---|---|---|---|---|---|---|---|---|---|---|---|
| 213 | 2jsa-8 | LYS_A_36_CA | LYS_A_36_C | 0.446 | -4.963 | -10.968 | 0.376 | -4.316 | -12.352 | 1.53 | 154.82 | 96.17 |
| 214 | 2jsa-8 | LYS_A_36_C | LYS_A_36_O | 0.376 | -4.316 | -12.352 | 1.381 | -4.234 | -13.055 | 1.23 | 124.88 | 4.66 |
| 215 | 2jsa-8 | LYS_A_36_C | LYS_A_37_N | 0.376 | -4.316 | -12.352 | -0.823 | -3.871 | -12.701 | 1.33 | 105.26 | 159.64 |
| 216 | 2jsa-8 | LYS_A_37_N | LYS_A_37_H | -0.823 | -3.871 | -12.701 | -1.606 | -3.971 | -12.071 | 1.01 | 51.41 | -172.72 |
| 217 | 2jsa-8 | LYS_A_37_N | LYS_A_37_CA | -0.823 | -3.871 | -12.701 | -1.038 | -3.233 | -13.989 | 1.45 | 152.40 | 108.62 |
| 218 | 2jsa-8 | LYS_A_37_CA | LYS_A_37_HA | -1.038 | -3.233 | -13.989 | -0.584 | -3.86 | -14.756 | 1.09 | 134.74 | -54.09 |
| 219 | 2jsa-8 | LYS_A_37_CA | LYS_A_37_C | -1.038 | -3.233 | -13.989 | -0.321 | -1.881 | -14.01 | 1.53 | 90.79 | 62.06 |
| 220 | 2jsa-8 | LYS_A_37_C | LYS_A_37_O | -0.321 | -1.881 | -14.01 | 0.169 | -1.45 | -15.053 | 1.23 | 147.97 | 41.33 |
| 221 | 2jsa-8 | LYS_A_37_C | LEU_A_38_N | -0.321 | -1.881 | -14.01 | -0.282 | -1.25 | -12.845 | 1.33 | 28.49 | 86.46 |
| 222 | 2jsa-8 | LEU_A_38_N | LEU_A_38_H | -0.282 | -1.25 | -12.845 | -0.705 | -1.664 | -12.027 | 1.01 | 35.89 | -135.62 |
| 223 | 2jsa-8 | LEU_A_38_N | LEU_A_38_CA | -0.282 | -1.25 | -12.845 | 0.367 | 0.044 | -12.717 | 1.45 | 84.95 | 63.36 |
| 224 | 2jsa-8 | LEU_A_38_CA | LEU_A_38_HA | 0.367 | 0.044 | -12.717 | -0.102 | 0.729 | -13.423 | 1.09 | 130.38 | 124.40 |
| 225 | 2jsa-8 | LEU_A_38_CA | LEU_A_38_C | 0.367 | 0.044 | -12.717 | 1.84 | -0.088 | -13.107 | 1.53 | 104.77 | -5.12 |
| 226 | 2jsa-8 | LEU_A_38_C | LEU_A_38_O | 1.84 | -0.088 | -13.107 | 2.315 | 0.613 | -14.0 | 1.23 | 136.52 | 55.88 |
| 227 | 2jsa-8 | LEU_A_38_C | PHE_A_39_N | 1.84 | -0.088 | -13.107 | 2.524 | -0.992 | -12.421 | 1.33 | 58.82 | -52.89 |
| 228 | 2jsa-8 | PHE_A_39_N | PHE_A_39_H | 2.524 | -0.992 | -12.421 | 2.069 | -1.534 | -11.7 | 1.01 | 44.47 | -130.01 |
| 229 | 2jsa-8 | PHE_A_39_N | PHE_A_39_CA | 2.524 | -0.992 | -12.421 | 3.933 | -1.225 | -12.685 | 1.45 | 100.47 | -9.39 |
| 230 | 2jsa-8 | PHE_A_39_CA | PHE_A_39_HA | 3.933 | -1.225 | -12.685 | 4.267 | -0.585 | -13.502 | 1.09 | 138.54 | 62.44 |
| 231 | 2jsa-8 | PHE_A_39_CA | PHE_A_39_C | 3.933 | -1.225 | -12.685 | 4.19 | -2.69 | -13.041 | 1.53 | 103.46 | -80.05 |
| 232 | 2jsa-8 | PHE_A_39_C | PHE_A_39_O | 4.19 | -2.69 | -13.041 | 4.977 | -3.365 | -12.381 | 1.23 | 57.52 | -40.62 |
| 233 | 2jsa-8 | PHE_A_39_C | HIS_A_40_N | 4.19 | -2.69 | -13.041 | 3.509 | -3.139 | -14.086 | 1.33 | 142.03 | -146.60 |
| 234 | 2jsa-8 | HIS_A_40_N | HIS_A_40_H | 3.509 | -3.139 | -14.086 | 2.878 | -2.527 | -14.582 | 1.01 | 119.43 | 135.88 |
| 235 | 2jsa-8 | HIS_A_40_N | HIS_A_40_CA | 3.509 | -3.139 | -14.086 | 3.653 | -4.513 | -14.539 | 1.45 | 108.15 | -84.02 |
| 236 | 2jsa-8 | HIS_A_40_CA | HIS_A_40_HA | 3.653 | -4.513 | -14.539 | 3.548 | -5.166 | -13.672 | 1.09 | 37.34 | -99.13 |
| 237 | 2jsa-8 | HIS_A_40_CA | HIS_A_40_C | 3.653 | -4.513 | -14.539 | 5.055 | -4.719 | -15.114 | 1.53 | 112.09 | -8.36 |
| 238 | 2jsa-8 | HIS_A_40_C | HIS_A_40_O | 5.055 | -4.719 | -15.114 | 5.815 | -5.555 | -14.625 | 1.23 | 66.60 | -47.73 |
| 239 | 2jsa-8 | HIS_A_40_C | THR_A_41_N | 5.055 | -4.719 | -15.114 | 5.357 | -3.943 | -16.144 | 1.32 | 141.05 | 68.74 |
| 240 | 2jsa-8 | THR_A_41_N | THR_A_41_H | 5.357 | -3.943 | -16.144 | 4.684 | -3.276 | -16.493 | 1.01 | 110.22 | 135.26 |
| 241 | 2jsa-8 | THR_A_41_N | THR_A_41_CA | 5.357 | -3.943 | -16.144 | 6.655 | -4.031 | -16.792 | 1.45 | 116.48 | -3.88 |
| 242 | 2jsa-8 | THR_A_41_CA | THR_A_41_HA | 6.655 | -4.031 | -16.792 | 6.745 | -5.009 | -17.265 | 1.09 | 115.72 | -84.74 |
| 243 | 2jsa-8 | THR_A_41_CA | THR_A_41_C | 6.655 | -4.031 | -16.792 | 7.777 | -3.902 | -15.759 | 1.53 | 47.55 | 6.56 |
| 244 | 2jsa-8 | THR_A_41_C | THR_A_41_O | 7.777 | -3.902 | -15.759 | 8.767 | -4.629 | -15.82 | 1.23 | 92.84 | -36.29 |
| 245 | 2jsa-8 | THR_A_41_C | ILE_A_42_N | 7.777 | -3.902 | -15.759 | 7.581 | -2.974 | -14.834 | 1.32 | 45.72 | 101.93 |
| 246 | 2jsa-8 | ILE_A_42_N | ILE_A_42_H | 7.581 | -2.974 | -14.834 | 6.74 | -2.415 | -14.844 | 1.01 | 90.57 | 146.39 |
| 247 | 2jsa-8 | ILE_A_42_N | ILE_A_42_CA | 7.581 | -2.974 | -14.834 | 8.564 | -2.741 | -13.789 | 1.45 | 44.03 | 13.33 |
| 248 | 2jsa-8 | ILE_A_42_CA | ILE_A_42_HA | 8.564 | -2.741 | -13.789 | 9.375 | -2.149 | -14.214 | 1.09 | 112.94 | 36.13 |
| 249 | 2jsa-8 | ILE_A_42_CA | ILE_A_42_C | 8.564 | -2.741 | -13.789 | 9.152 | -4.08 | -13.341 | 1.53 | 72.97 | -66.29 |
| 250 | 2jsa-8 | ILE_A_42_C | ILE_A_42_O | 9.152 | -4.08 | -13.341 | 8.436 | -5.076 | -13.244 | 1.23 | 85.48 | -125.71 |
| 251 | 2jsa-8 | ILE_A_42_C | PRO_A_43_N | 9.152 | -4.08 | -13.341 | 10.486 | -4.061 | -13.073 | 1.36 | 78.64 | 0.82 |
| 252 | 2jsa-8 | PRO_A_43_N | PRO_A_43_CA | 10.486 | -4.061 | -13.073 | 11.179 | -5.262 | -12.637 | 1.45 | 72.54 | -60.01 |
| 253 | 2jsa-8 | PRO_A_43_CA | PRO_A_43_HA | 11.179 | -5.262 | -12.637 | 10.879 | -6.11 | -13.252 | 1.09 | 124.36 | -109.48 |
| 254 | 2jsa-8 | PRO_A_43_CA | PRO_A_43_C | 11.179 | -5.262 | -12.637 | 10.853 | -5.583 | -11.177 | 1.53 | 17.40 | -135.44 |
| 255 | 2jsa-8 | PRO_A_43_C | PRO_A_43_O | 10.853 | -5.583 | -11.177 | 10.515 | -6.719 | -10.847 | 1.23 | 74.44 | -106.57 |
| 256 | 2jsa-8 | PRO_A_43_C | PHE_A_44_N | 10.853 | -5.583 | -11.177 | 10.969 | -4.563 | -10.34 | 1.32 | 50.81 | 83.51 |
| 257 | 2jsa-8 | PHE_A_44_N | PHE_A_44_H | 10.969 | -4.563 | -10.34 | 11.255 | -3.656 | -10.68 | 1.01 | 109.67 | 72.50 |
| 258 | 2jsa-8 | PHE_A_44_N | PHE_A_44_CA | 10.969 | -4.563 | -10.34 | 10.691 | -4.722 | -8.923 | 1.45 | 12.74 | -150.23 |
| 259 | 2jsa-8 | PHE_A_44_CA | PHE_A_44_HA | 10.691 | -4.722 | -8.923 | 10.641 | -5.782 | -8.672 | 1.09 | 76.69 | -92.70 |
| 260 | 2jsa-8 | PHE_A_44_CA | PHE_A_44_C | 10.691 | -4.722 | -8.923 | 9.376 | -4.04 | -8.541 | 1.53 | 75.54 | 152.59 |
| 261 | 2jsa-8 | PHE_A_44_C | PHE_A_44_O | 9.376 | -4.04 | -8.541 | 9.223 | -3.564 | -7.417 | 1.23 | 23.98 | 107.82 |
| 262 | 2jsa-8 | PHE_A_44_C | GLY_A_45_N | 9.376 | -4.04 | -8.541 | 8.46 | -4.016 | -9.498 | 1.32 | 136.24 | 178.50 |
| 263 | 2jsa-8 | GLY_A_45_N | GLY_A_45_H | 8.46 | -4.016 | -9.498 | 8.656 | -4.428 | -10.399 | 1.01 | 153.14 | -64.56 |
| 264 | 2jsa-8 | GLY_A_45_N | GLY_A_45_CA | 8.46 | -4.016 | -9.498 | 7.163 | -3.401 | -9.277 | 1.45 | 81.25 | 154.63 |
| 265 | 2jsa-8 | GLY_A_45_CA | GLY_A_45_C | 7.163 | -3.401 | -9.277 | 6.175 | -4.404 | -8.676 | 1.53 | 66.88 | -134.57 |
| 266 | 2jsa-8 | GLY_A_45_C | GLY_A_45_O | 6.175 | -4.404 | -8.676 | 5.879 | -4.35 | -7.483 | 1.23 | 14.16 | 169.66 |
| 267 | 2jsa-8 | GLY_A_45_C | THR_A_46_N | 6.175 | -4.404 | -8.676 | 5.694 | -5.295 | -9.53 | 1.32 | 130.15 | -118.36 |
| 268 | 2jsa-8 | THR_A_46_N | THR_A_46_H | 5.694 | -5.295 | -9.53 | 5.983 | -5.28 | -10.498 | 1.01 | 163.36 | 2.97 |
| 269 | 2jsa-8 | THR_A_46_N | THR_A_46_CA | 5.694 | -5.295 | -9.53 | 4.747 | -6.309 | -9.098 | 1.45 | 72.71 | -133.04 |
| 270 | 2jsa-8 | THR_A_46_CA | THR_A_46_HA | 4.747 | -6.309 | -9.098 | 3.852 | -5.814 | -8.72 | 1.09 | 69.72 | 151.05 |
| 271 | 2jsa-8 | THR_A_46_CA | THR_A_46_C | 4.747 | -6.309 | -9.098 | 5.339 | -7.143 | -7.961 | 1.53 | 41.97 | -54.63 |
| 272 | 2jsa-8 | THR_A_46_C | THR_A_46_O | 5.339 | -7.143 | -7.961 | 4.619 | -7.568 | -7.058 | 1.23 | 42.80 | -149.45 |
| 273 | 2jsa-8 | THR_A_46_C | ARG_A_47_N | 5.339 | -7.143 | -7.961 | 6.644 | -7.351 | -8.041 | 1.32 | 93.46 | -9.06 |
| 274 | 2jsa-8 | ARG_A_47_N | ARG_A_47_H | 6.644 | -7.351 | -8.041 | 7.174 | -6.971 | -8.813 | 1.01 | 139.81 | 35.64 |
| 275 | 2jsa-8 | ARG_A_47_N | ARG_A_47_CA | 6.644 | -7.351 | -8.041 | 7.342 | -8.126 | -7.028 | 1.45 | 45.84 | -47.99 |
| 276 | 2jsa-8 | ARG_A_47_CA | ARG_A_47_HA | 7.342 | -8.126 | -7.028 | 6.901 | -9.122 | -6.992 | 1.09 | 88.11 | -113.88 |
| 277 | 2jsa-8 | ARG_A_47_CA | ARG_A_47_C | 7.342 | -8.126 | -7.028 | 7.191 | -7.465 | -5.656 | 1.53 | 26.30 | 102.87 |
| 278 | 2jsa-8 | ARG_A_47_C | ARG_A_47_O | 7.191 | -7.465 | -5.656 | 6.843 | -8.128 | -4.68 | 1.23 | 37.50 | -117.69 |
| 279 | 2jsa-8 | ARG_A_47_C | GLU_A_48_N | 7.191 | -7.465 | -5.656 | 7.461 | -6.169 | -5.627 | 1.32 | 88.75 | 78.23 |
| 280 | 2jsa-8 | GLU_A_48_N | GLU_A_48_H | 7.461 | -6.169 | -5.627 | 7.742 | -5.689 | -6.47 | 1.01 | 146.58 | 59.65 |
| 281 | 2jsa-8 | GLU_A_48_N | GLU_A_48_CA | 7.461 | -6.169 | -5.627 | 7.36 | -5.412 | -4.392 | 1.45 | 31.73 | 97.60 |
| 282 | 2jsa-8 | GLU_A_48_CA | GLU_A_48_HA | 7.36 | -5.412 | -4.392 | 7.886 | -5.965 | -3.614 | 1.09 | 44.45 | -46.43 |
| 283 | 2jsa-8 | GLU_A_48_CA | GLU_A_48_C | 7.36 | -5.412 | -4.392 | 5.895 | -5.27 | -3.974 | 1.53 | 74.15 | 174.46 |
| 284 | 2jsa-8 | GLU_A_48_C | GLU_A_48_O | 5.895 | -5.27 | -3.974 | 5.577 | -5.316 | -2.787 | 1.23 | 15.15 | -171.77 |
| 285 | 2jsa-8 | GLU_A_48_C | CYS_A_49_N | 5.895 | -5.27 | -3.974 | 5.043 | -5.098 | -4.974 | 1.32 | 139.00 | 168.59 |
| 286 | 2jsa-8 | CYS_A_49_N | CYS_A_49_H | 5.043 | -5.098 | -4.974 | 5.376 | -5.07 | -5.927 | 1.01 | 160.68 | 4.81 |
| 287 | 2jsa-8 | CYS_A_49_N | CYS_A_49_CA | 5.043 | -5.098 | -4.974 | 3.619 | -4.948 | -4.726 | 1.45 | 80.17 | 173.99 |
| 288 | 2jsa-8 | CYS_A_49_CA | CYS_A_49_HA | 3.619 | -4.948 | -4.726 | 3.458 | -3.997 | -4.218 | 1.09 | 62.23 | 99.61 |
| 289 | 2jsa-8 | CYS_A_49_CA | CYS_A_49_C | 3.619 | -4.948 | -4.726 | 3.172 | -6.079 | -3.797 | 1.53 | 52.62 | -111.57 |
| 290 | 2jsa-8 | CYS_A_49_C | CYS_A_49_O | 3.172 | -6.079 | -3.797 | 2.754 | -5.831 | -2.668 | 1.23 | 23.29 | 149.32 |
| 291 | 2jsa-8 | CYS_A_49_C | ASP_A_50_N | 3.172 | -6.079 | -3.797 | 3.274 | -7.298 | -4.308 | 1.33 | 112.67 | -85.22 |
| 292 | 2jsa-8 | ASP_A_50_N | ASP_A_50_H | 3.274 | -7.298 | -4.308 | 3.625 | -7.429 | -5.246 | 1.01 | 158.23 | -20.47 |
| 293 | 2jsa-8 | ASP_A_50_N | ASP_A_50_CA | 3.274 | -7.298 | -4.308 | 2.886 | -8.467 | -3.538 | 1.45 | 57.99 | -108.36 |
| 294 | 2jsa-8 | ASP_A_50_CA | ASP_A_50_HA | 2.886 | -8.467 | -3.538 | 1.814 | -8.428 | -3.348 | 1.09 | 79.96 | 177.92 |
| 295 | 2jsa-8 | ASP_A_50_CA | ASP_A_50_C | 2.886 | -8.467 | -3.538 | 3.648 | -8.476 | -2.212 | 1.53 | 29.89 | -0.68 |
| 296 | 2jsa-8 | ASP_A_50_C | ASP_A_50_O | 3.648 | -8.476 | -2.212 | 3.168 | -9.019 | -1.218 | 1.23 | 36.10 | -131.48 |
| 297 | 2jsa-8 | ASP_A_50_C | HIS_A_51_N | 3.648 | -8.476 | -2.212 | 4.825 | -7.867 | -2.238 | 1.33 | 91.12 | 27.36 |
| 298 | 2jsa-8 | HIS_A_51_N | HIS_A_51_H | 4.825 | -7.867 | -2.238 | 5.159 | -7.441 | -3.091 | 1.01 | 147.60 | 51.90 |
| 299 | 2jsa-8 | HIS_A_51_N | HIS_A_51_CA | 4.825 | -7.867 | -2.238 | 5.658 | -7.798 | -1.05 | 1.45 | 35.13 | 4.74 |
| 300 | 2jsa-8 | HIS_A_51_CA | HIS_A_51_HA | 5.658 | -7.798 | -1.05 | 5.453 | -8.679 | -0.443 | 1.09 | 56.14 | -103.10 |
| 301 | 2jsa-8 | HIS_A_51_CA | HIS_A_51_C | 5.658 | -7.798 | -1.05 | 5.281 | -6.563 | -0.229 | 1.53 | 57.55 | 106.98 |
| 302 | 2jsa-8 | HIS_A_51_C | HIS_A_51_O | 5.281 | -6.563 | -0.229 | 5.973 | -6.212 | 0.726 | 1.23 | 39.09 | 26.90 |
| 303 | 2jsa-8 | HIS_A_51_C | TYR_A_52_N | 5.281 | -6.563 | -0.229 | 4.184 | -5.938 | -0.631 | 1.33 | 107.66 | 150.33 |
| 304 | 2jsa-8 | TYR_A_52_N | TYR_A_52_H | 4.184 | -5.938 | -0.631 | 3.665 | -6.282 | -1.426 | 1.01 | 141.93 | -146.46 |
| 305 | 2jsa-8 | TYR_A_52_N | TYR_A_52_CA | 4.184 | -5.938 | -0.631 | 3.706 | -4.75 | 0.056 | 1.45 | 61.79 | 111.92 |
| 306 | 2jsa-8 | TYR_A_52_CA | TYR_A_52_HA | 3.706 | -4.75 | 0.056 | 4.434 | -4.426 | 0.801 | 1.09 | 46.93 | 23.99 |
| 307 | 2jsa-8 | TYR_A_52_CA | TYR_A_52_C | 3.706 | -4.75 | 0.056 | 2.356 | -5.01 | 0.727 | 1.53 | 63.98 | -169.10 |
| 308 | 2jsa-8 | TYR_A_52_C | TYR_A_52_O | 2.356 | -5.01 | 0.727 | 2.168 | -4.68 | 1.897 | 1.23 | 17.98 | 119.67 |

| | | | | | | | | | | | | |
|---|---|---|---|---|---|---|---|---|---|---|---|---|
| 309 | 2jsa-8 | TYR_A_52_C | VAL_A_53_N | 2.356 | -5.01 | 0.727 | 1.453 | -5.6 | -0.042 | 1.32 | 125.49 | -146.84 |
| 310 | 2jsa-8 | VAL_A_53_N | VAL_A_53_H | 1.453 | -5.6 | -0.042 | 1.679 | -5.844 | -0.995 | 1.01 | 160.76 | -47.19 |
| 311 | 2jsa-8 | VAL_A_53_N | VAL_A_53_CA | 1.453 | -5.6 | -0.042 | 0.126 | -5.908 | 0.463 | 1.45 | 69.66 | -166.93 |
| 312 | 2jsa-8 | VAL_A_53_CA | VAL_A_53_HA | 0.126 | -5.908 | 0.463 | -0.363 | -4.974 | 0.739 | 1.09 | 75.33 | 117.63 |
| 313 | 2jsa-8 | VAL_A_53_CA | VAL_A_53_C | 0.126 | -5.908 | 0.463 | 0.253 | -6.769 | 1.722 | 1.53 | 34.66 | -81.61 |
| 314 | 2jsa-8 | VAL_A_53_C | VAL_A_53_O | 0.253 | -6.769 | 1.722 | -0.663 | -6.813 | 2.543 | 1.23 | 48.16 | -177.25 |
| 315 | 2jsa-8 | VAL_A_53_C | ASN_A_54_N | 0.253 | -6.769 | 1.722 | 1.394 | -7.431 | 1.836 | 1.32 | 85.06 | -30.12 |
| 316 | 2jsa-8 | ASN_A_54_N | ASN_A_54_H | 1.394 | -7.431 | 1.836 | 2.104 | -7.347 | 1.122 | 1.01 | 134.96 | 6.75 |
| 317 | 2jsa-8 | ASN_A_54_N | ASN_A_54_CA | 1.394 | -7.431 | 1.836 | 1.652 | -8.287 | 2.981 | 1.45 | 37.98 | -73.23 |
| 318 | 2jsa-8 | ASN_A_54_CA | ASN_A_54_HA | 1.652 | -8.287 | 2.981 | 0.919 | -8.073 | 3.759 | 1.09 | 44.46 | 163.72 |
| 319 | 2jsa-8 | ASN_A_54_CA | ASN_A_54_C | 1.652 | -8.287 | 2.981 | 3.063 | -8.017 | 3.508 | 1.53 | 69.86 | 10.83 |
| 320 | 2jsa-8 | ASN_A_54_C | ASN_A_54_O | 3.063 | -8.017 | 3.508 | 3.891 | -8.924 | 3.571 | 1.23 | 87.06 | -47.61 |
| 321 | 2jsa-8 | ASN_A_54_C | SER_A_55_N | 3.063 | -8.017 | 3.508 | 3.294 | -6.764 | 3.872 | 1.33 | 74.06 | 79.55 |
| 322 | 2jsa-8 | SER_A_55_N | SER_A_55_H | 3.294 | -6.764 | 3.872 | 2.566 | -6.069 | 3.793 | 1.01 | 94.49 | 136.33 |
| 323 | 2jsa-8 | SER_A_55_N | SER_A_55_CA | 3.294 | -6.764 | 3.872 | 4.59 | -6.362 | 4.391 | 1.45 | 69.07 | 17.23 |
| 324 | 2jsa-8 | SER_A_55_CA | SER_A_55_HA | 4.59 | -6.362 | 4.391 | 4.843 | -7.018 | 5.224 | 1.09 | 40.17 | -68.91 |
| 325 | 2jsa-8 | SER_A_55_CA | SER_A_55_C | 4.59 | -6.362 | 4.391 | 4.526 | -4.921 | 4.903 | 1.53 | 70.46 | 92.54 |
| 326 | 2jsa-8 | SER_A_55_C | SER_A_55_O | 4.526 | -4.921 | 4.903 | 5.079 | -4.608 | 5.956 | 1.23 | 31.11 | 29.51 |
| 327 | 2jsa-8 | SER_A_55_C | LYS_A_56_N | 4.526 | -4.921 | 4.903 | 3.846 | -4.084 | 4.133 | 1.33 | 125.53 | 129.09 |
| 328 | 2jsa-8 | LYS_A_56_N | LYS_A_56_H | 3.846 | -4.084 | 4.133 | 3.418 | -4.412 | 3.279 | 1.01 | 147.73 | -142.54 |
| 329 | 2jsa-8 | LYS_A_56_N | LYS_A_56_CA | 3.846 | -4.084 | 4.133 | 3.701 | -2.684 | 4.496 | 1.45 | 75.54 | 95.91 |
| 330 | 2jsa-8 | LYS_A_56_CA | LYS_A_56_HA | 3.701 | -2.684 | 4.496 | 4.104 | -2.552 | 5.5 | 1.09 | 22.90 | 18.14 |
| 331 | 2jsa-8 | LYS_A_56_CA | LYS_A_56_C | 3.701 | -2.684 | 4.496 | 2.215 | -2.324 | 4.541 | 1.53 | 88.31 | 166.38 |
| 332 | 2jsa-8 | LYS_A_56_C | LYS_A_56_O | 2.215 | -2.324 | 4.541 | 1.858 | -1.146 | 4.545 | 1.23 | 89.81 | 106.86 |
| 333 | 2jsa-8 | LYS_A_56_C | VAL_A_57_N | 2.215 | -2.324 | 4.541 | 1.389 | -3.359 | 4.573 | 1.32 | 88.62 | -128.59 |
| 334 | 2jsa-8 | VAL_A_57_N | VAL_A_57_H | 1.389 | -3.359 | 4.573 | 1.754 | -4.301 | 4.567 | 1.01 | 90.34 | -68.82 |
| 335 | 2jsa-8 | VAL_A_57_N | VAL_A_57_CA | 1.389 | -3.359 | 4.573 | -0.05 | -3.166 | 4.617 | 1.45 | 88.26 | 172.36 |
| 336 | 2jsa-8 | VAL_A_57_CA | VAL_A_57_HA | -0.05 | -3.166 | 4.617 | -0.273 | -2.159 | 4.264 | 1.09 | 108.89 | 102.49 |
| 337 | 2jsa-8 | VAL_A_57_CA | VAL_A_57_C | -0.05 | -3.166 | 4.617 | -0.533 | -3.279 | 6.064 | 1.53 | 18.92 | -166.83 |
| 338 | 2jsa-8 | VAL_A_57_C | VAL_A_57_O | -0.533 | -3.279 | 6.064 | -1.312 | -2.449 | 6.53 | 1.23 | 67.74 | 133.18 |
| 339 | 2jsa-8 | VAL_A_57_C | ASP_A_58_N | -0.533 | -3.279 | 6.064 | -0.047 | -4.312 | 6.737 | 1.33 | 59.48 | -64.80 |
| 340 | 2jsa-8 | ASP_A_58_N | ASP_A_58_H | -0.047 | -4.312 | 6.737 | 0.592 | -4.955 | 6.291 | 1.01 | 116.20 | -45.18 |
| 341 | 2jsa-8 | ASP_A_58_N | ASP_A_58_CA | -0.047 | -4.312 | 6.737 | -0.418 | -4.544 | 8.122 | 1.45 | 17.53 | -147.98 |
| 342 | 2jsa-8 | ASP_A_58_CA | ASP_A_58_HA | -0.418 | -4.544 | 8.122 | -1.468 | -4.833 | 8.166 | 1.09 | 87.69 | -164.61 |
| 343 | 2jsa-8 | ASP_A_58_CA | ASP_A_58_C | -0.418 | -4.544 | 8.122 | -0.198 | -3.261 | 8.927 | 1.53 | 58.27 | 80.27 |
| 344 | 2jsa-8 | ASP_A_58_C | ASP_A_58_O | -0.198 | -3.261 | 8.927 | -1.05 | -2.87 | 9.723 | 1.23 | 49.66 | 155.35 |
| 345 | 2jsa-8 | ASP_A_58_C | PRO_A_59_N | -0.198 | -3.261 | 8.927 | 0.979 | -2.625 | 8.684 | 1.36 | 100.29 | 28.38 |
| 346 | 2jsa-8 | PRO_A_59_N | PRO_A_59_CA | 0.979 | -2.625 | 8.684 | 1.322 | -1.394 | 9.375 | 1.45 | 61.60 | 74.43 |
| 347 | 2jsa-8 | PRO_A_59_CA | PRO_A_59_HA | 1.322 | -1.394 | 9.375 | 1.1 | -1.489 | 10.438 | 1.09 | 12.80 | -156.83 |
| 348 | 2jsa-8 | PRO_A_59_CA | PRO_A_59_C | 1.322 | -1.394 | 9.375 | 0.526 | -0.212 | 8.817 | 1.53 | 111.38 | 123.96 |
| 349 | 2jsa-8 | PRO_A_59_C | PRO_A_59_O | 0.526 | -0.212 | 8.817 | -0.175 | 0.475 | 9.558 | 1.23 | 52.95 | 135.58 |
| 350 | 2jsa-8 | PRO_A_59_C | ILE_A_60_N | 0.526 | -0.212 | 8.817 | 0.664 | -0.01 | 7.514 | 1.33 | 169.37 | 55.66 |
| 351 | 2jsa-8 | ILE_A_60_N | ILE_A_60_H | 0.664 | -0.01 | 7.514 | 1.258 | -0.615 | 6.965 | 1.01 | 122.92 | -45.53 |
| 352 | 2jsa-8 | ILE_A_60_N | ILE_A_60_CA | 0.664 | -0.01 | 7.514 | -0.032 | 1.078 | 6.849 | 1.45 | 117.24 | 122.61 |
| 353 | 2jsa-8 | ILE_A_60_CA | ILE_A_60_HA | -0.032 | 1.078 | 6.849 | 0.434 | 2.015 | 7.157 | 1.09 | 73.60 | 63.56 |
| 354 | 2jsa-8 | ILE_A_60_CA | ILE_A_60_C | -0.032 | 1.078 | 6.849 | -1.488 | 1.107 | 7.315 | 1.53 | 72.26 | 178.86 |
| 355 | 2jsa-8 | ILE_A_60_C | ILE_A_60_O | -1.488 | 1.107 | 7.315 | -1.952 | 2.111 | 7.854 | 1.23 | 64.02 | 114.80 |
| 356 | 2jsa-8 | ILE_A_60_C | ILE_A_61_N | -1.488 | 1.107 | 7.315 | -2.172 | -0.006 | 7.091 | 1.33 | 99.73 | -121.57 |
| 357 | 2jsa-8 | ILE_A_61_N | ILE_A_61_H | -2.172 | -0.006 | 7.091 | -1.727 | -0.794 | 6.643 | 1.01 | 116.34 | -60.55 |
| 358 | 2jsa-8 | ILE_A_61_N | ILE_A_61_CA | -2.172 | -0.006 | 7.091 | -3.566 | -0.121 | 7.482 | 1.45 | 74.38 | -175.28 |
| 359 | 2jsa-8 | ILE_A_61_CA | ILE_A_61_HA | -3.566 | -0.121 | 7.482 | -4.124 | 0.667 | 6.976 | 1.09 | 117.66 | 125.30 |
| 360 | 2jsa-8 | ILE_A_61_CA | ILE_A_61_C | -3.566 | -0.121 | 7.482 | -3.69 | 0.115 | 8.988 | 1.53 | 10.04 | 117.72 |
| 361 | 2jsa-8 | ILE_A_61_C | ILE_A_61_O | -3.69 | 0.115 | 8.988 | -4.758 | 0.48 | 9.478 | 1.23 | 66.53 | 161.13 |
| 362 | 2jsa-8 | ILE_A_61_C | HIS_A_62_N | -3.69 | 0.115 | 8.988 | -2.582 | -0.102 | 9.682 | 1.33 | 58.42 | -11.08 |
| 363 | 2jsa-8 | HIS_A_62_N | HIS_A_62_H | -2.582 | -0.102 | 9.682 | -1.738 | -0.4 | 9.214 | 1.01 | 117.60 | -19.45 |
| 364 | 2jsa-8 | HIS_A_62_N | HIS_A_62_CA | -2.582 | -0.102 | 9.682 | -2.554 | 0.082 | 11.123 | 1.45 | 7.36 | 81.35 |
| 365 | 2jsa-8 | HIS_A_62_CA | HIS_A_62_HA | -2.554 | 0.082 | 11.123 | -3.379 | -0.487 | 11.552 | 1.09 | 66.83 | -145.41 |
| 366 | 2jsa-8 | HIS_A_62_CA | HIS_A_62_C | -2.554 | 0.082 | 11.123 | -2.777 | 1.558 | 11.456 | 1.53 | 77.42 | 98.59 |
| 367 | 2jsa-8 | HIS_A_62_C | HIS_A_62_O | -2.777 | 1.558 | 11.456 | -3.656 | 1.893 | 12.249 | 1.23 | 49.87 | 159.14 |
| 368 | 2jsa-8 | HIS_A_62_C | GLU_A_63_N | -2.777 | 1.558 | 11.456 | -1.968 | 2.402 | 10.834 | 1.32 | 118.01 | 46.21 |
| 369 | 2jsa-8 | GLU_A_63_N | GLU_A_63_H | -1.968 | 2.402 | 10.834 | -1.267 | 2.056 | 10.194 | 1.01 | 129.31 | -26.27 |
| 370 | 2jsa-8 | GLU_A_63_N | GLU_A_63_CA | -1.968 | 2.402 | 10.834 | -2.067 | 3.835 | 11.054 | 1.45 | 81.29 | 93.95 |
| 371 | 2jsa-8 | GLU_A_63_CA | GLU_A_63_HA | -2.067 | 3.835 | 11.054 | -1.824 | 4.034 | 12.098 | 1.09 | 16.74 | 39.32 |
| 372 | 2jsa-8 | GLU_A_63_CA | GLU_A_63_C | -2.067 | 3.835 | 11.054 | -3.496 | 4.314 | 10.794 | 1.53 | 99.79 | 161.47 |
| 373 | 2jsa-8 | GLU_A_63_C | GLU_A_63_O | -3.496 | 4.314 | 10.794 | -4.078 | 5.017 | 11.619 | 1.23 | 47.89 | 129.62 |
| 374 | 2jsa-8 | GLU_A_63_C | LEU_A_64_N | -3.496 | 4.314 | 10.794 | -4.02 | 3.916 | 9.644 | 1.32 | 150.22 | -142.78 |
| 375 | 2jsa-8 | LEU_A_64_N | LEU_A_64_H | -4.02 | 3.916 | 9.644 | -3.482 | 3.339 | 9.013 | 1.01 | 128.65 | -47.00 |
| 376 | 2jsa-8 | LEU_A_64_N | LEU_A_64_CA | -4.02 | 3.916 | 9.644 | -5.37 | 4.297 | 9.265 | 1.45 | 105.12 | 164.24 |
| 377 | 2jsa-8 | LEU_A_64_CA | LEU_A_64_HA | -5.37 | 4.297 | 9.265 | -5.428 | 5.386 | 9.27 | 1.09 | 89.74 | 93.05 |
| 378 | 2jsa-8 | LEU_A_64_CA | LEU_A_64_C | -5.37 | 4.297 | 9.265 | -6.355 | 3.774 | 10.312 | 1.53 | 46.81 | -152.03 |
| 379 | 2jsa-8 | LEU_A_64_C | LEU_A_64_O | -6.355 | 3.774 | 10.312 | -7.303 | 4.465 | 10.678 | 1.23 | 72.67 | 143.91 |
| 380 | 2jsa-8 | LEU_A_64_C | GLU_A_65_N | -6.355 | 3.774 | 10.312 | -6.097 | 2.554 | 10.763 | 1.33 | 70.12 | -78.06 |
| 381 | 2jsa-8 | GLU_A_65_N | GLU_A_65_H | -6.097 | 2.554 | 10.763 | -5.298 | 2.042 | 10.417 | 1.01 | 110.03 | -32.65 |
| 382 | 2jsa-8 | GLU_A_65_N | GLU_A_65_CA | -6.097 | 2.554 | 10.763 | -6.95 | 1.929 | 11.76 | 1.45 | 46.69 | -143.77 |
| 383 | 2jsa-8 | GLU_A_65_CA | GLU_A_65_HA | -6.95 | 1.929 | 11.76 | -7.949 | 1.832 | 11.337 | 1.09 | 112.85 | -174.45 |
| 384 | 2jsa-8 | GLU_A_65_CA | GLU_A_65_C | -6.95 | 1.929 | 11.76 | -7.039 | 2.81 | 13.009 | 1.53 | 35.33 | 95.77 |
| 385 | 2jsa-8 | GLU_A_65_C | GLU_A_65_O | -7.039 | 2.81 | 13.009 | -8.131 | 3.19 | 13.427 | 1.23 | 70.12 | 160.81 |
| 386 | 2jsa-8 | GLU_A_65_C | GLY_A_66_N | -7.039 | 2.81 | 13.009 | -5.876 | 3.109 | 13.568 | 1.32 | 65.04 | 14.42 |
| 387 | 2jsa-8 | GLY_A_66_N | GLY_A_66_H | -5.876 | 3.109 | 13.568 | -5.015 | 2.761 | 13.169 | 1.01 | 113.25 | -22.01 |
| 388 | 2jsa-8 | GLY_A_66_N | GLY_A_66_CA | -5.876 | 3.109 | 13.568 | -5.809 | 3.938 | 14.76 | 1.45 | 34.90 | 85.38 |
| 389 | 2jsa-8 | GLY_A_66_CA | GLY_A_66_C | -5.809 | 3.938 | 14.76 | -6.505 | 5.281 | 14.533 | 1.53 | 98.53 | 117.40 |
| 390 | 2jsa-8 | GLY_A_66_C | GLY_A_66_O | -6.505 | 5.281 | 14.533 | -7.275 | 5.736 | 15.377 | 1.23 | 46.66 | 149.42 |
| 391 | 2jsa-8 | GLY_A_66_C | GLY_A_67_N | -6.505 | 5.281 | 14.533 | -6.209 | 5.879 | 13.387 | 1.33 | 149.79 | 63.67 |
| 392 | 2jsa-8 | GLY_A_67_N | GLY_A_67_H | -6.209 | 5.879 | 13.387 | -5.566 | 5.447 | 12.739 | 1.01 | 129.91 | -33.90 |
| 393 | 2jsa-8 | GLY_A_67_N | GLY_A_67_CA | -6.209 | 5.879 | 13.387 | -6.797 | 7.161 | 13.039 | 1.45 | 103.86 | 114.64 |
| 394 | 2jsa-8 | GLY_A_67_CA | GLY_A_67_C | -6.797 | 7.161 | 13.039 | -5.743 | 8.108 | 12.463 | 1.53 | 112.12 | 41.94 |
| 395 | 2jsa-8 | GLY_A_67_C | GLY_A_67_O | -5.743 | 8.108 | 12.463 | -5.688 | 9.28 | 12.831 | 1.23 | 72.59 | 87.31 |
| 396 | 2jsa-8 | GLY_A_67_C | THR_A_68_N | -5.743 | 8.108 | 12.463 | -4.931 | 7.564 | 11.568 | 1.33 | 132.48 | -33.82 |
| 397 | 2jsa-8 | THR_A_68_N | THR_A_68_H | -4.931 | 7.564 | 11.568 | -5.033 | 6.592 | 11.313 | 1.01 | 104.62 | -95.99 |
| 398 | 2jsa-8 | THR_A_68_N | THR_A_68_CA | -4.931 | 7.564 | 11.568 | -3.882 | 8.347 | 10.936 | 1.45 | 115.77 | 36.74 |
| 399 | 2jsa-8 | THR_A_68_CA | THR_A_68_HA | -3.882 | 8.347 | 10.936 | -3.84 | 9.326 | 11.413 | 1.09 | 64.04 | 87.54 |
| 400 | 2jsa-8 | THR_A_68_CA | THR_A_68_C | -3.882 | 8.347 | 10.936 | -4.193 | 8.559 | 9.454 | 1.53 | 165.75 | 145.72 |
| 401 | 2jsa-8 | THR_A_68_C | THR_A_68_O | -4.193 | 8.559 | 9.454 | -4.64 | 7.637 | 8.772 | 1.23 | 123.65 | -115.86 |
| 402 | 2jsa-8 | THR_A_68_C | ALA_A_69_N | -4.193 | 8.559 | 9.454 | -3.946 | 9.778 | 8.998 | 1.32 | 110.13 | 78.55 |
| 403 | 2jsa-8 | ALA_A_69_N | ALA_A_69_H | -3.946 | 9.778 | 8.998 | -3.579 | 10.487 | 9.617 | 1.01 | 52.21 | 62.63 |
| 404 | 2jsa-8 | ALA_A_69_N | ALA_A_69_CA | -3.946 | 9.778 | 8.998 | -4.195 | 10.123 | 7.609 | 1.45 | 162.97 | 125.82 |

| | | | | | | | | | | | |
|---|---|---|---|---|---|---|---|---|---|---|---|
| 405 | 2jsa-8 | ALA_A_69_CA | ALA_A_69_HA | -4.195 | 10.123 | 7.609 | -5.257 | 9.994 | 7.401 | 1.09 | 101.00 | -173.07 |
| 406 | 2jsa-8 | ALA_A_69_CA | ALA_A_69_C | -4.195 | 10.123 | 7.609 | -3.405 | 9.175 | 6.703 | 1.53 | 126.29 | -50.19 |
| 407 | 2jsa-8 | ALA_A_69_C | ALA_A_69_O | -3.405 | 9.175 | 6.703 | -2.464 | 8.524 | 7.152 | 1.23 | 68.57 | -34.68 |
| 408 | 2jsa-8 | ALA_A_69_C | PRO_A_70_N | -3.405 | 9.175 | 6.703 | -3.829 | 9.127 | 5.412 | 1.36 | 161.71 | -173.54 |
| 409 | 2jsa-8 | PRO_A_70_N | PRO_A_70_CA | -3.829 | 9.127 | 5.412 | -3.172 | 8.27 | 4.44 | 1.45 | 131.99 | -52.53 |
| 410 | 2jsa-8 | PRO_A_70_CA | PRO_A_70_HA | -3.172 | 8.27 | 4.44 | -2.997 | 7.284 | 4.87 | 1.09 | 66.76 | -79.94 |
| 411 | 2jsa-8 | PRO_A_70_CA | PRO_A_70_C | -3.172 | 8.27 | 4.44 | -1.825 | 8.858 | 4.013 | 1.53 | 106.20 | 23.58 |
| 412 | 2jsa-8 | PRO_A_70_C | PRO_A_70_O | -1.825 | 8.858 | 4.013 | -1.024 | 8.181 | 3.37 | 1.23 | 121.51 | -40.20 |
| 413 | 2jsa-8 | PRO_A_70_C | LYS_A_71_N | -1.825 | 8.858 | 4.013 | -1.618 | 10.112 | 4.387 | 1.32 | 73.60 | 80.63 |
| 414 | 2jsa-8 | LYS_A_71_N | LYS_A_71_H | -1.618 | 10.112 | 4.387 | -2.322 | 10.609 | 4.914 | 1.01 | 58.55 | 144.78 |
| 415 | 2jsa-8 | LYS_A_71_N | LYS_A_71_CA | -1.618 | 10.112 | 4.387 | -0.382 | 10.799 | 4.051 | 1.45 | 103.37 | 29.07 |
| 416 | 2jsa-8 | LYS_A_71_CA | LYS_A_71_HA | -0.382 | 10.799 | 4.051 | 0.107 | 10.243 | 3.251 | 1.09 | 137.21 | -48.67 |
| 417 | 2jsa-8 | LYS_A_71_CA | LYS_A_71_C | -0.382 | 10.799 | 4.051 | 0.551 | 10.784 | 5.263 | 1.53 | 37.59 | -0.92 |
| 418 | 2jsa-8 | LYS_A_71_C | LYS_A_71_O | 0.551 | 10.784 | 5.263 | 1.761 | 10.952 | 5.121 | 1.23 | 96.63 | 7.90 |
| 419 | 2jsa-8 | LYS_A_71_C | ASP_A_72_N | 0.551 | 10.784 | 5.263 | -0.047 | 10.581 | 6.427 | 1.32 | 28.48 | -161.25 |
| 420 | 2jsa-8 | ASP_A_72_N | ASP_A_72_H | -0.047 | 10.581 | 6.427 | -1.047 | 10.451 | 6.469 | 1.01 | 87.62 | -172.59 |
| 421 | 2jsa-8 | ASP_A_72_N | ASP_A_72_CA | -0.047 | 10.581 | 6.427 | 0.715 | 10.542 | 7.663 | 1.45 | 31.69 | -2.93 |
| 422 | 2jsa-8 | ASP_A_72_CA | ASP_A_72_HA | 0.715 | 10.542 | 7.663 | 1.475 | 11.323 | 7.637 | 1.09 | 91.37 | 45.78 |
| 423 | 2jsa-8 | ASP_A_72_CA | ASP_A_72_C | 0.715 | 10.542 | 7.663 | 1.382 | 9.172 | 7.804 | 1.53 | 84.71 | -64.04 |
| 424 | 2jsa-8 | ASP_A_72_C | ASP_A_72_O | 1.382 | 9.172 | 7.804 | 2.439 | 9.053 | 8.419 | 1.23 | 59.96 | -6.42 |
| 425 | 2jsa-8 | ASP_A_72_C | VAL_A_73_N | 1.382 | 9.172 | 7.804 | 0.735 | 8.173 | 7.223 | 1.32 | 116.02 | -122.93 |
| 426 | 2jsa-8 | VAL_A_73_N | VAL_A_73_H | 0.735 | 8.173 | 7.223 | -0.131 | 8.345 | 6.734 | 1.01 | 118.98 | 168.77 |
| 427 | 2jsa-8 | VAL_A_73_N | VAL_A_73_CA | 0.735 | 8.173 | 7.223 | 1.251 | 6.816 | 7.275 | 1.45 | 87.95 | -69.18 |
| 428 | 2jsa-8 | VAL_A_73_CA | VAL_A_73_HA | 1.251 | 6.816 | 7.275 | 1.48 | 6.574 | 8.313 | 1.09 | 17.80 | -46.58 |
| 429 | 2jsa-8 | VAL_A_73_CA | VAL_A_73_C | 1.251 | 6.816 | 7.275 | 2.551 | 6.738 | 6.471 | 1.53 | 121.69 | -3.43 |
| 430 | 2jsa-8 | VAL_A_73_C | VAL_A_73_O | 2.551 | 6.738 | 6.471 | 3.549 | 6.199 | 6.948 | 1.23 | 67.19 | -28.37 |
| 431 | 2jsa-8 | VAL_A_73_C | CYS_A_74_N | 2.551 | 6.738 | 6.471 | 2.498 | 7.284 | 5.265 | 1.32 | 155.54 | 95.54 |
| 432 | 2jsa-8 | CYS_A_74_N | CYS_A_74_H | 2.498 | 7.284 | 5.265 | 1.644 | 7.711 | 4.937 | 1.01 | 108.96 | 153.43 |
| 433 | 2jsa-8 | CYS_A_74_N | CYS_A_74_CA | 2.498 | 7.284 | 5.265 | 3.658 | 7.284 | 4.391 | 1.45 | 127.00 | 148.00 |
| 434 | 2jsa-8 | CYS_A_74_CA | CYS_A_74_HA | 3.658 | 7.284 | 4.391 | 4.128 | 6.302 | 4.445 | 1.09 | 87.16 | -64.42 |
| 435 | 2jsa-8 | CYS_A_74_CA | CYS_A_74_C | 3.658 | 7.284 | 4.391 | 4.65 | 8.326 | 4.909 | 1.53 | 70.20 | 46.41 |
| 436 | 2jsa-8 | CYS_A_74_C | CYS_A_74_O | 4.65 | 8.326 | 4.909 | 4.96 | 9.293 | 4.214 | 1.23 | 124.39 | 72.23 |
| 437 | 2jsa-8 | CYS_A_74_C | THR_A_75_N | 4.65 | 8.326 | 4.909 | 5.121 | 8.096 | 6.126 | 1.33 | 23.30 | -26.03 |
| 438 | 2jsa-8 | THR_A_75_N | THR_A_75_H | 5.121 | 8.096 | 6.126 | 4.822 | 7.28 | 6.641 | 1.01 | 59.35 | -110.12 |
| 439 | 2jsa-8 | THR_A_75_N | THR_A_75_CA | 5.121 | 8.096 | 6.126 | 6.072 | 9.003 | 6.745 | 1.45 | 64.78 | 43.64 |
| 440 | 2jsa-8 | THR_A_75_CA | THR_A_75_HA | 6.072 | 9.003 | 6.745 | 6.976 | 9.033 | 6.136 | 1.09 | 123.95 | 1.90 |
| 441 | 2jsa-8 | THR_A_75_CA | THR_A_75_C | 6.072 | 9.003 | 6.745 | 6.459 | 8.501 | 8.138 | 1.53 | 24.47 | -52.37 |
| 442 | 2jsa-8 | THR_A_75_C | THR_A_75_O | 6.459 | 8.501 | 8.138 | 7.641 | 8.388 | 8.455 | 1.23 | 75.05 | -5.46 |
| 443 | 2jsa-8 | THR_A_75_C | LYS_A_76_N | 6.459 | 8.501 | 8.138 | 5.437 | 8.214 | 8.933 | 1.33 | 53.17 | -164.31 |
| 444 | 2jsa-8 | LYS_A_76_N | LYS_A_76_H | 5.437 | 8.214 | 8.933 | 4.489 | 8.332 | 8.606 | 1.01 | 108.90 | 172.90 |
| 445 | 2jsa-8 | LYS_A_76_N | LYS_A_76_CA | 5.437 | 8.214 | 8.933 | 5.655 | 7.727 | 10.285 | 1.45 | 21.54 | -65.88 |
| 446 | 2jsa-8 | LYS_A_76_CA | LYS_A_76_HA | 5.655 | 7.727 | 10.285 | 6.253 | 8.467 | 10.818 | 1.09 | 60.74 | 51.06 |
| 447 | 2jsa-8 | LYS_A_76_CA | LYS_A_76_C | 5.655 | 7.727 | 10.285 | 6.463 | 6.429 | 10.231 | 1.53 | 92.02 | -58.10 |
| 448 | 2jsa-8 | LYS_A_76_C | LYS_A_76_O | 6.463 | 6.429 | 10.231 | 7.408 | 6.249 | 10.997 | 1.23 | 51.47 | -10.78 |
| 449 | 2jsa-8 | LYS_A_76_C | LEU_A_77_N | 6.463 | 6.429 | 10.231 | 6.062 | 5.558 | 9.316 | 1.33 | 133.66 | -114.72 |
| 450 | 2jsa-8 | LEU_A_77_N | LEU_A_77_H | 6.062 | 5.558 | 9.316 | 5.278 | 5.773 | 8.718 | 1.01 | 126.34 | 164.66 |
| 451 | 2jsa-8 | LEU_A_77_N | LEU_A_77_CA | 6.062 | 5.558 | 9.316 | 6.737 | 4.281 | 9.152 | 1.45 | 96.48 | -62.14 |
| 452 | 2jsa-8 | LEU_A_77_CA | LEU_A_77_HA | 6.737 | 4.281 | 9.152 | 7.25 | 4.047 | 10.086 | 1.09 | 31.12 | -24.52 |
| 453 | 2jsa-8 | LEU_A_77_CA | LEU_A_77_C | 6.737 | 4.281 | 9.152 | 7.799 | 4.409 | 8.058 | 1.53 | 135.64 | 6.87 |
| 454 | 2jsa-8 | LEU_A_77_C | LEU_A_77_O | 7.799 | 4.409 | 8.058 | 8.046 | 3.46 | 7.314 | 1.23 | 127.19 | -75.41 |
| 455 | 2jsa-8 | LEU_A_77_C | ASN_A_78_N | 7.799 | 4.409 | 8.058 | 8.4 | 5.588 | 7.994 | 1.32 | 92.77 | 62.99 |
| 456 | 2jsa-8 | ASN_A_78_N | ASN_A_78_H | 8.4 | 5.588 | 7.994 | 8.148 | 6.323 | 8.639 | 1.01 | 50.30 | 108.92 |
| 457 | 2jsa-8 | ASN_A_78_N | ASN_A_78_CA | 8.4 | 5.588 | 7.994 | 9.429 | 5.85 | 7.004 | 1.45 | 132.99 | 14.28 |
| 458 | 2jsa-8 | ASN_A_78_CA | ASN_A_78_HA | 9.429 | 5.85 | 7.004 | 9.556 | 6.928 | 6.899 | 1.09 | 95.53 | 83.28 |
| 459 | 2jsa-8 | ASN_A_78_CA | ASN_A_78_C | 9.429 | 5.85 | 7.004 | 9.009 | 5.243 | 5.664 | 1.53 | 151.15 | -124.68 |
| 460 | 2jsa-8 | ASN_A_78_C | ASN_A_78_O | 9.009 | 5.243 | 5.664 | 9.852 | 4.786 | 4.894 | 1.23 | 128.76 | -28.46 |
| 461 | 2jsa-8 | ASN_A_78_C | GLU_A_79_N | 9.009 | 5.243 | 5.664 | 7.706 | 5.258 | 5.426 | 1.32 | 100.35 | 179.34 |
| 462 | 2jsa-8 | GLU_A_79_N | GLU_A_79_H | 7.706 | 5.258 | 5.426 | 7.07 | 5.651 | 6.105 | 1.01 | 47.75 | 148.29 |
| 463 | 2jsa-8 | GLU_A_79_N | GLU_A_79_CA | 7.706 | 5.258 | 5.426 | 7.164 | 4.715 | 4.193 | 1.45 | 148.11 | -134.95 |
| 464 | 2jsa-8 | GLU_A_79_CA | GLU_A_79_HA | 7.164 | 4.715 | 4.193 | 7.736 | 3.823 | 3.938 | 1.09 | 103.53 | -57.33 |
| 465 | 2jsa-8 | GLU_A_79_CA | GLU_A_79_C | 7.164 | 4.715 | 4.193 | 7.317 | 5.726 | 3.054 | 1.53 | 138.08 | 81.39 |
| 466 | 2jsa-8 | GLU_A_79_C | GLU_A_79_O | 7.317 | 5.726 | 3.054 | 6.82 | 5.505 | 1.951 | 1.23 | 153.75 | -156.03 |
| 467 | 2jsa-8 | GLU_A_79_C | CYS_A_80_N | 7.317 | 5.726 | 3.054 | 8.007 | 6.815 | 3.361 | 1.33 | 76.61 | 57.64 |
| 468 | 2jsa-8 | CYS_A_80_N | CYS_A_80_H | 8.007 | 6.815 | 3.361 | 8.388 | 6.931 | 4.289 | 1.01 | 23.23 | 16.93 |
| 469 | 2jsa-8 | CYS_A_80_N | CYS_A_80_CA | 8.007 | 6.815 | 3.361 | 8.232 | 7.86 | 2.379 | 1.45 | 132.57 | 77.85 |
| 470 | 2jsa-8 | CYS_A_80_CA | CYS_A_80_HA | 8.232 | 7.86 | 2.379 | 8.547 | 7.391 | 1.447 | 1.09 | 148.78 | -56.11 |
| 471 | 2jsa-8 | CYS_A_80_CA | CYS_A_80_C | 8.232 | 7.86 | 2.379 | 9.366 | 8.756 | 2.88 | 1.53 | 70.88 | 38.31 |
| 472 | 2jsa-8 | CYS_A_80_C | CYS_A_80_O | 9.366 | 8.756 | 2.88 | 9.415 | 9.1 | 4.061 | 1.23 | 16.39 | 81.89 |
| 473 | 2jsa-8 | CYS_A_80_C | PRO_A_81_N | 9.366 | 8.756 | 2.88 | 10.274 | 9.118 | 1.935 | 1.36 | 134.03 | 21.74 |
| 474 | 2jsa-8 | PRO_A_81_N | PRO_A_81_CA | 10.274 | 9.118 | 1.935 | 11.406 | 9.967 | 2.27 | 1.45 | 76.68 | 36.87 |
| 475 | 2jsa-8 | PRO_A_81_CA | PRO_A_81_HA | 11.406 | 9.967 | 2.27 | 11.867 | 9.627 | 3.197 | 1.09 | 31.71 | -36.41 |
| 476 | 2jsa-8 | PRO_A_81_CA | PRO_A_81_C | 11.406 | 9.967 | 2.27 | 10.965 | 11.42 | 2.452 | 1.53 | 83.17 | 106.88 |
| 477 | 2jsa-8 | PRO_A_81_C | PRO_A_81_O | 10.965 | 11.42 | 2.452 | 11.775 | 12.338 | 2.333 | 1.23 | 95.55 | 48.58 |
| 1 | 2jsa-9 | ARG_A_1_N | ARG_A_1_H | 13.527 | 0.363 | -3.759 | 14.347 | 0.948 | -3.834 | 1.01 | 94.26 | 35.50 |
| 2 | 2jsa-9 | ARG_A_1_N | ARG_A_1_CA | 13.527 | 0.363 | -3.759 | 12.492 | 0.736 | -2.81 | 1.45 | 49.22 | 160.18 |
| 3 | 2jsa-9 | ARG_A_1_CA | ARG_A_1_HA | 12.492 | 0.736 | -2.81 | 11.567 | 0.907 | -3.36 | 1.09 | 120.31 | 169.53 |
| 4 | 2jsa-9 | ARG_A_1_CA | ARG_A_1_C | 12.492 | 0.736 | -2.81 | 12.882 | 2.023 | -2.079 | 1.53 | 61.47 | 73.14 |
| 5 | 2jsa-9 | ARG_A_1_C | ARG_A_1_O | 12.882 | 2.023 | -2.079 | 13.427 | 1.974 | -0.976 | 1.23 | 26.39 | -5.14 |
| 6 | 2jsa-9 | ARG_A_1_C | SER_A_2_N | 12.882 | 2.023 | -2.079 | 12.587 | 3.144 | -2.721 | 1.33 | 118.98 | 104.74 |
| 7 | 2jsa-9 | SER_A_2_N | SER_A_2_H | 12.587 | 3.144 | -2.721 | 12.137 | 3.109 | -3.625 | 1.01 | 153.47 | -175.55 |
| 8 | 2jsa-9 | SER_A_2_N | SER_A_2_CA | 12.587 | 3.144 | -2.721 | 12.9 | 4.441 | -2.145 | 1.45 | 66.65 | 76.43 |
| 9 | 2jsa-9 | SER_A_2_CA | SER_A_2_HA | 12.9 | 4.441 | -2.145 | 12.545 | 4.452 | -1.114 | 1.09 | 19.01 | 178.23 |
| 10 | 2jsa-9 | SER_A_2_CA | SER_A_2_C | 12.9 | 4.441 | -2.145 | 12.181 | 5.545 | -2.923 | 1.53 | 120.56 | 123.07 |
| 11 | 2jsa-9 | SER_A_2_C | SER_A_2_O | 12.181 | 5.545 | -2.923 | 12.77 | 6.584 | -3.22 | 1.23 | 103.96 | 60.45 |
| 12 | 2jsa-9 | SER_A_2_C | ALA_A_3_N | 12.181 | 5.545 | -2.923 | 10.919 | 5.283 | -3.231 | 1.33 | 103.44 | -168.27 |
| 13 | 2jsa-9 | ALA_A_3_N | ALA_A_3_H | 10.919 | 5.283 | -3.231 | 10.499 | 4.407 | -2.955 | 1.01 | 74.14 | -115.62 |
| 14 | 2jsa-9 | ALA_A_3_N | ALA_A_3_CA | 10.919 | 5.283 | -3.231 | 10.116 | 6.243 | -3.969 | 1.45 | 120.53 | 129.91 |
| 15 | 2jsa-9 | ALA_A_3_CA | ALA_A_3_HA | 10.116 | 6.243 | -3.969 | 9.883 | 7.082 | -3.314 | 1.09 | 53.05 | 105.52 |
| 16 | 2jsa-9 | ALA_A_3_CA | ALA_A_3_C | 10.116 | 6.243 | -3.969 | 8.803 | 5.581 | -4.395 | 1.53 | 106.16 | -153.24 |
| 17 | 2jsa-9 | ALA_A_3_C | ALA_A_3_O | 8.803 | 5.581 | -4.395 | 7.728 | 5.98 | -3.949 | 1.23 | 68.75 | 159.64 |
| 18 | 2jsa-9 | ALA_A_3_C | LEU_A_4_N | 8.803 | 5.581 | -4.395 | 8.934 | 4.579 | -5.252 | 1.32 | 130.30 | -82.55 |
| 19 | 2jsa-9 | LEU_A_4_N | LEU_A_4_H | 8.934 | 4.579 | -5.252 | 9.85 | 4.303 | -5.576 | 1.01 | 108.71 | -16.77 |
| 20 | 2jsa-9 | LEU_A_4_N | LEU_A_4_CA | 8.934 | 4.579 | -5.252 | 7.772 | 3.858 | -5.743 | 1.45 | 109.75 | -148.18 |
| 21 | 2jsa-9 | LEU_A_4_CA | LEU_A_4_HA | 7.772 | 3.858 | -5.743 | 7.1 | 4.579 | -6.208 | 1.09 | 115.26 | 132.99 |
| 22 | 2jsa-9 | LEU_A_4_CA | LEU_A_4_C | 7.772 | 3.858 | -5.743 | 7.032 | 3.227 | -4.561 | 1.53 | 39.45 | -139.55 |
| 23 | 2jsa-9 | LEU_A_4_C | LEU_A_4_O | 7.032 | 3.227 | -4.561 | 5.845 | 2.918 | -4.661 | 1.23 | 94.66 | -165.41 |

| | | | | | | | | | | | |
|---|---|---|---|---|---|---|---|---|---|---|---|
| 24 | 2jsa-9 | LEU_A_4_C | SER_A_5_N | 7.032 | 3.227 | -4.561 | 7.763 | 3.056 | -3.47 | 1.32 | 34.53 | -13.17 |
| 25 | 2jsa-9 | SER_A_5_N | SER_A_5_H | 7.763 | 3.056 | -3.47 | 8.734 | 3.335 | -3.459 | 1.01 | 89.38 | 16.03 |
| 26 | 2jsa-9 | SER_A_5_N | SER_A_5_CA | 7.763 | 3.056 | -3.47 | 7.19 | 2.468 | -2.271 | 1.45 | 34.40 | -134.26 |
| 27 | 2jsa-9 | SER_A_5_CA | SER_A_5_HA | 7.19 | 2.468 | -2.271 | 6.656 | 1.562 | -2.556 | 1.09 | 105.16 | -120.52 |
| 28 | 2jsa-9 | SER_A_5_CA | SER_A_5_C | 7.19 | 2.468 | -2.271 | 6.198 | 3.442 | -1.633 | 1.53 | 65.35 | 135.52 |
| 29 | 2jsa-9 | SER_A_5_C | SER_A_5_O | 6.198 | 3.442 | -1.633 | 5.173 | 3.026 | -1.094 | 1.23 | 64.02 | -157.91 |
| 30 | 2jsa-9 | SER_A_5_C | CYS_A_6_N | 6.198 | 3.442 | -1.633 | 6.537 | 4.721 | -1.715 | 1.33 | 93.55 | 75.16 |
| 31 | 2jsa-9 | CYS_A_6_N | CYS_A_6_H | 6.537 | 4.721 | -1.715 | 7.395 | 4.992 | -2.174 | 1.01 | 117.03 | 17.53 |
| 32 | 2jsa-9 | CYS_A_6_N | CYS_A_6_CA | 6.537 | 4.721 | -1.715 | 5.688 | 5.756 | -1.152 | 1.45 | 67.19 | 129.36 |
| 33 | 2jsa-9 | CYS_A_6_CA | CYS_A_6_HA | 5.688 | 5.756 | -1.152 | 5.559 | 5.553 | -0.089 | 1.09 | 12.75 | -122.43 |
| 34 | 2jsa-9 | CYS_A_6_CA | CYS_A_6_C | 5.688 | 5.756 | -1.152 | 4.32 | 5.679 | -1.832 | 1.53 | 116.39 | -176.78 |
| 35 | 2jsa-9 | CYS_A_6_C | CYS_A_6_O | 4.32 | 5.679 | -1.832 | 3.301 | 5.502 | -1.167 | 1.23 | 57.26 | -170.15 |
| 36 | 2jsa-9 | CYS_A_6_C | GLN_A_7_N | 4.32 | 5.679 | -1.832 | 4.341 | 5.816 | -3.15 | 1.33 | 174.00 | 81.29 |
| 37 | 2jsa-9 | GLN_A_7_N | GLN_A_7_H | 4.341 | 5.816 | -3.15 | 5.217 | 5.959 | -3.632 | 1.01 | 118.50 | 9.27 |
| 38 | 2jsa-9 | GLN_A_7_N | GLN_A_7_CA | 4.341 | 5.816 | -3.15 | 3.115 | 5.765 | -3.928 | 1.45 | 122.38 | -177.62 |
| 39 | 2jsa-9 | GLN_A_7_CA | GLN_A_7_HA | 3.115 | 5.765 | -3.928 | 2.558 | 6.683 | -3.74 | 1.09 | 80.07 | 121.25 |
| 40 | 2jsa-9 | GLN_A_7_CA | GLN_A_7_C | 3.115 | 5.765 | -3.928 | 2.255 | 4.579 | -3.484 | 1.53 | 73.14 | -125.95 |
| 41 | 2jsa-9 | GLN_A_7_C | GLN_A_7_O | 2.255 | 4.579 | -3.484 | 1.067 | 4.739 | -3.207 | 1.23 | 76.99 | 172.33 |
| 42 | 2jsa-9 | GLN_A_7_C | MET_A_8_N | 2.255 | 4.579 | -3.484 | 2.888 | 3.416 | -3.431 | 1.33 | 87.71 | -61.44 |
| 43 | 2jsa-9 | MET_A_8_N | MET_A_8_H | 2.888 | 3.416 | -3.431 | 3.867 | 3.358 | -3.675 | 1.01 | 103.97 | -3.39 |
| 44 | 2jsa-9 | MET_A_8_N | MET_A_8_CA | 2.888 | 3.416 | -3.431 | 2.196 | 2.205 | -3.025 | 1.45 | 73.77 | -119.74 |
| 45 | 2jsa-9 | MET_A_8_CA | MET_A_8_HA | 2.196 | 2.205 | -3.025 | 1.526 | 1.899 | -3.828 | 1.09 | 137.47 | -155.45 |
| 46 | 2jsa-9 | MET_A_8_CA | MET_A_8_C | 2.196 | 2.205 | -3.025 | 1.375 | 2.442 | -1.756 | 1.53 | 33.96 | 163.90 |
| 47 | 2jsa-9 | MET_A_8_C | MET_A_8_O | 1.375 | 2.442 | -1.756 | 0.267 | 1.925 | -1.625 | 1.23 | 83.88 | -154.99 |
| 48 | 2jsa-9 | MET_A_8_C | CYS_A_9_N | 1.375 | 2.442 | -1.756 | 1.95 | 3.223 | -0.855 | 1.32 | 47.11 | 53.64 |
| 49 | 2jsa-9 | CYS_A_9_N | CYS_A_9_H | 1.95 | 3.223 | -0.855 | 2.865 | 3.614 | -1.031 | 1.01 | 100.03 | 23.14 |
| 50 | 2jsa-9 | CYS_A_9_N | CYS_A_9_CA | 1.95 | 3.223 | -0.855 | 1.285 | 3.536 | 0.4 | 1.45 | 30.35 | 154.79 |
| 51 | 2jsa-9 | CYS_A_9_CA | CYS_A_9_HA | 1.285 | 3.536 | 0.4 | 1.075 | 2.601 | 0.919 | 1.09 | 61.56 | -102.66 |
| 52 | 2jsa-9 | CYS_A_9_CA | CYS_A_9_C | 1.285 | 3.536 | 0.4 | -0.043 | 4.224 | 0.08 | 1.53 | 102.08 | 152.61 |
| 53 | 2jsa-9 | CYS_A_9_C | CYS_A_9_O | -0.043 | 4.224 | 0.08 | -1.035 | 4.023 | 0.781 | 1.23 | 55.29 | -168.55 |
| 54 | 2jsa-9 | CYS_A_9_C | GLU_A_10_N | -0.043 | 4.224 | 0.08 | -0.02 | 5.022 | -0.977 | 1.32 | 142.94 | 88.35 |
| 55 | 2jsa-9 | GLU_A_10_N | GLU_A_10_H | -0.02 | 5.022 | -0.977 | 0.832 | 5.14 | -1.507 | 1.01 | 121.64 | 7.89 |
| 56 | 2jsa-9 | GLU_A_10_N | GLU_A_10_CA | -0.02 | 5.022 | -0.977 | -1.211 | 5.742 | -1.398 | 1.45 | 106.83 | 148.85 |
| 57 | 2jsa-9 | GLU_A_10_CA | GLU_A_10_HA | -1.211 | 5.742 | -1.398 | -1.726 | 6.093 | -0.504 | 1.09 | 34.88 | 145.72 |
| 58 | 2jsa-9 | GLU_A_10_CA | GLU_A_10_C | -1.211 | 5.742 | -1.398 | -2.154 | 4.809 | -2.159 | 1.53 | 119.84 | -135.31 |
| 59 | 2jsa-9 | GLU_A_10_C | GLU_A_10_O | -2.154 | 4.809 | -2.159 | -3.275 | 5.191 | -2.493 | 1.23 | 105.75 | 161.18 |
| 60 | 2jsa-9 | GLU_A_10_C | LEU_A_11_N | -2.154 | 4.809 | -2.159 | -1.664 | 3.604 | -2.414 | 1.33 | 101.09 | -67.87 |
| 61 | 2jsa-9 | LEU_A_11_N | LEU_A_11_H | -1.664 | 3.604 | -2.414 | -0.732 | 3.359 | -2.11 | 1.01 | 72.49 | -14.73 |
| 62 | 2jsa-9 | LEU_A_11_N | LEU_A_11_CA | -1.664 | 3.604 | -2.414 | -2.449 | 2.614 | -3.132 | 1.45 | 119.61 | -128.41 |
| 63 | 2jsa-9 | LEU_A_11_CA | LEU_A_11_HA | -2.449 | 2.614 | -3.132 | -3.148 | 3.143 | -3.78 | 1.09 | 126.47 | 142.88 |
| 64 | 2jsa-9 | LEU_A_11_CA | LEU_A_11_C | -2.449 | 2.614 | -3.132 | -3.269 | 1.797 | -2.132 | 1.53 | 49.18 | -135.11 |
| 65 | 2jsa-9 | LEU_A_11_C | LEU_A_11_O | -3.269 | 1.797 | -2.132 | -4.471 | 1.61 | -2.315 | 1.23 | 98.56 | -171.16 |
| 66 | 2jsa-9 | LEU_A_11_C | VAL_A_12_N | -3.269 | 1.797 | -2.132 | -2.586 | 1.333 | -1.095 | 1.33 | 38.53 | -34.19 |
| 67 | 2jsa-9 | VAL_A_12_N | VAL_A_12_H | -2.586 | 1.333 | -1.095 | -1.598 | 1.527 | -1.008 | 1.01 | 85.06 | 11.11 |
| 68 | 2jsa-9 | VAL_A_12_N | VAL_A_12_CA | -2.586 | 1.333 | -1.095 | -3.236 | 0.54 | -0.065 | 1.45 | 44.87 | -129.34 |
| 69 | 2jsa-9 | VAL_A_12_CA | VAL_A_12_HA | -3.236 | 0.54 | -0.065 | -3.744 | -0.297 | -0.544 | 1.09 | 116.07 | -121.25 |
| 70 | 2jsa-9 | VAL_A_12_CA | VAL_A_12_C | -3.236 | 0.54 | -0.065 | -4.29 | 1.395 | 0.642 | 1.53 | 62.48 | 140.95 |
| 71 | 2jsa-9 | VAL_A_12_C | VAL_A_12_O | -4.29 | 1.395 | 0.642 | -5.445 | 0.987 | 0.765 | 1.23 | 84.27 | -160.54 |
| 72 | 2jsa-9 | VAL_A_12_C | VAL_A_13_N | -4.29 | 1.395 | 0.642 | -3.856 | 2.565 | 1.086 | 1.32 | 70.41 | 69.65 |
| 73 | 2jsa-9 | VAL_A_13_N | VAL_A_13_H | -3.856 | 2.565 | 1.086 | -2.892 | 2.836 | 0.949 | 1.01 | 97.79 | 15.70 |
| 74 | 2jsa-9 | VAL_A_13_N | VAL_A_13_CA | -3.856 | 2.565 | 1.086 | -4.749 | 3.481 | 1.776 | 1.45 | 61.66 | 134.27 |
| 75 | 2jsa-9 | VAL_A_13_CA | VAL_A_13_HA | -4.749 | 3.481 | 1.776 | -5.136 | 2.981 | 2.664 | 1.09 | 35.45 | -127.74 |
| 76 | 2jsa-9 | VAL_A_13_CA | VAL_A_13_C | -4.749 | 3.481 | 1.776 | -5.932 | 3.818 | 0.866 | 1.53 | 126.49 | 164.10 |
| 77 | 2jsa-9 | VAL_A_13_C | VAL_A_13_O | -5.932 | 3.818 | 0.866 | -7.083 | 3.584 | 1.227 | 1.23 | 72.92 | -168.51 |
| 78 | 2jsa-9 | VAL_A_13_C | LYS_A_14_N | -5.932 | 3.818 | 0.866 | -5.605 | 4.362 | -0.298 | 1.33 | 151.40 | 58.99 |
| 79 | 2jsa-9 | LYS_A_14_N | LYS_A_14_H | -5.605 | 4.362 | -0.298 | -4.635 | 4.526 | -0.529 | 1.01 | 103.21 | 9.60 |
| 80 | 2jsa-9 | LYS_A_14_N | LYS_A_14_CA | -5.605 | 4.362 | -0.298 | -6.627 | 4.733 | -1.262 | 1.45 | 131.56 | 160.05 |
| 81 | 2jsa-9 | LYS_A_14_CA | LYS_A_14_HA | -6.627 | 4.733 | -1.262 | -7.139 | 5.619 | -0.887 | 1.09 | 69.87 | 120.02 |
| 82 | 2jsa-9 | LYS_A_14_CA | LYS_A_14_C | -6.627 | 4.733 | -1.262 | -7.658 | 3.609 | -1.365 | 1.53 | 93.86 | -132.53 |
| 83 | 2jsa-9 | LYS_A_14_C | LYS_A_14_O | -7.658 | 3.609 | -1.365 | -8.839 | 3.864 | -1.598 | 1.23 | 100.92 | 167.82 |
| 84 | 2jsa-9 | LYS_A_14_C | LYS_A_15_N | -7.658 | 3.609 | -1.365 | -7.176 | 2.388 | -1.185 | 1.32 | 82.19 | -68.46 |
| 85 | 2jsa-9 | LYS_A_15_N | LYS_A_15_H | -7.176 | 2.388 | -1.185 | -6.193 | 2.252 | -0.996 | 1.01 | 79.22 | -7.88 |
| 86 | 2jsa-9 | LYS_A_15_N | LYS_A_15_CA | -7.176 | 2.388 | -1.185 | -8.042 | 1.223 | -1.254 | 1.45 | 92.72 | -126.63 |
| 87 | 2jsa-9 | LYS_A_15_CA | LYS_A_15_HA | -8.042 | 1.223 | -1.254 | -8.766 | 1.388 | -2.052 | 1.09 | 137.06 | 167.16 |
| 88 | 2jsa-9 | LYS_A_15_CA | LYS_A_15_C | -8.042 | 1.223 | -1.254 | -8.818 | 1.091 | 0.058 | 1.53 | 30.96 | -170.35 |
| 89 | 2jsa-9 | LYS_A_15_C | LYS_A_15_O | -8.818 | 1.091 | 0.058 | -10.01 | 0.789 | 0.05 | 1.23 | 90.37 | -165.78 |
| 90 | 2jsa-9 | LYS_A_15_C | TYR_A_16_N | -8.818 | 1.091 | 0.058 | -8.109 | 1.324 | 1.153 | 1.33 | 34.28 | 18.19 |
| 91 | 2jsa-9 | TYR_A_16_N | TYR_A_16_H | -8.109 | 1.324 | 1.153 | -7.131 | 1.567 | 1.084 | 1.01 | 93.92 | 13.95 |
| 92 | 2jsa-9 | TYR_A_16_N | TYR_A_16_CA | -8.109 | 1.324 | 1.153 | -8.717 | 1.235 | 2.47 | 1.45 | 25.01 | -171.67 |
| 93 | 2jsa-9 | TYR_A_16_CA | TYR_A_16_HA | -8.717 | 1.235 | 2.47 | -9.296 | 0.316 | 2.557 | 1.09 | 85.42 | -122.21 |
| 94 | 2jsa-9 | TYR_A_16_CA | TYR_A_16_C | -8.717 | 1.235 | 2.47 | -9.616 | 2.444 | 2.742 | 1.53 | 79.77 | 126.63 |
| 95 | 2jsa-9 | TYR_A_16_C | TYR_A_16_O | -9.616 | 2.444 | 2.742 | -10.833 | 2.305 | 2.852 | 1.23 | 84.87 | -173.48 |
| 96 | 2jsa-9 | TYR_A_16_C | GLU_A_17_N | -9.616 | 2.444 | 2.742 | -8.981 | 3.603 | 2.843 | 1.33 | 85.63 | 61.28 |
| 97 | 2jsa-9 | GLU_A_17_N | GLU_A_17_H | -8.981 | 3.603 | 2.843 | -7.977 | 3.643 | 2.741 | 1.01 | 95.80 | 2.28 |
| 98 | 2jsa-9 | GLU_A_17_N | GLU_A_17_CA | -8.981 | 3.603 | 2.843 | -9.708 | 4.835 | 3.1 | 1.45 | 79.82 | 120.54 |
| 99 | 2jsa-9 | GLU_A_17_CA | GLU_A_17_HA | -9.708 | 4.835 | 3.1 | -9.959 | 4.864 | 4.16 | 1.09 | 13.41 | 173.41 |
| 100 | 2jsa-9 | GLU_A_17_CA | GLU_A_17_C | -9.708 | 4.835 | 3.1 | -11.009 | 4.86 | 2.296 | 1.53 | 121.71 | 178.90 |
| 101 | 2jsa-9 | GLU_A_17_C | GLU_A_17_O | -11.009 | 4.86 | 2.296 | -12.049 | 5.274 | 2.807 | 1.23 | 65.46 | 158.29 |
| 102 | 2jsa-9 | GLU_A_17_C | GLY_A_18_N | -11.009 | 4.86 | 2.296 | -10.91 | 4.413 | 1.053 | 1.32 | 159.78 | -77.51 |
| 103 | 2jsa-9 | GLY_A_18_N | GLY_A_18_H | -10.91 | 4.413 | 1.053 | -10.021 | 4.088 | 0.699 | 1.01 | 110.51 | -20.08 |
| 104 | 2jsa-9 | GLY_A_18_N | GLY_A_18_CA | -10.91 | 4.413 | 1.053 | -12.066 | 4.38 | 0.175 | 1.45 | 127.21 | -178.36 |
| 105 | 2jsa-9 | GLY_A_18_CA | GLY_A_18_C | -12.066 | 4.38 | 0.175 | -12.418 | 2.942 | -0.215 | 1.53 | 104.76 | -103.75 |
| 106 | 2jsa-9 | GLY_A_18_C | GLY_A_18_O | -12.418 | 2.942 | -0.215 | -12.722 | 2.666 | -1.374 | 1.23 | 160.49 | -137.76 |
| 107 | 2jsa-9 | GLY_A_18_C | SER_A_19_N | -12.418 | 2.942 | -0.215 | -12.364 | 2.065 | 0.777 | 1.33 | 41.53 | -86.48 |
| 108 | 2jsa-9 | SER_A_19_N | SER_A_19_H | -12.364 | 2.065 | 0.777 | -12.106 | 2.363 | 1.707 | 1.01 | 22.97 | 49.11 |
| 109 | 2jsa-9 | SER_A_19_N | SER_A_19_CA | -12.364 | 2.065 | 0.777 | -12.673 | 0.662 | 0.552 | 1.45 | 98.90 | -102.42 |
| 110 | 2jsa-9 | SER_A_19_CA | SER_A_19_HA | -12.673 | 0.662 | 0.552 | -12.156 | 0.339 | -0.351 | 1.09 | 145.98 | -32.00 |
| 111 | 2jsa-9 | SER_A_19_CA | SER_A_19_C | -12.673 | 0.662 | 0.552 | -14.179 | 0.485 | 0.346 | 1.53 | 97.74 | -173.30 |
| 112 | 2jsa-9 | SER_A_19_C | SER_A_19_O | -14.179 | 0.485 | 0.346 | -14.606 | -0.134 | -0.628 | 1.23 | 142.33 | -124.60 |
| 113 | 2jsa-9 | SER_A_19_C | ALA_A_20_N | -14.179 | 0.485 | 0.346 | -14.941 | 1.039 | 1.277 | 1.32 | 45.34 | 143.98 |
| 114 | 2jsa-9 | ALA_A_20_N | ALA_A_20_H | -14.941 | 1.039 | 1.277 | -14.52 | 1.533 | 2.051 | 1.01 | 39.98 | 49.56 |
| 115 | 2jsa-9 | ALA_A_20_N | ALA_A_20_CA | -14.941 | 1.039 | 1.277 | -16.389 | 0.949 | 1.209 | 1.45 | 92.68 | -176.44 |
| 116 | 2jsa-9 | ALA_A_20_CA | ALA_A_20_HA | -16.389 | 0.949 | 1.209 | -16.818 | 1.774 | 1.778 | 1.09 | 58.54 | 117.47 |
| 117 | 2jsa-9 | ALA_A_20_CA | ALA_A_20_C | -16.389 | 0.949 | 1.209 | -16.845 | -0.366 | 1.844 | 1.53 | 65.48 | -109.12 |
| 118 | 2jsa-9 | ALA_A_20_C | ALA_A_20_O | -16.845 | -0.366 | 1.844 | -18.031 | -0.55 | 2.115 | 1.23 | 77.28 | -171.18 |
| 119 | 2jsa-9 | ALA_A_20_C | ASP_A_21_N | -16.845 | -0.366 | 1.844 | -15.879 | -1.247 | 2.062 | 1.33 | 80.53 | -42.37 |

| | | | | | | | | | | | | |
|---|---|---|---|---|---|---|---|---|---|---|---|---|
| 120 | 2jsa-9 | ASP_A_21_N | ASP_A_21_H | -15.879 | -1.247 | 2.062 | -14.925 | -1.029 | 1.812 | 1.01 | 104.33 | 12.87 |
| 121 | 2jsa-9 | ASP_A_21_N | ASP_A_21_CA | -15.879 | -1.247 | 2.062 | -16.167 | -2.539 | 2.661 | 1.45 | 65.65 | -102.57 |
| 122 | 2jsa-9 | ASP_A_21_CA | ASP_A_21_HA | -16.167 | -2.539 | 2.661 | -17.244 | -2.639 | 2.791 | 1.09 | 83.15 | -174.70 |
| 123 | 2jsa-9 | ASP_A_21_CA | ASP_A_21_C | -16.167 | -2.539 | 2.661 | -15.468 | -2.634 | 4.018 | 1.53 | 27.47 | -7.74 |
| 124 | 2jsa-9 | ASP_A_21_C | ASP_A_21_O | -15.468 | -2.634 | 4.018 | -15.94 | -3.331 | 4.916 | 1.23 | 43.15 | -124.11 |
| 125 | 2jsa-9 | ASP_A_21_C | LYS_A_22_N | -15.468 | -2.634 | 4.018 | -14.354 | -1.924 | 4.126 | 1.33 | 85.33 | 32.51 |
| 126 | 2jsa-9 | LYS_A_22_N | LYS_A_22_H | -14.354 | -1.924 | 4.126 | -14.023 | -1.374 | 3.346 | 1.01 | 140.55 | 58.96 |
| 127 | 2jsa-9 | LYS_A_22_N | LYS_A_22_CA | -14.354 | -1.924 | 4.126 | -13.586 | -1.92 | 5.36 | 1.45 | 31.90 | 0.30 |
| 128 | 2jsa-9 | LYS_A_22_CA | LYS_A_22_HA | -13.586 | -1.92 | 5.36 | -12.749 | -1.232 | 5.235 | 1.09 | 96.58 | 39.42 |
| 129 | 2jsa-9 | LYS_A_22_CA | LYS_A_22_C | -13.586 | -1.92 | 5.36 | -13.008 | -3.316 | 5.599 | 1.53 | 81.01 | -67.51 |
| 130 | 2jsa-9 | LYS_A_22_C | LYS_A_22_O | -13.008 | -3.316 | 5.599 | -13.332 | -3.963 | 6.594 | 1.23 | 36.03 | -116.60 |
| 131 | 2jsa-9 | LYS_A_22_C | ASP_A_23_N | -13.008 | -3.316 | 5.599 | -12.163 | -3.739 | 4.672 | 1.32 | 134.45 | -26.59 |
| 132 | 2jsa-9 | ASP_A_23_N | ASP_A_23_H | -12.163 | -3.739 | 4.672 | -11.943 | -3.153 | 3.879 | 1.01 | 141.72 | 69.42 |
| 133 | 2jsa-9 | ASP_A_23_N | ASP_A_23_CA | -12.163 | -3.739 | 4.672 | -11.536 | -5.046 | 4.77 | 1.45 | 86.13 | -64.37 |
| 134 | 2jsa-9 | ASP_A_23_CA | ASP_A_23_HA | -11.536 | -5.046 | 4.77 | -11.521 | -5.356 | 5.815 | 1.09 | 16.54 | -87.23 |
| 135 | 2jsa-9 | ASP_A_23_CA | ASP_A_23_C | -11.536 | -5.046 | 4.77 | -10.106 | -4.964 | 4.233 | 1.53 | 110.55 | 3.28 |
| 136 | 2jsa-9 | ASP_A_23_C | ASP_A_23_O | -10.106 | -4.964 | 4.233 | -9.898 | -4.702 | 3.049 | 1.23 | 164.22 | 51.55 |
| 137 | 2jsa-9 | ASP_A_23_C | ALA_A_24_N | -10.106 | -4.964 | 4.233 | -9.157 | -5.192 | 5.129 | 1.32 | 47.45 | -13.51 |
| 138 | 2jsa-9 | ALA_A_24_N | ALA_A_24_H | -9.157 | -5.192 | 5.129 | -9.401 | -5.4 | 6.087 | 1.01 | 18.50 | -139.55 |
| 139 | 2jsa-9 | ALA_A_24_N | ALA_A_24_CA | -9.157 | -5.192 | 5.129 | -7.753 | -5.148 | 4.76 | 1.45 | 104.72 | 1.80 |
| 140 | 2jsa-9 | ALA_A_24_CA | ALA_A_24_HA | -7.753 | -5.148 | 4.76 | -7.541 | -4.181 | 4.303 | 1.09 | 114.78 | 77.63 |
| 141 | 2jsa-9 | ALA_A_24_CA | ALA_A_24_C | -7.753 | -5.148 | 4.76 | -7.464 | -6.243 | 3.731 | 1.53 | 132.26 | -75.22 |
| 142 | 2jsa-9 | ALA_A_24_C | ALA_A_24_O | -7.464 | -6.243 | 3.731 | -6.391 | -6.269 | 3.132 | 1.23 | 119.17 | -1.39 |
| 143 | 2jsa-9 | ALA_A_24_C | ASN_A_25_N | -7.464 | -6.243 | 3.731 | -8.442 | -7.121 | 3.56 | 1.33 | 97.41 | -138.08 |
| 144 | 2jsa-9 | ASN_A_25_N | ASN_A_25_H | -8.442 | -7.121 | 3.56 | -9.298 | -7.038 | 4.089 | 1.01 | 58.40 | 174.46 |
| 145 | 2jsa-9 | ASN_A_25_N | ASN_A_25_CA | -8.442 | -7.121 | 3.56 | -8.307 | -8.216 | 2.614 | 1.45 | 130.61 | -82.97 |
| 146 | 2jsa-9 | ASN_A_25_CA | ASN_A_25_HA | -8.307 | -8.216 | 2.614 | -7.293 | -8.613 | 2.67 | 1.09 | 87.06 | -21.38 |
| 147 | 2jsa-9 | ASN_A_25_CA | ASN_A_25_C | -8.307 | -8.216 | 2.614 | -8.587 | -7.701 | 1.201 | 1.53 | 157.47 | 118.53 |
| 148 | 2jsa-9 | ASN_A_25_C | ASN_A_25_O | -8.587 | -7.701 | 1.201 | -7.882 | -8.054 | 0.256 | 1.23 | 140.16 | -26.60 |
| 149 | 2jsa-9 | ASN_A_25_C | VAL_A_26_N | -8.587 | -7.701 | 1.201 | -9.617 | -6.874 | 1.099 | 1.32 | 94.42 | 141.24 |
| 150 | 2jsa-9 | VAL_A_26_N | VAL_A_26_H | -9.617 | -6.874 | 1.099 | -10.152 | -6.626 | 1.919 | 1.01 | 35.72 | 155.13 |
| 151 | 2jsa-9 | VAL_A_26_N | VAL_A_26_CA | -9.617 | -6.874 | 1.099 | -9.999 | -6.307 | -0.183 | 1.45 | 151.93 | 123.97 |
| 152 | 2jsa-9 | VAL_A_26_CA | VAL_A_26_HA | -9.999 | -6.307 | -0.183 | -9.801 | -7.046 | -0.959 | 1.09 | 135.41 | -75.00 |
| 153 | 2jsa-9 | VAL_A_26_CA | VAL_A_26_C | -9.999 | -6.307 | -0.183 | -9.136 | -5.076 | -0.469 | 1.53 | 100.77 | 54.97 |
| 154 | 2jsa-9 | VAL_A_26_C | VAL_A_26_O | -9.136 | -5.076 | -0.469 | -9.187 | -4.518 | -1.563 | 1.23 | 152.88 | 95.22 |
| 155 | 2jsa-9 | VAL_A_26_C | ILE_A_27_N | -9.136 | -5.076 | -0.469 | -8.363 | -4.689 | 0.535 | 1.32 | 40.73 | 26.59 |
| 156 | 2jsa-9 | ILE_A_27_N | ILE_A_27_H | -8.363 | -4.689 | 0.535 | -8.371 | -5.197 | 1.408 | 1.01 | 30.20 | -90.90 |
| 157 | 2jsa-9 | ILE_A_27_N | ILE_A_27_CA | -8.363 | -4.689 | 0.535 | -7.49 | -3.535 | 0.406 | 1.45 | 95.09 | 52.89 |
| 158 | 2jsa-9 | ILE_A_27_CA | ILE_A_27_HA | -7.49 | -3.535 | 0.406 | -7.909 | -2.878 | -0.356 | 1.09 | 134.36 | 122.53 |
| 159 | 2jsa-9 | ILE_A_27_CA | ILE_A_27_C | -7.49 | -3.535 | 0.406 | -6.115 | -3.993 | -0.082 | 1.53 | 108.61 | -18.42 |
| 160 | 2jsa-9 | ILE_A_27_C | ILE_A_27_O | -6.115 | -3.993 | -0.082 | -5.697 | -3.646 | -1.186 | 1.23 | 153.80 | 39.70 |
| 161 | 2jsa-9 | ILE_A_27_C | LYS_A_28_N | -6.115 | -3.993 | -0.082 | -5.449 | -4.765 | 0.764 | 1.32 | 50.32 | -49.22 |
| 162 | 2jsa-9 | LYS_A_28_N | LYS_A_28_H | -5.449 | -4.765 | 0.764 | -5.854 | -5.01 | 1.656 | 1.01 | 27.95 | -148.83 |
| 163 | 2jsa-9 | LYS_A_28_N | LYS_A_28_CA | -5.449 | -4.765 | 0.764 | -4.129 | -5.274 | 0.433 | 1.45 | 103.17 | -21.09 |
| 164 | 2jsa-9 | LYS_A_28_CA | LYS_A_28_HA | -4.129 | -5.274 | 0.433 | -3.441 | -4.429 | 0.393 | 1.09 | 92.10 | 50.85 |
| 165 | 2jsa-9 | LYS_A_28_CA | LYS_A_28_C | -4.129 | -5.274 | 0.433 | -4.167 | -5.91 | -0.959 | 1.53 | 155.41 | -93.42 |
| 166 | 2jsa-9 | LYS_A_28_C | LYS_A_28_O | -4.167 | -5.91 | -0.959 | -3.234 | -5.749 | -1.744 | 1.23 | 129.66 | 9.79 |
| 167 | 2jsa-9 | LYS_A_28_C | LYS_A_29_N | -4.167 | -5.91 | -0.959 | -5.255 | -6.618 | -1.221 | 1.32 | 101.41 | -146.95 |
| 168 | 2jsa-9 | LYS_A_29_N | LYS_A_29_H | -5.255 | -6.618 | -1.221 | -5.983 | -6.707 | -0.527 | 1.01 | 46.58 | -173.03 |
| 169 | 2jsa-9 | LYS_A_29_N | LYS_A_29_CA | -5.255 | -6.618 | -1.221 | -5.427 | -7.279 | -2.504 | 1.45 | 151.97 | -104.59 |
| 170 | 2jsa-9 | LYS_A_29_CA | LYS_A_29_HA | -5.427 | -7.279 | -2.504 | -4.504 | -7.812 | -2.735 | 1.09 | 102.23 | -30.00 |
| 171 | 2jsa-9 | LYS_A_29_CA | LYS_A_29_C | -5.427 | -7.279 | -2.504 | -5.639 | -6.224 | -3.592 | 1.53 | 135.32 | 101.36 |
| 172 | 2jsa-9 | LYS_A_29_C | LYS_A_29_O | -5.639 | -6.224 | -3.592 | -4.899 | -6.182 | -4.573 | 1.23 | 142.93 | 3.25 |
| 173 | 2jsa-9 | LYS_A_29_C | ASP_A_30_N | -5.639 | -6.224 | -3.592 | -6.654 | -5.397 | -3.38 | 1.33 | 80.80 | 140.83 |
| 174 | 2jsa-9 | ASP_A_30_N | ASP_A_30_H | -6.654 | -5.397 | -3.38 | -7.222 | -5.49 | -2.55 | 1.01 | 34.74 | -170.70 |
| 175 | 2jsa-9 | ASP_A_30_N | ASP_A_30_CA | -6.654 | -5.397 | -3.38 | -6.972 | -4.345 | -4.33 | 1.45 | 130.84 | 106.82 |
| 176 | 2jsa-9 | ASP_A_30_CA | ASP_A_30_HA | -6.972 | -4.345 | -4.33 | -7.497 | -4.78 | -5.18 | 1.09 | 141.27 | -140.36 |
| 177 | 2jsa-9 | ASP_A_30_CA | ASP_A_30_C | -6.972 | -4.345 | -4.33 | -5.676 | -3.681 | -4.804 | 1.53 | 108.03 | 27.13 |
| 178 | 2jsa-9 | ASP_A_30_C | ASP_A_30_O | -5.676 | -3.681 | -4.804 | -5.426 | -3.585 | -6.004 | 1.23 | 167.42 | 21.01 |
| 179 | 2jsa-9 | ASP_A_30_C | PHE_A_31_N | -5.676 | -3.681 | -4.804 | -4.887 | -3.241 | -3.834 | 1.33 | 42.96 | 29.15 |
| 180 | 2jsa-9 | PHE_A_31_N | PHE_A_31_H | -4.887 | -3.241 | -3.834 | -5.156 | -3.355 | -2.867 | 1.01 | 16.81 | -157.03 |
| 181 | 2jsa-9 | PHE_A_31_N | PHE_A_31_CA | -4.887 | -3.241 | -3.834 | -3.624 | -2.589 | -4.137 | 1.45 | 102.03 | 27.30 |
| 182 | 2jsa-9 | PHE_A_31_CA | PHE_A_31_HA | -3.624 | -2.589 | -4.137 | -3.803 | -1.557 | -4.441 | 1.09 | 106.18 | 99.84 |
| 183 | 2jsa-9 | PHE_A_31_CA | PHE_A_31_C | -3.624 | -2.589 | -4.137 | -2.877 | -3.33 | -5.248 | 1.53 | 136.56 | -44.77 |
| 184 | 2jsa-9 | PHE_A_31_C | PHE_A_31_O | -2.877 | -3.33 | -5.248 | -2.652 | -2.779 | -6.323 | 1.23 | 151.03 | 67.79 |
| 185 | 2jsa-9 | PHE_A_31_C | ASP_A_32_N | -2.877 | -3.33 | -5.248 | -2.513 | -4.568 | -4.947 | 1.33 | 76.87 | -73.62 |
| 186 | 2jsa-9 | ASP_A_32_N | ASP_A_32_H | -2.513 | -4.568 | -4.947 | -2.73 | -4.957 | -4.04 | 1.01 | 26.16 | -119.15 |
| 187 | 2jsa-9 | ASP_A_32_N | ASP_A_32_CA | -2.513 | -4.568 | -4.947 | -1.796 | -5.39 | -5.908 | 1.45 | 131.38 | -48.90 |
| 188 | 2jsa-9 | ASP_A_32_CA | ASP_A_32_HA | -1.796 | -5.39 | -5.908 | -0.759 | -5.055 | -5.959 | 1.09 | 92.68 | 17.90 |
| 189 | 2jsa-9 | ASP_A_32_CA | ASP_A_32_C | -1.796 | -5.39 | -5.908 | -2.454 | -5.255 | -7.281 | 1.53 | 153.93 | 168.41 |
| 190 | 2jsa-9 | ASP_A_32_C | ASP_A_32_O | -2.454 | -5.255 | -7.281 | -1.773 | -5.28 | -8.306 | 1.23 | 146.38 | -2.10 |
| 191 | 2jsa-9 | ASP_A_32_C | ALA_A_33_N | -2.454 | -5.255 | -7.281 | -3.772 | -5.114 | -7.259 | 1.33 | 89.05 | 173.89 |
| 192 | 2jsa-9 | ALA_A_33_N | ALA_A_33_H | -3.772 | -5.114 | -7.259 | -4.267 | -5.102 | -6.379 | 1.01 | 29.36 | 178.61 |
| 193 | 2jsa-9 | ALA_A_33_N | ALA_A_33_CA | -3.772 | -5.114 | -7.259 | -4.53 | -4.975 | -8.491 | 1.45 | 147.97 | 169.61 |
| 194 | 2jsa-9 | ALA_A_33_CA | ALA_A_33_HA | -4.53 | -4.975 | -8.491 | -4.346 | -5.849 | -9.116 | 1.09 | 124.98 | -78.11 |
| 195 | 2jsa-9 | ALA_A_33_CA | ALA_A_33_C | -4.53 | -4.975 | -8.491 | -4.05 | -3.732 | -9.243 | 1.53 | 119.44 | 68.89 |
| 196 | 2jsa-9 | ALA_A_33_C | ALA_A_33_O | -4.05 | -3.732 | -9.243 | -3.625 | -3.824 | -10.393 | 1.23 | 159.29 | -12.21 |
| 197 | 2jsa-9 | ALA_A_33_C | GLU_A_34_N | -4.05 | -3.732 | -9.243 | -4.135 | -2.599 | -8.562 | 1.32 | 59.06 | 94.29 |
| 198 | 2jsa-9 | GLU_A_34_N | GLU_A_34_H | -4.135 | -2.599 | -8.562 | -4.495 | -2.598 | -7.618 | 1.01 | 20.87 | 179.84 |
| 199 | 2jsa-9 | GLU_A_34_N | GLU_A_34_CA | -4.135 | -2.599 | -8.562 | -3.714 | -1.339 | -9.151 | 1.45 | 113.91 | 71.52 |
| 200 | 2jsa-9 | GLU_A_34_CA | GLU_A_34_HA | -3.714 | -1.339 | -9.151 | -3.874 | -1.399 | -10.227 | 1.09 | 170.98 | -159.44 |
| 201 | 2jsa-9 | GLU_A_34_CA | GLU_A_34_C | -3.714 | -1.339 | -9.151 | -2.224 | -1.107 | -8.9 | 1.53 | 80.55 | 8.85 |
| 202 | 2jsa-9 | GLU_A_34_C | GLU_A_34_O | -2.224 | -1.107 | -8.9 | -1.769 | 0.035 | -8.847 | 1.23 | 87.53 | 68.28 |
| 203 | 2jsa-9 | GLU_A_34_C | CYS_A_35_N | -2.224 | -1.107 | -8.9 | -1.502 | -2.209 | -8.753 | 1.33 | 83.63 | -56.77 |
| 204 | 2jsa-9 | CYS_A_35_N | CYS_A_35_H | -1.502 | -2.209 | -8.753 | -1.942 | -3.116 | -8.81 | 1.01 | 93.24 | -115.88 |
| 205 | 2jsa-9 | CYS_A_35_N | CYS_A_35_CA | -1.502 | -2.209 | -8.753 | -0.072 | -2.141 | -8.509 | 1.45 | 80.33 | 2.72 |
| 206 | 2jsa-9 | CYS_A_35_CA | CYS_A_35_HA | -0.072 | -2.141 | -8.509 | 0.232 | -1.094 | -8.539 | 1.09 | 91.58 | 73.81 |
| 207 | 2jsa-9 | CYS_A_35_CA | CYS_A_35_C | -0.072 | -2.141 | -8.509 | 0.644 | -2.888 | -9.636 | 1.53 | 137.44 | -46.21 |
| 208 | 2jsa-9 | CYS_A_35_C | CYS_A_35_O | 0.644 | -2.888 | -9.636 | 1.498 | -2.321 | -10.316 | 1.23 | 123.56 | 33.58 |
| 209 | 2jsa-9 | CYS_A_35_C | LYS_A_36_N | 0.644 | -2.888 | -9.636 | 0.27 | -4.148 | -9.798 | 1.32 | 97.03 | -106.53 |
| 210 | 2jsa-9 | LYS_A_36_N | LYS_A_36_H | 0.27 | -4.148 | -9.798 | -0.439 | -4.549 | -9.201 | 1.01 | 53.76 | -150.51 |
| 211 | 2jsa-9 | LYS_A_36_N | LYS_A_36_CA | 0.27 | -4.148 | -9.798 | 0.866 | -4.978 | -10.831 | 1.45 | 135.31 | -54.32 |
| 212 | 2jsa-9 | LYS_A_36_CA | LYS_A_36_HA | 0.866 | -4.978 | -10.831 | 1.95 | -4.909 | -10.736 | 1.09 | 85.00 | 3.64 |
| 213 | 2jsa-9 | LYS_A_36_CA | LYS_A_36_C | 0.866 | -4.978 | -10.831 | 0.484 | -4.427 | -12.206 | 1.53 | 154.01 | 124.73 |
| 214 | 2jsa-9 | LYS_A_36_C | LYS_A_36_O | 0.484 | -4.427 | -12.206 | 1.291 | -4.448 | -13.134 | 1.23 | 138.98 | -1.49 |
| 215 | 2jsa-9 | LYS_A_36_C | LYS_A_37_N | 0.484 | -4.427 | -12.206 | -0.748 | -3.946 | -12.294 | 1.33 | 93.81 | 158.67 |

| | | | | | | | | | | | |
|---|---|---|---|---|---|---|---|---|---|---|---|
| 216 | 2jsa-9 | LYS_A_37_N | LYS_A_37_H | -0.748 | -3.946 | -12.294 | -1.358 | -3.961 | -11.489 | 1.01 | 37.16 | -178.59 |
| 217 | 2jsa-9 | LYS_A_37_N | LYS_A_37_CA | -0.748 | -3.946 | -12.294 | -1.247 | -3.39 | -13.54 | 1.45 | 149.05 | 131.91 |
| 218 | 2jsa-9 | LYS_A_37_CA | LYS_A_37_HA | -1.247 | -3.39 | -13.54 | -0.902 | -4.025 | -14.356 | 1.09 | 138.47 | -61.48 |
| 219 | 2jsa-9 | LYS_A_37_CA | LYS_A_37_C | -1.247 | -3.39 | -13.54 | -0.647 | -1.998 | -13.749 | 1.53 | 97.85 | 66.68 |
| 220 | 2jsa-9 | LYS_A_37_C | LYS_A_37_O | -0.647 | -1.998 | -13.749 | -0.241 | -1.654 | -14.858 | 1.23 | 154.37 | 40.27 |
| 221 | 2jsa-9 | LYS_A_37_C | LEU_A_38_N | -0.647 | -1.998 | -13.749 | -0.61 | -1.235 | -12.666 | 1.33 | 35.20 | 87.22 |
| 222 | 2jsa-9 | LEU_A_38_N | LEU_A_38_H | -0.61 | -1.235 | -12.666 | -0.962 | -1.583 | -11.786 | 1.01 | 29.36 | -135.33 |
| 223 | 2jsa-9 | LEU_A_38_N | LEU_A_38_CA | -0.61 | -1.235 | -12.666 | -0.066 | 0.111 | -12.718 | 1.45 | 92.05 | 67.99 |
| 224 | 2jsa-9 | LEU_A_38_CA | LEU_A_38_HA | -0.066 | 0.111 | -12.718 | -0.486 | 0.612 | -13.59 | 1.09 | 143.14 | 129.97 |
| 225 | 2jsa-9 | LEU_A_38_CA | LEU_A_38_C | -0.066 | 0.111 | -12.718 | 1.45 | 0.037 | -12.91 | 1.53 | 97.21 | -2.79 |
| 226 | 2jsa-9 | LEU_A_38_C | LEU_A_38_O | 1.45 | 0.037 | -12.91 | 1.984 | 0.577 | -13.877 | 1.23 | 141.86 | 45.32 |
| 227 | 2jsa-9 | LEU_A_38_C | PHE_A_39_N | 1.45 | 0.037 | -12.91 | 2.101 | -0.636 | -11.972 | 1.33 | 44.95 | -45.95 |
| 228 | 2jsa-9 | PHE_A_39_N | PHE_A_39_H | 2.101 | -0.636 | -11.972 | 1.597 | -1.054 | -11.203 | 1.01 | 40.41 | -140.33 |
| 229 | 2jsa-9 | PHE_A_39_N | PHE_A_39_CA | 2.101 | -0.636 | -11.972 | 3.545 | -0.788 | -12.026 | 1.45 | 92.13 | -6.01 |
| 230 | 2jsa-9 | PHE_A_39_CA | PHE_A_39_HA | 3.545 | -0.788 | -12.026 | 3.996 | 0.107 | -12.454 | 1.09 | 113.13 | 63.26 |
| 231 | 2jsa-9 | PHE_A_39_CA | PHE_A_39_C | 3.545 | -0.788 | -12.026 | 3.937 | -2.01 | -12.859 | 1.53 | 122.99 | -72.21 |
| 232 | 2jsa-9 | PHE_A_39_C | PHE_A_39_O | 3.937 | -2.01 | -12.859 | 5.015 | -2.574 | -12.672 | 1.23 | 81.26 | -27.62 |
| 233 | 2jsa-9 | PHE_A_39_C | HIS_A_40_N | 3.937 | -2.01 | -12.859 | 3.041 | -2.384 | -13.761 | 1.33 | 132.89 | -157.34 |
| 234 | 2jsa-9 | HIS_A_40_N | HIS_A_40_H | 3.041 | -2.384 | -13.761 | 2.175 | -1.874 | -13.86 | 1.01 | 95.63 | 149.51 |
| 235 | 2jsa-9 | HIS_A_40_N | HIS_A_40_CA | 3.041 | -2.384 | -13.761 | 3.281 | -3.528 | -14.622 | 1.45 | 126.37 | -78.15 |
| 236 | 2jsa-9 | HIS_A_40_CA | HIS_A_40_HA | 3.281 | -3.528 | -14.622 | 3.425 | -4.402 | -13.987 | 1.09 | 54.36 | -80.64 |
| 237 | 2jsa-9 | HIS_A_40_CA | HIS_A_40_C | 3.281 | -3.528 | -14.622 | 4.568 | -3.311 | -15.421 | 1.53 | 121.47 | 9.57 |
| 238 | 2jsa-9 | HIS_A_40_C | HIS_A_40_O | 4.568 | -3.311 | -15.421 | 5.144 | -4.26 | -15.948 | 1.23 | 115.39 | -58.74 |
| 239 | 2jsa-9 | HIS_A_40_C | THR_A_41_N | 4.568 | -3.311 | -15.421 | 4.981 | -2.054 | -15.483 | 1.32 | 92.68 | 71.81 |
| 240 | 2jsa-9 | THR_A_41_N | THR_A_41_H | 4.981 | -2.054 | -15.483 | 4.457 | -1.323 | -15.023 | 1.01 | 62.91 | 125.63 |
| 241 | 2jsa-9 | THR_A_41_N | THR_A_41_CA | 4.981 | -2.054 | -15.483 | 6.189 | -1.698 | -16.208 | 1.45 | 119.93 | 16.42 |
| 242 | 2jsa-9 | THR_A_41_CA | THR_A_41_HA | 6.189 | -1.698 | -16.208 | 6.333 | -2.412 | -17.019 | 1.09 | 138.07 | -78.60 |
| 243 | 2jsa-9 | THR_A_41_CA | THR_A_41_C | 6.189 | -1.698 | -16.208 | 7.408 | -1.78 | -15.287 | 1.53 | 52.99 | -3.85 |
| 244 | 2jsa-9 | THR_A_41_C | THR_A_41_O | 7.408 | -1.78 | -15.287 | 8.365 | -2.496 | -15.579 | 1.23 | 103.73 | -36.80 |
| 245 | 2jsa-9 | THR_A_41_C | ILE_A_42_N | 7.408 | -1.78 | -15.287 | 7.333 | -1.038 | -14.191 | 1.33 | 34.23 | 95.77 |
| 246 | 2jsa-9 | ILE_A_42_N | ILE_A_42_H | 7.333 | -1.038 | -14.191 | 6.514 | -0.474 | -14.013 | 1.01 | 79.85 | 145.45 |
| 247 | 2jsa-9 | ILE_A_42_N | ILE_A_42_CA | 7.333 | -1.038 | -14.191 | 8.417 | -1.018 | -13.225 | 1.45 | 48.30 | 1.06 |
| 248 | 2jsa-9 | ILE_A_42_CA | ILE_A_42_HA | 8.417 | -1.018 | -13.225 | 9.125 | -0.245 | -13.527 | 1.09 | 106.07 | 47.51 |
| 249 | 2jsa-9 | ILE_A_42_CA | ILE_A_42_C | 8.417 | -1.018 | -13.225 | 9.151 | -2.36 | -13.261 | 1.53 | 91.35 | -61.32 |
| 250 | 2jsa-9 | ILE_A_42_C | ILE_A_42_O | 9.151 | -2.36 | -13.261 | 8.523 | -3.412 | -13.374 | 1.23 | 95.27 | -120.84 |
| 251 | 2jsa-9 | ILE_A_42_C | PRO_A_43_N | 9.151 | -2.36 | -13.261 | 10.505 | -2.277 | -13.159 | 1.36 | 85.70 | 3.51 |
| 252 | 2jsa-9 | PRO_A_43_N | PRO_A_43_CA | 10.505 | -2.277 | -13.159 | 11.331 | -3.473 | -13.179 | 1.45 | 90.79 | -55.37 |
| 253 | 2jsa-9 | PRO_A_43_CA | PRO_A_43_HA | 11.331 | -3.473 | -13.179 | 11.002 | -4.141 | -13.974 | 1.09 | 136.87 | -116.22 |
| 254 | 2jsa-9 | PRO_A_43_CA | PRO_A_43_C | 11.331 | -3.473 | -13.179 | 11.242 | -4.222 | -11.847 | 1.53 | 29.52 | -96.78 |
| 255 | 2jsa-9 | PRO_A_43_C | PRO_A_43_O | 11.242 | -4.222 | -11.847 | 11.864 | -3.825 | -10.864 | 1.23 | 36.89 | 32.55 |
| 256 | 2jsa-9 | PRO_A_43_C | PHE_A_44_N | 11.242 | -4.222 | -11.847 | 10.463 | -5.294 | -11.86 | 1.33 | 90.56 | -126.01 |
| 257 | 2jsa-9 | PHE_A_44_N | PHE_A_44_H | 10.463 | -5.294 | -11.86 | 9.982 | -5.563 | -12.706 | 1.01 | 146.92 | -150.78 |
| 258 | 2jsa-9 | PHE_A_44_N | PHE_A_44_CA | 10.463 | -5.294 | -11.86 | 10.285 | -6.103 | -10.665 | 1.45 | 34.73 | -102.41 |
| 259 | 2jsa-9 | PHE_A_44_CA | PHE_A_44_HA | 10.285 | -6.103 | -10.665 | 9.876 | -7.076 | -10.935 | 1.09 | 104.35 | -112.80 |
| 260 | 2jsa-9 | PHE_A_44_CA | PHE_A_44_C | 10.285 | -6.103 | -10.665 | 9.353 | -5.411 | -9.669 | 1.53 | 49.37 | 143.41 |
| 261 | 2jsa-9 | PHE_A_44_C | PHE_A_44_O | 9.353 | -5.411 | -9.669 | 9.045 | -5.964 | -8.615 | 1.23 | 30.99 | -119.12 |
| 262 | 2jsa-9 | PHE_A_44_C | GLY_A_45_N | 9.353 | -5.411 | -9.669 | 8.93 | -4.211 | -10.038 | 1.32 | 106.17 | 109.42 |
| 263 | 2jsa-9 | GLY_A_45_N | GLY_A_45_H | 8.93 | -4.211 | -10.038 | 9.225 | -3.817 | -10.92 | 1.01 | 150.84 | 53.18 |
| 264 | 2jsa-9 | GLY_A_45_N | GLY_A_45_CA | 8.93 | -4.211 | -10.038 | 8.039 | -3.438 | -9.189 | 1.45 | 54.26 | 139.06 |
| 265 | 2jsa-9 | GLY_A_45_CA | GLY_A_45_C | 8.039 | -3.438 | -9.189 | 6.903 | -4.309 | -8.65 | 1.53 | 69.37 | -142.52 |
| 266 | 2jsa-9 | GLY_A_45_C | GLY_A_45_O | 6.903 | -4.309 | -8.65 | 6.546 | -4.213 | -7.475 | 1.23 | 17.46 | 164.95 |
| 267 | 2jsa-9 | GLY_A_45_C | THR_A_46_N | 6.903 | -4.309 | -8.65 | 6.366 | -5.139 | -9.532 | 1.32 | 131.74 | -122.90 |
| 268 | 2jsa-9 | THR_A_46_N | THR_A_46_H | 6.366 | -5.139 | -9.532 | 6.711 | -5.165 | -10.481 | 1.01 | 159.97 | -4.31 |
| 269 | 2jsa-9 | THR_A_46_N | THR_A_46_CA | 6.366 | -5.139 | -9.532 | 5.277 | -6.026 | -9.159 | 1.45 | 75.13 | -140.84 |
| 270 | 2jsa-9 | THR_A_46_CA | THR_A_46_HA | 5.277 | -6.026 | -9.159 | 4.398 | -5.424 | -8.931 | 1.09 | 77.92 | 145.59 |
| 271 | 2jsa-9 | THR_A_46_CA | THR_A_46_C | 5.277 | -6.026 | -9.159 | 5.643 | -6.82 | -7.903 | 1.53 | 34.84 | -65.25 |
| 272 | 2jsa-9 | THR_A_46_C | THR_A_46_O | 5.643 | -6.82 | -7.903 | 4.848 | -6.913 | -6.97 | 1.23 | 40.63 | -173.33 |
| 273 | 2jsa-9 | THR_A_46_C | ARG_A_47_N | 5.643 | -6.82 | -7.903 | 6.848 | -7.372 | -7.921 | 1.33 | 90.78 | -24.61 |
| 274 | 2jsa-9 | ARG_A_47_N | ARG_A_47_H | 6.848 | -7.372 | -7.921 | 7.45 | -7.254 | -8.723 | 1.01 | 142.59 | 11.09 |
| 275 | 2jsa-9 | ARG_A_47_N | ARG_A_47_CA | 6.848 | -7.372 | -7.921 | 7.329 | -8.156 | -6.796 | 1.45 | 39.27 | -58.47 |
| 276 | 2jsa-9 | ARG_A_47_CA | ARG_A_47_HA | 7.329 | -8.156 | -6.796 | 6.726 | -9.061 | -6.723 | 1.09 | 86.16 | -123.68 |
| 277 | 2jsa-9 | ARG_A_47_CA | ARG_A_47_C | 7.329 | -8.156 | -6.796 | 7.191 | -7.358 | -5.496 | 1.53 | 31.92 | 99.81 |
| 278 | 2jsa-9 | ARG_A_47_C | ARG_A_47_O | 7.191 | -7.358 | -5.496 | 6.847 | -7.916 | -4.455 | 1.23 | 32.20 | -121.65 |
| 279 | 2jsa-9 | ARG_A_47_C | GLU_A_48_N | 7.191 | -7.358 | -5.496 | 7.466 | -6.067 | -5.6 | 1.32 | 94.51 | 77.97 |
| 280 | 2jsa-9 | GLU_A_48_N | GLU_A_48_H | 7.466 | -6.067 | -5.6 | 7.743 | -5.676 | -6.489 | 1.01 | 151.68 | 54.68 |
| 281 | 2jsa-9 | GLU_A_48_N | GLU_A_48_CA | 7.466 | -6.067 | -5.6 | 7.376 | -5.188 | -4.447 | 1.45 | 37.46 | 95.85 |
| 282 | 2jsa-9 | GLU_A_48_CA | GLU_A_48_HA | 7.376 | -5.188 | -4.447 | 7.863 | -5.683 | -3.607 | 1.09 | 39.58 | -45.47 |
| 283 | 2jsa-9 | GLU_A_48_CA | GLU_A_48_C | 7.376 | -5.188 | -4.447 | 5.911 | -4.943 | -4.078 | 1.53 | 76.05 | 170.51 |
| 284 | 2jsa-9 | GLU_A_48_C | GLU_A_48_O | 5.911 | -4.943 | -4.078 | 5.576 | -4.826 | -2.901 | 1.23 | 16.78 | 160.75 |
| 285 | 2jsa-9 | GLU_A_48_C | CYS_A_49_N | 5.911 | -4.943 | -4.078 | 5.079 | -4.872 | -5.107 | 1.33 | 140.94 | 175.12 |
| 286 | 2jsa-9 | CYS_A_49_N | CYS_A_49_H | 5.079 | -4.872 | -5.107 | 5.426 | -4.977 | -6.05 | 1.01 | 158.97 | -16.84 |
| 287 | 2jsa-9 | CYS_A_49_N | CYS_A_49_CA | 5.079 | -4.872 | -5.107 | 3.658 | -4.644 | -4.906 | 1.45 | 82.05 | 170.88 |
| 288 | 2jsa-9 | CYS_A_49_CA | CYS_A_49_HA | 3.658 | -4.644 | -4.906 | 3.531 | -3.674 | -4.425 | 1.09 | 63.82 | 97.46 |
| 289 | 2jsa-9 | CYS_A_49_CA | CYS_A_49_C | 3.658 | -4.644 | -4.906 | 3.126 | -5.726 | -3.965 | 1.53 | 52.03 | -116.18 |
| 290 | 2jsa-9 | CYS_A_49_C | CYS_A_49_O | 3.126 | -5.726 | -3.965 | 2.58 | -5.419 | -2.906 | 1.23 | 30.60 | 150.65 |
| 291 | 2jsa-9 | CYS_A_49_C | ASP_A_50_N | 3.126 | -5.726 | -3.965 | 3.304 | -6.97 | -4.385 | 1.32 | 108.48 | -81.86 |
| 292 | 2jsa-9 | ASP_A_50_N | ASP_A_50_H | 3.304 | -6.97 | -4.385 | 3.762 | -7.149 | -5.268 | 1.01 | 150.89 | -21.35 |
| 293 | 2jsa-9 | ASP_A_50_N | ASP_A_50_CA | 3.304 | -6.97 | -4.385 | 2.848 | -8.1 | -3.592 | 1.45 | 56.94 | -111.98 |
| 294 | 2jsa-9 | ASP_A_50_CA | ASP_A_50_HA | 2.848 | -8.1 | -3.592 | 1.77 | -8.024 | -3.45 | 1.09 | 82.51 | 175.97 |
| 295 | 2jsa-9 | ASP_A_50_CA | ASP_A_50_C | 2.848 | -8.1 | -3.592 | 3.554 | -8.083 | -2.235 | 1.53 | 27.49 | 1.38 |
| 296 | 2jsa-9 | ASP_A_50_C | ASP_A_50_O | 3.554 | -8.083 | -2.235 | 3.014 | -8.568 | -1.243 | 1.23 | 36.19 | -138.07 |
| 297 | 2jsa-9 | ASP_A_50_C | HIS_A_51_N | 3.554 | -8.083 | -2.235 | 4.754 | -7.52 | -2.234 | 1.33 | 89.96 | 25.13 |
| 298 | 2jsa-9 | HIS_A_51_N | HIS_A_51_H | 4.754 | -7.52 | -2.234 | 5.139 | -7.142 | -3.088 | 1.01 | 147.72 | 44.47 |
| 299 | 2jsa-9 | HIS_A_51_N | HIS_A_51_CA | 4.754 | -7.52 | -2.234 | 5.54 | -7.433 | -1.015 | 1.45 | 32.97 | 6.32 |
| 300 | 2jsa-9 | HIS_A_51_CA | HIS_A_51_HA | 5.54 | -7.433 | -1.015 | 5.364 | -8.339 | -0.435 | 1.09 | 57.85 | -100.99 |
| 301 | 2jsa-9 | HIS_A_51_CA | HIS_A_51_C | 5.54 | -7.433 | -1.015 | 5.064 | -6.242 | -0.181 | 1.53 | 56.97 | 111.78 |
| 302 | 2jsa-9 | HIS_A_51_C | HIS_A_51_O | 5.064 | -6.242 | -0.181 | 5.588 | -5.99 | 0.903 | 1.23 | 28.21 | 25.68 |
| 303 | 2jsa-9 | HIS_A_51_C | TYR_A_52_N | 5.064 | -6.242 | -0.181 | 4.076 | -5.54 | -0.718 | 1.33 | 113.90 | 144.61 |
| 304 | 2jsa-9 | TYR_A_52_N | TYR_A_52_H | 4.076 | -5.54 | -0.718 | 3.695 | -5.804 | -1.615 | 1.01 | 152.67 | -145.28 |
| 305 | 2jsa-9 | TYR_A_52_N | TYR_A_52_CA | 4.076 | -5.54 | -0.718 | 3.523 | -4.383 | -0.037 | 1.45 | 62.03 | 115.55 |
| 306 | 2jsa-9 | TYR_A_52_CA | TYR_A_52_HA | 3.523 | -4.383 | -0.037 | 4.228 | -4.01 | 0.706 | 1.09 | 47.03 | 27.88 |
| 307 | 2jsa-9 | TYR_A_52_CA | TYR_A_52_C | 3.523 | -4.383 | -0.037 | 2.193 | -4.725 | 0.636 | 1.53 | 63.89 | -165.58 |
| 308 | 2jsa-9 | TYR_A_52_C | TYR_A_52_O | 2.193 | -4.725 | 0.636 | 1.838 | -4.134 | 1.654 | 1.23 | 34.11 | 120.99 |
| 309 | 2jsa-9 | TYR_A_52_C | VAL_A_53_N | 2.193 | -4.725 | 0.636 | 1.493 | -5.679 | 0.039 | 1.33 | 116.77 | -126.27 |
| 310 | 2jsa-9 | VAL_A_53_N | VAL_A_53_H | 1.493 | -5.679 | 0.039 | 1.847 | -6.122 | -0.797 | 1.01 | 145.85 | -51.37 |
| 311 | 2jsa-9 | VAL_A_53_N | VAL_A_53_CA | 1.493 | -5.679 | 0.039 | 0.209 | -6.108 | 0.568 | 1.45 | 68.66 | -161.52 |

| | | | | | | | | | | |
|---|---|---|---|---|---|---|---|---|---|---|
| 312 | 2jsa-9 | VAL_A_53_CA | VAL_A_53_HA | 0.209 | -6.108 | 0.568 | -0.235 | -5.274 | 1.112 | 1.09 | 60.07 | 118.03 |
| 313 | 2jsa-9 | VAL_A_53_CA | VAL_A_53_C | 0.209 | -6.108 | 0.568 | 0.427 | -7.257 | 1.555 | 1.53 | 49.84 | -79.26 |
| 314 | 2jsa-9 | VAL_A_53_C | VAL_A_53_O | 0.427 | -7.257 | 1.555 | -0.395 | -8.167 | 1.644 | 1.23 | 85.85 | -132.09 |
| 315 | 2jsa-9 | VAL_A_53_C | ASN_A_54_N | 0.427 | -7.257 | 1.555 | 1.538 | -7.175 | 2.272 | 1.32 | 57.23 | 4.22 |
| 316 | 2jsa-9 | ASN_A_54_N | ASN_A_54_H | 1.538 | -7.175 | 2.272 | 2.168 | -6.396 | 2.147 | 1.01 | 97.11 | 51.04 |
| 317 | 2jsa-9 | ASN_A_54_N | ASN_A_54_CA | 1.538 | -7.175 | 2.272 | 1.874 | -8.196 | 3.249 | 1.45 | 47.73 | -71.78 |
| 318 | 2jsa-9 | ASN_A_54_CA | ASN_A_54_HA | 1.874 | -8.196 | 3.249 | 1.078 | -8.256 | 3.991 | 1.09 | 47.09 | -175.69 |
| 319 | 2jsa-9 | ASN_A_54_CA | ASN_A_54_C | 1.874 | -8.196 | 3.249 | 3.193 | -7.829 | 3.932 | 1.53 | 63.49 | 15.55 |
| 320 | 2jsa-9 | ASN_A_54_C | ASN_A_54_O | 3.193 | -7.829 | 3.932 | 4.08 | -8.67 | 4.073 | 1.23 | 83.42 | -43.48 |
| 321 | 2jsa-9 | ASN_A_54_C | SER_A_55_N | 3.193 | -7.829 | 3.932 | 3.282 | -6.571 | 4.339 | 1.33 | 72.11 | 85.95 |
| 322 | 2jsa-9 | SER_A_55_N | SER_A_55_H | 3.282 | -6.571 | 4.339 | 2.513 | -5.933 | 4.191 | 1.01 | 98.43 | 140.32 |
| 323 | 2jsa-9 | SER_A_55_N | SER_A_55_CA | 3.282 | -6.571 | 4.339 | 4.477 | -6.081 | 5.004 | 1.45 | 62.76 | 22.30 |
| 324 | 2jsa-9 | SER_A_55_CA | SER_A_55_HA | 4.477 | -6.081 | 5.004 | 4.513 | -6.513 | 6.004 | 1.09 | 23.44 | -85.24 |
| 325 | 2jsa-9 | SER_A_55_CA | SER_A_55_C | 4.477 | -6.081 | 5.004 | 4.419 | -4.558 | 5.13 | 1.53 | 85.27 | 92.18 |
| 326 | 2jsa-9 | SER_A_55_C | SER_A_55_O | 4.419 | -4.558 | 5.13 | 4.953 | -3.99 | 6.082 | 1.23 | 39.31 | 46.77 |
| 327 | 2jsa-9 | SER_A_55_C | LYS_A_56_N | 4.419 | -4.558 | 5.13 | 3.765 | -3.939 | 4.158 | 1.33 | 137.19 | 136.57 |
| 328 | 2jsa-9 | LYS_A_56_N | LYS_A_56_H | 3.765 | -3.939 | 4.158 | 3.35 | -4.47 | 3.406 | 1.01 | 138.13 | -128.01 |
| 329 | 2jsa-9 | LYS_A_56_N | LYS_A_56_CA | 3.765 | -3.939 | 4.158 | 3.63 | -2.493 | 4.149 | 1.45 | 90.36 | 95.33 |
| 330 | 2jsa-9 | LYS_A_56_CA | LYS_A_56_HA | 3.63 | -2.493 | 4.149 | 4.136 | -2.101 | 5.032 | 1.09 | 35.94 | 37.77 |
| 331 | 2jsa-9 | LYS_A_56_CA | LYS_A_56_C | 3.63 | -2.493 | 4.149 | 2.15 | -2.122 | 4.259 | 1.53 | 85.88 | 165.93 |
| 332 | 2jsa-9 | LYS_A_56_C | LYS_A_56_O | 2.15 | -2.122 | 4.259 | 1.807 | -0.946 | 4.371 | 1.23 | 84.78 | 106.26 |
| 333 | 2jsa-9 | LYS_A_56_C | VAL_A_57_N | 2.15 | -2.122 | 4.259 | 1.312 | -3.148 | 4.223 | 1.33 | 91.56 | -129.24 |
| 334 | 2jsa-9 | VAL_A_57_N | VAL_A_57_H | 1.312 | -3.148 | 4.223 | 1.665 | -4.089 | 4.129 | 1.01 | 95.34 | -69.44 |
| 335 | 2jsa-9 | VAL_A_57_N | VAL_A_57_CA | 1.312 | -3.148 | 4.223 | -0.124 | -2.946 | 4.317 | 1.45 | 86.29 | 171.99 |
| 336 | 2jsa-9 | VAL_A_57_CA | VAL_A_57_HA | -0.124 | -2.946 | 4.317 | -0.343 | -1.911 | 4.053 | 1.09 | 104.01 | 101.95 |
| 337 | 2jsa-9 | VAL_A_57_CA | VAL_A_57_C | -0.124 | -2.946 | 4.317 | -0.573 | -3.172 | 5.761 | 1.53 | 19.19 | -153.28 |
| 338 | 2jsa-9 | VAL_A_57_C | VAL_A_57_O | -0.573 | -3.172 | 5.761 | -1.312 | -2.363 | 6.32 | 1.23 | 62.97 | 132.41 |
| 339 | 2jsa-9 | VAL_A_57_C | ASP_A_58_N | -0.573 | -3.172 | 5.761 | -0.108 | -4.277 | 6.326 | 1.33 | 64.77 | -67.18 |
| 340 | 2jsa-9 | ASP_A_58_N | ASP_A_58_H | -0.108 | -4.277 | 6.326 | 0.497 | -4.898 | 5.807 | 1.01 | 120.91 | -45.75 |
| 341 | 2jsa-9 | ASP_A_58_N | ASP_A_58_CA | -0.108 | -4.277 | 6.326 | -0.453 | -4.621 | 7.694 | 1.45 | 19.60 | -135.08 |
| 342 | 2jsa-9 | ASP_A_58_CA | ASP_A_58_HA | -0.453 | -4.621 | 7.694 | -1.512 | -4.871 | 7.742 | 1.09 | 87.47 | -166.72 |
| 343 | 2jsa-9 | ASP_A_58_CA | ASP_A_58_C | -0.453 | -4.621 | 7.694 | -0.158 | -3.427 | 8.605 | 1.53 | 53.47 | 76.12 |
| 344 | 2jsa-9 | ASP_A_58_C | ASP_A_58_O | -0.158 | -3.427 | 8.605 | -0.962 | -3.09 | 9.473 | 1.23 | 45.12 | 157.26 |
| 345 | 2jsa-9 | ASP_A_58_C | PRO_A_59_N | -0.158 | -3.427 | 8.605 | 1.028 | -2.803 | 8.37 | 1.36 | 99.95 | 27.75 |
| 346 | 2jsa-9 | PRO_A_59_N | PRO_A_59_CA | 1.028 | -2.803 | 8.37 | 1.438 | -1.655 | 9.159 | 1.45 | 57.09 | 70.35 |
| 347 | 2jsa-9 | PRO_A_59_CA | PRO_A_59_HA | 1.438 | -1.655 | 9.159 | 1.257 | -1.845 | 10.217 | 1.09 | 13.93 | -133.61 |
| 348 | 2jsa-9 | PRO_A_59_CA | PRO_A_59_C | 1.438 | -1.655 | 9.159 | 0.66 | -0.402 | 8.751 | 1.53 | 105.46 | 121.84 |
| 349 | 2jsa-9 | PRO_A_59_C | PRO_A_59_O | 0.66 | -0.402 | 8.751 | 0.02 | 0.235 | 9.585 | 1.23 | 47.27 | 135.13 |
| 350 | 2jsa-9 | PRO_A_59_C | ILE_A_60_N | 0.66 | -0.402 | 8.751 | 0.741 | -0.087 | 7.465 | 1.33 | 165.81 | 75.58 |
| 351 | 2jsa-9 | ILE_A_60_N | ILE_A_60_H | 0.741 | -0.087 | 7.465 | 1.286 | -0.658 | 6.835 | 1.01 | 128.59 | -46.33 |
| 352 | 2jsa-9 | ILE_A_60_N | ILE_A_60_CA | 0.741 | -0.087 | 7.465 | 0.052 | 1.077 | 6.936 | 1.45 | 111.36 | 120.62 |
| 353 | 2jsa-9 | ILE_A_60_CA | ILE_A_60_HA | 0.052 | 1.077 | 6.936 | 0.567 | 1.967 | 7.298 | 1.09 | 70.61 | 59.94 |
| 354 | 2jsa-9 | ILE_A_60_CA | ILE_A_60_C | 0.052 | 1.077 | 6.936 | -1.377 | 1.111 | 7.483 | 1.53 | 69.06 | 178.64 |
| 355 | 2jsa-9 | ILE_A_60_C | ILE_A_60_O | -1.377 | 1.111 | 7.483 | -1.774 | 2.077 | 8.134 | 1.23 | 58.06 | 112.34 |
| 356 | 2jsa-9 | ILE_A_60_C | ILE_A_61_N | -1.377 | 1.111 | 7.483 | -2.112 | 0.046 | 7.198 | 1.33 | 102.42 | -124.61 |
| 357 | 2jsa-9 | ILE_A_61_N | ILE_A_61_H | -2.112 | 0.046 | 7.198 | -1.72 | -0.712 | 6.657 | 1.01 | 122.37 | -62.65 |
| 358 | 2jsa-9 | ILE_A_61_N | ILE_A_61_CA | -2.112 | 0.046 | 7.198 | -3.487 | -0.059 | 7.653 | 1.45 | 71.74 | -175.63 |
| 359 | 2jsa-9 | ILE_A_61_CA | ILE_A_61_HA | -3.487 | -0.059 | 7.653 | -4.034 | 0.803 | 7.271 | 1.09 | 110.51 | 122.40 |
| 360 | 2jsa-9 | ILE_A_61_CA | ILE_A_61_C | -3.487 | -0.059 | 7.653 | -3.522 | 0.009 | 9.181 | 1.53 | 2.87 | 117.24 |
| 361 | 2jsa-9 | ILE_A_61_C | ILE_A_61_O | -3.522 | 0.009 | 9.181 | -4.548 | 0.348 | 9.768 | 1.23 | 61.49 | 161.72 |
| 362 | 2jsa-9 | ILE_A_61_C | HIS_A_62_N | -3.522 | 0.009 | 9.181 | -2.386 | -0.318 | 9.782 | 1.33 | 63.05 | -16.06 |
| 363 | 2jsa-9 | HIS_A_62_N | HIS_A_62_H | -2.386 | -0.318 | 9.782 | -1.58 | -0.586 | 9.236 | 1.01 | 122.73 | -18.39 |
| 364 | 2jsa-9 | HIS_A_62_N | HIS_A_62_CA | -2.386 | -0.318 | 9.782 | -2.274 | -0.299 | 11.23 | 1.45 | 4.49 | 9.63 |
| 365 | 2jsa-9 | HIS_A_62_CA | HIS_A_62_HA | -2.274 | -0.299 | 11.23 | -3.094 | -0.889 | 11.639 | 1.09 | 67.96 | -144.26 |
| 366 | 2jsa-9 | HIS_A_62_CA | HIS_A_62_C | -2.274 | -0.299 | 11.23 | -2.428 | 1.136 | 11.738 | 1.53 | 70.61 | 96.13 |
| 367 | 2jsa-9 | HIS_A_62_C | HIS_A_62_O | -2.428 | 1.136 | 11.738 | -2.914 | 1.358 | 12.845 | 1.23 | 25.76 | 155.45 |
| 368 | 2jsa-9 | HIS_A_62_C | GLU_A_63_N | -2.428 | 1.136 | 11.738 | -2.003 | 2.074 | 10.903 | 1.33 | 129.04 | 65.63 |
| 369 | 2jsa-9 | GLU_A_63_N | GLU_A_63_H | -2.003 | 2.074 | 10.903 | -1.613 | 1.818 | 10.007 | 1.01 | 152.50 | -33.28 |
| 370 | 2jsa-9 | GLU_A_63_N | GLU_A_63_CA | -2.003 | 2.074 | 10.903 | -2.087 | 3.481 | 11.253 | 1.45 | 76.05 | 93.42 |
| 371 | 2jsa-9 | GLU_A_63_CA | GLU_A_63_HA | -2.087 | 3.481 | 11.253 | -1.916 | 3.572 | 12.326 | 1.09 | 10.23 | 28.02 |
| 372 | 2jsa-9 | GLU_A_63_CA | GLU_A_63_C | -2.087 | 3.481 | 11.253 | -3.483 | 4.024 | 10.94 | 1.53 | 101.80 | 158.75 |
| 373 | 2jsa-9 | GLU_A_63_C | GLU_A_63_O | -3.483 | 4.024 | 10.94 | -4.169 | 4.531 | 11.826 | 1.23 | 43.91 | 143.53 |
| 374 | 2jsa-9 | GLU_A_63_C | LEU_A_64_N | -3.483 | 4.024 | 10.94 | -3.863 | 3.898 | 9.677 | 1.32 | 162.41 | -161.66 |
| 375 | 2jsa-9 | LEU_A_64_N | LEU_A_64_H | -3.863 | 3.898 | 9.677 | -3.247 | 3.47 | 9.001 | 1.01 | 132.03 | -34.79 |
| 376 | 2jsa-9 | LEU_A_64_N | LEU_A_64_CA | -3.863 | 3.898 | 9.677 | -5.164 | 4.369 | 9.237 | 1.45 | 107.64 | 160.10 |
| 377 | 2jsa-9 | LEU_A_64_CA | LEU_A_64_HA | -5.164 | 4.369 | 9.237 | -5.181 | 5.455 | 9.336 | 1.09 | 84.79 | 90.90 |
| 378 | 2jsa-9 | LEU_A_64_CA | LEU_A_64_C | -5.164 | 4.369 | 9.237 | -6.246 | 3.801 | 10.157 | 1.53 | 53.03 | -152.30 |
| 379 | 2jsa-9 | LEU_A_64_C | LEU_A_64_O | -6.246 | 3.801 | 10.157 | -7.133 | 4.529 | 10.602 | 1.23 | 68.80 | 140.62 |
| 380 | 2jsa-9 | LEU_A_64_C | GLU_A_65_N | -6.246 | 3.801 | 10.157 | -6.138 | 2.506 | 10.415 | 1.32 | 78.77 | -85.23 |
| 381 | 2jsa-9 | GLU_A_65_N | GLU_A_65_H | -6.138 | 2.506 | 10.415 | -5.382 | 1.969 | 10.014 | 1.01 | 113.39 | -35.39 |
| 382 | 2jsa-9 | GLU_A_65_N | GLU_A_65_CA | -6.138 | 2.506 | 10.415 | -7.096 | 1.833 | 11.275 | 1.45 | 53.70 | -144.91 |
| 383 | 2jsa-9 | GLU_A_65_CA | GLU_A_65_HA | -7.096 | 1.833 | 11.275 | -8.091 | 1.989 | 10.859 | 1.09 | 112.44 | 171.09 |
| 384 | 2jsa-9 | GLU_A_65_CA | GLU_A_65_C | -7.096 | 1.833 | 11.275 | -7.063 | 2.436 | 12.681 | 1.53 | 23.24 | 86.87 |
| 385 | 2jsa-9 | GLU_A_65_C | GLU_A_65_O | -7.063 | 2.436 | 12.681 | -8.102 | 2.575 | 13.325 | 1.23 | 58.44 | 172.38 |
| 386 | 2jsa-9 | GLU_A_65_C | GLY_A_66_N | -7.063 | 2.436 | 12.681 | -5.859 | 2.775 | 13.117 | 1.32 | 70.78 | 15.73 |
| 387 | 2jsa-9 | GLY_A_66_N | GLY_A_66_H | -5.859 | 2.775 | 13.117 | -5.047 | 2.63 | 12.533 | 1.01 | 125.30 | -10.12 |
| 388 | 2jsa-9 | GLY_A_66_N | GLY_A_66_CA | -5.859 | 2.775 | 13.117 | -5.676 | 3.359 | 14.434 | 1.45 | 24.92 | 72.60 |
| 389 | 2jsa-9 | GLY_A_66_CA | GLY_A_66_C | -5.676 | 3.359 | 14.434 | -6.346 | 4.732 | 14.523 | 1.53 | 86.67 | 116.01 |
| 390 | 2jsa-9 | GLY_A_66_C | GLY_A_66_O | -6.346 | 4.732 | 14.523 | -6.744 | 5.164 | 15.604 | 1.23 | 28.52 | 132.65 |
| 391 | 2jsa-9 | GLY_A_66_C | GLY_A_67_N | -6.346 | 4.732 | 14.523 | -6.451 | 5.379 | 13.372 | 1.32 | 150.34 | 99.22 |
| 392 | 2jsa-9 | GLY_A_67_N | GLY_A_67_H | -6.451 | 5.379 | 13.372 | -6.101 | 4.963 | 12.521 | 1.01 | 147.43 | -49.92 |
| 393 | 2jsa-9 | GLY_A_67_N | GLY_A_67_CA | -6.451 | 5.379 | 13.372 | -7.067 | 6.694 | 13.305 | 1.45 | 92.64 | 115.10 |
| 394 | 2jsa-9 | GLY_A_67_CA | GLY_A_67_C | -7.067 | 6.694 | 13.305 | -6.156 | 7.69 | 12.583 | 1.53 | 118.14 | 47.55 |
| 395 | 2jsa-9 | GLY_A_67_C | GLY_A_67_O | -6.156 | 7.69 | 12.583 | -6.523 | 8.849 | 12.394 | 1.23 | 98.84 | 107.57 |
| 396 | 2jsa-9 | GLY_A_67_C | THR_A_68_N | -6.156 | 7.69 | 12.583 | -4.986 | 7.201 | 12.199 | 1.32 | 106.85 | -22.68 |
| 397 | 2jsa-9 | THR_A_68_N | THR_A_68_H | -4.986 | 7.201 | 12.199 | -4.75 | 6.237 | 12.388 | 1.01 | 79.22 | -76.24 |
| 398 | 2jsa-9 | THR_A_68_N | THR_A_68_CA | -4.986 | 7.201 | 12.199 | -4.02 | 8.033 | 11.501 | 1.45 | 118.70 | 40.74 |
| 399 | 2jsa-9 | THR_A_68_CA | THR_A_68_HA | -4.02 | 8.033 | 11.501 | -3.92 | 8.977 | 12.036 | 1.09 | 60.60 | 83.95 |
| 400 | 2jsa-9 | THR_A_68_CA | THR_A_68_C | -4.02 | 8.033 | 11.501 | -4.508 | 8.344 | 10.085 | 1.53 | 157.77 | 147.49 |
| 401 | 2jsa-9 | THR_A_68_C | THR_A_68_O | -4.508 | 8.344 | 10.085 | -5.339 | 7.621 | 9.538 | 1.23 | 116.41 | -138.98 |
| 402 | 2jsa-9 | THR_A_68_C | ALA_A_69_N | -4.508 | 8.344 | 10.085 | -3.97 | 9.42 | 9.531 | 1.32 | 114.73 | 63.43 |
| 403 | 2jsa-9 | ALA_A_69_N | ALA_A_69_H | -3.97 | 9.42 | 9.531 | -3.29 | 9.968 | 10.04 | 1.01 | 59.77 | 38.86 |
| 404 | 2jsa-9 | ALA_A_69_N | ALA_A_69_CA | -3.97 | 9.42 | 9.531 | -4.341 | 9.837 | 8.189 | 1.45 | 157.42 | 131.66 |
| 405 | 2jsa-9 | ALA_A_69_CA | ALA_A_69_HA | -4.341 | 9.837 | 8.189 | -5.382 | 9.568 | 8.014 | 1.09 | 99.24 | -165.51 |
| 406 | 2jsa-9 | ALA_A_69_CA | ALA_A_69_C | -4.341 | 9.837 | 8.189 | -3.471 | 9.094 | 7.172 | 1.53 | 131.63 | -40.50 |
| 407 | 2jsa-9 | ALA_A_69_C | ALA_A_69_O | -3.471 | 9.094 | 7.172 | -2.44 | 8.526 | 7.529 | 1.23 | 73.13 | -28.85 |

| | | | | | | | | | | | | |
|---|---|---|---|---|---|---|---|---|---|---|---|---|
| 408 | 2jsa-9 | ALA_A_69_C | PRO_A_70_N | -3.471 | 9.094 | 7.172 | -3.93 | 9.123 | 5.893 | 1.36 | 160.22 | 176.38 |
| 409 | 2jsa-9 | PRO_A_70_N | PRO_A_70_CA | -3.93 | 9.123 | 5.893 | -3.206 | 8.461 | 4.821 | 1.45 | 137.54 | -42.44 |
| 410 | 2jsa-9 | PRO_A_70_CA | PRO_A_70_HA | -3.206 | 8.461 | 4.821 | -2.894 | 7.467 | 5.141 | 1.09 | 72.93 | -72.57 |
| 411 | 2jsa-9 | PRO_A_70_CA | PRO_A_70_C | -3.206 | 8.461 | 4.821 | -1.96 | 9.256 | 4.428 | 1.53 | 104.89 | 32.54 |
| 412 | 2jsa-9 | PRO_A_70_C | PRO_A_70_O | -1.96 | 9.256 | 4.428 | -1.207 | 8.843 | 3.547 | 1.23 | 135.73 | -28.74 |
| 413 | 2jsa-9 | PRO_A_70_C | LYS_A_71_N | -1.96 | 9.256 | 4.428 | -1.779 | 10.383 | 5.102 | 1.33 | 59.44 | 80.88 |
| 414 | 2jsa-9 | LYS_A_71_N | LYS_A_71_H | -1.779 | 10.383 | 5.102 | -2.438 | 10.661 | 5.815 | 1.01 | 45.09 | 157.13 |
| 415 | 2jsa-9 | LYS_A_71_N | LYS_A_71_CA | -1.779 | 10.383 | 5.102 | -0.637 | 11.24 | 4.834 | 1.45 | 100.63 | 36.89 |
| 416 | 2jsa-9 | LYS_A_71_CA | LYS_A_71_HA | -0.637 | 11.24 | 4.834 | -0.107 | 10.836 | 3.971 | 1.09 | 142.32 | -37.32 |
| 417 | 2jsa-9 | LYS_A_71_CA | LYS_A_71_C | -0.637 | 11.24 | 4.834 | 0.319 | 11.197 | 6.028 | 1.53 | 38.71 | -2.58 |
| 418 | 2jsa-9 | LYS_A_71_C | LYS_A_71_O | 0.319 | 11.197 | 6.028 | 1.426 | 11.728 | 5.961 | 1.23 | 93.12 | 25.63 |
| 419 | 2jsa-9 | LYS_A_71_C | ASP_A_72_N | 0.319 | 11.197 | 6.028 | -0.144 | 10.559 | 7.093 | 1.33 | 36.51 | -125.97 |
| 420 | 2jsa-9 | ASP_A_72_N | ASP_A_72_H | -0.144 | 10.559 | 7.093 | -1.065 | 10.146 | 7.078 | 1.01 | 90.85 | -155.85 |
| 421 | 2jsa-9 | ASP_A_72_N | ASP_A_72_CA | -0.144 | 10.559 | 7.093 | 0.655 | 10.439 | 8.301 | 1.45 | 33.78 | -8.54 |
| 422 | 2jsa-9 | ASP_A_72_CA | ASP_A_72_HA | 0.655 | 10.439 | 8.301 | 1.422 | 11.214 | 8.299 | 1.09 | 90.11 | 45.30 |
| 423 | 2jsa-9 | ASP_A_72_CA | ASP_A_72_C | 0.655 | 10.439 | 8.301 | 1.312 | 9.057 | 8.338 | 1.53 | 88.61 | -64.57 |
| 424 | 2jsa-9 | ASP_A_72_C | ASP_A_72_O | 1.312 | 9.057 | 8.338 | 2.43 | 8.91 | 8.83 | 1.23 | 66.43 | -7.49 |
| 425 | 2jsa-9 | ASP_A_72_C | VAL_A_73_N | 1.312 | 9.057 | 8.338 | 0.59 | 8.08 | 7.811 | 1.32 | 113.45 | -126.46 |
| 426 | 2jsa-9 | VAL_A_73_N | VAL_A_73_H | 0.59 | 8.08 | 7.811 | -0.322 | 8.274 | 7.424 | 1.01 | 112.54 | 167.99 |
| 427 | 2jsa-9 | VAL_A_73_N | VAL_A_73_CA | 0.59 | 8.08 | 7.811 | 1.088 | 6.715 | 7.777 | 1.45 | 91.34 | -69.96 |
| 428 | 2jsa-9 | VAL_A_73_CA | VAL_A_73_HA | 1.088 | 6.715 | 7.777 | 1.3 | 6.4 | 8.799 | 1.09 | 20.38 | -56.06 |
| 429 | 2jsa-9 | VAL_A_73_CA | VAL_A_73_C | 1.088 | 6.715 | 7.777 | 2.397 | 6.675 | 6.987 | 1.53 | 121.10 | -1.75 |
| 430 | 2jsa-9 | VAL_A_73_C | VAL_A_73_O | 2.397 | 6.675 | 6.987 | 3.405 | 6.165 | 7.474 | 1.23 | 66.68 | -26.84 |
| 431 | 2jsa-9 | VAL_A_73_C | CYS_A_74_N | 2.397 | 6.675 | 6.987 | 2.339 | 7.217 | 5.78 | 1.32 | 155.70 | 96.11 |
| 432 | 2jsa-9 | CYS_A_74_N | CYS_A_74_H | 2.339 | 7.217 | 5.78 | 1.476 | 7.619 | 5.444 | 1.01 | 109.44 | 155.02 |
| 433 | 2jsa-9 | CYS_A_74_N | CYS_A_74_CA | 2.339 | 7.217 | 5.78 | 3.508 | 7.248 | 4.917 | 1.45 | 126.43 | 1.52 |
| 434 | 2jsa-9 | CYS_A_74_CA | CYS_A_74_HA | 3.508 | 7.248 | 4.917 | 4.028 | 6.294 | 5.009 | 1.09 | 85.16 | -61.41 |
| 435 | 2jsa-9 | CYS_A_74_CA | CYS_A_74_C | 3.508 | 7.248 | 4.917 | 4.438 | 8.36 | 5.408 | 1.53 | 71.29 | 50.09 |
| 436 | 2jsa-9 | CYS_A_74_C | CYS_A_74_O | 4.438 | 8.36 | 5.408 | 4.585 | 9.389 | 4.748 | 1.23 | 122.41 | 81.87 |
| 437 | 2jsa-9 | CYS_A_74_C | THR_A_75_N | 4.438 | 8.36 | 5.408 | 5.042 | 8.116 | 6.562 | 1.33 | 29.44 | -22.00 |
| 438 | 2jsa-9 | THR_A_75_N | THR_A_75_H | 5.042 | 8.116 | 6.562 | 4.875 | 7.248 | 7.05 | 1.01 | 61.10 | -100.89 |
| 439 | 2jsa-9 | THR_A_75_N | THR_A_75_CA | 5.042 | 8.116 | 6.562 | 5.953 | 9.084 | 7.149 | 1.45 | 66.17 | 46.74 |
| 440 | 2jsa-9 | THR_A_75_CA | THR_A_75_HA | 5.953 | 9.084 | 7.149 | 6.834 | 9.167 | 6.512 | 1.09 | 125.75 | 5.38 |
| 441 | 2jsa-9 | THR_A_75_CA | THR_A_75_C | 5.953 | 9.084 | 7.149 | 6.414 | 8.613 | 8.53 | 1.53 | 25.51 | -45.61 |
| 442 | 2jsa-9 | THR_A_75_C | THR_A_75_O | 6.414 | 8.613 | 8.53 | 7.582 | 8.761 | 8.882 | 1.23 | 73.35 | 7.22 |
| 443 | 2jsa-9 | THR_A_75_C | LYS_A_76_N | 6.414 | 8.613 | 8.53 | 5.471 | 8.054 | 9.274 | 1.32 | 55.84 | -149.34 |
| 444 | 2jsa-9 | LYS_A_76_N | LYS_A_76_H | 5.471 | 8.054 | 9.274 | 4.53 | 7.964 | 8.919 | 1.01 | 110.58 | -174.54 |
| 445 | 2jsa-9 | LYS_A_76_N | LYS_A_76_CA | 5.471 | 8.054 | 9.274 | 5.764 | 7.561 | 10.608 | 1.45 | 23.26 | -59.28 |
| 446 | 2jsa-9 | LYS_A_76_CA | LYS_A_76_HA | 5.764 | 7.561 | 10.608 | 6.413 | 8.286 | 11.101 | 1.09 | 63.13 | 48.17 |
| 447 | 2jsa-9 | LYS_A_76_CA | LYS_A_76_C | 5.764 | 7.561 | 10.608 | 6.536 | 6.243 | 10.501 | 1.53 | 94.01 | -59.64 |
| 448 | 2jsa-9 | LYS_A_76_C | LYS_A_76_O | 6.536 | 6.243 | 10.501 | 7.54 | 6.051 | 11.186 | 1.23 | 56.17 | -10.83 |
| 449 | 2jsa-9 | LYS_A_76_C | LEU_A_77_N | 6.536 | 6.243 | 10.501 | 6.037 | 5.371 | 9.64 | 1.32 | 130.60 | -119.78 |
| 450 | 2jsa-9 | LEU_A_77_N | LEU_A_77_H | 6.037 | 5.371 | 9.64 | 5.206 | 5.596 | 9.112 | 1.01 | 121.52 | 164.85 |
| 451 | 2jsa-9 | LEU_A_77_N | LEU_A_77_CA | 6.037 | 5.371 | 9.64 | 6.667 | 4.077 | 9.434 | 1.45 | 98.15 | -64.04 |
| 452 | 2jsa-9 | LEU_A_77_CA | LEU_A_77_HA | 6.667 | 4.077 | 9.434 | 7.153 | 3.786 | 10.366 | 1.09 | 31.29 | -30.91 |
| 453 | 2jsa-9 | LEU_A_77_CA | LEU_A_77_C | 6.667 | 4.077 | 9.434 | 7.753 | 4.208 | 8.365 | 1.53 | 134.34 | 6.88 |
| 454 | 2jsa-9 | LEU_A_77_C | LEU_A_77_O | 7.753 | 4.208 | 8.365 | 8.196 | 3.209 | 7.799 | 1.23 | 117.38 | -66.09 |
| 455 | 2jsa-9 | LEU_A_77_C | ASN_A_78_N | 7.753 | 4.208 | 8.365 | 8.152 | 5.447 | 8.12 | 1.32 | 100.66 | 72.15 |
| 456 | 2jsa-9 | ASN_A_78_N | ASN_A_78_H | 8.152 | 5.447 | 8.12 | 7.744 | 6.221 | 8.624 | 1.01 | 60.06 | 117.80 |
| 457 | 2jsa-9 | ASN_A_78_N | ASN_A_78_CA | 8.152 | 5.447 | 8.12 | 9.179 | 5.721 | 7.129 | 1.45 | 132.99 | 14.94 |
| 458 | 2jsa-9 | ASN_A_78_CA | ASN_A_78_HA | 9.179 | 5.721 | 7.129 | 9.227 | 6.796 | 6.953 | 1.09 | 99.29 | 87.44 |
| 459 | 2jsa-9 | ASN_A_78_CA | ASN_A_78_C | 9.179 | 5.721 | 7.129 | 8.832 | 4.996 | 5.828 | 1.53 | 148.29 | -115.58 |
| 460 | 2jsa-9 | ASN_A_78_C | ASN_A_78_O | 8.832 | 4.996 | 5.828 | 9.712 | 4.456 | 5.159 | 1.23 | 122.94 | -31.53 |
| 461 | 2jsa-9 | ASN_A_78_C | GLU_A_79_N | 8.832 | 4.996 | 5.828 | 7.546 | 5.007 | 5.507 | 1.33 | 104.01 | 179.51 |
| 462 | 2jsa-9 | GLU_A_79_N | GLU_A_79_H | 7.546 | 5.007 | 5.507 | 6.877 | 5.472 | 6.104 | 1.01 | 53.77 | 145.20 |
| 463 | 2jsa-9 | GLU_A_79_N | GLU_A_79_CA | 7.546 | 5.007 | 5.507 | 7.071 | 4.356 | 4.297 | 1.45 | 146.34 | -126.12 |
| 464 | 2jsa-9 | GLU_A_79_CA | GLU_A_79_HA | 7.071 | 4.356 | 4.297 | 7.914 | 3.838 | 3.84 | 1.09 | 114.79 | -31.57 |
| 465 | 2jsa-9 | GLU_A_79_CA | GLU_A_79_C | 7.071 | 4.356 | 4.297 | 6.55 | 5.398 | 3.304 | 1.53 | 130.44 | 116.57 |
| 466 | 2jsa-9 | GLU_A_79_C | GLU_A_79_O | 6.55 | 5.398 | 3.304 | 5.358 | 5.434 | 3.005 | 1.23 | 104.08 | 178.27 |
| 467 | 2jsa-9 | GLU_A_79_C | CYS_A_80_N | 6.55 | 5.398 | 3.304 | 7.471 | 6.219 | 2.82 | 1.33 | 111.42 | 41.71 |
| 468 | 2jsa-9 | CYS_A_80_N | CYS_A_80_H | 7.471 | 6.219 | 2.82 | 8.435 | 6.13 | 3.11 | 1.01 | 73.32 | -5.27 |
| 469 | 2jsa-9 | CYS_A_80_N | CYS_A_80_CA | 7.471 | 6.219 | 2.82 | 7.121 | 7.258 | 1.867 | 1.45 | 131.00 | 108.62 |
| 470 | 2jsa-9 | CYS_A_80_CA | CYS_A_80_HA | 7.121 | 7.258 | 1.867 | 6.747 | 6.78 | 0.962 | 1.09 | 146.15 | -128.04 |
| 471 | 2jsa-9 | CYS_A_80_CA | CYS_A_80_C | 7.121 | 7.258 | 1.867 | 8.39 | 8.036 | 1.516 | 1.53 | 103.27 | 31.51 |
| 472 | 2jsa-9 | CYS_A_80_C | CYS_A_80_O | 8.39 | 8.036 | 1.516 | 8.484 | 9.232 | 1.791 | 1.23 | 77.09 | 85.51 |
| 473 | 2jsa-9 | CYS_A_80_C | PRO_A_81_N | 8.39 | 8.036 | 1.516 | 9.359 | 7.309 | 0.899 | 1.36 | 116.99 | -36.88 |
| 474 | 2jsa-9 | PRO_A_81_N | PRO_A_81_CA | 9.359 | 7.309 | 0.899 | 10.618 | 7.919 | 0.509 | 1.45 | 105.58 | 25.85 |
| 475 | 2jsa-9 | PRO_A_81_CA | PRO_A_81_HA | 10.618 | 7.919 | 0.509 | 11.0 | 8.539 | 1.321 | 1.09 | 41.89 | 58.36 |
| 476 | 2jsa-9 | PRO_A_81_CA | PRO_A_81_C | 10.618 | 7.919 | 0.509 | 10.441 | 8.798 | -0.731 | 1.53 | 144.13 | 101.39 |
| 477 | 2jsa-9 | PRO_A_81_C | PRO_A_81_O | 10.441 | 8.798 | -0.731 | 9.569 | 9.665 | -0.762 | 1.23 | 91.44 | 135.16 |
| 1 | 2jsa-10 | ARG_A_1_N | ARG_A_1_H | 16.471 | 2.049 | 0.959 | 17.049 | 2.625 | 0.364 | 1.01 | 126.10 | 44.90 |
| 2 | 2jsa-10 | ARG_A_1_N | ARG_A_1_CA | 16.471 | 2.049 | 0.959 | 15.098 | 2.458 | 1.199 | 1.45 | 80.49 | 163.41 |
| 3 | 2jsa-10 | ARG_A_1_CA | ARG_A_1_HA | 15.098 | 2.458 | 1.199 | 14.437 | 1.672 | 0.834 | 1.09 | 109.57 | -130.06 |
| 4 | 2jsa-10 | ARG_A_1_CA | ARG_A_1_C | 15.098 | 2.458 | 1.199 | 14.79 | 3.756 | 0.449 | 1.53 | 119.34 | 103.35 |
| 5 | 2jsa-10 | ARG_A_1_C | ARG_A_1_O | 14.79 | 3.756 | 0.449 | 15.627 | 4.255 | -0.304 | 1.23 | 127.69 | 30.80 |
| 6 | 2jsa-10 | ARG_A_1_C | SER_A_2_N | 14.79 | 3.756 | 0.449 | 13.59 | 4.266 | 0.68 | 1.32 | 79.95 | 156.97 |
| 7 | 2jsa-10 | SER_A_2_N | SER_A_2_H | 13.59 | 4.266 | 0.68 | 12.954 | 3.802 | 1.313 | 1.01 | 51.20 | -143.89 |
| 8 | 2jsa-10 | SER_A_2_N | SER_A_2_CA | 13.59 | 4.266 | 0.68 | 13.162 | 5.496 | 0.036 | 1.45 | 116.31 | 109.19 |
| 9 | 2jsa-10 | SER_A_2_CA | SER_A_2_HA | 13.162 | 5.496 | 0.036 | 12.179 | 5.762 | 0.425 | 1.09 | 69.09 | 164.86 |
| 10 | 2jsa-10 | SER_A_2_CA | SER_A_2_C | 13.162 | 5.496 | 0.036 | 13.051 | 5.283 | -1.475 | 1.53 | 170.97 | -117.53 |
| 11 | 2jsa-10 | SER_A_2_C | SER_A_2_O | 13.051 | 5.283 | -1.475 | 14.061 | 5.26 | -2.178 | 1.23 | 124.83 | -1.30 |
| 12 | 2jsa-10 | SER_A_2_C | ALA_A_3_N | 13.051 | 5.283 | -1.475 | 11.816 | 5.134 | -1.93 | 1.32 | 110.09 | -173.12 |
| 13 | 2jsa-10 | ALA_A_3_N | ALA_A_3_H | 11.816 | 5.134 | -1.93 | 11.033 | 5.164 | -1.293 | 1.01 | 50.89 | 177.81 |
| 14 | 2jsa-10 | ALA_A_3_N | ALA_A_3_CA | 11.816 | 5.134 | -1.93 | 11.56 | 4.924 | -3.346 | 1.45 | 166.84 | -140.64 |
| 15 | 2jsa-10 | ALA_A_3_CA | ALA_A_3_HA | 11.56 | 4.924 | -3.346 | 12.114 | 5.669 | -3.917 | 1.09 | 121.59 | 53.36 |
| 16 | 2jsa-10 | ALA_A_3_CA | ALA_A_3_C | 11.56 | 4.924 | -3.346 | 10.067 | 5.114 | -3.622 | 1.53 | 100.39 | 172.75 |
| 17 | 2jsa-10 | ALA_A_3_C | ALA_A_3_O | 10.067 | 5.114 | -3.622 | 9.386 | 5.843 | -2.903 | 1.23 | 54.22 | 133.05 |
| 18 | 2jsa-10 | ALA_A_3_C | LEU_A_4_N | 10.067 | 5.114 | -3.622 | 9.602 | 4.445 | -4.668 | 1.33 | 142.08 | -124.80 |
| 19 | 2jsa-10 | LEU_A_4_N | LEU_A_4_H | 9.602 | 4.445 | -4.668 | 10.22 | 3.864 | -5.216 | 1.01 | 122.86 | -43.23 |
| 20 | 2jsa-10 | LEU_A_4_N | LEU_A_4_CA | 9.602 | 4.445 | -4.668 | 8.203 | 4.531 | -5.049 | 1.45 | 105.21 | 176.48 |
| 21 | 2jsa-10 | LEU_A_4_CA | LEU_A_4_HA | 8.203 | 4.531 | -5.049 | 7.929 | 5.585 | -5.092 | 1.09 | 92.26 | 104.57 |
| 22 | 2jsa-10 | LEU_A_4_CA | LEU_A_4_C | 8.203 | 4.531 | -5.049 | 7.344 | 3.87 | -3.969 | 1.53 | 45.10 | -142.42 |
| 23 | 2jsa-10 | LEU_A_4_C | LEU_A_4_O | 7.344 | 3.87 | -3.969 | 6.119 | 3.981 | -3.992 | 1.23 | 91.07 | 174.82 |
| 24 | 2jsa-10 | LEU_A_4_C | SER_A_5_N | 7.344 | 3.87 | -3.969 | 8.02 | 3.197 | -3.049 | 1.33 | 46.04 | -44.87 |
| 25 | 2jsa-10 | SER_A_5_N | SER_A_5_H | 8.02 | 3.197 | -3.049 | 9.028 | 3.148 | -3.092 | 1.01 | 92.44 | -2.78 |
| 26 | 2jsa-10 | SER_A_5_N | SER_A_5_CA | 8.02 | 3.197 | -3.049 | 7.333 | 2.519 | -1.963 | 1.45 | 41.63 | -135.38 |

| | | | | | | | | | | | |
|---|---|---|---|---|---|---|---|---|---|---|---|
| 27 | 2jsa-10 | SER_A_5_CA | SER_A_5_HA | 7.333 | 2.519 | -1.963 | 6.852 | 1.629 | -2.367 | 1.09 | 111.77 | -118.39 |
| 28 | 2jsa-10 | SER_A_5_CA | SER_A_5_C | 7.333 | 2.519 | -1.963 | 6.257 | 3.433 | -1.373 | 1.53 | 67.32 | 139.65 |
| 29 | 2jsa-10 | SER_A_5_C | SER_A_5_O | 6.257 | 3.433 | -1.373 | 5.212 | 2.961 | -0.928 | 1.23 | 68.79 | -155.69 |
| 30 | 2jsa-10 | SER_A_5_C | CYS_A_6_N | 6.257 | 3.433 | -1.373 | 6.551 | 4.725 | -1.386 | 1.33 | 90.56 | 77.18 |
| 31 | 2jsa-10 | CYS_A_6_N | CYS_A_6_H | 6.551 | 4.725 | -1.386 | 7.431 | 5.043 | -1.768 | 1.01 | 112.21 | 19.87 |
| 32 | 2jsa-10 | CYS_A_6_N | CYS_A_6_CA | 6.551 | 4.725 | -1.386 | 5.623 | 5.709 | -0.857 | 1.45 | 68.64 | 133.32 |
| 33 | 2jsa-10 | CYS_A_6_CA | CYS_A_6_HA | 5.623 | 5.709 | -0.857 | 5.452 | 5.487 | 0.197 | 1.09 | 14.89 | -127.61 |
| 34 | 2jsa-10 | CYS_A_6_CA | CYS_A_6_C | 5.623 | 5.709 | -0.857 | 4.295 | 5.569 | -1.606 | 1.53 | 119.29 | -173.98 |
| 35 | 2jsa-10 | CYS_A_6_C | CYS_A_6_O | 4.295 | 5.569 | -1.606 | 3.279 | 5.211 | -1.014 | 1.23 | 61.21 | -160.59 |
| 36 | 2jsa-10 | CYS_A_6_C | GLN_A_7_N | 4.295 | 5.569 | -1.606 | 4.348 | 5.861 | -2.898 | 1.33 | 167.06 | 79.71 |
| 37 | 2jsa-10 | GLN_A_7_N | GLN_A_7_H | 4.348 | 5.861 | -2.898 | 5.219 | 6.152 | -3.318 | 1.01 | 114.58 | 18.47 |
| 38 | 2jsa-10 | GLN_A_7_N | GLN_A_7_CA | 4.348 | 5.861 | -2.898 | 3.164 | 5.772 | -3.734 | 1.45 | 125.15 | -175.70 |
| 39 | 2jsa-10 | GLN_A_7_CA | GLN_A_7_HA | 3.164 | 5.772 | -3.734 | 2.558 | 6.66 | -3.555 | 1.09 | 80.55 | 124.31 |
| 40 | 2jsa-10 | GLN_A_7_CA | GLN_A_7_C | 3.164 | 5.772 | -3.734 | 2.337 | 4.542 | -3.355 | 1.53 | 75.66 | -123.92 |
| 41 | 2jsa-10 | GLN_A_7_C | GLN_A_7_O | 2.337 | 4.542 | -3.355 | 1.142 | 4.652 | -3.087 | 1.23 | 77.41 | 174.74 |
| 42 | 2jsa-10 | GLN_A_7_C | MET_A_8_N | 2.337 | 4.542 | -3.355 | 3.007 | 3.399 | -3.344 | 1.32 | 89.52 | -59.62 |
| 43 | 2jsa-10 | MET_A_8_N | MET_A_8_H | 3.007 | 3.399 | -3.344 | 3.99 | 3.383 | -3.576 | 1.01 | 103.28 | -0.93 |
| 44 | 2jsa-10 | MET_A_8_N | MET_A_8_CA | 3.007 | 3.399 | -3.344 | 2.349 | 2.149 | -3.002 | 1.45 | 76.39 | -117.76 |
| 45 | 2jsa-10 | MET_A_8_CA | MET_A_8_HA | 2.349 | 2.149 | -3.002 | 1.726 | 1.841 | -3.841 | 1.09 | 140.36 | -153.69 |
| 46 | 2jsa-10 | MET_A_8_CA | MET_A_8_C | 2.349 | 2.149 | -3.002 | 1.467 | 2.316 | -1.763 | 1.53 | 35.92 | 169.28 |
| 47 | 2jsa-10 | MET_A_8_C | MET_A_8_O | 1.467 | 2.316 | -1.763 | 0.395 | 1.717 | -1.674 | 1.23 | 85.85 | -150.80 |
| 48 | 2jsa-10 | MET_A_8_C | CYS_A_9_N | 1.467 | 2.316 | -1.763 | 1.948 | 3.132 | -0.838 | 1.32 | 45.68 | 59.48 |
| 49 | 2jsa-10 | CYS_A_9_N | CYS_A_9_H | 1.948 | 3.132 | -0.838 | 2.836 | 3.592 | -0.979 | 1.01 | 98.03 | 27.38 |
| 50 | 2jsa-10 | CYS_A_9_N | CYS_A_9_CA | 1.948 | 3.132 | -0.838 | 1.217 | 3.386 | 0.393 | 1.45 | 32.16 | 160.84 |
| 51 | 2jsa-10 | CYS_A_9_CA | CYS_A_9_HA | 1.217 | 3.386 | 0.393 | 1.005 | 2.429 | 0.87 | 1.09 | 64.05 | -102.49 |
| 52 | 2jsa-10 | CYS_A_9_CA | CYS_A_9_C | 1.217 | 3.386 | 0.393 | -0.111 | 4.055 | 0.032 | 1.53 | 103.65 | 153.26 |
| 53 | 2jsa-10 | CYS_A_9_C | CYS_A_9_O | -0.111 | 4.055 | 0.032 | -1.15 | 3.731 | 0.606 | 1.23 | 62.19 | -162.68 |
| 54 | 2jsa-10 | CYS_A_9_C | GLU_A_10_N | -0.111 | 4.055 | 0.032 | -0.035 | 4.977 | -0.917 | 1.33 | 135.73 | 85.29 |
| 55 | 2jsa-10 | GLU_A_10_N | GLU_A_10_H | -0.035 | 4.977 | -0.917 | 0.854 | 5.192 | -1.346 | 1.01 | 115.13 | 13.60 |
| 56 | 2jsa-10 | GLU_A_10_N | GLU_A_10_CA | -0.035 | 4.977 | -0.917 | -1.217 | 5.694 | -1.36 | 1.45 | 107.77 | 148.76 |
| 57 | 2jsa-10 | GLU_A_10_CA | GLU_A_10_HA | -1.217 | 5.694 | -1.36 | -1.745 | 6.053 | -0.476 | 1.09 | 35.84 | 145.79 |
| 58 | 2jsa-10 | GLU_A_10_CA | GLU_A_10_C | -1.217 | 5.694 | -1.36 | -2.149 | 4.757 | -2.13 | 1.53 | 120.23 | -134.85 |
| 59 | 2jsa-10 | GLU_A_10_C | GLU_A_10_O | -2.149 | 4.757 | -2.13 | -3.261 | 5.141 | -2.492 | 1.23 | 107.10 | 160.95 |
| 60 | 2jsa-10 | GLU_A_10_C | LEU_A_11_N | -2.149 | 4.757 | -2.13 | -1.663 | 3.547 | -2.359 | 1.32 | 99.96 | -68.12 |
| 61 | 2jsa-10 | LEU_A_11_N | LEU_A_11_H | -1.663 | 3.547 | -2.359 | -0.739 | 3.3 | -2.032 | 1.01 | 71.12 | -14.97 |
| 62 | 2jsa-10 | LEU_A_11_N | LEU_A_11_CA | -1.663 | 3.547 | -2.359 | -2.438 | 2.552 | -3.079 | 1.45 | 119.72 | -127.91 |
| 63 | 2jsa-10 | LEU_A_11_CA | LEU_A_11_HA | -2.438 | 2.552 | -3.079 | -3.177 | 3.075 | -3.685 | 1.09 | 123.80 | 144.71 |
| 64 | 2jsa-10 | LEU_A_11_CA | LEU_A_11_C | -2.438 | 2.552 | -3.079 | -3.194 | 1.677 | -2.077 | 1.53 | 49.09 | -130.83 |
| 65 | 2jsa-10 | LEU_A_11_C | LEU_A_11_O | -3.194 | 1.677 | -2.077 | -4.349 | 1.319 | -2.307 | 1.23 | 100.77 | -162.78 |
| 66 | 2jsa-10 | LEU_A_11_C | VAL_A_12_N | -3.194 | 1.677 | -2.077 | -2.512 | 1.358 | -0.987 | 1.32 | 34.63 | -25.07 |
| 67 | 2jsa-10 | VAL_A_12_N | VAL_A_12_H | -2.512 | 1.358 | -0.987 | -1.565 | 1.688 | -0.864 | 1.01 | 83.01 | 19.21 |
| 68 | 2jsa-10 | VAL_A_12_N | VAL_A_12_CA | -2.512 | 1.358 | -0.987 | -3.104 | 0.532 | 0.051 | 1.45 | 44.39 | -125.63 |
| 69 | 2jsa-10 | VAL_A_12_CA | VAL_A_12_HA | -3.104 | 0.532 | 0.051 | -3.639 | -0.289 | -0.426 | 1.09 | 115.96 | -123.09 |
| 70 | 2jsa-10 | VAL_A_12_CA | VAL_A_12_C | -3.104 | 0.532 | 0.051 | -4.117 | 1.363 | 0.842 | 1.53 | 58.88 | 140.64 |
| 71 | 2jsa-10 | VAL_A_12_C | VAL_A_12_O | -4.117 | 1.363 | 0.842 | -5.209 | 0.887 | 1.152 | 1.23 | 75.41 | -156.45 |
| 72 | 2jsa-10 | VAL_A_12_C | VAL_A_13_N | -4.117 | 1.363 | 0.842 | -3.719 | 2.589 | 1.147 | 1.32 | 76.69 | 72.01 |
| 73 | 2jsa-10 | VAL_A_13_N | VAL_A_13_H | -3.719 | 2.589 | 1.147 | -2.807 | 2.914 | 0.859 | 1.01 | 106.57 | 19.61 |
| 74 | 2jsa-10 | VAL_A_13_N | VAL_A_13_CA | -3.719 | 2.589 | 1.147 | -4.577 | 3.49 | 1.897 | 1.45 | 58.92 | 133.60 |
| 75 | 2jsa-10 | VAL_A_13_CA | VAL_A_13_HA | -4.577 | 3.49 | 1.897 | -4.911 | 2.977 | 2.799 | 1.09 | 34.16 | -123.07 |
| 76 | 2jsa-10 | VAL_A_13_CA | VAL_A_13_C | -4.577 | 3.49 | 1.897 | -5.811 | 3.828 | 1.058 | 1.53 | 123.25 | 164.68 |
| 77 | 2jsa-10 | VAL_A_13_C | VAL_A_13_O | -5.811 | 3.828 | 1.058 | -6.937 | 3.545 | 1.461 | 1.23 | 70.86 | -165.89 |
| 78 | 2jsa-10 | VAL_A_13_C | LYS_A_14_N | -5.811 | 3.828 | 1.058 | -5.556 | 4.43 | -0.096 | 1.33 | 150.47 | 67.04 |
| 79 | 2jsa-10 | LYS_A_14_N | LYS_A_14_H | -5.556 | 4.43 | -0.096 | -4.603 | 4.633 | -0.364 | 1.01 | 105.38 | 12.02 |
| 80 | 2jsa-10 | LYS_A_14_N | LYS_A_14_CA | -5.556 | 4.43 | -0.096 | -6.632 | 4.809 | -0.995 | 1.45 | 128.24 | 160.60 |
| 81 | 2jsa-10 | LYS_A_14_CA | LYS_A_14_HA | -6.632 | 4.809 | -0.995 | -7.164 | 5.65 | -0.549 | 1.09 | 65.86 | 122.32 |
| 82 | 2jsa-10 | LYS_A_14_CA | LYS_A_14_C | -6.632 | 4.809 | -0.995 | -7.62 | 3.648 | -1.121 | 1.53 | 94.72 | -130.40 |
| 83 | 2jsa-10 | LYS_A_14_C | LYS_A_14_O | -7.62 | 3.648 | -1.121 | -8.817 | 3.864 | -1.297 | 1.23 | 98.23 | 169.77 |
| 84 | 2jsa-10 | LYS_A_14_C | LYS_A_15_N | -7.62 | 3.648 | -1.121 | -7.081 | 2.441 | -1.024 | 1.33 | 85.80 | -65.94 |
| 85 | 2jsa-10 | LYS_A_15_N | LYS_A_15_H | -7.081 | 2.441 | -1.024 | -6.087 | 2.338 | -0.878 | 1.01 | 81.69 | -5.92 |
| 86 | 2jsa-10 | LYS_A_15_N | LYS_A_15_CA | -7.081 | 2.441 | -1.024 | -7.9 | 1.246 | -1.124 | 1.45 | 93.95 | -124.43 |
| 87 | 2jsa-10 | LYS_A_15_CA | LYS_A_15_HA | -7.9 | 1.246 | -1.124 | -8.682 | 1.434 | -1.859 | 1.09 | 132.42 | 166.48 |
| 88 | 2jsa-10 | LYS_A_15_CA | LYS_A_15_C | -7.9 | 1.246 | -1.124 | -8.58 | 0.983 | 0.221 | 1.53 | 28.46 | -158.86 |
| 89 | 2jsa-10 | LYS_A_15_C | LYS_A_15_O | -8.58 | 0.983 | 0.221 | -9.714 | 0.51 | 0.266 | 1.23 | 87.90 | -157.36 |
| 90 | 2jsa-10 | LYS_A_15_C | TYR_A_16_N | -8.58 | 0.983 | 0.221 | -7.858 | 1.304 | 1.285 | 1.33 | 36.60 | 23.97 |
| 91 | 2jsa-10 | TYR_A_16_N | TYR_A_16_H | -7.858 | 1.304 | 1.285 | -6.931 | 1.689 | 1.173 | 1.01 | 96.37 | 22.55 |
| 92 | 2jsa-10 | TYR_A_16_N | TYR_A_16_CA | -7.858 | 1.304 | 1.285 | -8.377 | 1.11 | 2.629 | 1.45 | 22.40 | -159.50 |
| 93 | 2jsa-10 | TYR_A_16_CA | TYR_A_16_HA | -8.377 | 1.11 | 2.629 | -8.866 | 0.139 | 2.708 | 1.09 | 85.84 | -116.73 |
| 94 | 2jsa-10 | TYR_A_16_CA | TYR_A_16_C | -8.377 | 1.11 | 2.629 | -9.362 | 2.22 | 3.002 | 1.53 | 75.89 | 131.59 |
| 95 | 2jsa-10 | TYR_A_16_C | TYR_A_16_O | -9.362 | 2.22 | 3.002 | -10.562 | 1.977 | 3.116 | 1.23 | 84.68 | -168.55 |
| 96 | 2jsa-10 | TYR_A_16_C | GLU_A_17_N | -9.362 | 2.22 | 3.002 | -8.817 | 3.414 | 3.18 | 1.32 | 82.28 | 65.47 |
| 97 | 2jsa-10 | GLU_A_17_N | GLU_A_17_H | -8.817 | 3.414 | 3.18 | -7.821 | 3.541 | 3.067 | 1.01 | 96.42 | 7.27 |
| 98 | 2jsa-10 | GLU_A_17_N | GLU_A_17_CA | -8.817 | 3.414 | 3.18 | -9.632 | 4.562 | 3.539 | 1.45 | 75.69 | 125.37 |
| 99 | 2jsa-10 | GLU_A_17_CA | GLU_A_17_HA | -9.632 | 4.562 | 3.539 | -10.154 | 4.328 | 4.467 | 1.09 | 31.65 | -155.85 |
| 100 | 2jsa-10 | GLU_A_17_CA | GLU_A_17_C | -9.632 | 4.562 | 3.539 | -10.677 | 4.831 | 2.453 | 1.53 | 135.18 | 165.56 |
| 101 | 2jsa-10 | GLU_A_17_C | GLU_A_17_O | -10.677 | 4.831 | 2.453 | -11.653 | 5.542 | 2.688 | 1.23 | 78.99 | 143.93 |
| 102 | 2jsa-10 | GLU_A_17_C | GLY_A_18_N | -10.677 | 4.831 | 2.453 | -10.437 | 4.247 | 1.288 | 1.33 | 151.54 | -67.66 |
| 103 | 2jsa-10 | GLY_A_18_N | GLY_A_18_H | -10.437 | 4.247 | 1.288 | -9.614 | 3.674 | 1.167 | 1.01 | 96.88 | -34.85 |
| 104 | 2jsa-10 | GLY_A_18_N | GLY_A_18_CA | -10.437 | 4.247 | 1.288 | -11.344 | 4.415 | 0.165 | 1.45 | 140.60 | 169.51 |
| 105 | 2jsa-10 | GLY_A_18_CA | GLY_A_18_C | -11.344 | 4.415 | 0.165 | -11.871 | 3.063 | -0.319 | 1.53 | 108.45 | -111.30 |
| 106 | 2jsa-10 | GLY_A_18_C | GLY_A_18_O | -11.871 | 3.063 | -0.319 | -12.163 | 2.894 | -1.503 | 1.23 | 164.10 | -149.94 |
| 107 | 2jsa-10 | GLY_A_18_C | SER_A_19_N | -11.871 | 3.063 | -0.319 | -11.979 | 2.134 | 0.621 | 1.33 | 44.86 | -96.63 |
| 108 | 2jsa-10 | SER_A_19_N | SER_A_19_H | -11.979 | 2.134 | 0.621 | -11.722 | 2.343 | 1.575 | 1.01 | 19.15 | 39.12 |
| 109 | 2jsa-10 | SER_A_19_N | SER_A_19_CA | -11.979 | 2.134 | 0.621 | -12.467 | 0.802 | 0.304 | 1.45 | 102.60 | -110.12 |
| 110 | 2jsa-10 | SER_A_19_CA | SER_A_19_HA | -12.467 | 0.802 | 0.304 | -12.102 | 0.531 | -0.687 | 1.09 | 155.36 | -36.59 |
| 111 | 2jsa-10 | SER_A_19_CA | SER_A_19_C | -12.467 | 0.802 | 0.304 | -13.996 | 0.798 | 0.281 | 1.53 | 90.86 | -179.85 |
| 112 | 2jsa-10 | SER_A_19_C | SER_A_19_O | -13.996 | 0.798 | 0.281 | -14.605 | 0.389 | -0.707 | 1.23 | 143.41 | -146.11 |
| 113 | 2jsa-10 | SER_A_19_C | ALA_A_20_N | -13.996 | 0.798 | 0.281 | -14.574 | 1.256 | 1.381 | 1.32 | 33.84 | 141.61 |
| 114 | 2jsa-10 | ALA_A_20_N | ALA_A_20_H | -14.574 | 1.256 | 1.381 | -14.01 | 1.575 | 2.156 | 1.01 | 39.90 | 29.49 |
| 115 | 2jsa-10 | ALA_A_20_N | ALA_A_20_CA | -14.574 | 1.256 | 1.381 | -16.022 | 1.309 | 1.5 | 1.45 | 85.30 | 177.90 |
| 116 | 2jsa-10 | ALA_A_20_CA | ALA_A_20_HA | -16.022 | 1.309 | 1.5 | -16.285 | 2.053 | 2.252 | 1.09 | 46.38 | 109.47 |
| 117 | 2jsa-10 | ALA_A_20_CA | ALA_A_20_C | -16.022 | 1.309 | 1.5 | -16.542 | -0.053 | 1.964 | 1.53 | 72.35 | -110.90 |
| 118 | 2jsa-10 | ALA_A_20_C | ALA_A_20_O | -16.542 | -0.053 | 1.964 | -17.727 | -0.202 | 2.257 | 1.23 | 76.22 | -172.83 |
| 119 | 2jsa-10 | ALA_A_20_C | ASP_A_21_N | -16.542 | -0.053 | 1.964 | -15.629 | -1.012 | 2.018 | 1.33 | 87.66 | -46.41 |
| 120 | 2jsa-10 | ASP_A_21_N | ASP_A_21_H | -15.629 | -1.012 | 2.018 | -14.671 | -0.818 | 1.762 | 1.01 | 104.68 | 11.45 |
| 121 | 2jsa-10 | ASP_A_21_N | ASP_A_21_CA | -15.629 | -1.012 | 2.018 | -15.98 | -2.357 | 2.443 | 1.45 | 73.00 | -104.63 |
| 122 | 2jsa-10 | ASP_A_21_CA | ASP_A_21_HA | -15.98 | -2.357 | 2.443 | -17.065 | -2.44 | 2.502 | 1.09 | 86.90 | -175.63 |

| | | | | | | | | | | | | |
|---|---|---|---|---|---|---|---|---|---|---|---|---|
| 123 | 2jsa-10 | ASP_A_21_CA | ASP_A_21_C | -15.98 | -2.357 | 2.443 | -15.362 | -2.631 | 3.815 | 1.53 | 26.23 | -23.91 |
| 124 | 2jsa-10 | ASP_A_21_C | ASP_A_21_O | -15.362 | -2.631 | 3.815 | -15.828 | -3.504 | 4.546 | 1.23 | 53.55 | -118.09 |
| 125 | 2jsa-10 | ASP_A_21_C | LYS_A_22_N | -15.362 | -2.631 | 3.815 | -14.323 | -1.87 | 4.124 | 1.32 | 76.51 | 36.22 |
| 126 | 2jsa-10 | LYS_A_22_N | LYS_A_22_H | -14.323 | -1.87 | 4.124 | -13.993 | -1.171 | 3.474 | 1.01 | 130.06 | 64.73 |
| 127 | 2jsa-10 | LYS_A_22_N | LYS_A_22_CA | -14.323 | -1.87 | 4.124 | -13.637 | -2.021 | 5.396 | 1.45 | 28.91 | -12.41 |
| 128 | 2jsa-10 | LYS_A_22_CA | LYS_A_22_HA | -13.637 | -2.021 | 5.396 | -12.776 | -1.352 | 5.396 | 1.09 | 90.00 | 37.85 |
| 129 | 2jsa-10 | LYS_A_22_CA | LYS_A_22_C | -13.637 | -2.021 | 5.396 | -13.112 | -3.452 | 5.522 | 1.53 | 85.27 | -69.85 |
| 130 | 2jsa-10 | LYS_A_22_C | LYS_A_22_O | -13.112 | -3.452 | 5.522 | -13.557 | -4.207 | 6.385 | 1.23 | 45.44 | -120.52 |
| 131 | 2jsa-10 | LYS_A_22_C | ASP_A_23_N | -13.112 | -3.452 | 5.522 | -12.173 | -3.784 | 4.647 | 1.33 | 131.30 | -19.47 |
| 132 | 2jsa-10 | ASP_A_23_N | ASP_A_23_H | -12.173 | -3.784 | 4.647 | -11.854 | -3.112 | 3.963 | 1.01 | 132.60 | 64.61 |
| 133 | 2jsa-10 | ASP_A_23_N | ASP_A_23_CA | -12.173 | -3.784 | 4.647 | -11.584 | -5.112 | 4.649 | 1.45 | 89.92 | -66.08 |
| 134 | 2jsa-10 | ASP_A_23_CA | ASP_A_23_HA | -11.584 | -5.112 | 4.649 | -11.612 | -5.512 | 5.663 | 1.09 | 21.58 | -94.00 |
| 135 | 2jsa-10 | ASP_A_23_CA | ASP_A_23_C | -11.584 | -5.112 | 4.649 | -10.135 | -5.022 | 4.165 | 1.53 | 108.44 | 3.55 |
| 136 | 2jsa-10 | ASP_A_23_C | ASP_A_23_O | -10.135 | -5.022 | 4.165 | -9.885 | -4.777 | 2.986 | 1.23 | 163.46 | 44.42 |
| 137 | 2jsa-10 | ASP_A_23_C | ALA_A_24_N | -10.135 | -5.022 | 4.165 | -9.218 | -5.228 | 5.1 | 1.33 | 45.15 | -12.66 |
| 138 | 2jsa-10 | ALA_A_24_N | ALA_A_24_H | -9.218 | -5.228 | 5.1 | -9.496 | -5.425 | 6.051 | 1.01 | 19.71 | -144.68 |
| 139 | 2jsa-10 | ALA_A_24_N | ALA_A_24_CA | -9.218 | -5.228 | 5.1 | -7.802 | -5.174 | 4.783 | 1.45 | 102.61 | 2.18 |
| 140 | 2jsa-10 | ALA_A_24_CA | ALA_A_24_HA | -7.802 | -5.174 | 4.783 | -7.575 | -4.196 | 4.358 | 1.09 | 112.94 | 76.93 |
| 141 | 2jsa-10 | ALA_A_24_CA | ALA_A_24_C | -7.802 | -5.174 | 4.783 | -7.475 | -6.244 | 3.739 | 1.53 | 133.02 | -73.01 |
| 142 | 2jsa-10 | ALA_A_24_C | ALA_A_24_O | -7.475 | -6.244 | 3.739 | -6.395 | -6.234 | 3.152 | 1.23 | 118.52 | 0.53 |
| 143 | 2jsa-10 | ALA_A_24_C | ASN_A_25_N | -7.475 | -6.244 | 3.739 | -8.43 | -7.141 | 3.54 | 1.33 | 98.64 | -136.79 |
| 144 | 2jsa-10 | ASN_A_25_N | ASN_A_25_H | -8.43 | -7.141 | 3.54 | -9.294 | -7.087 | 4.059 | 1.01 | 59.06 | 176.42 |
| 145 | 2jsa-10 | ASN_A_25_N | ASN_A_25_CA | -8.43 | -7.141 | 3.54 | -8.258 | -8.216 | 2.577 | 1.45 | 131.49 | -80.91 |
| 146 | 2jsa-10 | ASN_A_25_CA | ASN_A_25_HA | -8.258 | -8.216 | 2.577 | -7.246 | -8.613 | 2.663 | 1.09 | 85.48 | -21.42 |
| 147 | 2jsa-10 | ASN_A_25_CA | ASN_A_25_C | -8.258 | -8.216 | 2.577 | -8.484 | -7.67 | 1.166 | 1.53 | 157.28 | 112.49 |
| 148 | 2jsa-10 | ASN_A_25_C | ASN_A_25_O | -8.484 | -7.67 | 1.166 | -7.619 | -7.798 | 0.301 | 1.23 | 134.69 | -8.42 |
| 149 | 2jsa-10 | ASN_A_25_C | VAL_A_26_N | -8.484 | -7.67 | 1.166 | -9.652 | -7.074 | 0.976 | 1.32 | 98.24 | 152.97 |
| 150 | 2jsa-10 | VAL_A_26_N | VAL_A_26_H | -9.652 | -7.074 | 0.976 | -10.316 | -7.005 | 1.734 | 1.01 | 41.37 | 174.07 |
| 151 | 2jsa-10 | VAL_A_26_N | VAL_A_26_CA | -9.652 | -7.074 | 0.976 | -10.004 | -6.51 | -0.316 | 1.45 | 152.77 | 121.97 |
| 152 | 2jsa-10 | VAL_A_26_CA | VAL_A_26_HA | -10.004 | -6.51 | -0.316 | -9.739 | -7.23 | -1.09 | 1.09 | 135.25 | -69.79 |
| 153 | 2jsa-10 | VAL_A_26_CA | VAL_A_26_C | -10.004 | -6.51 | -0.316 | -9.187 | -5.237 | -0.55 | 1.53 | 98.79 | 57.31 |
| 154 | 2jsa-10 | VAL_A_26_C | VAL_A_26_O | -9.187 | -5.237 | -0.55 | -9.255 | -4.641 | -1.624 | 1.23 | 150.82 | 96.51 |
| 155 | 2jsa-10 | VAL_A_26_C | ILE_A_27_N | -9.187 | -5.237 | -0.55 | -8.433 | -4.859 | 0.472 | 1.33 | 39.53 | 26.63 |
| 156 | 2jsa-10 | ILE_A_27_N | ILE_A_27_H | -8.433 | -4.859 | 0.472 | -8.425 | -5.398 | 1.326 | 1.01 | 32.26 | -89.15 |
| 157 | 2jsa-10 | ILE_A_27_N | ILE_A_27_CA | -8.433 | -4.859 | 0.472 | -7.603 | -3.669 | 0.39 | 1.45 | 93.23 | 55.11 |
| 158 | 2jsa-10 | ILE_A_27_CA | ILE_A_27_HA | -7.603 | -3.669 | 0.39 | -8.073 | -2.977 | -0.308 | 1.09 | 129.84 | 124.18 |
| 159 | 2jsa-10 | ILE_A_27_CA | ILE_A_27_C | -7.603 | -3.669 | 0.39 | -6.234 | -4.045 | -0.18 | 1.53 | 111.88 | -15.36 |
| 160 | 2jsa-10 | ILE_A_27_C | ILE_A_27_O | -6.234 | -4.045 | -0.18 | -5.888 | -3.643 | -1.289 | 1.23 | 154.44 | 49.28 |
| 161 | 2jsa-10 | ILE_A_27_C | LYS_A_28_N | -6.234 | -4.045 | -0.18 | -5.493 | -4.811 | 0.606 | 1.32 | 53.59 | -45.95 |
| 162 | 2jsa-10 | LYS_A_28_N | LYS_A_28_H | -5.493 | -4.811 | 0.606 | -5.842 | -5.104 | 1.507 | 1.01 | 26.83 | -139.99 |
| 163 | 2jsa-10 | LYS_A_28_N | LYS_A_28_CA | -5.493 | -4.811 | 0.606 | -4.168 | -5.246 | 0.193 | 1.45 | 106.50 | -18.18 |
| 164 | 2jsa-10 | LYS_A_28_CA | LYS_A_28_HA | -4.168 | -5.246 | 0.193 | -3.53 | -4.365 | 0.122 | 1.09 | 93.73 | 54.09 |
| 165 | 2jsa-10 | LYS_A_28_CA | LYS_A_28_C | -4.168 | -5.246 | 0.193 | -4.253 | -5.873 | -1.2 | 1.53 | 155.57 | -97.72 |
| 166 | 2jsa-10 | LYS_A_28_C | LYS_A_28_O | -4.253 | -5.873 | -1.2 | -3.354 | -5.694 | -2.02 | 1.23 | 131.81 | 11.26 |
| 167 | 2jsa-10 | LYS_A_28_C | LYS_A_29_N | -4.253 | -5.873 | -1.2 | -5.341 | -6.594 | -1.424 | 1.32 | 99.74 | -146.47 |
| 168 | 2jsa-10 | LYS_A_29_N | LYS_A_29_H | -5.341 | -6.594 | -1.424 | -6.039 | -6.698 | -0.702 | 1.01 | 44.35 | -171.53 |
| 169 | 2jsa-10 | LYS_A_29_N | LYS_A_29_CA | -5.341 | -6.594 | -1.424 | -5.555 | -7.249 | -2.704 | 1.45 | 151.70 | -108.09 |
| 170 | 2jsa-10 | LYS_A_29_CA | LYS_A_29_HA | -5.555 | -7.249 | -2.704 | -4.651 | -7.805 | -2.954 | 1.09 | 103.25 | -31.59 |
| 171 | 2jsa-10 | LYS_A_29_CA | LYS_A_29_C | -5.555 | -7.249 | -2.704 | -5.762 | -6.187 | -3.786 | 1.53 | 135.00 | 101.03 |
| 172 | 2jsa-10 | LYS_A_29_C | LYS_A_29_O | -5.762 | -6.187 | -3.786 | -5.145 | -6.25 | -4.849 | 1.23 | 149.74 | -5.83 |
| 173 | 2jsa-10 | LYS_A_29_C | ASP_A_30_N | -5.762 | -6.187 | -3.786 | -6.633 | -5.236 | -3.478 | 1.33 | 76.57 | 132.49 |
| 174 | 2jsa-10 | ASP_A_30_N | ASP_A_30_H | -6.633 | -5.236 | -3.478 | -7.105 | -5.248 | -2.585 | 1.01 | 27.87 | -178.54 |
| 175 | 2jsa-10 | ASP_A_30_N | ASP_A_30_CA | -6.633 | -5.236 | -3.478 | -6.928 | -4.162 | -4.411 | 1.45 | 129.95 | 105.36 |
| 176 | 2jsa-10 | ASP_A_30_CA | ASP_A_30_HA | -6.928 | -4.162 | -4.411 | -7.494 | -4.565 | -5.25 | 1.09 | 140.37 | -144.55 |
| 177 | 2jsa-10 | ASP_A_30_CA | ASP_A_30_C | -6.928 | -4.162 | -4.411 | -5.618 | -3.553 | -4.914 | 1.53 | 109.20 | 24.93 |
| 178 | 2jsa-10 | ASP_A_30_C | ASP_A_30_O | -5.618 | -3.553 | -4.914 | -5.399 | -3.455 | -6.121 | 1.23 | 168.76 | 24.11 |
| 179 | 2jsa-10 | ASP_A_30_C | PHE_A_31_N | -5.618 | -3.553 | -4.914 | -4.782 | -3.162 | -3.964 | 1.32 | 44.17 | 25.07 |
| 180 | 2jsa-10 | PHE_A_31_N | PHE_A_31_H | -4.782 | -3.162 | -3.964 | -5.028 | -3.275 | -2.991 | 1.01 | 15.55 | -155.33 |
| 181 | 2jsa-10 | PHE_A_31_N | PHE_A_31_CA | -4.782 | -3.162 | -3.964 | -3.499 | -2.565 | -4.296 | 1.45 | 103.20 | 24.95 |
| 182 | 2jsa-10 | PHE_A_31_CA | PHE_A_31_HA | -3.499 | -2.565 | -4.296 | -3.639 | -1.525 | -4.591 | 1.09 | 105.70 | 97.67 |
| 183 | 2jsa-10 | PHE_A_31_CA | PHE_A_31_C | -3.499 | -2.565 | -4.296 | -2.814 | -3.333 | -5.427 | 1.53 | 137.70 | -48.27 |
| 184 | 2jsa-10 | PHE_A_31_C | PHE_A_31_O | -2.814 | -3.333 | -5.427 | -2.502 | -2.763 | -6.471 | 1.23 | 148.10 | 61.31 |
| 185 | 2jsa-10 | PHE_A_31_C | ASP_A_32_N | -2.814 | -3.333 | -5.427 | -2.598 | -4.618 | -5.181 | 1.33 | 79.31 | -80.46 |
| 186 | 2jsa-10 | ASP_A_32_N | ASP_A_32_H | -2.598 | -4.618 | -5.181 | -2.88 | -5.022 | -4.299 | 1.01 | 29.19 | -124.92 |
| 187 | 2jsa-10 | ASP_A_32_N | ASP_A_32_CA | -2.598 | -4.618 | -5.181 | -1.954 | -5.47 | -6.166 | 1.45 | 132.68 | -52.92 |
| 188 | 2jsa-10 | ASP_A_32_CA | ASP_A_32_HA | -1.954 | -5.47 | -6.166 | -0.897 | -5.21 | -6.226 | 1.09 | 93.16 | 13.82 |
| 189 | 2jsa-10 | ASP_A_32_CA | ASP_A_32_C | -1.954 | -5.47 | -6.166 | -2.62 | -5.261 | -7.529 | 1.53 | 152.88 | 162.58 |
| 190 | 2jsa-10 | ASP_A_32_C | ASP_A_32_O | -2.62 | -5.261 | -7.529 | -1.981 | -5.425 | -8.567 | 1.23 | 147.56 | -14.39 |
| 191 | 2jsa-10 | ASP_A_32_C | ALA_A_33_N | -2.62 | -5.261 | -7.529 | -3.895 | -4.903 | -7.48 | 1.33 | 87.88 | 164.32 |
| 192 | 2jsa-10 | ALA_A_33_N | ALA_A_33_H | -3.895 | -4.903 | -7.48 | -4.357 | -4.788 | -6.589 | 1.01 | 28.12 | 166.02 |
| 193 | 2jsa-10 | ALA_A_33_N | ALA_A_33_CA | -3.895 | -4.903 | -7.48 | -4.654 | -4.67 | -8.696 | 1.45 | 146.86 | 162.93 |
| 194 | 2jsa-10 | ALA_A_33_CA | ALA_A_33_HA | -4.654 | -4.67 | -8.696 | -4.527 | -5.527 | -9.357 | 1.09 | 127.34 | -81.57 |
| 195 | 2jsa-10 | ALA_A_33_CA | ALA_A_33_C | -4.654 | -4.67 | -8.696 | -4.107 | -3.428 | -9.403 | 1.53 | 117.52 | 66.23 |
| 196 | 2jsa-10 | ALA_A_33_C | ALA_A_33_O | -4.107 | -3.428 | -9.403 | -3.631 | -3.512 | -10.534 | 1.23 | 156.86 | -10.01 |
| 197 | 2jsa-10 | ALA_A_33_C | GLU_A_34_N | -4.107 | -3.428 | -9.403 | -4.195 | -2.303 | -8.707 | 1.33 | 58.33 | 94.47 |
| 198 | 2jsa-10 | GLU_A_34_N | GLU_A_34_H | -4.195 | -2.303 | -8.707 | -4.599 | -2.308 | -7.781 | 1.01 | 23.57 | -179.29 |
| 199 | 2jsa-10 | GLU_A_34_N | GLU_A_34_CA | -4.195 | -2.303 | -8.707 | -3.714 | -1.046 | -9.254 | 1.45 | 112.12 | 69.06 |
| 200 | 2jsa-10 | GLU_A_34_CA | GLU_A_34_HA | -3.714 | -1.046 | -9.254 | -3.9 | -1.052 | -10.328 | 1.09 | 170.17 | -178.15 |
| 201 | 2jsa-10 | GLU_A_34_CA | GLU_A_34_C | -3.714 | -1.046 | -9.254 | -2.207 | -0.91 | -9.029 | 1.53 | 81.54 | 5.16 |
| 202 | 2jsa-10 | GLU_A_34_C | GLU_A_34_O | -2.207 | -0.91 | -9.029 | -1.681 | 0.2 | -8.978 | 1.23 | 87.62 | 64.64 |
| 203 | 2jsa-10 | GLU_A_34_C | CYS_A_35_N | -2.207 | -0.91 | -9.029 | -1.554 | -2.057 | -8.901 | 1.33 | 84.46 | -60.35 |
| 204 | 2jsa-10 | CYS_A_35_N | CYS_A_35_H | -1.554 | -2.057 | -8.901 | -2.051 | -2.934 | -8.954 | 1.01 | 93.01 | -119.54 |
| 205 | 2jsa-10 | CYS_A_35_N | CYS_A_35_CA | -1.554 | -2.057 | -8.901 | -0.118 | -2.08 | -8.684 | 1.45 | 81.41 | -0.92 |
| 206 | 2jsa-10 | CYS_A_35_CA | CYS_A_35_HA | -0.118 | -2.08 | -8.684 | 0.258 | -1.06 | -8.764 | 1.09 | 94.21 | 69.76 |
| 207 | 2jsa-10 | CYS_A_35_CA | CYS_A_35_C | -0.118 | -2.08 | -8.684 | 0.522 | -2.922 | -9.788 | 1.53 | 136.23 | -52.76 |
| 208 | 2jsa-10 | CYS_A_35_C | CYS_A_35_O | 0.522 | -2.922 | -9.788 | 1.506 | -2.507 | -10.399 | 1.23 | 119.78 | 22.87 |
| 209 | 2jsa-10 | CYS_A_35_C | LYS_A_36_N | 0.522 | -2.922 | -9.788 | -0.061 | -4.091 | -10.011 | 1.33 | 99.69 | -116.51 |
| 210 | 2jsa-10 | LYS_A_36_N | LYS_A_36_H | -0.061 | -4.091 | -10.011 | -0.867 | -4.368 | -9.469 | 1.01 | 57.55 | -161.03 |
| 211 | 2jsa-10 | LYS_A_36_N | LYS_A_36_CA | -0.061 | -4.091 | -10.011 | 0.44 | -4.995 | -11.031 | 1.45 | 134.62 | -61.00 |
| 212 | 2jsa-10 | LYS_A_36_CA | LYS_A_36_HA | 0.44 | -4.995 | -11.031 | 1.504 | -5.152 | -10.85 | 1.09 | 80.45 | -8.39 |
| 213 | 2jsa-10 | LYS_A_36_CA | LYS_A_36_C | 0.44 | -4.995 | -11.031 | 0.294 | -4.338 | -12.405 | 1.53 | 153.90 | 102.53 |
| 214 | 2jsa-10 | LYS_A_36_C | LYS_A_36_O | 0.294 | -4.338 | -12.405 | 1.165 | -4.482 | -13.262 | 1.23 | 134.15 | -9.39 |
| 215 | 2jsa-10 | LYS_A_36_C | LYS_A_37_N | 0.294 | -4.338 | -12.405 | -0.815 | -3.631 | -12.572 | 1.33 | 97.24 | 147.48 |
| 216 | 2jsa-10 | LYS_A_37_N | LYS_A_37_H | -0.815 | -3.631 | -12.572 | -1.488 | -3.556 | -11.823 | 1.01 | 42.12 | 173.64 |
| 217 | 2jsa-10 | LYS_A_37_N | LYS_A_37_CA | -0.815 | -3.631 | -12.572 | -1.085 | -2.952 | -13.827 | 1.45 | 149.79 | 111.68 |
| 218 | 2jsa-10 | LYS_A_37_CA | LYS_A_37_HA | -1.085 | -2.952 | -13.827 | -0.649 | -3.544 | -14.631 | 1.09 | 137.56 | -53.63 |

| | | | | | | | | | | | |
|---|---|---|---|---|---|---|---|---|---|---|---|
| 219 | 2jsa-10 | LYS_A_37_CA | LYS_A_37_C | -1.085 | -2.952 | -13.827 | -0.391 | -1.588 | -13.826 | 1.53 | 89.96 | 63.03 |
| 220 | 2jsa-10 | LYS_A_37_C | LYS_A_37_O | -0.391 | -1.588 | -13.826 | 0.104 | -1.139 | -14.858 | 1.23 | 147.07 | 42.21 |
| 221 | 2jsa-10 | LYS_A_37_C | LEU_A_38_N | -0.391 | -1.588 | -13.826 | -0.376 | -0.97 | -12.655 | 1.32 | 27.83 | 88.61 |
| 222 | 2jsa-10 | LEU_A_38_N | LEU_A_38_H | -0.376 | -0.97 | -12.655 | -0.803 | -1.4 | -11.847 | 1.01 | 36.87 | -134.80 |
| 223 | 2jsa-10 | LEU_A_38_N | LEU_A_38_CA | -0.376 | -0.97 | -12.655 | 0.251 | 0.333 | -12.505 | 1.45 | 84.08 | 64.30 |
| 224 | 2jsa-10 | LEU_A_38_CA | LEU_A_38_HA | 0.251 | 0.333 | -12.505 | -0.142 | 0.988 | -13.282 | 1.09 | 135.49 | 120.96 |
| 225 | 2jsa-10 | LEU_A_38_CA | LEU_A_38_C | 0.251 | 0.333 | -12.505 | 1.758 | 0.199 | -12.73 | 1.53 | 98.46 | -5.08 |
| 226 | 2jsa-10 | LEU_A_38_C | LEU_A_38_O | 1.758 | 0.199 | -12.73 | 2.294 | 0.731 | -13.7 | 1.23 | 142.10 | 44.79 |
| 227 | 2jsa-10 | LEU_A_38_C | PHE_A_39_N | 1.758 | 0.199 | -12.73 | 2.399 | -0.516 | -11.816 | 1.33 | 46.41 | -48.12 |
| 228 | 2jsa-10 | PHE_A_39_N | PHE_A_39_H | 2.399 | -0.516 | -11.816 | 1.894 | -0.925 | -11.043 | 1.01 | 40.05 | -141.00 |
| 229 | 2jsa-10 | PHE_A_39_N | PHE_A_39_CA | 2.399 | -0.516 | -11.816 | 3.834 | -0.727 | -11.903 | 1.45 | 93.43 | -8.36 |
| 230 | 2jsa-10 | PHE_A_39_CA | PHE_A_39_HA | 3.834 | -0.727 | -11.903 | 4.31 | 0.144 | -12.355 | 1.09 | 114.48 | 61.34 |
| 231 | 2jsa-10 | PHE_A_39_CA | PHE_A_39_C | 3.834 | -0.727 | -11.903 | 4.156 | -1.975 | -12.727 | 1.53 | 122.59 | -75.53 |
| 232 | 2jsa-10 | PHE_A_39_C | PHE_A_39_O | 4.156 | -1.975 | -12.727 | 5.192 | -2.605 | -12.526 | 1.23 | 80.59 | -31.30 |
| 233 | 2jsa-10 | PHE_A_39_C | HIS_A_40_N | 4.156 | -1.975 | -12.727 | 3.248 | -2.292 | -13.639 | 1.33 | 133.48 | -160.75 |
| 234 | 2jsa-10 | HIS_A_40_N | HIS_A_40_H | 3.248 | -2.292 | -13.639 | 2.419 | -1.726 | -13.749 | 1.01 | 96.25 | 145.68 |
| 235 | 2jsa-10 | HIS_A_40_N | HIS_A_40_CA | 3.248 | -2.292 | -13.639 | 3.422 | -3.453 | -14.495 | 1.45 | 126.10 | -81.48 |
| 236 | 2jsa-10 | HIS_A_40_CA | HIS_A_40_HA | 3.422 | -3.453 | -14.495 | 3.451 | -4.338 | -13.859 | 1.09 | 54.31 | -88.12 |
| 237 | 2jsa-10 | HIS_A_40_CA | HIS_A_40_C | 3.422 | -3.453 | -14.495 | 4.763 | -3.356 | -15.226 | 1.53 | 118.53 | 4.14 |
| 238 | 2jsa-10 | HIS_A_40_C | HIS_A_40_O | 4.763 | -3.356 | -15.226 | 5.557 | -4.294 | -15.204 | 1.23 | 88.97 | -49.75 |
| 239 | 2jsa-10 | HIS_A_40_C | THR_A_41_N | 4.763 | -3.356 | -15.226 | 4.974 | -2.21 | -15.857 | 1.33 | 118.44 | 79.57 |
| 240 | 2jsa-10 | THR_A_41_N | THR_A_41_H | 4.974 | -2.21 | -15.857 | 4.277 | -1.48 | -15.832 | 1.01 | 88.58 | 133.68 |
| 241 | 2jsa-10 | THR_A_41_N | THR_A_41_CA | 4.974 | -2.21 | -15.857 | 6.205 | -1.977 | -16.594 | 1.45 | 120.47 | 10.72 |
| 242 | 2jsa-10 | THR_A_41_CA | THR_A_41_HA | 6.205 | -1.977 | -16.594 | 6.264 | -2.694 | -17.413 | 1.09 | 138.70 | -85.30 |
| 243 | 2jsa-10 | THR_A_41_CA | THR_A_41_C | 6.205 | -1.977 | -16.594 | 7.418 | -2.196 | -15.688 | 1.53 | 53.68 | -10.23 |
| 244 | 2jsa-10 | THR_A_41_C | THR_A_41_O | 7.418 | -2.196 | -15.688 | 8.374 | -2.866 | -16.076 | 1.23 | 108.38 | -35.02 |
| 245 | 2jsa-10 | THR_A_41_C | ILE_A_42_N | 7.418 | -2.196 | -15.688 | 7.34 | -1.617 | -14.498 | 1.33 | 26.15 | 97.67 |
| 246 | 2jsa-10 | ILE_A_42_N | ILE_A_42_H | 7.34 | -1.617 | -14.498 | 6.523 | -1.08 | -14.246 | 1.01 | 75.55 | 146.68 |
| 247 | 2jsa-10 | ILE_A_42_N | ILE_A_42_CA | 7.34 | -1.617 | -14.498 | 8.42 | -1.741 | -13.534 | 1.45 | 48.43 | -6.55 |
| 248 | 2jsa-10 | ILE_A_42_CA | ILE_A_42_HA | 8.42 | -1.741 | -13.534 | 9.25 | -1.121 | -13.875 | 1.09 | 108.22 | 36.76 |
| 249 | 2jsa-10 | ILE_A_42_CA | ILE_A_42_C | 8.42 | -1.741 | -13.534 | 8.906 | -3.192 | -13.502 | 1.53 | 88.80 | -71.48 |
| 250 | 2jsa-10 | ILE_A_42_C | ILE_A_42_O | 8.906 | -3.192 | -13.502 | 8.108 | -4.12 | -13.62 | 1.23 | 95.51 | -130.69 |
| 251 | 2jsa-10 | ILE_A_42_C | PRO_A_43_N | 8.906 | -3.192 | -13.502 | 10.247 | -3.344 | -13.339 | 1.36 | 83.11 | -6.47 |
| 252 | 2jsa-10 | PRO_A_43_N | PRO_A_43_CA | 10.247 | -3.344 | -13.339 | 10.85 | -4.666 | -13.291 | 1.45 | 88.11 | -65.48 |
| 253 | 2jsa-10 | PRO_A_43_CA | PRO_A_43_HA | 10.85 | -4.666 | -13.291 | 10.442 | -5.291 | -14.085 | 1.09 | 136.77 | -123.14 |
| 254 | 2jsa-10 | PRO_A_43_CA | PRO_A_43_C | 10.85 | -4.666 | -13.291 | 10.574 | -5.345 | -11.948 | 1.53 | 28.62 | -112.12 |
| 255 | 2jsa-10 | PRO_A_43_C | PRO_A_43_O | 10.574 | -5.345 | -11.948 | 10.073 | -6.468 | -11.907 | 1.23 | 88.09 | -114.04 |
| 256 | 2jsa-10 | PRO_A_43_C | PHE_A_44_N | 10.574 | -5.345 | -11.948 | 10.913 | -4.636 | -10.882 | 1.32 | 36.40 | 64.45 |
| 257 | 2jsa-10 | PHE_A_44_N | PHE_A_44_H | 10.913 | -4.636 | -10.882 | 11.321 | -3.718 | -10.99 | 1.01 | 96.14 | 66.04 |
| 258 | 2jsa-10 | PHE_A_44_N | PHE_A_44_CA | 10.913 | -4.636 | -10.882 | 10.708 | -5.157 | -9.54 | 1.45 | 22.65 | -111.48 |
| 259 | 2jsa-10 | PHE_A_44_CA | PHE_A_44_HA | 10.708 | -5.157 | -9.54 | 10.515 | -6.229 | -9.584 | 1.09 | 92.31 | -100.21 |
| 260 | 2jsa-10 | PHE_A_44_CA | PHE_A_44_C | 10.708 | -5.157 | -9.54 | 9.539 | -4.451 | -8.85 | 1.53 | 63.19 | 148.87 |
| 261 | 2jsa-10 | PHE_A_44_C | PHE_A_44_O | 9.539 | -4.451 | -8.85 | 9.552 | -4.266 | -7.635 | 1.23 | 8.68 | 85.98 |
| 262 | 2jsa-10 | PHE_A_44_C | GLY_A_45_N | 9.539 | -4.451 | -8.85 | 8.558 | -4.074 | -9.657 | 1.33 | 127.52 | 158.98 |
| 263 | 2jsa-10 | GLY_A_45_N | GLY_A_45_H | 8.558 | -4.074 | -9.657 | 8.616 | -4.258 | -10.648 | 1.01 | 168.98 | -72.50 |
| 264 | 2jsa-10 | GLY_A_45_N | GLY_A_45_CA | 8.558 | -4.074 | -9.657 | 7.384 | -3.392 | -9.14 | 1.45 | 69.15 | 149.85 |
| 265 | 2jsa-10 | GLY_A_45_CA | GLY_A_45_C | 7.384 | -3.392 | -9.14 | 6.341 | -4.396 | -8.645 | 1.53 | 71.12 | -136.09 |
| 266 | 2jsa-10 | GLY_A_45_C | GLY_A_45_O | 6.341 | -4.396 | -8.645 | 5.909 | -4.33 | -7.495 | 1.23 | 20.81 | 171.31 |
| 267 | 2jsa-10 | GLY_A_45_C | THR_A_46_N | 6.341 | -4.396 | -8.645 | 5.967 | -5.302 | -9.536 | 1.32 | 132.27 | -112.43 |
| 268 | 2jsa-10 | THR_A_46_N | THR_A_46_H | 5.967 | -5.302 | -9.536 | 6.366 | -5.297 | -10.464 | 1.01 | 156.73 | 0.72 |
| 269 | 2jsa-10 | THR_A_46_N | THR_A_46_CA | 5.967 | -5.302 | -9.536 | 4.983 | -6.317 | -9.204 | 1.45 | 76.78 | -134.11 |
| 270 | 2jsa-10 | THR_A_46_CA | THR_A_46_HA | 4.983 | -6.317 | -9.204 | 4.06 | -5.824 | -8.9 | 1.09 | 73.80 | 151.89 |
| 271 | 2jsa-10 | THR_A_46_CA | THR_A_46_C | 4.983 | -6.317 | -9.204 | 5.471 | -7.169 | -8.03 | 1.53 | 39.91 | -60.20 |
| 272 | 2jsa-10 | THR_A_46_C | THR_A_46_O | 5.471 | -7.169 | -8.03 | 4.698 | -7.493 | -7.131 | 1.23 | 42.99 | -157.26 |
| 273 | 2jsa-10 | THR_A_46_C | ARG_A_47_N | 5.471 | -7.169 | -8.03 | 6.751 | -7.507 | -8.078 | 1.32 | 92.08 | -14.79 |
| 274 | 2jsa-10 | ARG_A_47_N | ARG_A_47_H | 6.751 | -7.507 | -8.078 | 7.329 | -7.205 | -8.85 | 1.01 | 139.81 | 27.59 |
| 275 | 2jsa-10 | ARG_A_47_N | ARG_A_47_CA | 6.751 | -7.507 | -8.078 | 7.351 | -8.315 | -7.031 | 1.45 | 43.87 | -53.40 |
| 276 | 2jsa-10 | ARG_A_47_CA | ARG_A_47_HA | 7.351 | -8.315 | -7.031 | 6.815 | -9.263 | -6.978 | 1.09 | 87.21 | -119.48 |
| 277 | 2jsa-10 | ARG_A_47_CA | ARG_A_47_C | 7.351 | -8.315 | -7.031 | 7.239 | -7.603 | -5.681 | 1.53 | 28.10 | 98.94 |
| 278 | 2jsa-10 | ARG_A_47_C | ARG_A_47_O | 7.239 | -7.603 | -5.681 | 6.923 | -8.227 | -4.67 | 1.23 | 34.68 | -116.86 |
| 279 | 2jsa-10 | ARG_A_47_C | GLU_A_48_N | 7.239 | -7.603 | -5.681 | 7.502 | -6.305 | -5.709 | 1.32 | 91.21 | 78.55 |
| 280 | 2jsa-10 | GLU_A_48_N | GLU_A_48_H | 7.502 | -6.305 | -5.709 | 7.754 | -5.856 | -6.578 | 1.01 | 149.35 | 60.70 |
| 281 | 2jsa-10 | GLU_A_48_N | GLU_A_48_CA | 7.502 | -6.305 | -5.709 | 7.435 | -5.501 | -4.501 | 1.45 | 33.74 | 94.76 |
| 282 | 2jsa-10 | GLU_A_48_CA | GLU_A_48_HA | 7.435 | -5.501 | -4.501 | 7.937 | -6.048 | -3.703 | 1.09 | 42.93 | -47.46 |
| 283 | 2jsa-10 | GLU_A_48_CA | GLU_A_48_C | 7.435 | -5.501 | -4.501 | 5.977 | -5.281 | -4.091 | 1.53 | 74.46 | 171.42 |
| 284 | 2jsa-10 | GLU_A_48_C | GLU_A_48_O | 5.977 | -5.281 | -4.091 | 5.68 | -5.114 | -2.91 | 1.23 | 16.09 | 150.65 |
| 285 | 2jsa-10 | GLU_A_48_C | CYS_A_49_N | 5.977 | -5.281 | -4.091 | 5.107 | -5.286 | -5.09 | 1.32 | 138.95 | -179.67 |
| 286 | 2jsa-10 | CYS_A_49_N | CYS_A_49_H | 5.107 | -5.286 | -5.09 | 5.424 | -5.428 | -6.038 | 1.01 | 159.88 | -24.13 |
| 287 | 2jsa-10 | CYS_A_49_N | CYS_A_49_CA | 5.107 | -5.286 | -5.09 | 3.689 | -5.09 | -4.849 | 1.45 | 80.44 | 172.13 |
| 288 | 2jsa-10 | CYS_A_49_CA | CYS_A_49_HA | 3.689 | -5.09 | -4.849 | 3.549 | -4.097 | -4.422 | 1.09 | 66.94 | 98.03 |
| 289 | 2jsa-10 | CYS_A_49_CA | CYS_A_49_C | 3.689 | -5.09 | -4.849 | 3.227 | -6.129 | -3.825 | 1.53 | 48.00 | -113.97 |
| 290 | 2jsa-10 | CYS_A_49_C | CYS_A_49_O | 3.227 | -6.129 | -3.825 | 2.85 | -5.78 | -2.707 | 1.23 | 24.68 | 137.21 |
| 291 | 2jsa-10 | CYS_A_49_C | ASP_A_50_N | 3.227 | -6.129 | -3.825 | 3.273 | -7.386 | -4.243 | 1.33 | 108.38 | -87.90 |
| 292 | 2jsa-10 | ASP_A_50_N | ASP_A_50_H | 3.273 | -7.386 | -4.243 | 3.596 | -7.601 | -5.176 | 1.01 | 157.42 | -33.65 |
| 293 | 2jsa-10 | ASP_A_50_N | ASP_A_50_CA | 3.273 | -7.386 | -4.243 | 2.863 | -8.477 | -3.377 | 1.45 | 53.39 | -110.60 |
| 294 | 2jsa-10 | ASP_A_50_CA | ASP_A_50_HA | 2.863 | -8.477 | -3.377 | 1.805 | -8.365 | -3.139 | 1.09 | 77.39 | 173.96 |
| 295 | 2jsa-10 | ASP_A_50_CA | ASP_A_50_C | 2.863 | -8.477 | -3.377 | 3.692 | -8.441 | -2.091 | 1.53 | 32.83 | 2.49 |
| 296 | 2jsa-10 | ASP_A_50_C | ASP_A_50_O | 3.692 | -8.441 | -2.091 | 3.242 | -8.907 | -1.045 | 1.23 | 31.77 | -134.00 |
| 297 | 2jsa-10 | ASP_A_50_C | HIS_A_51_N | 3.692 | -8.441 | -2.091 | 4.887 | -7.881 | -2.211 | 1.33 | 95.20 | 25.11 |
| 298 | 2jsa-10 | HIS_A_51_N | HIS_A_51_H | 4.887 | -7.881 | -2.211 | 5.192 | -7.517 | -3.103 | 1.01 | 151.97 | 50.04 |
| 299 | 2jsa-10 | HIS_A_51_N | HIS_A_51_CA | 4.887 | -7.881 | -2.211 | 5.782 | -7.778 | -1.071 | 1.45 | 38.32 | 6.56 |
| 300 | 2jsa-10 | HIS_A_51_CA | HIS_A_51_HA | 5.782 | -7.778 | -1.071 | 5.647 | -8.667 | -0.454 | 1.09 | 55.54 | -98.63 |
| 301 | 2jsa-10 | HIS_A_51_CA | HIS_A_51_C | 5.782 | -7.778 | -1.071 | 5.402 | -6.559 | -0.229 | 1.53 | 56.60 | 107.31 |
| 302 | 2jsa-10 | HIS_A_51_C | HIS_A_51_O | 5.402 | -6.559 | -0.229 | 6.036 | -6.28 | 0.788 | 1.23 | 34.26 | 23.75 |
| 303 | 2jsa-10 | HIS_A_51_C | TYR_A_52_N | 5.402 | -6.559 | -0.229 | 4.367 | -5.865 | -0.682 | 1.33 | 109.98 | 146.16 |
| 304 | 2jsa-10 | TYR_A_52_N | TYR_A_52_H | 4.367 | -5.865 | -0.682 | 3.893 | -6.153 | -1.526 | 1.01 | 146.69 | -148.72 |
| 305 | 2jsa-10 | TYR_A_52_N | TYR_A_52_CA | 4.367 | -5.865 | -0.682 | 3.895 | -4.683 | 0.016 | 1.45 | 61.26 | 111.77 |
| 306 | 2jsa-10 | TYR_A_52_CA | TYR_A_52_HA | 3.895 | -4.683 | 0.016 | 4.599 | -4.402 | 0.8 | 1.09 | 44.03 | 21.76 |
| 307 | 2jsa-10 | TYR_A_52_CA | TYR_A_52_C | 3.895 | -4.683 | 0.016 | 2.509 | -4.922 | 0.622 | 1.53 | 66.69 | -170.22 |
| 308 | 2jsa-10 | TYR_A_52_C | TYR_A_52_O | 2.509 | -4.922 | 0.622 | 2.285 | -4.638 | 1.798 | 1.23 | 17.10 | 128.26 |
| 309 | 2jsa-10 | TYR_A_52_C | VAL_A_53_N | 2.509 | -4.922 | 0.622 | 1.617 | -5.439 | -0.208 | 1.32 | 128.84 | -149.90 |
| 310 | 2jsa-10 | VAL_A_53_N | VAL_A_53_H | 1.617 | -5.439 | -0.208 | 1.872 | -5.646 | -1.163 | 1.01 | 161.02 | -39.07 |
| 311 | 2jsa-10 | VAL_A_53_N | VAL_A_53_CA | 1.617 | -5.439 | -0.208 | 0.26 | -5.719 | 0.23 | 1.45 | 72.46 | -168.34 |
| 312 | 2jsa-10 | VAL_A_53_CA | VAL_A_53_HA | 0.26 | -5.719 | 0.23 | -0.222 | -4.774 | 0.482 | 1.09 | 76.64 | 117.02 |
| 313 | 2jsa-10 | VAL_A_53_CA | VAL_A_53_C | 0.26 | -5.719 | 0.23 | 0.305 | -6.581 | 1.493 | 1.53 | 34.35 | -87.01 |
| 314 | 2jsa-10 | VAL_A_53_C | VAL_A_53_O | 0.305 | -6.581 | 1.493 | -0.662 | -6.626 | 2.252 | 1.23 | 51.90 | -177.34 |

| | | | | | | | | | | |
|---|---|---|---|---|---|---|---|---|---|---|
| 315 | 2jsa-10 | VAL_A_53_C | ASN_A_54_N | 0.305 | -6.581 | 1.493 | 1.436 | -7.245 | 1.679 | 1.32 | 81.93 | -30.42 |
| 316 | 2jsa-10 | ASN_A_54_N | ASN_A_54_H | 1.436 | -7.245 | 1.679 | 2.19 | -7.161 | 1.012 | 1.01 | 131.32 | 6.36 |
| 317 | 2jsa-10 | ASN_A_54_N | ASN_A_54_CA | 1.436 | -7.245 | 1.679 | 1.62 | -8.103 | 2.837 | 1.45 | 37.15 | -77.90 |
| 318 | 2jsa-10 | ASN_A_54_CA | ASN_A_54_HA | 1.62 | -8.103 | 2.837 | 0.875 | -7.849 | 3.591 | 1.09 | 46.23 | 161.17 |
| 319 | 2jsa-10 | ASN_A_54_CA | ASN_A_54_C | 1.62 | -8.103 | 2.837 | 3.025 | -7.892 | 3.405 | 1.53 | 68.21 | 8.54 |
| 320 | 2jsa-10 | ASN_A_54_C | ASN_A_54_O | 3.025 | -7.892 | 3.405 | 3.881 | -8.769 | 3.292 | 1.23 | 95.27 | -45.69 |
| 321 | 2jsa-10 | ASN_A_54_C | SER_A_55_N | 3.025 | -7.892 | 3.405 | 3.219 | -6.727 | 4.002 | 1.32 | 63.18 | 80.55 |
| 322 | 2jsa-10 | SER_A_55_N | SER_A_55_H | 3.219 | -6.727 | 4.002 | 2.469 | -6.053 | 4.058 | 1.01 | 86.82 | 138.06 |
| 323 | 2jsa-10 | SER_A_55_N | SER_A_55_CA | 3.219 | -6.727 | 4.002 | 4.506 | -6.391 | 4.589 | 1.45 | 66.19 | 14.63 |
| 324 | 2jsa-10 | SER_A_55_CA | SER_A_55_HA | 4.506 | -6.391 | 4.589 | 4.647 | -7.003 | 5.48 | 1.09 | 35.18 | -77.03 |
| 325 | 2jsa-10 | SER_A_55_CA | SER_A_55_C | 4.506 | -6.391 | 4.589 | 4.525 | -4.917 | 4.998 | 1.53 | 74.49 | 89.26 |
| 326 | 2jsa-10 | SER_A_55_C | SER_A_55_O | 4.525 | -4.917 | 4.998 | 5.163 | -4.552 | 5.985 | 1.23 | 36.68 | 29.77 |
| 327 | 2jsa-10 | SER_A_55_C | LYS_A_56_N | 4.525 | -4.917 | 4.998 | 3.82 | -4.11 | 4.22 | 1.32 | 125.98 | 131.14 |
| 328 | 2jsa-10 | LYS_A_56_N | LYS_A_56_H | 3.82 | -4.11 | 4.22 | 3.321 | -4.48 | 3.424 | 1.01 | 142.03 | -143.44 |
| 329 | 2jsa-10 | LYS_A_56_N | LYS_A_56_CA | 3.82 | -4.11 | 4.22 | 3.748 | -2.684 | 4.49 | 1.45 | 79.29 | 92.89 |
| 330 | 2jsa-10 | LYS_A_56_CA | LYS_A_56_HA | 3.748 | -2.684 | 4.49 | 4.175 | -2.505 | 5.477 | 1.09 | 25.13 | 22.74 |
| 331 | 2jsa-10 | LYS_A_56_CA | LYS_A_56_C | 3.748 | -2.684 | 4.49 | 2.282 | -2.25 | 4.531 | 1.53 | 88.46 | 163.51 |
| 332 | 2jsa-10 | LYS_A_56_C | LYS_A_56_O | 2.282 | -2.25 | 4.531 | 1.985 | -1.066 | 4.68 | 1.23 | 83.04 | 104.08 |
| 333 | 2jsa-10 | LYS_A_56_C | VAL_A_57_N | 2.282 | -2.25 | 4.531 | 1.403 | -3.233 | 4.398 | 1.33 | 95.76 | -131.80 |
| 334 | 2jsa-10 | VAL_A_57_N | VAL_A_57_H | 1.403 | -3.233 | 4.398 | 1.719 | -4.185 | 4.28 | 1.01 | 96.71 | -71.64 |
| 335 | 2jsa-10 | VAL_A_57_N | VAL_A_57_CA | 1.403 | -3.233 | 4.398 | -0.026 | -2.967 | 4.418 | 1.45 | 89.21 | 169.46 |
| 336 | 2jsa-10 | VAL_A_57_CA | VAL_A_57_HA | -0.026 | -2.967 | 4.418 | -0.183 | -1.914 | 4.185 | 1.09 | 102.34 | 98.48 |
| 337 | 2jsa-10 | VAL_A_57_CA | VAL_A_57_C | -0.026 | -2.967 | 4.418 | -0.569 | -3.229 | 5.824 | 1.53 | 23.21 | -154.24 |
| 338 | 2jsa-10 | VAL_A_57_C | VAL_A_57_O | -0.569 | -3.229 | 5.824 | -1.36 | -2.443 | 6.344 | 1.23 | 65.00 | 135.18 |
| 339 | 2jsa-10 | VAL_A_57_C | ASP_A_58_N | -0.569 | -3.229 | 5.824 | -0.121 | -4.335 | 6.401 | 1.33 | 64.19 | -67.95 |
| 340 | 2jsa-10 | ASP_A_58_N | ASP_A_58_H | -0.121 | -4.335 | 6.401 | 0.53 | -4.936 | 5.915 | 1.01 | 118.75 | -42.71 |
| 341 | 2jsa-10 | ASP_A_58_N | ASP_A_58_CA | -0.121 | -4.335 | 6.401 | -0.552 | -4.708 | 7.737 | 1.45 | 23.11 | -139.13 |
| 342 | 2jsa-10 | ASP_A_58_CA | ASP_A_58_HA | -0.552 | -4.708 | 7.737 | -1.618 | -4.931 | 7.718 | 1.09 | 91.00 | -168.18 |
| 343 | 2jsa-10 | ASP_A_58_CA | ASP_A_58_C | -0.552 | -4.708 | 7.737 | -0.281 | -3.55 | 8.698 | 1.53 | 51.06 | 76.83 |
| 344 | 2jsa-10 | ASP_A_58_C | ASP_A_58_O | -0.281 | -3.55 | 8.698 | -1.129 | -3.213 | 9.524 | 1.23 | 47.85 | 158.33 |
| 345 | 2jsa-10 | ASP_A_58_C | PRO_A_59_N | -0.281 | -3.55 | 8.698 | 0.934 | -2.957 | 8.557 | 1.36 | 95.95 | 26.02 |
| 346 | 2jsa-10 | PRO_A_59_N | PRO_A_59_CA | 0.934 | -2.957 | 8.557 | 1.327 | -1.843 | 9.404 | 1.45 | 54.36 | 70.57 |
| 347 | 2jsa-10 | PRO_A_59_CA | PRO_A_59_HA | 1.327 | -1.843 | 9.404 | 1.073 | -2.056 | 10.443 | 1.09 | 17.70 | -140.02 |
| 348 | 2jsa-10 | PRO_A_59_CA | PRO_A_59_C | 1.327 | -1.843 | 9.404 | 0.613 | -0.557 | 8.984 | 1.53 | 105.94 | 119.04 |
| 349 | 2jsa-10 | PRO_A_59_C | PRO_A_59_O | 0.613 | -0.557 | 8.984 | -0.083 | 0.061 | 9.788 | 1.23 | 49.18 | 138.40 |
| 350 | 2jsa-10 | PRO_A_59_C | ILE_A_60_N | 0.613 | -0.557 | 8.984 | 0.81 | -0.193 | 7.725 | 1.33 | 161.80 | 61.58 |
| 351 | 2jsa-10 | ILE_A_60_N | ILE_A_60_H | 0.81 | -0.193 | 7.725 | 1.395 | -0.751 | 7.119 | 1.01 | 126.85 | -43.65 |
| 352 | 2jsa-10 | ILE_A_60_N | ILE_A_60_CA | 0.81 | -0.193 | 7.725 | 0.192 | 1.009 | 7.19 | 1.45 | 111.60 | 117.21 |
| 353 | 2jsa-10 | ILE_A_60_CA | ILE_A_60_HA | 0.192 | 1.009 | 7.19 | 0.688 | 1.871 | 7.637 | 1.09 | 65.80 | 60.08 |
| 354 | 2jsa-10 | ILE_A_60_CA | ILE_A_60_C | 0.192 | 1.009 | 7.19 | -1.279 | 1.051 | 7.607 | 1.53 | 74.18 | 178.36 |
| 355 | 2jsa-10 | ILE_A_60_C | ILE_A_60_O | -1.279 | 1.051 | 7.607 | -1.735 | 2.034 | 8.188 | 1.23 | 61.80 | 114.89 |
| 356 | 2jsa-10 | ILE_A_60_C | ILE_A_61_N | -1.279 | 1.051 | 7.607 | -1.981 | -0.029 | 7.294 | 1.33 | 103.66 | -123.02 |
| 357 | 2jsa-10 | ILE_A_61_N | ILE_A_61_H | -1.981 | -0.029 | 7.294 | -1.539 | -0.802 | 6.817 | 1.01 | 118.18 | -60.24 |
| 358 | 2jsa-10 | ILE_A_61_N | ILE_A_61_CA | -1.981 | -0.029 | 7.294 | -3.392 | -0.127 | 7.628 | 1.45 | 76.71 | -176.03 |
| 359 | 2jsa-10 | ILE_A_61_CA | ILE_A_61_HA | -3.392 | -0.127 | 7.628 | -3.901 | 0.736 | 7.199 | 1.09 | 113.18 | 120.53 |
| 360 | 2jsa-10 | ILE_A_61_CA | ILE_A_61_C | -3.392 | -0.127 | 7.628 | -3.556 | -0.056 | 9.148 | 1.53 | 6.71 | 156.59 |
| 361 | 2jsa-10 | ILE_A_61_C | ILE_A_61_O | -3.556 | -0.056 | 9.148 | -4.62 | 0.313 | 9.644 | 1.23 | 66.23 | 160.87 |
| 362 | 2jsa-10 | ILE_A_61_C | HIS_A_62_N | -3.556 | -0.056 | 9.148 | -2.489 | -0.416 | 9.845 | 1.32 | 58.24 | -18.64 |
| 363 | 2jsa-10 | HIS_A_62_N | HIS_A_62_H | -2.489 | -0.416 | 9.845 | -1.647 | -0.71 | 9.37 | 1.01 | 118.04 | -19.25 |
| 364 | 2jsa-10 | HIS_A_62_N | HIS_A_62_CA | -2.489 | -0.416 | 9.845 | -2.501 | -0.398 | 11.298 | 1.45 | 0.85 | 123.69 |
| 365 | 2jsa-10 | HIS_A_62_CA | HIS_A_62_HA | -2.501 | -0.398 | 11.298 | -3.362 | -0.974 | 11.635 | 1.09 | 71.98 | -146.22 |
| 366 | 2jsa-10 | HIS_A_62_CA | HIS_A_62_C | -2.501 | -0.398 | 11.298 | -2.676 | 1.041 | 11.791 | 1.53 | 71.22 | 96.93 |
| 367 | 2jsa-10 | HIS_A_62_C | HIS_A_62_O | -2.676 | 1.041 | 11.791 | -3.49 | 1.303 | 12.674 | 1.23 | 44.08 | 162.16 |
| 368 | 2jsa-10 | HIS_A_62_C | GLU_A_63_N | -2.676 | 1.041 | 11.791 | -1.896 | 1.933 | 11.199 | 1.32 | 116.55 | 48.83 |
| 369 | 2jsa-10 | GLU_A_63_N | GLU_A_63_H | -1.896 | 1.933 | 11.199 | -1.248 | 1.645 | 10.479 | 1.01 | 135.44 | -23.96 |
| 370 | 2jsa-10 | GLU_A_63_N | GLU_A_63_CA | -1.896 | 1.933 | 11.199 | -1.954 | 3.338 | 11.568 | 1.45 | 75.30 | 92.36 |
| 371 | 2jsa-10 | GLU_A_63_CA | GLU_A_63_HA | -1.954 | 3.338 | 11.568 | -1.744 | 3.416 | 12.635 | 1.09 | 11.86 | 20.38 |
| 372 | 2jsa-10 | GLU_A_63_CA | GLU_A_63_C | -1.954 | 3.338 | 11.568 | -3.355 | 3.897 | 11.311 | 1.53 | 99.67 | 158.25 |
| 373 | 2jsa-10 | GLU_A_63_C | GLU_A_63_O | -3.355 | 3.897 | 11.311 | -3.963 | 4.49 | 12.2 | 1.23 | 43.69 | 135.72 |
| 374 | 2jsa-10 | GLU_A_63_C | LEU_A_64_N | -3.355 | 3.897 | 11.311 | -3.826 | 3.687 | 10.09 | 1.33 | 157.10 | -155.97 |
| 375 | 2jsa-10 | LEU_A_64_N | LEU_A_64_H | -3.826 | 3.687 | 10.09 | -3.27 | 3.19 | 9.408 | 1.01 | 132.44 | -41.79 |
| 376 | 2jsa-10 | LEU_A_64_N | LEU_A_64_CA | -3.826 | 3.687 | 10.09 | -5.144 | 4.164 | 9.706 | 1.45 | 105.32 | 160.10 |
| 377 | 2jsa-10 | LEU_A_64_CA | LEU_A_64_HA | -5.144 | 4.164 | 9.706 | -5.16 | 5.247 | 9.829 | 1.09 | 83.52 | 90.85 |
| 378 | 2jsa-10 | LEU_A_64_CA | LEU_A_64_C | -5.144 | 4.164 | 9.706 | -6.191 | 3.572 | 10.651 | 1.53 | 51.84 | -150.52 |
| 379 | 2jsa-10 | LEU_A_64_C | LEU_A_64_O | -6.191 | 3.572 | 10.651 | -7.174 | 4.231 | 10.986 | 1.23 | 74.19 | 146.16 |
| 380 | 2jsa-10 | LEU_A_64_C | GLU_A_65_N | -6.191 | 3.572 | 10.651 | -5.946 | 2.334 | 11.054 | 1.32 | 72.29 | -78.81 |
| 381 | 2jsa-10 | GLU_A_65_N | GLU_A_65_H | -5.946 | 2.334 | 11.054 | -5.117 | 1.851 | 10.738 | 1.01 | 108.23 | -30.23 |
| 382 | 2jsa-10 | GLU_A_65_N | GLU_A_65_CA | -5.946 | 2.334 | 11.054 | -6.855 | 1.646 | 11.954 | 1.45 | 51.71 | -142.88 |
| 383 | 2jsa-10 | GLU_A_65_CA | GLU_A_65_HA | -6.855 | 1.646 | 11.954 | -7.854 | 1.688 | 11.521 | 1.09 | 113.42 | 177.59 |
| 384 | 2jsa-10 | GLU_A_65_CA | GLU_A_65_C | -6.855 | 1.646 | 11.954 | -6.891 | 2.349 | 13.312 | 1.53 | 27.40 | 92.93 |
| 385 | 2jsa-10 | GLU_A_65_C | GLU_A_65_O | -6.891 | 2.349 | 13.312 | -7.875 | 2.245 | 14.044 | 1.23 | 53.51 | -173.97 |
| 386 | 2jsa-10 | GLU_A_65_C | GLY_A_66_N | -6.891 | 2.349 | 13.312 | -5.806 | 3.048 | 13.61 | 1.32 | 77.00 | 32.79 |
| 387 | 2jsa-10 | GLY_A_66_N | GLY_A_66_H | -5.806 | 3.048 | 13.61 | -5.033 | 3.088 | 12.961 | 1.01 | 129.98 | 2.96 |
| 388 | 2jsa-10 | GLY_A_66_N | GLY_A_66_CA | -5.806 | 3.048 | 13.61 | -5.7 | 3.768 | 14.867 | 1.45 | 30.07 | 81.62 |
| 389 | 2jsa-10 | GLY_A_66_CA | GLY_A_66_C | -5.7 | 3.768 | 14.867 | -6.346 | 5.152 | 14.763 | 1.53 | 93.90 | 115.02 |
| 390 | 2jsa-10 | GLY_A_66_C | GLY_A_66_O | -6.346 | 5.152 | 14.763 | -6.76 | 5.725 | 15.77 | 1.23 | 35.07 | 125.85 |
| 391 | 2jsa-10 | GLY_A_66_C | GLY_A_67_N | -6.346 | 5.152 | 14.763 | -6.413 | 5.648 | 13.537 | 1.32 | 157.79 | 97.69 |
| 392 | 2jsa-10 | GLY_A_67_N | GLY_A_67_H | -6.413 | 5.648 | 13.537 | -6.053 | 5.12 | 12.755 | 1.01 | 140.74 | -55.71 |
| 393 | 2jsa-10 | GLY_A_67_N | GLY_A_67_CA | -6.413 | 5.648 | 13.537 | -7.001 | 6.953 | 13.288 | 1.45 | 99.87 | 114.26 |
| 394 | 2jsa-10 | GLY_A_67_CA | GLY_A_67_C | -7.001 | 6.953 | 13.288 | -6.028 | 7.857 | 12.53 | 1.53 | 119.71 | 42.89 |
| 395 | 2jsa-10 | GLY_A_67_C | GLY_A_67_O | -6.028 | 7.857 | 12.53 | -6.331 | 9.02 | 12.266 | 1.23 | 102.39 | 104.60 |
| 396 | 2jsa-10 | GLY_A_67_C | THR_A_68_N | -6.028 | 7.857 | 12.53 | -4.877 | 7.29 | 12.2 | 1.32 | 104.42 | -26.23 |
| 397 | 2jsa-10 | THR_A_68_N | THR_A_68_H | -4.877 | 7.29 | 12.2 | -4.694 | 6.328 | 12.449 | 1.01 | 75.73 | -79.23 |
| 398 | 2jsa-10 | THR_A_68_N | THR_A_68_CA | -4.877 | 7.29 | 12.2 | -3.856 | 8.031 | 11.477 | 1.45 | 119.82 | 35.97 |
| 399 | 2jsa-10 | THR_A_68_CA | THR_A_68_HA | -3.856 | 8.031 | 11.477 | -3.721 | 9.001 | 11.957 | 1.09 | 63.89 | 82.08 |
| 400 | 2jsa-10 | THR_A_68_CA | THR_A_68_C | -3.856 | 8.031 | 11.477 | -4.299 | 8.277 | 10.033 | 1.53 | 160.66 | 150.96 |
| 401 | 2jsa-10 | THR_A_68_C | THR_A_68_O | -4.299 | 8.277 | 10.033 | -4.905 | 7.408 | 9.409 | 1.23 | 120.50 | -124.89 |
| 402 | 2jsa-10 | THR_A_68_C | ALA_A_69_N | -4.299 | 8.277 | 10.033 | -3.979 | 9.467 | 9.545 | 1.33 | 111.60 | 74.95 |
| 403 | 2jsa-10 | ALA_A_69_N | ALA_A_69_H | -3.979 | 9.467 | 9.545 | -3.479 | 10.134 | 10.116 | 1.01 | 55.59 | 53.14 |
| 404 | 2jsa-10 | ALA_A_69_N | ALA_A_69_CA | -3.979 | 9.467 | 9.545 | -4.337 | 9.839 | 8.187 | 1.45 | 159.18 | 133.90 |
| 405 | 2jsa-10 | ALA_A_69_CA | ALA_A_69_HA | -4.337 | 9.839 | 8.187 | -5.394 | 9.624 | 8.031 | 1.09 | 98.23 | -168.50 |
| 406 | 2jsa-10 | ALA_A_69_CA | ALA_A_69_C | -4.337 | 9.839 | 8.187 | -3.521 | 9.0 | 7.201 | 1.53 | 130.11 | -45.80 |
| 407 | 2jsa-10 | ALA_A_69_C | ALA_A_69_O | -3.521 | 9.0 | 7.201 | -2.517 | 8.397 | 7.577 | 1.23 | 72.20 | -30.99 |
| 408 | 2jsa-10 | ALA_A_69_C | PRO_A_70_N | -3.521 | 9.0 | 7.201 | -3.996 | 8.986 | 5.927 | 1.36 | 159.54 | -178.31 |
| 409 | 2jsa-10 | PRO_A_70_N | PRO_A_70_CA | -3.996 | 8.986 | 5.927 | -3.323 | 8.231 | 4.885 | 1.45 | 135.85 | -48.29 |
| 410 | 2jsa-10 | PRO_A_70_CA | PRO_A_70_HA | -3.323 | 8.231 | 4.885 | -3.06 | 7.24 | 5.254 | 1.09 | 70.21 | -75.14 |

| | | | | | | | | | | | | |
|---|---|---|---|---|---|---|---|---|---|---|---|---|
| 411 | 2jsa-10 | PRO_A_70_CA | PRO_A_70_C | -3.323 | 8.231 | 4.885 | -2.042 | 8.936 | 4.436 | 1.53 | 107.07 | 28.83 |
| 412 | 2jsa-10 | PRO_A_70_C | PRO_A_70_O | -2.042 | 8.936 | 4.436 | -1.15 | 8.309 | 3.868 | 1.23 | 117.52 | -35.10 |
| 413 | 2jsa-10 | PRO_A_70_C | LYS_A_71_N | -2.042 | 8.936 | 4.436 | -1.993 | 10.232 | 4.708 | 1.33 | 78.16 | 87.83 |
| 414 | 2jsa-10 | LYS_A_71_N | LYS_A_71_H | -1.993 | 10.232 | 4.708 | -2.765 | 10.683 | 5.178 | 1.01 | 62.27 | 149.71 |
| 415 | 2jsa-10 | LYS_A_71_N | LYS_A_71_CA | -1.993 | 10.232 | 4.708 | -0.835 | 11.031 | 4.34 | 1.45 | 104.66 | 34.61 |
| 416 | 2jsa-10 | LYS_A_71_CA | LYS_A_71_HA | -0.835 | 11.031 | 4.34 | -0.295 | 10.502 | 3.554 | 1.09 | 136.12 | -44.41 |
| 417 | 2jsa-10 | LYS_A_71_CA | LYS_A_71_C | -0.835 | 11.031 | 4.34 | 0.1 | 11.152 | 5.545 | 1.53 | 38.04 | 7.37 |
| 418 | 2jsa-10 | LYS_A_71_C | LYS_A_71_O | 0.1 | 11.152 | 5.545 | 1.118 | 11.838 | 5.477 | 1.23 | 93.17 | 33.97 |
| 419 | 2jsa-10 | LYS_A_71_C | ASP_A_72_N | 0.1 | 11.152 | 5.545 | -0.281 | 10.476 | 6.619 | 1.32 | 35.85 | -119.41 |
| 420 | 2jsa-10 | ASP_A_72_N | ASP_A_72_H | -0.281 | 10.476 | 6.619 | -1.132 | 9.933 | 6.604 | 1.01 | 90.85 | -147.46 |
| 421 | 2jsa-10 | ASP_A_72_N | ASP_A_72_CA | -0.281 | 10.476 | 6.619 | 0.511 | 10.5 | 7.837 | 1.45 | 33.05 | 1.74 |
| 422 | 2jsa-10 | ASP_A_72_CA | ASP_A_72_HA | 0.511 | 10.5 | 7.837 | 1.236 | 11.312 | 7.775 | 1.09 | 93.26 | 48.24 |
| 423 | 2jsa-10 | ASP_A_72_CA | ASP_A_72_C | 0.511 | 10.5 | 7.837 | 1.239 | 9.163 | 7.993 | 1.53 | 84.15 | -61.43 |
| 424 | 2jsa-10 | ASP_A_72_C | ASP_A_72_O | 1.239 | 9.163 | 7.993 | 2.342 | 9.113 | 8.535 | 1.23 | 63.85 | -2.60 |
| 425 | 2jsa-10 | ASP_A_72_C | VAL_A_73_N | 1.239 | 9.163 | 7.993 | 0.593 | 8.114 | 7.509 | 1.32 | 111.45 | -121.63 |
| 426 | 2jsa-10 | VAL_A_73_N | VAL_A_73_H | 0.593 | 8.114 | 7.509 | -0.312 | 8.23 | 7.077 | 1.01 | 115.34 | 172.70 |
| 427 | 2jsa-10 | VAL_A_73_N | VAL_A_73_CA | 0.593 | 8.114 | 7.509 | 1.165 | 6.78 | 7.587 | 1.45 | 86.92 | -66.79 |
| 428 | 2jsa-10 | VAL_A_73_CA | VAL_A_73_HA | 1.165 | 6.78 | 7.587 | 1.419 | 6.574 | 8.627 | 1.09 | 17.46 | -39.04 |
| 429 | 2jsa-10 | VAL_A_73_CA | VAL_A_73_C | 1.165 | 6.78 | 7.587 | 2.456 | 6.735 | 6.766 | 1.53 | 122.44 | -2.00 |
| 430 | 2jsa-10 | VAL_A_73_C | VAL_A_73_O | 2.456 | 6.735 | 6.766 | 3.453 | 6.162 | 7.201 | 1.23 | 69.28 | -29.89 |
| 431 | 2jsa-10 | VAL_A_73_C | CYS_A_74_N | 2.456 | 6.735 | 6.766 | 2.393 | 7.346 | 5.592 | 1.32 | 152.38 | 95.89 |
| 432 | 2jsa-10 | CYS_A_74_N | CYS_A_74_H | 2.393 | 7.346 | 5.592 | 1.539 | 7.797 | 5.298 | 1.01 | 106.93 | 152.16 |
| 433 | 2jsa-10 | CYS_A_74_N | CYS_A_74_CA | 2.393 | 7.346 | 5.592 | 3.545 | 7.382 | 4.705 | 1.45 | 127.58 | 1.79 |
| 434 | 2jsa-10 | CYS_A_74_CA | CYS_A_74_HA | 3.545 | 7.382 | 4.705 | 4.07 | 6.43 | 4.784 | 1.09 | 85.84 | -61.12 |
| 435 | 2jsa-10 | CYS_A_74_CA | CYS_A_74_C | 3.545 | 7.382 | 4.705 | 4.482 | 8.495 | 5.18 | 1.53 | 71.92 | 49.91 |
| 436 | 2jsa-10 | CYS_A_74_C | CYS_A_74_O | 4.482 | 8.495 | 5.18 | 4.665 | 9.494 | 4.487 | 1.23 | 124.31 | 79.62 |
| 437 | 2jsa-10 | CYS_A_74_C | THR_A_75_N | 4.482 | 8.495 | 5.18 | 5.049 | 8.284 | 6.358 | 1.32 | 27.18 | -20.41 |
| 438 | 2jsa-10 | THR_A_75_N | THR_A_75_H | 5.049 | 8.284 | 6.358 | 4.852 | 7.438 | 6.874 | 1.01 | 59.29 | -103.11 |
| 439 | 2jsa-10 | THR_A_75_N | THR_A_75_CA | 5.049 | 8.284 | 6.358 | 5.963 | 9.256 | 6.933 | 1.45 | 66.69 | 46.76 |
| 440 | 2jsa-10 | THR_A_75_CA | THR_A_75_HA | 5.963 | 9.256 | 6.933 | 6.867 | 9.29 | 6.324 | 1.09 | 123.95 | 2.15 |
| 441 | 2jsa-10 | THR_A_75_CA | THR_A_75_C | 5.963 | 9.256 | 6.933 | 6.365 | 8.839 | 8.349 | 1.53 | 22.25 | -46.05 |
| 442 | 2jsa-10 | THR_A_75_C | THR_A_75_O | 6.365 | 8.839 | 8.349 | 7.516 | 8.478 | 8.591 | 1.23 | 78.66 | -17.41 |
| 443 | 2jsa-10 | THR_A_75_C | LYS_A_76_N | 6.365 | 8.839 | 8.349 | 5.394 | 8.899 | 9.248 | 1.32 | 47.26 | 176.46 |
| 444 | 2jsa-10 | LYS_A_76_N | LYS_A_76_H | 5.394 | 8.899 | 9.248 | 4.47 | 9.203 | 8.978 | 1.01 | 105.51 | 161.79 |
| 445 | 2jsa-10 | LYS_A_76_N | LYS_A_76_CA | 5.394 | 8.899 | 9.248 | 5.632 | 8.532 | 10.633 | 1.45 | 17.53 | -57.04 |
| 446 | 2jsa-10 | LYS_A_76_CA | LYS_A_76_HA | 5.632 | 8.532 | 10.633 | 6.44 | 9.158 | 11.014 | 1.09 | 69.56 | 37.77 |
| 447 | 2jsa-10 | LYS_A_76_CA | LYS_A_76_C | 5.632 | 8.532 | 10.633 | 6.101 | 7.077 | 10.697 | 1.53 | 87.60 | -72.13 |
| 448 | 2jsa-10 | LYS_A_76_C | LYS_A_76_O | 6.101 | 7.077 | 10.697 | 6.945 | 6.728 | 11.521 | 1.23 | 47.94 | -22.47 |
| 449 | 2jsa-10 | LYS_A_76_C | LEU_A_77_N | 6.101 | 7.077 | 10.697 | 5.533 | 6.266 | 9.815 | 1.33 | 131.69 | -125.01 |
| 450 | 2jsa-10 | LEU_A_77_N | LEU_A_77_H | 5.533 | 6.266 | 9.815 | 4.846 | 6.622 | 9.166 | 1.01 | 129.99 | 152.61 |
| 451 | 2jsa-10 | LEU_A_77_N | LEU_A_77_CA | 5.533 | 6.266 | 9.815 | 5.882 | 4.857 | 9.761 | 1.45 | 92.13 | -76.09 |
| 452 | 2jsa-10 | LEU_A_77_CA | LEU_A_77_HA | 5.882 | 4.857 | 9.761 | 5.878 | 4.469 | 10.78 | 1.09 | 20.85 | -90.59 |
| 453 | 2jsa-10 | LEU_A_77_CA | LEU_A_77_C | 5.882 | 4.857 | 9.761 | 7.302 | 4.709 | 9.212 | 1.53 | 111.03 | -5.95 |
| 454 | 2jsa-10 | LEU_A_77_C | LEU_A_77_O | 7.302 | 4.709 | 9.212 | 7.85 | 3.608 | 9.189 | 1.23 | 91.07 | -63.54 |
| 455 | 2jsa-10 | LEU_A_77_C | ASN_A_78_N | 7.302 | 4.709 | 9.212 | 7.858 | 5.833 | 8.785 | 1.32 | 108.80 | 63.68 |
| 456 | 2jsa-10 | ASN_A_78_N | ASN_A_78_H | 7.858 | 5.833 | 8.785 | 7.349 | 6.704 | 8.836 | 1.01 | 87.11 | 120.30 |
| 457 | 2jsa-10 | ASN_A_78_N | ASN_A_78_CA | 7.858 | 5.833 | 8.785 | 9.203 | 5.842 | 8.238 | 1.45 | 112.13 | 0.38 |
| 458 | 2jsa-10 | ASN_A_78_CA | ASN_A_78_HA | 9.203 | 5.842 | 8.238 | 9.528 | 6.875 | 8.108 | 1.09 | 96.85 | 72.54 |
| 459 | 2jsa-10 | ASN_A_78_CA | ASN_A_78_C | 9.203 | 5.842 | 8.238 | 9.207 | 5.122 | 6.888 | 1.53 | 151.93 | -89.68 |
| 460 | 2jsa-10 | ASN_A_78_C | ASN_A_78_O | 9.207 | 5.122 | 6.888 | 10.214 | 4.534 | 6.497 | 1.23 | 108.54 | -30.28 |
| 461 | 2jsa-10 | ASN_A_78_C | GLU_A_79_N | 9.207 | 5.122 | 6.888 | 8.068 | 5.191 | 6.215 | 1.32 | 120.53 | 176.53 |
| 462 | 2jsa-10 | GLU_A_79_N | GLU_A_79_H | 8.068 | 5.191 | 6.215 | 7.28 | 5.693 | 6.599 | 1.01 | 67.66 | 147.50 |
| 463 | 2jsa-10 | GLU_A_79_N | GLU_A_79_CA | 8.068 | 5.191 | 6.215 | 7.926 | 4.553 | 4.918 | 1.45 | 153.25 | -102.55 |
| 464 | 2jsa-10 | GLU_A_79_CA | GLU_A_79_HA | 7.926 | 4.553 | 4.918 | 8.651 | 3.741 | 4.86 | 1.09 | 93.05 | -48.24 |
| 465 | 2jsa-10 | GLU_A_79_CA | GLU_A_79_C | 7.926 | 4.553 | 4.918 | 8.231 | 5.551 | 3.799 | 1.53 | 137.00 | 73.01 |
| 466 | 2jsa-10 | GLU_A_79_C | GLU_A_79_O | 8.231 | 5.551 | 3.799 | 9.366 | 5.635 | 3.33 | 1.23 | 112.40 | 4.23 |
| 467 | 2jsa-10 | GLU_A_79_C | CYS_A_80_N | 8.231 | 5.551 | 3.799 | 7.2 | 6.283 | 3.404 | 1.32 | 107.35 | 144.63 |
| 468 | 2jsa-10 | CYS_A_80_N | CYS_A_80_H | 7.2 | 6.283 | 3.404 | 6.296 | 6.162 | 3.836 | 1.01 | 64.66 | -172.38 |
| 469 | 2jsa-10 | CYS_A_80_N | CYS_A_80_CA | 7.2 | 6.283 | 3.404 | 7.344 | 7.271 | 2.349 | 1.45 | 136.58 | 81.71 |
| 470 | 2jsa-10 | CYS_A_80_CA | CYS_A_80_HA | 7.344 | 7.271 | 2.349 | 7.765 | 6.778 | 1.473 | 1.09 | 143.50 | -49.50 |
| 471 | 2jsa-10 | CYS_A_80_CA | CYS_A_80_C | 7.344 | 7.271 | 2.349 | 8.328 | 8.342 | 2.825 | 1.53 | 71.88 | 47.42 |
| 472 | 2jsa-10 | CYS_A_80_C | CYS_A_80_O | 8.328 | 8.342 | 2.825 | 8.295 | 8.75 | 3.985 | 1.23 | 19.44 | 94.62 |
| 473 | 2jsa-10 | CYS_A_80_C | PRO_A_81_N | 8.328 | 8.342 | 2.825 | 9.205 | 8.779 | 1.881 | 1.36 | 133.93 | 26.49 |
| 474 | 2jsa-10 | PRO_A_81_N | PRO_A_81_CA | 9.205 | 8.779 | 1.881 | 10.197 | 9.793 | 2.193 | 1.45 | 77.60 | 45.63 |
| 475 | 2jsa-10 | PRO_A_81_CA | PRO_A_81_HA | 10.197 | 9.793 | 2.193 | 10.673 | 9.569 | 3.148 | 1.09 | 28.85 | -25.20 |
| 476 | 2jsa-10 | PRO_A_81_CA | PRO_A_81_C | 10.197 | 9.793 | 2.193 | 9.554 | 11.178 | 2.284 | 1.53 | 86.59 | 114.90 |
| 477 | 2jsa-10 | PRO_A_81_C | PRO_A_81_O | 9.554 | 11.178 | 2.284 | 9.673 | 11.855 | 3.304 | 1.23 | 33.98 | 80.03 |
| 1 | 2jsa-11 | ARG_A_1_N | ARG_A_1_H | 13.063 | -0.16 | -2.146 | 14.039 | 0.095 | -2.083 | 1.01 | 86.43 | 14.64 |
| 2 | 2jsa-11 | ARG_A_1_N | ARG_A_1_CA | 13.063 | -0.16 | -2.146 | 12.061 | 0.852 | -1.859 | 1.45 | 78.61 | 134.72 |
| 3 | 2jsa-11 | ARG_A_1_CA | ARG_A_1_HA | 12.061 | 0.852 | -1.859 | 11.449 | 0.993 | -2.75 | 1.09 | 144.82 | 167.03 |
| 4 | 2jsa-11 | ARG_A_1_CA | ARG_A_1_C | 12.061 | 0.852 | -1.859 | 12.734 | 2.179 | -1.501 | 1.53 | 76.47 | 63.11 |
| 5 | 2jsa-11 | ARG_A_1_C | ARG_A_1_O | 12.734 | 2.179 | -1.501 | 12.977 | 2.46 | -0.329 | 1.23 | 17.59 | 49.15 |
| 6 | 2jsa-11 | ARG_A_1_C | SER_A_2_N | 12.734 | 2.179 | -1.501 | 13.017 | 2.959 | -2.534 | 1.32 | 141.23 | 70.06 |
| 7 | 2jsa-11 | SER_A_2_N | SER_A_2_H | 13.017 | 2.959 | -2.534 | 12.789 | 2.664 | -3.472 | 1.01 | 158.32 | -127.70 |
| 8 | 2jsa-11 | SER_A_2_N | SER_A_2_CA | 13.017 | 2.959 | -2.534 | 13.657 | 4.25 | -2.343 | 1.45 | 82.45 | 63.63 |
| 9 | 2jsa-11 | SER_A_2_CA | SER_A_2_HA | 13.657 | 4.25 | -2.343 | 13.836 | 4.389 | -1.277 | 1.09 | 12.00 | 37.83 |
| 10 | 2jsa-11 | SER_A_2_CA | SER_A_2_C | 13.657 | 4.25 | -2.343 | 12.737 | 5.369 | -2.836 | 1.53 | 108.79 | 129.43 |
| 11 | 2jsa-11 | SER_A_2_C | SER_A_2_O | 12.737 | 5.369 | -2.836 | 13.205 | 6.442 | -3.209 | 1.23 | 107.67 | 66.44 |
| 12 | 2jsa-11 | SER_A_2_C | ALA_A_3_N | 12.737 | 5.369 | -2.836 | 11.445 | 5.078 | -2.82 | 1.32 | 89.31 | -167.31 |
| 13 | 2jsa-11 | ALA_A_3_N | ALA_A_3_H | 11.445 | 5.078 | -2.82 | 11.132 | 4.173 | -2.501 | 1.01 | 71.58 | -109.08 |
| 14 | 2jsa-11 | ALA_A_3_N | ALA_A_3_CA | 11.445 | 5.078 | -2.82 | 10.454 | 6.046 | -3.259 | 1.45 | 107.58 | 135.67 |
| 15 | 2jsa-11 | ALA_A_3_CA | ALA_A_3_HA | 10.454 | 6.046 | -3.259 | 10.283 | 6.762 | -2.455 | 1.09 | 42.48 | 103.43 |
| 16 | 2jsa-11 | ALA_A_3_CA | ALA_A_3_C | 10.454 | 6.046 | -3.259 | 9.138 | 5.324 | -3.552 | 1.53 | 101.05 | -151.25 |
| 17 | 2jsa-11 | ALA_A_3_C | ALA_A_3_O | 9.138 | 5.324 | -3.552 | 8.081 | 5.733 | -3.073 | 1.23 | 67.09 | 158.85 |
| 18 | 2jsa-11 | ALA_A_3_C | LEU_A_4_N | 9.138 | 5.324 | -3.552 | 9.243 | 4.263 | -4.339 | 1.33 | 126.43 | -84.35 |
| 19 | 2jsa-11 | LEU_A_4_N | LEU_A_4_H | 9.243 | 4.263 | -4.339 | 10.145 | 3.984 | -4.698 | 1.01 | 110.82 | -17.19 |
| 20 | 2jsa-11 | LEU_A_4_N | LEU_A_4_CA | 9.243 | 4.263 | -4.339 | 8.073 | 3.481 | -4.702 | 1.45 | 104.46 | -146.24 |
| 21 | 2jsa-11 | LEU_A_4_CA | LEU_A_4_HA | 8.073 | 3.481 | -4.702 | 7.49 | 4.06 | -5.418 | 1.09 | 131.07 | 135.20 |
| 22 | 2jsa-11 | LEU_A_4_CA | LEU_A_4_C | 8.073 | 3.481 | -4.702 | 7.205 | 3.265 | -3.462 | 1.53 | 35.80 | -166.03 |
| 23 | 2jsa-11 | LEU_A_4_C | LEU_A_4_O | 7.205 | 3.265 | -3.462 | 6.027 | 3.618 | -3.455 | 1.23 | 89.67 | 163.32 |
| 24 | 2jsa-11 | LEU_A_4_C | SER_A_5_N | 7.205 | 3.265 | -3.462 | 7.82 | 2.686 | -2.441 | 1.33 | 39.60 | -43.27 |
| 25 | 2jsa-11 | SER_A_5_N | SER_A_5_H | 7.82 | 2.686 | -2.441 | 8.793 | 2.423 | -2.516 | 1.01 | 94.26 | -15.13 |
| 26 | 2jsa-11 | SER_A_5_N | SER_A_5_CA | 7.82 | 2.686 | -2.441 | 7.118 | 2.418 | -1.198 | 1.45 | 31.15 | -159.10 |
| 27 | 2jsa-11 | SER_A_5_CA | SER_A_5_HA | 7.118 | 2.418 | -1.198 | 6.538 | 1.504 | -1.323 | 1.09 | 96.59 | -122.40 |
| 28 | 2jsa-11 | SER_A_5_CA | SER_A_5_C | 7.118 | 2.418 | -1.198 | 6.16 | 3.569 | -0.88 | 1.53 | 78.01 | 129.77 |
| 29 | 2jsa-11 | SER_A_5_C | SER_A_5_O | 6.16 | 3.569 | -0.88 | 5.057 | 3.343 | -0.383 | 1.23 | 66.18 | -168.42 |

| | | | | | | | | | | | | |
|---|---|---|---|---|---|---|---|---|---|---|---|---|
| 30 | 2jsa-11 | SER_A_5_C | CYS_A_6_N | 6.16 | 3.569 | -0.88 | 6.615 | 4.776 | -1.18 | 1.32 | 103.09 | 69.35 |
| 31 | 2jsa-11 | CYS_A_6_N | CYS_A_6_H | 6.615 | 4.776 | -1.18 | 7.533 | 4.887 | -1.587 | 1.01 | 113.76 | 6.89 |
| 32 | 2jsa-11 | CYS_A_6_N | CYS_A_6_CA | 6.615 | 4.776 | -1.18 | 5.811 | 5.961 | -0.933 | 1.45 | 80.21 | 124.16 |
| 33 | 2jsa-11 | CYS_A_6_CA | CYS_A_6_HA | 5.811 | 5.961 | -0.933 | 5.641 | 6.042 | 0.141 | 1.09 | 9.94 | 154.52 |
| 34 | 2jsa-11 | CYS_A_6_CA | CYS_A_6_C | 5.811 | 5.961 | -0.933 | 4.461 | 5.78 | -1.627 | 1.53 | 117.00 | -172.36 |
| 35 | 2jsa-11 | CYS_A_6_C | CYS_A_6_O | 4.461 | 5.78 | -1.627 | 3.443 | 5.574 | -0.968 | 1.23 | 57.61 | -168.56 |
| 36 | 2jsa-11 | CYS_A_6_C | GLN_A_7_N | 4.461 | 5.78 | -1.627 | 4.495 | 5.862 | -2.949 | 1.32 | 176.16 | 67.48 |
| 37 | 2jsa-11 | GLN_A_7_N | GLN_A_7_H | 4.495 | 5.862 | -2.949 | 5.369 | 6.034 | -3.425 | 1.01 | 118.12 | 11.13 |
| 38 | 2jsa-11 | GLN_A_7_N | GLN_A_7_CA | 4.495 | 5.862 | -2.949 | 3.286 | 5.709 | -3.74 | 1.45 | 122.99 | -172.79 |
| 39 | 2jsa-11 | GLN_A_7_CA | GLN_A_7_HA | 3.286 | 5.709 | -3.74 | 2.68 | 6.605 | -3.605 | 1.09 | 82.89 | 124.07 |
| 40 | 2jsa-11 | GLN_A_7_CA | GLN_A_7_C | 3.286 | 5.709 | -3.74 | 2.48 | 4.505 | -3.249 | 1.53 | 71.28 | -123.80 |
| 41 | 2jsa-11 | GLN_A_7_C | GLN_A_7_O | 2.48 | 4.505 | -3.249 | 1.283 | 4.619 | -2.987 | 1.23 | 77.71 | 174.56 |
| 42 | 2jsa-11 | GLN_A_7_C | MET_A_8_N | 2.48 | 4.505 | -3.249 | 3.167 | 3.378 | -3.141 | 1.32 | 85.32 | -58.63 |
| 43 | 2jsa-11 | MET_A_8_N | MET_A_8_H | 3.167 | 3.378 | -3.141 | 4.149 | 3.357 | -3.376 | 1.01 | 103.46 | -1.23 |
| 44 | 2jsa-11 | MET_A_8_N | MET_A_8_CA | 3.167 | 3.378 | -3.141 | 2.53 | 2.154 | -2.687 | 1.45 | 71.79 | -117.49 |
| 45 | 2jsa-11 | MET_A_8_CA | MET_A_8_HA | 2.53 | 2.154 | -2.687 | 1.898 | 1.772 | -3.488 | 1.09 | 137.33 | -148.85 |
| 46 | 2jsa-11 | MET_A_8_CA | MET_A_8_C | 2.53 | 2.154 | -2.687 | 1.666 | 2.411 | -1.452 | 1.53 | 36.13 | 163.43 |
| 47 | 2jsa-11 | MET_A_8_C | MET_A_8_O | 1.666 | 2.411 | -1.452 | 0.527 | 1.952 | -1.38 | 1.23 | 86.64 | -158.05 |
| 48 | 2jsa-11 | MET_A_8_C | CYS_A_9_N | 1.666 | 2.411 | -1.452 | 2.239 | 3.144 | -0.51 | 1.32 | 44.64 | 51.98 |
| 49 | 2jsa-11 | CYS_A_9_N | CYS_A_9_H | 2.239 | 3.144 | -0.51 | 3.18 | 3.49 | -0.637 | 1.01 | 97.22 | 20.19 |
| 50 | 2jsa-11 | CYS_A_9_N | CYS_A_9_CA | 2.239 | 3.144 | -0.51 | 1.536 | 3.468 | 0.72 | 1.45 | 32.18 | 155.26 |
| 51 | 2jsa-11 | CYS_A_9_CA | CYS_A_9_HA | 1.536 | 3.468 | 0.72 | 1.279 | 2.536 | 1.224 | 1.09 | 62.47 | -105.42 |
| 52 | 2jsa-11 | CYS_A_9_CA | CYS_A_9_C | 1.536 | 3.468 | 0.72 | 0.242 | 4.199 | 0.354 | 1.53 | 103.83 | 150.54 |
| 53 | 2jsa-11 | CYS_A_9_C | CYS_A_9_O | 0.242 | 4.199 | 0.354 | -0.764 | 4.071 | 1.05 | 1.23 | 55.54 | -172.75 |
| 54 | 2jsa-11 | CYS_A_9_C | GLU_A_10_N | 0.242 | 4.199 | 0.354 | 0.31 | 4.947 | -0.737 | 1.32 | 145.46 | 84.81 |
| 55 | 2jsa-11 | GLU_A_10_N | GLU_A_10_H | 0.31 | 4.947 | -0.737 | 1.172 | 5.004 | -1.261 | 1.01 | 121.24 | 3.78 |
| 56 | 2jsa-11 | GLU_A_10_N | GLU_A_10_CA | 0.31 | 4.947 | -0.737 | -0.843 | 5.697 | -1.203 | 1.45 | 108.72 | 146.96 |
| 57 | 2jsa-11 | GLU_A_10_CA | GLU_A_10_HA | -0.843 | 5.697 | -1.203 | -1.39 | 6.052 | -0.329 | 1.09 | 36.73 | 147.02 |
| 58 | 2jsa-11 | GLU_A_10_CA | GLU_A_10_C | -0.843 | 5.697 | -1.203 | -1.771 | 4.793 | -2.016 | 1.53 | 122.11 | -135.75 |
| 59 | 2jsa-11 | GLU_A_10_C | GLU_A_10_O | -1.771 | 4.793 | -2.016 | -2.838 | 5.223 | -2.453 | 1.23 | 110.80 | 158.05 |
| 60 | 2jsa-11 | GLU_A_10_C | LEU_A_11_N | -1.771 | 4.793 | -2.016 | -1.332 | 3.556 | -2.196 | 1.32 | 97.81 | -70.46 |
| 61 | 2jsa-11 | LEU_A_11_N | LEU_A_11_H | -1.332 | 3.556 | -2.196 | -0.444 | 3.27 | -1.808 | 1.01 | 67.42 | -17.85 |
| 62 | 2jsa-11 | LEU_A_11_N | LEU_A_11_CA | -1.332 | 3.556 | -2.196 | -2.11 | 2.587 | -2.949 | 1.45 | 121.21 | -128.76 |
| 63 | 2jsa-11 | LEU_A_11_CA | LEU_A_11_HA | -2.11 | 2.587 | -2.949 | -2.775 | 3.134 | -3.617 | 1.09 | 127.80 | 140.56 |
| 64 | 2jsa-11 | LEU_A_11_CA | LEU_A_11_C | -2.11 | 2.587 | -2.949 | -2.981 | 1.778 | -1.985 | 1.53 | 50.96 | -137.11 |
| 65 | 2jsa-11 | LEU_A_11_C | LEU_A_11_O | -2.981 | 1.778 | -1.985 | -4.16 | 1.549 | -2.252 | 1.23 | 102.53 | -169.01 |
| 66 | 2jsa-11 | LEU_A_11_C | VAL_A_12_N | -2.981 | 1.778 | -1.985 | -2.367 | 1.369 | -0.885 | 1.32 | 33.85 | -33.67 |
| 67 | 2jsa-11 | VAL_A_12_N | VAL_A_12_H | -2.367 | 1.369 | -0.885 | -1.395 | 1.595 | -0.731 | 1.01 | 81.23 | 13.09 |
| 68 | 2jsa-11 | VAL_A_12_N | VAL_A_12_CA | -2.367 | 1.369 | -0.885 | -3.072 | 0.59 | 0.12 | 1.45 | 46.27 | -132.15 |
| 69 | 2jsa-11 | VAL_A_12_CA | VAL_A_12_HA | -3.072 | 0.59 | 0.12 | -3.553 | -0.254 | -0.373 | 1.09 | 116.91 | -119.68 |
| 70 | 2jsa-11 | VAL_A_12_CA | VAL_A_12_C | -3.072 | 0.59 | 0.12 | -4.163 | 1.456 | 0.755 | 1.53 | 65.49 | 141.56 |
| 71 | 2jsa-11 | VAL_A_12_C | VAL_A_12_O | -4.163 | 1.456 | 0.755 | -5.249 | 0.966 | 1.059 | 1.23 | 75.69 | -155.72 |
| 72 | 2jsa-11 | VAL_A_12_C | VAL_A_13_N | -4.163 | 1.456 | 0.755 | -3.835 | 2.727 | 0.937 | 1.33 | 82.11 | 75.53 |
| 73 | 2jsa-11 | VAL_A_13_N | VAL_A_13_H | -3.835 | 2.727 | 0.937 | -2.922 | 3.061 | 0.664 | 1.01 | 105.69 | 20.09 |
| 74 | 2jsa-11 | VAL_A_13_N | VAL_A_13_CA | -3.835 | 2.727 | 0.937 | -4.774 | 3.663 | 1.53 | 1.45 | 65.90 | 135.09 |
| 75 | 2jsa-11 | VAL_A_13_CA | VAL_A_13_HA | -4.774 | 3.663 | 1.53 | -5.167 | 3.223 | 2.446 | 1.09 | 32.78 | -131.77 |
| 76 | 2jsa-11 | VAL_A_13_CA | VAL_A_13_C | -4.774 | 3.663 | 1.53 | -5.944 | 3.884 | 0.57 | 1.53 | 128.88 | 169.30 |
| 77 | 2jsa-11 | VAL_A_13_C | VAL_A_13_O | -5.944 | 3.884 | 0.57 | -7.095 | 3.628 | 0.919 | 1.23 | 73.51 | -167.46 |
| 78 | 2jsa-11 | VAL_A_13_C | LYS_A_14_N | -5.944 | 3.884 | 0.57 | -5.608 | 4.357 | -0.622 | 1.33 | 154.05 | 54.61 |
| 79 | 2jsa-11 | LYS_A_14_N | LYS_A_14_H | -5.608 | 4.357 | -0.622 | -4.64 | 4.544 | -0.841 | 1.01 | 102.52 | 10.93 |
| 80 | 2jsa-11 | LYS_A_14_N | LYS_A_14_CA | -5.608 | 4.357 | -0.622 | -6.617 | 4.616 | -1.635 | 1.45 | 134.20 | 165.60 |
| 81 | 2jsa-11 | LYS_A_14_CA | LYS_A_14_HA | -6.617 | 4.616 | -1.635 | -7.204 | 5.477 | -1.315 | 1.09 | 72.93 | 124.28 |
| 82 | 2jsa-11 | LYS_A_14_CA | LYS_A_14_C | -6.617 | 4.616 | -1.635 | -7.562 | 3.415 | -1.726 | 1.53 | 93.41 | -128.20 |
| 83 | 2jsa-11 | LYS_A_14_C | LYS_A_14_O | -7.562 | 3.415 | -1.726 | -8.745 | 3.573 | -2.021 | 1.23 | 103.88 | 172.39 |
| 84 | 2jsa-11 | LYS_A_14_C | LYS_A_15_N | -7.562 | 3.415 | -1.726 | -7.004 | 2.242 | -1.466 | 1.32 | 78.68 | -64.56 |
| 85 | 2jsa-11 | LYS_A_15_N | LYS_A_15_H | -7.004 | 2.242 | -1.466 | -6.023 | 2.187 | -1.229 | 1.01 | 76.44 | -3.21 |
| 86 | 2jsa-11 | LYS_A_15_N | LYS_A_15_CA | -7.004 | 2.242 | -1.466 | -7.781 | 1.016 | -1.514 | 1.45 | 91.89 | -122.37 |
| 87 | 2jsa-11 | LYS_A_15_CA | LYS_A_15_HA | -7.781 | 1.016 | -1.514 | -8.513 | 1.112 | -2.316 | 1.09 | 137.37 | 172.53 |
| 88 | 2jsa-11 | LYS_A_15_CA | LYS_A_15_C | -7.781 | 1.016 | -1.514 | -8.549 | 0.852 | -0.201 | 1.53 | 30.88 | -167.95 |
| 89 | 2jsa-11 | LYS_A_15_C | LYS_A_15_O | -8.549 | 0.852 | -0.201 | -9.708 | 0.438 | -0.203 | 1.23 | 90.09 | -160.34 |
| 90 | 2jsa-11 | LYS_A_15_C | TYR_A_16_N | -8.549 | 0.852 | -0.201 | -7.872 | 1.183 | 0.888 | 1.32 | 34.68 | 26.06 |
| 91 | 2jsa-11 | TYR_A_16_N | TYR_A_16_H | -7.872 | 1.183 | 0.888 | -6.92 | 1.514 | 0.814 | 1.01 | 94.20 | 19.17 |
| 92 | 2jsa-11 | TYR_A_16_N | TYR_A_16_CA | -7.872 | 1.183 | 0.888 | -8.475 | 1.078 | 2.205 | 1.45 | 24.93 | -170.12 |
| 93 | 2jsa-11 | TYR_A_16_CA | TYR_A_16_HA | -8.475 | 1.078 | 2.205 | -8.973 | 0.115 | 2.316 | 1.09 | 84.15 | -117.35 |
| 94 | 2jsa-11 | TYR_A_16_CA | TYR_A_16_C | -8.475 | 1.078 | 2.205 | -9.476 | 2.21 | 2.443 | 1.53 | 81.05 | 131.49 |
| 95 | 2jsa-11 | TYR_A_16_C | TYR_A_16_O | -9.476 | 2.21 | 2.443 | -10.681 | 1.973 | 2.518 | 1.23 | 86.51 | -168.87 |
| 96 | 2jsa-11 | TYR_A_16_C | GLU_A_17_N | -9.476 | 2.21 | 2.443 | -8.94 | 3.416 | 2.555 | 1.32 | 85.15 | 66.04 |
| 97 | 2jsa-11 | GLU_A_17_N | GLU_A_17_H | -8.94 | 3.416 | 2.555 | -7.939 | 3.537 | 2.484 | 1.01 | 94.03 | 6.89 |
| 98 | 2jsa-11 | GLU_A_17_N | GLU_A_17_CA | -8.94 | 3.416 | 2.555 | -9.772 | 4.587 | 2.781 | 1.45 | 81.06 | 125.39 |
| 99 | 2jsa-11 | GLU_A_17_CA | GLU_A_17_HA | -9.772 | 4.587 | 2.781 | -10.106 | 4.568 | 3.819 | 1.09 | 17.86 | -176.74 |
| 100 | 2jsa-11 | GLU_A_17_CA | GLU_A_17_C | -9.772 | 4.587 | 2.781 | -11.006 | 4.544 | 1.879 | 1.53 | 126.15 | -178.00 |
| 101 | 2jsa-11 | GLU_A_17_C | GLU_A_17_O | -11.006 | 4.544 | 1.879 | -12.113 | 4.846 | 2.321 | 1.23 | 68.93 | 164.74 |
| 102 | 2jsa-11 | GLU_A_17_C | GLY_A_18_N | -11.006 | 4.544 | 1.879 | -10.773 | 4.167 | 0.63 | 1.33 | 160.46 | -58.28 |
| 103 | 2jsa-11 | GLY_A_18_N | GLY_A_18_H | -10.773 | 4.167 | 0.63 | -9.835 | 3.934 | 0.336 | 1.01 | 106.92 | -13.95 |
| 104 | 2jsa-11 | GLY_A_18_N | GLY_A_18_CA | -10.773 | 4.167 | 0.63 | -11.853 | 4.081 | -0.339 | 1.45 | 131.81 | -175.45 |
| 105 | 2jsa-11 | GLY_A_18_CA | GLY_A_18_C | -11.853 | 4.081 | -0.339 | -12.065 | 2.637 | -0.796 | 1.53 | 107.39 | -98.35 |
| 106 | 2jsa-11 | GLY_A_18_C | GLY_A_18_O | -12.065 | 2.637 | -0.796 | -12.184 | 2.371 | -1.992 | 1.23 | 166.31 | -114.10 |
| 107 | 2jsa-11 | GLY_A_18_C | SER_A_19_N | -12.065 | 2.637 | -0.796 | -12.107 | 1.741 | 0.179 | 1.32 | 42.61 | -92.68 |
| 108 | 2jsa-11 | SER_A_19_N | SER_A_19_H | -12.107 | 1.741 | 0.179 | -12.003 | 2.031 | 1.141 | 1.01 | 17.76 | 70.27 |
| 109 | 2jsa-11 | SER_A_19_N | SER_A_19_CA | -12.107 | 1.741 | 0.179 | -12.303 | 0.33 | -0.109 | 1.45 | 101.43 | -97.91 |
| 110 | 2jsa-11 | SER_A_19_CA | SER_A_19_HA | -12.303 | 0.33 | -0.109 | -11.761 | 0.09 | -1.024 | 1.09 | 147.06 | -23.88 |
| 111 | 2jsa-11 | SER_A_19_CA | SER_A_19_C | -12.303 | 0.33 | -0.109 | -13.789 | 0.042 | -0.329 | 1.53 | 98.27 | -169.03 |
| 112 | 2jsa-11 | SER_A_19_C | SER_A_19_O | -13.789 | 0.042 | -0.329 | -14.164 | -0.588 | -1.318 | 1.23 | 143.45 | -120.76 |
| 113 | 2jsa-11 | SER_A_19_C | ALA_A_20_N | -13.789 | 0.042 | -0.329 | -14.596 | 0.518 | 0.608 | 1.33 | 45.00 | 149.47 |
| 114 | 2jsa-11 | ALA_A_20_N | ALA_A_20_H | -14.596 | 0.518 | 0.608 | -14.218 | 1.027 | 1.394 | 1.01 | 38.89 | 53.40 |
| 115 | 2jsa-11 | ALA_A_20_N | ALA_A_20_CA | -14.596 | 0.518 | 0.608 | -16.034 | 0.32 | 0.529 | 1.45 | 93.12 | -172.16 |
| 116 | 2jsa-11 | ALA_A_20_CA | ALA_A_20_HA | -16.034 | 0.32 | 0.529 | -16.527 | 1.092 | 1.119 | 1.09 | 57.21 | 122.56 |
| 117 | 2jsa-11 | ALA_A_20_CA | ALA_A_20_C | -16.034 | 0.32 | 0.529 | -16.389 | -1.044 | 1.123 | 1.53 | 67.15 | -104.59 |
| 118 | 2jsa-11 | ALA_A_20_C | ALA_A_20_O | -16.389 | -1.044 | 1.123 | -16.97 | -1.889 | 0.443 | 1.23 | 123.55 | -124.51 |
| 119 | 2jsa-11 | ALA_A_20_C | ASP_A_21_N | -16.389 | -1.044 | 1.123 | -16.026 | -1.218 | 2.385 | 1.32 | 17.69 | -25.61 |
| 120 | 2jsa-11 | ASP_A_21_N | ASP_A_21_H | -16.026 | -1.218 | 2.385 | -15.55 | -0.479 | 2.884 | 1.01 | 60.42 | 57.21 |
| 121 | 2jsa-11 | ASP_A_21_N | ASP_A_21_CA | -16.026 | -1.218 | 2.385 | -16.299 | -2.465 | 3.078 | 1.45 | 61.50 | -102.35 |
| 122 | 2jsa-11 | ASP_A_21_CA | ASP_A_21_HA | -16.299 | -2.465 | 3.078 | -17.351 | -2.491 | 3.362 | 1.09 | 74.90 | -178.58 |
| 123 | 2jsa-11 | ASP_A_21_CA | ASP_A_21_C | -16.299 | -2.465 | 3.078 | -15.421 | -2.552 | 4.328 | 1.53 | 35.22 | -5.66 |
| 124 | 2jsa-11 | ASP_A_21_C | ASP_A_21_O | -15.421 | -2.552 | 4.328 | -15.791 | -3.199 | 5.307 | 1.23 | 37.28 | -119.76 |
| 125 | 2jsa-11 | ASP_A_21_C | LYS_A_22_N | -15.421 | -2.552 | 4.328 | -14.275 | -1.892 | 4.254 | 1.32 | 93.20 | 29.94 |

| | | | | | | | | | | | | |
|---|---|---|---|---|---|---|---|---|---|---|---|---|
| 126 | 2jsa-11 | LYS_A_22_N | LYS_A_22_H | -14.275 | -1.892 | 4.254 | -14.035 | -1.381 | 3.417 | 1.01 | 146.00 | 64.84 |
| 127 | 2jsa-11 | LYS_A_22_N | LYS_A_22_CA | -14.275 | -1.892 | 4.254 | -13.342 | -1.887 | 5.367 | 1.45 | 39.97 | 0.31 |
| 128 | 2jsa-11 | LYS_A_22_CA | LYS_A_22_HA | -13.342 | -1.887 | 5.367 | -12.5 | -1.248 | 5.099 | 1.09 | 104.23 | 37.20 |
| 129 | 2jsa-11 | LYS_A_22_CA | LYS_A_22_C | -13.342 | -1.887 | 5.367 | -12.8 | -3.302 | 5.581 | 1.53 | 81.96 | -69.04 |
| 130 | 2jsa-11 | LYS_A_22_C | LYS_A_22_O | -12.8 | -3.302 | 5.581 | -13.019 | -3.902 | 6.632 | 1.23 | 31.29 | -110.05 |
| 131 | 2jsa-11 | LYS_A_22_C | ASP_A_23_N | -12.8 | -3.302 | 5.581 | -12.105 | -3.795 | 4.566 | 1.33 | 139.99 | -35.35 |
| 132 | 2jsa-11 | ASP_A_23_N | ASP_A_23_H | -12.105 | -3.795 | 4.566 | -11.965 | -3.245 | 3.73 | 1.01 | 145.83 | 75.72 |
| 133 | 2jsa-11 | ASP_A_23_N | ASP_A_23_CA | -12.105 | -3.795 | 4.566 | -11.532 | -5.129 | 4.629 | 1.45 | 87.52 | -66.75 |
| 134 | 2jsa-11 | ASP_A_23_CA | ASP_A_23_HA | -11.532 | -5.129 | 4.629 | -11.541 | -5.472 | 5.664 | 1.09 | 18.34 | -91.50 |
| 135 | 2jsa-11 | ASP_A_23_CA | ASP_A_23_C | -11.532 | -5.129 | 4.629 | -10.094 | -5.086 | 4.109 | 1.53 | 109.87 | 1.71 |
| 136 | 2jsa-11 | ASP_A_23_C | ASP_A_23_O | -10.094 | -5.086 | 4.109 | -9.868 | -5.031 | 2.901 | 1.23 | 169.10 | 13.68 |
| 137 | 2jsa-11 | ASP_A_23_C | ALA_A_24_N | -10.094 | -5.086 | 4.109 | -9.158 | -5.112 | 5.047 | 1.33 | 44.95 | -1.59 |
| 138 | 2jsa-11 | ALA_A_24_N | ALA_A_24_H | -9.158 | -5.112 | 5.047 | -9.416 | -5.156 | 6.023 | 1.01 | 15.01 | -170.32 |
| 139 | 2jsa-11 | ALA_A_24_N | ALA_A_24_CA | -9.158 | -5.112 | 5.047 | -7.748 | -5.077 | 4.698 | 1.45 | 103.90 | 1.42 |
| 140 | 2jsa-11 | ALA_A_24_CA | ALA_A_24_HA | -7.748 | -5.077 | 4.698 | -7.521 | -4.11 | 4.25 | 1.09 | 114.28 | 76.79 |
| 141 | 2jsa-11 | ALA_A_24_CA | ALA_A_24_C | -7.748 | -5.077 | 4.698 | -7.455 | -6.169 | 3.667 | 1.53 | 132.36 | -74.98 |
| 142 | 2jsa-11 | ALA_A_24_C | ALA_A_24_O | -7.455 | -6.169 | 3.667 | -6.432 | -6.124 | 2.984 | 1.23 | 123.70 | 2.52 |
| 143 | 2jsa-11 | ALA_A_24_C | ASN_A_25_N | -7.455 | -6.169 | 3.667 | -8.371 | -7.124 | 3.587 | 1.33 | 93.46 | -133.81 |
| 144 | 2jsa-11 | ASN_A_25_N | ASN_A_25_H | -8.371 | -7.124 | 3.587 | -9.188 | -7.096 | 4.18 | 1.01 | 54.04 | 178.04 |
| 145 | 2jsa-11 | ASN_A_25_N | ASN_A_25_CA | -8.371 | -7.124 | 3.587 | -8.223 | -8.225 | 2.651 | 1.45 | 130.12 | -82.34 |
| 146 | 2jsa-11 | ASN_A_25_CA | ASN_A_25_HA | -8.223 | -8.225 | 2.651 | -7.213 | -8.627 | 2.731 | 1.09 | 85.79 | -21.70 |
| 147 | 2jsa-11 | ASN_A_25_CA | ASN_A_25_C | -8.223 | -8.225 | 2.651 | -8.469 | -7.717 | 1.229 | 1.53 | 158.35 | 115.84 |
| 148 | 2jsa-11 | ASN_A_25_C | ASN_A_25_O | -8.469 | -7.717 | 1.229 | -7.629 | -7.892 | 0.348 | 1.23 | 135.76 | -11.77 |
| 149 | 2jsa-11 | ASN_A_25_C | VAL_A_26_N | -8.469 | -7.717 | 1.229 | -9.628 | -7.098 | 1.049 | 1.33 | 97.80 | 151.89 |
| 150 | 2jsa-11 | VAL_A_26_N | VAL_A_26_H | -9.628 | -7.098 | 1.049 | -10.272 | -6.99 | 1.819 | 1.01 | 40.30 | 170.48 |
| 151 | 2jsa-11 | VAL_A_26_N | VAL_A_26_CA | -9.628 | -7.098 | 1.049 | -9.996 | -6.564 | -0.25 | 1.45 | 153.47 | 124.57 |
| 152 | 2jsa-11 | VAL_A_26_CA | VAL_A_26_HA | -9.996 | -6.564 | -0.25 | -9.765 | -7.313 | -1.008 | 1.09 | 134.04 | -72.86 |
| 153 | 2jsa-11 | VAL_A_26_CA | VAL_A_26_C | -9.996 | -6.564 | -0.25 | -9.156 | -5.318 | -0.542 | 1.53 | 101.00 | 56.01 |
| 154 | 2jsa-11 | VAL_A_26_C | VAL_A_26_O | -9.156 | -5.318 | -0.542 | -9.209 | -4.773 | -1.644 | 1.23 | 153.58 | 95.55 |
| 155 | 2jsa-11 | VAL_A_26_C | ILE_A_27_N | -9.156 | -5.318 | -0.542 | -8.401 | -4.904 | 0.465 | 1.32 | 40.53 | 28.74 |
| 156 | 2jsa-11 | ILE_A_27_N | ILE_A_27_H | -8.401 | -4.904 | 0.465 | -8.406 | -5.402 | 1.344 | 1.01 | 29.54 | -90.58 |
| 157 | 2jsa-11 | ILE_A_27_N | ILE_A_27_CA | -8.401 | -4.904 | 0.465 | -7.552 | -3.733 | 0.331 | 1.45 | 95.29 | 54.06 |
| 158 | 2jsa-11 | ILE_A_27_CA | ILE_A_27_HA | -7.552 | -3.733 | 0.331 | -7.988 | -3.085 | -0.429 | 1.09 | 134.22 | 123.93 |
| 159 | 2jsa-11 | ILE_A_27_CA | ILE_A_27_C | -7.552 | -3.733 | 0.331 | -6.17 | -4.164 | -0.164 | 1.53 | 108.88 | -17.32 |
| 160 | 2jsa-11 | ILE_A_27_C | ILE_A_27_O | -6.17 | -4.164 | -0.164 | -5.823 | -3.939 | -1.322 | 1.23 | 160.35 | 32.96 |
| 161 | 2jsa-11 | ILE_A_27_C | LYS_A_28_N | -6.17 | -4.164 | -0.164 | -5.418 | -4.777 | 0.74 | 1.33 | 47.02 | -39.19 |
| 162 | 2jsa-11 | LYS_A_28_N | LYS_A_28_H | -5.418 | -4.777 | 0.74 | -5.767 | -4.928 | 1.675 | 1.01 | 22.13 | -156.60 |
| 163 | 2jsa-11 | LYS_A_28_N | LYS_A_28_CA | -5.418 | -4.777 | 0.74 | -4.082 | -5.243 | 0.409 | 1.45 | 103.17 | -19.23 |
| 164 | 2jsa-11 | LYS_A_28_CA | LYS_A_28_HA | -4.082 | -5.243 | 0.409 | -3.424 | -4.375 | 0.361 | 1.09 | 92.52 | 52.84 |
| 165 | 2jsa-11 | LYS_A_28_CA | LYS_A_28_C | -4.082 | -5.243 | 0.409 | -4.1 | -5.889 | -0.978 | 1.53 | 155.02 | -91.60 |
| 166 | 2jsa-11 | LYS_A_28_C | LYS_A_28_O | -4.1 | -5.889 | -0.978 | -3.169 | -5.71 | -1.761 | 1.23 | 129.55 | 10.88 |
| 167 | 2jsa-11 | LYS_A_28_C | LYS_A_29_N | -4.1 | -5.889 | -0.978 | -5.169 | -6.626 | -1.237 | 1.32 | 101.28 | -145.42 |
| 168 | 2jsa-11 | LYS_A_29_N | LYS_A_29_H | -5.169 | -6.626 | -1.237 | -5.896 | -6.728 | -0.544 | 1.01 | 46.65 | -172.01 |
| 169 | 2jsa-11 | LYS_A_29_N | LYS_A_29_CA | -5.169 | -6.626 | -1.237 | -5.32 | -7.301 | -2.516 | 1.45 | 151.60 | -102.61 |
| 170 | 2jsa-11 | LYS_A_29_CA | LYS_A_29_HA | -5.32 | -7.301 | -2.516 | -4.381 | -7.805 | -2.745 | 1.09 | 102.13 | -28.22 |
| 171 | 2jsa-11 | LYS_A_29_CA | LYS_A_29_C | -5.32 | -7.301 | -2.516 | -5.569 | -6.261 | -3.61 | 1.53 | 135.65 | 103.46 |
| 172 | 2jsa-11 | LYS_A_29_C | LYS_A_29_O | -5.569 | -6.261 | -3.61 | -4.961 | -6.321 | -4.678 | 1.23 | 150.23 | -5.64 |
| 173 | 2jsa-11 | LYS_A_29_C | ASP_A_30_N | -5.569 | -6.261 | -3.61 | -6.463 | -5.331 | -3.306 | 1.33 | 76.74 | 133.87 |
| 174 | 2jsa-11 | ASP_A_30_N | ASP_A_30_H | -6.463 | -5.331 | -3.306 | -6.924 | -5.343 | -2.408 | 1.01 | 27.18 | -178.51 |
| 175 | 2jsa-11 | ASP_A_30_N | ASP_A_30_CA | -6.463 | -5.331 | -3.306 | -6.799 | -4.28 | -4.25 | 1.45 | 130.55 | 107.73 |
| 176 | 2jsa-11 | ASP_A_30_CA | ASP_A_30_HA | -6.799 | -4.28 | -4.25 | -7.319 | -4.719 | -5.101 | 1.09 | 141.35 | -139.83 |
| 177 | 2jsa-11 | ASP_A_30_CA | ASP_A_30_C | -6.799 | -4.28 | -4.25 | -5.515 | -3.594 | -4.723 | 1.53 | 108.00 | 28.11 |
| 178 | 2jsa-11 | ASP_A_30_C | ASP_A_30_O | -5.515 | -3.594 | -4.723 | -5.303 | -3.426 | -5.923 | 1.23 | 167.30 | 38.40 |
| 179 | 2jsa-11 | ASP_A_30_C | PHE_A_31_N | -5.515 | -3.594 | -4.723 | -4.693 | -3.219 | -3.755 | 1.32 | 43.03 | 24.52 |
| 180 | 2jsa-11 | PHE_A_31_N | PHE_A_31_H | -4.693 | -3.219 | -3.755 | -4.932 | -3.391 | -2.789 | 1.01 | 16.95 | -144.26 |
| 181 | 2jsa-11 | PHE_A_31_N | PHE_A_31_CA | -4.693 | -3.219 | -3.755 | -3.435 | -2.556 | -4.056 | 1.45 | 101.95 | 27.79 |
| 182 | 2jsa-11 | PHE_A_31_CA | PHE_A_31_HA | -3.435 | -2.556 | -4.056 | -3.624 | -1.534 | -4.383 | 1.09 | 107.46 | 100.48 |
| 183 | 2jsa-11 | PHE_A_31_CA | PHE_A_31_C | -3.435 | -2.556 | -4.056 | -2.665 | -3.309 | -5.143 | 1.53 | 135.27 | -44.36 |
| 184 | 2jsa-11 | PHE_A_31_C | PHE_A_31_O | -2.665 | -3.309 | -5.143 | -2.223 | -2.713 | -6.123 | 1.23 | 142.87 | 53.44 |
| 185 | 2jsa-11 | PHE_A_31_C | ASP_A_32_N | -2.665 | -3.309 | -5.143 | -2.528 | -4.611 | -4.931 | 1.33 | 80.80 | -83.99 |
| 186 | 2jsa-11 | ASP_A_32_N | ASP_A_32_H | -2.528 | -4.611 | -4.931 | -2.917 | -5.035 | -4.101 | 1.01 | 34.73 | -132.53 |
| 187 | 2jsa-11 | ASP_A_32_N | ASP_A_32_CA | -2.528 | -4.611 | -4.931 | -1.819 | -5.452 | -5.88 | 1.45 | 130.79 | -49.87 |
| 188 | 2jsa-11 | ASP_A_32_CA | ASP_A_32_HA | -1.819 | -5.452 | -5.88 | -0.772 | -5.147 | -5.914 | 1.09 | 91.79 | 16.24 |
| 189 | 2jsa-11 | ASP_A_32_CA | ASP_A_32_C | -1.819 | -5.452 | -5.88 | -2.451 | -5.297 | -7.265 | 1.53 | 154.83 | 166.22 |
| 190 | 2jsa-11 | ASP_A_32_C | ASP_A_32_O | -2.451 | -5.297 | -7.265 | -1.792 | -5.517 | -8.28 | 1.23 | 145.61 | -18.46 |
| 191 | 2jsa-11 | ASP_A_32_C | ALA_A_33_N | -2.451 | -5.297 | -7.265 | -3.721 | -4.92 | -7.262 | 1.32 | 89.87 | 163.47 |
| 192 | 2jsa-11 | ALA_A_33_N | ALA_A_33_H | -3.721 | -4.92 | -7.262 | -4.201 | -4.758 | -6.388 | 1.01 | 30.10 | 161.35 |
| 193 | 2jsa-11 | ALA_A_33_N | ALA_A_33_CA | -3.721 | -4.92 | -7.262 | -4.449 | -4.733 | -8.505 | 1.45 | 148.84 | 165.59 |
| 194 | 2jsa-11 | ALA_A_33_CA | ALA_A_33_HA | -4.449 | -4.733 | -8.505 | -4.304 | -5.613 | -9.132 | 1.09 | 125.11 | -80.64 |
| 195 | 2jsa-11 | ALA_A_33_CA | ALA_A_33_C | -4.449 | -4.733 | -8.505 | -3.887 | -3.515 | -9.242 | 1.53 | 118.79 | 65.23 |
| 196 | 2jsa-11 | ALA_A_33_C | ALA_A_33_O | -3.887 | -3.515 | -9.242 | -3.169 | -3.66 | -10.23 | 1.23 | 143.45 | -11.42 |
| 197 | 2jsa-11 | ALA_A_33_C | GLU_A_34_N | -3.887 | -3.515 | -9.242 | -4.234 | -2.342 | -8.732 | 1.33 | 67.37 | 106.48 |
| 198 | 2jsa-11 | GLU_A_34_N | GLU_A_34_H | -4.234 | -2.342 | -8.732 | -4.826 | -2.298 | -7.915 | 1.01 | 36.00 | 175.75 |
| 199 | 2jsa-11 | GLU_A_34_N | GLU_A_34_CA | -4.234 | -2.342 | -8.732 | -3.774 | -1.101 | -9.329 | 1.45 | 114.28 | 69.66 |
| 200 | 2jsa-11 | GLU_A_34_CA | GLU_A_34_HA | -3.774 | -1.101 | -9.329 | -4.252 | -0.997 | -10.303 | 1.09 | 153.33 | 167.73 |
| 201 | 2jsa-11 | GLU_A_34_CA | GLU_A_34_C | -3.774 | -1.101 | -9.329 | -2.259 | -1.144 | -9.54 | 1.53 | 97.93 | -1.63 |
| 202 | 2jsa-11 | GLU_A_34_C | GLU_A_34_O | -2.259 | -1.144 | -9.54 | -1.736 | -0.493 | -10.443 | 1.23 | 137.24 | 51.22 |
| 203 | 2jsa-11 | GLU_A_34_C | CYS_A_35_N | -2.259 | -1.144 | -9.54 | -1.597 | -1.916 | -8.693 | 1.32 | 50.21 | -49.39 |
| 204 | 2jsa-11 | CYS_A_35_N | CYS_A_35_H | -1.597 | -1.916 | -8.693 | -2.092 | -2.424 | -7.974 | 1.01 | 44.61 | -134.26 |
| 205 | 2jsa-11 | CYS_A_35_N | CYS_A_35_CA | -1.597 | -1.916 | -8.693 | -0.152 | -2.052 | -8.775 | 1.45 | 93.23 | -5.38 |
| 206 | 2jsa-11 | CYS_A_35_CA | CYS_A_35_HA | -0.152 | -2.052 | -8.775 | 0.278 | -1.063 | -8.936 | 1.09 | 98.49 | 66.50 |
| 207 | 2jsa-11 | CYS_A_35_CA | CYS_A_35_C | -0.152 | -2.052 | -8.775 | 0.179 | -2.929 | -9.984 | 1.53 | 142.21 | -69.32 |
| 208 | 2jsa-11 | CYS_A_35_C | CYS_A_35_O | 0.179 | -2.929 | -9.984 | 1.161 | -2.684 | -10.682 | 1.23 | 124.59 | 14.01 |
| 209 | 2jsa-11 | CYS_A_35_C | LYS_A_36_N | 0.179 | -2.929 | -9.984 | -0.659 | -3.933 | -10.193 | 1.32 | 99.08 | -129.85 |
| 210 | 2jsa-11 | LYS_A_36_N | LYS_A_36_H | -0.659 | -3.933 | -10.193 | -1.446 | -4.075 | -9.577 | 1.01 | 52.39 | -169.77 |
| 211 | 2jsa-11 | LYS_A_36_N | LYS_A_36_CA | -0.659 | -3.933 | -10.193 | -0.467 | -4.848 | -11.306 | 1.45 | 139.97 | -78.15 |
| 212 | 2jsa-11 | LYS_A_36_CA | LYS_A_36_HA | -0.467 | -4.848 | -11.306 | 0.599 | -5.061 | -11.388 | 1.09 | 94.31 | -11.30 |
| 213 | 2jsa-11 | LYS_A_36_CA | LYS_A_36_C | -0.467 | -4.848 | -11.306 | -0.907 | -4.166 | -12.604 | 1.53 | 147.98 | 122.83 |
| 214 | 2jsa-11 | LYS_A_36_C | LYS_A_36_O | -0.907 | -4.166 | -12.604 | -0.818 | -4.754 | -13.68 | 1.23 | 151.07 | -81.39 |
| 215 | 2jsa-11 | LYS_A_36_C | LYS_A_37_N | -0.907 | -4.166 | -12.604 | -1.372 | -2.934 | -12.459 | 1.32 | 83.72 | 110.68 |
| 216 | 2jsa-11 | LYS_A_37_N | LYS_A_37_H | -1.372 | -2.934 | -12.459 | -1.417 | -2.513 | -11.542 | 1.01 | 24.78 | 96.10 |
| 217 | 2jsa-11 | LYS_A_37_N | LYS_A_37_CA | -1.372 | -2.934 | -12.459 | -1.824 | -2.164 | -13.605 | 1.45 | 142.08 | 120.41 |
| 218 | 2jsa-11 | LYS_A_37_CA | LYS_A_37_HA | -1.824 | -2.164 | -13.605 | -1.717 | -2.787 | -14.493 | 1.09 | 144.55 | -80.25 |
| 219 | 2jsa-11 | LYS_A_37_CA | LYS_A_37_C | -1.824 | -2.164 | -13.605 | -0.921 | -0.943 | -13.784 | 1.53 | 96.72 | 53.51 |
| 220 | 2jsa-11 | LYS_A_37_C | LYS_A_37_O | -0.921 | -0.943 | -13.784 | -0.433 | -0.682 | -14.883 | 1.23 | 153.27 | 28.14 |
| 221 | 2jsa-11 | LYS_A_37_C | LEU_A_38_N | -0.921 | -0.943 | -13.784 | -0.726 | -0.225 | -12.687 | 1.33 | 34.15 | 74.81 |

| | | | | | | | | | | | |
|---|---|---|---|---|---|---|---|---|---|---|---|
| 222 | 2jsa-11 | LEU_A_38_N | LEU_A_38_H | -0.726 | -0.225 | -12.687 | -1.159 | -0.498 | -11.816 | 1.01 | 30.44 | -147.77 |
| 223 | 2jsa-11 | LEU_A_38_N | LEU_A_38_CA | -0.726 | -0.225 | -12.687 | 0.11 | 0.963 | -12.709 | 1.45 | 90.87 | 54.87 |
| 224 | 2jsa-11 | LEU_A_38_CA | LEU_A_38_HA | 0.11 | 0.963 | -12.709 | -0.046 | 1.469 | -13.662 | 1.09 | 150.94 | 107.13 |
| 225 | 2jsa-11 | LEU_A_38_CA | LEU_A_38_C | 0.11 | 0.963 | -12.709 | 1.581 | 0.547 | -12.64 | 1.53 | 87.42 | -15.79 |
| 226 | 2jsa-11 | LEU_A_38_C | LEU_A_38_O | 1.581 | 0.547 | -12.64 | 2.4 | 1.017 | -13.428 | 1.23 | 129.84 | 29.85 |
| 227 | 2jsa-11 | LEU_A_38_C | PHE_A_39_N | 1.581 | 0.547 | -12.64 | 1.871 | -0.329 | -11.69 | 1.32 | 44.17 | -71.68 |
| 228 | 2jsa-11 | PHE_A_39_N | PHE_A_39_H | 1.871 | -0.329 | -11.69 | 1.146 | -0.673 | -11.077 | 1.01 | 52.62 | -154.62 |
| 229 | 2jsa-11 | PHE_A_39_N | PHE_A_39_CA | 1.871 | -0.329 | -11.69 | 3.229 | -0.813 | -11.507 | 1.45 | 82.77 | -19.62 |
| 230 | 2jsa-11 | PHE_A_39_CA | PHE_A_39_HA | 3.229 | -0.813 | -11.507 | 3.933 | 0.013 | -11.605 | 1.09 | 95.16 | 49.56 |
| 231 | 2jsa-11 | PHE_A_39_CA | PHE_A_39_C | 3.229 | -0.813 | -11.507 | 3.565 | -1.901 | -12.53 | 1.53 | 131.94 | -72.84 |
| 232 | 2jsa-11 | PHE_A_39_C | PHE_A_39_O | 3.565 | -1.901 | -12.53 | 4.623 | -2.524 | -12.453 | 1.23 | 86.41 | -30.49 |
| 233 | 2jsa-11 | PHE_A_39_C | HIS_A_40_N | 3.565 | -1.901 | -12.53 | 2.645 | -2.095 | -13.464 | 1.33 | 134.81 | -168.09 |
| 234 | 2jsa-11 | HIS_A_40_N | HIS_A_40_H | 2.645 | -2.095 | -13.464 | 1.798 | -1.546 | -13.465 | 1.01 | 90.06 | 147.05 |
| 235 | 2jsa-11 | HIS_A_40_N | HIS_A_40_CA | 2.645 | -2.095 | -13.464 | 2.832 | -3.096 | -14.501 | 1.45 | 135.52 | -79.42 |
| 236 | 2jsa-11 | HIS_A_40_CA | HIS_A_40_HA | 2.832 | -3.096 | -14.501 | 2.967 | -4.062 | -14.015 | 1.09 | 63.51 | -82.04 |
| 237 | 2jsa-11 | HIS_A_40_CA | HIS_A_40_C | 2.832 | -3.096 | -14.501 | 4.105 | -2.787 | -15.291 | 1.53 | 121.09 | 13.64 |
| 238 | 2jsa-11 | HIS_A_40_C | HIS_A_40_O | 4.105 | -2.787 | -15.291 | 4.97 | -3.647 | -15.442 | 1.23 | 97.06 | -44.83 |
| 239 | 2jsa-11 | HIS_A_40_C | THR_A_41_N | 4.105 | -2.787 | -15.291 | 4.178 | -1.555 | -15.774 | 1.33 | 111.37 | 86.61 |
| 240 | 2jsa-11 | THR_A_41_N | THR_A_41_H | 4.178 | -1.555 | -15.774 | 3.428 | -0.898 | -15.61 | 1.01 | 80.66 | 138.78 |
| 241 | 2jsa-11 | THR_A_41_N | THR_A_41_CA | 4.178 | -1.555 | -15.774 | 5.331 | -1.122 | -16.545 | 1.45 | 122.05 | 20.58 |
| 242 | 2jsa-11 | THR_A_41_CA | THR_A_41_HA | 5.331 | -1.122 | -16.545 | 5.387 | -1.716 | -17.457 | 1.09 | 146.81 | -84.61 |
| 243 | 2jsa-11 | THR_A_41_CA | THR_A_41_C | 5.331 | -1.122 | -16.545 | 6.619 | -1.353 | -15.753 | 1.53 | 58.82 | -10.17 |
| 244 | 2jsa-11 | THR_A_41_C | THR_A_41_O | 6.619 | -1.353 | -15.753 | 7.707 | -1.389 | -16.327 | 1.23 | 117.80 | -1.90 |
| 245 | 2jsa-11 | THR_A_41_C | ILE_A_42_N | 6.619 | -1.353 | -15.753 | 6.455 | -1.505 | -14.448 | 1.32 | 9.72 | -137.17 |
| 246 | 2jsa-11 | ILE_A_42_N | ILE_A_42_H | 6.455 | -1.505 | -14.448 | 5.531 | -1.465 | -14.044 | 1.01 | 66.40 | 177.52 |
| 247 | 2jsa-11 | ILE_A_42_N | ILE_A_42_CA | 6.455 | -1.505 | -14.448 | 7.591 | -1.732 | -13.572 | 1.45 | 52.90 | -11.30 |
| 248 | 2jsa-11 | ILE_A_42_CA | ILE_A_42_HA | 7.591 | -1.732 | -13.572 | 8.423 | -1.127 | -13.934 | 1.09 | 109.39 | 36.02 |
| 249 | 2jsa-11 | ILE_A_42_CA | ILE_A_42_C | 7.591 | -1.732 | -13.572 | 8.012 | -3.2 | -13.66 | 1.53 | 93.30 | -74.00 |
| 250 | 2jsa-11 | ILE_A_42_C | ILE_A_42_O | 8.012 | -3.2 | -13.66 | 7.165 | -4.088 | -13.737 | 1.23 | 93.59 | -133.65 |
| 251 | 2jsa-11 | ILE_A_42_C | PRO_A_43_N | 8.012 | -3.2 | -13.66 | 9.355 | -3.416 | -13.644 | 1.36 | 89.33 | -9.14 |
| 252 | 2jsa-11 | PRO_A_43_N | PRO_A_43_CA | 9.355 | -3.416 | -13.644 | 9.897 | -4.762 | -13.721 | 1.45 | 93.04 | -68.07 |
| 253 | 2jsa-11 | PRO_A_43_CA | PRO_A_43_HA | 9.897 | -4.762 | -13.721 | 9.383 | -5.328 | -14.497 | 1.09 | 135.43 | -132.24 |
| 254 | 2jsa-11 | PRO_A_43_CA | PRO_A_43_C | 9.897 | -4.762 | -13.721 | 9.728 | -5.498 | -12.39 | 1.53 | 29.57 | -102.93 |
| 255 | 2jsa-11 | PRO_A_43_C | PRO_A_43_O | 9.728 | -5.498 | -12.39 | 8.984 | -6.473 | -12.306 | 1.23 | 86.08 | -127.35 |
| 256 | 2jsa-11 | PRO_A_43_C | PHE_A_44_N | 9.728 | -5.498 | -12.39 | 10.433 | -5.003 | -11.383 | 1.33 | 40.54 | 35.07 |
| 257 | 2jsa-11 | PHE_A_44_N | PHE_A_44_H | 10.433 | -5.003 | -11.383 | 11.026 | -4.198 | -11.525 | 1.01 | 98.08 | 53.62 |
| 258 | 2jsa-11 | PHE_A_44_N | PHE_A_44_CA | 10.433 | -5.003 | -11.383 | 10.371 | -5.601 | -10.061 | 1.45 | 24.45 | -95.92 |
| 259 | 2jsa-11 | PHE_A_44_CA | PHE_A_44_HA | 10.371 | -5.601 | -10.061 | 10.272 | -6.683 | -10.149 | 1.09 | 94.63 | -95.23 |
| 260 | 2jsa-11 | PHE_A_44_CA | PHE_A_44_C | 10.371 | -5.601 | -10.061 | 9.194 | -5.039 | -9.259 | 1.53 | 58.41 | 154.48 |
| 261 | 2jsa-11 | PHE_A_44_C | PHE_A_44_O | 9.194 | -5.039 | -9.259 | 9.041 | -5.345 | -8.077 | 1.23 | 16.14 | -116.57 |
| 262 | 2jsa-11 | PHE_A_44_C | GLY_A_45_N | 9.194 | -5.039 | -9.259 | 8.394 | -4.228 | -9.934 | 1.32 | 120.65 | 134.61 |
| 263 | 2jsa-11 | GLY_A_45_N | GLY_A_45_H | 8.394 | -4.228 | -9.934 | 8.582 | -4.02 | -10.904 | 1.01 | 163.88 | 47.89 |
| 264 | 2jsa-11 | GLY_A_45_N | GLY_A_45_CA | 8.394 | -4.228 | -9.934 | 7.236 | -3.621 | -9.3 | 1.45 | 64.13 | 152.34 |
| 265 | 2jsa-11 | GLY_A_45_CA | GLY_A_45_C | 7.236 | -3.621 | -9.3 | 6.273 | -4.689 | -8.779 | 1.53 | 70.08 | -132.04 |
| 266 | 2jsa-11 | GLY_A_45_C | GLY_A_45_O | 6.273 | -4.689 | -8.779 | 6.22 | -4.951 | -7.578 | 1.23 | 12.55 | -101.44 |
| 267 | 2jsa-11 | GLY_A_45_C | THR_A_46_N | 6.273 | -4.689 | -8.779 | 5.534 | -5.278 | -9.709 | 1.33 | 134.54 | -141.44 |
| 268 | 2jsa-11 | THR_A_46_N | THR_A_46_H | 5.534 | -5.278 | -9.709 | 5.63 | -5.012 | -10.679 | 1.01 | 163.75 | 70.16 |
| 269 | 2jsa-11 | THR_A_46_N | THR_A_46_CA | 5.534 | -5.278 | -9.709 | 4.576 | -6.313 | -9.359 | 1.45 | 76.06 | -132.79 |
| 270 | 2jsa-11 | THR_A_46_CA | THR_A_46_HA | 4.576 | -6.313 | -9.359 | 3.674 | -5.838 | -8.974 | 1.09 | 69.31 | 152.23 |
| 271 | 2jsa-11 | THR_A_46_CA | THR_A_46_C | 4.576 | -6.313 | -9.359 | 5.141 | -7.213 | -8.258 | 1.53 | 43.98 | -57.88 |
| 272 | 2jsa-11 | THR_A_46_C | THR_A_46_O | 5.141 | -7.213 | -8.258 | 4.4 | -7.693 | -7.402 | 1.23 | 45.89 | -147.07 |
| 273 | 2jsa-11 | THR_A_46_C | ARG_A_47_N | 5.141 | -7.213 | -8.258 | 6.449 | -7.417 | -8.318 | 1.33 | 92.60 | -8.86 |
| 274 | 2jsa-11 | ARG_A_47_N | ARG_A_47_H | 6.449 | -7.417 | -8.318 | 6.997 | -6.992 | -9.052 | 1.01 | 136.63 | 37.80 |
| 275 | 2jsa-11 | ARG_A_47_N | ARG_A_47_CA | 6.449 | -7.417 | -8.318 | 7.122 | -8.251 | -7.338 | 1.45 | 47.56 | -51.10 |
| 276 | 2jsa-11 | ARG_A_47_CA | ARG_A_47_HA | 7.122 | -8.251 | -7.338 | 6.62 | -9.218 | -7.307 | 1.09 | 88.37 | -117.44 |
| 277 | 2jsa-11 | ARG_A_47_CA | ARG_A_47_C | 7.122 | -8.251 | -7.338 | 7.049 | -7.607 | -5.953 | 1.53 | 25.08 | 96.47 |
| 278 | 2jsa-11 | ARG_A_47_C | ARG_A_47_O | 7.049 | -7.607 | -5.953 | 6.663 | -8.256 | -4.981 | 1.23 | 37.84 | -120.74 |
| 279 | 2jsa-11 | ARG_A_47_C | GLU_A_48_N | 7.049 | -7.607 | -5.953 | 7.425 | -6.337 | -5.904 | 1.33 | 87.88 | 73.51 |
| 280 | 2jsa-11 | GLU_A_48_N | GLU_A_48_H | 7.425 | -6.337 | -5.904 | 7.731 | -5.865 | -6.743 | 1.01 | 146.16 | 57.04 |
| 281 | 2jsa-11 | GLU_A_48_N | GLU_A_48_CA | 7.425 | -6.337 | -5.904 | 7.406 | -5.598 | -4.654 | 1.45 | 30.60 | 91.47 |
| 282 | 2jsa-11 | GLU_A_48_CA | GLU_A_48_HA | 7.406 | -5.598 | -4.654 | 7.914 | -6.199 | -3.899 | 1.09 | 46.19 | -49.79 |
| 283 | 2jsa-11 | GLU_A_48_CA | GLU_A_48_C | 7.406 | -5.598 | -4.654 | 5.965 | -5.367 | -4.195 | 1.53 | 72.54 | 170.89 |
| 284 | 2jsa-11 | GLU_A_48_C | GLU_A_48_O | 5.965 | -5.367 | -4.195 | 5.707 | -5.21 | -3.002 | 1.23 | 14.21 | 148.68 |
| 285 | 2jsa-11 | GLU_A_48_C | CYS_A_49_N | 5.965 | -5.367 | -4.195 | 5.064 | -5.353 | -5.166 | 1.32 | 137.14 | 179.11 |
| 286 | 2jsa-11 | CYS_A_49_N | CYS_A_49_H | 5.064 | -5.353 | -5.166 | 5.348 | -5.49 | -6.125 | 1.01 | 161.80 | -25.75 |
| 287 | 2jsa-11 | CYS_A_49_N | CYS_A_49_CA | 5.064 | -5.353 | -5.166 | 3.656 | -5.143 | -4.877 | 1.45 | 78.52 | 171.52 |
| 288 | 2jsa-11 | CYS_A_49_CA | CYS_A_49_HA | 3.656 | -5.143 | -4.877 | 3.537 | -4.141 | -4.463 | 1.09 | 67.69 | 96.77 |
| 289 | 2jsa-11 | CYS_A_49_CA | CYS_A_49_C | 3.656 | -5.143 | -4.877 | 3.226 | -6.162 | -3.819 | 1.53 | 46.27 | -112.88 |
| 290 | 2jsa-11 | CYS_A_49_C | CYS_A_49_O | 3.226 | -6.162 | -3.819 | 2.946 | -5.797 | -2.678 | 1.23 | 21.96 | 127.49 |
| 291 | 2jsa-11 | CYS_A_49_C | ASP_A_50_N | 3.226 | -6.162 | -3.819 | 3.186 | -7.42 | -4.236 | 1.33 | 108.33 | -91.82 |
| 292 | 2jsa-11 | ASP_A_50_N | ASP_A_50_H | 3.186 | -7.42 | -4.236 | 3.429 | -7.648 | -5.19 | 1.01 | 160.75 | -43.18 |
| 293 | 2jsa-11 | ASP_A_50_N | ASP_A_50_CA | 3.186 | -7.42 | -4.236 | 2.795 | -8.493 | -3.339 | 1.45 | 51.85 | -110.02 |
| 294 | 2jsa-11 | ASP_A_50_CA | ASP_A_50_HA | 2.795 | -8.493 | -3.339 | 1.756 | -8.348 | -3.043 | 1.09 | 74.24 | 172.06 |
| 295 | 2jsa-11 | ASP_A_50_CA | ASP_A_50_C | 2.795 | -8.493 | -3.339 | 3.696 | -8.473 | -2.102 | 1.53 | 36.08 | 1.27 |
| 296 | 2jsa-11 | ASP_A_50_C | ASP_A_50_O | 3.696 | -8.473 | -2.102 | 3.307 | -8.951 | -1.038 | 1.23 | 30.08 | -129.14 |
| 297 | 2jsa-11 | ASP_A_50_C | HIS_A_51_N | 3.696 | -8.473 | -2.102 | 4.883 | -7.914 | -2.285 | 1.32 | 97.94 | 25.22 |
| 298 | 2jsa-11 | HIS_A_51_N | HIS_A_51_H | 4.883 | -7.914 | -2.285 | 5.137 | -7.541 | -3.189 | 1.01 | 153.47 | 55.75 |
| 299 | 2jsa-11 | HIS_A_51_N | HIS_A_51_CA | 4.883 | -7.914 | -2.285 | 5.843 | -7.825 | -1.197 | 1.45 | 41.55 | 5.30 |
| 300 | 2jsa-11 | HIS_A_51_CA | HIS_A_51_HA | 5.843 | -7.825 | -1.197 | 5.729 | -8.712 | -0.574 | 1.09 | 55.14 | -97.32 |
| 301 | 2jsa-11 | HIS_A_51_CA | HIS_A_51_C | 5.843 | -7.825 | -1.197 | 5.528 | -6.603 | -0.334 | 1.53 | 55.63 | 104.45 |
| 302 | 2jsa-11 | HIS_A_51_C | HIS_A_51_O | 5.528 | -6.603 | -0.334 | 6.231 | -6.327 | 0.637 | 1.23 | 37.88 | 21.44 |
| 303 | 2jsa-11 | HIS_A_51_C | TYR_A_52_N | 5.528 | -6.603 | -0.334 | 4.472 | -5.902 | -0.719 | 1.32 | 106.90 | 146.42 |
| 304 | 2jsa-11 | TYR_A_52_N | TYR_A_52_H | 4.472 | -5.902 | -0.719 | 3.941 | -6.186 | -1.529 | 1.01 | 143.37 | -151.86 |
| 305 | 2jsa-11 | TYR_A_52_N | TYR_A_52_CA | 4.472 | -5.902 | -0.719 | 4.056 | -4.715 | 0.008 | 1.45 | 59.97 | 109.31 |
| 306 | 2jsa-11 | TYR_A_52_CA | TYR_A_52_HA | 4.056 | -4.715 | 0.008 | 4.806 | -4.447 | 0.753 | 1.09 | 46.91 | 19.66 |
| 307 | 2jsa-11 | TYR_A_52_CA | TYR_A_52_C | 4.056 | -4.715 | 0.008 | 2.703 | -4.934 | 0.688 | 1.53 | 63.61 | -170.81 |
| 308 | 2jsa-11 | TYR_A_52_C | TYR_A_52_O | 2.703 | -4.934 | 0.688 | 2.546 | -4.646 | 1.873 | 1.23 | 15.47 | 118.60 |
| 309 | 2jsa-11 | TYR_A_52_C | VAL_A_53_N | 2.703 | -4.934 | 0.688 | 1.761 | -5.442 | -0.092 | 1.32 | 126.08 | -151.66 |
| 310 | 2jsa-11 | VAL_A_53_N | VAL_A_53_H | 1.761 | -5.442 | -0.092 | 1.962 | -5.655 | -1.059 | 1.01 | 163.15 | -46.66 |
| 311 | 2jsa-11 | VAL_A_53_N | VAL_A_53_CA | 1.761 | -5.442 | -0.092 | 0.426 | -5.704 | 0.42 | 1.45 | 69.38 | -168.90 |
| 312 | 2jsa-11 | VAL_A_53_CA | VAL_A_53_HA | 0.426 | -5.704 | 0.42 | -0.019 | -4.756 | 0.722 | 1.09 | 73.91 | 115.15 |
| 313 | 2jsa-11 | VAL_A_53_CA | VAL_A_53_C | 0.426 | -5.704 | 0.42 | 0.525 | -6.596 | 1.658 | 1.53 | 35.94 | -83.67 |
| 314 | 2jsa-11 | VAL_A_53_C | VAL_A_53_O | 0.525 | -6.596 | 1.658 | -0.413 | -6.672 | 2.45 | 1.23 | 49.92 | -175.37 |
| 315 | 2jsa-11 | VAL_A_53_C | ASN_A_54_N | 0.525 | -6.596 | 1.658 | 1.671 | -7.251 | 1.787 | 1.33 | 84.42 | -29.75 |
| 316 | 2jsa-11 | ASN_A_54_N | ASN_A_54_H | 1.671 | -7.251 | 1.787 | 2.4 | -7.141 | 1.097 | 1.01 | 133.10 | 8.58 |
| 317 | 2jsa-11 | ASN_A_54_N | ASN_A_54_CA | 1.671 | -7.251 | 1.787 | 1.904 | -8.135 | 2.915 | 1.45 | 39.02 | -75.23 |

| | | | | | | | | | | | |
|---|---|---|---|---|---|---|---|---|---|---|---|
| 318 | 2jsa-11 | ASN_A_54_CA | ASN_A_54_HA | 1.904 | -8.135 | 2.915 | 1.174 | -7.919 | 3.695 | 1.09 | 44.30 | 163.52 |
| 319 | 2jsa-11 | ASN_A_54_CA | ASN_A_54_C | 1.904 | -8.135 | 2.915 | 3.319 | -7.907 | 3.451 | 1.53 | 69.50 | 9.15 |
| 320 | 2jsa-11 | ASN_A_54_C | ASN_A_54_O | 3.319 | -7.907 | 3.451 | 4.201 | -8.742 | 3.261 | 1.23 | 98.89 | -43.43 |
| 321 | 2jsa-11 | ASN_A_54_C | SER_A_55_N | 3.319 | -7.907 | 3.451 | 3.49 | -6.772 | 4.113 | 1.33 | 60.03 | 81.43 |
| 322 | 2jsa-11 | SER_A_55_N | SER_A_55_H | 3.49 | -6.772 | 4.113 | 2.718 | -6.132 | 4.234 | 1.01 | 83.12 | 140.34 |
| 323 | 2jsa-11 | SER_A_55_N | SER_A_55_CA | 3.49 | -6.772 | 4.113 | 4.783 | -6.423 | 4.678 | 1.45 | 67.13 | 15.11 |
| 324 | 2jsa-11 | SER_A_55_CA | SER_A_55_HA | 4.783 | -6.423 | 4.678 | 4.937 | -7.022 | 5.575 | 1.09 | 34.59 | -75.58 |
| 325 | 2jsa-11 | SER_A_55_CA | SER_A_55_C | 4.783 | -6.423 | 4.678 | 4.802 | -4.944 | 5.068 | 1.53 | 75.23 | 89.26 |
| 326 | 2jsa-11 | SER_A_55_C | SER_A_55_O | 4.802 | -4.944 | 5.068 | 5.505 | -4.553 | 5.999 | 1.23 | 40.83 | 29.08 |
| 327 | 2jsa-11 | SER_A_55_C | LYS_A_56_N | 4.802 | -4.944 | 5.068 | 4.021 | -4.162 | 4.337 | 1.33 | 123.48 | 134.96 |
| 328 | 2jsa-11 | LYS_A_56_N | LYS_A_56_H | 4.021 | -4.162 | 4.337 | 3.47 | -4.553 | 3.587 | 1.01 | 137.99 | -144.64 |
| 329 | 2jsa-11 | LYS_A_56_N | LYS_A_56_CA | 4.021 | -4.162 | 4.337 | 3.939 | -2.735 | 4.594 | 1.45 | 79.81 | 93.29 |
| 330 | 2jsa-11 | LYS_A_56_CA | LYS_A_56_HA | 3.939 | -2.735 | 4.594 | 4.364 | -2.545 | 5.58 | 1.09 | 25.27 | 24.09 |
| 331 | 2jsa-11 | LYS_A_56_CA | LYS_A_56_C | 3.939 | -2.735 | 4.594 | 2.47 | -2.311 | 4.632 | 1.53 | 88.58 | 163.90 |
| 332 | 2jsa-11 | LYS_A_56_C | LYS_A_56_O | 2.47 | -2.311 | 4.632 | 2.164 | -1.119 | 4.661 | 1.23 | 88.65 | 104.40 |
| 333 | 2jsa-11 | LYS_A_56_C | VAL_A_57_N | 2.47 | -2.311 | 4.632 | 1.599 | -3.309 | 4.629 | 1.32 | 90.13 | -131.11 |
| 334 | 2jsa-11 | VAL_A_57_N | VAL_A_57_H | 1.599 | -3.309 | 4.629 | 1.922 | -4.265 | 4.603 | 1.01 | 91.48 | -71.33 |
| 335 | 2jsa-11 | VAL_A_57_N | VAL_A_57_CA | 1.599 | -3.309 | 4.629 | 0.169 | -3.055 | 4.663 | 1.45 | 88.66 | 169.93 |
| 336 | 2jsa-11 | VAL_A_57_CA | VAL_A_57_HA | 0.169 | -3.055 | 4.663 | -0.006 | -2.03 | 4.338 | 1.09 | 107.36 | 99.69 |
| 337 | 2jsa-11 | VAL_A_57_CA | VAL_A_57_C | 0.169 | -3.055 | 4.663 | -0.335 | -3.189 | 6.101 | 1.53 | 19.93 | -165.11 |
| 338 | 2jsa-11 | VAL_A_57_C | VAL_A_57_O | -0.335 | -3.189 | 6.101 | -1.035 | -2.31 | 6.602 | 1.23 | 65.97 | 128.53 |
| 339 | 2jsa-11 | VAL_A_57_C | ASP_A_58_N | -0.335 | -3.189 | 6.101 | 0.041 | -4.294 | 6.726 | 1.32 | 61.83 | -71.21 |
| 340 | 2jsa-11 | ASP_A_58_N | ASP_A_58_H | 0.041 | -4.294 | 6.726 | 0.618 | -4.975 | 6.252 | 1.01 | 117.97 | -49.73 |
| 341 | 2jsa-11 | ASP_A_58_N | ASP_A_58_CA | 0.041 | -4.294 | 6.726 | -0.363 | -4.554 | 8.098 | 1.45 | 19.30 | -147.24 |
| 342 | 2jsa-11 | ASP_A_58_CA | ASP_A_58_HA | -0.363 | -4.554 | 8.098 | -1.431 | -4.771 | 8.119 | 1.09 | 88.90 | -168.51 |
| 343 | 2jsa-11 | ASP_A_58_CA | ASP_A_58_C | -0.363 | -4.554 | 8.098 | -0.066 | -3.323 | 8.954 | 1.53 | 55.94 | 76.44 |
| 344 | 2jsa-11 | ASP_A_58_C | ASP_A_58_O | -0.066 | -3.323 | 8.954 | -0.889 | -2.919 | 9.775 | 1.23 | 48.16 | 153.85 |
| 345 | 2jsa-11 | ASP_A_58_C | PRO_A_59_N | -0.066 | -3.323 | 8.954 | 1.144 | -2.744 | 8.728 | 1.36 | 99.56 | 25.57 |
| 346 | 2jsa-11 | PRO_A_59_N | PRO_A_59_CA | 1.144 | -2.744 | 8.728 | 1.559 | -1.566 | 9.471 | 1.45 | 59.25 | 70.59 |
| 347 | 2jsa-11 | PRO_A_59_CA | PRO_A_59_HA | 1.559 | -1.566 | 9.471 | 1.337 | -1.695 | 10.53 | 1.09 | 13.63 | -149.84 |
| 348 | 2jsa-11 | PRO_A_59_CA | PRO_A_59_C | 1.559 | -1.566 | 9.471 | 0.83 | -0.317 | 8.971 | 1.53 | 109.07 | 120.27 |
| 349 | 2jsa-11 | PRO_A_59_C | PRO_A_59_O | 0.83 | -0.317 | 8.971 | 0.175 | 0.376 | 9.747 | 1.23 | 50.86 | 133.39 |
| 350 | 2jsa-11 | PRO_A_59_C | ILE_A_60_N | 0.83 | -0.317 | 8.971 | 0.97 | -0.067 | 7.677 | 1.33 | 167.51 | 60.75 |
| 351 | 2jsa-11 | ILE_A_60_N | ILE_A_60_H | 0.97 | -0.067 | 7.677 | 1.525 | -0.681 | 7.097 | 1.01 | 125.02 | -47.89 |
| 352 | 2jsa-11 | ILE_A_60_N | ILE_A_60_CA | 0.97 | -0.067 | 7.677 | 0.333 | 1.086 | 7.065 | 1.45 | 114.92 | 118.92 |
| 353 | 2jsa-11 | ILE_A_60_CA | ILE_A_60_HA | 0.333 | 1.086 | 7.065 | 0.851 | 1.981 | 7.411 | 1.09 | 71.50 | 59.94 |
| 354 | 2jsa-11 | ILE_A_60_CA | ILE_A_60_C | 0.333 | 1.086 | 7.065 | -1.118 | 1.174 | 7.541 | 1.53 | 71.87 | 176.53 |
| 355 | 2jsa-11 | ILE_A_60_C | ILE_A_60_O | -1.118 | 1.174 | 7.541 | -1.52 | 2.171 | 8.14 | 1.23 | 60.87 | 111.96 |
| 356 | 2jsa-11 | ILE_A_60_C | ILE_A_61_N | -1.118 | 1.174 | 7.541 | -1.867 | 0.118 | 7.257 | 1.33 | 102.37 | -125.35 |
| 357 | 2jsa-11 | ILE_A_61_N | ILE_A_61_H | -1.867 | 0.118 | 7.257 | -1.471 | -0.668 | 6.761 | 1.01 | 119.40 | -63.26 |
| 358 | 2jsa-11 | ILE_A_61_N | ILE_A_61_CA | -1.867 | 0.118 | 7.257 | -3.264 | 0.064 | 7.651 | 1.45 | 74.26 | -177.79 |
| 359 | 2jsa-11 | ILE_A_61_CA | ILE_A_61_HA | -3.264 | 0.064 | 7.651 | -3.774 | 0.916 | 7.202 | 1.09 | 114.33 | 120.90 |
| 360 | 2jsa-11 | ILE_A_61_CA | ILE_A_61_C | -3.264 | 0.064 | 7.651 | -3.369 | 0.209 | 9.17 | 1.53 | 6.72 | 125.91 |
| 361 | 2jsa-11 | ILE_A_61_C | ILE_A_61_O | -3.369 | 0.209 | 9.17 | -4.425 | 0.565 | 9.691 | 1.23 | 64.94 | 161.37 |
| 362 | 2jsa-11 | ILE_A_61_C | HIS_A_62_N | -3.369 | 0.209 | 9.17 | -2.26 | -0.074 | 9.837 | 1.32 | 59.77 | -14.32 |
| 363 | 2jsa-11 | HIS_A_62_N | HIS_A_62_H | -2.26 | -0.074 | 9.837 | -1.427 | -0.361 | 9.342 | 1.01 | 119.33 | -19.01 |
| 364 | 2jsa-11 | HIS_A_62_N | HIS_A_62_CA | -2.26 | -0.074 | 9.837 | -2.214 | 0.02 | 11.286 | 1.45 | 4.13 | 63.92 |
| 365 | 2jsa-11 | HIS_A_62_CA | HIS_A_62_HA | -2.214 | 0.02 | 11.286 | -3.051 | -0.551 | 11.687 | 1.09 | 68.41 | -145.70 |
| 366 | 2jsa-11 | HIS_A_62_CA | HIS_A_62_C | -2.214 | 0.02 | 11.286 | -2.393 | 1.48 | 11.711 | 1.53 | 73.88 | 96.99 |
| 367 | 2jsa-11 | HIS_A_62_C | HIS_A_62_O | -2.393 | 1.48 | 11.711 | -2.905 | 1.756 | 12.795 | 1.23 | 28.22 | 151.67 |
| 368 | 2jsa-11 | HIS_A_62_C | GLU_A_63_N | -2.393 | 1.48 | 11.711 | -1.96 | 2.375 | 10.836 | 1.32 | 131.35 | 64.18 |
| 369 | 2jsa-11 | GLU_A_63_N | GLU_A_63_H | -1.96 | 2.375 | 10.836 | -1.548 | 2.075 | 9.964 | 1.01 | 149.70 | -36.06 |
| 370 | 2jsa-11 | GLU_A_63_N | GLU_A_63_CA | -1.96 | 2.375 | 10.836 | -2.066 | 3.798 | 11.106 | 1.45 | 79.29 | 94.26 |
| 371 | 2jsa-11 | GLU_A_63_CA | GLU_A_63_HA | -2.066 | 3.798 | 11.106 | -1.905 | 3.951 | 12.173 | 1.09 | 11.76 | 43.54 |
| 372 | 2jsa-11 | GLU_A_63_CA | GLU_A_63_C | -2.066 | 3.798 | 11.106 | -3.467 | 4.304 | 10.753 | 1.53 | 103.33 | 160.14 |
| 373 | 2jsa-11 | GLU_A_63_C | GLU_A_63_O | -3.467 | 4.304 | 10.753 | -4.174 | 4.832 | 11.609 | 1.23 | 45.87 | 143.25 |
| 374 | 2jsa-11 | GLU_A_63_C | LEU_A_64_N | -3.467 | 4.304 | 10.753 | -3.825 | 4.123 | 9.491 | 1.32 | 162.37 | -153.18 |
| 375 | 2jsa-11 | LEU_A_64_N | LEU_A_64_H | -3.825 | 4.123 | 9.491 | -3.191 | 3.681 | 8.841 | 1.01 | 130.06 | -34.88 |
| 376 | 2jsa-11 | LEU_A_64_N | LEU_A_64_CA | -3.825 | 4.123 | 9.491 | -5.128 | 4.553 | 9.014 | 1.45 | 109.17 | 161.74 |
| 377 | 2jsa-11 | LEU_A_64_CA | LEU_A_64_HA | -5.128 | 4.553 | 9.014 | -5.164 | 5.641 | 9.067 | 1.09 | 87.21 | 91.90 |
| 378 | 2jsa-11 | LEU_A_64_CA | LEU_A_64_C | -5.128 | 4.553 | 9.014 | -6.213 | 4.007 | 9.944 | 1.53 | 52.56 | -153.29 |
| 379 | 2jsa-11 | LEU_A_64_C | LEU_A_64_O | -6.213 | 4.007 | 9.944 | -7.136 | 4.729 | 10.319 | 1.23 | 72.25 | 141.97 |
| 380 | 2jsa-11 | LEU_A_64_C | GLU_A_65_N | -6.213 | 4.007 | 9.944 | -6.067 | 2.737 | 10.289 | 1.32 | 74.90 | -83.44 |
| 381 | 2jsa-11 | GLU_A_65_N | GLU_A_65_H | -6.067 | 2.737 | 10.289 | -5.283 | 2.203 | 9.942 | 1.01 | 110.09 | -34.26 |
| 382 | 2jsa-11 | GLU_A_65_N | GLU_A_65_CA | -6.067 | 2.737 | 10.289 | -7.022 | 2.086 | 11.169 | 1.45 | 52.71 | -145.72 |
| 383 | 2jsa-11 | GLU_A_65_CA | GLU_A_65_HA | -7.022 | 2.086 | 11.169 | -8.019 | 2.241 | 10.758 | 1.09 | 112.16 | 171.16 |
| 384 | 2jsa-11 | GLU_A_65_CA | GLU_A_65_C | -7.022 | 2.086 | 11.169 | -6.975 | 2.714 | 12.563 | 1.53 | 24.31 | 85.72 |
| 385 | 2jsa-11 | GLU_A_65_C | GLU_A_65_O | -6.975 | 2.714 | 12.563 | -7.989 | 2.772 | 13.257 | 1.23 | 55.66 | 176.73 |
| 386 | 2jsa-11 | GLU_A_65_C | GLY_A_66_N | -6.975 | 2.714 | 12.563 | -5.786 | 3.168 | 12.932 | 1.33 | 73.83 | 20.90 |
| 387 | 2jsa-11 | GLY_A_66_N | GLY_A_66_H | -5.786 | 3.168 | 12.932 | -4.994 | 3.086 | 12.31 | 1.01 | 128.00 | -5.91 |
| 388 | 2jsa-11 | GLY_A_66_N | GLY_A_66_CA | -5.786 | 3.168 | 12.932 | -5.593 | 3.791 | 14.232 | 1.45 | 26.64 | 72.79 |
| 389 | 2jsa-11 | GLY_A_66_CA | GLY_A_66_C | -5.593 | 3.791 | 14.232 | -6.264 | 5.164 | 14.287 | 1.53 | 87.94 | 116.05 |
| 390 | 2jsa-11 | GLY_A_66_C | GLY_A_66_O | -6.264 | 5.164 | 14.287 | -6.837 | 5.538 | 15.308 | 1.23 | 33.83 | 146.87 |
| 391 | 2jsa-11 | GLY_A_66_C | GLY_A_67_N | -6.264 | 5.164 | 14.287 | -6.171 | 5.878 | 13.175 | 1.32 | 147.08 | 82.58 |
| 392 | 2jsa-11 | GLY_A_67_N | GLY_A_67_H | -6.171 | 5.878 | 13.175 | -5.684 | 5.507 | 12.372 | 1.01 | 142.68 | -37.30 |
| 393 | 2jsa-11 | GLY_A_67_N | GLY_A_67_CA | -6.171 | 5.878 | 13.175 | -6.762 | 7.203 | 13.083 | 1.45 | 93.63 | 114.04 |
| 394 | 2jsa-11 | GLY_A_67_CA | GLY_A_67_C | -6.762 | 7.203 | 13.083 | -5.909 | 8.125 | 12.211 | 1.53 | 124.77 | 47.23 |
| 395 | 2jsa-11 | GLY_A_67_C | GLY_A_67_O | -5.909 | 8.125 | 12.211 | -6.366 | 9.187 | 11.791 | 1.23 | 109.96 | 113.28 |
| 396 | 2jsa-11 | GLY_A_67_C | THR_A_68_N | -5.909 | 8.125 | 12.211 | -4.683 | 7.686 | 11.963 | 1.33 | 100.78 | -19.70 |
| 397 | 2jsa-11 | THR_A_68_N | THR_A_68_H | -4.683 | 7.686 | 11.963 | -4.373 | 6.802 | 12.341 | 1.01 | 68.03 | -70.68 |
| 398 | 2jsa-11 | THR_A_68_N | THR_A_68_CA | -4.683 | 7.686 | 11.963 | -3.762 | 8.459 | 11.148 | 1.45 | 124.13 | 40.01 |
| 399 | 2jsa-11 | THR_A_68_CA | THR_A_68_HA | -3.762 | 8.459 | 11.148 | -3.65 | 9.449 | 11.59 | 1.09 | 66.08 | 83.55 |
| 400 | 2jsa-11 | THR_A_68_CA | THR_A_68_C | -3.762 | 8.459 | 11.148 | -4.319 | 8.634 | 9.734 | 1.53 | 157.56 | 162.56 |
| 401 | 2jsa-11 | THR_A_68_C | THR_A_68_O | -4.319 | 8.634 | 9.734 | -5.113 | 7.817 | 9.27 | 1.23 | 112.16 | -134.18 |
| 402 | 2jsa-11 | THR_A_68_C | ALA_A_69_N | -4.319 | 8.634 | 9.734 | -3.879 | 9.704 | 9.087 | 1.33 | 119.22 | 67.65 |
| 403 | 2jsa-11 | ALA_A_69_N | ALA_A_69_H | -3.879 | 9.704 | 9.087 | -3.225 | 10.334 | 9.53 | 1.01 | 64.00 | 43.93 |
| 404 | 2jsa-11 | ALA_A_69_N | ALA_A_69_CA | -3.879 | 9.704 | 9.087 | -4.324 | 9.996 | 7.735 | 1.45 | 158.51 | 146.73 |
| 405 | 2jsa-11 | ALA_A_69_CA | ALA_A_69_HA | -4.324 | 9.996 | 7.735 | -5.38 | 9.74 | 7.651 | 1.09 | 94.42 | -166.37 |
| 406 | 2jsa-11 | ALA_A_69_CA | ALA_A_69_C | -4.324 | 9.996 | 7.735 | -3.536 | 9.133 | 6.746 | 1.53 | 130.24 | -47.60 |
| 407 | 2jsa-11 | ALA_A_69_C | ALA_A_69_O | -3.536 | 9.133 | 6.746 | -2.554 | 8.494 | 7.12 | 1.23 | 72.30 | -33.05 |
| 408 | 2jsa-11 | ALA_A_69_C | PRO_A_70_N | -3.536 | 9.133 | 6.746 | -4.009 | 9.143 | 5.471 | 1.36 | 159.64 | 178.79 |
| 409 | 2jsa-11 | PRO_A_70_N | PRO_A_70_CA | -4.009 | 9.143 | 5.471 | -3.361 | 8.37 | 4.426 | 1.45 | 136.01 | -50.03 |
| 410 | 2jsa-11 | PRO_A_70_CA | PRO_A_70_HA | -3.361 | 8.37 | 4.426 | -3.133 | 7.368 | 4.79 | 1.09 | 70.49 | -77.18 |
| 411 | 2jsa-11 | PRO_A_70_CA | PRO_A_70_C | -3.361 | 8.37 | 4.426 | -2.054 | 9.033 | 3.983 | 1.53 | 106.82 | 26.90 |
| 412 | 2jsa-11 | PRO_A_70_C | PRO_A_70_O | -2.054 | 9.033 | 3.983 | -1.228 | 8.407 | 3.322 | 1.23 | 122.53 | -37.16 |
| 413 | 2jsa-11 | PRO_A_70_C | LYS_A_71_N | -2.054 | 9.033 | 3.983 | -1.909 | 10.294 | 4.367 | 1.33 | 73.17 | 83.44 |

| | | | | | | | | | | | |
|---|---|---|---|---|---|---|---|---|---|---|---|
| 414 | 2jsa-11 | LYS_A_71_N | LYS_A_71_H | -1.909 | 10.294 | 4.367 | -2.63 | 10.747 | 4.911 | 1.01 | 57.43 | 147.86 |
| 415 | 2jsa-11 | LYS_A_71_N | LYS_A_71_CA | -1.909 | 10.294 | 4.367 | -0.719 | 11.049 | 4.018 | 1.45 | 103.91 | 32.39 |
| 416 | 2jsa-11 | LYS_A_71_CA | LYS_A_71_HA | -0.719 | 11.049 | 4.018 | -0.13 | 10.449 | 3.324 | 1.09 | 129.54 | -45.53 |
| 417 | 2jsa-11 | LYS_A_71_CA | LYS_A_71_C | -0.719 | 11.049 | 4.018 | 0.126 | 11.271 | 5.274 | 1.53 | 34.82 | 14.72 |
| 418 | 2jsa-11 | LYS_A_71_C | LYS_A_71_O | 0.126 | 11.271 | 5.274 | 1.035 | 12.1 | 5.277 | 1.23 | 89.86 | 42.36 |
| 419 | 2jsa-11 | LYS_A_71_C | ASP_A_72_N | 0.126 | 11.271 | 5.274 | -0.204 | 10.514 | 6.311 | 1.33 | 38.53 | -113.55 |
| 420 | 2jsa-11 | ASP_A_72_N | ASP_A_72_H | -0.204 | 10.514 | 6.311 | -0.964 | 9.853 | 6.238 | 1.01 | 94.15 | -138.99 |
| 421 | 2jsa-11 | ASP_A_72_N | ASP_A_72_CA | -0.204 | 10.514 | 6.311 | 0.514 | 10.617 | 7.57 | 1.45 | 29.95 | 8.16 |
| 422 | 2jsa-11 | ASP_A_72_CA | ASP_A_72_HA | 0.514 | 10.617 | 7.57 | 1.346 | 11.312 | 7.45 | 1.09 | 96.32 | 39.87 |
| 423 | 2jsa-11 | ASP_A_72_CA | ASP_A_72_C | 0.514 | 10.617 | 7.57 | 1.042 | 9.237 | 7.966 | 1.53 | 75.00 | -69.06 |
| 424 | 2jsa-11 | ASP_A_72_C | ASP_A_72_O | 1.042 | 9.237 | 7.966 | 1.689 | 9.089 | 9.0 | 1.23 | 32.70 | -12.88 |
| 425 | 2jsa-11 | ASP_A_72_C | VAL_A_73_N | 1.042 | 9.237 | 7.966 | 0.745 | 8.26 | 7.121 | 1.33 | 129.61 | -106.91 |
| 426 | 2jsa-11 | VAL_A_73_N | VAL_A_73_H | 0.745 | 8.26 | 7.121 | 0.207 | 8.454 | 6.289 | 1.01 | 145.50 | 160.17 |
| 427 | 2jsa-11 | VAL_A_73_N | VAL_A_73_CA | 0.745 | 8.26 | 7.121 | 1.182 | 6.897 | 7.368 | 1.45 | 80.21 | -72.22 |
| 428 | 2jsa-11 | VAL_A_73_CA | VAL_A_73_HA | 1.182 | 6.897 | 7.368 | 1.276 | 6.756 | 8.445 | 1.09 | 8.94 | -56.31 |
| 429 | 2jsa-11 | VAL_A_73_CA | VAL_A_73_C | 1.182 | 6.897 | 7.368 | 2.561 | 6.686 | 6.741 | 1.53 | 114.20 | -8.70 |
| 430 | 2jsa-11 | VAL_A_73_C | VAL_A_73_O | 2.561 | 6.686 | 6.741 | 3.414 | 6.012 | 7.317 | 1.23 | 62.08 | -38.31 |
| 431 | 2jsa-11 | VAL_A_73_C | CYS_A_74_N | 2.561 | 6.686 | 6.741 | 2.737 | 7.273 | 5.566 | 1.33 | 152.46 | 73.31 |
| 432 | 2jsa-11 | CYS_A_74_N | CYS_A_74_H | 2.737 | 7.273 | 5.566 | 1.991 | 7.812 | 5.15 | 1.01 | 114.32 | 144.15 |
| 433 | 2jsa-11 | CYS_A_74_N | CYS_A_74_CA | 2.737 | 7.273 | 5.566 | 3.998 | 7.158 | 4.853 | 1.45 | 119.38 | -5.21 |
| 434 | 2jsa-11 | CYS_A_74_CA | CYS_A_74_HA | 3.998 | 7.158 | 4.853 | 4.305 | 6.112 | 4.866 | 1.09 | 89.32 | -73.64 |
| 435 | 2jsa-11 | CYS_A_74_CA | CYS_A_74_C | 3.998 | 7.158 | 4.853 | 5.049 | 7.983 | 5.599 | 1.53 | 60.82 | 38.13 |
| 436 | 2jsa-11 | CYS_A_74_C | CYS_A_74_O | 5.049 | 7.983 | 5.599 | 6.23 | 7.946 | 5.255 | 1.23 | 106.23 | -1.79 |
| 437 | 2jsa-11 | CYS_A_74_C | THR_A_75_N | 5.049 | 7.983 | 5.599 | 4.582 | 8.706 | 6.605 | 1.32 | 40.55 | 122.86 |
| 438 | 2jsa-11 | THR_A_75_N | THR_A_75_H | 4.582 | 8.706 | 6.605 | 3.598 | 8.687 | 6.831 | 1.01 | 77.07 | -178.89 |
| 439 | 2jsa-11 | THR_A_75_N | THR_A_75_CA | 4.582 | 8.706 | 6.605 | 5.467 | 9.537 | 7.403 | 1.45 | 56.68 | 43.20 |
| 440 | 2jsa-11 | THR_A_75_CA | THR_A_75_HA | 5.467 | 9.537 | 7.403 | 6.471 | 9.484 | 6.982 | 1.09 | 112.72 | -3.02 |
| 441 | 2jsa-11 | THR_A_75_CA | THR_A_75_C | 5.467 | 9.537 | 7.403 | 5.533 | 9.021 | 8.841 | 1.53 | 19.89 | -82.71 |
| 442 | 2jsa-11 | THR_A_75_C | THR_A_75_O | 5.533 | 9.021 | 8.841 | 6.468 | 9.336 | 9.576 | 1.23 | 53.32 | 18.62 |
| 443 | 2jsa-11 | THR_A_75_C | LYS_A_76_N | 5.533 | 9.021 | 8.841 | 4.528 | 8.236 | 9.201 | 1.33 | 74.24 | -142.01 |
| 444 | 2jsa-11 | LYS_A_76_N | LYS_A_76_H | 4.528 | 8.236 | 9.201 | 3.792 | 8.019 | 8.545 | 1.01 | 130.53 | -163.57 |
| 445 | 2jsa-11 | LYS_A_76_N | LYS_A_76_CA | 4.528 | 8.236 | 9.201 | 4.46 | 7.672 | 10.538 | 1.45 | 23.02 | -96.87 |
| 446 | 2jsa-11 | LYS_A_76_CA | LYS_A_76_HA | 4.46 | 7.672 | 10.538 | 4.619 | 8.48 | 11.253 | 1.09 | 49.03 | 78.87 |
| 447 | 2jsa-11 | LYS_A_76_CA | LYS_A_76_C | 4.46 | 7.672 | 10.538 | 5.593 | 6.662 | 10.72 | 1.53 | 83.16 | -41.72 |
| 448 | 2jsa-11 | LYS_A_76_C | LYS_A_76_O | 5.593 | 6.662 | 10.72 | 6.546 | 6.916 | 11.455 | 1.23 | 53.31 | 14.92 |
| 449 | 2jsa-11 | LYS_A_76_C | LEU_A_77_N | 5.593 | 6.662 | 10.72 | 5.452 | 5.534 | 10.037 | 1.33 | 121.00 | -97.13 |
| 450 | 2jsa-11 | LEU_A_77_N | LEU_A_77_H | 5.452 | 5.534 | 10.037 | 4.642 | 5.392 | 9.451 | 1.01 | 125.47 | -170.06 |
| 451 | 2jsa-11 | LEU_A_77_N | LEU_A_77_CA | 5.452 | 5.534 | 10.037 | 6.453 | 4.483 | 10.114 | 1.45 | 86.96 | -46.40 |
| 452 | 2jsa-11 | LEU_A_77_CA | LEU_A_77_HA | 6.453 | 4.483 | 10.114 | 6.908 | 4.522 | 11.104 | 1.09 | 24.76 | 4.90 |
| 453 | 2jsa-11 | LEU_A_77_CA | LEU_A_77_C | 6.453 | 4.483 | 10.114 | 7.553 | 4.756 | 9.085 | 1.53 | 132.24 | 13.94 |
| 454 | 2jsa-11 | LEU_A_77_C | LEU_A_77_O | 7.553 | 4.756 | 9.085 | 8.133 | 3.825 | 8.531 | 1.23 | 116.80 | -58.08 |
| 455 | 2jsa-11 | LEU_A_77_C | ASN_A_78_N | 7.553 | 4.756 | 9.085 | 7.804 | 6.037 | 8.86 | 1.32 | 99.78 | 78.91 |
| 456 | 2jsa-11 | ASN_A_78_N | ASN_A_78_H | 7.804 | 6.037 | 8.86 | 7.286 | 6.751 | 9.352 | 1.01 | 60.85 | 125.96 |
| 457 | 2jsa-11 | ASN_A_78_N | ASN_A_78_CA | 7.804 | 6.037 | 8.86 | 8.823 | 6.444 | 7.909 | 1.45 | 130.92 | 21.77 |
| 458 | 2jsa-11 | ASN_A_78_CA | ASN_A_78_HA | 8.823 | 6.444 | 7.909 | 8.615 | 7.461 | 7.577 | 1.09 | 107.74 | 101.56 |
| 459 | 2jsa-11 | ASN_A_78_CA | ASN_A_78_C | 8.823 | 6.444 | 7.909 | 8.805 | 5.492 | 6.71 | 1.53 | 141.55 | -91.08 |
| 460 | 2jsa-11 | ASN_A_78_C | ASN_A_78_O | 8.805 | 5.492 | 6.71 | 9.841 | 5.24 | 6.097 | 1.23 | 119.90 | -13.67 |
| 461 | 2jsa-11 | ASN_A_78_C | GLU_A_79_N | 8.805 | 5.492 | 6.71 | 7.616 | 4.99 | 6.413 | 1.32 | 102.96 | -157.11 |
| 462 | 2jsa-11 | GLU_A_79_N | GLU_A_79_H | 7.616 | 4.99 | 6.413 | 6.807 | 5.243 | 6.961 | 1.01 | 57.12 | 162.63 |
| 463 | 2jsa-11 | GLU_A_79_N | GLU_A_79_CA | 7.616 | 4.99 | 6.413 | 7.448 | 4.071 | 5.299 | 1.45 | 140.02 | -100.36 |
| 464 | 2jsa-11 | GLU_A_79_CA | GLU_A_79_HA | 7.448 | 4.071 | 5.299 | 7.973 | 3.147 | 5.541 | 1.09 | 77.17 | -60.40 |
| 465 | 2jsa-11 | GLU_A_79_CA | GLU_A_79_C | 7.448 | 4.071 | 5.299 | 8.069 | 4.659 | 4.03 | 1.53 | 146.02 | 43.44 |
| 466 | 2jsa-11 | GLU_A_79_C | GLU_A_79_O | 8.069 | 4.659 | 4.03 | 9.079 | 4.155 | 3.539 | 1.23 | 113.51 | -26.52 |
| 467 | 2jsa-11 | GLU_A_79_C | CYS_A_80_N | 8.069 | 4.659 | 4.03 | 7.44 | 5.713 | 3.535 | 1.32 | 111.96 | 120.83 |
| 468 | 2jsa-11 | CYS_A_80_N | CYS_A_80_H | 7.44 | 5.713 | 3.535 | 6.615 | 6.073 | 3.992 | 1.01 | 63.08 | 156.43 |
| 469 | 2jsa-11 | CYS_A_80_N | CYS_A_80_CA | 7.44 | 5.713 | 3.535 | 7.918 | 6.374 | 2.332 | 1.45 | 145.86 | 54.13 |
| 470 | 2jsa-11 | CYS_A_80_CA | CYS_A_80_HA | 7.918 | 6.374 | 2.332 | 7.801 | 5.687 | 1.494 | 1.09 | 140.25 | -99.67 |
| 471 | 2jsa-11 | CYS_A_80_CA | CYS_A_80_C | 7.918 | 6.374 | 2.332 | 9.407 | 6.676 | 2.51 | 1.53 | 83.32 | 11.47 |
| 472 | 2jsa-11 | CYS_A_80_C | CYS_A_80_O | 9.407 | 6.676 | 2.51 | 9.777 | 7.573 | 3.266 | 1.23 | 52.08 | 67.58 |
| 473 | 2jsa-11 | CYS_A_80_C | PRO_A_81_N | 9.407 | 6.676 | 2.51 | 10.245 | 5.89 | 1.781 | 1.36 | 122.40 | -43.17 |
| 474 | 2jsa-11 | PRO_A_81_N | PRO_A_81_CA | 10.245 | 5.89 | 1.781 | 11.685 | 6.065 | 1.85 | 1.45 | 87.28 | 6.93 |
| 475 | 2jsa-11 | PRO_A_81_CA | PRO_A_81_HA | 11.685 | 6.065 | 1.85 | 11.998 | 6.176 | 2.888 | 1.09 | 17.74 | 19.53 |
| 476 | 2jsa-11 | PRO_A_81_CA | PRO_A_81_C | 11.685 | 6.065 | 1.85 | 12.124 | 7.308 | 1.077 | 1.53 | 120.39 | 70.55 |
| 477 | 2jsa-11 | PRO_A_81_C | PRO_A_81_O | 12.124 | 7.308 | 1.077 | 12.514 | 8.31 | 1.674 | 1.23 | 60.96 | 68.73 |
| 1 | 2jsa-12 | ARG_A_1_N | ARG_A_1_H | 13.725 | 5.005 | -1.373 | 13.659 | 5.752 | -2.049 | 1.01 | 132.03 | 95.05 |
| 2 | 2jsa-12 | ARG_A_1_N | ARG_A_1_CA | 13.725 | 5.005 | -1.373 | 12.511 | 4.553 | -0.716 | 1.45 | 63.11 | -159.58 |
| 3 | 2jsa-12 | ARG_A_1_CA | ARG_A_1_HA | 12.511 | 4.553 | -0.716 | 12.364 | 3.499 | -0.953 | 1.09 | 102.56 | -97.94 |
| 4 | 2jsa-12 | ARG_A_1_CA | ARG_A_1_C | 12.511 | 4.553 | -0.716 | 11.306 | 5.35 | -1.22 | 1.53 | 109.23 | 146.52 |
| 5 | 2jsa-12 | ARG_A_1_C | ARG_A_1_O | 11.306 | 5.35 | -1.22 | 10.262 | 4.775 | -1.527 | 1.23 | 104.44 | -151.16 |
| 6 | 2jsa-12 | ARG_A_1_C | SER_A_2_N | 11.306 | 5.35 | -1.22 | 11.49 | 6.66 | -1.288 | 1.32 | 92.94 | 82.00 |
| 7 | 2jsa-12 | SER_A_2_N | SER_A_2_H | 11.49 | 6.66 | -1.288 | 12.376 | 7.064 | -1.019 | 1.01 | 74.56 | 24.51 |
| 8 | 2jsa-12 | SER_A_2_N | SER_A_2_CA | 11.49 | 6.66 | -1.288 | 10.431 | 7.542 | -1.749 | 1.45 | 108.49 | 140.21 |
| 9 | 2jsa-12 | SER_A_2_CA | SER_A_2_HA | 10.431 | 7.542 | -1.749 | 9.499 | 7.23 | -1.279 | 1.09 | 64.44 | -161.49 |
| 10 | 2jsa-12 | SER_A_2_CA | SER_A_2_C | 10.431 | 7.542 | -1.749 | 10.278 | 7.427 | -3.267 | 1.53 | 172.81 | -143.07 |
| 11 | 2jsa-12 | SER_A_2_C | SER_A_2_O | 10.278 | 7.427 | -3.267 | 10.852 | 8.218 | -4.014 | 1.23 | 127.39 | 54.03 |
| 12 | 2jsa-12 | SER_A_2_C | ALA_A_3_N | 10.278 | 7.427 | -3.267 | 9.502 | 6.435 | -3.678 | 1.32 | 108.07 | -128.03 |
| 13 | 2jsa-12 | ALA_A_3_N | ALA_A_3_H | 9.502 | 6.435 | -3.678 | 9.063 | 5.822 | -3.007 | 1.01 | 48.33 | -125.61 |
| 14 | 2jsa-12 | ALA_A_3_N | ALA_A_3_CA | 9.502 | 6.435 | -3.678 | 9.267 | 6.206 | -5.094 | 1.45 | 166.95 | -135.74 |
| 15 | 2jsa-12 | ALA_A_3_CA | ALA_A_3_HA | 9.267 | 6.206 | -5.094 | 8.916 | 7.134 | -5.545 | 1.09 | 114.44 | 110.72 |
| 16 | 2jsa-12 | ALA_A_3_CA | ALA_A_3_C | 9.267 | 6.206 | -5.094 | 8.178 | 5.145 | -5.261 | 1.53 | 96.27 | -135.75 |
| 17 | 2jsa-12 | ALA_A_3_C | ALA_A_3_O | 8.178 | 5.145 | -5.261 | 7.098 | 5.433 | -5.775 | 1.23 | 114.70 | 165.07 |
| 18 | 2jsa-12 | ALA_A_3_C | LEU_A_4_N | 8.178 | 5.145 | -5.261 | 8.499 | 3.938 | -4.816 | 1.33 | 70.39 | -75.11 |
| 19 | 2jsa-12 | LEU_A_4_N | LEU_A_4_H | 8.499 | 3.938 | -4.816 | 9.406 | 3.772 | -4.403 | 1.01 | 65.87 | -10.37 |
| 20 | 2jsa-12 | LEU_A_4_N | LEU_A_4_CA | 8.499 | 3.938 | -4.816 | 7.561 | 2.832 | -4.91 | 1.45 | 93.71 | -130.30 |
| 21 | 2jsa-12 | LEU_A_4_CA | LEU_A_4_HA | 7.561 | 2.832 | -4.91 | 6.789 | 3.104 | -5.63 | 1.09 | 131.34 | 160.59 |
| 22 | 2jsa-12 | LEU_A_4_CA | LEU_A_4_C | 7.561 | 2.832 | -4.91 | 6.882 | 2.629 | -3.554 | 1.53 | 27.59 | -163.35 |
| 23 | 2jsa-12 | LEU_A_4_C | LEU_A_4_O | 6.882 | 2.629 | -3.554 | 5.661 | 2.735 | -3.446 | 1.23 | 84.96 | 175.04 |
| 24 | 2jsa-12 | LEU_A_4_C | SER_A_5_N | 6.882 | 2.629 | -3.554 | 7.702 | 2.34 | -2.554 | 1.33 | 41.00 | -19.41 |
| 25 | 2jsa-12 | SER_A_5_N | SER_A_5_H | 7.702 | 2.34 | -2.554 | 8.697 | 2.268 | -2.716 | 1.01 | 99.22 | -4.14 |
| 26 | 2jsa-12 | SER_A_5_N | SER_A_5_CA | 7.702 | 2.34 | -2.554 | 7.196 | 2.122 | -1.211 | 1.45 | 22.31 | -156.69 |
| 27 | 2jsa-12 | SER_A_5_CA | SER_A_5_HA | 7.196 | 2.122 | -1.211 | 6.637 | 1.187 | -1.205 | 1.09 | 89.68 | -120.87 |
| 28 | 2jsa-12 | SER_A_5_CA | SER_A_5_C | 7.196 | 2.122 | -1.211 | 6.252 | 3.26 | -0.815 | 1.53 | 75.01 | 129.68 |
| 29 | 2jsa-12 | SER_A_5_C | SER_A_5_O | 6.252 | 3.26 | -0.815 | 5.27 | 3.039 | -0.107 | 1.23 | 54.88 | -167.32 |
| 30 | 2jsa-12 | SER_A_5_C | CYS_A_6_N | 6.252 | 3.26 | -0.815 | 6.584 | 4.452 | -1.288 | 1.32 | 110.92 | 74.44 |
| 31 | 2jsa-12 | CYS_A_6_N | CYS_A_6_H | 6.584 | 4.452 | -1.288 | 7.407 | 4.56 | -1.864 | 1.01 | 124.76 | 7.48 |
| 32 | 2jsa-12 | CYS_A_6_N | CYS_A_6_CA | 6.584 | 4.452 | -1.288 | 5.778 | 5.625 | -0.993 | 1.45 | 78.29 | 124.49 |

| | | | | | | | | | | | | |
|---|---|---|---|---|---|---|---|---|---|---|---|---|
| 33 | 2jsa-12 | CYS_A_6_CA | CYS_A_6_HA | 5.778 | 5.625 | -0.993 | 5.587 | 5.648 | 0.08 | 1.09 | 10.16 | 173.13 |
| 34 | 2jsa-12 | CYS_A_6_CA | CYS_A_6_C | 5.778 | 5.625 | -0.993 | 4.441 | 5.486 | -1.723 | 1.53 | 118.51 | -174.06 |
| 35 | 2jsa-12 | CYS_A_6_C | CYS_A_6_O | 4.441 | 5.486 | -1.723 | 3.423 | 5.171 | -1.108 | 1.23 | 60.01 | -162.81 |
| 36 | 2jsa-12 | CYS_A_6_C | GLN_A_7_N | 4.441 | 5.486 | -1.723 | 4.487 | 5.726 | -3.025 | 1.32 | 169.37 | 79.15 |
| 37 | 2jsa-12 | GLN_A_7_N | GLN_A_7_H | 4.487 | 5.726 | -3.025 | 5.361 | 5.98 | -3.464 | 1.01 | 115.75 | 16.20 |
| 38 | 2jsa-12 | GLN_A_7_N | GLN_A_7_CA | 4.487 | 5.726 | -3.025 | 3.292 | 5.632 | -3.847 | 1.45 | 124.44 | -175.50 |
| 39 | 2jsa-12 | GLN_A_7_CA | GLN_A_7_HA | 3.292 | 5.632 | -3.847 | 2.71 | 6.541 | -3.699 | 1.09 | 82.19 | 122.63 |
| 40 | 2jsa-12 | GLN_A_7_CA | GLN_A_7_C | 3.292 | 5.632 | -3.847 | 2.44 | 4.438 | -3.409 | 1.53 | 73.37 | -125.51 |
| 41 | 2jsa-12 | GLN_A_7_C | GLN_A_7_O | 2.44 | 4.438 | -3.409 | 1.246 | 4.583 | -3.157 | 1.23 | 78.17 | 173.08 |
| 42 | 2jsa-12 | GLN_A_7_C | MET_A_8_N | 2.44 | 4.438 | -3.409 | 3.09 | 3.285 | -3.334 | 1.33 | 86.76 | -60.59 |
| 43 | 2jsa-12 | MET_A_8_N | MET_A_8_H | 3.09 | 3.285 | -3.334 | 4.074 | 3.24 | -3.558 | 1.01 | 102.81 | -2.62 |
| 44 | 2jsa-12 | MET_A_8_N | MET_A_8_CA | 3.09 | 3.285 | -3.334 | 2.407 | 2.068 | -2.932 | 1.45 | 73.93 | -119.30 |
| 45 | 2jsa-12 | MET_A_8_CA | MET_A_8_HA | 2.407 | 2.068 | -2.932 | 1.759 | 1.745 | -3.747 | 1.09 | 138.38 | -153.51 |
| 46 | 2jsa-12 | MET_A_8_CA | MET_A_8_C | 2.407 | 2.068 | -2.932 | 1.556 | 2.303 | -1.683 | 1.53 | 35.25 | 164.56 |
| 47 | 2jsa-12 | MET_A_8_C | MET_A_8_O | 1.556 | 2.303 | -1.683 | 0.437 | 1.798 | -1.586 | 1.23 | 85.48 | -155.71 |
| 48 | 2jsa-12 | MET_A_8_C | CYS_A_9_N | 1.556 | 2.303 | -1.683 | 2.116 | 3.069 | -0.759 | 1.32 | 45.76 | 53.83 |
| 49 | 2jsa-12 | CYS_A_9_N | CYS_A_9_H | 2.116 | 3.069 | -0.759 | 3.04 | 3.451 | -0.906 | 1.01 | 98.36 | 22.46 |
| 50 | 2jsa-12 | CYS_A_9_N | CYS_A_9_CA | 2.116 | 3.069 | -0.759 | 1.421 | 3.378 | 0.479 | 1.45 | 31.57 | 156.03 |
| 51 | 2jsa-12 | CYS_A_9_CA | CYS_A_9_HA | 1.421 | 3.378 | 0.479 | 1.191 | 2.441 | 0.986 | 1.09 | 62.28 | -103.79 |
| 52 | 2jsa-12 | CYS_A_9_CA | CYS_A_9_C | 1.421 | 3.378 | 0.479 | 0.106 | 4.079 | 0.131 | 1.53 | 103.14 | 151.94 |
| 53 | 2jsa-12 | CYS_A_9_C | CYS_A_9_O | 0.106 | 4.079 | 0.131 | -0.894 | 3.911 | 0.827 | 1.23 | 55.54 | -170.46 |
| 54 | 2jsa-12 | CYS_A_9_C | GLU_A_10_N | 0.106 | 4.079 | 0.131 | 0.15 | 4.849 | -0.946 | 1.32 | 144.39 | 86.73 |
| 55 | 2jsa-12 | GLU_A_10_N | GLU_A_10_H | 0.15 | 4.849 | -0.946 | 1.008 | 4.939 | -1.471 | 1.01 | 121.32 | 5.99 |
| 56 | 2jsa-12 | GLU_A_10_N | GLU_A_10_CA | 0.15 | 4.849 | -0.946 | -1.025 | 5.576 | -1.395 | 1.45 | 108.00 | 148.25 |
| 57 | 2jsa-12 | GLU_A_10_CA | GLU_A_10_HA | -1.025 | 5.576 | -1.395 | -1.554 | 5.938 | -0.514 | 1.09 | 36.04 | 145.62 |
| 58 | 2jsa-12 | GLU_A_10_CA | GLU_A_10_C | -1.025 | 5.576 | -1.395 | -1.96 | 4.646 | -2.17 | 1.53 | 120.44 | -135.15 |
| 59 | 2jsa-12 | GLU_A_10_C | GLU_A_10_O | -1.96 | 4.646 | -2.17 | -3.065 | 5.039 | -2.541 | 1.23 | 107.55 | 160.42 |
| 60 | 2jsa-12 | GLU_A_10_C | LEU_A_11_N | -1.96 | 4.646 | -2.17 | -1.483 | 3.43 | -2.392 | 1.32 | 99.65 | -68.58 |
| 61 | 2jsa-12 | LEU_A_11_N | LEU_A_11_H | -1.483 | 3.43 | -2.392 | -0.564 | 3.175 | -2.058 | 1.01 | 70.70 | -15.51 |
| 62 | 2jsa-12 | LEU_A_11_N | LEU_A_11_CA | -1.483 | 3.43 | -2.392 | -2.263 | 2.44 | -3.116 | 1.45 | 119.87 | -128.23 |
| 63 | 2jsa-12 | LEU_A_11_CA | LEU_A_11_HA | -2.263 | 2.44 | -3.116 | -2.958 | 2.969 | -3.768 | 1.09 | 126.74 | 142.72 |
| 64 | 2jsa-12 | LEU_A_11_CA | LEU_A_11_C | -2.263 | 2.44 | -3.116 | -3.09 | 1.622 | -2.122 | 1.53 | 49.49 | -135.31 |
| 65 | 2jsa-12 | LEU_A_11_C | LEU_A_11_O | -3.09 | 1.622 | -2.122 | -4.27 | 1.367 | -2.354 | 1.23 | 100.88 | -167.81 |
| 66 | 2jsa-12 | LEU_A_11_C | VAL_A_12_N | -3.09 | 1.622 | -2.122 | -2.436 | 1.234 | -1.037 | 1.32 | 35.03 | -30.68 |
| 67 | 2jsa-12 | VAL_A_12_N | VAL_A_12_H | -2.436 | 1.234 | -1.037 | -1.464 | 1.482 | -0.912 | 1.01 | 82.90 | 14.31 |
| 68 | 2jsa-12 | VAL_A_12_N | VAL_A_12_CA | -2.436 | 1.234 | -1.037 | -3.095 | 0.449 | -0.008 | 1.45 | 44.89 | -130.01 |
| 69 | 2jsa-12 | VAL_A_12_CA | VAL_A_12_HA | -3.095 | 0.449 | -0.008 | -3.603 | -0.388 | -0.486 | 1.09 | 116.02 | -121.25 |
| 70 | 2jsa-12 | VAL_A_12_CA | VAL_A_12_C | -3.095 | 0.449 | -0.008 | -4.151 | 1.313 | 0.688 | 1.53 | 62.97 | 140.71 |
| 71 | 2jsa-12 | VAL_A_12_C | VAL_A_12_O | -4.151 | 1.313 | 0.688 | -5.302 | 0.901 | 0.824 | 1.23 | 83.65 | -160.31 |
| 72 | 2jsa-12 | VAL_A_12_C | VAL_A_13_N | -4.151 | 1.313 | 0.688 | -3.721 | 2.493 | 1.109 | 1.32 | 71.47 | 69.98 |
| 73 | 2jsa-12 | VAL_A_13_N | VAL_A_13_H | -3.721 | 2.493 | 1.109 | -2.759 | 2.766 | 0.963 | 1.01 | 98.31 | 15.84 |
| 74 | 2jsa-12 | VAL_A_13_N | VAL_A_13_CA | -3.721 | 2.493 | 1.109 | -4.614 | 3.417 | 1.786 | 1.45 | 62.22 | 134.02 |
| 75 | 2jsa-12 | VAL_A_13_CA | VAL_A_13_HA | -4.614 | 3.417 | 1.786 | -5.036 | 2.914 | 2.656 | 1.09 | 37.04 | -130.00 |
| 76 | 2jsa-12 | VAL_A_13_CA | VAL_A_13_C | -4.614 | 3.417 | 1.786 | -5.764 | 3.788 | 0.848 | 1.53 | 127.82 | 162.12 |
| 77 | 2jsa-12 | VAL_A_13_C | VAL_A_13_O | -5.764 | 3.788 | 0.848 | -6.929 | 3.554 | 1.165 | 1.23 | 75.06 | -168.64 |
| 78 | 2jsa-12 | VAL_A_13_C | LYS_A_14_N | -5.764 | 3.788 | 0.848 | -5.397 | 4.36 | -0.29 | 1.33 | 149.15 | 57.32 |
| 79 | 2jsa-12 | LYS_A_14_N | LYS_A_14_H | -5.397 | 4.36 | -0.29 | -4.419 | 4.522 | -0.485 | 1.01 | 101.13 | 9.41 |
| 80 | 2jsa-12 | LYS_A_14_N | LYS_A_14_CA | -5.397 | 4.36 | -0.29 | -6.384 | 4.765 | -1.277 | 1.45 | 132.77 | 157.69 |
| 81 | 2jsa-12 | LYS_A_14_CA | LYS_A_14_HA | -6.384 | 4.765 | -1.277 | -6.906 | 5.641 | -0.893 | 1.09 | 69.37 | 120.79 |
| 82 | 2jsa-12 | LYS_A_14_CA | LYS_A_14_C | -6.384 | 4.765 | -1.277 | -7.415 | 3.648 | -1.45 | 1.53 | 96.49 | -132.71 |
| 83 | 2jsa-12 | LYS_A_14_C | LYS_A_14_O | -7.415 | 3.648 | -1.45 | -8.593 | 3.917 | -1.681 | 1.23 | 100.82 | 167.14 |
| 84 | 2jsa-12 | LYS_A_14_C | LYS_A_15_N | -7.415 | 3.648 | -1.45 | -6.934 | 2.419 | -1.329 | 1.33 | 84.76 | -68.63 |
| 85 | 2jsa-12 | LYS_A_15_N | LYS_A_15_H | -6.934 | 2.419 | -1.329 | -5.953 | 2.273 | -1.137 | 1.01 | 79.04 | -8.47 |
| 86 | 2jsa-12 | LYS_A_15_N | LYS_A_15_CA | -6.934 | 2.419 | -1.329 | -7.8 | 1.26 | -1.469 | 1.45 | 95.53 | -126.77 |
| 87 | 2jsa-12 | LYS_A_15_CA | LYS_A_15_HA | -7.8 | 1.26 | -1.469 | -8.497 | 1.456 | -2.283 | 1.09 | 138.35 | 164.29 |
| 88 | 2jsa-12 | LYS_A_15_CA | LYS_A_15_C | -7.8 | 1.26 | -1.469 | -8.618 | 1.083 | -0.189 | 1.53 | 33.18 | -167.79 |
| 89 | 2jsa-12 | LYS_A_15_C | LYS_A_15_O | -8.618 | 1.083 | -0.189 | -9.807 | 0.773 | -0.245 | 1.23 | 92.61 | -165.39 |
| 90 | 2jsa-12 | LYS_A_15_C | TYR_A_16_N | -8.618 | 1.083 | -0.189 | -7.948 | 1.287 | 0.937 | 1.33 | 31.88 | 16.93 |
| 91 | 2jsa-12 | TYR_A_16_N | TYR_A_16_H | -7.948 | 1.287 | 0.937 | -6.97 | 1.539 | 0.909 | 1.01 | 91.59 | 14.45 |
| 92 | 2jsa-12 | TYR_A_16_N | TYR_A_16_CA | -7.948 | 1.287 | 0.937 | -8.598 | 1.153 | 2.229 | 1.45 | 27.19 | -168.35 |
| 93 | 2jsa-12 | TYR_A_16_CA | TYR_A_16_HA | -8.598 | 1.153 | 2.229 | -9.228 | 0.264 | 2.244 | 1.09 | 89.21 | -125.32 |
| 94 | 2jsa-12 | TYR_A_16_CA | TYR_A_16_C | -8.598 | 1.153 | 2.229 | -9.438 | 2.391 | 2.551 | 1.53 | 77.85 | 124.16 |
| 95 | 2jsa-12 | TYR_A_16_C | TYR_A_16_O | -9.438 | 2.391 | 2.551 | -10.666 | 2.336 | 2.525 | 1.23 | 91.21 | -177.44 |
| 96 | 2jsa-12 | TYR_A_16_C | GLU_A_17_N | -9.438 | 2.391 | 2.551 | -8.74 | 3.478 | 2.849 | 1.33 | 77.01 | 57.29 |
| 97 | 2jsa-12 | GLU_A_17_N | GLU_A_17_H | -8.74 | 3.478 | 2.849 | -7.73 | 3.45 | 2.851 | 1.01 | 89.89 | -1.59 |
| 98 | 2jsa-12 | GLU_A_17_N | GLU_A_17_CA | -8.74 | 3.478 | 2.849 | -9.406 | 4.727 | 3.176 | 1.45 | 76.99 | 118.07 |
| 99 | 2jsa-12 | GLU_A_17_CA | GLU_A_17_HA | -9.406 | 4.727 | 3.176 | -9.71 | 4.683 | 4.222 | 1.09 | 16.37 | -171.76 |
| 100 | 2jsa-12 | GLU_A_17_CA | GLU_A_17_C | -9.406 | 4.727 | 3.176 | -10.661 | 4.901 | 2.317 | 1.53 | 124.14 | 172.11 |
| 101 | 2jsa-12 | GLU_A_17_C | GLU_A_17_O | -10.661 | 4.901 | 2.317 | -11.699 | 5.339 | 2.811 | 1.23 | 66.32 | 157.12 |
| 102 | 2jsa-12 | GLU_A_17_C | GLY_A_18_N | -10.661 | 4.901 | 2.317 | -10.522 | 4.55 | 1.048 | 1.32 | 163.43 | -68.40 |
| 103 | 2jsa-12 | GLY_A_18_N | GLY_A_18_H | -10.522 | 4.55 | 1.048 | -9.637 | 4.199 | 0.711 | 1.01 | 109.49 | -21.63 |
| 104 | 2jsa-12 | GLY_A_18_N | GLY_A_18_CA | -10.522 | 4.55 | 1.048 | -11.632 | 4.661 | 0.116 | 1.45 | 129.88 | 174.29 |
| 105 | 2jsa-12 | GLY_A_18_CA | GLY_A_18_C | -11.632 | 4.661 | 0.116 | -11.986 | 3.298 | -0.482 | 1.53 | 113.01 | -104.56 |
| 106 | 2jsa-12 | GLY_A_18_C | GLY_A_18_O | -11.986 | 3.298 | -0.482 | -12.05 | 3.149 | -1.701 | 1.23 | 172.42 | -113.24 |
| 107 | 2jsa-12 | GLY_A_18_C | SER_A_19_N | -11.986 | 3.298 | -0.482 | -12.205 | 2.338 | 0.405 | 1.33 | 47.99 | -102.85 |
| 108 | 2jsa-12 | SER_A_19_N | SER_A_19_H | -12.205 | 2.338 | 0.405 | -12.134 | 2.533 | 1.394 | 1.01 | 11.85 | 69.99 |
| 109 | 2jsa-12 | SER_A_19_N | SER_A_19_CA | -12.205 | 2.338 | 0.405 | -12.55 | 0.992 | -0.02 | 1.45 | 107.01 | -104.38 |
| 110 | 2jsa-12 | SER_A_19_CA | SER_A_19_HA | -12.55 | 0.992 | -0.02 | -12.139 | 0.833 | -1.017 | 1.09 | 156.15 | -21.15 |
| 111 | 2jsa-12 | SER_A_19_CA | SER_A_19_C | -12.55 | 0.992 | -0.02 | -14.07 | 0.838 | -0.087 | 1.53 | 92.51 | -174.21 |
| 112 | 2jsa-12 | SER_A_19_C | SER_A_19_O | -14.07 | 0.838 | -0.087 | -14.602 | 0.302 | -1.058 | 1.23 | 142.13 | -134.79 |
| 113 | 2jsa-12 | SER_A_19_C | ALA_A_20_N | -14.07 | 0.838 | -0.087 | -14.728 | 1.318 | 0.959 | 1.33 | 37.91 | 143.89 |
| 114 | 2jsa-12 | ALA_A_20_N | ALA_A_20_H | -14.728 | 1.318 | 0.959 | -14.225 | 1.746 | 1.723 | 1.01 | 40.84 | 40.39 |
| 115 | 2jsa-12 | ALA_A_20_N | ALA_A_20_CA | -14.728 | 1.318 | 0.959 | -16.177 | 1.24 | 1.032 | 1.45 | 87.12 | -176.92 |
| 116 | 2jsa-12 | ALA_A_20_CA | ALA_A_20_HA | -16.177 | 1.24 | 1.032 | -16.535 | 2.005 | 1.721 | 1.09 | 50.79 | 115.08 |
| 117 | 2jsa-12 | ALA_A_20_CA | ALA_A_20_C | -16.177 | 1.24 | 1.032 | -16.583 | -0.13 | 1.578 | 1.53 | 69.09 | -106.51 |
| 118 | 2jsa-12 | ALA_A_20_C | ALA_A_20_O | -16.583 | -0.13 | 1.578 | -17.756 | -0.367 | 1.862 | 1.23 | 76.65 | -168.58 |
| 119 | 2jsa-12 | ALA_A_20_C | ASP_A_21_N | -16.583 | -0.13 | 1.578 | -15.59 | -0.998 | 1.709 | 1.33 | 84.33 | -41.16 |
| 120 | 2jsa-12 | ASP_A_21_N | ASP_A_21_H | -15.59 | -0.998 | 1.709 | -14.648 | -0.735 | 1.455 | 1.01 | 104.56 | 15.60 |
| 121 | 2jsa-12 | ASP_A_21_N | ASP_A_21_CA | -15.59 | -0.998 | 1.709 | -15.829 | -2.338 | 2.216 | 1.45 | 69.57 | -100.11 |
| 122 | 2jsa-12 | ASP_A_21_CA | ASP_A_21_HA | -15.829 | -2.338 | 2.216 | -16.902 | -2.485 | 2.34 | 1.09 | 83.47 | -172.20 |
| 123 | 2jsa-12 | ASP_A_21_CA | ASP_A_21_C | -15.829 | -2.338 | 2.216 | -15.122 | -2.503 | 3.563 | 1.53 | 28.32 | -13.14 |
| 124 | 2jsa-12 | ASP_A_21_C | ASP_A_21_O | -15.122 | -2.503 | 3.563 | -15.491 | -3.363 | 4.361 | 1.23 | 49.54 | -113.22 |
| 125 | 2jsa-12 | ASP_A_21_C | LYS_A_22_N | -15.122 | -2.503 | 3.563 | -14.118 | -1.665 | 3.774 | 1.32 | 80.83 | 39.85 |
| 126 | 2jsa-12 | LYS_A_22_N | LYS_A_22_H | -14.118 | -1.665 | 3.774 | -13.869 | -0.982 | 3.073 | 1.01 | 133.96 | 69.97 |
| 127 | 2jsa-12 | LYS_A_22_N | LYS_A_22_CA | -14.118 | -1.665 | 3.774 | -13.355 | -1.708 | 5.01 | 1.45 | 31.73 | -3.23 |
| 128 | 2jsa-12 | LYS_A_22_CA | LYS_A_22_HA | -13.355 | -1.708 | 5.01 | -12.475 | -1.076 | 4.886 | 1.09 | 96.53 | 35.69 |

| | | | | | | | | | | | | |
|---|---|---|---|---|---|---|---|---|---|---|---|---|
| 129 | 2jsa-12 | LYS_A_22_CA | LYS_A_22_C | -13.355 | -1.708 | 5.01 | -12.869 | -3.137 | 5.256 | 1.53 | 80.74 | -71.22 |
| 130 | 2jsa-12 | LYS_A_22_C | LYS_A_22_O | -12.869 | -3.137 | 5.256 | -13.301 | -3.788 | 6.205 | 1.23 | 39.46 | -123.57 |
| 131 | 2jsa-12 | LYS_A_22_C | ASP_A_23_N | -12.869 | -3.137 | 5.256 | -11.976 | -3.582 | 4.384 | 1.33 | 131.15 | -26.49 |
| 132 | 2jsa-12 | ASP_A_23_N | ASP_A_23_H | -11.976 | -3.582 | 4.384 | -11.665 | -2.99 | 3.626 | 1.01 | 138.58 | 62.29 |
| 133 | 2jsa-12 | ASP_A_23_N | ASP_A_23_CA | -11.976 | -3.582 | 4.384 | -11.426 | -4.922 | 4.495 | 1.45 | 85.62 | -67.68 |
| 134 | 2jsa-12 | ASP_A_23_CA | ASP_A_23_HA | -11.426 | -4.922 | 4.495 | -11.445 | -5.23 | 5.541 | 1.09 | 16.44 | -93.53 |
| 135 | 2jsa-12 | ASP_A_23_CA | ASP_A_23_C | -11.426 | -4.922 | 4.495 | -9.985 | -4.922 | 3.978 | 1.53 | 109.74 | 0.00 |
| 136 | 2jsa-12 | ASP_A_23_C | ASP_A_23_O | -9.985 | -4.922 | 3.978 | -9.744 | -4.658 | 2.801 | 1.23 | 163.11 | 47.61 |
| 137 | 2jsa-12 | ASP_A_23_C | ALA_A_24_N | -9.985 | -4.922 | 3.978 | -9.066 | -5.22 | 4.884 | 1.32 | 46.84 | -17.97 |
| 138 | 2jsa-12 | ALA_A_24_N | ALA_A_24_H | -9.066 | -5.22 | 4.884 | -9.337 | -5.425 | 5.835 | 1.01 | 19.66 | -142.89 |
| 139 | 2jsa-12 | ALA_A_24_N | ALA_A_24_CA | -9.066 | -5.22 | 4.884 | -7.656 | -5.258 | 4.536 | 1.45 | 103.86 | -1.54 |
| 140 | 2jsa-12 | ALA_A_24_CA | ALA_A_24_HA | -7.656 | -5.258 | 4.536 | -7.392 | -4.324 | 4.041 | 1.09 | 117.02 | 74.22 |
| 141 | 2jsa-12 | ALA_A_24_CA | ALA_A_24_C | -7.656 | -5.258 | 4.536 | -7.407 | -6.41 | 3.561 | 1.53 | 129.60 | -77.80 |
| 142 | 2jsa-12 | ALA_A_24_C | ALA_A_24_O | -7.407 | -6.41 | 3.561 | -6.315 | -6.537 | 3.009 | 1.23 | 116.66 | -6.63 |
| 143 | 2jsa-12 | ALA_A_24_C | ASN_A_25_N | -7.407 | -6.41 | 3.561 | -8.438 | -7.223 | 3.378 | 1.33 | 97.93 | -141.74 |
| 144 | 2jsa-12 | ASN_A_25_N | ASN_A_25_H | -8.438 | -7.223 | 3.378 | -9.307 | -7.059 | 3.865 | 1.01 | 61.16 | 169.31 |
| 145 | 2jsa-12 | ASN_A_25_N | ASN_A_25_CA | -8.438 | -7.223 | 3.378 | -8.344 | -8.361 | 2.479 | 1.45 | 128.21 | -85.28 |
| 146 | 2jsa-12 | ASN_A_25_CA | ASN_A_25_HA | -8.344 | -8.361 | 2.479 | -7.347 | -8.797 | 2.558 | 1.09 | 85.85 | -23.62 |
| 147 | 2jsa-12 | ASN_A_25_CA | ASN_A_25_C | -8.344 | -8.361 | 2.479 | -8.595 | -7.894 | 1.045 | 1.53 | 159.71 | 118.26 |
| 148 | 2jsa-12 | ASN_A_25_C | ASN_A_25_O | -8.595 | -7.894 | 1.045 | -7.824 | -8.211 | 0.141 | 1.23 | 137.32 | -22.35 |
| 149 | 2jsa-12 | ASN_A_25_C | VAL_A_26_N | -8.595 | -7.894 | 1.045 | -9.678 | -7.148 | 0.88 | 1.33 | 97.15 | 145.44 |
| 150 | 2jsa-12 | VAL_A_26_N | VAL_A_26_H | -9.678 | -7.148 | 0.88 | -10.268 | -6.927 | 1.669 | 1.01 | 38.61 | 159.47 |
| 151 | 2jsa-12 | VAL_A_26_N | VAL_A_26_CA | -9.678 | -7.148 | 0.88 | -10.04 | -6.634 | -0.43 | 1.45 | 154.36 | 125.16 |
| 152 | 2jsa-12 | VAL_A_26_CA | VAL_A_26_HA | -10.04 | -6.634 | -0.43 | -9.819 | -7.4 | -1.173 | 1.09 | 132.98 | -73.91 |
| 153 | 2jsa-12 | VAL_A_26_CA | VAL_A_26_C | -10.04 | -6.634 | -0.43 | -9.186 | -5.406 | -0.746 | 1.53 | 101.93 | 55.18 |
| 154 | 2jsa-12 | VAL_A_26_C | VAL_A_26_O | -9.186 | -5.406 | -0.746 | -9.178 | -4.924 | -1.878 | 1.23 | 156.93 | 89.05 |
| 155 | 2jsa-12 | VAL_A_26_C | ILE_A_27_N | -9.186 | -5.406 | -0.746 | -8.487 | -4.932 | 0.276 | 1.33 | 39.57 | 34.14 |
| 156 | 2jsa-12 | ILE_A_27_N | ILE_A_27_H | -8.487 | -4.932 | 0.276 | -8.541 | -5.378 | 1.181 | 1.01 | 26.40 | -96.90 |
| 157 | 2jsa-12 | ILE_A_27_N | ILE_A_27_CA | -8.487 | -4.932 | 0.276 | -7.63 | -3.768 | 0.121 | 1.45 | 96.12 | 53.64 |
| 158 | 2jsa-12 | ILE_A_27_CA | ILE_A_27_HA | -7.63 | -3.768 | 0.121 | -8.032 | -3.157 | -0.687 | 1.09 | 137.85 | 123.34 |
| 159 | 2jsa-12 | ILE_A_27_CA | ILE_A_27_C | -7.63 | -3.768 | 0.121 | -6.229 | -4.223 | -0.294 | 1.53 | 105.73 | -17.99 |
| 160 | 2jsa-12 | ILE_A_27_C | ILE_A_27_O | -6.229 | -4.223 | -0.294 | -5.838 | -4.067 | -1.45 | 1.23 | 159.99 | 21.75 |
| 161 | 2jsa-12 | ILE_A_27_C | LYS_A_28_N | -6.229 | -4.223 | -0.294 | -5.511 | -4.777 | 0.673 | 1.33 | 43.16 | -37.65 |
| 162 | 2jsa-12 | LYS_A_28_N | LYS_A_28_H | -5.511 | -4.777 | 0.673 | -5.896 | -4.873 | 1.602 | 1.01 | 23.13 | -166.00 |
| 163 | 2jsa-12 | LYS_A_28_N | LYS_A_28_CA | -5.511 | -4.777 | 0.673 | -4.163 | -5.255 | 0.423 | 1.45 | 99.91 | -19.52 |
| 164 | 2jsa-12 | LYS_A_28_CA | LYS_A_28_HA | -4.163 | -5.255 | 0.423 | -3.498 | -4.392 | 0.4 | 1.09 | 91.21 | 52.38 |
| 165 | 2jsa-12 | LYS_A_28_CA | LYS_A_28_C | -4.163 | -5.255 | 0.423 | -4.108 | -5.921 | -0.955 | 1.53 | 154.13 | -85.28 |
| 166 | 2jsa-12 | LYS_A_28_C | LYS_A_28_O | -4.108 | -5.921 | -0.955 | -3.182 | -5.682 | -1.727 | 1.23 | 128.91 | 14.47 |
| 167 | 2jsa-12 | LYS_A_28_C | LYS_A_29_N | -4.108 | -5.921 | -0.955 | -5.115 | -6.741 | -1.219 | 1.33 | 101.49 | -140.84 |
| 168 | 2jsa-12 | LYS_A_29_N | LYS_A_29_H | -5.115 | -6.741 | -1.219 | -5.843 | -6.889 | -0.535 | 1.01 | 47.36 | -168.51 |
| 169 | 2jsa-12 | LYS_A_29_N | LYS_A_29_CA | -5.115 | -6.741 | -1.219 | -5.194 | -7.442 | -2.489 | 1.45 | 150.95 | -96.43 |
| 170 | 2jsa-12 | LYS_A_29_CA | LYS_A_29_HA | -5.194 | -7.442 | -2.489 | -4.243 | -7.948 | -2.655 | 1.09 | 98.76 | -28.02 |
| 171 | 2jsa-12 | LYS_A_29_CA | LYS_A_29_C | -5.194 | -7.442 | -2.489 | -5.382 | -6.426 | -3.617 | 1.53 | 137.51 | 100.48 |
| 172 | 2jsa-12 | LYS_A_29_C | LYS_A_29_O | -5.382 | -6.426 | -3.617 | -4.634 | -6.432 | -4.594 | 1.23 | 142.56 | -0.46 |
| 173 | 2jsa-12 | LYS_A_29_C | ASP_A_30_N | -5.382 | -6.426 | -3.617 | -6.385 | -5.578 | -3.445 | 1.32 | 82.54 | 139.79 |
| 174 | 2jsa-12 | ASP_A_30_N | ASP_A_30_H | -6.385 | -5.578 | -3.445 | -6.96 | -5.631 | -2.616 | 1.01 | 34.86 | -174.73 |
| 175 | 2jsa-12 | ASP_A_30_N | ASP_A_30_CA | -6.385 | -5.578 | -3.445 | -6.681 | -4.558 | -4.437 | 1.45 | 133.05 | 106.18 |
| 176 | 2jsa-12 | ASP_A_30_CA | ASP_A_30_HA | -6.681 | -4.558 | -4.437 | -7.162 | -5.025 | -5.296 | 1.09 | 142.03 | -135.85 |
| 177 | 2jsa-12 | ASP_A_30_CA | ASP_A_30_C | -6.681 | -4.558 | -4.437 | -5.379 | -3.884 | -4.873 | 1.53 | 106.56 | 27.37 |
| 178 | 2jsa-12 | ASP_A_30_C | ASP_A_30_O | -5.379 | -3.884 | -4.873 | -5.064 | -3.843 | -6.062 | 1.23 | 165.04 | 7.42 |
| 179 | 2jsa-12 | ASP_A_30_C | PHE_A_31_N | -5.379 | -3.884 | -4.873 | -4.656 | -3.372 | -3.888 | 1.32 | 41.97 | 35.30 |
| 180 | 2jsa-12 | PHE_A_31_N | PHE_A_31_H | -4.656 | -3.372 | -3.888 | -4.976 | -3.443 | -2.933 | 1.01 | 18.94 | -167.49 |
| 181 | 2jsa-12 | PHE_A_31_N | PHE_A_31_CA | -4.656 | -3.372 | -3.888 | -3.395 | -2.701 | -4.154 | 1.45 | 100.55 | 28.02 |
| 182 | 2jsa-12 | PHE_A_31_CA | PHE_A_31_HA | -3.395 | -2.701 | -4.154 | -3.582 | -1.684 | -4.5 | 1.09 | 108.50 | 100.42 |
| 183 | 2jsa-12 | PHE_A_31_CA | PHE_A_31_C | -3.395 | -2.701 | -4.154 | -2.582 | -3.461 | -5.205 | 1.53 | 133.36 | -43.07 |
| 184 | 2jsa-12 | PHE_A_31_C | PHE_A_31_O | -2.582 | -3.461 | -5.205 | -2.142 | -2.879 | -6.195 | 1.23 | 143.61 | 52.91 |
| 185 | 2jsa-12 | PHE_A_31_C | ASP_A_32_N | -2.582 | -3.461 | -5.205 | -2.406 | -4.75 | -4.953 | 1.33 | 79.04 | -82.22 |
| 186 | 2jsa-12 | ASP_A_32_N | ASP_A_32_H | -2.406 | -4.75 | -4.953 | -2.795 | -5.162 | -4.117 | 1.01 | 34.13 | -133.36 |
| 187 | 2jsa-12 | ASP_A_32_N | ASP_A_32_CA | -2.406 | -4.75 | -4.953 | -1.654 | -5.596 | -5.864 | 1.45 | 128.83 | -48.37 |
| 188 | 2jsa-12 | ASP_A_32_CA | ASP_A_32_HA | -1.654 | -5.596 | -5.864 | -0.604 | -5.301 | -5.839 | 1.09 | 88.69 | 15.69 |
| 189 | 2jsa-12 | ASP_A_32_CA | ASP_A_32_C | -1.654 | -5.596 | -5.864 | -2.207 | -5.43 | -7.28 | 1.53 | 157.82 | 163.29 |
| 190 | 2jsa-12 | ASP_A_32_C | ASP_A_32_O | -2.207 | -5.43 | -7.28 | -1.446 | -5.26 | -8.232 | 1.23 | 140.68 | 12.59 |
| 191 | 2jsa-12 | ASP_A_32_C | ALA_A_33_N | -2.207 | -5.43 | -7.28 | -3.527 | -5.486 | -7.376 | 1.32 | 94.16 | -177.57 |
| 192 | 2jsa-12 | ALA_A_33_N | ALA_A_33_H | -3.527 | -5.486 | -7.376 | -4.09 | -5.631 | -6.551 | 1.01 | 35.17 | -165.56 |
| 193 | 2jsa-12 | ALA_A_33_N | ALA_A_33_CA | -3.527 | -5.486 | -7.376 | -4.192 | -5.343 | -8.661 | 1.45 | 152.11 | 167.86 |
| 194 | 2jsa-12 | ALA_A_33_CA | ALA_A_33_HA | -4.192 | -5.343 | -8.661 | -3.902 | -6.177 | -9.3 | 1.09 | 125.89 | -70.83 |
| 195 | 2jsa-12 | ALA_A_33_CA | ALA_A_33_C | -4.192 | -5.343 | -8.661 | -3.735 | -4.044 | -9.325 | 1.53 | 115.74 | 70.62 |
| 196 | 2jsa-12 | ALA_A_33_C | ALA_A_33_O | -3.735 | -4.044 | -9.325 | -3.104 | -4.071 | -10.381 | 1.23 | 149.12 | -2.45 |
| 197 | 2jsa-12 | ALA_A_33_C | GLU_A_34_N | -3.735 | -4.044 | -9.325 | -4.071 | -2.937 | -8.679 | 1.33 | 60.82 | 106.88 |
| 198 | 2jsa-12 | GLU_A_34_N | GLU_A_34_H | -4.071 | -2.937 | -8.679 | -4.589 | -2.99 | -7.814 | 1.01 | 31.05 | -174.16 |
| 199 | 2jsa-12 | GLU_A_34_N | GLU_A_34_CA | -4.071 | -2.937 | -8.679 | -3.703 | -1.629 | -9.195 | 1.45 | 110.79 | 74.29 |
| 200 | 2jsa-12 | GLU_A_34_CA | GLU_A_34_HA | -3.703 | -1.629 | -9.195 | -4.329 | -1.42 | -10.062 | 1.09 | 142.72 | 161.54 |
| 201 | 2jsa-12 | GLU_A_34_CA | GLU_A_34_C | -3.703 | -1.629 | -9.195 | -2.241 | -1.628 | -9.646 | 1.53 | 107.14 | 0.04 |
| 202 | 2jsa-12 | GLU_A_34_C | GLU_A_34_O | -2.241 | -1.628 | -9.646 | -1.918 | -1.113 | -10.716 | 1.23 | 150.40 | 57.90 |
| 203 | 2jsa-12 | GLU_A_34_C | CYS_A_35_N | -2.241 | -1.628 | -9.646 | -1.396 | -2.21 | -8.808 | 1.32 | 50.76 | -34.56 |
| 204 | 2jsa-12 | CYS_A_35_N | CYS_A_35_H | -1.396 | -2.21 | -8.808 | -1.732 | -2.613 | -7.945 | 1.01 | 31.30 | -129.82 |
| 205 | 2jsa-12 | CYS_A_35_N | CYS_A_35_CA | -1.396 | -2.21 | -8.808 | 0.024 | -2.283 | -9.107 | 1.45 | 101.88 | -2.94 |
| 206 | 2jsa-12 | CYS_A_35_CA | CYS_A_35_HA | 0.024 | -2.283 | -9.107 | 0.413 | -1.267 | -9.177 | 1.09 | 93.68 | 69.05 |
| 207 | 2jsa-12 | CYS_A_35_CA | CYS_A_35_C | 0.024 | -2.283 | -9.107 | 0.194 | -2.964 | -10.466 | 1.53 | 152.68 | -75.98 |
| 208 | 2jsa-12 | CYS_A_35_C | CYS_A_35_O | 0.194 | -2.964 | -10.466 | 1.001 | -2.528 | -11.287 | 1.23 | 131.83 | 28.38 |
| 209 | 2jsa-12 | CYS_A_35_C | LYS_A_36_N | 0.194 | -2.964 | -10.466 | -0.578 | -4.022 | -10.663 | 1.32 | 98.55 | -126.12 |
| 210 | 2jsa-12 | LYS_A_36_N | LYS_A_36_H | -0.578 | -4.022 | -10.663 | -1.218 | -4.325 | -9.943 | 1.01 | 44.52 | -154.67 |
| 211 | 2jsa-12 | LYS_A_36_N | LYS_A_36_CA | -0.578 | -4.022 | -10.663 | -0.523 | -4.768 | -11.909 | 1.45 | 149.02 | -85.78 |
| 212 | 2jsa-12 | LYS_A_36_CA | LYS_A_36_HA | -0.523 | -4.768 | -11.909 | 0.524 | -4.873 | -12.193 | 1.09 | 105.10 | -5.73 |
| 213 | 2jsa-12 | LYS_A_36_CA | LYS_A_36_C | -0.523 | -4.768 | -11.909 | -1.225 | -3.969 | -13.008 | 1.53 | 135.94 | 131.30 |
| 214 | 2jsa-12 | LYS_A_36_C | LYS_A_36_O | -1.225 | -3.969 | -13.008 | -1.295 | -4.411 | -14.154 | 1.23 | 158.67 | -99.00 |
| 215 | 2jsa-12 | LYS_A_36_C | LYS_A_37_N | -1.225 | -3.969 | -13.008 | -1.727 | -2.805 | -12.622 | 1.33 | 73.06 | 113.33 |
| 216 | 2jsa-12 | LYS_A_37_N | LYS_A_37_H | -1.727 | -2.805 | -12.622 | -1.632 | -2.504 | -11.662 | 1.01 | 18.20 | 72.48 |
| 217 | 2jsa-12 | LYS_A_37_N | LYS_A_37_CA | -1.727 | -2.805 | -12.622 | -2.42 | -1.939 | -13.559 | 1.45 | 130.19 | 128.67 |
| 218 | 2jsa-12 | LYS_A_37_CA | LYS_A_37_HA | -2.42 | -1.939 | -13.559 | -2.594 | -2.504 | -14.474 | 1.09 | 147.13 | -107.12 |
| 219 | 2jsa-12 | LYS_A_37_CA | LYS_A_37_C | -2.42 | -1.939 | -13.559 | -1.522 | -0.752 | -13.911 | 1.53 | 103.31 | 52.89 |
| 220 | 2jsa-12 | LYS_A_37_C | LYS_A_37_O | -1.522 | -0.752 | -13.911 | -1.367 | -0.413 | -15.084 | 1.23 | 162.37 | 65.43 |
| 221 | 2jsa-12 | LYS_A_37_C | LEU_A_38_N | -1.522 | -0.752 | -13.911 | -0.955 | -0.153 | -12.874 | 1.33 | 38.50 | 46.57 |
| 222 | 2jsa-12 | LEU_A_38_N | LEU_A_38_H | -0.955 | -0.153 | -12.874 | -1.128 | -0.486 | -11.936 | 1.01 | 21.80 | -117.45 |
| 223 | 2jsa-12 | LEU_A_38_N | LEU_A_38_CA | -0.955 | -0.153 | -12.874 | -0.076 | 0.99 | -13.059 | 1.45 | 97.31 | 52.44 |
| 224 | 2jsa-12 | LEU_A_38_CA | LEU_A_38_HA | -0.076 | 0.99 | -13.059 | -0.241 | 1.384 | -14.062 | 1.09 | 156.93 | 112.72 |

| | | | | | | | | | | | | |
|---|---|---|---|---|---|---|---|---|---|---|---|---|
| 225 | 2jsa-12 | LEU_A_38_CA | LEU_A_38_C | -0.076 | 0.99 | -13.059 | 1.38 | 0.528 | -12.976 | 1.53 | 86.89 | -17.60 |
| 226 | 2jsa-12 | LEU_A_38_C | LEU_A_38_O | 1.38 | 0.528 | -12.976 | 2.193 | 0.867 | -13.834 | 1.23 | 134.25 | 22.63 |
| 227 | 2jsa-12 | LEU_A_38_C | PHE_A_39_N | 1.38 | 0.528 | -12.976 | 1.665 | -0.241 | -11.935 | 1.33 | 38.23 | -69.66 |
| 228 | 2jsa-12 | PHE_A_39_N | PHE_A_39_H | 1.665 | -0.241 | -11.935 | 0.946 | -0.481 | -11.268 | 1.01 | 48.65 | -161.54 |
| 229 | 2jsa-12 | PHE_A_39_N | PHE_A_39_CA | 1.665 | -0.241 | -11.935 | 3.008 | -0.754 | -11.728 | 1.45 | 81.81 | -20.91 |
| 230 | 2jsa-12 | PHE_A_39_CA | PHE_A_39_HA | 3.008 | -0.754 | -11.728 | 3.741 | -0.013 | -12.05 | 1.09 | 107.17 | 45.31 |
| 231 | 2jsa-12 | PHE_A_39_CA | PHE_A_39_C | 3.008 | -0.754 | -11.728 | 3.222 | -2.058 | -12.5 | 1.53 | 120.29 | -80.68 |
| 232 | 2jsa-12 | PHE_A_39_C | PHE_A_39_O | 3.222 | -2.058 | -12.5 | 4.185 | -2.781 | -12.25 | 1.23 | 78.27 | -36.90 |
| 233 | 2jsa-12 | PHE_A_39_C | HIS_A_40_N | 3.222 | -2.058 | -12.5 | 2.306 | -2.321 | -13.421 | 1.33 | 134.02 | -163.98 |
| 234 | 2jsa-12 | HIS_A_40_N | HIS_A_40_H | 2.306 | -2.321 | -13.421 | 1.539 | -1.682 | -13.572 | 1.01 | 98.60 | 140.20 |
| 235 | 2jsa-12 | HIS_A_40_N | HIS_A_40_CA | 2.306 | -2.321 | -13.421 | 2.382 | -3.525 | -14.229 | 1.45 | 123.81 | -86.39 |
| 236 | 2jsa-12 | HIS_A_40_CA | HIS_A_40_HA | 2.382 | -3.525 | -14.229 | 2.472 | -4.377 | -13.555 | 1.09 | 51.81 | -83.97 |
| 237 | 2jsa-12 | HIS_A_40_CA | HIS_A_40_C | 2.382 | -3.525 | -14.229 | 3.639 | -3.48 | -15.1 | 1.53 | 124.70 | 2.05 |
| 238 | 2jsa-12 | HIS_A_40_C | HIS_A_40_O | 3.639 | -3.48 | -15.1 | 4.493 | -4.361 | -15.009 | 1.23 | 85.76 | -45.89 |
| 239 | 2jsa-12 | HIS_A_40_C | THR_A_41_N | 3.639 | -3.48 | -15.1 | 3.713 | -2.446 | -15.924 | 1.32 | 128.48 | 85.91 |
| 240 | 2jsa-12 | THR_A_41_N | THR_A_41_H | 3.713 | -2.446 | -15.924 | 2.973 | -1.759 | -15.945 | 1.01 | 91.19 | 137.13 |
| 241 | 2jsa-12 | THR_A_41_N | THR_A_41_CA | 3.713 | -2.446 | -15.924 | 4.852 | -2.274 | -16.81 | 1.45 | 127.57 | 8.59 |
| 242 | 2jsa-12 | THR_A_41_CA | THR_A_41_HA | 4.852 | -2.274 | -16.81 | 4.78 | -3.003 | -17.617 | 1.09 | 137.77 | -95.64 |
| 243 | 2jsa-12 | THR_A_41_CA | THR_A_41_C | 4.852 | -2.274 | -16.81 | 6.158 | -2.534 | -16.057 | 1.53 | 60.51 | -11.26 |
| 244 | 2jsa-12 | THR_A_41_C | THR_A_41_O | 6.158 | -2.534 | -16.057 | 7.092 | -3.114 | -16.607 | 1.23 | 116.58 | -31.84 |
| 245 | 2jsa-12 | THR_A_41_C | ILE_A_42_N | 6.158 | -2.534 | -16.057 | 6.181 | -2.091 | -14.808 | 1.33 | 19.55 | 87.03 |
| 246 | 2jsa-12 | ILE_A_42_N | ILE_A_42_H | 6.181 | -2.091 | -14.808 | 5.375 | -1.623 | -14.419 | 1.01 | 67.35 | 149.86 |
| 247 | 2jsa-12 | ILE_A_42_N | ILE_A_42_CA | 6.181 | -2.091 | -14.808 | 7.358 | -2.267 | -13.973 | 1.45 | 54.95 | -8.50 |
| 248 | 2jsa-12 | ILE_A_42_CA | ILE_A_42_HA | 7.358 | -2.267 | -13.973 | 8.236 | -2.011 | -14.567 | 1.09 | 123.00 | 16.26 |
| 249 | 2jsa-12 | ILE_A_42_CA | ILE_A_42_C | 7.358 | -2.267 | -13.973 | 7.478 | -3.741 | -13.576 | 1.53 | 74.97 | -85.35 |
| 250 | 2jsa-12 | ILE_A_42_C | ILE_A_42_O | 7.478 | -3.741 | -13.576 | 6.481 | -4.385 | -13.257 | 1.23 | 74.96 | -147.14 |
| 251 | 2jsa-12 | ILE_A_42_C | PRO_A_43_N | 7.478 | -3.741 | -13.576 | 8.742 | -4.243 | -13.608 | 1.36 | 91.35 | -21.66 |
| 252 | 2jsa-12 | PRO_A_43_N | PRO_A_43_CA | 8.742 | -4.243 | -13.608 | 9.006 | -5.627 | -13.256 | 1.45 | 75.97 | -79.20 |
| 253 | 2jsa-12 | PRO_A_43_CA | PRO_A_43_HA | 9.006 | -5.627 | -13.256 | 8.274 | -6.279 | -13.731 | 1.09 | 115.85 | -138.31 |
| 254 | 2jsa-12 | PRO_A_43_CA | PRO_A_43_C | 9.006 | -5.627 | -13.256 | 8.928 | -5.831 | -11.741 | 1.53 | 8.20 | -110.92 |
| 255 | 2jsa-12 | PRO_A_43_C | PRO_A_43_O | 8.928 | -5.831 | -11.741 | 7.958 | -6.396 | -11.237 | 1.23 | 65.82 | -149.78 |
| 256 | 2jsa-12 | PRO_A_43_C | PHE_A_44_N | 8.928 | -5.831 | -11.741 | 9.961 | -5.361 | -11.059 | 1.32 | 59.00 | 24.46 |
| 257 | 2jsa-12 | PHE_A_44_N | PHE_A_44_H | 9.961 | -5.361 | -11.059 | 10.725 | -4.907 | -11.54 | 1.01 | 118.42 | 30.72 |
| 258 | 2jsa-12 | PHE_A_44_N | PHE_A_44_CA | 9.961 | -5.361 | -11.059 | 10.021 | -5.485 | -9.612 | 1.45 | 5.44 | -64.18 |
| 259 | 2jsa-12 | PHE_A_44_CA | PHE_A_44_HA | 10.021 | -5.485 | -9.612 | 10.028 | -6.538 | -9.331 | 1.09 | 75.06 | -89.62 |
| 260 | 2jsa-12 | PHE_A_44_CA | PHE_A_44_C | 10.021 | -5.485 | -9.612 | 8.833 | -4.784 | -8.952 | 1.53 | 64.43 | 149.46 |
| 261 | 2jsa-12 | PHE_A_44_C | PHE_A_44_O | 8.833 | -4.784 | -8.952 | 8.498 | -5.069 | -7.804 | 1.23 | 20.96 | -139.61 |
| 262 | 2jsa-12 | PHE_A_44_C | GLY_A_45_N | 8.833 | -4.784 | -8.952 | 8.227 | -3.878 | -9.706 | 1.33 | 124.67 | 123.78 |
| 263 | 2jsa-12 | GLY_A_45_N | GLY_A_45_H | 8.227 | -3.878 | -9.706 | 8.557 | -3.694 | -10.643 | 1.01 | 158.04 | 29.14 |
| 264 | 2jsa-12 | GLY_A_45_N | GLY_A_45_CA | 8.227 | -3.878 | -9.706 | 7.083 | -3.133 | -9.209 | 1.45 | 70.00 | 146.93 |
| 265 | 2jsa-12 | GLY_A_45_CA | GLY_A_45_C | 7.083 | -3.133 | -9.209 | 5.999 | -4.077 | -8.687 | 1.53 | 70.04 | -138.95 |
| 266 | 2jsa-12 | GLY_A_45_C | GLY_A_45_O | 5.999 | -4.077 | -8.687 | 5.465 | -3.875 | -7.598 | 1.23 | 27.67 | 159.28 |
| 267 | 2jsa-12 | GLY_A_45_C | THR_A_46_N | 5.999 | -4.077 | -8.687 | 5.704 | -5.089 | -9.49 | 1.33 | 127.30 | -106.25 |
| 268 | 2jsa-12 | THR_A_46_N | THR_A_46_H | 5.704 | -5.089 | -9.49 | 6.184 | -5.193 | -10.373 | 1.01 | 150.92 | -12.23 |
| 269 | 2jsa-12 | THR_A_46_N | THR_A_46_CA | 5.704 | -5.089 | -9.49 | 4.693 | -6.066 | -9.124 | 1.45 | 75.41 | -135.98 |
| 270 | 2jsa-12 | THR_A_46_CA | THR_A_46_HA | 4.693 | -6.066 | -9.124 | 3.822 | -5.539 | -8.736 | 1.09 | 69.14 | 148.82 |
| 271 | 2jsa-12 | THR_A_46_CA | THR_A_46_C | 4.693 | -6.066 | -9.124 | 5.216 | -6.987 | -8.02 | 1.53 | 43.81 | -60.41 |
| 272 | 2jsa-12 | THR_A_46_C | THR_A_46_O | 5.216 | -6.987 | -8.02 | 4.478 | -7.343 | -7.102 | 1.23 | 41.75 | -154.25 |
| 273 | 2jsa-12 | THR_A_46_C | ARG_A_47_N | 5.216 | -6.987 | -8.02 | 6.485 | -7.347 | -8.146 | 1.33 | 95.46 | -15.84 |
| 274 | 2jsa-12 | ARG_A_47_N | ARG_A_47_H | 6.485 | -7.347 | -8.146 | 7.033 | -7.017 | -8.928 | 1.01 | 140.72 | 31.06 |
| 275 | 2jsa-12 | ARG_A_47_N | ARG_A_47_CA | 6.485 | -7.347 | -8.146 | 7.115 | -8.22 | -7.17 | 1.45 | 47.81 | -54.18 |
| 276 | 2jsa-12 | ARG_A_47_CA | ARG_A_47_HA | 7.115 | -8.22 | -7.17 | 6.525 | -9.133 | -7.093 | 1.09 | 85.95 | -122.87 |
| 277 | 2jsa-12 | ARG_A_47_CA | ARG_A_47_C | 7.115 | -8.22 | -7.17 | 7.161 | -7.539 | -5.8 | 1.53 | 26.48 | 86.14 |
| 278 | 2jsa-12 | ARG_A_47_C | ARG_A_47_O | 7.161 | -7.539 | -5.8 | 6.848 | -8.158 | -4.785 | 1.23 | 34.35 | -116.82 |
| 279 | 2jsa-12 | ARG_A_47_C | GLU_A_48_N | 7.161 | -7.539 | -5.8 | 7.555 | -6.275 | -5.817 | 1.32 | 90.74 | 72.69 |
| 280 | 2jsa-12 | GLU_A_48_N | GLU_A_48_H | 7.555 | -6.275 | -5.817 | 7.798 | -5.829 | -6.69 | 1.01 | 149.81 | 61.42 |
| 281 | 2jsa-12 | GLU_A_48_N | GLU_A_48_CA | 7.555 | -6.275 | -5.817 | 7.646 | -5.504 | -4.588 | 1.45 | 32.28 | 83.27 |
| 282 | 2jsa-12 | GLU_A_48_CA | GLU_A_48_HA | 7.646 | -5.504 | -4.588 | 8.159 | -6.114 | -3.844 | 1.09 | 46.97 | -49.94 |
| 283 | 2jsa-12 | GLU_A_48_CA | GLU_A_48_C | 7.646 | -5.504 | -4.588 | 6.247 | -5.178 | -4.061 | 1.53 | 69.85 | 166.88 |
| 284 | 2jsa-12 | GLU_A_48_C | GLU_A_48_O | 6.247 | -5.178 | -4.061 | 6.009 | -5.22 | -2.855 | 1.23 | 11.33 | -169.99 |
| 285 | 2jsa-12 | GLU_A_48_C | CYS_A_49_N | 6.247 | -5.178 | -4.061 | 5.358 | -4.862 | -4.99 | 1.32 | 134.56 | 160.43 |
| 286 | 2jsa-12 | CYS_A_49_N | CYS_A_49_H | 5.358 | -4.862 | -4.99 | 5.624 | -4.849 | -5.964 | 1.01 | 164.71 | 2.80 |
| 287 | 2jsa-12 | CYS_A_49_N | CYS_A_49_CA | 5.358 | -4.862 | -4.99 | 3.989 | -4.529 | -4.634 | 1.45 | 75.82 | 166.33 |
| 288 | 2jsa-12 | CYS_A_49_CA | CYS_A_49_HA | 3.989 | -4.529 | -4.634 | 4.009 | -3.679 | -3.951 | 1.09 | 51.22 | 88.65 |
| 289 | 2jsa-12 | CYS_A_49_CA | CYS_A_49_C | 3.989 | -4.529 | -4.634 | 3.378 | -5.727 | -3.904 | 1.53 | 61.51 | -117.02 |
| 290 | 2jsa-12 | CYS_A_49_C | CYS_A_49_O | 3.378 | -5.727 | -3.904 | 2.719 | -5.563 | -2.878 | 1.23 | 33.50 | 166.03 |
| 291 | 2jsa-12 | CYS_A_49_C | ASP_A_50_N | 3.378 | -5.727 | -3.904 | 3.618 | -6.905 | -4.462 | 1.33 | 114.90 | -78.48 |
| 292 | 2jsa-12 | ASP_A_50_N | ASP_A_50_H | 3.618 | -6.905 | -4.462 | 4.167 | -6.965 | -5.308 | 1.01 | 146.86 | -6.24 |
| 293 | 2jsa-12 | ASP_A_50_N | ASP_A_50_CA | 3.618 | -6.905 | -4.462 | 3.101 | -8.129 | -3.876 | 1.45 | 66.20 | -112.90 |
| 294 | 2jsa-12 | ASP_A_50_CA | ASP_A_50_HA | 3.101 | -8.129 | -3.876 | 2.03 | -8.017 | -3.706 | 1.09 | 81.03 | 174.03 |
| 295 | 2jsa-12 | ASP_A_50_CA | ASP_A_50_C | 3.101 | -8.129 | -3.876 | 3.813 | -8.395 | -2.548 | 1.53 | 29.78 | -20.49 |
| 296 | 2jsa-12 | ASP_A_50_C | ASP_A_50_O | 3.813 | -8.395 | -2.548 | 3.286 | -9.099 | -1.687 | 1.23 | 45.61 | -126.82 |
| 297 | 2jsa-12 | ASP_A_50_C | HIS_A_51_N | 3.813 | -8.395 | -2.548 | 4.999 | -7.818 | -2.423 | 1.32 | 84.59 | 25.94 |
| 298 | 2jsa-12 | HIS_A_51_N | HIS_A_51_H | 4.999 | -7.818 | -2.423 | 5.371 | -7.252 | -3.172 | 1.01 | 137.88 | 56.69 |
| 299 | 2jsa-12 | HIS_A_51_N | HIS_A_51_CA | 4.999 | -7.818 | -2.423 | 5.788 | -7.984 | -1.214 | 1.45 | 33.70 | -11.88 |
| 300 | 2jsa-12 | HIS_A_51_CA | HIS_A_51_HA | 5.788 | -7.984 | -1.214 | 5.445 | -8.887 | -0.709 | 1.09 | 62.40 | -110.80 |
| 301 | 2jsa-12 | HIS_A_51_CA | HIS_A_51_C | 5.788 | -7.984 | -1.214 | 5.546 | -6.8 | -0.276 | 1.53 | 52.18 | 101.55 |
| 302 | 2jsa-12 | HIS_A_51_C | HIS_A_51_O | 5.546 | -6.8 | -0.276 | 6.204 | -6.677 | 0.755 | 1.23 | 32.99 | 10.59 |
| 303 | 2jsa-12 | HIS_A_51_C | TYR_A_52_N | 5.546 | -6.8 | -0.276 | 4.6 | -5.96 | -0.669 | 1.32 | 107.26 | 138.40 |
| 304 | 2jsa-12 | TYR_A_52_N | TYR_A_52_H | 4.6 | -5.96 | -0.669 | 4.1 | -6.124 | -1.531 | 1.01 | 148.60 | -161.84 |
| 305 | 2jsa-12 | TYR_A_52_N | TYR_A_52_CA | 4.6 | -5.96 | -0.669 | 4.263 | -4.79 | 0.124 | 1.45 | 56.92 | 106.07 |
| 306 | 2jsa-12 | TYR_A_52_CA | TYR_A_52_HA | 4.263 | -4.79 | 0.124 | 5.064 | -4.569 | 0.83 | 1.09 | 49.65 | 15.42 |
| 307 | 2jsa-12 | TYR_A_52_CA | TYR_A_52_C | 4.263 | -4.79 | 0.124 | 2.949 | -4.999 | 0.878 | 1.53 | 60.46 | -170.96 |
| 308 | 2jsa-12 | TYR_A_52_C | TYR_A_52_O | 2.949 | -4.999 | 0.878 | 2.8 | -4.546 | 2.012 | 1.23 | 22.81 | 108.21 |
| 309 | 2jsa-12 | TYR_A_52_C | VAL_A_53_N | 2.949 | -4.999 | 0.878 | 2.027 | -5.685 | 0.217 | 1.33 | 119.91 | -143.35 |
| 310 | 2jsa-12 | VAL_A_53_N | VAL_A_53_H | 2.027 | -5.685 | 0.217 | 2.218 | -6.027 | -0.714 | 1.01 | 157.18 | -60.82 |
| 311 | 2jsa-12 | VAL_A_53_N | VAL_A_53_CA | 2.027 | -5.685 | 0.217 | 0.73 | -5.959 | 0.811 | 1.45 | 65.86 | -168.07 |
| 312 | 2jsa-12 | VAL_A_53_CA | VAL_A_53_HA | 0.73 | -5.959 | 0.811 | 0.442 | -5.102 | 1.42 | 1.09 | 56.04 | 108.58 |
| 313 | 2jsa-12 | VAL_A_53_CA | VAL_A_53_C | 0.73 | -5.959 | 0.811 | 0.841 | -7.178 | 1.729 | 1.53 | 53.13 | -84.80 |
| 314 | 2jsa-12 | VAL_A_53_C | VAL_A_53_O | 0.841 | -7.178 | 1.729 | 0.055 | -8.117 | 1.618 | 1.23 | 95.18 | -129.93 |
| 315 | 2jsa-12 | VAL_A_53_C | ASN_A_54_N | 0.841 | -7.178 | 1.729 | 1.825 | -7.124 | 2.615 | 1.33 | 48.04 | 3.14 |
| 316 | 2jsa-12 | ASN_A_54_N | ASN_A_54_H | 1.825 | -7.124 | 2.615 | 2.435 | -6.319 | 2.649 | 1.01 | 88.07 | 52.85 |
| 317 | 2jsa-12 | ASN_A_54_N | ASN_A_54_CA | 1.825 | -7.124 | 2.615 | 2.049 | -8.212 | 3.552 | 1.45 | 49.85 | -78.37 |
| 318 | 2jsa-12 | ASN_A_54_CA | ASN_A_54_HA | 2.049 | -8.212 | 3.552 | 1.16 | -8.341 | 4.169 | 1.09 | 55.52 | -171.74 |
| 319 | 2jsa-12 | ASN_A_54_CA | ASN_A_54_C | 2.049 | -8.212 | 3.552 | 3.251 | -7.876 | 4.437 | 1.53 | 54.66 | 15.62 |
| 320 | 2jsa-12 | ASN_A_54_C | ASN_A_54_O | 3.251 | -7.876 | 4.437 | 4.14 | -8.706 | 4.625 | 1.23 | 81.21 | -43.03 |

| | | | | | | | | | | | | |
|---|---|---|---|---|---|---|---|---|---|---|---|---|
| 321 | 2jsa-12 | ASN_A_54_C | SER_A_55_N | 3.251 | -7.876 | 4.437 | 3.239 | -6.658 | 4.959 | 1.33 | 66.80 | 90.56 |
| 322 | 2jsa-12 | SER_A_55_N | SER_A_55_H | 3.239 | -6.658 | 4.959 | 2.475 | -6.027 | 4.763 | 1.01 | 101.19 | 140.45 |
| 323 | 2jsa-12 | SER_A_55_N | SER_A_55_CA | 3.239 | -6.658 | 4.959 | 4.317 | -6.204 | 5.821 | 1.45 | 53.61 | 22.84 |
| 324 | 2jsa-12 | SER_A_55_CA | SER_A_55_HA | 4.317 | -6.204 | 5.821 | 4.173 | -6.642 | 6.809 | 1.09 | 25.02 | -108.20 |
| 325 | 2jsa-12 | SER_A_55_CA | SER_A_55_C | 4.317 | -6.204 | 5.821 | 4.276 | -4.68 | 5.95 | 1.53 | 85.16 | 91.54 |
| 326 | 2jsa-12 | SER_A_55_C | SER_A_55_O | 4.276 | -4.68 | 5.95 | 4.709 | -4.126 | 6.959 | 1.23 | 34.87 | 51.99 |
| 327 | 2jsa-12 | SER_A_55_C | LYS_A_56_N | 4.276 | -4.68 | 5.95 | 3.753 | -4.044 | 4.911 | 1.33 | 141.60 | 129.43 |
| 328 | 2jsa-12 | LYS_A_56_N | LYS_A_56_H | 3.753 | -4.044 | 4.911 | 3.418 | -4.563 | 4.113 | 1.01 | 142.26 | -122.84 |
| 329 | 2jsa-12 | LYS_A_56_N | LYS_A_56_CA | 3.753 | -4.044 | 4.911 | 3.65 | -2.594 | 4.896 | 1.45 | 90.59 | 94.06 |
| 330 | 2jsa-12 | LYS_A_56_CA | LYS_A_56_HA | 3.65 | -2.594 | 4.896 | 4.026 | -2.218 | 5.848 | 1.09 | 29.19 | 45.00 |
| 331 | 2jsa-12 | LYS_A_56_CA | LYS_A_56_C | 3.65 | -2.594 | 4.896 | 2.178 | -2.193 | 4.785 | 1.53 | 94.16 | 164.76 |
| 332 | 2jsa-12 | LYS_A_56_C | LYS_A_56_O | 2.178 | -2.193 | 4.785 | 1.854 | -1.007 | 4.778 | 1.23 | 90.33 | 105.28 |
| 333 | 2jsa-12 | LYS_A_56_C | VAL_A_57_N | 2.178 | -2.193 | 4.785 | 1.327 | -3.205 | 4.702 | 1.32 | 93.59 | -130.06 |
| 334 | 2jsa-12 | VAL_A_57_N | VAL_A_57_H | 1.327 | -3.205 | 4.702 | 1.666 | -4.156 | 4.714 | 1.01 | 89.32 | -70.38 |
| 335 | 2jsa-12 | VAL_A_57_N | VAL_A_57_CA | 1.327 | -3.205 | 4.702 | -0.104 | -2.972 | 4.592 | 1.45 | 94.34 | 170.75 |
| 336 | 2jsa-12 | VAL_A_57_CA | VAL_A_57_HA | -0.104 | -2.972 | 4.592 | -0.262 | -1.942 | 4.273 | 1.09 | 107.02 | 98.72 |
| 337 | 2jsa-12 | VAL_A_57_CA | VAL_A_57_C | -0.104 | -2.972 | 4.592 | -0.751 | -3.148 | 5.966 | 1.53 | 26.01 | -164.78 |
| 338 | 2jsa-12 | VAL_A_57_C | VAL_A_57_O | -0.751 | -3.148 | 5.966 | -1.598 | -2.35 | 6.364 | 1.23 | 71.12 | 136.71 |
| 339 | 2jsa-12 | VAL_A_57_C | ASP_A_58_N | -0.751 | -3.148 | 5.966 | -0.327 | -4.198 | 6.655 | 1.33 | 58.68 | -68.01 |
| 340 | 2jsa-12 | ASP_A_58_N | ASP_A_58_H | -0.327 | -4.198 | 6.655 | 0.373 | -4.813 | 6.265 | 1.01 | 112.71 | -41.30 |
| 341 | 2jsa-12 | ASP_A_58_N | ASP_A_58_CA | -0.327 | -4.198 | 6.655 | -0.855 | -4.489 | 7.977 | 1.45 | 24.51 | -151.14 |
| 342 | 2jsa-12 | ASP_A_58_CA | ASP_A_58_HA | -0.855 | -4.489 | 7.977 | -1.924 | -4.687 | 7.898 | 1.09 | 94.16 | -169.51 |
| 343 | 2jsa-12 | ASP_A_58_CA | ASP_A_58_C | -0.855 | -4.489 | 7.977 | -0.616 | -3.287 | 8.893 | 1.53 | 53.22 | 78.75 |
| 344 | 2jsa-12 | ASP_A_58_C | ASP_A_58_O | -0.616 | -3.287 | 8.893 | -1.501 | -2.892 | 9.649 | 1.23 | 52.04 | 155.95 |
| 345 | 2jsa-12 | ASP_A_58_C | PRO_A_59_N | -0.616 | -3.287 | 8.893 | 0.619 | -2.725 | 8.793 | 1.36 | 94.22 | 24.47 |
| 346 | 2jsa-12 | PRO_A_59_N | PRO_A_59_CA | 0.619 | -2.725 | 8.793 | 0.986 | -1.577 | 9.603 | 1.45 | 56.10 | 72.27 |
| 347 | 2jsa-12 | PRO_A_59_CA | PRO_A_59_HA | 0.986 | -1.577 | 9.603 | 0.669 | -1.731 | 10.634 | 1.09 | 18.87 | -154.09 |
| 348 | 2jsa-12 | PRO_A_59_CA | PRO_A_59_C | 0.986 | -1.577 | 9.603 | 0.325 | -0.3 | 9.078 | 1.53 | 110.06 | 117.37 |
| 349 | 2jsa-12 | PRO_A_59_C | PRO_A_59_O | 0.325 | -0.3 | 9.078 | -0.303 | 0.436 | 9.838 | 1.23 | 51.85 | 130.47 |
| 350 | 2jsa-12 | PRO_A_59_C | ILE_A_60_N | 0.325 | -0.3 | 9.078 | 0.488 | -0.078 | 7.781 | 1.33 | 168.01 | 53.71 |
| 351 | 2jsa-12 | ILE_A_60_N | ILE_A_60_H | 0.488 | -0.078 | 7.781 | 1.016 | -0.728 | 7.216 | 1.01 | 124.01 | -50.91 |
| 352 | 2jsa-12 | ILE_A_60_N | ILE_A_60_CA | 0.488 | -0.078 | 7.781 | -0.085 | 1.096 | 7.146 | 1.45 | 115.92 | 116.02 |
| 353 | 2jsa-12 | ILE_A_60_CA | ILE_A_60_HA | -0.085 | 1.096 | 7.146 | 0.436 | 1.974 | 7.528 | 1.09 | 69.49 | 59.32 |
| 354 | 2jsa-12 | ILE_A_60_CA | ILE_A_60_C | -0.085 | 1.096 | 7.146 | -1.558 | 1.218 | 7.544 | 1.53 | 74.93 | 175.27 |
| 355 | 2jsa-12 | ILE_A_60_C | ILE_A_60_O | -1.558 | 1.218 | 7.544 | -2.023 | 2.303 | 7.889 | 1.23 | 73.71 | 113.20 |
| 356 | 2jsa-12 | ILE_A_60_C | ILE_A_61_N | -1.558 | 1.218 | 7.544 | -2.25 | 0.09 | 7.481 | 1.32 | 92.73 | -121.53 |
| 357 | 2jsa-12 | ILE_A_61_N | ILE_A_61_H | -2.25 | 0.09 | 7.481 | -1.801 | -0.766 | 7.188 | 1.01 | 106.86 | -62.32 |
| 358 | 2jsa-12 | ILE_A_61_N | ILE_A_61_CA | -2.25 | 0.09 | 7.481 | -3.66 | 0.057 | 7.831 | 1.45 | 76.06 | -178.66 |
| 359 | 2jsa-12 | ILE_A_61_CA | ILE_A_61_HA | -3.66 | 0.057 | 7.831 | -4.14 | 0.926 | 7.38 | 1.09 | 114.43 | 118.91 |
| 360 | 2jsa-12 | ILE_A_61_CA | ILE_A_61_C | -3.66 | 0.057 | 7.831 | -3.808 | 0.18 | 9.349 | 1.53 | 7.22 | 140.27 |
| 361 | 2jsa-12 | ILE_A_61_C | ILE_A_61_O | -3.808 | 0.18 | 9.349 | -4.883 | 0.513 | 9.846 | 1.23 | 66.17 | 162.79 |
| 362 | 2jsa-12 | ILE_A_61_C | HIS_A_62_N | -3.808 | 0.18 | 9.349 | -2.713 | -0.096 | 10.042 | 1.32 | 58.46 | -14.15 |
| 363 | 2jsa-12 | HIS_A_62_N | HIS_A_62_H | -2.713 | -0.096 | 10.042 | -1.864 | -0.365 | 9.566 | 1.01 | 118.12 | -17.58 |
| 364 | 2jsa-12 | HIS_A_62_N | HIS_A_62_CA | -2.713 | -0.096 | 10.042 | -2.707 | -0.019 | 11.494 | 1.45 | 3.04 | 85.54 |
| 365 | 2jsa-12 | HIS_A_62_CA | HIS_A_62_HA | -2.707 | -0.019 | 11.494 | -3.596 | -0.531 | 11.862 | 1.09 | 70.27 | -150.06 |
| 366 | 2jsa-12 | HIS_A_62_CA | HIS_A_62_C | -2.707 | -0.019 | 11.494 | -2.794 | 1.443 | 11.931 | 1.53 | 73.39 | 93.41 |
| 367 | 2jsa-12 | HIS_A_62_C | HIS_A_62_O | -2.794 | 1.443 | 11.931 | -3.301 | 1.744 | 13.012 | 1.23 | 28.61 | 149.30 |
| 368 | 2jsa-12 | HIS_A_62_C | GLU_A_63_N | -2.794 | 1.443 | 11.931 | -2.292 | 2.316 | 11.07 | 1.32 | 130.53 | 60.10 |
| 369 | 2jsa-12 | GLU_A_63_N | GLU_A_63_H | -2.292 | 2.316 | 11.07 | -1.891 | 1.998 | 10.199 | 1.01 | 149.56 | -38.42 |
| 370 | 2jsa-12 | GLU_A_63_N | GLU_A_63_CA | -2.292 | 2.316 | 11.07 | -2.306 | 3.741 | 11.354 | 1.45 | 78.73 | 90.56 |
| 371 | 2jsa-12 | GLU_A_63_CA | GLU_A_63_HA | -2.306 | 3.741 | 11.354 | -2.128 | 3.873 | 12.421 | 1.09 | 11.73 | 36.56 |
| 372 | 2jsa-12 | GLU_A_63_CA | GLU_A_63_C | -2.306 | 3.741 | 11.354 | -3.674 | 4.337 | 11.017 | 1.53 | 102.73 | 156.46 |
| 373 | 2jsa-12 | GLU_A_63_C | GLU_A_63_O | -3.674 | 4.337 | 11.017 | -4.286 | 5.006 | 11.847 | 1.23 | 47.53 | 132.45 |
| 374 | 2jsa-12 | GLU_A_63_C | LEU_A_64_N | -3.674 | 4.337 | 11.017 | -4.115 | 4.071 | 9.795 | 1.33 | 157.15 | -148.90 |
| 375 | 2jsa-12 | LEU_A_64_N | LEU_A_64_H | -4.115 | 4.071 | 9.795 | -3.558 | 3.512 | 9.164 | 1.01 | 128.65 | -45.10 |
| 376 | 2jsa-12 | LEU_A_64_N | LEU_A_64_CA | -4.115 | 4.071 | 9.795 | -5.398 | 4.571 | 9.336 | 1.45 | 108.44 | 158.71 |
| 377 | 2jsa-12 | LEU_A_64_CA | LEU_A_64_HA | -5.398 | 4.571 | 9.336 | -5.392 | 5.656 | 9.437 | 1.09 | 84.68 | 89.68 |
| 378 | 2jsa-12 | LEU_A_64_CA | LEU_A_64_C | -5.398 | 4.571 | 9.336 | -6.506 | 4.026 | 10.241 | 1.53 | 53.76 | -153.81 |
| 379 | 2jsa-12 | LEU_A_64_C | LEU_A_64_O | -6.506 | 4.026 | 10.241 | -7.437 | 4.749 | 10.593 | 1.23 | 73.37 | 142.17 |
| 380 | 2jsa-12 | LEU_A_64_C | GLU_A_65_N | -6.506 | 4.026 | 10.241 | -6.367 | 2.757 | 10.593 | 1.32 | 74.58 | -83.75 |
| 381 | 2jsa-12 | GLU_A_65_N | GLU_A_65_H | -6.367 | 2.757 | 10.593 | -5.573 | 2.223 | 10.268 | 1.01 | 108.76 | -33.92 |
| 382 | 2jsa-12 | GLU_A_65_N | GLU_A_65_CA | -6.367 | 2.757 | 10.593 | -7.345 | 2.107 | 11.45 | 1.45 | 53.88 | -146.39 |
| 383 | 2jsa-12 | GLU_A_65_CA | GLU_A_65_HA | -7.345 | 2.107 | 11.45 | -8.338 | 2.335 | 11.063 | 1.09 | 110.80 | 167.07 |
| 384 | 2jsa-12 | GLU_A_65_CA | GLU_A_65_C | -7.345 | 2.107 | 11.45 | -7.247 | 2.653 | 12.875 | 1.53 | 21.27 | 79.82 |
| 385 | 2jsa-12 | GLU_A_65_C | GLU_A_65_O | -7.247 | 2.653 | 12.875 | -8.221 | 2.615 | 13.627 | 1.23 | 52.35 | -177.77 |
| 386 | 2jsa-12 | GLU_A_65_C | GLY_A_66_N | -7.247 | 2.653 | 12.875 | -6.063 | 3.149 | 13.205 | 1.33 | 75.58 | 22.73 |
| 387 | 2jsa-12 | GLY_A_66_N | GLY_A_66_H | -6.063 | 3.149 | 13.205 | -5.306 | 3.148 | 12.536 | 1.01 | 131.47 | -0.08 |
| 388 | 2jsa-12 | GLY_A_66_N | GLY_A_66_CA | -6.063 | 3.149 | 13.205 | -5.827 | 3.703 | 14.528 | 1.45 | 24.47 | 66.93 |
| 389 | 2jsa-12 | GLY_A_66_CA | GLY_A_66_C | -5.827 | 3.703 | 14.528 | -6.435 | 5.101 | 14.654 | 1.53 | 85.28 | 113.50 |
| 390 | 2jsa-12 | GLY_A_66_C | GLY_A_66_O | -6.435 | 5.101 | 14.654 | -6.972 | 5.456 | 15.702 | 1.23 | 31.56 | 146.53 |
| 391 | 2jsa-12 | GLY_A_66_C | GLY_A_67_N | -6.435 | 5.101 | 14.654 | -6.332 | 5.857 | 13.57 | 1.33 | 144.86 | 82.24 |
| 392 | 2jsa-12 | GLY_A_67_N | GLY_A_67_H | -6.332 | 5.857 | 13.57 | -5.878 | 5.499 | 12.742 | 1.01 | 145.07 | -38.26 |
| 393 | 2jsa-12 | GLY_A_67_N | GLY_A_67_CA | -6.332 | 5.857 | 13.57 | -6.865 | 7.208 | 13.547 | 1.45 | 90.91 | 111.53 |
| 394 | 2jsa-12 | GLY_A_67_CA | GLY_A_67_C | -6.865 | 7.208 | 13.547 | -6.003 | 8.123 | 12.674 | 1.53 | 124.78 | 46.71 |
| 395 | 2jsa-12 | GLY_A_67_C | GLY_A_67_O | -6.003 | 8.123 | 12.674 | -6.441 | 9.202 | 12.278 | 1.23 | 108.78 | 112.09 |
| 396 | 2jsa-12 | GLY_A_67_C | THR_A_68_N | -6.003 | 8.123 | 12.674 | -4.793 | 7.658 | 12.4 | 1.32 | 101.94 | -21.02 |
| 397 | 2jsa-12 | THR_A_68_N | THR_A_68_H | -4.793 | 7.658 | 12.4 | -4.499 | 6.761 | 12.76 | 1.01 | 69.12 | -71.85 |
| 398 | 2jsa-12 | THR_A_68_N | THR_A_68_CA | -4.793 | 7.658 | 12.4 | -3.866 | 8.42 | 11.582 | 1.45 | 124.28 | 39.42 |
| 399 | 2jsa-12 | THR_A_68_CA | THR_A_68_HA | -3.866 | 8.42 | 11.582 | -3.755 | 9.415 | 12.014 | 1.09 | 66.66 | 83.63 |
| 400 | 2jsa-12 | THR_A_68_CA | THR_A_68_C | -3.866 | 8.42 | 11.582 | -4.415 | 8.581 | 10.163 | 1.53 | 158.04 | 163.66 |
| 401 | 2jsa-12 | THR_A_68_C | THR_A_68_O | -4.415 | 8.581 | 10.163 | -5.225 | 7.772 | 9.712 | 1.23 | 111.50 | -135.04 |
| 402 | 2jsa-12 | THR_A_68_C | ALA_A_69_N | -4.415 | 8.581 | 10.163 | -3.952 | 9.629 | 9.499 | 1.32 | 120.09 | 66.16 |
| 403 | 2jsa-12 | ALA_A_69_N | ALA_A_69_H | -3.952 | 9.629 | 9.499 | -3.287 | 10.253 | 9.934 | 1.01 | 64.50 | 43.18 |
| 404 | 2jsa-12 | ALA_A_69_N | ALA_A_69_CA | -3.952 | 9.629 | 9.499 | -4.386 | 9.906 | 8.139 | 1.45 | 159.26 | 147.45 |
| 405 | 2jsa-12 | ALA_A_69_CA | ALA_A_69_HA | -4.386 | 9.906 | 8.139 | -5.42 | 9.58 | 8.028 | 1.09 | 95.85 | -162.50 |
| 406 | 2jsa-12 | ALA_A_69_CA | ALA_A_69_C | -4.386 | 9.906 | 8.139 | -3.521 | 9.111 | 7.16 | 1.53 | 129.80 | -42.59 |
| 407 | 2jsa-12 | ALA_A_69_C | ALA_A_69_O | -3.521 | 9.111 | 7.16 | -2.499 | 8.547 | 7.547 | 1.23 | 71.66 | -28.89 |
| 408 | 2jsa-12 | ALA_A_69_C | PRO_A_70_N | -3.521 | 9.111 | 7.16 | -3.975 | 9.09 | 5.878 | 1.36 | 160.48 | -177.35 |
| 409 | 2jsa-12 | PRO_A_70_N | PRO_A_70_CA | -3.975 | 9.09 | 5.878 | -3.253 | 8.374 | 4.84 | 1.45 | 135.59 | -44.76 |
| 410 | 2jsa-12 | PRO_A_70_CA | PRO_A_70_HA | -3.253 | 8.374 | 4.84 | -2.954 | 7.391 | 5.204 | 1.09 | 70.49 | -73.08 |
| 411 | 2jsa-12 | PRO_A_70_CA | PRO_A_70_C | -3.253 | 8.374 | 4.84 | -1.995 | 9.137 | 4.42 | 1.53 | 105.93 | 31.24 |
| 412 | 2jsa-12 | PRO_A_70_C | PRO_A_70_O | -1.995 | 9.137 | 4.42 | -1.092 | 8.564 | 3.813 | 1.23 | 119.58 | -32.40 |
| 413 | 2jsa-12 | PRO_A_70_C | LYS_A_71_N | -1.995 | 9.137 | 4.42 | -1.976 | 10.417 | 4.76 | 1.32 | 75.13 | 89.15 |
| 414 | 2jsa-12 | LYS_A_71_N | LYS_A_71_H | -1.976 | 10.417 | 4.76 | -2.755 | 10.822 | 5.26 | 1.01 | 60.34 | 152.53 |
| 415 | 2jsa-12 | LYS_A_71_N | LYS_A_71_CA | -1.976 | 10.417 | 4.76 | -0.844 | 11.264 | 4.426 | 1.45 | 103.29 | 36.81 |
| 416 | 2jsa-12 | LYS_A_71_CA | LYS_A_71_HA | -0.844 | 11.264 | 4.426 | -0.269 | 10.767 | 3.644 | 1.09 | 135.82 | -40.84 |

| | | | | | | | | | | | |
|---|---|---|---|---|---|---|---|---|---|---|---|
| 417 | 2jsa-12 | LYS_A_71_CA | LYS_A_71_C | -0.844 | 11.264 | 4.426 | 0.062 | 11.404 | 5.651 | 1.53 | 36.81 | 8.78 |
| 418 | 2jsa-12 | LYS_A_71_C | LYS_A_71_O | 0.062 | 11.404 | 5.651 | 0.978 | 12.224 | 5.659 | 1.23 | 89.63 | 41.83 |
| 419 | 2jsa-12 | LYS_A_71_C | ASP_A_72_N | 0.062 | 11.404 | 5.651 | -0.226 | 10.591 | 6.657 | 1.33 | 40.61 | -109.51 |
| 420 | 2jsa-12 | ASP_A_72_N | ASP_A_72_H | -0.226 | 10.591 | 6.657 | -0.995 | 9.941 | 6.581 | 1.01 | 94.32 | -139.79 |
| 421 | 2jsa-12 | ASP_A_72_N | ASP_A_72_CA | -0.226 | 10.591 | 6.657 | 0.55 | 10.614 | 7.884 | 1.45 | 32.32 | 1.70 |
| 422 | 2jsa-12 | ASP_A_72_CA | ASP_A_72_HA | 0.55 | 10.614 | 7.884 | 1.283 | 11.419 | 7.827 | 1.09 | 93.00 | 47.68 |
| 423 | 2jsa-12 | ASP_A_72_CA | ASP_A_72_C | 0.55 | 10.614 | 7.884 | 1.265 | 9.272 | 8.054 | 1.53 | 83.62 | -61.95 |
| 424 | 2jsa-12 | ASP_A_72_C | ASP_A_72_O | 1.265 | 9.272 | 8.054 | 2.407 | 9.227 | 8.51 | 1.23 | 68.25 | -2.26 |
| 425 | 2jsa-12 | ASP_A_72_C | VAL_A_73_N | 1.265 | 9.272 | 8.054 | 0.564 | 8.213 | 7.68 | 1.32 | 106.41 | -123.50 |
| 426 | 2jsa-12 | VAL_A_73_N | VAL_A_73_H | 0.564 | 8.213 | 7.68 | -0.37 | 8.325 | 7.314 | 1.01 | 111.26 | 173.16 |
| 427 | 2jsa-12 | VAL_A_73_N | VAL_A_73_CA | 0.564 | 8.213 | 7.68 | 1.118 | 6.873 | 7.786 | 1.45 | 85.82 | -67.54 |
| 428 | 2jsa-12 | VAL_A_73_CA | VAL_A_73_HA | 1.118 | 6.873 | 7.786 | 1.331 | 6.67 | 8.836 | 1.09 | 15.65 | -43.62 |
| 429 | 2jsa-12 | VAL_A_73_CA | VAL_A_73_C | 1.118 | 6.873 | 7.786 | 2.437 | 6.808 | 7.014 | 1.53 | 120.31 | -2.82 |
| 430 | 2jsa-12 | VAL_A_73_C | VAL_A_73_O | 2.437 | 6.808 | 7.014 | 3.434 | 6.301 | 7.524 | 1.23 | 65.49 | -26.95 |
| 431 | 2jsa-12 | VAL_A_73_C | CYS_A_74_N | 2.437 | 6.808 | 7.014 | 2.399 | 7.329 | 5.796 | 1.33 | 156.79 | 94.17 |
| 432 | 2jsa-12 | CYS_A_74_N | CYS_A_74_H | 2.399 | 7.329 | 5.796 | 1.544 | 7.731 | 5.441 | 1.01 | 110.59 | 154.82 |
| 433 | 2jsa-12 | CYS_A_74_N | CYS_A_74_CA | 2.399 | 7.329 | 5.796 | 3.579 | 7.336 | 4.949 | 1.45 | 125.67 | 0.34 |
| 434 | 2jsa-12 | CYS_A_74_CA | CYS_A_74_HA | 3.579 | 7.336 | 4.949 | 4.138 | 6.417 | 5.131 | 1.09 | 80.40 | -58.69 |
| 435 | 2jsa-12 | CYS_A_74_CA | CYS_A_74_C | 3.579 | 7.336 | 4.949 | 4.454 | 8.526 | 5.353 | 1.53 | 74.70 | 53.67 |
| 436 | 2jsa-12 | CYS_A_74_C | CYS_A_74_O | 4.454 | 8.526 | 5.353 | 4.762 | 9.383 | 4.527 | 1.23 | 132.21 | 70.23 |
| 437 | 2jsa-12 | CYS_A_74_C | THR_A_75_N | 4.454 | 8.526 | 5.353 | 4.828 | 8.539 | 6.623 | 1.32 | 16.42 | 1.99 |
| 438 | 2jsa-12 | THR_A_75_N | THR_A_75_H | 4.828 | 8.539 | 6.623 | 4.537 | 7.799 | 7.246 | 1.01 | 51.92 | -111.47 |
| 439 | 2jsa-12 | THR_A_75_N | THR_A_75_CA | 4.828 | 8.539 | 6.623 | 5.66 | 9.608 | 7.148 | 1.45 | 68.82 | 52.11 |
| 440 | 2jsa-12 | THR_A_75_CA | THR_A_75_HA | 5.66 | 9.608 | 7.148 | 6.4 | 9.878 | 6.394 | 1.09 | 133.75 | 20.05 |
| 441 | 2jsa-12 | THR_A_75_CA | THR_A_75_C | 5.66 | 9.608 | 7.148 | 6.41 | 9.138 | 8.395 | 1.53 | 35.37 | -32.07 |
| 442 | 2jsa-12 | THR_A_75_C | THR_A_75_O | 6.41 | 9.138 | 8.395 | 7.639 | 9.16 | 8.43 | 1.23 | 88.37 | 1.03 |
| 443 | 2jsa-12 | THR_A_75_C | LYS_A_76_N | 6.41 | 9.138 | 8.395 | 5.638 | 8.724 | 9.39 | 1.33 | 41.36 | -151.80 |
| 444 | 2jsa-12 | LYS_A_76_N | LYS_A_76_H | 5.638 | 8.724 | 9.39 | 4.633 | 8.734 | 9.292 | 1.01 | 95.57 | 179.43 |
| 445 | 2jsa-12 | LYS_A_76_N | LYS_A_76_CA | 5.638 | 8.724 | 9.39 | 6.214 | 8.249 | 10.636 | 1.45 | 30.93 | -39.51 |
| 446 | 2jsa-12 | LYS_A_76_CA | LYS_A_76_HA | 6.214 | 8.249 | 10.636 | 7.088 | 8.862 | 10.859 | 1.09 | 78.20 | 35.04 |
| 447 | 2jsa-12 | LYS_A_76_CA | LYS_A_76_C | 6.214 | 8.249 | 10.636 | 6.692 | 6.807 | 10.456 | 1.53 | 96.76 | -71.66 |
| 448 | 2jsa-12 | LYS_A_76_C | LYS_A_76_O | 6.692 | 6.807 | 10.456 | 7.736 | 6.425 | 10.982 | 1.23 | 64.68 | -20.10 |
| 449 | 2jsa-12 | LYS_A_76_C | LEU_A_77_N | 6.692 | 6.807 | 10.456 | 5.904 | 6.046 | 9.71 | 1.33 | 124.25 | -136.00 |
| 450 | 2jsa-12 | LEU_A_77_N | LEU_A_77_H | 5.904 | 6.046 | 9.71 | 5.06 | 6.428 | 9.308 | 1.01 | 113.46 | 155.65 |
| 451 | 2jsa-12 | LEU_A_77_N | LEU_A_77_CA | 5.904 | 6.046 | 9.71 | 6.233 | 4.654 | 9.455 | 1.45 | 100.11 | -76.70 |
| 452 | 2jsa-12 | LEU_A_77_CA | LEU_A_77_HA | 6.233 | 4.654 | 9.455 | 6.46 | 4.18 | 10.41 | 1.09 | 28.82 | -64.41 |
| 453 | 2jsa-12 | LEU_A_77_CA | LEU_A_77_C | 6.233 | 4.654 | 9.455 | 7.493 | 4.583 | 8.589 | 1.53 | 124.46 | -3.23 |
| 454 | 2jsa-12 | LEU_A_77_C | LEU_A_77_O | 7.493 | 4.583 | 8.589 | 8.031 | 3.502 | 8.357 | 1.23 | 100.88 | -63.54 |
| 455 | 2jsa-12 | LEU_A_77_C | ASN_A_78_N | 7.493 | 4.583 | 8.589 | 7.927 | 5.751 | 8.135 | 1.33 | 110.02 | 69.62 |
| 456 | 2jsa-12 | ASN_A_78_N | ASN_A_78_H | 7.927 | 5.751 | 8.135 | 7.435 | 6.602 | 8.368 | 1.01 | 76.67 | 120.03 |
| 457 | 2jsa-12 | ASN_A_78_N | ASN_A_78_CA | 7.927 | 5.751 | 8.135 | 9.112 | 5.834 | 7.299 | 1.45 | 125.14 | 4.01 |
| 458 | 2jsa-12 | ASN_A_78_CA | ASN_A_78_HA | 9.112 | 5.834 | 7.299 | 9.324 | 6.881 | 7.083 | 1.09 | 101.43 | 78.55 |
| 459 | 2jsa-12 | ASN_A_78_CA | ASN_A_78_C | 9.112 | 5.834 | 7.299 | 8.867 | 5.074 | 5.994 | 1.53 | 148.54 | -107.87 |
| 460 | 2jsa-12 | ASN_A_78_C | ASN_A_78_O | 8.867 | 5.074 | 5.994 | 9.781 | 4.458 | 5.449 | 1.23 | 116.31 | -33.98 |
| 461 | 2jsa-12 | ASN_A_78_C | GLU_A_79_N | 8.867 | 5.074 | 5.994 | 7.628 | 5.143 | 5.531 | 1.32 | 110.46 | 176.81 |
| 462 | 2jsa-12 | GLU_A_79_N | GLU_A_79_H | 7.628 | 5.143 | 5.531 | 6.927 | 5.67 | 6.032 | 1.01 | 60.26 | 143.06 |
| 463 | 2jsa-12 | GLU_A_79_N | GLU_A_79_CA | 7.628 | 5.143 | 5.531 | 7.25 | 4.469 | 4.3 | 1.45 | 147.88 | -119.29 |
| 464 | 2jsa-12 | GLU_A_79_CA | GLU_A_79_HA | 7.25 | 4.469 | 4.3 | 8.141 | 3.996 | 3.888 | 1.09 | 112.22 | -27.96 |
| 465 | 2jsa-12 | GLU_A_79_CA | GLU_A_79_C | 7.25 | 4.469 | 4.3 | 6.728 | 5.483 | 3.279 | 1.53 | 131.84 | 117.24 |
| 466 | 2jsa-12 | GLU_A_79_C | GLU_A_79_O | 6.728 | 5.483 | 3.279 | 5.539 | 5.496 | 2.966 | 1.23 | 104.75 | 179.37 |
| 467 | 2jsa-12 | GLU_A_79_C | CYS_A_80_N | 6.728 | 5.483 | 3.279 | 7.643 | 6.306 | 2.788 | 1.33 | 111.75 | 41.97 |
| 468 | 2jsa-12 | CYS_A_80_N | CYS_A_80_H | 7.643 | 6.306 | 2.788 | 8.604 | 6.237 | 3.091 | 1.01 | 72.54 | -4.11 |
| 469 | 2jsa-12 | CYS_A_80_N | CYS_A_80_CA | 7.643 | 6.306 | 2.788 | 7.29 | 7.32 | 1.809 | 1.45 | 132.36 | 109.19 |
| 470 | 2jsa-12 | CYS_A_80_CA | CYS_A_80_HA | 7.29 | 7.32 | 1.809 | 7.07 | 6.823 | 0.864 | 1.09 | 150.09 | -113.88 |
| 471 | 2jsa-12 | CYS_A_80_CA | CYS_A_80_C | 7.29 | 7.32 | 1.809 | 8.499 | 8.235 | 1.608 | 1.53 | 97.55 | 37.12 |
| 472 | 2jsa-12 | CYS_A_80_C | CYS_A_80_O | 8.499 | 8.235 | 1.608 | 8.966 | 8.415 | 0.484 | 1.23 | 156.00 | 21.08 |
| 473 | 2jsa-12 | CYS_A_80_C | PRO_A_81_N | 8.499 | 8.235 | 1.608 | 8.985 | 8.804 | 2.744 | 1.36 | 33.37 | 49.50 |
| 474 | 2jsa-12 | PRO_A_81_N | PRO_A_81_CA | 8.985 | 8.804 | 2.744 | 10.131 | 9.696 | 2.702 | 1.45 | 91.66 | 37.90 |
| 475 | 2jsa-12 | PRO_A_81_CA | PRO_A_81_HA | 10.131 | 9.696 | 2.702 | 10.023 | 10.405 | 1.881 | 1.09 | 138.86 | 98.66 |
| 476 | 2jsa-12 | PRO_A_81_CA | PRO_A_81_C | 10.131 | 9.696 | 2.702 | 11.429 | 8.913 | 2.498 | 1.53 | 97.66 | -31.10 |
| 477 | 2jsa-12 | PRO_A_81_C | PRO_A_81_O | 11.429 | 8.913 | 2.498 | 12.249 | 8.814 | 3.41 | 1.23 | 42.17 | -6.88 |
| 1 | 2jsa-13 | ARG_A_1_N | ARG_A_1_H | 14.991 | 3.291 | 3.617 | 14.475 | 4.007 | 4.108 | 1.01 | 60.91 | 125.78 |
| 2 | 2jsa-13 | ARG_A_1_N | ARG_A_1_CA | 14.991 | 3.291 | 3.617 | 14.724 | 3.088 | 2.203 | 1.45 | 166.66 | -142.75 |
| 3 | 2jsa-13 | ARG_A_1_CA | ARG_A_1_HA | 14.724 | 3.088 | 2.203 | 15.645 | 3.27 | 1.649 | 1.09 | 120.55 | 11.18 |
| 4 | 2jsa-13 | ARG_A_1_CA | ARG_A_1_C | 14.724 | 3.088 | 2.203 | 13.65 | 4.065 | 1.72 | 1.53 | 108.40 | 137.71 |
| 5 | 2jsa-13 | ARG_A_1_C | ARG_A_1_O | 13.65 | 4.065 | 1.72 | 12.634 | 4.256 | 2.388 | 1.23 | 57.13 | 169.35 |
| 6 | 2jsa-13 | ARG_A_1_C | SER_A_2_N | 13.65 | 4.065 | 1.72 | 13.91 | 4.657 | 0.565 | 1.32 | 150.76 | 66.29 |
| 7 | 2jsa-13 | SER_A_2_N | SER_A_2_H | 13.91 | 4.657 | 0.565 | 14.766 | 4.451 | 0.069 | 1.01 | 119.39 | -13.53 |
| 8 | 2jsa-13 | SER_A_2_N | SER_A_2_CA | 13.91 | 4.657 | 0.565 | 12.979 | 5.61 | -0.015 | 1.45 | 113.53 | 134.33 |
| 9 | 2jsa-13 | SER_A_2_CA | SER_A_2_HA | 12.979 | 5.61 | -0.015 | 12.005 | 5.461 | 0.449 | 1.09 | 64.78 | -171.30 |
| 10 | 2jsa-13 | SER_A_2_CA | SER_A_2_C | 12.979 | 5.61 | -0.015 | 12.848 | 5.364 | -1.519 | 1.53 | 169.50 | -118.04 |
| 11 | 2jsa-13 | SER_A_2_C | SER_A_2_O | 12.848 | 5.364 | -1.519 | 13.839 | 5.399 | -2.247 | 1.23 | 126.28 | 2.02 |
| 12 | 2jsa-13 | SER_A_2_C | ALA_A_3_N | 12.848 | 5.364 | -1.519 | 11.616 | 5.12 | -1.942 | 1.33 | 108.61 | -168.80 |
| 13 | 2jsa-13 | ALA_A_3_N | ALA_A_3_H | 11.616 | 5.12 | -1.942 | 10.848 | 5.105 | -1.287 | 1.01 | 49.55 | -178.88 |
| 14 | 2jsa-13 | ALA_A_3_N | ALA_A_3_CA | 11.616 | 5.12 | -1.942 | 11.343 | 4.869 | -3.347 | 1.45 | 165.21 | -137.40 |
| 15 | 2jsa-13 | ALA_A_3_CA | ALA_A_3_HA | 11.343 | 4.869 | -3.347 | 11.679 | 5.728 | -3.929 | 1.09 | 122.25 | 68.64 |
| 16 | 2jsa-13 | ALA_A_3_CA | ALA_A_3_C | 11.343 | 4.869 | -3.347 | 9.834 | 4.708 | -3.547 | 1.53 | 97.51 | -173.91 |
| 17 | 2jsa-13 | ALA_A_3_C | ALA_A_3_O | 9.834 | 4.708 | -3.547 | 9.041 | 5.247 | -2.777 | 1.23 | 51.23 | 145.80 |
| 18 | 2jsa-13 | ALA_A_3_C | LEU_A_4_N | 9.834 | 4.708 | -3.547 | 9.484 | 3.965 | -4.587 | 1.33 | 141.70 | -115.22 |
| 19 | 2jsa-13 | LEU_A_4_N | LEU_A_4_H | 9.484 | 3.965 | -4.587 | 10.191 | 3.554 | -5.179 | 1.01 | 125.90 | -30.17 |
| 20 | 2jsa-13 | LEU_A_4_N | LEU_A_4_CA | 9.484 | 3.965 | -4.587 | 8.085 | 3.726 | -4.898 | 1.45 | 102.36 | -170.31 |
| 21 | 2jsa-13 | LEU_A_4_CA | LEU_A_4_HA | 8.085 | 3.726 | -4.898 | 7.613 | 4.69 | -5.088 | 1.09 | 100.04 | 116.09 |
| 22 | 2jsa-13 | LEU_A_4_CA | LEU_A_4_C | 8.085 | 3.726 | -4.898 | 7.396 | 3.103 | -3.684 | 1.53 | 37.42 | -137.88 |
| 23 | 2jsa-13 | LEU_A_4_C | LEU_A_4_O | 7.396 | 3.103 | -3.684 | 6.17 | 3.136 | -3.575 | 1.23 | 84.92 | 178.46 |
| 24 | 2jsa-13 | LEU_A_4_C | SER_A_5_N | 7.396 | 3.103 | -3.684 | 8.212 | 2.547 | -2.8 | 1.33 | 48.16 | -34.27 |
| 25 | 2jsa-13 | SER_A_5_N | SER_A_5_H | 8.212 | 2.547 | -2.8 | 9.211 | 2.555 | -2.954 | 1.01 | 98.76 | 0.46 |
| 26 | 2jsa-13 | SER_A_5_N | SER_A_5_CA | 8.212 | 2.547 | -2.8 | 7.696 | 1.918 | -1.596 | 1.45 | 34.05 | -129.36 |
| 27 | 2jsa-13 | SER_A_5_CA | SER_A_5_HA | 7.696 | 1.918 | -1.596 | 7.217 | 0.981 | -1.879 | 1.09 | 105.05 | -117.08 |
| 28 | 2jsa-13 | SER_A_5_CA | SER_A_5_C | 7.696 | 1.918 | -1.596 | 6.651 | 2.822 | -0.938 | 1.53 | 64.54 | 139.14 |
| 29 | 2jsa-13 | SER_A_5_C | SER_A_5_O | 6.651 | 2.822 | -0.938 | 5.616 | 2.346 | -0.477 | 1.23 | 67.97 | -155.30 |
| 30 | 2jsa-13 | SER_A_5_C | CYS_A_6_N | 6.651 | 2.822 | -0.938 | 6.96 | 4.11 | -0.917 | 1.32 | 89.09 | 76.51 |
| 31 | 2jsa-13 | CYS_A_6_N | CYS_A_6_H | 6.96 | 4.11 | -0.917 | 7.831 | 4.431 | -1.316 | 1.01 | 113.26 | 20.23 |
| 32 | 2jsa-13 | CYS_A_6_N | CYS_A_6_CA | 6.96 | 4.11 | -0.917 | 6.061 | 5.085 | -0.324 | 1.45 | 65.91 | 132.68 |
| 33 | 2jsa-13 | CYS_A_6_CA | CYS_A_6_HA | 6.061 | 5.085 | -0.324 | 5.861 | 4.785 | 0.705 | 1.09 | 19.31 | -123.69 |
| 34 | 2jsa-13 | CYS_A_6_CA | CYS_A_6_C | 6.061 | 5.085 | -0.324 | 4.745 | 5.062 | -1.103 | 1.53 | 120.62 | -179.00 |
| 35 | 2jsa-13 | CYS_A_6_C | CYS_A_6_O | 4.745 | 5.062 | -1.103 | 3.7 | 4.719 | -0.554 | 1.23 | 63.47 | -161.83 |

| | | | | | | | | | | | | |
|---|---|---|---|---|---|---|---|---|---|---|---|---|
| 36 | 2jsa-13 | CYS_A_6_C | GLN_A_7_N | 4.745 | 5.062 | -1.103 | 4.839 | 5.432 | -2.372 | 1.33 | 163.26 | 75.75 |
| 37 | 2jsa-13 | GLN_A_7_N | GLN_A_7_H | 4.839 | 5.432 | -2.372 | 5.733 | 5.703 | -2.758 | 1.01 | 112.45 | 16.86 |
| 38 | 2jsa-13 | GLN_A_7_N | GLN_A_7_CA | 4.839 | 5.432 | -2.372 | 3.669 | 5.458 | -3.233 | 1.45 | 126.34 | 178.73 |
| 39 | 2jsa-13 | GLN_A_7_CA | GLN_A_7_HA | 3.669 | 5.458 | -3.233 | 3.108 | 6.366 | -3.011 | 1.09 | 78.25 | 121.71 |
| 40 | 2jsa-13 | GLN_A_7_CA | GLN_A_7_C | 3.669 | 5.458 | -3.233 | 2.77 | 4.254 | -2.944 | 1.53 | 79.11 | -126.75 |
| 41 | 2jsa-13 | GLN_A_7_C | GLN_A_7_O | 2.77 | 4.254 | -2.944 | 1.557 | 4.399 | -2.8 | 1.23 | 83.28 | 173.18 |
| 42 | 2jsa-13 | GLN_A_7_C | MET_A_8_N | 2.77 | 4.254 | -2.944 | 3.4 | 3.091 | -2.867 | 1.32 | 86.67 | -61.56 |
| 43 | 2jsa-13 | MET_A_8_N | MET_A_8_H | 3.4 | 3.091 | -2.867 | 4.401 | 3.046 | -2.997 | 1.01 | 97.39 | -2.57 |
| 44 | 2jsa-13 | MET_A_8_N | MET_A_8_CA | 3.4 | 3.091 | -2.867 | 2.673 | 1.861 | -2.597 | 1.45 | 79.30 | -120.59 |
| 45 | 2jsa-13 | MET_A_8_CA | MET_A_8_HA | 2.673 | 1.861 | -2.597 | 2.045 | 1.63 | -3.457 | 1.09 | 142.11 | -159.80 |
| 46 | 2jsa-13 | MET_A_8_CA | MET_A_8_C | 2.673 | 1.861 | -2.597 | 1.785 | 2.011 | -1.36 | 1.53 | 36.06 | 170.41 |
| 47 | 2jsa-13 | MET_A_8_C | MET_A_8_O | 1.785 | 2.011 | -1.36 | 0.642 | 1.557 | -1.354 | 1.23 | 89.72 | -158.34 |
| 48 | 2jsa-13 | MET_A_8_C | CYS_A_9_N | 1.785 | 2.011 | -1.36 | 2.345 | 2.648 | -0.342 | 1.33 | 39.80 | 48.68 |
| 49 | 2jsa-13 | CYS_A_9_N | CYS_A_9_H | 2.345 | 2.648 | -0.342 | 3.291 | 2.996 | -0.417 | 1.01 | 94.26 | 20.20 |
| 50 | 2jsa-13 | CYS_A_9_N | CYS_A_9_CA | 2.345 | 2.648 | -0.342 | 1.618 | 2.861 | 0.898 | 1.45 | 31.42 | 163.67 |
| 51 | 2jsa-13 | CYS_A_9_CA | CYS_A_9_HA | 1.618 | 2.861 | 0.898 | 1.315 | 1.89 | 1.29 | 1.09 | 68.92 | -107.33 |
| 52 | 2jsa-13 | CYS_A_9_CA | CYS_A_9_C | 1.618 | 2.861 | 0.898 | 0.36 | 3.674 | 0.583 | 1.53 | 101.88 | 147.13 |
| 53 | 2jsa-13 | CYS_A_9_C | CYS_A_9_O | 0.36 | 3.674 | 0.583 | -0.665 | 3.517 | 1.244 | 1.23 | 57.48 | -171.29 |
| 54 | 2jsa-13 | CYS_A_9_C | GLU_A_10_N | 0.36 | 3.674 | 0.583 | 0.48 | 4.524 | -0.427 | 1.33 | 139.64 | 81.96 |
| 55 | 2jsa-13 | GLU_A_10_N | GLU_A_10_H | 0.48 | 4.524 | -0.427 | 1.354 | 4.599 | -0.927 | 1.01 | 119.68 | 4.90 |
| 56 | 2jsa-13 | GLU_A_10_N | GLU_A_10_CA | 0.48 | 4.524 | -0.427 | -0.635 | 5.361 | -0.836 | 1.45 | 106.35 | 143.11 |
| 57 | 2jsa-13 | GLU_A_10_CA | GLU_A_10_HA | -0.635 | 5.361 | -0.836 | -1.162 | 5.684 | 0.062 | 1.09 | 34.54 | 148.50 |
| 58 | 2jsa-13 | GLU_A_10_CA | GLU_A_10_C | -0.635 | 5.361 | -0.836 | -1.607 | 4.561 | -1.705 | 1.53 | 124.62 | -140.54 |
| 59 | 2jsa-13 | GLU_A_10_C | GLU_A_10_O | -1.607 | 4.561 | -1.705 | -2.695 | 5.038 | -2.026 | 1.23 | 105.12 | 156.33 |
| 60 | 2jsa-13 | GLU_A_10_C | LEU_A_11_N | -1.607 | 4.561 | -1.705 | -1.181 | 3.358 | -2.061 | 1.32 | 105.59 | -70.50 |
| 61 | 2jsa-13 | LEU_A_11_N | LEU_A_11_H | -1.181 | 3.358 | -2.061 | -0.273 | 3.03 | -1.761 | 1.01 | 72.74 | -19.86 |
| 62 | 2jsa-13 | LEU_A_11_N | LEU_A_11_CA | -1.181 | 3.358 | -2.061 | -2.002 | 2.487 | -2.885 | 1.45 | 124.54 | -133.31 |
| 63 | 2jsa-13 | LEU_A_11_CA | LEU_A_11_HA | -2.002 | 2.487 | -2.885 | -2.667 | 3.113 | -3.479 | 1.09 | 123.04 | 136.73 |
| 64 | 2jsa-13 | LEU_A_11_CA | LEU_A_11_C | -2.002 | 2.487 | -2.885 | -2.872 | 1.61 | -1.984 | 1.53 | 53.89 | -134.77 |
| 65 | 2jsa-13 | LEU_A_11_C | LEU_A_11_O | -2.872 | 1.61 | -1.984 | -4.054 | 1.409 | -2.26 | 1.23 | 102.96 | -170.35 |
| 66 | 2jsa-13 | LEU_A_11_C | VAL_A_12_N | -2.872 | 1.61 | -1.984 | -2.254 | 1.108 | -0.924 | 1.33 | 36.91 | -39.09 |
| 67 | 2jsa-13 | VAL_A_12_N | VAL_A_12_H | -2.254 | 1.108 | -0.924 | -1.279 | 1.314 | -0.759 | 1.01 | 80.60 | 11.93 |
| 68 | 2jsa-13 | VAL_A_12_N | VAL_A_12_CA | -2.254 | 1.108 | -0.924 | -2.957 | 0.256 | 0.019 | 1.45 | 49.51 | -129.53 |
| 69 | 2jsa-13 | VAL_A_12_CA | VAL_A_12_HA | -2.957 | 0.256 | 0.019 | -3.485 | -0.515 | -0.541 | 1.09 | 120.93 | -124.40 |
| 70 | 2jsa-13 | VAL_A_12_CA | VAL_A_12_C | -2.957 | 0.256 | 0.019 | -3.994 | 1.088 | 0.776 | 1.53 | 60.34 | 141.26 |
| 71 | 2jsa-13 | VAL_A_12_C | VAL_A_12_O | -3.994 | 1.088 | 0.776 | -5.152 | 0.687 | 0.893 | 1.23 | 84.55 | -160.90 |
| 72 | 2jsa-13 | VAL_A_12_C | VAL_A_13_N | -3.994 | 1.088 | 0.776 | -3.542 | 2.231 | 1.271 | 1.33 | 68.06 | 68.42 |
| 73 | 2jsa-13 | VAL_A_13_N | VAL_A_13_H | -3.542 | 2.231 | 1.271 | -2.577 | 2.497 | 1.137 | 1.01 | 97.62 | 15.41 |
| 74 | 2jsa-13 | VAL_A_13_N | VAL_A_13_CA | -3.542 | 2.231 | 1.271 | -4.416 | 3.123 | 2.013 | 1.45 | 59.28 | 134.42 |
| 75 | 2jsa-13 | VAL_A_13_CA | VAL_A_13_HA | -4.416 | 3.123 | 2.013 | -4.841 | 2.57 | 2.851 | 1.09 | 39.77 | -127.54 |
| 76 | 2jsa-13 | VAL_A_13_CA | VAL_A_13_C | -4.416 | 3.123 | 2.013 | -5.566 | 3.572 | 1.109 | 1.53 | 126.21 | 158.67 |
| 77 | 2jsa-13 | VAL_A_13_C | VAL_A_13_O | -5.566 | 3.572 | 1.109 | -6.733 | 3.353 | 1.427 | 1.23 | 75.01 | -169.37 |
| 78 | 2jsa-13 | VAL_A_13_C | LYS_A_14_N | -5.566 | 3.572 | 1.109 | -5.194 | 4.195 | -0.001 | 1.33 | 146.83 | 59.16 |
| 79 | 2jsa-13 | LYS_A_14_N | LYS_A_14_H | -5.194 | 4.195 | -0.001 | -4.214 | 4.342 | -0.198 | 1.01 | 101.24 | 8.53 |
| 80 | 2jsa-13 | LYS_A_14_N | LYS_A_14_CA | -5.194 | 4.195 | -0.001 | -6.18 | 4.678 | -0.953 | 1.45 | 130.93 | 153.90 |
| 81 | 2jsa-13 | LYS_A_14_CA | LYS_A_14_HA | -6.18 | 4.678 | -0.953 | -6.663 | 5.556 | -0.523 | 1.09 | 66.78 | 118.82 |
| 82 | 2jsa-13 | LYS_A_14_CA | LYS_A_14_C | -6.18 | 4.678 | -0.953 | -7.254 | 3.608 | -1.155 | 1.53 | 97.59 | -135.11 |
| 83 | 2jsa-13 | LYS_A_14_C | LYS_A_14_O | -7.254 | 3.608 | -1.155 | -8.418 | 3.929 | -1.398 | 1.23 | 101.38 | 164.58 |
| 84 | 2jsa-13 | LYS_A_14_C | LYS_A_15_N | -7.254 | 3.608 | -1.155 | -6.829 | 2.359 | -1.047 | 1.32 | 85.32 | -71.21 |
| 85 | 2jsa-13 | LYS_A_15_N | LYS_A_15_H | -6.829 | 2.359 | -1.047 | -5.857 | 2.169 | -0.845 | 1.01 | 78.47 | -11.06 |
| 86 | 2jsa-13 | LYS_A_15_N | LYS_A_15_CA | -6.829 | 2.359 | -1.047 | -7.74 | 1.239 | -1.214 | 1.45 | 96.60 | -129.12 |
| 87 | 2jsa-13 | LYS_A_15_CA | LYS_A_15_HA | -7.74 | 1.239 | -1.214 | -8.468 | 1.508 | -1.979 | 1.09 | 134.59 | 159.72 |
| 88 | 2jsa-13 | LYS_A_15_CA | LYS_A_15_C | -7.74 | 1.239 | -1.214 | -8.503 | 1.009 | 0.092 | 1.53 | 31.39 | -163.23 |
| 89 | 2jsa-13 | LYS_A_15_C | LYS_A_15_O | -8.503 | 1.009 | 0.092 | -9.701 | 0.73 | 0.074 | 1.23 | 90.84 | -166.89 |
| 90 | 2jsa-13 | LYS_A_15_C | TYR_A_16_N | -8.503 | 1.009 | 0.092 | -7.777 | 1.132 | 1.193 | 1.32 | 33.77 | 9.62 |
| 91 | 2jsa-13 | TYR_A_16_N | TYR_A_16_H | -7.777 | 1.132 | 1.193 | -6.795 | 1.361 | 1.132 | 1.01 | 93.46 | 13.13 |
| 92 | 2jsa-13 | TYR_A_16_N | TYR_A_16_CA | -7.777 | 1.132 | 1.193 | -8.37 | 0.941 | 2.506 | 1.45 | 25.38 | -162.15 |
| 93 | 2jsa-13 | TYR_A_16_CA | TYR_A_16_HA | -8.37 | 0.941 | 2.506 | -8.937 | 0.011 | 2.532 | 1.09 | 88.63 | -121.37 |
| 94 | 2jsa-13 | TYR_A_16_CA | TYR_A_16_C | -8.37 | 0.941 | 2.506 | -9.281 | 2.114 | 2.872 | 1.53 | 76.16 | 127.83 |
| 95 | 2jsa-13 | TYR_A_16_C | TYR_A_16_O | -9.281 | 2.114 | 2.872 | -10.496 | 1.951 | 2.98 | 1.23 | 84.97 | -172.36 |
| 96 | 2jsa-13 | TYR_A_16_C | GLU_A_17_N | -9.281 | 2.114 | 2.872 | -8.659 | 3.27 | 3.054 | 1.33 | 82.11 | 61.72 |
| 97 | 2jsa-13 | GLU_A_17_N | GLU_A_17_H | -8.659 | 3.27 | 3.054 | -7.656 | 3.33 | 2.948 | 1.01 | 96.02 | 3.42 |
| 98 | 2jsa-13 | GLU_A_17_N | GLU_A_17_CA | -8.659 | 3.27 | 3.054 | -9.398 | 4.47 | 3.407 | 1.45 | 75.94 | 121.63 |
| 99 | 2jsa-13 | GLU_A_17_CA | GLU_A_17_HA | -9.398 | 4.47 | 3.407 | -9.737 | 4.367 | 4.438 | 1.09 | 18.97 | -163.10 |
| 100 | 2jsa-13 | GLU_A_17_CA | GLU_A_17_C | -9.398 | 4.47 | 3.407 | -10.627 | 4.619 | 2.508 | 1.53 | 125.99 | 173.09 |
| 101 | 2jsa-13 | GLU_A_17_C | GLU_A_17_O | -10.627 | 4.619 | 2.508 | -11.69 | 5.034 | 2.967 | 1.23 | 68.09 | 158.67 |
| 102 | 2jsa-13 | GLU_A_17_C | GLY_A_18_N | -10.627 | 4.619 | 2.508 | -10.441 | 4.273 | 1.242 | 1.33 | 162.76 | -61.74 |
| 103 | 2jsa-13 | GLY_A_18_N | GLY_A_18_H | -10.441 | 4.273 | 1.242 | -9.538 | 3.94 | 0.934 | 1.01 | 107.75 | -20.24 |
| 104 | 2jsa-13 | GLY_A_18_N | GLY_A_18_CA | -10.441 | 4.273 | 1.242 | -11.521 | 4.363 | 0.274 | 1.45 | 131.77 | 175.24 |
| 105 | 2jsa-13 | GLY_A_18_CA | GLY_A_18_C | -11.521 | 4.363 | 0.274 | -12.036 | 2.974 | -0.104 | 1.53 | 104.31 | -110.34 |
| 106 | 2jsa-13 | GLY_A_18_C | GLY_A_18_O | -12.036 | 2.974 | -0.104 | -12.384 | 2.73 | -1.259 | 1.23 | 159.80 | -144.96 |
| 107 | 2jsa-13 | GLY_A_18_C | SER_A_19_N | -12.036 | 2.974 | -0.104 | -12.069 | 2.099 | 0.891 | 1.33 | 41.35 | -92.16 |
| 108 | 2jsa-13 | SER_A_19_N | SER_A_19_H | -12.069 | 2.099 | 0.891 | -11.768 | 2.369 | 1.817 | 1.01 | 23.59 | 41.89 |
| 109 | 2jsa-13 | SER_A_19_N | SER_A_19_CA | -12.069 | 2.099 | 0.891 | -12.536 | 0.74 | 0.676 | 1.45 | 98.51 | -108.96 |
| 110 | 2jsa-13 | SER_A_19_CA | SER_A_19_HA | -12.536 | 0.74 | 0.676 | -12.295 | 0.454 | -0.348 | 1.09 | 159.94 | -49.88 |
| 111 | 2jsa-13 | SER_A_19_CA | SER_A_19_C | -12.536 | 0.74 | 0.676 | -14.053 | 0.673 | 0.86 | 1.53 | 83.09 | -177.47 |
| 112 | 2jsa-13 | SER_A_19_C | SER_A_19_O | -14.053 | 0.673 | 0.86 | -14.774 | 0.254 | -0.045 | 1.23 | 137.34 | -149.84 |
| 113 | 2jsa-13 | SER_A_19_C | ALA_A_20_N | -14.053 | 0.673 | 0.86 | -14.494 | 1.092 | 2.037 | 1.32 | 27.33 | 136.47 |
| 114 | 2jsa-13 | ALA_A_20_N | ALA_A_20_H | -14.494 | 1.092 | 2.037 | -13.843 | 1.424 | 2.735 | 1.01 | 46.31 | 27.02 |
| 115 | 2jsa-13 | ALA_A_20_N | ALA_A_20_CA | -14.494 | 1.092 | 2.037 | -15.912 | 1.084 | 2.351 | 1.45 | 77.51 | -179.68 |
| 116 | 2jsa-13 | ALA_A_20_CA | ALA_A_20_HA | -15.912 | 1.084 | 2.351 | -16.089 | 1.763 | 3.185 | 1.09 | 40.08 | 104.61 |
| 117 | 2jsa-13 | ALA_A_20_CA | ALA_A_20_C | -15.912 | 1.084 | 2.351 | -16.329 | -0.325 | 2.777 | 1.53 | 73.83 | -106.49 |
| 118 | 2jsa-13 | ALA_A_20_C | ALA_A_20_O | -16.329 | -0.325 | 2.777 | -17.487 | -0.558 | 3.12 | 1.23 | 73.81 | -168.62 |
| 119 | 2jsa-13 | ALA_A_20_C | ASP_A_21_N | -16.329 | -0.325 | 2.777 | -15.362 | -1.23 | 2.739 | 1.32 | 91.64 | -43.10 |
| 120 | 2jsa-13 | ASP_A_21_N | ASP_A_21_H | -15.362 | -1.23 | 2.739 | -14.431 | -0.969 | 2.445 | 1.01 | 106.91 | 15.66 |
| 121 | 2jsa-13 | ASP_A_21_N | ASP_A_21_CA | -15.362 | -1.23 | 2.739 | -15.614 | -2.611 | 3.116 | 1.45 | 74.97 | -100.34 |
| 122 | 2jsa-13 | ASP_A_21_CA | ASP_A_21_HA | -15.614 | -2.611 | 3.116 | -16.685 | -2.751 | 3.258 | 1.09 | 82.51 | -172.55 |
| 123 | 2jsa-13 | ASP_A_21_CA | ASP_A_21_C | -15.614 | -2.611 | 3.116 | -14.871 | -2.923 | 4.416 | 1.53 | 31.79 | -22.78 |
| 124 | 2jsa-13 | ASP_A_21_C | ASP_A_21_O | -14.871 | -2.923 | 4.416 | -15.16 | -3.92 | 5.077 | 1.23 | 57.51 | -106.17 |
| 125 | 2jsa-13 | ASP_A_21_C | LYS_A_22_N | -14.871 | -2.923 | 4.416 | -13.927 | -2.053 | 4.746 | 1.33 | 75.58 | 42.66 |
| 126 | 2jsa-13 | LYS_A_22_N | LYS_A_22_H | -13.927 | -2.053 | 4.746 | -13.742 | -1.255 | 4.155 | 1.01 | 125.81 | 76.95 |
| 127 | 2jsa-13 | LYS_A_22_N | LYS_A_22_CA | -13.927 | -2.053 | 4.746 | -13.141 | -2.223 | 5.956 | 1.45 | 33.61 | -12.20 |
| 128 | 2jsa-13 | LYS_A_22_CA | LYS_A_22_HA | -13.141 | -2.223 | 5.956 | -12.302 | -1.528 | 5.915 | 1.09 | 92.16 | 39.64 |
| 129 | 2jsa-13 | LYS_A_22_CA | LYS_A_22_C | -13.141 | -2.223 | 5.956 | -12.568 | -3.642 | 5.991 | 1.53 | 88.69 | -68.01 |
| 130 | 2jsa-13 | LYS_A_22_C | LYS_A_22_O | -12.568 | -3.642 | 5.991 | -12.798 | -4.385 | 6.944 | 1.23 | 39.22 | -107.20 |
| 131 | 2jsa-13 | LYS_A_22_C | ASP_A_23_N | -12.568 | -3.642 | 5.991 | -11.833 | -3.974 | 4.94 | 1.32 | 142.50 | -24.31 |

| | | | | | | | | | | | |
|---|---|---|---|---|---|---|---|---|---|---|---|
| 132 | 2jsa-13 | ASP_A_23_N | ASP_A_23_H | -11.833 | -3.974 | 4.94 | -11.686 | -3.311 | 4.192 | 1.01 | 137.76 | 77.50 |
| 133 | 2jsa-13 | ASP_A_23_N | ASP_A_23_CA | -11.833 | -3.974 | 4.94 | -11.225 | -5.29 | 4.838 | 1.45 | 94.02 | -65.20 |
| 134 | 2jsa-13 | ASP_A_23_CA | ASP_A_23_HA | -11.225 | -5.29 | 4.838 | -11.17 | -5.735 | 5.832 | 1.09 | 24.28 | -82.95 |
| 135 | 2jsa-13 | ASP_A_23_CA | ASP_A_23_C | -11.225 | -5.29 | 4.838 | -9.819 | -5.154 | 4.252 | 1.53 | 112.53 | 5.52 |
| 136 | 2jsa-13 | ASP_A_23_C | ASP_A_23_O | -9.819 | -5.154 | 4.252 | -9.661 | -4.785 | 3.089 | 1.23 | 160.96 | 66.82 |
| 137 | 2jsa-13 | ASP_A_23_C | ALA_A_24_N | -9.819 | -5.154 | 4.252 | -8.834 | -5.459 | 5.083 | 1.32 | 51.13 | -17.20 |
| 138 | 2jsa-13 | ALA_A_24_N | ALA_A_24_H | -8.834 | -5.459 | 5.083 | -9.038 | -5.755 | 6.027 | 1.01 | 20.85 | -124.57 |
| 139 | 2jsa-13 | ALA_A_24_N | ALA_A_24_CA | -8.834 | -5.459 | 5.083 | -7.446 | -5.375 | 4.662 | 1.45 | 106.84 | 3.46 |
| 140 | 2jsa-13 | ALA_A_24_CA | ALA_A_24_HA | -7.446 | -5.375 | 4.662 | -7.285 | -4.413 | 4.176 | 1.09 | 116.49 | 80.50 |
| 141 | 2jsa-13 | ALA_A_24_CA | ALA_A_24_C | -7.446 | -5.375 | 4.662 | -7.156 | -6.485 | 3.649 | 1.53 | 131.44 | -75.36 |
| 142 | 2jsa-13 | ALA_A_24_C | ALA_A_24_O | -7.156 | -6.485 | 3.649 | -6.065 | -6.545 | 3.083 | 1.23 | 117.38 | -3.15 |
| 143 | 2jsa-13 | ALA_A_24_C | ASN_A_25_N | -7.156 | -6.485 | 3.649 | -8.153 | -7.334 | 3.45 | 1.32 | 98.64 | -139.58 |
| 144 | 2jsa-13 | ASN_A_25_N | ASN_A_25_H | -8.153 | -7.334 | 3.45 | -9.023 | -7.222 | 3.95 | 1.01 | 60.32 | 172.66 |
| 145 | 2jsa-13 | ASN_A_25_N | ASN_A_25_CA | -8.153 | -7.334 | 3.45 | -8.02 | -8.439 | 2.515 | 1.45 | 130.03 | -83.14 |
| 146 | 2jsa-13 | ASN_A_25_CA | ASN_A_25_HA | -8.02 | -8.439 | 2.515 | -7.0 | -8.823 | 2.559 | 1.09 | 87.69 | -20.63 |
| 147 | 2jsa-13 | ASN_A_25_CA | ASN_A_25_C | -8.02 | -8.439 | 2.515 | -8.328 | -7.943 | 1.1 | 1.53 | 157.58 | 121.84 |
| 148 | 2jsa-13 | ASN_A_25_C | ASN_A_25_O | -8.328 | -7.943 | 1.1 | -7.546 | -8.164 | 0.177 | 1.23 | 138.64 | -15.78 |
| 149 | 2jsa-13 | ASN_A_25_C | VAL_A_26_N | -8.328 | -7.943 | 1.1 | -9.469 | -7.281 | 0.975 | 1.33 | 95.41 | 149.88 |
| 150 | 2jsa-13 | VAL_A_26_N | VAL_A_26_H | -9.469 | -7.281 | 0.975 | -10.063 | -7.134 | 1.779 | 1.01 | 37.27 | 166.10 |
| 151 | 2jsa-13 | VAL_A_26_N | VAL_A_26_CA | -9.469 | -7.281 | 0.975 | -9.89 | -6.754 | -0.311 | 1.45 | 152.32 | 128.62 |
| 152 | 2jsa-13 | VAL_A_26_CA | VAL_A_26_HA | -9.89 | -6.754 | -0.311 | -9.673 | -7.499 | -1.076 | 1.09 | 134.59 | -73.76 |
| 153 | 2jsa-13 | VAL_A_26_CA | VAL_A_26_C | -9.89 | -6.754 | -0.311 | -9.081 | -5.495 | -0.631 | 1.53 | 102.07 | 57.28 |
| 154 | 2jsa-13 | VAL_A_26_C | VAL_A_26_O | -9.081 | -5.495 | -0.631 | -9.122 | -4.994 | -1.754 | 1.23 | 155.89 | 94.68 |
| 155 | 2jsa-13 | VAL_A_26_C | ILE_A_27_N | -9.081 | -5.495 | -0.631 | -8.363 | -5.02 | 0.377 | 1.33 | 40.50 | 33.49 |
| 156 | 2jsa-13 | ILE_A_27_N | ILE_A_27_H | -8.363 | -5.02 | 0.377 | -8.376 | -5.484 | 1.274 | 1.01 | 27.36 | -91.60 |
| 157 | 2jsa-13 | ILE_A_27_N | ILE_A_27_CA | -8.363 | -5.02 | 0.377 | -7.544 | -3.83 | 0.218 | 1.45 | 96.28 | 55.46 |
| 158 | 2jsa-13 | ILE_A_27_CA | ILE_A_27_HA | -7.544 | -3.83 | 0.218 | -8.027 | -3.183 | -0.514 | 1.09 | 132.20 | 126.74 |
| 159 | 2jsa-13 | ILE_A_27_CA | ILE_A_27_C | -7.544 | -3.83 | 0.218 | -6.178 | -4.227 | -0.344 | 1.53 | 111.56 | -16.21 |
| 160 | 2jsa-13 | ILE_A_27_C | ILE_A_27_O | -6.178 | -4.227 | -0.344 | -5.899 | -4.007 | -1.521 | 1.23 | 163.20 | 38.26 |
| 161 | 2jsa-13 | ILE_A_27_C | LYS_A_28_N | -6.178 | -4.227 | -0.344 | -5.362 | -4.806 | 0.525 | 1.33 | 49.02 | -35.36 |
| 162 | 2jsa-13 | LYS_A_28_N | LYS_A_28_H | -5.362 | -4.806 | 0.525 | -5.658 | -4.956 | 1.479 | 1.01 | 19.18 | -153.13 |
| 163 | 2jsa-13 | LYS_A_28_N | LYS_A_28_CA | -5.362 | -4.806 | 0.525 | -4.032 | -5.236 | 0.13 | 1.45 | 105.78 | -17.92 |
| 164 | 2jsa-13 | LYS_A_28_CA | LYS_A_28_HA | -4.032 | -5.236 | 0.13 | -3.403 | -4.35 | 0.043 | 1.09 | 94.58 | 54.63 |
| 165 | 2jsa-13 | LYS_A_28_CA | LYS_A_28_C | -4.032 | -5.236 | 0.13 | -4.101 | -5.895 | -1.249 | 1.53 | 154.34 | -95.98 |
| 166 | 2jsa-13 | LYS_A_28_C | LYS_A_28_O | -4.101 | -5.895 | -1.249 | -3.189 | -5.743 | -2.061 | 1.23 | 131.29 | 9.46 |
| 167 | 2jsa-13 | LYS_A_28_C | LYS_A_29_N | -4.101 | -5.895 | -1.249 | -5.192 | -6.612 | -1.473 | 1.32 | 99.74 | -146.69 |
| 168 | 2jsa-13 | LYS_A_29_N | LYS_A_29_H | -5.192 | -6.612 | -1.473 | -5.902 | -6.693 | -0.759 | 1.01 | 45.02 | -173.49 |
| 169 | 2jsa-13 | LYS_A_29_N | LYS_A_29_CA | -5.192 | -6.612 | -1.473 | -5.393 | -7.294 | -2.74 | 1.45 | 150.70 | -106.42 |
| 170 | 2jsa-13 | LYS_A_29_CA | LYS_A_29_HA | -5.393 | -7.294 | -2.74 | -4.486 | -7.854 | -2.969 | 1.09 | 102.12 | -31.69 |
| 171 | 2jsa-13 | LYS_A_29_CA | LYS_A_29_C | -5.393 | -7.294 | -2.74 | -5.59 | -6.257 | -3.847 | 1.53 | 136.36 | 100.76 |
| 172 | 2jsa-13 | LYS_A_29_C | LYS_A_29_O | -5.59 | -6.257 | -3.847 | -4.971 | -6.35 | -4.906 | 1.23 | 149.41 | -8.54 |
| 173 | 2jsa-13 | LYS_A_29_C | ASP_A_30_N | -5.59 | -6.257 | -3.847 | -6.454 | -5.293 | -3.565 | 1.32 | 77.71 | 131.87 |
| 174 | 2jsa-13 | ASP_A_30_N | ASP_A_30_H | -6.454 | -5.293 | -3.565 | -6.928 | -5.28 | -2.673 | 1.01 | 27.99 | 178.43 |
| 175 | 2jsa-13 | ASP_A_30_N | ASP_A_30_CA | -6.454 | -5.293 | -3.565 | -6.739 | -4.24 | -4.523 | 1.45 | 131.29 | 105.14 |
| 176 | 2jsa-13 | ASP_A_30_CA | ASP_A_30_HA | -6.739 | -4.24 | -4.523 | -7.211 | -4.678 | -5.402 | 1.09 | 143.78 | -137.14 |
| 177 | 2jsa-13 | ASP_A_30_CA | ASP_A_30_C | -6.739 | -4.24 | -4.523 | -5.433 | -3.551 | -4.923 | 1.53 | 105.16 | 27.81 |
| 178 | 2jsa-13 | ASP_A_30_C | ASP_A_30_O | -5.433 | -3.551 | -4.923 | -5.174 | -3.341 | -6.107 | 1.23 | 164.27 | 39.04 |
| 179 | 2jsa-13 | ASP_A_30_C | PHE_A_31_N | -5.433 | -3.551 | -4.923 | -4.644 | -3.217 | -3.911 | 1.33 | 40.25 | 22.94 |
| 180 | 2jsa-13 | PHE_A_31_N | PHE_A_31_H | -4.644 | -3.217 | -3.911 | -4.921 | -3.419 | -2.961 | 1.01 | 19.84 | -143.90 |
| 181 | 2jsa-13 | PHE_A_31_N | PHE_A_31_CA | -4.644 | -3.217 | -3.911 | -3.371 | -2.557 | -4.142 | 1.45 | 99.15 | 27.40 |
| 182 | 2jsa-13 | PHE_A_31_CA | PHE_A_31_HA | -3.371 | -2.557 | -4.142 | -3.537 | -1.509 | -4.393 | 1.09 | 103.31 | 99.00 |
| 183 | 2jsa-13 | PHE_A_31_CA | PHE_A_31_C | -3.371 | -2.557 | -4.142 | -2.595 | -3.24 | -5.27 | 1.53 | 137.50 | -41.35 |
| 184 | 2jsa-13 | PHE_A_31_C | PHE_A_31_O | -2.595 | -3.24 | -5.27 | -2.202 | -2.592 | -6.238 | 1.23 | 141.94 | 58.76 |
| 185 | 2jsa-13 | PHE_A_31_C | ASP_A_32_N | -2.595 | -3.24 | -5.27 | -2.399 | -4.54 | -5.108 | 1.32 | 82.98 | -81.43 |
| 186 | 2jsa-13 | ASP_A_32_N | ASP_A_32_H | -2.399 | -4.54 | -5.108 | -2.751 | -5.009 | -4.286 | 1.01 | 35.50 | -126.89 |
| 187 | 2jsa-13 | ASP_A_32_N | ASP_A_32_CA | -2.399 | -4.54 | -5.108 | -1.677 | -5.318 | -6.101 | 1.45 | 133.09 | -47.14 |
| 188 | 2jsa-13 | ASP_A_32_CA | ASP_A_32_HA | -1.677 | -5.318 | -6.101 | -0.656 | -4.941 | -6.176 | 1.09 | 93.94 | 20.27 |
| 189 | 2jsa-13 | ASP_A_32_CA | ASP_A_32_C | -1.677 | -5.318 | -6.101 | -2.38 | -5.188 | -7.453 | 1.53 | 152.13 | 169.52 |
| 190 | 2jsa-13 | ASP_A_32_C | ASP_A_32_O | -2.38 | -5.188 | -7.453 | -1.76 | -5.37 | -8.5 | 1.23 | 148.32 | -16.36 |
| 191 | 2jsa-13 | ASP_A_32_C | ALA_A_33_N | -2.38 | -5.188 | -7.453 | -3.665 | -4.873 | -7.388 | 1.32 | 87.19 | 166.23 |
| 192 | 2jsa-13 | ALA_A_33_N | ALA_A_33_H | -3.665 | -4.873 | -7.388 | -4.11 | -4.737 | -6.492 | 1.01 | 27.44 | 163.01 |
| 193 | 2jsa-13 | ALA_A_33_N | ALA_A_33_CA | -3.665 | -4.873 | -7.388 | -4.46 | -4.718 | -8.595 | 1.45 | 146.14 | 168.97 |
| 194 | 2jsa-13 | ALA_A_33_CA | ALA_A_33_HA | -4.46 | -4.718 | -8.595 | -4.317 | -5.596 | -9.224 | 1.09 | 125.26 | -80.75 |
| 195 | 2jsa-13 | ALA_A_33_CA | ALA_A_33_C | -4.46 | -4.718 | -8.595 | -3.974 | -3.487 | -9.364 | 1.53 | 120.16 | 68.46 |
| 196 | 2jsa-13 | ALA_A_33_C | ALA_A_33_O | -3.974 | -3.487 | -9.364 | -3.321 | -3.614 | -10.398 | 1.23 | 147.24 | -11.01 |
| 197 | 2jsa-13 | ALA_A_33_C | GLU_A_34_N | -3.974 | -3.487 | -9.364 | -4.312 | -2.323 | -8.828 | 1.33 | 66.14 | 106.19 |
| 198 | 2jsa-13 | GLU_A_34_N | GLU_A_34_H | -4.312 | -2.323 | -8.828 | -4.85 | -2.293 | -7.974 | 1.01 | 32.25 | 176.81 |
| 199 | 2jsa-13 | GLU_A_34_N | GLU_A_34_CA | -4.312 | -2.323 | -8.828 | -3.919 | -1.071 | -9.451 | 1.45 | 115.40 | 72.57 |
| 200 | 2jsa-13 | GLU_A_34_CA | GLU_A_34_HA | -3.919 | -1.071 | -9.451 | -4.425 | -1.0 | -10.414 | 1.09 | 152.05 | 172.01 |
| 201 | 2jsa-13 | GLU_A_34_CA | GLU_A_34_C | -3.919 | -1.071 | -9.451 | -2.409 | -1.051 | -9.699 | 1.53 | 99.33 | 0.76 |
| 202 | 2jsa-13 | GLU_A_34_C | GLU_A_34_O | -2.409 | -1.051 | -9.699 | -1.939 | -0.414 | -10.639 | 1.23 | 139.90 | 53.58 |
| 203 | 2jsa-13 | GLU_A_34_C | CYS_A_35_N | -2.409 | -1.051 | -9.699 | -1.692 | -1.759 | -8.838 | 1.33 | 49.49 | -44.64 |
| 204 | 2jsa-13 | CYS_A_35_N | CYS_A_35_H | -1.692 | -1.759 | -8.838 | -2.146 | -2.26 | -8.088 | 1.01 | 42.03 | -132.18 |
| 205 | 2jsa-13 | CYS_A_35_N | CYS_A_35_CA | -1.692 | -1.759 | -8.838 | -0.245 | -1.832 | -8.951 | 1.45 | 94.46 | -2.89 |
| 206 | 2jsa-13 | CYS_A_35_CA | CYS_A_35_HA | -0.245 | -1.832 | -8.951 | 0.144 | -0.817 | -9.038 | 1.09 | 94.58 | 69.03 |
| 207 | 2jsa-13 | CYS_A_35_CA | CYS_A_35_C | -0.245 | -1.832 | -8.951 | 0.098 | -2.597 | -10.231 | 1.53 | 146.78 | -65.85 |
| 208 | 2jsa-13 | CYS_A_35_C | CYS_A_35_O | 0.098 | -2.597 | -10.231 | 0.837 | -2.097 | -11.078 | 1.23 | 133.51 | 34.08 |
| 209 | 2jsa-13 | CYS_A_35_C | LYS_A_36_N | 0.098 | -2.597 | -10.231 | -0.454 | -3.797 | -10.332 | 1.32 | 94.37 | -114.70 |
| 210 | 2jsa-13 | LYS_A_36_N | LYS_A_36_H | -0.454 | -3.797 | -10.332 | -1.052 | -4.145 | -9.596 | 1.01 | 43.23 | -149.80 |
| 211 | 2jsa-13 | LYS_A_36_N | LYS_A_36_CA | -0.454 | -3.797 | -10.332 | -0.215 | -4.636 | -11.493 | 1.45 | 143.08 | -74.10 |
| 212 | 2jsa-13 | LYS_A_36_CA | LYS_A_36_HA | -0.215 | -4.636 | -11.493 | 0.837 | -4.922 | -11.492 | 1.09 | 89.95 | -15.21 |
| 213 | 2jsa-13 | LYS_A_36_CA | LYS_A_36_C | -0.215 | -4.636 | -11.493 | -0.477 | -3.825 | -12.765 | 1.53 | 146.18 | 107.90 |
| 214 | 2jsa-13 | LYS_A_36_C | LYS_A_36_O | -0.477 | -3.825 | -12.765 | -0.042 | -4.208 | -13.849 | 1.23 | 151.87 | -41.36 |
| 215 | 2jsa-13 | LYS_A_36_C | LYS_A_37_N | -0.477 | -3.825 | -12.765 | -1.188 | -2.721 | -12.588 | 1.33 | 82.32 | 122.78 |
| 216 | 2jsa-13 | LYS_A_37_N | LYS_A_37_H | -1.188 | -2.721 | -12.588 | -1.514 | -2.468 | -11.666 | 1.01 | 24.11 | 142.19 |
| 217 | 2jsa-13 | LYS_A_37_N | LYS_A_37_CA | -1.188 | -2.721 | -12.588 | -1.513 | -1.854 | -13.708 | 1.45 | 140.42 | 110.55 |
| 218 | 2jsa-13 | LYS_A_37_CA | LYS_A_37_HA | -1.513 | -1.854 | -13.708 | -1.319 | -2.404 | -14.629 | 1.09 | 147.66 | -70.57 |
| 219 | 2jsa-13 | LYS_A_37_CA | LYS_A_37_C | -1.513 | -1.854 | -13.708 | -0.586 | -0.636 | -13.69 | 1.53 | 89.33 | 52.73 |
| 220 | 2jsa-13 | LYS_A_37_C | LYS_A_37_O | -0.586 | -0.636 | -13.69 | 0.262 | -0.484 | -14.567 | 1.23 | 135.51 | 10.16 |
| 221 | 2jsa-13 | LYS_A_37_C | LEU_A_38_N | -0.586 | -0.636 | -13.69 | -0.78 | 0.2 | -12.681 | 1.32 | 40.38 | 103.06 |
| 222 | 2jsa-13 | LEU_A_38_N | LEU_A_38_H | -0.78 | 0.2 | -12.681 | -1.497 | 0.012 | -11.995 | 1.01 | 47.22 | -165.31 |
| 223 | 2jsa-13 | LEU_A_38_N | LEU_A_38_CA | -0.78 | 0.2 | -12.681 | 0.029 | 1.4 | -12.537 | 1.45 | 84.32 | 56.01 |
| 224 | 2jsa-13 | LEU_A_38_CA | LEU_A_38_HA | 0.029 | 1.4 | -12.537 | -0.153 | 2.036 | -13.403 | 1.09 | 142.62 | 105.97 |
| 225 | 2jsa-13 | LEU_A_38_CA | LEU_A_38_C | 0.029 | 1.4 | -12.537 | 1.509 | 1.016 | -12.546 | 1.53 | 90.34 | -14.55 |
| 226 | 2jsa-13 | LEU_A_38_C | LEU_A_38_O | 1.509 | 1.016 | -12.546 | 2.294 | 1.572 | -13.314 | 1.23 | 128.60 | 35.31 |
| 227 | 2jsa-13 | LEU_A_38_C | PHE_A_39_N | 1.509 | 1.016 | -12.546 | 1.847 | 0.067 | -11.685 | 1.33 | 49.48 | -70.40 |

| | | | | | | | | | | |
|---|---|---|---|---|---|---|---|---|---|---|
| 228 | 2jsa-13 | PHE_A_39_N | PHE_A_39_H | 1.847 | 0.067 | -11.685 | 1.149 | -0.347 | -11.084 | 1.01 | 53.48 | -149.33 |
| 229 | 2jsa-13 | PHE_A_39_N | PHE_A_39_CA | 1.847 | 0.067 | -11.685 | 3.22 | -0.397 | -11.585 | 1.45 | 86.05 | -18.67 |
| 230 | 2jsa-13 | PHE_A_39_CA | PHE_A_39_HA | 3.22 | -0.397 | -11.585 | 3.905 | 0.426 | -11.79 | 1.09 | 100.84 | 50.23 |
| 231 | 2jsa-13 | PHE_A_39_CA | PHE_A_39_C | 3.22 | -0.397 | -11.585 | 3.484 | -1.543 | -12.563 | 1.53 | 129.75 | -77.03 |
| 232 | 2jsa-13 | PHE_A_39_C | PHE_A_39_O | 3.484 | -1.543 | -12.563 | 4.227 | -2.472 | -12.251 | 1.23 | 75.30 | -51.35 |
| 233 | 2jsa-13 | PHE_A_39_C | HIS_A_40_N | 3.484 | -1.543 | -12.563 | 2.862 | -1.439 | -13.728 | 1.32 | 151.57 | 170.51 |
| 234 | 2jsa-13 | HIS_A_40_N | HIS_A_40_H | 2.862 | -1.439 | -13.728 | 2.266 | -0.646 | -13.917 | 1.01 | 100.79 | 126.93 |
| 235 | 2jsa-13 | HIS_A_40_N | HIS_A_40_CA | 2.862 | -1.439 | -13.728 | 3.021 | -2.455 | -14.755 | 1.45 | 134.96 | -81.11 |
| 236 | 2jsa-13 | HIS_A_40_CA | HIS_A_40_HA | 3.021 | -2.455 | -14.755 | 3.281 | -3.393 | -14.263 | 1.09 | 63.19 | -74.51 |
| 237 | 2jsa-13 | HIS_A_40_CA | HIS_A_40_C | 3.021 | -2.455 | -14.755 | 4.179 | -2.074 | -15.679 | 1.53 | 127.16 | 18.21 |
| 238 | 2jsa-13 | HIS_A_40_C | HIS_A_40_O | 4.179 | -2.074 | -15.679 | 3.961 | -1.71 | -16.835 | 1.23 | 159.85 | 120.92 |
| 239 | 2jsa-13 | HIS_A_40_C | THR_A_41_N | 4.179 | -2.074 | -15.679 | 5.384 | -2.169 | -15.138 | 1.32 | 65.89 | -4.51 |
| 240 | 2jsa-13 | THR_A_41_N | THR_A_41_H | 5.384 | -2.169 | -15.138 | 5.489 | -2.475 | -14.181 | 1.01 | 18.68 | -71.06 |
| 241 | 2jsa-13 | THR_A_41_N | THR_A_41_CA | 5.384 | -2.169 | -15.138 | 6.576 | -1.839 | -15.899 | 1.45 | 121.60 | 15.47 |
| 242 | 2jsa-13 | THR_A_41_CA | THR_A_41_HA | 6.576 | -1.839 | -15.899 | 6.799 | -2.664 | -16.575 | 1.09 | 128.34 | -74.87 |
| 243 | 2jsa-13 | THR_A_41_CA | THR_A_41_C | 6.576 | -1.839 | -15.899 | 7.774 | -1.66 | -14.965 | 1.53 | 52.37 | 8.50 |
| 244 | 2jsa-13 | THR_A_41_C | THR_A_41_O | 7.774 | -1.66 | -14.965 | 8.881 | -2.095 | -15.278 | 1.23 | 104.74 | -21.45 |
| 245 | 2jsa-13 | THR_A_41_C | ILE_A_42_N | 7.774 | -1.66 | -14.965 | 7.511 | -1.019 | -13.835 | 1.33 | 31.51 | 112.31 |
| 246 | 2jsa-13 | ILE_A_42_N | ILE_A_42_H | 7.511 | -1.019 | -13.835 | 6.576 | -0.69 | -13.642 | 1.01 | 78.98 | 160.61 |
| 247 | 2jsa-13 | ILE_A_42_N | ILE_A_42_CA | 7.511 | -1.019 | -13.835 | 8.555 | -0.777 | -12.852 | 1.45 | 47.47 | 13.05 |
| 248 | 2jsa-13 | ILE_A_42_CA | ILE_A_42_HA | 8.555 | -0.777 | -12.852 | 9.172 | 0.048 | -13.21 | 1.09 | 109.16 | 53.21 |
| 249 | 2jsa-13 | ILE_A_42_CA | ILE_A_42_C | 8.555 | -0.777 | -12.852 | 9.448 | -2.015 | -12.748 | 1.53 | 86.10 | -54.20 |
| 250 | 2jsa-13 | ILE_A_42_C | ILE_A_42_O | 9.448 | -2.015 | -12.748 | 8.965 | -3.143 | -12.834 | 1.23 | 94.01 | -113.18 |
| 251 | 2jsa-13 | ILE_A_42_C | PRO_A_43_N | 9.448 | -2.015 | -12.748 | 10.77 | -1.755 | -12.561 | 1.36 | 82.10 | 11.13 |
| 252 | 2jsa-13 | PRO_A_43_N | PRO_A_43_CA | 10.77 | -1.755 | -12.561 | 11.735 | -2.834 | -12.444 | 1.45 | 85.38 | -48.19 |
| 253 | 2jsa-13 | PRO_A_43_CA | PRO_A_43_HA | 11.735 | -2.834 | -12.444 | 11.552 | -3.583 | -13.214 | 1.09 | 134.96 | -103.73 |
| 254 | 2jsa-13 | PRO_A_43_CA | PRO_A_43_C | 11.735 | -2.834 | -12.444 | 11.638 | -3.511 | -11.076 | 1.53 | 26.56 | -98.15 |
| 255 | 2jsa-13 | PRO_A_43_C | PRO_A_43_O | 11.638 | -3.511 | -11.076 | 12.614 | -4.082 | -10.591 | 1.23 | 66.78 | -30.33 |
| 256 | 2jsa-13 | PRO_A_43_C | PHE_A_44_N | 11.638 | -3.511 | -11.076 | 10.452 | -3.426 | -10.491 | 1.33 | 63.80 | 175.90 |
| 257 | 2jsa-13 | PHE_A_44_N | PHE_A_44_H | 10.452 | -3.426 | -10.491 | 9.694 | -2.941 | -10.948 | 1.01 | 116.92 | 147.39 |
| 258 | 2jsa-13 | PHE_A_44_N | PHE_A_44_CA | 10.452 | -3.426 | -10.491 | 10.215 | -4.024 | -9.188 | 1.45 | 26.27 | -111.62 |
| 259 | 2jsa-13 | PHE_A_44_CA | PHE_A_44_HA | 10.215 | -4.024 | -9.188 | 10.108 | -5.104 | -9.29 | 1.09 | 95.37 | -95.66 |
| 260 | 2jsa-13 | PHE_A_44_CA | PHE_A_44_C | 10.215 | -4.024 | -9.188 | 8.959 | -3.44 | -8.538 | 1.53 | 64.86 | 155.06 |
| 261 | 2jsa-13 | PHE_A_44_C | PHE_A_44_O | 8.959 | -3.44 | -8.538 | 8.937 | -3.193 | -7.333 | 1.23 | 11.63 | 95.09 |
| 262 | 2jsa-13 | PHE_A_44_C | GLY_A_45_N | 8.959 | -3.44 | -8.538 | 7.943 | -3.237 | -9.364 | 1.33 | 128.56 | 168.70 |
| 263 | 2jsa-13 | GLY_A_45_N | GLY_A_45_H | 7.943 | -3.237 | -9.364 | 8.032 | -3.464 | -10.344 | 1.01 | 166.03 | -68.59 |
| 264 | 2jsa-13 | GLY_A_45_N | GLY_A_45_CA | 7.943 | -3.237 | -9.364 | 6.687 | -2.688 | -8.885 | 1.45 | 70.74 | 156.39 |
| 265 | 2jsa-13 | GLY_A_45_CA | GLY_A_45_C | 6.687 | -2.688 | -8.885 | 5.689 | -3.802 | -8.56 | 1.53 | 77.74 | -131.86 |
| 266 | 2jsa-13 | GLY_A_45_C | GLY_A_45_O | 5.689 | -3.802 | -8.56 | 5.276 | -3.955 | -7.412 | 1.23 | 20.99 | -159.67 |
| 267 | 2jsa-13 | GLY_A_45_C | THR_A_46_N | 5.689 | -3.802 | -8.56 | 5.332 | -4.552 | -9.593 | 1.33 | 141.20 | -115.45 |
| 268 | 2jsa-13 | THR_A_46_N | THR_A_46_H | 5.332 | -4.552 | -9.593 | 5.714 | -4.369 | -10.51 | 1.01 | 155.21 | 25.60 |
| 269 | 2jsa-13 | THR_A_46_N | THR_A_46_CA | 5.332 | -4.552 | -9.593 | 4.391 | -5.648 | -9.432 | 1.45 | 83.64 | -130.65 |
| 270 | 2jsa-13 | THR_A_46_CA | THR_A_46_HA | 4.391 | -5.648 | -9.432 | 3.47 | -5.257 | -9.0 | 1.09 | 66.65 | 157.00 |
| 271 | 2jsa-13 | THR_A_46_CA | THR_A_46_C | 4.391 | -5.648 | -9.432 | 4.953 | -6.697 | -8.472 | 1.53 | 51.11 | -61.82 |
| 272 | 2jsa-13 | THR_A_46_C | THR_A_46_O | 4.953 | -6.697 | -8.472 | 4.199 | -7.374 | -7.775 | 1.23 | 55.48 | -138.08 |
| 273 | 2jsa-13 | THR_A_46_C | ARG_A_47_N | 4.953 | -6.697 | -8.472 | 6.275 | -6.799 | -8.465 | 1.33 | 89.70 | -4.41 |
| 274 | 2jsa-13 | ARG_A_47_N | ARG_A_47_H | 6.275 | -6.799 | -8.465 | 6.834 | -6.209 | -9.065 | 1.01 | 126.44 | 46.55 |
| 275 | 2jsa-13 | ARG_A_47_N | ARG_A_47_CA | 6.275 | -6.799 | -8.465 | 6.947 | -7.754 | -7.602 | 1.45 | 53.53 | -54.87 |
| 276 | 2jsa-13 | ARG_A_47_CA | ARG_A_47_HA | 6.947 | -7.754 | -7.602 | 6.446 | -8.717 | -7.699 | 1.09 | 95.11 | -117.49 |
| 277 | 2jsa-13 | ARG_A_47_CA | ARG_A_47_C | 6.947 | -7.754 | -7.602 | 6.871 | -7.298 | -6.143 | 1.53 | 17.58 | 99.46 |
| 278 | 2jsa-13 | ARG_A_47_C | ARG_A_47_O | 6.871 | -7.298 | -6.143 | 6.211 | -7.935 | -5.323 | 1.23 | 48.20 | -136.02 |
| 279 | 2jsa-13 | ARG_A_47_C | GLU_A_48_N | 6.871 | -7.298 | -6.143 | 7.554 | -6.197 | -5.864 | 1.33 | 77.85 | 58.19 |
| 280 | 2jsa-13 | GLU_A_48_N | GLU_A_48_H | 7.554 | -6.197 | -5.864 | 8.073 | -5.723 | -6.59 | 1.01 | 135.93 | 42.41 |
| 281 | 2jsa-13 | GLU_A_48_N | GLU_A_48_CA | 7.554 | -6.197 | -5.864 | 7.573 | -5.648 | -4.52 | 1.45 | 22.23 | 88.02 |
| 282 | 2jsa-13 | GLU_A_48_CA | GLU_A_48_HA | 7.573 | -5.648 | -4.52 | 8.091 | -6.357 | -3.873 | 1.09 | 53.62 | -53.85 |
| 283 | 2jsa-13 | GLU_A_48_CA | GLU_A_48_C | 7.573 | -5.648 | -4.52 | 6.144 | -5.474 | -3.998 | 1.53 | 70.07 | 173.06 |
| 284 | 2jsa-13 | GLU_A_48_C | GLU_A_48_O | 6.144 | -5.474 | -3.998 | 5.912 | -5.506 | -2.79 | 1.23 | 10.97 | -172.15 |
| 285 | 2jsa-13 | GLU_A_48_C | CYS_A_49_N | 6.144 | -5.474 | -3.998 | 5.225 | -5.293 | -4.934 | 1.32 | 134.98 | 168.86 |
| 286 | 2jsa-13 | CYS_A_49_N | CYS_A_49_H | 5.225 | -5.293 | -4.934 | 5.489 | -5.278 | -5.909 | 1.01 | 164.83 | 3.25 |
| 287 | 2jsa-13 | CYS_A_49_N | CYS_A_49_CA | 5.225 | -5.293 | -4.934 | 3.826 | -5.114 | -4.585 | 1.45 | 76.10 | 172.71 |
| 288 | 2jsa-13 | CYS_A_49_CA | CYS_A_49_HA | 3.826 | -5.114 | -4.585 | 3.751 | -4.276 | -3.891 | 1.09 | 50.48 | 95.11 |
| 289 | 2jsa-13 | CYS_A_49_CA | CYS_A_49_C | 3.826 | -5.114 | -4.585 | 3.342 | -6.38 | -3.874 | 1.53 | 62.32 | -110.92 |
| 290 | 2jsa-13 | CYS_A_49_C | CYS_A_49_O | 3.342 | -6.38 | -3.874 | 2.827 | -6.311 | -2.759 | 1.23 | 24.99 | 172.37 |
| 291 | 2jsa-13 | CYS_A_49_C | ASP_A_50_N | 3.342 | -6.38 | -3.874 | 3.525 | -7.505 | -4.549 | 1.32 | 120.63 | -80.76 |
| 292 | 2jsa-13 | ASP_A_50_N | ASP_A_50_H | 3.525 | -7.505 | -4.549 | 3.955 | -7.486 | -5.463 | 1.01 | 154.78 | 2.53 |
| 293 | 2jsa-13 | ASP_A_50_N | ASP_A_50_CA | 3.525 | -7.505 | -4.549 | 3.114 | -8.785 | -3.995 | 1.45 | 67.60 | -107.80 |
| 294 | 2jsa-13 | ASP_A_50_CA | ASP_A_50_HA | 3.114 | -8.785 | -3.995 | 2.033 | -8.782 | -3.855 | 1.09 | 82.62 | 179.84 |
| 295 | 2jsa-13 | ASP_A_50_CA | ASP_A_50_C | 3.114 | -8.785 | -3.995 | 3.811 | -9.001 | -2.651 | 1.53 | 28.50 | -17.22 |
| 296 | 2jsa-13 | ASP_A_50_C | ASP_A_50_O | 3.811 | -9.001 | -2.651 | 3.389 | -9.842 | -1.857 | 1.23 | 49.84 | -116.65 |
| 297 | 2jsa-13 | ASP_A_50_C | HIS_A_51_N | 3.811 | -9.001 | -2.651 | 4.867 | -8.23 | -2.436 | 1.33 | 80.66 | 36.13 |
| 298 | 2jsa-13 | HIS_A_51_N | HIS_A_51_H | 4.867 | -8.23 | -2.436 | 5.159 | -7.56 | -3.134 | 1.01 | 133.68 | 66.45 |
| 299 | 2jsa-13 | HIS_A_51_N | HIS_A_51_CA | 4.867 | -8.23 | -2.436 | 5.626 | -8.328 | -1.202 | 1.45 | 31.81 | -7.36 |
| 300 | 2jsa-13 | HIS_A_51_CA | HIS_A_51_HA | 5.626 | -8.328 | -1.202 | 5.309 | -9.232 | -0.682 | 1.09 | 61.51 | -109.32 |
| 301 | 2jsa-13 | HIS_A_51_CA | HIS_A_51_C | 5.626 | -8.328 | -1.202 | 5.303 | -7.131 | -0.305 | 1.53 | 54.11 | 105.10 |
| 302 | 2jsa-13 | HIS_A_51_C | HIS_A_51_O | 5.303 | -7.131 | -0.305 | 5.909 | -6.961 | 0.752 | 1.23 | 30.77 | 15.67 |
| 303 | 2jsa-13 | HIS_A_51_C | TYR_A_52_N | 5.303 | -7.131 | -0.305 | 4.35 | -6.331 | -0.76 | 1.32 | 110.09 | 139.99 |
| 304 | 2jsa-13 | TYR_A_52_N | TYR_A_52_H | 4.35 | -6.331 | -0.76 | 3.896 | -6.531 | -1.639 | 1.01 | 150.56 | -156.23 |
| 305 | 2jsa-13 | TYR_A_52_N | TYR_A_52_CA | 4.35 | -6.331 | -0.76 | 3.94 | -5.155 | -0.012 | 1.45 | 59.01 | 109.22 |
| 306 | 2jsa-13 | TYR_A_52_CA | TYR_A_52_HA | 3.94 | -5.155 | -0.012 | 4.701 | -4.891 | 0.723 | 1.09 | 47.62 | 19.13 |
| 307 | 2jsa-13 | TYR_A_52_CA | TYR_A_52_C | 3.94 | -5.155 | -0.012 | 2.599 | -5.392 | 0.688 | 1.53 | 62.80 | -169.98 |
| 308 | 2jsa-13 | TYR_A_52_C | TYR_A_52_O | 2.599 | -5.392 | 0.688 | 2.477 | -5.169 | 1.891 | 1.23 | 11.93 | 118.68 |
| 309 | 2jsa-13 | TYR_A_52_C | VAL_A_53_N | 2.599 | -5.392 | 0.688 | 1.629 | -5.838 | -0.095 | 1.32 | 126.26 | -155.31 |
| 310 | 2jsa-13 | VAL_A_53_N | VAL_A_53_H | 1.629 | -5.838 | -0.095 | 1.802 | -5.998 | -1.077 | 1.01 | 166.51 | -42.76 |
| 311 | 2jsa-13 | VAL_A_53_N | VAL_A_53_CA | 1.629 | -5.838 | -0.095 | 0.303 | -6.106 | 0.434 | 1.45 | 68.64 | -168.57 |
| 312 | 2jsa-13 | VAL_A_53_CA | VAL_A_53_HA | 0.303 | -6.106 | 0.434 | -0.153 | -5.157 | 0.717 | 1.09 | 74.96 | 115.66 |
| 313 | 2jsa-13 | VAL_A_53_CA | VAL_A_53_C | 0.303 | -6.106 | 0.434 | 0.426 | -6.968 | 1.692 | 1.53 | 34.69 | -81.88 |
| 314 | 2jsa-13 | VAL_A_53_C | VAL_A_53_O | 0.426 | -6.968 | 1.692 | -0.346 | -6.806 | 2.636 | 1.23 | 39.88 | 168.15 |
| 315 | 2jsa-13 | VAL_A_53_C | ASN_A_54_N | 0.426 | -6.968 | 1.692 | 1.401 | -7.864 | 1.664 | 1.32 | 91.21 | -42.58 |
| 316 | 2jsa-13 | ASN_A_54_N | ASN_A_54_H | 1.401 | -7.864 | 1.664 | 1.998 | -7.941 | 0.853 | 1.01 | 143.42 | -7.35 |
| 317 | 2jsa-13 | ASN_A_54_N | ASN_A_54_CA | 1.401 | -7.864 | 1.664 | 1.635 | -8.751 | 2.791 | 1.45 | 39.14 | -75.22 |
| 318 | 2jsa-13 | ASN_A_54_CA | ASN_A_54_HA | 1.635 | -8.751 | 2.791 | 0.769 | -8.725 | 3.452 | 1.09 | 52.66 | 178.28 |
| 319 | 2jsa-13 | ASN_A_54_CA | ASN_A_54_C | 1.635 | -8.751 | 2.791 | 2.88 | -8.287 | 3.549 | 1.53 | 60.30 | 20.44 |
| 320 | 2jsa-13 | ASN_A_54_C | ASN_A_54_O | 2.88 | -8.287 | 3.549 | 3.602 | -9.104 | 4.121 | 1.23 | 62.32 | -48.53 |
| 321 | 2jsa-13 | ASN_A_54_C | SER_A_55_N | 2.88 | -8.287 | 3.549 | 3.094 | -6.981 | 3.529 | 1.32 | 90.87 | 80.69 |
| 322 | 2jsa-13 | SER_A_55_N | SER_A_55_H | 3.094 | -6.981 | 3.529 | 2.458 | -6.369 | 3.038 | 1.01 | 119.09 | 136.10 |
| 323 | 2jsa-13 | SER_A_55_N | SER_A_55_CA | 3.094 | -6.981 | 3.529 | 4.24 | -6.399 | 4.208 | 1.45 | 62.15 | 26.92 |

| | | | | | | | | | | | |
|---|---|---|---|---|---|---|---|---|---|---|---|
| 324 | 2jsa-13 | SER_A_55_CA | SER_A_55_HA | 4.24 | -6.399 | 4.208 | 4.206 | -6.699 | 5.255 | 1.09 | 16.09 | -96.47 |
| 325 | 2jsa-13 | SER_A_55_CA | SER_A_55_C | 4.24 | -6.399 | 4.208 | 4.173 | -4.872 | 4.134 | 1.53 | 92.77 | 92.51 |
| 326 | 2jsa-13 | SER_A_55_C | SER_A_55_O | 4.173 | -4.872 | 4.134 | 4.555 | -4.279 | 3.127 | 1.23 | 144.99 | 57.21 |
| 327 | 2jsa-13 | SER_A_55_C | LYS_A_56_N | 4.173 | -4.872 | 4.134 | 3.684 | -4.28 | 5.213 | 1.32 | 35.44 | 129.56 |
| 328 | 2jsa-13 | LYS_A_56_N | LYS_A_56_H | 3.684 | -4.28 | 5.213 | 3.388 | -4.831 | 6.006 | 1.01 | 38.26 | -118.24 |
| 329 | 2jsa-13 | LYS_A_56_N | LYS_A_56_CA | 3.684 | -4.28 | 5.213 | 3.562 | -2.833 | 5.283 | 1.45 | 87.24 | 94.82 |
| 330 | 2jsa-13 | LYS_A_56_CA | LYS_A_56_HA | 3.562 | -2.833 | 5.283 | 3.842 | -2.518 | 6.289 | 1.09 | 22.73 | 48.37 |
| 331 | 2jsa-13 | LYS_A_56_CA | LYS_A_56_C | 3.562 | -2.833 | 5.283 | 2.101 | -2.438 | 5.062 | 1.53 | 98.31 | 164.87 |
| 332 | 2jsa-13 | LYS_A_56_C | LYS_A_56_O | 2.101 | -2.438 | 5.062 | 1.766 | -1.255 | 5.071 | 1.23 | 89.58 | 105.81 |
| 333 | 2jsa-13 | LYS_A_56_C | VAL_A_57_N | 2.101 | -2.438 | 5.062 | 1.27 | -3.451 | 4.868 | 1.32 | 98.42 | -129.36 |
| 334 | 2jsa-13 | VAL_A_57_N | VAL_A_57_H | 1.27 | -3.451 | 4.868 | 1.616 | -4.4 | 4.874 | 1.01 | 89.66 | -69.97 |
| 335 | 2jsa-13 | VAL_A_57_N | VAL_A_57_CA | 1.27 | -3.451 | 4.868 | -0.148 | -3.225 | 4.644 | 1.45 | 98.87 | 170.94 |
| 336 | 2jsa-13 | VAL_A_57_CA | VAL_A_57_HA | -0.148 | -3.225 | 4.644 | -0.286 | -2.19 | 4.331 | 1.09 | 106.69 | 97.59 |
| 337 | 2jsa-13 | VAL_A_57_CA | VAL_A_57_C | -0.148 | -3.225 | 4.644 | -0.906 | -3.429 | 5.956 | 1.53 | 30.89 | -164.94 |
| 338 | 2jsa-13 | VAL_A_57_C | VAL_A_57_O | -0.906 | -3.429 | 5.956 | -2.013 | -2.918 | 6.124 | 1.23 | 82.15 | 155.22 |
| 339 | 2jsa-13 | VAL_A_57_C | ASP_A_58_N | -0.906 | -3.429 | 5.956 | -0.281 | -4.176 | 6.855 | 1.33 | 47.29 | -50.08 |
| 340 | 2jsa-13 | ASP_A_58_N | ASP_A_58_H | -0.281 | -4.176 | 6.855 | 0.629 | -4.565 | 6.65 | 1.01 | 101.70 | -23.15 |
| 341 | 2jsa-13 | ASP_A_58_N | ASP_A_58_CA | -0.281 | -4.176 | 6.855 | -0.883 | -4.453 | 8.147 | 1.45 | 27.15 | -155.29 |
| 342 | 2jsa-13 | ASP_A_58_CA | ASP_A_58_HA | -0.883 | -4.453 | 8.147 | -1.942 | -4.667 | 8.008 | 1.09 | 97.33 | -168.58 |
| 343 | 2jsa-13 | ASP_A_58_CA | ASP_A_58_C | -0.883 | -4.453 | 8.147 | -0.713 | -3.234 | 9.057 | 1.53 | 53.52 | 82.06 |
| 344 | 2jsa-13 | ASP_A_58_C | ASP_A_58_O | -0.713 | -3.234 | 9.057 | -1.639 | -2.855 | 9.77 | 1.23 | 54.53 | 157.74 |
| 345 | 2jsa-13 | ASP_A_58_C | PRO_A_59_N | -0.713 | -3.234 | 9.057 | 0.509 | -2.64 | 8.998 | 1.36 | 92.49 | 25.92 |
| 346 | 2jsa-13 | PRO_A_59_N | PRO_A_59_CA | 0.509 | -2.64 | 8.998 | 0.811 | -1.472 | 9.808 | 1.45 | 56.12 | 75.50 |
| 347 | 2jsa-13 | PRO_A_59_CA | PRO_A_59_HA | 0.811 | -1.472 | 9.808 | 0.457 | -1.623 | 10.828 | 1.09 | 20.67 | -156.90 |
| 348 | 2jsa-13 | PRO_A_59_CA | PRO_A_59_C | 0.811 | -1.472 | 9.808 | 0.138 | -0.221 | 9.24 | 1.53 | 111.79 | 118.28 |
| 349 | 2jsa-13 | PRO_A_59_C | PRO_A_59_O | 0.138 | -0.221 | 9.24 | -0.559 | 0.493 | 9.958 | 1.23 | 54.26 | 134.31 |
| 350 | 2jsa-13 | PRO_A_59_C | ILE_A_60_N | 0.138 | -0.221 | 9.24 | 0.369 | 0.005 | 7.955 | 1.33 | 165.88 | 44.37 |
| 351 | 2jsa-13 | ILE_A_60_N | ILE_A_60_H | 0.369 | 0.005 | 7.955 | 0.953 | -0.626 | 7.425 | 1.01 | 121.65 | -47.22 |
| 352 | 2jsa-13 | ILE_A_60_N | ILE_A_60_CA | 0.369 | 0.005 | 7.955 | -0.207 | 1.156 | 7.282 | 1.45 | 117.60 | 116.58 |
| 353 | 2jsa-13 | ILE_A_60_CA | ILE_A_60_HA | -0.207 | 1.156 | 7.282 | 0.292 | 2.05 | 7.657 | 1.09 | 69.88 | 60.83 |
| 354 | 2jsa-13 | ILE_A_60_CA | ILE_A_60_C | -0.207 | 1.156 | 7.282 | -1.69 | 1.265 | 7.645 | 1.53 | 76.28 | 175.80 |
| 355 | 2jsa-13 | ILE_A_60_C | ILE_A_60_O | -1.69 | 1.265 | 7.645 | -2.167 | 2.339 | 8.006 | 1.23 | 72.92 | 113.95 |
| 356 | 2jsa-13 | ILE_A_60_C | ILE_A_61_N | -1.69 | 1.265 | 7.645 | -2.377 | 0.137 | 7.535 | 1.33 | 94.76 | -121.34 |
| 357 | 2jsa-13 | ILE_A_61_N | ILE_A_61_H | -2.377 | 0.137 | 7.535 | -1.918 | -0.71 | 7.232 | 1.01 | 107.46 | -61.55 |
| 358 | 2jsa-13 | ILE_A_61_N | ILE_A_61_CA | -2.377 | 0.137 | 7.535 | -3.795 | 0.092 | 7.847 | 1.45 | 77.60 | -178.18 |
| 359 | 2jsa-13 | ILE_A_61_CA | ILE_A_61_HA | -3.795 | 0.092 | 7.847 | -4.285 | 0.901 | 7.305 | 1.09 | 119.81 | 121.20 |
| 360 | 2jsa-13 | ILE_A_61_CA | ILE_A_61_C | -3.795 | 0.092 | 7.847 | -3.993 | 0.347 | 9.341 | 1.53 | 12.19 | 127.83 |
| 361 | 2jsa-13 | ILE_A_61_C | ILE_A_61_O | -3.993 | 0.347 | 9.341 | -5.038 | 0.844 | 9.757 | 1.23 | 70.23 | 154.56 |
| 362 | 2jsa-13 | ILE_A_61_C | HIS_A_62_N | -3.993 | 0.347 | 9.341 | -2.972 | -0.003 | 10.11 | 1.33 | 54.53 | -18.92 |
| 363 | 2jsa-13 | HIS_A_62_N | HIS_A_62_H | -2.972 | -0.003 | 10.11 | -2.142 | -0.406 | 9.7 | 1.01 | 113.96 | -25.90 |
| 364 | 2jsa-13 | HIS_A_62_N | HIS_A_62_CA | -2.972 | -0.003 | 10.11 | -3.02 | 0.182 | 11.55 | 1.45 | 7.56 | 104.55 |
| 365 | 2jsa-13 | HIS_A_62_CA | HIS_A_62_HA | -3.02 | 0.182 | 11.55 | -3.985 | -0.181 | 11.904 | 1.09 | 71.05 | -159.39 |
| 366 | 2jsa-13 | HIS_A_62_CA | HIS_A_62_C | -3.02 | 0.182 | 11.55 | -2.93 | 1.673 | 11.879 | 1.53 | 77.58 | 86.55 |
| 367 | 2jsa-13 | HIS_A_62_C | HIS_A_62_O | -2.93 | 1.673 | 11.879 | -3.212 | 2.084 | 13.003 | 1.23 | 23.92 | 124.46 |
| 368 | 2jsa-13 | HIS_A_62_C | GLU_A_63_N | -2.93 | 1.673 | 11.879 | -2.534 | 2.445 | 10.877 | 1.33 | 139.11 | 62.84 |
| 369 | 2jsa-13 | GLU_A_63_N | GLU_A_63_H | -2.534 | 2.445 | 10.877 | -2.318 | 2.039 | 9.978 | 1.01 | 152.91 | -61.99 |
| 370 | 2jsa-13 | GLU_A_63_N | GLU_A_63_CA | -2.534 | 2.445 | 10.877 | -2.402 | 3.882 | 11.045 | 1.45 | 83.36 | 84.75 |
| 371 | 2jsa-13 | GLU_A_63_CA | GLU_A_63_HA | -2.402 | 3.882 | 11.045 | -2.05 | 4.072 | 12.059 | 1.09 | 21.53 | 28.36 |
| 372 | 2jsa-13 | GLU_A_63_CA | GLU_A_63_C | -2.402 | 3.882 | 11.045 | -3.76 | 4.563 | 10.871 | 1.53 | 96.53 | 153.37 |
| 373 | 2jsa-13 | GLU_A_63_C | GLU_A_63_O | -3.76 | 4.563 | 10.871 | -4.171 | 5.361 | 11.711 | 1.23 | 46.90 | 117.25 |
| 374 | 2jsa-13 | GLU_A_63_C | LEU_A_64_N | -3.76 | 4.563 | 10.871 | -4.421 | 4.223 | 9.774 | 1.33 | 145.88 | -152.78 |
| 375 | 2jsa-13 | LEU_A_64_N | LEU_A_64_H | -4.421 | 4.223 | 9.774 | -4.022 | 3.559 | 9.126 | 1.01 | 129.91 | -59.00 |
| 376 | 2jsa-13 | LEU_A_64_N | LEU_A_64_CA | -4.421 | 4.223 | 9.774 | -5.725 | 4.792 | 9.478 | 1.45 | 101.75 | 156.43 |
| 377 | 2jsa-13 | LEU_A_64_CA | LEU_A_64_HA | -5.725 | 4.792 | 9.478 | -5.669 | 5.868 | 9.646 | 1.09 | 81.14 | 87.02 |
| 378 | 2jsa-13 | LEU_A_64_CA | LEU_A_64_C | -5.725 | 4.792 | 9.478 | -6.758 | 4.222 | 10.452 | 1.53 | 50.46 | -151.11 |
| 379 | 2jsa-13 | LEU_A_64_C | LEU_A_64_O | -6.758 | 4.222 | 10.452 | -7.761 | 4.87 | 10.748 | 1.23 | 76.08 | 147.14 |
| 380 | 2jsa-13 | LEU_A_64_C | GLU_A_65_N | -6.758 | 4.222 | 10.452 | -6.477 | 3.017 | 10.925 | 1.32 | 69.08 | -76.87 |
| 381 | 2jsa-13 | GLU_A_65_N | GLU_A_65_H | -6.477 | 3.017 | 10.925 | -5.632 | 2.543 | 10.639 | 1.01 | 106.45 | -29.29 |
| 382 | 2jsa-13 | GLU_A_65_N | GLU_A_65_CA | -6.477 | 3.017 | 10.925 | -7.369 | 2.353 | 11.86 | 1.45 | 49.94 | -143.34 |
| 383 | 2jsa-13 | GLU_A_65_CA | GLU_A_65_HA | -7.369 | 2.353 | 11.86 | -8.377 | 2.392 | 11.449 | 1.09 | 112.17 | 177.78 |
| 384 | 2jsa-13 | GLU_A_65_CA | GLU_A_65_C | -7.369 | 2.353 | 11.86 | -7.369 | 3.084 | 13.204 | 1.53 | 28.54 | 90.00 |
| 385 | 2jsa-13 | GLU_A_65_C | GLU_A_65_O | -7.369 | 3.084 | 13.204 | -8.409 | 3.206 | 13.849 | 1.23 | 58.37 | 173.31 |
| 386 | 2jsa-13 | GLU_A_65_C | GLY_A_66_N | -7.369 | 3.084 | 13.204 | -6.189 | 3.55 | 13.588 | 1.33 | 73.16 | 21.55 |
| 387 | 2jsa-13 | GLY_A_66_N | GLY_A_66_H | -6.189 | 3.55 | 13.588 | -5.374 | 3.413 | 13.007 | 1.01 | 125.11 | -9.54 |
| 388 | 2jsa-13 | GLY_A_66_N | GLY_A_66_CA | -6.189 | 3.55 | 13.588 | -6.039 | 4.264 | 14.844 | 1.45 | 30.15 | 78.14 |
| 389 | 2jsa-13 | GLY_A_66_CA | GLY_A_66_C | -6.039 | 4.264 | 14.844 | -6.586 | 5.69 | 14.732 | 1.53 | 94.19 | 110.99 |
| 390 | 2jsa-13 | GLY_A_66_C | GLY_A_66_O | -6.586 | 5.69 | 14.732 | -7.399 | 6.112 | 15.553 | 1.23 | 48.13 | 152.57 |
| 391 | 2jsa-13 | GLY_A_66_C | GLY_A_67_N | -6.586 | 5.69 | 14.732 | -6.118 | 6.39 | 13.71 | 1.32 | 140.51 | 56.23 |
| 392 | 2jsa-13 | GLY_A_67_N | GLY_A_67_H | -6.118 | 6.39 | 13.71 | -5.45 | 5.978 | 13.075 | 1.01 | 128.98 | -31.66 |
| 393 | 2jsa-13 | GLY_A_67_N | GLY_A_67_CA | -6.118 | 6.39 | 13.71 | -6.55 | 7.759 | 13.479 | 1.45 | 99.14 | 107.51 |
| 394 | 2jsa-13 | GLY_A_67_CA | GLY_A_67_C | -6.55 | 7.759 | 13.479 | -5.48 | 8.551 | 12.723 | 1.53 | 119.59 | 36.51 |
| 395 | 2jsa-13 | GLY_A_67_C | GLY_A_67_O | -5.48 | 8.551 | 12.723 | -5.208 | 9.703 | 13.054 | 1.23 | 74.38 | 76.72 |
| 396 | 2jsa-13 | GLY_A_67_C | THR_A_68_N | -5.48 | 8.551 | 12.723 | -4.906 | 7.901 | 11.723 | 1.32 | 139.07 | -48.55 |
| 397 | 2jsa-13 | THR_A_68_N | THR_A_68_H | -4.906 | 7.901 | 11.723 | -5.182 | 6.954 | 11.507 | 1.01 | 102.35 | -106.25 |
| 398 | 2jsa-13 | THR_A_68_N | THR_A_68_CA | -4.906 | 7.901 | 11.723 | -3.873 | 8.53 | 10.917 | 1.45 | 123.68 | 31.34 |
| 399 | 2jsa-13 | THR_A_68_CA | THR_A_68_HA | -3.873 | 8.53 | 10.917 | -3.731 | 9.553 | 11.266 | 1.09 | 71.33 | 82.10 |
| 400 | 2jsa-13 | THR_A_68_CA | THR_A_68_C | -3.873 | 8.53 | 10.917 | -4.303 | 8.587 | 9.449 | 1.53 | 163.54 | 172.45 |
| 401 | 2jsa-13 | THR_A_68_C | THR_A_68_O | -4.303 | 8.587 | 9.449 | -4.533 | 7.552 | 8.826 | 1.23 | 120.44 | -102.53 |
| 402 | 2jsa-13 | THR_A_68_C | ALA_A_69_N | -4.303 | 8.587 | 9.449 | -4.398 | 9.807 | 8.94 | 1.33 | 112.59 | 94.45 |
| 403 | 2jsa-13 | ALA_A_69_N | ALA_A_69_H | -4.398 | 9.807 | 8.94 | -4.194 | 10.614 | 9.512 | 1.01 | 55.50 | 75.81 |
| 404 | 2jsa-13 | ALA_A_69_N | ALA_A_69_CA | -4.398 | 9.807 | 8.94 | -4.795 | 10.012 | 7.558 | 1.45 | 162.08 | 152.69 |
| 405 | 2jsa-13 | ALA_A_69_CA | ALA_A_69_HA | -4.795 | 10.012 | 7.558 | -5.83 | 9.692 | 7.44 | 1.09 | 96.22 | -162.82 |
| 406 | 2jsa-13 | ALA_A_69_CA | ALA_A_69_C | -4.795 | 10.012 | 7.558 | -3.917 | 9.153 | 6.646 | 1.53 | 126.59 | -44.37 |
| 407 | 2jsa-13 | ALA_A_69_C | ALA_A_69_O | -3.917 | 9.153 | 6.646 | -2.865 | 8.672 | 7.065 | 1.23 | 70.09 | -24.57 |
| 408 | 2jsa-13 | ALA_A_69_C | PRO_A_70_N | -3.917 | 9.153 | 6.646 | -4.394 | 8.982 | 5.384 | 1.36 | 158.12 | -160.28 |
| 409 | 2jsa-13 | PRO_A_70_N | PRO_A_70_CA | -4.394 | 8.982 | 5.384 | -3.664 | 8.189 | 4.41 | 1.45 | 132.10 | -47.37 |
| 410 | 2jsa-13 | PRO_A_70_CA | PRO_A_70_HA | -3.664 | 8.189 | 4.41 | -3.319 | 7.261 | 4.867 | 1.09 | 65.22 | -69.61 |
| 411 | 2jsa-13 | PRO_A_70_CA | PRO_A_70_C | -3.664 | 8.189 | 4.41 | -2.446 | 8.952 | 3.883 | 1.53 | 110.14 | 32.06 |
| 412 | 2jsa-13 | PRO_A_70_C | PRO_A_70_O | -2.446 | 8.952 | 3.883 | -1.521 | 8.352 | 3.337 | 1.23 | 116.35 | -32.97 |
| 413 | 2jsa-13 | PRO_A_70_C | LYS_A_71_N | -2.446 | 8.952 | 3.883 | -2.485 | 10.264 | 4.067 | 1.33 | 82.02 | 91.70 |
| 414 | 2jsa-13 | LYS_A_71_N | LYS_A_71_H | -2.485 | 10.264 | 4.067 | -3.278 | 10.691 | 4.525 | 1.01 | 63.05 | 151.70 |
| 415 | 2jsa-13 | LYS_A_71_N | LYS_A_71_CA | -2.485 | 10.264 | 4.067 | -1.396 | 11.114 | 3.618 | 1.45 | 108.01 | 37.97 |
| 416 | 2jsa-13 | LYS_A_71_CA | LYS_A_71_HA | -1.396 | 11.114 | 3.618 | -0.84 | 10.574 | 2.852 | 1.09 | 134.66 | -44.16 |
| 417 | 2jsa-13 | LYS_A_71_CA | LYS_A_71_C | -1.396 | 11.114 | 3.618 | -0.443 | 11.376 | 4.786 | 1.53 | 40.24 | 15.37 |
| 418 | 2jsa-13 | LYS_A_71_C | LYS_A_71_O | -0.443 | 11.376 | 4.786 | 0.442 | 12.225 | 4.692 | 1.23 | 94.38 | 43.81 |
| 419 | 2jsa-13 | LYS_A_71_C | ASP_A_72_N | -0.443 | 11.376 | 4.786 | -0.657 | 10.632 | 5.861 | 1.32 | 35.76 | -106.05 |

| | | | | | | | | | | | | |
|---|---|---|---|---|---|---|---|---|---|---|---|---|
| 420 | 2jsa-13 | ASP_A_72_N | ASP_A_72_H | -0.657 | 10.632 | 5.861 | -1.405 | 9.954 | 5.869 | 1.01 | 89.55 | -137.81 |
| 421 | 2jsa-13 | ASP_A_72_N | ASP_A_72_CA | -0.657 | 10.632 | 5.861 | 0.171 | 10.771 | 7.047 | 1.45 | 35.30 | 9.53 |
| 422 | 2jsa-13 | ASP_A_72_CA | ASP_A_72_HA | 0.171 | 10.771 | 7.047 | 0.86 | 11.604 | 6.905 | 1.09 | 97.48 | 50.40 |
| 423 | 2jsa-13 | ASP_A_72_CA | ASP_A_72_C | 0.171 | 10.771 | 7.047 | 0.958 | 9.479 | 7.269 | 1.53 | 81.65 | -58.65 |
| 424 | 2jsa-13 | ASP_A_72_C | ASP_A_72_O | 0.958 | 9.479 | 7.269 | 2.122 | 9.518 | 7.666 | 1.23 | 71.18 | 1.92 |
| 425 | 2jsa-13 | ASP_A_72_C | VAL_A_73_N | 0.958 | 9.479 | 7.269 | 0.294 | 8.365 | 7.002 | 1.32 | 101.63 | -120.80 |
| 426 | 2jsa-13 | VAL_A_73_N | VAL_A_73_H | 0.294 | 8.365 | 7.002 | -0.661 | 8.409 | 6.677 | 1.01 | 108.78 | 177.36 |
| 427 | 2jsa-13 | VAL_A_73_N | VAL_A_73_CA | 0.294 | 8.365 | 7.002 | 0.916 | 7.063 | 7.169 | 1.45 | 83.40 | -64.46 |
| 428 | 2jsa-13 | VAL_A_73_CA | VAL_A_73_HA | 0.916 | 7.063 | 7.169 | 1.113 | 6.909 | 8.23 | 1.09 | 13.26 | -38.02 |
| 429 | 2jsa-13 | VAL_A_73_CA | VAL_A_73_C | 0.916 | 7.063 | 7.169 | 2.256 | 7.046 | 6.429 | 1.53 | 118.91 | -0.73 |
| 430 | 2jsa-13 | VAL_A_73_C | VAL_A_73_O | 2.256 | 7.046 | 6.429 | 3.288 | 6.718 | 7.013 | 1.23 | 61.66 | -17.63 |
| 431 | 2jsa-13 | VAL_A_73_C | CYS_A_74_N | 2.256 | 7.046 | 6.429 | 2.197 | 7.403 | 5.155 | 1.32 | 164.14 | 99.38 |
| 432 | 2jsa-13 | CYS_A_74_N | CYS_A_74_H | 2.197 | 7.403 | 5.155 | 1.314 | 7.661 | 4.74 | 1.01 | 114.28 | 163.71 |
| 433 | 2jsa-13 | CYS_A_74_N | CYS_A_74_CA | 2.197 | 7.403 | 5.155 | 3.392 | 7.433 | 4.329 | 1.45 | 124.64 | 1.44 |
| 434 | 2jsa-13 | CYS_A_74_CA | CYS_A_74_HA | 3.392 | 7.433 | 4.329 | 3.988 | 6.549 | 4.556 | 1.09 | 77.98 | -56.01 |
| 435 | 2jsa-13 | CYS_A_74_CA | CYS_A_74_C | 3.392 | 7.433 | 4.329 | 4.203 | 8.675 | 4.702 | 1.53 | 75.89 | 56.86 |
| 436 | 2jsa-13 | CYS_A_74_C | CYS_A_74_O | 4.203 | 8.675 | 4.702 | 4.241 | 9.645 | 3.947 | 1.23 | 127.87 | 87.76 |
| 437 | 2jsa-13 | CYS_A_74_C | THR_A_75_N | 4.203 | 8.675 | 4.702 | 4.831 | 8.606 | 5.867 | 1.33 | 28.47 | -6.27 |
| 438 | 2jsa-13 | THR_A_75_N | THR_A_75_H | 4.831 | 8.606 | 5.867 | 4.755 | 7.776 | 6.437 | 1.01 | 55.63 | -95.23 |
| 439 | 2jsa-13 | THR_A_75_N | THR_A_75_CA | 4.831 | 8.606 | 5.867 | 5.639 | 9.713 | 6.35 | 1.45 | 70.59 | 53.87 |
| 440 | 2jsa-13 | THR_A_75_CA | THR_A_75_HA | 5.639 | 9.713 | 6.35 | 6.465 | 9.872 | 5.656 | 1.09 | 129.52 | 10.90 |
| 441 | 2jsa-13 | THR_A_75_CA | THR_A_75_C | 5.639 | 9.713 | 6.35 | 6.233 | 9.382 | 7.72 | 1.53 | 26.40 | -29.13 |
| 442 | 2jsa-13 | THR_A_75_C | THR_A_75_O | 6.233 | 9.382 | 7.72 | 7.451 | 9.296 | 7.868 | 1.23 | 83.09 | -4.04 |
| 443 | 2jsa-13 | THR_A_75_C | LYS_A_76_N | 6.233 | 9.382 | 7.72 | 5.344 | 9.205 | 8.686 | 1.32 | 43.18 | -168.74 |
| 444 | 2jsa-13 | LYS_A_76_N | LYS_A_76_H | 5.344 | 9.205 | 8.686 | 4.357 | 9.29 | 8.492 | 1.01 | 101.08 | 175.08 |
| 445 | 2jsa-13 | LYS_A_76_N | LYS_A_76_CA | 5.344 | 9.205 | 8.686 | 5.766 | 8.886 | 10.04 | 1.45 | 21.34 | -37.09 |
| 446 | 2jsa-13 | LYS_A_76_CA | LYS_A_76_HA | 5.766 | 8.886 | 10.04 | 6.614 | 9.525 | 10.289 | 1.09 | 76.80 | 37.00 |
| 447 | 2jsa-13 | LYS_A_76_CA | LYS_A_76_C | 5.766 | 8.886 | 10.04 | 6.247 | 7.434 | 10.091 | 1.53 | 88.09 | -71.67 |
| 448 | 2jsa-13 | LYS_A_76_C | LYS_A_76_O | 6.247 | 7.434 | 10.091 | 7.089 | 7.084 | 10.916 | 1.23 | 47.86 | -22.57 |
| 449 | 2jsa-13 | LYS_A_76_C | LEU_A_77_N | 6.247 | 7.434 | 10.091 | 5.689 | 6.629 | 9.198 | 1.33 | 132.36 | -124.73 |
| 450 | 2jsa-13 | LEU_A_77_N | LEU_A_77_H | 5.689 | 6.629 | 9.198 | 5.002 | 6.986 | 8.549 | 1.01 | 129.97 | 152.54 |
| 451 | 2jsa-13 | LEU_A_77_N | LEU_A_77_CA | 5.689 | 6.629 | 9.198 | 6.051 | 5.223 | 9.132 | 1.45 | 92.60 | -75.56 |
| 452 | 2jsa-13 | LEU_A_77_CA | LEU_A_77_HA | 6.051 | 5.223 | 9.132 | 6.088 | 4.836 | 10.15 | 1.09 | 20.90 | -84.54 |
| 453 | 2jsa-13 | LEU_A_77_CA | LEU_A_77_C | 6.051 | 5.223 | 9.132 | 7.452 | 5.089 | 8.532 | 1.53 | 113.09 | -5.46 |
| 454 | 2jsa-13 | LEU_A_77_C | LEU_A_77_O | 7.452 | 5.089 | 8.532 | 7.98 | 3.983 | 8.418 | 1.23 | 95.31 | -64.48 |
| 455 | 2jsa-13 | LEU_A_77_C | ASN_A_78_N | 7.452 | 5.089 | 8.532 | 8.015 | 6.231 | 8.164 | 1.33 | 106.12 | 63.76 |
| 456 | 2jsa-13 | ASN_A_78_N | ASN_A_78_H | 8.015 | 6.231 | 8.164 | 7.523 | 7.104 | 8.287 | 1.01 | 83.00 | 119.40 |
| 457 | 2jsa-13 | ASN_A_78_N | ASN_A_78_CA | 8.015 | 6.231 | 8.164 | 9.345 | 6.254 | 7.579 | 1.45 | 113.74 | 0.99 |
| 458 | 2jsa-13 | ASN_A_78_CA | ASN_A_78_HA | 9.345 | 6.254 | 7.579 | 9.638 | 7.289 | 7.4 | 1.09 | 99.45 | 74.19 |
| 459 | 2jsa-13 | ASN_A_78_CA | ASN_A_78_C | 9.345 | 6.254 | 7.579 | 9.329 | 5.483 | 6.257 | 1.53 | 149.74 | -91.19 |
| 460 | 2jsa-13 | ASN_A_78_C | ASN_A_78_O | 9.329 | 5.483 | 6.257 | 10.365 | 4.993 | 5.809 | 1.23 | 111.35 | -25.31 |
| 461 | 2jsa-13 | ASN_A_78_C | GLU_A_79_N | 9.329 | 5.483 | 6.257 | 8.144 | 5.399 | 5.672 | 1.32 | 116.22 | -175.95 |
| 462 | 2jsa-13 | GLU_A_79_N | GLU_A_79_H | 8.144 | 5.399 | 5.672 | 7.335 | 5.827 | 6.099 | 1.01 | 64.99 | 152.12 |
| 463 | 2jsa-13 | GLU_A_79_N | GLU_A_79_CA | 8.144 | 5.399 | 5.672 | 7.981 | 4.695 | 4.411 | 1.45 | 150.18 | -103.04 |
| 464 | 2jsa-13 | GLU_A_79_CA | GLU_A_79_HA | 7.981 | 4.695 | 4.411 | 8.84 | 4.037 | 4.278 | 1.09 | 97.01 | -37.45 |
| 465 | 2jsa-13 | GLU_A_79_CA | GLU_A_79_C | 7.981 | 4.695 | 4.411 | 7.953 | 5.689 | 3.249 | 1.53 | 139.44 | 91.61 |
| 466 | 2jsa-13 | GLU_A_79_C | GLU_A_79_O | 7.953 | 5.689 | 3.249 | 7.942 | 5.289 | 2.085 | 1.23 | 161.03 | -91.58 |
| 467 | 2jsa-13 | GLU_A_79_C | CYS_A_80_N | 7.953 | 5.689 | 3.249 | 7.942 | 6.965 | 3.603 | 1.32 | 74.49 | 90.49 |
| 468 | 2jsa-13 | CYS_A_80_N | CYS_A_80_H | 7.942 | 6.965 | 3.603 | 7.953 | 7.222 | 4.58 | 1.01 | 14.75 | 87.55 |
| 469 | 2jsa-13 | CYS_A_80_N | CYS_A_80_CA | 7.942 | 6.965 | 3.603 | 7.916 | 8.02 | 2.603 | 1.45 | 133.46 | 91.41 |
| 470 | 2jsa-13 | CYS_A_80_CA | CYS_A_80_HA | 7.916 | 8.02 | 2.603 | 8.419 | 7.656 | 1.707 | 1.09 | 145.28 | -35.89 |
| 471 | 2jsa-13 | CYS_A_80_CA | CYS_A_80_C | 7.916 | 8.02 | 2.603 | 8.699 | 9.217 | 3.147 | 1.53 | 69.18 | 56.81 |
| 472 | 2jsa-13 | CYS_A_80_C | CYS_A_80_O | 8.699 | 9.217 | 3.147 | 8.534 | 9.595 | 4.305 | 1.23 | 19.60 | 113.58 |
| 473 | 2jsa-13 | CYS_A_80_C | PRO_A_81_N | 8.699 | 9.217 | 3.147 | 9.554 | 9.795 | 2.262 | 1.36 | 130.61 | 34.06 |
| 474 | 2jsa-13 | PRO_A_81_N | PRO_A_81_CA | 9.554 | 9.795 | 2.262 | 10.362 | 10.942 | 2.642 | 1.45 | 74.85 | 54.84 |
| 475 | 2jsa-13 | PRO_A_81_CA | PRO_A_81_HA | 10.362 | 10.942 | 2.642 | 10.809 | 10.776 | 3.623 | 1.09 | 25.92 | -20.37 |
| 476 | 2jsa-13 | PRO_A_81_CA | PRO_A_81_C | 10.362 | 10.942 | 2.642 | 9.513 | 12.214 | 2.704 | 1.53 | 87.68 | 123.72 |
| 477 | 2jsa-13 | PRO_A_81_C | PRO_A_81_O | 9.513 | 12.214 | 2.704 | 8.286 | 12.143 | 2.759 | 1.23 | 87.44 | -176.69 |
| 1 | 2jsa-14 | ARG_A_1_N | ARG_A_1_H | 13.785 | 3.394 | -4.024 | 14.475 | 4.121 | -4.153 | 1.01 | 97.33 | 46.50 |
| 2 | 2jsa-14 | ARG_A_1_N | ARG_A_1_CA | 13.785 | 3.394 | -4.024 | 12.457 | 3.77 | -3.568 | 1.45 | 71.72 | 164.19 |
| 3 | 2jsa-14 | ARG_A_1_CA | ARG_A_1_HA | 12.457 | 3.77 | -3.568 | 11.735 | 3.47 | -4.327 | 1.09 | 134.15 | -157.44 |
| 4 | 2jsa-14 | ARG_A_1_CA | ARG_A_1_C | 12.457 | 3.77 | -3.568 | 12.372 | 5.284 | -3.37 | 1.53 | 82.56 | 93.21 |
| 5 | 2jsa-14 | ARG_A_1_C | ARG_A_1_O | 12.372 | 5.284 | -3.37 | 12.68 | 5.791 | -2.293 | 1.23 | 28.85 | 58.72 |
| 6 | 2jsa-14 | ARG_A_1_C | SER_A_2_N | 12.372 | 5.284 | -3.37 | 11.953 | 5.964 | -4.427 | 1.32 | 142.92 | 121.64 |
| 7 | 2jsa-14 | SER_A_2_N | SER_A_2_H | 11.953 | 5.964 | -4.427 | 11.718 | 5.481 | -5.282 | 1.01 | 147.86 | -115.94 |
| 8 | 2jsa-14 | SER_A_2_N | SER_A_2_CA | 11.953 | 5.964 | -4.427 | 11.822 | 7.41 | -4.383 | 1.45 | 88.26 | 95.18 |
| 9 | 2jsa-14 | SER_A_2_CA | SER_A_2_HA | 11.822 | 7.41 | -4.383 | 12.039 | 7.742 | -3.367 | 1.09 | 21.32 | 56.83 |
| 10 | 2jsa-14 | SER_A_2_CA | SER_A_2_C | 11.822 | 7.41 | -4.383 | 10.391 | 7.819 | -4.736 | 1.53 | 103.34 | 164.05 |
| 11 | 2jsa-14 | SER_A_2_C | SER_A_2_O | 10.391 | 7.819 | -4.736 | 10.151 | 8.945 | -5.167 | 1.23 | 110.52 | 102.03 |
| 12 | 2jsa-14 | SER_A_2_C | ALA_A_3_N | 10.391 | 7.819 | -4.736 | 9.477 | 6.88 | -4.54 | 1.32 | 81.49 | -134.23 |
| 13 | 2jsa-14 | ALA_A_3_N | ALA_A_3_H | 9.477 | 6.88 | -4.54 | 9.747 | 5.974 | -4.183 | 1.01 | 69.31 | -73.41 |
| 14 | 2jsa-14 | ALA_A_3_N | ALA_A_3_CA | 9.477 | 6.88 | -4.54 | 8.075 | 7.128 | -4.832 | 1.45 | 101.59 | 169.97 |
| 15 | 2jsa-14 | ALA_A_3_CA | ALA_A_3_HA | 8.075 | 7.128 | -4.832 | 7.726 | 7.948 | -4.204 | 1.09 | 54.83 | 113.06 |
| 16 | 2jsa-14 | ALA_A_3_CA | ALA_A_3_C | 8.075 | 7.128 | -4.832 | 7.26 | 5.877 | -4.494 | 1.53 | 77.24 | -123.08 |
| 17 | 2jsa-14 | ALA_A_3_C | ALA_A_3_O | 7.26 | 5.877 | -4.494 | 6.319 | 5.941 | -3.703 | 1.23 | 50.01 | 176.11 |
| 18 | 2jsa-14 | ALA_A_3_C | LEU_A_4_N | 7.26 | 5.877 | -4.494 | 7.649 | 4.771 | -5.11 | 1.32 | 117.72 | -70.62 |
| 19 | 2jsa-14 | LEU_A_4_N | LEU_A_4_H | 7.649 | 4.771 | -5.11 | 8.432 | 4.793 | -5.748 | 1.01 | 129.16 | 1.61 |
| 20 | 2jsa-14 | LEU_A_4_N | LEU_A_4_CA | 7.649 | 4.771 | -5.11 | 6.966 | 3.509 | -4.885 | 1.45 | 81.09 | -118.42 |
| 21 | 2jsa-14 | LEU_A_4_CA | LEU_A_4_HA | 6.966 | 3.509 | -4.885 | 6.049 | 3.512 | -5.473 | 1.09 | 122.67 | 179.81 |
| 22 | 2jsa-14 | LEU_A_4_CA | LEU_A_4_C | 6.966 | 3.509 | -4.885 | 6.573 | 3.403 | -3.409 | 1.53 | 15.42 | -164.91 |
| 23 | 2jsa-14 | LEU_A_4_C | LEU_A_4_O | 6.573 | 3.403 | -3.409 | 5.431 | 3.079 | -3.089 | 1.23 | 74.91 | -164.16 |
| 24 | 2jsa-14 | LEU_A_4_C | SER_A_5_N | 6.573 | 3.403 | -3.409 | 7.543 | 3.683 | -2.55 | 1.33 | 49.61 | 16.10 |
| 25 | 2jsa-14 | SER_A_5_N | SER_A_5_H | 7.543 | 3.683 | -2.55 | 8.46 | 3.944 | -2.884 | 1.01 | 109.31 | 15.89 |
| 26 | 2jsa-14 | SER_A_5_N | SER_A_5_CA | 7.543 | 3.683 | -2.55 | 7.312 | 3.623 | -1.117 | 1.45 | 9.46 | -165.44 |
| 27 | 2jsa-14 | SER_A_5_CA | SER_A_5_HA | 7.312 | 3.623 | -1.117 | 7.188 | 2.578 | -0.834 | 1.09 | 74.95 | -96.77 |
| 28 | 2jsa-14 | SER_A_5_CA | SER_A_5_C | 7.312 | 3.623 | -1.117 | 6.032 | 4.382 | -0.76 | 1.53 | 76.51 | 149.33 |
| 29 | 2jsa-14 | SER_A_5_C | SER_A_5_O | 6.032 | 4.382 | -0.76 | 5.064 | 3.787 | -0.29 | 1.23 | 67.53 | -148.42 |
| 30 | 2jsa-14 | SER_A_5_C | CYS_A_6_N | 6.032 | 4.382 | -0.76 | 6.07 | 5.685 | -0.997 | 1.32 | 100.30 | 88.33 |
| 31 | 2jsa-14 | CYS_A_6_N | CYS_A_6_H | 6.07 | 5.685 | -0.997 | 6.901 | 6.108 | -1.385 | 1.01 | 112.59 | 26.98 |
| 32 | 2jsa-14 | CYS_A_6_N | CYS_A_6_CA | 6.07 | 5.685 | -0.997 | 4.925 | 6.532 | -0.707 | 1.45 | 78.49 | 143.51 |
| 33 | 2jsa-14 | CYS_A_6_CA | CYS_A_6_HA | 4.925 | 6.532 | -0.707 | 4.668 | 6.413 | 0.346 | 1.09 | 15.05 | -155.15 |
| 34 | 2jsa-14 | CYS_A_6_CA | CYS_A_6_C | 4.925 | 6.532 | -0.707 | 3.745 | 6.05 | -1.554 | 1.53 | 123.60 | -157.78 |
| 35 | 2jsa-14 | CYS_A_6_C | CYS_A_6_O | 3.745 | 6.05 | -1.554 | 2.77 | 5.521 | -1.022 | 1.23 | 64.38 | -151.52 |
| 36 | 2jsa-14 | CYS_A_6_C | GLN_A_7_N | 3.745 | 6.05 | -1.554 | 3.871 | 6.253 | -2.858 | 1.33 | 169.62 | 58.17 |
| 37 | 2jsa-14 | GLN_A_7_N | GLN_A_7_H | 3.871 | 6.253 | -2.858 | 4.701 | 6.697 | -3.226 | 1.01 | 111.35 | 28.14 |
| 38 | 2jsa-14 | GLN_A_7_N | GLN_A_7_CA | 3.871 | 6.253 | -2.858 | 2.827 | 5.847 | -3.783 | 1.45 | 129.55 | -158.75 |

| | | | | | | | | | | | | |
|---|---|---|---|---|---|---|---|---|---|---|---|---|
| 39 | 2jsa-14 | GLN_A_7_CA | GLN_A_7_HA | 2.827 | 5.847 | -3.783 | 2.089 | 6.647 | -3.83 | 1.09 | 92.47 | 132.69 |
| 40 | 2jsa-14 | GLN_A_7_CA | GLN_A_7_C | 2.827 | 5.847 | -3.783 | 2.133 | 4.581 | -3.276 | 1.53 | 70.65 | -118.73 |
| 41 | 2jsa-14 | GLN_A_7_C | GLN_A_7_O | 2.133 | 4.581 | -3.276 | 0.939 | 4.6 | -2.978 | 1.23 | 75.99 | 179.09 |
| 42 | 2jsa-14 | GLN_A_7_C | MET_A_8_N | 2.133 | 4.581 | -3.276 | 2.909 | 3.51 | -3.195 | 1.33 | 86.50 | -54.07 |
| 43 | 2jsa-14 | MET_A_8_N | MET_A_8_H | 2.909 | 3.51 | -3.195 | 3.882 | 3.566 | -3.459 | 1.01 | 105.16 | 3.29 |
| 44 | 2jsa-14 | MET_A_8_N | MET_A_8_CA | 2.909 | 3.51 | -3.195 | 2.383 | 2.238 | -2.729 | 1.45 | 71.30 | -112.47 |
| 45 | 2jsa-14 | MET_A_8_CA | MET_A_8_HA | 2.383 | 2.238 | -2.729 | 1.768 | 1.804 | -3.517 | 1.09 | 136.31 | -144.79 |
| 46 | 2jsa-14 | MET_A_8_CA | MET_A_8_C | 2.383 | 2.238 | -2.729 | 1.526 | 2.426 | -1.476 | 1.53 | 35.00 | 167.63 |
| 47 | 2jsa-14 | MET_A_8_C | MET_A_8_O | 1.526 | 2.426 | -1.476 | 0.423 | 1.887 | -1.388 | 1.23 | 85.90 | -153.96 |
| 48 | 2jsa-14 | MET_A_8_C | CYS_A_9_N | 1.526 | 2.426 | -1.476 | 2.064 | 3.19 | -0.537 | 1.32 | 44.86 | 54.85 |
| 49 | 2jsa-14 | CYS_A_9_N | CYS_A_9_H | 2.064 | 3.19 | -0.537 | 2.976 | 3.602 | -0.677 | 1.01 | 97.96 | 24.31 |
| 50 | 2jsa-14 | CYS_A_9_N | CYS_A_9_CA | 2.064 | 3.19 | -0.537 | 1.363 | 3.454 | 0.707 | 1.45 | 31.05 | 159.36 |
| 51 | 2jsa-14 | CYS_A_9_CA | CYS_A_9_HA | 1.363 | 3.454 | 0.707 | 1.136 | 2.499 | 1.182 | 1.09 | 64.18 | -103.37 |
| 52 | 2jsa-14 | CYS_A_9_CA | CYS_A_9_C | 1.363 | 3.454 | 0.707 | 0.046 | 4.159 | 0.376 | 1.53 | 102.49 | 151.84 |
| 53 | 2jsa-14 | CYS_A_9_C | CYS_A_9_O | 0.046 | 4.159 | 0.376 | -0.954 | 3.97 | 1.066 | 1.23 | 55.86 | -169.30 |
| 54 | 2jsa-14 | CYS_A_9_C | GLU_A_10_N | 0.046 | 4.159 | 0.376 | 0.089 | 4.958 | -0.68 | 1.32 | 142.85 | 86.92 |
| 55 | 2jsa-14 | GLU_A_10_N | GLU_A_10_H | 0.089 | 4.958 | -0.68 | 0.948 | 5.066 | -1.201 | 1.01 | 121.04 | 7.17 |
| 56 | 2jsa-14 | GLU_A_10_N | GLU_A_10_CA | 0.089 | 4.958 | -0.68 | -1.088 | 5.692 | -1.112 | 1.45 | 107.30 | 148.05 |
| 57 | 2jsa-14 | GLU_A_10_CA | GLU_A_10_HA | -1.088 | 5.692 | -1.112 | -1.627 | 6.018 | -0.222 | 1.09 | 35.29 | 148.83 |
| 58 | 2jsa-14 | GLU_A_10_CA | GLU_A_10_C | -1.088 | 5.692 | -1.112 | -2.011 | 4.784 | -1.927 | 1.53 | 122.19 | -135.47 |
| 59 | 2jsa-14 | GLU_A_10_C | GLU_A_10_O | -2.011 | 4.784 | -1.927 | -3.107 | 5.191 | -2.31 | 1.23 | 108.14 | 159.63 |
| 60 | 2jsa-14 | GLU_A_10_C | LEU_A_11_N | -2.011 | 4.784 | -1.927 | -1.534 | 3.572 | -2.169 | 1.32 | 100.53 | -68.52 |
| 61 | 2jsa-14 | LEU_A_11_N | LEU_A_11_H | -1.534 | 3.572 | -2.169 | -0.623 | 3.306 | -1.823 | 1.01 | 69.97 | -16.28 |
| 62 | 2jsa-14 | LEU_A_11_N | LEU_A_11_CA | -1.534 | 3.572 | -2.169 | -2.302 | 2.603 | -2.931 | 1.45 | 121.64 | -128.40 |
| 63 | 2jsa-14 | LEU_A_11_CA | LEU_A_11_HA | -2.302 | 2.603 | -2.931 | -3.001 | 3.15 | -3.564 | 1.09 | 125.50 | 141.96 |
| 64 | 2jsa-14 | LEU_A_11_CA | LEU_A_11_C | -2.302 | 2.603 | -2.931 | -3.123 | 1.74 | -1.971 | 1.53 | 51.13 | -133.57 |
| 65 | 2jsa-14 | LEU_A_11_C | LEU_A_11_O | -3.123 | 1.74 | -1.971 | -4.276 | 1.418 | -2.251 | 1.23 | 103.16 | -164.40 |
| 66 | 2jsa-14 | LEU_A_11_C | VAL_A_12_N | -3.123 | 1.74 | -1.971 | -2.494 | 1.389 | -0.859 | 1.32 | 32.93 | -29.16 |
| 67 | 2jsa-14 | VAL_A_12_N | VAL_A_12_H | -2.494 | 1.389 | -0.859 | -1.544 | 1.69 | -0.695 | 1.01 | 80.65 | 17.58 |
| 68 | 2jsa-14 | VAL_A_12_N | VAL_A_12_CA | -2.494 | 1.389 | -0.859 | -3.151 | 0.568 | 0.145 | 1.45 | 46.32 | -128.67 |
| 69 | 2jsa-14 | VAL_A_12_CA | VAL_A_12_HA | -3.151 | 0.568 | 0.145 | -3.667 | -0.247 | -0.361 | 1.09 | 117.68 | -122.34 |
| 70 | 2jsa-14 | VAL_A_12_CA | VAL_A_12_C | -3.151 | 0.568 | 0.145 | -4.197 | 1.408 | 0.88 | 1.53 | 61.28 | 141.23 |
| 71 | 2jsa-14 | VAL_A_12_C | VAL_A_12_O | -4.197 | 1.408 | 0.88 | -5.345 | 0.991 | 1.023 | 1.23 | 83.32 | -160.04 |
| 72 | 2jsa-14 | VAL_A_12_C | VAL_A_13_N | -4.197 | 1.408 | 0.88 | -3.763 | 2.578 | 1.326 | 1.33 | 70.33 | 69.65 |
| 73 | 2jsa-14 | VAL_A_13_N | VAL_A_13_H | -3.763 | 2.578 | 1.326 | -2.804 | 2.857 | 1.171 | 1.01 | 98.82 | 16.22 |
| 74 | 2jsa-14 | VAL_A_13_N | VAL_A_13_CA | -3.763 | 2.578 | 1.326 | -4.647 | 3.481 | 2.043 | 1.45 | 60.43 | 134.39 |
| 75 | 2jsa-14 | VAL_A_13_CA | VAL_A_13_HA | -4.647 | 3.481 | 2.043 | -5.06 | 2.95 | 2.9 | 1.09 | 38.13 | -127.87 |
| 76 | 2jsa-14 | VAL_A_13_CA | VAL_A_13_C | -4.647 | 3.481 | 2.043 | -5.806 | 3.885 | 1.129 | 1.53 | 126.67 | 160.78 |
| 77 | 2jsa-14 | VAL_A_13_C | VAL_A_13_O | -5.806 | 3.885 | 1.129 | -6.969 | 3.655 | 1.459 | 1.23 | 74.45 | -168.81 |
| 78 | 2jsa-14 | VAL_A_13_C | LYS_A_14_N | -5.806 | 3.885 | 1.129 | -5.449 | 4.478 | -0.001 | 1.33 | 148.51 | 58.95 |
| 79 | 2jsa-14 | LYS_A_14_N | LYS_A_14_H | -5.449 | 4.478 | -0.001 | -4.472 | 4.634 | -0.207 | 1.01 | 101.76 | 9.07 |
| 80 | 2jsa-14 | LYS_A_14_N | LYS_A_14_CA | -5.449 | 4.478 | -0.001 | -6.445 | 4.915 | -0.964 | 1.45 | 131.52 | 156.31 |
| 81 | 2jsa-14 | LYS_A_14_CA | LYS_A_14_HA | -6.445 | 4.915 | -0.964 | -6.955 | 5.785 | -0.551 | 1.09 | 67.73 | 120.38 |
| 82 | 2jsa-14 | LYS_A_14_CA | LYS_A_14_C | -6.445 | 4.915 | -0.964 | -7.487 | 3.81 | -1.15 | 1.53 | 96.98 | -133.32 |
| 83 | 2jsa-14 | LYS_A_14_C | LYS_A_14_O | -7.487 | 3.81 | -1.15 | -8.664 | 4.093 | -1.367 | 1.23 | 100.16 | 166.48 |
| 84 | 2jsa-14 | LYS_A_14_C | LYS_A_15_N | -7.487 | 3.81 | -1.15 | -7.017 | 2.575 | -1.057 | 1.32 | 85.97 | -69.16 |
| 85 | 2jsa-14 | LYS_A_15_N | LYS_A_15_H | -7.017 | 2.575 | -1.057 | -6.036 | 2.416 | -0.875 | 1.01 | 79.62 | -9.21 |
| 86 | 2jsa-14 | LYS_A_15_N | LYS_A_15_CA | -7.017 | 2.575 | -1.057 | -7.894 | 1.427 | -1.213 | 1.45 | 96.16 | -127.38 |
| 87 | 2jsa-14 | LYS_A_15_CA | LYS_A_15_HA | -7.894 | 1.427 | -1.213 | -8.605 | 1.651 | -2.008 | 1.09 | 136.84 | 162.51 |
| 88 | 2jsa-14 | LYS_A_15_CA | LYS_A_15_C | -7.894 | 1.427 | -1.213 | -8.689 | 1.22 | 0.077 | 1.53 | 32.49 | -165.41 |
| 89 | 2jsa-14 | LYS_A_15_C | LYS_A_15_O | -8.689 | 1.22 | 0.077 | -9.878 | 0.905 | 0.034 | 1.23 | 92.00 | -165.16 |
| 90 | 2jsa-14 | LYS_A_15_C | TYR_A_16_N | -8.689 | 1.22 | 0.077 | -8.003 | 1.406 | 1.195 | 1.32 | 32.45 | 15.17 |
| 91 | 2jsa-14 | TYR_A_16_N | TYR_A_16_H | -8.003 | 1.406 | 1.195 | -7.027 | 1.664 | 1.155 | 1.01 | 92.27 | 14.81 |
| 92 | 2jsa-14 | TYR_A_16_N | TYR_A_16_CA | -8.003 | 1.406 | 1.195 | -8.63 | 1.244 | 2.495 | 1.45 | 26.48 | -165.51 |
| 93 | 2jsa-14 | TYR_A_16_CA | TYR_A_16_HA | -8.63 | 1.244 | 2.495 | -9.2 | 0.316 | 2.525 | 1.09 | 88.42 | -121.56 |
| 94 | 2jsa-14 | TYR_A_16_CA | TYR_A_16_C | -8.63 | 1.244 | 2.495 | -9.547 | 2.427 | 2.814 | 1.53 | 77.97 | 127.78 |
| 95 | 2jsa-14 | TYR_A_16_C | TYR_A_16_O | -9.547 | 2.427 | 2.814 | -10.766 | 2.274 | 2.873 | 1.23 | 87.25 | -172.85 |
| 96 | 2jsa-14 | TYR_A_16_C | GLU_A_17_N | -9.547 | 2.427 | 2.814 | -8.924 | 3.581 | 3.008 | 1.33 | 81.59 | 61.64 |
| 97 | 2jsa-14 | GLU_A_17_N | GLU_A_17_H | -8.924 | 3.581 | 3.008 | -7.917 | 3.634 | 2.941 | 1.01 | 93.80 | 3.01 |
| 98 | 2jsa-14 | GLU_A_17_N | GLU_A_17_CA | -8.924 | 3.581 | 3.008 | -9.669 | 4.789 | 3.319 | 1.45 | 77.64 | 121.66 |
| 99 | 2jsa-14 | GLU_A_17_CA | GLU_A_17_HA | -9.669 | 4.789 | 3.319 | -9.941 | 4.756 | 4.374 | 1.09 | 14.56 | -173.08 |
| 100 | 2jsa-14 | GLU_A_17_CA | GLU_A_17_C | -9.669 | 4.789 | 3.319 | -10.955 | 4.847 | 2.491 | 1.53 | 122.75 | 177.42 |
| 101 | 2jsa-14 | GLU_A_17_C | GLU_A_17_O | -10.955 | 4.847 | 2.491 | -12.006 | 5.236 | 2.999 | 1.23 | 65.62 | 159.69 |
| 102 | 2jsa-14 | GLU_A_17_C | GLY_A_18_N | -10.955 | 4.847 | 2.491 | -10.83 | 4.455 | 1.232 | 1.32 | 161.90 | -72.31 |
| 103 | 2jsa-14 | GLY_A_18_N | GLY_A_18_H | -10.83 | 4.455 | 1.232 | -9.934 | 4.148 | 0.882 | 1.01 | 110.28 | -18.91 |
| 104 | 2jsa-14 | GLY_A_18_N | GLY_A_18_CA | -10.83 | 4.455 | 1.232 | -11.969 | 4.457 | 0.332 | 1.45 | 128.31 | 179.90 |
| 105 | 2jsa-14 | GLY_A_18_CA | GLY_A_18_C | -11.969 | 4.457 | 0.332 | -12.265 | 3.047 | -0.183 | 1.53 | 109.67 | -101.86 |
| 106 | 2jsa-14 | GLY_A_18_C | GLY_A_18_O | -12.265 | 3.047 | -0.183 | -12.429 | 2.842 | -1.385 | 1.23 | 167.68 | -128.66 |
| 107 | 2jsa-14 | GLY_A_18_C | SER_A_19_N | -12.265 | 3.047 | -0.183 | -12.323 | 2.11 | 0.752 | 1.32 | 45.12 | -93.54 |
| 108 | 2jsa-14 | SER_A_19_N | SER_A_19_H | -12.323 | 2.11 | 0.752 | -12.177 | 2.351 | 1.722 | 1.01 | 16.20 | 58.79 |
| 109 | 2jsa-14 | SER_A_19_N | SER_A_19_CA | -12.323 | 2.11 | 0.752 | -12.596 | 0.725 | 0.409 | 1.45 | 103.66 | -101.15 |
| 110 | 2jsa-14 | SER_A_19_CA | SER_A_19_HA | -12.596 | 0.725 | 0.409 | -12.083 | 0.498 | -0.526 | 1.09 | 149.04 | -23.87 |
| 111 | 2jsa-14 | SER_A_19_CA | SER_A_19_C | -12.596 | 0.725 | 0.409 | -14.099 | 0.522 | 0.206 | 1.53 | 97.62 | -172.31 |
| 112 | 2jsa-14 | SER_A_19_C | SER_A_19_O | -14.099 | 0.522 | 0.206 | -14.528 | 0.039 | -0.839 | 1.23 | 148.28 | -131.61 |
| 113 | 2jsa-14 | SER_A_19_C | ALA_A_20_N | -14.099 | 0.522 | 0.206 | -14.857 | 0.902 | 1.225 | 1.33 | 39.76 | 153.37 |
| 114 | 2jsa-14 | ALA_A_20_N | ALA_A_20_H | -14.857 | 0.902 | 1.225 | -14.434 | 1.291 | 2.056 | 1.01 | 34.67 | 42.60 |
| 115 | 2jsa-14 | ALA_A_20_N | ALA_A_20_CA | -14.857 | 0.902 | 1.225 | -16.303 | 0.768 | 1.172 | 1.45 | 92.09 | -174.71 |
| 116 | 2jsa-14 | ALA_A_20_CA | ALA_A_20_HA | -16.303 | 0.768 | 1.172 | -16.745 | 1.462 | 1.886 | 1.09 | 49.05 | 122.49 |
| 117 | 2jsa-14 | ALA_A_20_CA | ALA_A_20_C | -16.303 | 0.768 | 1.172 | -16.694 | -0.655 | 1.575 | 1.53 | 74.73 | -105.36 |
| 118 | 2jsa-14 | ALA_A_20_C | ALA_A_20_O | -16.694 | -0.655 | 1.575 | -17.875 | -0.955 | 1.739 | 1.23 | 82.33 | -165.75 |
| 119 | 2jsa-14 | ALA_A_20_C | ASP_A_21_N | -16.694 | -0.655 | 1.575 | -15.679 | -1.494 | 1.722 | 1.33 | 83.63 | -39.58 |
| 120 | 2jsa-14 | ASP_A_21_N | ASP_A_21_H | -15.679 | -1.494 | 1.722 | -14.731 | -1.179 | 1.569 | 1.01 | 98.71 | 18.38 |
| 121 | 2jsa-14 | ASP_A_21_N | ASP_A_21_CA | -15.679 | -1.494 | 1.722 | -15.902 | -2.878 | 2.103 | 1.45 | 74.80 | -99.15 |
| 122 | 2jsa-14 | ASP_A_21_CA | ASP_A_21_HA | -15.902 | -2.878 | 2.103 | -16.974 | -3.06 | 2.169 | 1.09 | 86.53 | -170.36 |
| 123 | 2jsa-14 | ASP_A_21_CA | ASP_A_21_C | -15.902 | -2.878 | 2.103 | -15.245 | -3.143 | 3.459 | 1.53 | 27.58 | -21.97 |
| 124 | 2jsa-14 | ASP_A_21_C | ASP_A_21_O | -15.245 | -3.143 | 3.459 | -15.449 | -4.198 | 4.056 | 1.23 | 60.94 | -100.94 |
| 125 | 2jsa-14 | ASP_A_21_C | LYS_A_22_N | -15.245 | -3.143 | 3.459 | -14.47 | -2.164 | 3.905 | 1.33 | 70.34 | 51.63 |
| 126 | 2jsa-14 | LYS_A_22_N | LYS_A_22_H | -14.47 | -2.164 | 3.905 | -14.348 | -1.323 | 3.36 | 1.01 | 122.67 | 81.75 |
| 127 | 2jsa-14 | LYS_A_22_N | LYS_A_22_CA | -14.47 | -2.164 | 3.905 | -13.782 | -2.277 | 5.18 | 1.45 | 28.67 | -9.33 |
| 128 | 2jsa-14 | LYS_A_22_CA | LYS_A_22_HA | -13.782 | -2.277 | 5.18 | -12.958 | -1.563 | 5.183 | 1.09 | 89.84 | 40.91 |
| 129 | 2jsa-14 | LYS_A_22_CA | LYS_A_22_C | -13.782 | -2.277 | 5.18 | -13.183 | -3.679 | 5.309 | 1.53 | 85.16 | -66.87 |
| 130 | 2jsa-14 | LYS_A_22_C | LYS_A_22_O | -13.183 | -3.679 | 5.309 | -13.655 | -4.489 | 6.106 | 1.23 | 49.63 | -120.23 |
| 131 | 2jsa-14 | LYS_A_22_C | ASP_A_23_N | -13.183 | -3.679 | 5.309 | -12.153 | -3.924 | 4.512 | 1.33 | 126.97 | -13.38 |
| 132 | 2jsa-14 | ASP_A_23_N | ASP_A_23_H | -12.153 | -3.924 | 4.512 | -11.818 | -3.211 | 3.88 | 1.01 | 128.74 | 64.83 |
| 133 | 2jsa-14 | ASP_A_23_N | ASP_A_23_CA | -12.153 | -3.924 | 4.512 | -11.485 | -5.214 | 4.527 | 1.45 | 89.41 | -62.62 |
| 134 | 2jsa-14 | ASP_A_23_CA | ASP_A_23_HA | -11.485 | -5.214 | 4.527 | -11.499 | -5.61 | 5.543 | 1.09 | 21.31 | -92.02 |

| | | | | | | | | | | | |
|---|---|---|---|---|---|---|---|---|---|---|---|
| 135 | 2jsa-14 | ASP_A_23_CA | ASP_A_23_C | -11.485 | -5.214 | 4.527 | -10.04 | -5.041 | 4.056 | 1.53 | 107.93 | 6.83 |
| 136 | 2jsa-14 | ASP_A_23_C | ASP_A_23_O | -10.04 | -5.041 | 4.056 | -9.792 | -4.789 | 2.877 | 1.23 | 163.31 | 45.46 |
| 137 | 2jsa-14 | ASP_A_23_C | ALA_A_24_N | -10.04 | -5.041 | 4.056 | -9.121 | -5.182 | 5.001 | 1.33 | 44.53 | -8.72 |
| 138 | 2jsa-14 | ALA_A_24_N | ALA_A_24_H | -9.121 | -5.182 | 5.001 | -9.396 | -5.388 | 5.951 | 1.01 | 19.88 | -143.16 |
| 139 | 2jsa-14 | ALA_A_24_N | ALA_A_24_CA | -9.121 | -5.182 | 5.001 | -7.707 | -5.044 | 4.697 | 1.45 | 102.08 | 5.57 |
| 140 | 2jsa-14 | ALA_A_24_CA | ALA_A_24_HA | -7.707 | -5.044 | 4.697 | -7.537 | -4.063 | 4.253 | 1.09 | 114.03 | 80.17 |
| 141 | 2jsa-14 | ALA_A_24_CA | ALA_A_24_C | -7.707 | -5.044 | 4.697 | -7.302 | -6.112 | 3.681 | 1.53 | 131.65 | -69.23 |
| 142 | 2jsa-14 | ALA_A_24_C | ALA_A_24_O | -7.302 | -6.112 | 3.681 | -6.219 | -6.044 | 3.1 | 1.23 | 118.17 | 3.59 |
| 143 | 2jsa-14 | ALA_A_24_C | ASN_A_25_N | -7.302 | -6.112 | 3.681 | -8.192 | -7.075 | 3.494 | 1.32 | 98.12 | -132.74 |
| 144 | 2jsa-14 | ASN_A_25_N | ASN_A_25_H | -8.192 | -7.075 | 3.494 | -9.062 | -7.068 | 4.006 | 1.01 | 59.52 | 179.54 |
| 145 | 2jsa-14 | ASN_A_25_N | ASN_A_25_CA | -8.192 | -7.075 | 3.494 | -7.941 | -8.156 | 2.556 | 1.45 | 130.21 | -76.93 |
| 146 | 2jsa-14 | ASN_A_25_CA | ASN_A_25_HA | -7.941 | -8.156 | 2.556 | -6.902 | -8.474 | 2.648 | 1.09 | 85.16 | -17.02 |
| 147 | 2jsa-14 | ASN_A_25_CA | ASN_A_25_C | -7.941 | -8.156 | 2.556 | -8.21 | -7.664 | 1.132 | 1.53 | 158.51 | 118.67 |
| 148 | 2jsa-14 | ASN_A_25_C | ASN_A_25_O | -8.21 | -7.664 | 1.132 | -7.47 | -7.996 | 0.208 | 1.23 | 138.72 | -24.16 |
| 149 | 2jsa-14 | ASN_A_25_C | VAL_A_26_N | -8.21 | -7.664 | 1.132 | -9.27 | -6.879 | 1.002 | 1.33 | 95.63 | 143.48 |
| 150 | 2jsa-14 | VAL_A_26_N | VAL_A_26_H | -9.27 | -6.879 | 1.002 | -9.833 | -6.647 | 1.808 | 1.01 | 37.07 | 157.60 |
| 151 | 2jsa-14 | VAL_A_26_N | VAL_A_26_CA | -9.27 | -6.879 | 1.002 | -9.645 | -6.338 | -0.293 | 1.45 | 153.06 | 124.73 |
| 152 | 2jsa-14 | VAL_A_26_CA | VAL_A_26_HA | -9.645 | -6.338 | -0.293 | -9.361 | -7.057 | -1.062 | 1.09 | 134.85 | -68.45 |
| 153 | 2jsa-14 | VAL_A_26_CA | VAL_A_26_C | -9.645 | -6.338 | -0.293 | -8.869 | -5.043 | -0.544 | 1.53 | 99.44 | 59.07 |
| 154 | 2jsa-14 | VAL_A_26_C | VAL_A_26_O | -8.869 | -5.043 | -0.544 | -8.965 | -4.456 | -1.62 | 1.23 | 151.07 | 99.29 |
| 155 | 2jsa-14 | VAL_A_26_C | ILE_A_27_N | -8.869 | -5.043 | -0.544 | -8.116 | -4.636 | 0.468 | 1.33 | 40.22 | 28.39 |
| 156 | 2jsa-14 | ILE_A_27_N | ILE_A_27_H | -8.116 | -4.636 | 0.468 | -8.082 | -5.17 | 1.325 | 1.01 | 31.98 | -86.36 |
| 157 | 2jsa-14 | ILE_A_27_N | ILE_A_27_CA | -8.116 | -4.636 | 0.468 | -7.325 | -3.422 | 0.371 | 1.45 | 93.83 | 56.91 |
| 158 | 2jsa-14 | ILE_A_27_CA | ILE_A_27_HA | -7.325 | -3.422 | 0.371 | -7.771 | -2.789 | -0.396 | 1.09 | 134.73 | 125.17 |
| 159 | 2jsa-14 | ILE_A_27_CA | ILE_A_27_C | -7.325 | -3.422 | 0.371 | -5.909 | -3.777 | -0.085 | 1.53 | 107.35 | -14.07 |
| 160 | 2jsa-14 | ILE_A_27_C | ILE_A_27_O | -5.909 | -3.777 | -0.085 | -5.39 | -3.184 | -1.031 | 1.23 | 140.20 | 48.81 |
| 161 | 2jsa-14 | ILE_A_27_C | LYS_A_28_N | -5.909 | -3.777 | -0.085 | -5.322 | -4.742 | 0.607 | 1.32 | 58.51 | -58.69 |
| 162 | 2jsa-14 | LYS_A_28_N | LYS_A_28_H | -5.322 | -4.742 | 0.607 | -5.808 | -5.186 | 1.373 | 1.01 | 40.67 | -137.59 |
| 163 | 2jsa-14 | LYS_A_28_N | LYS_A_28_CA | -5.322 | -4.742 | 0.607 | -3.976 | -5.183 | 0.285 | 1.45 | 102.81 | -18.14 |
| 164 | 2jsa-14 | LYS_A_28_CA | LYS_A_28_HA | -3.976 | -5.183 | 0.285 | -3.342 | -4.3 | 0.204 | 1.09 | 94.26 | 54.32 |
| 165 | 2jsa-14 | LYS_A_28_CA | LYS_A_28_C | -3.976 | -5.183 | 0.285 | -3.984 | -5.877 | -1.079 | 1.53 | 153.03 | -90.66 |
| 166 | 2jsa-14 | LYS_A_28_C | LYS_A_28_O | -3.984 | -5.877 | -1.079 | -2.995 | -5.834 | -1.808 | 1.23 | 126.37 | 2.49 |
| 167 | 2jsa-14 | LYS_A_28_C | LYS_A_29_N | -3.984 | -5.877 | -1.079 | -5.113 | -6.501 | -1.383 | 1.33 | 103.26 | -151.07 |
| 168 | 2jsa-14 | LYS_A_29_N | LYS_A_29_H | -5.113 | -6.501 | -1.383 | -5.887 | -6.495 | -0.735 | 1.01 | 50.06 | 179.56 |
| 169 | 2jsa-14 | LYS_A_29_N | LYS_A_29_CA | -5.113 | -6.501 | -1.383 | -5.263 | -7.203 | -2.646 | 1.45 | 150.39 | -102.06 |
| 170 | 2jsa-14 | LYS_A_29_CA | LYS_A_29_HA | -5.263 | -7.203 | -2.646 | -4.337 | -7.745 | -2.843 | 1.09 | 100.40 | -30.34 |
| 171 | 2jsa-14 | LYS_A_29_CA | LYS_A_29_C | -5.263 | -7.203 | -2.646 | -5.454 | -6.184 | -3.772 | 1.53 | 137.36 | 100.62 |
| 172 | 2jsa-14 | LYS_A_29_C | LYS_A_29_O | -5.454 | -6.184 | -3.772 | -4.78 | -6.254 | -4.799 | 1.23 | 146.58 | -5.93 |
| 173 | 2jsa-14 | LYS_A_29_C | ASP_A_30_N | -5.454 | -6.184 | -3.772 | -6.375 | -5.261 | -3.54 | 1.32 | 79.91 | 134.94 |
| 174 | 2jsa-14 | ASP_A_30_N | ASP_A_30_H | -6.375 | -5.261 | -3.54 | -6.891 | -5.264 | -2.672 | 1.01 | 30.73 | -179.67 |
| 175 | 2jsa-14 | ASP_A_30_N | ASP_A_30_CA | -6.375 | -5.261 | -3.54 | -6.664 | -4.228 | -4.522 | 1.45 | 132.47 | 105.63 |
| 176 | 2jsa-14 | ASP_A_30_CA | ASP_A_30_HA | -6.664 | -4.228 | -4.522 | -7.12 | -4.687 | -5.399 | 1.09 | 143.58 | -134.81 |
| 177 | 2jsa-14 | ASP_A_30_CA | ASP_A_30_C | -6.664 | -4.228 | -4.522 | -5.361 | -3.531 | -4.919 | 1.53 | 105.04 | 28.14 |
| 178 | 2jsa-14 | ASP_A_30_C | ASP_A_30_O | -5.361 | -3.531 | -4.919 | -5.065 | -3.389 | -6.105 | 1.23 | 164.53 | 25.63 |
| 179 | 2jsa-14 | ASP_A_30_C | PHE_A_31_N | -5.361 | -3.531 | -4.919 | -4.617 | -3.115 | -3.905 | 1.32 | 40.05 | 29.21 |
| 180 | 2jsa-14 | PHE_A_31_N | PHE_A_31_H | -4.617 | -3.115 | -3.905 | -4.922 | -3.266 | -2.954 | 1.01 | 19.69 | -153.66 |
| 181 | 2jsa-14 | PHE_A_31_N | PHE_A_31_CA | -4.617 | -3.115 | -3.905 | -3.353 | -2.436 | -4.134 | 1.45 | 99.07 | 28.24 |
| 182 | 2jsa-14 | PHE_A_31_CA | PHE_A_31_HA | -3.353 | -2.436 | -4.134 | -3.534 | -1.455 | -4.572 | 1.09 | 113.70 | 100.45 |
| 183 | 2jsa-14 | PHE_A_31_CA | PHE_A_31_C | -3.353 | -2.436 | -4.134 | -2.451 | -3.254 | -5.06 | 1.53 | 127.25 | -42.20 |
| 184 | 2jsa-14 | PHE_A_31_C | PHE_A_31_O | -2.451 | -3.254 | -5.06 | -1.749 | -2.695 | -5.902 | 1.23 | 133.18 | 38.53 |
| 185 | 2jsa-14 | PHE_A_31_C | ASP_A_32_N | -2.451 | -3.254 | -5.06 | -2.498 | -4.565 | -4.872 | 1.33 | 81.84 | -92.05 |
| 186 | 2jsa-14 | ASP_A_32_N | ASP_A_32_H | -2.498 | -4.565 | -4.872 | -3.098 | -4.956 | -4.159 | 1.01 | 45.13 | -146.91 |
| 187 | 2jsa-14 | ASP_A_32_N | ASP_A_32_CA | -2.498 | -4.565 | -4.872 | -1.693 | -5.466 | -5.679 | 1.45 | 123.74 | -48.22 |
| 188 | 2jsa-14 | ASP_A_32_CA | ASP_A_32_HA | -1.693 | -5.466 | -5.679 | -0.65 | -5.152 | -5.629 | 1.09 | 87.37 | 16.75 |
| 189 | 2jsa-14 | ASP_A_32_CA | ASP_A_32_C | -1.693 | -5.466 | -5.679 | -2.182 | -5.42 | -7.129 | 1.53 | 161.29 | 174.63 |
| 190 | 2jsa-14 | ASP_A_32_C | ASP_A_32_O | -2.182 | -5.42 | -7.129 | -1.401 | -5.615 | -8.058 | 1.23 | 139.09 | -14.02 |
| 191 | 2jsa-14 | ASP_A_32_C | ALA_A_33_N | -2.182 | -5.42 | -7.129 | -3.474 | -5.16 | -7.275 | 1.33 | 96.32 | 168.62 |
| 192 | 2jsa-14 | ALA_A_33_N | ALA_A_33_H | -3.474 | -5.16 | -7.275 | -4.057 | -5.009 | -6.464 | 1.01 | 36.60 | 165.48 |
| 193 | 2jsa-14 | ALA_A_33_N | ALA_A_33_CA | -3.474 | -5.16 | -7.275 | -4.077 | -5.087 | -8.595 | 1.45 | 155.29 | 173.10 |
| 194 | 2jsa-14 | ALA_A_33_CA | ALA_A_33_HA | -4.077 | -5.087 | -8.595 | -3.8 | -5.979 | -9.157 | 1.09 | 121.04 | -72.75 |
| 195 | 2jsa-14 | ALA_A_33_CA | ALA_A_33_C | -4.077 | -5.087 | -8.595 | -3.531 | -3.861 | -9.332 | 1.53 | 118.77 | 65.99 |
| 196 | 2jsa-14 | ALA_A_33_C | ALA_A_33_O | -3.531 | -3.861 | -9.332 | -2.732 | -3.993 | -10.257 | 1.23 | 138.80 | -9.38 |
| 197 | 2jsa-14 | ALA_A_33_C | GLU_A_34_N | -3.531 | -3.861 | -9.332 | -3.987 | -2.696 | -8.895 | 1.33 | 70.75 | 111.38 |
| 198 | 2jsa-14 | GLU_A_34_N | GLU_A_34_H | -3.987 | -2.696 | -8.895 | -4.644 | -2.663 | -8.129 | 1.01 | 40.66 | 177.12 |
| 199 | 2jsa-14 | GLU_A_34_N | GLU_A_34_CA | -3.987 | -2.696 | -8.895 | -3.556 | -1.448 | -9.501 | 1.45 | 114.65 | 70.95 |
| 200 | 2jsa-14 | GLU_A_34_CA | GLU_A_34_HA | -3.556 | -1.448 | -9.501 | -3.989 | -1.39 | -10.499 | 1.09 | 156.36 | 172.37 |
| 201 | 2jsa-14 | GLU_A_34_CA | GLU_A_34_C | -3.556 | -1.448 | -9.501 | -2.032 | -1.423 | -9.636 | 1.53 | 95.06 | 0.94 |
| 202 | 2jsa-14 | GLU_A_34_C | GLU_A_34_O | -2.032 | -1.423 | -9.636 | -1.502 | -0.966 | -10.648 | 1.23 | 145.34 | 40.77 |
| 203 | 2jsa-14 | GLU_A_34_C | CYS_A_35_N | -2.032 | -1.423 | -9.636 | -1.37 | -1.918 | -8.6 | 1.33 | 38.59 | -36.79 |
| 204 | 2jsa-14 | CYS_A_35_N | CYS_A_35_H | -1.37 | -1.918 | -8.6 | -1.871 | -2.276 | -7.8 | 1.01 | 37.59 | -144.45 |
| 205 | 2jsa-14 | CYS_A_35_N | CYS_A_35_CA | -1.37 | -1.918 | -8.6 | 0.082 | -1.959 | -8.59 | 1.45 | 89.61 | -1.62 |
| 206 | 2jsa-14 | CYS_A_35_CA | CYS_A_35_HA | 0.082 | -1.959 | -8.59 | 0.454 | -0.942 | -8.715 | 1.09 | 96.58 | 69.91 |
| 207 | 2jsa-14 | CYS_A_35_CA | CYS_A_35_C | 0.082 | -1.959 | -8.59 | 0.549 | -2.797 | -9.782 | 1.53 | 141.17 | -60.87 |
| 208 | 2jsa-14 | CYS_A_35_C | CYS_A_35_O | 0.549 | -2.797 | -9.782 | 1.33 | -2.328 | -10.608 | 1.23 | 132.20 | 30.99 |
| 209 | 2jsa-14 | CYS_A_35_C | LYS_A_36_N | 0.549 | -2.797 | -9.782 | 0.049 | -4.023 | -9.835 | 1.33 | 92.29 | -112.19 |
| 210 | 2jsa-14 | LYS_A_36_N | LYS_A_36_H | 0.049 | -4.023 | -9.835 | -0.589 | -4.342 | -9.121 | 1.01 | 44.97 | -153.43 |
| 211 | 2jsa-14 | LYS_A_36_N | LYS_A_36_CA | 0.049 | -4.023 | -9.835 | 0.404 | -4.931 | -10.912 | 1.45 | 137.85 | -68.65 |
| 212 | 2jsa-14 | LYS_A_36_CA | LYS_A_36_HA | 0.404 | -4.931 | -10.912 | 1.488 | -5.048 | -10.912 | 1.09 | 90.00 | -6.16 |
| 213 | 2jsa-14 | LYS_A_36_CA | LYS_A_36_C | 0.404 | -4.931 | -10.912 | 0.004 | -4.308 | -12.251 | 1.53 | 151.06 | 122.70 |
| 214 | 2jsa-14 | LYS_A_36_C | LYS_A_36_O | 0.004 | -4.308 | -12.251 | 0.296 | -4.863 | -13.309 | 1.23 | 149.34 | -62.25 |
| 215 | 2jsa-14 | LYS_A_36_C | LYS_A_37_N | 0.004 | -4.308 | -12.251 | -0.66 | -3.164 | -12.16 | 1.33 | 86.06 | 120.13 |
| 216 | 2jsa-14 | LYS_A_37_N | LYS_A_37_H | -0.66 | -3.164 | -12.16 | -0.866 | -2.766 | -11.255 | 1.01 | 26.34 | 117.37 |
| 217 | 2jsa-14 | LYS_A_37_N | LYS_A_37_CA | -0.66 | -3.164 | -12.16 | -1.104 | -2.461 | -13.351 | 1.45 | 145.08 | 122.28 |
| 218 | 2jsa-14 | LYS_A_37_CA | LYS_A_37_HA | -1.104 | -2.461 | -13.351 | -1.057 | -3.155 | -14.19 | 1.09 | 140.34 | -86.13 |
| 219 | 2jsa-14 | LYS_A_37_CA | LYS_A_37_C | -1.104 | -2.461 | -13.351 | -0.141 | -1.309 | -13.647 | 1.53 | 101.15 | 50.11 |
| 220 | 2jsa-14 | LYS_A_37_C | LYS_A_37_O | -0.141 | -1.309 | -13.647 | 0.675 | -1.396 | -14.563 | 1.23 | 138.14 | -6.09 |
| 221 | 2jsa-14 | LYS_A_37_C | LEU_A_38_N | -0.141 | -1.309 | -13.647 | -0.267 | -0.256 | -12.853 | 1.32 | 53.18 | 96.82 |
| 222 | 2jsa-14 | LEU_A_38_N | LEU_A_38_H | -0.267 | -0.256 | -12.853 | -0.96 | -0.253 | -12.118 | 1.01 | 43.32 | 179.75 |
| 223 | 2jsa-14 | LEU_A_38_N | LEU_A_38_CA | -0.267 | -0.256 | -12.853 | 0.581 | 0.912 | -13.017 | 1.45 | 96.48 | 54.02 |
| 224 | 2jsa-14 | LEU_A_38_CA | LEU_A_38_HA | 0.581 | 0.912 | -13.017 | 0.481 | 1.261 | -14.045 | 1.09 | 160.55 | 105.99 |
| 225 | 2jsa-14 | LEU_A_38_CA | LEU_A_38_C | 0.581 | 0.912 | -13.017 | 2.041 | 0.509 | -12.804 | 1.53 | 81.99 | -15.43 |
| 226 | 2jsa-14 | LEU_A_38_C | LEU_A_38_O | 2.041 | 0.509 | -12.804 | 2.907 | 0.848 | -13.61 | 1.23 | 130.91 | 21.38 |
| 227 | 2jsa-14 | LEU_A_38_C | PHE_A_39_N | 2.041 | 0.509 | -12.804 | 2.272 | -0.207 | -11.713 | 1.33 | 34.59 | -72.12 |
| 228 | 2jsa-14 | PHE_A_39_N | PHE_A_39_H | 2.272 | -0.207 | -11.713 | 1.512 | -0.448 | -11.093 | 1.01 | 52.13 | -162.41 |
| 229 | 2jsa-14 | PHE_A_39_N | PHE_A_39_CA | 2.272 | -0.207 | -11.713 | 3.612 | -0.66 | -11.383 | 1.45 | 76.87 | -18.68 |
| 230 | 2jsa-14 | PHE_A_39_CA | PHE_A_39_HA | 3.612 | -0.66 | -11.383 | 4.341 | 0.097 | -11.674 | 1.09 | 105.48 | 46.08 |

| | | | | | | | | | | | |
|---|---|---|---|---|---|---|---|---|---|---|---|
| 231 | 2jsa-14 | PHE_A_39_CA | PHE_A_39_C | 3.612 | -0.66 | -11.383 | 3.934 | -1.98 | -12.086 | 1.53 | 117.36 | -76.29 |
| 232 | 2jsa-14 | PHE_A_39_C | PHE_A_39_O | 3.934 | -1.98 | -12.086 | 4.644 | -2.822 | -11.538 | 1.23 | 63.55 | -49.86 |
| 233 | 2jsa-14 | PHE_A_39_C | HIS_A_40_N | 3.934 | -1.98 | -12.086 | 3.396 | -2.119 | -13.289 | 1.33 | 155.21 | -165.51 |
| 234 | 2jsa-14 | HIS_A_40_N | HIS_A_40_H | 3.396 | -2.119 | -13.289 | 2.822 | -1.385 | -13.677 | 1.01 | 112.61 | 128.03 |
| 235 | 2jsa-14 | HIS_A_40_N | HIS_A_40_CA | 3.396 | -2.119 | -13.289 | 3.618 | -3.323 | -14.072 | 1.45 | 122.60 | -79.55 |
| 236 | 2jsa-14 | HIS_A_40_CA | HIS_A_40_HA | 3.618 | -3.323 | -14.072 | 3.794 | -4.147 | -13.38 | 1.09 | 50.60 | -77.94 |
| 237 | 2jsa-14 | HIS_A_40_CA | HIS_A_40_C | 3.618 | -3.323 | -14.072 | 4.874 | -3.154 | -14.93 | 1.53 | 124.10 | 7.66 |
| 238 | 2jsa-14 | HIS_A_40_C | HIS_A_40_O | 4.874 | -3.154 | -14.93 | 5.631 | -4.104 | -15.121 | 1.23 | 98.94 | -51.45 |
| 239 | 2jsa-14 | HIS_A_40_C | THR_A_41_N | 4.874 | -3.154 | -14.93 | 5.057 | -1.937 | -15.421 | 1.33 | 111.75 | 81.45 |
| 240 | 2jsa-14 | THR_A_41_N | THR_A_41_H | 5.057 | -1.937 | -15.421 | 4.394 | -1.202 | -15.22 | 1.01 | 78.52 | 132.05 |
| 241 | 2jsa-14 | THR_A_41_N | THR_A_41_CA | 5.057 | -1.937 | -15.421 | 6.208 | -1.631 | -16.253 | 1.45 | 124.94 | 14.89 |
| 242 | 2jsa-14 | THR_A_41_CA | THR_A_41_HA | 6.208 | -1.631 | -16.253 | 6.237 | -2.336 | -17.084 | 1.09 | 139.67 | -87.64 |
| 243 | 2jsa-14 | THR_A_41_CA | THR_A_41_C | 6.208 | -1.631 | -16.253 | 7.502 | -1.794 | -15.453 | 1.53 | 58.48 | -7.18 |
| 244 | 2jsa-14 | THR_A_41_C | THR_A_41_O | 7.502 | -1.794 | -15.453 | 8.565 | -2.029 | -16.025 | 1.23 | 117.72 | -12.47 |
| 245 | 2jsa-14 | THR_A_41_C | ILE_A_42_N | 7.502 | -1.794 | -15.453 | 7.369 | -1.662 | -14.141 | 1.33 | 8.13 | 135.22 |
| 246 | 2jsa-14 | ILE_A_42_N | ILE_A_42_H | 7.369 | -1.662 | -14.141 | 6.463 | -1.47 | -13.738 | 1.01 | 66.48 | 168.03 |
| 247 | 2jsa-14 | ILE_A_42_N | ILE_A_42_CA | 7.369 | -1.662 | -14.141 | 8.515 | -1.791 | -13.257 | 1.45 | 52.53 | -6.42 |
| 248 | 2jsa-14 | ILE_A_42_CA | ILE_A_42_HA | 8.515 | -1.791 | -13.257 | 9.284 | -1.098 | -13.598 | 1.09 | 108.23 | 42.02 |
| 249 | 2jsa-14 | ILE_A_42_CA | ILE_A_42_C | 8.515 | -1.791 | -13.257 | 9.083 | -3.207 | -13.368 | 1.53 | 94.16 | -68.14 |
| 250 | 2jsa-14 | ILE_A_42_C | ILE_A_42_O | 9.083 | -3.207 | -13.368 | 8.331 | -4.176 | -13.457 | 1.23 | 94.15 | -127.81 |
| 251 | 2jsa-14 | ILE_A_42_C | PRO_A_43_N | 9.083 | -3.207 | -13.368 | 10.441 | -3.284 | -13.36 | 1.36 | 89.66 | -3.25 |
| 252 | 2jsa-14 | PRO_A_43_N | PRO_A_43_CA | 10.441 | -3.284 | -13.36 | 11.117 | -4.567 | -13.459 | 1.45 | 93.91 | -62.22 |
| 253 | 2jsa-14 | PRO_A_43_CA | PRO_A_43_HA | 11.117 | -4.567 | -13.459 | 10.659 | -5.171 | -14.242 | 1.09 | 135.93 | -127.17 |
| 254 | 2jsa-14 | PRO_A_43_CA | PRO_A_43_C | 11.117 | -4.567 | -13.459 | 11.031 | -5.336 | -12.139 | 1.53 | 30.38 | -96.38 |
| 255 | 2jsa-14 | PRO_A_43_C | PRO_A_43_O | 11.031 | -5.336 | -12.139 | 11.556 | -6.442 | -12.025 | 1.23 | 84.68 | -64.61 |
| 256 | 2jsa-14 | PRO_A_43_C | PHE_A_44_N | 11.031 | -5.336 | -12.139 | 10.364 | -4.719 | -11.174 | 1.33 | 43.28 | 137.23 |
| 257 | 2jsa-14 | PHE_A_44_N | PHE_A_44_H | 10.364 | -4.719 | -11.174 | 9.958 | -3.809 | -11.338 | 1.01 | 99.35 | 114.04 |
| 258 | 2jsa-14 | PHE_A_44_N | PHE_A_44_CA | 10.364 | -4.719 | -11.174 | 10.202 | -5.332 | -9.867 | 1.45 | 25.88 | -104.80 |
| 259 | 2jsa-14 | PHE_A_44_CA | PHE_A_44_HA | 10.202 | -5.332 | -9.867 | 10.116 | -6.413 | -9.973 | 1.09 | 95.58 | -94.55 |
| 260 | 2jsa-14 | PHE_A_44_CA | PHE_A_44_C | 10.202 | -5.332 | -9.867 | 8.964 | -4.784 | -9.154 | 1.53 | 62.23 | 156.12 |
| 261 | 2jsa-14 | PHE_A_44_C | PHE_A_44_O | 8.964 | -4.784 | -9.154 | 8.845 | -4.895 | -7.936 | 1.23 | 7.61 | -136.99 |
| 262 | 2jsa-14 | PHE_A_44_C | GLY_A_45_N | 8.964 | -4.784 | -9.154 | 8.073 | -4.206 | -9.946 | 1.32 | 126.71 | 147.03 |
| 263 | 2jsa-14 | GLY_A_45_N | GLY_A_45_H | 8.073 | -4.206 | -9.946 | 8.237 | -4.15 | -10.941 | 1.01 | 170.12 | 18.85 |
| 264 | 2jsa-14 | GLY_A_45_N | GLY_A_45_CA | 8.073 | -4.206 | -9.946 | 6.848 | -3.642 | -9.406 | 1.45 | 68.18 | 155.28 |
| 265 | 2jsa-14 | GLY_A_45_CA | GLY_A_45_C | 6.848 | -3.642 | -9.406 | 5.934 | -4.739 | -8.856 | 1.53 | 68.93 | -129.80 |
| 266 | 2jsa-14 | GLY_A_45_C | GLY_A_45_O | 5.934 | -4.739 | -8.856 | 5.585 | -4.728 | -7.676 | 1.23 | 16.48 | 178.19 |
| 267 | 2jsa-14 | GLY_A_45_C | THR_A_46_N | 5.934 | -4.739 | -8.856 | 5.572 | -5.66 | -9.736 | 1.32 | 131.65 | -111.46 |
| 268 | 2jsa-14 | THR_A_46_N | THR_A_46_H | 5.572 | -5.66 | -9.736 | 5.898 | -5.606 | -10.691 | 1.01 | 160.91 | 9.41 |
| 269 | 2jsa-14 | THR_A_46_N | THR_A_46_CA | 5.572 | -5.66 | -9.736 | 4.706 | -6.762 | -9.354 | 1.45 | 74.75 | -128.16 |
| 270 | 2jsa-14 | THR_A_46_CA | THR_A_46_HA | 4.706 | -6.762 | -9.354 | 3.74 | -6.359 | -9.049 | 1.09 | 73.75 | 157.35 |
| 271 | 2jsa-14 | THR_A_46_CA | THR_A_46_C | 4.706 | -6.762 | -9.354 | 5.299 | -7.517 | -8.162 | 1.53 | 38.85 | -51.85 |
| 272 | 2jsa-14 | THR_A_46_C | THR_A_46_O | 5.299 | -7.517 | -8.162 | 4.564 | -8.013 | -7.309 | 1.23 | 46.11 | -145.99 |
| 273 | 2jsa-14 | THR_A_46_C | ARG_A_47_N | 5.299 | -7.517 | -8.162 | 6.621 | -7.579 | -8.14 | 1.32 | 89.05 | -2.69 |
| 274 | 2jsa-14 | ARG_A_47_N | ARG_A_47_H | 6.621 | -7.579 | -8.14 | 7.163 | -7.148 | -8.875 | 1.01 | 136.71 | 38.49 |
| 275 | 2jsa-14 | ARG_A_47_N | ARG_A_47_CA | 6.621 | -7.579 | -8.14 | 7.321 | -8.264 | -7.067 | 1.45 | 42.39 | -44.38 |
| 276 | 2jsa-14 | ARG_A_47_CA | ARG_A_47_HA | 7.321 | -8.264 | -7.067 | 6.924 | -9.276 | -6.989 | 1.09 | 85.90 | -111.42 |
| 277 | 2jsa-14 | ARG_A_47_CA | ARG_A_47_C | 7.321 | -8.264 | -7.067 | 7.098 | -7.538 | -5.739 | 1.53 | 29.77 | 107.07 |
| 278 | 2jsa-14 | ARG_A_47_C | ARG_A_47_O | 7.098 | -7.538 | -5.739 | 6.75 | -8.162 | -4.737 | 1.23 | 35.49 | -119.15 |
| 279 | 2jsa-14 | ARG_A_47_C | GLU_A_48_N | 7.098 | -7.538 | -5.739 | 7.307 | -6.23 | -5.774 | 1.33 | 91.51 | 80.92 |
| 280 | 2jsa-14 | GLU_A_48_N | GLU_A_48_H | 7.307 | -6.23 | -5.774 | 7.592 | -5.783 | -6.634 | 1.01 | 148.35 | 57.48 |
| 281 | 2jsa-14 | GLU_A_48_N | GLU_A_48_CA | 7.307 | -6.23 | -5.774 | 7.133 | -5.412 | -4.586 | 1.45 | 35.14 | 102.01 |
| 282 | 2jsa-14 | GLU_A_48_CA | GLU_A_48_HA | 7.133 | -5.412 | -4.586 | 7.646 | -5.905 | -3.759 | 1.09 | 40.71 | -43.86 |
| 283 | 2jsa-14 | GLU_A_48_CA | GLU_A_48_C | 7.133 | -5.412 | -4.586 | 5.649 | -5.299 | -4.232 | 1.53 | 76.62 | 175.65 |
| 284 | 2jsa-14 | GLU_A_48_C | GLU_A_48_O | 5.649 | -5.299 | -4.232 | 5.298 | -5.072 | -3.075 | 1.23 | 19.86 | 147.11 |
| 285 | 2jsa-14 | GLU_A_48_C | CYS_A_49_N | 5.649 | -5.299 | -4.232 | 4.816 | -5.462 | -5.25 | 1.33 | 140.18 | -168.93 |
| 286 | 2jsa-14 | CYS_A_49_N | CYS_A_49_H | 4.816 | -5.462 | -5.25 | 5.175 | -5.646 | -6.176 | 1.01 | 156.46 | -27.14 |
| 287 | 2jsa-14 | CYS_A_49_N | CYS_A_49_CA | 4.816 | -5.462 | -5.25 | 3.378 | -5.381 | -5.06 | 1.45 | 82.49 | 176.78 |
| 288 | 2jsa-14 | CYS_A_49_CA | CYS_A_49_HA | 3.378 | -5.381 | -5.06 | 3.131 | -4.367 | -4.744 | 1.09 | 73.15 | 103.69 |
| 289 | 2jsa-14 | CYS_A_49_CA | CYS_A_49_C | 3.378 | -5.381 | -5.06 | 2.985 | -6.349 | -3.943 | 1.53 | 43.09 | -112.10 |
| 290 | 2jsa-14 | CYS_A_49_C | CYS_A_49_O | 2.985 | -6.349 | -3.943 | 2.369 | -5.947 | -2.958 | 1.23 | 36.75 | 146.87 |
| 291 | 2jsa-14 | CYS_A_49_C | ASP_A_50_N | 2.985 | -6.349 | -3.943 | 3.358 | -7.607 | -4.134 | 1.33 | 98.28 | -73.48 |
| 292 | 2jsa-14 | ASP_A_50_N | ASP_A_50_H | 3.358 | -7.607 | -4.134 | 3.864 | -7.866 | -4.969 | 1.01 | 145.75 | -27.11 |
| 293 | 2jsa-14 | ASP_A_50_N | ASP_A_50_CA | 3.358 | -7.607 | -4.134 | 3.051 | -8.635 | -3.156 | 1.45 | 47.65 | -106.63 |
| 294 | 2jsa-14 | ASP_A_50_CA | ASP_A_50_HA | 3.051 | -8.635 | -3.156 | 2.014 | -8.527 | -2.839 | 1.09 | 73.09 | 174.05 |
| 295 | 2jsa-14 | ASP_A_50_CA | ASP_A_50_C | 3.051 | -8.635 | -3.156 | 3.983 | -8.48 | -1.951 | 1.53 | 38.10 | 9.44 |
| 296 | 2jsa-14 | ASP_A_50_C | ASP_A_50_O | 3.983 | -8.48 | -1.951 | 3.721 | -9.032 | -0.884 | 1.23 | 29.80 | -115.39 |
| 297 | 2jsa-14 | ASP_A_50_C | HIS_A_51_N | 3.983 | -8.48 | -1.951 | 5.051 | -7.724 | -2.163 | 1.33 | 99.20 | 35.29 |
| 298 | 2jsa-14 | HIS_A_51_N | HIS_A_51_H | 5.051 | -7.724 | -2.163 | 5.203 | -7.299 | -3.067 | 1.01 | 153.47 | 70.32 |
| 299 | 2jsa-14 | HIS_A_51_N | HIS_A_51_CA | 5.051 | -7.724 | -2.163 | 6.023 | -7.49 | -1.109 | 1.45 | 43.49 | 13.54 |
| 300 | 2jsa-14 | HIS_A_51_CA | HIS_A_51_HA | 6.023 | -7.49 | -1.109 | 6.037 | -8.368 | -0.462 | 1.09 | 53.62 | -89.09 |
| 301 | 2jsa-14 | HIS_A_51_CA | HIS_A_51_C | 6.023 | -7.49 | -1.109 | 5.584 | -6.291 | -0.265 | 1.53 | 56.54 | 110.11 |
| 302 | 2jsa-14 | HIS_A_51_C | HIS_A_51_O | 5.584 | -6.291 | -0.265 | 6.286 | -5.89 | 0.662 | 1.23 | 41.09 | 29.74 |
| 303 | 2jsa-14 | HIS_A_51_C | TYR_A_52_N | 5.584 | -6.291 | -0.265 | 4.426 | -5.752 | -0.616 | 1.32 | 105.37 | 155.04 |
| 304 | 2jsa-14 | TYR_A_52_N | TYR_A_52_H | 4.426 | -5.752 | -0.616 | 3.902 | -6.133 | -1.391 | 1.01 | 140.11 | -143.98 |
| 305 | 2jsa-14 | TYR_A_52_N | TYR_A_52_CA | 4.426 | -5.752 | -0.616 | 3.886 | -4.608 | 0.097 | 1.45 | 60.59 | 115.27 |
| 306 | 2jsa-14 | TYR_A_52_CA | TYR_A_52_HA | 3.886 | -4.608 | 0.097 | 4.613 | -4.235 | 0.819 | 1.09 | 48.54 | 27.16 |
| 307 | 2jsa-14 | TYR_A_52_CA | TYR_A_52_C | 3.886 | -4.608 | 0.097 | 2.584 | -4.972 | 0.812 | 1.53 | 62.13 | -164.38 |
| 308 | 2jsa-14 | TYR_A_52_C | TYR_A_52_O | 2.584 | -4.972 | 0.812 | 2.37 | -4.579 | 1.958 | 1.23 | 21.33 | 118.57 |
| 309 | 2jsa-14 | TYR_A_52_C | VAL_A_53_N | 2.584 | -4.972 | 0.812 | 1.746 | -5.718 | 0.106 | 1.33 | 122.18 | -138.32 |
| 310 | 2jsa-14 | VAL_A_53_N | VAL_A_53_H | 1.746 | -5.718 | 0.106 | 1.987 | -6.004 | -0.832 | 1.01 | 158.26 | -49.88 |
| 311 | 2jsa-14 | VAL_A_53_N | VAL_A_53_CA | 1.746 | -5.718 | 0.106 | 0.471 | -6.139 | 0.66 | 1.45 | 67.58 | -161.73 |
| 312 | 2jsa-14 | VAL_A_53_CA | VAL_A_53_HA | 0.471 | -6.139 | 0.66 | 0.081 | -5.331 | 1.28 | 1.09 | 55.35 | 115.77 |
| 313 | 2jsa-14 | VAL_A_53_CA | VAL_A_53_C | 0.471 | -6.139 | 0.66 | 0.686 | -7.362 | 1.553 | 1.53 | 54.28 | -80.03 |
| 314 | 2jsa-14 | VAL_A_53_C | VAL_A_53_O | 0.686 | -7.362 | 1.553 | -0.025 | -8.358 | 1.43 | 1.23 | 95.74 | -125.52 |
| 315 | 2jsa-14 | VAL_A_53_C | ASN_A_54_N | 0.686 | -7.362 | 1.553 | 1.671 | -7.246 | 2.432 | 1.33 | 48.45 | 6.72 |
| 316 | 2jsa-14 | ASN_A_54_N | ASN_A_54_H | 1.671 | -7.246 | 2.432 | 2.216 | -6.396 | 2.477 | 1.01 | 87.45 | 57.33 |
| 317 | 2jsa-14 | ASN_A_54_N | ASN_A_54_CA | 1.671 | -7.246 | 2.432 | 1.989 | -8.33 | 3.346 | 1.45 | 51.02 | -73.65 |
| 318 | 2jsa-14 | ASN_A_54_CA | ASN_A_54_HA | 1.989 | -8.33 | 3.346 | 1.146 | -8.489 | 4.018 | 1.09 | 51.93 | -169.32 |
| 319 | 2jsa-14 | ASN_A_54_CA | ASN_A_54_C | 1.989 | -8.33 | 3.346 | 3.235 | -7.961 | 4.154 | 1.53 | 58.13 | 16.50 |
| 320 | 2jsa-14 | ASN_A_54_C | ASN_A_54_O | 3.235 | -7.961 | 4.154 | 4.158 | -8.763 | 4.281 | 1.23 | 84.07 | -40.99 |
| 321 | 2jsa-14 | ASN_A_54_C | SER_A_55_N | 3.235 | -7.961 | 4.154 | 3.221 | -6.745 | 4.681 | 1.33 | 66.57 | 90.66 |
| 322 | 2jsa-14 | SER_A_55_N | SER_A_55_H | 3.221 | -6.745 | 4.681 | 2.427 | -6.138 | 4.537 | 1.01 | 98.20 | 142.60 |
| 323 | 2jsa-14 | SER_A_55_N | SER_A_55_CA | 3.221 | -6.745 | 4.681 | 4.337 | -6.259 | 5.473 | 1.45 | 56.95 | 23.53 |
| 324 | 2jsa-14 | SER_A_55_CA | SER_A_55_HA | 4.337 | -6.259 | 5.473 | 4.224 | -6.636 | 6.49 | 1.09 | 21.16 | -106.69 |
| 325 | 2jsa-14 | SER_A_55_CA | SER_A_55_C | 4.337 | -6.259 | 5.473 | 4.325 | -4.73 | 5.514 | 1.53 | 88.46 | 90.45 |
| 326 | 2jsa-14 | SER_A_55_C | SER_A_55_O | 4.325 | -4.73 | 5.514 | 4.875 | -4.125 | 6.433 | 1.23 | 41.66 | 47.73 |

| | | | | | | | | | | |
|---|---|---|---|---|---|---|---|---|---|---|
| 327 | 2jsa-14 | SER_A_55_C | LYS_A_56_N | 4.325 | -4.73 | 5.514 | 3.692 | -4.149 | 4.505 | 1.33 | 139.58 | 137.45 |
| 328 | 2jsa-14 | LYS_A_56_N | LYS_A_56_H | 3.692 | -4.149 | 4.505 | 3.263 | -4.71 | 3.783 | 1.01 | 135.63 | -127.41 |
| 329 | 2jsa-14 | LYS_A_56_N | LYS_A_56_CA | 3.692 | -4.149 | 4.505 | 3.601 | -2.701 | 4.414 | 1.45 | 93.59 | 93.60 |
| 330 | 2jsa-14 | LYS_A_56_CA | LYS_A_56_HA | 3.601 | -2.701 | 4.414 | 4.081 | -2.276 | 5.296 | 1.09 | 36.01 | 41.52 |
| 331 | 2jsa-14 | LYS_A_56_CA | LYS_A_56_C | 3.601 | -2.701 | 4.414 | 2.128 | -2.285 | 4.439 | 1.53 | 89.06 | 164.23 |
| 332 | 2jsa-14 | LYS_A_56_C | LYS_A_56_O | 2.128 | -2.285 | 4.439 | 1.817 | -1.098 | 4.518 | 1.23 | 86.32 | 104.68 |
| 333 | 2jsa-14 | LYS_A_56_C | VAL_A_57_N | 2.128 | -2.285 | 4.439 | 1.262 | -3.285 | 4.371 | 1.32 | 92.94 | -130.89 |
| 334 | 2jsa-14 | VAL_A_57_N | VAL_A_57_H | 1.262 | -3.285 | 4.371 | 1.59 | -4.238 | 4.308 | 1.01 | 93.58 | -71.01 |
| 335 | 2jsa-14 | VAL_A_57_N | VAL_A_57_CA | 1.262 | -3.285 | 4.371 | -0.17 | -3.038 | 4.386 | 1.45 | 89.41 | 170.21 |
| 336 | 2jsa-14 | VAL_A_57_CA | VAL_A_57_HA | -0.17 | -3.038 | 4.386 | -0.343 | -1.999 | 4.105 | 1.09 | 104.94 | 99.45 |
| 337 | 2jsa-14 | VAL_A_57_CA | VAL_A_57_C | -0.17 | -3.038 | 4.386 | -0.703 | -3.24 | 5.805 | 1.53 | 21.88 | -159.24 |
| 338 | 2jsa-14 | VAL_A_57_C | VAL_A_57_O | -0.703 | -3.24 | 5.805 | -1.435 | -2.398 | 6.323 | 1.23 | 65.10 | 131.00 |
| 339 | 2jsa-14 | VAL_A_57_C | ASP_A_58_N | -0.703 | -3.24 | 5.805 | -0.318 | -4.363 | 6.393 | 1.32 | 63.65 | -71.08 |
| 340 | 2jsa-14 | ASP_A_58_N | ASP_A_58_H | -0.318 | -4.363 | 6.393 | 0.283 | -5.012 | 5.906 | 1.01 | 118.84 | -47.20 |
| 341 | 2jsa-14 | ASP_A_58_N | ASP_A_58_CA | -0.318 | -4.363 | 6.393 | -0.748 | -4.687 | 7.743 | 1.45 | 21.74 | -143.00 |
| 342 | 2jsa-14 | ASP_A_58_CA | ASP_A_58_HA | -0.748 | -4.687 | 7.743 | -1.818 | -4.895 | 7.736 | 1.09 | 90.37 | -169.00 |
| 343 | 2jsa-14 | ASP_A_58_CA | ASP_A_58_C | -0.748 | -4.687 | 7.743 | -0.455 | -3.502 | 8.665 | 1.53 | 52.94 | 76.11 |
| 344 | 2jsa-14 | ASP_A_58_C | ASP_A_58_O | -0.455 | -3.502 | 8.665 | -1.293 | -3.126 | 9.484 | 1.23 | 48.28 | 155.83 |
| 345 | 2jsa-14 | ASP_A_58_C | PRO_A_59_N | -0.455 | -3.502 | 8.665 | 0.768 | -2.932 | 8.498 | 1.36 | 97.06 | 24.99 |
| 346 | 2jsa-14 | PRO_A_59_N | PRO_A_59_CA | 0.768 | -2.932 | 8.498 | 1.183 | -1.797 | 9.305 | 1.45 | 56.27 | 69.92 |
| 347 | 2jsa-14 | PRO_A_59_CA | PRO_A_59_HA | 1.183 | -1.797 | 9.305 | 0.935 | -1.973 | 10.352 | 1.09 | 16.20 | -144.64 |
| 348 | 2jsa-14 | PRO_A_59_CA | PRO_A_59_C | 1.183 | -1.797 | 9.305 | 0.483 | -0.515 | 8.849 | 1.53 | 107.34 | 118.64 |
| 349 | 2jsa-14 | PRO_A_59_C | PRO_A_59_O | 0.483 | -0.515 | 8.849 | -0.183 | 0.15 | 9.642 | 1.23 | 49.88 | 135.04 |
| 350 | 2jsa-14 | PRO_A_59_C | ILE_A_60_N | 0.483 | -0.515 | 8.849 | 0.658 | -0.205 | 7.573 | 1.32 | 164.41 | 60.55 |
| 351 | 2jsa-14 | ILE_A_60_N | ILE_A_60_H | 0.658 | -0.205 | 7.573 | 1.219 | -0.798 | 6.978 | 1.01 | 126.09 | -46.59 |
| 352 | 2jsa-14 | ILE_A_60_N | ILE_A_60_CA | 0.658 | -0.205 | 7.573 | 0.052 | 0.986 | 7.002 | 1.45 | 113.14 | 116.97 |
| 353 | 2jsa-14 | ILE_A_60_CA | ILE_A_60_HA | 0.052 | 0.986 | 7.002 | 0.599 | 1.854 | 7.372 | 1.09 | 70.17 | 57.78 |
| 354 | 2jsa-14 | ILE_A_60_CA | ILE_A_60_C | 0.052 | 0.986 | 7.002 | -1.392 | 1.103 | 7.493 | 1.53 | 71.28 | 175.37 |
| 355 | 2jsa-14 | ILE_A_60_C | ILE_A_60_O | -1.392 | 1.103 | 7.493 | -1.775 | 2.118 | 8.072 | 1.23 | 61.91 | 110.67 |
| 356 | 2jsa-14 | ILE_A_60_C | ILE_A_61_N | -1.392 | 1.103 | 7.493 | -2.156 | 0.049 | 7.242 | 1.33 | 100.91 | -125.94 |
| 357 | 2jsa-14 | ILE_A_61_N | ILE_A_61_H | -2.156 | 0.049 | 7.242 | -1.774 | -0.754 | 6.762 | 1.01 | 118.36 | -64.56 |
| 358 | 2jsa-14 | ILE_A_61_N | ILE_A_61_CA | -2.156 | 0.049 | 7.242 | -3.551 | 0.023 | 7.651 | 1.45 | 73.66 | -178.93 |
| 359 | 2jsa-14 | ILE_A_61_CA | ILE_A_61_HA | -3.551 | 0.023 | 7.651 | -4.055 | 0.87 | 7.187 | 1.09 | 115.21 | 120.75 |
| 360 | 2jsa-14 | ILE_A_61_CA | ILE_A_61_C | -3.551 | 0.023 | 7.651 | -3.637 | 0.204 | 9.167 | 1.53 | 7.53 | 115.41 |
| 361 | 2jsa-14 | ILE_A_61_C | ILE_A_61_O | -3.637 | 0.204 | 9.167 | -4.653 | 0.664 | 9.685 | 1.23 | 65.09 | 155.64 |
| 362 | 2jsa-14 | ILE_A_61_C | HIS_A_62_N | -3.637 | 0.204 | 9.167 | -2.555 | -0.165 | 9.838 | 1.33 | 59.59 | -18.83 |
| 363 | 2jsa-14 | HIS_A_62_N | HIS_A_62_H | -2.555 | -0.165 | 9.838 | -1.754 | -0.536 | 9.347 | 1.01 | 119.08 | -24.85 |
| 364 | 2jsa-14 | HIS_A_62_N | HIS_A_62_CA | -2.555 | -0.165 | 9.838 | -2.495 | -0.047 | 11.285 | 1.45 | 5.23 | 63.05 |
| 365 | 2jsa-14 | HIS_A_62_CA | HIS_A_62_HA | -2.495 | -0.047 | 11.285 | -3.355 | -0.571 | 11.701 | 1.09 | 67.56 | -148.65 |
| 366 | 2jsa-14 | HIS_A_62_CA | HIS_A_62_C | -2.495 | -0.047 | 11.285 | -2.604 | 1.425 | 11.685 | 1.53 | 74.84 | 94.23 |
| 367 | 2jsa-14 | HIS_A_62_C | HIS_A_62_O | -2.604 | 1.425 | 11.685 | -3.093 | 1.744 | 12.768 | 1.23 | 28.33 | 146.88 |
| 368 | 2jsa-14 | HIS_A_62_C | GLU_A_63_N | -2.604 | 1.425 | 11.685 | -2.139 | 2.285 | 10.789 | 1.33 | 132.50 | 61.60 |
| 369 | 2jsa-14 | GLU_A_63_N | GLU_A_63_H | -2.139 | 2.285 | 10.789 | -1.751 | 1.952 | 9.918 | 1.01 | 149.59 | -40.64 |
| 370 | 2jsa-14 | GLU_A_63_N | GLU_A_63_CA | -2.139 | 2.285 | 10.789 | -2.177 | 3.716 | 11.036 | 1.45 | 80.21 | 91.52 |
| 371 | 2jsa-14 | GLU_A_63_CA | GLU_A_63_HA | -2.177 | 3.716 | 11.036 | -1.904 | 3.886 | 12.078 | 1.09 | 17.15 | 31.91 |
| 372 | 2jsa-14 | GLU_A_63_CA | GLU_A_63_C | -2.177 | 3.716 | 11.036 | -3.592 | 4.255 | 10.816 | 1.53 | 98.27 | 159.15 |
| 373 | 2jsa-14 | GLU_A_63_C | GLU_A_63_O | -3.592 | 4.255 | 10.816 | -4.136 | 4.95 | 11.673 | 1.23 | 45.84 | 128.05 |
| 374 | 2jsa-14 | GLU_A_63_C | LEU_A_64_N | -3.592 | 4.255 | 10.816 | -4.147 | 3.914 | 9.662 | 1.33 | 150.56 | -148.43 |
| 375 | 2jsa-14 | LEU_A_64_N | LEU_A_64_H | -4.147 | 3.914 | 9.662 | -3.64 | 3.34 | 9.003 | 1.01 | 130.71 | -48.55 |
| 376 | 2jsa-14 | LEU_A_64_N | LEU_A_64_CA | -4.147 | 3.914 | 9.662 | -5.488 | 4.354 | 9.319 | 1.45 | 103.66 | 161.83 |
| 377 | 2jsa-14 | LEU_A_64_CA | LEU_A_64_HA | -5.488 | 4.354 | 9.319 | -5.502 | 5.444 | 9.346 | 1.09 | 88.58 | 90.74 |
| 378 | 2jsa-14 | LEU_A_64_CA | LEU_A_64_C | -5.488 | 4.354 | 9.319 | -6.473 | 3.85 | 10.375 | 1.53 | 46.34 | -152.90 |
| 379 | 2jsa-14 | LEU_A_64_C | LEU_A_64_O | -6.473 | 3.85 | 10.375 | -7.418 | 4.552 | 10.733 | 1.23 | 73.09 | 143.39 |
| 380 | 2jsa-14 | LEU_A_64_C | GLU_A_65_N | -6.473 | 3.85 | 10.375 | -6.219 | 2.638 | 10.845 | 1.32 | 69.22 | -78.16 |
| 381 | 2jsa-14 | GLU_A_65_N | GLU_A_65_H | -6.219 | 2.638 | 10.845 | -5.423 | 2.117 | 10.505 | 1.01 | 109.67 | -33.21 |
| 382 | 2jsa-14 | GLU_A_65_N | GLU_A_65_CA | -6.219 | 2.638 | 10.845 | -7.071 | 2.033 | 11.854 | 1.45 | 46.00 | -144.62 |
| 383 | 2jsa-14 | GLU_A_65_CA | GLU_A_65_HA | -7.071 | 2.033 | 11.854 | -8.106 | 2.143 | 11.532 | 1.09 | 107.19 | 173.93 |
| 384 | 2jsa-14 | GLU_A_65_CA | GLU_A_65_C | -7.071 | 2.033 | 11.854 | -6.905 | 2.755 | 13.193 | 1.53 | 28.95 | 77.05 |
| 385 | 2jsa-14 | GLU_A_65_C | GLU_A_65_O | -6.905 | 2.755 | 13.193 | -7.83 | 2.789 | 14.003 | 1.23 | 48.81 | 177.89 |
| 386 | 2jsa-14 | GLU_A_65_C | GLY_A_66_N | -6.905 | 2.755 | 13.193 | -5.72 | 3.317 | 13.383 | 1.33 | 81.76 | 25.37 |
| 387 | 2jsa-14 | GLY_A_66_N | GLY_A_66_H | -5.72 | 3.317 | 13.383 | -5.005 | 3.25 | 12.672 | 1.01 | 134.71 | -5.35 |
| 388 | 2jsa-14 | GLY_A_66_N | GLY_A_66_CA | -5.72 | 3.317 | 13.383 | -5.422 | 4.037 | 14.609 | 1.45 | 32.44 | 67.52 |
| 389 | 2jsa-14 | GLY_A_66_CA | GLY_A_66_C | -5.422 | 4.037 | 14.609 | -6.044 | 5.434 | 14.589 | 1.53 | 90.75 | 114.00 |
| 390 | 2jsa-14 | GLY_A_66_C | GLY_A_66_O | -6.044 | 5.434 | 14.589 | -6.569 | 5.899 | 15.6 | 1.23 | 34.75 | 138.47 |
| 391 | 2jsa-14 | GLY_A_66_C | GLY_A_67_N | -6.044 | 5.434 | 14.589 | -5.964 | 6.067 | 13.428 | 1.32 | 151.21 | 82.80 |
| 392 | 2jsa-14 | GLY_A_67_N | GLY_A_67_H | -5.964 | 6.067 | 13.428 | -5.516 | 5.624 | 12.638 | 1.01 | 141.43 | -44.68 |
| 393 | 2jsa-14 | GLY_A_67_N | GLY_A_67_CA | -5.964 | 6.067 | 13.428 | -6.512 | 7.402 | 13.262 | 1.45 | 96.56 | 112.32 |
| 394 | 2jsa-14 | GLY_A_67_CA | GLY_A_67_C | -6.512 | 7.402 | 13.262 | -5.629 | 8.248 | 12.345 | 1.53 | 126.87 | 43.77 |
| 395 | 2jsa-14 | GLY_A_67_C | GLY_A_67_O | -5.629 | 8.248 | 12.345 | -6.04 | 9.315 | 11.89 | 1.23 | 111.70 | 111.07 |
| 396 | 2jsa-14 | GLY_A_67_C | THR_A_68_N | -5.629 | 8.248 | 12.345 | -4.429 | 7.741 | 12.098 | 1.33 | 100.74 | -22.90 |
| 397 | 2jsa-14 | THR_A_68_N | THR_A_68_H | -4.429 | 7.741 | 12.098 | -4.156 | 6.858 | 12.505 | 1.01 | 66.23 | -72.82 |
| 398 | 2jsa-14 | THR_A_68_N | THR_A_68_CA | -4.429 | 7.741 | 12.098 | -3.484 | 8.438 | 11.242 | 1.45 | 126.09 | 36.41 |
| 399 | 2jsa-14 | THR_A_68_CA | THR_A_68_HA | -3.484 | 8.438 | 11.242 | -3.318 | 9.438 | 11.645 | 1.09 | 68.32 | 80.57 |
| 400 | 2jsa-14 | THR_A_68_CA | THR_A_68_C | -3.484 | 8.438 | 11.242 | -4.053 | 8.587 | 9.83 | 1.53 | 157.39 | 165.33 |
| 401 | 2jsa-14 | THR_A_68_C | THR_A_68_O | -4.053 | 8.587 | 9.83 | -4.723 | 7.685 | 9.328 | 1.23 | 114.07 | -126.60 |
| 402 | 2jsa-14 | THR_A_68_C | ALA_A_69_N | -4.053 | 8.587 | 9.83 | -3.766 | 9.732 | 9.229 | 1.32 | 116.98 | 75.93 |
| 403 | 2jsa-14 | ALA_A_69_N | ALA_A_69_H | -3.766 | 9.732 | 9.229 | -3.21 | 10.43 | 9.703 | 1.01 | 62.02 | 51.46 |
| 404 | 2jsa-14 | ALA_A_69_N | ALA_A_69_CA | -3.766 | 9.732 | 9.229 | -4.24 | 10.011 | 7.884 | 1.45 | 157.76 | 149.52 |
| 405 | 2jsa-14 | ALA_A_69_CA | ALA_A_69_HA | -4.24 | 10.011 | 7.884 | -5.297 | 9.753 | 7.825 | 1.09 | 93.10 | -166.28 |
| 406 | 2jsa-14 | ALA_A_69_CA | ALA_A_69_C | -4.24 | 10.011 | 7.884 | -3.473 | 9.141 | 6.886 | 1.53 | 130.71 | -48.60 |
| 407 | 2jsa-14 | ALA_A_69_C | ALA_A_69_O | -3.473 | 9.141 | 6.886 | -2.468 | 8.526 | 7.238 | 1.23 | 73.37 | -31.46 |
| 408 | 2jsa-14 | ALA_A_69_C | PRO_A_70_N | -3.473 | 9.141 | 6.886 | -3.99 | 9.117 | 5.628 | 1.36 | 157.64 | -177.34 |
| 409 | 2jsa-14 | PRO_A_70_N | PRO_A_70_CA | -3.99 | 9.117 | 5.628 | -3.365 | 8.333 | 4.577 | 1.45 | 136.35 | -51.44 |
| 410 | 2jsa-14 | PRO_A_70_CA | PRO_A_70_HA | -3.365 | 8.333 | 4.577 | -3.108 | 7.342 | 4.953 | 1.09 | 69.83 | -75.46 |
| 411 | 2jsa-14 | PRO_A_70_CA | PRO_A_70_C | -3.365 | 8.333 | 4.577 | -2.086 | 9.009 | 4.076 | 1.53 | 109.10 | 27.86 |
| 412 | 2jsa-14 | PRO_A_70_C | PRO_A_70_O | -2.086 | 9.009 | 4.076 | -1.218 | 8.353 | 3.502 | 1.23 | 117.81 | -37.08 |
| 413 | 2jsa-14 | PRO_A_70_C | LYS_A_71_N | -2.086 | 9.009 | 4.076 | -2.011 | 10.31 | 4.312 | 1.32 | 79.74 | 86.70 |
| 414 | 2jsa-14 | LYS_A_71_N | LYS_A_71_H | -2.011 | 10.31 | 4.312 | -2.763 | 10.785 | 4.79 | 1.01 | 61.75 | 147.72 |
| 415 | 2jsa-14 | LYS_A_71_N | LYS_A_71_CA | -2.011 | 10.31 | 4.312 | -0.853 | 11.081 | 3.892 | 1.45 | 106.80 | 33.66 |
| 416 | 2jsa-14 | LYS_A_71_CA | LYS_A_71_HA | -0.853 | 11.081 | 3.892 | -0.341 | 10.522 | 3.109 | 1.09 | 135.93 | -47.51 |
| 417 | 2jsa-14 | LYS_A_71_CA | LYS_A_71_C | -0.853 | 11.081 | 3.892 | 0.114 | 11.223 | 5.069 | 1.53 | 39.71 | 8.35 |
| 418 | 2jsa-14 | LYS_A_71_C | LYS_A_71_O | 0.114 | 11.223 | 5.069 | 1.072 | 11.992 | 5.001 | 1.23 | 93.17 | 38.75 |
| 419 | 2jsa-14 | LYS_A_71_C | ASP_A_72_N | 0.114 | 11.223 | 5.069 | -0.17 | 10.469 | 6.121 | 1.33 | 37.45 | -110.64 |
| 420 | 2jsa-14 | ASP_A_72_N | ASP_A_72_H | -0.17 | 10.469 | 6.121 | -0.975 | 9.859 | 6.108 | 1.01 | 90.74 | -142.85 |
| 421 | 2jsa-14 | ASP_A_72_N | ASP_A_72_CA | -0.17 | 10.469 | 6.121 | 0.663 | 10.501 | 7.312 | 1.45 | 34.99 | 2.20 |
| 422 | 2jsa-14 | ASP_A_72_CA | ASP_A_72_HA | 0.663 | 10.501 | 7.312 | 1.418 | 11.279 | 7.198 | 1.09 | 96.00 | 45.86 |

| | | | | | | | | | | | |
|---|---|---|---|---|---|---|---|---|---|---|---|
| 423 | 2jsa-14 | ASP_A_72_CA | ASP_A_72_C | 0.663 | 10.501 | 7.312 | 1.34 | 9.141 | 7.492 | 1.53 | 83.24 | -63.54 |
| 424 | 2jsa-14 | ASP_A_72_C | ASP_A_72_O | 1.34 | 9.141 | 7.492 | 2.481 | 9.067 | 7.946 | 1.23 | 68.34 | -3.71 |
| 425 | 2jsa-14 | ASP_A_72_C | VAL_A_73_N | 1.34 | 9.141 | 7.492 | 0.61 | 8.098 | 7.125 | 1.32 | 106.08 | -124.99 |
| 426 | 2jsa-14 | VAL_A_73_N | VAL_A_73_H | 0.61 | 8.098 | 7.125 | -0.321 | 8.233 | 6.759 | 1.01 | 111.26 | 171.75 |
| 427 | 2jsa-14 | VAL_A_73_N | VAL_A_73_CA | 0.61 | 8.098 | 7.125 | 1.127 | 6.745 | 7.24 | 1.45 | 85.46 | -69.09 |
| 428 | 2jsa-14 | VAL_A_73_CA | VAL_A_73_HA | 1.127 | 6.745 | 7.24 | 1.374 | 6.558 | 8.285 | 1.09 | 16.51 | -37.13 |
| 429 | 2jsa-14 | VAL_A_73_CA | VAL_A_73_C | 1.127 | 6.745 | 7.24 | 2.412 | 6.623 | 6.42 | 1.53 | 122.43 | -5.42 |
| 430 | 2jsa-14 | VAL_A_73_C | VAL_A_73_O | 2.412 | 6.623 | 6.42 | 3.376 | 5.996 | 6.859 | 1.23 | 69.11 | -33.04 |
| 431 | 2jsa-14 | VAL_A_73_C | CYS_A_74_N | 2.412 | 6.623 | 6.42 | 2.386 | 7.232 | 5.243 | 1.33 | 152.62 | 92.44 |
| 432 | 2jsa-14 | CYS_A_74_N | CYS_A_74_H | 2.386 | 7.232 | 5.243 | 1.56 | 7.731 | 4.947 | 1.01 | 107.05 | 148.86 |
| 433 | 2jsa-14 | CYS_A_74_N | CYS_A_74_CA | 2.386 | 7.232 | 5.243 | 3.537 | 7.199 | 4.357 | 1.45 | 127.58 | -1.64 |
| 434 | 2jsa-14 | CYS_A_74_CA | CYS_A_74_HA | 3.537 | 7.199 | 4.357 | 3.94 | 6.186 | 4.356 | 1.09 | 90.05 | -68.31 |
| 435 | 2jsa-14 | CYS_A_74_CA | CYS_A_74_C | 3.537 | 7.199 | 4.357 | 4.601 | 8.146 | 4.917 | 1.53 | 68.54 | 41.67 |
| 436 | 2jsa-14 | CYS_A_74_C | CYS_A_74_O | 4.601 | 8.146 | 4.917 | 4.96 | 9.131 | 4.273 | 1.23 | 121.56 | 69.97 |
| 437 | 2jsa-14 | CYS_A_74_C | THR_A_75_N | 4.601 | 8.146 | 4.917 | 5.074 | 7.813 | 6.108 | 1.32 | 25.91 | -35.15 |
| 438 | 2jsa-14 | THR_A_75_N | THR_A_75_H | 5.074 | 7.813 | 6.108 | 4.733 | 6.987 | 6.579 | 1.01 | 62.21 | -112.43 |
| 439 | 2jsa-14 | THR_A_75_N | THR_A_75_CA | 5.074 | 7.813 | 6.108 | 6.09 | 8.622 | 6.761 | 1.45 | 63.31 | 38.53 |
| 440 | 2jsa-14 | THR_A_75_CA | THR_A_75_HA | 6.09 | 8.622 | 6.761 | 6.99 | 8.619 | 6.145 | 1.09 | 124.39 | -0.19 |
| 441 | 2jsa-14 | THR_A_75_CA | THR_A_75_C | 6.09 | 8.622 | 6.761 | 6.45 | 8.027 | 8.124 | 1.53 | 27.03 | -58.82 |
| 442 | 2jsa-14 | THR_A_75_C | THR_A_75_O | 6.45 | 8.027 | 8.124 | 7.603 | 7.677 | 8.369 | 1.23 | 78.51 | -16.89 |
| 443 | 2jsa-14 | THR_A_75_C | LYS_A_76_N | 6.45 | 8.027 | 8.124 | 5.44 | 7.932 | 8.977 | 1.33 | 49.94 | -174.63 |
| 444 | 2jsa-14 | LYS_A_76_N | LYS_A_76_H | 5.44 | 7.932 | 8.977 | 4.517 | 8.24 | 8.708 | 1.01 | 105.45 | 161.55 |
| 445 | 2jsa-14 | LYS_A_76_N | LYS_A_76_CA | 5.44 | 7.932 | 8.977 | 5.635 | 7.386 | 10.31 | 1.45 | 23.51 | -70.35 |
| 446 | 2jsa-14 | LYS_A_76_CA | LYS_A_76_HA | 5.635 | 7.386 | 10.31 | 6.502 | 7.878 | 10.753 | 1.09 | 66.04 | 29.57 |
| 447 | 2jsa-14 | LYS_A_76_CA | LYS_A_76_C | 5.635 | 7.386 | 10.31 | 5.953 | 5.893 | 10.203 | 1.53 | 94.01 | -77.98 |
| 448 | 2jsa-14 | LYS_A_76_C | LYS_A_76_O | 5.953 | 5.893 | 10.203 | 6.561 | 5.318 | 11.104 | 1.23 | 42.89 | -43.40 |
| 449 | 2jsa-14 | LYS_A_76_C | LEU_A_77_N | 5.953 | 5.893 | 10.203 | 5.525 | 5.308 | 9.094 | 1.32 | 146.83 | -126.19 |
| 450 | 2jsa-14 | LEU_A_77_N | LEU_A_77_H | 5.525 | 5.308 | 9.094 | 5.031 | 5.844 | 8.395 | 1.01 | 133.80 | 132.66 |
| 451 | 2jsa-14 | LEU_A_77_N | LEU_A_77_CA | 5.525 | 5.308 | 9.094 | 5.755 | 3.892 | 8.857 | 1.45 | 99.38 | -80.77 |
| 452 | 2jsa-14 | LEU_A_77_CA | LEU_A_77_HA | 5.755 | 3.892 | 8.857 | 5.77 | 3.386 | 9.823 | 1.09 | 27.66 | -88.30 |
| 453 | 2jsa-14 | LEU_A_77_CA | LEU_A_77_C | 5.755 | 3.892 | 8.857 | 7.131 | 3.705 | 8.213 | 1.53 | 114.88 | -7.74 |
| 454 | 2jsa-14 | LEU_A_77_C | LEU_A_77_O | 7.131 | 3.705 | 8.213 | 7.447 | 2.623 | 7.721 | 1.23 | 113.58 | -73.72 |
| 455 | 2jsa-14 | LEU_A_77_C | ASN_A_78_N | 7.131 | 3.705 | 8.213 | 7.912 | 4.775 | 8.239 | 1.32 | 88.88 | 53.87 |
| 456 | 2jsa-14 | ASN_A_78_N | ASN_A_78_H | 7.912 | 4.775 | 8.239 | 7.587 | 5.632 | 8.663 | 1.01 | 65.17 | 110.77 |
| 457 | 2jsa-14 | ASN_A_78_N | ASN_A_78_CA | 7.912 | 4.775 | 8.239 | 9.246 | 4.742 | 7.664 | 1.45 | 113.31 | -1.42 |
| 458 | 2jsa-14 | ASN_A_78_CA | ASN_A_78_HA | 9.246 | 4.742 | 7.664 | 9.644 | 5.756 | 7.627 | 1.09 | 91.95 | 68.57 |
| 459 | 2jsa-14 | ASN_A_78_CA | ASN_A_78_C | 9.246 | 4.742 | 7.664 | 9.173 | 4.161 | 6.251 | 1.53 | 157.49 | -97.16 |
| 460 | 2jsa-14 | ASN_A_78_C | ASN_A_78_O | 9.173 | 4.161 | 6.251 | 10.127 | 3.542 | 5.781 | 1.23 | 112.45 | -32.98 |
| 461 | 2jsa-14 | ASN_A_78_C | GLU_A_79_N | 9.173 | 4.161 | 6.251 | 8.032 | 4.378 | 5.613 | 1.33 | 118.78 | 169.23 |
| 462 | 2jsa-14 | GLU_A_79_N | GLU_A_79_H | 8.032 | 4.378 | 5.613 | 7.289 | 4.894 | 6.062 | 1.01 | 63.60 | 145.22 |
| 463 | 2jsa-14 | GLU_A_79_N | GLU_A_79_CA | 8.032 | 4.378 | 5.613 | 7.824 | 3.884 | 4.263 | 1.45 | 158.34 | -112.83 |
| 464 | 2jsa-14 | GLU_A_79_CA | GLU_A_79_HA | 7.824 | 3.884 | 4.263 | 8.329 | 2.922 | 4.174 | 1.09 | 94.68 | -62.30 |
| 465 | 2jsa-14 | GLU_A_79_CA | GLU_A_79_C | 7.824 | 3.884 | 4.263 | 8.439 | 4.845 | 3.245 | 1.53 | 131.74 | 57.38 |
| 466 | 2jsa-14 | GLU_A_79_C | GLU_A_79_O | 8.439 | 4.845 | 3.245 | 9.38 | 4.487 | 2.538 | 1.23 | 125.08 | -20.83 |
| 467 | 2jsa-14 | GLU_A_79_C | CYS_A_80_N | 8.439 | 4.845 | 3.245 | 7.881 | 6.046 | 3.201 | 1.33 | 91.90 | 114.92 |
| 468 | 2jsa-14 | CYS_A_80_N | CYS_A_80_H | 7.881 | 6.046 | 3.201 | 7.109 | 6.271 | 3.812 | 1.01 | 52.77 | 163.75 |
| 469 | 2jsa-14 | CYS_A_80_N | CYS_A_80_CA | 7.881 | 6.046 | 3.201 | 8.363 | 7.062 | 2.281 | 1.45 | 129.29 | 64.62 |
| 470 | 2jsa-14 | CYS_A_80_CA | CYS_A_80_HA | 8.363 | 7.062 | 2.281 | 8.101 | 6.759 | 1.268 | 1.09 | 158.42 | -130.85 |
| 471 | 2jsa-14 | CYS_A_80_CA | CYS_A_80_C | 8.363 | 7.062 | 2.281 | 9.888 | 7.125 | 2.39 | 1.53 | 85.92 | 2.37 |
| 472 | 2jsa-14 | CYS_A_80_C | CYS_A_80_O | 9.888 | 7.125 | 2.39 | 10.463 | 6.665 | 3.375 | 1.23 | 36.78 | -38.66 |
| 473 | 2jsa-14 | CYS_A_80_C | PRO_A_81_N | 9.888 | 7.125 | 2.39 | 10.516 | 7.712 | 1.336 | 1.36 | 140.80 | 43.07 |
| 474 | 2jsa-14 | PRO_A_81_N | PRO_A_81_CA | 10.516 | 7.712 | 1.336 | 11.963 | 7.84 | 1.305 | 1.45 | 91.22 | 5.06 |
| 475 | 2jsa-14 | PRO_A_81_CA | PRO_A_81_HA | 11.963 | 7.84 | 1.305 | 12.428 | 6.903 | 1.612 | 1.09 | 73.64 | -63.61 |
| 476 | 2jsa-14 | PRO_A_81_CA | PRO_A_81_C | 11.963 | 7.84 | 1.305 | 12.436 | 8.948 | 2.247 | 1.53 | 51.98 | 66.88 |
| 477 | 2jsa-14 | PRO_A_81_C | PRO_A_81_O | 12.436 | 8.948 | 2.247 | 13.633 | 9.21 | 2.352 | 1.23 | 85.10 | 12.35 |
| 1 | 2jsa-15 | ARG_A_1_N | ARG_A_1_H | 13.763 | 1.432 | 2.232 | 14.415 | 2.167 | 1.997 | 1.01 | 103.45 | 48.42 |
| 2 | 2jsa-15 | ARG_A_1_N | ARG_A_1_CA | 13.763 | 1.432 | 2.232 | 12.354 | 1.765 | 2.35 | 1.45 | 85.34 | 166.70 |
| 3 | 2jsa-15 | ARG_A_1_CA | ARG_A_1_HA | 12.354 | 1.765 | 2.35 | 11.8 | 1.181 | 1.615 | 1.09 | 132.40 | -133.49 |
| 4 | 2jsa-15 | ARG_A_1_CA | ARG_A_1_C | 12.354 | 1.765 | 2.35 | 12.135 | 3.253 | 2.072 | 1.53 | 100.47 | 98.37 |
| 5 | 2jsa-15 | ARG_A_1_C | ARG_A_1_O | 12.135 | 3.253 | 2.072 | 11.525 | 3.956 | 2.876 | 1.23 | 49.18 | 130.95 |
| 6 | 2jsa-15 | ARG_A_1_C | SER_A_2_N | 12.135 | 3.253 | 2.072 | 12.646 | 3.69 | 0.929 | 1.33 | 149.53 | 40.54 |
| 7 | 2jsa-15 | SER_A_2_N | SER_A_2_H | 12.646 | 3.69 | 0.929 | 13.136 | 3.055 | 0.315 | 1.01 | 127.43 | -52.34 |
| 8 | 2jsa-15 | SER_A_2_N | SER_A_2_CA | 12.646 | 3.69 | 0.929 | 12.514 | 5.082 | 0.534 | 1.45 | 105.77 | 95.42 |
| 9 | 2jsa-15 | SER_A_2_CA | SER_A_2_HA | 12.514 | 5.082 | 0.534 | 11.56 | 5.454 | 0.907 | 1.09 | 69.98 | 158.70 |
| 10 | 2jsa-15 | SER_A_2_CA | SER_A_2_C | 12.514 | 5.082 | 0.534 | 12.522 | 5.196 | -0.992 | 1.53 | 175.72 | 85.99 |
| 11 | 2jsa-15 | SER_A_2_C | SER_A_2_O | 12.522 | 5.196 | -0.992 | 13.551 | 5.506 | -1.589 | 1.23 | 119.05 | 16.77 |
| 12 | 2jsa-15 | SER_A_2_C | ALA_A_3_N | 12.522 | 5.196 | -0.992 | 11.362 | 4.94 | -1.578 | 1.32 | 116.26 | -167.55 |
| 13 | 2jsa-15 | ALA_A_3_N | ALA_A_3_H | 11.362 | 4.94 | -1.578 | 10.556 | 4.691 | -1.023 | 1.01 | 56.66 | -162.83 |
| 14 | 2jsa-15 | ALA_A_3_N | ALA_A_3_CA | 11.362 | 4.94 | -1.578 | 11.222 | 5.01 | -3.023 | 1.45 | 173.82 | 153.43 |
| 15 | 2jsa-15 | ALA_A_3_CA | ALA_A_3_HA | 11.222 | 5.01 | -3.023 | 11.589 | 5.978 | -3.364 | 1.09 | 108.23 | 69.24 |
| 16 | 2jsa-15 | ALA_A_3_CA | ALA_A_3_C | 11.222 | 5.01 | -3.023 | 9.742 | 4.895 | -3.394 | 1.53 | 104.03 | -175.56 |
| 17 | 2jsa-15 | ALA_A_3_C | ALA_A_3_O | 9.742 | 4.895 | -3.394 | 8.881 | 5.439 | -2.704 | 1.23 | 55.88 | 147.71 |
| 18 | 2jsa-15 | ALA_A_3_C | LEU_A_4_N | 9.742 | 4.895 | -3.394 | 9.492 | 4.183 | -4.483 | 1.32 | 145.28 | -109.35 |
| 19 | 2jsa-15 | LEU_A_4_N | LEU_A_4_H | 9.492 | 4.183 | -4.483 | 10.251 | 3.764 | -5.002 | 1.01 | 120.91 | -28.90 |
| 20 | 2jsa-15 | LEU_A_4_N | LEU_A_4_CA | 9.492 | 4.183 | -4.483 | 8.131 | 3.989 | -4.954 | 1.45 | 108.91 | -171.89 |
| 21 | 2jsa-15 | LEU_A_4_CA | LEU_A_4_HA | 8.131 | 3.989 | -4.954 | 7.714 | 4.968 | -5.189 | 1.09 | 102.45 | 113.07 |
| 22 | 2jsa-15 | LEU_A_4_CA | LEU_A_4_C | 8.131 | 3.989 | -4.954 | 7.288 | 3.382 | -3.831 | 1.53 | 42.77 | -144.24 |
| 23 | 2jsa-15 | LEU_A_4_C | LEU_A_4_O | 7.288 | 3.382 | -3.831 | 6.06 | 3.452 | -3.864 | 1.23 | 91.54 | 176.74 |
| 24 | 2jsa-15 | LEU_A_4_C | SER_A_5_N | 7.288 | 3.382 | -3.831 | 7.981 | 2.801 | -2.862 | 1.33 | 43.02 | -39.98 |
| 25 | 2jsa-15 | SER_A_5_N | SER_A_5_H | 7.981 | 2.801 | -2.862 | 8.991 | 2.783 | -2.899 | 1.01 | 92.10 | -1.02 |
| 26 | 2jsa-15 | SER_A_5_N | SER_A_5_CA | 7.981 | 2.801 | -2.862 | 7.312 | 2.182 | -1.731 | 1.45 | 38.86 | -137.22 |
| 27 | 2jsa-15 | SER_A_5_CA | SER_A_5_HA | 7.312 | 2.182 | -1.731 | 6.804 | 1.285 | -2.084 | 1.09 | 108.90 | -119.52 |
| 28 | 2jsa-15 | SER_A_5_CA | SER_A_5_C | 7.312 | 2.182 | -1.731 | 6.269 | 3.141 | -1.152 | 1.53 | 67.77 | 137.40 |
| 29 | 2jsa-15 | SER_A_5_C | SER_A_5_O | 6.269 | 3.141 | -1.152 | 5.117 | 2.761 | -0.951 | 1.23 | 80.59 | -161.74 |
| 30 | 2jsa-15 | SER_A_5_C | CYS_A_6_N | 6.269 | 3.141 | -1.152 | 6.711 | 4.364 | -0.901 | 1.32 | 79.08 | 70.13 |
| 31 | 2jsa-15 | CYS_A_6_N | CYS_A_6_H | 6.711 | 4.364 | -0.901 | 7.673 | 4.607 | -1.092 | 1.01 | 100.90 | 14.18 |
| 32 | 2jsa-15 | CYS_A_6_N | CYS_A_6_CA | 6.711 | 4.364 | -0.901 | 5.83 | 5.379 | -0.349 | 1.45 | 67.67 | 130.96 |
| 33 | 2jsa-15 | CYS_A_6_CA | CYS_A_6_HA | 5.83 | 5.379 | -0.349 | 5.593 | 5.105 | 0.679 | 1.09 | 19.41 | -130.86 |
| 34 | 2jsa-15 | CYS_A_6_CA | CYS_A_6_C | 5.83 | 5.379 | -0.349 | 4.535 | 5.385 | -1.163 | 1.53 | 122.15 | 179.73 |
| 35 | 2jsa-15 | CYS_A_6_C | CYS_A_6_O | 4.535 | 5.385 | -1.163 | 3.444 | 5.302 | -0.601 | 1.23 | 62.81 | -175.65 |
| 36 | 2jsa-15 | CYS_A_6_C | GLN_A_7_N | 4.535 | 5.385 | -1.163 | 4.698 | 5.486 | -2.475 | 1.33 | 171.68 | 31.78 |
| 37 | 2jsa-15 | GLN_A_7_N | GLN_A_7_H | 4.698 | 5.486 | -2.475 | 5.626 | 5.552 | -2.868 | 1.01 | 112.90 | 4.07 |
| 38 | 2jsa-15 | GLN_A_7_N | GLN_A_7_CA | 4.698 | 5.486 | -2.475 | 3.555 | 5.505 | -3.372 | 1.45 | 128.12 | 179.05 |
| 39 | 2jsa-15 | GLN_A_7_CA | GLN_A_7_HA | 3.555 | 5.505 | -3.372 | 3.009 | 6.433 | -3.202 | 1.09 | 81.03 | 120.47 |
| 40 | 2jsa-15 | GLN_A_7_CA | GLN_A_7_C | 3.555 | 5.505 | -3.372 | 2.621 | 4.332 | -3.064 | 1.53 | 78.39 | -128.53 |
| 41 | 2jsa-15 | GLN_A_7_C | GLN_A_7_O | 2.621 | 4.332 | -3.064 | 1.424 | 4.525 | -2.857 | 1.23 | 80.31 | 170.84 |

| | | | | | | | | | | | | |
|---|---|---|---|---|---|---|---|---|---|---|---|---|
| 42 | 2jsa-15 | GLN_A_7_C | MET_A_8_N | 2.621 | 4.332 | -3.064 | 3.202 | 3.142 | -3.045 | 1.32 | 89.18 | -63.98 |
| 43 | 2jsa-15 | MET_A_8_N | MET_A_8_H | 3.202 | 3.142 | -3.045 | 4.192 | 3.057 | -3.226 | 1.01 | 100.32 | -4.91 |
| 44 | 2jsa-15 | MET_A_8_N | MET_A_8_CA | 3.202 | 3.142 | -3.045 | 2.437 | 1.939 | -2.767 | 1.45 | 78.97 | -122.45 |
| 45 | 2jsa-15 | MET_A_8_CA | MET_A_8_HA | 2.437 | 1.939 | -2.767 | 1.786 | 1.735 | -3.617 | 1.09 | 141.25 | -162.60 |
| 46 | 2jsa-15 | MET_A_8_CA | MET_A_8_C | 2.437 | 1.939 | -2.767 | 1.577 | 2.113 | -1.513 | 1.53 | 34.98 | 168.56 |
| 47 | 2jsa-15 | MET_A_8_C | MET_A_8_O | 1.577 | 2.113 | -1.513 | 0.438 | 1.653 | -1.468 | 1.23 | 87.90 | -158.01 |
| 48 | 2jsa-15 | MET_A_8_C | CYS_A_9_N | 1.577 | 2.113 | -1.513 | 2.157 | 2.78 | -0.526 | 1.32 | 41.85 | 48.99 |
| 49 | 2jsa-15 | CYS_A_9_N | CYS_A_9_H | 2.157 | 2.78 | -0.526 | 3.098 | 3.132 | -0.633 | 1.01 | 96.08 | 20.51 |
| 50 | 2jsa-15 | CYS_A_9_N | CYS_A_9_CA | 2.157 | 2.78 | -0.526 | 1.458 | 3.02 | 0.725 | 1.45 | 30.57 | 161.05 |
| 51 | 2jsa-15 | CYS_A_9_CA | CYS_A_9_HA | 1.458 | 3.02 | 0.725 | 1.165 | 2.058 | 1.145 | 1.09 | 67.33 | -106.94 |
| 52 | 2jsa-15 | CYS_A_9_CA | CYS_A_9_C | 1.458 | 3.02 | 0.725 | 0.193 | 3.825 | 0.422 | 1.53 | 101.42 | 147.53 |
| 53 | 2jsa-15 | CYS_A_9_C | CYS_A_9_O | 0.193 | 3.825 | 0.422 | -0.82 | 3.677 | 1.104 | 1.23 | 56.33 | -171.69 |
| 54 | 2jsa-15 | CYS_A_9_C | GLU_A_10_N | 0.193 | 3.825 | 0.422 | 0.292 | 4.66 | -0.603 | 1.33 | 140.64 | 83.24 |
| 55 | 2jsa-15 | GLU_A_10_N | GLU_A_10_H | 0.292 | 4.66 | -0.603 | 1.157 | 4.73 | -1.12 | 1.01 | 120.78 | 4.63 |
| 56 | 2jsa-15 | GLU_A_10_N | GLU_A_10_CA | 0.292 | 4.66 | -0.603 | -0.832 | 5.487 | -1.005 | 1.45 | 106.07 | 143.66 |
| 57 | 2jsa-15 | GLU_A_10_CA | GLU_A_10_HA | -0.832 | 5.487 | -1.005 | -1.346 | 5.819 | -0.103 | 1.09 | 34.15 | 147.14 |
| 58 | 2jsa-15 | GLU_A_10_CA | GLU_A_10_C | -0.832 | 5.487 | -1.005 | -1.816 | 4.674 | -1.847 | 1.53 | 123.41 | -140.44 |
| 59 | 2jsa-15 | GLU_A_10_C | GLU_A_10_O | -1.816 | 4.674 | -1.847 | -2.897 | 5.156 | -2.184 | 1.23 | 105.89 | 155.97 |
| 60 | 2jsa-15 | GLU_A_10_C | LEU_A_11_N | -1.816 | 4.674 | -1.847 | -1.407 | 3.454 | -2.163 | 1.32 | 103.80 | -71.47 |
| 61 | 2jsa-15 | LEU_A_11_N | LEU_A_11_H | -1.407 | 3.454 | -2.163 | -0.505 | 3.123 | -1.852 | 1.01 | 72.06 | -20.15 |
| 62 | 2jsa-15 | LEU_A_11_N | LEU_A_11_CA | -1.407 | 3.454 | -2.163 | -2.24 | 2.569 | -2.959 | 1.45 | 123.22 | -133.27 |
| 63 | 2jsa-15 | LEU_A_11_CA | LEU_A_11_HA | -2.24 | 2.569 | -2.959 | -2.927 | 3.184 | -3.54 | 1.09 | 122.22 | 138.17 |
| 64 | 2jsa-15 | LEU_A_11_CA | LEU_A_11_C | -2.24 | 2.569 | -2.959 | -3.077 | 1.688 | -2.029 | 1.53 | 52.57 | -133.53 |
| 65 | 2jsa-15 | LEU_A_11_C | LEU_A_11_O | -3.077 | 1.688 | -2.029 | -4.239 | 1.406 | -2.316 | 1.23 | 103.50 | -166.36 |
| 66 | 2jsa-15 | LEU_A_11_C | VAL_A_12_N | -3.077 | 1.688 | -2.029 | -2.454 | 1.28 | -0.934 | 1.32 | 34.22 | -33.22 |
| 67 | 2jsa-15 | VAL_A_12_N | VAL_A_12_H | -2.454 | 1.28 | -0.934 | -1.496 | 1.552 | -0.763 | 1.01 | 80.26 | 15.85 |
| 68 | 2jsa-15 | VAL_A_12_N | VAL_A_12_CA | -2.454 | 1.28 | -0.934 | -3.127 | 0.438 | 0.041 | 1.45 | 47.87 | -128.63 |
| 69 | 2jsa-15 | VAL_A_12_CA | VAL_A_12_HA | -3.127 | 0.438 | 0.041 | -3.675 | -0.337 | -0.494 | 1.09 | 119.41 | -125.26 |
| 70 | 2jsa-15 | VAL_A_12_CA | VAL_A_12_C | -3.127 | 0.438 | 0.041 | -4.136 | 1.279 | 0.824 | 1.53 | 59.20 | 140.19 |
| 71 | 2jsa-15 | VAL_A_12_C | VAL_A_12_O | -4.136 | 1.279 | 0.824 | -5.193 | 0.784 | 1.213 | 1.23 | 71.57 | -154.91 |
| 72 | 2jsa-15 | VAL_A_12_C | VAL_A_13_N | -4.136 | 1.279 | 0.824 | -3.775 | 2.536 | 1.032 | 1.32 | 80.96 | 73.98 |
| 73 | 2jsa-15 | VAL_A_13_N | VAL_A_13_H | -3.775 | 2.536 | 1.032 | -2.89 | 2.876 | 0.683 | 1.01 | 110.21 | 21.02 |
| 74 | 2jsa-15 | VAL_A_13_N | VAL_A_13_CA | -3.775 | 2.536 | 1.032 | -4.637 | 3.451 | 1.763 | 1.45 | 59.82 | 133.29 |
| 75 | 2jsa-15 | VAL_A_13_CA | VAL_A_13_HA | -4.637 | 3.451 | 1.763 | -4.983 | 2.951 | 2.668 | 1.09 | 33.90 | -124.68 |
| 76 | 2jsa-15 | VAL_A_13_CA | VAL_A_13_C | -4.637 | 3.451 | 1.763 | -5.86 | 3.785 | 0.907 | 1.53 | 124.03 | 164.73 |
| 77 | 2jsa-15 | VAL_A_13_C | VAL_A_13_O | -5.86 | 3.785 | 0.907 | -6.995 | 3.547 | 1.319 | 1.23 | 70.44 | -168.16 |
| 78 | 2jsa-15 | VAL_A_13_C | LYS_A_14_N | -5.86 | 3.785 | 0.907 | -5.589 | 4.332 | -0.269 | 1.33 | 152.57 | 63.64 |
| 79 | 2jsa-15 | LYS_A_14_N | LYS_A_14_H | -5.589 | 4.332 | -0.269 | -4.631 | 4.5 | -0.543 | 1.01 | 105.73 | 9.95 |
| 80 | 2jsa-15 | LYS_A_14_N | LYS_A_14_CA | -5.589 | 4.332 | -0.269 | -6.653 | 4.701 | -1.186 | 1.45 | 129.15 | 160.87 |
| 81 | 2jsa-15 | LYS_A_14_CA | LYS_A_14_HA | -6.653 | 4.701 | -1.186 | -7.202 | 5.534 | -0.747 | 1.09 | 66.25 | 123.39 |
| 82 | 2jsa-15 | LYS_A_14_CA | LYS_A_14_C | -6.653 | 4.701 | -1.186 | -7.626 | 3.529 | -1.332 | 1.53 | 95.47 | -129.70 |
| 83 | 2jsa-15 | LYS_A_14_C | LYS_A_14_O | -7.626 | 3.529 | -1.332 | -8.826 | 3.732 | -1.507 | 1.23 | 98.18 | 170.40 |
| 84 | 2jsa-15 | LYS_A_14_C | LYS_A_15_N | -7.626 | 3.529 | -1.332 | -7.07 | 2.329 | -1.253 | 1.32 | 86.58 | -65.14 |
| 85 | 2jsa-15 | LYS_A_15_N | LYS_A_15_H | -7.07 | 2.329 | -1.253 | -6.074 | 2.238 | -1.107 | 1.01 | 81.69 | -5.22 |
| 86 | 2jsa-15 | LYS_A_15_N | LYS_A_15_CA | -7.07 | 2.329 | -1.253 | -7.874 | 1.123 | -1.373 | 1.45 | 94.73 | -123.69 |
| 87 | 2jsa-15 | LYS_A_15_CA | LYS_A_15_HA | -7.874 | 1.123 | -1.373 | -8.628 | 1.295 | -2.14 | 1.09 | 134.76 | 167.15 |
| 88 | 2jsa-15 | LYS_A_15_CA | LYS_A_15_C | -7.874 | 1.123 | -1.373 | -8.603 | 0.868 | -0.053 | 1.53 | 30.33 | -160.72 |
| 89 | 2jsa-15 | LYS_A_15_C | LYS_A_15_O | -8.603 | 0.868 | -0.053 | -9.77 | 0.48 | -0.05 | 1.23 | 89.86 | -161.61 |
| 90 | 2jsa-15 | LYS_A_15_C | TYR_A_16_N | -8.603 | 0.868 | -0.053 | -7.885 | 1.097 | 1.037 | 1.33 | 34.66 | 17.69 |
| 91 | 2jsa-15 | TYR_A_16_N | TYR_A_16_H | -7.885 | 1.097 | 1.037 | -6.929 | 1.414 | 0.959 | 1.01 | 94.43 | 18.35 |
| 92 | 2jsa-15 | TYR_A_16_N | TYR_A_16_CA | -7.885 | 1.097 | 1.037 | -8.45 | 0.897 | 2.361 | 1.45 | 24.36 | -160.51 |
| 93 | 2jsa-15 | TYR_A_16_CA | TYR_A_16_HA | -8.45 | 0.897 | 2.361 | -9.018 | -0.033 | 2.392 | 1.09 | 88.37 | -121.41 |
| 94 | 2jsa-15 | TYR_A_16_CA | TYR_A_16_C | -8.45 | 0.897 | 2.361 | -9.351 | 2.069 | 2.756 | 1.53 | 75.04 | 127.55 |
| 95 | 2jsa-15 | TYR_A_16_C | TYR_A_16_O | -9.351 | 2.069 | 2.756 | -10.572 | 1.929 | 2.805 | 1.23 | 87.72 | -173.46 |
| 96 | 2jsa-15 | TYR_A_16_C | GLU_A_17_N | -9.351 | 2.069 | 2.756 | -8.714 | 3.198 | 3.027 | 1.32 | 78.19 | 60.57 |
| 97 | 2jsa-15 | GLU_A_17_N | GLU_A_17_H | -8.714 | 3.198 | 3.027 | -7.706 | 3.241 | 2.965 | 1.01 | 93.52 | 2.44 |
| 98 | 2jsa-15 | GLU_A_17_N | GLU_A_17_CA | -8.714 | 3.198 | 3.027 | -9.441 | 4.394 | 3.416 | 1.45 | 74.47 | 121.29 |
| 99 | 2jsa-15 | GLU_A_17_CA | GLU_A_17_HA | -9.441 | 4.394 | 3.416 | -9.68 | 4.313 | 4.476 | 1.09 | 13.39 | -161.28 |
| 100 | 2jsa-15 | GLU_A_17_CA | GLU_A_17_C | -9.441 | 4.394 | 3.416 | -10.752 | 4.495 | 2.633 | 1.53 | 120.77 | 175.59 |
| 101 | 2jsa-15 | GLU_A_17_C | GLU_A_17_O | -10.752 | 4.495 | 2.633 | -11.786 | 4.859 | 3.192 | 1.23 | 62.98 | 160.61 |
| 102 | 2jsa-15 | GLU_A_17_C | GLY_A_18_N | -10.752 | 4.495 | 2.633 | -10.665 | 4.168 | 1.352 | 1.32 | 165.20 | -75.10 |
| 103 | 2jsa-15 | GLY_A_18_N | GLY_A_18_H | -10.665 | 4.168 | 1.352 | -9.78 | 3.878 | 0.96 | 1.01 | 112.83 | -18.14 |
| 104 | 2jsa-15 | GLY_A_18_N | GLY_A_18_CA | -10.665 | 4.168 | 1.352 | -11.832 | 4.218 | 0.486 | 1.45 | 126.55 | 177.55 |
| 105 | 2jsa-15 | GLY_A_18_CA | GLY_A_18_C | -11.832 | 4.218 | 0.486 | -12.197 | 2.823 | -0.023 | 1.53 | 109.44 | -104.66 |
| 106 | 2jsa-15 | GLY_A_18_C | GLY_A_18_O | -12.197 | 2.823 | -0.023 | -12.315 | 2.609 | -1.228 | 1.23 | 168.54 | -118.87 |
| 107 | 2jsa-15 | GLY_A_18_C | SER_A_19_N | -12.197 | 2.823 | -0.023 | -12.366 | 1.908 | 0.922 | 1.33 | 44.56 | -100.46 |
| 108 | 2jsa-15 | SER_A_19_N | SER_A_19_H | -12.366 | 1.908 | 0.922 | -12.253 | 2.155 | 1.895 | 1.01 | 15.60 | 65.42 |
| 109 | 2jsa-15 | SER_A_19_N | SER_A_19_CA | -12.366 | 1.908 | 0.922 | -12.715 | 0.54 | 0.584 | 1.45 | 103.46 | -104.31 |
| 110 | 2jsa-15 | SER_A_19_CA | SER_A_19_HA | -12.715 | 0.54 | 0.584 | -12.327 | 0.324 | -0.411 | 1.09 | 155.95 | -29.10 |
| 111 | 2jsa-15 | SER_A_19_CA | SER_A_19_C | -12.715 | 0.54 | 0.584 | -14.237 | 0.378 | 0.564 | 1.53 | 90.75 | -173.92 |
| 112 | 2jsa-15 | SER_A_19_C | SER_A_19_O | -14.237 | 0.378 | 0.564 | -14.796 | -0.15 | -0.396 | 1.23 | 141.31 | -136.63 |
| 113 | 2jsa-15 | SER_A_19_C | ALA_A_20_N | -14.237 | 0.378 | 0.564 | -14.864 | 0.842 | 1.635 | 1.32 | 36.07 | 143.50 |
| 114 | 2jsa-15 | ALA_A_20_N | ALA_A_20_H | -14.864 | 0.842 | 1.635 | -14.339 | 1.264 | 2.388 | 1.01 | 41.81 | 38.79 |
| 115 | 2jsa-15 | ALA_A_20_N | ALA_A_20_CA | -14.864 | 0.842 | 1.635 | -16.309 | 0.755 | 1.753 | 1.45 | 85.34 | -176.55 |
| 116 | 2jsa-15 | ALA_A_20_CA | ALA_A_20_HA | -16.309 | 0.755 | 1.753 | -16.65 | 1.517 | 2.454 | 1.09 | 49.98 | 114.11 |
| 117 | 2jsa-15 | ALA_A_20_CA | ALA_A_20_C | -16.309 | 0.755 | 1.753 | -16.69 | -0.62 | 2.308 | 1.53 | 68.74 | -105.49 |
| 118 | 2jsa-15 | ALA_A_20_C | ALA_A_20_O | -16.69 | -0.62 | 2.308 | -17.841 | -0.847 | 2.676 | 1.23 | 72.58 | -168.84 |
| 119 | 2jsa-15 | ALA_A_20_C | ASP_A_21_N | -16.69 | -0.62 | 2.308 | -15.7 | -1.5 | 2.35 | 1.33 | 88.18 | -41.63 |
| 120 | 2jsa-15 | ASP_A_21_N | ASP_A_21_H | -15.7 | -1.5 | 2.35 | -14.777 | -1.244 | 2.03 | 1.01 | 108.47 | 15.50 |
| 121 | 2jsa-15 | ASP_A_21_N | ASP_A_21_CA | -15.7 | -1.5 | 2.35 | -15.917 | -2.845 | 2.853 | 1.45 | 69.74 | -99.17 |
| 122 | 2jsa-15 | ASP_A_21_CA | ASP_A_21_HA | -15.917 | -2.845 | 2.853 | -16.975 | -2.975 | 3.077 | 1.09 | 78.13 | -172.99 |
| 123 | 2jsa-15 | ASP_A_21_CA | ASP_A_21_C | -15.917 | -2.845 | 2.853 | -15.088 | -3.05 | 4.122 | 1.53 | 33.94 | -13.89 |
| 124 | 2jsa-15 | ASP_A_21_C | ASP_A_21_O | -15.088 | -3.05 | 4.122 | -15.279 | -4.03 | 4.842 | 1.23 | 54.20 | -101.03 |
| 125 | 2jsa-15 | ASP_A_21_C | LYS_A_22_N | -15.088 | -3.05 | 4.122 | -14.185 | -2.111 | 4.359 | 1.32 | 79.69 | 46.12 |
| 126 | 2jsa-15 | LYS_A_22_N | LYS_A_22_H | -14.185 | -2.111 | 4.359 | -14.082 | -1.333 | 3.723 | 1.01 | 129.02 | 82.46 |
| 127 | 2jsa-15 | LYS_A_22_N | LYS_A_22_CA | -14.185 | -2.111 | 4.359 | -13.326 | -2.175 | 5.529 | 1.45 | 36.36 | -4.26 |
| 128 | 2jsa-15 | LYS_A_22_CA | LYS_A_22_HA | -13.326 | -2.175 | 5.529 | -12.474 | -1.516 | 5.36 | 1.09 | 98.92 | 37.72 |
| 129 | 2jsa-15 | LYS_A_22_CA | LYS_A_22_C | -13.326 | -2.175 | 5.529 | -12.788 | -3.6 | 5.683 | 1.53 | 84.23 | -69.32 |
| 130 | 2jsa-15 | LYS_A_22_C | LYS_A_22_O | -12.788 | -3.6 | 5.683 | -13.081 | -4.274 | 6.669 | 1.23 | 36.70 | -113.50 |
| 131 | 2jsa-15 | LYS_A_22_C | ASP_A_23_N | -12.788 | -3.6 | 5.683 | -12.012 | -4.015 | 4.693 | 1.32 | 138.37 | -28.14 |
| 132 | 2jsa-15 | ASP_A_23_N | ASP_A_23_H | -12.012 | -4.015 | 4.693 | -11.815 | -3.407 | 3.911 | 1.01 | 140.74 | 72.05 |
| 133 | 2jsa-15 | ASP_A_23_N | ASP_A_23_CA | -12.012 | -4.015 | 4.693 | -11.43 | -5.347 | 4.707 | 1.45 | 89.45 | -66.40 |
| 134 | 2jsa-15 | ASP_A_23_CA | ASP_A_23_HA | -11.43 | -5.347 | 4.707 | -11.456 | -5.737 | 5.725 | 1.09 | 21.00 | -93.81 |
| 135 | 2jsa-15 | ASP_A_23_CA | ASP_A_23_C | -11.43 | -5.347 | 4.707 | -9.983 | -5.272 | 4.216 | 1.53 | 108.72 | 2.97 |
| 136 | 2jsa-15 | ASP_A_23_C | ASP_A_23_O | -9.983 | -5.272 | 4.216 | -9.736 | -5.027 | 3.036 | 1.23 | 163.57 | 44.77 |
| 137 | 2jsa-15 | ASP_A_23_C | ALA_A_24_N | -9.983 | -5.272 | 4.216 | -9.065 | -5.487 | 5.146 | 1.32 | 45.39 | -13.18 |

| | | | | | | | | | | | | |
|---|---|---|---|---|---|---|---|---|---|---|---|---|
| 138 | 2jsa-15 | ALA_A_24_N | ALA_A_24_H | -9.065 | -5.487 | 5.146 | -9.34 | -5.682 | 6.098 | 1.01 | 19.50 | -144.66 |
| 139 | 2jsa-15 | ALA_A_24_N | ALA_A_24_CA | -9.065 | -5.487 | 5.146 | -7.649 | -5.447 | 4.822 | 1.45 | 102.88 | 1.62 |
| 140 | 2jsa-15 | ALA_A_24_CA | ALA_A_24_HA | -7.649 | -5.447 | 4.822 | -7.411 | -4.467 | 4.409 | 1.09 | 112.27 | 76.35 |
| 141 | 2jsa-15 | ALA_A_24_CA | ALA_A_24_C | -7.649 | -5.447 | 4.822 | -7.343 | -6.506 | 3.762 | 1.53 | 133.88 | -73.88 |
| 142 | 2jsa-15 | ALA_A_24_C | ALA_A_24_O | -7.343 | -6.506 | 3.762 | -6.275 | -6.486 | 3.15 | 1.23 | 119.81 | 1.07 |
| 143 | 2jsa-15 | ALA_A_24_C | ASN_A_25_N | -7.343 | -6.506 | 3.762 | -8.296 | -7.406 | 3.577 | 1.32 | 98.03 | -136.64 |
| 144 | 2jsa-15 | ASN_A_25_N | ASN_A_25_H | -8.296 | -7.406 | 3.577 | -9.148 | -7.361 | 4.117 | 1.01 | 57.67 | 176.98 |
| 145 | 2jsa-15 | ASN_A_25_N | ASN_A_25_CA | -8.296 | -7.406 | 3.577 | -8.142 | -8.471 | 2.602 | 1.45 | 132.18 | -81.77 |
| 146 | 2jsa-15 | ASN_A_25_CA | ASN_A_25_HA | -8.142 | -8.471 | 2.602 | -7.118 | -8.844 | 2.642 | 1.09 | 87.90 | -20.01 |
| 147 | 2jsa-15 | ASN_A_25_CA | ASN_A_25_C | -8.142 | -8.471 | 2.602 | -8.442 | -7.926 | 1.204 | 1.53 | 156.01 | 118.83 |
| 148 | 2jsa-15 | ASN_A_25_C | ASN_A_25_O | -8.442 | -7.926 | 1.204 | -7.589 | -7.968 | 0.32 | 1.23 | 135.99 | -2.82 |
| 149 | 2jsa-15 | ASN_A_25_C | VAL_A_26_N | -8.442 | -7.926 | 1.204 | -9.659 | -7.424 | 1.049 | 1.33 | 96.72 | 157.58 |
| 150 | 2jsa-15 | VAL_A_26_N | VAL_A_26_H | -9.659 | -7.424 | 1.049 | -10.309 | -7.421 | 1.821 | 1.01 | 40.10 | 179.74 |
| 151 | 2jsa-15 | VAL_A_26_N | VAL_A_26_CA | -9.659 | -7.424 | 1.049 | -10.084 | -6.87 | -0.226 | 1.45 | 151.29 | 127.49 |
| 152 | 2jsa-15 | VAL_A_26_CA | VAL_A_26_HA | -10.084 | -6.87 | -0.226 | -9.82 | -7.576 | -1.013 | 1.09 | 136.24 | -69.50 |
| 153 | 2jsa-15 | VAL_A_26_CA | VAL_A_26_C | -10.084 | -6.87 | -0.226 | -9.329 | -5.565 | -0.485 | 1.53 | 99.75 | 59.95 |
| 154 | 2jsa-15 | VAL_A_26_C | VAL_A_26_O | -9.329 | -5.565 | -0.485 | -9.432 | -4.99 | -1.568 | 1.23 | 151.66 | 100.16 |
| 155 | 2jsa-15 | VAL_A_26_C | ILE_A_27_N | -9.329 | -5.565 | -0.485 | -8.588 | -5.135 | 0.526 | 1.33 | 40.28 | 30.13 |
| 156 | 2jsa-15 | ILE_A_27_N | ILE_A_27_H | -8.588 | -5.135 | 0.526 | -8.55 | -5.66 | 1.388 | 1.01 | 31.41 | -85.86 |
| 157 | 2jsa-15 | ILE_A_27_N | ILE_A_27_CA | -8.588 | -5.135 | 0.526 | -7.816 | -3.908 | 0.42 | 1.45 | 94.18 | 57.82 |
| 158 | 2jsa-15 | ILE_A_27_CA | ILE_A_27_HA | -7.816 | -3.908 | 0.42 | -8.298 | -3.268 | -0.319 | 1.09 | 132.69 | 126.98 |
| 159 | 2jsa-15 | ILE_A_27_CA | ILE_A_27_C | -7.816 | -3.908 | 0.42 | -6.413 | -4.237 | -0.094 | 1.53 | 109.63 | -13.20 |
| 160 | 2jsa-15 | ILE_A_27_C | ILE_A_27_O | -6.413 | -4.237 | -0.094 | -6.0 | -3.743 | -1.142 | 1.23 | 148.43 | 50.10 |
| 161 | 2jsa-15 | ILE_A_27_C | LYS_A_28_N | -6.413 | -4.237 | -0.094 | -5.717 | -5.068 | 0.669 | 1.33 | 54.86 | -50.05 |
| 162 | 2jsa-15 | LYS_A_28_N | LYS_A_28_H | -5.717 | -5.068 | 0.669 | -6.119 | -5.436 | 1.519 | 1.01 | 32.67 | -137.53 |
| 163 | 2jsa-15 | LYS_A_28_N | LYS_A_28_CA | -5.717 | -5.068 | 0.669 | -4.369 | -5.466 | 0.304 | 1.45 | 104.56 | -16.45 |
| 164 | 2jsa-15 | LYS_A_28_CA | LYS_A_28_HA | -4.369 | -5.466 | 0.304 | -3.73 | -4.584 | 0.356 | 1.09 | 87.27 | 54.08 |
| 165 | 2jsa-15 | LYS_A_28_CA | LYS_A_28_C | -4.369 | -5.466 | 0.304 | -4.362 | -5.961 | -1.144 | 1.53 | 161.13 | -89.19 |
| 166 | 2jsa-15 | LYS_A_28_C | LYS_A_28_O | -4.362 | -5.961 | -1.144 | -3.438 | -5.666 | -1.9 | 1.23 | 127.93 | 17.71 |
| 167 | 2jsa-15 | LYS_A_28_C | LYS_A_29_N | -4.362 | -5.961 | -1.144 | -5.404 | -6.703 | -1.488 | 1.32 | 105.05 | -144.55 |
| 168 | 2jsa-15 | LYS_A_29_N | LYS_A_29_H | -5.404 | -6.703 | -1.488 | -6.128 | -6.905 | -0.814 | 1.01 | 48.12 | -164.41 |
| 169 | 2jsa-15 | LYS_A_29_N | LYS_A_29_CA | -5.404 | -6.703 | -1.488 | -5.53 | -7.24 | -2.832 | 1.45 | 157.69 | -103.20 |
| 170 | 2jsa-15 | LYS_A_29_CA | LYS_A_29_HA | -5.53 | -7.24 | -2.832 | -4.581 | -7.705 | -3.1 | 1.09 | 104.23 | -26.10 |
| 171 | 2jsa-15 | LYS_A_29_CA | LYS_A_29_C | -5.53 | -7.24 | -2.832 | -5.779 | -6.094 | -3.813 | 1.53 | 129.91 | 102.26 |
| 172 | 2jsa-15 | LYS_A_29_C | LYS_A_29_O | -5.779 | -6.094 | -3.813 | -5.105 | -5.992 | -4.838 | 1.23 | 146.37 | 8.61 |
| 173 | 2jsa-15 | LYS_A_29_C | ASP_A_30_N | -5.779 | -6.094 | -3.813 | -6.748 | -5.26 | -3.466 | 1.32 | 74.81 | 139.28 |
| 174 | 2jsa-15 | ASP_A_30_N | ASP_A_30_H | -6.748 | -5.26 | -3.466 | -7.261 | -5.405 | -2.608 | 1.01 | 31.85 | -164.22 |
| 175 | 2jsa-15 | ASP_A_30_N | ASP_A_30_CA | -6.748 | -5.26 | -3.466 | -7.094 | -4.125 | -4.304 | 1.45 | 125.23 | 106.95 |
| 176 | 2jsa-15 | ASP_A_30_CA | ASP_A_30_HA | -7.094 | -4.125 | -4.304 | -7.647 | -4.48 | -5.173 | 1.09 | 142.90 | -147.30 |
| 177 | 2jsa-15 | ASP_A_30_CA | ASP_A_30_C | -7.094 | -4.125 | -4.304 | -5.814 | -3.419 | -4.755 | 1.53 | 107.15 | 28.88 |
| 178 | 2jsa-15 | ASP_A_30_C | ASP_A_30_O | -5.814 | -3.419 | -4.755 | -5.633 | -3.153 | -5.941 | 1.23 | 164.82 | 55.77 |
| 179 | 2jsa-15 | ASP_A_30_C | PHE_A_31_N | -5.814 | -3.419 | -4.755 | -4.957 | -3.138 | -3.783 | 1.33 | 42.86 | 18.15 |
| 180 | 2jsa-15 | PHE_A_31_N | PHE_A_31_H | -4.957 | -3.138 | -3.783 | -5.171 | -3.388 | -2.828 | 1.01 | 19.01 | -130.56 |
| 181 | 2jsa-15 | PHE_A_31_N | PHE_A_31_CA | -4.957 | -3.138 | -3.783 | -3.699 | -2.47 | -4.065 | 1.45 | 101.20 | 27.97 |
| 182 | 2jsa-15 | PHE_A_31_CA | PHE_A_31_HA | -3.699 | -2.47 | -4.065 | -3.879 | -1.415 | -4.271 | 1.09 | 100.90 | 99.68 |
| 183 | 2jsa-15 | PHE_A_31_CA | PHE_A_31_C | -3.699 | -2.47 | -4.065 | -2.991 | -3.114 | -5.26 | 1.53 | 141.31 | -42.29 |
| 184 | 2jsa-15 | PHE_A_31_C | PHE_A_31_O | -2.991 | -3.114 | -5.26 | -2.791 | -2.47 | -6.288 | 1.23 | 146.74 | 72.75 |
| 185 | 2jsa-15 | PHE_A_31_C | ASP_A_32_N | -2.991 | -3.114 | -5.26 | -2.631 | -4.376 | -5.082 | 1.32 | 82.28 | -74.08 |
| 186 | 2jsa-15 | ASP_A_32_N | ASP_A_32_H | -2.631 | -4.376 | -5.082 | -2.826 | -4.842 | -4.208 | 1.01 | 30.03 | -112.71 |
| 187 | 2jsa-15 | ASP_A_32_N | ASP_A_32_CA | -2.631 | -4.376 | -5.082 | -1.95 | -5.115 | -6.132 | 1.45 | 136.26 | -47.34 |
| 188 | 2jsa-15 | ASP_A_32_CA | ASP_A_32_HA | -1.95 | -5.115 | -6.132 | -0.912 | -4.784 | -6.185 | 1.09 | 92.79 | 17.69 |
| 189 | 2jsa-15 | ASP_A_32_CA | ASP_A_32_C | -1.95 | -5.115 | -6.132 | -2.647 | -4.854 | -7.468 | 1.53 | 150.88 | 159.47 |
| 190 | 2jsa-15 | ASP_A_32_C | ASP_A_32_O | -2.647 | -4.854 | -7.468 | -1.99 | -4.694 | -8.494 | 1.23 | 146.61 | 13.69 |
| 191 | 2jsa-15 | ASP_A_32_C | ALA_A_33_N | -2.647 | -4.854 | -7.468 | -3.971 | -4.82 | -7.411 | 1.33 | 87.54 | 178.53 |
| 192 | 2jsa-15 | ALA_A_33_N | ALA_A_33_H | -3.971 | -4.82 | -7.411 | -4.445 | -4.962 | -6.531 | 1.01 | 29.35 | -163.32 |
| 193 | 2jsa-15 | ALA_A_33_N | ALA_A_33_CA | -3.971 | -4.82 | -7.411 | -4.766 | -4.581 | -8.604 | 1.45 | 145.17 | 163.27 |
| 194 | 2jsa-15 | ALA_A_33_CA | ALA_A_33_HA | -4.766 | -4.581 | -8.604 | -4.684 | -5.448 | -9.259 | 1.09 | 126.95 | -84.60 |
| 195 | 2jsa-15 | ALA_A_33_CA | ALA_A_33_C | -4.766 | -4.581 | -8.604 | -4.212 | -3.362 | -9.344 | 1.53 | 118.93 | 65.56 |
| 196 | 2jsa-15 | ALA_A_33_C | ALA_A_33_O | -4.212 | -3.362 | -9.344 | -3.754 | -3.476 | -10.481 | 1.23 | 157.46 | -13.98 |
| 197 | 2jsa-15 | ALA_A_33_C | GLU_A_34_N | -4.212 | -3.362 | -9.344 | -4.274 | -2.223 | -8.671 | 1.32 | 59.46 | 93.12 |
| 198 | 2jsa-15 | GLU_A_34_N | GLU_A_34_H | -4.274 | -2.223 | -8.671 | -4.665 | -2.203 | -7.74 | 1.01 | 22.81 | 177.07 |
| 199 | 2jsa-15 | GLU_A_34_N | GLU_A_34_CA | -4.274 | -2.223 | -8.671 | -3.784 | -0.984 | -9.25 | 1.45 | 113.49 | 68.42 |
| 200 | 2jsa-15 | GLU_A_34_CA | GLU_A_34_HA | -3.784 | -0.984 | -9.25 | -4.008 | -0.999 | -10.316 | 1.09 | 168.11 | -176.17 |
| 201 | 2jsa-15 | GLU_A_34_CA | GLU_A_34_C | -3.784 | -0.984 | -9.25 | -2.268 | -0.88 | -9.081 | 1.53 | 83.65 | 3.92 |
| 202 | 2jsa-15 | GLU_A_34_C | GLU_A_34_O | -2.268 | -0.88 | -9.081 | -1.691 | 0.194 | -9.252 | 1.23 | 97.98 | 61.75 |
| 203 | 2jsa-15 | GLU_A_34_C | CYS_A_35_N | -2.268 | -0.88 | -9.081 | -1.663 | -2.011 | -8.745 | 1.33 | 75.32 | -61.86 |
| 204 | 2jsa-15 | CYS_A_35_N | CYS_A_35_H | -1.663 | -2.011 | -8.745 | -2.199 | -2.857 | -8.62 | 1.01 | 82.89 | -122.36 |
| 205 | 2jsa-15 | CYS_A_35_N | CYS_A_35_CA | -1.663 | -2.011 | -8.745 | -0.225 | -2.061 | -8.55 | 1.45 | 82.28 | -1.99 |
| 206 | 2jsa-15 | CYS_A_35_CA | CYS_A_35_HA | -0.225 | -2.061 | -8.55 | 0.139 | -1.042 | -8.416 | 1.09 | 82.94 | 70.34 |
| 207 | 2jsa-15 | CYS_A_35_CA | CYS_A_35_C | -0.225 | -2.061 | -8.55 | 0.412 | -2.642 | -9.814 | 1.53 | 145.70 | -42.37 |
| 208 | 2jsa-15 | CYS_A_35_C | CYS_A_35_O | 0.412 | -2.642 | -9.814 | 1.179 | -1.965 | -10.496 | 1.23 | 123.69 | 41.43 |
| 209 | 2jsa-15 | CYS_A_35_C | LYS_A_36_N | 0.412 | -2.642 | -9.814 | 0.07 | -3.893 | -10.089 | 1.33 | 101.97 | -105.29 |
| 210 | 2jsa-15 | LYS_A_36_N | LYS_A_36_H | 0.07 | -3.893 | -10.089 | -0.567 | -4.39 | -9.483 | 1.01 | 53.13 | -142.04 |
| 211 | 2jsa-15 | LYS_A_36_N | LYS_A_36_CA | 0.07 | -3.893 | -10.089 | 0.598 | -4.574 | -11.258 | 1.45 | 143.60 | -52.21 |
| 212 | 2jsa-15 | LYS_A_36_CA | LYS_A_36_HA | 0.598 | -4.574 | -11.258 | 1.687 | -4.557 | -11.195 | 1.09 | 86.69 | 0.89 |
| 213 | 2jsa-15 | LYS_A_36_CA | LYS_A_36_C | 0.598 | -4.574 | -11.258 | 0.193 | -3.805 | -12.517 | 1.53 | 145.38 | 117.77 |
| 214 | 2jsa-15 | LYS_A_36_C | LYS_A_36_O | 0.193 | -3.805 | -12.517 | 0.907 | -3.824 | -13.518 | 1.23 | 144.49 | -1.52 |
| 215 | 2jsa-15 | LYS_A_36_C | LYS_A_37_N | 0.193 | -3.805 | -12.517 | -0.954 | -3.146 | -12.426 | 1.33 | 86.06 | 150.12 |
| 216 | 2jsa-15 | LYS_A_37_N | LYS_A_37_H | -0.954 | -3.146 | -12.426 | -1.49 | -3.175 | -11.571 | 1.01 | 32.12 | -176.90 |
| 217 | 2jsa-15 | LYS_A_37_N | LYS_A_37_CA | -0.954 | -3.146 | -12.426 | -1.463 | -2.372 | -13.545 | 1.45 | 140.38 | 123.33 |
| 218 | 2jsa-15 | LYS_A_37_CA | LYS_A_37_HA | -1.463 | -2.372 | -13.545 | -1.226 | -2.91 | -14.463 | 1.09 | 147.36 | -66.23 |
| 219 | 2jsa-15 | LYS_A_37_CA | LYS_A_37_C | -1.463 | -2.372 | -13.545 | -0.741 | -1.024 | -13.598 | 1.53 | 91.99 | 61.83 |
| 220 | 2jsa-15 | LYS_A_37_C | LYS_A_37_O | -0.741 | -1.024 | -13.598 | -0.173 | -0.659 | -14.625 | 1.23 | 146.68 | 32.73 |
| 221 | 2jsa-15 | LYS_A_37_C | LEU_A_38_N | -0.741 | -1.024 | -13.598 | -0.788 | -0.32 | -12.476 | 1.33 | 32.16 | 93.82 |
| 222 | 2jsa-15 | LEU_A_38_N | LEU_A_38_H | -0.788 | -0.32 | -12.476 | -1.275 | -0.682 | -11.668 | 1.01 | 36.91 | -143.38 |
| 223 | 2jsa-15 | LEU_A_38_N | LEU_A_38_CA | -0.788 | -0.32 | -12.476 | -0.145 | 0.98 | -12.38 | 1.45 | 86.21 | 63.68 |
| 224 | 2jsa-15 | LEU_A_38_CA | LEU_A_38_HA | -0.145 | 0.98 | -12.38 | -0.547 | 1.614 | -13.17 | 1.09 | 136.46 | 122.38 |
| 225 | 2jsa-15 | LEU_A_38_CA | LEU_A_38_C | -0.145 | 0.98 | -12.38 | 1.356 | 0.822 | -12.63 | 1.53 | 99.41 | -6.01 |
| 226 | 2jsa-15 | LEU_A_38_C | LEU_A_38_O | 1.356 | 0.822 | -12.63 | 1.908 | 1.448 | -13.534 | 1.23 | 137.29 | 48.59 |
| 227 | 2jsa-15 | LEU_A_38_C | PHE_A_39_N | 1.356 | 0.822 | -12.63 | 1.974 | -0.019 | -11.814 | 1.32 | 51.98 | -53.69 |
| 228 | 2jsa-15 | PHE_A_39_N | PHE_A_39_H | 1.974 | -0.019 | -11.814 | 1.457 | -0.501 | -11.093 | 1.01 | 44.43 | -137.01 |
| 229 | 2jsa-15 | PHE_A_39_N | PHE_A_39_CA | 1.974 | -0.019 | -11.814 | 3.401 | -0.267 | -11.935 | 1.45 | 94.78 | -9.86 |
| 230 | 2jsa-15 | PHE_A_39_CA | PHE_A_39_HA | 3.401 | -0.267 | -11.935 | 3.908 | 0.639 | -12.268 | 1.09 | 107.78 | 60.77 |
| 231 | 2jsa-15 | PHE_A_39_CA | PHE_A_39_C | 3.401 | -0.267 | -11.935 | 3.677 | -1.403 | -12.922 | 1.53 | 130.17 | -76.34 |
| 232 | 2jsa-15 | PHE_A_39_C | PHE_A_39_O | 3.677 | -1.403 | -12.922 | 4.659 | -2.13 | -12.777 | 1.23 | 83.23 | -36.51 |
| 233 | 2jsa-15 | PHE_A_39_C | HIS_A_40_N | 3.677 | -1.403 | -12.922 | 2.793 | -1.522 | -13.902 | 1.33 | 137.69 | -172.33 |

| | | | | | | | | | | | | |
|---|---|---|---|---|---|---|---|---|---|---|---|---|
| 234 | 2jsa-15 | HIS_A_40_N | HIS_A_40_H | 2.793 | -1.522 | -13.902 | 2.008 | -0.889 | -13.958 | 1.01 | 93.18 | 141.12 |
| 235 | 2jsa-15 | HIS_A_40_N | HIS_A_40_CA | 2.793 | -1.522 | -13.902 | 2.93 | -2.558 | -14.912 | 1.45 | 134.02 | -82.47 |
| 236 | 2jsa-15 | HIS_A_40_CA | HIS_A_40_HA | 2.93 | -2.558 | -14.912 | 2.915 | -3.524 | -14.407 | 1.09 | 62.40 | -90.89 |
| 237 | 2jsa-15 | HIS_A_40_CA | HIS_A_40_C | 2.93 | -2.558 | -14.912 | 4.282 | -2.415 | -15.614 | 1.53 | 117.31 | 6.04 |
| 238 | 2jsa-15 | HIS_A_40_C | HIS_A_40_O | 4.282 | -2.415 | -15.614 | 4.759 | -3.356 | -16.246 | 1.23 | 120.92 | -63.12 |
| 239 | 2jsa-15 | HIS_A_40_C | THR_A_41_N | 4.282 | -2.415 | -15.614 | 4.861 | -1.231 | -15.477 | 1.33 | 84.07 | 63.94 |
| 240 | 2jsa-15 | THR_A_41_N | THR_A_41_H | 4.861 | -1.231 | -15.477 | 4.411 | -0.503 | -14.941 | 1.01 | 57.94 | 121.72 |
| 241 | 2jsa-15 | THR_A_41_N | THR_A_41_CA | 4.861 | -1.231 | -15.477 | 6.15 | -0.953 | -16.09 | 1.45 | 114.93 | 12.17 |
| 242 | 2jsa-15 | THR_A_41_CA | THR_A_41_HA | 6.15 | -0.953 | -16.09 | 6.563 | -1.885 | -16.476 | 1.09 | 110.74 | -66.10 |
| 243 | 2jsa-15 | THR_A_41_CA | THR_A_41_C | 6.15 | -0.953 | -16.09 | 7.127 | -0.401 | -15.05 | 1.53 | 47.18 | 29.47 |
| 244 | 2jsa-15 | THR_A_41_C | THR_A_41_O | 7.127 | -0.401 | -15.05 | 7.449 | 0.786 | -15.065 | 1.23 | 90.70 | 74.82 |
| 245 | 2jsa-15 | THR_A_41_C | ILE_A_42_N | 7.127 | -0.401 | -15.05 | 7.572 | -1.289 | -14.173 | 1.33 | 48.56 | -63.38 |
| 246 | 2jsa-15 | ILE_A_42_N | ILE_A_42_H | 7.572 | -1.289 | -14.173 | 7.266 | -2.25 | -14.221 | 1.01 | 92.72 | -107.66 |
| 247 | 2jsa-15 | ILE_A_42_N | ILE_A_42_CA | 7.572 | -1.289 | -14.173 | 8.506 | -0.905 | -13.129 | 1.45 | 44.05 | 22.35 |
| 248 | 2jsa-15 | ILE_A_42_CA | ILE_A_42_HA | 8.506 | -0.905 | -13.129 | 9.146 | -0.114 | -13.522 | 1.09 | 111.12 | 51.02 |
| 249 | 2jsa-15 | ILE_A_42_CA | ILE_A_42_C | 8.506 | -0.905 | -13.129 | 9.4 | -2.098 | -12.786 | 1.53 | 77.04 | -53.15 |
| 250 | 2jsa-15 | ILE_A_42_C | ILE_A_42_O | 9.4 | -2.098 | -12.786 | 8.939 | -3.238 | -12.767 | 1.23 | 89.11 | -112.02 |
| 251 | 2jsa-15 | ILE_A_42_C | PRO_A_43_N | 9.4 | -2.098 | -12.786 | 10.696 | -1.787 | -12.517 | 1.36 | 78.59 | 13.49 |
| 252 | 2jsa-15 | PRO_A_43_N | PRO_A_43_CA | 10.696 | -1.787 | -12.517 | 11.659 | -2.821 | -12.177 | 1.45 | 76.47 | -47.04 |
| 253 | 2jsa-15 | PRO_A_43_CA | PRO_A_43_HA | 11.659 | -2.821 | -12.177 | 11.554 | -3.667 | -12.856 | 1.09 | 128.54 | -97.07 |
| 254 | 2jsa-15 | PRO_A_43_CA | PRO_A_43_C | 11.659 | -2.821 | -12.177 | 11.446 | -3.317 | -10.745 | 1.53 | 20.65 | -113.24 |
| 255 | 2jsa-15 | PRO_A_43_C | PRO_A_43_O | 11.446 | -3.317 | -10.745 | 11.763 | -2.616 | -9.787 | 1.23 | 38.77 | 65.67 |
| 256 | 2jsa-15 | PRO_A_43_C | PHE_A_44_N | 11.446 | -3.317 | -10.745 | 10.909 | -4.526 | -10.646 | 1.33 | 85.72 | -113.95 |
| 257 | 2jsa-15 | PHE_A_44_N | PHE_A_44_H | 10.909 | -4.526 | -10.646 | 10.673 | -5.047 | -11.478 | 1.01 | 145.49 | -114.37 |
| 258 | 2jsa-15 | PHE_A_44_N | PHE_A_44_CA | 10.909 | -4.526 | -10.646 | 10.65 | -5.125 | -9.349 | 1.45 | 26.71 | -113.38 |
| 259 | 2jsa-15 | PHE_A_44_CA | PHE_A_44_HA | 10.65 | -5.125 | -9.349 | 10.528 | -6.203 | -9.457 | 1.09 | 95.69 | -96.46 |
| 260 | 2jsa-15 | PHE_A_44_CA | PHE_A_44_C | 10.65 | -5.125 | -9.349 | 9.395 | -4.526 | -8.709 | 1.53 | 65.29 | 154.49 |
| 261 | 2jsa-15 | PHE_A_44_C | PHE_A_44_O | 9.395 | -4.526 | -8.709 | 8.974 | -4.959 | -7.637 | 1.23 | 29.40 | -134.19 |
| 262 | 2jsa-15 | PHE_A_44_C | GLY_A_45_N | 9.395 | -4.526 | -8.709 | 8.834 | -3.541 | -9.394 | 1.32 | 121.14 | 119.66 |
| 263 | 2jsa-15 | GLY_A_45_N | GLY_A_45_H | 8.834 | -3.541 | -9.394 | 9.236 | -3.237 | -10.269 | 1.01 | 150.06 | 37.10 |
| 264 | 2jsa-15 | GLY_A_45_N | GLY_A_45_CA | 8.834 | -3.541 | -9.394 | 7.636 | -2.879 | -8.906 | 1.45 | 70.38 | 151.08 |
| 265 | 2jsa-15 | GLY_A_45_CA | GLY_A_45_C | 7.636 | -2.879 | -8.906 | 6.564 | -3.899 | -8.518 | 1.53 | 75.31 | -136.42 |
| 266 | 2jsa-15 | GLY_A_45_C | GLY_A_45_O | 6.564 | -3.899 | -8.518 | 6.134 | -3.945 | -7.366 | 1.23 | 20.58 | -173.89 |
| 267 | 2jsa-15 | GLY_A_45_C | THR_A_46_N | 6.564 | -3.899 | -8.518 | 6.164 | -4.692 | -9.501 | 1.32 | 137.90 | -116.77 |
| 268 | 2jsa-15 | THR_A_46_N | THR_A_46_H | 6.164 | -4.692 | -9.501 | 6.563 | -4.598 | -10.424 | 1.01 | 156.05 | 13.26 |
| 269 | 2jsa-15 | THR_A_46_N | THR_A_46_CA | 6.164 | -4.692 | -9.501 | 5.151 | -5.709 | -9.276 | 1.45 | 81.09 | -134.89 |
| 270 | 2jsa-15 | THR_A_46_CA | THR_A_46_HA | 5.151 | -5.709 | -9.276 | 4.242 | -5.225 | -8.918 | 1.09 | 70.83 | 151.97 |
| 271 | 2jsa-15 | THR_A_46_CA | THR_A_46_C | 5.151 | -5.709 | -9.276 | 5.614 | -6.695 | -8.201 | 1.53 | 45.38 | -64.85 |
| 272 | 2jsa-15 | THR_A_46_C | THR_A_46_O | 5.614 | -6.695 | -8.201 | 4.803 | -7.196 | -7.423 | 1.23 | 50.78 | -148.29 |
| 273 | 2jsa-15 | THR_A_46_C | ARG_A_47_N | 5.614 | -6.695 | -8.201 | 6.915 | -6.944 | -8.193 | 1.32 | 89.65 | -10.83 |
| 274 | 2jsa-15 | ARG_A_47_N | ARG_A_47_H | 6.915 | -6.944 | -8.193 | 7.523 | -6.497 | -8.864 | 1.01 | 131.64 | 36.32 |
| 275 | 2jsa-15 | ARG_A_47_N | ARG_A_47_CA | 6.915 | -6.944 | -8.193 | 7.495 | -7.86 | -7.227 | 1.45 | 48.30 | -57.66 |
| 276 | 2jsa-15 | ARG_A_47_CA | ARG_A_47_HA | 7.495 | -7.86 | -7.227 | 6.986 | -8.82 | -7.314 | 1.09 | 94.58 | -117.93 |
| 277 | 2jsa-15 | ARG_A_47_CA | ARG_A_47_C | 7.495 | -7.86 | -7.227 | 7.307 | -7.324 | -5.806 | 1.53 | 21.79 | 109.33 |
| 278 | 2jsa-15 | ARG_A_47_C | ARG_A_47_O | 7.307 | -7.324 | -5.806 | 6.924 | -8.067 | -4.904 | 1.23 | 42.82 | -117.27 |
| 279 | 2jsa-15 | ARG_A_47_C | GLU_A_48_N | 7.307 | -7.324 | -5.806 | 7.585 | -6.037 | -5.652 | 1.33 | 83.33 | 77.81 |
| 280 | 2jsa-15 | GLU_A_48_N | GLU_A_48_H | 7.585 | -6.037 | -5.652 | 7.896 | -5.486 | -6.439 | 1.01 | 141.20 | 60.56 |
| 281 | 2jsa-15 | GLU_A_48_N | GLU_A_48_CA | 7.585 | -6.037 | -5.652 | 7.45 | -5.393 | -4.357 | 1.45 | 26.94 | 101.84 |
| 282 | 2jsa-15 | GLU_A_48_CA | GLU_A_48_HA | 7.45 | -5.393 | -4.357 | 7.954 | -6.015 | -3.617 | 1.09 | 47.25 | -50.98 |
| 283 | 2jsa-15 | GLU_A_48_CA | GLU_A_48_C | 7.45 | -5.393 | -4.357 | 5.974 | -5.283 | -3.969 | 1.53 | 75.31 | 175.74 |
| 284 | 2jsa-15 | GLU_A_48_C | GLU_A_48_O | 5.974 | -5.283 | -3.969 | 5.631 | -5.352 | -2.789 | 1.23 | 16.52 | -168.63 |
| 285 | 2jsa-15 | GLU_A_48_C | CYS_A_49_N | 5.974 | -5.283 | -3.969 | 5.139 | -5.115 | -4.984 | 1.33 | 140.00 | 168.62 |
| 286 | 2jsa-15 | CYS_A_49_N | CYS_A_49_H | 5.139 | -5.115 | -4.984 | 5.491 | -5.069 | -5.929 | 1.01 | 159.41 | 7.45 |
| 287 | 2jsa-15 | CYS_A_49_N | CYS_A_49_CA | 5.139 | -5.115 | -4.984 | 3.708 | -4.995 | -4.765 | 1.45 | 81.33 | 175.21 |
| 288 | 2jsa-15 | CYS_A_49_CA | CYS_A_49_HA | 3.708 | -4.995 | -4.765 | 3.517 | -4.048 | -4.26 | 1.09 | 62.40 | 101.40 |
| 289 | 2jsa-15 | CYS_A_49_CA | CYS_A_49_C | 3.708 | -4.995 | -4.765 | 3.266 | -6.135 | -3.845 | 1.53 | 53.04 | -111.19 |
| 290 | 2jsa-15 | CYS_A_49_C | CYS_A_49_O | 3.266 | -6.135 | -3.845 | 2.842 | -5.896 | -2.715 | 1.23 | 23.30 | 150.59 |
| 291 | 2jsa-15 | CYS_A_49_C | ASP_A_50_N | 3.266 | -6.135 | -3.845 | 3.381 | -7.349 | -4.362 | 1.32 | 112.98 | -84.59 |
| 292 | 2jsa-15 | ASP_A_50_N | ASP_A_50_H | 3.381 | -7.349 | -4.362 | 3.738 | -7.471 | -5.299 | 1.01 | 158.07 | -18.87 |
| 293 | 2jsa-15 | ASP_A_50_N | ASP_A_50_CA | 3.381 | -7.349 | -4.362 | 2.999 | -8.526 | -3.602 | 1.45 | 58.44 | -107.98 |
| 294 | 2jsa-15 | ASP_A_50_CA | ASP_A_50_HA | 2.999 | -8.526 | -3.602 | 1.927 | -8.493 | -3.41 | 1.09 | 79.85 | 178.24 |
| 295 | 2jsa-15 | ASP_A_50_CA | ASP_A_50_C | 2.999 | -8.526 | -3.602 | 3.764 | -8.543 | -2.277 | 1.53 | 30.01 | -1.27 |
| 296 | 2jsa-15 | ASP_A_50_C | ASP_A_50_O | 3.764 | -8.543 | -2.277 | 3.272 | -9.066 | -1.278 | 1.23 | 35.71 | -133.25 |
| 297 | 2jsa-15 | ASP_A_50_C | HIS_A_51_N | 3.764 | -8.543 | -2.277 | 4.955 | -7.964 | -2.311 | 1.32 | 91.47 | 25.93 |
| 298 | 2jsa-15 | HIS_A_51_N | HIS_A_51_H | 4.955 | -7.964 | -2.311 | 5.297 | -7.553 | -3.168 | 1.01 | 148.04 | 50.24 |
| 299 | 2jsa-15 | HIS_A_51_N | HIS_A_51_CA | 4.955 | -7.964 | -2.311 | 5.794 | -7.906 | -1.126 | 1.45 | 35.36 | 3.95 |
| 300 | 2jsa-15 | HIS_A_51_CA | HIS_A_51_HA | 5.794 | -7.906 | -1.126 | 5.602 | -8.8 | -0.533 | 1.09 | 57.04 | -102.12 |
| 301 | 2jsa-15 | HIS_A_51_CA | HIS_A_51_C | 5.794 | -7.906 | -1.126 | 5.405 | -6.69 | -0.282 | 1.53 | 56.53 | 107.74 |
| 302 | 2jsa-15 | HIS_A_51_C | HIS_A_51_O | 5.405 | -6.69 | -0.282 | 6.1 | -6.344 | 0.672 | 1.23 | 39.14 | 26.47 |
| 303 | 2jsa-15 | HIS_A_51_C | TYR_A_52_N | 5.405 | -6.69 | -0.282 | 4.294 | -6.076 | -0.662 | 1.33 | 106.67 | 151.07 |
| 304 | 2jsa-15 | TYR_A_52_N | TYR_A_52_H | 4.294 | -6.076 | -0.662 | 3.772 | -6.416 | -1.456 | 1.01 | 141.88 | -146.92 |
| 305 | 2jsa-15 | TYR_A_52_N | TYR_A_52_CA | 4.294 | -6.076 | -0.662 | 3.804 | -4.907 | 0.047 | 1.45 | 60.78 | 112.74 |
| 306 | 2jsa-15 | TYR_A_52_CA | TYR_A_52_HA | 3.804 | -4.907 | 0.047 | 4.531 | -4.587 | 0.794 | 1.09 | 46.76 | 23.76 |
| 307 | 2jsa-15 | TYR_A_52_CA | TYR_A_52_C | 3.804 | -4.907 | 0.047 | 2.462 | -5.197 | 0.721 | 1.53 | 63.85 | -167.81 |
| 308 | 2jsa-15 | TYR_A_52_C | TYR_A_52_O | 2.462 | -5.197 | 0.721 | 2.223 | -4.766 | 1.848 | 1.23 | 23.62 | 119.01 |
| 309 | 2jsa-15 | TYR_A_52_C | VAL_A_53_N | 2.462 | -5.197 | 0.721 | 1.62 | -5.926 | 0.001 | 1.33 | 122.88 | -139.11 |
| 310 | 2jsa-15 | VAL_A_53_N | VAL_A_53_H | 1.62 | -5.926 | 0.001 | 1.884 | -6.247 | -0.919 | 1.01 | 155.69 | -50.57 |
| 311 | 2jsa-15 | VAL_A_53_N | VAL_A_53_CA | 1.62 | -5.926 | 0.001 | 0.308 | -6.278 | 0.515 | 1.45 | 69.27 | -164.98 |
| 312 | 2jsa-15 | VAL_A_53_CA | VAL_A_53_HA | 0.308 | -6.278 | 0.515 | -0.054 | -5.453 | 1.129 | 1.09 | 55.72 | 113.69 |
| 313 | 2jsa-15 | VAL_A_53_CA | VAL_A_53_C | 0.308 | -6.278 | 0.515 | 0.429 | -7.516 | 1.406 | 1.53 | 54.39 | -84.42 |
| 314 | 2jsa-15 | VAL_A_53_C | VAL_A_53_O | 0.429 | -7.516 | 1.406 | -0.289 | -8.496 | 1.214 | 1.23 | 98.98 | -126.23 |
| 315 | 2jsa-15 | VAL_A_53_C | ASN_A_54_N | 0.429 | -7.516 | 1.406 | 1.343 | -7.43 | 2.362 | 1.33 | 43.84 | 5.38 |
| 316 | 2jsa-15 | ASN_A_54_N | ASN_A_54_H | 1.343 | -7.43 | 2.362 | 1.898 | -6.592 | 2.462 | 1.01 | 84.32 | 56.48 |
| 317 | 2jsa-15 | ASN_A_54_N | ASN_A_54_CA | 1.343 | -7.43 | 2.362 | 1.567 | -8.531 | 3.283 | 1.45 | 50.66 | -78.50 |
| 318 | 2jsa-15 | ASN_A_54_CA | ASN_A_54_HA | 1.567 | -8.531 | 3.283 | 0.684 | -8.658 | 3.909 | 1.09 | 54.94 | -171.82 |
| 319 | 2jsa-15 | ASN_A_54_CA | ASN_A_54_C | 1.567 | -8.531 | 3.283 | 2.784 | -8.219 | 4.157 | 1.53 | 55.18 | 14.38 |
| 320 | 2jsa-15 | ASN_A_54_C | ASN_A_54_O | 2.784 | -8.219 | 4.157 | 3.629 | -9.083 | 4.382 | 1.23 | 79.45 | -45.64 |
| 321 | 2jsa-15 | ASN_A_54_C | SER_A_55_N | 2.784 | -8.219 | 4.157 | 2.832 | -6.981 | 4.626 | 1.32 | 69.27 | 87.78 |
| 322 | 2jsa-15 | SER_A_55_N | SER_A_55_H | 2.832 | -6.981 | 4.626 | 2.101 | -6.322 | 4.4 | 1.01 | 102.93 | 137.97 |
| 323 | 2jsa-15 | SER_A_55_N | SER_A_55_CA | 2.832 | -6.981 | 4.626 | 3.931 | -6.544 | 5.472 | 1.45 | 54.42 | 21.68 |
| 324 | 2jsa-15 | SER_A_55_CA | SER_A_55_HA | 3.931 | -6.544 | 5.472 | 3.812 | -7.005 | 6.453 | 1.09 | 25.89 | -104.47 |
| 325 | 2jsa-15 | SER_A_55_CA | SER_A_55_C | 3.931 | -6.544 | 5.472 | 3.89 | -5.023 | 5.637 | 1.53 | 83.81 | 91.54 |
| 326 | 2jsa-15 | SER_A_55_C | SER_A_55_O | 3.89 | -5.023 | 5.637 | 4.296 | -4.497 | 6.671 | 1.23 | 32.73 | 52.34 |
| 327 | 2jsa-15 | SER_A_55_C | LYS_A_56_N | 3.89 | -5.023 | 5.637 | 3.398 | -4.36 | 4.601 | 1.32 | 141.45 | 126.58 |
| 328 | 2jsa-15 | LYS_A_56_N | LYS_A_56_H | 3.398 | -4.36 | 4.601 | 3.083 | -4.859 | 3.781 | 1.01 | 144.26 | -122.26 |
| 329 | 2jsa-15 | LYS_A_56_N | LYS_A_56_CA | 3.398 | -4.36 | 4.601 | 3.3 | -2.911 | 4.618 | 1.45 | 89.33 | 93.87 |

| | | | | | | | | | | | |
|---|---|---|---|---|---|---|---|---|---|---|---|
| 330 | 2jsa-15 | LYS_A_56_CA | LYS_A_56_HA | 3.3 | -2.911 | 4.618 | 3.695 | -2.557 | 5.571 | 1.09 | 29.10 | 41.87 |
| 331 | 2jsa-15 | LYS_A_56_CA | LYS_A_56_C | 3.3 | -2.911 | 4.618 | 1.826 | -2.503 | 4.544 | 1.53 | 92.77 | 164.53 |
| 332 | 2jsa-15 | LYS_A_56_C | LYS_A_56_O | 1.826 | -2.503 | 4.544 | 1.502 | -1.319 | 4.618 | 1.23 | 86.55 | 105.30 |
| 333 | 2jsa-15 | LYS_A_56_C | VAL_A_57_N | 1.826 | -2.503 | 4.544 | 0.974 | -3.506 | 4.399 | 1.32 | 96.29 | -130.35 |
| 334 | 2jsa-15 | VAL_A_57_N | VAL_A_57_H | 0.974 | -3.506 | 4.399 | 1.312 | -4.456 | 4.346 | 1.01 | 93.01 | -70.41 |
| 335 | 2jsa-15 | VAL_A_57_N | VAL_A_57_CA | 0.974 | -3.506 | 4.399 | -0.456 | -3.267 | 4.314 | 1.45 | 93.36 | 170.51 |
| 336 | 2jsa-15 | VAL_A_57_CA | VAL_A_57_HA | -0.456 | -3.267 | 4.314 | -0.615 | -2.224 | 4.041 | 1.09 | 104.51 | 98.67 |
| 337 | 2jsa-15 | VAL_A_57_CA | VAL_A_57_C | -0.456 | -3.267 | 4.314 | -1.091 | -3.501 | 5.687 | 1.53 | 26.24 | -159.77 |
| 338 | 2jsa-15 | VAL_A_57_C | VAL_A_57_O | -1.091 | -3.501 | 5.687 | -2.188 | -3.015 | 5.959 | 1.23 | 77.23 | 156.11 |
| 339 | 2jsa-15 | VAL_A_57_C | ASP_A_58_N | -1.091 | -3.501 | 5.687 | -0.373 | -4.245 | 6.516 | 1.33 | 51.28 | -46.02 |
| 340 | 2jsa-15 | ASP_A_58_N | ASP_A_58_H | -0.373 | -4.245 | 6.516 | 0.522 | -4.612 | 6.225 | 1.01 | 106.74 | -22.30 |
| 341 | 2jsa-15 | ASP_A_58_N | ASP_A_58_CA | -0.373 | -4.245 | 6.516 | -0.852 | -4.549 | 7.854 | 1.45 | 22.98 | -147.60 |
| 342 | 2jsa-15 | ASP_A_58_CA | ASP_A_58_HA | -0.852 | -4.549 | 7.854 | -1.915 | -4.786 | 7.807 | 1.09 | 92.47 | -167.43 |
| 343 | 2jsa-15 | ASP_A_58_CA | ASP_A_58_C | -0.852 | -4.549 | 7.854 | -0.63 | -3.335 | 8.758 | 1.53 | 53.78 | 79.64 |
| 344 | 2jsa-15 | ASP_A_58_C | ASP_A_58_O | -0.63 | -3.335 | 8.758 | -1.518 | -2.951 | 9.517 | 1.23 | 51.89 | 156.61 |
| 345 | 2jsa-15 | ASP_A_58_C | PRO_A_59_N | -0.63 | -3.335 | 8.758 | 0.593 | -2.751 | 8.645 | 1.36 | 94.77 | 25.53 |
| 346 | 2jsa-15 | PRO_A_59_N | PRO_A_59_CA | 0.593 | -2.751 | 8.645 | 0.944 | -1.588 | 9.444 | 1.45 | 56.67 | 73.21 |
| 347 | 2jsa-15 | PRO_A_59_CA | PRO_A_59_HA | 0.944 | -1.588 | 9.444 | 0.637 | -1.739 | 10.479 | 1.09 | 18.29 | -153.81 |
| 348 | 2jsa-15 | PRO_A_59_CA | PRO_A_59_C | 0.944 | -1.588 | 9.444 | 0.255 | -0.331 | 8.913 | 1.53 | 110.33 | 118.73 |
| 349 | 2jsa-15 | PRO_A_59_C | PRO_A_59_O | 0.255 | -0.331 | 8.913 | -0.377 | 0.403 | 9.672 | 1.23 | 51.92 | 130.73 |
| 350 | 2jsa-15 | PRO_A_59_C | ILE_A_60_N | 0.255 | -0.331 | 8.913 | 0.401 | -0.118 | 7.613 | 1.33 | 168.76 | 55.57 |
| 351 | 2jsa-15 | ILE_A_60_N | ILE_A_60_H | 0.401 | -0.118 | 7.613 | 0.935 | -0.763 | 7.049 | 1.01 | 123.96 | -50.38 |
| 352 | 2jsa-15 | ILE_A_60_N | ILE_A_60_CA | 0.401 | -0.118 | 7.613 | -0.2 | 1.039 | 6.972 | 1.45 | 116.18 | 117.45 |
| 353 | 2jsa-15 | ILE_A_60_CA | ILE_A_60_HA | -0.2 | 1.039 | 6.972 | 0.327 | 1.928 | 7.319 | 1.09 | 71.44 | 59.34 |
| 354 | 2jsa-15 | ILE_A_60_CA | ILE_A_60_C | -0.2 | 1.039 | 6.972 | -1.659 | 1.162 | 7.414 | 1.53 | 73.20 | 175.18 |
| 355 | 2jsa-15 | ILE_A_60_C | ILE_A_60_O | -1.659 | 1.162 | 7.414 | -2.101 | 2.236 | 7.82 | 1.23 | 70.73 | 112.37 |
| 356 | 2jsa-15 | ILE_A_60_C | ILE_A_61_N | -1.659 | 1.162 | 7.414 | -2.368 | 0.046 | 7.322 | 1.33 | 93.98 | -122.43 |
| 357 | 2jsa-15 | ILE_A_61_N | ILE_A_61_H | -2.368 | 0.046 | 7.322 | -1.939 | -0.803 | 6.981 | 1.01 | 109.72 | -63.19 |
| 358 | 2jsa-15 | ILE_A_61_N | ILE_A_61_CA | -2.368 | 0.046 | 7.322 | -3.769 | 0.016 | 7.708 | 1.45 | 74.60 | -178.77 |
| 359 | 2jsa-15 | ILE_A_61_CA | ILE_A_61_HA | -3.769 | 0.016 | 7.708 | -4.276 | 0.838 | 7.203 | 1.09 | 117.60 | 121.67 |
| 360 | 2jsa-15 | ILE_A_61_CA | ILE_A_61_C | -3.769 | 0.016 | 7.708 | -3.884 | 0.255 | 9.215 | 1.53 | 9.98 | 115.70 |
| 361 | 2jsa-15 | ILE_A_61_C | ILE_A_61_O | -3.884 | 0.255 | 9.215 | -4.91 | 0.735 | 9.694 | 1.23 | 67.08 | 154.93 |
| 362 | 2jsa-15 | ILE_A_61_C | HIS_A_62_N | -3.884 | 0.255 | 9.215 | -2.817 | -0.089 | 9.92 | 1.32 | 57.84 | -17.87 |
| 363 | 2jsa-15 | HIS_A_62_N | HIS_A_62_H | -2.817 | -0.089 | 9.92 | -2.007 | -0.478 | 9.459 | 1.01 | 117.16 | -25.65 |
| 364 | 2jsa-15 | HIS_A_62_N | HIS_A_62_CA | -2.817 | -0.089 | 9.92 | -2.784 | 0.082 | 11.362 | 1.45 | 6.89 | 79.08 |
| 365 | 2jsa-15 | HIS_A_62_CA | HIS_A_62_HA | -2.784 | 0.082 | 11.362 | -3.736 | -0.264 | 11.763 | 1.09 | 68.40 | -160.03 |
| 366 | 2jsa-15 | HIS_A_62_CA | HIS_A_62_C | -2.784 | 0.082 | 11.362 | -2.644 | 1.568 | 11.7 | 1.53 | 77.24 | 84.62 |
| 367 | 2jsa-15 | HIS_A_62_C | HIS_A_62_O | -2.644 | 1.568 | 11.7 | -2.928 | 1.983 | 12.822 | 1.23 | 24.14 | 124.39 |
| 368 | 2jsa-15 | HIS_A_62_C | GLU_A_63_N | -2.644 | 1.568 | 11.7 | -2.205 | 2.327 | 10.707 | 1.32 | 138.56 | 59.96 |
| 369 | 2jsa-15 | GLU_A_63_N | GLU_A_63_H | -2.205 | 2.327 | 10.707 | -1.991 | 1.917 | 9.809 | 1.01 | 152.75 | -62.44 |
| 370 | 2jsa-15 | GLU_A_63_N | GLU_A_63_CA | -2.205 | 2.327 | 10.707 | -2.022 | 3.758 | 10.884 | 1.45 | 83.01 | 82.71 |
| 371 | 2jsa-15 | GLU_A_63_CA | GLU_A_63_HA | -2.022 | 3.758 | 10.884 | -1.66 | 3.929 | 11.898 | 1.09 | 21.55 | 25.28 |
| 372 | 2jsa-15 | GLU_A_63_CA | GLU_A_63_C | -2.022 | 3.758 | 10.884 | -3.357 | 4.488 | 10.72 | 1.53 | 96.15 | 151.33 |
| 373 | 2jsa-15 | GLU_A_63_C | GLU_A_63_O | -3.357 | 4.488 | 10.72 | -3.794 | 5.202 | 11.622 | 1.23 | 42.86 | 121.47 |
| 374 | 2jsa-15 | GLU_A_63_C | LEU_A_64_N | -3.357 | 4.488 | 10.72 | -3.969 | 4.283 | 9.563 | 1.32 | 150.85 | -161.48 |
| 375 | 2jsa-15 | LEU_A_64_N | LEU_A_64_H | -3.969 | 4.283 | 9.563 | -3.551 | 3.682 | 8.867 | 1.01 | 133.55 | -55.18 |
| 376 | 2jsa-15 | LEU_A_64_N | LEU_A_64_CA | -3.969 | 4.283 | 9.563 | -5.246 | 4.912 | 9.269 | 1.45 | 101.67 | 153.78 |
| 377 | 2jsa-15 | LEU_A_64_CA | LEU_A_64_HA | -5.246 | 4.912 | 9.269 | -5.139 | 5.985 | 9.432 | 1.09 | 81.40 | 84.31 |
| 378 | 2jsa-15 | LEU_A_64_CA | LEU_A_64_C | -5.246 | 4.912 | 9.269 | -6.301 | 4.396 | 10.249 | 1.53 | 50.16 | -153.94 |
| 379 | 2jsa-15 | LEU_A_64_C | LEU_A_64_O | -6.301 | 4.396 | 10.249 | -7.049 | 5.179 | 10.83 | 1.23 | 61.78 | 133.69 |
| 380 | 2jsa-15 | LEU_A_64_C | GLU_A_65_N | -6.301 | 4.396 | 10.249 | -6.327 | 3.079 | 10.402 | 1.33 | 83.37 | -91.13 |
| 381 | 2jsa-15 | GLU_A_65_N | GLU_A_65_H | -6.327 | 3.079 | 10.402 | -5.68 | 2.495 | 9.892 | 1.01 | 120.33 | -42.07 |
| 382 | 2jsa-15 | GLU_A_65_N | GLU_A_65_CA | -6.327 | 3.079 | 10.402 | -7.278 | 2.449 | 11.301 | 1.45 | 51.76 | -146.48 |
| 383 | 2jsa-15 | GLU_A_65_CA | GLU_A_65_HA | -7.278 | 2.449 | 11.301 | -8.281 | 2.739 | 10.988 | 1.09 | 106.69 | 163.87 |
| 384 | 2jsa-15 | GLU_A_65_CA | GLU_A_65_C | -7.278 | 2.449 | 11.301 | -7.061 | 2.941 | 12.734 | 1.53 | 20.57 | 66.20 |
| 385 | 2jsa-15 | GLU_A_65_C | GLU_A_65_O | -7.061 | 2.941 | 12.734 | -7.994 | 2.966 | 13.533 | 1.23 | 49.43 | 178.47 |
| 386 | 2jsa-15 | GLU_A_65_C | GLY_A_66_N | -7.061 | 2.941 | 12.734 | -5.822 | 3.318 | 13.013 | 1.32 | 77.84 | 16.92 |
| 387 | 2jsa-15 | GLY_A_66_N | GLY_A_66_H | -5.822 | 3.318 | 13.013 | -5.103 | 3.269 | 12.305 | 1.01 | 134.49 | -3.90 |
| 388 | 2jsa-15 | GLY_A_66_N | GLY_A_66_CA | -5.822 | 3.318 | 13.013 | -5.469 | 3.807 | 14.335 | 1.45 | 24.52 | 54.18 |
| 389 | 2jsa-15 | GLY_A_66_CA | GLY_A_66_C | -5.469 | 3.807 | 14.335 | -5.957 | 5.242 | 14.537 | 1.53 | 82.41 | 108.78 |
| 390 | 2jsa-15 | GLY_A_66_C | GLY_A_66_O | -5.957 | 5.242 | 14.537 | -6.474 | 5.583 | 15.6 | 1.23 | 30.23 | 146.59 |
| 391 | 2jsa-15 | GLY_A_66_C | GLY_A_67_N | -5.957 | 5.242 | 14.537 | -5.777 | 6.047 | 13.5 | 1.33 | 141.50 | 77.40 |
| 392 | 2jsa-15 | GLY_A_67_N | GLY_A_67_H | -5.777 | 6.047 | 13.5 | -5.345 | 5.699 | 12.656 | 1.01 | 146.68 | -38.85 |
| 393 | 2jsa-15 | GLY_A_67_N | GLY_A_67_CA | -5.777 | 6.047 | 13.5 | -6.192 | 7.438 | 13.551 | 1.45 | 87.99 | 106.61 |
| 394 | 2jsa-15 | GLY_A_67_CA | GLY_A_67_C | -6.192 | 7.438 | 13.551 | -5.375 | 8.29 | 12.577 | 1.53 | 129.53 | 46.20 |
| 395 | 2jsa-15 | GLY_A_67_C | GLY_A_67_O | -5.375 | 8.29 | 12.577 | -5.798 | 9.381 | 12.195 | 1.23 | 108.08 | 111.19 |
| 396 | 2jsa-15 | GLY_A_67_C | THR_A_68_N | -5.375 | 8.29 | 12.577 | -4.219 | 7.76 | 12.203 | 1.33 | 106.39 | -24.63 |
| 397 | 2jsa-15 | THR_A_68_N | THR_A_68_H | -4.219 | 7.76 | 12.203 | -3.934 | 6.858 | 12.557 | 1.01 | 69.48 | -72.47 |
| 398 | 2jsa-15 | THR_A_68_N | THR_A_68_CA | -4.219 | 7.76 | 12.203 | -3.34 | 8.458 | 11.281 | 1.45 | 129.40 | 38.45 |
| 399 | 2jsa-15 | THR_A_68_CA | THR_A_68_HA | -3.34 | 8.458 | 11.281 | -3.076 | 9.424 | 11.713 | 1.09 | 66.67 | 74.71 |
| 400 | 2jsa-15 | THR_A_68_CA | THR_A_68_C | -3.34 | 8.458 | 11.281 | -4.054 | 8.716 | 9.953 | 1.53 | 150.24 | 160.13 |
| 401 | 2jsa-15 | THR_A_68_C | THR_A_68_O | -4.054 | 8.716 | 9.953 | -5.101 | 8.128 | 9.683 | 1.23 | 102.67 | -150.68 |
| 402 | 2jsa-15 | THR_A_68_C | ALA_A_69_N | -4.054 | 8.716 | 9.953 | -3.46 | 9.595 | 9.159 | 1.33 | 126.81 | 55.95 |
| 403 | 2jsa-15 | ALA_A_69_N | ALA_A_69_H | -3.46 | 9.595 | 9.159 | -2.6 | 10.041 | 9.447 | 1.01 | 73.44 | 27.41 |
| 404 | 2jsa-15 | ALA_A_69_N | ALA_A_69_CA | -3.46 | 9.595 | 9.159 | -4.026 | 9.937 | 7.866 | 1.45 | 152.91 | 148.86 |
| 405 | 2jsa-15 | ALA_A_69_CA | ALA_A_69_HA | -4.026 | 9.937 | 7.866 | -5.081 | 9.666 | 7.863 | 1.09 | 90.16 | -165.59 |
| 406 | 2jsa-15 | ALA_A_69_CA | ALA_A_69_C | -4.026 | 9.937 | 7.866 | -3.317 | 9.132 | 6.774 | 1.53 | 135.51 | -48.63 |
| 407 | 2jsa-15 | ALA_A_69_C | ALA_A_69_O | -3.317 | 9.132 | 6.774 | -2.257 | 8.556 | 7.012 | 1.23 | 78.84 | -28.52 |
| 408 | 2jsa-15 | ALA_A_69_C | PRO_A_70_N | -3.317 | 9.132 | 6.774 | -3.946 | 9.12 | 5.568 | 1.36 | 152.45 | -178.91 |
| 409 | 2jsa-15 | PRO_A_70_N | PRO_A_70_CA | -3.946 | 9.12 | 5.568 | -3.386 | 8.398 | 4.439 | 1.45 | 141.02 | -52.20 |
| 410 | 2jsa-15 | PRO_A_70_CA | PRO_A_70_HA | -3.386 | 8.398 | 4.439 | -3.051 | 7.411 | 4.757 | 1.09 | 73.03 | -71.25 |
| 411 | 2jsa-15 | PRO_A_70_CA | PRO_A_70_C | -3.386 | 8.398 | 4.439 | -2.191 | 9.147 | 3.846 | 1.53 | 112.81 | 32.08 |
| 412 | 2jsa-15 | PRO_A_70_C | PRO_A_70_O | -2.191 | 9.147 | 3.846 | -1.613 | 8.712 | 2.851 | 1.23 | 143.98 | -36.96 |
| 413 | 2jsa-15 | PRO_A_70_C | LYS_A_71_N | -2.191 | 9.147 | 3.846 | -1.857 | 10.26 | 4.483 | 1.33 | 61.27 | 73.30 |
| 414 | 2jsa-15 | LYS_A_71_N | LYS_A_71_H | -1.857 | 10.26 | 4.483 | -2.377 | 10.558 | 5.296 | 1.01 | 36.40 | 150.18 |
| 415 | 2jsa-15 | LYS_A_71_N | LYS_A_71_CA | -1.857 | 10.26 | 4.483 | -0.741 | 11.073 | 4.03 | 1.45 | 108.16 | 36.07 |
| 416 | 2jsa-15 | LYS_A_71_CA | LYS_A_71_HA | -0.741 | 11.073 | 4.03 | -0.251 | 10.548 | 3.21 | 1.09 | 138.79 | -46.97 |
| 417 | 2jsa-15 | LYS_A_71_CA | LYS_A_71_C | -0.741 | 11.073 | 4.03 | 0.275 | 11.214 | 5.166 | 1.53 | 42.08 | 7.90 |
| 418 | 2jsa-15 | LYS_A_71_C | LYS_A_71_O | 0.275 | 11.214 | 5.166 | 1.249 | 11.955 | 5.043 | 1.23 | 95.74 | 37.26 |
| 419 | 2jsa-15 | LYS_A_71_C | ASP_A_72_N | 0.275 | 11.214 | 5.166 | 0.012 | 10.492 | 6.245 | 1.32 | 35.46 | -110.01 |
| 420 | 2jsa-15 | ASP_A_72_N | ASP_A_72_H | 0.012 | 10.492 | 6.245 | -0.809 | 9.905 | 6.277 | 1.01 | 88.18 | -144.44 |
| 421 | 2jsa-15 | ASP_A_72_N | ASP_A_72_CA | 0.012 | 10.492 | 6.245 | 0.891 | 10.528 | 7.401 | 1.45 | 37.27 | 2.35 |
| 422 | 2jsa-15 | ASP_A_72_CA | ASP_A_72_HA | 0.891 | 10.528 | 7.401 | 1.667 | 11.276 | 7.236 | 1.09 | 98.70 | 43.95 |
| 423 | 2jsa-15 | ASP_A_72_CA | ASP_A_72_C | 0.891 | 10.528 | 7.401 | 1.529 | 9.151 | 7.593 | 1.53 | 82.79 | -65.14 |
| 424 | 2jsa-15 | ASP_A_72_C | ASP_A_72_O | 1.529 | 9.151 | 7.593 | 2.508 | 9.013 | 8.325 | 1.23 | 53.48 | -8.02 |
| 425 | 2jsa-15 | ASP_A_72_C | VAL_A_73_N | 1.529 | 9.151 | 7.593 | 0.949 | 8.166 | 6.924 | 1.32 | 120.34 | -120.49 |

| | | | | | | | | | | | | |
|---|---|---|---|---|---|---|---|---|---|---|---|---|
| 426 | 2jsa-15 | VAL_A_73_N | VAL_A_73_H | 0.949 | 8.166 | 6.924 | 0.146 | 8.352 | 6.341 | 1.01 | 125.27 | 166.96 |
| 427 | 2jsa-15 | VAL_A_73_N | VAL_A_73_CA | 0.949 | 8.166 | 6.924 | 1.448 | 6.804 | 7.011 | 1.45 | 86.57 | -69.88 |
| 428 | 2jsa-15 | VAL_A_73_CA | VAL_A_73_HA | 1.448 | 6.804 | 7.011 | 1.598 | 6.559 | 8.063 | 1.09 | 15.27 | -58.52 |
| 429 | 2jsa-15 | VAL_A_73_CA | VAL_A_73_C | 1.448 | 6.804 | 7.011 | 2.803 | 6.716 | 6.306 | 1.53 | 117.44 | -3.72 |
| 430 | 2jsa-15 | VAL_A_73_C | VAL_A_73_O | 2.803 | 6.716 | 6.306 | 3.758 | 6.17 | 6.855 | 1.23 | 63.48 | -29.76 |
| 431 | 2jsa-15 | VAL_A_73_C | CYS_A_74_N | 2.803 | 6.716 | 6.306 | 2.842 | 7.262 | 5.099 | 1.33 | 155.61 | 85.91 |
| 432 | 2jsa-15 | CYS_A_74_N | CYS_A_74_H | 2.842 | 7.262 | 5.099 | 2.017 | 7.695 | 4.71 | 1.01 | 112.66 | 152.31 |
| 433 | 2jsa-15 | CYS_A_74_N | CYS_A_74_CA | 2.842 | 7.262 | 5.099 | 4.064 | 7.252 | 4.312 | 1.45 | 122.78 | -0.47 |
| 434 | 2jsa-15 | CYS_A_74_CA | CYS_A_74_HA | 4.064 | 7.252 | 4.312 | 4.617 | 6.342 | 4.549 | 1.09 | 77.45 | -58.71 |
| 435 | 2jsa-15 | CYS_A_74_CA | CYS_A_74_C | 4.064 | 7.252 | 4.312 | 4.911 | 8.456 | 4.725 | 1.53 | 74.33 | 54.87 |
| 436 | 2jsa-15 | CYS_A_74_C | CYS_A_74_O | 4.911 | 8.456 | 4.725 | 5.354 | 9.229 | 3.876 | 1.23 | 133.62 | 60.18 |
| 437 | 2jsa-15 | CYS_A_74_C | THR_A_75_N | 4.911 | 8.456 | 4.725 | 5.111 | 8.58 | 6.03 | 1.33 | 10.22 | 31.80 |
| 438 | 2jsa-15 | THR_A_75_N | THR_A_75_H | 5.111 | 8.58 | 6.03 | 4.717 | 7.907 | 6.672 | 1.01 | 50.54 | -120.35 |
| 439 | 2jsa-15 | THR_A_75_N | THR_A_75_CA | 5.111 | 8.58 | 6.03 | 5.897 | 9.678 | 6.565 | 1.45 | 68.39 | 54.40 |
| 440 | 2jsa-15 | THR_A_75_CA | THR_A_75_HA | 5.897 | 9.678 | 6.565 | 6.769 | 9.827 | 5.928 | 1.09 | 125.76 | 9.70 |
| 441 | 2jsa-15 | THR_A_75_CA | THR_A_75_C | 5.897 | 9.678 | 6.565 | 6.394 | 9.341 | 7.972 | 1.53 | 23.11 | -34.14 |
| 442 | 2jsa-15 | THR_A_75_C | THR_A_75_O | 6.394 | 9.341 | 7.972 | 7.6 | 9.288 | 8.213 | 1.23 | 78.71 | -2.52 |
| 443 | 2jsa-15 | THR_A_75_C | LYS_A_76_N | 6.394 | 9.341 | 7.972 | 5.44 | 9.122 | 8.865 | 1.32 | 47.62 | -167.07 |
| 444 | 2jsa-15 | LYS_A_76_N | LYS_A_76_H | 5.44 | 9.122 | 8.865 | 4.469 | 9.183 | 8.597 | 1.01 | 105.40 | 176.41 |
| 445 | 2jsa-15 | LYS_A_76_N | LYS_A_76_CA | 5.44 | 9.122 | 8.865 | 5.766 | 8.792 | 10.242 | 1.45 | 18.62 | -45.35 |
| 446 | 2jsa-15 | LYS_A_76_CA | LYS_A_76_HA | 5.766 | 8.792 | 10.242 | 6.495 | 9.52 | 10.6 | 1.09 | 70.84 | 44.96 |
| 447 | 2jsa-15 | LYS_A_76_CA | LYS_A_76_C | 5.766 | 8.792 | 10.242 | 6.425 | 7.411 | 10.29 | 1.53 | 88.20 | -64.49 |
| 448 | 2jsa-15 | LYS_A_76_C | LYS_A_76_O | 6.425 | 7.411 | 10.29 | 7.26 | 7.147 | 11.152 | 1.23 | 45.45 | -17.55 |
| 449 | 2jsa-15 | LYS_A_76_C | LEU_A_77_N | 6.425 | 7.411 | 10.29 | 6.025 | 6.567 | 9.349 | 1.33 | 135.21 | -115.36 |
| 450 | 2jsa-15 | LEU_A_77_N | LEU_A_77_H | 6.025 | 6.567 | 9.349 | 5.335 | 6.853 | 8.67 | 1.01 | 132.27 | 157.49 |
| 451 | 2jsa-15 | LEU_A_77_N | LEU_A_77_CA | 6.025 | 6.567 | 9.349 | 6.566 | 5.221 | 9.272 | 1.45 | 93.04 | -68.10 |
| 452 | 2jsa-15 | LEU_A_77_CA | LEU_A_77_HA | 6.566 | 5.221 | 9.272 | 6.864 | 4.919 | 10.276 | 1.09 | 22.91 | -45.38 |
| 453 | 2jsa-15 | LEU_A_77_CA | LEU_A_77_C | 6.566 | 5.221 | 9.272 | 7.822 | 5.231 | 8.399 | 1.53 | 124.80 | 0.46 |
| 454 | 2jsa-15 | LEU_A_77_C | LEU_A_77_O | 7.822 | 5.231 | 8.399 | 8.281 | 4.179 | 7.954 | 1.23 | 111.19 | -66.43 |
| 455 | 2jsa-15 | LEU_A_77_C | ASN_A_78_N | 7.822 | 5.231 | 8.399 | 8.342 | 6.428 | 8.176 | 1.32 | 99.70 | 66.52 |
| 456 | 2jsa-15 | ASN_A_78_N | ASN_A_78_H | 8.342 | 6.428 | 8.176 | 7.911 | 7.251 | 8.572 | 1.01 | 66.91 | 117.64 |
| 457 | 2jsa-15 | ASN_A_78_N | ASN_A_78_CA | 8.342 | 6.428 | 8.176 | 9.535 | 6.588 | 7.364 | 1.45 | 124.00 | 7.64 |
| 458 | 2jsa-15 | ASN_A_78_CA | ASN_A_78_HA | 9.535 | 6.588 | 7.364 | 9.708 | 7.65 | 7.189 | 1.09 | 99.24 | 80.75 |
| 459 | 2jsa-15 | ASN_A_78_CA | ASN_A_78_C | 9.535 | 6.588 | 7.364 | 9.339 | 5.866 | 6.03 | 1.53 | 150.72 | -105.19 |
| 460 | 2jsa-15 | ASN_A_78_C | ASN_A_78_O | 9.339 | 5.866 | 6.03 | 10.31 | 5.511 | 5.363 | 1.23 | 122.83 | -20.08 |
| 461 | 2jsa-15 | ASN_A_78_C | GLU_A_79_N | 9.339 | 5.866 | 6.03 | 8.076 | 5.669 | 5.68 | 1.33 | 105.31 | -171.13 |
| 462 | 2jsa-15 | GLU_A_79_N | GLU_A_79_H | 8.076 | 5.669 | 5.68 | 7.327 | 5.988 | 6.278 | 1.01 | 53.70 | 156.93 |
| 463 | 2jsa-15 | GLU_A_79_N | GLU_A_79_CA | 8.076 | 5.669 | 5.68 | 7.74 | 4.995 | 4.437 | 1.45 | 148.79 | -116.50 |
| 464 | 2jsa-15 | GLU_A_79_CA | GLU_A_79_HA | 7.74 | 4.995 | 4.437 | 8.546 | 4.299 | 4.202 | 1.09 | 102.44 | -40.81 |
| 465 | 2jsa-15 | GLU_A_79_CA | GLU_A_79_C | 7.74 | 4.995 | 4.437 | 7.632 | 6.009 | 3.296 | 1.53 | 138.21 | 96.08 |
| 466 | 2jsa-15 | GLU_A_79_C | GLU_A_79_O | 7.632 | 6.009 | 3.296 | 6.725 | 5.924 | 2.47 | 1.23 | 132.20 | -174.65 |
| 467 | 2jsa-15 | GLU_A_79_C | CYS_A_80_N | 7.632 | 6.009 | 3.296 | 8.57 | 6.944 | 3.288 | 1.32 | 90.35 | 44.91 |
| 468 | 2jsa-15 | CYS_A_80_N | CYS_A_80_H | 8.57 | 6.944 | 3.288 | 9.287 | 6.953 | 4.0 | 1.01 | 45.20 | 0.72 |
| 469 | 2jsa-15 | CYS_A_80_N | CYS_A_80_CA | 8.57 | 6.944 | 3.288 | 8.592 | 7.973 | 2.261 | 1.45 | 134.94 | 88.78 |
| 470 | 2jsa-15 | CYS_A_80_CA | CYS_A_80_HA | 8.592 | 7.973 | 2.261 | 8.753 | 7.492 | 1.296 | 1.09 | 152.27 | -71.49 |
| 471 | 2jsa-15 | CYS_A_80_CA | CYS_A_80_C | 8.592 | 7.973 | 2.261 | 9.772 | 8.906 | 2.542 | 1.53 | 79.42 | 38.33 |
| 472 | 2jsa-15 | CYS_A_80_C | CYS_A_80_O | 9.772 | 8.906 | 2.542 | 9.995 | 9.303 | 3.685 | 1.23 | 21.72 | 60.68 |
| 473 | 2jsa-15 | CYS_A_80_C | PRO_A_81_N | 9.772 | 8.906 | 2.542 | 10.513 | 9.239 | 1.452 | 1.36 | 143.30 | 24.20 |
| 474 | 2jsa-15 | PRO_A_81_N | PRO_A_81_CA | 10.513 | 9.239 | 1.452 | 11.664 | 10.119 | 1.57 | 1.45 | 85.34 | 37.40 |
| 475 | 2jsa-15 | PRO_A_81_CA | PRO_A_81_HA | 11.664 | 10.119 | 1.57 | 12.271 | 9.827 | 2.427 | 1.09 | 38.17 | -25.69 |
| 476 | 2jsa-15 | PRO_A_81_CA | PRO_A_81_C | 11.664 | 10.119 | 1.57 | 11.225 | 11.574 | 1.758 | 1.53 | 82.95 | 106.79 |
| 477 | 2jsa-15 | PRO_A_81_C | PRO_A_81_O | 11.225 | 11.574 | 1.758 | 11.249 | 12.093 | 2.873 | 1.23 | 24.98 | 87.35 |

Table 3: A reversible spherical geometric conversion of Caenopore-5 backbone structure coordinate matrix to three independent vectors of $\rho$, $\theta$ and $\phi$ [6,7,9].

#### 2.6 An ABN-SCS Approach for Protein Peptide Bond Structure Feature Extraction: A Case Study with Cp5

| Bond ID | Model ID | Atom_Initial | Atom_Final | $X_i$ | $Y_i$ | $Z_i$ | $X_f$ | $Y_f$ | $Z_f$ | ρ (Å) | θ (°) | φ (°) |
|---|---|---|---|---|---|---|---|---|---|---|---|---|
| 1 | 2jsa-1 | ARG_A_1_C | SER_A_2_N | 13.875 | 5.84 | -0.967 | 12.838 | 6.548 | -0.544 | 1.32 | 71.38 | 145.68 |
| 2 | 2jsa-1 | SER_A_2_C | ALA_A_3_N | 12.226 | 7.489 | -2.715 | 11.022 | 7.04 | -3.037 | 1.32 | 104.07 | -159.55 |
| 3 | 2jsa-1 | ALA_A_3_C | LEU_A_4_N | 9.201 | 6.358 | -4.5 | 8.901 | 5.473 | -5.439 | 1.32 | 135.14 | -108.73 |
| 4 | 2jsa-1 | LEU_A_4_C | SER_A_5_N | 7.055 | 4.298 | -4.374 | 8.012 | 3.981 | -3.514 | 1.33 | 49.53 | -18.33 |
| 5 | 2jsa-1 | SER_A_5_C | CYS_A_6_N | 6.482 | 3.925 | -1.609 | 6.528 | 5.245 | -1.501 | 1.33 | 85.33 | 88.00 |
| 6 | 2jsa-1 | CYS_A_6_C | GLN_A_7_N | 4.155 | 5.681 | -1.669 | 4.218 | 5.929 | -2.969 | 1.32 | 168.86 | 75.75 |
| 7 | 2jsa-1 | GLN_A_7_C | MET_A_8_N | 2.299 | 4.459 | -3.356 | 2.995 | 3.332 | -3.33 | 1.32 | 88.88 | -58.30 |
| 8 | 2jsa-1 | MET_A_8_C | CYS_A_9_N | 1.584 | 2.284 | -1.611 | 2.224 | 2.929 | -0.647 | 1.32 | 43.31 | 45.22 |
| 9 | 2jsa-1 | CYS_A_9_C | GLU_A_10_N | 0.277 | 3.938 | 0.372 | 0.291 | 4.739 | -0.684 | 1.33 | 142.81 | 89.00 |
| 10 | 2jsa-1 | GLU_A_10_C | LEU_A_11_N | -1.841 | 4.661 | -1.885 | -1.395 | 3.451 | -2.188 | 1.32 | 103.22 | -69.77 |
| 11 | 2jsa-1 | LEU_A_11_C | VAL_A_12_N | -3.025 | 1.65 | -2.036 | -2.379 | 1.215 | -0.965 | 1.32 | 36.02 | -33.96 |
| 12 | 2jsa-1 | VAL_A_12_C | VAL_A_13_N | -4.052 | 1.199 | 0.802 | -3.584 | 2.341 | 1.285 | 1.33 | 68.63 | 67.72 |
| 13 | 2jsa-1 | VAL_A_13_C | LYS_A_14_N | -5.599 | 3.701 | 1.179 | -5.246 | 4.324 | 0.064 | 1.33 | 147.29 | 60.46 |
| 14 | 2jsa-1 | LYS_A_14_C | LYS_A_15_N | -7.323 | 3.755 | -1.069 | -6.89 | 2.505 | -1.009 | 1.32 | 87.40 | -70.89 |
| 15 | 2jsa-1 | LYS_A_15_C | TYR_A_16_N | -8.512 | 1.1 | 0.138 | -7.769 | 1.272 | 1.221 | 1.32 | 35.15 | 13.03 |
| 16 | 2jsa-1 | TYR_A_16_C | GLU_A_17_N | -9.301 | 2.135 | 2.94 | -8.762 | 3.332 | 3.124 | 1.33 | 82.02 | 65.76 |
| 17 | 2jsa-1 | GLU_A_17_C | GLY_A_18_N | -10.75 | 4.629 | 2.524 | -10.513 | 4.202 | 1.292 | 1.33 | 158.38 | -60.97 |
| 18 | 2jsa-1 | GLY_A_18_C | SER_A_19_N | -11.817 | 2.915 | -0.347 | -12.397 | 2.05 | 0.472 | 1.32 | 51.82 | -123.84 |
| 19 | 2jsa-1 | SER_A_19_C | ALA_A_20_N | -14.232 | 0.591 | -0.222 | -14.936 | 0.126 | 0.799 | 1.32 | 39.57 | -146.55 |
| 20 | 2jsa-1 | ALA_A_20_C | ASP_A_21_N | -16.849 | -1.012 | 1.781 | -16.008 | -2.0 | 2.049 | 1.32 | 78.33 | -49.60 |
| 21 | 2jsa-1 | ASP_A_21_C | LYS_A_22_N | -15.378 | -2.846 | 4.242 | -14.303 | -2.108 | 4.009 | 1.32 | 100.13 | 34.47 |
| 22 | 2jsa-1 | LYS_A_22_C | ASP_A_23_N | -12.574 | -3.179 | 5.345 | -12.424 | -3.986 | 4.305 | 1.32 | 141.72 | -79.47 |
| 23 | 2jsa-1 | ASP_A_23_C | ALA_A_24_N | -10.258 | -5.077 | 4.099 | -9.423 | -5.274 | 5.108 | 1.32 | 40.37 | -13.27 |
| 24 | 2jsa-1 | ALA_A_24_C | ASN_A_25_N | -7.485 | -6.235 | 3.993 | -8.369 | -7.183 | 3.718 | 1.33 | 101.98 | -133.00 |
| 25 | 2jsa-1 | ASN_A_25_C | VAL_A_26_N | -8.27 | -7.88 | 1.388 | -9.483 | -7.415 | 1.131 | 1.32 | 101.19 | 159.03 |
| 26 | 2jsa-1 | VAL_A_26_C | ILE_A_27_N | -9.229 | -5.618 | -0.49 | -8.604 | -5.064 | 0.539 | 1.33 | 39.06 | 41.55 |
| 27 | 2jsa-1 | ILE_A_27_C | LYS_A_28_N | -6.549 | -3.949 | -0.136 | -5.8 | -4.82 | 0.525 | 1.33 | 60.08 | -49.31 |
| 28 | 2jsa-1 | LYS_A_28_C | LYS_A_29_N | -4.442 | -5.715 | -1.284 | -5.567 | -6.33 | -1.618 | 1.32 | 104.60 | -151.34 |
| 29 | 2jsa-1 | LYS_A_29_C | ASP_A_30_N | -5.841 | -5.887 | -3.995 | -6.722 | -4.933 | -3.733 | 1.32 | 78.59 | 132.72 |
| 30 | 2jsa-1 | ASP_A_30_C | PHE_A_31_N | -5.601 | -3.237 | -5.069 | -4.735 | -3.088 | -4.077 | 1.33 | 41.53 | 9.76 |
| 31 | 2jsa-1 | PHE_A_31_C | ASP_A_32_N | -2.675 | -3.299 | -5.403 | -2.437 | -4.572 | -5.121 | 1.33 | 77.72 | -79.41 |
| 32 | 2jsa-1 | ASP_A_32_C | ALA_A_33_N | -2.482 | -5.415 | -7.403 | -3.751 | -5.038 | -7.337 | 1.33 | 87.15 | 163.45 |
| 33 | 2jsa-1 | ALA_A_33_C | GLU_A_34_N | -3.907 | -4.071 | -9.564 | -3.938 | -2.778 | -9.278 | 1.32 | 77.53 | 91.37 |
| 34 | 2jsa-1 | GLU_A_34_C | CYS_A_35_N | -1.814 | -1.866 | -10.085 | -1.329 | -2.056 | -8.867 | 1.32 | 23.15 | -21.39 |
| 35 | 2jsa-1 | CYS_A_35_C | LYS_A_36_N | 0.704 | -3.081 | -9.682 | -0.141 | -3.967 | -10.188 | 1.32 | 112.45 | -133.64 |
| 36 | 2jsa-1 | LYS_A_36_C | LYS_A_37_N | 0.202 | -4.282 | -12.577 | -1.013 | -3.891 | -12.935 | 1.33 | 105.67 | 162.16 |
| 37 | 2jsa-1 | LYS_A_37_C | LEU_A_38_N | -0.537 | -1.909 | -14.265 | -0.983 | -1.017 | -13.393 | 1.32 | 48.83 | 116.57 |
| 38 | 2jsa-1 | LEU_A_38_C | PHE_A_39_N | 1.117 | 0.21 | -13.48 | 1.695 | -0.732 | -12.75 | 1.32 | 56.55 | -58.47 |
| 39 | 2jsa-1 | PHE_A_39_C | HIS_A_40_N | 3.462 | -2.336 | -13.343 | 2.835 | -2.665 | -14.463 | 1.33 | 147.70 | -152.31 |
| 40 | 2jsa-1 | HIS_A_40_C | THR_A_41_N | 4.39 | -3.95 | -15.832 | 4.835 | -2.749 | -16.172 | 1.33 | 104.87 | 69.67 |
| 41 | 2jsa-1 | THR_A_41_C | ILE_A_42_N | 7.143 | -1.953 | -15.97 | 7.172 | -2.427 | -14.733 | 1.33 | 21.00 | -86.50 |
| 42 | 2jsa-1 | ILE_A_42_C | PRO_A_43_N | 9.052 | -3.044 | -13.317 | 10.341 | -2.671 | -13.096 | 1.36 | 80.65 | 16.14 |
| 43 | 2jsa-1 | PRO_A_43_C | PHE_A_44_N | 11.156 | -4.007 | -11.2 | 10.599 | -5.189 | -10.982 | 1.32 | 80.53 | -115.23 |
| 44 | 2jsa-1 | PHE_A_44_C | GLY_A_45_N | 9.153 | -4.975 | -9.005 | 8.552 | -4.087 | -9.784 | 1.33 | 126.00 | 124.09 |
| 45 | 2jsa-1 | GLY_A_45_C | THR_A_46_N | 6.3 | -4.305 | -8.821 | 5.774 | -5.089 | -9.75 | 1.32 | 134.54 | -123.86 |
| 46 | 2jsa-1 | THR_A_46_C | ARG_A_47_N | 5.218 | -7.016 | -8.342 | 6.518 | -7.27 | -8.365 | 1.32 | 90.99 | -11.06 |
| 47 | 2jsa-1 | ARG_A_47_C | GLU_A_48_N | 6.99 | -7.606 | -5.986 | 7.502 | -6.396 | -5.82 | 1.32 | 82.80 | 67.06 |
| 48 | 2jsa-1 | GLU_A_48_C | CYS_A_49_N | 5.998 | -5.451 | -4.135 | 5.17 | -5.255 | -5.15 | 1.32 | 140.03 | 166.68 |
| 49 | 2jsa-1 | CYS_A_49_C | ASP_A_50_N | 3.218 | -6.02 | -3.944 | 3.151 | -7.255 | -4.419 | 1.32 | 111.01 | -93.11 |
| 50 | 2jsa-1 | ASP_A_50_C | HIS_A_51_N | 3.474 | -8.423 | -2.309 | 4.698 | -7.922 | -2.393 | 1.33 | 93.63 | 22.26 |
| 51 | 2jsa-1 | HIS_A_51_C | TYR_A_52_N | 5.294 | -6.713 | -0.361 | 4.268 | -5.972 | -0.752 | 1.32 | 107.17 | 144.16 |
| 52 | 2jsa-1 | TYR_A_52_C | VAL_A_53_N | 2.505 | -4.985 | 0.649 | 1.58 | -5.52 | -0.134 | 1.32 | 126.23 | -149.96 |
| 53 | 2jsa-1 | VAL_A_53_C | ASN_A_54_N | 0.301 | -6.666 | 1.59 | 1.445 | -7.317 | 1.747 | 1.33 | 83.20 | -29.64 |
| 54 | 2jsa-1 | ASN_A_54_C | SER_A_55_N | 3.041 | -7.968 | 3.465 | 3.193 | -6.805 | 4.081 | 1.32 | 62.29 | 82.55 |
| 55 | 2jsa-1 | SER_A_55_C | LYS_A_56_N | 4.416 | -4.987 | 5.164 | 3.7 | -4.171 | 4.404 | 1.33 | 125.00 | 131.27 |
| 56 | 2jsa-1 | LYS_A_56_C | VAL_A_57_N | 2.087 | -2.381 | 4.741 | 1.245 | -3.403 | 4.699 | 1.32 | 91.82 | -129.48 |
| 57 | 2jsa-1 | VAL_A_57_C | ASP_A_58_N | -0.728 | -3.355 | 6.123 | -0.263 | -4.409 | 6.778 | 1.33 | 60.38 | -66.19 |
| 58 | 2jsa-1 | ASP_A_58_C | PRO_A_59_N | -0.444 | -3.455 | 9.009 | 0.754 | -2.841 | 8.815 | 1.36 | 98.20 | 27.14 |
| 59 | 2jsa-1 | PRO_A_59_C | ILE_A_60_N | 0.365 | -0.425 | 9.062 | 0.518 | -0.171 | 7.77 | 1.33 | 167.07 | 58.94 |
| 60 | 2jsa-1 | ILE_A_60_C | ILE_A_61_N | -1.604 | 1.007 | 7.59 | -2.289 | -0.104 | 7.36 | 1.33 | 99.99 | -121.66 |
| 61 | 2jsa-1 | ILE_A_61_C | HIS_A_62_N | -3.828 | -0.063 | 9.244 | -2.727 | -0.321 | 9.933 | 1.32 | 58.65 | -13.19 |
| 62 | 2jsa-1 | HIS_A_62_C | GLU_A_63_N | -2.935 | 1.23 | 11.802 | -2.237 | 2.124 | 11.117 | 1.33 | 121.13 | 52.02 |
| 63 | 2jsa-1 | GLU_A_63_C | LEU_A_64_N | -3.791 | 4.014 | 11.205 | -4.263 | 3.861 | 9.976 | 1.33 | 158.01 | -162.04 |
| 64 | 2jsa-1 | LEU_A_64_C | GLU_A_65_N | -6.606 | 3.506 | 10.527 | -6.11 | 2.437 | 11.132 | 1.32 | 62.82 | -65.11 |
| 65 | 2jsa-1 | GLU_A_65_C | GLY_A_66_N | -7.161 | 2.334 | 13.341 | -6.161 | 3.109 | 13.738 | 1.33 | 72.58 | 37.78 |
| 66 | 2jsa-1 | GLY_A_66_C | GLY_A_67_N | -6.909 | 5.206 | 14.781 | -6.627 | 5.802 | 13.632 | 1.32 | 150.15 | 64.68 |
| 67 | 2jsa-1 | GLY_A_67_C | THR_A_68_N | -6.14 | 8.017 | 12.683 | -5.35 | 7.441 | 11.788 | 1.33 | 132.47 | -36.10 |
| 68 | 2jsa-1 | THR_A_68_C | ALA_A_69_N | -4.673 | 8.439 | 9.656 | -4.196 | 9.561 | 9.137 | 1.33 | 113.06 | 66.97 |
| 69 | 2jsa-1 | ALA_A_69_C | PRO_A_70_N | -3.498 | 9.178 | 6.838 | -3.871 | 9.124 | 5.531 | 1.36 | 163.91 | -171.76 |
| 70 | 2jsa-1 | PRO_A_70_C | LYS_A_71_N | -1.813 | 9.285 | 4.195 | -1.778 | 10.488 | 4.748 | 1.32 | 65.32 | 88.33 |
| 71 | 2jsa-1 | LYS_A_71_C | ASP_A_72_N | 0.179 | 11.509 | 5.773 | -0.236 | 10.771 | 6.792 | 1.32 | 39.72 | -119.35 |
| 72 | 2jsa-1 | ASP_A_72_C | VAL_A_73_N | 1.211 | 9.455 | 8.239 | 0.694 | 8.432 | 7.575 | 1.32 | 120.08 | -116.81 |
| 73 | 2jsa-1 | VAL_A_73_C | CYS_A_74_N | 2.633 | 7.119 | 6.913 | 2.577 | 7.533 | 5.656 | 1.32 | 161.62 | 97.70 |
| 74 | 2jsa-1 | CYS_A_74_C | THR_A_75_N | 4.65 | 8.74 | 5.343 | 5.17 | 8.554 | 6.547 | 1.32 | 24.64 | -19.68 |
| 75 | 2jsa-1 | THR_A_75_C | LYS_A_76_N | 6.512 | 9.086 | 8.527 | 5.56 | 8.916 | 9.433 | 1.33 | 46.87 | -169.88 |
| 76 | 2jsa-1 | LYS_A_76_C | LEU_A_77_N | 6.209 | 6.975 | 10.749 | 5.66 | 6.299 | 9.75 | 1.33 | 138.92 | -129.08 |
| 77 | 2jsa-1 | LEU_A_77_C | ASN_A_78_N | 7.224 | 4.646 | 8.888 | 7.981 | 5.726 | 8.761 | 1.32 | 95.50 | 54.97 |
| 78 | 2jsa-1 | ASN_A_78_C | GLU_A_79_N | 9.189 | 4.72 | 6.904 | 8.019 | 4.713 | 6.283 | 1.32 | 117.96 | -179.66 |
| 79 | 2jsa-1 | GLU_A_79_C | CYS_A_80_N | 8.022 | 4.661 | 3.835 | 7.544 | 5.897 | 3.846 | 1.33 | 89.52 | 111.14 |
| 80 | 2jsa-1 | CYS_A_80_C | PRO_A_81_N | 9.16 | 6.755 | 2.264 | 10.051 | 6.958 | 3.27 | 1.36 | 42.25 | 12.83 |
| 1 | 2jsa-2 | ARG_A_1_C | SER_A_2_N | 13.384 | 4.161 | -0.108 | 14.105 | 5.075 | -0.74 | 1.32 | 118.50 | 51.73 |
| 2 | 2jsa-2 | SER_A_2_C | ALA_A_3_N | 13.216 | 5.968 | -2.835 | 11.964 | 5.663 | -3.139 | 1.32 | 103.27 | -166.31 |
| 3 | 2jsa-2 | ALA_A_3_C | LEU_A_4_N | 10.047 | 5.404 | -4.616 | 9.535 | 4.549 | -5.489 | 1.32 | 131.22 | -120.91 |
| 4 | 2jsa-2 | LEU_A_4_C | SER_A_5_N | 7.467 | 3.713 | -4.516 | 8.326 | 3.26 | -3.614 | 1.33 | 47.11 | -27.81 |
| 5 | 2jsa-2 | SER_A_5_C | CYS_A_6_N | 6.827 | 3.364 | -1.685 | 7.101 | 4.657 | -1.597 | 1.32 | 86.19 | 78.04 |
| 6 | 2jsa-2 | CYS_A_6_C | GLN_A_7_N | 4.833 | 5.495 | -1.585 | 4.85 | 5.65 | -2.9 | 1.32 | 173.24 | 83.74 |
| 7 | 2jsa-2 | GLN_A_7_C | MET_A_8_N | 2.773 | 4.404 | -3.256 | 3.415 | 3.245 | -3.231 | 1.33 | 88.92 | -61.02 |
| 8 | 2jsa-2 | MET_A_8_C | CYS_A_9_N | 1.919 | 2.22 | -1.57 | 2.539 | 2.888 | -0.61 | 1.32 | 43.51 | 47.13 |
| 9 | 2jsa-2 | CYS_A_9_C | GLU_A_10_N | 0.601 | 3.937 | 0.387 | 0.639 | 4.717 | -0.685 | 1.33 | 143.93 | 87.21 |
| 10 | 2jsa-2 | GLU_A_10_C | LEU_A_11_N | -1.506 | 4.676 | -1.865 | -1.082 | 3.461 | -2.181 | 1.33 | 103.80 | -70.76 |
| 11 | 2jsa-2 | LEU_A_11_C | VAL_A_12_N | -2.775 | 1.726 | -1.971 | -2.117 | 1.216 | -0.94 | 1.33 | 38.92 | -37.78 |
| 12 | 2jsa-2 | VAL_A_12_C | VAL_A_13_N | -3.825 | 1.278 | 0.793 | -3.344 | 2.401 | 1.307 | 1.33 | 67.18 | 66.81 |
| 13 | 2jsa-2 | VAL_A_13_C | LYS_A_14_N | -5.345 | 3.773 | 1.114 | -4.96 | 4.374 | -0.002 | 1.32 | 147.40 | 57.36 |
| 14 | 2jsa-2 | LYS_A_14_C | LYS_A_15_N | -6.998 | 3.779 | -1.192 | -6.569 | 2.533 | -1.058 | 1.32 | 84.19 | -71.00 |
| 15 | 2jsa-2 | LYS_A_15_C | TYR_A_16_N | -8.283 | 1.194 | 0.033 | -7.617 | 1.393 | 1.162 | 1.33 | 31.62 | 16.64 |

| | | | | | | | | | | | | |
|---|---|---|---|---|---|---|---|---|---|---|---|---|
| 16 | 2jsa-2 | TYR_A_16_C | GLU_A_17_N | -9.163 | 2.421 | 2.773 | -8.524 | 3.564 | 2.979 | 1.33 | 81.06 | 60.79 |
| 17 | 2jsa-2 | GLU_A_17_C | GLY_A_18_N | -10.429 | 4.96 | 2.33 | -10.199 | 4.567 | 1.086 | 1.32 | 159.90 | -59.66 |
| 18 | 2jsa-2 | GLY_A_18_C | SER_A_19_N | -11.643 | 3.298 | -0.446 | -11.906 | 2.407 | 0.499 | 1.33 | 44.51 | -106.45 |
| 19 | 2jsa-2 | SER_A_19_C | ALA_A_20_N | -13.836 | 0.929 | 0.247 | -14.367 | 1.331 | 1.392 | 1.32 | 30.19 | 142.87 |
| 20 | 2jsa-2 | ALA_A_20_C | ASP_A_21_N | -16.186 | -0.148 | 2.044 | -15.194 | -1.027 | 2.025 | 1.33 | 90.82 | -41.54 |
| 21 | 2jsa-2 | ASP_A_21_C | LYS_A_22_N | -14.792 | -2.661 | 3.783 | -13.949 | -1.725 | 4.195 | 1.33 | 71.89 | 47.99 |
| 22 | 2jsa-2 | LYS_A_22_C | ASP_A_23_N | -12.705 | -3.238 | 5.638 | -12.088 | -3.713 | 4.565 | 1.33 | 144.03 | -37.59 |
| 23 | 2jsa-2 | ASP_A_23_C | ALA_A_24_N | -10.02 | -4.929 | 4.16 | -9.151 | -5.303 | 5.086 | 1.32 | 45.61 | -23.29 |
| 24 | 2jsa-2 | ALA_A_24_C | ASN_A_25_N | -7.339 | -6.369 | 3.86 | -8.322 | -7.201 | 3.55 | 1.32 | 103.53 | -139.76 |
| 25 | 2jsa-2 | ASN_A_25_C | VAL_A_26_N | -8.346 | -7.834 | 1.201 | -9.498 | -7.21 | 1.004 | 1.32 | 98.55 | 151.56 |
| 26 | 2jsa-2 | VAL_A_26_C | ILE_A_27_N | -9.121 | -5.391 | -0.569 | -8.48 | -4.893 | 0.479 | 1.33 | 37.76 | 37.84 |
| 27 | 2jsa-2 | ILE_A_27_C | LYS_A_28_N | -6.332 | -3.946 | -0.158 | -5.556 | -4.647 | 0.656 | 1.33 | 52.10 | -42.09 |
| 28 | 2jsa-2 | LYS_A_28_C | LYS_A_29_N | -4.218 | -5.679 | -1.095 | -5.276 | -6.439 | -1.328 | 1.32 | 100.14 | -144.31 |
| 29 | 2jsa-2 | LYS_A_29_C | ASP_A_30_N | -5.611 | -6.129 | -3.719 | -6.432 | -5.127 | -3.443 | 1.32 | 77.97 | 129.33 |
| 30 | 2jsa-2 | ASP_A_30_C | PHE_A_31_N | -5.378 | -3.466 | -4.875 | -4.561 | -3.112 | -3.894 | 1.32 | 42.23 | 23.43 |
| 31 | 2jsa-2 | PHE_A_31_C | ASP_A_32_N | -2.542 | -3.265 | -5.289 | -2.409 | -4.567 | -5.087 | 1.32 | 81.23 | -84.17 |
| 32 | 2jsa-2 | ASP_A_32_C | ALA_A_33_N | -2.473 | -5.321 | -7.399 | -3.746 | -4.959 | -7.316 | 1.33 | 86.41 | 164.13 |
| 33 | 2jsa-2 | ALA_A_33_C | GLU_A_34_N | -4.141 | -3.582 | -9.283 | -4.432 | -2.432 | -8.694 | 1.32 | 63.59 | 104.20 |
| 34 | 2jsa-2 | GLU_A_34_C | CYS_A_35_N | -2.599 | -1.133 | -9.668 | -1.824 | -1.878 | -8.893 | 1.33 | 54.21 | -43.87 |
| 35 | 2jsa-2 | CYS_A_35_C | LYS_A_36_N | -0.133 | -2.784 | -10.365 | -0.817 | -3.916 | -10.43 | 1.32 | 92.81 | -121.14 |
| 36 | 2jsa-2 | LYS_A_36_C | LYS_A_37_N | -0.906 | -4.033 | -12.86 | -1.569 | -2.895 | -12.716 | 1.32 | 83.76 | 120.23 |
| 37 | 2jsa-2 | LYS_A_37_C | LEU_A_38_N | -0.968 | -0.809 | -13.815 | -0.858 | -0.241 | -12.622 | 1.33 | 25.87 | 79.04 |
| 38 | 2jsa-2 | LEU_A_38_C | PHE_A_39_N | 1.425 | 0.6 | -12.704 | 1.962 | -0.269 | -11.86 | 1.33 | 50.44 | -58.29 |
| 39 | 2jsa-2 | PHE_A_39_C | HIS_A_40_N | 3.461 | -1.903 | -12.923 | 2.574 | -1.944 | -13.907 | 1.33 | 137.94 | -177.35 |
| 40 | 2jsa-2 | HIS_A_40_C | THR_A_41_N | 3.825 | -2.979 | -15.725 | 4.399 | -1.787 | -15.794 | 1.32 | 92.99 | 64.29 |
| 41 | 2jsa-2 | THR_A_41_C | ILE_A_42_N | 6.769 | -1.178 | -15.672 | 7.101 | -2.074 | -14.755 | 1.32 | 46.18 | -69.67 |
| 42 | 2jsa-2 | ILE_A_42_C | PRO_A_43_N | 8.9 | -3.143 | -13.511 | 10.247 | -3.046 | -13.352 | 1.36 | 83.29 | 4.12 |
| 43 | 2jsa-2 | PRO_A_43_C | PHE_A_44_N | 10.874 | -4.646 | -11.593 | 10.151 | -5.741 | -11.414 | 1.32 | 82.23 | -123.44 |
| 44 | 2jsa-2 | PHE_A_44_C | GLY_A_45_N | 8.825 | -5.445 | -9.365 | 8.364 | -4.406 | -10.047 | 1.33 | 120.96 | 113.93 |
| 45 | 2jsa-2 | GLY_A_45_C | THR_A_46_N | 6.149 | -4.356 | -8.981 | 5.733 | -5.311 | -9.8 | 1.33 | 128.18 | -113.54 |
| 46 | 2jsa-2 | THR_A_46_C | ARG_A_47_N | 5.007 | -7.125 | -8.322 | 6.277 | -7.502 | -8.329 | 1.32 | 90.30 | -16.53 |
| 47 | 2jsa-2 | ARG_A_47_C | GLU_A_48_N | 6.748 | -7.756 | -5.939 | 7.476 | -6.655 | -5.821 | 1.33 | 84.89 | 56.53 |
| 48 | 2jsa-2 | GLU_A_48_C | CYS_A_49_N | 6.115 | -5.603 | -4.079 | 5.211 | -5.463 | -5.037 | 1.32 | 136.32 | 171.20 |
| 49 | 2jsa-2 | CYS_A_49_C | ASP_A_50_N | 3.261 | -6.3 | -3.877 | 3.395 | -7.506 | -4.409 | 1.32 | 113.67 | -83.66 |
| 50 | 2jsa-2 | ASP_A_50_C | HIS_A_51_N | 3.674 | -8.863 | -2.409 | 4.756 | -8.109 | -2.289 | 1.32 | 84.80 | 34.87 |
| 51 | 2jsa-2 | HIS_A_51_C | TYR_A_52_N | 5.338 | -6.934 | -0.234 | 4.411 | -6.106 | -0.694 | 1.33 | 110.31 | 138.23 |
| 52 | 2jsa-2 | TYR_A_52_C | VAL_A_53_N | 2.698 | -5.008 | 0.687 | 1.755 | -5.547 | -0.072 | 1.33 | 124.95 | -150.25 |
| 53 | 2jsa-2 | VAL_A_53_C | ASN_A_54_N | 0.441 | -6.554 | 1.711 | 1.555 | -7.246 | 1.897 | 1.32 | 81.93 | -31.85 |
| 54 | 2jsa-2 | ASN_A_54_C | SER_A_55_N | 3.116 | -7.825 | 3.672 | 3.381 | -6.556 | 3.948 | 1.33 | 77.98 | 78.20 |
| 55 | 2jsa-2 | SER_A_55_C | LYS_A_56_N | 4.565 | -4.749 | 5.088 | 4.066 | -3.845 | 4.258 | 1.32 | 128.79 | 118.90 |
| 56 | 2jsa-2 | LYS_A_56_C | VAL_A_57_N | 2.433 | -2.104 | 4.73 | 1.608 | -3.137 | 4.646 | 1.32 | 93.64 | -128.61 |
| 57 | 2jsa-2 | VAL_A_57_C | ASP_A_58_N | -0.322 | -3.17 | 6.129 | 0.092 | -4.293 | 6.696 | 1.32 | 64.65 | -69.76 |
| 58 | 2jsa-2 | ASP_A_58_C | PRO_A_59_N | -0.075 | -3.419 | 8.961 | 1.106 | -2.771 | 8.777 | 1.36 | 97.78 | 28.75 |
| 59 | 2jsa-2 | PRO_A_59_C | ILE_A_60_N | 0.663 | -0.374 | 9.102 | 0.756 | -0.108 | 7.807 | 1.33 | 167.72 | 70.73 |
| 60 | 2jsa-2 | ILE_A_60_C | ILE_A_61_N | -1.395 | 1.027 | 7.697 | -2.079 | -0.068 | 7.397 | 1.33 | 103.08 | -121.99 |
| 61 | 2jsa-2 | ILE_A_61_C | HIS_A_62_N | -3.592 | -0.088 | 9.303 | -2.476 | -0.338 | 9.973 | 1.33 | 59.64 | -12.63 |
| 62 | 2jsa-2 | HIS_A_62_C | GLU_A_63_N | -2.73 | 1.172 | 11.87 | -2.265 | 2.113 | 11.061 | 1.33 | 127.62 | 63.70 |
| 63 | 2jsa-2 | GLU_A_63_C | LEU_A_64_N | -3.881 | 3.951 | 10.972 | -4.204 | 3.736 | 9.705 | 1.33 | 162.97 | -146.35 |
| 64 | 2jsa-2 | LEU_A_64_C | GLU_A_65_N | -6.586 | 3.636 | 10.189 | -6.396 | 2.427 | 10.697 | 1.33 | 67.46 | -81.07 |
| 65 | 2jsa-2 | GLU_A_65_C | GLY_A_66_N | -7.366 | 2.706 | 12.928 | -6.187 | 3.16 | 13.33 | 1.33 | 72.35 | 21.06 |
| 66 | 2jsa-2 | GLY_A_66_C | GLY_A_67_N | -6.851 | 5.283 | 14.378 | -6.818 | 5.821 | 13.168 | 1.32 | 155.99 | 86.49 |
| 67 | 2jsa-2 | GLY_A_67_C | THR_A_68_N | -6.668 | 7.979 | 11.997 | -5.421 | 7.573 | 11.805 | 1.33 | 98.33 | -18.03 |
| 68 | 2jsa-2 | THR_A_68_C | ALA_A_69_N | -5.03 | 8.459 | 9.555 | -4.487 | 9.429 | 8.834 | 1.32 | 122.97 | 60.76 |
| 69 | 2jsa-2 | ALA_A_69_C | PRO_A_70_N | -4.131 | 8.695 | 6.541 | -4.674 | 8.528 | 5.305 | 1.36 | 155.32 | -162.90 |
| 70 | 2jsa-2 | PRO_A_70_C | LYS_A_71_N | -2.802 | 8.28 | 3.73 | -2.588 | 9.538 | 4.085 | 1.32 | 74.45 | 80.35 |
| 71 | 2jsa-2 | LYS_A_71_C | ASP_A_72_N | -0.538 | 10.659 | 4.77 | -0.935 | 10.2 | 5.949 | 1.33 | 27.24 | -130.86 |
| 72 | 2jsa-2 | ASP_A_72_C | VAL_A_73_N | 0.441 | 9.199 | 7.688 | -0.241 | 8.096 | 7.417 | 1.32 | 101.80 | -121.73 |
| 73 | 2jsa-2 | VAL_A_73_C | CYS_A_74_N | 1.728 | 6.672 | 7.559 | 2.02 | 6.095 | 6.403 | 1.32 | 150.78 | -63.16 |
| 74 | 2jsa-2 | CYS_A_74_C | THR_A_75_N | 4.212 | 7.116 | 6.466 | 3.662 | 8.296 | 6.216 | 1.33 | 100.87 | 114.99 |
| 75 | 2jsa-2 | THR_A_75_C | LYS_A_76_N | 5.002 | 9.353 | 7.973 | 4.177 | 9.259 | 9.005 | 1.32 | 38.82 | -173.50 |
| 76 | 2jsa-2 | LYS_A_76_C | LEU_A_77_N | 5.336 | 7.712 | 10.483 | 4.87 | 6.793 | 9.649 | 1.33 | 128.99 | -116.89 |
| 77 | 2jsa-2 | LEU_A_77_C | ASN_A_78_N | 6.722 | 5.407 | 8.896 | 7.213 | 6.596 | 8.577 | 1.33 | 103.93 | 67.56 |
| 78 | 2jsa-2 | ASN_A_78_C | GLU_A_79_N | 8.515 | 5.664 | 6.743 | 7.334 | 5.303 | 6.263 | 1.32 | 111.24 | -163.00 |
| 79 | 2jsa-2 | GLU_A_79_C | CYS_A_80_N | 6.85 | 4.985 | 3.884 | 7.775 | 5.787 | 3.379 | 1.32 | 112.42 | 40.93 |
| 80 | 2jsa-2 | CYS_A_80_C | PRO_A_81_N | 8.67 | 7.523 | 1.952 | 8.824 | 8.387 | 2.992 | 1.36 | 40.16 | 79.89 |
| 1 | 2jsa-3 | ARG_A_1_C | SER_A_2_N | 13.816 | 2.625 | -2.435 | 12.949 | 3.538 | -2.847 | 1.32 | 108.12 | 133.52 |
| 2 | 2jsa-3 | SER_A_2_C | ALA_A_3_N | 11.925 | 5.751 | -3.011 | 10.736 | 5.187 | -3.172 | 1.33 | 96.98 | -154.62 |
| 3 | 2jsa-3 | ALA_A_3_C | LEU_A_4_N | 8.433 | 4.981 | -3.93 | 8.599 | 3.9 | -4.678 | 1.32 | 124.37 | -81.27 |
| 4 | 2jsa-3 | LEU_A_4_C | SER_A_5_N | 6.904 | 2.615 | -3.496 | 7.745 | 2.106 | -2.607 | 1.33 | 47.88 | -31.18 |
| 5 | 2jsa-3 | SER_A_5_C | CYS_A_6_N | 6.262 | 2.762 | -0.778 | 6.615 | 4.032 | -0.91 | 1.32 | 95.72 | 74.47 |
| 6 | 2jsa-3 | CYS_A_6_C | GLN_A_7_N | 4.473 | 5.062 | -1.354 | 4.67 | 5.339 | -2.635 | 1.33 | 165.14 | 54.58 |
| 7 | 2jsa-3 | GLN_A_7_C | MET_A_8_N | 2.598 | 4.197 | -3.263 | 3.156 | 2.999 | -3.171 | 1.32 | 86.02 | -65.03 |
| 8 | 2jsa-3 | MET_A_8_C | CYS_A_9_N | 1.479 | 2.05 | -1.643 | 2.031 | 2.79 | -0.692 | 1.33 | 44.15 | 53.28 |
| 9 | 2jsa-3 | CYS_A_9_C | GLU_A_10_N | 0.038 | 3.853 | 0.172 | 0.147 | 4.657 | -0.876 | 1.33 | 142.25 | 82.28 |
| 10 | 2jsa-3 | GLU_A_10_C | LEU_A_11_N | -1.936 | 4.579 | -2.162 | -1.481 | 3.367 | -2.449 | 1.33 | 102.50 | -69.42 |
| 11 | 2jsa-3 | LEU_A_11_C | VAL_A_12_N | -3.136 | 1.589 | -2.282 | -2.526 | 1.202 | -1.172 | 1.32 | 33.06 | -32.39 |
| 12 | 2jsa-3 | VAL_A_12_C | VAL_A_13_N | -4.23 | 1.265 | 0.566 | -3.75 | 2.412 | 1.024 | 1.33 | 69.78 | 67.29 |
| 13 | 2jsa-3 | VAL_A_13_C | LYS_A_14_N | -5.791 | 3.73 | 0.877 | -5.475 | 4.296 | -0.278 | 1.32 | 150.70 | 60.83 |
| 14 | 2jsa-3 | LYS_A_14_C | LYS_A_15_N | -7.567 | 3.622 | -1.323 | -7.106 | 2.383 | -1.221 | 1.33 | 85.59 | -69.59 |
| 15 | 2jsa-3 | LYS_A_15_C | TYR_A_16_N | -8.748 | 1.083 | 0.017 | -8.019 | 1.306 | 1.101 | 1.33 | 35.12 | 17.01 |
| 16 | 2jsa-3 | TYR_A_16_C | GLU_A_17_N | -9.486 | 2.395 | 2.746 | -8.844 | 3.547 | 2.876 | 1.33 | 84.37 | 60.87 |
| 17 | 2jsa-3 | GLU_A_17_C | GLY_A_18_N | -10.881 | 4.821 | 2.402 | -10.814 | 4.38 | 1.155 | 1.32 | 160.32 | -81.36 |
| 18 | 2jsa-3 | GLY_A_18_C | SER_A_19_N | -12.343 | 2.94 | -0.124 | -12.367 | 2.051 | 0.859 | 1.33 | 42.14 | -91.55 |
| 19 | 2jsa-3 | SER_A_19_C | ALA_A_20_N | -14.202 | 0.477 | 0.501 | -14.886 | 0.898 | 1.555 | 1.33 | 37.31 | 148.39 |
| 20 | 2jsa-3 | ALA_A_20_C | ASP_A_21_N | -16.722 | -0.613 | 2.074 | -15.713 | -1.465 | 2.186 | 1.33 | 85.15 | -40.18 |
| 21 | 2jsa-3 | ASP_A_21_C | LYS_A_22_N | -15.238 | -3.048 | 3.972 | -14.405 | -2.082 | 4.333 | 1.33 | 74.20 | 49.23 |
| 22 | 2jsa-3 | LYS_A_22_C | ASP_A_23_N | -13.044 | -3.54 | 5.727 | -12.156 | -3.848 | 4.792 | 1.33 | 134.85 | -19.13 |
| 23 | 2jsa-3 | ASP_A_23_C | ALA_A_24_N | -10.043 | -4.949 | 4.298 | -9.099 | -5.308 | 5.156 | 1.33 | 49.65 | -20.82 |
| 24 | 2jsa-3 | ALA_A_24_C | ASN_A_25_N | -7.307 | -6.319 | 3.856 | -8.283 | -7.168 | 3.57 | 1.32 | 102.47 | -138.98 |
| 25 | 2jsa-3 | ASN_A_25_C | VAL_A_26_N | -8.29 | -7.838 | 1.231 | -9.396 | -7.136 | 1.034 | 1.32 | 98.55 | 147.60 |
| 26 | 2jsa-3 | VAL_A_26_C | ILE_A_27_N | -8.916 | -5.39 | -0.592 | -8.347 | -4.826 | 0.462 | 1.32 | 37.24 | 44.75 |
| 27 | 2jsa-3 | ILE_A_27_C | LYS_A_28_N | -6.158 | -4.016 | -0.23 | -5.401 | -4.724 | 0.594 | 1.32 | 51.52 | -43.08 |
| 28 | 2jsa-3 | LYS_A_28_C | LYS_A_29_N | -4.151 | -5.866 | -1.152 | -5.263 | -6.557 | -1.36 | 1.33 | 99.03 | -148.14 |
| 29 | 2jsa-3 | LYS_A_29_C | ASP_A_30_N | -5.635 | -6.277 | -3.749 | -6.707 | -5.504 | -3.67 | 1.32 | 86.58 | 144.21 |
| 30 | 2jsa-3 | ASP_A_30_C | PHE_A_31_N | -5.689 | -3.794 | -5.07 | -4.974 | -3.346 | -4.049 | 1.32 | 39.57 | 32.07 |
| 31 | 2jsa-3 | PHE_A_31_C | ASP_A_32_N | -2.886 | -3.34 | -5.347 | -2.6 | -4.611 | -5.11 | 1.32 | 79.69 | -77.32 |

| | | | | | | | | | | | |
|---|---|---|---|---|---|---|---|---|---|---|---|
| 32 | 2jsa-3 | ASP_A_32_C | ALA_A_33_N | -2.464 | -5.305 | -7.439 | -3.779 | -5.145 | -7.433 | 1.32 | 89.74 | 173.06 |
| 33 | 2jsa-3 | ALA_A_33_C | GLU_A_34_N | -4.057 | -3.814 | -9.452 | -4.078 | -2.68 | -8.767 | 1.32 | 58.87 | 91.06 |
| 34 | 2jsa-3 | GLU_A_34_C | CYS_A_35_N | -2.133 | -1.335 | -9.401 | -1.502 | -2.296 | -8.742 | 1.33 | 60.18 | -56.71 |
| 35 | 2jsa-3 | CYS_A_35_C | LYS_A_36_N | 0.472 | -2.876 | -10.013 | -0.086 | -4.009 | -10.415 | 1.33 | 107.66 | -116.22 |
| 36 | 2jsa-3 | LYS_A_36_C | LYS_A_37_N | -0.095 | -3.746 | -12.833 | -1.143 | -2.967 | -12.606 | 1.33 | 80.14 | 143.38 |
| 37 | 2jsa-3 | LYS_A_37_C | LEU_A_38_N | -0.796 | -0.79 | -13.637 | -0.728 | -0.157 | -12.475 | 1.32 | 28.72 | 83.87 |
| 38 | 2jsa-3 | LEU_A_38_C | PHE_A_39_N | 1.517 | 0.772 | -12.602 | 2.054 | -0.152 | -11.82 | 1.32 | 53.81 | -59.84 |
| 39 | 2jsa-3 | PHE_A_39_C | HIS_A_40_N | 3.614 | -1.62 | -13.025 | 2.725 | -1.583 | -14.007 | 1.33 | 137.82 | 177.62 |
| 40 | 2jsa-3 | HIS_A_40_C | THR_A_41_N | 4.115 | -2.483 | -15.795 | 4.694 | -1.291 | -15.78 | 1.33 | 89.35 | 64.09 |
| 41 | 2jsa-3 | THR_A_41_C | ILE_A_42_N | 7.013 | -0.609 | -15.384 | 7.392 | -1.531 | -14.512 | 1.32 | 48.82 | -67.65 |
| 42 | 2jsa-3 | ILE_A_42_C | PRO_A_43_N | 9.191 | -2.479 | -13.176 | 10.505 | -2.25 | -12.907 | 1.36 | 78.60 | 9.89 |
| 43 | 2jsa-3 | PRO_A_43_C | PHE_A_44_N | 11.175 | -3.874 | -11.187 | 10.441 | -4.974 | -11.113 | 1.32 | 86.80 | -123.71 |
| 44 | 2jsa-3 | PHE_A_44_C | GLY_A_45_N | 9.051 | -4.803 | -9.09 | 8.582 | -3.748 | -9.74 | 1.32 | 119.38 | 113.97 |
| 45 | 2jsa-3 | GLY_A_45_C | THR_A_46_N | 6.416 | -3.762 | -8.577 | 5.805 | -4.55 | -9.447 | 1.32 | 131.10 | -127.79 |
| 46 | 2jsa-3 | THR_A_46_C | ARG_A_47_N | 5.203 | -6.451 | -8.024 | 6.399 | -6.962 | -8.281 | 1.33 | 101.18 | -23.13 |
| 47 | 2jsa-3 | ARG_A_47_C | GLU_A_48_N | 7.015 | -7.448 | -5.962 | 7.245 | -6.149 | -5.83 | 1.33 | 84.29 | 79.96 |
| 48 | 2jsa-3 | GLU_A_48_C | CYS_A_49_N | 5.881 | -5.307 | -3.98 | 4.976 | -4.992 | -4.895 | 1.32 | 133.68 | 160.81 |
| 49 | 2jsa-3 | CYS_A_49_C | ASP_A_50_N | 3.079 | -5.981 | -3.767 | 3.452 | -7.148 | -4.27 | 1.32 | 112.32 | -72.27 |
| 50 | 2jsa-3 | ASP_A_50_C | HIS_A_51_N | 3.688 | -8.513 | -2.27 | 4.934 | -8.069 | -2.196 | 1.32 | 86.80 | 19.61 |
| 51 | 2jsa-3 | HIS_A_51_C | TYR_A_52_N | 5.367 | -6.869 | -0.119 | 4.338 | -6.152 | -0.548 | 1.33 | 108.88 | 145.13 |
| 52 | 2jsa-3 | TYR_A_52_C | VAL_A_53_N | 2.581 | -5.129 | 0.834 | 1.633 | -5.66 | 0.077 | 1.32 | 124.86 | -150.75 |
| 53 | 2jsa-3 | VAL_A_53_C | ASN_A_54_N | 0.401 | -6.725 | 1.885 | 1.54 | -7.388 | 2.035 | 1.33 | 83.51 | -30.20 |
| 54 | 2jsa-3 | ASN_A_54_C | SER_A_55_N | 3.188 | -7.954 | 3.733 | 3.369 | -6.763 | 4.283 | 1.32 | 65.46 | 81.36 |
| 55 | 2jsa-3 | SER_A_55_C | LYS_A_56_N | 4.649 | -4.895 | 5.204 | 3.903 | -4.126 | 4.426 | 1.32 | 125.99 | 134.13 |
| 56 | 2jsa-3 | LYS_A_56_C | VAL_A_57_N | 2.322 | -2.294 | 4.691 | 1.466 | -3.302 | 4.605 | 1.33 | 93.72 | -130.34 |
| 57 | 2jsa-3 | VAL_A_57_C | ASP_A_58_N | -0.499 | -3.241 | 6.04 | -0.047 | -4.309 | 6.681 | 1.33 | 61.07 | -67.06 |
| 58 | 2jsa-3 | ASP_A_58_C | PRO_A_59_N | -0.279 | -3.35 | 8.905 | 0.904 | -2.702 | 8.729 | 1.36 | 97.43 | 28.71 |
| 59 | 2jsa-3 | PRO_A_59_C | ILE_A_60_N | 0.44 | -0.298 | 8.948 | 0.625 | -0.038 | 7.662 | 1.32 | 166.06 | 54.57 |
| 60 | 2jsa-3 | ILE_A_60_C | ILE_A_61_N | -1.523 | 1.084 | 7.436 | -2.19 | -0.025 | 7.156 | 1.32 | 102.21 | -121.02 |
| 61 | 2jsa-3 | ILE_A_61_C | HIS_A_62_N | -3.77 | 0.014 | 9.007 | -2.687 | -0.238 | 9.727 | 1.32 | 57.08 | -13.10 |
| 62 | 2jsa-3 | HIS_A_62_C | GLU_A_63_N | -2.949 | 1.346 | 11.562 | -2.221 | 2.23 | 10.897 | 1.32 | 120.14 | 50.53 |
| 63 | 2jsa-3 | GLU_A_63_C | LEU_A_64_N | -3.776 | 4.12 | 10.88 | -4.255 | 3.775 | 9.693 | 1.33 | 153.56 | -144.24 |
| 64 | 2jsa-3 | LEU_A_64_C | GLU_A_65_N | -6.608 | 3.554 | 10.274 | -6.309 | 2.34 | 10.714 | 1.33 | 70.61 | -76.16 |
| 65 | 2jsa-3 | GLU_A_65_C | GLY_A_66_N | -7.321 | 2.464 | 12.94 | -6.179 | 2.871 | 13.475 | 1.33 | 66.19 | 19.62 |
| 66 | 2jsa-3 | GLY_A_66_C | GLY_A_67_N | -6.888 | 4.988 | 14.506 | -6.585 | 5.633 | 13.39 | 1.32 | 147.44 | 64.84 |
| 67 | 2jsa-3 | GLY_A_67_C | THR_A_68_N | -6.157 | 7.906 | 12.554 | -5.291 | 7.405 | 11.686 | 1.32 | 130.94 | -30.05 |
| 68 | 2jsa-3 | THR_A_68_C | ALA_A_69_N | -4.473 | 8.409 | 9.606 | -4.404 | 9.658 | 9.171 | 1.32 | 109.18 | 86.84 |
| 69 | 2jsa-3 | ALA_A_69_C | PRO_A_70_N | -3.6 | 9.178 | 6.925 | -3.904 | 9.087 | 5.603 | 1.36 | 166.50 | -163.34 |
| 70 | 2jsa-3 | PRO_A_70_C | LYS_A_71_N | -1.776 | 9.159 | 4.373 | -1.827 | 10.442 | 4.702 | 1.33 | 75.63 | 92.28 |
| 71 | 2jsa-3 | LYS_A_71_C | ASP_A_72_N | 0.138 | 11.431 | 5.742 | -0.277 | 10.674 | 6.747 | 1.32 | 40.66 | -118.73 |
| 72 | 2jsa-3 | ASP_A_72_C | VAL_A_73_N | 1.184 | 9.362 | 8.185 | 0.616 | 8.309 | 7.616 | 1.32 | 115.43 | -118.34 |
| 73 | 2jsa-3 | VAL_A_73_C | CYS_A_74_N | 2.516 | 6.993 | 6.858 | 2.401 | 7.522 | 5.649 | 1.32 | 155.88 | 102.26 |
| 74 | 2jsa-3 | CYS_A_74_C | THR_A_75_N | 4.412 | 8.773 | 5.161 | 5.013 | 8.647 | 6.335 | 1.32 | 27.61 | -11.84 |
| 75 | 2jsa-3 | THR_A_75_C | LYS_A_76_N | 6.496 | 9.268 | 8.184 | 5.628 | 8.972 | 9.14 | 1.32 | 43.81 | -161.17 |
| 76 | 2jsa-3 | LYS_A_76_C | LEU_A_77_N | 6.565 | 7.1 | 10.381 | 6.014 | 6.373 | 9.42 | 1.33 | 136.49 | -127.16 |
| 77 | 2jsa-3 | LEU_A_77_C | ASN_A_78_N | 7.631 | 4.9 | 8.354 | 8.242 | 6.057 | 8.142 | 1.33 | 99.20 | 62.16 |
| 78 | 2jsa-3 | ASN_A_78_C | GLU_A_79_N | 9.229 | 5.336 | 6.038 | 7.992 | 5.362 | 5.563 | 1.33 | 111.00 | 178.80 |
| 79 | 2jsa-3 | GLU_A_79_C | CYS_A_80_N | 7.373 | 5.642 | 3.211 | 7.754 | 6.888 | 3.453 | 1.33 | 79.48 | 73.00 |
| 80 | 2jsa-3 | CYS_A_80_C | PRO_A_81_N | 8.159 | 9.232 | 3.014 | 9.426 | 9.5 | 2.598 | 1.36 | 107.81 | 11.94 |
| 1 | 2jsa-4 | ARG_A_1_C | SER_A_2_N | 13.831 | 5.809 | 1.973 | 13.569 | 5.572 | 0.696 | 1.32 | 164.54 | -137.87 |
| 2 | 2jsa-4 | SER_A_2_C | ALA_A_3_N | 12.335 | 5.701 | -1.41 | 11.19 | 5.171 | -1.813 | 1.32 | 107.71 | -155.16 |
| 3 | 2jsa-4 | ALA_A_3_C | LEU_A_4_N | 9.563 | 4.281 | -3.389 | 9.387 | 3.389 | -4.353 | 1.33 | 136.68 | -101.16 |
| 4 | 2jsa-4 | LEU_A_4_C | SER_A_5_N | 7.318 | 2.598 | -3.348 | 8.059 | 2.103 | -2.367 | 1.33 | 42.25 | -33.74 |
| 5 | 2jsa-4 | SER_A_5_C | CYS_A_6_N | 6.496 | 2.9 | -0.667 | 7.006 | 4.123 | -0.632 | 1.33 | 88.49 | 67.36 |
| 6 | 2jsa-4 | CYS_A_6_C | GLN_A_7_N | 4.93 | 5.255 | -1.138 | 5.145 | 5.456 | -2.43 | 1.33 | 167.17 | 43.07 |
| 7 | 2jsa-4 | GLN_A_7_C | MET_A_8_N | 3.066 | 4.336 | -3.075 | 3.606 | 3.127 | -3.038 | 1.32 | 88.40 | -65.93 |
| 8 | 2jsa-4 | MET_A_8_C | CYS_A_9_N | 1.936 | 2.163 | -1.513 | 2.43 | 3.015 | -0.628 | 1.32 | 48.06 | 59.89 |
| 9 | 2jsa-4 | CYS_A_9_C | GLU_A_10_N | 0.429 | 4.05 | 0.249 | 0.509 | 4.84 | -0.811 | 1.32 | 143.16 | 84.22 |
| 10 | 2jsa-4 | GLU_A_10_C | LEU_A_11_N | -1.593 | 4.707 | -2.06 | -1.125 | 3.499 | -2.338 | 1.32 | 102.11 | -68.82 |
| 11 | 2jsa-4 | LEU_A_11_C | VAL_A_12_N | -2.764 | 1.715 | -2.115 | -2.123 | 1.295 | -1.033 | 1.33 | 35.31 | -33.23 |
| 12 | 2jsa-4 | VAL_A_12_C | VAL_A_13_N | -3.858 | 1.361 | 0.672 | -3.419 | 2.521 | 1.138 | 1.32 | 69.41 | 69.27 |
| 13 | 2jsa-4 | VAL_A_13_C | LYS_A_14_N | -5.46 | 3.821 | 0.883 | -5.087 | 4.367 | -0.264 | 1.32 | 150.04 | 55.66 |
| 14 | 2jsa-4 | LYS_A_14_C | LYS_A_15_N | -7.106 | 3.668 | -1.428 | -6.648 | 2.439 | -1.242 | 1.32 | 81.93 | -69.56 |
| 15 | 2jsa-4 | LYS_A_15_C | TYR_A_16_N | -8.301 | 1.088 | -0.075 | -7.616 | 1.339 | 1.032 | 1.33 | 33.39 | 20.12 |
| 16 | 2jsa-4 | TYR_A_16_C | GLU_A_17_N | -9.222 | 2.325 | 2.61 | -8.68 | 3.53 | 2.72 | 1.33 | 85.24 | 65.78 |
| 17 | 2jsa-4 | GLU_A_17_C | GLY_A_18_N | -10.74 | 4.686 | 2.077 | -10.515 | 4.334 | 0.819 | 1.33 | 161.63 | -57.41 |
| 18 | 2jsa-4 | GLY_A_18_C | SER_A_19_N | -11.824 | 2.843 | -0.633 | -11.966 | 1.942 | 0.328 | 1.32 | 43.51 | -98.96 |
| 19 | 2jsa-4 | SER_A_19_C | ALA_A_20_N | -13.686 | 0.267 | -0.132 | -14.438 | 0.733 | 0.853 | 1.32 | 41.93 | 148.21 |
| 20 | 2jsa-4 | ALA_A_20_C | ASP_A_21_N | -16.217 | -0.799 | 1.491 | -15.788 | -0.948 | 2.736 | 1.33 | 20.04 | -19.15 |
| 21 | 2jsa-4 | ASP_A_21_C | LYS_A_22_N | -15.082 | -2.253 | 4.664 | -13.947 | -1.587 | 4.517 | 1.32 | 96.37 | 30.40 |
| 22 | 2jsa-4 | LYS_A_22_C | ASP_A_23_N | -12.378 | -2.982 | 5.747 | -12.018 | -3.586 | 4.623 | 1.33 | 147.97 | -59.20 |
| 23 | 2jsa-4 | ASP_A_23_C | ALA_A_24_N | -9.999 | -4.875 | 4.2 | -9.118 | -5.145 | 5.153 | 1.33 | 44.04 | -17.04 |
| 24 | 2jsa-4 | ALA_A_24_C | ASN_A_25_N | -7.352 | -6.268 | 3.912 | -8.349 | -7.1 | 3.654 | 1.32 | 101.24 | -140.15 |
| 25 | 2jsa-4 | ASN_A_25_C | VAL_A_26_N | -8.401 | -7.765 | 1.314 | -9.577 | -7.196 | 1.091 | 1.33 | 99.69 | 154.18 |
| 26 | 2jsa-4 | VAL_A_26_C | ILE_A_27_N | -9.156 | -5.423 | -0.523 | -8.461 | -4.945 | 0.499 | 1.33 | 39.53 | 34.52 |
| 27 | 2jsa-4 | ILE_A_27_C | LYS_A_28_N | -6.286 | -4.082 | -0.168 | -5.462 | -4.614 | 0.723 | 1.33 | 47.75 | -32.85 |
| 28 | 2jsa-4 | LYS_A_28_C | LYS_A_29_N | -4.131 | -5.684 | -1.011 | -5.167 | -6.483 | -1.22 | 1.32 | 99.08 | -142.36 |
| 29 | 2jsa-4 | LYS_A_29_C | ASP_A_30_N | -5.527 | -6.196 | -3.609 | -6.417 | -5.247 | -3.36 | 1.32 | 79.17 | 133.16 |
| 30 | 2jsa-4 | ASP_A_30_C | PHE_A_31_N | -5.412 | -3.602 | -4.844 | -4.602 | -3.166 | -3.891 | 1.32 | 43.99 | 28.29 |
| 31 | 2jsa-4 | PHE_A_31_C | ASP_A_32_N | -2.593 | -3.346 | -5.298 | -2.461 | -4.64 | -5.049 | 1.32 | 79.16 | -84.18 |
| 32 | 2jsa-4 | ASP_A_32_C | ALA_A_33_N | -2.526 | -5.488 | -7.33 | -3.834 | -5.297 | -7.242 | 1.32 | 86.19 | 171.69 |
| 33 | 2jsa-4 | ALA_A_33_C | GLU_A_34_N | -4.24 | -4.053 | -9.294 | -4.52 | -2.864 | -8.781 | 1.32 | 67.22 | 103.25 |
| 34 | 2jsa-4 | GLU_A_34_C | CYS_A_35_N | -2.73 | -1.709 | -9.989 | -1.815 | -1.856 | -9.042 | 1.33 | 44.38 | -9.13 |
| 35 | 2jsa-4 | CYS_A_35_C | LYS_A_36_N | -0.238 | -2.89 | -10.555 | -0.726 | -4.107 | -10.365 | 1.32 | 81.75 | -111.85 |
| 36 | 2jsa-4 | LYS_A_36_C | LYS_A_37_N | -0.906 | -4.47 | -12.765 | -1.71 | -3.418 | -12.735 | 1.32 | 88.70 | 127.39 |
| 37 | 2jsa-4 | LYS_A_37_C | LEU_A_38_N | -1.173 | -1.453 | -14.068 | -1.217 | -0.64 | -13.022 | 1.33 | 37.90 | 93.10 |
| 38 | 2jsa-4 | LEU_A_38_C | PHE_A_39_N | 1.008 | 0.272 | -13.396 | 1.525 | -0.812 | -12.837 | 1.32 | 65.04 | -64.50 |
| 39 | 2jsa-4 | PHE_A_39_C | HIS_A_40_N | 2.905 | -2.596 | -13.815 | 2.042 | -2.736 | -14.81 | 1.32 | 138.70 | -170.79 |
| 40 | 2jsa-4 | HIS_A_40_C | THR_A_41_N | 3.234 | -4.196 | -16.355 | 3.797 | -3.089 | -16.818 | 1.33 | 110.45 | 63.04 |
| 41 | 2jsa-4 | THR_A_41_C | ILE_A_42_N | 6.229 | -3.357 | -16.681 | 6.243 | -2.606 | -15.589 | 1.33 | 34.52 | 88.93 |
| 42 | 2jsa-4 | ILE_A_42_C | PRO_A_43_N | 7.56 | -4.175 | -14.276 | 8.863 | -4.525 | -14.101 | 1.36 | 82.61 | -15.04 |
| 43 | 2jsa-4 | PRO_A_43_C | PHE_A_44_N | 8.919 | -6.195 | -12.298 | 9.932 | -6.03 | -11.461 | 1.32 | 50.80 | 9.25 |
| 44 | 2jsa-4 | PHE_A_44_C | GLY_A_45_N | 8.554 | -5.554 | -9.481 | 8.196 | -4.468 | -10.15 | 1.32 | 120.33 | 108.24 |
| 45 | 2jsa-4 | GLY_A_45_C | THR_A_46_N | 5.947 | -4.567 | -9.162 | 5.442 | -5.448 | -10.013 | 1.32 | 129.96 | -119.82 |
| 46 | 2jsa-4 | THR_A_46_C | ARG_A_47_N | 4.848 | -7.222 | -8.433 | 6.11 | -7.621 | -8.494 | 1.32 | 92.64 | -17.55 |
| 47 | 2jsa-4 | ARG_A_47_C | GLU_A_48_N | 6.726 | -7.676 | -6.125 | 7.421 | -6.548 | -6.154 | 1.33 | 91.25 | 58.36 |

| | | | | | | | | | | | |
|---|---|---|---|---|---|---|---|---|---|---|---|
| 48 | 2jsa-4 | GLU_A_48_C | CYS_A_49_N | 6.16 | -5.345 | -4.435 | 5.254 | -5.073 | -5.363 | 1.33 | 134.45 | 163.29 |
| 49 | 2jsa-4 | CYS_A_49_C | ASP_A_50_N | 3.257 | -5.894 | -4.275 | 3.505 | -7.079 | -4.814 | 1.33 | 114.00 | -78.18 |
| 50 | 2jsa-4 | ASP_A_50_C | HIS_A_51_N | 3.704 | -8.607 | -2.931 | 4.726 | -7.808 | -2.664 | 1.32 | 78.37 | 38.02 |
| 51 | 2jsa-4 | HIS_A_51_C | TYR_A_52_N | 5.311 | -6.799 | -0.522 | 4.447 | -5.892 | -0.954 | 1.33 | 109.03 | 133.61 |
| 52 | 2jsa-4 | TYR_A_52_C | VAL_A_53_N | 2.809 | -4.858 | 0.562 | 1.846 | -5.426 | -0.148 | 1.32 | 122.42 | -149.47 |
| 53 | 2jsa-4 | VAL_A_53_C | ASN_A_54_N | 0.643 | -6.459 | 1.697 | 1.79 | -7.106 | 1.841 | 1.32 | 83.76 | -29.43 |
| 54 | 2jsa-4 | ASN_A_54_C | SER_A_55_N | 3.46 | -7.662 | 3.522 | 3.669 | -6.434 | 3.973 | 1.32 | 70.10 | 80.34 |
| 55 | 2jsa-4 | SER_A_55_C | LYS_A_56_N | 4.897 | -4.617 | 5.052 | 4.173 | -3.767 | 4.337 | 1.33 | 122.63 | 130.42 |
| 56 | 2jsa-4 | LYS_A_56_C | VAL_A_57_N | 2.53 | -2.023 | 4.764 | 1.704 | -3.049 | 4.615 | 1.33 | 96.45 | -128.84 |
| 57 | 2jsa-4 | VAL_A_57_C | ASP_A_58_N | -0.285 | -3.096 | 6.015 | 0.154 | -4.197 | 6.607 | 1.32 | 63.46 | -68.26 |
| 58 | 2jsa-4 | ASP_A_58_C | PRO_A_59_N | -0.113 | -3.344 | 8.872 | 1.067 | -2.682 | 8.741 | 1.36 | 95.53 | 29.29 |
| 59 | 2jsa-4 | PRO_A_59_C | ILE_A_60_N | 0.586 | -0.293 | 9.067 | 0.798 | 0.038 | 7.802 | 1.32 | 162.74 | 57.36 |
| 60 | 2jsa-4 | ILE_A_60_C | ILE_A_61_N | -1.353 | 1.158 | 7.601 | -2.019 | 0.076 | 7.225 | 1.33 | 106.49 | -121.61 |
| 61 | 2jsa-4 | ILE_A_61_C | HIS_A_62_N | -3.634 | 0.047 | 9.046 | -2.564 | -0.223 | 9.779 | 1.32 | 56.41 | -14.16 |
| 62 | 2jsa-4 | HIS_A_62_C | GLU_A_63_N | -2.769 | 1.316 | 11.658 | -2.316 | 2.208 | 10.789 | 1.33 | 130.98 | 63.08 |
| 63 | 2jsa-4 | GLU_A_63_C | LEU_A_64_N | -3.831 | 4.131 | 10.85 | -4.367 | 3.806 | 9.683 | 1.32 | 151.76 | -148.77 |
| 64 | 2jsa-4 | LEU_A_64_C | GLU_A_65_N | -6.708 | 3.527 | 10.287 | -6.303 | 2.361 | 10.77 | 1.33 | 68.63 | -70.85 |
| 65 | 2jsa-4 | GLU_A_65_C | GLY_A_66_N | -7.29 | 2.337 | 13.011 | -6.157 | 2.784 | 13.534 | 1.33 | 66.76 | 21.53 |
| 66 | 2jsa-4 | GLY_A_66_C | GLY_A_67_N | -6.874 | 4.847 | 14.665 | -6.406 | 5.647 | 13.718 | 1.33 | 135.62 | 59.67 |
| 67 | 2jsa-4 | GLY_A_67_C | THR_A_68_N | -5.999 | 7.892 | 12.801 | -5.298 | 7.347 | 11.818 | 1.32 | 137.91 | -37.86 |
| 68 | 2jsa-4 | THR_A_68_C | ALA_A_69_N | -4.734 | 8.271 | 9.62 | -4.589 | 9.487 | 9.116 | 1.32 | 112.37 | 83.20 |
| 69 | 2jsa-4 | ALA_A_69_C | PRO_A_70_N | -4.014 | 9.008 | 6.8 | -4.369 | 9.032 | 5.488 | 1.36 | 164.83 | 176.13 |
| 70 | 2jsa-4 | PRO_A_70_C | LYS_A_71_N | -2.286 | 9.119 | 4.185 | -2.311 | 10.401 | 4.52 | 1.33 | 75.36 | 91.12 |
| 71 | 2jsa-4 | LYS_A_71_C | ASP_A_72_N | -0.315 | 11.317 | 5.569 | -0.864 | 10.744 | 6.63 | 1.32 | 36.79 | -133.77 |
| 72 | 2jsa-4 | ASP_A_72_C | VAL_A_73_N | 0.528 | 9.381 | 8.087 | -0.137 | 8.334 | 7.621 | 1.32 | 110.59 | -122.42 |
| 73 | 2jsa-4 | VAL_A_73_C | CYS_A_74_N | 1.847 | 6.993 | 7.189 | 1.989 | 7.476 | 5.964 | 1.32 | 157.66 | 73.62 |
| 74 | 2jsa-4 | CYS_A_74_C | THR_A_75_N | 4.015 | 8.791 | 5.834 | 4.114 | 8.889 | 7.151 | 1.32 | 6.04 | 44.71 |
| 75 | 2jsa-4 | THR_A_75_C | LYS_A_76_N | 5.504 | 9.592 | 9.041 | 4.744 | 9.001 | 9.95 | 1.32 | 46.64 | -142.13 |
| 76 | 2jsa-4 | LYS_A_76_C | LEU_A_77_N | 5.944 | 7.162 | 10.999 | 5.438 | 6.455 | 10.0 | 1.32 | 138.97 | -125.59 |
| 77 | 2jsa-4 | LEU_A_77_C | ASN_A_78_N | 7.272 | 5.264 | 8.933 | 7.688 | 6.509 | 8.749 | 1.33 | 97.98 | 71.52 |
| 78 | 2jsa-4 | ASN_A_78_C | GLU_A_79_N | 8.893 | 6.13 | 6.667 | 7.686 | 5.772 | 6.251 | 1.33 | 108.29 | -163.48 |
| 79 | 2jsa-4 | GLU_A_79_C | CYS_A_80_N | 7.018 | 6.162 | 3.928 | 7.812 | 7.212 | 3.771 | 1.33 | 96.80 | 52.90 |
| 80 | 2jsa-4 | CYS_A_80_C | PRO_A_81_N | 8.375 | 9.472 | 3.119 | 8.663 | 10.25 | 2.042 | 1.36 | 142.39 | 69.69 |
| 1 | 2jsa-5 | ARG_A_1_C | SER_A_2_N | 13.136 | 3.863 | -0.521 | 12.151 | 4.637 | -0.092 | 1.32 | 71.10 | 141.84 |
| 2 | 2jsa-5 | SER_A_2_C | ALA_A_3_N | 11.851 | 6.475 | -1.674 | 10.728 | 5.943 | -2.134 | 1.33 | 110.31 | -154.65 |
| 3 | 2jsa-5 | ALA_A_3_C | LEU_A_4_N | 8.937 | 5.501 | -3.722 | 9.054 | 4.236 | -4.095 | 1.32 | 106.36 | -84.72 |
| 4 | 2jsa-5 | LEU_A_4_C | SER_A_5_N | 7.213 | 3.052 | -3.033 | 8.049 | 2.789 | -2.04 | 1.32 | 41.43 | -17.46 |
| 5 | 2jsa-5 | SER_A_5_C | CYS_A_6_N | 6.516 | 3.442 | -0.252 | 6.799 | 4.704 | -0.539 | 1.32 | 102.51 | 77.36 |
| 6 | 2jsa-5 | CYS_A_6_C | GLN_A_7_N | 4.638 | 5.682 | -1.009 | 4.841 | 5.763 | -2.317 | 1.33 | 170.51 | 21.75 |
| 7 | 2jsa-5 | GLN_A_7_C | MET_A_8_N | 2.863 | 4.465 | -2.946 | 3.509 | 3.309 | -2.918 | 1.32 | 88.79 | -60.80 |
| 8 | 2jsa-5 | MET_A_8_C | CYS_A_9_N | 1.905 | 2.21 | -1.414 | 2.446 | 2.853 | -0.39 | 1.32 | 39.37 | 49.92 |
| 9 | 2jsa-5 | CYS_A_9_C | GLU_A_10_N | 0.428 | 3.845 | 0.503 | 0.538 | 4.675 | -0.524 | 1.33 | 140.81 | 82.45 |
| 10 | 2jsa-5 | GLU_A_10_C | LEU_A_11_N | -1.553 | 4.641 | -1.798 | -1.106 | 3.438 | -2.125 | 1.32 | 104.29 | -69.62 |
| 11 | 2jsa-5 | LEU_A_11_C | VAL_A_12_N | -2.755 | 1.654 | -1.995 | -2.131 | 1.233 | -0.904 | 1.33 | 34.60 | -34.01 |
| 12 | 2jsa-5 | VAL_A_12_C | VAL_A_13_N | -3.825 | 1.232 | 0.844 | -3.368 | 2.394 | 1.289 | 1.33 | 70.38 | 68.53 |
| 13 | 2jsa-5 | VAL_A_13_C | LYS_A_14_N | -5.401 | 3.724 | 1.166 | -5.068 | 4.35 | 0.046 | 1.33 | 147.66 | 61.99 |
| 14 | 2jsa-5 | LYS_A_14_C | LYS_A_15_N | -7.128 | 3.714 | -1.084 | -6.659 | 2.477 | -1.002 | 1.33 | 86.45 | -69.24 |
| 15 | 2jsa-5 | LYS_A_15_C | TYR_A_16_N | -8.333 | 1.11 | 0.116 | -7.644 | 1.275 | 1.235 | 1.32 | 32.34 | 13.47 |
| 16 | 2jsa-5 | TYR_A_16_C | GLU_A_17_N | -9.113 | 2.318 | 2.908 | -8.419 | 3.403 | 3.219 | 1.32 | 76.42 | 57.40 |
| 17 | 2jsa-5 | GLU_A_17_C | GLY_A_18_N | -10.309 | 4.872 | 2.704 | -10.16 | 4.502 | 1.44 | 1.33 | 162.49 | -68.07 |
| 18 | 2jsa-5 | GLY_A_18_C | SER_A_19_N | -11.682 | 3.314 | -0.082 | -11.863 | 2.363 | 0.823 | 1.33 | 46.93 | -100.78 |
| 19 | 2jsa-5 | SER_A_19_C | ALA_A_20_N | -13.808 | 0.916 | 0.515 | -14.337 | 1.305 | 1.666 | 1.33 | 29.70 | 143.67 |
| 20 | 2jsa-5 | ALA_A_20_C | ASP_A_21_N | -16.185 | -0.189 | 2.189 | -15.191 | -1.065 | 2.204 | 1.33 | 89.35 | -41.39 |
| 21 | 2jsa-5 | ASP_A_21_C | LYS_A_22_N | -14.816 | -2.838 | 3.828 | -14.006 | -1.923 | 4.339 | 1.32 | 67.31 | 48.48 |
| 22 | 2jsa-5 | LYS_A_22_C | ASP_A_23_N | -12.782 | -3.559 | 5.661 | -12.046 | -3.902 | 4.615 | 1.32 | 142.18 | -24.99 |
| 23 | 2jsa-5 | ASP_A_23_C | ALA_A_24_N | -9.997 | -5.088 | 4.052 | -9.081 | -5.32 | 4.982 | 1.33 | 45.46 | -14.21 |
| 24 | 2jsa-5 | ALA_A_24_C | ASN_A_25_N | -7.299 | -6.296 | 3.642 | -8.242 | -7.198 | 3.413 | 1.32 | 99.95 | -136.27 |
| 25 | 2jsa-5 | ASN_A_25_C | VAL_A_26_N | -8.289 | -7.745 | 1.043 | -9.434 | -7.096 | 0.885 | 1.33 | 96.85 | 150.45 |
| 26 | 2jsa-5 | VAL_A_26_C | ILE_A_27_N | -9.018 | -5.255 | -0.651 | -8.305 | -4.83 | 0.383 | 1.33 | 38.76 | 30.80 |
| 27 | 2jsa-5 | ILE_A_27_C | LYS_A_28_N | -6.108 | -3.976 | -0.22 | -5.389 | -4.735 | 0.596 | 1.33 | 52.03 | -46.55 |
| 28 | 2jsa-5 | LYS_A_28_C | LYS_A_29_N | -4.06 | -5.774 | -1.158 | -5.103 | -6.552 | -1.412 | 1.33 | 101.05 | -143.28 |
| 29 | 2jsa-5 | LYS_A_29_C | ASP_A_30_N | -5.443 | -6.154 | -3.789 | -6.341 | -5.22 | -3.509 | 1.33 | 77.81 | 133.87 |
| 30 | 2jsa-5 | ASP_A_30_C | PHE_A_31_N | -5.314 | -3.523 | -4.919 | -4.502 | -3.135 | -3.947 | 1.32 | 42.80 | 25.54 |
| 31 | 2jsa-5 | PHE_A_31_C | ASP_A_32_N | -2.487 | -3.285 | -5.349 | -2.291 | -4.572 | -5.104 | 1.32 | 79.34 | -81.34 |
| 32 | 2jsa-5 | ASP_A_32_C | ALA_A_33_N | -2.39 | -5.423 | -7.383 | -3.672 | -5.106 | -7.274 | 1.33 | 85.28 | 166.11 |
| 33 | 2jsa-5 | ALA_A_33_C | GLU_A_34_N | -4.162 | -3.858 | -9.306 | -4.346 | -2.68 | -8.73 | 1.32 | 64.21 | 98.88 |
| 34 | 2jsa-5 | GLU_A_34_C | CYS_A_35_N | -2.579 | -1.456 | -9.899 | -1.698 | -1.832 | -8.983 | 1.33 | 46.28 | -23.11 |
| 35 | 2jsa-5 | CYS_A_35_C | LYS_A_36_N | -0.076 | -2.849 | -10.461 | -0.694 | -4.016 | -10.342 | 1.33 | 84.85 | -117.90 |
| 36 | 2jsa-5 | LYS_A_36_C | LYS_A_37_N | -0.674 | -4.353 | -12.752 | -1.521 | -3.338 | -12.826 | 1.32 | 93.20 | 129.84 |
| 37 | 2jsa-5 | LYS_A_37_C | LEU_A_38_N | -0.791 | -1.376 | -14.069 | -0.933 | -0.574 | -13.024 | 1.32 | 37.93 | 100.04 |
| 38 | 2jsa-5 | LEU_A_38_C | PHE_A_39_N | 1.329 | 0.305 | -13.209 | 1.802 | -0.782 | -12.617 | 1.33 | 63.46 | -66.48 |
| 39 | 2jsa-5 | PHE_A_39_C | HIS_A_40_N | 3.22 | -2.591 | -13.49 | 2.353 | -2.768 | -14.477 | 1.33 | 138.12 | -168.46 |
| 40 | 2jsa-5 | HIS_A_40_C | THR_A_41_N | 3.562 | -4.221 | -16.016 | 4.209 | -3.104 | -16.315 | 1.33 | 103.04 | 59.92 |
| 41 | 2jsa-5 | THR_A_41_C | ILE_A_42_N | 6.65 | -3.243 | -16.164 | 6.622 | -2.442 | -15.109 | 1.32 | 37.22 | 92.00 |
| 42 | 2jsa-5 | ILE_A_42_C | PRO_A_43_N | 8.195 | -3.871 | -13.924 | 9.54 | -4.005 | -13.772 | 1.36 | 83.58 | -5.69 |
| 43 | 2jsa-5 | PRO_A_43_C | PHE_A_44_N | 9.9 | -5.796 | -12.127 | 10.222 | -4.929 | -11.178 | 1.33 | 44.26 | 69.63 |
| 44 | 2jsa-5 | PHE_A_44_C | GLY_A_45_N | 8.874 | -4.475 | -9.172 | 7.874 | -4.235 | -10.005 | 1.32 | 129.01 | 166.50 |
| 45 | 2jsa-5 | GLY_A_45_C | THR_A_46_N | 5.622 | -4.454 | -9.04 | 5.286 | -5.434 | -9.867 | 1.33 | 128.60 | -108.92 |
| 46 | 2jsa-5 | THR_A_46_C | ARG_A_47_N | 4.822 | -7.357 | -8.422 | 6.122 | -7.602 | -8.498 | 1.33 | 93.29 | -10.67 |
| 47 | 2jsa-5 | ARG_A_47_C | GLU_A_48_N | 6.807 | -7.831 | -6.159 | 7.421 | -6.658 | -6.105 | 1.33 | 87.66 | 62.37 |
| 48 | 2jsa-5 | GLU_A_48_C | CYS_A_49_N | 6.137 | -5.565 | -4.332 | 5.222 | -5.359 | -5.268 | 1.33 | 134.94 | 167.31 |
| 49 | 2jsa-5 | CYS_A_49_C | ASP_A_50_N | 3.233 | -6.193 | -4.173 | 3.334 | -7.358 | -4.795 | 1.32 | 118.01 | -85.05 |
| 50 | 2jsa-5 | ASP_A_50_C | HIS_A_51_N | 3.5 | -8.858 | -2.884 | 4.619 | -8.176 | -2.693 | 1.32 | 81.71 | 31.36 |
| 51 | 2jsa-5 | HIS_A_51_C | TYR_A_52_N | 5.191 | -7.146 | -0.558 | 4.346 | -6.233 | -1.014 | 1.32 | 110.13 | 132.78 |
| 52 | 2jsa-5 | TYR_A_52_C | VAL_A_53_N | 2.679 | -5.144 | 0.429 | 1.727 | -5.677 | -0.322 | 1.32 | 124.54 | -150.76 |
| 53 | 2jsa-5 | VAL_A_53_C | ASN_A_54_N | 0.416 | -6.669 | 1.471 | 1.529 | -7.364 | 1.655 | 1.33 | 82.02 | -31.98 |
| 54 | 2jsa-5 | ASN_A_54_C | SER_A_55_N | 3.123 | -8.009 | 3.377 | 3.363 | -6.822 | 3.915 | 1.33 | 66.05 | 78.57 |
| 55 | 2jsa-5 | SER_A_55_C | LYS_A_56_N | 4.669 | -5.072 | 5.012 | 3.945 | -4.209 | 4.314 | 1.33 | 121.78 | 129.99 |
| 56 | 2jsa-5 | LYS_A_56_C | VAL_A_57_N | 2.389 | -2.38 | 4.706 | 1.514 | -3.365 | 4.567 | 1.32 | 96.02 | -131.62 |
| 57 | 2jsa-5 | VAL_A_57_C | ASP_A_58_N | -0.493 | -3.331 | 5.942 | -0.072 | -4.431 | 6.548 | 1.32 | 62.77 | -69.06 |
| 58 | 2jsa-5 | ASP_A_58_C | PRO_A_59_N | -0.321 | -3.587 | 8.817 | 0.891 | -2.981 | 8.705 | 1.36 | 94.72 | 26.57 |
| 59 | 2jsa-5 | PRO_A_59_C | ILE_A_60_N | 0.52 | -0.575 | 9.049 | 0.693 | -0.288 | 7.767 | 1.33 | 165.35 | 58.92 |
| 60 | 2jsa-5 | ILE_A_60_C | ILE_A_61_N | -1.385 | 0.971 | 7.631 | -2.112 | -0.111 | 7.401 | 1.32 | 100.01 | -123.90 |
| 61 | 2jsa-5 | ILE_A_61_C | HIS_A_62_N | -3.631 | 0.048 | 9.295 | -2.556 | -0.285 | 9.994 | 1.32 | 58.15 | -17.21 |
| 62 | 2jsa-5 | HIS_A_62_C | GLU_A_63_N | -2.559 | 1.354 | 11.8 | -2.175 | 2.173 | 10.832 | 1.32 | 136.94 | 64.88 |
| 63 | 2jsa-5 | GLU_A_63_C | LEU_A_64_N | -3.522 | 4.213 | 10.703 | -4.087 | 3.748 | 9.599 | 1.32 | 146.46 | -140.55 |

| | | | | | | | | | | |
|---|---|---|---|---|---|---|---|---|---|---|
| 64 | 2jsa-5 | LEU_A_64_C | GLU_A_65_N | -6.45 | 3.804 | 10.178 | -6.319 | 2.57 | 10.643 | 1.33 | 69.46 | -83.94 |
| 65 | 2jsa-5 | GLU_A_65_C | GLY_A_66_N | -7.101 | 2.75 | 12.957 | -5.868 | 3.142 | 13.241 | 1.32 | 77.62 | 17.64 |
| 66 | 2jsa-5 | GLY_A_66_C | GLY_A_67_N | -6.2 | 5.231 | 14.495 | -5.978 | 5.955 | 13.407 | 1.33 | 145.16 | 72.95 |
| 67 | 2jsa-5 | GLY_A_67_C | THR_A_68_N | -5.772 | 8.087 | 12.201 | -4.54 | 7.668 | 11.95 | 1.33 | 100.92 | -18.78 |
| 68 | 2jsa-5 | THR_A_68_C | ALA_A_69_N | -4.451 | 8.379 | 9.607 | -4.144 | 9.417 | 8.844 | 1.32 | 125.18 | 73.52 |
| 69 | 2jsa-5 | ALA_A_69_C | PRO_A_70_N | -3.968 | 8.823 | 6.49 | -4.59 | 8.679 | 5.289 | 1.36 | 152.00 | -166.97 |
| 70 | 2jsa-5 | PRO_A_70_C | LYS_A_71_N | -2.847 | 8.811 | 3.562 | -2.689 | 10.017 | 4.091 | 1.33 | 66.49 | 82.54 |
| 71 | 2jsa-5 | LYS_A_71_C | ASP_A_72_N | -0.667 | 11.225 | 4.697 | -0.848 | 10.541 | 5.817 | 1.32 | 32.28 | -104.82 |
| 72 | 2jsa-5 | ASP_A_72_C | VAL_A_73_N | 0.804 | 9.423 | 7.211 | 0.102 | 8.313 | 7.028 | 1.33 | 97.93 | -122.31 |
| 73 | 2jsa-5 | VAL_A_73_C | CYS_A_74_N | 2.126 | 7.009 | 6.671 | 2.222 | 7.339 | 5.392 | 1.32 | 164.96 | 73.78 |
| 74 | 2jsa-5 | CYS_A_74_C | THR_A_75_N | 4.212 | 8.684 | 5.119 | 4.627 | 8.736 | 6.377 | 1.33 | 18.39 | 7.14 |
| 75 | 2jsa-5 | THR_A_75_C | LYS_A_76_N | 5.755 | 9.68 | 8.337 | 4.777 | 9.682 | 9.231 | 1.33 | 47.57 | 179.88 |
| 76 | 2jsa-5 | LYS_A_76_C | LEU_A_77_N | 5.301 | 7.985 | 10.895 | 4.495 | 7.164 | 10.237 | 1.33 | 119.77 | -134.47 |
| 77 | 2jsa-5 | LEU_A_77_C | ASN_A_78_N | 6.043 | 5.298 | 10.035 | 6.749 | 6.198 | 9.367 | 1.32 | 120.28 | 51.89 |
| 78 | 2jsa-5 | ASN_A_78_C | GLU_A_79_N | 8.124 | 5.015 | 7.745 | 7.451 | 5.475 | 6.7 | 1.33 | 142.04 | 145.65 |
| 79 | 2jsa-5 | GLU_A_79_C | CYS_A_80_N | 8.265 | 5.335 | 4.396 | 7.82 | 6.475 | 3.888 | 1.33 | 112.54 | 111.32 |
| 80 | 2jsa-5 | CYS_A_80_C | PRO_A_81_N | 9.983 | 7.436 | 3.388 | 10.043 | 7.948 | 4.647 | 1.36 | 22.27 | 83.32 |
| 1 | 2jsa-6 | ARG_A_1_C | SER_A_2_N | 13.524 | 3.618 | -1.098 | 12.724 | 4.398 | -1.81 | 1.32 | 122.51 | 135.73 |
| 2 | 2jsa-6 | SER_A_2_C | ALA_A_3_N | 11.665 | 6.481 | -2.525 | 10.501 | 5.857 | -2.625 | 1.32 | 94.33 | -151.81 |
| 3 | 2jsa-6 | ALA_A_3_C | LEU_A_4_N | 8.345 | 5.315 | -3.616 | 8.638 | 4.275 | -4.382 | 1.32 | 125.33 | -74.27 |
| 4 | 2jsa-6 | LEU_A_4_C | SER_A_5_N | 6.953 | 2.915 | -3.27 | 7.73 | 2.492 | -2.283 | 1.33 | 41.87 | -28.56 |
| 5 | 2jsa-6 | SER_A_5_C | CYS_A_6_N | 6.113 | 3.216 | -0.599 | 6.533 | 4.471 | -0.589 | 1.32 | 89.57 | 71.50 |
| 6 | 2jsa-6 | CYS_A_6_C | GLN_A_7_N | 4.391 | 5.451 | -1.137 | 4.601 | 5.698 | -2.423 | 1.33 | 165.85 | 49.63 |
| 7 | 2jsa-6 | GLN_A_7_C | MET_A_8_N | 2.605 | 4.44 | -3.076 | 3.215 | 3.265 | -3.041 | 1.32 | 88.49 | -62.56 |
| 8 | 2jsa-6 | MET_A_8_C | CYS_A_9_N | 1.532 | 2.238 | -1.572 | 1.933 | 3.129 | -0.677 | 1.33 | 47.51 | 65.77 |
| 9 | 2jsa-6 | CYS_A_9_C | GLU_A_10_N | -0.168 | 4.076 | 0.055 | -0.029 | 5.022 | -0.863 | 1.33 | 133.83 | 81.64 |
| 10 | 2jsa-6 | GLU_A_10_C | LEU_A_11_N | -2.077 | 4.788 | -2.183 | -1.563 | 3.587 | -2.405 | 1.33 | 99.64 | -66.83 |
| 11 | 2jsa-6 | LEU_A_11_C | VAL_A_12_N | -3.124 | 1.734 | -2.182 | -2.441 | 1.223 | -1.169 | 1.32 | 40.10 | -36.80 |
| 12 | 2jsa-6 | VAL_A_12_C | VAL_A_13_N | -4.149 | 1.225 | 0.565 | -3.738 | 2.415 | 0.976 | 1.32 | 71.92 | 70.95 |
| 13 | 2jsa-6 | VAL_A_13_C | LYS_A_14_N | -5.842 | 3.625 | 0.803 | -5.551 | 4.162 | -0.373 | 1.33 | 152.55 | 61.55 |
| 14 | 2jsa-6 | LYS_A_14_C | LYS_A_15_N | -7.631 | 3.382 | -1.368 | -7.132 | 2.163 | -1.232 | 1.32 | 84.11 | -67.74 |
| 15 | 2jsa-6 | LYS_A_15_C | TYR_A_16_N | -8.7 | 0.852 | 0.088 | -7.943 | 1.107 | 1.146 | 1.33 | 37.05 | 18.62 |
| 16 | 2jsa-6 | TYR_A_16_C | GLU_A_17_N | -9.348 | 2.228 | 2.823 | -8.681 | 3.358 | 3.006 | 1.32 | 82.06 | 59.45 |
| 17 | 2jsa-6 | GLU_A_17_C | GLY_A_18_N | -10.661 | 4.71 | 2.504 | -10.573 | 4.259 | 1.261 | 1.33 | 159.71 | -78.96 |
| 18 | 2jsa-6 | GLY_A_18_C | SER_A_19_N | -12.088 | 2.904 | -0.122 | -12.149 | 1.98 | 0.825 | 1.32 | 44.36 | -93.78 |
| 19 | 2jsa-6 | SER_A_19_C | ALA_A_20_N | -14.011 | 0.44 | 0.459 | -14.657 | 0.904 | 1.519 | 1.33 | 36.88 | 144.31 |
| 20 | 2jsa-6 | ALA_A_20_C | ASP_A_21_N | -16.495 | -0.564 | 2.145 | -15.497 | -1.432 | 2.237 | 1.33 | 86.02 | -41.01 |
| 21 | 2jsa-6 | ASP_A_21_C | LYS_A_22_N | -14.966 | -2.969 | 4.047 | -14.058 | -2.042 | 4.31 | 1.32 | 78.54 | 45.59 |
| 22 | 2jsa-6 | LYS_A_22_C | ASP_A_23_N | -12.689 | -3.501 | 5.695 | -11.776 | -3.843 | 4.798 | 1.32 | 132.62 | -20.54 |
| 23 | 2jsa-6 | ASP_A_23_C | ALA_A_24_N | -9.733 | -5.046 | 4.25 | -8.752 | -5.304 | 5.103 | 1.33 | 49.94 | -14.73 |
| 24 | 2jsa-6 | ALA_A_24_C | ASN_A_25_N | -7.093 | -6.347 | 3.66 | -8.078 | -7.22 | 3.505 | 1.33 | 96.72 | -138.45 |
| 25 | 2jsa-6 | ASN_A_25_C | VAL_A_26_N | -8.286 | -7.821 | 1.156 | -9.415 | -7.135 | 1.051 | 1.33 | 94.54 | 148.72 |
| 26 | 2jsa-6 | VAL_A_26_C | ILE_A_27_N | -9.03 | -5.35 | -0.556 | -8.267 | -4.904 | 0.432 | 1.33 | 41.81 | 30.31 |
| 27 | 2jsa-6 | ILE_A_27_C | LYS_A_28_N | -6.062 | -4.147 | -0.264 | -5.323 | -4.838 | 0.591 | 1.32 | 49.80 | -43.08 |
| 28 | 2jsa-6 | LYS_A_28_C | LYS_A_29_N | -4.045 | -5.925 | -1.171 | -5.082 | -6.717 | -1.396 | 1.32 | 99.78 | -142.63 |
| 29 | 2jsa-6 | LYS_A_29_C | ASP_A_30_N | -5.479 | -6.318 | -3.764 | -6.46 | -5.461 | -3.518 | 1.33 | 79.31 | 138.86 |
| 30 | 2jsa-6 | ASP_A_30_C | PHE_A_31_N | -5.492 | -3.72 | -4.915 | -4.733 | -3.249 | -3.937 | 1.32 | 42.41 | 31.82 |
| 31 | 2jsa-6 | PHE_A_31_C | ASP_A_32_N | -2.712 | -3.29 | -5.337 | -2.538 | -4.59 | -5.151 | 1.32 | 81.93 | -82.38 |
| 32 | 2jsa-6 | ASP_A_32_C | ALA_A_33_N | -2.541 | -5.299 | -7.479 | -3.864 | -5.271 | -7.412 | 1.32 | 87.10 | 178.79 |
| 33 | 2jsa-6 | ALA_A_33_C | GLU_A_34_N | -4.237 | -3.956 | -9.426 | -4.171 | -2.824 | -8.74 | 1.33 | 58.83 | 86.66 |
| 34 | 2jsa-6 | GLU_A_34_C | CYS_A_35_N | -2.279 | -1.616 | -9.717 | -1.472 | -1.699 | -8.671 | 1.32 | 37.80 | -5.87 |
| 35 | 2jsa-6 | CYS_A_35_C | LYS_A_36_N | 0.288 | -2.611 | -10.056 | -0.539 | -3.628 | -10.247 | 1.32 | 98.29 | -129.12 |
| 36 | 2jsa-6 | LYS_A_36_C | LYS_A_37_N | -0.438 | -3.755 | -12.676 | -1.664 | -3.431 | -13.061 | 1.33 | 106.89 | 165.20 |
| 37 | 2jsa-6 | LYS_A_37_C | LEU_A_38_N | -1.112 | -1.37 | -14.23 | -0.897 | -0.9 | -13.01 | 1.32 | 22.96 | 65.42 |
| 38 | 2jsa-6 | LEU_A_38_C | PHE_A_39_N | 1.283 | 0.157 | -13.243 | 1.925 | -0.837 | -12.648 | 1.32 | 63.31 | -57.14 |
| 39 | 2jsa-6 | PHE_A_39_C | HIS_A_40_N | 3.459 | -2.542 | -13.535 | 2.633 | -2.835 | -14.529 | 1.33 | 138.60 | -160.47 |
| 40 | 2jsa-6 | HIS_A_40_C | THR_A_41_N | 3.992 | -4.329 | -15.894 | 4.401 | -3.28 | -16.591 | 1.32 | 121.76 | 68.70 |
| 41 | 2jsa-6 | THR_A_41_C | ILE_A_42_N | 6.837 | -3.426 | -16.377 | 6.712 | -2.744 | -15.248 | 1.32 | 31.56 | 100.39 |
| 42 | 2jsa-6 | ILE_A_42_C | PRO_A_43_N | 8.156 | -4.204 | -13.943 | 9.487 | -4.411 | -13.755 | 1.36 | 82.05 | -8.84 |
| 43 | 2jsa-6 | PRO_A_43_C | PHE_A_44_N | 9.703 | -6.133 | -12.013 | 10.598 | -5.728 | -11.124 | 1.32 | 47.86 | 24.35 |
| 44 | 2jsa-6 | PHE_A_44_C | GLY_A_45_N | 9.277 | -5.244 | -9.107 | 8.651 | -4.435 | -9.949 | 1.32 | 129.46 | 127.73 |
| 45 | 2jsa-6 | GLY_A_45_C | THR_A_46_N | 6.434 | -4.493 | -8.889 | 5.887 | -5.379 | -9.709 | 1.33 | 128.22 | -121.69 |
| 46 | 2jsa-6 | THR_A_46_C | ARG_A_47_N | 5.341 | -7.149 | -8.104 | 6.63 | -7.454 | -8.159 | 1.33 | 92.38 | -13.31 |
| 47 | 2jsa-6 | ARG_A_47_C | GLU_A_48_N | 7.136 | -7.602 | -5.768 | 7.573 | -6.352 | -5.724 | 1.32 | 88.10 | 70.73 |
| 48 | 2jsa-6 | GLU_A_48_C | CYS_A_49_N | 6.08 | -5.265 | -4.118 | 5.282 | -5.004 | -5.143 | 1.32 | 140.68 | 161.89 |
| 49 | 2jsa-6 | CYS_A_49_C | ASP_A_50_N | 3.223 | -5.852 | -4.2 | 3.506 | -7.051 | -4.685 | 1.32 | 111.49 | -76.72 |
| 50 | 2jsa-6 | ASP_A_50_C | HIS_A_51_N | 3.617 | -8.492 | -2.729 | 4.753 | -7.838 | -2.538 | 1.32 | 81.71 | 29.93 |
| 51 | 2jsa-6 | HIS_A_51_C | TYR_A_52_N | 5.215 | -6.763 | -0.399 | 4.259 | -5.954 | -0.832 | 1.33 | 109.07 | 139.76 |
| 52 | 2jsa-6 | TYR_A_52_C | VAL_A_53_N | 2.519 | -4.942 | 0.581 | 1.646 | -5.654 | -0.118 | 1.33 | 121.82 | -140.80 |
| 53 | 2jsa-6 | VAL_A_53_C | ASN_A_54_N | 0.394 | -6.775 | 1.642 | 1.585 | -7.305 | 1.873 | 1.32 | 79.95 | -23.99 |
| 54 | 2jsa-6 | ASN_A_54_C | SER_A_55_N | 3.196 | -7.857 | 3.612 | 3.352 | -6.627 | 4.077 | 1.32 | 69.44 | 82.77 |
| 55 | 2jsa-6 | SER_A_55_C | LYS_A_56_N | 4.485 | -4.776 | 5.203 | 3.754 | -3.958 | 4.459 | 1.33 | 124.14 | 131.79 |
| 56 | 2jsa-6 | LYS_A_56_C | VAL_A_57_N | 2.065 | -2.223 | 4.701 | 1.249 | -3.263 | 4.618 | 1.32 | 93.59 | -128.12 |
| 57 | 2jsa-6 | VAL_A_57_C | ASP_A_58_N | -0.858 | -3.256 | 5.836 | -0.181 | -3.994 | 6.704 | 1.33 | 49.08 | -47.47 |
| 58 | 2jsa-6 | ASP_A_58_C | PRO_A_59_N | -0.46 | -3.014 | 8.914 | 0.765 | -2.442 | 8.778 | 1.36 | 95.74 | 25.03 |
| 59 | 2jsa-6 | PRO_A_59_C | ILE_A_60_N | 0.448 | -0.013 | 8.996 | 0.602 | 0.181 | 7.695 | 1.32 | 169.22 | 51.56 |
| 60 | 2jsa-6 | ILE_A_60_C | ILE_A_61_N | -1.468 | 1.432 | 7.427 | -2.129 | 0.284 | 7.421 | 1.32 | 90.26 | -119.93 |
| 61 | 2jsa-6 | ILE_A_61_C | HIS_A_62_N | -3.69 | 0.436 | 9.282 | -2.599 | 0.191 | 9.994 | 1.33 | 57.51 | -12.66 |
| 62 | 2jsa-6 | HIS_A_62_C | GLU_A_63_N | -2.652 | 1.83 | 11.796 | -2.249 | 2.652 | 10.839 | 1.32 | 136.27 | 63.88 |
| 63 | 2jsa-6 | GLU_A_63_C | LEU_A_64_N | -3.632 | 4.671 | 10.745 | -4.177 | 4.266 | 9.606 | 1.33 | 149.20 | -143.38 |
| 64 | 2jsa-6 | LEU_A_64_C | GLU_A_65_N | -6.543 | 4.204 | 10.172 | -6.308 | 2.988 | 10.642 | 1.32 | 69.22 | -79.06 |
| 65 | 2jsa-6 | GLU_A_65_C | GLY_A_66_N | -7.065 | 2.969 | 12.97 | -5.858 | 3.443 | 13.24 | 1.32 | 78.24 | 21.44 |
| 66 | 2jsa-6 | GLY_A_66_C | GLY_A_67_N | -6.125 | 5.472 | 14.601 | -5.907 | 6.219 | 13.528 | 1.33 | 144.05 | 73.73 |
| 67 | 2jsa-6 | GLY_A_67_C | THR_A_68_N | -5.53 | 8.422 | 12.504 | -4.322 | 7.934 | 12.265 | 1.32 | 100.40 | -22.00 |
| 68 | 2jsa-6 | THR_A_68_C | ALA_A_69_N | -4.004 | 8.747 | 9.976 | -3.665 | 9.846 | 9.317 | 1.33 | 119.81 | 72.86 |
| 69 | 2jsa-6 | ALA_A_69_C | PRO_A_70_N | -3.362 | 9.269 | 6.973 | -3.889 | 9.215 | 5.721 | 1.36 | 157.07 | -174.15 |
| 70 | 2jsa-6 | PRO_A_70_C | LYS_A_71_N | -1.997 | 9.204 | 4.15 | -1.83 | 10.422 | 4.643 | 1.32 | 68.15 | 82.19 |
| 71 | 2jsa-6 | LYS_A_71_C | ASP_A_72_N | 0.285 | 11.349 | 5.41 | -0.097 | 10.738 | 6.523 | 1.33 | 32.92 | -122.01 |
| 72 | 2jsa-6 | ASP_A_72_C | VAL_A_73_N | 1.438 | 9.406 | 7.861 | 0.745 | 8.365 | 7.421 | 1.33 | 109.38 | -123.65 |
| 73 | 2jsa-6 | VAL_A_73_C | CYS_A_74_N | 2.622 | 6.967 | 6.757 | 2.616 | 7.524 | 5.555 | 1.32 | 155.14 | 90.62 |
| 74 | 2jsa-6 | CYS_A_74_C | THR_A_75_N | 4.767 | 8.602 | 5.304 | 5.143 | 8.398 | 6.558 | 1.32 | 18.84 | -28.48 |
| 75 | 2jsa-6 | THR_A_75_C | LYS_A_76_N | 6.183 | 8.949 | 8.706 | 5.098 | 8.423 | 9.256 | 1.33 | 65.48 | -154.14 |
| 76 | 2jsa-6 | LYS_A_76_C | LEU_A_77_N | 5.955 | 6.778 | 10.831 | 5.764 | 5.827 | 9.927 | 1.33 | 132.98 | -101.36 |
| 77 | 2jsa-6 | LEU_A_77_C | ASN_A_78_N | 7.809 | 4.752 | 9.164 | 8.213 | 6.003 | 9.003 | 1.32 | 96.98 | 72.10 |
| 78 | 2jsa-6 | ASN_A_78_C | GLU_A_79_N | 9.331 | 5.677 | 6.866 | 8.106 | 5.358 | 6.474 | 1.33 | 107.21 | -165.40 |
| 79 | 2jsa-6 | GLU_A_79_C | CYS_A_80_N | 8.432 | 5.638 | 4.064 | 7.742 | 6.744 | 3.829 | 1.32 | 100.22 | 121.96 |

| | | | | | | | | | | | | |
|---|---|---|---|---|---|---|---|---|---|---|---|---|
| 80 | 2jsa-6 | CYS_A_80_C | PRO_A_81_N | 9.679 | 7.839 | 2.88 | 10.259 | 8.395 | 1.784 | 1.36 | 143.76 | 43.79 |
| 1 | 2jsa-7 | ARG_A_1_C | SER_A_2_N | 12.624 | 4.584 | -0.334 | 12.748 | 5.903 | -0.375 | 1.33 | 91.77 | 84.63 |
| 2 | 2jsa-7 | SER_A_2_C | ALA_A_3_N | 11.668 | 6.516 | -2.48 | 10.656 | 5.776 | -2.907 | 1.32 | 108.81 | -143.82 |
| 3 | 2jsa-7 | ALA_A_3_C | LEU_A_4_N | 9.063 | 4.984 | -4.568 | 8.962 | 3.988 | -5.436 | 1.33 | 130.93 | -95.79 |
| 4 | 2jsa-7 | LEU_A_4_C | SER_A_5_N | 7.067 | 2.772 | -4.515 | 7.907 | 2.623 | -3.501 | 1.33 | 40.08 | -10.06 |
| 5 | 2jsa-7 | SER_A_5_C | CYS_A_6_N | 6.459 | 2.949 | -1.559 | 6.828 | 4.218 | -1.467 | 1.32 | 86.02 | 73.79 |
| 6 | 2jsa-7 | CYS_A_6_C | GLN_A_7_N | 4.619 | 5.197 | -1.562 | 4.678 | 5.468 | -2.857 | 1.32 | 167.91 | 77.72 |
| 7 | 2jsa-7 | GLN_A_7_C | MET_A_8_N | 2.573 | 4.308 | -3.329 | 3.17 | 3.126 | -3.362 | 1.32 | 91.43 | -63.20 |
| 8 | 2jsa-7 | MET_A_8_C | CYS_A_9_N | 1.616 | 2.068 | -1.778 | 2.169 | 2.837 | -0.851 | 1.33 | 45.62 | 54.28 |
| 9 | 2jsa-7 | CYS_A_9_C | GLU_A_10_N | 0.191 | 3.816 | 0.138 | 0.254 | 4.722 | -0.828 | 1.33 | 136.77 | 86.02 |
| 10 | 2jsa-7 | GLU_A_10_C | LEU_A_11_N | -1.894 | 4.637 | -2.0 | -1.455 | 3.425 | -2.302 | 1.32 | 103.19 | -70.09 |
| 11 | 2jsa-7 | LEU_A_11_C | VAL_A_12_N | -3.07 | 1.616 | -2.088 | -2.4 | 1.227 | -1.012 | 1.33 | 35.75 | -30.14 |
| 12 | 2jsa-7 | VAL_A_12_C | VAL_A_13_N | -4.017 | 1.223 | 0.806 | -3.553 | 2.39 | 1.228 | 1.32 | 71.43 | 68.32 |
| 13 | 2jsa-7 | VAL_A_13_C | LYS_A_14_N | -5.57 | 3.749 | 1.158 | -5.244 | 4.393 | 0.047 | 1.32 | 146.99 | 63.15 |
| 14 | 2jsa-7 | LYS_A_14_C | LYS_A_15_N | -7.342 | 3.815 | -1.043 | -6.893 | 2.569 | -1.062 | 1.32 | 90.82 | -70.18 |
| 15 | 2jsa-7 | LYS_A_15_C | TYR_A_16_N | -8.522 | 1.18 | 0.093 | -7.776 | 1.324 | 1.178 | 1.32 | 35.00 | 10.93 |
| 16 | 2jsa-7 | TYR_A_16_C | GLU_A_17_N | -9.191 | 2.286 | 2.943 | -8.516 | 3.38 | 3.267 | 1.33 | 75.85 | 58.33 |
| 17 | 2jsa-7 | GLU_A_17_C | GLY_A_18_N | -10.446 | 4.825 | 2.841 | -10.351 | 4.397 | 1.591 | 1.32 | 160.67 | -77.49 |
| 18 | 2jsa-7 | GLY_A_18_C | SER_A_19_N | -11.927 | 3.208 | 0.125 | -11.972 | 2.237 | 1.025 | 1.32 | 47.20 | -92.65 |
| 19 | 2jsa-7 | SER_A_19_C | ALA_A_20_N | -13.93 | 0.849 | 0.566 | -14.572 | 1.258 | 1.649 | 1.32 | 35.10 | 147.50 |
| 20 | 2jsa-7 | ALA_A_20_C | ASP_A_21_N | -16.523 | -0.13 | 2.081 | -15.585 | -1.064 | 2.138 | 1.32 | 87.53 | -44.88 |
| 21 | 2jsa-7 | ASP_A_21_C | LYS_A_22_N | -15.285 | -2.78 | 3.837 | -14.302 | -1.977 | 4.217 | 1.32 | 73.33 | 39.24 |
| 22 | 2jsa-7 | LYS_A_22_C | ASP_A_23_N | -13.096 | -3.625 | 5.541 | -12.189 | -3.936 | 4.627 | 1.32 | 133.63 | -18.93 |
| 23 | 2jsa-7 | ASP_A_23_C | ALA_A_24_N | -10.166 | -5.17 | 4.077 | -9.243 | -5.375 | 5.005 | 1.32 | 45.53 | -12.52 |
| 24 | 2jsa-7 | ALA_A_24_C | ASN_A_25_N | -7.499 | -6.395 | 3.649 | -8.436 | -7.316 | 3.484 | 1.32 | 97.16 | -135.49 |
| 25 | 2jsa-7 | ASN_A_25_C | VAL_A_26_N | -8.562 | -7.898 | 1.125 | -9.573 | -7.046 | 1.04 | 1.32 | 93.68 | 139.88 |
| 26 | 2jsa-7 | VAL_A_26_C | ILE_A_27_N | -9.147 | -5.228 | -0.52 | -8.386 | -4.822 | 0.485 | 1.32 | 40.64 | 28.08 |
| 27 | 2jsa-7 | ILE_A_27_C | LYS_A_28_N | -6.159 | -4.057 | -0.128 | -5.515 | -4.901 | 0.666 | 1.33 | 53.21 | -52.66 |
| 28 | 2jsa-7 | LYS_A_28_C | LYS_A_29_N | -4.208 | -5.982 | -1.079 | -5.239 | -6.773 | -1.334 | 1.32 | 101.10 | -142.50 |
| 29 | 2jsa-7 | LYS_A_29_C | ASP_A_30_N | -5.659 | -6.346 | -3.693 | -6.593 | -5.462 | -3.374 | 1.32 | 76.07 | 136.58 |
| 30 | 2jsa-7 | ASP_A_30_C | PHE_A_31_N | -5.684 | -3.698 | -4.781 | -4.862 | -3.289 | -3.825 | 1.33 | 43.84 | 26.45 |
| 31 | 2jsa-7 | PHE_A_31_C | ASP_A_32_N | -2.897 | -3.313 | -5.302 | -2.64 | -4.598 | -5.106 | 1.33 | 81.49 | -78.69 |
| 32 | 2jsa-7 | ASP_A_32_C | ALA_A_33_N | -2.638 | -5.164 | -7.473 | -3.957 | -5.05 | -7.428 | 1.32 | 88.05 | 175.06 |
| 33 | 2jsa-7 | ALA_A_33_C | GLU_A_34_N | -4.127 | -3.669 | -9.423 | -4.042 | -2.528 | -8.756 | 1.32 | 59.76 | 85.74 |
| 34 | 2jsa-7 | GLU_A_34_C | CYS_A_35_N | -1.977 | -1.486 | -9.559 | -1.303 | -1.747 | -8.447 | 1.33 | 33.02 | -21.17 |
| 35 | 2jsa-7 | CYS_A_35_C | LYS_A_36_N | 0.515 | -2.692 | -9.733 | -0.399 | -3.556 | -10.151 | 1.33 | 108.38 | -136.61 |
| 36 | 2jsa-7 | LYS_A_36_C | LYS_A_37_N | -0.196 | -3.461 | -12.573 | -1.402 | -3.088 | -12.977 | 1.33 | 107.75 | 162.81 |
| 37 | 2jsa-7 | LYS_A_37_C | LEU_A_38_N | -0.815 | -0.919 | -13.912 | -0.479 | -0.676 | -12.653 | 1.33 | 18.23 | 35.87 |
| 38 | 2jsa-7 | LEU_A_38_C | PHE_A_39_N | 1.719 | 0.368 | -12.652 | 2.411 | -0.417 | -11.839 | 1.33 | 52.16 | -48.60 |
| 39 | 2jsa-7 | PHE_A_39_C | HIS_A_40_N | 4.037 | -1.93 | -12.894 | 3.076 | -2.192 | -13.768 | 1.33 | 131.27 | -164.75 |
| 40 | 2jsa-7 | HIS_A_40_C | THR_A_41_N | 4.519 | -3.428 | -15.295 | 4.832 | -2.375 | -16.036 | 1.33 | 124.00 | 73.45 |
| 41 | 2jsa-7 | THR_A_41_C | ILE_A_42_N | 7.176 | -1.694 | -15.82 | 7.577 | -2.477 | -14.829 | 1.33 | 41.60 | -62.88 |
| 42 | 2jsa-7 | ILE_A_42_C | PRO_A_43_N | 9.386 | -3.241 | -13.392 | 10.718 | -3.021 | -13.227 | 1.36 | 83.03 | 9.38 |
| 43 | 2jsa-7 | PRO_A_43_C | PHE_A_44_N | 11.418 | -4.326 | -11.263 | 10.886 | -5.497 | -10.948 | 1.32 | 76.24 | -114.43 |
| 44 | 2jsa-7 | PHE_A_44_C | GLY_A_45_N | 9.452 | -5.119 | -8.988 | 8.844 | -4.301 | -9.835 | 1.33 | 129.73 | 126.62 |
| 45 | 2jsa-7 | GLY_A_45_C | THR_A_46_N | 6.569 | -4.437 | -8.913 | 6.074 | -5.277 | -9.811 | 1.33 | 132.65 | -120.51 |
| 46 | 2jsa-7 | THR_A_46_C | ARG_A_47_N | 5.468 | -7.153 | -8.357 | 6.775 | -7.369 | -8.311 | 1.33 | 88.01 | -9.38 |
| 47 | 2jsa-7 | ARG_A_47_C | GLU_A_48_N | 7.206 | -7.64 | -5.916 | 7.497 | -6.35 | -5.835 | 1.32 | 86.49 | 77.29 |
| 48 | 2jsa-7 | GLU_A_48_C | CYS_A_49_N | 5.934 | -5.415 | -4.199 | 5.121 | -5.206 | -5.223 | 1.32 | 140.66 | 165.58 |
| 49 | 2jsa-7 | CYS_A_49_C | ASP_A_50_N | 3.16 | -6.097 | -4.125 | 3.546 | -7.319 | -4.462 | 1.33 | 104.73 | -72.47 |
| 50 | 2jsa-7 | ASP_A_50_C | HIS_A_51_N | 3.903 | -8.556 | -2.397 | 5.091 | -7.97 | -2.429 | 1.33 | 91.38 | 26.26 |
| 51 | 2jsa-7 | HIS_A_51_C | TYR_A_52_N | 5.656 | -6.733 | -0.405 | 4.53 | -6.102 | -0.707 | 1.33 | 103.17 | 150.73 |
| 52 | 2jsa-7 | TYR_A_52_C | VAL_A_53_N | 2.797 | -5.15 | 0.754 | 1.878 | -5.798 | 0.054 | 1.32 | 121.90 | -144.81 |
| 53 | 2jsa-7 | VAL_A_53_C | ASN_A_54_N | 0.688 | -7.302 | 1.553 | 1.616 | -7.199 | 2.493 | 1.32 | 44.81 | 6.33 |
| 54 | 2jsa-7 | ASN_A_54_C | SER_A_55_N | 3.035 | -7.924 | 4.332 | 3.1 | -6.657 | 4.714 | 1.32 | 73.24 | 87.06 |
| 55 | 2jsa-7 | SER_A_55_C | LYS_A_56_N | 4.045 | -4.679 | 5.794 | 3.649 | -3.98 | 4.74 | 1.33 | 142.68 | 119.53 |
| 56 | 2jsa-7 | LYS_A_56_C | VAL_A_57_N | 1.986 | -2.204 | 4.692 | 1.189 | -3.248 | 4.515 | 1.33 | 97.67 | -127.36 |
| 57 | 2jsa-7 | VAL_A_57_C | ASP_A_58_N | -0.921 | -3.372 | 5.72 | -0.227 | -4.15 | 6.537 | 1.32 | 51.92 | -48.27 |
| 58 | 2jsa-7 | ASP_A_58_C | PRO_A_59_N | -0.572 | -3.331 | 8.803 | 0.638 | -2.714 | 8.741 | 1.36 | 92.61 | 27.02 |
| 59 | 2jsa-7 | PRO_A_59_C | ILE_A_60_N | 0.235 | -0.315 | 9.097 | 0.412 | -0.042 | 7.813 | 1.32 | 165.78 | 57.04 |
| 60 | 2jsa-7 | ILE_A_60_C | ILE_A_61_N | -1.68 | 1.182 | 7.594 | -2.339 | 0.042 | 7.454 | 1.32 | 96.07 | -120.03 |
| 61 | 2jsa-7 | ILE_A_61_C | HIS_A_62_N | -3.926 | 0.103 | 9.299 | -2.858 | -0.204 | 10.021 | 1.33 | 56.99 | -16.04 |
| 62 | 2jsa-7 | HIS_A_62_C | GLU_A_63_N | -2.891 | 1.37 | 11.882 | -2.525 | 2.219 | 10.934 | 1.32 | 135.72 | 66.68 |
| 63 | 2jsa-7 | GLU_A_63_C | LEU_A_64_N | -3.873 | 4.262 | 10.962 | -4.442 | 3.956 | 9.804 | 1.33 | 150.84 | -151.73 |
| 64 | 2jsa-7 | LEU_A_64_C | GLU_A_65_N | -6.796 | 3.883 | 10.418 | -6.619 | 2.606 | 10.724 | 1.33 | 76.65 | -82.11 |
| 65 | 2jsa-7 | GLU_A_65_C | GLY_A_66_N | -7.459 | 2.537 | 13.022 | -6.249 | 2.917 | 13.407 | 1.33 | 73.11 | 17.43 |
| 66 | 2jsa-7 | GLY_A_66_C | GLY_A_67_N | -6.648 | 4.919 | 14.777 | -6.236 | 5.757 | 13.837 | 1.32 | 135.19 | 63.82 |
| 67 | 2jsa-7 | GLY_A_67_C | THR_A_68_N | -5.728 | 8.045 | 13.095 | -5.16 | 7.549 | 12.005 | 1.33 | 145.32 | -41.13 |
| 68 | 2jsa-7 | THR_A_68_C | ALA_A_69_N | -4.629 | 8.472 | 9.799 | -4.397 | 9.662 | 9.265 | 1.32 | 113.77 | 78.97 |
| 69 | 2jsa-7 | ALA_A_69_C | PRO_A_70_N | -3.892 | 9.127 | 6.946 | -4.356 | 9.028 | 5.671 | 1.36 | 159.59 | -167.96 |
| 70 | 2jsa-7 | PRO_A_70_C | LYS_A_71_N | -2.392 | 9.047 | 4.193 | -2.341 | 10.313 | 4.579 | 1.32 | 73.06 | 87.69 |
| 71 | 2jsa-7 | LYS_A_71_C | ASP_A_72_N | -0.275 | 11.309 | 5.394 | -0.665 | 10.691 | 6.499 | 1.32 | 33.48 | -122.25 |
| 72 | 2jsa-7 | ASP_A_72_C | VAL_A_73_N | 0.885 | 9.416 | 7.875 | 0.203 | 8.341 | 7.51 | 1.32 | 106.00 | -122.39 |
| 73 | 2jsa-7 | VAL_A_73_C | CYS_A_74_N | 2.15 | 7.005 | 6.917 | 2.138 | 7.409 | 5.656 | 1.32 | 162.23 | 91.70 |
| 74 | 2jsa-7 | CYS_A_74_C | THR_A_75_N | 4.143 | 8.703 | 5.273 | 4.604 | 8.701 | 6.515 | 1.32 | 20.36 | -0.25 |
| 75 | 2jsa-7 | THR_A_75_C | LYS_A_76_N | 6.089 | 9.429 | 8.322 | 5.297 | 9.06 | 9.317 | 1.32 | 41.29 | -155.02 |
| 76 | 2jsa-7 | LYS_A_76_C | LEU_A_77_N | 6.268 | 7.19 | 10.534 | 5.744 | 6.501 | 9.53 | 1.33 | 139.23 | -127.25 |
| 77 | 2jsa-7 | LEU_A_77_C | ASN_A_78_N | 7.383 | 4.968 | 8.591 | 7.899 | 6.112 | 8.167 | 1.32 | 108.67 | 65.72 |
| 78 | 2jsa-7 | ASN_A_78_C | GLU_A_79_N | 8.958 | 5.506 | 6.062 | 7.734 | 5.616 | 5.565 | 1.33 | 112.02 | 174.86 |
| 79 | 2jsa-7 | GLU_A_79_C | CYS_A_80_N | 6.872 | 6.152 | 3.337 | 7.662 | 7.209 | 3.21 | 1.33 | 95.50 | 53.23 |
| 80 | 2jsa-7 | CYS_A_80_C | PRO_A_81_N | 8.211 | 9.503 | 2.678 | 8.347 | 10.413 | 1.677 | 1.36 | 137.41 | 81.50 |
| 1 | 2jsa-8 | ARG_A_1_C | SER_A_2_N | 13.936 | 6.737 | 0.31 | 13.944 | 5.905 | -0.721 | 1.32 | 141.10 | -89.45 |
| 2 | 2jsa-8 | SER_A_2_C | ALA_A_3_N | 12.039 | 6.029 | -2.247 | 10.732 | 5.833 | -2.349 | 1.33 | 94.41 | -171.47 |
| 3 | 2jsa-8 | ALA_A_3_C | LEU_A_4_N | 8.64 | 5.77 | -3.589 | 8.78 | 4.569 | -4.131 | 1.33 | 114.14 | -83.35 |
| 4 | 2jsa-8 | LEU_A_4_C | SER_A_5_N | 6.948 | 3.349 | -3.096 | 7.732 | 3.395 | -2.029 | 1.32 | 36.35 | 3.36 |
| 5 | 2jsa-8 | SER_A_5_C | CYS_A_6_N | 6.038 | 3.954 | -0.358 | 6.121 | 5.194 | -0.818 | 1.33 | 110.31 | 86.17 |
| 6 | 2jsa-8 | CYS_A_6_C | GLN_A_7_N | 3.873 | 5.82 | -1.455 | 4.078 | 5.994 | -2.752 | 1.32 | 168.29 | 40.32 |
| 7 | 2jsa-8 | GLN_A_7_C | MET_A_8_N | 2.27 | 4.444 | -3.322 | 3.018 | 3.358 | -3.194 | 1.32 | 84.46 | -55.44 |
| 8 | 2jsa-8 | MET_A_8_C | CYS_A_9_N | 1.563 | 2.226 | -1.567 | 2.05 | 3.037 | -0.64 | 1.32 | 45.58 | 59.02 |
| 9 | 2jsa-8 | CYS_A_9_C | GLU_A_10_N | 0.034 | 4.006 | 0.28 | 0.127 | 4.904 | -0.69 | 1.33 | 137.05 | 84.09 |
| 10 | 2jsa-8 | GLU_A_10_C | LEU_A_11_N | -1.988 | 4.811 | -1.919 | -1.553 | 3.586 | -2.175 | 1.32 | 101.14 | -70.45 |
| 11 | 2jsa-8 | LEU_A_11_C | VAL_A_12_N | -3.195 | 1.8 | -1.983 | -2.523 | 1.274 | -0.969 | 1.33 | 40.08 | -38.05 |
| 12 | 2jsa-8 | VAL_A_12_C | VAL_A_13_N | -4.221 | 1.285 | 0.774 | -3.797 | 2.482 | 1.154 | 1.33 | 73.34 | 70.49 |
| 13 | 2jsa-8 | VAL_A_13_C | LYS_A_14_N | -5.848 | 3.779 | 0.986 | -5.516 | 4.359 | -0.158 | 1.32 | 149.71 | 60.21 |
| 14 | 2jsa-8 | LYS_A_14_C | LYS_A_15_N | -7.589 | 3.683 | -1.24 | -7.132 | 2.448 | -1.098 | 1.32 | 83.85 | -69.69 |
| 15 | 2jsa-8 | LYS_A_15_C | TYR_A_16_N | -8.752 | 1.111 | 0.129 | -8.009 | 1.304 | 1.209 | 1.33 | 35.40 | 14.56 |

| | | | | | | | | | | | | |
|---|---|---|---|---|---|---|---|---|---|---|---|---|
| 16 | 2jsa-8 | TYR_A_16_C | GLU_A_17_N | -9.514 | 2.313 | 2.871 | -8.93 | 3.497 | 2.98 | 1.32 | 85.28 | 63.75 |
| 17 | 2jsa-8 | GLU_A_17_C | GLY_A_18_N | -10.963 | 4.743 | 2.424 | -10.802 | 4.332 | 1.175 | 1.32 | 160.54 | -68.61 |
| 18 | 2jsa-8 | GLY_A_18_C | SER_A_19_N | -12.277 | 2.903 | -0.178 | -12.445 | 2.054 | 0.826 | 1.33 | 40.76 | -101.19 |
| 19 | 2jsa-8 | SER_A_19_C | ALA_A_20_N | -14.275 | 0.434 | 0.888 | -14.708 | 1.001 | 2.005 | 1.33 | 32.57 | 127.37 |
| 20 | 2jsa-8 | ALA_A_20_C | ASP_A_21_N | -16.376 | -0.598 | 2.767 | -15.305 | -1.375 | 2.83 | 1.32 | 87.27 | -35.96 |
| 21 | 2jsa-8 | ASP_A_21_C | LYS_A_22_N | -14.639 | -3.088 | 4.426 | -13.856 | -2.105 | 4.845 | 1.32 | 71.56 | 51.46 |
| 22 | 2jsa-8 | LYS_A_22_C | ASP_A_23_N | -12.42 | -3.643 | 6.067 | -11.933 | -4.054 | 4.906 | 1.32 | 151.24 | -40.16 |
| 23 | 2jsa-8 | ASP_A_23_C | ALA_A_24_N | -9.869 | -5.178 | 4.274 | -8.926 | -5.56 | 5.122 | 1.32 | 50.19 | -22.05 |
| 24 | 2jsa-8 | ALA_A_24_C | ASN_A_25_N | -7.163 | -6.523 | 3.749 | -8.149 | -7.353 | 3.443 | 1.32 | 103.36 | -139.91 |
| 25 | 2jsa-8 | ASN_A_25_C | VAL_A_26_N | -8.272 | -7.908 | 1.077 | -9.396 | -7.213 | 0.978 | 1.33 | 94.28 | 148.27 |
| 26 | 2jsa-8 | VAL_A_26_C | ILE_A_27_N | -9.114 | -5.326 | -0.533 | -8.399 | -4.883 | 0.49 | 1.32 | 39.43 | 31.78 |
| 27 | 2jsa-8 | ILE_A_27_C | LYS_A_28_N | -6.257 | -3.898 | -0.112 | -5.58 | -4.818 | 0.56 | 1.33 | 59.53 | -53.65 |
| 28 | 2jsa-8 | LYS_A_28_C | LYS_A_29_N | -4.241 | -5.807 | -1.215 | -5.327 | -6.51 | -1.499 | 1.32 | 102.38 | -147.08 |
| 29 | 2jsa-8 | LYS_A_29_C | ASP_A_30_N | -5.664 | -6.105 | -3.874 | -6.614 | -5.213 | -3.632 | 1.33 | 79.48 | 136.80 |
| 30 | 2jsa-8 | ASP_A_30_C | PHE_A_31_N | -5.591 | -3.466 | -4.982 | -4.807 | -3.117 | -3.973 | 1.32 | 40.38 | 24.00 |
| 31 | 2jsa-8 | PHE_A_31_C | ASP_A_32_N | -2.736 | -3.183 | -5.295 | -2.532 | -4.473 | -5.076 | 1.32 | 80.48 | -81.01 |
| 32 | 2jsa-8 | ASP_A_32_C | ALA_A_33_N | -2.443 | -5.176 | -7.404 | -3.709 | -4.785 | -7.394 | 1.33 | 89.57 | 162.84 |
| 33 | 2jsa-8 | ALA_A_33_C | GLU_A_34_N | -3.959 | -3.399 | -9.378 | -4.021 | -2.268 | -8.69 | 1.33 | 58.73 | 93.14 |
| 34 | 2jsa-8 | GLU_A_34_C | CYS_A_35_N | -2.067 | -0.86 | -9.127 | -1.417 | -1.978 | -8.84 | 1.32 | 77.49 | -59.83 |
| 35 | 2jsa-8 | CYS_A_35_C | LYS_A_36_N | 0.633 | -2.879 | -9.754 | -0.016 | -4.013 | -9.969 | 1.32 | 99.34 | -119.78 |
| 36 | 2jsa-8 | LYS_A_36_C | LYS_A_37_N | 0.376 | -4.316 | -12.352 | -0.823 | -3.871 | -12.701 | 1.33 | 105.26 | 159.64 |
| 37 | 2jsa-8 | LYS_A_37_C | LEU_A_38_N | -0.321 | -1.881 | -14.01 | -0.282 | -1.25 | -12.845 | 1.33 | 28.49 | 86.46 |
| 38 | 2jsa-8 | LEU_A_38_C | PHE_A_39_N | 1.84 | -0.088 | -13.107 | 2.524 | -0.992 | -12.421 | 1.33 | 58.82 | -52.89 |
| 39 | 2jsa-8 | PHE_A_39_C | HIS_A_40_N | 4.19 | -2.69 | -13.041 | 3.509 | -3.139 | -14.086 | 1.33 | 142.03 | -146.60 |
| 40 | 2jsa-8 | HIS_A_40_C | THR_A_41_N | 5.055 | -4.719 | -15.114 | 5.357 | -3.943 | -16.144 | 1.32 | 141.05 | 68.74 |
| 41 | 2jsa-8 | THR_A_41_C | ILE_A_42_N | 7.777 | -3.902 | -15.759 | 7.581 | -2.974 | -14.834 | 1.32 | 45.72 | 101.93 |
| 42 | 2jsa-8 | ILE_A_42_C | PRO_A_43_N | 9.152 | -4.08 | -13.341 | 10.486 | -4.061 | -13.073 | 1.36 | 78.64 | 0.82 |
| 43 | 2jsa-8 | PRO_A_43_C | PHE_A_44_N | 10.853 | -5.583 | -11.177 | 10.969 | -4.563 | -10.34 | 1.32 | 50.81 | 83.51 |
| 44 | 2jsa-8 | PHE_A_44_C | GLY_A_45_N | 9.376 | -4.04 | -8.541 | 8.46 | -4.016 | -9.498 | 1.32 | 136.24 | 178.50 |
| 45 | 2jsa-8 | GLY_A_45_C | THR_A_46_N | 6.175 | -4.404 | -8.676 | 5.694 | -5.295 | -9.53 | 1.32 | 130.15 | -118.36 |
| 46 | 2jsa-8 | THR_A_46_C | ARG_A_47_N | 5.339 | -7.143 | -7.961 | 6.644 | -7.351 | -8.041 | 1.32 | 93.46 | -9.06 |
| 47 | 2jsa-8 | ARG_A_47_C | GLU_A_48_N | 7.191 | -7.465 | -5.656 | 7.461 | -6.169 | -5.627 | 1.32 | 88.75 | 78.23 |
| 48 | 2jsa-8 | GLU_A_48_C | CYS_A_49_N | 5.895 | -5.27 | -3.974 | 5.043 | -5.098 | -4.974 | 1.32 | 139.00 | 168.59 |
| 49 | 2jsa-8 | CYS_A_49_C | ASP_A_50_N | 3.172 | -6.079 | -3.797 | 3.274 | -7.298 | -4.308 | 1.33 | 112.67 | -85.22 |
| 50 | 2jsa-8 | ASP_A_50_C | HIS_A_51_N | 3.648 | -8.476 | -2.212 | 4.825 | -7.867 | -2.238 | 1.33 | 91.12 | 27.36 |
| 51 | 2jsa-8 | HIS_A_51_C | TYR_A_52_N | 5.281 | -6.563 | -0.229 | 4.184 | -5.938 | -0.631 | 1.33 | 107.66 | 150.33 |
| 52 | 2jsa-8 | TYR_A_52_C | VAL_A_53_N | 2.356 | -5.01 | 0.727 | 1.453 | -5.6 | -0.042 | 1.32 | 125.49 | -146.84 |
| 53 | 2jsa-8 | VAL_A_53_C | ASN_A_54_N | 0.253 | -6.769 | 1.722 | 1.394 | -7.431 | 1.836 | 1.32 | 85.06 | -30.12 |
| 54 | 2jsa-8 | ASN_A_54_C | SER_A_55_N | 3.063 | -8.017 | 3.508 | 3.294 | -6.764 | 3.872 | 1.33 | 74.06 | 79.55 |
| 55 | 2jsa-8 | SER_A_55_C | LYS_A_56_N | 4.526 | -4.921 | 4.903 | 3.846 | -4.084 | 4.133 | 1.33 | 125.53 | 129.09 |
| 56 | 2jsa-8 | LYS_A_56_C | VAL_A_57_N | 2.215 | -2.324 | 4.541 | 1.389 | -3.359 | 4.573 | 1.32 | 88.62 | -128.59 |
| 57 | 2jsa-8 | VAL_A_57_C | ASP_A_58_N | -0.533 | -3.279 | 6.064 | -0.047 | -4.312 | 6.737 | 1.33 | 59.48 | -64.80 |
| 58 | 2jsa-8 | ASP_A_58_C | PRO_A_59_N | -0.198 | -3.261 | 8.927 | 0.979 | -2.625 | 8.684 | 1.36 | 100.29 | 28.38 |
| 59 | 2jsa-8 | PRO_A_59_C | ILE_A_60_N | 0.526 | -0.212 | 8.817 | 0.664 | -0.01 | 7.514 | 1.33 | 169.37 | 55.66 |
| 60 | 2jsa-8 | ILE_A_60_C | ILE_A_61_N | -1.488 | 1.107 | 7.315 | -2.172 | -0.006 | 7.091 | 1.33 | 99.73 | -121.57 |
| 61 | 2jsa-8 | ILE_A_61_C | HIS_A_62_N | -3.69 | 0.115 | 8.988 | -2.582 | -0.102 | 9.682 | 1.33 | 58.42 | -11.08 |
| 62 | 2jsa-8 | HIS_A_62_C | GLU_A_63_N | -2.777 | 1.558 | 11.456 | -1.968 | 2.402 | 10.834 | 1.32 | 118.01 | 46.21 |
| 63 | 2jsa-8 | GLU_A_63_C | LEU_A_64_N | -3.496 | 4.314 | 10.794 | -4.02 | 3.916 | 9.644 | 1.32 | 150.22 | -142.78 |
| 64 | 2jsa-8 | LEU_A_64_C | GLU_A_65_N | -6.355 | 3.774 | 10.312 | -6.097 | 2.554 | 10.763 | 1.33 | 70.12 | -78.06 |
| 65 | 2jsa-8 | GLU_A_65_C | GLY_A_66_N | -7.039 | 2.81 | 13.009 | -5.876 | 3.109 | 13.568 | 1.32 | 65.04 | 14.42 |
| 66 | 2jsa-8 | GLY_A_66_C | GLY_A_67_N | -6.505 | 5.281 | 14.533 | -6.209 | 5.879 | 13.387 | 1.33 | 149.79 | 63.67 |
| 67 | 2jsa-8 | GLY_A_67_C | THR_A_68_N | -5.743 | 8.108 | 12.463 | -4.931 | 7.564 | 11.568 | 1.33 | 132.48 | -33.82 |
| 68 | 2jsa-8 | THR_A_68_C | ALA_A_69_N | -4.193 | 8.559 | 9.454 | -3.946 | 9.778 | 8.998 | 1.32 | 110.13 | 78.55 |
| 69 | 2jsa-8 | ALA_A_69_C | PRO_A_70_N | -3.405 | 9.175 | 6.703 | -3.829 | 9.127 | 5.412 | 1.36 | 161.71 | -173.54 |
| 70 | 2jsa-8 | PRO_A_70_C | LYS_A_71_N | -1.825 | 8.858 | 4.013 | -1.618 | 10.112 | 4.387 | 1.32 | 73.60 | 80.63 |
| 71 | 2jsa-8 | LYS_A_71_C | ASP_A_72_N | 0.551 | 10.784 | 5.263 | -0.047 | 10.581 | 6.427 | 1.32 | 28.48 | -161.25 |
| 72 | 2jsa-8 | ASP_A_72_C | VAL_A_73_N | 1.382 | 9.172 | 7.804 | 0.735 | 8.173 | 7.223 | 1.32 | 116.02 | -122.93 |
| 73 | 2jsa-8 | VAL_A_73_C | CYS_A_74_N | 2.551 | 6.738 | 6.471 | 2.498 | 7.284 | 5.265 | 1.32 | 155.54 | 95.54 |
| 74 | 2jsa-8 | CYS_A_74_C | THR_A_75_N | 4.65 | 8.326 | 4.909 | 5.121 | 8.096 | 6.126 | 1.33 | 23.30 | -26.03 |
| 75 | 2jsa-8 | THR_A_75_C | LYS_A_76_N | 6.459 | 8.501 | 8.138 | 5.437 | 8.214 | 8.933 | 1.33 | 53.17 | -164.31 |
| 76 | 2jsa-8 | LYS_A_76_C | LEU_A_77_N | 6.463 | 6.429 | 10.231 | 6.062 | 5.558 | 9.316 | 1.33 | 133.66 | -114.72 |
| 77 | 2jsa-8 | LEU_A_77_C | ASN_A_78_N | 7.799 | 4.409 | 8.058 | 8.4 | 5.588 | 7.994 | 1.32 | 92.77 | 62.99 |
| 78 | 2jsa-8 | ASN_A_78_C | GLU_A_79_N | 9.009 | 5.243 | 5.664 | 7.706 | 5.258 | 5.426 | 1.32 | 100.35 | 179.34 |
| 79 | 2jsa-8 | GLU_A_79_C | CYS_A_80_N | 7.317 | 5.726 | 3.054 | 8.007 | 6.815 | 3.361 | 1.33 | 76.61 | 57.64 |
| 80 | 2jsa-8 | CYS_A_80_C | PRO_A_81_N | 9.366 | 8.756 | 2.88 | 10.274 | 9.118 | 1.935 | 1.36 | 134.03 | 21.74 |
| 1 | 2jsa-9 | ARG_A_1_C | SER_A_2_N | 12.882 | 2.023 | -2.079 | 12.587 | 3.144 | -2.721 | 1.33 | 118.98 | 104.74 |
| 2 | 2jsa-9 | SER_A_2_C | ALA_A_3_N | 12.181 | 5.545 | -2.923 | 10.919 | 5.283 | -3.231 | 1.33 | 103.44 | -168.27 |
| 3 | 2jsa-9 | ALA_A_3_C | LEU_A_4_N | 8.803 | 5.581 | -4.395 | 8.934 | 4.579 | -5.252 | 1.32 | 130.30 | -82.55 |
| 4 | 2jsa-9 | LEU_A_4_C | SER_A_5_N | 7.032 | 3.227 | -4.561 | 7.763 | 3.056 | -3.47 | 1.32 | 34.53 | -13.17 |
| 5 | 2jsa-9 | SER_A_5_C | CYS_A_6_N | 6.198 | 3.442 | -1.633 | 6.537 | 4.721 | -1.715 | 1.33 | 93.55 | 75.16 |
| 6 | 2jsa-9 | CYS_A_6_C | GLN_A_7_N | 4.32 | 5.679 | -1.832 | 4.341 | 5.816 | -3.15 | 1.33 | 174.00 | 81.29 |
| 7 | 2jsa-9 | GLN_A_7_C | MET_A_8_N | 2.255 | 4.579 | -3.484 | 2.888 | 3.416 | -3.431 | 1.33 | 87.71 | -61.44 |
| 8 | 2jsa-9 | MET_A_8_C | CYS_A_9_N | 1.375 | 2.442 | -1.756 | 1.95 | 3.223 | -0.855 | 1.32 | 47.11 | 53.64 |
| 9 | 2jsa-9 | CYS_A_9_C | GLU_A_10_N | -0.043 | 4.224 | 0.08 | -0.02 | 5.022 | -0.977 | 1.32 | 142.94 | 88.35 |
| 10 | 2jsa-9 | GLU_A_10_C | LEU_A_11_N | -2.154 | 4.809 | -2.159 | -1.664 | 3.604 | -2.414 | 1.33 | 101.09 | -67.87 |
| 11 | 2jsa-9 | LEU_A_11_C | VAL_A_12_N | -3.269 | 1.797 | -2.132 | -2.586 | 1.333 | -1.095 | 1.33 | 38.53 | -34.19 |
| 12 | 2jsa-9 | VAL_A_12_C | VAL_A_13_N | -4.29 | 1.395 | 0.642 | -3.856 | 2.565 | 1.086 | 1.32 | 70.41 | 69.65 |
| 13 | 2jsa-9 | VAL_A_13_C | LYS_A_14_N | -5.932 | 3.818 | 0.866 | -5.605 | 4.362 | -0.298 | 1.33 | 151.40 | 58.99 |
| 14 | 2jsa-9 | LYS_A_14_C | LYS_A_15_N | -7.658 | 3.609 | -1.365 | -7.176 | 2.388 | -1.185 | 1.32 | 82.19 | -68.46 |
| 15 | 2jsa-9 | LYS_A_15_C | TYR_A_16_N | -8.818 | 1.091 | 0.058 | -8.109 | 1.324 | 1.153 | 1.33 | 34.28 | 18.19 |
| 16 | 2jsa-9 | TYR_A_16_C | GLU_A_17_N | -9.616 | 2.444 | 2.742 | -8.981 | 3.603 | 2.843 | 1.33 | 85.63 | 61.28 |
| 17 | 2jsa-9 | GLU_A_17_C | GLY_A_18_N | -11.009 | 4.86 | 2.296 | -10.91 | 4.413 | 1.053 | 1.32 | 159.78 | -77.51 |
| 18 | 2jsa-9 | GLY_A_18_C | SER_A_19_N | -12.418 | 2.942 | -0.215 | -12.364 | 2.065 | 0.777 | 1.33 | 41.53 | -86.48 |
| 19 | 2jsa-9 | SER_A_19_C | ALA_A_20_N | -14.179 | 0.485 | 0.346 | -14.941 | 1.039 | 1.277 | 1.32 | 45.34 | 143.98 |
| 20 | 2jsa-9 | ALA_A_20_C | ASP_A_21_N | -16.845 | -0.366 | 1.844 | -15.879 | -1.247 | 2.062 | 1.33 | 80.53 | -42.37 |
| 21 | 2jsa-9 | ASP_A_21_C | LYS_A_22_N | -15.468 | -2.634 | 4.018 | -14.354 | -1.924 | 4.126 | 1.33 | 85.33 | 32.51 |
| 22 | 2jsa-9 | LYS_A_22_C | ASP_A_23_N | -13.008 | -3.316 | 5.599 | -12.163 | -3.739 | 4.672 | 1.32 | 134.45 | -26.59 |
| 23 | 2jsa-9 | ASP_A_23_C | ALA_A_24_N | -10.106 | -4.964 | 4.233 | -9.157 | -5.192 | 5.129 | 1.32 | 47.45 | -13.51 |
| 24 | 2jsa-9 | ALA_A_24_C | ASN_A_25_N | -7.464 | -6.243 | 3.731 | -8.442 | -7.121 | 3.56 | 1.33 | 97.41 | -138.08 |
| 25 | 2jsa-9 | ASN_A_25_C | VAL_A_26_N | -8.587 | -7.701 | 1.201 | -9.617 | -6.874 | 1.099 | 1.32 | 94.42 | 141.24 |
| 26 | 2jsa-9 | VAL_A_26_C | ILE_A_27_N | -9.136 | -5.076 | -0.469 | -8.363 | -4.689 | 0.535 | 1.32 | 40.73 | 26.59 |
| 27 | 2jsa-9 | ILE_A_27_C | LYS_A_28_N | -6.115 | -3.993 | -0.082 | -5.449 | -4.765 | 0.764 | 1.32 | 50.32 | -49.22 |
| 28 | 2jsa-9 | LYS_A_28_C | LYS_A_29_N | -4.167 | -5.91 | -0.959 | -5.255 | -6.618 | -1.221 | 1.33 | 101.41 | -146.95 |
| 29 | 2jsa-9 | LYS_A_29_C | ASP_A_30_N | -5.639 | -6.224 | -3.592 | -6.654 | -5.397 | -3.38 | 1.33 | 80.80 | 140.83 |
| 30 | 2jsa-9 | ASP_A_30_C | PHE_A_31_N | -5.676 | -3.681 | -4.804 | -4.887 | -3.241 | -3.834 | 1.33 | 42.96 | 29.15 |
| 31 | 2jsa-9 | PHE_A_31_C | ASP_A_32_N | -2.877 | -3.33 | -5.248 | -2.513 | -4.568 | -4.947 | 1.33 | 76.87 | -73.62 |

| | | | | | | | | | | | | |
|---|---|---|---|---|---|---|---|---|---|---|---|---|
| 32 | 2jsa-9 | ASP_A_32_C | ALA_A_33_N | -2.454 | -5.255 | -7.281 | -3.772 | -5.114 | -7.259 | 1.33 | 89.05 | 173.89 |
| 33 | 2jsa-9 | ALA_A_33_C | GLU_A_34_N | -4.05 | -3.732 | -9.243 | -4.135 | -2.599 | -8.562 | 1.32 | 59.06 | 94.29 |
| 34 | 2jsa-9 | GLU_A_34_C | CYS_A_35_N | -2.224 | -1.107 | -8.9 | -1.502 | -2.209 | -8.753 | 1.33 | 83.63 | -56.77 |
| 35 | 2jsa-9 | CYS_A_35_C | LYS_A_36_N | 0.644 | -2.888 | -9.636 | 0.27 | -4.148 | -9.798 | 1.32 | 97.03 | -106.53 |
| 36 | 2jsa-9 | LYS_A_36_C | LYS_A_37_N | 0.484 | -4.427 | -12.206 | -0.748 | -3.946 | -12.294 | 1.33 | 93.81 | 158.67 |
| 37 | 2jsa-9 | LYS_A_37_C | LEU_A_38_N | -0.647 | -1.998 | -13.749 | -0.61 | -1.235 | -12.666 | 1.33 | 35.20 | 87.22 |
| 38 | 2jsa-9 | LEU_A_38_C | PHE_A_39_N | 1.45 | 0.037 | -12.91 | 2.101 | -0.636 | -11.972 | 1.33 | 44.95 | -45.95 |
| 39 | 2jsa-9 | PHE_A_39_C | HIS_A_40_N | 3.937 | -2.01 | -12.859 | 3.041 | -2.384 | -13.761 | 1.33 | 132.89 | -157.34 |
| 40 | 2jsa-9 | HIS_A_40_C | THR_A_41_N | 4.568 | -3.311 | -15.421 | 4.981 | -2.054 | -15.483 | 1.32 | 92.68 | 71.81 |
| 41 | 2jsa-9 | THR_A_41_C | ILE_A_42_N | 7.408 | -1.78 | -15.287 | 7.333 | -1.038 | -14.191 | 1.33 | 34.23 | 95.77 |
| 42 | 2jsa-9 | ILE_A_42_C | PRO_A_43_N | 9.151 | -2.36 | -13.261 | 10.505 | -2.277 | -13.159 | 1.36 | 85.70 | 3.51 |
| 43 | 2jsa-9 | PRO_A_43_C | PHE_A_44_N | 11.242 | -4.222 | -11.847 | 10.463 | -5.294 | -11.86 | 1.33 | 90.56 | -126.01 |
| 44 | 2jsa-9 | PHE_A_44_C | GLY_A_45_N | 9.353 | -5.411 | -9.669 | 8.93 | -4.211 | -10.038 | 1.32 | 106.17 | 109.42 |
| 45 | 2jsa-9 | GLY_A_45_C | THR_A_46_N | 6.903 | -4.309 | -8.65 | 6.366 | -5.139 | -9.532 | 1.32 | 131.74 | -122.90 |
| 46 | 2jsa-9 | THR_A_46_C | ARG_A_47_N | 5.643 | -6.82 | -7.903 | 6.848 | -7.372 | -7.921 | 1.33 | 90.78 | -24.61 |
| 47 | 2jsa-9 | ARG_A_47_C | GLU_A_48_N | 7.191 | -7.358 | -5.496 | 7.466 | -6.067 | -5.6 | 1.32 | 94.51 | 77.97 |
| 48 | 2jsa-9 | GLU_A_48_C | CYS_A_49_N | 5.911 | -4.943 | -4.078 | 5.079 | -4.872 | -5.107 | 1.33 | 140.94 | 175.12 |
| 49 | 2jsa-9 | CYS_A_49_C | ASP_A_50_N | 3.126 | -5.726 | -3.965 | 3.304 | -6.97 | -4.385 | 1.32 | 108.48 | -81.86 |
| 50 | 2jsa-9 | ASP_A_50_C | HIS_A_51_N | 3.554 | -8.083 | -2.235 | 4.754 | -7.52 | -2.234 | 1.33 | 89.96 | 25.13 |
| 51 | 2jsa-9 | HIS_A_51_C | TYR_A_52_N | 5.064 | -6.242 | -0.181 | 4.076 | -5.54 | -0.718 | 1.33 | 113.90 | 144.61 |
| 52 | 2jsa-9 | TYR_A_52_C | VAL_A_53_N | 2.193 | -4.725 | 0.636 | 1.493 | -5.679 | 0.039 | 1.33 | 116.77 | -126.27 |
| 53 | 2jsa-9 | VAL_A_53_C | ASN_A_54_N | 0.427 | -7.257 | 1.555 | 1.538 | -7.175 | 2.272 | 1.32 | 57.23 | 4.22 |
| 54 | 2jsa-9 | ASN_A_54_C | SER_A_55_N | 3.193 | -7.829 | 3.932 | 3.282 | -6.571 | 4.339 | 1.33 | 72.11 | 85.95 |
| 55 | 2jsa-9 | SER_A_55_C | LYS_A_56_N | 4.419 | -4.558 | 5.13 | 3.765 | -3.939 | 4.158 | 1.33 | 137.19 | 136.57 |
| 56 | 2jsa-9 | LYS_A_56_C | VAL_A_57_N | 2.15 | -2.122 | 4.259 | 1.312 | -3.148 | 4.223 | 1.33 | 91.56 | -129.24 |
| 57 | 2jsa-9 | VAL_A_57_C | ASP_A_58_N | -0.573 | -3.172 | 5.761 | -0.108 | -4.277 | 6.326 | 1.33 | 64.77 | -67.18 |
| 58 | 2jsa-9 | ASP_A_58_C | PRO_A_59_N | -0.158 | -3.427 | 8.605 | 1.028 | -2.803 | 8.37 | 1.36 | 99.95 | 27.75 |
| 59 | 2jsa-9 | PRO_A_59_C | ILE_A_60_N | 0.66 | -0.402 | 8.751 | 0.741 | -0.087 | 7.465 | 1.33 | 165.81 | 75.58 |
| 60 | 2jsa-9 | ILE_A_60_C | ILE_A_61_N | -1.377 | 1.111 | 7.483 | -2.112 | 0.046 | 7.198 | 1.33 | 102.42 | -124.61 |
| 61 | 2jsa-9 | ILE_A_61_C | HIS_A_62_N | -3.522 | 0.009 | 9.181 | -2.386 | -0.318 | 9.782 | 1.33 | 63.05 | -16.06 |
| 62 | 2jsa-9 | HIS_A_62_C | GLU_A_63_N | -2.428 | 1.136 | 11.738 | -2.003 | 2.074 | 10.903 | 1.33 | 129.04 | 65.63 |
| 63 | 2jsa-9 | GLU_A_63_C | LEU_A_64_N | -3.483 | 4.024 | 10.94 | -3.863 | 3.898 | 9.677 | 1.32 | 162.41 | -161.66 |
| 64 | 2jsa-9 | LEU_A_64_C | GLU_A_65_N | -6.246 | 3.801 | 10.157 | -6.138 | 2.506 | 10.415 | 1.32 | 78.77 | -85.23 |
| 65 | 2jsa-9 | GLU_A_65_C | GLY_A_66_N | -7.063 | 2.436 | 12.681 | -5.859 | 2.775 | 13.117 | 1.32 | 70.78 | 15.73 |
| 66 | 2jsa-9 | GLY_A_66_C | GLY_A_67_N | -6.346 | 4.732 | 14.523 | -6.451 | 5.379 | 13.372 | 1.32 | 150.34 | 99.22 |
| 67 | 2jsa-9 | GLY_A_67_C | THR_A_68_N | -6.156 | 7.69 | 12.583 | -4.986 | 7.201 | 12.199 | 1.32 | 106.85 | -22.68 |
| 68 | 2jsa-9 | THR_A_68_C | ALA_A_69_N | -4.508 | 8.344 | 10.085 | -3.97 | 9.42 | 9.531 | 1.32 | 114.73 | 63.43 |
| 69 | 2jsa-9 | ALA_A_69_C | PRO_A_70_N | -3.471 | 9.094 | 7.172 | -3.93 | 9.123 | 5.893 | 1.36 | 160.22 | 176.38 |
| 70 | 2jsa-9 | PRO_A_70_C | LYS_A_71_N | -1.96 | 9.256 | 4.428 | -1.779 | 10.383 | 5.102 | 1.33 | 59.44 | 80.88 |
| 71 | 2jsa-9 | LYS_A_71_C | ASP_A_72_N | 0.319 | 11.197 | 6.028 | -0.144 | 10.559 | 7.093 | 1.33 | 36.51 | -125.97 |
| 72 | 2jsa-9 | ASP_A_72_C | VAL_A_73_N | 1.312 | 9.057 | 8.338 | 0.59 | 8.08 | 7.811 | 1.32 | 113.45 | -126.46 |
| 73 | 2jsa-9 | VAL_A_73_C | CYS_A_74_N | 2.397 | 6.675 | 6.987 | 2.339 | 7.217 | 5.78 | 1.32 | 155.70 | 96.11 |
| 74 | 2jsa-9 | CYS_A_74_C | THR_A_75_N | 4.438 | 8.36 | 5.408 | 5.042 | 8.116 | 6.562 | 1.33 | 29.44 | -22.00 |
| 75 | 2jsa-9 | THR_A_75_C | LYS_A_76_N | 6.414 | 8.613 | 8.53 | 5.471 | 8.054 | 9.274 | 1.32 | 55.84 | -149.34 |
| 76 | 2jsa-9 | LYS_A_76_C | LEU_A_77_N | 6.536 | 6.243 | 10.501 | 6.037 | 5.371 | 9.64 | 1.32 | 130.60 | -119.78 |
| 77 | 2jsa-9 | LEU_A_77_C | ASN_A_78_N | 7.753 | 4.208 | 8.365 | 8.152 | 5.447 | 8.12 | 1.32 | 100.66 | 72.15 |
| 78 | 2jsa-9 | ASN_A_78_C | GLU_A_79_N | 8.832 | 4.996 | 5.828 | 7.546 | 5.007 | 5.507 | 1.33 | 104.01 | 179.51 |
| 79 | 2jsa-9 | GLU_A_79_C | CYS_A_80_N | 6.55 | 5.398 | 3.304 | 7.471 | 6.219 | 2.82 | 1.33 | 111.42 | 41.71 |
| 80 | 2jsa-9 | CYS_A_80_C | PRO_A_81_N | 8.39 | 8.036 | 1.516 | 9.359 | 7.309 | 0.899 | 1.36 | 116.99 | -36.88 |
| 1 | 2jsa-10 | ARG_A_1_C | SER_A_2_N | 14.79 | 3.756 | 0.449 | 13.59 | 4.266 | 0.68 | 1.32 | 79.95 | 156.97 |
| 2 | 2jsa-10 | SER_A_2_C | ALA_A_3_N | 13.051 | 5.283 | -1.475 | 11.816 | 5.134 | -1.93 | 1.32 | 110.09 | -173.12 |
| 3 | 2jsa-10 | ALA_A_3_C | LEU_A_4_N | 10.067 | 5.114 | -3.622 | 9.602 | 4.445 | -4.668 | 1.33 | 142.08 | -124.80 |
| 4 | 2jsa-10 | LEU_A_4_C | SER_A_5_N | 7.344 | 3.87 | -3.969 | 8.02 | 3.197 | -3.049 | 1.33 | 46.04 | -44.87 |
| 5 | 2jsa-10 | SER_A_5_C | CYS_A_6_N | 6.257 | 3.433 | -1.373 | 6.551 | 4.725 | -1.386 | 1.33 | 90.56 | 77.18 |
| 6 | 2jsa-10 | CYS_A_6_C | GLN_A_7_N | 4.295 | 5.569 | -1.606 | 4.348 | 5.861 | -2.898 | 1.33 | 167.06 | 79.71 |
| 7 | 2jsa-10 | GLN_A_7_C | MET_A_8_N | 2.337 | 4.542 | -3.355 | 3.007 | 3.399 | -3.344 | 1.32 | 89.52 | -59.62 |
| 8 | 2jsa-10 | MET_A_8_C | CYS_A_9_N | 1.467 | 2.316 | -1.763 | 1.948 | 3.132 | -0.838 | 1.32 | 45.68 | 59.48 |
| 9 | 2jsa-10 | CYS_A_9_C | GLU_A_10_N | -0.111 | 4.055 | 0.032 | -0.035 | 4.977 | -0.917 | 1.33 | 135.73 | 85.29 |
| 10 | 2jsa-10 | GLU_A_10_C | LEU_A_11_N | -2.149 | 4.757 | -2.13 | -1.663 | 3.547 | -2.359 | 1.32 | 99.96 | -68.12 |
| 11 | 2jsa-10 | LEU_A_11_C | VAL_A_12_N | -3.194 | 1.677 | -2.077 | -2.512 | 1.358 | -0.987 | 1.32 | 34.63 | -25.07 |
| 12 | 2jsa-10 | VAL_A_12_C | VAL_A_13_N | -4.117 | 1.363 | 0.842 | -3.719 | 2.589 | 1.147 | 1.32 | 76.69 | 72.01 |
| 13 | 2jsa-10 | VAL_A_13_C | LYS_A_14_N | -5.811 | 3.828 | 1.058 | -5.556 | 4.43 | -0.096 | 1.33 | 150.47 | 67.04 |
| 14 | 2jsa-10 | LYS_A_14_C | LYS_A_15_N | -7.62 | 3.648 | -1.121 | -7.081 | 2.441 | -1.024 | 1.33 | 85.80 | -65.94 |
| 15 | 2jsa-10 | LYS_A_15_C | TYR_A_16_N | -8.58 | 0.983 | 0.221 | -7.858 | 1.304 | 1.285 | 1.33 | 36.60 | 23.97 |
| 16 | 2jsa-10 | TYR_A_16_C | GLU_A_17_N | -9.362 | 2.22 | 3.002 | -8.817 | 3.414 | 3.18 | 1.32 | 82.28 | 65.47 |
| 17 | 2jsa-10 | GLU_A_17_C | GLY_A_18_N | -10.677 | 4.831 | 2.453 | -10.437 | 4.247 | 1.288 | 1.33 | 151.54 | -67.66 |
| 18 | 2jsa-10 | GLY_A_18_C | SER_A_19_N | -11.871 | 3.063 | -0.319 | -11.979 | 2.134 | 0.621 | 1.33 | 44.86 | -96.63 |
| 19 | 2jsa-10 | SER_A_19_C | ALA_A_20_N | -13.996 | 0.798 | 0.281 | -14.574 | 1.256 | 1.381 | 1.32 | 33.84 | 141.61 |
| 20 | 2jsa-10 | ALA_A_20_C | ASP_A_21_N | -16.542 | -0.053 | 1.964 | -15.629 | -1.012 | 2.018 | 1.33 | 87.66 | -46.41 |
| 21 | 2jsa-10 | ASP_A_21_C | LYS_A_22_N | -15.362 | -2.631 | 3.815 | -14.323 | -1.87 | 4.124 | 1.32 | 76.51 | 36.22 |
| 22 | 2jsa-10 | LYS_A_22_C | ASP_A_23_N | -13.112 | -3.452 | 5.522 | -12.173 | -3.784 | 4.647 | 1.33 | 131.30 | -19.47 |
| 23 | 2jsa-10 | ASP_A_23_C | ALA_A_24_N | -10.135 | -5.022 | 4.165 | -9.218 | -5.228 | 5.1 | 1.33 | 45.15 | -12.66 |
| 24 | 2jsa-10 | ALA_A_24_C | ASN_A_25_N | -7.475 | -6.244 | 3.739 | -8.43 | -7.141 | 3.54 | 1.33 | 98.64 | -136.79 |
| 25 | 2jsa-10 | ASN_A_25_C | VAL_A_26_N | -8.484 | -7.67 | 1.166 | -9.652 | -7.074 | 0.976 | 1.32 | 98.24 | 152.97 |
| 26 | 2jsa-10 | VAL_A_26_C | ILE_A_27_N | -9.187 | -5.237 | -0.55 | -8.433 | -4.859 | 0.472 | 1.33 | 39.53 | 26.63 |
| 27 | 2jsa-10 | ILE_A_27_C | LYS_A_28_N | -6.234 | -4.045 | -0.18 | -5.493 | -4.811 | 0.606 | 1.32 | 53.59 | -45.95 |
| 28 | 2jsa-10 | LYS_A_28_C | LYS_A_29_N | -4.253 | -5.873 | -1.2 | -5.341 | -6.594 | -1.424 | 1.33 | 99.74 | -146.47 |
| 29 | 2jsa-10 | LYS_A_29_C | ASP_A_30_N | -5.762 | -6.187 | -3.786 | -6.633 | -5.236 | -3.478 | 1.33 | 76.57 | 132.49 |
| 30 | 2jsa-10 | ASP_A_30_C | PHE_A_31_N | -5.618 | -3.553 | -4.914 | -4.782 | -3.162 | -3.964 | 1.32 | 44.17 | 25.07 |
| 31 | 2jsa-10 | PHE_A_31_C | ASP_A_32_N | -2.814 | -3.333 | -5.427 | -2.598 | -4.618 | -5.181 | 1.33 | 79.31 | -80.46 |
| 32 | 2jsa-10 | ASP_A_32_C | ALA_A_33_N | -2.62 | -5.261 | -7.529 | -3.895 | -4.903 | -7.48 | 1.33 | 87.88 | 164.32 |
| 33 | 2jsa-10 | ALA_A_33_C | GLU_A_34_N | -4.107 | -3.428 | -9.403 | -4.195 | -2.303 | -8.707 | 1.33 | 58.33 | 94.47 |
| 34 | 2jsa-10 | GLU_A_34_C | CYS_A_35_N | -2.207 | -0.91 | -9.029 | -1.554 | -2.057 | -8.901 | 1.33 | 84.46 | -60.35 |
| 35 | 2jsa-10 | CYS_A_35_C | LYS_A_36_N | 0.522 | -2.922 | -9.788 | -0.061 | -4.091 | -10.011 | 1.33 | 99.69 | -116.51 |
| 36 | 2jsa-10 | LYS_A_36_C | LYS_A_37_N | 0.294 | -4.338 | -12.405 | -0.815 | -3.631 | -12.572 | 1.33 | 97.24 | 147.48 |
| 37 | 2jsa-10 | LYS_A_37_C | LEU_A_38_N | -0.391 | -1.588 | -13.826 | -0.376 | -0.97 | -12.655 | 1.32 | 27.83 | 88.61 |
| 38 | 2jsa-10 | LEU_A_38_C | PHE_A_39_N | 1.758 | 0.199 | -12.73 | 2.399 | -0.516 | -11.816 | 1.33 | 46.41 | -48.12 |
| 39 | 2jsa-10 | PHE_A_39_C | HIS_A_40_N | 4.156 | -1.975 | -12.727 | 3.248 | -2.292 | -13.639 | 1.33 | 133.48 | -160.75 |
| 40 | 2jsa-10 | HIS_A_40_C | THR_A_41_N | 4.763 | -3.356 | -15.226 | 4.974 | -2.21 | -15.857 | 1.33 | 118.44 | 79.57 |
| 41 | 2jsa-10 | THR_A_41_C | ILE_A_42_N | 7.418 | -2.196 | -15.688 | 7.34 | -1.617 | -14.498 | 1.33 | 26.15 | 97.67 |
| 42 | 2jsa-10 | ILE_A_42_C | PRO_A_43_N | 8.906 | -3.192 | -13.502 | 10.247 | -3.344 | -13.339 | 1.36 | 83.11 | -6.47 |
| 43 | 2jsa-10 | PRO_A_43_C | PHE_A_44_N | 10.574 | -5.345 | -11.948 | 10.913 | -4.636 | -10.882 | 1.32 | 36.40 | 64.45 |
| 44 | 2jsa-10 | PHE_A_44_C | GLY_A_45_N | 9.539 | -4.451 | -8.85 | 8.558 | -4.074 | -9.657 | 1.33 | 127.52 | 158.98 |
| 45 | 2jsa-10 | GLY_A_45_C | THR_A_46_N | 6.341 | -4.396 | -8.645 | 5.967 | -5.302 | -9.536 | 1.32 | 132.27 | -112.43 |
| 46 | 2jsa-10 | THR_A_46_C | ARG_A_47_N | 5.471 | -7.169 | -8.03 | 6.751 | -7.507 | -8.078 | 1.32 | 92.08 | -14.79 |
| 47 | 2jsa-10 | ARG_A_47_C | GLU_A_48_N | 7.239 | -7.603 | -5.681 | 7.502 | -6.305 | -5.709 | 1.32 | 91.21 | 78.55 |

| | | | | | | | | | | | |
|---|---|---|---|---|---|---|---|---|---|---|---|
| 48 | 2jsa-10 | GLU_A_48_C | CYS_A_49_N | 5.977 | -5.281 | -4.091 | 5.107 | -5.286 | -5.09 | 1.32 | 138.95 | -179.67 |
| 49 | 2jsa-10 | CYS_A_49_C | ASP_A_50_N | 3.227 | -6.129 | -3.825 | 3.273 | -7.386 | -4.243 | 1.33 | 108.38 | -87.90 |
| 50 | 2jsa-10 | ASP_A_50_C | HIS_A_51_N | 3.692 | -8.441 | -2.091 | 4.887 | -7.881 | -2.211 | 1.33 | 95.20 | 25.11 |
| 51 | 2jsa-10 | HIS_A_51_C | TYR_A_52_N | 5.402 | -6.559 | -0.229 | 4.367 | -5.865 | -0.682 | 1.33 | 109.98 | 146.16 |
| 52 | 2jsa-10 | TYR_A_52_C | VAL_A_53_N | 2.509 | -4.922 | 0.622 | 1.617 | -5.439 | -0.208 | 1.32 | 128.84 | -149.90 |
| 53 | 2jsa-10 | VAL_A_53_C | ASN_A_54_N | 0.305 | -6.581 | 1.493 | 1.436 | -7.245 | 1.679 | 1.32 | 81.93 | -30.42 |
| 54 | 2jsa-10 | ASN_A_54_C | SER_A_55_N | 3.025 | -7.892 | 3.405 | 3.219 | -6.727 | 4.002 | 1.32 | 63.18 | 80.55 |
| 55 | 2jsa-10 | SER_A_55_C | LYS_A_56_N | 4.525 | -4.917 | 4.998 | 3.82 | -4.11 | 4.22 | 1.32 | 125.98 | 131.14 |
| 56 | 2jsa-10 | LYS_A_56_C | VAL_A_57_N | 2.282 | -2.25 | 4.531 | 1.403 | -3.233 | 4.398 | 1.33 | 95.76 | -131.80 |
| 57 | 2jsa-10 | VAL_A_57_C | ASP_A_58_N | -0.569 | -3.229 | 5.824 | -0.121 | -4.335 | 6.401 | 1.33 | 64.19 | -67.95 |
| 58 | 2jsa-10 | ASP_A_58_C | PRO_A_59_N | -0.281 | -3.55 | 8.698 | 0.934 | -2.957 | 8.557 | 1.36 | 95.95 | 26.02 |
| 59 | 2jsa-10 | PRO_A_59_C | ILE_A_60_N | 0.613 | -0.557 | 8.984 | 0.81 | -0.193 | 7.725 | 1.33 | 161.80 | 61.58 |
| 60 | 2jsa-10 | ILE_A_60_C | ILE_A_61_N | -1.279 | 1.051 | 7.607 | -1.981 | -0.029 | 7.294 | 1.33 | 103.66 | -123.02 |
| 61 | 2jsa-10 | ILE_A_61_C | HIS_A_62_N | -3.556 | -0.056 | 9.148 | -2.489 | -0.416 | 9.845 | 1.32 | 58.24 | -18.64 |
| 62 | 2jsa-10 | HIS_A_62_C | GLU_A_63_N | -2.676 | 1.041 | 11.791 | -1.896 | 1.933 | 11.199 | 1.32 | 116.55 | 48.83 |
| 63 | 2jsa-10 | GLU_A_63_C | LEU_A_64_N | -3.355 | 3.897 | 11.311 | -3.826 | 3.687 | 10.09 | 1.33 | 157.10 | -155.97 |
| 64 | 2jsa-10 | LEU_A_64_C | GLU_A_65_N | -6.191 | 3.572 | 10.651 | -5.946 | 2.334 | 11.054 | 1.32 | 72.29 | -78.81 |
| 65 | 2jsa-10 | GLU_A_65_C | GLY_A_66_N | -6.891 | 2.349 | 13.312 | -5.806 | 3.048 | 13.61 | 1.32 | 77.00 | 32.79 |
| 66 | 2jsa-10 | GLY_A_66_C | GLY_A_67_N | -6.346 | 5.152 | 14.763 | -6.413 | 5.648 | 13.537 | 1.32 | 157.79 | 97.69 |
| 67 | 2jsa-10 | GLY_A_67_C | THR_A_68_N | -6.028 | 7.857 | 12.53 | -4.877 | 7.29 | 12.2 | 1.32 | 104.42 | -26.23 |
| 68 | 2jsa-10 | THR_A_68_C | ALA_A_69_N | -4.299 | 8.277 | 10.033 | -3.979 | 9.467 | 9.545 | 1.33 | 111.60 | 74.95 |
| 69 | 2jsa-10 | ALA_A_69_C | PRO_A_70_N | -3.521 | 9.0 | 7.201 | -3.996 | 8.986 | 5.927 | 1.36 | 159.54 | -178.31 |
| 70 | 2jsa-10 | PRO_A_70_C | LYS_A_71_N | -2.042 | 8.936 | 4.436 | -1.993 | 10.232 | 4.708 | 1.33 | 78.16 | 87.83 |
| 71 | 2jsa-10 | LYS_A_71_C | ASP_A_72_N | 0.1 | 11.152 | 5.545 | -0.281 | 10.476 | 6.619 | 1.32 | 35.85 | -119.41 |
| 72 | 2jsa-10 | ASP_A_72_C | VAL_A_73_N | 1.239 | 9.163 | 7.993 | 0.593 | 8.114 | 7.509 | 1.32 | 111.45 | -121.63 |
| 73 | 2jsa-10 | VAL_A_73_C | CYS_A_74_N | 2.456 | 6.735 | 6.766 | 2.393 | 7.346 | 5.592 | 1.32 | 152.38 | 95.89 |
| 74 | 2jsa-10 | CYS_A_74_C | THR_A_75_N | 4.482 | 8.495 | 5.18 | 5.049 | 8.284 | 6.358 | 1.32 | 27.18 | -20.41 |
| 75 | 2jsa-10 | THR_A_75_C | LYS_A_76_N | 6.365 | 8.839 | 8.349 | 5.394 | 8.899 | 9.248 | 1.32 | 47.26 | 176.46 |
| 76 | 2jsa-10 | LYS_A_76_C | LEU_A_77_N | 6.101 | 7.077 | 10.697 | 5.533 | 6.266 | 9.815 | 1.33 | 131.69 | -125.01 |
| 77 | 2jsa-10 | LEU_A_77_C | ASN_A_78_N | 7.302 | 4.709 | 9.212 | 7.858 | 5.833 | 8.785 | 1.32 | 108.80 | 63.68 |
| 78 | 2jsa-10 | ASN_A_78_C | GLU_A_79_N | 9.207 | 5.122 | 6.888 | 8.068 | 5.191 | 6.215 | 1.32 | 120.53 | 176.53 |
| 79 | 2jsa-10 | GLU_A_79_C | CYS_A_80_N | 8.231 | 5.551 | 3.799 | 7.2 | 6.283 | 3.404 | 1.32 | 107.35 | 144.63 |
| 80 | 2jsa-10 | CYS_A_80_C | PRO_A_81_N | 8.328 | 8.342 | 2.825 | 9.205 | 8.779 | 1.881 | 1.36 | 133.93 | 26.49 |
| 1 | 2jsa-11 | ARG_A_1_C | SER_A_2_N | 12.734 | 2.179 | -1.501 | 13.017 | 2.959 | -2.534 | 1.32 | 141.23 | 70.06 |
| 2 | 2jsa-11 | SER_A_2_C | ALA_A_3_N | 12.737 | 5.369 | -2.836 | 11.445 | 5.078 | -2.82 | 1.32 | 89.31 | -167.31 |
| 3 | 2jsa-11 | ALA_A_3_C | LEU_A_4_N | 9.138 | 5.324 | -3.552 | 9.243 | 4.263 | -4.339 | 1.33 | 126.43 | -84.35 |
| 4 | 2jsa-11 | LEU_A_4_C | SER_A_5_N | 7.205 | 3.265 | -3.462 | 7.82 | 2.686 | -2.441 | 1.33 | 39.60 | -43.27 |
| 5 | 2jsa-11 | SER_A_5_C | CYS_A_6_N | 6.16 | 3.569 | -0.88 | 6.615 | 4.776 | -1.18 | 1.32 | 103.09 | 69.35 |
| 6 | 2jsa-11 | CYS_A_6_C | GLN_A_7_N | 4.461 | 5.78 | -1.627 | 4.495 | 5.862 | -2.949 | 1.32 | 176.16 | 67.48 |
| 7 | 2jsa-11 | GLN_A_7_C | MET_A_8_N | 2.48 | 4.505 | -3.249 | 3.167 | 3.378 | -3.141 | 1.32 | 85.32 | -58.63 |
| 8 | 2jsa-11 | MET_A_8_C | CYS_A_9_N | 1.666 | 2.411 | -1.452 | 2.239 | 3.144 | -0.51 | 1.32 | 44.64 | 51.98 |
| 9 | 2jsa-11 | CYS_A_9_C | GLU_A_10_N | 0.242 | 4.199 | 0.354 | 0.31 | 4.947 | -0.737 | 1.32 | 145.46 | 84.81 |
| 10 | 2jsa-11 | GLU_A_10_C | LEU_A_11_N | -1.771 | 4.793 | -2.016 | -1.332 | 3.556 | -2.196 | 1.32 | 97.81 | -70.46 |
| 11 | 2jsa-11 | LEU_A_11_C | VAL_A_12_N | -2.981 | 1.778 | -1.985 | -2.367 | 1.369 | -0.885 | 1.32 | 33.85 | -33.67 |
| 12 | 2jsa-11 | VAL_A_12_C | VAL_A_13_N | -4.163 | 1.456 | 0.755 | -3.835 | 2.727 | 0.937 | 1.33 | 82.11 | 75.53 |
| 13 | 2jsa-11 | VAL_A_13_C | LYS_A_14_N | -5.944 | 3.884 | 0.57 | -5.608 | 4.357 | -0.622 | 1.33 | 154.05 | 54.61 |
| 14 | 2jsa-11 | LYS_A_14_C | LYS_A_15_N | -7.562 | 3.415 | -1.726 | -7.004 | 2.242 | -1.466 | 1.32 | 78.68 | -64.56 |
| 15 | 2jsa-11 | LYS_A_15_C | TYR_A_16_N | -8.549 | 0.852 | -0.201 | -7.872 | 1.183 | 0.888 | 1.32 | 34.68 | 26.06 |
| 16 | 2jsa-11 | TYR_A_16_C | GLU_A_17_N | -9.476 | 2.21 | 2.443 | -8.94 | 3.416 | 2.555 | 1.32 | 85.15 | 66.04 |
| 17 | 2jsa-11 | GLU_A_17_C | GLY_A_18_N | -11.006 | 4.544 | 1.879 | -10.773 | 4.167 | 0.63 | 1.33 | 160.46 | -58.28 |
| 18 | 2jsa-11 | GLY_A_18_C | SER_A_19_N | -12.065 | 2.637 | -0.796 | -12.107 | 1.741 | 0.179 | 1.32 | 42.61 | -92.68 |
| 19 | 2jsa-11 | SER_A_19_C | ALA_A_20_N | -13.789 | 0.042 | -0.329 | -14.596 | 0.518 | 0.608 | 1.33 | 45.00 | 149.47 |
| 20 | 2jsa-11 | ALA_A_20_C | ASP_A_21_N | -16.389 | -1.044 | 1.123 | -16.026 | -1.218 | 2.385 | 1.32 | 17.69 | -25.61 |
| 21 | 2jsa-11 | ASP_A_21_C | LYS_A_22_N | -15.421 | -2.552 | 4.328 | -14.275 | -1.892 | 4.254 | 1.32 | 93.20 | 29.94 |
| 22 | 2jsa-11 | LYS_A_22_C | ASP_A_23_N | -12.8 | -3.302 | 5.581 | -12.105 | -3.795 | 4.566 | 1.33 | 139.99 | -35.35 |
| 23 | 2jsa-11 | ASP_A_23_C | ALA_A_24_N | -10.094 | -5.086 | 4.109 | -9.158 | -5.112 | 5.047 | 1.33 | 44.95 | -1.59 |
| 24 | 2jsa-11 | ALA_A_24_C | ASN_A_25_N | -7.455 | -6.169 | 3.667 | -8.371 | -7.124 | 3.587 | 1.33 | 93.46 | -133.81 |
| 25 | 2jsa-11 | ASN_A_25_C | VAL_A_26_N | -8.469 | -7.717 | 1.229 | -9.628 | -7.098 | 1.049 | 1.33 | 97.80 | 151.89 |
| 26 | 2jsa-11 | VAL_A_26_C | ILE_A_27_N | -9.156 | -5.318 | -0.542 | -8.401 | -4.904 | 0.465 | 1.32 | 40.53 | 28.74 |
| 27 | 2jsa-11 | ILE_A_27_C | LYS_A_28_N | -6.17 | -4.164 | -0.164 | -5.418 | -4.777 | 0.74 | 1.33 | 47.02 | -39.19 |
| 28 | 2jsa-11 | LYS_A_28_C | LYS_A_29_N | -4.1 | -5.889 | -0.978 | -5.169 | -6.626 | -1.237 | 1.32 | 101.28 | -145.42 |
| 29 | 2jsa-11 | LYS_A_29_C | ASP_A_30_N | -5.569 | -6.261 | -3.61 | -6.463 | -5.331 | -3.306 | 1.33 | 76.74 | 133.87 |
| 30 | 2jsa-11 | ASP_A_30_C | PHE_A_31_N | -5.515 | -3.594 | -4.723 | -4.693 | -3.219 | -3.755 | 1.32 | 43.03 | 24.52 |
| 31 | 2jsa-11 | PHE_A_31_C | ASP_A_32_N | -2.665 | -3.309 | -5.143 | -2.528 | -4.611 | -4.931 | 1.33 | 80.80 | -83.99 |
| 32 | 2jsa-11 | ASP_A_32_C | ALA_A_33_N | -2.451 | -5.297 | -7.265 | -3.721 | -4.92 | -7.262 | 1.32 | 89.87 | 163.47 |
| 33 | 2jsa-11 | ALA_A_33_C | GLU_A_34_N | -3.887 | -3.515 | -9.242 | -4.234 | -2.342 | -8.732 | 1.33 | 67.37 | 106.48 |
| 34 | 2jsa-11 | GLU_A_34_C | CYS_A_35_N | -2.259 | -1.144 | -9.54 | -1.597 | -1.916 | -8.693 | 1.32 | 50.21 | -49.39 |
| 35 | 2jsa-11 | CYS_A_35_C | LYS_A_36_N | 0.179 | -2.929 | -9.984 | -0.659 | -3.933 | -10.193 | 1.32 | 99.08 | -129.85 |
| 36 | 2jsa-11 | LYS_A_36_C | LYS_A_37_N | -0.907 | -4.166 | -12.604 | -1.372 | -2.934 | -12.459 | 1.32 | 83.72 | 110.68 |
| 37 | 2jsa-11 | LYS_A_37_C | LEU_A_38_N | -0.921 | -0.943 | -13.784 | -0.726 | -0.225 | -12.687 | 1.33 | 34.15 | 74.81 |
| 38 | 2jsa-11 | LEU_A_38_C | PHE_A_39_N | 1.581 | 0.547 | -12.64 | 1.871 | -0.329 | -11.69 | 1.32 | 44.17 | -71.68 |
| 39 | 2jsa-11 | PHE_A_39_C | HIS_A_40_N | 3.565 | -1.901 | -12.53 | 2.645 | -2.095 | -13.464 | 1.33 | 134.81 | -168.09 |
| 40 | 2jsa-11 | HIS_A_40_C | THR_A_41_N | 4.105 | -2.787 | -15.291 | 4.178 | -1.555 | -15.774 | 1.33 | 111.37 | 86.61 |
| 41 | 2jsa-11 | THR_A_41_C | ILE_A_42_N | 6.619 | -1.353 | -15.753 | 6.455 | -1.505 | -14.448 | 1.32 | 9.72 | -137.17 |
| 42 | 2jsa-11 | ILE_A_42_C | PRO_A_43_N | 8.012 | -3.2 | -13.66 | 9.355 | -3.416 | -13.644 | 1.36 | 89.33 | -9.14 |
| 43 | 2jsa-11 | PRO_A_43_C | PHE_A_44_N | 9.728 | -5.498 | -12.39 | 10.433 | -5.003 | -11.383 | 1.33 | 40.54 | 35.07 |
| 44 | 2jsa-11 | PHE_A_44_C | GLY_A_45_N | 9.194 | -5.039 | -9.259 | 8.394 | -4.228 | -9.934 | 1.32 | 120.65 | 134.61 |
| 45 | 2jsa-11 | GLY_A_45_C | THR_A_46_N | 6.273 | -4.689 | -8.779 | 5.534 | -5.278 | -9.709 | 1.33 | 134.54 | -141.44 |
| 46 | 2jsa-11 | THR_A_46_C | ARG_A_47_N | 5.141 | -7.213 | -8.258 | 6.449 | -7.417 | -8.318 | 1.33 | 92.60 | -8.86 |
| 47 | 2jsa-11 | ARG_A_47_C | GLU_A_48_N | 7.049 | -7.607 | -5.953 | 7.425 | -6.337 | -5.904 | 1.33 | 87.88 | 73.51 |
| 48 | 2jsa-11 | GLU_A_48_C | CYS_A_49_N | 5.965 | -5.367 | -4.195 | 5.064 | -5.353 | -5.166 | 1.32 | 137.14 | 179.11 |
| 49 | 2jsa-11 | CYS_A_49_C | ASP_A_50_N | 3.226 | -6.162 | -3.819 | 3.186 | -7.42 | -4.236 | 1.33 | 108.33 | -91.82 |
| 50 | 2jsa-11 | ASP_A_50_C | HIS_A_51_N | 3.696 | -8.473 | -2.102 | 4.883 | -7.914 | -2.285 | 1.32 | 97.94 | 25.22 |
| 51 | 2jsa-11 | HIS_A_51_C | TYR_A_52_N | 5.528 | -6.603 | -0.334 | 4.472 | -5.902 | -0.719 | 1.32 | 106.90 | 146.42 |
| 52 | 2jsa-11 | TYR_A_52_C | VAL_A_53_N | 2.703 | -4.934 | 0.688 | 1.761 | -5.442 | -0.092 | 1.32 | 126.08 | -151.66 |
| 53 | 2jsa-11 | VAL_A_53_C | ASN_A_54_N | 0.525 | -6.596 | 1.658 | 1.671 | -7.251 | 1.787 | 1.33 | 84.42 | -29.75 |
| 54 | 2jsa-11 | ASN_A_54_C | SER_A_55_N | 3.319 | -7.907 | 3.451 | 3.49 | -6.772 | 4.113 | 1.33 | 60.03 | 81.43 |
| 55 | 2jsa-11 | SER_A_55_C | LYS_A_56_N | 4.802 | -4.944 | 5.068 | 4.021 | -4.162 | 4.337 | 1.33 | 123.48 | 134.96 |
| 56 | 2jsa-11 | LYS_A_56_C | VAL_A_57_N | 2.47 | -2.311 | 4.632 | 1.599 | -3.309 | 4.629 | 1.32 | 90.13 | -131.11 |
| 57 | 2jsa-11 | VAL_A_57_C | ASP_A_58_N | -0.335 | -3.189 | 6.101 | 0.041 | -4.294 | 6.726 | 1.32 | 61.83 | -71.21 |
| 58 | 2jsa-11 | ASP_A_58_C | PRO_A_59_N | -0.066 | -3.323 | 8.954 | 1.144 | -2.744 | 8.728 | 1.36 | 99.56 | 25.57 |
| 59 | 2jsa-11 | PRO_A_59_C | ILE_A_60_N | 0.83 | -0.317 | 8.971 | 0.97 | -0.067 | 7.677 | 1.33 | 167.51 | 60.75 |
| 60 | 2jsa-11 | ILE_A_60_C | ILE_A_61_N | -1.118 | 1.174 | 7.541 | -1.867 | 0.118 | 7.257 | 1.33 | 102.37 | -125.35 |
| 61 | 2jsa-11 | ILE_A_61_C | HIS_A_62_N | -3.369 | 0.209 | 9.17 | -2.26 | -0.074 | 9.837 | 1.32 | 59.77 | -14.32 |
| 62 | 2jsa-11 | HIS_A_62_C | GLU_A_63_N | -2.393 | 1.48 | 11.711 | -1.96 | 2.375 | 10.836 | 1.32 | 131.35 | 64.18 |
| 63 | 2jsa-11 | GLU_A_63_C | LEU_A_64_N | -3.467 | 4.304 | 10.753 | -3.825 | 4.123 | 9.491 | 1.32 | 162.37 | -153.18 |

| | | | | | | | | | | | | |
|---|---|---|---|---|---|---|---|---|---|---|---|---|
| 64 | 2jsa-11 | LEU_A_64_C | GLU_A_65_N | -6.213 | 4.007 | 9.944 | -6.067 | 2.737 | 10.289 | 1.32 | 74.90 | -83.44 |
| 65 | 2jsa-11 | GLU_A_65_C | GLY_A_66_N | -6.975 | 2.714 | 12.563 | -5.786 | 3.168 | 12.932 | 1.33 | 73.83 | 20.90 |
| 66 | 2jsa-11 | GLY_A_66_C | GLY_A_67_N | -6.264 | 5.164 | 14.287 | -6.171 | 5.878 | 13.175 | 1.32 | 147.08 | 82.58 |
| 67 | 2jsa-11 | GLY_A_67_C | THR_A_68_N | -5.909 | 8.125 | 12.211 | -4.683 | 7.686 | 11.963 | 1.33 | 100.78 | -19.70 |
| 68 | 2jsa-11 | THR_A_68_C | ALA_A_69_N | -4.319 | 8.634 | 9.734 | -3.879 | 9.704 | 9.087 | 1.33 | 119.22 | 67.65 |
| 69 | 2jsa-11 | ALA_A_69_C | PRO_A_70_N | -3.536 | 9.133 | 6.746 | -4.009 | 9.143 | 5.471 | 1.36 | 159.64 | 178.79 |
| 70 | 2jsa-11 | PRO_A_70_C | LYS_A_71_N | -2.054 | 9.033 | 3.983 | -1.909 | 10.294 | 4.367 | 1.33 | 73.17 | 83.44 |
| 71 | 2jsa-11 | LYS_A_71_C | ASP_A_72_N | 0.126 | 11.271 | 5.274 | -0.204 | 10.514 | 6.311 | 1.33 | 38.53 | -113.55 |
| 72 | 2jsa-11 | ASP_A_72_C | VAL_A_73_N | 1.042 | 9.237 | 7.966 | 0.745 | 8.26 | 7.121 | 1.33 | 129.61 | -106.91 |
| 73 | 2jsa-11 | VAL_A_73_C | CYS_A_74_N | 2.561 | 6.686 | 6.741 | 2.737 | 7.273 | 5.566 | 1.33 | 152.46 | 73.31 |
| 74 | 2jsa-11 | CYS_A_74_C | THR_A_75_N | 5.049 | 7.983 | 5.599 | 4.582 | 8.706 | 6.605 | 1.32 | 40.55 | 122.86 |
| 75 | 2jsa-11 | THR_A_75_C | LYS_A_76_N | 5.533 | 9.021 | 8.841 | 4.528 | 8.236 | 9.201 | 1.33 | 74.24 | -142.01 |
| 76 | 2jsa-11 | LYS_A_76_C | LEU_A_77_N | 5.593 | 6.662 | 10.72 | 5.452 | 5.534 | 10.037 | 1.33 | 121.00 | -97.13 |
| 77 | 2jsa-11 | LEU_A_77_C | ASN_A_78_N | 7.553 | 4.756 | 9.085 | 7.804 | 6.037 | 8.86 | 1.32 | 99.78 | 78.91 |
| 78 | 2jsa-11 | ASN_A_78_C | GLU_A_79_N | 8.805 | 5.492 | 6.71 | 7.616 | 4.99 | 6.413 | 1.32 | 102.96 | -157.11 |
| 79 | 2jsa-11 | GLU_A_79_C | CYS_A_80_N | 8.069 | 4.659 | 4.03 | 7.44 | 5.713 | 3.535 | 1.32 | 111.96 | 120.83 |
| 80 | 2jsa-11 | CYS_A_80_C | PRO_A_81_N | 9.407 | 6.676 | 2.51 | 10.245 | 5.89 | 1.781 | 1.36 | 122.40 | -43.17 |
| 1 | 2jsa-12 | ARG_A_1_C | SER_A_2_N | 11.306 | 5.35 | -1.22 | 11.49 | 6.66 | -1.288 | 1.32 | 92.94 | 82.00 |
| 2 | 2jsa-12 | SER_A_2_C | ALA_A_3_N | 10.278 | 7.427 | -3.267 | 9.502 | 6.435 | -3.678 | 1.32 | 108.07 | -128.03 |
| 3 | 2jsa-12 | ALA_A_3_C | LEU_A_4_N | 8.178 | 5.145 | -5.261 | 8.499 | 3.938 | -4.816 | 1.33 | 70.39 | -75.11 |
| 4 | 2jsa-12 | LEU_A_4_C | SER_A_5_N | 6.882 | 2.629 | -3.554 | 7.702 | 2.34 | -2.554 | 1.33 | 41.00 | -19.41 |
| 5 | 2jsa-12 | SER_A_5_C | CYS_A_6_N | 6.252 | 3.26 | -0.815 | 6.584 | 4.452 | -1.288 | 1.32 | 110.92 | 74.44 |
| 6 | 2jsa-12 | CYS_A_6_C | GLN_A_7_N | 4.441 | 5.486 | -1.723 | 4.487 | 5.726 | -3.025 | 1.32 | 169.37 | 79.15 |
| 7 | 2jsa-12 | GLN_A_7_C | MET_A_8_N | 2.44 | 4.438 | -3.409 | 3.09 | 3.285 | -3.334 | 1.33 | 86.76 | -60.59 |
| 8 | 2jsa-12 | MET_A_8_C | CYS_A_9_N | 1.556 | 2.303 | -1.683 | 2.116 | 3.069 | -0.759 | 1.32 | 45.76 | 53.83 |
| 9 | 2jsa-12 | CYS_A_9_C | GLU_A_10_N | 0.106 | 4.079 | 0.131 | 0.15 | 4.849 | -0.946 | 1.32 | 144.39 | 86.73 |
| 10 | 2jsa-12 | GLU_A_10_C | LEU_A_11_N | -1.96 | 4.646 | -2.17 | -1.483 | 3.43 | -2.392 | 1.32 | 99.65 | -68.58 |
| 11 | 2jsa-12 | LEU_A_11_C | VAL_A_12_N | -3.09 | 1.622 | -2.122 | -2.436 | 1.234 | -1.037 | 1.32 | 35.03 | -30.68 |
| 12 | 2jsa-12 | VAL_A_12_C | VAL_A_13_N | -4.151 | 1.313 | 0.688 | -3.721 | 2.493 | 1.109 | 1.32 | 71.47 | 69.98 |
| 13 | 2jsa-12 | VAL_A_13_C | LYS_A_14_N | -5.764 | 3.788 | 0.848 | -5.397 | 4.36 | -0.29 | 1.33 | 149.15 | 57.32 |
| 14 | 2jsa-12 | LYS_A_14_C | LYS_A_15_N | -7.415 | 3.648 | -1.45 | -6.934 | 2.419 | -1.329 | 1.33 | 84.76 | -68.63 |
| 15 | 2jsa-12 | LYS_A_15_C | TYR_A_16_N | -8.618 | 1.083 | -0.189 | -7.948 | 1.287 | 0.937 | 1.33 | 31.88 | 16.93 |
| 16 | 2jsa-12 | TYR_A_16_C | GLU_A_17_N | -9.438 | 2.391 | 2.551 | -8.74 | 3.478 | 2.849 | 1.33 | 77.01 | 57.29 |
| 17 | 2jsa-12 | GLU_A_17_C | GLY_A_18_N | -10.661 | 4.901 | 2.317 | -10.522 | 4.55 | 1.048 | 1.32 | 163.43 | -68.40 |
| 18 | 2jsa-12 | GLY_A_18_C | SER_A_19_N | -11.986 | 3.298 | -0.482 | -12.205 | 2.338 | 0.405 | 1.33 | 47.99 | -102.85 |
| 19 | 2jsa-12 | SER_A_19_C | ALA_A_20_N | -14.07 | 0.838 | -0.087 | -14.728 | 1.318 | 0.959 | 1.33 | 37.91 | 143.89 |
| 20 | 2jsa-12 | ALA_A_20_C | ASP_A_21_N | -16.583 | -0.13 | 1.578 | -15.59 | -0.998 | 1.709 | 1.33 | 84.33 | -41.16 |
| 21 | 2jsa-12 | ASP_A_21_C | LYS_A_22_N | -15.122 | -2.503 | 3.563 | -14.118 | -1.665 | 3.774 | 1.32 | 80.83 | 39.85 |
| 22 | 2jsa-12 | LYS_A_22_C | ASP_A_23_N | -12.869 | -3.137 | 5.256 | -11.976 | -3.582 | 4.384 | 1.33 | 131.15 | -26.49 |
| 23 | 2jsa-12 | ASP_A_23_C | ALA_A_24_N | -9.985 | -4.922 | 3.978 | -9.066 | -5.22 | 4.884 | 1.32 | 46.84 | -17.97 |
| 24 | 2jsa-12 | ALA_A_24_C | ASN_A_25_N | -7.407 | -6.41 | 3.561 | -8.438 | -7.223 | 3.378 | 1.33 | 97.93 | -141.74 |
| 25 | 2jsa-12 | ASN_A_25_C | VAL_A_26_N | -8.595 | -7.894 | 1.045 | -9.678 | -7.148 | 0.88 | 1.33 | 97.15 | 145.44 |
| 26 | 2jsa-12 | VAL_A_26_C | ILE_A_27_N | -9.186 | -5.406 | -0.746 | -8.487 | -4.932 | 0.276 | 1.33 | 39.57 | 34.14 |
| 27 | 2jsa-12 | ILE_A_27_C | LYS_A_28_N | -6.229 | -4.223 | -0.294 | -5.511 | -4.777 | 0.673 | 1.33 | 43.16 | -37.65 |
| 28 | 2jsa-12 | LYS_A_28_C | LYS_A_29_N | -4.108 | -5.921 | -0.955 | -5.115 | -6.741 | -1.219 | 1.33 | 101.49 | -140.84 |
| 29 | 2jsa-12 | LYS_A_29_C | ASP_A_30_N | -5.382 | -6.426 | -3.617 | -6.385 | -5.578 | -3.445 | 1.32 | 82.54 | 139.79 |
| 30 | 2jsa-12 | ASP_A_30_C | PHE_A_31_N | -5.379 | -3.884 | -4.873 | -4.656 | -3.372 | -3.888 | 1.32 | 41.97 | 35.30 |
| 31 | 2jsa-12 | PHE_A_31_C | ASP_A_32_N | -2.582 | -3.461 | -5.205 | -2.406 | -4.75 | -4.953 | 1.33 | 79.04 | -82.22 |
| 32 | 2jsa-12 | ASP_A_32_C | ALA_A_33_N | -2.207 | -5.43 | -7.28 | -3.527 | -5.486 | -7.376 | 1.32 | 94.16 | -177.57 |
| 33 | 2jsa-12 | ALA_A_33_C | GLU_A_34_N | -3.735 | -4.044 | -9.325 | -4.071 | -2.937 | -8.679 | 1.33 | 60.82 | 106.88 |
| 34 | 2jsa-12 | GLU_A_34_C | CYS_A_35_N | -2.241 | -1.628 | -9.646 | -1.396 | -2.21 | -8.808 | 1.32 | 50.76 | -34.56 |
| 35 | 2jsa-12 | CYS_A_35_C | LYS_A_36_N | 0.194 | -2.964 | -10.466 | -0.578 | -4.022 | -10.663 | 1.32 | 98.55 | -126.12 |
| 36 | 2jsa-12 | LYS_A_36_C | LYS_A_37_N | -1.225 | -3.969 | -13.008 | -1.727 | -2.805 | -12.622 | 1.33 | 73.06 | 113.33 |
| 37 | 2jsa-12 | LYS_A_37_C | LEU_A_38_N | -1.522 | -0.752 | -13.911 | -0.955 | -0.153 | -12.874 | 1.33 | 38.50 | 46.57 |
| 38 | 2jsa-12 | LEU_A_38_C | PHE_A_39_N | 1.38 | 0.528 | -12.976 | 1.665 | -0.241 | -11.935 | 1.33 | 38.23 | -69.66 |
| 39 | 2jsa-12 | PHE_A_39_C | HIS_A_40_N | 3.222 | -2.058 | -12.5 | 2.306 | -2.321 | -13.421 | 1.33 | 134.02 | -163.98 |
| 40 | 2jsa-12 | HIS_A_40_C | THR_A_41_N | 3.639 | -3.48 | -15.1 | 3.713 | -2.446 | -15.924 | 1.32 | 128.48 | 85.91 |
| 41 | 2jsa-12 | THR_A_41_C | ILE_A_42_N | 6.158 | -2.534 | -16.057 | 6.181 | -2.091 | -14.808 | 1.33 | 19.55 | 87.03 |
| 42 | 2jsa-12 | ILE_A_42_C | PRO_A_43_N | 7.478 | -3.741 | -13.576 | 8.742 | -4.243 | -13.608 | 1.36 | 91.35 | -21.66 |
| 43 | 2jsa-12 | PRO_A_43_C | PHE_A_44_N | 8.928 | -5.831 | -11.741 | 9.961 | -5.361 | -11.059 | 1.32 | 59.00 | 24.46 |
| 44 | 2jsa-12 | PHE_A_44_C | GLY_A_45_N | 8.833 | -4.784 | -8.952 | 8.227 | -3.878 | -9.706 | 1.33 | 124.67 | 123.78 |
| 45 | 2jsa-12 | GLY_A_45_C | THR_A_46_N | 5.999 | -4.077 | -8.687 | 5.704 | -5.089 | -9.49 | 1.33 | 127.30 | -106.25 |
| 46 | 2jsa-12 | THR_A_46_C | ARG_A_47_N | 5.216 | -6.987 | -8.02 | 6.485 | -7.347 | -8.146 | 1.33 | 95.46 | -15.84 |
| 47 | 2jsa-12 | ARG_A_47_C | GLU_A_48_N | 7.161 | -7.539 | -5.8 | 7.555 | -6.275 | -5.817 | 1.32 | 90.74 | 72.69 |
| 48 | 2jsa-12 | GLU_A_48_C | CYS_A_49_N | 6.247 | -5.178 | -4.061 | 5.358 | -4.862 | -4.99 | 1.32 | 134.56 | 160.43 |
| 49 | 2jsa-12 | CYS_A_49_C | ASP_A_50_N | 3.378 | -5.727 | -3.904 | 3.618 | -6.905 | -4.462 | 1.33 | 114.90 | -78.48 |
| 50 | 2jsa-12 | ASP_A_50_C | HIS_A_51_N | 3.813 | -8.395 | -2.548 | 4.999 | -7.818 | -2.423 | 1.32 | 84.59 | 25.94 |
| 51 | 2jsa-12 | HIS_A_51_C | TYR_A_52_N | 5.546 | -6.8 | -0.276 | 4.6 | -5.96 | -0.669 | 1.32 | 107.26 | 138.40 |
| 52 | 2jsa-12 | TYR_A_52_C | VAL_A_53_N | 2.949 | -4.999 | 0.878 | 2.027 | -5.685 | 0.217 | 1.33 | 119.91 | -143.35 |
| 53 | 2jsa-12 | VAL_A_53_C | ASN_A_54_N | 0.841 | -7.178 | 1.729 | 1.825 | -7.124 | 2.615 | 1.33 | 48.04 | 3.14 |
| 54 | 2jsa-12 | ASN_A_54_C | SER_A_55_N | 3.251 | -7.876 | 4.437 | 3.239 | -6.658 | 4.959 | 1.33 | 66.80 | 90.56 |
| 55 | 2jsa-12 | SER_A_55_C | LYS_A_56_N | 4.276 | -4.68 | 5.95 | 3.753 | -4.044 | 4.911 | 1.33 | 141.60 | 129.43 |
| 56 | 2jsa-12 | LYS_A_56_C | VAL_A_57_N | 2.178 | -2.193 | 4.785 | 1.327 | -3.205 | 4.702 | 1.32 | 93.59 | -130.06 |
| 57 | 2jsa-12 | VAL_A_57_C | ASP_A_58_N | -0.751 | -3.148 | 5.966 | -0.327 | -4.198 | 6.655 | 1.33 | 58.68 | -68.01 |
| 58 | 2jsa-12 | ASP_A_58_C | PRO_A_59_N | -0.616 | -3.287 | 8.893 | 0.619 | -2.725 | 8.793 | 1.36 | 94.22 | 24.47 |
| 59 | 2jsa-12 | PRO_A_59_C | ILE_A_60_N | 0.325 | -0.3 | 9.078 | 0.488 | -0.078 | 7.781 | 1.33 | 168.01 | 53.71 |
| 60 | 2jsa-12 | ILE_A_60_C | ILE_A_61_N | -1.558 | 1.218 | 7.544 | -2.25 | 0.09 | 7.481 | 1.32 | 92.73 | -121.53 |
| 61 | 2jsa-12 | ILE_A_61_C | HIS_A_62_N | -3.808 | 0.18 | 9.349 | -2.713 | -0.096 | 10.042 | 1.32 | 58.46 | -14.15 |
| 62 | 2jsa-12 | HIS_A_62_C | GLU_A_63_N | -2.794 | 1.443 | 11.931 | -2.292 | 2.316 | 11.07 | 1.32 | 130.53 | 60.10 |
| 63 | 2jsa-12 | GLU_A_63_C | LEU_A_64_N | -3.674 | 4.337 | 11.017 | -4.115 | 4.071 | 9.795 | 1.33 | 157.15 | -148.90 |
| 64 | 2jsa-12 | LEU_A_64_C | GLU_A_65_N | -6.506 | 4.026 | 10.241 | -6.367 | 2.757 | 10.593 | 1.32 | 74.58 | -83.75 |
| 65 | 2jsa-12 | GLU_A_65_C | GLY_A_66_N | -7.247 | 2.653 | 12.875 | -6.063 | 3.149 | 13.205 | 1.33 | 75.58 | 22.73 |
| 66 | 2jsa-12 | GLY_A_66_C | GLY_A_67_N | -6.435 | 5.101 | 14.654 | -6.332 | 5.857 | 13.57 | 1.33 | 144.86 | 82.24 |
| 67 | 2jsa-12 | GLY_A_67_C | THR_A_68_N | -6.003 | 8.123 | 12.674 | -4.793 | 7.658 | 12.4 | 1.32 | 101.94 | -21.02 |
| 68 | 2jsa-12 | THR_A_68_C | ALA_A_69_N | -4.415 | 8.581 | 10.163 | -3.952 | 9.629 | 9.499 | 1.32 | 120.09 | 66.16 |
| 69 | 2jsa-12 | ALA_A_69_C | PRO_A_70_N | -3.521 | 9.111 | 7.16 | -3.975 | 9.09 | 5.878 | 1.36 | 160.48 | -177.35 |
| 70 | 2jsa-12 | PRO_A_70_C | LYS_A_71_N | -1.995 | 9.137 | 4.42 | -1.976 | 10.417 | 4.76 | 1.32 | 75.13 | 89.15 |
| 71 | 2jsa-12 | LYS_A_71_C | ASP_A_72_N | 0.062 | 11.404 | 5.651 | -0.226 | 10.591 | 6.657 | 1.33 | 40.61 | -109.51 |
| 72 | 2jsa-12 | ASP_A_72_C | VAL_A_73_N | 1.265 | 9.272 | 8.054 | 0.564 | 8.213 | 7.68 | 1.32 | 106.41 | -123.50 |
| 73 | 2jsa-12 | VAL_A_73_C | CYS_A_74_N | 2.437 | 6.808 | 7.014 | 2.399 | 7.329 | 5.796 | 1.33 | 156.79 | 94.17 |
| 74 | 2jsa-12 | CYS_A_74_C | THR_A_75_N | 4.454 | 8.526 | 5.353 | 4.828 | 8.539 | 6.623 | 1.32 | 16.42 | 1.99 |
| 75 | 2jsa-12 | THR_A_75_C | LYS_A_76_N | 6.41 | 9.138 | 8.395 | 5.638 | 8.724 | 9.39 | 1.33 | 41.36 | -151.80 |
| 76 | 2jsa-12 | LYS_A_76_C | LEU_A_77_N | 6.692 | 6.807 | 10.456 | 5.904 | 6.046 | 9.71 | 1.33 | 124.25 | -136.00 |
| 77 | 2jsa-12 | LEU_A_77_C | ASN_A_78_N | 7.493 | 4.583 | 8.589 | 7.927 | 5.751 | 8.135 | 1.33 | 110.02 | 69.62 |
| 78 | 2jsa-12 | ASN_A_78_C | GLU_A_79_N | 8.867 | 5.074 | 5.994 | 7.628 | 5.143 | 5.531 | 1.32 | 110.46 | 176.81 |
| 79 | 2jsa-12 | GLU_A_79_C | CYS_A_80_N | 6.728 | 5.483 | 3.279 | 7.643 | 6.306 | 2.788 | 1.33 | 111.75 | 41.97 |

| | | | | | | | | | | | | |
|---|---|---|---|---|---|---|---|---|---|---|---|---|
| 80 | 2jsa-12 | CYS_A_80_C | PRO_A_81_N | 8.499 | 8.235 | 1.608 | 8.985 | 8.804 | 2.744 | 1.36 | 33.37 | 49.50 |
| 1 | 2jsa-13 | ARG_A_1_C | SER_A_2_N | 13.65 | 4.065 | 1.72 | 13.91 | 4.657 | 0.565 | 1.32 | 150.76 | 66.29 |
| 2 | 2jsa-13 | SER_A_2_C | ALA_A_3_N | 12.848 | 5.364 | -1.519 | 11.616 | 5.12 | -1.942 | 1.33 | 108.61 | -168.80 |
| 3 | 2jsa-13 | ALA_A_3_C | LEU_A_4_N | 9.834 | 4.708 | -3.547 | 9.484 | 3.965 | -4.587 | 1.33 | 141.70 | -115.22 |
| 4 | 2jsa-13 | LEU_A_4_C | SER_A_5_N | 7.396 | 3.103 | -3.684 | 8.212 | 2.547 | -2.8 | 1.33 | 48.16 | -34.27 |
| 5 | 2jsa-13 | SER_A_5_C | CYS_A_6_N | 6.651 | 2.822 | -0.938 | 6.96 | 4.11 | -0.917 | 1.32 | 89.09 | 76.51 |
| 6 | 2jsa-13 | CYS_A_6_C | GLN_A_7_N | 4.745 | 5.062 | -1.103 | 4.839 | 5.432 | -2.372 | 1.33 | 163.26 | 75.75 |
| 7 | 2jsa-13 | GLN_A_7_C | MET_A_8_N | 2.77 | 4.254 | -2.944 | 3.4 | 3.091 | -2.867 | 1.32 | 86.67 | -61.56 |
| 8 | 2jsa-13 | MET_A_8_C | CYS_A_9_N | 1.785 | 2.011 | -1.36 | 2.345 | 2.648 | -0.342 | 1.33 | 39.80 | 48.68 |
| 9 | 2jsa-13 | CYS_A_9_C | GLU_A_10_N | 0.36 | 3.674 | 0.583 | 0.48 | 4.524 | -0.427 | 1.33 | 139.64 | 81.96 |
| 10 | 2jsa-13 | GLU_A_10_C | LEU_A_11_N | -1.607 | 4.561 | -1.705 | -1.181 | 3.358 | -2.061 | 1.32 | 105.59 | -70.50 |
| 11 | 2jsa-13 | LEU_A_11_C | VAL_A_12_N | -2.872 | 1.61 | -1.984 | -2.254 | 1.108 | -0.924 | 1.33 | 36.91 | -39.09 |
| 12 | 2jsa-13 | VAL_A_12_C | VAL_A_13_N | -3.994 | 1.088 | 0.776 | -3.542 | 2.231 | 1.271 | 1.33 | 68.06 | 68.42 |
| 13 | 2jsa-13 | VAL_A_13_C | LYS_A_14_N | -5.566 | 3.572 | 1.109 | -5.194 | 4.195 | -0.001 | 1.33 | 146.83 | 59.16 |
| 14 | 2jsa-13 | LYS_A_14_C | LYS_A_15_N | -7.254 | 3.608 | -1.155 | -6.829 | 2.359 | -1.047 | 1.32 | 85.32 | -71.21 |
| 15 | 2jsa-13 | LYS_A_15_C | TYR_A_16_N | -8.503 | 1.009 | 0.092 | -7.777 | 1.132 | 1.193 | 1.32 | 33.77 | 9.62 |
| 16 | 2jsa-13 | TYR_A_16_C | GLU_A_17_N | -9.281 | 2.114 | 2.872 | -8.659 | 3.27 | 3.054 | 1.33 | 82.11 | 61.72 |
| 17 | 2jsa-13 | GLU_A_17_C | GLY_A_18_N | -10.627 | 4.619 | 2.508 | -10.441 | 4.273 | 1.242 | 1.33 | 162.76 | -61.74 |
| 18 | 2jsa-13 | GLY_A_18_C | SER_A_19_N | -12.036 | 2.974 | -0.104 | -12.069 | 2.099 | 0.891 | 1.33 | 41.35 | -92.16 |
| 19 | 2jsa-13 | SER_A_19_C | ALA_A_20_N | -14.053 | 0.673 | 0.86 | -14.494 | 1.092 | 2.037 | 1.32 | 27.33 | 136.47 |
| 20 | 2jsa-13 | ALA_A_20_C | ASP_A_21_N | -16.329 | -0.325 | 2.777 | -15.362 | -1.23 | 2.739 | 1.32 | 91.64 | -43.10 |
| 21 | 2jsa-13 | ASP_A_21_C | LYS_A_22_N | -14.871 | -2.923 | 4.416 | -13.927 | -2.053 | 4.746 | 1.33 | 75.58 | 42.66 |
| 22 | 2jsa-13 | LYS_A_22_C | ASP_A_23_N | -12.568 | -3.642 | 5.991 | -11.833 | -3.974 | 4.94 | 1.32 | 142.50 | -24.31 |
| 23 | 2jsa-13 | ASP_A_23_C | ALA_A_24_N | -9.819 | -5.154 | 4.252 | -8.834 | -5.459 | 5.083 | 1.32 | 51.13 | -17.20 |
| 24 | 2jsa-13 | ALA_A_24_C | ASN_A_25_N | -7.156 | -6.485 | 3.649 | -8.153 | -7.334 | 3.45 | 1.32 | 98.64 | -139.58 |
| 25 | 2jsa-13 | ASN_A_25_C | VAL_A_26_N | -8.328 | -7.943 | 1.1 | -9.469 | -7.281 | 0.975 | 1.33 | 95.41 | 149.88 |
| 26 | 2jsa-13 | VAL_A_26_C | ILE_A_27_N | -9.081 | -5.495 | -0.631 | -8.363 | -5.02 | 0.377 | 1.33 | 40.50 | 33.49 |
| 27 | 2jsa-13 | ILE_A_27_C | LYS_A_28_N | -6.178 | -4.227 | -0.344 | -5.362 | -4.806 | 0.525 | 1.33 | 49.02 | -35.36 |
| 28 | 2jsa-13 | LYS_A_28_C | LYS_A_29_N | -4.101 | -5.895 | -1.249 | -5.192 | -6.612 | -1.473 | 1.32 | 99.74 | -146.69 |
| 29 | 2jsa-13 | LYS_A_29_C | ASP_A_30_N | -5.59 | -6.257 | -3.847 | -6.454 | -5.293 | -3.565 | 1.32 | 77.71 | 131.87 |
| 30 | 2jsa-13 | ASP_A_30_C | PHE_A_31_N | -5.433 | -3.551 | -4.923 | -4.644 | -3.217 | -3.911 | 1.33 | 40.25 | 22.94 |
| 31 | 2jsa-13 | PHE_A_31_C | ASP_A_32_N | -2.595 | -3.24 | -5.27 | -2.399 | -4.54 | -5.108 | 1.32 | 82.98 | -81.43 |
| 32 | 2jsa-13 | ASP_A_32_C | ALA_A_33_N | -2.38 | -5.188 | -7.453 | -3.665 | -4.873 | -7.388 | 1.32 | 87.19 | 166.23 |
| 33 | 2jsa-13 | ALA_A_33_C | GLU_A_34_N | -3.974 | -3.487 | -9.364 | -4.312 | -2.323 | -8.828 | 1.33 | 66.14 | 106.19 |
| 34 | 2jsa-13 | GLU_A_34_C | CYS_A_35_N | -2.409 | -1.051 | -9.699 | -1.692 | -1.759 | -8.838 | 1.33 | 49.49 | -44.64 |
| 35 | 2jsa-13 | CYS_A_35_C | LYS_A_36_N | 0.098 | -2.597 | -10.231 | -0.454 | -3.797 | -10.332 | 1.32 | 94.37 | -114.70 |
| 36 | 2jsa-13 | LYS_A_36_C | LYS_A_37_N | -0.477 | -3.825 | -12.765 | -1.188 | -2.721 | -12.588 | 1.33 | 82.32 | 122.78 |
| 37 | 2jsa-13 | LYS_A_37_C | LEU_A_38_N | -0.586 | -0.636 | -13.69 | -0.78 | 0.2 | -12.681 | 1.32 | 40.38 | 103.06 |
| 38 | 2jsa-13 | LEU_A_38_C | PHE_A_39_N | 1.509 | 1.016 | -12.546 | 1.847 | 0.067 | -11.685 | 1.33 | 49.48 | -70.40 |
| 39 | 2jsa-13 | PHE_A_39_C | HIS_A_40_N | 3.484 | -1.543 | -12.563 | 2.862 | -1.439 | -13.728 | 1.32 | 151.57 | 170.51 |
| 40 | 2jsa-13 | HIS_A_40_C | THR_A_41_N | 4.179 | -2.074 | -15.679 | 5.384 | -2.169 | -15.138 | 1.32 | 65.89 | -4.51 |
| 41 | 2jsa-13 | THR_A_41_C | ILE_A_42_N | 7.774 | -1.66 | -14.965 | 7.511 | -1.019 | -13.835 | 1.33 | 31.51 | 112.31 |
| 42 | 2jsa-13 | ILE_A_42_C | PRO_A_43_N | 9.448 | -2.015 | -12.748 | 10.77 | -1.755 | -12.561 | 1.36 | 82.10 | 11.13 |
| 43 | 2jsa-13 | PRO_A_43_C | PHE_A_44_N | 11.638 | -3.511 | -11.076 | 10.452 | -3.426 | -10.491 | 1.33 | 63.80 | 175.90 |
| 44 | 2jsa-13 | PHE_A_44_C | GLY_A_45_N | 8.959 | -3.44 | -8.538 | 7.943 | -3.237 | -9.364 | 1.33 | 128.56 | 168.70 |
| 45 | 2jsa-13 | GLY_A_45_C | THR_A_46_N | 5.689 | -3.802 | -8.56 | 5.332 | -4.552 | -9.593 | 1.33 | 141.20 | -115.45 |
| 46 | 2jsa-13 | THR_A_46_C | ARG_A_47_N | 4.953 | -6.697 | -8.472 | 6.275 | -6.799 | -8.465 | 1.33 | 89.70 | -4.41 |
| 47 | 2jsa-13 | ARG_A_47_C | GLU_A_48_N | 6.871 | -7.298 | -6.143 | 7.554 | -6.197 | -5.864 | 1.33 | 77.85 | 58.19 |
| 48 | 2jsa-13 | GLU_A_48_C | CYS_A_49_N | 6.144 | -5.474 | -3.998 | 5.225 | -5.293 | -4.934 | 1.32 | 134.98 | 168.86 |
| 49 | 2jsa-13 | CYS_A_49_C | ASP_A_50_N | 3.342 | -6.38 | -3.874 | 3.525 | -7.505 | -4.549 | 1.32 | 120.63 | -80.76 |
| 50 | 2jsa-13 | ASP_A_50_C | HIS_A_51_N | 3.811 | -9.001 | -2.651 | 4.867 | -8.23 | -2.436 | 1.33 | 80.66 | 36.13 |
| 51 | 2jsa-13 | HIS_A_51_C | TYR_A_52_N | 5.303 | -7.131 | -0.305 | 4.35 | -6.331 | -0.76 | 1.32 | 110.09 | 139.99 |
| 52 | 2jsa-13 | TYR_A_52_C | VAL_A_53_N | 2.599 | -5.392 | 0.688 | 1.629 | -5.838 | -0.095 | 1.32 | 126.26 | -155.31 |
| 53 | 2jsa-13 | VAL_A_53_C | ASN_A_54_N | 0.426 | -6.968 | 1.692 | 1.401 | -7.864 | 1.664 | 1.32 | 91.21 | -42.58 |
| 54 | 2jsa-13 | ASN_A_54_C | SER_A_55_N | 2.88 | -8.287 | 3.549 | 3.094 | -6.981 | 3.529 | 1.32 | 90.87 | 80.69 |
| 55 | 2jsa-13 | SER_A_55_C | LYS_A_56_N | 4.173 | -4.872 | 4.134 | 3.684 | -4.28 | 5.213 | 1.32 | 35.44 | 129.56 |
| 56 | 2jsa-13 | LYS_A_56_C | VAL_A_57_N | 2.101 | -2.438 | 5.062 | 1.27 | -3.451 | 4.868 | 1.32 | 98.42 | -129.36 |
| 57 | 2jsa-13 | VAL_A_57_C | ASP_A_58_N | -0.906 | -3.429 | 5.956 | -0.281 | -4.176 | 6.855 | 1.33 | 47.29 | -50.08 |
| 58 | 2jsa-13 | ASP_A_58_C | PRO_A_59_N | -0.713 | -3.234 | 9.057 | 0.509 | -2.64 | 8.998 | 1.36 | 92.49 | 25.92 |
| 59 | 2jsa-13 | PRO_A_59_C | ILE_A_60_N | 0.138 | -0.221 | 9.24 | 0.369 | 0.005 | 7.955 | 1.33 | 165.88 | 44.37 |
| 60 | 2jsa-13 | ILE_A_60_C | ILE_A_61_N | -1.69 | 1.265 | 7.645 | -2.377 | 0.137 | 7.535 | 1.33 | 94.76 | -121.34 |
| 61 | 2jsa-13 | ILE_A_61_C | HIS_A_62_N | -3.993 | 0.347 | 9.341 | -2.972 | -0.003 | 10.11 | 1.33 | 54.53 | -18.92 |
| 62 | 2jsa-13 | HIS_A_62_C | GLU_A_63_N | -2.93 | 1.673 | 11.879 | -2.534 | 2.445 | 10.877 | 1.33 | 139.11 | 62.84 |
| 63 | 2jsa-13 | GLU_A_63_C | LEU_A_64_N | -3.76 | 4.563 | 10.871 | -4.421 | 4.223 | 9.774 | 1.33 | 145.88 | -152.78 |
| 64 | 2jsa-13 | LEU_A_64_C | GLU_A_65_N | -6.758 | 4.222 | 10.452 | -6.477 | 3.017 | 10.925 | 1.32 | 69.08 | -76.87 |
| 65 | 2jsa-13 | GLU_A_65_C | GLY_A_66_N | -7.369 | 3.084 | 13.204 | -6.189 | 3.55 | 13.588 | 1.33 | 73.16 | 21.55 |
| 66 | 2jsa-13 | GLY_A_66_C | GLY_A_67_N | -6.586 | 5.69 | 14.732 | -6.118 | 6.39 | 13.71 | 1.32 | 140.51 | 56.23 |
| 67 | 2jsa-13 | GLY_A_67_C | THR_A_68_N | -5.48 | 8.551 | 12.723 | -4.906 | 7.901 | 11.723 | 1.32 | 139.07 | -48.55 |
| 68 | 2jsa-13 | THR_A_68_C | ALA_A_69_N | -4.303 | 8.587 | 9.449 | -4.398 | 9.807 | 8.94 | 1.33 | 112.59 | 94.45 |
| 69 | 2jsa-13 | ALA_A_69_C | PRO_A_70_N | -3.917 | 9.153 | 6.646 | -4.394 | 8.982 | 5.384 | 1.36 | 158.12 | -160.28 |
| 70 | 2jsa-13 | PRO_A_70_C | LYS_A_71_N | -2.446 | 8.952 | 3.883 | -2.485 | 10.264 | 4.067 | 1.33 | 82.02 | 91.70 |
| 71 | 2jsa-13 | LYS_A_71_C | ASP_A_72_N | -0.443 | 11.376 | 4.786 | -0.657 | 10.632 | 5.861 | 1.32 | 35.76 | -106.05 |
| 72 | 2jsa-13 | ASP_A_72_C | VAL_A_73_N | 0.958 | 9.479 | 7.269 | 0.294 | 8.365 | 7.002 | 1.32 | 101.63 | -120.80 |
| 73 | 2jsa-13 | VAL_A_73_C | CYS_A_74_N | 2.256 | 7.046 | 6.429 | 2.197 | 7.403 | 5.155 | 1.32 | 164.14 | 99.38 |
| 74 | 2jsa-13 | CYS_A_74_C | THR_A_75_N | 4.203 | 8.675 | 4.702 | 4.831 | 8.606 | 5.867 | 1.33 | 28.47 | -6.27 |
| 75 | 2jsa-13 | THR_A_75_C | LYS_A_76_N | 6.233 | 9.382 | 7.72 | 5.344 | 9.205 | 8.686 | 1.32 | 43.18 | -168.74 |
| 76 | 2jsa-13 | LYS_A_76_C | LEU_A_77_N | 6.247 | 7.434 | 10.091 | 5.689 | 6.629 | 9.198 | 1.33 | 132.36 | -124.73 |
| 77 | 2jsa-13 | LEU_A_77_C | ASN_A_78_N | 7.452 | 5.089 | 8.532 | 8.015 | 6.231 | 8.164 | 1.33 | 106.12 | 63.76 |
| 78 | 2jsa-13 | ASN_A_78_C | GLU_A_79_N | 9.329 | 5.483 | 6.257 | 8.144 | 5.399 | 5.672 | 1.32 | 116.22 | -175.95 |
| 79 | 2jsa-13 | GLU_A_79_C | CYS_A_80_N | 7.953 | 5.689 | 3.249 | 7.942 | 6.965 | 3.603 | 1.32 | 74.49 | 90.49 |
| 80 | 2jsa-13 | CYS_A_80_C | PRO_A_81_N | 8.699 | 9.217 | 3.147 | 9.554 | 9.795 | 2.262 | 1.36 | 130.61 | 34.06 |
| 1 | 2jsa-14 | ARG_A_1_C | SER_A_2_N | 12.372 | 5.284 | -3.37 | 11.953 | 5.964 | -4.427 | 1.32 | 142.92 | 121.64 |
| 2 | 2jsa-14 | SER_A_2_C | ALA_A_3_N | 10.391 | 7.819 | -4.736 | 9.477 | 6.88 | -4.54 | 1.32 | 81.49 | -134.23 |
| 3 | 2jsa-14 | ALA_A_3_C | LEU_A_4_N | 7.26 | 5.877 | -4.494 | 7.649 | 4.771 | -5.11 | 1.32 | 117.72 | -70.62 |
| 4 | 2jsa-14 | LEU_A_4_C | SER_A_5_N | 6.573 | 3.403 | -3.409 | 7.543 | 3.683 | -2.55 | 1.33 | 49.61 | 16.10 |
| 5 | 2jsa-14 | SER_A_5_C | CYS_A_6_N | 6.032 | 4.382 | -0.76 | 6.07 | 5.685 | -0.997 | 1.32 | 100.30 | 88.33 |
| 6 | 2jsa-14 | CYS_A_6_C | GLN_A_7_N | 3.745 | 6.05 | -1.554 | 3.871 | 6.253 | -2.858 | 1.33 | 169.62 | 58.17 |
| 7 | 2jsa-14 | GLN_A_7_C | MET_A_8_N | 2.133 | 4.581 | -3.276 | 2.909 | 3.51 | -3.195 | 1.33 | 86.50 | -54.07 |
| 8 | 2jsa-14 | MET_A_8_C | CYS_A_9_N | 1.526 | 2.426 | -1.476 | 2.064 | 3.19 | -0.537 | 1.32 | 44.86 | 54.85 |
| 9 | 2jsa-14 | CYS_A_9_C | GLU_A_10_N | 0.046 | 4.159 | 0.376 | 0.089 | 4.958 | -0.68 | 1.32 | 142.85 | 86.92 |
| 10 | 2jsa-14 | GLU_A_10_C | LEU_A_11_N | -2.011 | 4.784 | -1.927 | -1.534 | 3.572 | -2.169 | 1.32 | 100.53 | -68.52 |
| 11 | 2jsa-14 | LEU_A_11_C | VAL_A_12_N | -3.123 | 1.74 | -1.971 | -2.494 | 1.389 | -0.859 | 1.32 | 32.93 | -29.16 |
| 12 | 2jsa-14 | VAL_A_12_C | VAL_A_13_N | -4.197 | 1.408 | 0.88 | -3.763 | 2.578 | 1.326 | 1.33 | 70.33 | 69.65 |
| 13 | 2jsa-14 | VAL_A_13_C | LYS_A_14_N | -5.806 | 3.885 | 1.129 | -5.449 | 4.478 | -0.001 | 1.33 | 148.51 | 58.95 |
| 14 | 2jsa-14 | LYS_A_14_C | LYS_A_15_N | -7.487 | 3.81 | -1.15 | -7.017 | 2.575 | -1.057 | 1.32 | 85.97 | -69.16 |
| 15 | 2jsa-14 | LYS_A_15_C | TYR_A_16_N | -8.689 | 1.22 | 0.077 | -8.003 | 1.406 | 1.195 | 1.32 | 32.45 | 15.17 |

| | | | | | | | | | | | | |
|---|---|---|---|---|---|---|---|---|---|---|---|---|
| 16 | 2jsa-14 | TYR_A_16_C | GLU_A_17_N | -9.547 | 2.427 | 2.814 | -8.924 | 3.581 | 3.008 | 1.33 | 81.59 | 61.64 |
| 17 | 2jsa-14 | GLU_A_17_C | GLY_A_18_N | -10.955 | 4.847 | 2.491 | -10.83 | 4.455 | 1.232 | 1.32 | 161.90 | -72.31 |
| 18 | 2jsa-14 | GLY_A_18_C | SER_A_19_N | -12.265 | 3.047 | -0.183 | -12.323 | 2.11 | 0.752 | 1.32 | 45.12 | -93.54 |
| 19 | 2jsa-14 | SER_A_19_C | ALA_A_20_N | -14.099 | 0.522 | 0.206 | -14.857 | 0.902 | 1.225 | 1.33 | 39.76 | 153.37 |
| 20 | 2jsa-14 | ALA_A_20_C | ASP_A_21_N | -16.694 | -0.655 | 1.575 | -15.679 | -1.494 | 1.722 | 1.33 | 83.63 | -39.58 |
| 21 | 2jsa-14 | ASP_A_21_C | LYS_A_22_N | -15.245 | -3.143 | 3.459 | -14.47 | -2.164 | 3.905 | 1.33 | 70.34 | 51.63 |
| 22 | 2jsa-14 | LYS_A_22_C | ASP_A_23_N | -13.183 | -3.679 | 5.309 | -12.153 | -3.924 | 4.512 | 1.33 | 126.97 | -13.38 |
| 23 | 2jsa-14 | ASP_A_23_C | ALA_A_24_N | -10.04 | -5.041 | 4.056 | -9.121 | -5.182 | 5.001 | 1.33 | 44.53 | -8.72 |
| 24 | 2jsa-14 | ALA_A_24_C | ASN_A_25_N | -7.302 | -6.112 | 3.681 | -8.192 | -7.075 | 3.494 | 1.32 | 98.12 | -132.74 |
| 25 | 2jsa-14 | ASN_A_25_C | VAL_A_26_N | -8.21 | -7.664 | 1.132 | -9.27 | -6.879 | 1.002 | 1.33 | 95.63 | 143.48 |
| 26 | 2jsa-14 | VAL_A_26_C | ILE_A_27_N | -8.869 | -5.043 | -0.544 | -8.116 | -4.636 | 0.468 | 1.33 | 40.22 | 28.39 |
| 27 | 2jsa-14 | ILE_A_27_C | LYS_A_28_N | -5.909 | -3.777 | -0.085 | -5.322 | -4.742 | 0.607 | 1.32 | 58.51 | -58.69 |
| 28 | 2jsa-14 | LYS_A_28_C | LYS_A_29_N | -3.984 | -5.877 | -1.079 | -5.113 | -6.501 | -1.383 | 1.33 | 103.26 | -151.07 |
| 29 | 2jsa-14 | LYS_A_29_C | ASP_A_30_N | -5.454 | -6.184 | -3.772 | -6.375 | -5.261 | -3.54 | 1.32 | 79.91 | 134.94 |
| 30 | 2jsa-14 | ASP_A_30_C | PHE_A_31_N | -5.361 | -3.531 | -4.919 | -4.617 | -3.115 | -3.905 | 1.32 | 40.05 | 29.21 |
| 31 | 2jsa-14 | PHE_A_31_C | ASP_A_32_N | -2.451 | -3.254 | -5.06 | -2.498 | -4.565 | -4.872 | 1.33 | 81.84 | -92.05 |
| 32 | 2jsa-14 | ASP_A_32_C | ALA_A_33_N | -2.182 | -5.42 | -7.129 | -3.474 | -5.16 | -7.275 | 1.33 | 96.32 | 168.62 |
| 33 | 2jsa-14 | ALA_A_33_C | GLU_A_34_N | -3.531 | -3.861 | -9.332 | -3.987 | -2.696 | -8.895 | 1.33 | 70.75 | 111.38 |
| 34 | 2jsa-14 | GLU_A_34_C | CYS_A_35_N | -2.032 | -1.423 | -9.636 | -1.37 | -1.918 | -8.6 | 1.33 | 38.59 | -36.79 |
| 35 | 2jsa-14 | CYS_A_35_C | LYS_A_36_N | 0.549 | -2.797 | -9.782 | 0.049 | -4.023 | -9.835 | 1.33 | 92.29 | -112.19 |
| 36 | 2jsa-14 | LYS_A_36_C | LYS_A_37_N | 0.004 | -4.308 | -12.251 | -0.66 | -3.164 | -12.16 | 1.33 | 86.06 | 120.13 |
| 37 | 2jsa-14 | LYS_A_37_C | LEU_A_38_N | -0.141 | -1.309 | -13.647 | -0.267 | -0.256 | -12.853 | 1.32 | 53.18 | 96.82 |
| 38 | 2jsa-14 | LEU_A_38_C | PHE_A_39_N | 2.041 | 0.509 | -12.804 | 2.272 | -0.207 | -11.713 | 1.33 | 34.59 | -72.12 |
| 39 | 2jsa-14 | PHE_A_39_C | HIS_A_40_N | 3.934 | -1.98 | -12.086 | 3.396 | -2.119 | -13.289 | 1.33 | 155.21 | -165.51 |
| 40 | 2jsa-14 | HIS_A_40_C | THR_A_41_N | 4.874 | -3.154 | -14.93 | 5.057 | -1.937 | -15.421 | 1.33 | 111.75 | 81.45 |
| 41 | 2jsa-14 | THR_A_41_C | ILE_A_42_N | 7.502 | -1.794 | -15.453 | 7.369 | -1.662 | -14.141 | 1.33 | 8.13 | 135.22 |
| 42 | 2jsa-14 | ILE_A_42_C | PRO_A_43_N | 9.083 | -3.207 | -13.368 | 10.441 | -3.284 | -13.36 | 1.36 | 89.66 | -3.25 |
| 43 | 2jsa-14 | PRO_A_43_C | PHE_A_44_N | 11.031 | -5.336 | -12.139 | 10.364 | -4.719 | -11.174 | 1.33 | 43.28 | 137.23 |
| 44 | 2jsa-14 | PHE_A_44_C | GLY_A_45_N | 8.964 | -4.784 | -9.154 | 8.073 | -4.206 | -9.946 | 1.32 | 126.71 | 147.03 |
| 45 | 2jsa-14 | GLY_A_45_C | THR_A_46_N | 5.934 | -4.739 | -8.856 | 5.572 | -5.66 | -9.736 | 1.32 | 131.65 | -111.46 |
| 46 | 2jsa-14 | THR_A_46_C | ARG_A_47_N | 5.299 | -7.517 | -8.162 | 6.621 | -7.579 | -8.14 | 1.32 | 89.05 | -2.69 |
| 47 | 2jsa-14 | ARG_A_47_C | GLU_A_48_N | 7.098 | -7.538 | -5.739 | 7.307 | -6.23 | -5.774 | 1.33 | 91.51 | 80.92 |
| 48 | 2jsa-14 | GLU_A_48_C | CYS_A_49_N | 5.649 | -5.299 | -4.232 | 4.816 | -5.462 | -5.25 | 1.33 | 140.18 | -168.93 |
| 49 | 2jsa-14 | CYS_A_49_C | ASP_A_50_N | 2.985 | -6.349 | -3.943 | 3.358 | -7.607 | -4.134 | 1.33 | 98.28 | -73.48 |
| 50 | 2jsa-14 | ASP_A_50_C | HIS_A_51_N | 3.983 | -8.48 | -1.951 | 5.051 | -7.724 | -2.163 | 1.33 | 99.20 | 35.29 |
| 51 | 2jsa-14 | HIS_A_51_C | TYR_A_52_N | 5.584 | -6.291 | -0.265 | 4.426 | -5.752 | -0.616 | 1.32 | 105.37 | 155.04 |
| 52 | 2jsa-14 | TYR_A_52_C | VAL_A_53_N | 2.584 | -4.972 | 0.812 | 1.746 | -5.718 | 0.106 | 1.33 | 122.18 | -138.32 |
| 53 | 2jsa-14 | VAL_A_53_C | ASN_A_54_N | 0.686 | -7.362 | 1.553 | 1.671 | -7.246 | 2.432 | 1.33 | 48.45 | 6.72 |
| 54 | 2jsa-14 | ASN_A_54_C | SER_A_55_N | 3.235 | -7.961 | 4.154 | 3.221 | -6.745 | 4.681 | 1.33 | 66.57 | 90.66 |
| 55 | 2jsa-14 | SER_A_55_C | LYS_A_56_N | 4.325 | -4.73 | 5.514 | 3.692 | -4.149 | 4.505 | 1.33 | 139.58 | 137.45 |
| 56 | 2jsa-14 | LYS_A_56_C | VAL_A_57_N | 2.128 | -2.285 | 4.439 | 1.262 | -3.285 | 4.371 | 1.32 | 92.94 | -130.89 |
| 57 | 2jsa-14 | VAL_A_57_C | ASP_A_58_N | -0.703 | -3.24 | 5.805 | -0.318 | -4.363 | 6.393 | 1.32 | 63.65 | -71.08 |
| 58 | 2jsa-14 | ASP_A_58_C | PRO_A_59_N | -0.455 | -3.502 | 8.665 | 0.768 | -2.932 | 8.498 | 1.36 | 97.06 | 24.99 |
| 59 | 2jsa-14 | PRO_A_59_C | ILE_A_60_N | 0.483 | -0.515 | 8.849 | 0.658 | -0.205 | 7.573 | 1.32 | 164.41 | 60.55 |
| 60 | 2jsa-14 | ILE_A_60_C | ILE_A_61_N | -1.392 | 1.103 | 7.493 | -2.156 | 0.049 | 7.242 | 1.33 | 100.91 | -125.94 |
| 61 | 2jsa-14 | ILE_A_61_C | HIS_A_62_N | -3.637 | 0.204 | 9.167 | -2.555 | -0.165 | 9.838 | 1.33 | 59.59 | -18.83 |
| 62 | 2jsa-14 | HIS_A_62_C | GLU_A_63_N | -2.604 | 1.425 | 11.685 | -2.139 | 2.285 | 10.789 | 1.33 | 132.50 | 61.60 |
| 63 | 2jsa-14 | GLU_A_63_C | LEU_A_64_N | -3.592 | 4.255 | 10.816 | -4.147 | 3.914 | 9.662 | 1.33 | 150.56 | -148.43 |
| 64 | 2jsa-14 | LEU_A_64_C | GLU_A_65_N | -6.473 | 3.85 | 10.375 | -6.219 | 2.638 | 10.845 | 1.32 | 69.22 | -78.16 |
| 65 | 2jsa-14 | GLU_A_65_C | GLY_A_66_N | -6.905 | 2.755 | 13.193 | -5.72 | 3.317 | 13.383 | 1.33 | 81.76 | 25.37 |
| 66 | 2jsa-14 | GLY_A_66_C | GLY_A_67_N | -6.044 | 5.434 | 14.589 | -5.964 | 6.067 | 13.428 | 1.32 | 151.21 | 82.80 |
| 67 | 2jsa-14 | GLY_A_67_C | THR_A_68_N | -5.629 | 8.248 | 12.345 | -4.429 | 7.741 | 12.098 | 1.33 | 100.74 | -22.90 |
| 68 | 2jsa-14 | THR_A_68_C | ALA_A_69_N | -4.053 | 8.587 | 9.83 | -3.766 | 9.732 | 9.229 | 1.32 | 116.98 | 75.93 |
| 69 | 2jsa-14 | ALA_A_69_C | PRO_A_70_N | -3.473 | 9.141 | 6.886 | -3.99 | 9.117 | 5.628 | 1.36 | 157.64 | -177.34 |
| 70 | 2jsa-14 | PRO_A_70_C | LYS_A_71_N | -2.086 | 9.009 | 4.076 | -2.011 | 10.31 | 4.312 | 1.32 | 79.74 | 86.70 |
| 71 | 2jsa-14 | LYS_A_71_C | ASP_A_72_N | 0.114 | 11.223 | 5.069 | -0.17 | 10.469 | 6.121 | 1.33 | 37.45 | -110.64 |
| 72 | 2jsa-14 | ASP_A_72_C | VAL_A_73_N | 1.34 | 9.141 | 7.492 | 0.61 | 8.098 | 7.125 | 1.32 | 106.08 | -124.99 |
| 73 | 2jsa-14 | VAL_A_73_C | CYS_A_74_N | 2.412 | 6.623 | 6.42 | 2.386 | 7.232 | 5.243 | 1.33 | 152.62 | 92.44 |
| 74 | 2jsa-14 | CYS_A_74_C | THR_A_75_N | 4.601 | 8.146 | 4.917 | 5.074 | 7.813 | 6.108 | 1.32 | 25.91 | -35.15 |
| 75 | 2jsa-14 | THR_A_75_C | LYS_A_76_N | 6.45 | 8.027 | 8.124 | 5.44 | 7.932 | 8.977 | 1.33 | 49.94 | -174.63 |
| 76 | 2jsa-14 | LYS_A_76_C | LEU_A_77_N | 5.953 | 5.893 | 10.203 | 5.525 | 5.308 | 9.094 | 1.32 | 146.83 | -126.19 |
| 77 | 2jsa-14 | LEU_A_77_C | ASN_A_78_N | 7.131 | 3.705 | 8.213 | 7.912 | 4.775 | 8.239 | 1.32 | 88.88 | 53.87 |
| 78 | 2jsa-14 | ASN_A_78_C | GLU_A_79_N | 9.173 | 4.161 | 6.251 | 8.032 | 4.378 | 5.613 | 1.33 | 118.78 | 169.23 |
| 79 | 2jsa-14 | GLU_A_79_C | CYS_A_80_N | 8.439 | 4.845 | 3.245 | 7.881 | 6.046 | 3.201 | 1.33 | 91.90 | 114.92 |
| 80 | 2jsa-14 | CYS_A_80_C | PRO_A_81_N | 9.888 | 7.125 | 2.39 | 10.516 | 7.712 | 1.336 | 1.36 | 140.80 | 43.07 |
| 1 | 2jsa-15 | ARG_A_1_C | SER_A_2_N | 12.135 | 3.253 | 2.072 | 12.646 | 3.69 | 0.929 | 1.33 | 149.53 | 40.54 |
| 2 | 2jsa-15 | SER_A_2_C | ALA_A_3_N | 12.522 | 5.196 | -0.992 | 11.362 | 4.94 | -1.578 | 1.32 | 116.26 | -167.55 |
| 3 | 2jsa-15 | ALA_A_3_C | LEU_A_4_N | 9.742 | 4.895 | -3.394 | 9.492 | 4.183 | -4.483 | 1.32 | 145.28 | -109.35 |
| 4 | 2jsa-15 | LEU_A_4_C | SER_A_5_N | 7.288 | 3.382 | -3.831 | 7.981 | 2.801 | -2.862 | 1.33 | 43.02 | -39.98 |
| 5 | 2jsa-15 | SER_A_5_C | CYS_A_6_N | 6.269 | 3.141 | -1.152 | 6.711 | 4.364 | -0.901 | 1.32 | 79.08 | 70.13 |
| 6 | 2jsa-15 | CYS_A_6_C | GLN_A_7_N | 4.535 | 5.385 | -1.163 | 4.698 | 5.486 | -2.475 | 1.33 | 171.68 | 31.78 |
| 7 | 2jsa-15 | GLN_A_7_C | MET_A_8_N | 2.621 | 4.332 | -3.064 | 3.202 | 3.142 | -3.045 | 1.32 | 89.18 | -63.98 |
| 8 | 2jsa-15 | MET_A_8_C | CYS_A_9_N | 1.577 | 2.113 | -1.513 | 2.157 | 2.78 | -0.526 | 1.32 | 41.85 | 48.99 |
| 9 | 2jsa-15 | CYS_A_9_C | GLU_A_10_N | 0.193 | 3.825 | 0.422 | 0.292 | 4.66 | -0.603 | 1.33 | 140.64 | 83.24 |
| 10 | 2jsa-15 | GLU_A_10_C | LEU_A_11_N | -1.816 | 4.674 | -1.847 | -1.407 | 3.454 | -2.163 | 1.32 | 103.80 | -71.47 |
| 11 | 2jsa-15 | LEU_A_11_C | VAL_A_12_N | -3.077 | 1.688 | -2.029 | -2.454 | 1.28 | -0.934 | 1.32 | 34.22 | -33.22 |
| 12 | 2jsa-15 | VAL_A_12_C | VAL_A_13_N | -4.136 | 1.279 | 0.824 | -3.775 | 2.536 | 1.032 | 1.32 | 80.96 | 73.98 |
| 13 | 2jsa-15 | VAL_A_13_C | LYS_A_14_N | -5.86 | 3.785 | 0.907 | -5.589 | 4.332 | -0.269 | 1.33 | 152.57 | 63.64 |
| 14 | 2jsa-15 | LYS_A_14_C | LYS_A_15_N | -7.626 | 3.529 | -1.332 | -7.07 | 2.329 | -1.253 | 1.32 | 86.58 | -65.14 |
| 15 | 2jsa-15 | LYS_A_15_C | TYR_A_16_N | -8.603 | 0.868 | -0.053 | -7.885 | 1.097 | 1.037 | 1.33 | 34.66 | 17.69 |
| 16 | 2jsa-15 | TYR_A_16_C | GLU_A_17_N | -9.351 | 2.069 | 2.756 | -8.714 | 3.198 | 3.027 | 1.32 | 78.19 | 60.57 |
| 17 | 2jsa-15 | GLU_A_17_C | GLY_A_18_N | -10.752 | 4.495 | 2.633 | -10.665 | 4.168 | 1.352 | 1.32 | 165.20 | -75.10 |
| 18 | 2jsa-15 | GLY_A_18_C | SER_A_19_N | -12.197 | 2.823 | -0.023 | -12.366 | 1.908 | 0.922 | 1.33 | 44.56 | -100.46 |
| 19 | 2jsa-15 | SER_A_19_C | ALA_A_20_N | -14.237 | 0.378 | 0.564 | -14.864 | 0.842 | 1.635 | 1.32 | 36.07 | 143.50 |
| 20 | 2jsa-15 | ALA_A_20_C | ASP_A_21_N | -16.69 | -0.62 | 2.308 | -15.7 | -1.5 | 2.35 | 1.33 | 88.18 | -41.63 |
| 21 | 2jsa-15 | ASP_A_21_C | LYS_A_22_N | -15.088 | -3.05 | 4.122 | -14.185 | -2.111 | 4.359 | 1.32 | 79.69 | 46.12 |
| 22 | 2jsa-15 | LYS_A_22_C | ASP_A_23_N | -12.788 | -3.6 | 5.683 | -12.012 | -4.015 | 4.693 | 1.32 | 138.37 | -28.14 |
| 23 | 2jsa-15 | ASP_A_23_C | ALA_A_24_N | -9.983 | -5.272 | 4.216 | -9.065 | -5.487 | 5.146 | 1.32 | 45.39 | -13.18 |
| 24 | 2jsa-15 | ALA_A_24_C | ASN_A_25_N | -7.343 | -6.506 | 3.762 | -8.296 | -7.406 | 3.577 | 1.32 | 98.03 | -136.64 |
| 25 | 2jsa-15 | ASN_A_25_C | VAL_A_26_N | -8.442 | -7.926 | 1.204 | -9.659 | -7.424 | 1.049 | 1.33 | 96.72 | 157.58 |
| 26 | 2jsa-15 | VAL_A_26_C | ILE_A_27_N | -9.329 | -5.565 | -0.485 | -8.588 | -5.135 | 0.526 | 1.33 | 40.28 | 30.13 |
| 27 | 2jsa-15 | ILE_A_27_C | LYS_A_28_N | -6.413 | -4.237 | -0.094 | -5.717 | -5.068 | 0.669 | 1.33 | 54.86 | -50.05 |
| 28 | 2jsa-15 | LYS_A_28_C | LYS_A_29_N | -4.362 | -5.961 | -1.144 | -5.404 | -6.703 | -1.488 | 1.32 | 105.05 | -144.55 |
| 29 | 2jsa-15 | LYS_A_29_C | ASP_A_30_N | -5.779 | -6.094 | -3.813 | -6.748 | -5.26 | -3.466 | 1.32 | 74.81 | 139.28 |
| 30 | 2jsa-15 | ASP_A_30_C | PHE_A_31_N | -5.814 | -3.419 | -4.755 | -4.957 | -3.138 | -3.783 | 1.33 | 42.86 | 18.15 |
| 31 | 2jsa-15 | PHE_A_31_C | ASP_A_32_N | -2.991 | -3.114 | -5.26 | -2.631 | -4.376 | -5.082 | 1.32 | 82.28 | -74.08 |

| | | | | | | | | | | | | |
|---|---|---|---|---|---|---|---|---|---|---|---|---|
| 32 | 2jsa-15 | ASP_A_32_C | ALA_A_33_N | -2.647 | -4.854 | -7.468 | -3.971 | -4.82 | -7.411 | 1.33 | 87.54 | 178.53 |
| 33 | 2jsa-15 | ALA_A_33_C | GLU_A_34_N | -4.212 | -3.362 | -9.344 | -4.274 | -2.223 | -8.671 | 1.32 | 59.46 | 93.12 |
| 34 | 2jsa-15 | GLU_A_34_C | CYS_A_35_N | -2.268 | -0.88 | -9.081 | -1.663 | -2.011 | -8.745 | 1.33 | 75.32 | -61.86 |
| 35 | 2jsa-15 | CYS_A_35_C | LYS_A_36_N | 0.412 | -2.642 | -9.814 | 0.07 | -3.893 | -10.089 | 1.33 | 101.97 | -105.29 |
| 36 | 2jsa-15 | LYS_A_36_C | LYS_A_37_N | 0.193 | -3.805 | -12.517 | -0.954 | -3.146 | -12.426 | 1.33 | 86.06 | 150.12 |
| 37 | 2jsa-15 | LYS_A_37_C | LEU_A_38_N | -0.741 | -1.024 | -13.598 | -0.788 | -0.32 | -12.476 | 1.33 | 32.16 | 93.82 |
| 38 | 2jsa-15 | LEU_A_38_C | PHE_A_39_N | 1.356 | 0.822 | -12.63 | 1.974 | -0.019 | -11.814 | 1.32 | 51.98 | -53.69 |
| 39 | 2jsa-15 | PHE_A_39_C | HIS_A_40_N | 3.677 | -1.403 | -12.922 | 2.793 | -1.522 | -13.902 | 1.33 | 137.69 | -172.33 |
| 40 | 2jsa-15 | HIS_A_40_C | THR_A_41_N | 4.282 | -2.415 | -15.614 | 4.861 | -1.231 | -15.477 | 1.33 | 84.07 | 63.94 |
| 41 | 2jsa-15 | THR_A_41_C | ILE_A_42_N | 7.127 | -0.401 | -15.05 | 7.572 | -1.289 | -14.173 | 1.33 | 48.56 | -63.38 |
| 42 | 2jsa-15 | ILE_A_42_C | PRO_A_43_N | 9.4 | -2.098 | -12.786 | 10.696 | -1.787 | -12.517 | 1.36 | 78.59 | 13.49 |
| 43 | 2jsa-15 | PRO_A_43_C | PHE_A_44_N | 11.446 | -3.317 | -10.745 | 10.909 | -4.526 | -10.646 | 1.33 | 85.72 | -113.95 |
| 44 | 2jsa-15 | PHE_A_44_C | GLY_A_45_N | 9.395 | -4.526 | -8.709 | 8.834 | -3.541 | -9.394 | 1.32 | 121.14 | 119.66 |
| 45 | 2jsa-15 | GLY_A_45_C | THR_A_46_N | 6.564 | -3.899 | -8.518 | 6.164 | -4.692 | -9.501 | 1.32 | 137.90 | -116.77 |
| 46 | 2jsa-15 | THR_A_46_C | ARG_A_47_N | 5.614 | -6.695 | -8.201 | 6.915 | -6.944 | -8.193 | 1.32 | 89.65 | -10.83 |
| 47 | 2jsa-15 | ARG_A_47_C | GLU_A_48_N | 7.307 | -7.324 | -5.806 | 7.585 | -6.037 | -5.652 | 1.33 | 83.33 | 77.81 |
| 48 | 2jsa-15 | GLU_A_48_C | CYS_A_49_N | 5.974 | -5.283 | -3.969 | 5.139 | -5.115 | -4.984 | 1.33 | 140.00 | 168.62 |
| 49 | 2jsa-15 | CYS_A_49_C | ASP_A_50_N | 3.266 | -6.135 | -3.845 | 3.381 | -7.349 | -4.362 | 1.32 | 112.98 | -84.59 |
| 50 | 2jsa-15 | ASP_A_50_C | HIS_A_51_N | 3.764 | -8.543 | -2.277 | 4.955 | -7.964 | -2.311 | 1.32 | 91.47 | 25.93 |
| 51 | 2jsa-15 | HIS_A_51_C | TYR_A_52_N | 5.405 | -6.69 | -0.282 | 4.294 | -6.076 | -0.662 | 1.33 | 106.67 | 151.07 |
| 52 | 2jsa-15 | TYR_A_52_C | VAL_A_53_N | 2.462 | -5.197 | 0.721 | 1.62 | -5.926 | 0.001 | 1.33 | 122.88 | -139.11 |
| 53 | 2jsa-15 | VAL_A_53_C | ASN_A_54_N | 0.429 | -7.516 | 1.406 | 1.343 | -7.43 | 2.362 | 1.33 | 43.84 | 5.38 |
| 54 | 2jsa-15 | ASN_A_54_C | SER_A_55_N | 2.784 | -8.219 | 4.157 | 2.832 | -6.981 | 4.626 | 1.32 | 69.27 | 87.78 |
| 55 | 2jsa-15 | SER_A_55_C | LYS_A_56_N | 3.89 | -5.023 | 5.637 | 3.398 | -4.36 | 4.601 | 1.32 | 141.45 | 126.58 |
| 56 | 2jsa-15 | LYS_A_56_C | VAL_A_57_N | 1.826 | -2.503 | 4.544 | 0.974 | -3.506 | 4.399 | 1.32 | 96.29 | -130.35 |
| 57 | 2jsa-15 | VAL_A_57_C | ASP_A_58_N | -1.091 | -3.501 | 5.687 | -0.373 | -4.245 | 6.516 | 1.33 | 51.28 | -46.02 |
| 58 | 2jsa-15 | ASP_A_58_C | PRO_A_59_N | -0.63 | -3.335 | 8.758 | 0.593 | -2.751 | 8.645 | 1.36 | 94.77 | 25.53 |
| 59 | 2jsa-15 | PRO_A_59_C | ILE_A_60_N | 0.255 | -0.331 | 8.913 | 0.401 | -0.118 | 7.613 | 1.33 | 168.76 | 55.57 |
| 60 | 2jsa-15 | ILE_A_60_C | ILE_A_61_N | -1.659 | 1.162 | 7.414 | -2.368 | 0.046 | 7.322 | 1.33 | 93.98 | -122.43 |
| 61 | 2jsa-15 | ILE_A_61_C | HIS_A_62_N | -3.884 | 0.255 | 9.215 | -2.817 | -0.089 | 9.92 | 1.32 | 57.84 | -17.87 |
| 62 | 2jsa-15 | HIS_A_62_C | GLU_A_63_N | -2.644 | 1.568 | 11.7 | -2.205 | 2.327 | 10.707 | 1.32 | 138.56 | 59.96 |
| 63 | 2jsa-15 | GLU_A_63_C | LEU_A_64_N | -3.357 | 4.488 | 10.72 | -3.969 | 4.283 | 9.563 | 1.32 | 150.85 | -161.48 |
| 64 | 2jsa-15 | LEU_A_64_C | GLU_A_65_N | -6.301 | 4.396 | 10.249 | -6.327 | 3.079 | 10.402 | 1.33 | 83.37 | -91.13 |
| 65 | 2jsa-15 | GLU_A_65_C | GLY_A_66_N | -7.061 | 2.941 | 12.734 | -5.822 | 3.318 | 13.013 | 1.32 | 77.84 | 16.92 |
| 66 | 2jsa-15 | GLY_A_66_C | GLY_A_67_N | -5.957 | 5.242 | 14.537 | -5.777 | 6.047 | 13.5 | 1.33 | 141.50 | 77.40 |
| 67 | 2jsa-15 | GLY_A_67_C | THR_A_68_N | -5.375 | 8.29 | 12.577 | -4.219 | 7.76 | 12.203 | 1.33 | 106.39 | -24.63 |
| 68 | 2jsa-15 | THR_A_68_C | ALA_A_69_N | -4.054 | 8.716 | 9.953 | -3.46 | 9.595 | 9.159 | 1.33 | 126.81 | 55.95 |
| 69 | 2jsa-15 | ALA_A_69_C | PRO_A_70_N | -3.317 | 9.132 | 6.774 | -3.946 | 9.12 | 5.568 | 1.36 | 152.45 | -178.91 |
| 70 | 2jsa-15 | PRO_A_70_C | LYS_A_71_N | -2.191 | 9.147 | 3.846 | -1.857 | 10.26 | 4.483 | 1.33 | 61.27 | 73.30 |
| 71 | 2jsa-15 | LYS_A_71_C | ASP_A_72_N | 0.275 | 11.214 | 5.166 | 0.012 | 10.492 | 6.245 | 1.32 | 35.46 | -110.01 |
| 72 | 2jsa-15 | ASP_A_72_C | VAL_A_73_N | 1.529 | 9.151 | 7.593 | 0.949 | 8.166 | 6.924 | 1.32 | 120.34 | -120.49 |
| 73 | 2jsa-15 | VAL_A_73_C | CYS_A_74_N | 2.803 | 6.716 | 6.306 | 2.842 | 7.262 | 5.099 | 1.33 | 155.61 | 85.91 |
| 74 | 2jsa-15 | CYS_A_74_C | THR_A_75_N | 4.911 | 8.456 | 4.725 | 5.111 | 8.58 | 6.03 | 1.33 | 10.22 | 31.80 |
| 75 | 2jsa-15 | THR_A_75_C | LYS_A_76_N | 6.394 | 9.341 | 7.972 | 5.44 | 9.122 | 8.865 | 1.32 | 47.62 | -167.07 |
| 76 | 2jsa-15 | LYS_A_76_C | LEU_A_77_N | 6.425 | 7.411 | 10.29 | 6.025 | 6.567 | 9.349 | 1.33 | 135.21 | -115.36 |
| 77 | 2jsa-15 | LEU_A_77_C | ASN_A_78_N | 7.822 | 5.231 | 8.399 | 8.342 | 6.428 | 8.176 | 1.32 | 99.70 | 66.52 |
| 78 | 2jsa-15 | ASN_A_78_C | GLU_A_79_N | 9.339 | 5.866 | 6.03 | 8.076 | 5.669 | 5.68 | 1.33 | 105.31 | -171.13 |
| 79 | 2jsa-15 | GLU_A_79_C | CYS_A_80_N | 7.632 | 6.009 | 3.296 | 8.57 | 6.944 | 3.288 | 1.32 | 90.35 | 44.91 |
| 80 | 2jsa-15 | CYS_A_80_C | PRO_A_81_N | 9.772 | 8.906 | 2.542 | 10.513 | 9.239 | 1.452 | 1.36 | 143.30 | 24.20 |

Table 4: A reversible spherical geometric conversion of Caenopore-5 backbone peptide bond coordinate matrix to three independent vectors of $\rho$, $\theta$ and $\phi$ [6, 7, 9].

### 3 Supplementary Discussion

#### 3.1 Atomic bond length in ABN-SCS: is $\rho$ constant in practice?

By definition, Cartesian Coordinate System (CCS) requires three parameters, i.e., [$x$, $y$, $z$], to describe protein 3D structure. Here, in this article, the atomic bonding network-based spherical coordinate system (ABN-SCS) also requires three parameters, i.e., [$\rho$, $\theta$, $\phi$], for protein structure description, where $\rho$ is defined as the equilibrium bond length—a physical constant representing inter-nuclear distance that minimizes system energy [1, 2].

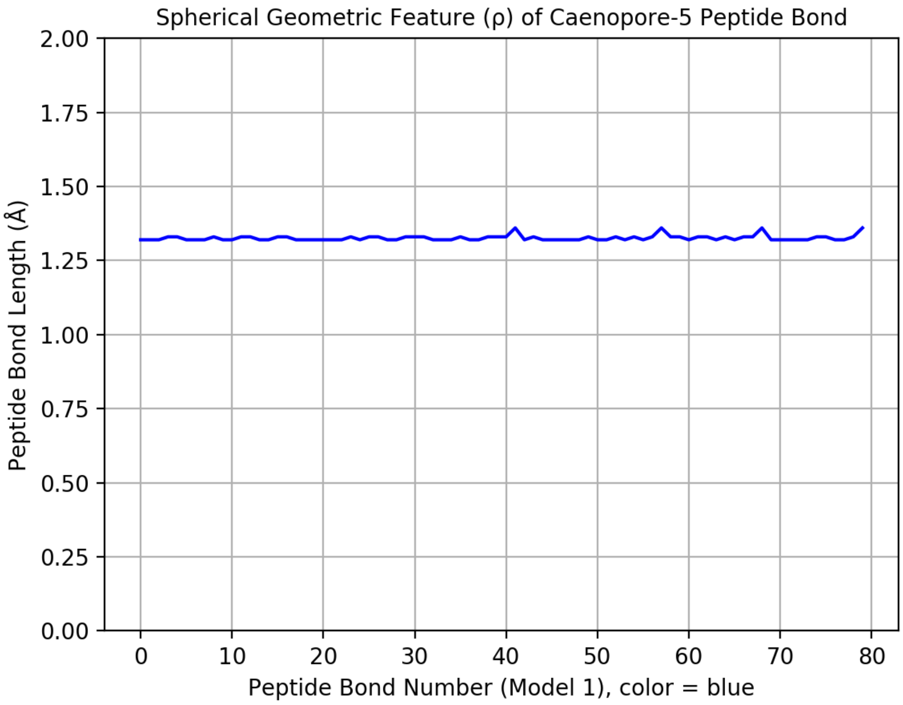


Figure 13: Distribution of the lengths ($\rho$) of 80 peptide bonds in Caenopore-5 [6–8]. This data corresponds to the first structural model in the NMR ensemble (PDB ID: 2JSA), depicted as the top-left subfigure in Figure 7.

In practice, however, $\rho$ is not of a truly fixed value [10]. By definition in the ABN-SCS framework, $\rho$ is the equilibrium bond length, i.e., the distance at which the attractive and repulsive forces between two nuclei are balanced, resulting in the lowest energy configuration [1, 2]. In theory, nonetheless, the value of the equilibrium bond length is highly sensitive to environmental conditions [10, 11], in a manner which is similar to how the precession frequency (i.e., chemical shifts in NMR spectroscopy [8, 12]) of atomic nucleus is sensitive to chemical environmental conditions, where atoms within a molecule are not stationary, but are instead constantly vibrating, twisting, etc [11]. For example, factors such as temperature [13, 14] can influence atomic vibrations or bond stretching, while pH variations can alter protonation states and electrostatic interactions [8, 15–18]. Moreover, the composition of the buffer and the ionic strength [19] of the surrounding medium can affect the local environment, influencing the forces between atoms and slightly altering the bond lengths ($\rho$).

In this article here, with Caenopore-5 [6–8] an example, its 80 peptide bond lengths ($\rho$) appear stable from the looks of it as shown in Figure 13. Yet, there is also slight variations of the values of $\rho$ to a certain degree across its 81 amino acid residues, as shown in Figure 13. These variations are represented by the distribution pattern (Figure 13) and quantitatively described by a statistics summary for the lengths ($\rho$) of 80 peptide bonds in Caenopore-5 (Table 5). Specifically, the 80 bond lengths ($\rho$) possess an average value of approximately 1.326 Å [20, 21] (Figure 13 and Table 5) and a standard deviation of ∼0.009 Å, yielding a ratio of **Std/Mean** at ∼ 0.0069-0.007 (Table 5).

| Model | Mean (Å) | Std (Å) | Min (Å) | Max (Å) | Std/Mean |
|---|---|---|---|---|---|
| 1 | 1.3260 | 0.0092 | 1.3200 | 1.3600 | 0.0070 |
| 2 | 1.3266 | 0.0091 | 1.3200 | 1.3600 | 0.0069 |
| 3 | 1.3265 | 0.0092 | 1.3200 | 1.3600 | 0.0069 |
| 4 | 1.3261 | 0.0092 | 1.3200 | 1.3600 | 0.0069 |
| 5 | 1.3272 | 0.0090 | 1.3200 | 1.3600 | 0.0068 |
| 6 | 1.3261 | 0.0092 | 1.3200 | 1.3600 | 0.0069 |
| 7 | 1.3262 | 0.0092 | 1.3200 | 1.3600 | 0.0069 |
| 8 | 1.3262 | 0.0092 | 1.3200 | 1.3600 | 0.0069 |
| 9 | 1.3274 | 0.0090 | 1.3200 | 1.3600 | 0.0068 |
| 10 | 1.3267 | 0.0091 | 1.3200 | 1.3600 | 0.0069 |
| 11 | 1.3265 | 0.0092 | 1.3200 | 1.3600 | 0.0069 |
| 12 | 1.3272 | 0.0090 | 1.3200 | 1.3600 | 0.0068 |
| 13 | 1.3270 | 0.0091 | 1.3200 | 1.3600 | 0.0068 |
| 14 | 1.3275 | 0.0089 | 1.3200 | 1.3600 | 0.0067 |
| 15 | 1.3267 | 0.0091 | 1.3200 | 1.3600 | 0.0069 |

Table 5: A summary of the statistics for the lengths ($\rho$) of 1200 (80 × 15) peptide bonds in the NMR ensemble of Caenopore-5 (PDB ID: 2JSA). Each row represents one structural model from the NMR ensemble, with Mean, Std, Min, and Max indicating average bond length, standard deviation, minimum, and maximum values of the lengths ($\rho$) of the 1200 peptide bonds, respectively.

Assume that the distribution of $\rho$ is a standard normal one, a data range between **Mean** ± **3** × **Std** encompasses 99.74% of the NMR-observed lengths ($\rho$) of 1200 peptide bonds (Table 5). Given that [$\rho$, $\theta$, $\phi$] is a three-dimension description of protein 3D structure, where each spherical parameter ($\rho$, $\theta$, $\phi$) represents one degree of dimensionality, the variation of $\rho$ represents 3 × **Std/Mean** (Table 5), i.e., 3 × 0.007 = 0.021 degree of dimensionality for the description of 1200 peptide bond in the backbone of Caenopore-5 (PDB ID: 2JSA) [6–8]. Thus, for Caenopore-5, its structure description does not require a three-dimension description with [$x$, $y$, $z$]. Instead, with the ABN-SCS framework proposed here, its structure description requires a 2.021-dimension description with [Std of $\rho$, $\theta$, $\phi$].

Taken together, since $\rho$ is not of a truly fixed value [11] in practice, the ABN-SCS framework proposed here could not induce a dimensionality reduction from three to two. Yet, the ABN-SCS framework does establish a dimensionality reduction from 3 to 2.021 in the case of the 1200 peptide bonds (Table 5) of Caenopore-5 [6–8]. In light of the fact that bonds cannot be compressed or elongated at will, and that equilibrium inter-atomic distances have been comprehensively studied, characterized and tabulated [11], this article argues that this dimensionality reduction (e.g., from 3 to 2.021) is applicable not just to the entire structure of Caenopore-5 [6–8], but also to the entire collection of protein structures [11, 20–22] inside PDB [23–26], and as a result, that this dimensionality reduction in protein structure description with [$\rho$, $\theta$ and $\phi$] could boost computational efficiency in fields such as molecular dynamics simulations [27, 28] in future.

#### 3.2 Explicit geometric features from the structural template in AF2

AlphaFold2 explicitly extracts a series of geometric features from structural templates, storing them in the `FeatureDict` for subsequent use by the Evoformer module of AF2 in protein structure prediction. These explicitly extracted features include:

1. `template_all_atom_positions` — Shape: $[N_{\text{template}}, L, 37, 3]$ (`float32`).
   Defined in
   `alphafold/data/template_features.py`
   and
   `alphafold/model/features.py`;
   used in `alphafold/model/model.py`.
   Stores 3D coordinates of all atoms for each template residue.
2. `template_all_atom_mask` — Shape: $[N_{\text{template}}, L, 37]$ (`float32`).
   Defined in `alphafold/data/template_features.py`;
   used in `alphafold/model/model.py`.
   Indicates which atoms have valid coordinates.
3. `template_aatype` — Shape: $[N_{\text{template}}, L]$ (`int64`).
   Defined in `alphafold/data/template_features.py`;
   used in
   `alphafold/model/features.py`
   and
   `alphafold/model/modules.py`.
   Encodes amino acid types.
4. `template_domain_name` — Shape: $[N_{\text{template}}]$ (`string`).
   Defined in `alphafold/data/template_features.py`;
   used mainly for logging and debugging.
5. `template_sum_probs` — Shape: $[N_{\text{template}}, L, L]$ (`float32`).
   Defined in `alphafold/data/template_features.py`;
   used in `alphafold/model/template.py`
   Stores summed probability scores for residue–residue contacts.
6. `template_pair_residue_mask` — Shape: $[N_{\text{template}}, L, L]$ (`float32`).
   Defined in `alphafold/data/template_features.py`;
   used in `alphafold/model/template_embedding.py`.
   Specifies valid residue–residue pairs.

7. `template_backbone_positions` — Shape: $[N_{\text{template}}, L, 4, 3]$ (`float32`).
   Computed from all-atom positions;
   used in
   `alphafold/model/geometry/vector.py`
   and
   `alphafold/model/folding.py`.
   Contains 3D coordinates for N, $C_\alpha$, C, and O atoms.

8. `template_torsion_angles_sine_cosine` — Shape: $[N_{\text{template}}, L, 7, 2]$ (`float32`).
   Derived internally;
   used in `alphafold/model/all_atom.py`.
   Stores sine and cosine of torsion angles.

9. `template_torsion_angles_mask` — Shape: $[N_{\text{template}}, L, 7]$ (`float32`).
   Derived internally;
   used in `alphafold/model/all_atom.py`.
   Indicates validity of torsion angles.

10. `template_features_mask` — Shape: $[N_{\text{template}}, L]$ (`float32`).
    Generated during filtering;
    used in `alphafold/model/features.py`.
    Marks which template features are valid for each residue.

#### 3.3 Implicitly extracted geometric features by Evoformer of AF2

In addition to template-derived features, AlphaFold2's Evoformer computes a set of implicit geometric quantities on-the-fly during model inference. These features are not stored as static inputs but are generated from sequence and structural context to refine the model's internal representations:

1. **Inter-residue distances** — Shape: $[L, L]$ (`float32`).

   Computed from predicted or template-based $C_\beta$ coordinates ($C_\alpha$ for glycine);

   used in attention biasing and loss functions.

   Represents Euclidean distances between residue pairs.

   Computation logic mainly in `alphafold/model/folding.py` and `alphafold/model/features.py`.

2. **Backbone torsion angles** — Shape: $[L, 7, 2]$ (`float32`, sine–cosine form).

   Derived from backbone N, $C_\alpha$, C, and O positions;

   used in angle prediction heads and structure modules.

   Encodes $\phi$, $\psi$, $\omega$, and side-chain $\chi$ angles.

   Computed in `alphafold/model/all_atom.py`.

3. **Pairwise orientation features** — Shape: $[L, L, 3]$ (`float32`).

   Encodes geometric orientation between residue pairs,

   typically as three angles describing relative frame alignment;

   used in attention biasing.

   Computed in `alphafold/model/geometry/rigid_matrix_vector.py`.

4. **Residue–residue unit vectors** — Shape: $[L, L, 3]$ (`float32`).

   Unit vectors pointing from one residue's $C_\beta$ to another's;

   used in orientation scoring and pair representation updates.

   Computed in `alphafold/model/geometry/vector.py`.

5. **Rigid-body frames** — Shape: $[L, 4, 4]$ (`float32`).

   Homogeneous transformation matrices representing each residue's local coordinate frame;

   used to transform atomic coordinates and compute relative orientations.

   Computed in `alphafold/model/geometry/rigid_matrix_vector.py`.

6. **Frame-to-frame transforms** — Shape: $[L, L, 4, 4]$ (`float32`).

   Relative rigid-body transformations between residue frames;

   supports equivariant reasoning over structure space.

   Calculated within

   `alphafold/model/geometry/rigid_matrix_vector.py`

   and

   `alphafold/model/folding.py`.

7. **Distance and orientation biases** — Shape: variable, `float32`.

   Applied in Evoformer attention modules;

   biases attention maps towards geometrically plausible residue interactions.

   Implemented across `alphafold/model/layer_stack.py` and `alphafold/model/modules.py`.

8. **Clash and geometry validity masks** — Shape: matching geometric feature shapes (`float32`/`bool`).

   Masks out physically impossible or undefined geometric relations;

   used in loss functions and structural filtering.

   Logic found in

   `alphafold/model/all_atom.py`

   and

   `alphafold/model/folding.py`.

Below is a list with complete directory paths for the python scripts inside subfolders of AF2 (AlphaFold2, Version = 2.3.2)), along with their full relative locations:

- `alphafold/model/all_atom.py`
- `alphafold/model/features.py`
- `alphafold/model/folding.py`
- `alphafold/model/geometry/rigid_matrix_vector.py`
- `alphafold/model/geometry/rotation_matrix.py`
- `alphafold/model/geometry/vector.py`
- `alphafold/model/model.py`
- `alphafold/model/modules.py`
- `alphafold/data/feature_processing.py`
- `alphafold/data/templates.py`

---

Below is a table of the Python scripts of AlphaFold2 (Version = 2.3.2) involved in protein structure feature extraction and processing.

| **Python Script Name** | **Role** |
|---|---|
| `all_atom.py` | Both |
| `features.py` | Both |
| `folding.py` | Both |
| `rigid_matrix_vector.py` | Implicit |
| `rotation_matrix.py` | Implicit |
| `geometry/vector.py` | Both |
| `model.py` | Explicit |
| `modules.py` | Explicit |
| `feature_processing.py` | Implicit |
| `templates.py` | Explicit |

Table 6: AlphaFold2's (Version = 2.3.2) Python scripts involved in geometric feature processing. The *Role* column indicates whether the script is used exclusively for template-derived features (Explicit), implicit Evoformer-computed features (Implicit), or in both contexts (Both).

[21] Protein peptide bond structure;. Available from: https://www.chem.uzh.ch/dam/jcr:ffffffff-9520-0a4a-0000-0000776b0b35/lecture_2.pdf.
