## Supplementary file 3 for "Protein Structure Description with *ρ, θ* and *ϕ*: A Case Study with Caenopore-5"

```
import os
import math
import time



def dis(x,y):
	return ((x[0]-y[0])**2+(x[1]-y[1])**2+(x[2]-y[2])**2)**0.5

def dis2(x,y,z):
	return (x**2+y**2-2*x*y*math.cos(z))**(0.5)



def conv_1(x,y,z):
	return [x*math.sin(y)*math.cos(z),x*math.sin(y)*math.sin(z), x*math.cos(y)]

def conv_2(coor_initial,coor_final):
	x=coor_final[0]-coor_initial[0]
	y=coor_final[1]-coor_initial[1]
	z=coor_final[2]-coor_initial[2]
	r=(x**2+y**2+z**2)**(0.5)
	temp=(x**2+y**2)**(0.5)
	theta=math.atan2(temp,z)*180/math.pi
	phi=math.atan2(y,x)*180/math.pi
	return [r, theta, phi]



files=[]



for i in range(1,16):
	f='2jsa-' + str(i) + '.pdb'
	files.append(f)

backboneAtoms=[]
backboneAtoms.append('H')#N-terminal hydrogen, first atom
backboneAtoms.append('N')#N-terminal nitrogen
backboneAtoms.append('CA')#carbon alfa
backboneAtoms.append('HA')#hydrogen alfa
backboneAtoms.append('C')#carbonyl carbon
backboneAtoms.append('O')#carbonyl oxygen
```

```
for f in files:
	reslist=[]
	dicCoor={}
	dicAtomCoor={}
	dicAtom={}
#	print f
	w=open(f).readlines()
	for l in w:
		if 'ATOM' == l[:4]:
#			print l[:-1]
			r=l.split()[3]+'_'+l.split()[4]+'_'+l.split()[5]
			if not r in reslist:
				reslist.append(r)
	for r in reslist:
#		print r
		dicCoor[r]=[]
		dicAtom[r]=[]
	for l in w:
		if 'ATOM' == l[:4] and l.split()[2] in backboneAtoms:
#			print l[:-1]
			r=l.split()[3]+'_'+l.split()[4]+'_'+l.split()[5]
			x=l[30:38]
			y=l[38:46]
			z=l[46:54]
			c=[float(x),float(y),float(z)]
			dicCoor[r].append(c)
			dicAtomCoor[r+'_'+l.split()[2]]=c
			dicAtom[r].append(l.split()[2])
	for r in reslist:
		for j in range(len(dicCoor[r])):
#			print r,dicCoor[r][j],dicAtom[r][j]
			pass


	newDic={}
	for r in reslist:
		for j in range(len(dicCoor[r])):
			newDic[r+'_'+dicAtom[r][j]]=dicCoor[r][j]
	for r in newDic:
		#print f+'_'+r,newDic[r][0],newDic[r][1],newDic[r][2]
		pass
```

```
    covalentbondList=[]
#   print len(reslist)
    for i in range(len(reslist)-1):
        r=reslist[i]
        rn=reslist[i+1]
        if backboneAtoms[1] in dicAtom[r] and backboneAtoms[0] in dicAtom[r]:
            covalentbondList.append([r + '_'+backboneAtoms[1],r + '_'+backboneAtoms[0]])
        if backboneAtoms[1] in dicAtom[r] and backboneAtoms[2] in dicAtom[r]:
            covalentbondList.append([r + '_'+backboneAtoms[1],r + '_'+backboneAtoms[2]])
        if backboneAtoms[2] in dicAtom[r] and backboneAtoms[3] in dicAtom[r]:
            covalentbondList.append([r + '_'+backboneAtoms[2],r + '_'+backboneAtoms[3]])
        if backboneAtoms[2] in dicAtom[r] and backboneAtoms[4] in dicAtom[r]:
            covalentbondList.append([r + '_'+backboneAtoms[2],r + '_'+backboneAtoms[4]])
        if backboneAtoms[4] in dicAtom[r] and backboneAtoms[5] in dicAtom[r]:
            covalentbondList.append([r + '_'+backboneAtoms[4],r + '_'+backboneAtoms[5]])
        if backboneAtoms[4] in dicAtom[r] and backboneAtoms[1] in dicAtom[rn]:
            covalentbondList.append([r + '_'+backboneAtoms[4],rn + '_'+backboneAtoms[1]])



    r=reslist[80]
    if backboneAtoms[1] in dicAtom[r] and backboneAtoms[0] in dicAtom[r]:
        covalentbondList.append([r + '_'+backboneAtoms[1],r + '_'+backboneAtoms[0]])
    if backboneAtoms[1] in dicAtom[r] and backboneAtoms[2] in dicAtom[r]:
        covalentbondList.append([r + '_'+backboneAtoms[1],r + '_'+backboneAtoms[2]])
    if backboneAtoms[2] in dicAtom[r] and backboneAtoms[3] in dicAtom[r]:
        covalentbondList.append([r + '_'+backboneAtoms[2],r + '_'+backboneAtoms[3]])
    if backboneAtoms[2] in dicAtom[r] and backboneAtoms[4] in dicAtom[r]:
        covalentbondList.append([r + '_'+backboneAtoms[2],r + '_'+backboneAtoms[4]])
    if backboneAtoms[4] in dicAtom[r] and backboneAtoms[5] in dicAtom[r]:
        covalentbondList.append([r + '_'+backboneAtoms[4],r + '_'+backboneAtoms[5]])


#   print dicAtomCoor
    t=1
    for bond in covalentbondList:
        coor1=dicAtomCoor[bond[0]]
        coor2=dicAtomCoor[bond[1]]
```

```
            sphere=conv_2(coor1,coor2)
            if t % 2==1:
                  print
t,'\t&\t',f.split('.')[0],'\t&\t',bond[0],'\t&\t',bond[1],'\t&\t',coor1[0],'\t&\t',coor1[1],'\t&\t',coor1
[2],'\t&\t',coor2[0],'\t&\t',coor2[1],'\t&\t',coor2[2],'\t&\t','%.2f'%(sphere[0]),'\t&\t','%.2f'%(spher
e[1]),'\t&\t','%.2f'%(sphere[2]),'\\\\\\hline\\rowcolor{gray}'
            else:
                  print
t,'\t&\t',f.split('.')[0],'\t&\t',bond[0],'\t&\t',bond[1],'\t&\t',coor1[0],'\t&\t',coor1[1],'\t&\t',coor1
[2],'\t&\t',coor2[0],'\t&\t',coor2[1],'\t&\t',coor2[2],'\t&\t','%.2f'%(sphere[0]),'\t&\t','%.2f'%(spher
e[1]),'\t&\t','%.2f'%(sphere[2]),'\\\\\\hline'
            t+=1



      print '\n\n\nkiwikiwikiwi\n\n\n'
```
